# Remnants of repetitive DNA shape the earliest stages of *de novo* gene evolution in insects

**DOI:** 10.64898/2026.07.07.736878

**Authors:** Ryuto Sanno, Kazuhiro Satomura, Yuka Azami, Shota Hayakawa, Kazuya Hirata, Ken Naito, Takeshi Suzuki, Atsushi Ogura, Kei Yura, Toru Asahi, Cassandra G. Extavour, Kosuke Kataoka

**Author notes:** Equal contribution: Ryuto Sanno and Kazuhiro Satomura; These authors contributed equally to this work. Corresponding authors: Kazuhiro Satomura,; Kosuke Kataoka.

## Abstract

A fundamental unresolved question in genome evolution is how novel genes arise from noncoding DNA and become fixed within stable gene repertoires. Here, we performed comparative genomic analyses across evolutionary timescales in insects using chromosome-scale genome assemblies of two cricket species, *Teleogryllus occipitalis* and *Tarbinskiellus portentosus*. Using conservative criteria, we identified 41 *de novo* gene candidates derived from intergenic regions in the *Te. occipitalis* lineage. These genes are simple and compact, exhibit hallmarks of evolutionarily young genes, and frequently contain fragments of transposable elements and simple sequence repeats. Across insects, repetitive sequence fragments retained in older genes show positional correspondence despite extensive nucleotide divergence and lack of conserved sequence motifs. Approximately 24% of gene families retained since the last common ancestor of insects contain such repetitive sequence fragments, whereas recent insertions of repetitive sequence into established exons appear to be strongly constrained. These findings support a model in which evolutionary constraints on sequence incorporation change during gene evolution: repetitive DNA supplies mutable sequence material, rather than pre-existing functional modules, at the earliest stages of *de novo* gene emergence, but its incorporation becomes increasingly constrained once genes are established.

## INTRODUCTION

The ultimate origin of eukaryotic genes remains one of the central unresolved questions in evolutionary biology. Genes are widely thought to arise through gene duplication followed by functional divergence, a process that generates gene families across genomes (Breathnach and Chambon 1981; Nei and Rooney 2005). Yet a substantial fraction of genes in many eukaryotic genomes lacks detectable homologues and cannot be readily explained by duplication alone (Khalturin et al. 2009; Wissler et al. 2013; Palmieri et al. 2014; Tautz and Domazet-Lošo 2011). Mounting evidence across diverse eukaryotes suggests that some of these orphan genes emerge as *de novo* genes from non-genic DNA within ancestral genomes (Knowles and McLysaght 2009; Carvunis et al. 2012; Wu et al. 2011; Vakirlis et al. 2018; Van Oss and Carvunis 2019; Zhang et al. 2019).

That said, estimates of *de novo* gene numbers vary widely across studies (McLysaght and Hurst 2016; Blevins et al. 2021), reflecting methodological differences as well as a fundamental ambiguity in defining the boundary between non-coding and coding sequences (McLysaght and Hurst 2016; Van Oss and Carvunis 2019; Grandchamp et al. 2025). Intergenic DNA is not a reservoir of entirely novel material but instead comprises an evolutionarily dynamic mosaic of accumulated transposable elements (TEs), pseudogenes, and other remnants (Kazazian 2004). One influential model for *de novo* gene evolution is the proto-gene hypothesis, which proposes that translation of short and unstable open reading frames from intergenic DNA may generate transient polypeptides with limited structural stability and high rates of evolutionary turnover (Carvunis et al. 2012). These immature gene-like entities—proto-genes—are considered intermediate stages between non-genic DNA and fully established functional genes, suggesting that gene evolution may proceed as a continuum from unstable proto-genes to evolutionarily stable genes (McLysaght and Hurst 2016; Van Oss and Carvunis 2019; Grandchamp et al. 2025). Consistent with this view, *de novo* genes typically exhibit characteristics of young genes, including short sequence lengths, fewer introns, and restricted gene expression (Carvunis et al. 2012; Heinen et al. 2009; Levine et al. 2006). However, whether *de novo* genes represent transient evolutionary trials or can persist long enough to contribute to deeply conserved gene families remains unclear (Bornberg-Bauer et al. 2021).

Most insights into *de novo* gene evolution derive from a limited number of model organisms, particularly *Drosophila melanogaster* in insects (Levine et al. 2006; Neme and Tautz 2016; Durand et al. 2016b; Mikina et al. 2024). In *Drosophila*, intergenic regions are typically short, constraining the amount of non-genic DNA available as potential raw material for gene emergence (Roy 2004; Rajon and Masel 2011). In contrast, crickets (Polyneoptera) are phylogenetically distant from holometabolous insects such as *Drosophila* and possess expansive intergenic regions. Incorporating such divergent genomes enables a less biased comparative analysis of gene evolution across insects. Here, we investigate whether genes originated from intergenic regions in the cricket *Teleogryllus occipitalis* (Supplemental Fig. S1) exhibit hallmarks of evolutionarily young genes and whether deeply conserved insect genes retain detectable traces of intergenic sequence material. By examining sequence features, including the remnants of sequence material used during *de novo* gene birth, across genes of different evolutionary ages, we test whether gene evolution represents a continuous process from non-genic DNA to evolutionarily stable genes.

## RESULTS

### Chromosome-Scale Genome Assembly of Crickets

To establish chromosome-scale genomic resources for analyses of *de novo* gene birth, we sequenced and assembled the genomes of *Te. occipitalis* and a phylogenetic outgroup within the subfamily Gryllinae, *Tarbinskiellus portentosus.* After estimating genome size by k-mer analysis (Supplemental Fig. S2), we integrated long-read, short-read, and Omni-C sequencing and adopted the best-quality genome using multiple assemblers (Supplemental Tables S1–S2). After polishing, haplotype removal, and elimination of mitochondrial and bacterial genome contamination, clustered super-scaffolds covered 95.02% (*Te. occipitalis*) and 86.80% (*Ta. portentosus*) of the whole genome (Table 1, Supplemental Fig. S3, Supplemental Table S3). Scaffold numbers (14 and 6, respectively) matched karyotypic data for *Te. occipitalis* and a close relative of *Ta. portentosus* (You and Xie 2003; Manna and Bhattacharjee 1964) (Fig. 1A, B). Mapping depth of short-read sequencing for *Te. occipitalis* indicated that Chr 1 is the X chromosome (Fig. 1C). Synteny analyses with *Ta. portentosus* and the published *Acheta domesticus* genome revealed chromosome-wide collinearity (Dossey et al. 2023) (Fig. 1D). Syntenic blocks on the X chromosome tended to be larger than those on autosomes, although the difference was not statistically significant (vs. *A. domesticus*, P = 0.096; vs. *Ta. portentosus*, P = 0.082; Pearson’s χ^2^ test) (Fig. 1E). The Benchmarking Universal Single-Copy Orthologs (v5; Arthropoda odb10; Manni et al. 2021) recovered 99.2% and 90.4% of expected genes, respectively (Simão et al. 2015).

**Fig. 1.**
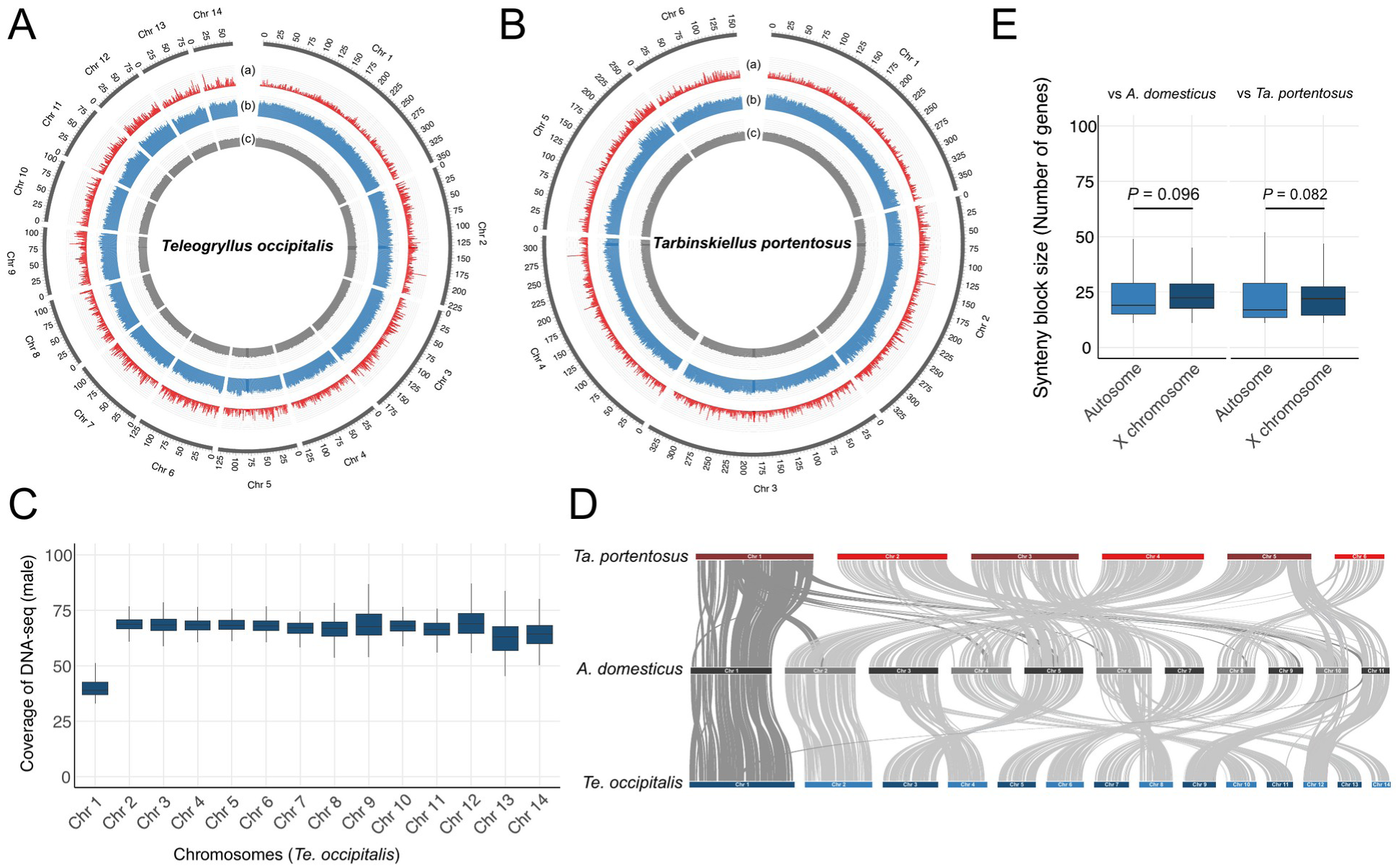
Genome-wide features and synteny of Gryllidea crickets. (**A**) Circos plot of the *Teleogryllus occipitalis* genome showing chromosomes, gene density, repeat density, and GC content. (**B**) Equivalent Circos plot for *Tarbinskiellus portentosus*. (**C**) Male DNA-seq coverage along *Te. occipitalis* chromosomes. Coverage on Chr 1 (X) is ≈½ that of autosomes, consistent with an XO system. (**D**) Pairwise synteny among *Te. occipitalis*, *Ta. portentosus*, and *A. domesticus*. Dark-gray links denote X-chromosome blocks; light-gray links denote autosomal blocks. (**E**) Box plots of syntenic block sizes (number of genes per block) on autosomes and the X chromosome for the two species pairs. X-chromosome blocks tended to be larger than autosomal blocks, but the differences were not significant (Pearson’s χ^2^ test).

**Table 1.**
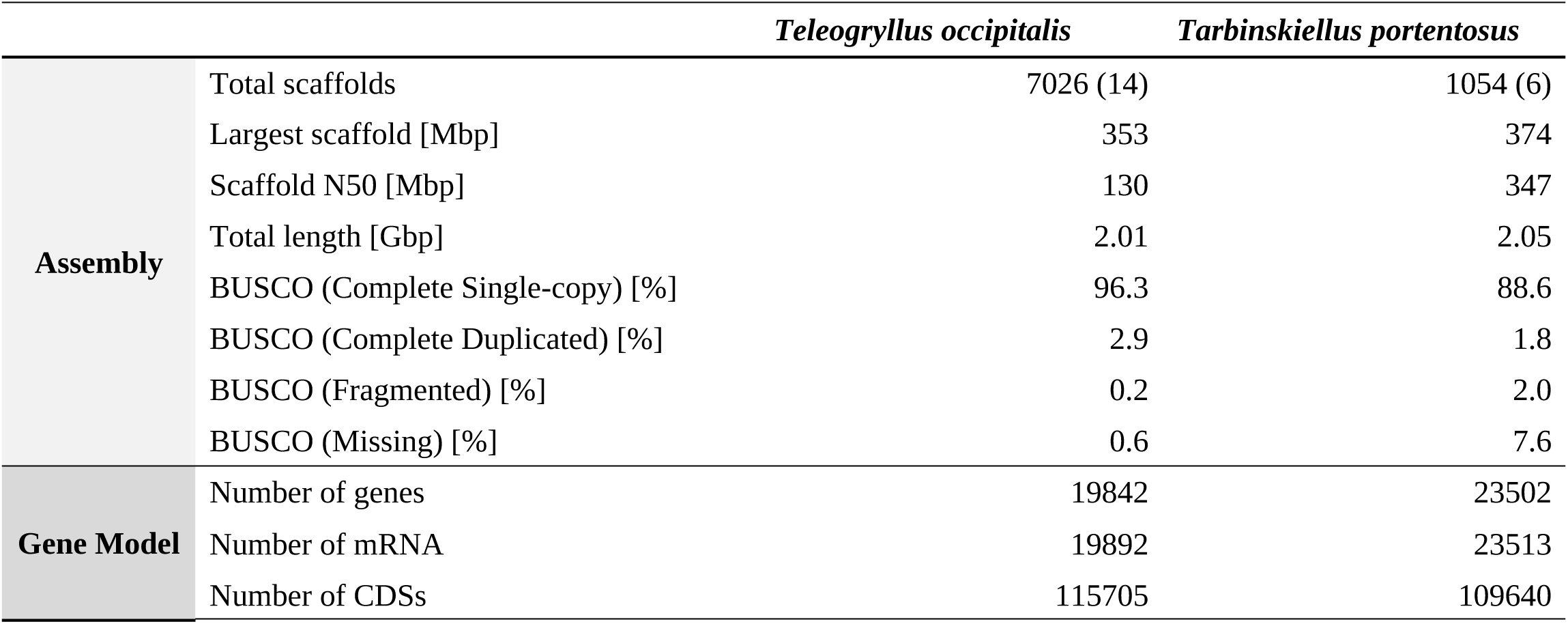
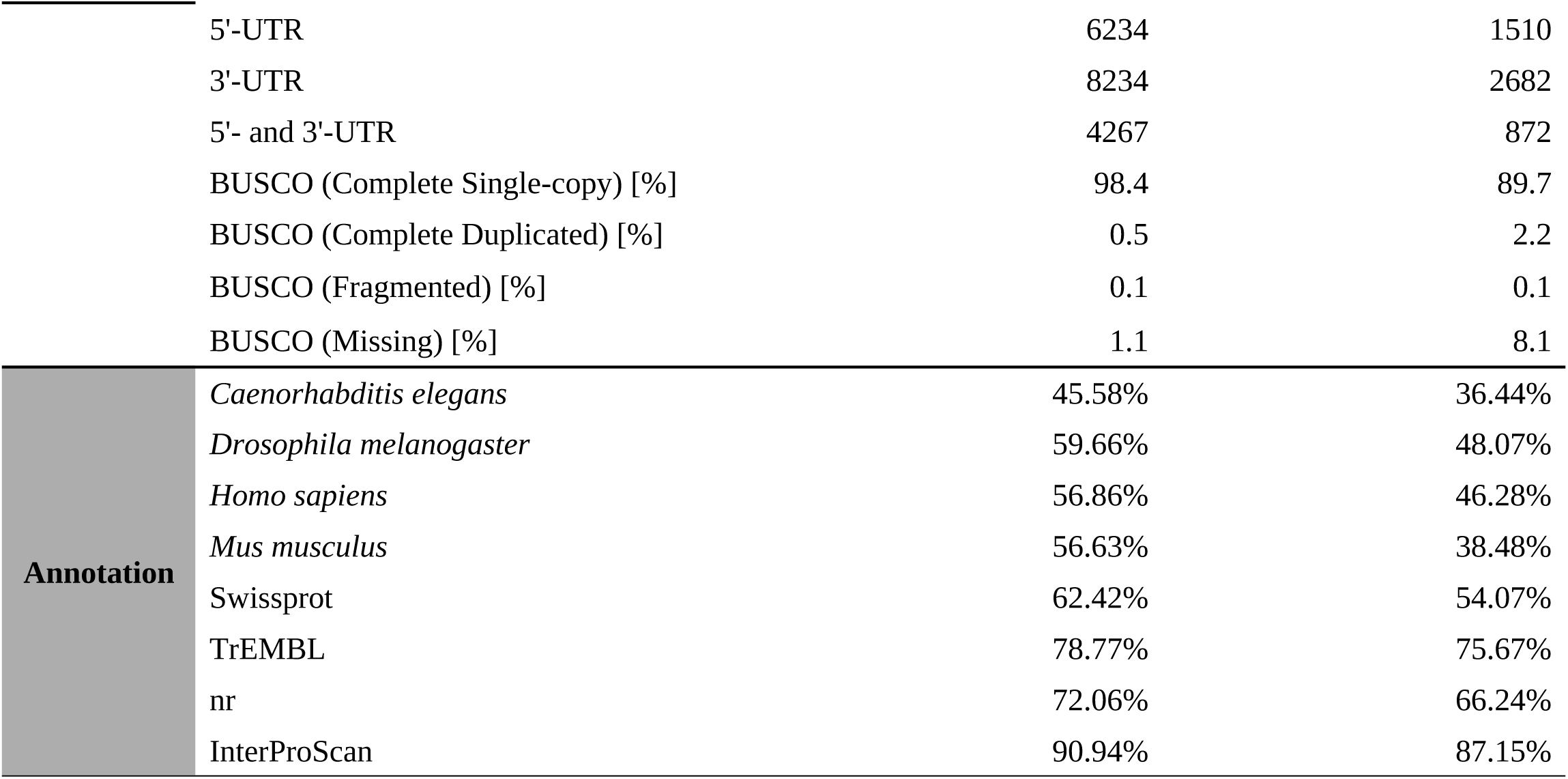
Genome assembly, annotation, and functional assignment statistics. Summary metrics for chromosome-scale assemblies and annotated gene models of *Teleogryllus occipitalis* and *Tarbinskiellus portentosus*. Assembly statistics include scaffold count (total and Omni-C–anchored, in parentheses), largest scaffold length, scaffold N50, total assembly size, and BUSCO completeness scores. Gene-model statistics list the numbers of predicted genes, mRNAs, coding sequences (CDSs), and UTR features, together with BUSCO assessments of the annotated proteomes. Functional assignment statistics report the percentage of predicted proteins with significant sequence similarity to reference proteomes (*C. elegans*, *D. melanogaster*, *H. sapiens*, *M. musculus*), UniProt/Swiss-Prot, UniProt/TrEMBL, and NCBI nr (BLASTP; E-value < 1 × 10^−6^), and the percentage with InterProScan domain signatures

|  |  | *Teleogryllus occipitalis* | *Tarbinskiellus portentosus* |
|---|---|---|---|
| **Assembly** | Total scaffolds | 7026 (14) | 1054 (6) |
|  | Largest scaffold [Mbp] | 353 | 374 |
|  | Scaffold N50 [Mbp] | 130 | 347 |
|  | Total length [Gbp] | 2.01 | 2.05 |
|  | BUSCO (Complete Single-copy) [%] | 96.3 | 88.6 |
|  | BUSCO (Complete Duplicated) [%] | 2.9 | 1.8 |
|  | BUSCO (Fragmented) [%] | 0.2 | 2.0 |
|  | BUSCO (Missing) [%] | 0.6 | 7.6 |
| **Gene Model** | Number of genes | 19842 | 23502 |
|  | Number of mRNA | 19892 | 23513 |
|  | Number of CDSs | 115705 | 109640 |
|  | 5'-UTR | 6234 | 1510 |
|  | 3'-UTR | 8234 | 2682 |
|  | 5'- and 3'-UTR | 4267 | 872 |
|  | BUSCO (Complete Single-copy) [%] | 98.4 | 89.7 |
|  | BUSCO (Complete Duplicated) [%] | 0.5 | 2.2 |
|  | BUSCO (Fragmented) [%] | 0.1 | 0.1 |
|  | BUSCO (Missing) [%] | 1.1 | 8.1 |
| **Annotation** | *Caenorhabditis elegans* | 45.58% | 36.44% |
|  | *Drosophila melanogaster* | 59.66% | 48.07% |
|  | *Homo sapiens* | 56.86% | 46.28% |
|  | *Mus musculus* | 56.63% | 38.48% |
|  | Swissprot | 62.42% | 54.07% |
|  | TrEMBL | 78.77% | 75.67% |
|  | nr | 72.06% | 66.24% |
|  | InterProScan | 90.94% | 87.15% |

We then generated comprehensive annotations for robust *de novo* gene discovery. Integrating *ab initio*, homology-based, and RNA-seq evidence-based gene prediction identified 19,842 and 23,502 protein-coding genes in *Te. occipitalis* and *Ta. portentosus*, respectively. Functional annotation assigned significant hits (E-value < 1e−6) to 15,707 (79.16%) and 17,913 (76.22%) genes, respectively, using eggNOG-mapper, InterProScan, or BLASTP searches against the *Caenorhabditis elegans*, *D. melanogaster*, *Homo sapiens*, and *Mus musculus* proteomes, the UniProt Swiss-Prot databases, and the NCBI non-redundant database (Cantalapiedra et al. 2021; Jones et al. 2014; Altschul et al. 1990; UniProt Consortium 2023; Sayers et al. 2021) (Table 1). Furthermore, RepeatModeler/RepeatMasker identified repetitive elements occupying 38.16% and 43.97% of the assemblies (Flynn et al. 2020; Chen 2004) (Supplemental Fig. S4).

### Identification of Species-Specific Gene Candidates

To investigate how gene properties change across evolutionary timescales, we first classified genes into evolutionary age categories and identified lineage-specific genes that may represent the youngest stage of gene evolution. Using genome assemblies and annotations, we constructed a *Te. occipitalis*-specific gene set for *de novo* gene discovery by profiling homology across 29 insect genomes from species with clearly defined phylogenetic relationships (Misof et al. 2014) (Fig. 2A; Supplemental Table S4). OrthoFinder clustered 1,470,750 proteins encoded by the 29 genomes (19,892 from *Te. occipitalis*) into 30,709 orthogroups (Emms and Kelly 2019), 10,670 of which contained at least one *Te. occipitalis* gene; 37,825 proteins (1,847 from *Te. occipitalis*) were not assigned to an orthogroup. Each *Te. occipitalis* protein, including those not assigned to an orthogroup, was then assigned to an evolutionary age category based on the deepest phylogenetic node at which an orthologue was detected, yielding nine age categories (Fig. 2B; Supplemental Table S5): Category 9 (root, 14,345 genes), Category 8 (719), Category 7 (181), Category 6 (83), Category 5 (450), Category 4 (1,448), Category 3 (317), Category 2 (217), and Category 1 (2,132 genes with no detectable orthologue in any other examined insect genome). Gene expression analysis showed that testis-expressed genes were abundant across categories, and approximately half of the genes in Category 1 did not exhibit biased expression in specific tissues (Fig. 2C). Category 1 genes therefore represent candidate lineage-specific genes, among which we hypothesized that genes newly originating from intergenic regions would occur.

**Fig. 2.**
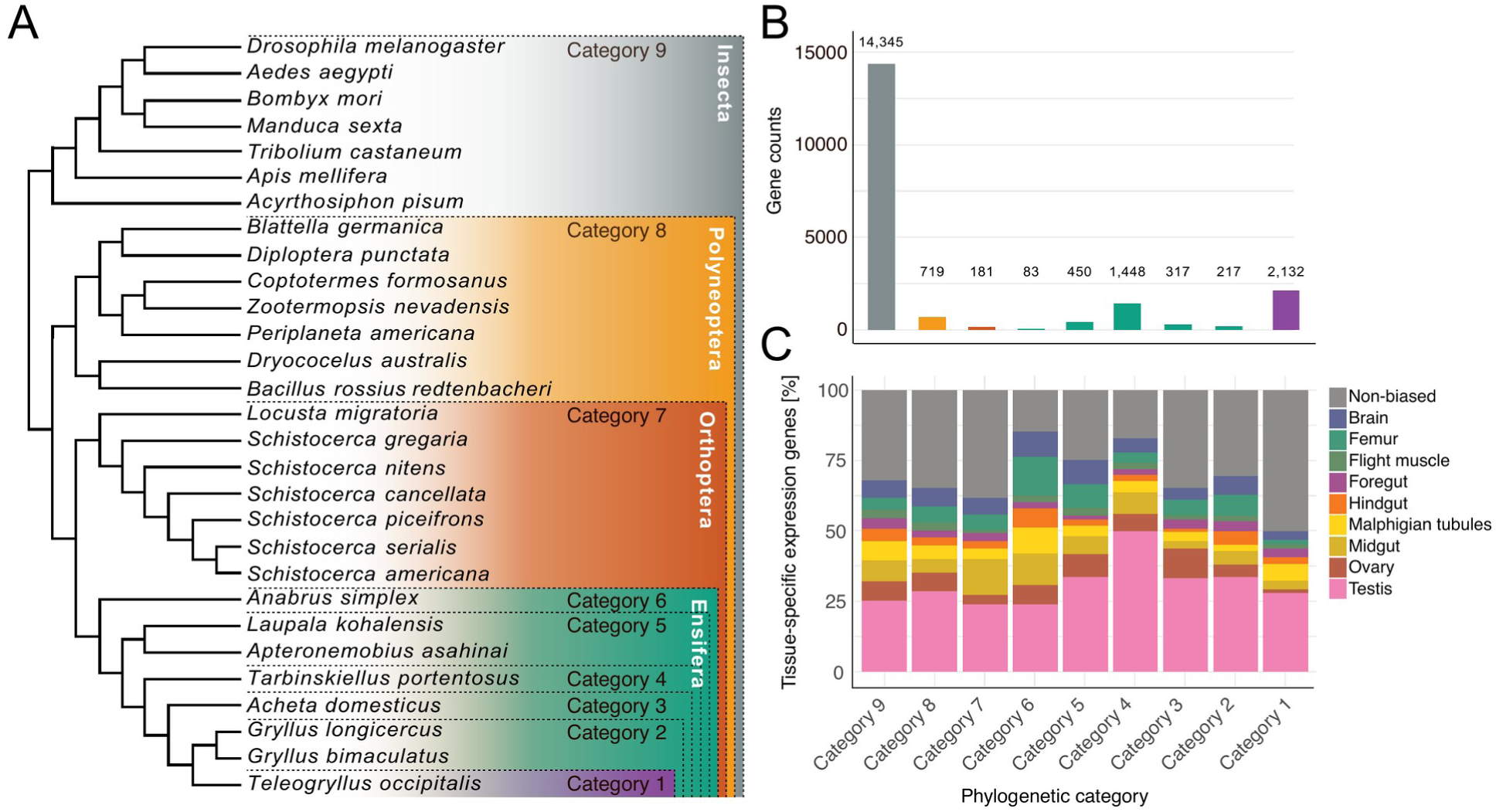
Evolutionary age classes of *Te. occipitalis* genes. (**A**) Phylogenetic framework used for gene-age inference (schematic; branch lengths not scaled). (**B**) Counts of *Te. occipitalis* genes assigned to Categories 2–9 (deepest node with an orthologue) and Category 1 (no orthologue in any other species). (**C**) Tissue-bias profile for each age category. Stacked bars show the proportion of genes with DESeq2-defined bias toward one of eight tissues.

### Identification and Characterization of *De Novo* Gene Candidates

To identify *de novo* genes in *Te. occipitalis*, we applied a series of strict filtering steps (Fig. 3A). From 2,132 Category-1 genes, we further excluded potential orthologs, xenologs, and both out- and in-paralogs (i.e., genes with intraspecific duplicates) using BLASTP v2.16.0 searches against the cricket genome and the NCBI-nr database, yielding 284 candidates. Among these, we retained 41 loci that (i) showed no detectable similarity to any protein in both NCBI-nr and cricket genomes, and (ii) had their best nucleotide matches in syntenic intergenic regions of closely related crickets. These candidates were distributed across the genome (5 on the X chromosome, 35 on autosomes, and 1 on an unplaced scaffold) (Table 2, Supplemental Table S6), had an average GC content of 70% and 62% hydrophobic amino acids (Supplemental Fig. S5).

**Fig. 3.**
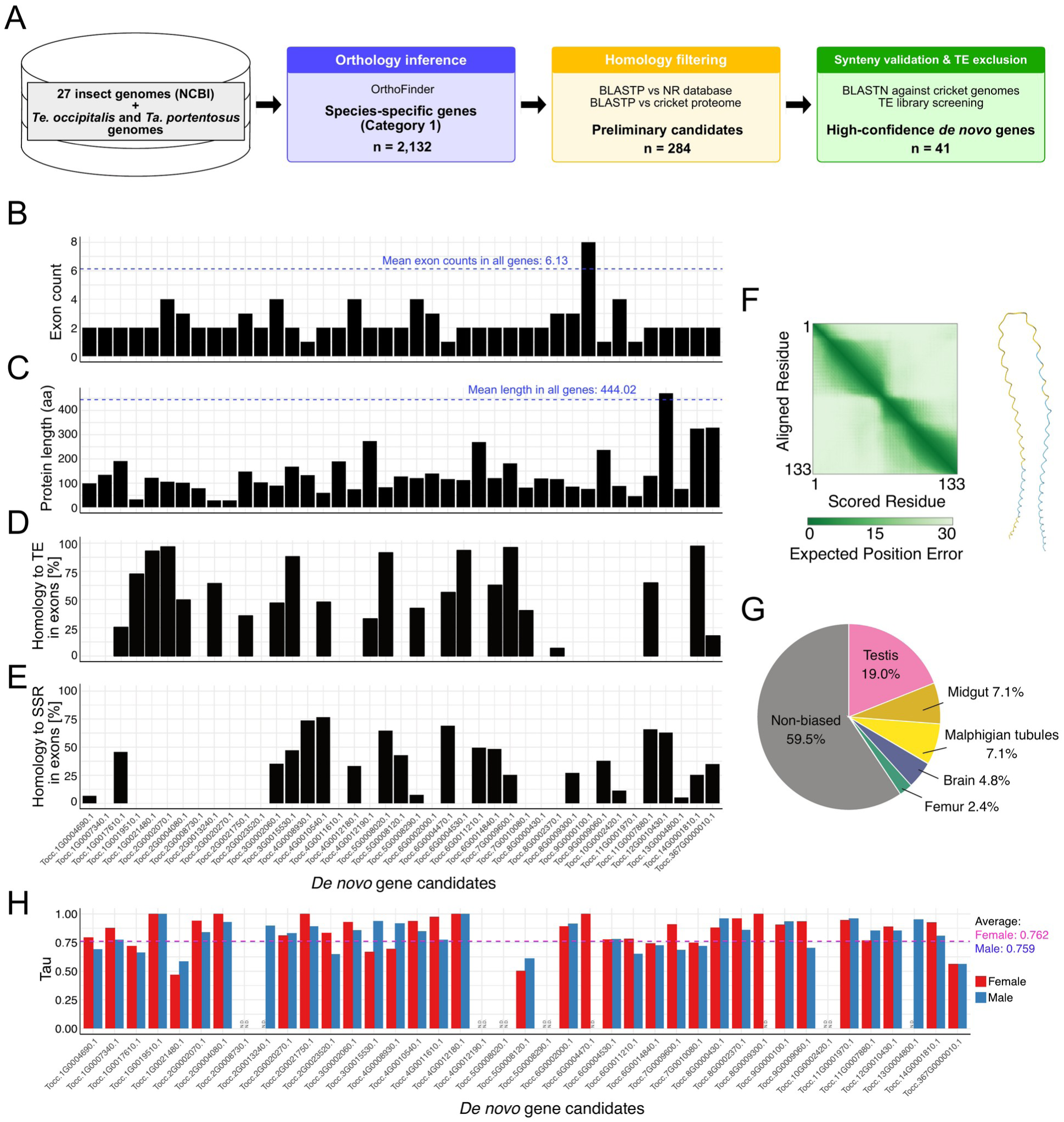
Identification and molecular signatures of *Te. occipitalis de novo* gene candidates. (**A**) Three-step screen for candidate genes. Step 1: OrthoFinder clustering across 29 insect genomes yielded 2,132 *Te. occipitalis*-specific proteins. Step 2: BLASTP removal of orthologs/paralogs. Step 3: synteny comparisons retained 41 final candidates. (**B**) Exon counts per candidate; blue dashed line = genome mean (6.13). (**C**) Protein lengths per candidate; blue dashed line = genome mean (444 aa). (**D**) Fraction of each candidate’s exon length homologous to TransposonPSI fragments; the purple wavy line marks 80% coverage. (**E**) Fraction of exon length occupied by *de novo* simple repeat sequences predicted with Krait2; purple line denotes the 80% benchmark. (**F**) Structural evidence for the protein-coding potential of candidate *Tocc.4G0008930.1*. Left, residue index versus aligned-residue confidence plot; depth of green shading denotes expected position error (Å). Right, AlphaFold3-predicted tertiary structure colored by pLDDT (blue = high confidence, red = low). (**G**) Tissue bias of candidates (DESeq2). Colors match the tissues in Fig. 2. (**H**) Tissue-specificity index (τ) per candidate for females (pink) and males (blue); dashed lines = genome-wide means.

**Table 2.**
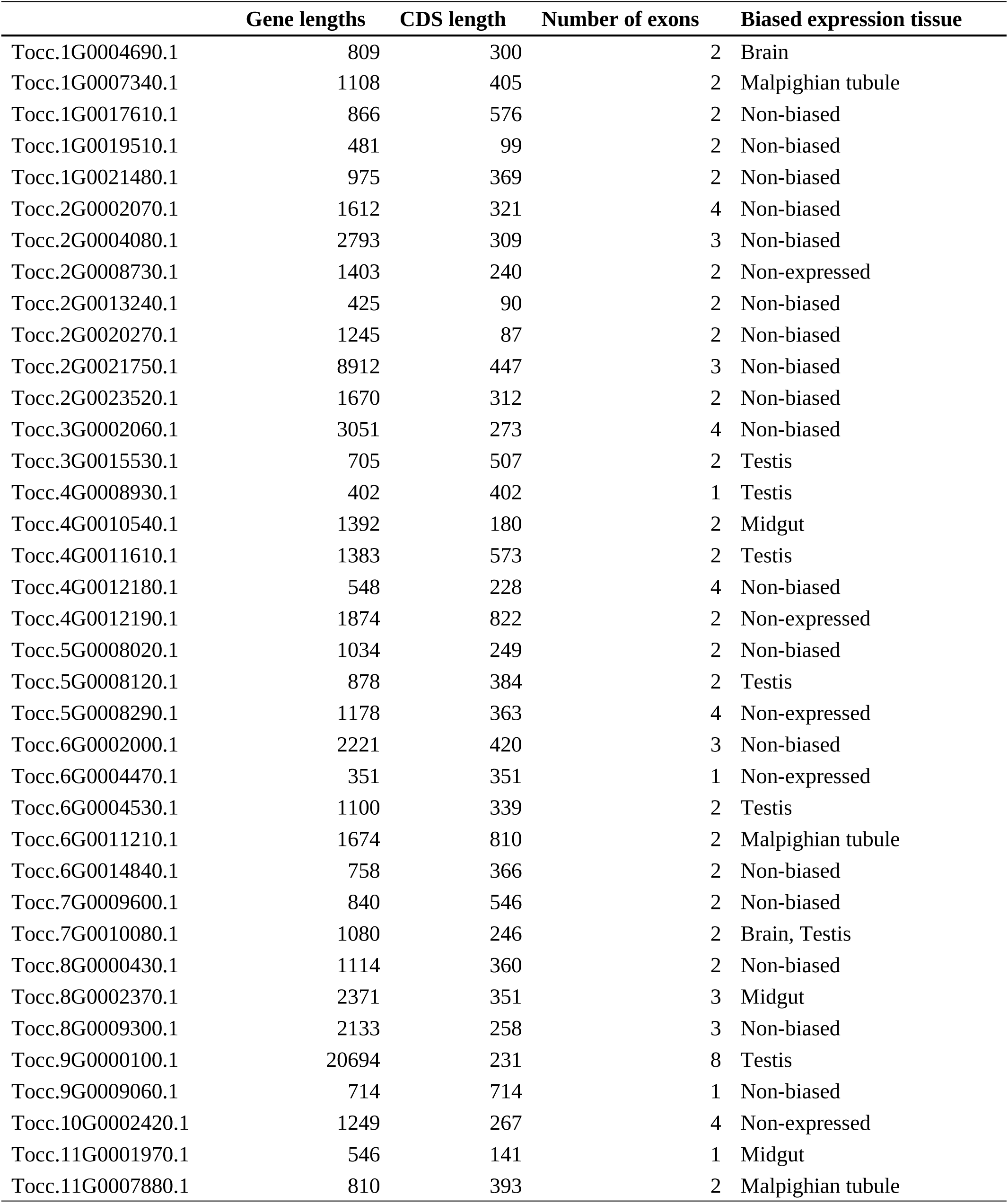
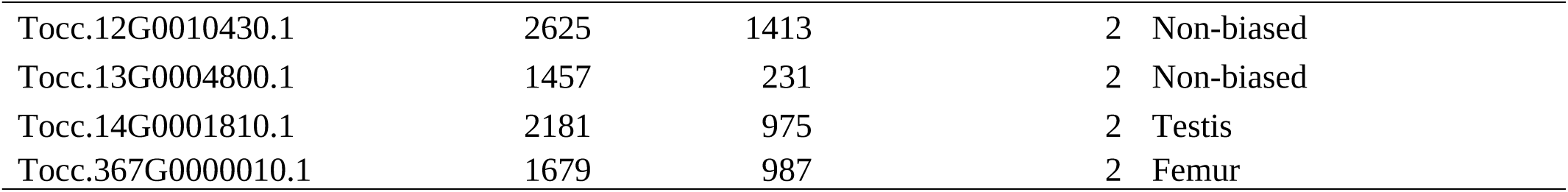
Characteristics of *Teleogryllus occipitalis de novo* gene candidates. Detailed statistics for each locus, including total gene length, coding-sequence (CDS) length, exon count, and the tissue in which expression is significantly biased (based on DESeq2 analysis). “Non-biased” indicates no significant tissue preference.

These 41 candidates generally had fewer exons than the genomic average (Fig. 3B) and encoded shorter predicted proteins (Fig. 3C). Syntenic alignments of each candidate plus 300 bp of flanking sequence against syntenic regions of closely related crickets revealed that 14 genes (*Tocc.3G0015530.1*, *Tocc.4G0008930.1*, *Tocc.4G0012180.1*, *Tocc.4G0012190.1*, *Tocc.5G0008290.1*, *Tocc.6G0004530.1*, *Tocc.6G0011210.1*, *Tocc.7G0009600.1*, *Tocc.9G0009060.1*, *Tocc.10G0002420.1*, *Tocc.11G0007880.1*, *Tocc.12G0010430.1*, *Tocc.13G0004800.1*, *Tocc.367G0000010.1*) shared conserved flanks but contained unique start/stop codons or exon sequences specific to *Te. occipitalis* (Supplemental Fig. S5).

Furthermore, to assess the contribution of TE-derived sequences and to exclude misannotated TE genes, we compared each exon with TransposonPSI (https://transposonpsi.sourceforge.net/, last accessed July 24, 2025) calls. BLASTN analyses detected TE fragments in 22 genes, seven of which had >80% of their exon length covered by TE fragments (Fig. 3D; Supplemental Fig. S5). Full-length searches against the TransposonPSI consensus library and the Dfam database (E ≤ 1 × 10^−6^) revealed significant matches for four genes (*Tocc.1G0004690.1*, *Tocc.1G0007340.1*, *Tocc.1G0017610.1*, *Tocc.1G0019510.1*) only in the Dfam search, and none aligned with TransposonPSI consensus sequences. No recognizable protein domains were detected by Pfam scanning, consistent with these loci lacking detectable signatures of known protein-coding genes, including annotated protein-coding sequences within TEs.

We then asked which sequence features might contribute to early coding potential and protein structure. Pairwise syntenic alignments revealed newly elongated simple-repeat segments absent from the orthologous non-coding loci of other crickets (Supplemental Fig. S5). Consistently, systematic screening with Krait2 identified simple sequence repeats within the exons of 22 candidates (Du et al. 2025) (Fig. 3E; Supplemental Fig. S5). Predicted protein structures were dominated by short α-helices (Fig. 3F; Supplemental Fig. S6), suggesting that repeat expansion could provide both extended open reading frames and foldable structural elements during *de novo* gene birth. Together, TE fragments and simple repeats coincide with segments lacking homology in syntenic alignments to related genomes, supporting their recent origin and highlighting repetitive sequence material as a substrate for protein-coding gene emergence.

Finally, we examined transcriptional properties of these candidates. Expression profiling revealed pronounced tissue bias for a subset of loci. Eight genes were testis-specific, three midgut-specific, three Malpighian-tubule-specific, two brain-specific, and one femur-specific (Fig. 3G), whereas the others showed no significant bias toward a single tissue. Overall, all candidates except for five genes were expressed in at least one tissue, and 30 showed a tissue specificity index (τ) above the genome-wide average in at least one sex (Yanai et al. 2005) (Fig. 3H). Overall expression levels tended to be lower than those of typical annotated genes (Supplemental Table S5), consistent with these loci representing recently emerged, weakly expressed genes (Schlötterer 2015).

### Transposable Element Fragments in *De Novo* Gene Candidates

To assess whether TE fragments within *de novo* gene candidates were unique to these loci or widespread across insect genes, we queried TE fragments from *Te. occipitalis de novo* gene candidates against other insect genomes. BLASTN searches detected homologous regions in gene sequences across multiple species (Supplemental Table S7), spanning diverse TE classes and showing no enrichment for any specific class. Homologs of TE fragments located in *Te. occipitalis de novo* gene introns were frequently detected in genomes of closely related crickets, whereas homologs of exonic TE fragments were also found in the exons of more distantly related species (Supplemental Table S7). Across insect species, genes containing TE fragments often had orthologues in other species that lacked the corresponding TE-derived regions, indicating that these fragments were inserted independently in different lineages (Supplemental Tables S7–S8). Similar TE fragments were inserted independently into different genes, including both duplicated and single-copy genes, across taxa, suggesting that their insertion is not restricted to specific gene classes (Supplemental Tables S7–S8). Although orthologues of these genes often retained traces of related repetitive sequences (Supplemental Table S9), the TE-derived regions lacked conserved motifs across taxa, suggesting that they do not encode conserved protein functions.

### The Sequence Properties of *De Novo G*enes in an Evolutionary Context

To test whether features characteristic of *de novo* gene candidates were enriched in younger genes and attenuated with age, we classified *Te. occipitalis* genes into six evolutionary age classes (see Methods) and examined gene structure and expression specificity for each category. From younger to older genes, the number of exons per gene increased (Spearman’s ρ = –0.505, P < 0.01; Jonckheere–Terpstra P < 0.01) (Fig. 4A), as did exon length (Spearman’s ρ = –0.440, P < 0.01; Jonckheere–Terpstra P < 0.01) and intron length (Spearman’s ρ = –0.374, P < 0.01; Jonckheere– Terpstra P < 0.01) (Fig. 4B, C). In contrast, τ decreased with evolutionary age (Spearman’s ρ = 0.432 (male), 0.359 (female), both P < 0.01; Jonckheere–Terpstra both P < 0.01) (Fig. 4D), showing that younger genes exhibit more restricted expression profiles. These monotonic trends remained significant after excluding the *de novo* category (all Jonckheere–Terpstra P < 0.01).

**Fig. 4.**
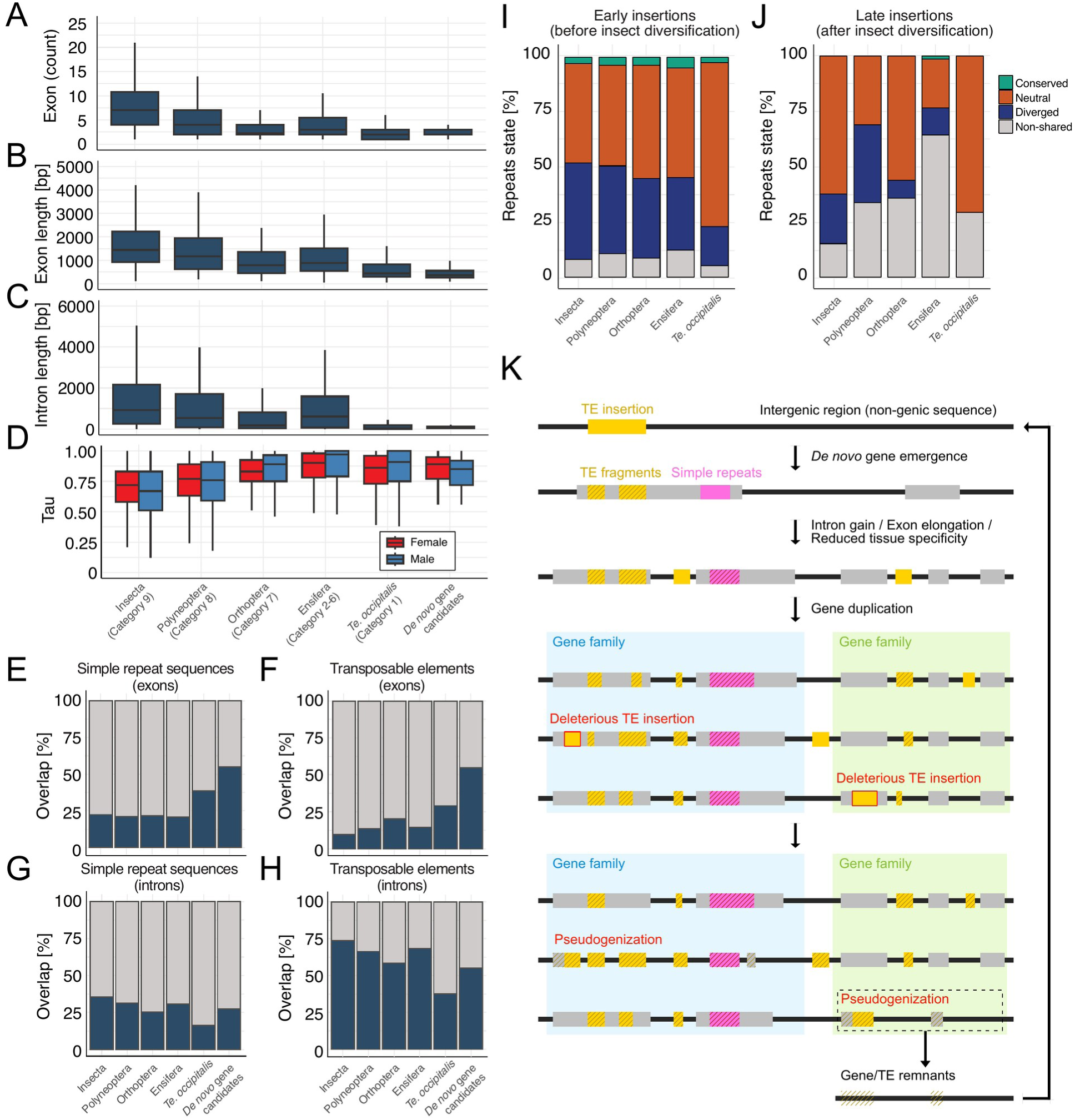
Age-class dependence of genomic and expression features in *Te. occipitalis*, and cross-species conservation of repeat tracts. (**A**) Relationship between exon counts and expected gene age. (**B**) Relationship between exon length and expected gene age. (**C**) Relationship between intron length and expected gene age. (**D**) Relationship between tissue specificity index (τ) in each sex and expected gene age. (**E**)–(**H**) Proportion of “overlap-rich” orthogroups (>50% of members containing the indicated repeat) in each age class: (**E**) simple repeat sequences in exons, (**F**) TEs in exons, (**G**) simple repeat sequences in introns, and (**H**) TEs in introns. (**I**) Conservation status of repetitive elements inferred to have been inserted during early stages of gene evolution (early insertions; acquired in the ancestral lineage leading to Insecta), evaluated across insect species grouped by clade. (**J**) Conservation status of repetitive elements inferred to have been inserted after the diversification of insect taxa (late insertions), evaluated across the same clades. For panels (**I**) and (**J**), repetitive elements were classified as Conserved, Neutral, Diverged, or Non-shared based on relative alignment identity compared with background regions. (**K**) Schematic model of gene evolution from intergenic sequences to evolutionarily stable genes. During the evolution, repetitive sequence material—such as TE-derived fragments and simple sequence repeats—can provide readily mutable sequence substrate that contributes to early ORF formation and nascent gene architecture. Newly emerged de novo genes may subsequently undergo progressive maturation over evolutionary time, including the acquisition of introns and additional structural features. Once stabilized, some loci may expand through gene duplication to form gene families. During this phase, additional TE insertions can occur; many are expected to be deleterious when they disrupt coding regions, resulting in gene loss or pseudogenization. Surviving duplicates are retained as gene families ultimately derived from a *de novo* gene ancestor. Remnants of pseudogenes and fragmented repeats left in intergenic regions can in turn serve as sequence material for subsequent rounds of *de novo* gene birth, linking gene gain and loss in a continuing cycle.

Next, we asked whether the prevalence of repetitive sequence fragments within genes changes across evolutionary age classes. Here, we used orthogroups inferred by OrthoFinder as operational units of gene families across insects. From younger to older, gene families in which >50% of the members contained exonic simple sequence repeats decreased (Cochran–Armitage χ^2^_1_ = 118.8, P < 0.01; OR = 1.14, 95% CI 1.12–1.17) (Fig. 4E, Supplemental Table S10), as did families with exonic TE segments (χ^2^_1_ = 420.5, P < 0.01; OR = 1.34, 95% CI 1.31–1.38) (Fig. 4F, Supplemental Table S10). Conversely, the proportion of families with intronic simple sequence repeats increased (χ^2^_1_ = 208.7, P < 0.01; OR = 0.83, 95% CI 0.81–0.86) (Fig. 4G, Supplemental Table S10), and this increase was even more pronounced for intronic TE fragments (χ^2^_1_ = 653.8, P < 0.01; OR = 0.75, 95% CI 0.73–0.77) (Fig. 4H, Supplemental Table S10). These trends were unchanged after excluding the *de novo* category (all Cochran–Armitage χ^2^_1_ P < 0.01).

To determine whether these trends reflected changes in repeat abundance within genes, we next quantified repeat coverage per gene across age classes (Supplemental Fig. S7). Younger genes showed a higher fraction of exon length occupied by simple sequence repeats (Spearman’s ρ = 0.120, P < 0.01; Jonckheere–Terpstra P < 0.01), whereas the fraction of intron length occupied by simple sequence repeats showed no significant monotonic trend (Spearman’s ρ = –0.018, P = 0.18). For TE fragments, both exonic (Spearman’s ρ = 0.167, P < 0.01; Jonckheere–Terpstra P < 0.01) and intronic (Spearman’s ρ = 0.056, P < 0.01; Jonckheere–Terpstra P < 0.01) coverage fractions declined toward older age classes. Thus, while the number of intron repeat-bearing genes increases with age, the per-gene share of intronic sequence covered by repeats does not increase in parallel, indicating a shift toward broader incidence across genes rather than expanding in copy or length within individual genes.

Finally, we examined whether the evolutionary persistence of repeat elements differed depending on their inferred timing of insertion. We first defined a set of 9,898 gene families inferred to have been maintained since the insect last common ancestor. This set included the *Te. occipitalis* genes assigned to Category 9 (deeply conserved genes shared across insects), in which we identified repetitive elements and traced their histories across insects. For each repetitive element detected within *Te. occipitalis* Category 9 genes, we inferred insertion timing from presence–absence patterns at orthologous loci in other insect genomes. Insertions were classified as early insertions if the element was inferred to have been acquired in the ancestral lineage leading to Insecta, and as late insertions if it was inferred to have occurred after diversification of insect taxa. Using this framework, early insertions were detected in 2,358 gene families, whereas late insertions were detected in only 11 families. Within the 2,358 gene families with early insertions, repetitive elements were identified in 91,499 genes (Supplemental Table S11). This accounts for ~90% of alignable genes and ~12% of the entire genome across insects. We then classified each *Te. occipitalis*-derived repeat element by its cross-species state (Conserved, Neutral, Diverged, or Non-shared). State frequencies were summarized across five phylogenetic nodes; values at each node represent how often the *Te. occipitalis*-detected elements are observed in that state among orthologous genes from the corresponding clade within the same orthogroups (Fig. 4I, J). Among early insertions, a total of 3,515 repeat elements detected in *Te. occipitalis* were classified as “Conserved” when summarized across phylogenetic nodes (in Insecta: 804, in Polyneoptera: 598, in Orthoptera: 1072, in Ensifera: 949, and in *Te. occipitalis*: 92) (Fig. 4I, Supplemental Table S11), while only two late-inserted repetitive elements were classified as “Conserved” (in Ensifera: 2) (Fig. 4J, Supplemental Table S11). Across insect taxa, TE fragments acquired by genes prior to the insect radiation showed no bias in insertion positions within genes (Supplemental Fig. S8). Neutrally conserved TE fragments acquired by genes after the insect radiation originated from seven independent evolutionary events, six of which involved insertions at the 5’ or 3’ ends of gene sequences (Supplemental Table S12). These analyses indicate that repetitive sequence fragments located within exons of insect-conserved genes tend to persist over evolutionary time, even though the underlying nucleotide or amino acid sequences themselves are poorly conserved.

## DISCUSSION

A central unresolved question in genome evolution is whether and how *de novo* genes transition from intergenic origins to stable components of long-lived gene repertoires (Carvunis et al. 2012; Neme and Tautz 2016; Van Oss and Carvunis 2019). Addressing this requires comparative analyses across evolutionary timescales anchored by conservative identification of *de novo* genes. Here, using such a comparative framework in insects, we provide empirical support for a view of gene evolution as a continuum from intergenic regions to mature genes and propose an evolutionary model centered on the sequence material used in molecular tinkering (Fig. 4K). In insects, genes can be constructed from intergenic regions through the incorporation of abundant and mutable sequence materials, such as TE fragments. However, once a gene becomes stabilized, purifying selection likely shifts to restrict further incorporation of such sequences into exons.

Based on chromosome-scale genomes and transcriptome resources for crickets, we identified 41 *Te. occipitalis* lineage-specific gene candidates for which non-genic homologous sequences were detected in other cricket genomes. Although this number is likely an underestimate due to strict filtering and limited transcriptomic sampling, and although alternative scenarios such as extreme sequence divergence cannot be fully excluded, the number of these candidates falls within the range reported in yeast and *D. melanogaster* (Blevins et al. 2021; Grandchamp et al. 2023). We therefore use them as a conservative set to examine the earliest stages of gene evolution in insects.

Using evolutionary age as a comparative framework, we found that multiple gene properties vary across insect genes, despite potential biases from gene loss or extreme divergence in outgroups (Weisman et al. 2020). These patterns indicate a gradual increase in gene length and structural complexity with increasing evolutionary age (Fig. 4A–C). Within this continuum, *de novo* gene candidates from *Te. occipitalis* fall within the spectrum of these youngest classes, consistent with a recent evolutionary origin. These results support a model of gene evolution based on a continuum from intergenic regions to mature genes (Palmieri et al. 2014; Knowles and McLysaght 2009). Consistent with this, expression specificity generally increases toward younger age classes, and most *de novo* gene candidates in *Te. occipitalis* exhibit restricted expression patterns (Fig. 3H), although only a subset show significant bias toward the testis or other single tissues (Fig. 3G). In *Drosophila*, a hypothesis has been proposed that *de novo* genes evolve under natural selection due to biased expression in the testes (Levine et al. 2006). However, it remains unclear at which evolutionary stage these candidate genes are located in this scenario. The evolutionary fate of these candidates therefore remains unresolved, including whether they will be eliminated, transition toward testis-biased expression, or become integrated into existing gene networks through alternative evolutionary routes.

The *de novo* gene candidates we identify here contain TE fragments and simple sequence repeats, and genes with such sequences in their exons are abundant in evolutionarily young classes (Fig. 4E, F). By distinguishing repetitive sequence insertions within exons traceable to the last common ancestor (LCA) from lineage-specific insertions, we found that LCA-derived repetitive sequences exhibit higher sequence conservation than lineage-specific insertions (Fig. 4I, J). Among insects, approximately 24% of LCA-derived gene families contain repetitive sequences. Their persistence over long evolutionary timescales despite the absence of conserved motifs suggests that they contribute as sequence material rather than as conserved functional elements. Conversely, the scarcity of recent insertions into exons is consistent with purifying selection against insertions into coding regions, which are generally deleterious (Brennecke et al. 2007; Cordaux and Batzer 2009; Lisch 2013), indicating that the evolutionary impact of repetitive sequence insertions is strongly dependent on the stage of gene evolution.

In contrast to exonic regions, repetitive sequences overlapping introns were less frequent in evolutionarily younger genes (Fig. 4G, H). Consistently, intronic TE fragments predominantly reflected lineage-specific insertions (Supplemental Table S11). This pattern is consistent with recurrent insertion of active TEs into introns during gene evolution, a common feature of eukaryotic genomes (Roy 2004; Mikina et al. 2024), suggesting intronic TE accumulation reflects continuous lineage-specific activity of TEs across evolutionary time independent of gene age.

The predicted protein structures of *de novo* gene candidates are dominated by α-helical conformations and disordered regions, a structural signature commonly observed in young genes originating from intergenic regions (Palmieri et al. 2014). Accidentally translated disordered proteins are generally expected to be selectively neutral or deleterious and predicted to be eliminated near the time of *de novo* gene birth (Rajon and Masel 2011). Nevertheless, functionally important α-helical cores are exemplified by *goddard*, a *Drosophila* gene originating from an ancestral intron and conserved across the genus (Gubala et al. 2017; Lange et al. 2021). Recent studies further suggest that the emergence of foldable *de novo* genes is shaped by genomic background and biochemical properties of DNA sequences (Grandchamp et al. 2025), including the propensity to form α-helical transmembrane domains that could reduce unintended intermolecular interactions (Vakirlis et al. 2020; Vakirlis and Fuqua 2025). Consistent with this view, *de novo* gene candidates in *Te. occipitalis* are GC-rich and enriched in hydrophobic amino acids. These candidates may originate through the extension of simple repeats, which rapidly change in copy number and can simultaneously provide extended open reading frames and foldable protein structures (Buschiazzo and Gemmell 2006).

Together, our results support a continuum model in which intergenic loci can give rise to *de novo*genes and, in some cases, seed long-lived gene repertoires through time. The observation that ~24% of LCA-derived gene families across insects retain repetitive sequences argues against the idea that such insertions are inherently lethal and is unlikely to be explained solely by survivorship bias (Lee 2022). The contrasting evolutionary dynamics of repetitive sequences between exonic and intronic regions further suggest that stages of gene evolution can be operationally distinguished by the changing receptivity of coding sequences to repetitive insertions. Further analyses integrating functional characterization and comparative genomics will be required to clarify the evolutionary fate of *de novo* gene candidates in *Te. occipitalis* and to test how intergenic region–derived genes are retained or eliminated over evolutionary time.

## METHODS

### Biological Samples, DNA and RNA Isolation

A schematic diagram of the process from DNA extraction to genome assembly is shown in Supplemental Fig. S9.

For constructing long-read and short-read sequencing libraries, high molecular weight DNA was extracted with the NucleoBond HMW kit (MACHEREY-NAGEL, GmbH & Co. KG, Düren, Germany) from the head and hind legs of a single adult female *Te. occipitalis* (lab-reared at Waseda University) and the hind legs of a single adult female *Ta. portentosus* (collected at Kampong Thom, Cambodia). The lysis step was extended to 3 h, the centrifugation step was omitted, and DNA pellets were transferred by pipette during the precipitation/wash steps. For Omni-C scaffolding, DNA from the head and hind legs of an additional adult single female *Te. occipitalis* and the hind legs of an additional adult single female *Ta. portentosus* was isolated with the Dovetail™ Omni-C™ kit (Dovetail Genomics, California, USA).

For gene prediction, RNA of *Te. occipitalis* was isolated from the heads, thoraxes, and abdomens of individual adult males and females, and whole-body larvae. Also, RNA of *Ta. portentosus* was isolated from the head and fat bodies of an adult male. For the differential expression analysis, RNA was extracted from the brain, femur, flight muscle, foregut, gonad, hindgut, Malpighian tubules, and midgut of eight males and eight females of *Te. occipitalis*. All RNA was extracted with TRIzol reagent (Invitrogen; Thermo Fisher Scientific, Inc., Waltham, MA, USA) following homogenization, chloroform phase separation, isopropanol precipitation, ethanol wash, and buffer elution. At least four biological replicates were obtained for each tissue and sex.

### Library Construction and Sequencing

For long-read sequencing of *Ta. portentosus*, genomic DNA was size-selected with the Short Read Eliminator XL kit (Pacific Biosciences of California, Inc., CA, USA) to remove fragments < 10 kbp. A 2.5 µg aliquot of the size-selected DNA was end-repaired with the NEBNext FFPE DNA Repair Mix and NEBNext Ultra II End Repair/dA-Tailing Module (New England Biolabs, MA, USA), then ligated to sequencing adapters from the SQK-LSK109 kit (Oxford Nanopore Technologies KK, Tokyo, Japan) using the NEBNext Quick Ligation Module. DNA was purified after each step with AMPure XP beads (Beckman Coulter, CA, USA). The finished library was loaded on an R9.4.1 Flowcell and run on a PromethION 24 (Oxford Nanopore Technologies PLC, Oxford, UK). Raw signals were base-called with Guppy v4.3.4 in high-accuracy mode. In addition, an Omni-C library was prepared using the Dovetail™ Omni-C™ kit (Cantata Bio LLC, California, USA), sequenced on an Illumina NovaSeq 6000, and utilized for scaffolding. Bionano optical mapping data (Bionano Genomics, CA, USA) were also generated for scaffolding, and Illumina paired-end reads from a NovaSeq 6000 were obtained for polishing the draft assembly. Furthermore, RNA-seq for gene prediction was performed on a HiSeq 2500. All auxiliary sequencing services were provided by Rhelixa, Inc. (Tokyo, Japan).

For long-read sequencing of *Te. occipitalis*, we followed a similar workflow. Genomic DNA was size-selected with the Short Read Eliminator XL kit, and 2.5 µg of the recovered DNA underwent end-repair (NEBNext FFPE DNA Repair Mix + Ultra II End Repair/dA-Tailing), adapter ligation with the SQK-LSK112 kit (Oxford Nanopore Technologies KK), and AMPure XP purification at every step. The library was loaded on an R9.4.1 Flowcell and sequenced on a PromethION 24. Raw signals were base-called with Guppy v5.0.11 in high-accuracy mode. PacBio continuous long reads (CLR) were produced on a Sequel II system, Omni-C reads were generated with a NovaSeq 6000, and Illumina paired-end reads (HiSeq 2500 for polishing; NovaSeq 6000 for X-chromosome identification) were also obtained. RNA-seq for gene prediction and differential expression analysis was performed on a HiSeq 2500. All auxiliary sequencing services were provided by Rhelixa, Inc. (Tokyo, Japan).

### Genome Assembly

The genome size of crickets was estimated using GenomeScope v2.3.0, with k-mers counted using Jellyfish v2.3.0 (Marçais and Kingsford 2011; Ranallo-Benavidez et al. 2020). For genome assembly of *Ta. portentosus*, long reads generated from the PromethION sequencer were assembled using de novo assemblers, Flye v2.8.1, NECAT v0.0.1, Miniasm v0.3, Raven v1.5.0, SMARTdenovo v3.29 (Liu et al. 2021), Shasta v0.7.0, and wtdbg2 v2.5. Additionally, MaSuRCA v3.4.2 was employed for hybrid de novo assembly (Kolmogorov et al. 2019; Chen et al. 2021; Li 2016; Vaser and Šikić 2021; Shafin et al. 2020; Ruan and Li 2019; Zimin et al. 2013), integrating short Illumina paired-end reads with the long PromethION reads. The performance of each assembler was evaluated based on assembly continuity by QUAST (Gurevich et al. 2013). Additionally, the gene completeness of the assembly is also assessed by BUSCO using the Arthropoda database. The genome assemblies generated by Flye and wtdbg2, which demonstrated superior performance among long-read assemblers, were polished with short reads and used to compare their performance to the assembly by MaSuRCA.

For genome assembly of *Te. occipitalis*, a similar approach was taken. Long reads generated from the PromethION sequencer and the PacBio Sequel II system (CLR) were assembled together using Flye v2.8.1 and wtdbg2 v2.5. MaSuRCA v3.4.2 was also employed for hybrid *de novo* assembly using both long-read datasets together with Illumina paired-end reads. The assemblies were evaluated by QUAST and BUSCO.

After selecting the genome assemblies in both species, the resulting contigs were subjected to error correction (polishing) using POLCA v1.4.1 with Illumina paired-end data (Zimin and Salzberg 2020). Contaminants were also removed based on unexpected coverage, GC content, or homology with bacterial and other contaminant sequences using BlobtoolKit v1.1.1 (Challis et al. 2020). The sequence coverage was obtained by mapping the Illumina reads with BWA. Similarity analysis was performed with BLASTN v2.12.0 using NCBI NT database v5 with the options “-task megablast culling_limit 10 -evalue 1e-25 -outfmt ’6 qseqid staxids bitscore std sscinames sskingdoms stitle’”. Allelic haplotigs were removed by Purge Haplotigs v1.1.2, using the long-read mapping data by bwa v0.7.17-r1188 (Li 2013; Roach et al. 2018). Followed by additional rounds of polishing using Hapo-G v1.3.6 for both genome assemblies (Aury and Istace 2021), BioNano optical mapping data were also integrated to further improve the assembly for *Ta. portentosus*.

The assembled contigs of both *Ta. portentosus* and *Te. occipitalis* were corrected for misjoins, ordered, oriented, and anchored into candidate chromosome-scale assemblies using Omni-C™ data with Juicer v1.9.9 and 3D-DNA v180419 (Dudchenko et al. 2017; Durand et al. 2016b). Finally, the candidate assemblies were reviewed and refined for quality control using Juicebox v1.11.08 for quality control and interactive corrections (Durand et al. 2016a). The contact maps of two crickets were visualized using Juicebox as the interactive signals between each pair of 2 bins. Additionally, a Circos plot, illustrating the distribution of genomic elements, was generated using Circos v0.69-9 (Krzywinski et al. 2009).

### Repeat Annotation and Gene Prediction

A schematic diagram of gene prediction is shown in Supplemental Fig. S10.

For *de novo* repeat prediction, RepeatModeler v2.0.5 was initially employed to construct a candidate database of repetitive elements. The identified repetitive sequences were then soft-masked using RepeatMasker v4.1.5. Structural annotation of protein-coding genes was performed on the soft-masked genome utilizing three approaches: *ab initio*, homology-based, and RNA-seq evidence-based prediction. Initially, to minimize noise from the RNA-seq reads for gene prediction, a *de novo* transcriptome assembly was carried out using Trinity v2.15.1 to generate contigs (Grabherr et al. 2011). The original RNA-seq reads were then mapped back to these contigs using HISAT2 v2.2.1 with default parameters (Kim et al. 2019). This step filtered out reads that did not map correctly in the proper orientation as paired-end reads. The filtered reads were subsequently utilized for gene prediction with BRAKER3 (Gabriel et al. 2024), GeMoMa (Keilwagen et al. 2019), and StringTie2 (Kovaka et al. 2019). The *ab initio* prediction of each cricket was carried out using BRAKER v3.0.2, which incorporated both protein data from OrthoDB 11 arthropoda dataset (https://www.orthodb.org/, last accessed August 8, 2024) and the filtered RNA-seq alignments. For the homology-based prediction, GeMoMa v1.9.0 used gene sets from *Apis mellifera*, *D. melanogaster*, and *Tribolium castaneum*; for *Ta. portentosus*, the *Te. occipitalis* gene set was added. RNA-seq-based prediction was conducted using StringTie2 v2.2.1. Predictions generated by BRAKER3 served as the reference backbone; gene models from GeMoMa v1.9.0 and StringTie2 v2.2.1 were then merged and deduplicated against this backbone with GffCompare v0.12.6, producing the final consensus annotation (Pertea and Pertea 2020). EggNOG-mapper (http://eggnog-mapper.embl.de/, last accessed August 8, 2024), InterProScan, and BLASTP v2.16.0 (E-value < 1 × 10^−6^) searches against proteomes of *C. elegans*, *D. melanogaster*, *H. sapiens*, *M. musculus*, UniProt Swiss-Prot, and the NCBI-nr database (downloaded on November 28, 2021) provided gene functional annotation. To assess the completeness of the gene structure annotation, BUSCO v5 (Manni et al. 2021) with the arthropoda database was utilized.

### X-chromosome Identification of *Te. occipitalis*

In the XO sex determination system, the X chromosome in males should have half the read depth of autosomes. An adult male was sequenced on an Illumina platform to ~65× genomic coverage. Reads were aligned to the *Te. occipitalis* assembly with BWA (default parameters). The SAM file was converted to BAM with SAMtools v1.9, and Mosdepth v0.3.8 computed windowed coverage (-t 10 --no-per-base --fast-mode --by 500000) (Pedersen and Quinlan 2018; Li et al. 2009).

### Synteny Analysis

Chromosome-scale assemblies for *Ta. portentosus*, *Te. occipitalis*, and *A. domesticus* from a previous study were compared pairwise (Dossey et al. 2023). Predicted proteins underwent all-versus-all BLASTP v2.16.0 searches (-evalue 1e-10 -outfmt 6). Gene collinearity was detected with MCScanX v1.1 (-b 2 -m 25 -g -1 -s 10) and visualized in SynVisio (https://synvisio.github.io) (Wang et al. 2012). For each comparison, MCScanX block sizes (gene counts per block) were tabulated separately for autosomes and the X chromosome and evaluated with Pearson’s χ^2^ tests.

### Identification of Species-Specific Gene Candidates

Protein-coding genes from *Te. occipitalis*, *Ta. portentosus*, and 27 additional insects (from NCBI; Supplemental Table S4) were clustered homologous amino acid sequences by OrthoFinder v2.5.5 (default settings). Using the molecular phylogenetic tree reconstructed by OrthoFinder, the species sets were divided into nine groups based on their divergence from *Te. occipitalis*. The genes specific to each species set were classified into the corresponding category (Categories 1–9).

### Transcriptome Statistics

Raw RNA-seq reads were trimmed and filtered with fastp v0.23.2 (Chen et al. 2018). Transcript abundances were then quantified as TPM (transcripts per million) with Salmon v1.10.2 under default settings (Patro et al. 2017). Log-transformed TPMs for all genes were analyzed in R v4.2.3. Genes expressed with tissue-bias were detected with DESeq2 v1.44 by contrasting each tissue against the pooled set of the other seven tissues (one-versus-all design) (Love et al. 2014). Genes with fewer than 10 counts were excluded, and significance was set at an adjusted P < 0.05 and |log_2_ fold change| > 1.

### Identification of *De Novo* Gene Candidates

A composite BLASTP database was assembled from (i) all predicted protein coding genes of the cricket genomes analysed here and (ii) the NCBI-nr collection. Each Category-1 gene was queried against this database (E ≤ 1 × 10^−6^). Hits to genes in the non-*Te. occipitalis* genome were scored as orthologs/out-paralogs; hits to additional genes in the *Te. occipitalis* genome were scored as in-paralogs. All sequences with either type of hit were discarded, leaving initial targets with no detectable similarity to any known protein-coding gene.

For each target, the coding sequence plus its 5′ and 3′ intergenic segments (nearest gene boundary to boundary) were queried (BLASTN, E ≤ 1 × 10^−6^) against every cricket genome (Supplemental Table S4). A gene was retained only if (i) the best coding-sequence alignment mapped to an intergenic region of the sister genome and (ii) both flanking segments aligned immediately upstream and downstream of that locus, preserving collinearity. Loci passing both tests were designated final *de novo* gene candidates (n = 41).

The predicted protein products of all 41 loci were subsequently modelled with AlphaFold 3 (DeepMind, default settings) to obtain putative tertiary structures (Abramson et al. 2024).

### Alignment *De Novo* Gene Candidates with Syntenic Regions

For every final candidate, the full gene plus 300-bp flanks and each exon’s protein sequence were aligned to their orthologous regions. Nucleotide alignments used MAFFT v7 (--auto) (Katoh and Standley 2013). Syntenic regions in other crickets were six-frame translated, considering strand and open reading frame shifts. The longest contiguous match for each exon was then selected, permitting different optimal frames among exons. These alignments verified (i) sequence continuity and frame integrity in *Te. occipitalis* and (ii) persistence of homologous yet non-coding DNA in other crickets.

### Detection of Repetitive Sequence Overlap

TE signatures were predicted with TransposonPSI. Simple sequence repeats were identified with Krait2 v2.0.3. For the TE homology search, all TransposonPSI calls were compiled into a BLASTN database. Each candidate gene was searched against this database (BLASTN v2.16.0, E ≤ 1 × 10^−6^). To uncover deeper similarity, consensus TE sequences were extracted and queried with HMMER3 nhmmscan (-E 1e-6) (Eddy 2011). Full-length candidates were also searched against the Dfam 3.8 nucleotide consensus set (BLASTN, E ≤ 1 × 10^−6^) (Storer et al. 2021). Protein domains were sought by HMMER3 hmmsearch (-E 1e-6) against Pfam 35.0 (Mistry et al. 2021). Coordinates from Krait2 and TE homology were merged and visualized on candidate alignments (TEs, green; simple repeat sequences, blue).

### Gene expression specificity across tissues

In each tissue and sex, a measure of gene expression specificity, was calculated using the average TPM of multiple samples according to the following formula:

where is the number of tissues, is the expression level of a gene in tissue i, and is the highest expression level of a gene detected in all tissues examined. values theoretically range from 0 to 1, with lower values indicating an expression pattern that is evenly distributed across all tissues examined and higher values indicating greater variation in expression levels among tissues and higher tissue specificity.

### Quantifying conservation of transposable elements across insect species

The classification of TE fragments was based on the output of TransposonPSI. Each TE fragment detected within *de novo* gene candidates was assigned to a TE class according to the TransposonPSI annotation. To investigate the evolutionary conservation of these fragments, BLASTN searches (E ≤ 1 × 10^−6^) were performed using TE-derived sequences from *Te. occipitalis de novo* gene candidates as queries against the genomes of other insects. Genes containing TE fragments were examined for gene duplication by intra-genomic BLASTP searches (E ≤ 1 × 10^−6^), in which any intraspecific hit other than the self-match was defined as a duplicated gene. Furthermore, to confirm the presence or absence of TE segments within the detected duplicate genes, we performed BLASTX searches (E ≤ 1 × 10^−6^) using the TE segments as queries and the protein sequences of the duplicate genes as the database.

### Evolutionary Age Classification and Statistical Analysis

All *Te. occipitalis* genes were assigned to one of six evolutionary age classes, based on the orthogroup framework described above (see “Identification of Species-Specific Gene Candidates”): (i) Category 9 genes conserved across insects (Insect-common), (ii) Category 8 genes shared within Polyneoptera (Polyneoptera-common), (iii) Category 7 genes restricted to Orthoptera (Orthoptera-common), (iv) Categories 2–6 genes restricted to Ensifera (Ensifera-common), (v) Category 1 genes excluding *de novo* gene candidates (species-specific), and (vi) *de novo* gene candidates.

Across these classes, we analyzed exon number, protein sequence length, tissue specificity index (τ), and four types of repetitive sequence overlap (TEs in introns, simple repeat sequences in introns, TEs in exons, and simple repeat sequences in exons). For exon number, protein length, and τ, values were calculated on an orthogroup basis, with duplicated genes averaged before computation. For TE and simple sequence repeats overlaps, orthogroups in which > 50% of genes contained a given overlap were defined as “overlap-rich,” and for each evolutionary age class, we calculated the proportion of overlap-rich orthogroups relative to all orthogroups in that class. Statistical analyses were tailored to the property examined: exon number, protein length, and τ were tested for correlations with evolutionary age (age classes (i)–(vi) coded from oldest to youngest) using Spearman’s rank correlation coefficient and the Jonckheere–Terpstra test, whereas age-dependent changes in the proportion of overlap-rich orthogroups were evaluated with the two-tailed Cochran–Armitage trend test and ordinal logistic regression.

### Quantifying conservation of repetitive segments in all *Te. occipitalis* genes

To evaluate the evolutionary conservation of repetitive sequences in *Te. occipitalis* genes relative to their orthologs, exon regions showing sequence similarity to TEs were defined as TE-homologous regions. Overlapping TE-homologous regions within the same gene were merged and treated as a single region. In parallel, simple repetitive sequences detected within exons of *Te. occipitalis* genes were identified, and overlapping regions within each gene were likewise merged and treated as single regions.

For each *Te. occipitalis* gene in which a repetitive sequence region was identified, all orthologous genes assigned by OrthoFinder were retrieved, including additional *Te. occipitalis* genes belonging to the same orthogroup when applicable. Pairwise alignments were then performed between the focal *Te. occipitalis* gene and each corresponding ortholog. For each pairwise alignment, two classes of regions were defined: (i) regions corresponding to the repetitive sequence region identified in the focal *Te. occipitalis* gene, and (ii) background regions outside these regions. When repetitive sequence regions were detected in multiple *Te. occipitalis* genes within the same orthogroup, mapped to overlapping but non-identical alignment coordinates relative to the focal *Te. occipitalis* gene, these regions were combined by taking their union and treated as a single repetitive sequence region for downstream analyses. Repetitive sequence regions shorter than 12 amino acids were excluded from subsequent analyses.

To quantify conservation, alignment identity values of repetitive sequence regions were compared with those of background regions for each *Te. occipitalis*–ortholog gene pair. Regions were excluded from analysis if the background-region identity was ≤35%, if the gap proportion exceeded 60%, or if the background-region length was <96 bp, as these conditions indicated insufficient alignment quality. Because conservation states may differ among species, each orthologous comparison was evaluated independently. For each comparison, the mean and standard deviation of alignment identity were calculated from background regions, with standard deviation defined as the square root of the variance of identity values across background-region alignment columns. Repetitive sequence regions were classified based on their relative alignment identity compared with background regions. Regions with identity values within ±3 standard deviations of the background-region identity were classified as Neutral, those with higher identity values as Conserved, and those with lower identity values as Diverged. Regions for which the orthologous counterpart exhibited a gap proportion greater than 10% relative to the repetitive sequence region in *Te. occipitalis* were classified as Non-shared. All thresholds used in this analysis were determined using false-positive tests based on randomized sequences, with parameter settings chosen to maintain a false-positive rate below 5%.

To further examine the evolutionary conservation of repetitive sequences specifically identified in *de novo* genes, TE-homologous regions and simple sequence repeat regions detected within *de novo* genes were used as queries for homology searches against each insect genome included in the dataset used in this study. For genes exhibiting detectable homology to these repetitive sequence regions, orthologs were identified and aligned, and conservation of the corresponding regions was evaluated using the same criteria described above.

### Software availability

The scripts used for the analyses in this study are available in GitHub (https://github.com/kataokaklab/DeNovoGene_Identification).

## DATA ACCESS

The raw sequencing data generated in this study have been submitted to the NCBI Sequence Read Archive (SRA; https://www.ncbi.nlm.nih.gov/sra) under BioProject accession number PRJNA1348230. The genome assemblies and gene annotations of *Te. occipitalis* and *Ta. portentosus* generated in this study are available at figshare (https://doi.org/10.6084/m9.figshare.31337614).

## COMPETING INTEREST STATEMENT

The authors declare no competing interests.

## ACKNOWLEDGMENTS

We thank Akihiko Ichikawa for species identification.

## Funding

This study was supported by the Cabinet Office, Government of Japan, Moonshot Research and Development Program for Agriculture, Forestry and Fisheries (funding agency: Biooriented Technology Research Advancement Institution) (JPJ009237), the Japan Society for the Promotion of Science (JSPS) Research Fellowships for Young Scientists (24KJ2092), and Japan Science and Technology Agency (JST) BOOST Program (JPMJBY24C1). CGE is an investigator of the Howard Hughes Medical Institute.

## Author Contributions

Conceptualization: R.S., K.S., K.K.

Methodology: R.S., K.S., K.K.

Resources: R.S., Y.A., S.H., K.H., K.N., K.K.

Visualization: R.S., K.S., K.K.

Writing—original draft: R.S., K.S., K.K.

Writing—review and editing: R.S., K.S., K.N., T.S., A.O., K.Y., T.A., C.G.E., K.K.

Supervision: K.K.

## Notes

### Competing Interest Statement

The authors have declared no competing interest.

### Summary of Updates

This version has been revised as follows. The title has been changed to "Remnants of repetitive DNA shape the earliest stages of de novo gene evolution in insects". The Abstract, Introduction and Discussion have been revised to clarify the role of repeat-derived sequences in de novo gene origination. Figures 1 to 4 and the Supplemental Material have been updated and corrected, including Table 1 and the Methods describing sequencing and software versions. Data availability has been updated with the figshare DOI and the NCBI BioProject accession. Minor errors in the text have been corrected throughout.

## REFERENCES

Abramson J, Adler J, Dunger J, Evans R, Green T, Pritzel A, Ronneberger O, Willmore L, Ballard AJ, Bambrick J, et al. 2024. Accurate structure prediction of biomolecular interactions with AlphaFold 3. Nature 630: 493–500. 10.1038/s41586-024-07487-w.

Altschul SF, Gish W, Miller W, Myers EW, Lipman DJ. 1990. Basic local alignment search tool. J Mol Biol 215: 403–410. 10.1016/S0022-2836(05)80360-2.

Aury J-M, Istace B. 2021. Hapo-G, haplotype-aware polishing of genome assemblies with accurate reads. NAR Genom Bioinform 3: lqab034. 10.1093/nargab/lqab034.

Blevins WR, Ruiz-Orera J, Messeguer X, Blasco-Moreno B, Villanueva-Cañas JL, Espinar L, Díez J, Carey LB, Albà MM. 2021. Uncovering de novo gene birth in yeast using deep transcriptomics. Nat Commun 12: 604. 10.1038/s41467-021-20911-3.

Bornberg-Bauer E, Hlouchova K, Lange A. 2021. Structure and function of naturally evolved de novo proteins. Curr Opin Struct Biol 68: 175–183. 10.1016/j.sbi.2020.11.010.

Breathnach R, Chambon P. 1981. Organization and expression of eucaryotic split genes coding for proteins. Annu Rev Biochem 50: 349–383. 10.1146/annurev.bi.50.070181.002025.

Brennecke J, Aravin AA, Stark A, Dus M, Kellis M, Sachidanandam R, Hannon GJ. 2007. Discrete small RNA-generating loci as master regulators of transposon activity in Drosophila. Cell 128: 1089–1103. 10.1016/j.cell.2007.01.043.

Buschiazzo E, Gemmell NJ. 2006. The rise, fall and renaissance of microsatellites in eukaryotic genomes. Bioessays 28: 1040–1050. 10.1002/bies.20470.

Cantalapiedra CP, Hernández-Plaza A, Letunic I, Bork P, Huerta-Cepas J. 2021. EggNOG-mapper v2: Functional annotation, orthology assignments, and domain prediction at the metagenomic scale. Mol Biol Evol 38: 5825–5829. 10.1093/molbev/msab293.

Carvunis A-R, Rolland T, Wapinski I, Calderwood MA, Yildirim MA, Simonis N, Charloteaux B, Hidalgo CA, Barbette J, Santhanam B, et al. 2012. Proto-genes and de novo gene birth. Nature 487: 370–374. 10.1038/nature11184.

Challis R, Richards E, Rajan J, Cochrane G, Blaxter M. 2020. BlobToolKit - interactive quality assessment of genome assemblies. G3 (Bethesda) 10: 1361–1374. 10.1534/g3.119.400908.

Chen N. 2004. Using RepeatMasker to identify repetitive elements in genomic sequences. Curr Protoc Bioinformatics **Chapter** 4: Unit 4.10. 10.1002/0471250953.bi0410s05.

Chen S, Zhou Y, Chen Y, Gu J. 2018. fastp: an ultra-fast all-in-one FASTQ preprocessor. Bioinformatics 34: i884–i890. 10.1093/bioinformatics/bty560.

Chen Y, Nie F, Xie S-Q, Zheng Y-F, Dai Q, Bray T, Wang Y-X, Xing J-F, Huang Z-J, Wang D-P, et al. 2021. Efficient assembly of nanopore reads via highly accurate and intact error correction. Nat Commun 12: 60. 10.1038/s41467-020-20236-7.

Cordaux R, Batzer MA. 2009. The impact of retrotransposons on human genome evolution. Nat Rev Genet 10: 691–703. 10.1038/nrg2640.

Dossey AT, Oppert B, Chu F-C, Lorenzen MD, Scheffler B, Simpson S, Koren S, Johnston JS, Kataoka K, Ide K. 2023. Genome and Genetic Engineering of the House Cricket (Acheta domesticus): A Resource for Sustainable Agriculture. Biomolecules 13. 10.3390/biom13040589.

Du L, Chen J, Sun D, Zhao K, Zeng Q, Yang N. 2025. Krait2: a versatile software for microsatellite investigation, visualization and marker development. BMC Genomics 26: 72. 10.1186/s12864-025-11252-2.

Dudchenko O, Batra SS, Omer AD, Nyquist SK, Hoeger M, Durand NC, Shamim MS, Machol I, Lander ES, Aiden AP, et al. 2017. De novo assembly of the Aedes aegypti genome using Hi-C yields chromosome-length scaffolds. Science 356: 92–95. 10.1126/science.aal3327.

Durand NC, Robinson JT, Shamim MS, Machol I, Mesirov JP, Lander ES, Aiden EL. 2016a. Juicebox provides a visualization system for hi-C contact maps with unlimited zoom. Cell Syst 3: 99–101. 10.1016/j.cels.2015.07.012.

Durand NC, Shamim MS, Machol I, Rao SSP, Huntley MH, Lander ES, Aiden EL. 2016b. Juicer provides a one-click system for analyzing loop-resolution hi-C experiments. Cell Syst 3: 95–98. 10.1016/j.cels.2016.07.002.

Eddy SR. 2011. Accelerated profile HMM searches. PLoS Comput Biol 7: e1002195. 10.1371/journal.pcbi.1002195.

Emms DM, Kelly S. 2019. OrthoFinder: phylogenetic orthology inference for comparative genomics. Genome Biol 20: 238. 10.1186/s13059-019-1832-y.

Flynn JM, Hubley R, Goubert C, Rosen J, Clark AG, Feschotte C, Smit AF. 2020. RepeatModeler2 for automated genomic discovery of transposable element families. Proc Natl Acad Sci U S A 117: 9451–9457. 10.1073/pnas.1921046117.

Gabriel L, Brůna T, Hoff KJ, Ebel M, Lomsadze A, Borodovsky M, Stanke M. 2024. BRAKER3: Fully automated genome annotation using RNA-seq and protein evidence with GeneMark-ETP, AUGUSTUS, and TSEBRA. Genome Res 34: 769–777. 10.1101/gr.278090.123.

Grabherr MG, Haas BJ, Yassour M, Levin JZ, Thompson DA, Amit I, Adiconis X, Fan L, Raychowdhury R, Zeng Q, et al. 2011. Full-length transcriptome assembly from RNA-Seq data without a reference genome. Nat Biotechnol 29: 644–652. 10.1038/nbt.1883.

Grandchamp A, Aubel M, Eicholt LA, Roginski P, Luria V, Karger A, Dohmen E. 2025. De Novo gene emergence: Summary, classification, and challenges of current methods. Genome Biol Evol 17. 10.1093/gbe/evaf197.

Grandchamp A, Kühl L, Lebherz M, Brüggemann K, Parsch J, Bornberg-Bauer E. 2023. Population genomics reveals mechanisms and dynamics of de novo expressed open reading frame emergence in Drosophila melanogaster. Genome Res 33: 872–890. 10.1101/gr.277482.122.

Gubala AM, Schmitz JF, Kearns MJ, Vinh TT, Bornberg-Bauer E, Wolfner MF, Findlay GD. 2017. The goddard and saturn genes are essential for Drosophila male fertility and may have arisen de novo. Mol Biol Evol msx057. 10.1093/molbev/msx057.

Gurevich A, Saveliev V, Vyahhi N, Tesler G. 2013. QUAST: quality assessment tool for genome assemblies. Bioinformatics 29: 1072–1075. 10.1093/bioinformatics/btt086.

Heinen TJAJ, Staubach F, Häming D, Tautz D. 2009. Emergence of a new gene from an intergenic region. Curr Biol 19: 1527–1531. 10.1016/j.cub.2009.07.049.

Jones P, Binns D, Chang H-Y, Fraser M, Li W, McAnulla C, McWilliam H, Maslen J, Mitchell A, Nuka G, et al. 2014. InterProScan 5: genome-scale protein function classification. Bioinformatics 30: 1236–1240. 10.1093/bioinformatics/btu031.

Katoh K, Standley DM. 2013. MAFFT multiple sequence alignment software version 7: improvements in performance and usability. Mol Biol Evol 30: 772–780. 10.1093/molbev/mst010.

Kazazian HH Jr. 2004. Mobile elements: drivers of genome evolution. Science 303: 1626–1632. 10.1126/science.1089670.

Keilwagen J, Hartung F, Grau J. 2019. GeMoMa: Homology-based gene prediction utilizing intron position conservation and RNA-seq data. Methods Mol Biol 1962: 161–177. 10.1007/978-1-4939-9173-0_9.

Khalturin K, Hemmrich G, Fraune S, Augustin R, Bosch TCG. 2009. More than just orphans: are taxonomically-restricted genes important in evolution? Trends Genet 25: 404–413. 10.1016/j.tig.2009.07.006.

Kim D, Paggi JM, Park C, Bennett C, Salzberg SL. 2019. Graph-based genome alignment and genotyping with HISAT2 and HISAT-genotype. Nat Biotechnol 37: 907–915. 10.1038/s41587-019-0201-4.

Knowles DG, McLysaght A. 2009. Recent de novo origin of human protein-coding genes. Genome Res 19: 1752–1759. 10.1101/gr.095026.109.

Kolmogorov M, Yuan J, Lin Y, Pevzner PA. 2019. Assembly of long, error-prone reads using repeat graphs. Nat Biotechnol 37: 540–546. 10.1038/s41587-019-0072-8.

Kovaka S, Zimin AV, Pertea GM, Razaghi R, Salzberg SL, Pertea M. 2019. Transcriptome assembly from long-read RNA-seq alignments with StringTie2. Genome Biol 20: 278. 10.1186/s13059-019-1910-1.

Krzywinski M, Schein J, Birol I, Connors J, Gascoyne R, Horsman D, Jones SJ, Marra MA. 2009. Circos: an information aesthetic for comparative genomics. Genome Res 19: 1639–1645. 10.1101/gr.092759.109.

Lange A, Patel PH, Heames B, Damry AM, Saenger T, Jackson CJ, Findlay GD, Bornberg-Bauer E. 2021. Structural and functional characterization of a putative de novo gene in Drosophila. Nat Commun 12: 1667. 10.1038/s41467-021-21667-6.

Lee YCG. 2022. Synergistic epistasis of the deleterious effects of transposable elements. Genetics 220: iyab211. 10.1093/genetics/iyab211.

Levine MT, Jones CD, Kern AD, Lindfors HA, Begun DJ. 2006. Novel genes derived from noncoding DNA in Drosophila melanogaster are frequently X-linked and exhibit testis-biased expression. Proc Natl Acad Sci U S A 103: 9935–9939. 10.1073/pnas.0509809103.

Li H. 2013. Aligning sequence reads, clone sequences and assembly contigs with BWA-MEM. *arXiv:1303.3997 [q-bio.GN].* http://arxiv.org/abs/1303.3997.

Li H. 2016. Minimap and miniasm: fast mapping and de novo assembly for noisy long sequences. Bioinformatics 32: 2103–2110. 10.1093/bioinformatics/btw152.

Li H, Handsaker B, Wysoker A, Fennell T, Ruan J, Homer N, Marth G, Abecasis G, Durbin R, 1000 Genome Project Data Processing Subgroup. 2009. The Sequence Alignment/Map format and SAMtools. Bioinformatics 25: 2078–2079. 10.1093/bioinformatics/btp352.

Lisch D. 2013. How important are transposons for plant evolution? Nat Rev Genet 14: 49–61. 10.1038/nrg3374.

Liu H, Wu S, Li A, Ruan J. 2021. SMARTdenovo: a de novo assembler using long noisy reads. Gigabyte 2021: 1–9. 10.46471/gigabyte.15.

Love MI, Huber W, Anders S. 2014. Moderated estimation of fold change and dispersion for RNA-seq data with DESeq2. Genome Biol 15: 550. 10.1186/s13059-014-0550-8.

Manna GK, Bhattacharjee TK. 1964. Studies of Gryllid Chromosomes. Cytologia (Tokyo) 29: 196–206. 10.1508/cytologia.29.196.

Manni M, Berkeley MR, Seppey M, Simão FA, Zdobnov EM. 2021. BUSCO update: novel and streamlined workflows along with broader and deeper phylogenetic coverage for scoring of eukaryotic, prokaryotic, and viral genomes. Mol Biol Evol 38: 4647–4654. 10.1093/molbev/msab199.

Marçais G, Kingsford C. 2011. A fast, lock-free approach for efficient parallel counting of occurrences of k-mers. Bioinformatics 27: 764–770. 10.1093/bioinformatics/btr011.

McLysaght A, Hurst LD. 2016. Open questions in the study of de novo genes: what, how and why. Nat Rev Genet 17: 567–578. 10.1038/nrg.2016.78.

Mikina W, Hałakuc P, Milanowski R. 2024. Transposon-derived introns as an element shaping the structure of eukaryotic genomes. Mob DNA 15: 15. 10.1186/s13100-024-00325-w.

Misof B, Liu S, Meusemann K, Peters RS, Donath A, Mayer C, Frandsen PB, Ware J, Flouri T, Beutel RG, et al. 2014. Phylogenomics resolves the timing and pattern of insect evolution. Science 346: 763–767. 10.1126/science.1257570.

Mistry J, Chuguransky S, Williams L, Qureshi M, Salazar GA, Sonnhammer ELL, Tosatto SCE, Paladin L, Raj S, Richardson LJ, et al. 2021. Pfam: The protein families database in 2021. Nucleic Acids Res 49: D412–D419. 10.1093/nar/gkaa913.

Nei M, Rooney AP. 2005. Concerted and birth-and-death evolution of multigene families. Annu Rev Genet 39: 121–152. 10.1146/annurev.genet.39.073003.112240.

Neme R, Tautz D. 2016. Fast turnover of genome transcription across evolutionary time exposes entire non-coding DNA to de novo gene emergence. Elife 5: e09977. 10.7554/eLife.09977.

Palmieri N, Kosiol C, Schlötterer C. 2014. The life cycle of Drosophila orphan genes. Elife 3: e01311. 10.7554/eLife.01311.

Patro R, Duggal G, Love MI, Irizarry RA, Kingsford C. 2017. Salmon provides fast and bias-aware quantification of transcript expression. Nat Methods 14: 417–419. 10.1038/nmeth.4197.

Pedersen BS, Quinlan AR. 2018. Mosdepth: quick coverage calculation for genomes and exomes. Bioinformatics 34: 867–868. 10.1093/bioinformatics/btx699.

Pertea G, Pertea M. 2020. GFF utilities: GffRead and GffCompare. F1000Res 9: 304. 10.12688/f1000research.23297.2.

Rajon E, Masel J. 2011. Evolution of molecular error rates and the consequences for evolvability. Proc Natl Acad Sci U S A 108: 1082–1087. 10.1073/pnas.1012918108.

Ranallo-Benavidez TR, Jaron KS, Schatz MC. 2020. GenomeScope 2.0 and Smudgeplot for reference-free profiling of polyploid genomes. Nat Commun 11: 1432. 10.1038/s41467-020-14998-3.

Roach MJ, Schmidt SA, Borneman AR. 2018. Purge Haplotigs: allelic contig reassignment for third-gen diploid genome assemblies. BMC Bioinformatics 19: 460. 10.1186/s12859-018-2485-7.

Roy SW. 2004. The origin of recent introns: transposons? Genome Biol 5: 251. 10.1186/gb-2004-5-12-251.

Ruan J, Li H. 2019. Fast and accurate long-read assembly with wtdbg2. Nat Methods 17: 155–158. 10.1038/s41592-019-0669-3.

Sayers EW, Beck J, Bolton EE, Bourexis D, Brister JR, Canese K, Comeau DC, Funk K, Kim S, Klimke W, et al. 2021. Database resources of the National Center for Biotechnology Information. Nucleic Acids Res 49: D10–D17. 10.1093/nar/gkaa892.

Schlötterer C. 2015. Genes from scratch--the evolutionary fate of de novo genes. Trends Genet 31: 215–219. 10.1016/j.tig.2015.02.007.

Shafin K, Pesout T, Lorig-Roach R, Haukness M, Olsen HE, Bosworth C, Armstrong J, Tigyi K, Maurer N, Koren S, et al. 2020. Nanopore sequencing and the Shasta toolkit enable efficient de novo assembly of eleven human genomes. Nat Biotechnol 38: 1044–1053. 10.1038/s41587-020-0503-6.

Simão FA, Waterhouse RM, Ioannidis P, Kriventseva EV, Zdobnov EM. 2015. BUSCO: assessing genome assembly and annotation completeness with single-copy orthologs. Bioinformatics 31: 3210–3212. 10.1093/bioinformatics/btv351.

Storer J, Hubley R, Rosen J, Wheeler TJ, Smit AF. 2021. The Dfam community resource of transposable element families, sequence models, and genome annotations. Mob DNA 12: 2. 10.1186/s13100-020-00230-y.

Tautz D, Domazet-Lošo T. 2011. The evolutionary origin of orphan genes. Nat Rev Genet 12: 692–702. 10.1038/nrg3053.

UniProt Consortium. 2023. UniProt: The universal protein knowledgebase in 2023. Nucleic Acids Res 51: D523–D531. 10.1093/nar/gkac1052.

Vakirlis N, Acar O, Hsu B, Castilho Coelho N, Van Oss SB, Wacholder A, Medetgul-Ernar K, Bowman RW 2nd, Hines CP, Iannotta J, et al. 2020. De novo emergence of adaptive membrane proteins from thymine-rich genomic sequences. Nat Commun 11: 781. 10.1038/s41467-020-14500-z.

Vakirlis N, Fuqua T. 2025. Intergenic polyA/T tracts explain the propensity of yeast de novo genes to encode transmembrane domains. J Evol Biol 38: 1272–1277. 10.1093/jeb/voaf089.

Vakirlis N, Hebert AS, Opulente DA, Achaz G, Hittinger CT, Fischer G, Coon JJ, Lafontaine I. 2018. A molecular portrait of DE Novo genes in yeasts. Mol Biol Evol 35: 631–645. 10.1093/molbev/msx315.

Van Oss SB, Carvunis A-R. 2019. De novo gene birth. PLoS Genet 15: e1008160. 10.1371/journal.pgen.1008160.

Vaser R, Šikić M. 2021. Time- and memory-efficient genome assembly with Raven. Nat Comput Sci 1: 332*–*336. 10.1038/s43588-021-00073-4.

Wang Y, Tang H, Debarry JD, Tan X, Li J, Wang X, Lee T-H, Jin H, Marler B, Guo H, et al. 2012. MCScanX: a toolkit for detection and evolutionary analysis of gene synteny and collinearity. Nucleic Acids Res 40: e49. 10.1093/nar/gkr1293.

Weisman CM, Murray AW, Eddy SR. 2020. Many, but not all, lineage-specific genes can be explained by homology detection failure. PLoS Biol 18: e3000862. 10.1371/journal.pbio.3000862.

Wissler L, Gadau J, Simola DF, Helmkampf M, Bornberg-Bauer E. 2013. Mechanisms and dynamics of orphan gene emergence in insect genomes. Genome Biol Evol 5: 439–455. 10.1093/gbe/evt009.

Wu D-D, Irwin DM, Zhang Y-P. 2011. De novo origin of human protein-coding genes. PLoS Genet 7: e1002379. 10.1371/journal.pgen.1002379.

Yanai I, Benjamin H, Shmoish M, Chalifa-Caspi V, Shklar M, Ophir R, Bar-Even A, Horn-Saban S, Safran M, Domany E, et al. 2005. Genome-wide midrange transcription profiles reveal expression level relationships in human tissue specification. Bioinformatics 21: 650–659. 10.1093/bioinformatics/bti042.

You P, Xie Z-H. 2003. A study on chromosomes in five crickets. Yi Chuan 25: 529–532. https://europepmc.org/article/med/15639923.

Zhang L, Ren Y, Yang T, Li G, Chen J, Gschwend AR, Yu Y, Hou G, Zi J, Zhou R, et al. 2019. Rapid evolution of protein diversity by de novo origination in Oryza. Nat Ecol Evol 3: 679–690. 10.1038/s41559-019-0822-5.

Zimin AV, Marçais G, Puiu D, Roberts M, Salzberg SL, Yorke JA. 2013. The MaSuRCA genome assembler. Bioinformatics 29: 2669–2677. 10.1093/bioinformatics/btt476.

Zimin AV, Salzberg SL. 2020. The genome polishing tool POLCA makes fast and accurate corrections in genome assemblies. PLoS Comput Biol 16: e1007981. 10.1371/journal.pcbi.1007981.

