## Supplemental Material for "Remnants of repetitive DNA shape the earliest stages of *de novo* gene evolution in insects"

### Contents

Supplemental Fig. S1. Representative Gryllidea crickets used in this study. 2

Supplemental Fig. S2. K-mer–based genome size estimation of crickets. 3

Supplemental Fig. S3. BlobToolKit-based contaminant screening of the cricket genome assemblies. 4

Supplemental Fig. S4. Interspersed repeat landscapes of the cricket genomes. 5

Supplemental Fig. S5. Nucleotide and exon-level protein alignments of *de novo* gene candidates in *Te. occipitalis* across cricket genomes. 6

Supplemental Fig. S6. AlphaFold3-predicted structures of *de novo* gene candidate proteins in *Te. occipitalis*. 140

Supplemental Fig. S7. Enrichment of transposable element fragments and simple repeats in exons and introns across gene age classes. 144

Supplemental Fig. S8. Distribution of TE midpoints along genes across conservation categories. 145

Supplemental Fig. S9. Genome sequencing and assembly workflow for *Ta. portentosus* and *Te. occipitalis*. 146

Supplemental Fig. S10. Transcriptome-based genome annotation workflow for *Ta. portentosus* and *Te. occipitalis*. 147

Supplemental Table legends (Supplemental Tables S1–S12) 148

Supplemental Tables S1–S12 are provided as separate Excel files (Supplemental_Table_S1.xlsx–Supplemental_Table_S12.xlsx).

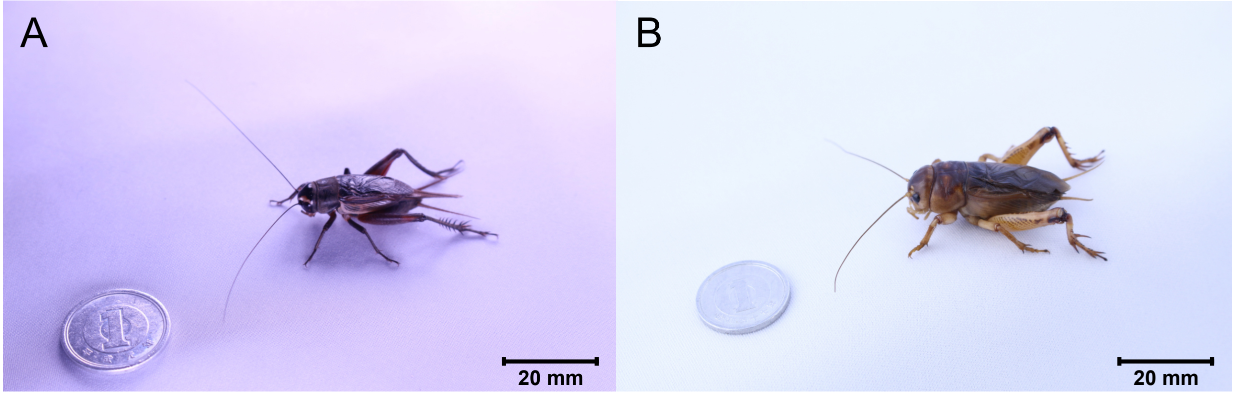


**Supplemental Fig. S1. Representative Gryllidea crickets used in this study.** (**A**) Adult male of *Te. occipitalis*. (**B**) Adult male of *Ta. portentosus*.

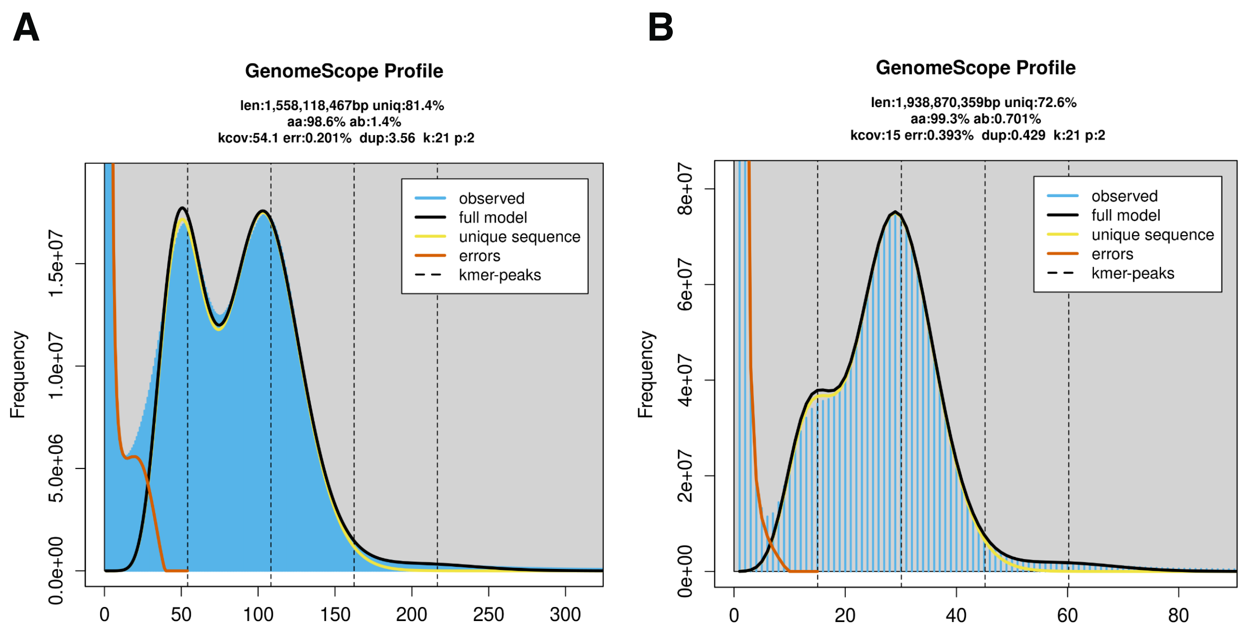


**Supplemental Fig. S2. K-mer–based genome size estimation of crickets.** Genome size was estimated from Illumina short-read data using GenomeScope v2.3.0 based on 21-mer frequency distributions, with k-mers counted by Jellyfish v2.3.0. (**A**) GenomeScope profile for *Ta. portentosus* (estimated genome length, 1,558,118,467 bp; unique sequence, 81.4%; kcov, 54.1; err, 0.201%; dup, 3.56). (**B**) GenomeScope profile for *Te. occipitalis* (estimated genome length, 1,938,870,359 bp; unique sequence, 72.6%; kcov, 15; err, 0.393%; dup, 0.429). Blue histograms show observed k-mer frequencies (y-axis) across k-mer coverage/multiplicity (x-axis); black curves indicate the fitted GenomeScope model; yellow curves indicate the unique-sequence component; orange curves indicate error k-mers; dashed vertical lines mark inferred k-mer peaks.

A

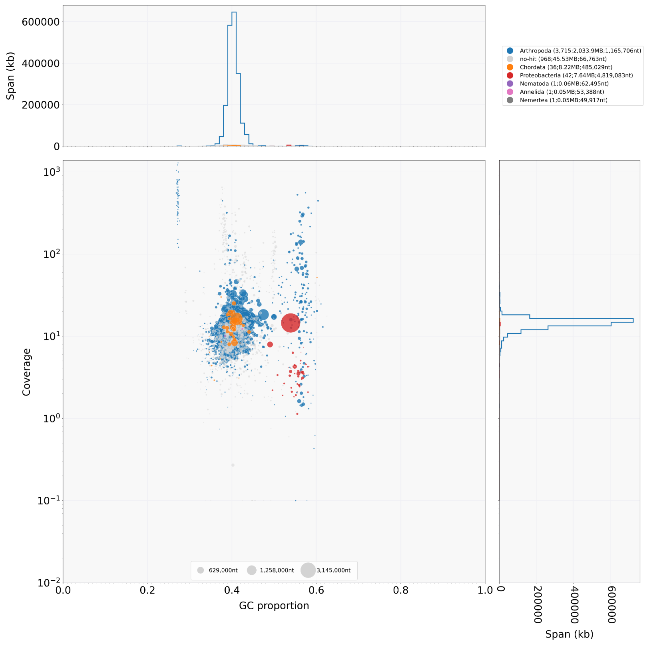


B

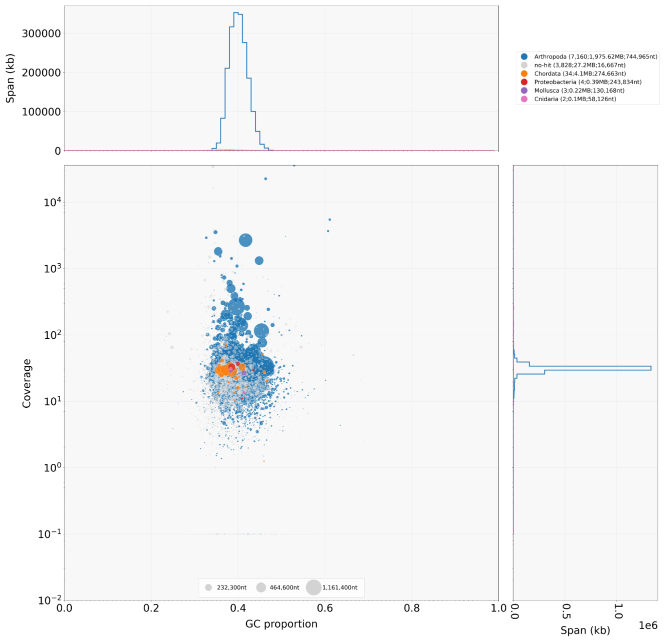


**Supplemental Fig. S3. BlobToolKit-based contaminant screening of the cricket genome assemblies.** GC–coverage "blobplots" were generated with BlobToolKit v1.1.1 to identify and remove contaminant scaffolds/contigs based on atypical sequencing coverage, GC proportion, and sequence homology. Each point represents a scaffold/contig plotted by GC proportion (x-axis) and Illumina read coverage (y-axis); point size is proportional to sequence span, and marginal histograms summarize the span distributions. Coverage was calculated by mapping Illumina reads with BWA, and taxonomic assignments were inferred from BLASTN v2.12.0 searches against the NCBI nt database (v5) using megablast (culling_limit 10; E-value threshold 1e–25; tabular output including taxonomy fields). (**A**) *Ta. portentosus*. (**B**) *Te. occipitalis*.

**A**

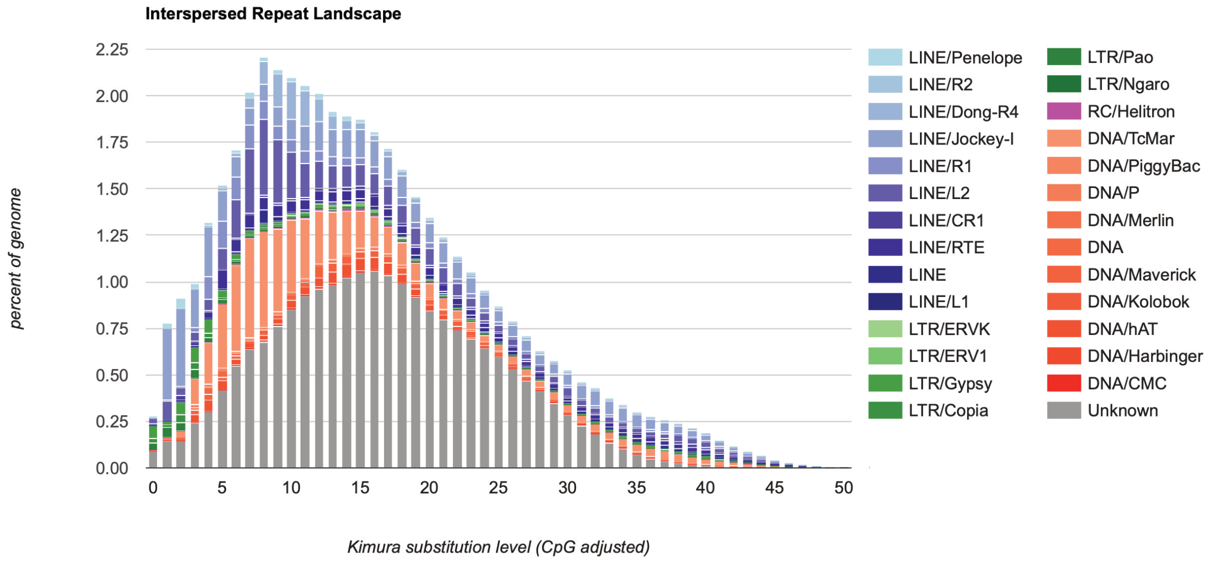


**B**

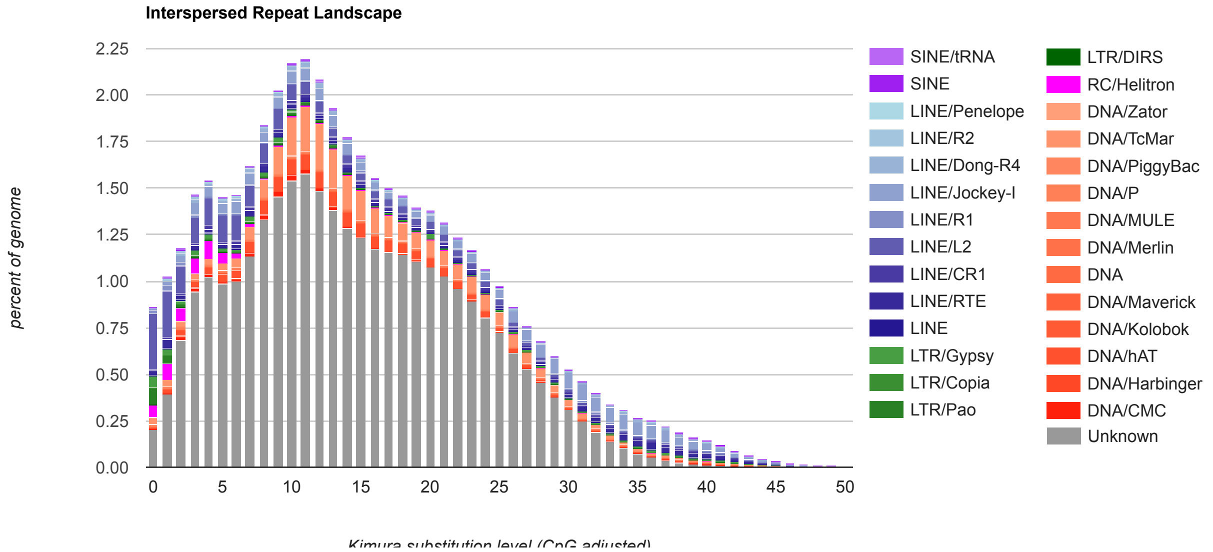


**Supplemental Fig. S4. Interspersed repeat landscapes of the cricket genomes.** Repeat landscapes were generated from a *de novo* repeat library built with RepeatModeler v2.0.5 and summarized as the fraction of the genome occupied by interspersed repeats across CpG-adjusted Kimura divergence from the repeat consensus (x-axis; percent of genome on y-axis). (**A**) *Ta. portentosus*. (**B**) *Te. occipitalis*. Lower Kimura values indicate less-diverged (putatively more recent) insertions.

### Supplemental Fig. S5

**Supplemental Fig. S5. Nucleotide and exon-level protein alignments of *de novo* gene candidates in *Te. occipitalis* across cricket genomes.** For each final candidate (gene IDs shown), the nucleotide sequence spanning the full gene plus 300-bp flanks and the amino-acid sequence of each exon were aligned to orthologous syntenic regions in other cricket genomes. Nucleotide alignments were generated with MAFFT v7 using the --auto option. Syntenic regions from other crickets were six-frame translated while accounting for strand and potential open reading frame shifts; for each exon, the longest contiguous match was selected, allowing different optimal frames among exons. These alignments were used to confirm sequence continuity and frame integrity in *Te. occipitalis* and the presence of homologous but non-coding sequence in other crickets.

**Tocc.1G0004690.1**

```
Teleogryllus_occipitalis    ATTCAAATATCCCAACTCCAAATGACAAAGCTGAGGGATACAGTAGTTCATAAAAGCATTCACGAGATACGAAATGAATGGAAGGACGTCGCCCCATTAAAAATATTAAAATATAAAAGA
Acheta_domesticus           ATTCAAAAATCCCAACTCCAAATGACAGAGCTGAGAGATACAGTGGGTTATCAAAGCATTCACGAGATATGTGATCAATGGCAGGAGGTCGCTCCATTAAAAAGATTAAAATATAAAAGA
Consensus                   ******* ****************** ******* ******** * * ** **************** *  ** ***** **** ***** ********** ****************

Teleogryllus_occipitalis    GAAAAATTCTGAGGCCCTATTAAGAGTAAAAGGGCGTGTACTCTGACTCTACCATCCCATGCCGGGCGCGTCGTCCACAATCCTCCCGTCGCATAGGCCCAGAGTTTCCGGCCGCCTGCG
Acheta_domesticus           GAAAAATGCTGAGTCCCTATTAAAAGCAAAATGGCGTGTGCTCTGACTCTACCATCCCATCTCGCTCGCGTCGTCCGCAATCCTC----------------GAGTTTCCGGCCGCCCGCG
Consensus                   ******* ***** ********* ** **** ******* ********************  **  ********** ********                **************** ***

Teleogryllus_occipitalis    CTGCAGCCCCCCTCTCTGAAGCGGTCTGCTCGCCT---CTTCGGCGAGCGTAATCGCGCAATATGTTTCCATTAGCAGTTAATACTCGTGTTTTGTCGCTGCTTGTGGTGTAAATGCGA
Acheta_domesticus           CTGCAG----CCGTCTCCGAAGCGGTCGGCTCGCCTCCACTTCGGCGAGCGTAATCGCGCAATATGTTTCCATTAGCAGTTAATACTCTTGTTTTGTCGCTGCTTGTGGTGTAAATGCGA
Consensus                   ******    ** **** ********* ********   ************************************************* ******************************

Teleogryllus_occipitalis    CGTCTTTCTTGCCGAAGGGCGGCCTCCACTAAGACTTATGAATACGGATATCGTCGAGAAAATTCCGCCATTATTTGTCTCGGTGCATTTCTCTTGTTTCGGCTGCGGGCCTCGAGGACA
Acheta_domesticus           CGTCTTTCTTGCCGAAGGGCGGCCTCCTCTAAGACTTATGAATACGGATATCGTCGAGAAAATTCCGCCATTATTTGTCTCGGTGCATTTCTCTTGTTTCGGCTGCGAGCCTCGAGGACA
Consensus                   *************************** ******************************************************************************** ***********

Teleogryllus_occipitalis    TTATATTCATTCATCAAAAACAAAGCTTGTTGCGTTGGCGTGACGCGCCGACGGCGGCGGCGGCGATGGCGATAGAGGCAGCGGCGGCACTGGCGGTGGTGGCGGTGGTGGCGGTGGTGG
Acheta_domesticus           TTATATTCATTCATCAAAAACAAAGCTTGTTGCGTTGGCGTGACGCGCCGGCGGCGGCGGTGGAG----------------------------------------------------------
Consensus                   ************************************************** ********* ** *

Teleogryllus_occipitalis    CGGTGGTGGCGGTCGTGGTGGTGGTGGTGGTGGTGGTGGCAGTAGCGGTAACGGTAACGGTGGCGGTGGCGGTGGCGGTGGCGGTGGCGGTGGCGGTGGCGGTGGCGGTGGCGGTGGCGGTGG
Acheta_domesticus           ------------------------------------------------------------------------------------------------------------------------
Consensus

Teleogryllus_occipitalis    TGGTGATGCTGGTGATGCTGGTGCTGGTGCTGGTGCTGGCTGCGGCGGTGGCGGTGGCGGTGGCGGTGGCGGTGGCGGTGGCGGCGGTGGCGGTGGCGGCGGAGGCGGCGGTGGCGGTGG
Acheta_domesticus           ----------------------------------------------------------------------------------------------------------------GGTTGTGG
Consensus                                                                                                                   **  ****

Teleogryllus_occipitalis    CGGTGGCGGTGGCGGTGGCGGTGGCGGTGGCGGTGGCGGCGACGGCGGCGGCGGCGGCGGCGGCGGTGGCGGTGGCGGCGGCGGCGGTGGCGGCGGTGGCGGCGGTGGCGGTGGCGG
Acheta_domesticus           CGGTGGCGGTGGCGGTGGCGGTGGCGGTGGCGGTGGCGGTGGCGGCGGCGGCGGCGGCGGGGGGCGGAGGCGGCGGCGGCGGCGGCAGCGGCAGCGGCGGTGGCGGTGGGGG
Consensus                   *************************************** * *** ********************** ***** ************** *** ****  *************** **

Teleogryllus_occipitalis    CGGTGGCGGCGGCGGTGGTGGCGGCGGTGGCGGCGGCGGTGGTGGCGGCGGTGGCGGCGGCGGTGGTGGCGGTGGCGGCGGTGGCGGCGGTAGCGGCGCTGGCTGCGGTGGCGGCGCCAG
Acheta_domesticus           C-----------------------------------------------------------------------------------------------------------------------
Consensus                   *

Teleogryllus_occipitalis    CGGCGGCGGCGGCGGCAGCGGTGGCGAGCTGAGCAGAGAGAAGGGGTCTGTTTAATTATTCAAATTGCGGGATGCTGTGGGGCTCGGCAAGGACGAGCTCGCCCGGGCGGGGGAGAGCAC
Acheta_domesticus           ---CGGCGGTGGCGGCAGTGGTGGCGAGCTGAGCAGAGAGAAGGGGTCTGTTTAATTATTCAAATTGCGGGATGCTGTGGGGCTCGGCAAGGACGCTCTGCATTGGGGCGGGGAGAGAGC
Consensus                      ****** ******** **********************************************************************  **     ***  ********  *

Teleogryllus_occipitalis    GCGGG---CGGGTGGAGGCGGTCCGCCAAGCGGCGTGGGCTCGAGCCGCGAGTCGCCGAGTTAGGAGACGGTGTGACGCTGGATCTTGAGGATGGATCAGGAGAGATGGTGAACGGGAAC
Acheta_domesticus           GCGGGGACGGGGCGGAGACGGTCCGCGAAACGGCGTGGGCTCGAGCCACGAGCCGCCAAGTTAGGGAACGGTGCGACGCAGGATCTTGAGGAGGAACCAGAGGAGATGGGGGACGGGAAC
Consensus                   *****    *** **** ******** ** ***************** **** **** *******  ****** ***** ************ * * ***  ******* * ********

Teleogryllus_occipitalis    ACGGCGCATTCGC----------GGGCGAAGACGTGGGCGGGCGTGGGCGGGCGTGGGCGGGCGT--------------------GGGCGGGCGTGGGCGGGCGTGGGCGGGCG------
Acheta_domesticus           ATGGCGCATTCGCGAGCGGAGGTGGGCGAAGACGGGGGCAGACGAGAGCAGGAGTGGGCGAGCATTGAGGCGGGGTCTGGCGCAGGTGCGGGCGTCGGCGAGGATGGTCGGACGGGTGGT
Consensus                   * ***********          *********** **** * ** * ** ** ******* ** *                    * ******** **** *  *** *** **

Teleogryllus_occipitalis    ------------------------------------------------------------------------------------------------------------------------
Acheta_domesticus           CAGGCGGGCTGGCGGACGGACGGAAGAGCGGGCGGACGGAAGAGCGGGCGGACGGAAGAGCGGGCGGACGGAAGAGCGGGCGGACGGAAGAGCGGGCGGACGGAAGAGCGGGCGGACGGA
Consensus

Teleogryllus_occipitalis    ------------------------------------------------------------------------------------------------------------------------
Acheta_domesticus           AGAGCGGGCGGACGGAAGAGCGGGCGGACGGAAGAGCGGGCGGACGGAGGAGCGGGCGGACGGAAGAGCGGGCGGACGGAAGAGCGGGCGGACGGAAGAGCGGGCGGACGGAAGAGCGGG
Consensus

Teleogryllus_occipitalis    ------------------------------------------------------------------------------------------------------------------------
Acheta_domesticus           CGGACGGAAGAGCGGGCGGACGGAAGAGCGGGCGGACGGAAGAGCGGGCGGAAGAGCGGACGGACGGGCGGACGGGCGGACGGGCGGACGGGCGGACGGGCGGACGGGCGGACGGGCGGA
Consensus

Teleogryllus_occipitalis    --------------------------------------------------------------------------------TGGGCGGGCGT-----------------------------
Acheta_domesticus           CGGGCGGACGGGCGGACGGGCGGACGGGCGGACGGGCGGACGGGCGGACGGGCGGACGGGCGGACGGGCGGACGAGCTGACGGGCGGACGTGCTGACGGGCGGACGAGCTGACGGGCGGA
Consensus                                                                                    ****** ***

Teleogryllus_occipitalis    ------------------------------------------------------------------------------------------------------------------------
Acheta_domesticus           CGAGCTGACGGGCGGACGAGCTGACGGGCGGACGAGCTGACGGGCGGACGAGCTGACGGGCGGACGAGCTGACGGGCGGACGAGCTGACGGGCGGACGAGCTGACGGGCGGACGAGCTGA
Consensus

Teleogryllus_occipitalis    ------------------------------------------------------------------------------------------------------------------------
Acheta_domesticus           CGGGCGGACGAGCTGACGGGCGGACGAGCTGACGGGCGGACGAGCTGACGGGCGGACGAGCTGACCGGCGGGCGGGCGGGCGGGCAGGCGGACGAGGCGAAGCGTTCATAAAGACGTCAA
Consensus

Teleogryllus_occipitalis    -------------------------------------------------------
Acheta_domesticus           ATAGTCTTATTAGCAATTAGTACGGAGTGAAAGACGCATGACGCATATACTCCTA
Consensus
```

**Tocc.1G0004690.1_exon1**

```
Teleogryllus_occipitalis      ------------------------------------------------------------------------------------------------------------------------
Acheta_domesticus             IHLSTHKHEKF*KNT*KTSNQ*YIAERFVYLFI*LKSKEYILCVTNRLWREVFSVQVFLNSGIWDKNGQF*ILITNVIFLIKVNHNIVTIF*IVKFEI*AKEENYVGKNMENSSVHTLLS
Consensus

Teleogryllus_occipitalis      ------------------------------------------------------------------------------------------------------------------------
Acheta_domesticus             MHSYIIVNNALLYRNLDWCARERKLLWSLVPFV**MLPRKVMVVSSPLNDEIRCLPTRRFRGIPFSGISPNGALAQSGQALRSRLVSRRVWMWSACKCRPS*CRKHSTLGNQTQGPHVQR
Consensus

Teleogryllus_occipitalis      ------------------------------------------------------------------------------------------------------------------------
Acheta_domesticus             RCRNSGNAGSELKSAGTETRTRLPQIIYVP*ICNEFKNPNSK*QS*EIQWVIKAFTRYVINGRRSLH*KD*NIKEKNAESLLKAKWRVL*LYHPISLASSAILEFPAARAAAVSEAVGSP
Consensus

Teleogryllus_occipitalis      ---------MFPL-----------------------------------------------------------------------------------------------------------
Acheta_domesticus             PLRRA*SRNMFPLAVNTLVLSLLVV*MRRLSCRRAASSKTYEYGYRRENSAIICLGAFLLFRLRASRTLYSFIKNKACCVGVTRRRRRWRVVAVAVAVAVAVAVAVAVAAAAAAAGAEAA
Consensus                               ****

Teleogryllus_occipitalis      ------------------------------------------------------------------------------------------------------------------------
Acheta_domesticus             AAAAAAAAVAVGAGGGGSGGELSREKGSV*LFKLRDAVGLGKDALHWGGERARGRGGDGPRNGVGSSHEPPS*GTVRRRILRRNQRRWGTGTWRIRERRWAKTGADESRSGRALRRGLAQ
Consensus

Teleogryllus_occipitalis      ------------------------------------------------------------------------------------------------------------------------
Acheta_domesticus             VRASARMVGRVVRRAGGRTEERADGRAGGRKSGRTEERADGRAGGRKSGRTEERADGRAGGRKSGRTEERADGRAGGRKSGRTEERADGRAGGRKSGRTEERADGRAGGRADGRADGRTG
Consensus

Teleogryllus_occipitalis      ------------------------------------------------------------------------------------------------------------------------
Acheta_domesticus             GRADGRTGGRADGRTGGRADGRTGGRADGRTGGRADELTGGRADGRTS*RADELTGGRADGRTS*RADELTGGRADGRTS*RADELTGGRADGRTS*RADELTGGRADGRTS*PAGGRAG
Consensus

Teleogryllus_occipitalis      ------------------------------
Acheta_domesticus             RRTRRSVHKDVK*SY*QLVRSERRMTHILL
Consensus
```

**Tocc.1G0004690.1_exon2**

```
Teleogryllus_occipitalis                   ------------------------------------------------------------------------------------------------------------------------
Acheta_domesticus                          IHLSTHKHEKF*KNT*KTSNQ*YIAERFVYLFI*LKSKEYILCVTNRLWREVFSVQVFLNSGIWDKNGQF*ILITNVIFLIKVNHNIVTIF*IVKFEI*AKEENYVGKNMENSSVHTLLS
Consensus                                  

Teleogryllus_occipitalis                   ------------------------------------------------------------------------------------------------------------------------
Acheta_domesticus                          MHSYIIVNNALLYRNLDWCARERKLLWSLVPFV**MLPRKVMVVSSPLNDEIRCLPTRRFRGIPFSGISPNGALAQSGQALRSRLVSRRVWMWSACKCRPS*CRKHSTLGNQTQGPHVQR
Consensus                                  

Teleogryllus_occipitalis                   --------------------------------------------------------------------------------------------------------------AAVA------
Acheta_domesticus                          RCRNSGNAGSELKSAGTETRTRLPQIIYVP*ICNEFKNPNSK*QS*EIQWVIKAFTRYVINGRRSLH*KD*NIKEKNAESLLKAKWRVL*LYHPISLASSAILEFPAARAAAVSEAVGSP
Consensus                                                                                                                                                           ***

Teleogryllus_occipitalis                   -------------------------------------------------------------------------------------------VAVAVAVAVAVAVAATAAAAAAAAVAVAA
Acheta_domesticus                          PLRRA*SRNMFPLAVNTLVLSLLVV*MRRLSCRRAASSKTYEYGYRRENSAIICLGAFLLFRLRASRTLYSFIKNKACCVGVTRRRRRWRVVAVAVAVAVAVAVAVAVAAAAAAAGAEAA
Consensus                                                                                                                             **************   ******* * **

Teleogryllus_occipitalis                   AAVAAVAAVAVAA------------VAAAVVAAVAAAV-------------------------------------------------------------------------------VAA
Acheta_domesticus                          AAAAAAAAVAVGAGGGGSGGELSREKGSV*LFKLRDAVGLGKDALHWGGERARGRGGDGPRNGVGSSHEPPS*GTVRRRILRRNQRRWGTGTWRIERRWAKTGADESRSGRALRRGLAQ
Consensus                                  ** ** ***** *                       **                                                                                *

Teleogryllus_occipitalis                   VAAAVVAVAAVAAVA---------------------------------------------------------------------------------------------------------
Acheta_domesticus                          VRASARMVGRVVRRAGGRTEERADGRAGGRKSGRTEERADGRAGGRKSGRTEERADGRAGGRKSGRTEERADGRAGGRKSGRTEERADGRAGGRKSGRTEERADGRAGGRADGRADGRTG
Consensus                                  * *    *  *   *

Teleogryllus_occipitalis                   ----------------------------------------------------------------------------------------------------------ALAAVAAPAAAAAA
Acheta_domesticus                          GRADGRTGGRADGRTGGRADGRTGGRADGRTGGRADELTGGRADGRTS*RADELTGGRADGRTS*RADELTGGRADGRTS*RADELTGGRADGRTS*RADELTGGRADGRTS*PAGGRAG
Consensus                                                                                                                                                         *      **   *

Teleogryllus_occipitalis                   A-----------------VAS*--------
Acheta_domesticus                          RRTRRSVHKDVK*SY*QLVRSERRMTHILL
Consensus                                                    * *
```

**Tocc.1G0007340.1**

```
Teleogryllus_occipitalis       TCTCATTTTCGCTCTACATAAAAAAATTTCGAAAATTGCTTAAGTTATTGATAATGGGAGAGATCACTATAGTTTTAAGAGTCAGCCAAATGAGCTCCGTAGAACGATTGCGAGTTGTGG
Gryllus_longicercus            TCTCATTTTCATTGTAGACACAAAATTTTCGAAAATTGAATACGTTAGAGA-ATTGGGCGAGATCACTATATTTTTACGTGCCAGTCAAGTGAGTCCCGTCGAACGACTGCGGGTTGTAG
Consensus                      **********  * ** * * **** ************  ** ****  ** * **** ************ ***** * * *** *** ****  **** ****** **** ***** *

Teleogryllus_occipitalis       AGGTAAAGCGTGTGGCGAGTGATGTCCTGTGTTCGAA---TCTCGACGCGGAG--AAAAAATTCTGATTGAAAACACTTTTGCAGATTCATAAAGTAGTATAATATGATGTAGCTATTAC
Gryllus_longicercus            CGGTAAAGCGCGTGGCGAGTAAAGTCCTGTGCCAGTGTTCTCTCGACGCGGAGGAAAAAAATTCTGGTTTGAAAAACTTTTGCATATTCATGAAGTAATGTAATACCACTTAGACATTAC
Consensus                       ********* ********* * ********   *     *************  *********** **  *** ********* ****** ***** * *****  *  ***  *****

Teleogryllus_occipitalis       AAAAAATTCGAGAATGCCAAACACTTTGACCACACG-TAGAGAAATTACTTGTTTCAATGTCTTTTATGCAGTCGCATGCATTAATGGTTGACGCCCT-TCCCCACCCCCCCCATGGGA
Gryllus_longicercus            AAATAATGTGAGAATGCAAAACTCTTCGGCCACACATTAAAGCAATTGCTTTGTTC-ACGATCTCAGTGTAGTCGTAGGCATTGATGGTTGCGACCCTGTCCCCTCCGCACCCAATGGAA
Consensus                      *** ***  ******** **** *** * ******  ** ** **** ***  *** * *   *   ** ***** * ***** *******   **** ***** ** * *** **** *

Teleogryllus_occipitalis       TTCTTAGCAAAAGTTCACAGACAGTCGCTCTTCTTTCCAGGAGACAAAGTCGAACACAGTTGGCAGACATCCAGCGGGTTGTCGCAGTTCTCATCTGCCGAGCTAACATTTGACGTCGTT
Gryllus_longicercus            TTCGTATCAAAGGTTCGCAAACAGTCGATCTTCTTTCCAGGAGACGTAGTCGAACACCGTTGGCAAACATCCATCGACTTGTCGCAGTTCTCCTCTGCCGAGTTAACAGTTGACGTCGTT
Consensus                      *** ** **** **** ** ******* ****************  ********** ******* ******* **  ************** ********* ***** ***********

Teleogryllus_occipitalis       GAGCGATCCGTGACGGTCGTAGGGTAGAAGTTTCTAAATTCTCTCTGAGTCGTGAGACTGTACCAGTTCGTGCCACGTTGTGGGTGTAAACGAACACTATCATCTGACATCACATTTATA
Gryllus_longicercus            GAGCGATCCGTGACGGTCATGCGGTAGAAGTTTCCAAACTCTCTCTCAGCCGTGAGACTGTACCAGTTCGTTCCACGTCGTGGGTGTTAACAAACACTATCTTCTGACGTCACATTTATA
Consensus                      ****************** *  ************ *** ******* ** ******************** ****** ******** *** ********* ****** ***********

Teleogryllus_occipitalis       TCGCCGTTTTCTCCAGCGA---------------------------------TTAATGCCAGCGATTAATGAAAATGCACCAGAGTGCAGATACAGACCCACATGCAATTCTCTTTGTA
Gryllus_longicercus            TCGCCGTTCTCCCAGCTCAAACTGTTCCTCTGTACTCAATTCTACCTTCCAATTTGGCGTCAGCGGTTAATGAAAATGCGCTTAAGCGCAGGCACAGACACACATGCAACTCTCTTTATA
Consensus                      ******** ** *     *                                 **   * ***** ************* *   ** ****  ****** ********* ******* **

Teleogryllus_occipitalis       GTCACCCAGTCACTTTCGTTTCAAAAACGCCGAAA-AAATACAAGTAAGAACTCAATTTACTTTAATTTCCATTACAAATCGCAAAAACCCTAAGTGACAGAAAAACTGTTTCCATTTAC
Gryllus_longicercus            TTCGCCCAGTCATTTCGCTTTCAAAAACACGGAAATAGATACAATTA--GACTCTATTTACCTTAATGTCAATTACAAATTTCGAAAACCATATATAACAG-AAACCAAATTACATTTAC
Consensus                       ** ******** **   ********** * **** * ****** **   **** ****** ***** ** *********  * ****** **  * **** *** *   ** *******

Teleogryllus_occipitalis       TTTAAGGAAGAAATCTGTGGAAGGAAGAAAACTAGTAAAGCTGCAACAGCACATACTTTCTCGCTTGTTTATAACTGCTTCTGTGTATGGTAGTAGTAGTAGTAGTAGCAGTAGTAGCAG
Gryllus_longicercus            TTCAAGAAAGAAATCTGTGGAAGAAAGAAAACTAGTAATGCTACAACAGCACTTATTTTCTCGCTTCTTTATAACTAATCCACTGTATTATAGTACCATATAAACCTGCAAAAGTGTTTC
Consensus                      ** *** **************** ************** *** ********* ** ********** ********* *********  * *  *****  *****  *     *   ***  ***

Teleogryllus_occipitalis       CAGCAGCAGTAGTAGTATTAGTAGTAGTAGTAGTAGTAGTAGTAGTAGTAGT---------------------------------AGTAGTAGTAG------------------------
Gryllus_longicercus            CAACCAGCAAAATGTTTTTTCTCGCCGAGACACGAAACAAAGTCCTAAGAGTTTACTCGCCATGAGCCTTCCCTACATGCTATCGAATAGCAGTGAATTTCAAACTCGGGCTTGAAGTTT
Consensus                      ** *      * *  * **  * *  *    *  *     ***  **  ***                                 * *** ***

Teleogryllus_occipitalis       ------------------------------------------TAGTAGTAGTAGTAGC-------------------------AGTAGTAGCAGCAGC------------------AGCA
Gryllus_longicercus            TGAAATATTATTTTTTTGAATTTTCGAAATACGGAGAGTTTGCCAAAAATGAGAATGGCGCCAAGTTTTTACCGTCCTTTGTTAAACACGAGGAGCAGCTTTAAGGAAAAACTATCGAGCG
Consensus                                                                   *  * *   * * **                         *  *  ** ******                 ***

Teleogryllus_occipitalis       GTAGTAGTAGTAGTAGT--------AGTAGTAGTAGTAGTAG-----------------------------------------------------------TAGTATTAGAAGAAGTA
Gryllus_longicercus            GTTTTTGAGAAAATAATTTTTTTATGAACTGCATAAATTTCAGACAAATTACGCACAATCTACTCTTCTTTCAAACCCTTTTACATGACCTTCTTTAAAGAGCTTATTTTTAGAACGGACT
Consensus                      **  * *    * ** *        *   * *  * *   **                                                           ** * ******

Teleogryllus_occipitalis       GTAGTAGAAGTAGTAGT------------------------------------------------AGTAGTAGTAGTAGTAGTAGTAGTAGTAGTAGTAGTAGTAGTAGTAGTAGTAGTA
Gryllus_longicercus            TTGTTTAAGACAGAAATTTCTAAATTAATTTTGAAGTTTTGGAATGTCAACTCTTTGCAATTTTCGACAATGAGACTAGCACTTTTTACCGTGATACATACGAGGAGATAATTTAAAGAA
Consensus                       *  *  *   ** * *                                                  * *   * *** * *  *    **  **           *  * *   ** *

Teleogryllus_occipitalis       GTAGTAGTAGTAGTAGTATTAGAAGTAGTAGTAGTAGTAGTAGTAGTAAT---------------------------------------AGTAGTAGTAGTAGTAGTAGTAGTAGCAGCA
Gryllus_longicercus            AAAATTGGAACAGTTTTTTGAGAAGAAATTGTTTAAACA--AGTTAAAATTTCCTGAACCATTTTCTTAGCCCTCTTTTATGCTCTCCAAGGATGAAGTTTAATCAAAAATTTGGCGTCA
Consensus                        * * *  ***  * * ***** * * **   *  *  ***   ***                                       ** *  *    ** *   *    * **  **

Teleogryllus_occipitalis       GTAGTAGGAG---------------TAGCAGTAGCAGTAGCAGTAGCAGCAGCAGTAGCAGCAGTAGCAGCAGTAGCAGCAGTAGCAGTAGCAGTAGCAGTAGCAGTAGCAGGAGCAGTAGC
Gryllus_longicercus            TTTTTGAGAAAATCCTGAAAATATTTTTGAGGCTTCATGCAAAGTAGCCTAAAAAATCAAATTCCATTTGAAACTCCCAAAGCCTTGG-----------GTTGTTTCAGATCGATCGGG
Consensus                       *  *  **                 * *  *   **     ******   *  *     * *  **   * * *  *   ***  * *           *  **  ***    *   *

Teleogryllus_occipitalis       AGTAGCAGTAGCAGTAGCAGCAGCAGCAGCAGCAGCAGCAGTAGCAGCAGCAGTAATAGTAGTAGTAGAAGTGTAGTAGTGTAGTATCACTACTACTACCA----CCACCACCACTACTA
Gryllus_longicercus            GATGTCAAT----TTTGCAGAAAATCGACTGCTAGCGTTGATCACAGACGGACGAACGGACATGCTGATGTTTTGAGAAAATTCTATTTAGAATTCTATCAAATTCTGTCAGCCTTCATG
Consensus                        *  ** *     * **** *     *     ***     *  ***  * *  **  *   *  *     * *   *   *  ***    * * *** **    *   ** *  *  *

Teleogryllus_occipitalis       CTACTACTACTACTACTACTACTACTACTACTACTACTACTACTACTACCACCACTACTACGACCACTACTACTACTACT-----------ACTACCACCACTACTACGACCACTACTACTA
Gryllus_longicercus            CAGCTTCAGAAACGAAAATGACTATT-TTTCAAATATTGCTAATCTTGGGATTTTTAAAAT-----TTATTTTTATTTATTTATGTTAAACTTTTACTCATCTTTCAACTATT-------
Consensus                      *  ** *    ** *  *  **** *  * * * ** * *** *      * *  **  *       ** *  ** *            ***   **   *  * * ** * *

Teleogryllus_occipitalis       CCACTACTACTACTACCACTACTACGACCACTACTACTACCACCACTACGACCAC---TACTACTACCACTACTACGACCACTACTACTACCACTACTACTACCACAACTACGACCACTA
Gryllus_longicercus            ----TATAGTTAAAATTACGGGAAGGGATTCTGGAACT--CAAGAAAACGTCCGCAGATAATTTTTTCCCCTGTACGTCACATACAGTTTTTACTTTTTTTTGACTGAATAAATATAAAT
Consensus                          **    **  *  **    * *    **   ***  **  *  *** ** *   ** *  *  * *   **** *   ***   *   ***  *  *      * **      *

Teleogryllus_occipitalis       CTACTACCACTACTACCACCACTACTACCACTACCGCTGCCAC
Gryllus_longicercus            CTACTTCTCTA--------------TACTACCTAAG----
Consensus                      *****  * ***              ******   *
```

**Tocc.1G0007340.1_exon1**

```
Teleogryllus_occipitalis     ------------------------------------------------------------------------------------------------------------------------
Gryllus_longicercus          KFTSLKRSFTYTVYTIFLRSRKMTKKHKHSWISLLCHYCLRLMQPFYGVTLCEV*VTFFCSKTFISVFVTT*NRQSLFLNFVYCFKFINFVKNKQVIYKTLPKVAQSNPVLQHSFPQIEK
Consensus

Teleogryllus_occipitalis     ------------------------------------------------------------------------------------------------------------------------
Gryllus_longicercus          YTT*LRTPFITNRENVYRLWQYLCELLHNITALAPGNDLLDYE*CDELSLAKNL*HWN*YLTDETDEIRDDRKLR*NVFAQFRMKYWQILNIFLEEQLAFLFLFSPLIYQMTVKNCRSSQ
Consensus

Teleogryllus_occipitalis     ------------------------------------------------------------------------------------------------------------------------
Gryllus_longicercus          LGRCLKT*FPHPLFLTCNFREFRYFNENEFSF*ASLVFSSTKFKENN*SCF*YDVFNNCINFKHKLQNIFYKNMYITYLKYLFF*TDFSTKISIFHFVVSNIFLTAPNSFLHFLLRIEQR
Consensus

Teleogryllus_occipitalis     ------------------------------------------------------------------------------------------------------------------------
Gryllus_longicercus          TIYKNGASLIFIVDTKFSKIEYVRELGEITIFLRASQVSPVERLRVVAVKRVASKVLCQCSLDAEEKNSGLKNFCIFMK*CNTT*TLQIM*ECKTLRPHIKAIALFTISV*S*ALMVATL
Consensus

Teleogryllus_occipitalis     ------------------------------------------------------------------------------------------------------------------------
Gryllus_longicercus          SPPHPMEFVSKVRKQSIFFPGDVVEHRWQTSIDLSQFSSAELTVDVVERSVTVMR*KFPNSLSAVRLYQFVPRRGC*QTLSSDVTFISPFSQLKLFLCTQFYLPIWRQRLMKMRLSAGTD
Consensus

Teleogryllus_occipitalis     ------------------------------------------------------------------------------------------------------------------------
Gryllus_longicercus          THATLFIFAQSFRFQKHGNRYN*TLFTLMSITNFENHI*QKPNYIYFKKEICGRKKTSNATTALIFSLLYN*STVL*YHINLQKCFQPAKCFFSPRHETKS*EFTRHEPSLHAIE*Q*IS
Consensus

Teleogryllus_occipitalis     ------------------------------------------------------------------------------------------------------------------------
Gryllus_longicercus          NSGLKF*NIIF*IFEIRRVCQK*EWRQVFTVLC*TRGAALRKNYRAVFEKIIFYELHKFQTNYAQSTLLSNPFT*PSLKSLFLERTLFKTEISKLILKFWNVNSLQFSTMRLALFTVIHT
Consensus

Teleogryllus_occipitalis     ------------------------------------------------------------------------------------------------------------------------
Gryllus_longicercus          RR*FKEKIGTVF*EEIV*TS*NFLNHFLSPLLCSPRMKFNQKFGVIFEKILKIFLRLHAK*PKKSKFHLKLPKPWVVSDRSGMSILQKIDC*R*SQTDERTC*CFEKILFRILSNSVSLH
Consensus

Teleogryllus_occipitalis     ------M--------------------------------------------------------------------------
Gryllus_longicercus          AASETKMTIFQILLILGFLKFIFIYLC*TFTHLSTIYS*NYGKGFWNSRKRPQIIFSPVRHIQFLLFFD*ININLLSLYYL
Consensus                          *
```

**Tocc.1G0007340.1_exon2**

```
Teleogryllus_occipitalis                 ------------------------------------------------------------------------------------------------------------------------
Gryllus_longicercus                      SLPV*NGVLHIQYIPFFSVQGR*RKNTNIRGFLYFVTTAYALCSHSMG*LCVKFR*HFFVQRHLFQYLSPHKIDKACF*ISFTALNLLIS*KINKLFTKLFQKWHKATPFYSIRFLRSKN
Consensus                                

Teleogryllus_occipitalis                 -------HS---------------------------------------------------------------------------------------------------------------
Gryllus_longicercus                      TLHNFERHS*QIVRMSIVYGNICVNFCIISRPWHLETICSIMNDVMSYHWQRICSIGIDT*PMKQTKLETTAN*DETFLHNFA*NIGKF*TFFSKNSWHFYFYFLLLSIK*R*KIVVRPN
Consensus                                       **

Teleogryllus_occipitalis                 ---------------------SSSSSSSSS-------------------------------SSSS-------------------------------------------------------
Gryllus_longicercus                      WVDA*KPNSRTHYF*HVIFGNSDTSTKMSSPFKRPLFFPRQNLKKTIKAVFDMMFLIIALISSTSYKTFFTKICT*LILNICFFEQILVQKFLFFIL*FRTFS*RLQIRSCISFYG*SKE
Consensus                                                     *  *   **                               ** *

Teleogryllus_occipitalis                 -------------------------------------------------------------SIRR-------------------------------------------------------
Gryllus_longicercus                      RFTKMALVSFSL*TQNFRKLNTLENWARSLYFYVPVK*VPSNDCGL*R*SAWRVKSCASVLSTRRKKILV*KTFAYS*SNVIPLRHYK*CENAKLFGHTLKQLLCSRSQCSRRH*WLRPC
Consensus                                                                                             * **

Teleogryllus_occipitalis                 ------------SSSRSSSSSSSSSSSSSSSSSSSSS-------SSSSSSIRS--------------SSS-------------------SSSSNSSSSSSSS------------------
Gryllus_longicercus                      PLRTQWNSYQRFANSRSSFQET*SNTVGKHPSTCRSSPLPS*QLTSLSDP*RSCGRSFQTLSQP*DCTSSFHVVGVNKHYLLTSHLYRRSPSSNCSSVLNSTFQFGVSG**KCA*AQAQT
Consensus                                              ****     *       *   **        * *   **               **                   * *** **   *

Teleogryllus_occipitalis                 -------------------------------------------SSSSRSSS-------------------------SSSSSSSSSSSSSSSSSSSSSSSSSRS----------SSSSSSS
Gryllus_longicercus                      HMQLSLYSPSHFAFKNTEIDTIRLYLP*CQLQISKTIYNRNQITFTSRKKSVEERKLVMLQQHLFSRFFITNPLYYSTI*TCKSVSNQQNVFSRRDTKQSPKSLLAMSLPYMLSNSSEFQ
Consensus                                                                              **  *                         *      * *      *      *  *          * **

Teleogryllus_occipitalis                 SSSSS-------------------------------------------------------------------------------------------------------------------
Gryllus_longicercus                      TRA*SFEILFFEFSKYGEFAKNENGAKFLPSFVKHEEQL*GKTIERFLRK*FFMNCINFRQITHNLLFFQTLLHDLL*RAYF*NGLCLRQKFLN*F*SFGMSTLCNFRQ*D*HFLP*YIR
Consensus                                    *

Teleogryllus_occipitalis                 -----------------------------------------------------------SSSSSSSSNSSSRSVVV*-------------------------------------------
Gryllus_longicercus                      GDNLKKKLEQFFEKKLFKQVKIS*TIFLALFYALQG*SLIKNLASFLRKS*KYF*GFMQSSLKNPNSI*NSQSLGLFQIDRGCQFCRKSTASVDHRRTNGHADVLRKFYLEFYQILSAFM
Consensus                                                                                           **     *   * *

Teleogryllus_occipitalis                 ---------------------------------------------------------------------------------
Gryllus_longicercus                      QLQKRK*LFFKYC*SWDF*NLFLEIYVKLLLIFQLFIVKITGRDSGTQENVRR*FFPLYVTYSFYFFLTE*I*IYFLYTT*
Consensus                                
```

**Tocc.1G0017610.1**

```
Teleogryllus_occipitalis        TTAGTTACAAT-TCGTGT--------GCTGGAAT---GCAGTTAACCTTGTTCCTTAATCTTGCTACTTGCAATTTTCTATT--ATTATAAGAACACA-----GATTTGGAACTAGAGAA
Tarbinskiellus_portentosus      TTAATTATAATGACGTGT------GTGGTGGTATTTAGGCGGTCACCCTACCGGATAAACCATCTACTTGCTGTTCACAGTTCGATCAGAACATCGGA--AATTGTTTGGTTCAAGAGGA
Acheta_domesticus               TTAGTTTCAAT-TCGTGTTTGTATATGCTGGAAT---GAAGCTAACCTTCTTCCTTAATTTCGCTACTTGCGAATTTCTATT--ATTATAAGAACCCAAAGAAGATTTGGTAGTAGAAGA
Gryllus_bimaculatus             TTAGTTTCAAT-TCGTGTTTGAATGAGCTGGAAT---GAAATATACCTTGCTCCTTAATCTTGCTACTCACAAATTTCTATT--ATTATAATAACACA---GAAGTTTAGTACAAGACGA
Consensus                       *** **  ***  *****        * *** **   *      *** *      ***     *****  *   *  *  **  ** * ** * *  *       *** *    ***  *

Teleogryllus_occipitalis        TACGTGCGCGGAAATGTAGCTCTTGCT-TCACTCCTGTTAACACTCGTGATCTCTAACGTGCGTGCAATATGGGTTTTATACTTGTCGTGCATATTTTTATTCAAAAATCGCCGTGAAGT
Tarbinskiellus_portentosus      ------CGCGGAAATGCAGCTGTTGCCGTTATTCTTGTAAACGCCCGTGATCTCAAACTTGCGTGCAATATGGGATTTTTACCCCACCTGCATATTTTCGCTAAAAACGTGCTGTGAATT
Acheta_domesticus               TTGGTGCGCGGAAATAATGCTCTTG------CTCCTGTTAACACTCGTGATCTCTAATGTGCGTGCAATATGGGTTTTATATTCCTCGTGCATATTTTTGTTGAAAAATCGCCGTGAAGT
Gryllus_bimaculatus             TACGTGCGCGGAAATGTTGCTCTTGCT-CTACTCCTATTAACCCTCGAGATCTCTAATGTGCGTGCAATATGGGCTTTATACTCGCCGTGCATATTTTTGTAGAAAAATCGCCGTGAAGT
Consensus                             *********   *** ***       ** * * *** * ** ****** **  *************** *** **     * **********     ****   ** ***** *

Teleogryllus_occipitalis        TTGCGTATAATGGAGAAAGATCACGAATTTGTTGCGGCATTTGTGAGCGTCGGCCGTCTCATAATCATGCAATCGAGCAAATGTGCGGTTGTCAGCAAATAGACAATTATGAGAAAAGGC
Tarbinskiellus_portentosus      TTGCGTATAAT----AAAGATCACCAATTTGTTGTGGCTTTTGTAAGCGCCGGCCGACTCCTAATCATACAATCGAGGAAATGTGTGGTTGTCAACAAATAGACAATTATGAGGAAAAGC
Acheta_domesticus               TTGCGTATAATGGAGAAAGATCACGAATTTGTTGCGGCATTTGTGAGCGTTGGCCGTCTCATAATCATACAATCGAGCAAATGTGCGGTTGTCAGCAAATAGACAATTATGAGGAAAGGC
Gryllus_bimaculatus             TTGCGTATAATGGCGAAAGATCACGAATTTGTTGCGGCATTTGTGAGCGTCGGCCGTCTCATAATCATACAATCGAGCAAATGTGCGGTTGTCAGCAAATAGACAATTATGAGGAAAGGC
Consensus                       ***********    ********* ********* *** ***** ****  ***** *** ******* ******** ******* ******** *****************  *** **

Teleogryllus_occipitalis        GATATATACTGGTACATACTAAAA-TTTTTTTGCACGCGTAAGCTTGTAATAAGCCGTGTTTGAATCAGATATATATCTGAGCACGCTGTTACAATAATCGCGCGTGAATCAGATATCTA
Tarbinskiellus_portentosus      GATATATACTGGTACATTGTAAAACATTTTTTGCATGCGTGAGCTTGTAATAAGCCATGTTTGAATCAGATATCTATCTGCGCACGCTGTTACAATAATCGCGCGTGAATCAGATATCTA
Acheta_domesticus               GATATATACTGGTACATAGTAAA--ATTTTTTGCACGCGTAAGCTTGTAATAAGCCGTGTTTGAATCAGATATATATCTGCGCACGCTGTTACAATAATCGCGCGTGAATCAGATATCTA
Gryllus_bimaculatus             GATATATACTGGTACATAGTAAAG-GTTTTTTGCACGCGTAAGCTTGTAATAAGCCGTGTTTGAATCAGATATATATCTGCGCACGCTGTTACAATAATCGCGCGTGAATCAGATATCTA
Consensus                       *****************  ****   ********* **** *************** **************** ****** ***************************************

Teleogryllus_occipitalis        TCTCCATACAGAGTTCCCGTGCGCCACCACTCTTTCCCTTCCCGACGTGGGGCTGGAACCGTGACCAAACAGGGGCCGCCGCCGCGGGCATAGCGGCGTCATAATGCGCTCTTTGTAGCG
Tarbinskiellus_portentosus      TGTCCATACAGAGTTCCCGTGCGCCACTACTTTCT--TTCCCCGACGCGGAGCTGGAGCCGTGACCAAACAGGGGCCGTCGCCGCAGGCAGCGCGGTGTCATAATGCGCTCTTTGTAGCG
Acheta_domesticus               TCTCCATACAGAGTTCCCGTGCGCCACCACTCTTTCCCCTCCCGTCGTGGGGCTGGAACCGTGACCAAACAGGGGCCGCCGCCGCGGGCACGGCGGCGTCATAATGCGCTCTTTGTAGCG
Gryllus_bimaculatus             TCTCCATACAGAGTTCCCGTGCGCCACCACTCTTT-CCCTCCCGACGTGGAGCTGGAACCGTGACCAAACAGGGGCCGCCGCTGCGGGCATGGCGGCGTCATAATGCGCTCTTTGTAGCG
Consensus                       * ************************* *** * *     **** ** ** ****** ******************** *** ** ****  **** ***********************

Teleogryllus_occipitalis        CACGAGCCTCCACAGCCGCTGCCGTCGCCGTCGCCGTCGCCACCGTCGCCGTCGCCACTGTCGTCGTCGCCACCGTCGTCGTCGCCACCGTCGTCGTCGTCCCCGCCCCCGTCCCCGTCC
Tarbinskiellus_portentosus      CCCGAGCCT---------------------------------------------CCGCCCCCGCCCTCCCCACCGCCACCGCCACAACCGGCGTTGT-----------------------
Acheta_domesticus               CACGAGCCTCCACCGCCGGTGCCGCCGTCGTCGCAGCCACCGTCGCCGCCACCGCCACCGACGCCGTCGCCACTGTCGCCGTCGCCACCGTCGCCGT-----------------------
Gryllus_bimaculatus             CACGAGCCT------------------------------CCGCCGCCGCCGCCGCCGCCGCCGCCTCCGCCTCCGCCGCCGTTGCCGTTGCCGCTGT-----------------------
Consensus                       * *******                                             ** *   ** *  * ** * * *  **   *    * **  **

Teleogryllus_occipitalis        CCGTCCCCGTCCCCGTCCCCGTCCCCGTCCCCGTCCCCGTCCCCGTTCCCGTCCCCGTCCCCGTCCCTGTCCCCGTCCCCGTCCCCGTCCCCGTCCCCGTCCCCGTCCCCGTCCCCGTCC
Tarbinskiellus_portentosus      ------------------------------------------------------------------------------------------------------------------------
Acheta_domesticus               ------------------------------------------------------------------------------------------------------------------------
Gryllus_bimaculatus             ------------------------------------------------------------------------------------------------------------------------
Consensus

Teleogryllus_occipitalis        CCGTCCCCGTCCCCGTCCCCGTCCCCGTCCCCGTCCCCGTCCCCGTCCCCGTCCCCGTCCCTGTCCCTGTCCCTGTCCCTGTCCCTGTCCCTGTACCCGTCCCTGTACCCGTCCCCGTCC
Tarbinskiellus_portentosus      ------------------------------------------------------------------------------------------------------------------------
Acheta_domesticus               ------------------------------------------------------------------------------------------------------------------------
Gryllus_bimaculatus             ------------------------------------------------------------------------------------------------------------------------
Consensus

Teleogryllus_occipitalis        CCGTCCCCGTCCCCGTCCCCGTCCCCGTTCCCGTTCCCGTTCCCGTTCCAGTTCCCGTTCCCGGTCCCGTCCCCGTAGACGTCGACGTCGACGTCGCCGTTGTCTGGGTCGACGACGGTA
Tarbinskiellus_portentosus      ------------------------------------------------------------------------------------------------------------------------
Acheta_domesticus               -------------------------------------------------------------------------------CGTCGTCGTTGTCGTCGCCGTTGTTTTAGTCGACGGTGGCT
Gryllus_bimaculatus             -------------------------------------------------------------------------------TGTCGTCGTCCTCGTCGTCGTTGTCTCGGTCGGCGGCGG--
Consensus

Teleogryllus_occipitalis        TTGGCGACGGCTACGGCAACAGCAACAGCGGCGACGG------------------------------------------------CAACAGCGACAGCAGCAGCGACTACGGTGATAGCG
Tarbinskiellus_portentosus      -------------------------------------------------------------------------------------CGGCGGCAGCGGCGGCGACGCCAACAGAGCTGGCG
Acheta_domesticus               ACGGCGACGGCTACGACGACAGCGACGGCTACGGCAACGGCTATGACGACAGCGAAGGCTACGGCGATGGCGCCGGCTAAGGCAACAGCGACTATGGCGACAGCGACTATGGCGACAGCG
Gryllus_bimaculatus             -------------------------------------------------------------------------------------AAACGGCTACAGCGACAGCGACTTCGGCA------
Consensus                                                                                               *  *    **  *  ** *    *

Teleogryllus_occipitalis        GCGGCA--------------ACAGCAGTAGTAAATGCGACTGCAAC-TGAGACAGTGGCAGGTACCCGCGAGCAGCTCGTGTAGCGA-CGGAGACGGTTACGGTGATGATGGCGAAAGTG
Tarbinskiellus_portentosus      --------------------ACGGCTTCAGCCACAGCGGCTGCTGC-AGAGAC--------GCACCCTTCAGCATCTCTTGCAGCGG-CGGAAACGGTCGTGGTGACGATGGTGATGGCA
Acheta_domesticus               ACTACAGTGACAGCGACGGCACAGCAGTAGCAAATGCGACTGCCAG-AGAGGTCCCGGCAGGCACCCGCGAGCAGCCTGTGTAGCAA-GGGAGACGGTTGCGATGATGATGGCGAAGGTG
Gryllus_bimaculatus             --------------------ACAGCAGTATCAAACGCGACTGCAAACAGAGGCATTGGCAGGCACCCGCGAGCAGCTCCTGTAGCGACGGGAGGCTGTTGAAGTGATGATGGCGAAGGTG
Consensus                                           ** **   *   *  *** ****     ***          * ****   **** *   ** ***    ***  * **     *** ***** **  *

Teleogryllus_occipitalis        AGGTCTACAATGACGATAGCACAGTCCGCCCCCTCACGTGCAGCTTGGAAGA-GGGAGGGGGGCGAAATGTACGGTA-CCGCGCCGCCGGAAAGGAATTAACGTAAAGGGTGGCGGAGCG
Tarbinskiellus_portentosus      GGG------ACGACGGGAGCGTGGTCTGCCCCGTCACGTTTGGCTCAGAA---GGGGGCGTTGCGGAACGTACGGCA------GCGCCGGAAAGGAATTAACGTAAAGGGTAGCGAAGCG
Acheta_domesticus               AGGTCTAGAATGACGATAGCACGGTCCG-CCCCTCACGTGCAGCTTGGAAGA-GGGGGAGTAGCGAAATGTACGGTACCCGCGCCGCCGGAAAGGAATTAACGTAAAGGGTGGCGGAGCG
Gryllus_bimaculatus             AGGTGTAGAATGACGATAGCACGGTCCGCCCCCTCTCGCGCAGCTTGGAAGAGGGGGGAGTGGTGAAATGTACGGTA-CCGCGCCGCCGGAAAGGAATTAACGTAAAGGGTGGCGGAGCG
Consensus                        **      * ****  ***   *** * *** ** **    ***  ***   *** * *  * * ** ****** *       *************************** *** ****

Teleogryllus_occipitalis        GCATGGGGAGCTGCGTGGAGAGCGTGGGGCTTTTGGCATACGTGCACGCGTACCGCACGCTCACGCTCCATCCCCTCGCTCTCGTTGCTGCTGAAATTTCGCTCCTTCATGCG-------
Tarbinskiellus_portentosus      GTGTGGAGAGCTGTATAGAGAGCGTGGGGCATTTGGAACACGTGCACGCGCTCTGTACGTCC---CGCCACCCCCTCGTCGTCGTTGCCAGCGTTTCCCTCCTTCCGCATGCGAAATCCC
Acheta_domesticus               GCGTGGGGAGCTGCGTGGAGAGCGTGGGGCGTTTGGCATACGTGCACGCGCACCGCACGCTCACACTCC-GCCCCTCGTTCTCGTTGCTGCTGAAATTTCCCTCCTTCATGCGAAATGCA
Gryllus_bimaculatus             ------------GCGTGGAGAGCGTGGGGCTTTTGGCATACGTGCGCGCGCACCCTCCAT-----------CCCCTCGTTCTCGTTGCTGCTGAAATGTCCTTCCTTCATGCGAAATGCA
Consensus                                   *  * ************* ***** * ****** ****  *    *             *******   *******    *         * *  ******

Teleogryllus_occipitalis        ------------------------------------------------------------------------------------------------------------------------
Tarbinskiellus_portentosus      AGGGCTACGTTTCGTTCCCTAGCAA---CCGCTCAACGCCGATAGCCGATTTCTTACA-------------CCGTCAAGGTACGACTGTATTATCGGTCTTTATGATTACGCAC-----T
Acheta_domesticus               GCGGCTGGCTTGGAGTTCGCTGCCACACCGTTCCCACATTCATGGACTATCACTTATATCACCCATGTCTCCCGACATTATATTCATTCACTCCGAATTATTATAATCACAGACCTTCAT
Gryllus_bimaculatus             GCGGCTGGTTTGGAGTTCTCTGCCACG-CGTTCCCACATTCATGGACTATCACTTGTAGCGCCCATGCCACCCGTCATTATAATAATTCACCCTTAGTTATTATAATCACAAACCTTCAT
Consensus

Teleogryllus_occipitalis        ------------------------------------------------------------------------------------------------------------------------
Tarbinskiellus_portentosus      GCATACTTACGCACGCATTGCTTTGAGTCAACCCAGTCTTATTACATCAATATTTGCATCCTCTGTGTGATGAACTTTTCCACCATGTTTCGTGTCGCGTTTTCATTCGGCACTTTTATT
Acheta_domesticus               TCTCGTCTCTACTTGCAT--------GCGGACTTTCTCTTATTACCCTC--------CTATTTTGGGTGACAA--TTAACCTCCAGGTTTTCTGTCTCGTTTTCATTCACC---------
Gryllus_bimaculatus             ACTCTTTTACG-GTACAT--------GAAGACTTTTTCTTACTACCCTCA------GCTCTTCTGGGTGATAA--ATAACCAGCAGATTTTTGTCTCGTTTTTCATTCGTC---------
Consensus

Teleogryllus_occipitalis        ------------------------------------------------------------------------------------------------------------------------
Tarbinskiellus_portentosus      GCAATGGCAACCCAAGACCACTACTTGCATTTT----------------TCTTGATTGGCACT-------AGACATGTCTGTCACATTTATTT-----AGTCTCCCTTTGCCCTGTACCA
Acheta_domesticus               -CATTGGAAACATTAAGAACCTCCTTCCATTATGATTTCCATTGCAATGTTTTCAATCGCACA-GCATTCAGAAATGCCCGATGTGTTCCTTCGTCTAGGTCC-CGAGTGCTCTGCTCCA
Gryllus_bimaculatus             -CATTGGAAATATTAATAACCTCCTTGCATTTTGATTTCTCTTGC-ATATTTTCAATCGCACATCCATTCAGAAATGTCCGATGTGTTCCTTCGTCTGGGTCCGCGAGGGCTCTGCTCCG
Consensus

Teleogryllus_occipitalis        ------------------------------------------------------------------------------------------------------------------------
Tarbinskiellus_portentosus      CCTAGCCGCCACGCA--CCATCTTCGCTATGAAATCCTCTGGTGGAGCGTTGAAGCG-ACGGTAATGGACGTTTCGGACGGGATGT-GCTTTCGTCGCCGGAGCGAC-----TGTGCGAG
Acheta_domesticus               CCCAGCCCCCAGGCAC-CCGTCTTTGCTCCGCCGCCCCCTCGCGGGGTGTTGAAGCGAACGGTAATGGACGCTTTGGACGGGATGTAGTTTTCGTCGCCGGCGCGAC----TTGCGCCGG
Gryllus_bimaculatus             CCCAGCCCCCAGGCACACCGTCTTTGCTCCGCCGCCCTCTCGCGGGGCGTTGAAGCG-ACGGTAATGGACGCTTTGGACGGGATGTAGTTTTCGTCGCCGGCGCGGCGCGACTGCGCGGG
Consensus

Teleogryllus_occipitalis        ------------------------------------------------------------------------------------------------------------------------
Tarbinskiellus_portentosus      GGCGAAGG-----GCTGGGCTGGGCTGGGGATCTTGTATCGTTCGTCGAC-GAACGCG-TCGCTCCTCCGCCCACAC-----CGCCGCCTCCCGCGTCGTTGTAGCACTGGCTCCTCGAC
Acheta_domesticus               GGGCGAGGG----GCTGGGCTGGGTTGGGGAGCCTGTATCGTTCGTCGACGGCGAACGTTTGCTTCTCCGCCCTCGCCACCGCGCCGCCTCCCGCGTCGTTGTAGCTCTGGCTCCT----
Gryllus_bimaculatus             GGAGAGGGGCTGCGCTGAGATGGGTTGGGGAGCCTGTATCGTTCGTCGAC-GCGAGCG-TCGCTTCCCCGCCCTCGCCACTGCGCCGCCTCCCGCGTCGTTGTAGCCCTGGCTCCTCGAC
Consensus

Teleogryllus_occipitalis        ------------------------------------------------------------------------------------------------------------------------
Tarbinskiellus_portentosus      GACGGCGGGGGCGAGGGCGGAGGCAGAGGCGGGGGTGCCAGCGGAGACAGCAATTAGAGCGGCGGGCAGGGCGGAGGCTGATTTC-AGGGAGGGGTGATCGGAGCGCCGCGGGAAGGTCG
Acheta_domesticus               --CGGCGGCGGCGGGGGCGGG------------------AGCGGAGACAGCAATTAGAGCGGCGCGCAGGGCGGAGGCTGATTTCGGGGGAGGGGTGATCGGCGCGCCGCGGGGAGG---
Gryllus_bimaculatus             GACGACGGCAGCGGGGGCGGG------------------AGCGGAGACAGCAATTAGAGCGGCGCGCAGGGCGGAGGCTGATTTCGGGGGAGGGGTGATCGGCGCGCCGCAGGGAGGTCG
Consensus

Teleogryllus_occipitalis        ------------------------------------------------------------------------------------------------------------------------
Tarbinskiellus_portentosus      TCGGCTCTGCCGGAAGGAAATTACCACCTCCCGGGGGGTGAAGAAGGCGTTGCCGCTGGCGGTGGCATCAGGGATGGCGGCGGTGCAGGCGGT----CCCCGGCCCGTCGTTCCCTCCTT
Acheta_domesticus               ------------------------------------------------------------------------------------------------------------------------
Gryllus_bimaculatus             -AGGCTCTGCCGGAAGGAAATTACCACCTCCCGGGGGGTGAAGAAGGCGTTGCCGCGGGCGGTGGCGTCATGGCTGGCGGCGGCGGCGGCGGTGGCCCCTCTCCCCGTCGTTCCCACCAT
Consensus

Teleogryllus_occipitalis        --------------
Tarbinskiellus_portentosus      CGCAGCAGCCTTC-
Acheta_domesticus               --------------
Gryllus_bimaculatus             GGCAGTCCGCTCCC
Consensus
```

**Tocc.1G0017610.1_exon1**

```
Teleogryllus_occipitalis        -----------------------------------------------------------------------------------------------------------------------
Tarbinskiellus_portentosus      SKFLFWHDCCSETLDPLARRVHFTPICMDLPRVQRRRLRRWRRRWPRLRLRRQPALARQRQISGIK*NRLLHSGAPKLCRFAEYGLMLALIKHFLLLSC----------GCG-----A
Acheta_domesticus               -----------------------------------AVAQWRA----------------------------EVVP----FCGVRFDVSVDK-TFPLT*LRLRRWRP*RRGCGRWRGRGA
Gryllus_bimaculatus             -------------------------------------SAARR----------------------------SCAP----FRGVRLMLALKNISFLLSC----------GCG-----AG
Consensus

Teleogryllus_occipitalis        -----------------------------------------------------------------------------------------------------------------------
Tarbinskiellus_portentosus      GAGVCGGGGGGGWGLGRGR-------WRSVAWRARRSPPPLPPRCLFPLI*LGNSAGRR------GSGVPCTRPGLRGETWPVPTSHPPGI---AAATVPISQLDSWARR---------H
Acheta_domesticus               GAGGDGRGQLLG---GRAEARRYRRDVYSRSFDSEIARGGAGVEF---LVPGRASAGRR------GPPPP-----------PRPTSPAPPR------PAPLRSAPRFSRQIAARLAGSFE
Gryllus_bimaculatus             GAGVAGRRRRGG---GDG--------VRSVCFAGE-PKPAATAEFIPLILTRK*RAGREWKFPVPGPGPPR-------ETWPRPDRGPTPRVARPASPGPALHPP--NRRSARRLFRAFV
Consensus

Teleogryllus_occipitalis        -----------------------------------------------------------------------------------------------------------------------
Tarbinskiellus_portentosus      LSWSDGMGLFGLRKTVQ*LRV---LVTTNAVGV----------YLFCA*--FV----------------DLFVDE*V*VLLYFHPY**Q*CSIGEIGTFTLRRLAAI*REIKYLRYRRN
Acheta_domesticus               HLFSSGG--MAL*RVAH-------NWATSCLPLRDSRGRRYLPSV*CIDL-FLKIVCTYPSVRNLSRPLLSLSIDHIVVSLYERRSY---LC-----GTIT-KKLSLN*R*IKSRHFGRN
Gryllus_bimaculatus             LEWWHG----AVMRCAQLLRHRTCH*ETVCSGV----------SSFCMSCRFI*KLHIVSSESNVSRPPPSVFID---VSLNVWQRY---FC-----STST-SKFGLN*I*IKS*YFRRT
Consensus

Teleogryllus_occipitalis        -----------------------------------------------------------------------------------------------------------------------
Tarbinskiellus_portentosus      I*L---IIMTCVVVFRRSPYRIN----------------------------------HLLAVHSSI--------------------------RTSEIVWFKRTRKCSCCRYSCKRP*S
Acheta_domesticus               VQLENYIRFGAYI*FHNDQKQVNI---------YKSFNSIHNTSVTKSISII------HLVSIRVCICWNEAN-------LLP*FRYLRISIIIRTQRRFGSRRLVRGNNALAPVNTRDL
Gryllus_bimaculatus             GLLEN-CRFVVYTLFINDQKQALI*TFSHITDQHFCFTKLTQTT*THTRHLCNKKC*KY*FSFNSCLNELE*NIPCSLILLLTNFYYYN-----NTEV*YKTIRARKCCSCSTPINPRDL
Consensus

Teleogryllus_occipitalis        ----------------------------------------------------------M----------------------------------------------------------
Tarbinskiellus_portentosus      QTCVQYGIFTPPAYFR*KRAVNF-AYNKDHQFVVAFVSAGRLLIIQSRKCVVVNK*TIMRKSDIYWYIVKHFLHA*ACNKPCLNQI--------SICARCYNNRA*IRYL----------
Acheta_domesticus               *CACNMGFIFLVHIFVEKSP*SLRIMEKDHEFVAAFVSVGRLIIIQSSKCAVVSK*TIMRKGDIYWYIVKFF-----------ARV--------SL**AVFESDIYLRTLLQ*SRVNQIS
Gryllus_bimaculatus             *CACNMGFILAVHIFVEKSP*SLRIMAKDHEFVAAFVSVGRLIIIQSSKCAVVSK*TIMRKGDIYWYIVKVFC----TRKLVISRV*IRYISAHAVTIIARESDIYLHT-----------
Consensus                                                                                 *

Teleogryllus_occipitalis        -----------------------------------------------------------------------------------------------------------------------
Tarbinskiellus_portentosus      -------CPYRVPVRHYFLSPTRSWSRDQTGAVAAGSAVS*CALCSARASAPA---------------LPTATATTGVV----------GGSG------------------------G
Acheta_domesticus               ISIQSSRAPPLFP------LPSWGWNRDQTGAAAAGTAAS*CALCSARASTAGAAVVAATVAATATDAVATVAVATVAVVVVVVAVVLVDGGYGDGYDDSDGYGNGYDDSEGYGDGAG*G
Gryllus_bimaculatus             ----------EFPCATTLSLPTWSWNRDQTGAAAAGMAAS*CALCSARASAAAAAAAAS----------ASAAVAVAAVVVVLVVVVSVGGGNG-------------------------
Consensus

Teleogryllus_occipitalis        -----------------------------------------------------------------------------------------------------------------------
Tarbinskiellus_portentosus      DANRAGDGFSHSGCCRD-------APFSISCSG----------GNGRGDDGDGRDDG--------SVVCPVTFGS----------------EGGVAERTAAPERN*RKG*RS-GVES--C
Acheta_domesticus               NSDYGDSDYGDS----DYSDSDGTAVANATAREVPAGTREQPV*QGRRLR**WRR*GLE*R*HGPPLTCSLEEGE*RNVRYPRRRKGINVKGGGAAWGAA---------WRAWGVWHT-C
Gryllus_bimaculatus             ---YSDSDFGNSSIKRDCKQRHWQAPASSSCSD------------GRLLK**WRR*GVE*R*HGPPPLAQLGRG-----------------GSGEMYGTAPPERN*RKGWRS-GVESVGL
Consensus

Teleogryllus_occipitalis        -----------------------------------------------------------------------------------------------------------------------
Tarbinskiellus_portentosus      IESV-GHLEHVHALCTSRHPLVVVASVSLLPHAKSQ--GYVSFPSNRS----------TPIADFLHRQGTTVLSVFMITHCILTHALL*VNP---VLLHQYLHPLCDELFHHVSCRVFIR
Acheta_domesticus               TRTARSH-SAPRSRCC----------*NFPPSCEMQRLAWSSLPHRSHIHGLSLISPMSPDIIFIHSELL*SQTFILVSTCMRTFSYY------------------PPILGDN*PPGFLS
Gryllus_bimaculatus             LAYVRAHPPSPRSRCC----------*NVLPSCEMQRLVWSSLPRVPTFMDYHL*RPCHPSL**F------------------TLSYYNHKPSYSFTVHEDFFLLPSALLGDK*PADFCL
Consensus

Teleogryllus_occipitalis        -----------------------------------------------------------------------------------------------------------------------
Tarbinskiellus_portentosus      HFYCN-GNPRPLLAF-FLIGTRHVCHIYLVSLCPV-----------PPSRHAPSSL*NPLVER*S-------DGNGRFGRDVLSSP-ERLCEG----EGLGWAGDLVSFVDERVAPPPTP
Acheta_domesticus               RFHSPIGNIKNLLPL*FPL---QCFQSHSIQKCPMCSFV*VPSALLHPAPRHPSLLRRPLAGC*SER*WTLWTGCS------FRR--RRDLRRGRG-AGLGWGACIVRRR-RTF---ASP
Gryllus_bimaculatus             VFHSSIGNINNLLAF*FLL---HIFNRTSIQKCPMCSFVWVREGSAPPSPQAHRLCSAALSRGVEATVMDALDGM*------FSSPARRDCAGERGCAEMGWGACIVRRRERRFPALATA
Consensus

Teleogryllus_occipitalis        ---------------------------------------------------------------------------------------------------
Tarbinskiellus_portentosus      PPPASL*HWLLDDGGGEGGGRGGGASGDSN*SGGQGGG*FQGGVIGAPREGRRL------CRKEITTSRGVKKALPLAVASGMAAVQAV-PGPSFPPSQQP-
Acheta_domesticus               PSPPRRLP------------RRCSSGSSAAAGA--GAETAI------RAARRA---------EADFGGGVIGA-PRG-----------------------
Gryllus_bimaculatus             PPPASL*PWLLDDDG------SGGGSGDSN*SGAQGGG*FRG------RGDRRAAGRSRLCRKEITTSRGVKKALPRAVASWLAAAAAVAPLPVVPTMAVRS
Consensus
```

**Tocc.1G0017610.1_exon2**

```
Teleogryllus_occipitalis      ------------------------------------------------------------------------SAAVAVA-VATVAVATVVVATVVVATVVV-------------------
Tarbinskiellus_portentosus    SKFLFWHDCCSETLDPLARRVHFTPICMDLPRVQRRRLRRWRRRWPRLRLRRQPALARQRQISGIK*NRLLHSGAPKLCRFAEYGLMLALIKHFLLLSC-----------GCG------A
Acheta_domesticus             ------------------------------------AVAQWRA-----------------------------EVVP----FCGVRFDVSVDK-TFPLT*LRLRRWRP*RRGCGRWRGRGA
Gryllus_bimaculatus           --------------------------------------SAARR-----------------------------SCAP----FRGVRLMLALKNISFLLSC-----------GCG-----AG
Consensus                     

Teleogryllus_occipitalis      ------------------------------------VPAPVPVPV-----------------PVPVPVPV-----------PVPVPVPVPV------PVPV-------------------
Tarbinskiellus_portentosus    GAGVCGGGGGGGWGLGRGR-------WRSVAWRARRSPPPLPPRCLFPLI*LGNSAGRR------GSGVPCTRPGLRGETWPVPTSHPPGI---AAATVPISQLDSWARR---------H
Acheta_domesticus             GAGGDGRGQLLG---GRAEARRYRRDVYSRSFDSEIARGGAGVEF---LVPGRASAGRR------GPPPP-----------PRPTSPAPPR------PAPLRSAPRFSRQIAARLAGSFE
Gryllus_bimaculatus           GAGVAGRRRRGG---GDG--------VRSVCFAGE-PKPAATAEFIPLILTRK*RAGREWKFPVPGPGPPR-------ETWPRPDRGPTPRVARPASPGPALHPP--NRRSARRLFRAFV
Consensus                                                                                                    * *               *

Teleogryllus_occipitalis      ------------------------------------------------------------------------------------------------------------------------
Tarbinskiellus_portentosus    LSWSDGMGLFGLRKTVQ*LRV---LVTTNAVGV----------YLFCA*--FV-----------------DLFVDE*V*VLLYFHPY**Q*CSIGEIGTFTLRRLAAI*REIKYLRYRRN
Acheta_domesticus             HLFSSGG--MAL*RVAH-------NWATSCLPLRDSRGRRYLPSV*CIDL-FLKIVCTYPSVRNLSRPLLSLSIDHIVVSLYERRSY---LC-----GTIT-KKLSLN*R*IKSRHFGRN
Gryllus_bimaculatus           LEWWHG----AVMRCAQLLRHRTCH*ETVCSGV----------SSFCMSCRFI*KLHIVSSESNVSRPPPSVFID---VSLNVWQRY---FC-----STST-SKFGLN*I*IKS*YFRRT
Consensus                     

Teleogryllus_occipitalis      --------------------------------------------------------------------------P---------------------------------------------
Tarbinskiellus_portentosus    I*L---IIMTCVVVFRRSPYRIN-----------------------------------HLLAVHSSI---------------------------RTSEIVWFKRTRKCSCCRYSCKRP*S
Acheta_domesticus             VQLENYIRFGAYI*FHNDQKQVNI---------YKSFNSIHNTSVTKSISII------HLVSIRVCICWNEAN-------LLP*FRYLRISIIIRTQRRFGSRRLVRGNNALAPVNTRDL
Gryllus_bimaculatus           GLLEN-CRFVVYTLFINDQKQALI*TFSHITDQHFCFTKLTQTT*THTRHLCNKKC*KY*FSFNSCLNELE*NIPCSLILLLTNFYYYN-----NTEV*YKTIRARKCCSCSTPINPRDL
Consensus                     

Teleogryllus_occipitalis      ------------------------------------------------------------------------------------------------------------------------
Tarbinskiellus_portentosus    QTCVQYGIFTPPAYFR*KRAVNF-AYNKDHQFVVAFVSAGRLLIIQSRKCVVVNK*TIMRKSDIYWYIVKHFLHA*ACNKPCLNQI--------SICARCYNNRA*IRYL----------
Acheta_domesticus             *CACNMGFIFLVHIFVEKSP*SLRIMEKDHEFVAAFVSVGRLIIIQSSKCAVVSK*TIMRKGDIYWYIVKFF-----------ARV--------SL**AVFESDIYLRTLLQ*SRVNQIS
Gryllus_bimaculatus           *CACNMGFILAVHIFVEKSP*SLRIMAKDHEFVAAFVSVGRLIIIQSSKCAVVSK*TIMRKGDIYWYIVKVFC----TRKLVISRV*IRYISAHAVTIIARESDIYLHT-----------
Consensus                     

Teleogryllus_occipitalis      -------VPVPVPVPVPVPVP------------------------------------------------VPVPVPVPVPVPVPVPV----------------------------------
Tarbinskiellus_portentosus    -------CPYRVPVRHYFLSPTRSWSRDQTGAVAAGSAVS*CALCSARASAPA----------------LPTATATTGVV----------GGSG-------------------------G
Acheta_domesticus             ISIQSSRAPPLFP------LPSWGWNRDQTGAAAAGTAAS*CALCSARASTAGAAVVAATVAATATDAVATVAVATVAVVVVVVAVVLVDGGYGDGYDDSDGYGNGYDDSEGYGDGAG*G
Gryllus_bimaculatus           ----------EFPCATTLSLPTWSWNRDQTGAAAAGMAAS*CALCSARASAAAAAAAAS----------ASAAVAVAAVVVVLVVVVSVGGGNG--------------------------
Consensus                                 *       *                                                          *

Teleogryllus_occipitalis      -----------------------------------------------------------------PVPVPV-------------------------------------------------
Tarbinskiellus_portentosus    DANRAGDGFSHSGCCRD-------APFSISCSG----------GNGRGDDGDGRDDG--------SVVCPVTFGS----------------EGGVAERTAAPERN*RKG*RS-GVES--C
Acheta_domesticus             NSDYGDSDYGDS----DYSDSDGTAVANATAREVPAGTREQPV*QGRRLR**WRR*GLE*R*HGPPLTCSLEEGE*RNVRYPRRRKGINVKGGGAAWGAA---------WRAWGVWHT-C
Gryllus_bimaculatus           ---YSDSDFGNSSIKRDCKQRHWQAPASSSCSD-----------GRLLK**WRR*GVE*R*HGPPPLAQLGRG------------------GSGEMYGTAPPERN*RKGWRS-GVESVGL
Consensus                     

Teleogryllus_occipitalis      -------------------------------------------------------------------------------------------P----------------------------
Tarbinskiellus_portentosus    IESV-GHLEHVHALCTSRHPLVVVASVSLLPHAKSQ--GYVSFPSNRS----------TPIADFLHRQGTTVLSVFMITHCILTHALL*VNP---VLLHQYLHPLCDELFHHVSCRVFIR
Acheta_domesticus             TRTARSH-SAPRSRCC----------*NFPPSCEMQRLAWSSLPHRSHIHGLSLISPMSPDIIFIHSELL*SQTFILVSTCMRTFSYY------------------PPILGDN*PPGFLS
Gryllus_bimaculatus           LAYVRAHPPSPRSRCC----------*NVLPSCEMQRLVWSSLPRVPTFMDYHL*RPCHPSL**F------------------TLSYYNHKPSYSFTVHEDFFLLPSALLGDK*PADFCL
Consensus                     

Teleogryllus_occipitalis      ------------VPVPVPV-------------PVPVPVPVPVPVPVPVP---------------------------------VPVP----------------------------VPVPVP
Tarbinskiellus_portentosus    HFYCN-GNPRPLLAF-FLIGTRHVCHIYLVSLCPV-----------PPSRHAPSSL*NPLVER*S-------DGNGRFGRDVLSSP-ERLCEG----EGLGWAGDLVSFVDERVAPPPTP
Acheta_domesticus             RFHSPIGNIKNLLPL*FPL---QCFQSHSIQKCPMCSFV*VPSALLHPAPRHPSLLRRPLAGC*SER*WTLWTGCS------FRR--RRDLRRGRG-AGLGWGACIVRRR-RTF---ASP
Gryllus_bimaculatus           VFHSSIGNINNLLAF*FLL---HIFNRTSIQKCPMCSFVWVREGSAPPSPQAHRLCSAALSRGVEATVMDALDGM*------FSSPARRDCAGERGCAEMGWGACIVRRRERRFPALATA
Consensus                                                      *             *

Teleogryllus_occipitalis      VPVPGPVPVDVDVDVAVVWVDDG-IGDGYGNSNSGDGNSDSSSDYGDSGG------NSSSKCD-----CN*---------------------------------------
Tarbinskiellus_portentosus    PPPASL*HWLLD--------DGGGEGGGRGGGASGDSN*SGGQGGG*FQGGVIGAPREGRRL------CRKEITTSRGVKKALPLAVASGMAAVQAV-PGPSFPPSQQP-
Acheta_domesticus             PSPPRRLP---------------------RRCSSGSSAAAGA--GAETAI------RAARRA---------EADFGGGVIGA-PRG------------------------
Gryllus_bimaculatus           PPPASL*PWLLD--------DDG------SGGGSGDSN*SGAQGGG*FRG------RGDRRAAGRSRLCRKEITTSRGVKKALPRAVASWLAAAAAVAPLPVVPTMAVRS
Consensus                                                      **
```

**Tocc.1G0019510.1**

```
Teleogryllus_occipitalis        CTTTTTTAAAATGACACAG--------------------GAAAAGCAAAGGATATTTCTGTACTATTGATAGCCAAACAATAATCATGATTA----------------------------
Tarbinskiellus_portentosus      CACTTTTGAAAAGTTATAACTGTGTTGAAACCTTCTTGCGGTTAAGGATAGATGTTTAAAAAGAAATTATCTTGAAAAAACGTTCATCATTGTATTACAAATATGCTAAATGCTTTTGAA
Consensus                       *  **** *** *  * *                     *   *   *  *** ***    *  * * **    *** **   **** ***

Teleogryllus_occipitalis        ---------------------------------------------------------AAAGTTAGTAAATAATTGCAAGGACATGGAAGAAACTACTAAATAAAAA--------------
Tarbinskiellus_portentosus      TTGTATTTAATTTGTCTGGATGACAAACGTTTTAAAAGCGGAAATGACTTGGCCTTTATAATTGTGTAATAGATGCGAGTATTTGATAG----TATTAGGTAAGAAGAATTGCGCGAGGA
Consensus                                                                                * * **    ****  *** ** *  **  **    ** **  *** **

Teleogryllus_occipitalis        -----TCAAATATTTT---------------TCCAATGAATTGAA----------ACTGCTCTAG-------------------------------------------------------
Tarbinskiellus_portentosus      ACTTTTGAAAAGTTTTGACTGTGTTGAAACCTCCAATAAATGAGAGTATAATGCTATTATTTTAGGCTTATCTGAGTGGCTCCCTTGAGTAAATTCCATTACGTACGCATCCTGCATTCC
Consensus                            * ***  ****               ****** ***   *          * *  * ***

Teleogryllus_occipitalis        ----------------------TAATTTTAGACATCCAATAATATTAAAAA---------------------------------------------------------------------
Tarbinskiellus_portentosus      TGTAGATTAAAGCGCCGGGCGACAATATTTCAAGTTCAATAGCGGTGAAGAGGAATCACAAGACGTGATCATCAATCGATTACCTATACTCGGGCCCGGTCGCTCATGCCTGCTCTCGCT
Consensus                                              *** **  *  * *****    * ** *

Teleogryllus_occipitalis        ----TCCATCATTTCG----------------------------------------------------------------------ATAAATTGTATCGT--------------------
Tarbinskiellus_portentosus      ACCCTCCATCACCCCGTTGCCGCCTTTGCCTCCACCTCCACCTCCGCGTCCGTCCACCTCTTCGCTTCCTCCGCCGGCTCCCTTGAGTAAATTCCATTACGTACGCATCCTGCATTCCTG
Consensus                           *******   **                                                                     ******  **

Teleogryllus_occipitalis        --GAGAGAAGTATTGCGTGAATGGTGAACAAAGTTATTAGACCCTCTGTTGCAGATGAACAGTTAGATTCGCGCATGAACAGAATATTTTGATGCGGCTCCGTTCGCCCACGCC--CTCT
Tarbinskiellus_portentosus      TAGATTAAAGCGCCGGGCGACAATATTTCAAGTTCAATAG------CGGTGAAGAGGAATCACAAGACGTGATCATCAATCGATTACCTATACTCGGGCCCGGTCGCTCATGCCTGCTCT
Consensus                         **   ***    * * **        ***  * * ***       * ** *** ***     ***   *  *** **  ** **  *  *  ***  *** **** ** ***  ****

Teleogryllus_occipitalis        CCACCGCGTCCATCGCCCCTCCTCCGCGTCCGCCTCCGCTCCCGCCTCCGTTCCCGCCTTCCTAGCCCGCTTCCTCCGCCGTCAGGGGTGTCCACGGCGCCGCGGGCCGCGCTTGCCGTG
Tarbinskiellus_portentosus      CGCTACCCTCCATCACCCCGCTGCCGCCTTCACCTCCACCTCCACCTCCGCGTCCGTCCACCT-CTTCGCTTCCTCCGCCGGCAGGGGTGTCCAGCATGCCGCCGGCCGCACCGATCTCG
Consensus                       *     * ****** **** *  **** * * ***** *  ** ******   *** *  ***    ************** ************    ***** ****** *    *  *

Teleogryllus_occipitalis        CTTGCCCCCCTAGCTGGTTGTATTTTTAGGTTGCCGACGCTGCGGGTGACCTGCAACGTTGACGCATGGTCAGTGACGACAGCCGCGCCGGAGTGTAGACAGTGTACACAGCTGGACGGA
Tarbinskiellus_portentosus      CAT------CTAGCTGGTTGTATTTTTAGGTCGCCGACGCTGCGGGTGACCTGCAACGTTGACGCATGGTCAGTGACGACAGCTGCGCCGGAGTGTAGACAGTGTACACAGCTGGACGGA
Consensus                       * *      ********************** *************************************************** ***********************************

Teleogryllus_occipitalis        TTTAGCGGGCTTCTCTGGGGCTCCGCGCGCGCC---------------------------GGCTCGTCCGTCCGCCCGCGACGCCGC-------------CGCCGCCGCCCCCCGC----
Tarbinskiellus_portentosus      TTTAGCGGGCTTCTCTGGGGCTCCGCGCTCGCCCGCTACCCATTCGCCCCCTCACCCGCTCGCTCGCTCGCTCGCTCGCCCCCTCGCGCCTACGTTCCCTCGCCCTCGTCCCTCGCCCCC
Consensus                       **************************** ****                             *****  **  *** ***  *  ***             ****  ** *** ***

Teleogryllus_occipitalis        ----------------------------------CAGTTACGAAACACGAGGAGGAGGAGGCGTTGGAGGCGATGGAGG-----------CGATGGAGATGGCGACGGC---------GG
Tarbinskiellus_portentosus      TCACCCAAGCCCCCTCGCCTCCTCGCCCGCCCGACGGTTACGAAACAGGAGGAGGCGGAGGTGACGGTGTCGGGGGTGGACAGTAGCGTCGGGCTGAGATACAGGCGACTCACCATGAAC
Consensus                                                         * *********** ******* ***** *  ** * **  ** **           *   *****   * ** *

Teleogryllus_occipitalis        CAAAGAC---------------------------GGCGATTGCGGC-------------GACGGCGACGGCAGCGGCGACAGCG--TCGGAGGCGGGGGCGACCACAGGGATGACGGCGG
Tarbinskiellus_portentosus      CAAATATCTCAATGATCCTCAATCTCGAATGTCAAGCAATCGCTACTTTCAGACAAACAAACAAAAAAGTCCGTGCTAATGATGAATAAAAGACCGGAGTAATATTTATGATAATTTTCT
Consensus                       **** *                             ** ** **  *              **    * * * * *   *    *  *   ** * ** *  *       *** *

Teleogryllus_occipitalis        GAACGGCGACGGAGGCGGCAACGG--AGGTGGAGATGGGG---------------------------------------CGGCGGCGGGGGTGGCGACAGGGACAGCGGCGGGAGCGGCG
Tarbinskiellus_portentosus      AATGAAAGATTGAAGTGTTATTTGTCGGATCGTATTGAGGTTTATTAGTATAAACATTTTGTTGCCATTCGCTACTCGCCATCCGCAAAACACGAGGTAGAAAAGGTGATGGTGCCGGGG
Consensus                        *     **  ** * *  *   *   * * *   ** **                                        *  * **       * *  **  *  * *  **   *** *

Teleogryllus_occipitalis        ACGGGGGGCGGCGACGGGGGCGGCGACGGGGGCGGCGACGGGGGCGGCGACGGGG----TCGGCGACGGGGGCGGCGACGGGGGCGGCGACGGGGGTGGCGACGGGGGCGG---CGGCGG
Tarbinskiellus_portentosus      ATGCATAGCAGCGCCGGGCCGACGTACAGGGAGCTCGTCCTTAACCGGAACTCGAAACTCTAGCAACAGGCGCTCTCCCACAAACGAGAAAATAAATGGTAACGGTAAATAATTCTTTAA
Consensus                       * *    ** *** ****       ** ***    ** *     * *  **  *        ** ** ** **     *     **   *      ***  ****         *

Teleogryllus_occipitalis        GGGCGGCGACGGGGGGCGGCGACGGGGGCGGCGACGGGGGCGGCGACGGGGGCGGCGACGGGGGCGGCGACGGGGG--------------------------------------------
Tarbinskiellus_portentosus      GATCGGGAGTACGGCGTTGTACTTGCGGTCATTTCATAGGAAATGA-------------------AATGTTGTGGGATCATATTGTGTTTTATAAGCTTTCAGTTTTTTTTGACCGCTAT
Consensus                       *  ***      ** *  *     * **      *   **    **                      *  * ***

Teleogryllus_occipitalis        ---------------CGGCGACGGGGGCGGCGACGGGGGGT---------------------------------------------------------------------------
Tarbinskiellus_portentosus      TAGCCATCTGCCATACAATATTAAATGTGGGTTTCAGGGATTTCAATATCTCATAATGCATAAGTCCACTCCACACTTTTAATCTGACGACATATATTACAGAACGCTTGCGACCTA
Consensus                                      *          * * *      *** *
```

**Tocc.1G0019510.1_exon1**

```
Teleogryllus_occipitalis        -------------MRLR-------------------------------------------------------------------------------------------------------
Tarbinskiellus_portentosus      EALLKSYNCVETFLRLRIDV*KEIILKKRSSLYYKYAKCF*IVFNLSG*QTF*KRK*LGLYNCVIDASI**Y*VRRIARGTFEKF*LC*NLQ*MRV*CYYFRLI*VAPLSKFHYVRILHS
Consensus                                      ***

Teleogryllus_occipitalis        ------------------------------------------------------------------------------------------------------------------------
Tarbinskiellus_portentosus      CRLKRRATIFQVQ*R*RGITRRDHQSITYTRARSLMPALATLHHPVAAFASTSTSASVHLFASSAGSLE*IPLRTHPAFL*IKAPGDNISSSIAVKRNHKT*SSIDYLYSGPVAHACSRY
Consensus

Teleogryllus_occipitalis        ------------------------------------------------------------------------------------------------------------------------
Tarbinskiellus_portentosus      PPSPRCRLHLHLHLRVRPPLRFLRRQGCPACRRPHRSRI*LVVFLGRRRCG*PATLTHGQ*RQLRRSVDSVHSWTDLAGFSGAPRSPATHSPPHPLARSLARPLAPTFPRPRPSPPHPSP
Consensus

Teleogryllus_occipitalis        ------------------------------------------------------------------------------------------------------------------------
Tarbinskiellus_portentosus      LASSPARRLRNRRRRR*RCRGWTVASG*DTGDSP*TKYLNDPQSRMSSNRYFQTNKQKSPC***IKDRSNIYDNFLMKD*SVICRIVLRFISINILLPFATRHPQNTR*KR*WCRGCIAA
Consensus

Teleogryllus_occipitalis        ------------------------------------------------------------------------------------------------------------
Tarbinskiellus_portentosus      PGRRTGSSSLTGTRNSSNRRSPTNEKINGNGK*FFKIGSTALYLRSFHRK*NVVGSYCVL*AFSFF*PLLAICHTILNVPFRDFNIS*CISPLHTFNLTTYITERLRP
Consensus
```

**Tocc.1G0019510.1_exon2**

```
Teleogryllus_occipitalis                 -----SW-----------------------------------------------------------------------------------------------------------------
Tarbinskiellus_portentosus               EALLKSYNCVETFLRLRIDV*KEIILKKRSSLYYKYAKCF*IVFNLSG*QTF*KRK*LGLYNCVIDASI**Y*VRRIARGTFEKF*LC*NLQ*MRV*CYYFRLI*VAPLSKFHYVRILHS
Consensus                                     *

Teleogryllus_occipitalis                 ------------------------------------------------------------------------------------------------------------------------
Tarbinskiellus_portentosus               CRLKRRATIFQVQ*R*RGITRRDHQSITYTRARSLMPALATLHHPVAAFASTSTSASVHLFASSAGSLE*IPLRTHPAFL*IKAPGDNISSSIAVKRNHKT*SSIDYLYSGPVAHACSRY
Consensus

Teleogryllus_occipitalis                 ----------------------------------------------RRR----------QRRRLRRR-----------------------------------------------------
Tarbinskiellus_portentosus               PPSPRCRLHLHLHLRVRPPLRFLRRQGCPACRRPHRSRI*LVVFLGRRRCG*PATLTHGQ*RQLRRSVDSVHSWTDLAGFSGAPRSPATHSPPHPLARSLARPLAPTFPRPRPSPPHPSP
Consensus                                                                              ***          * * ***

Teleogryllus_occipitalis                 ------RRQRRQRRRRGRPQG*--------------------------------------------------------------------------------------------------
Tarbinskiellus_portentosus               LASSPARRLRNRRRRR*RCRGWTVASG*DTGDSP*TKYLNDPQSRMSSNRYFQTNKQKSPC***IKDRSNIYDNFLMKD*SVICRIVLRFISINILLPFATRHPQNTR*KR*WCRGCIAA
Consensus                                      ** *  **** *  *

Teleogryllus_occipitalis                 ------------------------------------------------------------------------------------------------------------
Tarbinskiellus_portentosus               PGRRTGSSSLTGTRNSSNRRSPTNEKINGNGK*FFKIGSTALYLRSFHRK*NVVGSYCVL*AFSFF*PLLAICHTILNVPFRDFNIS*CISPLHTFNLTTYITERLRP
Consensus
```

**Tocc.1G0021480.1**

```
Teleogryllus_occipitalis      TATTTTAAACTGACAGGGTCTTG-----------AATTCCTTCAAATCATTCC-------------------------------------------------------------------
Tarbinskiellus_portentosus    TTCTTTGAAATATTAGTATCATGCCACGTGACACCTATTCCTGCAGCACGTTCCTCCTTCCAGTCCCATCCTCCATCGCTAATCGCCCTGCCTCTCGCCCCGCGGAAAAGGTACGTAACA
Consensus                     *  *** ** *   **  ** **            ****** **   * ****

Teleogryllus_occipitalis      AGGAATATGATTTTTTGAGACAAT---------CGGCTTAACGGTTTTGGTGCTAGCGT-----------------TGATCACAGACGGACAGATAGACGTTTTGCCGTCAAACGAT
Tarbinskiellus_portentosus    ATAATTTTGAAATTGCAACACAATAATAAATTCACTATTCAATAGTTATAAGTACAGCATATTAATTAAATAGTTGTAAATATAACAAAATAACACACAAAAATATTGAATAATAACAAA
Consensus                     *  * * ***  **   * *****          *   * **  *** *      *** *                     ** *** *   *** * * *  * ***      *** *

Teleogryllus_occipitalis      TGTTCGGTTTGATTGTTCGGT------------------TCATAAAACGTGGATTCTGAGTAGACGTGTTAGAGGAAGTAATTTAATA-TACATTGGTACAAGATTTATCAATT----
Tarbinskiellus_portentosus    TTAATTGGCTAACTCTTGAATAAATGAAAACTATTTTATAACAATAAAAAGTGAAAGTTAATAAAACATACTAACAAAATTAAATAAAAACTATAAAAAATACAACATGCTTCAGGCGGAT
Consensus                     *     *  * * * **   *                      ****** *** *   * *  * ** *  **    ** *** * ** * ** *    ***** **   ***

Teleogryllus_occipitalis      ----------------------------------TTTGTGTTGATAAGAGTACCTCAATCGTAAAATGGCCGTT----------------------------------------------
Tarbinskiellus_portentosus    GGTAGGCCCACTCTTCCAGTACTCGGATTACCAATCCAAACCAATGACGACACGATGGTGTTATGATGGTTGTTGAAAAATTAATGAATGGAAATGTATTGTTTTTCTTTATGCTTTACC
Consensus                                                     *        ** *    **     *  **  ****  ***

Teleogryllus_occipitalis      ----TCCTGCGCACGTTTTCAAT-----------------------------------ACATTCTCT-------------------------------------------------------
Tarbinskiellus_portentosus    ACTGTCTTCCATATGTTTCCAGTCGTAAATTAAGAACTGAATGTGATGTGGTTCTTTTATCATCCTCTGACACCGGGTAACCTGGAGTGAAGTTATTAGTGTAGGACAAATCTACGCACA
Consensus                         ** * *  * **** ** *                                    *** ****

Teleogryllus_occipitalis      ------------------------------TTTTATGAAAGCGAGAGTCTAAACAATATCGCCAGGGCAAAATAATACAAAATCGATACATAGACAACAG--------------CTGGC
Tarbinskiellus_portentosus    CACATACATACACACACATATGTGTGATAATTTTGCGATACCGAAATCATACATTTAGTAATATATTTAATAAAACATAGATTTAATATTTATGTTGCGAAGAAATTGTCTCTCTCGAGT
Consensus                                                     ****  ** * *** *   ** *     *         ** * ** * * * *  ***  **     *                 *  *

Teleogryllus_occipitalis      TCAATT--------------------------------ACTCCATCAAATT---------------------------------------------------------------AAAG
Tarbinskiellus_portentosus    TCAGTTATTTGCTGCAAAAATTGAGAATAAAATACAAAAATTCTAGTAATTTAAACAGAAGAAATAAACTTCCATGCTCAGAGTTCTGAAGTATCAACAAAAAGTAATGCTTGTAGAAAA
Consensus                     *** **                                * ** *  ** **                                                              ***

Teleogryllus_occipitalis      GATATCAATTTCTC------CACGAACTTTC---------------------------TTCATTTTGACACA---------------------------------------TTCTGTA
Tarbinskiellus_portentosus    GATCTCTATTTCCCTAACCACACGATTTTTTTCATCCAGGAAAAATTTGTTATATATCATCTATTACGACGCAACGTTAGAATTTCTTGGAAACATTTATTTAATACGCACATTCTTCTA
Consensus                     *** ** ***** *      *****  ***                              *  ***  *** **                                         *  * **

Teleogryllus_occipitalis      GTCCGCATCCTCCTGAAACATTATTTCGACGCCTGTTGGCACATCGCCGCTCGGTAGA---CCATTATTATTTTTTCTTTCATTTCTCCTTGTTGCGCCGCCATCATACTACTAACGTTG
Tarbinskiellus_portentosus    GTTTTCGATTTTCTGAAACGCTATTGTGGTACTT-------------CGCTTGGCAGAGATTCTCCGTGAAGTTCTCTTTCTTTTTTCCC-----CACCGGCATCATACCCCCAAAGTT-
Consensus                     **    *     * *******   ****   *    * *              **** ** ***     *     * *   ** ****** *** ***       * *** ********  * ** ***

Teleogryllus_occipitalis      CCGCCCTCCACCGCCACCGCCTCCGCAGTTGCGGCCGTGGCCGCGCCTGGGTGGAGCTGCAAATAACTACAACGTGGCTAATGTTTAGTAGATTTTAATATAGAGCCGCCAGCTGTATCA
Tarbinskiellus_portentosus    ---TCCTCCACTTCTACCGCCTCCGCATCCTCCACCAGAATTGCGCCTGGGCGGAGCTGCAAATAACTACAACGTGGCTAATGTTTAGTAGATTTTAATATAGAGACGCCAGCTGTATCA
Consensus                         *******  * * ************     *  **       ********* ***************************************************** *************

Teleogryllus_occipitalis      TTTTTCAAAACGCGCTTTCGGCCAAGAAACCGGGCGGAAGG---------GAAGGCGAGCGAAGGCCGAGGCGGGGCGAGGCGGGGCGAGGTGGGGTGC-------------------
Tarbinskiellus_portentosus    TTTTTCAAAACGAGCTTTCGGCCAAGAAAGCGGGCGGAAGGCGAGCGGGCAGGGGGCGGGCAGGGGGCGGGCAGGGGGCGGGCAGGGGGCGGGCAGGGGGCGGGCAAGGGGCGGGCAAGGG
Consensus                     ************ *************** ***********          *  **** **   ** ** *  ****** *  ****   **  *** **

Teleogryllus_occipitalis      -------------------GGTGGGGTGGGGTGGGGGAGGGGGC--TGGCTGAATTCCCGCCCCGCGCAC-CCGACAAACGCGCGGAATTCCTGAGCGCGCCCCGCCGCGCTTCGCCG
Tarbinskiellus_portentosus    ACGGGCAAGGGGCGGGCAAGGGGCGGGCAGGGGCGGGGAGGAGAGCGTTGGCGGAATTCCCGCCCCGCGCACGTCGACAAACGCCTGGAATTCCCGTGAGCATCGCGCCTCAC-------
Consensus                                        ** ***  **** ****  * * **  **** ******************  **********  ******** * * **  * **** * *

Teleogryllus_occipitalis      ACGATGAAAGAGCCCCGTGCGCGATGCGT------GATGCGTGAATGGGCGCCAGGCCGGCGCCGCCTCGCACTTTCCTTCATCTTTGGCGACTGGATCATCTTCACCGTCGTCATCTTC
Tarbinskiellus_portentosus    -CGATGAAAGTGTCCTCT-CGCGACGCGTCTCTTCGCTGCTCTTGTTCGTTTCCTTCTTCTCCTTCTTCTCCCTTTCCTCCTCCCCTTGTACTTCTGCC-TCTTCTTCTCCTCTCCCTTC
Consensus                      ********* * **  * ***** ****      * ***     *  *    *   *     *    *  * ** * ******* *  *   * *    *    * *****  *  *     ****

Teleogryllus_occipitalis      GCCGTCGTCATCTTCGCCCT-CGTCATCTTCGCCGTCGTCATCTTCGCCGTCGTCATCTTCGC-CGTCGTCATCTTCGCCGTC---------------------------------GT
Tarbinskiellus_portentosus    TCTATCTTCTTCTTAACCTTGGGTTATTCGCGTCGC-----TCGCTCCCATAATTGTATCAGCGCGCTGGAATTTTTATCATTAGATGGTTTTTAAAAGAATTTTAGAACATTTATCAAT
Consensus                      *  ** ** ****  ** *  ** **   ** **      **    ** *  *   * *  ** **  *  ** **    * *                                   *

Teleogryllus_occipitalis      CATCTTCGCCGTCGTCATCTTCGCCGTCGTCATCTTCGCCGTCGTCATCTTCG---------------CCGTCGTCATCTTCGCCGTCGTCA----TCTTCGCCGTCGTCTTCTTCGCC
Tarbinskiellus_portentosus    GATATTTTACGCTCGTGATTTCATAGGAATTATTTATTTTGATTCCATTTTGGTTTATGTGCTTGGTAACTTTCTTTTTATTTGCCATTTTTATTATTTTCCCTCTTCATTTTCTAATTA
Consensus                      ** **    **        ***    *   * ** *     *     *** ** *                *   ** *   * ** *** *   * *     * * *   * ** * ****

Teleogryllus_occipitalis      GTCGTCTTCTTCGCCGTCGTCTTCTTCGCC--GTCGTCATCTTCGCCGTCGTCATCTTCGCCGTCGTCATCTTCGCCGTCGTCATCATCCGCATCTTCATA------TTTTTCTTCGTCT
Tarbinskiellus_portentosus    TATATTTTTATTATTTATGATTTTTTTCGTCTGGTTGTCCTTTTCTGCTCCCTCCATATTTTCGTCTTGATCCTATTTCTAGTTTATGTTCGTCTTTTATTACCCTCCTTCTTCCTCGTTT
Consensus                         * **  *        *  ** **** *  ** *** * ***  *  * **    *   **** * *** *      * **      * **  * **  **       ** *** **** *

Teleogryllus_occipitalis      TAAACTTTCGCAACTTCAATTTTTCTTCACAATCATCTCATTTTGTT-TTGCACACCATTATCTAC---TTTTGCTTCTACATCTTCACCATTTCCTTTCTTTTCTACATCTGCTTCAT-
Tarbinskiellus_portentosus    TTATCTTTTTC--TCTTACTTTTTTCTCATTCTTCTATCATTTTCTTCTTGCACCTCCTAGTTTACTAATTCTCCTTTTCCCTTTTCACTTCTTTTTATATTTTCCTTTTCTTTTCCGTT
Consensus                     * * ****  *     * * *****  ***    *   * ******* ** ******  * *  * ***     ** * *** * * * *****    **   * * *****     ***   * * *

Teleogryllus_occipitalis      ------------------------------------------------------------------------------------------------------------------------
Tarbinskiellus_portentosus    GAAACTCGTGATTTCTGGTTGTGAAAGTACCCATTTTTAACGACTATAGTACTCGTACGTCACTGATCTGAAATACTGATCTTTCGGAAGACAAATGATCTTGCTGTGGTTCTTGACTTG
Consensus

Teleogryllus_occipitalis      ------------------------------ATTCTTCTTCTTCT-----------------------------------------------------GCTGCTGCTGCGGCGTCTT
Tarbinskiellus_portentosus    CCAAGTTTTGTTCCACTGCTGCCAAAATTAATTTCTCTCCTTTTTTCCGTTTGCGTGAACGTACACAATCACGTTTATAAGGCTGAGTTTAATGACAACGAGTTCCTGGAACTGGTTACA
Consensus                                                   ***   *** *** *                                                     * * ***    * *   *

Teleogryllus_occipitalis      CTGATACTGACTCTGCATCATCTAC-----TGTTGCTACTGCTTCTGAATCTTCTTCTGCTTCTTCTGCATTTTC-------------------------------------------
Tarbinskiellus_portentosus    TTTACACTGTTTGTGGGCCATCTTCTAACATGTAACTCATAACTTCATATCTTATTTTTTTTATTTATTCACATTCAGTGTGTGTTTGTTCAAACGATTGGCCATAGCATGAACTGTTTAT
Consensus                      * * ****   * **    ***** *       ***   **   *    *     ***** ** *   *   ** * * **   ***

Teleogryllus_occipitalis      --TTCTATTGCTGCTTCTGCATCTTC------------------AGCTGCTTCTGGTACGGAAACTTCTGCT-----------------------GTAGCATTTTCTTCTCCTGTTGC
Tarbinskiellus_portentosus    GTTTTTAATACTTTACTTAAATCTTTACGCCGAAGCTATGCAATAAACCGCATCAAAAATACACATTTATATTCTTTTTCTCTATTTTAAAGTACCGATGTTCCTATATTTCTCTGCTGT
Consensus                       ** ** * **      *   *****                    * * ** **     *     * * ** *   *                         *      * * **    *** **

Teleogryllus_occipitalis      CACCAGAGCG---
Tarbinskiellus_portentosus    TCTTGGAATATGG
Consensus                          **
```

**Tocc.1G0021480.1_exon1**

```
Teleogryllus_occipitalis          ------------------------------------------------------------------------------------------------------------------------
Tarbinskiellus_portentosus        RSAIVAQFPDRNRIAI*RWWG*YSAANCPAVRG*STFLNLFIIGRRRAVPAF**VTPDVL*NISIMPRDTYSCSTFLLPVPSSIANRPASRPAEKVRNNNFEIATQ**IHYSIVISTAY*
Consensus

Teleogryllus_occipitalis          --------------------------MK--------------------------------------------------------------------------------------------
Tarbinskiellus_portentosus        LNSCKYNKITHKNIE**QINWLTLE*MKTIL*Q*KVKVNKTY*QN*IKTIKIQHASGGW*AHSSSTRITNPNQ*RHDGVMMVVEKLMNGNVLFFFMLYHCLPYVSSRKLRTECDVVLLSS
Consensus                                                   **

Teleogryllus_occipitalis          ------------------------------------------------------------------------------------------------------------------------
Tarbinskiellus_portentosus        SDTG*PGVKLLV*DKSTHTHTYTHICVIILRYRNHTFSNIFNKT*I*YLCCEEIVSLEFSYLLQKLRIKYKNSSNLNRRNKLPCSEF*SINKK*CL*KRSLFP*PHDFFHPGKICYISSI
Consensus

Teleogryllus_occipitalis          ------------------------------------------------------------------------------------------------------------------------
Tarbinskiellus_portentosus        TTQR*NFLETFI*YAHSSSFRFSETLLWYFAWQRFSVKFSFFFPHRHHTPKVSSTSTASASSTRIAPGRSCK*LQRG*CLVDFNIETPAVSFFKTSFRPRKRAEGERAGGGQGAGRGGQG
Consensus

Teleogryllus_occipitalis          ------------------------------------------------------------------------------------------------------------------------
Tarbinskiellus_portentosus        AGRGRARGGQGTGKGRARGGQGRGGERWRNSRPAHVDKRLEFP*ASRLTDESVLSRRVSSLLLFVSFFSFFSLSSSPCTSASSSPLPSLSSS*PWVIRVARSHNCISALEFLSLDGF*KN
Consensus

Teleogryllus_occipitalis          ------------------------------------------------------------------------------------------------------------------------
Tarbinskiellus_portentosus        FRTFINDILRS*FHRNYLF*FHFGLCAW*LSFYLPFLLFSLFIF*LYIFIIYDFFVWLSFSAPSIFSS*SYF*FMFVFYYPPSSSFLSFSLTFFSFFYHFLLAPPSLLILLFPFHFFLYF
Consensus

Teleogryllus_occipitalis          ------------------------------------------------------------------------------------------------------------------------
Tarbinskiellus_portentosus        PFLFR*NS*FLVVKVPIFNDYSTRTSLI*NTDLSEDK*SCCGS*LAKFCSTAAKINFSPFFRLRERTQSRL*G*V**QRVPGTGYIYTVCGPSSNM*LITSYLIFLFIHIQCVFVQTIGH
Consensus

Teleogryllus_occipitalis          -------------------------------------------------
Tarbinskiellus_portentosus        SMNCLCF*YFT*IFTPKLCNKPHQKYTFIFFFSILKYRCSYISLLFLEY
Consensus
```

**Tocc.1G0021480.1_exon2**

```
Teleogryllus_occipitalis      ----------------AGAA--------------SHF----------------------------------PSSLATGSSSPSS------------------------------------
Tarbinskiellus_portentosus    ALQSLLSSQIAIELRSSGGGDDTQLPIALLFVGKAHFLIYS*SGDDEQFQLSDESHPMFFEILVSCHVTPIPAARSSFQSHPPSLIALPLAPRKRYVTIILKLQHNNKFTIQ*L*VQHIN
Consensus                                      *                 **                                  *       * * *

Teleogryllus_occipitalis      ------------------------------------------------------------------------------------------------------------------------
Tarbinskiellus_portentosus    *IVVNITK*HTKILNNNKLIG*LLNK*KLFYNNKK*KLIKHTNKIK*KL*KYNMLQADGRPTLPVLGLPIQTNDDTMVL*WLLKN**MEMYCFSLCFTTVFHMFPVVN*ELNVMWFFYHP
Consensus

Teleogryllus_occipitalis      -------------------------------------------------------------------------------------SSPSSSSPSS-------------------------
Tarbinskiellus_portentosus    LTPGNLE*SY*CRTNLRTHIHTHTYV**FCDTEIIHLVIYLIKHRFNIYVAKKLSLSSSVICCKN*E*NTKILVI*TEEINFHAQSSEVSTKSNACRKDLYFPNHTIFFIQEKFVIYHLL
Consensus                                                                                                  **  *

Teleogryllus_occipitalis      ----------------------------SSPSSSSPSSS---SPSSSSPSSSSP------------------------------------------------------------------
Tarbinskiellus_portentosus    RRNVRISWKHLFNTHILLVFDFLKRYCGTSLGRDSP*SSLSFFPTGIIPPKFPPLLPPPHPPPELRLGGAANNYNVANV**ILI*RRQLYHFSKRAFGQESGRKASGQGAGRGRAGAGRG
Consensus                                                  *    ** **    *    *    *

Teleogryllus_occipitalis      ----------------------------------SSSSPSSSSPSSSSPSSSSPSSSSPSSSSPSSSSPSSSSP-----------------------SSSSPSSSS--------------
Tarbinskiellus_portentosus    RAGGGQGAGKGRARGGQGAGRGGEESVGGIPAPRTSTNAWNSREHRASPMKVSSRDASLRCSCSFPSSPSSPFPPPLVLLPLLLLSLLYLLLNLGLFASLAPIIVSARWNFYH*MVFKRI
Consensus                                                 *     *     **   *    *   *    *****  *                        *  *   *

Teleogryllus_occipitalis      ---------------------------------------PSSSS-------------------------PSSSSASSY--------------FSSS*-----------------------
Tarbinskiellus_portentosus    LEHLSMIFYARDFIGIIYFDSILVYVLGNFLFICHFYYFPSSFSNYIFLLFMIFSSGCPFLLPPYFRLDPISSLCSSFITLLLPRFYLFLLLFSHSSIIFFLHLLVY*FSFSLFTSFYIF
Consensus                                                          *** *                         * **  **               ** *

Teleogryllus_occipitalis      ------------------------------------------------------------------------------------------------------------------------
Tarbinskiellus_portentosus    LFFSVETRDFWL*KYPFLTTIVLVRH*SEILIFRKTNDLAVVLDLPSFVPLLPKLISLLFSVCVNVHNHVYKAEFNDNEFLELVTFTLFVGHLLTCNS*LHILFFYLFTFSVCLFKRLAI
Consensus

Teleogryllus_occipitalis      -------------------------------------------------
Tarbinskiellus_portentosus    A*TVYVFNTLLKSLRRSYAINRIKNTHLYSFSLF*STDVPIFLCCSWNM
Consensus
```

**Tocc.2G0002070.1**

```
Teleogryllus_occipitalis        CGGGGCCTCTTAAAATATTAAGCAATTTCAAAACTCTCTCCAAATTACATAAAACGAAAACTCAGACATTTTTAAAAGGCCAGTTTAAATTTAGGCCCGCGGTTAATTATGGGAACCGAA
Acheta_domesticus               CACGCTCTCTTAAAATATTAAGCAATTTCAAAACCCTTTCCAAATTACATAAAATGAAAACTAAGACA-TTTTAAAACGCCGGTTTAAATTTAGGCCCGCGGTTAATTATAGGAACCGAA
Gryllus_bimaculatus             CAGGGCCTCTTAAAATATTAAGCAATTTCGAAACCCTCTCCGAATTACATAAAATGAAAACTCGGACA-TTTTAAAACGCCAGTTTAAATTTAAGCCCGCGGTTAATTATGAAAACCCAA
Gryllus_longicercus             CAGAGCCTCTTAAAATATTAAGCAATTTCGAAACCCTCGCCGAATTACATAAAATGAAAACTCGGACA-TTTTAAAACGCCAGTTTAAATTTAGACCCGCGGTTAATTATGAAAACCCAA
Consensus                       *     *********************** **** **  ** ************ *******  **** ******** *** ***********  ***************   **** **

Teleogryllus_occipitalis        CGGCTTGTAATTAAACTGAAATGCGGAGTCTGCCGTTTCTGAAAGACGCTTATGCGAAATACATCAATGAACTTAATTTTAACACTAACAGCTGCGAACAGCACACACTGTGTTGGCTCG
Acheta_domesticus               CGGCTTGTAATTAAACTGAAATGCGGAGTCAGCCGTTTCTGAAAGACGCTTATGCGAAATACATCAATGAACTTAATTTTAACACTAACAGCTGCGAACAGCGTACACTGTGTTGGCTCG
Gryllus_bimaculatus             CGGCTTGTAATTAAACTGAAATGCCGAGTCGGCAGTTTCTGAAAGACGCTTATGCGAAATACATCAATGAACTTAATTTTAACCCTAACAGCTGCAAACAGCGTACACTGTGTTGGCTCG
Gryllus_longicercus             CGGCTTGTAATTAAACTGAAATGCCGAGTCGGCAGTTTCTGAAAGACGCTTATGCGAAATACATCAATGAACTTAATTTTAACACTAACAGCTGCGAACAGCGTACACTGTGTTGGCTCG
Consensus                       *********************** ***** ** ***************************************************** *********** ******  ****************

Teleogryllus_occipitalis        AAAATTTGGCTGCGGAAGAATGAATTTAATTGCGAAGACAAACATAATAAATAGGAGAGAATGGAGGAGCCGGAGAATTGTGTACGTAATTAATAATTACATTGGTGTGACACGTCGACG
Acheta_domesticus               AAAATTTGGCTTCAGTAGAATGAATTTAATTGCGAAGACAAACACAATAAATAGGAGAGAATGGAGAAGCCGGAGAACTGTGTGGGTAATTAATAATCACATTTGTGTGATTCGCCGACG
Gryllus_bimaculatus             AAAATTTGGCTTCAGAAGAATGAATTTAATTGCGAAGACAAACACAATAAATAGGAGAGAATGGAGAAGCCGGAGAATTGTGTAGGTAATTAATAATTCCATTTGTGTGACACGCCGAAG
Gryllus_longicercus             AAAATTTGGCTTCAGAAGAATGAATTTAATTGCGAAGACAAACACAATAAATAGGAGAGAATGGAGAAGCCGGAGAATTGTGTAGGTAATTTATAATTCCATTTGCGTGACACGCCGAAG
Consensus                       *********** * * **************************** ******************** ********** *****  ****** *****  **** * ****  ** *** *

Teleogryllus_occipitalis        AGTTTTGGCTTAACAATGGCTCCGAAAAAACGGACAGTTTAAATCCA-GTTTAAAATAT-------------------------------TACAGGTCATGTTAATACTTCAATGTTAAG
Acheta_domesticus               AG-TTTGCCTTAACAATGTCGTTGAAGAAA-GGGCAGTTTAAATCCAGGTTTAAAATGTGTTAA-------------GGGGAGTTCATTCTGCAGATCACGTTAATATTTCAATGTTAAG
Gryllus_bimaculatus             AGTATTGGCTTAACACAGGCTCTAAAAAAA-GGGCAGTTTAAATTCA-GTTTTAAATATTTTAATTACATATCTCAGCGTGCGTTCATATTATACGTCATGTTAATACGTCAATGTTAAG
Gryllus_longicercus             AGTTTTGGCTTAACACAGGCTCTATAAAAA-GGGCAGTTTAAATTCA-GTTTTAAATATTTTAATTATATATCTCAGCGTGCGTTCATATTATACATCATGTTAATACTTCAATGTTAAG
Consensus                       **  *** *******  * *     * *** ** ********** ** **** **** *                              *  *  *** *******  ***********

Teleogryllus_occipitalis        CATTAACATTTCAATGTTCAATCTCGAATACTTTGCGTGAGAATTTAATTTCTACCCCAAAAGAAAACAAAAAAATCAGATCTTGAGTACAAGAG---AAACATGAAATATATTGAGAAA
Acheta_domesticus               CGTTAATATCTCAATGTTTAATCTCGAATTCTTCG-GTAAAAATGTAATTTCTATGCC-----AAAACAAAAAAATCAGATCTTTAATACAAAAGAAAAAACGAAAAATATATTGAGAAA
Gryllus_bimaculatus             CGTTTATATTTTAATGTTCAATCTCGAATACTTGTCGTGAAAATTTAATTTCTACGCCAAAAGAAAAT-CAAAATTCATATCCTCAATACAAGAG-AAACGCGAAAAATTATTTAATCAG
Gryllus_longicercus             CTTTTATATTTCAATGTTTAATCTCGAATACTTGGCGTGAAAATTTAATTTCTACGCCAAAAGAAAAC-CACAATTCATATCCTCAATACAAGAG-AAACGCGAAAAATTATTTAATCAT
Consensus                       * ** * ** * ****** ********** ***   ** * *** *********  **     ****   * ** *** *** * * ***** **   *  *   ****   ** *  *

Teleogryllus_occipitalis        AAA-TCCACATTATTAAT----------------------AAGAAAGGAGTATATATTTATTCACATCCATATCTTAATCATACAGTTGAGTACAAGCGATCAACATGGGTGTTTTTCTA
Acheta_domesticus               AAATTTTGCATTACTGA--------------------------AAGAAAGTAGATATTCATTCATAACCATATCTTAATCATACAGT----CACATGCGATCAAAAT-AGTGTTTTTCTA
Gryllus_bimaculatus             AAATTCCACTTTCTTGAT-AAGAAACATATTATTAATACGAATAAAAACGAGTCTATTCATTCACAACT-TGACTTAATCATTCATC----------CGATCAATAT-AGATGTTTTCTA
Gryllus_longicercus             AAATTCCACTTTATTGATAAAAAAACATATTATGGATACGAATAAAAACGAGTCTATTCATTCACAACC-TGACTTAATCATTCATC----------CGATCAATAT-AGAGGTTTTCTA
Consensus                       *** *   * **  * *                       **    *    **** ***** * *  *  ********* **           ******* **  *   *******

Teleogryllus_occipitalis        GTGAATTCAACAATTTTACAGAAAATAATTTAACGAGCTAATTTCATTGAAAGAAACGTCATTATGGAAAAGGAATTCAAATAGATCAAGGTTATTATTTAGACAATTCCCACATTTCTA
Acheta_domesticus               GTGATTTCAAAAATTTTACAGAAAA-AATCTACGAGAAATATTCCATTCAAAGAAGCGACGTTATTGAAAAGGAATTCAAATGAATAAAAACTCTTATTTACATAATTCCCACATTTCTA
Gryllus_bimaculatus             GTGAATTGAAAAATGTTACAGAAAAAAATTCTACATGCTGTATTTAATAAGAAGAAACGAGACTATTGAGAAG------------------CTCTTTTTTACACAATTCTATAATTTCTA
Gryllus_longicercus             GTGAATTGAAAAATGTTACAGAAAAAAATTCTACATGCTGTATTTAATAAGAAGAAACAAGACTATTGAGAAG------------------CTCTTTTTTACACAATTCCATAATTTCTA
Consensus                       **** ** ** *** ********** * * **        ***  **   ***** *     *** ** ***                 * ** **** * *****    *******

Teleogryllus_occipitalis        ATTTCCTCCACAACCGTAATTATTATATTTTTTAAAACCACATGTACAAGAGTGGAGAAAGTTTAACAAATTAATTAGAAATTGATTTCATTCACAAGGGAATGTATTACAATTTATATT
Acheta_domesticus               ACTTCTTCCACAATCGTAATTATCTGATTTTCTAAAACCACATGTAAAGGAGTGGAAAAGGTTTAAAACTTCAATTCGAAATTGATTTCGTTCACCAGGAGATGTATTAAAATTGATATT
Gryllus_bimaculatus             ATATCTTCCACAATCGTAATTATTTGATTTTCTAAAATCACATGTAAAATGTGGAATAAAGTTT--------AAGTCGAAATTGATTTAGTTCACGAGGGAATGCATTAAAATTTATATT
Gryllus_longicercus             ATATCTTCCACAATCGTAATTATTTGATTTTCTAAATTCACATGTAAAGCGTGGAATAAAGTTT--------AACTCGAAATTGATTTGGTTCACGAAGGAATGCATTAAAATCCATAAT
Consensus                       *  ** ******* *********   ***** ****  ******** *     * * ** ****        ** * ***********  ***** * *  *** **** ***  *** *

Teleogryllus_occipitalis        ATTCTCCCCTCTGAATGATAAACTCTAATATGAAAAACAATAGATTCGAATCAACTAATAGTATGTATTCAATAAAAACAACGAACTTCTGAAGAAATATTATCGACTACATTTCTTGTA
Acheta_domesticus               ATCTTAATTCCTGAATGATAAACTGTAATACGAGAAACAATAGGTTC-------------------------------------------AAAGAAACGCT---------ACTTCCTGTA
Gryllus_bimaculatus             ATTCTCATACTTGAATGATAAACTTTAATAC-AAAAACAATACATTCGAACC---------AATATATTCAATAAAAATAACAAAATTTTTAAGAAATATTCTCGAC---GCTACTTCTA
Gryllus_longicercus             ATTCTCATACTAGAATGATAAACTTTAATAC-AAAAACAATACATTTGAATT---------CATATATTCAATCAAAATAACTAACTTCTTAAGAAATATTCTCGACGCTACTTCTTCTA
Consensus                       **  *       ************ *****  * ********  **                                                   ******   *           * * * **

Teleogryllus_occipitalis        CGAGCCCGCTTGTGTAAAAGTCTAAACAAATAGCTTCTCTGTCTGAGAATTC--CGTTTAATTTACAAACCGTCCAGGAGCACTTTACAACTGCTCAGTATTATATATTTGGCACTTCTGTT
Acheta_domesticus               TGAGATCGTTTCTATAAAAGCCTG----------TCTCTGACTGAGGAC------TTAAATTCACCAACCGTCCTGTAGCATTTTAGGACTACTCAGTATTATATATTTGGCACTTCTGTT
Gryllus_bimaculatus             CGAGACCGCAACTATGAAAGTCTAAACAAATCGCTTCTCAGTCTGAGAATTCCACACTTAATTCACAACCCGTCTTATAGCACTTTACAACTCCTGAGTATTATACTTGGCACTTCTGTT
Gryllus_longicercus             CGAGATTGTAGCTATGGAAGTCTAAACAAATCGCTTCTCAGTCTGAGAATTCCACACTTAATTCCCAACCCGCATTATAGCACTTTACAACTCCTGAGTATTATATATTTGGCACTTCTGTT
Consensus                        ***   *    * *  *** **            **** * ***** *        * ****  * * ***      **** ****  *** ** ********* *************

Teleogryllus_occipitalis        TCACATACTTCGCAAAATAACTTCGAACTGTTAGAAAAGACAAAAACCCTAACTAG-AAAAACTTGCTTACTACCGTTTTCGCTCGGAACGCTGTTAACTGCAGCATCACAATCATCCAT
Acheta_domesticus               TCCCATACTTCGCAAAATAACTTCGAAGTGTTAGAAAAGACAAAAACCTCAACAAGAAAAAACTTGCTTACTACCGTTTTCGCTCGGAACACTATTCGGTGCAGCATCACAATCGTCCAT
Gryllus_bimaculatus             TCCCTTACTATGCAAA--AACTTCGATCAGTTAGAAAAGAC------CTCAACTAG-AAAAGCTAGCTTACTACTGTTTTCGCTCGGAACGCTAGTAAGTGCAGCATCACAATCGCCCAT
Gryllus_longicercus             TCCCTTACTTTGCAAA--AACTTCGATCAGTTAGAAAGGAC-AAAACCTCGACAAG-AAAAGCTTGCTTACTGCCGTTTTCGCTCGGAACGCTAGTAAGTGCAGCATCACAATCGTCCAT
Consensus                       ** * ****  *****  ********   ******** ***      *   ** ** **** ** ******* * *************** **  *   **************  ****

Teleogryllus_occipitalis        CCGCCCCTT-CCCGATAACCCGAGGGCCAATCCTGGGCGCCGTTTGCCGGAGGAGCGGCTCCTGGGGAATCGTGAGCGGAGTGCGGGACGCGG---------------------------
Acheta_domesticus               TCGCCCCCTACCCGATAATTCGAGGGCCAATCCTCGGCGCCGTTCGCCGGAGGAGCGGCTCATGGGGAATTGGGAGCGGAGTGCGGGGCGCGGGGCGCG---------------------
Gryllus_bimaculatus             CCGCCCCTT-CCCGATAACTCGAGGGCCAATCCTCGGCGCCGTTTGCCGGAGGAGCGGCTCGTGGGGAATTGGAAGCGGGGAGCGGGGCGCGG---------------------------
Gryllus_longicercus             CCGCTCCTT-CCCGATAACTCGAGGGCCAATCCTCGGCGCCGTTAGCCGGAGGAGCGGCTCGTGGGGAATTGGAAGCGGGGAGCGGGGAGCGGGGAGCGGGGCCCGGGGAGCGGGGCGCG
Consensus                        *** ** * ********  ************** ********* **************** ******** *  ***** * *****  ****

Teleogryllus_occipitalis        --------------------------------------GCGGGCGCCCACAGGGCCACAAAGCGGCGGGCGCTATGTTTAGCGCACGGAATGGGCACAACAGCCTACGGGCGGCGCGCCA
Acheta_domesticus               ----------------------------------GGGCGCGGGCGCCCACAGGGCCACAAAGC----GGCGCTATGTTTAGCGCACGGAATGGGCACAGCAGCCTACGGGCGGCGCGCCA
Gryllus_bimaculatus             -----------------------------------GGCGCGGGCGCCCACAGGGCCACAAAGC----GGCGCTATGTTTAGCGCACGGAATGGGCACAACAGCCTACGGGCGGCGCGCCA
Gryllus_longicercus             GGGAGCGGGGCGCGGGGAGCGGGGAGCGGGGCGCGGGCGCGGGCGCCCACAGGGCCACAAAGC----GGCGCTATGTTTAGCGCACGGAATGGGCACAACAGCCTACGGGCGGCGCGCCA
Consensus                                                             *************************    ******************************* ********************

Teleogryllus_occipitalis        GGGGCGGCGACGGCG--------------GCGGATGTGGAGAGGCGCACACTGGCGGAGGCGGTGCGAGAGGAAAGACGGCGGAGGCGAAGGAGGCGGCGGCAGCGGCGGCGGTGGCAGG
Acheta_domesticus               GGGGCGGCGGAGGCGGCGCGGCGGAGGGAGCGGATGTGGAGAGGCGCACAGTGGCGACGGCGGCGCAAGAGGGAAGGAGACGGCGGCGGCGGCGGCGGCGGCGATGGCATTGGTGGTGGC
Gryllus_bimaculatus             GGGGCGGCGACGGCG-----GCGGAGGGAGCGGATGTGAAGAGGAGCACAGCGGCGAAGGCGGCGCGCGAGGGA----GGCGGAGGCGGTGGCGGTGGCGATGGCAGCGGTAGAGGTGGC
Gryllus_longicercus             GGGGCGGCGTCGGTA-----GCGGAGGGAGCGGATGTGGAGAGGAGCACAGCGGCGAAGGCGGCGCGCGAGGGA----GGCGGAGGCGGAGGCGGTGGCGATGGCGGCGGTAGAGGTGGC
Consensus                       *********  **              ********* ***** *****  ****  ***** **  **** *    * *** ****  ** ** ****      **    * **  *

Teleogryllus_occipitalis        GGCGACGGAGGCGACAGAGGCGGTGGCGGTGGCGGTGGCGGCAGAAGCGACGGCGGCAGAAGCGGCAGCGGCAGCGGCAGAAGCGGCAGCGGCGGCGGCGGCGGCGGCGGCGGCGGC---
Acheta_domesticus               GGTGGTGATGGTGGCAGTGGCAGTGGTGGTAGTGGTGGTGGTGGCAGTGGCAGTGATGGTGGTGGTGGCAATGGCAGTGGTAGTAGTGGTGGTGGTGGTAGTGGTGGTGGTGGCAAT
Gryllus_bimaculatus             GGCGGCGACGGTNNNNNNNNNNNNNNNNNNNNNN--------------------------------------------------------------------------------------
Gryllus_longicercus             GGCGGCAGTGGCGGTGGTGACGGTGGAGGCGGCGGTGGTGTCGGCGGAGGCGGCG---------------------------GCGGCGGTGGTGACGGCGGAGGCGGCGGCGGTGGCTTC
Consensus                       ** *     **

Teleogryllus_occipitalis        GGCGGCGGTGGCAGGGGCGGCGGCAGAGGCGGCGGCGGCAGAAGCGGCGGCGGTGGTGGTGGTGGCGGCGGCGGCGGCGACAGAAGCGGCGGCGGCGGCGGCGGCAGAAGCGGC------
Acheta_domesticus               GGCAGTGGTAGTAGTGGTGGTGGTGGAAGTGGTGGTGGTGGTGGTGGTGGTGTTGACGGCGGCGACGGCGACGGCGGCGACGGCGGCGGCGGCGGCGGCGACGGCGGCGGCGGCGGCGGT
Gryllus_bimaculatus             ------------------------------------------------------------------------------------------------------------------------
Gryllus_longicercus             GGCGATGGCGGTGGTGGTGGCGGTGGCGGTGGCGGTGGCGGAGGCGGAGGCGGAGGCGGAGGCGGAGGCGGAGGCGGAGGCGGAGGCGGAGGCGGAGGCGGAGGCGGAGGCGGA------
Consensus

Teleogryllus_occipitalis        ------------GCAGAGGCGGCGGCCGAGACGGCGGCGGTGGTGGCGGCAGAGGCGGCGGCGGTGGAGGCGGCGGCAGTCGCAGCAGTGGCAGAGACGGCTGCGA-------GGAGGC
Acheta_domesticus               GGCGGTGGCGGTGGCAGTGGCGGCGGCGGCGGCGGCGGCGGCGGCGGCGGCGGCGGCGGCGGCGGCGGCGGCGGCGGCGGCGGCGGCGGTGGCAGTGGCGGTGGCGGTGGCGGTGGCGGC
Gryllus_bimaculatus             ------------NNCAAGGGAAGGGAT---------------------------------------------------------------------------------------------
Gryllus_longicercus             ------------GGCAGAGGCGGAGGCAGAGGCGGAGGCAGAGGCGGAGGCAGAGGCAGAGGCAGAGGCAGAGGTGGCTGAAGCAGAGGTGGCTGTTGCCGCGA----------GGAGAC
Consensus                                     **  **  * *

Teleogryllus_occipitalis        TGTGGCGGAAGTGGTGGTGGTGGTGGTGGTGGTTGTGGTTGAA---------------------------GTTCGGGGTGGTAGGAGCGGCGTCGGAGGAGA-----------------T
Acheta_domesticus               GGCGACGGCGGCGGCGGCGGCAGTGGCGGCAGTGGCGGCTGCAAGGAGGCGGTGGCGGTACGTAGAAGTTGTTCGGAGTGGTAGAAACGGCGTCGGAAAAGA-----------------T
Gryllus_bimaculatus             --------------------------------------------------------------------TTTTTTGGGGGGTTCGAAACAACACAACAAAAGGCCCTTTTTTAATTTTTTT
Gryllus_longicercus             TGTGGCGGTAGCGGCGGTGGTGGCAGTGGTTG---------------------------------ACGTTGTACGGAATGGTAGGAACGGTGTTGGAAAGAG-----------------T
Consensus                                                                                     *  **   * * * * *        *                 *

Teleogryllus_occipitalis        CGAGTGATAGTGTGTGTACGAGGTGCTGCAGCGTTGAGAGATACGAAACCGAAATAT--------------GTGGCAAATGTGCATCCCATTCCAGGAC-------------------TT
Acheta_domesticus               TGAGTGGTAGTGAGTGTACGAGGTGATGCAGCGTTGAGAGACTTGAAACTGAAATAC--------------GTGGCA-----GCATTCCATTCGGGAG---------------------T
Gryllus_bimaculatus             CAATTAAAAGTTAAAAAATAAAATA---------TTAAAAAAATGAACCTGAAAAATTTTCAATACCGAAAGGGACAATACAAAATTTAATAACGAAGACAGAAAAAAAAACGCTAATGT
Gryllus_longicercus             CGAGTGCTAGTGATTGTACGAGGTGAGGCTGTGTTCAGAGATATGAAACCGAAATTT--------------GCGGCAATCGTGCATCCGATTCCGGGACT----------------TTTT
Consensus                         * *   ***      *  *  *          * * * *   *** * ****                 * * **       **   **                            *

Teleogryllus_occipitalis        TTTTCCAGATTGGAAGAAAGATAATCGTTGGAG--------------AAAGAAAAGCAAAACGAAAACGGACAAAATCCTAAGGACTCGCAGAGGTAAAGATT----------------C
Acheta_domesticus               TTTCCAAGATTGGAAGAAAAATAATTTTTGAAA--------------AACGGGAAGTAA---------GGTACTACACTAAGGTATCTCGGAAGTGAAG---------------------
Gryllus_bimaculatus             TTTTTTAGACTATTGAAACAACGTCCCCCGTAATAATATATTTTCCCCAAAAAAAAATAATA-------GGAAAGAACAAAAAATAAATACAAGAGCACAAATTATAAATAAAATTGGGC
Gryllus_longicercus             TTTTCCAGAATGGAAGAAATATATTTTTTGGAG--------------AAAGGAGAGTAA------GGTAGAGGCGGTCCTAAGGACACGCAGAGGTGCAGATT----------------C
Consensus                       ***   *** *     **  *        * *                **     *  **           *          **        *    *   *

Teleogryllus_occipitalis        ATCGTGGACACGTTCCCAGGACAAAGAATTATTGTGTG---------------------------------------------------------------------TGTGTGAGTGAGA
Acheta_domesticus               ----------------------AAAGGTTTATTTTGTGTTTGTGTATGCGTGTGTGGAGTGCGCTCGCTCGCTTGCGCGAGCGCGTACCAGCATGTATGCGAGCCTATGTGCGAGCGAGA
Gryllus_bimaculatus             GCCTTAATTTTAATTATCGTGCAAAGATTTTTCTGGCGTTTC-----------------------------------------------------------------GAAATAAAAATAA
Gryllus_longicercus             ATAGTGGTCATATTGGTATGACGAAAAATTAGCTTGTGTTTGTGTATGCGTGTGTGGTGAGTGTGCGGGGGCGCGCGCATGCGAGCGATAGAGAGAGTAAGATATGGAGAGAGGGGGATA
Consensus                                                **   **     * *                                                                                 *

Teleogryllus_occipitalis        GAGAGAGAGCGAGAGCGA---------------------------------------------------------------------------------------------------GAG
Acheta_domesticus               AAGAGGTAACGAAAGCGAA-----------------------------------------------------------------AGGAGAAAGAGATTTAAAAACTGA-------ACGAG
Gryllus_bimaculatus             ACAATACAATGAATGTGAAACTTTTTTTC------------------------------------------------------------------ATGCTAAGATTGTACACAATTTGCG
Gryllus_longicercus             GAGACGAATAGAGAGGGAGGATAGAGACGTATAGAGAGGGAGGATAGAGACGTATAGAGAGGGAGGATAAAGACGTATAGAGAGGGAAAGGAGAGATATAAAAATTG---------CGCG
Consensus                          *   *  **  * **                                                                                                   * *

Teleogryllus_occipitalis        AGAGCGA-----------------------------------------------------------------------------------------------------------------
Acheta_domesticus               AGAGCGACAATAGGCGACTGACTAGGCAACAAGCACAA--------------------ACTGAGCGTACGAGCTGATGTGTGTCCAAGGGACAAAGAGAGGACACACAGGTTCAAAGACA
Gryllus_bimaculatus             AAATTTTTCATTAGCC-------------------CAATTTGGTTTTTCCTCATTTCCACCCAAAACCCAAGCCGGCGCTTTTCCAAGTTGCACCCGGCTAAAACTCGGTCTCAA-----
Gryllus_longicercus             AGAGCGATAATTGGCA-------------------------------------------------CTAGGAGCGGACGTGTGTCTAAGGTG-AGAGAAGGGACACACAGGTTCAAAGACA
Consensus                       * *

Teleogryllus_occipitalis        ----------------------------------------------------------------------------------------------------------------GAGCGAAA
Acheta_domesticus               GCGAACAGACAGAAAGAGGCGGACAAGGTAC-----------AGAAGGGGAT----------------------------TCACGGGAGAATTGTTGGAAGAGAATCAACCTGAACGAGA
Gryllus_bimaculatus             ---------------TGTTGATCATTGCACTTTTTTAATGTAAAAGTGAACCTTTACATTTTACTTTACGCATTCACTACCATAATAAAACTGCTGCAA---------------------
Gryllus_longicercus             GTGAACAGACAGAAAGAGGCGGTCAAGGTACACGCAC-ATAAATAAGTGGAT----------------------------TCAAAGGAGAATTGGTGAAA---AGTCAACCTGAACGAGA
Consensus

Teleogryllus_occipitalis        GA----------------------------------------------------------------------------------------------------------------------
Acheta_domesticus               GGAGGCTTCCCGAAACGTTTTCCGG--------ACCTCAAATGCTATCTCCCGGGGAACATCCGTCACCAAATGCGGGATATCGTCTCCATATCAATGAACGTCGTAGAGAGATGAGCA-
Gryllus_bimaculatus             ------------------------------ACATTCCAAACAATATTCTTTGTA------TTGGCAGCAAAAGTACTCCGTTACCTTTCCACTA--------------------------
Gryllus_longicercus             AGCGGCTTCTGGAAATGTTATCGGGACGGCAGCACCTCAGCAGCCATCTCCTGGGAAATGTTCGCCACCGAATGCGGTCTATCGTCTCCATGTAAATGAACTTCGTACAGAGATGAACAC
Consensus

Teleogryllus_occipitalis        ------------------------------------------------------------------------------------------------------------------------
Acheta_domesticus               ACGAAGAAAACCCTTTCTAGATT-TTCGAACGAAAGCTCATGTGTAG---CATGGAATAATAACG----------CTCGTTTG-------------------------------------
Gryllus_bimaculatus             ATCAAGAAAAAAGAATTTGGATATTTTTCACAATAGATCAATTGGAGACCCTCGGGATAATAATAGCAAGATCCTACCGACCTATTTTCAGTTTTTAGAATA----------GATGGAG
Gryllus_longicercus             ACGAAGAAAATCCATTCTAGATTCTCCGAACGAAAGTTAATGTGTAG---CATGTAACAATAACG----------CTTCTCTGTATTTAGTTGTCGAGAGACAGTAAAGGAGCATTGAA
Consensus

Teleogryllus_occipitalis        ------------------------------------------------------------------------------------------------------------------------
Acheta_domesticus               -----------------CATTTCAAGTGAATTTGACGAAAGTATACAAATGTAAATAAGTTAATAATTGCATTATAA-----------------GAAATTATGAAAACCACACTGCCCAC
Gryllus_bimaculatus             AACGCCTGCCAAATAGCTATCGTATCTTGAAACTATCAAAAGAATTGGATATTGGCAAACATGAAATTGCACCTTAAAATGCTGATCCAGAGTGAAAGTGAACAAAAACAGGCCAATTTT
Gryllus_longicercus             TGCATTTTTTAAGTA-TCATTTCATATTAATTTTACGAGAGGATA-AACTGGAAGCTAGCATGTA---GGAGCATAA-----------------CAAATTATGAAAAACACACTGCTCAC
Consensus

Teleogryllus_occipitalis        ------------------------------------------------------------------------------------------------------------------------
Acheta_domesticus               ACGAGTCCAATGACATAACATGGAGAGCTCTTCCCTTACACTGCGCAATATTATTTTTAATTTTGTTAGTCGGACGGTACCTTAATGAATTTTTAGTGTTTCAAATATTTTCTCCCAATT
Gryllus_bimaculatus             CTCAATACAA------------------ACTTCGGTTGGATCTGCTGTTATTATTCTAAGCGGCATCAAATATATATCGCATCACTGAATT------------------------AAATT
Gryllus_longicercus             ACTTGTTCAATACTATAACATGGAGAGCTCTTCCCCCACACTGCCCAATATTATTTTTAATTTCGTTAGACGAACAGTACTTTACTAAATTTTTAGAGTTTCAAATATTTTCTCGTAGTT
Consensus

Teleogryllus_occipitalis        ------------------------------------------------------------------------------------------------------------------------
Acheta_domesticus               AAA--AAAAGGCAGTGG-AAGTACAAAAGCAACGAGAAATTTCGCAGAGAAGACAACTTGTGAAATACAGAAGTGCTAAAGCGAGTTTTGGCCTAATGCCATGAAGTCTAATTACACTAA
Gryllus_bimaculatus             TTAACAACACCCAACCG----------------------TTTTCACCAGTAACAAAACTCCT------------TCTTGGAAAAGCCTAGCCCCAAAAATACATTAACTAACTATTTCCA
Gryllus_longicercus             AGAATAAACGCCAGCGGAAAGTACAAATGGAACGAGAATGTTCGCAGAGAAGATAACTTGTGAAATACAGAAGGGCTAAAGCGAGTTTTGGCCTAACGCCATGAAGTCTAATTACACTAA
Consensus

Teleogryllus_occipitalis        ------------------------------------------------------------------------------------------------------------------------
Acheta_domesticus               CGACAATTTAGGGTGAGAAGAGGCGTAGTCCCTGTACTCTACAGCATGCACAACTCTTGACGTTTCACGCGTTATT----------GCCACGTGTAACTTGCCCGGCTTACAGA------
Gryllus_bimaculatus             AAGGAACCCA--------------------TTCAAACTCCACA-TACAAATGAATTTTCGTGCCTTACTAATTTTTTTAAACTCTCATTATACATTATTTGGTCCGCTAACTGAAATTCC
Gryllus_longicercus             TGACAATTTAGAGTGAGAAAAGGCGTAGGCCCTGTACTCTACAGCGCGCACAACCCTTTCCGTTTCACGCGTTGTT----------GCCACGTGTAACTCGCTCGGCTTACAGA------
Consensus

Teleogryllus_occipitalis        ------------------------------------------------------------------------------------------------------------------------
Acheta_domesticus               --------------CGTACGAGGATACTCACAAACAAACGTTGGAATTCTCCCAAAGCGAACAGTGTAAACAGCACTGTCCTCTTCATCTTAGA-------TCCCCAAATCGTATATTTA
Gryllus_bimaculatus             TTCCTTTAAGCAATGGCGTAAAAGCATCTCTTTAGTTGCTTCGCGGCATTCATGATGCAGAAAGTACTAGCATTTCCCTCAACACTGCTTGGAAATGTGCTTTCCCTAATTTTTTATAAA
Gryllus_longicercus             --------------CGTACGAGGACACTCGCAAACAAACGCTGGAATTCTCCCAACACGAATAGAGTAAACAGCTCCGTCCCCTTAATCTTAGA-------TCTCCAAACCGTACATTTA
Consensus

Teleogryllus_occipitalis        ------------------------------------------------------------------------------------------------------------------------
Acheta_domesticus               ATTTCACCTCTTATAATATTTCCATTGGCACCATCATCGCTGTC-----GCGTGATTCGATTGGTAAAT---------------------------------------------------
Gryllus_bimaculatus             AAATTGGATGGCATGTGAGTAACACTTACGTAAGTATCTATGCCTTTATGCATGAATGTATTAATTTATGTGTGTGTGAGCTCTGTGTGTGCGTGCGCGCGCGCGCGTGTGTGTGTGT
Gryllus_longicercus             ATTTCTGTTCTTGTAATATTTTCATTGGCGCCGTCATCGATGTC-----GCCCGATTTGATTGGGATAT---------------------------------------------------
Consensus

Teleogryllus_occipitalis        ------------------------------------------------------------------------------------------------------------------------
Acheta_domesticus               ------------------------------------------------------CGAAGGCA-GTTCGTGAAATCGATCTTGACGAGAATAGTACTGGAATTCCAAGAAGTTGGAGGCAC
Gryllus_bimaculatus             GTGTGCTGTGTGGTGTGTGTGTGTGTGTGTGTGGTGGTGGTGTGTGTGTGTGTGTGTAAGTA-TTTTGTTATATCATCACTTTCAATCATT--AGTCGATTTTTAGGAAGCAAAAAGAAA
Gryllus_longicercus             ------------------------------------------------------CGAAGACAGGTCCATGAAATGAACTTTTACAAGAATAG-ACGAGAATACCAAGAAGTACGAGG--C
Consensus

Teleogryllus_occipitalis        ------------------------------------------------------------------------------------------------------------------------
Acheta_domesticus               ACAAAATAACGCCATTATCATAAAACGGCATTAAGTTTCAGAAGACGTTTCGTGATCAGATATTTTAAATTTGGATATATGAAGTTGCATTTCAGCTAGTTTAATTTCTAGAAAAAATAT
Gryllus_bimaculatus             AAAAAAAAAAGC--------------------GGGTTTCGAGGATCCA------ATACGTTACGCAAATATTGGATACA--------------------ATAAAACGTCACCATAATTAT
Gryllus_longicercus             ACAAAATAACGCCATTATCATAATATGGCATCGAGTATCAGAAGTCCATTCTTGACAAAGTATTTTAAATTTGGATGTGTAAAGTAGAAGTACAATTATTTTAAGTAATAGAGGAATTAT
Consensus

Teleogryllus_occipitalis        ------------------------------------------------------------------------------------------------------------------------
Acheta_domesticus               ATAATTC---TTCCATTTCATTTAATTCGTTATCTA--------ATAGCGCTTTCGTTACTAGCGCAAGAAC--GTCCATAAGATGACCTGTATACA-AAATTGCTAATCGCTCATAGGA
Gryllus_bimaculatus             GCACTTCGCAATGCATCTAAGGTAAT--GTTGCCCGGCGAACGTGCGGCGGTTGCTGGGCGGAGCCCGGCACAAGCGTGCGAGGCGAGGCGAGGCGA-----------GCAGGGGCAGGA
Gryllus_longicercus             ATATTTC---TTTTATTTAACTCAATTTGTTAATT---------------TTAAACTTTCAGCCCGAGAACATGTGCACACGAAGACCAGAGCACAGAAATTGTT--TCACTAATAAAA
Consensus

Teleogryllus_occipitalis        ------------------------------------------------------------------------------------------------------------------------
Acheta_domesticus               AACAAGTTACTTATGAAA---------------------------------AATATTTCTTAATGAAATTCTTTTTAAAAATAGAAAGCCAA----CAAGTGGAAATAAATGACCAAATT
Gryllus_bimaculatus             GGGTCGAGCGATATGAAGGCGAGCGGGGGTTGGAAGGGCAATCGGGAAGGCGGAGGTTCATAAT-AGATGATGGTGATAATTAAATACCCGGGCAACGCCCGGGAATCAT--GCCAAATA
Gryllus_longicercus             AATTTCCGCGTCATTAAA---------------------------------AAATATTTTTA---AAGTCCTCTTTAAAAATAGAGTGCCGATCACCGAGTTGAAATAATTAACCAAATT
Consensus

Teleogryllus_occipitalis        ------------------------------------------------------------------------------------------------------------------------
Acheta_domesticus               TCCATGTTGATTACCATCTTCATTACTTACTAGAAATTTTCACGGTCTTCCTAAATGATAGTTCATTGGAATGCATCTTCCAATACAATGTTACAAAACTTTGCAACATTAGAGAAAAGG
Gryllus_bimaculatus             TCTTT--------TAATCTTTGTT-CCCGTAACAAACTTCCATAACGTCC-------------------------------------ATGTTTTGGAG------AACACTAACAAGAAGA
Gryllus_longicercus             T------------CCATCTTCATTACCTTAAAGAAATTTTCA------------------------------------------------------------------------------
Consensus

Teleogryllus_occipitalis        ----------------------------------------------------------
Acheta_domesticus               TGGTACAGCCTTTTCATTGAAATGTTTGTTTATTCTTTGATATATGGCGGGTCTGAAA
Gryllus_bimaculatus             GAATGCTCTTCATTCAATTAGCGATGAGGTTTTTC-----------------------
Gryllus_longicercus             -ATTACGCTCTT----------------------------------------------
Consensus
```

**Tocc.2G0002070.1_exon1**

```
Teleogryllus_occipitalis        ------------------------------------------------------------------------------------------------------------------------
Acheta_domesticus               ----------------------SRVH-----------------------ASES*E-----------------------------------------------------------------
Gryllus_bimaculatus             PLFAPPFRHFHLRRNYRGQD*E*QLHTHVPRTGGLQSSMTARSPSPRARTATEALRAFLALLRTGADVCLHPRGTERPTRRHPCTPPLQPRRGLWGTSVLCTWGQHDIFAMNLDRFHRKN
Gryllus_longicercus             ------------------------------------------------------------------------------------------------------------------------
Consensus

Teleogryllus_occipitalis        ------------------------------------------------------------------------------------------------------------------------
Acheta_domesticus               ------------------FIYRN------------------------CC----------------------------------------------------------------------I
Gryllus_bimaculatus             IIK*KQHINIFA*PLMRLF*Y*NMSLDMSNTGSMMRYYENITNKMPKCVLMVI*IYNLPVRTDF**HN*EIKLLRINKVFAFFERIMHRALRNTAYAFS*FPGSMHPRVKGVIYLNVIYI
Gryllus_longicercus             ------------------------------------------------------------------------------------------------------------------------
Consensus

Teleogryllus_occipitalis        ------------------------------------------------------------------------------------------------------------------------
Acheta_domesticus               HFTYIIYVIN---TL*VMV*VTCDITFILKALSFKCSFFLFRIPYTI---SLYILRQKTNKTTTNKNT*NKFSSTRGDNGRLRSRVIRSNYTS--SNLNKEKGQINL*KLDINEEKKKTG
Gryllus_bimaculatus             HFIHIMCVTNTEPTL*VMV*VT------------LLLFLKKVPSNFDFPSFEFCKQSLSLHSSAKNN-NTHQPTKHVK*VLE*--TKW*HLQQPSSF*E*LDQITRVEL*IRRKDK*LF
Gryllus_longicercus             ------------------------------------------------------------------------------NGLLRS---KSNYIK--SH------E*NF------------
Consensus

Teleogryllus_occipitalis        ------------------------------------------------------------------------------------------------------------------------
Acheta_domesticus               *TTL*KLKIKGVKEEKRINNSLETRDKMKKKKLGQITF**NRDK*VKVFLFLFQFFPQQNEELATQNIPCNPKATS-SAHLGSARQRNRP--*PPRAIRKLCKGRRANAGRGAKTPSN*I
Gryllus_bimaculatus             TN*IER------GKEDRINTSLRN*R-*SEKKIR---------------FIF-SSPQK*KR-HPKTSPATPTQPRVRIYGASARQRNRP-*PPPRNPKIMQREARRRPAERPRRLRIKL
Gryllus_longicercus             -----------K*EERINNHLKTRN--NEEKTGEIPL*ETRDKVRKNSFYI--FFATKIKT-SSQNIPCNPNAAS-SAHLRSERAPAQPPITAPTQSENYAKGSAQTPGREAKTPSN*I
Consensus

Teleogryllus_occipitalis        ------------------------------------------------------------------------------------------------------------------------
Acheta_domesticus               IGK-PSN*RETLIKINLAHNMRATMTGTRRAETICPVALARSQPMLPRNCTLLTLS*NIKQFQNPFQIT*NEN*DILKRRFKFRPAVNYRNRTACN*TE---MRSQPF---LKDAYAKYI
Gryllus_bimaculatus             LGSLPTEERRS*K*I------*HTICERRRQAPVEQKPFAPSPWLARHRC-----S*GIVRCSGPLKILSNFETLSELHKMKTRTF*N----ASLNLSPRLIMKTQRLVIKLKCRVGSF*
Gryllus_longicercus             IGK-PSN*REALIKINLAHNMRATTTGPRRAETICPVALARSPLMLLRNCALLRAS*NIKQFRNPRRIT*NENSDILKRQFKFRPAVNYENPTACN*TE---MPSRQF---LKDAYAKYI
Consensus

Teleogryllus_occipitalis        ----------------------------ME------------------------------------------------------------------------------------------
Acheta_domesticus               NELNFNTN-------SCEQRTLC-----WLE-NLASVE*I*LRRQTQ*IGENGEAGELC-G*LIITFV*FADEFAL-TMSLKKG---QFKSRFK----MC*GEFILQITLIFQC*ALISQ
Gryllus_bimaculatus             KTLMRNTSMNLILTLTAANSVHCVGSKIWLQKNEFNCE------DKHNK*ERMEKPENCVGN**FHLCDTPKSIGL-TQALKKG---QFKFSFKYFNYISQRAFILYVMLIRQC*AFIF*
Gryllus_longicercus             NELNFNTN-------SCEQRTLC-----WLE-NLASEE*I*LRRQTQ*IGENGEAGELC-R*FIIPFA*HAEEFWLNTGSIKRAV*IQF*I-----------F*LYIS---ACVHIIHH
Consensus

Teleogryllus_occipitalis        ------------------------------------------------------------------------------------------------------------------------
Acheta_domesticus               CLISNSSV---------------------------------KM*FLCQNK-KIRSLIQKKKRKIY*EKI-LHY*KKVDIHS---------*PYLNHTVTCDQNSVFLVISKILQKKSTR
Gryllus_bimaculatus             CSISNTCR-------ENLISTPKENQNSYPQYKRNAKNYLIRNSTFLIRNI-----LLIRIKTSLFIHNLT*SFIRSI*MFSSELKNV---TEKILH-------------AVFNKK---
Gryllus_longicercus             V---NTSMLSFYISMFNL------------------EYLA*KFNFYAKRKPQFISSIQEKREKLFNHK--FHFIDKKTYYGYE*KRVYSFTT*LNH--SSDQYRGFLVN*KMLQKK---
Consensus

Teleogryllus_occipitalis        ------------------------------------------------------------------------------------------------------------------------
Acheta_domesticus               NIPFKEATLLKRNSNE*KLLFT*FPHF*LLPQS*LSDFLKPHVKEWKRFKTSIRN*FRSPGDVLK--LILS*F--------LNDKL*YEKQ*V------QRN-----ATSCMRSFL*KPV
Gryllus_bimaculatus             ----------KRDY*EALFYTIL-*FLISSTIVII*FSKITCKMWNKV*VEI--------DLVHEGMH*NLYYSHT*MINFNTKTIHSNQYIQ*K---*QNF*EIFSTLLLRDRNYESL
Gryllus_longicercus             ---FYMLYLIRRNKTIEKLFFTQFHNF*YLPQS*LFDFLNSHVKRGIKFNSKLIWFTK---ECIK--IHNILI--------LE**TLIQKQYI*IHIFNQNN*--LLKKYSRRYFFYEIV
Consensus

Teleogryllus_occipitalis        ------------------------------------------------------------------------------------------------------------------------
Acheta_domesticus               S----------D*GLKFTNRPVAF*DYSVLYLALLFPILRKITSKC*KRQKPQQEKTCLLPF----------------------SLGTLFGAASQSSIR----PLPDNSRANPRRRSP
Gryllus_bimaculatus             NKSLLSLRI-----------PHLIHNPSYSTLQLLSIILGTSVSLTMQKLRSVRKDL--------------N*KS*--------LTTVFARNASKCSITIAHPPLPDNSRANPRRRLP
Gryllus_longicercus             AMEV*TNRFSV*EFHT*FPTRIIALYNS*VLYLALLFPLLCKNFDQ-LERTKPRQEKLAYCRFRSER**VQHHNRPSAPSR*LEGQSSAPLAGGAARGELE----A-----------GS
Consensus

Teleogryllus_occipitalis        ------------------------------------------------------------------------------------------------------------------------
Acheta_domesticus               EERLMGNWERSAGRGARGAGAHRATKRRYV*RTEWAQQPTGGAPGAAEAAR---------------RRERMWRGAQWRRRRKREGDGGGGGGGDGIGGGGGDGGSG-----------SG
Gryllus_bimaculatus             EERLVGNWKRGAGRG-------------------ARAPTGPQSGAMFSARNGHNSLRAARQGRRRRRERM*RGAQRRRRRAREAE------AVAVAMAAVEVAAA----TVXXXXXXX
Gryllus_longicercus             GERGAGPGERGAGSGARGAGSGAR-----------ARAPTGPQSGAMFSARNGHNSLRAARQGRRR*RRERMWRGAQRRRRRAREAE------AEAVAMAAVEVAAAVAVVTVEAAVVSA
Consensus

Teleogryllus_occipitalis        ------------------------------------------------------------------------------------------------------------------------
Acheta_domesticus               GSGGGGSGSDGGGGGNGSG---------SSGGGGSGGGGNGSGSSGGGGSGGGGGGGVDGGDGDGGDGGGGGDGGGGGGGGGGGSGGGGGGGGGGGGGGGGGGGGGGGGGSGGGGGGG
Gryllus_bimaculatus             XQGKGFF-----------------------------------------------------------------------------------------------------------------
Gryllus_longicercus             EAAAAVVTAEAAAVASAMAVVVAVAVAVAEAEAEAEAEAEAEAEAEAEAEAEAEAEAEAEAE----------------------------------------------------------
Consensus

Teleogryllus_occipitalis        ------------------------------------------------------------------------------------------------------------------------
Acheta_domesticus               GDGGGGGSGGSGGCKEAVAVRRSCSEW*KRRRKR-----------LSGSEC-----TR*CSVERLETEIRGS-IPFGSFPRLEEK*FL---KNGK*GTTLR------------------
Gryllus_bimaculatus             --------------------------WGVR-------------------------------------NNTTKGP---FLIFFN*KLKNKILKK*T*KIFNTERDNTKFNNEDRKKNANVFLD
Gryllus_longicercus             AEAEAEAEAEVAEAEVAVAARRL---WR*RRWWQWLTLYGMVGTVLERVEC**LYEVRLCS-EI*NRNLRQSCIRFRDFFFPEWKKYIFWRKESKVEAVLRT---------RRGADS*WS
Consensus

Teleogryllus_occipitalis        ------------------------------------------------------------------------------------------------------------------------
Acheta_domesticus               YLGSEERFILCLCMRVWSALARLRERVPACMRA----YVRARKR*RKRKEKEI*K--------------LNERATIGD*LGN---------------------------------------
Gryllus_bimaculatus             Y*NN--------------------------------VPRNNIFSPKKNNRKEQK--------------INTRAQIINKIGRLNFNYRAKIFLAFRNKNKQYNECETFFHAKIVHNLRNF
Gryllus_longicercus             YWYDEKLACVCVCVCGECAGAR------ACER*RE*DMERGG*RRIEREDRDV*RGRIETYREGG*RRIERERRDI--KIAR-----------------------------------------
Consensus

Teleogryllus_occipitalis        ------------------------------------------------------------------------------------------------------------------------
Acheta_domesticus               ----------------------KHKLSVRADVCPRDKER-------THR------FKDSEQTE---RGGQGTE------GDSRE--------NCWKR-------------INLNERRL
Gryllus_bimaculatus             SLAQFGFSSFPPKTQAGAFPSCTRLKLGLNVDHCTFLM*K*TFTFYFTHSLP**NCCKHSKQYSLYWQQKYSVT---FPLIKKKEFGYFSQ*IN-WRPSG***QDPTDLFSVFRIDGERL
Gryllus_longicercus             ----------------------ER*LALGADVCLR-*EK-------GH--------TGSKTVN---RQKEAVKVHAHK*VDSKE--------N-W*K-------------VNLNEKRL
Consensus

Teleogryllus_occipitalis        ------------------------------------------------------------------------------------------------------------------------
Acheta_domesticus               PETFS----------GPQMLSP--------GEHPSPNAGYRLH------INERRREMSNEENPF*IFERKLMCSME**RSFAFQVNLTKVYKCK*VNNCIIRNYENHTAHTSPMT*HGEL
Gryllus_bimaculatus             PNSYRILKLSKELDIGKHEIAP*NADPE*K*TKTGQFSQYKLR------LDLLLL-F*AASNIYRITELNFNNTQPF--SPVTKLLLGKA*PQKYIN-*LFPKEPIQTPHTN--------
Gryllus_longicercus             LEMLS----------GRQHLS----------SHLLGNVRHRMRSIVSM*MNFVQR-*THEENPF*ILRTKVN---------V*HVTITLL--------CI*LSRD--------------
Consensus

Teleogryllus_occipitalis        ------------------------------------------------------------------------------------------------------------------------
Acheta_domesticus               FPYTAQYYF*FC*SDGTLMNF*CFKYFLPIKKRQWKYKSN------------------------EKFRRE--DNL*NTEVLKRV-----------LA*CHEV*LH--------*RQFRV
Gryllus_bimaculatus             -------------EFSCLTNF--FKLSLYII-----WSAN*NSFL*AMA*KHLFSCFAAFMMQKVLAFPSTLLGNVLSLIFYKKLDGM*VTLT*VSMPLCMNVLIYVCVSSVCACARARV
Gryllus_longicercus             -------------SKGALNAF--FKYHFILILREDKLEAS--------------------M*EHNKL*KTHCSHLFNTITWRAL-----------PPHC-------------------
Consensus

Teleogryllus_occipitalis        ------------------------------------------------------------------------------------------------------------------------
Acheta_domesticus               RRGVVPVL---------------YSMH-NS*RFTRYCH-------------V*LARLTDVRGY------SQTNVGIL-----------P-----------KRTV*TALSSSS*IPKS
Gryllus_bimaculatus             CVCVCCVVCVCVCVVVVCVCVCKYFVIS-SLSIISRFLG-------SKKKKKKSGFRGSNTLRKYW-----IQ*NVTIIMHFAMHLR*CCPANVRRLLGGARHKRARRGEASRG------
Gryllus_longicercus             -----PIL---------------FLISLDEQYFTKFLEFQIFSRS*NKRQRKVQMERECSQRR*LVKYRRAKASFGLT-----------P*SLITLMTI*SEKRRRPCTLQRAQPFP--
Consensus

Teleogryllus_occipitalis        ------------------------------------------------------------------------------------------------------------------------
Acheta_domesticus               YI*FHLL*YFHWHHHRCRVIRLVNRRQFVKSILTR--------------IVLEFQEVGGTQNNAIIIKRH*VSEDVS*SDILNLDI*SCISASLI--------------SRKNI*FFH
Gryllus_bimaculatus             ----------------RRVERYEGERGLEGQSGRRR-------------FIID------DGDN*IPGQRPGIMPNI----F*SL-FP*QTSITSM------------------------
Gryllus_longicercus             ---FHALL--------PRVTRSAYRR--TRTLANKRWNSPNTNRVNSSVPLILDLQTV-----------HLISVLV----IFSL-APSSMSPDLIGISKTGP*NELLQE*TRIPRSTRH
Consensus

Teleogryllus_occipitalis        ------------------------------------------------------------------------------------------------------------------------
Acheta_domesticus               FI*FVI**RFRY*RKNVHKMTCIQNC-------------*SLIGNKLLMKNIS**N--------------SF*K*KANKWK*MTKFPC*LPSSLLTRNFHGLPK**FIGMHLPIQCYK
Gryllus_bimaculatus             --------------------------------------------------------------------------------FWRTLTRREC--SSFN*R*GF------------------
Gryllus_longicercus             KITPLS*YGIEYQKSILDKVF*IWMCKVEVQLF*VIEELYISFI*LNLLILNLFQPENMCTRRPEHRNCFTNKKFPRH*KIFLKSSLKIECRSPS*NN*PNFH------------------
Consensus

Teleogryllus_occipitalis        ---------------------------
Acheta_domesticus               TLQH*RKGGTAFSLKCLFIL*YMAGLK
Gryllus_bimaculatus             ---------------------------
Gryllus_longicercus             -LHYLKF------------IFNYAL-
Consensus
```

**Tocc.2G0002070.1_exon2**

```
Teleogryllus_occipitalis    ------------------------------------------------------------------------------------------------------------------------
Acheta_domesticus           ---------------------------------------------------------------------------------------------------------------------SRV
Gryllus_bimaculatus         HYSHRHFGIFIYVEITAGKTESDSCIRMCQEPAACNRA*PRDHLRRAPGQQQRPSERFSRCCERGLTCVCIREEQSGPHGATRAPPPCSPDEASGALRCYAHGGNMTSSQ*I*IDFTARI
Gryllus_longicercus         ------------------------------------------------------------------------------------------------------------------------
Consensus

Teleogryllus_occipitalis    ------------------------------------------------------------------------------------------------------------------------
Acheta_domesticus           HASES*EFIY-----------RNC----------CI---------------HFTYIIYVINTL*VMV*VTCDITF-----------ILKALSF---------------------------
Gryllus_bimaculatus         *SNRSNISIFLPNH*CDYSDIRICL*ICLIREA*CVIMKTSLIKCQSVY*WLFKYIIY-----------Q*ELTFNDIIKKLSYYV*TKYLHFSRGLCTGP*ETPHTHSHSFPGPCIQGL
Gryllus_longicercus         ---RSS-------------------------------------------------------------------SF------------*K*LH----------------------------
Consensus

Teleogryllus_occipitalis    ------------------------------------------------------------------------------------------------------------------------
Acheta_domesticus           KCSF------------------------------------FLFRIPY-------TISLYILRQKTNKTTTNKNT*NKFSSTRGDNGRLRSRVIRSN---YTSSNLNKEKGQINL*KLDIN
Gryllus_bimaculatus         KGSFT*T*YIYILYILCVSPTQNRLCKLWCR*HYFYS*RKFLLILIFPLSNSVNNLSLYILLQKTTTHINQQNT*NKF--SSKPSGDISNSLLRSKSN*IKSHE*N--------------
Gryllus_longicercus         ---------------------------------------------------QITRVEL*I--------------------RRKDK*PFKN*KQRGK------------------------
Consensus

Teleogryllus_occipitalis    ------------------------------------------------------------------------------------------------------------------------
Acheta_domesticus           EEKKKTG*TTL*KLKIKGVKEEKRINNSLETRDKMKKKKLGQITF**NRDK*VK--VFLFLFQ----------FFPQQNEELATQNIPCNPKATSSAHLGSARQRNRP*PPRAIRKLCKG
Gryllus_bimaculatus         ------------------FK*EERINNYLQTK-*NEEKKTG*IPL*ETRDKVRKKSVLYFLRHKNKNVTPKHPLQPQRSLECASTE---RARASATAH-------NR---PHAIRKLCKG
Gryllus_longicercus         ----------------------DRRNTSLRN*-R*SEKKF----------------VLYFLRHKN*NVIPEHPLQPQRSLECASTE---RARASATAH-------NR---PHAIRKLCKG
Consensus

Teleogryllus_occipitalis    ------------------------------------------------------------------------------------------------------------------------
Acheta_domesticus           RRANA------------------GRGA-----KTPSN*IIGKPSN*RETLIKINLAHNMRATMTGTRRAETICPV-------ALARSQPMLPRNCTLLTLS*NIKQFQNPFQI-------
Gryllus_bimaculatus         KRADARPRGQDAFELNYWEAFQLKRGAHKNKFSTQYASDDDRPP*SRNHLPR-------RLGSPATDAPKELCAAQGLLKY*AISKPSPNY--------IK*KLGHFKTPV*I*ARG*L*
Gryllus_longicercus         KRADARPRGQDAFELNYWETFQLKRGAHKNKFSTQYASDDDRPP*SRNHLPR-------RLGSVATDAPKELCAAQSLLKY*AISKPSPNY--------IK*KLGHFKTPV*I*TRG*L*
Consensus

Teleogryllus_occipitalis    ------------------------------------------------------------------------------------------------------------------------
Acheta_domesticus           -----T*NEN*D---------------------------ILKRRFKFRPAVNYRNRTACN*TEMRSQPFLKDAYAKYINELNFNTNSCEQRTLCWLENLASVE*I*LRRQTQ*IGENGEA
Gryllus_bimaculatus         KPNGL*LN*NAESAVSERRLCEIHQ*T*F*P*QLQTAYTVLARKFGFRRMNLIAKTNTINRREWRS-----------------------RRIV*VINNSICVT----RRRV---------
Gryllus_longicercus         KPNGL*LN*NAESAVSERRLCEIHQ*T*F*H*QLRTAYTVLARKFGFRRMNLIAKTNTINRREWRS-----------------------RRIV*VIYNSICVT----RRRV---------
Consensus

Teleogryllus_occipitalis    ------------------------------------------------------------------------------------------------------------------------
Acheta_domesticus           GELCG*LIITFV*FADEFALTMSLKKGQFKSRFKMC*GEFILQITLIF-QC*ALISQCLISNSS------------VKM*FLCQNK-KIRSLIQKKKRKI--------Y*EKILHY*KKV
Gryllus_bimaculatus         -----------------LA*HRL*KKGSLNS---------VLNILITYLSVRSYYTSC*YVNVKRLYFNVQSRILVVKI*FLRQKKIKIHILNTRETRKII*SEIPLS**ETYY*YE*KR
Gryllus_longicercus         -----------------LA*HRLYKKGSLNS---------VLNILIIYLSVRSYYTSC*YFNVKLLYFNV*SRILGVKI*FLRQKKTTIHILNTRETRKII*S*IPLY**KNILWIRIKT
Consensus

Teleogryllus_occipitalis    ------------------------------------------------------------------------------------------------------------------------
Acheta_domesticus           D--IHS*PY-LNHTVTCDQNSVFLVISKILQKKSTRNIPFKEATLLKRNSNE*KLLFT*FPHF*LLPQS*LSDFLKPHVKEWKRFKTSIRN*FRSPGDV-LKLILS*FL-----------
Gryllus_bimaculatus         ---VYSFTT*LNHSS--DQYRCFLVN*KMLQKK------FYMLYLIRRNETIEKLFFTQFYNF*YLPQS*LFDFLKSHVKCGIKFKSKLI*FTRECIKIYIILILE**TLIQKQYIRTNI
Gryllus_longicercus         SLFIHNLT*SFIRSI--*RFSSELKN--VTEKI------LHAVFNKKKQDY*EALFYT-------IP-----*FLISSTIVII*F-SKFTCKAWNKV*LEIDLVHEGMH-*NP*YSHTRM
Consensus

Teleogryllus_occipitalis    ------------------------------------------------------------------------------------------------------------------------
Acheta_domesticus           ---NDKL*YEKQ*----------VQRNATSCMRSFL*KPVSD*GLKFTNRPV-------AF*DYSVLYLALLFPILRKI--------------TS--------KC*KRQKPQQEKTCLLP
Gryllus_bimaculatus         FNKNNKIFKKYSRRYFYETATMKV*TNRFSV*EFH---------T*FTTRLI-------ALYNS*VLYLALLFPLLCKN--FDQLE-----------------------KTSTRKASLLL
Gryllus_longicercus         INFNTKTIHLNSY----------IQSK*LTS*EI------------FSTLLLLRDCSYGSLNKSL---LSLRIPHLIPNPHYSTLQLLSIIFGTSVSLTLQKLRSVRKDKTSTRKACLLP
Consensus

Teleogryllus_occipitalis    ------------------------------------------------------------------------------------------------------------------------
Acheta_domesticus           FSLGTLFGAASQSSIRPLPDNSRANPRRRSP----EERLMGNW-----ERSAGRGAR----------------GAGAHRATKRRYV*RTEWAQQPTGGAPGAAEAARRRERMWRGAQWRR
Gryllus_bimaculatus         FSLGTLVSAASQSPIRPFPIT-------RGPILGAVCRRSGSWGIGSGERGAG---RGRPQGHKAALCLAHGMGTTAY------------------------------------------
Gryllus_longicercus         FSLGTLVSAASQSSIRSFPIT-------RGPILGAVSRRSGSWGIGSGERGAGSGARGAGRGERGAGSGERGAGAGAHRATKRRYV*RTEWAQQPTGGAPGAASVA-------EGADVER
Consensus

Teleogryllus_occipitalis    ---------------------------------------EGGDGG--------------------------------------------------------------------------G
Acheta_domesticus           ----RRKREGDGGGGGGGDGIGGGGGDGGSGSGGSGGGGSGSDGGGGGNGSGSSGGGGSGGGGNGSGSSGGGGSGGGGGGGVDGGDGDGGDGGGGGDGGGGGGGGGGGSGGGGGGGGG
Gryllus_bimaculatus         -----------------------------------GRRARGGDGGG-----------------------------------------------------------------------GSG
Gryllus_longicercus         STAAKAAREG-GGGGGGGD-----------GGGRGGGGSGGGDGGGG---------GGVGGGG------GGGGDGGGGGGGGF-------GDGGGG---GGGGGGGGGGGGGGGGGGGGGG
Consensus                                                            * ***                                                                          *

Teleogryllus_occipitalis    CGEAHTGGG---------------------------------------------------------------------------------------------------------------
Acheta_domesticus           GGGGGGGGGGGGGGGSGGGGGGGGGDGGGG-----------GSGGSGGC--KEAVAVRRSCSEW*KRRRKRLSGSECTR*CSVERLETEIRGS----------IPFGS------------
Gryllus_bimaculatus         CEEEHSGEGGARGRRRRWRWRWQR*RWRRRRXXXXXXXXXKGRDFFGGF---ETTQQKALF*FFSI----------KS*KIKY*KNEP------EKFS-----IPKGTIQNLITKTEKKT
Gryllus_longicercus         GGGGGGGGGGGRGGGRGGGRGGGRGRGRGR-----------GG*SRGGCCREETVAVAAVVAVVDVVRNGR-NGVGKSRVLVIVRGEAVFRDMKPKFAAIVHPIPG--------------
Consensus                         * *

Teleogryllus_occipitalis    ------------------------------------------------------------------------------------------------------------------------
Acheta_domesticus           ---FPRLEE-----K*FLKNGK*GT----------------------TLRYLGSEE-------RFILCLCMRVWSAL------------------ARLRERVP----ACMRAYV------
Gryllus_bimaculatus         LMFF*TIETTSPVIIYFPQKKIIGK-NKK*IQEHKL*IKLGALILIIVQRFFWRFEIKINNTMNVKLFFMLRLYTICEIFH*PNLVFPHFHPKPKPALFQVAPG*NSVSMLIIALF*CKS
Gryllus_longicercus         -LFFSRMEE-----IYFLEKGE*GRGGPKDTQRCRF------IVVILV*R-------------KISLCLCMRVW*VC----------------------------GGARMRAIE------
Consensus

Teleogryllus_occipitalis    ------------------------------------------------------------------------------------------------------------------------
Acheta_domesticus           -----RARKR*RKRKEKEI*KLNERATIG----------------D*LGNKHKLSVRADVCPRD--------------------------------------KERTHRFKDSEQTERGGQ
Gryllus_bimaculatus         EPLHFTLRIHYHNKTAANIP--NNILCIGSKSTPLPFH*SRKKNLDIFHNRSIG------DPRDNNSKILPTYFQFLE*MENACQIAIVS*NYQKNWILANMKLHLKMLIQSESEQKQAN
Gryllus_longicercus         -------RVRYGER-------------------------------GIETNREGG*RRIEREDRDV*RGRIKTYREGKERYKN-CARAIIGTRSGR---VSKVREGTHRFKDSEQTERGGQ
Consensus

Teleogryllus_occipitalis    ------------------------------------------------------------------------------------------------------------------------
Acheta_domesticus           GTEGDSRE-NCWKRINLNERRLPETFSG-------PQMLSPGEHPSPNAGYRLHIN-------ERRREMSNEENP------------F*IFERKLMCSME**RSFAFQVNLTK-------
Gryllus_bimaculatus         FLNTNF----GWICCYYSKRHQIYIASLN*ILTT------------PN---RFH---------Q*QNSFLEKPSP---------------------------------------------
Gryllus_longicercus         GTRT*ISGFKGELVKSQPEREA---ASGNVIGTAAPQQPSPGKCSPPNAVYRLHVNELRTEMNTRRKSILDSPNES*CVACNNNASLYLVVERQ*R-SIECIF*VSFHINFTRG*TGS*H
Consensus

Teleogryllus_occipitalis    ------------------------------------------------------------------------------------------------------------------------
Acheta_domesticus           VYKCK*VNNCIIRNYENHTAH-TSPMT*HGELFPYTAQYYF*FC*SDGTLMNF----------*CFKYFLPIKK----RQWKYKSNEKFRREDNL*NTEVLKRVLA*CHEV---------
Gryllus_bimaculatus         -------KNTLTNYFQ------RNPFKLH-------------------IQMNF--RALLIFLNSHYTLFGPLTEIPSFKQWR-------------KSISLVASRHS*CRKY*HFPQHCLE
Gryllus_longicercus         VGA*QIMKNTLLTLVQYYNMESSSPTLPN-------------------IIFNFVRRTVLY*IFRVSNIFS*LE*TPA--EST-------------NGTRMFAEKIT-CEI-----QKG*S
Consensus

Teleogryllus_occipitalis    ------------------------------------------------------------------------------------------------------------------------
Acheta_domesticus           --------------*LH*RQFRVRRGVVPVLYSMHNS*RFTRYCHV*LARLTDVRGYSQTNVGILPKRTV*-------TALSSSS*IPKSYI*FHLL*YFHWHHHRCRV------IRLVN
Gryllus_bimaculatus         MCFP*FFIKNWMACE*HLRKYL-------CLYA*M-------Y*FMCV*ALC-VRARARACVCVCAVWCVCVCVWWWCVCVCVSILLYHH---FQSLVDF*EAKRKKKKAGFEDPIRYAN
Gryllus_longicercus         EFWPNAMKSNYTNDNLE*EKA*-------ALYST-----------------------ARTTLSVSRV-----------VATCNSLGLQTYEDTRKQTLEFSQHE*SKQ-------LRPLN
Consensus

Teleogryllus_occipitalis    ------------------------------------------------------------------------------------------------------------------------
Acheta_domesticus           ------------------------RRQ----------FVKSILTRIVL------------EFQEVGGTQNNAIIIKRH*VSEDVS*SDILNLDI*SCISASLISRKNI*FFH---FI*FV
Gryllus_bimaculatus         IGYNKTSP*LCTSQCI*G---NVARRTCGGCWAEPGTSVRGEARRAGAGGSSDMKASGGWKGNREGGGS**MMVIIKYPGNARESCQISFNL--CSRN--------------------K
Gryllus_longicercus         LRSPNRTFNFCSCNIFIGAVIDVARFD----WDIEDRSMK*TFTRID-------------ENTKKYEAQNNAIIIIWHRVSEVHS*QSILNLDV*SRSTIILSNRGIIYFFYLTQFVNFK
Consensus

Teleogryllus_occipitalis    -------------------------------------------------------------------------------------------------------------------
Acheta_domesticus           I**R--FRY*RKNVHKMTCIQNC*SLIGNKLLMKNIS**NSF*K*KANKWK*MTKFPC*LPSSLLTRNFHGLPK**FIGMHLPIQCYKTLQH*RKGGTAFSLKCLFIL*YMAGLK
Gryllus_bimaculatus         LP*RPCF---GEH*QEEN-------------ALHSISDEVF--------------------------------------------------------------------------
Gryllus_longicercus         LSAR-------EHVHTKTRAQKLEH**KISASLKNIFKVLFKNRVPITELK*LTKFPSSLP----*RNFQ-------------------------------LRS-----------
Consensus
```

**Tocc.2G0002070.1_exon3**

```
Teleogryllus_occipitalis    ------------------------------------------------------------------------------------------------------------------------
Acheta_domesticus           ------------------------------------------------------------------SRVHASE-----------------------------------------------
Gryllus_bimaculatus         PTIRTAISAFSFTSKLPRARLRVTAAYACAKNRRLAIEHDREITFAARPDSNRGPQSVSRVAANGG*RVFASERNRAAHTAPPVHPPPAAPTRPLGHFGAMHMGAT*HLRNEFR*ISPQE
Gryllus_longicercus         ------------------------------------------------------------------------------------------------------------------------
Consensus

Teleogryllus_occipitalis    ------------------------------------------------------------------------------------------------------------------------
Acheta_domesticus           --S*EFIYRNCCIHFTYIIYVI------------------------------------------------NTL*VMV*VTC---------------------------------------
Gryllus_bimaculatus         YNQIEATYQYFCLTINAIILILEYVFRYV*YGKHDALL*KHH**NAKVCTNGYLNI*FTSKN*LLMT*LRN*VITYKQSICIFREDYAQGLKKHRIRILIVSRVHASKG*RGHLPERDIY
Gryllus_longicercus         ------------------------------------------------------------------------------------------------------------------------
Consensus

Teleogryllus_occipitalis    ------------------------------------------------------------------------------------------------------------------------
Acheta_domesticus           ---------------------DITFILKALSFKCSFFLFRIPYTISL-YILRQKTNKTTTNKNT*NKFSSTRGDNGRLRS-----------RVIRSNYTSSNLNKEKGQINL*KLDINEE
Gryllus_bimaculatus         TFYTYYVCHQHRTDFVSYGVGDITFILEESSF*F*FSLFRIL*TISLSTFFCKKQQHTSTNK--------TREISSRVNQVVTSPTAFFVLRVIRSNHTSRTLNKKKG*ITIYKLNRTRK
Gryllus_longicercus         --------------------------------------------------------------------------------------------------RSSSF*K*LHQITRVEL*IRRK
Consensus

Teleogryllus_occipitalis    ------------------------------------------------------------------------------------------------------------------------
Acheta_domesticus           KKKTG*TTL*KLKIKGVKEEKRINNSLETRDKMKKKKLGQITF**NRDK*VKVFLFLFQFFPQQNEELATQNIPCNPK-----ATSSAHLGSARQRNRP--*PPRAIRKLCKGRRANA--
Gryllus_bimaculatus         RRQDK-YLFKKLEIK*-----------------EKNP--------------------FYIFFATKIKTSPQNIPCNPN-----AASSAHLRSERAPAQPPITAPTQSENYAKGSAQTP--
Gryllus_longicercus         DK----*PFKN*KQRG---KDRRNTSLRN*R*SEKK-------------------FVLYFLRHKN*NVIPEH-PLQPQRSLECASTERARASATAHNR-----PHAIRKLCKGKRADARP
Consensus

Teleogryllus_occipitalis    ------------------------------------------------------------------------------------------------------------------------
Acheta_domesticus           ----------------GRGA-----KTPSN*IIGKPSN*RETLIKINLAHNMRATMTGTRRAETICPV-------ALARSQPMLPRNCTLLTLS*NIKQFQNPFQI------------T*
Gryllus_bimaculatus         ----------------GREA-----KTPSN*IIGKPSN*REALIKINLAHNMRATTTGPRRAETICPV-------ALARPPPMLLRNCALLRAS*NIKQFRNPLRI------------T*
Gryllus_longicercus         RGQDAFELNYWETFQLKRGAHKNKFSTQYASDDDRPP*SRNHLPR-------RLGSVATDAPKELCAAQSLLKY*AISKPSPNY--------IK*KLGHFKTPV*I*TRG*L*KPNGL*L
Consensus

Teleogryllus_occipitalis    ------------------------------------------------------------------------------------------------------------------------
Acheta_domesticus           NEN*D---------------------------ILKRRFKFRPAVNYRNRTACN*TEMRSQPFLKDAYAKYINELNFNTNSCEQRTLCWLENLASVE*I*LRRQTQ*IGENGEAGELCG*L
Gryllus_bimaculatus         NENSD---------------------------ILKRQFKFKPAVNYENPTACN*TEMPSRQFLKDAYAKYINELNFNPNSCKQRTLCWLENLASEE*I*LRRQTQ*IGENGEAGELCR*L
Gryllus_longicercus         N*NAESAVSERRLCEIHQ*T*F*H*QLRTAYTVLARKFGFRRMNLIAKTNTINRREWRSRRIV*VIY----------------NSIC-----------VTRRRV----------------
Consensus

Teleogryllus_occipitalis    ------------------------------------------------------------------------------------------------------------------------
Acheta_domesticus           IITFV*FADEFALTMSLKKGQFKSRFKMC*GEFILQITLIF-QC*ALISQCLISNSS-----------------VKM*FLCQNK-KIRSLIQKKKRKIY*EKILHY*KK--------VDI
Gryllus_bimaculatus         IIPFV*HAEEYWLNTGSKKRAV-----------*IQF*IF*LHISACVHIIRHVNTSMLSVYILMFNL--EYLS*KFNFYAKRKSKFISSIQEKREKLFNQKFHFLDKKHIINTNKNESI
Gryllus_longicercus         ----------LA*HRLYKKGSLNS---------VLNILIIYLSVRSYYTSC*YFNVKL-----LYFNV*SRILGVKI*FLRQKKTTIHILNTRETRKII*S*IPLY**KNIL----WIRI
Consensus

Teleogryllus_occipitalis    ------------------------------------------------------------------------------------------------------------------------
Acheta_domesticus           HS*P-YLNHTVTCDQNSVFLVISKILQKKST-RNIPFK--EATLLKRNSNE*KLLFT*FPHF*LLPQS*LSDFLKPHVKEWKRFKTSI--------RN*FRSPGDVLKLILS*FLNDKL*
Gryllus_bimaculatus         HSQLDLIIHPINID---VF**IEKCYRKNST-CCI**E------------ETRLLRSSFLHNSIISNI-------FHNRNYLIF*NHM*NVE*SLSRN*FSSRGNALKFILFSYLNDKL*
Gryllus_longicercus         KTSL-FI-HNLT*S------FIRSI*RFSSELKNVTEKILHAVFNKKKQDY*EALFYTIP*FLISSTIVII*FSKFTCKAWNKV*LEIDLVHEGMH*NP*YSHTRMIN------FNTKTI
Consensus

Teleogryllus_occipitalis    ------------------------------------------------------------------------------------------------------------------------
Acheta_domesticus           YEKQ*VQ-----------RNATSCMRSF----L*KPVSD*GLKFTNRP---------VAF*DYSVL-YLALLFP------------ILRKITSKC*KRQKPQQEKTCLLPFSLGTLFGAA
Gryllus_bimaculatus         YKNNTFEPIYSIKITKFLRNILDATSTRPQL*KSKQIASQSE---NSTLNSQPVL*HFTTPEYYTW---HFCFPYYAKTSIS*KRPQLEKLAYYCFRSER**V-----------------
Gryllus_longicercus         HLNSYIQ-----------SK*LTS*EIFSTLLLLRDCSYGSL---NKSLLSLRIPHLIPNPHYSTLQLLSIIFG----TSVS---LTLQKLRS--VRKDKTSTRKACLLPFSLGTLVSAA
Consensus

Teleogryllus_occipitalis    ------------------------------------------------------------------------------------------------------------------------
Acheta_domesticus           SQSSIRPLPDNSRANPRRRSP----EERLMGNW-----ERSAGRGAR----------------GAGAHRATKRRYV*RTEWAQQPTGGAPGAAEAARRRERMWRGAQWRR----RRKREG
Gryllus_bimaculatus         -QHHNRPSAPSR*LEGQSSAPFAGGAAR---------GELEAGSGAR----------------GAGAHRATKRRYV*RTEWAQQPTGGAPGAATAA-------EGADVKRSTAAKAARE-
Gryllus_longicercus         SQSSIRSFPIT-------RGPILGAVSRRSGSWGIGSGERGAGSGARGAGRGERGAGSGERGAGAGAHRATKRRYV*RTEWAQQPTGGAPGAASVA-------EGADVERSTAAKAAREG
Consensus

Teleogryllus_occipitalis    ----------------------------------------------------------------------------------------GGGGGGGGRSDGGRSGSGSGRSGSGGGGGGGG
Acheta_domesticus           DGGGGGGGDGIGGGGGDGGSGSGGSGGGGSGSDGGGGGNGSGSSGGGGSGGGGNGSGSSGGGGSGGGGGGGVDGGDGDGGDGGGGGGDGGGGGGGGGGSGGGGGGGGGGGGGGGGGGG
Gryllus_bimaculatus         -----------------------------------------------------------------------------------------------------------------GGGGGGG
Gryllus_longicercus         -GGGGGGGDG-------GGRGGGGSGGG----DGG------------------------GGGGVGGGGGGG-----GDGGGGGGGFDGGGGGGGGGGGGGGGGGGGGGGGGGGGGGGGGG
Consensus                                                                                                                    *******

Teleogryllus_occipitalis    GGGGGRGG----------------------------------------------------------------------------------------------------------------
Acheta_domesticus           GGGGGSGG--GGGGGGGDGGGGGSGG--SGGC--KEAVAVRRSCSEW*KRRRKRLSGSECTR*CSVERLETEIRG----------SIPFGSFPRLE------EK*FLKNGK*GTTLRYLG
Gryllus_bimaculatus         GDGSGRGG--GGDGXXXXXXXXXXREGIFLGGS---------------KQHNKR---------------------------------PFFNFFQLKVKK*NIKKMNLKNFQYRKG-----
Gryllus_longicercus         GGGGGRGGGRGGGRGGGRGRGRGRGG*SRGGCCREETVAVAAVVAVVDVVRNGR-NGVGKSRVLVIVRGEAVFRDMKPKFAAIVHPIPGLFFSRME------EIYFLEKGE*GRG----G
Consensus                   * * * **

Teleogryllus_occipitalis    ------------------------------------------------------------------------------------------------------------------------
Acheta_domesticus           SEE----RFI----------LCLCMRVWSAL--ARLR--ERVPACMRAYVRARKR*RKRKEKE--------I*KLNE---------RATIGD*LGNKHKLSVRADVCP------RDKERT
Gryllus_bimaculatus         -------QYKI**RRQKKKR*CF----------FRLLK-QRPP---**YIFPKKK**ERTKNK--------YKSTNY-------------------------K*NWAP*F*LSCKDFSGV
Gryllus_longicercus         PKDTQRCRFIVVILV*RKISLCLCMRVW*VCGGARMRAIERVRYGERGIETNREGG*RRIEREDRDV*RGRIKTYREGKERYKNCARAIIGTRSGRVSKV----------------REGT
Consensus

Teleogryllus_occipitalis    ------------------------------------------------------------------------------------------------------------------------
Acheta_domesticus           HRFKDSEQTERGGQGT-------EGDSRENCWKRINLNERRLPETFSG-------PQMLSP------GEHPSPNAGYRLHINERRREM-------------------SNEENPF*IFERK
Gryllus_bimaculatus         SK*K*TIQ--------------------------------*M*NFFSC*DCTQFAKFFISPIWFFLISTQNPSRRFSKLHPAKTRSQC*SLHFF-------------NVKVNLYILL---
Gryllus_longicercus         HRFKDSEQTERGGQGTRT*ISGFKGELVKSQPER---------EAASGNVIGTAAPQQPSP------GKCSPPNAVYRLHVNELRTEMNTRRKSILDSPNES*CVACNNNASLYLVVERQ
Consensus

Teleogryllus_occipitalis    ------------------------------------------------------------------------------------------------------------------------
Acheta_domesticus           LMCSME**RSFAFQVNLTK-------VYKCK*VNNCIIRNYENHTAHTSPMT*HGELFPYTAQ------------YYF*FC*S------------------------------DGTLMNF
Gryllus_bimaculatus         ----------YAF---------------------TTIIKLLQ---------TFQTIFFVLAAKVLRYLSTNQEKRIWIFFTIDQLETLGIIIARSYRPIFSF*NRWRTPA----------
Gryllus_longicercus         *R-SIECIF*VSFHINFTRG*TGS*HVGA*QIMKNTLLTLVQYYNMESSSPTLPNIIFNFVRRT----------VLY*IFRVSN--------------IFS*LE--*TPAESTNGTRM--
Consensus

Teleogryllus_occipitalis    ------------------------------------------------------------------------------------------------------------------------
Acheta_domesticus           *CFKYFLPIKK-RQWKY-----KSNEKFRREDNL------------*NTEVLKRVLA*CHEV*LH-----------*RQFRVRRGVVPV-LYSMHNS*RFTRYCHV*LARLTDVRGYSQT
Gryllus_bimaculatus         K*LSYLETIKRIGYWQT*NCTLKC*SRVKVNKNR------PIFSIQTSVGSAVIILSGIKYISHH--------*IKF*QH-------PT-VFTSNKTPSWKS------------------
Gryllus_longicercus         FAEKITCEIQK-G*SEFWPNAMKSNYT---NDNLE*EKA*ALYSTARTTLSVSRVVATCNSLGLQTYEDTRKQTLEFSQHE*SKQLRPLNLRSPNRTFNFCS-CNIFIGAVIDV------
Consensus

Teleogryllus_occipitalis    ------------------------------------------------------------------------------------------------------------------------
Acheta_domesticus           NVGILPK---RTV*TALSSSS*IPKSYI*FHLL*YFHWHHHRCRVIRLVNRRQFVKSIL-----------TRIVLEFQEVGGTQNNAIIIKRH*VSEDVS----*SDILNLDI*SCISAS
Gryllus_bimaculatus         ---LAPKIH*LTISKGTHSNSTYK*IFVPY*FF*TLIIH-YLVR*LKFLPLSNGVKASL*LLRGIHDAESTSISLNTAWKCAFPNFL*KIGWHVSNTYVSIYAFMHECINLCVCE-LCVC
Gryllus_longicercus         ---------------------------------ARFDWD---------IEDRS-MK*TF-----------TRID-ENTKKYEAQNNAIIIIWHRVSEVHS----*QSILNLDV*SRSTII
Consensus

Teleogryllus_occipitalis    ------------------------------------------------------------------------------------------------------------------------
Acheta_domesticus           LISRKNI----------------------*FFHFI*FVI**RFRY-*RKNVHKMTCIQNC*SLIGN-----KLLMKNIS**NSF*K----------------------------------
Gryllus_bimaculatus         VRARARVCVCVLCGVCVCVCGGGVCVCV*VFCYIITFNH*SIFRKQKEKKKKRVSRIQYVTQILDTIKRHHNYALRNASKVMLPGERAAVAGRSPAQACEARRGEQGQEGRAI*RRAGVG
Gryllus_longicercus         LSNRGII----------------------YFFYLTQFV---NFKLSAREHVHTKTRAQKLFH**KI-----SASLKNIFKVLFKNR----------------------------------
Consensus

Teleogryllus_occipitalis    -------------------------------------------------------------------------------------------
Acheta_domesticus           --------------------*KANKWK*MTKFPC*LPSSLLTRNFHGLPK**FIGMHLPIQCYKTLQH*RKGGTAFSLKCLFIL*YMAGLK
Gryllus_bimaculatus         RAIGKAEVHNR*W**LNTRATPGNHAKYLLIF---VP----VTNFH--------NVHV-------LENTNK------KRMLFIQLAMRFF-
Gryllus_longicercus         --------------------VPITELK*LTKFPSSLP----*RNFQ-------------------------------LRS-----------
Consensus
```

**Tocc.2G0002070.1_exon4**

```
Teleogryllus_occipitalis        ------------------------------------------------------------------------------------------------------------------------
Acheta_domesticus               ------------------------------------------------------------------SRVHASES----------------------------------------------
Gryllus_bimaculatus             PTIRTAISAFSFTSKLPRARLRVTAAYACAKNRRLAIEHDREITFAARPDSNRGPQSVSRVAANGG*RVFASERNRAAHTAPPVHPPPAAPTRPLGHFGAMHMGAT*HLRNEFR*ISPQE
Gryllus_longicercus             ------------------------------------------------------------------------------------------------------------------------
Consensus

Teleogryllus_occipitalis        ------------------------------------------------------------------------------------------------------------------------
Acheta_domesticus               ---------------------*EFIYR---------------------------------------------------NCCIH------------------------------------F
Gryllus_bimaculatus             YNQIEATYQYFCLTINAIILILEYVFRYV*YGKHDALL*KHH**NAKVCTNGYLNI*FTSKN*LLMT*LRN*VITYKQSICIFREDYAQGLKKHRIRILIVSRVHASKG*RGHLPERDIY
Gryllus_longicercus             ------------------------------------------------------------------------------------------------------------------------
Consensus

Teleogryllus_occipitalis        ------------------------------------------------------------------------------------------------------------------------
Acheta_domesticus               TYIIYVI----NTL*VMV*VTCDITFILKALSFKCSFFLFRIPYTISL-YILRQKTNKTTTNKNT*NKFSSTRGDNGRLRS-----------RVIRSNYTSSNLNKEKGQINL*KLDINE
Gryllus_bimaculatus             TFYTYYVCHQHRTDFVSYGVG-DITFILEESSF*F*FSLFRIL*TISLSTFFCKKQQHTSTNK--------TREISSRVNQVVTSPTAFFVLRVIRSNHTSRTLNKKKG*ITIYKLNRT-
Gryllus_longicercus             ----------------------TVFFVLK----------------------------------------------------------------VITSNHTSRTLNKKKG*ITI*KLETT-
Consensus

Teleogryllus_occipitalis        ----------------------------------------------------------------------------------------AAQRRRP-------------------------
Acheta_domesticus               EKKKTG*TTL*KLKIKGVKEEKRINNSLETRDKMKKKKLGQITF**NRDK*VKVFLFLFQFFPQQNEELATQNIPCNPKATSSAHLGSARQRNRP*P---PRAIRKLCK--GRRANAGRG
Gryllus_bimaculatus             RKRRQDKYLFKKLEIK----*EKNPFYIFFATKIKTS-------------------------P--------QNIPCNPNAASSAHLRSERAPAQP-PITAPTQSENYAK--GSAQTPGRE
Gryllus_longicercus             -RKRQEKYLFKKLEIK----*EKIRFIFSSPQKLKRH-------------------------PRTSPATPTQ-----PRV--RIYGASARQRNRP*P---PPRNPKIMQREARR----RP
Consensus                                                                                                                     *

Teleogryllus_occipitalis        ------------------------------------------------------------------------------------------------------------------------
Acheta_domesticus               AKTPSN*IIGKPSN*RETLIKINLAHNMRATMTGTRRA-----ETIC------PVALARSQPMLP------RNCTLLTLS*NIKQFQNPFQIT*NEN*DILKRRFKFRPAVNYRNRTACN
Gryllus_bimaculatus             AKTPSN*IIGKPSN*REALIKINLAHNMRATTTGPRRA-----ETIC------PVALARPPPMLL------RNCALLRAS*NIKQFRNPLRIT*NENSDILKRQFKFKPAVNYENPTACN
Gryllus_longicercus             AERPRR------------LRIKLLGNLP---TEERRS*K*I*HTICERRRQAPV---EQKPFAPSPWLGRH*C-----S*GIVRCSEPLKIL--SNFETLAELHKM------KTRTF*N
Consensus

Teleogryllus_occipitalis        ------------------------------------------------------------------------------------------------------------------------
Acheta_domesticus               *TE-------MRSQPF---LKDAYAKYINELNFNTNSCEQRTLCWLENLASV-E*I*LRRQ------TQ*IGENGEAGELC-G*LIITFV*FADEFALTMSLKKGQFKSRFK----MC*G
Gryllus_bimaculatus             *TE-------MPSRQF---LKDAYAKYINELNFNPNSCKQRTLCWLENLASE-E*I*LRRQ------TQ*IGENGEAGELC-R*LIIPFV*HAEEYWLNTGSKKRAV*IQF*I-------
Gryllus_longicercus             ASLNLDPRLIMKTQRLVIKLKCRVGSF*KTLMRNTSMNLILTLTAANSVHCVGSKIWLQKNEFNCEDKHNK*ERMEKPENCVGNL*FHLRDTPKSFGLTQAL*KGQFKFSFKYFNYISQR
Consensus

Teleogryllus_occipitalis        ------------------------------------------------------------------------------------------------------------------------
Acheta_domesticus               EFILQITLIFQC*ALISQCLISN----------SSVKM*FLCQNKKIRSLIQKKKRK------------IY*EKILHY*KKVDIHS*PYL-NHTVTCDQNSVFLVISKILQKKSTRNIPF
Gryllus_bimaculatus             -F*LHISACVHIIRHVNTSMLSVYILMFNLEYLS*KFNFYAKRKSKFISSIQEKREK----LFNQKFHFLDKKHIINTNKNESIHSQLDLIIHPINID---VF**IEKCYRKNSTCCI**
Gryllus_longicercus             AFILYIMLILQC*AFIFQCLISN----------TWRENLISTPKENHNSYPQYKRNAKNYLIINST--LLIKKHIMDTNKNESIHSQPDLIIHPINIE---VF**IEKCYRKNSTCCI**
Consensus

Teleogryllus_occipitalis        ------------------------------------------------------------------------------------------------------------------------
Acheta_domesticus               KEATLLKRNSNE*KLLFT*FPHF*LLPQS*LSDFLKPHVKEWKRFKTSI--------RN*FRSPGDVLKLILS*FLNDKL*YEKQ*VQ-----------RN---ATSCMRSFL*KPV---
Gryllus_bimaculatus             EETRLLRSS---------------FLHNSIISNIF--HNRNYLIF*NHM*NVE*SLSRN*FSSRGNALKFILFSYLNDKL*YKNNTFEPIYSIKITKFLRNILDATS-TRPQL*KSKQIA
Gryllus_longicercus             EETRLLRSS---------------FLHNSIISNIF--HNRNYLIF*IHM*SVE*SLTRN*FGSRRNALKSIIFSY*NDKL*YKNNTFEFIYSIKITNFLRNILDATSSTRL*LWKSKQIA
Consensus

Teleogryllus_occipitalis        ------------------------------------------------------------------------------------------------------------------------
Acheta_domesticus               ---------SD*GLKFTNRPVAF*DYSVLYLALLFPILRKITSKC*KRQKPQQEKTCLLPFSLGTLFGAASQSSIRP-LPDNSRANPRRRSPEERLMGNWE-------------RSAGR
Gryllus_bimaculatus             SQSENSTLNSQPVL*HFTTP----EYYTWHFC--FPYYAK-TSIS*KR--PQLEKLAYYCF-----------RSER**VQHHNRPSAPSR*LEGQSSAPFAGGAARGE-------LEAGS
Gryllus_longicercus             SQSENSTLNSQPAL*HFTTP----EYYIWHFC--FPYFAK-TSIS*KGQ--NLDKKSLLTAVFARNASKCSITIVHPLLPDNSRANPRRR*PEERLVGNWKRGAGSGERGPGSGARGAGR
Consensus

Teleogryllus_occipitalis        ----------------------------------------------------------------------------------------RRRRW--------------WRQRRRRW-----
Acheta_domesticus               GARGAG-------AHRAT--------------------------------------------------------------KRRYV---*RTEWAQQPTGGAPGAAEAARRRERMW-RGAQ
Gryllus_bimaculatus             GARGAG-------AHRAT--------------------------------------------------------------KRRYV---*RTEWAQQPTGGAPGAATAA---------EGAD
Gryllus_longicercus             GERGAGRGRGRPQGHKAALCLAHGMGTTAYGRRARGGVGSGGSGCGEEHSGEGGARGRRRRRRWRWRR*RWRRQWRW*RWRRRWCRRRRRRRW--------------*RRRRRRWLRRWR
Consensus                                                                                                                  *  *

Teleogryllus_occipitalis        ------------------------------------------------RRRQSQQW--QRRL--R-------------------------------------------------------
Acheta_domesticus               W-----------------------------------------------RRRRK-----------REGDGGGGGGGDGIGGGGGDGGSGSGGSGGGGSGSDGGGGGNGSGSSGGGGSGGGG
Gryllus_bimaculatus             V-----------------------------------------------KRSTAAKAA-------RE------------------------------------------------------
Gryllus_longicercus             WWWRWRWRWRRRRRRRRRRRRRRRRRRRRRRRRRRQRRRQRRRQRRRQRQRQRQRWLKQRWLLPR-------------------------------------------------------
Consensus                                                                       *

Teleogryllus_occipitalis        -------------------------------------------------------------------GGCGGSGGGGGGG----------------------------------------
Acheta_domesticus               NGSGSSGGGGSGGGGGGGVDGGDGDGGDGGGGGGDGGGGGGGGGGGSGGGGGGGGGGGGGGGGGGGGGGGGSGGGGGGGGGDGGGGGSGGS--GGCKEAVAVRRSCSEW*KRRRKRLSG
Gryllus_bimaculatus             ------------------------------------------------------------GGGGGGGGDGSGRGGGGDGXXXXXXXXXREGIFLGGSKQH--------------------
Gryllus_longicercus             -------------------------------------------------------------------GDCGGSGGGGSG*R--------------------------CTEW*ERCWKESSA
Consensus                                                                          *   * **** *

Teleogryllus_occipitalis        ------------------------------------------------------------------------------------------------------------------------
Acheta_domesticus               SECTR*CSVERLETEIRG--SIPFGSFPRLE------EK*FLKNGK--------------------------*GTTLRYLGSEERFILCLCMRVWSALARLRERVPACMR----------
Gryllus_bimaculatus             -------------------NKRPFFNFFQLKVKK*NIKKMNLKNFQ--------------------------------------------------------------------------
Gryllus_longicercus             SDCTR*GCVQRYETEICGNRASDSGTF-------------FFQNGRNIFFGERRVR*RRS*GHAEVQIHSGHIGMTKN*L-----------VFVYACVVSVRGRAHASDRESKIWREGDR
Consensus

Teleogryllus_occipitalis        ------------------------------------------------------------------------------------------------------------------------
Acheta_domesticus               --------AYVRARKR*RKRKEK-------EI*KLNERATIGD*LGNKHKLSVRADVCPRDKERTHRFKDSEQTERGGQGTEGDSRE-----NCWKRINLNERRLPE------TFSGP--
Gryllus_bimaculatus             ---------YRKGQYKI**RRQK--------------------------------------KKR*CFFRLLKQRPP**YIFPKKK**ERTK-NKYKSTNYK*NWAP*F*LSCKDFSGVSK
Gryllus_longicercus             DE*RGRIETYREGG*RRIEREDKDV*RGKGEI*KLRES-------DNWH------------*ERTCV*GERRDTQVQRQ*TDRKRRSRYTHINKWI-----QRRIGEK--ST*TRSGFWK
Consensus

Teleogryllus_occipitalis        -------------------CG*--------------------------------------------------------------------------------------------------
Acheta_domesticus               -------------QMLS--PGE-HPSPNAGYRLHINERRREMSNEENPF*IFERKLMCSME**R---------------------SFA-FQVNLTKVYKCK*VNNCIIRNYEN-HTAHTS
Gryllus_bimaculatus             ----*K*TIQ*M*NFFS--C*D-----------------------------------CT--------------------------QFAKFFISPIWFFLISTQNP--SRRFSKLHPAKTR
Gryllus_longicercus             CYRDGSTSAAISWEMFATECGLSSPCK*TSYR---DEHTKKIHSR-----FSERKLMCSM*Q*RFSVFSCRETVKEH*MHFLSIISY*-FYERINWKLACRS-----ITNYEK-HTAHTC
Consensus

Teleogryllus_occipitalis        ------------------------------------------------------------------------------------------------------------------------
Acheta_domesticus               PMT*HGELFPYTAQYY---F*FC*SDGTLMNF*CFKYFLPIK-KRQWKYKSNEKFRR----------------------------------EDNL*NTEVLKRV---------LA*CHEV
Gryllus_bimaculatus             SQC*SLHFFNVKVNLYILLYAFTTIIKLLQTFQTIFFVLAAK---VLRYLSTNQEKRIWIFFTIDQLETLGIIIARSYRPIFSF*NRWRTPAK*LSYLETIKRIGYWQT*NCTLKC*SRV
Gryllus_longicercus             SIL*HGELFPHTAQYY---F*FR*TNSTLLNF*SFKYFLVVRINASGKYKWNENVRR----------------------------------EDNL*NTEGLKRV---------LA*RHEV
Consensus

Teleogryllus_occipitalis        ------------------------------------------------------------------------------------------------------------------------
Acheta_domesticus               *LH*RQ--FRVRRGVVPVL-------YSMHNS*RFTRYCHV*LARLTDVRGYSQTNVGILPKRTV*TALSSSS*IPKSYI*FHLL*YFHWHHHRCRVIRLVNRRQFVKSILTRIVLEFQE
Gryllus_bimaculatus             KVNKNRPIFSIQTSVGSAVIILSGIKYISHH*IKF*QHPTVFTSNKTPSWKSLAPKIH*LTISKGTHSNSTYK*IFVPY*FF*TLI-IHYLVR*LKFLPLSNGVK-ASL*LLRGIHDAES
Gryllus_longicercus             *LH**Q--FRVRKGVGPVL-------YSAHNPFRFTRCCHV*LARLTDVRGHSQTNAGILPTRIE*TAPSP*S*ISKPYI*FLFL*YFHWRRHRCRPI*LGYRRQVHEMNFYKNRREYQE
Consensus

Teleogryllus_occipitalis        ------------------------------------------------------------------------------------------------------------------------
Acheta_domesticus               VGGTQNNA---------IIIKRH*VSEDVS*S-------DILNLDI*SCISA-------------------------------------SLISRKN---------I*FFHFI*FVI**RF
Gryllus_bimaculatus             TSISLNTAWKCAFPNFL*KIGWHVSNTYVSIYAFMHECINLCVCELCVCVRARARVCVCVLCGVCVCVCGGGVCVCV*VFCYIITFNH*SIFRKQKEKKKKRVSRIQYVTQILDTIKRHH
Gryllus_longicercus             VRGTK---------------*RHY-------------------HNMASSIRS-------------------------------------PFLTK-------------YFKF--GCVK*KY
Consensus

Teleogryllus_occipitalis        ------------------------------------------------------------------------------------------------------------------------
Acheta_domesticus               RY*RKNVHKMTCIQNC*SLIGNKLLMKNIS**NSF*K*KANKWK*MTKFP-------C*LPSSLLTRNFHGLPK**FIGMHL-------PIQCYKT---LQH*RKGGTAF-SLKCLFIL*
Gryllus_bimaculatus             NYALRNASKVM-------LPGERAAVAGRSPAQACEARRGEQGQEGRAI*RRAGVGRAIGKAEVHNR*W**LNTRATPGNHAKYLLIFVPVTNFHNVHVLENTNKKRMLFIQLAMRFF--
Gryllus_longicercus             NYFK**RNYIF------------LLFNSIC*F*TFSPRTCAHEDQSTEIV-------SLIKNFRVIKKYF*SPL*K*SADHRVE--IINQISIFIT---LKK-------F-SITL-----
Consensus

Teleogryllus_occipitalis        ------
Acheta_domesticus               YMAGLK
Gryllus_bimaculatus             ------
Gryllus_longicercus             ------
Consensus
```

**Tocc.2G0004080.1**

```
Teleogryllus_occipitalis   AACATGGATCTGCGCTATTTTTAAGCTTCCTTTGCTCGA---------------------------------------------------------------------------------
Laupala_kohalensis         ----------------------------TTTTTTTCTATACTA-----------------------------------------------------------------------------
Acheta_domesticus          AACACAAAATTGCAATTTAATAATTGATTCTTCACCAAGAATATTATAATCTCCAAATCATAGAAAAATGAACAAAATCATGAGAGCAATCTCTCACATCAAGATTGACATAACATAGAA
Gryllus_bimaculatus        AGCGATTAATTTAACTATAGCCATGTGACTATCACGCAC---------------------------------------------------------------------------------
Gryllus_longicercus        ------------------------------------------------------------------------------------------------------------------------
Consensus

Teleogryllus_occipitalis   -----------TCAAAAAAAAGTAATGTACAA----------------ATTTTGCGCCATT------TTGTAGAAAACTTTGAAAGTAGTAAGGCTTC----------------------
Laupala_kohalensis         ----------------------TATTGTGGAAT---------------ATTTTTTCTTGTT-----------------------------------------------------------
Acheta_domesticus          TATGTGGCAATTCAAAGAAGGGTAATGTTTAATAGCACATTTGGGACCATTTTTCCCCATTATGAAACTGAACAAAACATATCAACGAGCAAACTTCTGT--------------------
Gryllus_bimaculatus        -TTGCGGC----------------ACGTATAATCACGCAATTA---------------ACTAGTCAATTCTACTAAACGTGGTAATTAACAGTGCATCATTTAGTATGAGGTTTATTATC
Gryllus_longicercus        ------------------------------------------------------------TATGCTATTTGACGCAGTGAGAACAATACCAGAACTGTGTGTGGCGAAACTGACGCCGAT
Consensus                                                                              *

Teleogryllus_occipitalis   --------------------------TTGTAAAGTACCAAAAAATAAAAGTTAA-----------------------------------------------TTTGTCTTTCAAAATATAC
Laupala_kohalensis         --------------------------TTGGTAAATTATTAA-----------------------------------------------------------CTTTCCCATATATGAAATGC
Acheta_domesticus          -------GACGATATGTTACGGACAATCGTAAGATAATCAAGGGTATTGGTAAATAGAA---------------CGCAAAGAGCTTTATATTTTTATCTCTTTTTGATCTCGTCAAAGAC
Gryllus_bimaculatus        AACTCCATTCAGTGTGTGCCCACCATCTGGTAAGGAGTTATAACAACTCATTCCTTGCGAGAGCTCAGCCGGCGCCACGCAAACTG------------------------CATGGGGCAC
Gryllus_longicercus        ACGTGCAGTGGCCGCGTGTTAGCTAGTTATATCAAAACGATGAAAGATAATTAGAGGGAAGTGCGACGCCGAAGTACCAGGAGCCAAG--------------------CCCGAGGGATTC
Consensus                                                         *                                                                                *

Teleogryllus_occipitalis   GTTGCATCGTTAGCAGATA----------------------------------------------------------GAAACTGTTGGAAATAGAAAA----------------------
Laupala_kohalensis         GATGTTTTATTGTGTGATT--------------------------------------------------------------------AACTTAGTCAA----------------------
Acheta_domesticus          TCTACACTCTTGTCAGAAATTAGAACAGAAAATAAT---CATGATTTTT-TTAAGAACAATATGTTTTTGCTTTGAGAGAACCGTCTTATTCAATTAA----------------------
Gryllus_bimaculatus        TACGGGTCAACCGCAGATG--ACCCCAATGCGCAAGCGACACCAGCTGTGTTGGCGCGCCTGTGTTATCGCGCTGCGGAAACCAGCGTCTTCC-------CAAGCCACCGTTTATGACTG
Gryllus_longicercus        AAGAGGCCGTCAGCCGAGG--CGACCGCTACAC------CACCAGCCGC-------CGACTGCGCCAAGCTGCTGTGAAGGTCGGCGATACCTGCCGATGCGAGCCGTAATTCAAACAAC
Consensus                                 **

Teleogryllus_occipitalis   --ATGTCGTCAATAGAAC----------AACATTTAATCAAATTCATTGAGCCAA-------TAGTAAGAAAAATGTGTGCTTGTGACAA----------------ATTCTGTAGCTTAA
Laupala_kohalensis         --TCTGAGTGAATAAAAC--------TCAATTTTATTTGAATATC----------------TTAATAAAAGCGATTTGTGTTTTTGTTAT-------------ATTTATTTATATCTAAA
Acheta_domesticus          --ACATAGTTAATAAAA-----------AATAGTTTATTGCTGTCATCAAACAAAAACAATTCAGTATCAAAATTTTGTGTTTAGCACAACGCTAAATTTGTGATTACATGGTAGCAAAG
Gryllus_bimaculatus        TTATGTTTTTATTAAAGCGTTATATGTGAATTTTAGTGTAGTTTCTTTGTTTCAA------TGTGCG-TCATGATTCACCCTCCACGTAT----------------TTATGCCATTCATT
Gryllus_longicercus        GGAGGTATTTGTTGGGGC--------------------------------CCCAA------GGTATGATGAAAACTCATGTTAAACCTGA----------------TCTTGGTACCAAGG
Consensus                          *   *                                                                     *                               *

Teleogryllus_occipitalis   CAAAATTCAT---------------------------------------------------------------CTTCATATCT-------TCATATGGCAA---TCACAATTAGGCGTAA
Laupala_kohalensis         CAAAGTTTTT---------------------------------------TGTAATATTTTTCTCTGAGCATGTTGTTTTATGTTGCACTATTCCAGCCAAAGGGTTATATCTGGGATCAA
Acheta_domesticus          CGAATTACATTATAAGCCCTACAAGCAATTATAATAGTTGGTGTCGAAATCTAATATGTATAATTCGTTGTAGTTTTATATTTCGCCCTGTCATATACCGA---GAACAAACATTTGCAA
Gryllus_bimaculatus        CAGAGTTT----------------------------------------------------------------GTTTTAAATGTTTGGTAATGGTATATTA--------------------
Gryllus_longicercus        CAAAGTGTAT------------------------------------------------------------AGGTATTATAAATCATGTAAT-----------------------------
Consensus                  *  * *                                                                     *   *  *       *

Teleogryllus_occipitalis   CAGAAATGTAACAAC-----------------------------------------TTGCTTGTGTAA--------------------------------TTAATAAGTTA---------
Laupala_kohalensis         TATAATTCTAGTAGAGG---------------------------------------TAATTTATTTAT------------------------------TCTTTATTAATTATTTATTAAC
Acheta_domesticus          TATAATACAAAAGACAATATTACTTGAACTGTATATGACCGTCATGTTCTTGCCGATTGCTCTTTAAA--------------------------------TTTACGCGCTATCTTTCGAA
Gryllus_bimaculatus        --------------------------------------------------------TTACTGTTTTAAGCTCACCCGCACCAGTGTGTGTTTCTTTACACTTCACAAATTAT--------
Gryllus_longicercus        --------------------------------------------------------TAACTGATTTGG--------------------------------TTTGTGAATAAA--------
Consensus                                                                          *   *  *                                   **        *

Teleogryllus_occipitalis   -------------------------------------------------------------------------ATTTGCATCATAGAT-----------GATCTTGAAAC----------
Laupala_kohalensis         CT-----------TTAGTGTTATTCATTCATATTTTA---CACAACTGTTTTTTTTATAATTCTTGAATTCCGATTGACATGTAAGAATAGTATAAAAGAATTGCAAATTATGTGAAATT
Acheta_domesticus          TTGCGCATTCTGATTGGTGGTCCCCACTCCTAGATTGGGTCCCATCCCCACCCTTCATCGATT-----TTCTCATTTGCACCCGAGTTGCCCGCACGCAGCTCCTGAGGTCAGTGAAAGT
Gryllus_bimaculatus        -------------------------------------------------------------------------ACCCGTACGAGGTAT--------------------------------
Gryllus_longicercus        -------------------------------------------------------------------------TCCTGTTTATGATTG--------------------------------
Consensus

Teleogryllus_occipitalis   -------------------------------------------------------GATTTTACTGGTGGGTCAT------ATATGTGCTGATTCCAACTCTTTTTGAAAACCTATTTGAG
Laupala_kohalensis         ACGATAGACAATGCACATCACTTCATGCCAATGAACTTGGAGTTCTTTAGGACAGAAATAAATTATTGGAATATTTATATGTAATTCCTGATTCAAA---TTTTAAAAAATCCATACGAT
Acheta_domesticus          GCAAAATGCTGTGCAGGTCTGACTAT----------------------------ATATCATACTGCCTAATTAT------ATATTAATTGAAGTCAATCCTTCTGAATTACCTAAATGA-
Gryllus_bimaculatus        -------------------------------------------------------CACAATATCTCTTAATTAATTATACGTATTAATTCATGTCAAGTACTCTGAATTCTCCAAATGA-
Gryllus_longicercus        -------------------------------------------------------TACCACATTTTCTAATTAA-------------------------------ATTTCCCAATGCTA-
Consensus                                                                          *    *          *                                      * *    *

Teleogryllus_occipitalis   GTCTAGTGAAAACCCCTGA--------------------CATTTTCTTAAAC----------CAGAATCTTCTGTTTATATACAATTCTACTTCGAGTTAAATTTTCCACTAATACAAAA
Laupala_kohalensis         AACTTGCGATGATATCCAAACACTATGGTAAAAGAACGTTTTTTTTACAGT--------AAGAAGTAAAATCTTTTTACGGAGAAGCTTAATTTGTTTTATATTCCCT------------
Acheta_domesticus          ---CAGCAACTTCCCTCGAAT--------------TATGTATTTTATTTCATACTT-----GCAGTATCTTCACTACAAAAATCATACTGGCTCTTCTAGAAGCTATTGTTGGTAAGTGG
Gryllus_bimaculatus        ---TAACAACCTCCTTGGAAT--------------CACATTTTTATTCCAGAAATTTGAAAGCGTATGTATTCGCTTGCAATAAATACACTATTCCTATGAAGCTATTAAACGTAGGTGA
Gryllus_longicercus        ---TGTCGACTCCT------------------------------------------------CATAGGGACCCATTCACGAAAAAGATACCCTCCTTTAGTAATT---------------
Consensus                          *                                                                           *       *        *

Teleogryllus_occipitalis   T---------------------GCTTCCTTTTTATTTCGATTATTATT------AGACATC-----------------------------------------------------------
Laupala_kohalensis         -------------------AAGACGGCGTTGTGAGATAACTTG-----------------------------------------------------------------------------
Acheta_domesticus          TCTAGTTAGTTATCATAATAGGGCATCCCCCTGAATCAGTTTATTGCTTACGAATGCCATCTCTGC------------------------------------------------------
Gryllus_bimaculatus        TGGATTTTGTACTAATTATAGGGCATCCGCCTGAGCCGGTTCACTGCTAACTACACCAACTCTATCATATGTTTTAAAAATTACTCAGTTCGTCATCTTATCCAC-----------CGTG
Gryllus_longicercus        ----TCTTGCAGTG-----AAAGTACCGGCATCACCAAGTACGTCACAATATATATATATATATATATATA---TATATATGTGTGTGTGTGTCTGTGTGTGTGTGTGTGTGTGTGTGTG
Consensus                                            *    * *
```

[illegible]

```
Teleogryllus_occipitalis   ------------------------------------------------------------------------------------------------------------------------
Laupala_kohalensis         -------------------------------------------GTAAGCTGAGCTCAGAACAGAGAAACAGAATATCTAAAA----------------TATAAGGGTAACGCCAGATAAA
Acheta_domesticus          CAGGCCCACGGCGGCGCCCGCTTCTTATGGCTGCACCTTGGCAGTAAACCATGCTCAACTTGGCGCGGTGCTATATCGAATATCATACGCCGCGGGGTCGTGAGGATGCATCGCTCCAGA
Gryllus_bimaculatus        GATATCATATCCTAAACTTACTTTT------------------------CCAATCCAACTAATAGTGTACGTATTCATACTGATATTGTATACGTGA-TACGAATATGAAACATAACACA
Gryllus_longicercus        CAGGCCCACGGCGGCGCCCGCTTCTTATGGCTGCACCTCGGCAGTAAACCATGCTCAACTTGGGGCGGCGCTGTATGGAATATCATACGGCGCGGGG-CGTGAGGATGCATCGCACCAGA
Consensus

Teleogryllus_occipitalis   ------------------------------------------------------------------------------------------------------------------------
Laupala_kohalensis         A--------GTTTCCAAAGCCAT-------AATTTAATTGACAAATATTC-----------------GTCCCTATG----TCTAAAATACTTTTAGGAAAAGAAGGTCT-----------
Acheta_domesticus          A--------GCGCCCGAAACCATTTGC---AGCACATGCGGTCTACGCGAAGGTAACCTCGCAATAAGCCCCTGTGGCCTCTTAATAGAGCTGCACGAAAGGGAAATTTAGCACGAAGGA
Gryllus_bimaculatus        ATGTTTAATATACTTATATTCATCAACAAAAACATATTTACTGAATGTTTAAATTTAATCAACAT---CCCATTTGGTTTTTTCGTAAAATTTTTATAAA---ACACATTGTACGTCGCA
Gryllus_longicercus        A--------GCGCCCAAAACCATTCGC---AGCTCATGCAGTCTACGTGAAGGTAGACTCGCAATAAGCCCCCGTGATCTCTTAATAGAGTTGCAAGAAAGGGACATTTAGCATGAAGGG
Consensus

Teleogryllus_occipitalis   ------------------------------------------------------------------------------------------------------------------------
Laupala_kohalensis         -------------------------------------------------TTCCAAAT----------------TTCGTTCCTAAAACTGGCAAAATAGATCGTTT----TAATCATTTAA
Acheta_domesticus          TAAGAC------------------------------------------------------------------------------------------------------------------
Gryllus_bimaculatus        TTGTATTACTTCGAGAACAATGCACA---------------------------------AAAGGCAGAACTA------TCCAATAGTTATATTAG---------------ATAGATTTAT
Gryllus_longicercus        AAAGACTAAATAAAGAATAAAGGGTATGCAGGTACGAGAGTGTTTATGGTTCTAAATATAAAAACGTAACTGCTTTACTTCAAAACTTACACAAATGTATGGTTCGTGGTAATCACTTAT
Consensus

Teleogryllus_occipitalis   ------------------------------------------------------------------------------------------------------------------------
Laupala_kohalensis         TCATTTCAC---------------------TGTTTACAGTCAAAACTTCGAACCACGAATTGTGATA---------AAACGTTATCAAATATTATTAGA--TACAAAGCAACATTCTTAA
Acheta_domesticus          ------------------------------------------------------------------------------------------------------------------------
Gryllus_bimaculatus        TAAAGTTGGTTTTGAGCTTGGTGGCAAACATGATTACAATTAGAATTAGAACAATTACAAAACTAT-------TACAAACATTTAAAATTAGAATTAGAGGTTCGATATCCTATATCTAA
Gryllus_longicercus        TCAATGTCTTAAAATGTTTAGATACAAGTGTGCTAACGTGTTGAATGTAGTTTAAAGCAAAATTATACCTTGCTATGAACTTTTAGATTCATTTTCA----TTATACATCATAATTTTAC
Consensus

Teleogryllus_occipitalis   ------------------------------------------------------------------------------------
Laupala_kohalensis         CTGTTTATAAGGAA--------------------------------------AAA-----------------------------
Acheta_domesticus          ------------------------------------------------------------------------------------
Gryllus_bimaculatus        TTAATTTCT--------------------------------------------AATTATCTCTATATTTACTTCTATTTTAT--
Gryllus_longicercus        TTTCTTCCTACGGATTGGAGACAAACGTAAGTAATGCTGTGCGAAAAACGGCAAATTTTAGGCAGAAAAACTCTTATTACAGGC
Consensus
```

**Tocc.2G0004080.1_exon1**

```
Teleogryllus_occipitalis   -----------------MAITIR-------------------------------------------------------------------------------------------
Laupala_kohalensis         FFLINS*ECCFVSNNI**RFITIRGSKF*L*TVK*LND*NDLFCQF*ERNLERPSFPKSILDIGTNICQLNYGFGNFYLALPLYFRYSVSLF*AQLTFRKKLFH----------VHSKIE
Acheta_domesticus          ---------------------------------------------------------------------------------I*F*HKIAI**LILHQEYYNLQ---------IIEK*TK
Gryllus_bimaculatus        ------------------------------------------------------------------CE---------RLI*L*PCDYHALAARIITQLTSQFY*TW*LTVHHLV*GLLS
Gryllus_longicercus        ----------------MLFDAVR-----------------------------TIPE--------LC-----------VAKLTPIRAVAAC*LVISKR*K------------IIRGKCD
Consensus

Teleogryllus_occipitalis   ------------------------------------------------------------------------------------------------------------------------
Laupala_kohalensis         T*NHLARSEGSR-------AENIEKREIF---------TNIAS*I---LSSLCSYSSSM-RGNIISAPEDTRHCESTA*TWLP---------KSSESPLYYKLESTI*LFSRRSFS*LSN
Acheta_domesticus          S*----------------EQSLTSRLT*HRICGNSKKGNV**HIWDHFSPL*N*TKHI------------NEQTSVTICYGQS*DNQGYWLNRTQRALYFYL----FLISSKTLHSCQK
Gryllus_bimaculatus        TPFSVCPPSGKEL*QLIPCESSAGATQTAWGTTGQPQ-----------MTPMRKRHQLCWRACVIALRKPASSQATVYDCY------------VFIKALYVNF----SVVSL--FQCAS*
Gryllus_longicercus        AEVPGAKPEGFK-------RPSAEATATP---------------------PAADCAKLL*RSAIPADASRNSNNGGI--CWGP---------KV**K-LMLNL----ILVP---------
Consensus

Teleogryllus_occipitalis   ------------------------------------------------------------------------------------------------------------------------
Laupala_kohalensis         LTDFENNKNKKK*LILRFFMKLKDLLKVDINQEELI*YF*TVF*HLFQRR*VRRPRLPCF*SHFI*SS-----------PHLKQ*REEHRDSP-RCI----------------GNTENPS
Acheta_domesticus          L---------EQKIIMIFLRTICFCFERTV---------------LFN*T-------*LIKNSLLLSSNKNNSVSKFCV*HNAKFV----------ITW--------------*QSELHY
Gryllus_bimaculatus        F---------TLHVFMPFIQSLF*MFGNGI---------------LL-----------LF*AH----------------PHQCVFLYTSQIIPVRGITISLN*LYVLIHVKYSEFSK**Q
Gryllus_longicercus        ----------RQSV*VL*IM*LTDLVCE*I---------------LF-----------MIVPHFLIKF--PNAMS---TPHRDPF---TKKIP-------------------SFSNFLQ
Consensus

Teleogryllus_occipitalis   ------------------------------------------------------------------------------------------------------------------------
Laupala_kohalensis         VALRH------------*TIKVLSIMSLRCAVDS---------QTRSQSWCL*LSPGT*H*NM*QHNTQYYEELHLN*WLGGRNYVSRRRILKHSN*LFNLFCQF*ERNLVRA-----SF
Acheta_domesticus          KPYKQL**LVSKSNMY----NSL*FYISPC----------------------HIPRTNICNIIQ-KTILLEL----------------------------YMTVMFLPIAL*-----IY
Gryllus_bimaculatus        PPWNHIFIPEI*KRMYSLAINTLFL*SY*T*VMDFVLII-------------GHPPEPVHC*LHQLYHMF*KL------LSSSSYPPCVSKLI*RTMTFRILTPYL*LPITRSHLCSYLF
Gryllus_longicercus        *KYRH---HQVRHNIY------IYIYIYICVCVSVCVCVCVCV*QRSQN--AQQ*LDSSYCTHYQYD*ILLRLT----*IKYKSFPINS*******************PTHRG-----AY
Consensus

Teleogryllus_occipitalis   ------------------------------------------------------------------------------------------------------------------------
Laupala_kohalensis         PKSI---LDIGRI------------PES-----PLILLE*GIA*VTSVYLL*GV------------------T*MVLYSTDGTYNE-----KTLKCI--PEINKCLSA*HLETCGKGGCQ
Acheta_domesticus          ALSFELRILIGGPHS---*IGSHPHPSSIFSFAPE-LPARSS*GQ*KCKML-----CRSDYISYCLIIY*LKSILLNYLNDSNFPRIMYFISYLQYL------------HYKNHTGSSRS
Gryllus_bimaculatus        IIGVIIKVIIIKI-----IV----------------KFSRSLEEKSSCRLL*SI--------------------VLPGFSDQHQAN-EYLSSTLTNR--PQ-------------TARDLS
Gryllus_longicercus        PINIHKKINIQPINRDRWLVTN*CHSSS-----PNWKVLRSSSEND--RIIHTILSARSSSVAV*RI------NLVAGCSDR*YS----LASPISTRRIPE-------QHLNKCTNRARS
Consensus

Teleogryllus_occipitalis   ------------------------------------------------------------------------------------------------------------------------
Laupala_kohalensis         FF----IL*TVCSLPLSP-------YFVC---LYMILN-CYLFI*AIL*ILL--LP---------------------------VSIF*HVVVLYSVFCQYYLFVINK*ICLFKLCY*DF*
Acheta_domesticus          YCW*VV*LVIIIGHP-PE-------SVYC---LRMPS-----------------LP---------------------RKY*KMALF----VYQ**FCEDYSILDSRN---EETINTNV*
Gryllus_bimaculatus        YL--------LLALP-SPLH*HALHYYTC----------------RETGRELRTCPGWTGAPLVADTWLGKPPLA-GSRRASDLA------------------IERR-----KTTATIRA
Gryllus_longicercus        F-----------LP-PPCPTLTL-TLTCTTLLHMPGDRSEA*ILSRLGR----CP-VGGRYLVGETPASRQ*VGVGSRQAAEE--------------------DHR-----HLPNKNMH
Consensus

Teleogryllus_occipitalis   ------------------------------------------------------------------------------------------------------------------------
Laupala_kohalensis         *T--TMFISKEYI----------------SINDYDLPLHLIDESLKIDETLFL*YD*KFRQFCICIL*KHYSRTI*VRCISFLLLINEKFELHEYSTYVLVTFWSD------CTRIKCIL
Acheta_domesticus          QTYPSKFIN----VRYQALK*KSHFSI*HSIDS*YVIAN*IY*N-RTFLSVF---------VTFTIFENSESS-TK*TCILF*ASFARFFKKRTA*KNEQF--WSTF*KR*GCT---CKV
Gryllus_bimaculatus        NTYTTLTVLMT*VVDGPGITNKQ------SIQRIYLPPNSI-------LT*FR-----YRV*SHSTLPEAQTIQQQ*KISIFSIIICGKYIFRNGP*IEAINLWKI-------T---LSL
Gryllus_longicercus        YTYSTNDLSS*RIRNH*QTVNPTHNPTHLPIHQILF*PNAD--------TKF----------THSTYPETQTI-QQ*KISIFS-----RYS*RYFPK------W--------------TL
Consensus

Teleogryllus_occipitalis   ------------------------------------------------------------------------------------------------------------------------
Laupala_kohalensis         VLL*RISNQYI----------------------------------TRIYLLSTFRILSS*NF--------------------------------SNFREFIMHFCKHLP--HHETRT*CLPV
Acheta_domesticus          LKK*NIRFRL---PGYRV*LRKNMKKYQFWRKTYE*LRL*HLQ*IEELIYY-IDILQSTTF*YLGA-------------PKRGLWLHNTL*YTSYRNNI---CQLF---HNEVAT----I
Gryllus_bimaculatus        IYVNTIRRKTII*NGATSESRTSAVTSLTCRTMPARVSLRH--GLRPLLLRPRSVLRNNT----GA---------------------------------------------SSVSR----A
Gryllus_longicercus        IRSNKLMED*IIADPYKYHQKENDNLKKSYQVRPGASRCYH---IRSHLFN----LPDDA----GAGESSPRPPPSPPPPT----------------------QCFA**HGSVERF---A
Consensus

Teleogryllus_occipitalis   ------------------------------------------------------------------------------------------------------------------------
Laupala_kohalensis         FSRA*FT*GPLKWAPTRTLTISDSLSDPL-------------------SVVSGLRMSTSVNL*YVA----------------TLESAPNITHL--------------TYNIY--------
Acheta_domesticus          *TNSCYPA*LH*FIKNKNIA--------LRSG*FS*RCRLNIP*LVRCIVMSALSRDRLP------------KQ*S*CEV--NYMNEPVLLQNGTDFP---------ILVIY--------
Gryllus_bimaculatus        RARAQRP---HGWAEDKRRAHARTRAPEARGGGVEGEQGATPNSAPRRRTRRELRLGRAAFVWLAARGE---FLCSVC----SYFTSTNLLKA----------IK*IIRVVFREIQRLQS
Gryllus_longicercus        RARAQRP---HGWAEDKRRAHARTQAPGARGGGGRGRTRCNPKHRP--QAPHAARVSTWPGCFCLARGAR*IFMLGVFLL*EH*LTEGN*MNYKSGFS*NSTFAK**LQNVY--------
Consensus

Teleogryllus_occipitalis   ------------------------------------------------------------------------------------------------------------------------
Laupala_kohalensis         ------FHIS------------------------FSCL------*NKEKNINQ*YQIN**I-IKELNNNNNNNNNLIT*-RRHKTKSKRS-----------------------ELFTKY
Acheta_domesticus          ------FL**LFAPDPYSSELPCIAARST-----FYC-----------------HILNFSVF*L*-----NRNI-ILRILKNSKTSS*MAIFGKIWSRI*ENGV*ALL*I*NFIGLFHRK
Gryllus_bimaculatus        DN*KCLF*ILIFLNYTYFKKIVVQLARKY*KMELYFCLSIIDLRR**YSEVKK*YNTKFCRWIRTFDNHIDRN*YILRI-KH*NT*ICYM-----------AFYDATLDM*IFVLDYLVT
Gryllus_longicercus        ------FEILLFENYTYLK*L*YNSQENTEKWNYFFV-----------------YQ**FCE---------DDN--ILKA-RN-DTIRNFA-----------DGYESLTTISIAINTY*E*
Consensus

Teleogryllus_occipitalis   ------------------------------------------------------------------------------------------------------------------------
Laupala_kohalensis         KFVH*RS*VLFYL-------FN*II*L----TLIVVEY*IKFNI-------IYL*CH-DLKSCLVINTSGHYQ-------SFTFQLQKK*EQKVLYL----------QILN---------
Acheta_domesticus          VLSFRK-TSLEFI*STSTYKYSSIDKLLEICTLTVGEIFQKNYICVIFEKNILLSCTERTRIYFDVNNI*EPQSAVPASFST*VQFKRSSTQKPTSLFRDSTVRPV-SIFRASVASFAET
Gryllus_bimaculatus        RVTSKDLTVTHYL------------------*LLQSRIFRVFT---------------EINLRCF*ASFA*SEKKINAHISSHFEENFESSKVPCYVPRNINVNLF-------------
Gryllus_longicercus        STKIRKFATLYF---------------------MM*QLISEFSCWII----**LGLRQQTSL*LIICNFCNPE-----YSESSLK*ICALSKHPLLSLRKKLMHSFRAIFK---------
Consensus

Teleogryllus_occipitalis   ------------------------------------------------------------------------------------------------------------------------
Laupala_kohalensis         -------IVRFLKMQFE---*LMISKQDY*CQFIKNI*YSC--DEKLCNSIKTHRP-------------------N*KFK*CISI------KLGYFYFCDLINIISNMSDRVPSC-----
Acheta_domesticus          KFSRDGLLYKMVAVHVY---IALLAKSN--YW**RDDADRTRFNETH*TNRKKMYT-------------------------CNDL------*LHIIHLYTAITILYSRNVTT*-------
Gryllus_bimaculatus        -------*LRLYEIH-----IA*LTESN--CWE*RNDANRARFNQNH*ANGKKM*PFYMLCSLTEFNFISS*DNHNYA*QKCN--------NLH-LYLTNLVSLDHE*S*NDF--TGGLY
Gryllus_longicercus        ------KILKVLRLHVMCPEISTLICFD--CAFTKYTLND---*QKVTVGNRGMTP-------TEQDLIKTIEP---MVRKCDPLTNVA*QNST-SYLHKAITIMSNGYVTTYICT*ATW
Consensus

Teleogryllus_occipitalis   ------------------------------------------------------------------------------------------------------------------------
Laupala_kohalensis         INLQ*NCIEMTLFNVFLVTSSMS-----------RELLVL-------------------AYS*CGFS-------SFVNFLISCLS------------------YFLLYSSP---------
Acheta_domesticus          SI*MMDDLEKILLVVQIFAD*RTLLENL------SEL**---*HLTFS*TSIMFFFTN-PLLRSTFHL*ETASFFILNTIMNVI----VIKMLLVDDYFKFCLYFFLYSFP*KVFSTTLF
Gryllus_bimaculatus        I**FMNSF*KTYLR---SIRNF*HYPKP------SSLVF----------T*FHF*IPLSPVKKGSF*-------FVL*ISL*LLS*KTYLKIIAVNFV*LFCLYTFL*GG----FFYHLI
Gryllus_longicercus        VAWIMNDLETILLVGYISNDS*TLFEKPI*GL*ETADIFLNQAHLFLHNSNFEFHFPL--*KRVLFN-------FYYEYLYDCYHKKAI*RLLL*ILFNYFSCILFFEVG----FSTTLF
Consensus

Teleogryllus_occipitalis   ------------------------------------------------------------------------------------------------------------------------
Laupala_kohalensis         -------------SSCR--------HFVARVQS-----------------------------------------------------RARKSKFYELKIIRNRLQMH--------LH---
Acheta_domesticus          IE**MKTF-----------DFLIKYH*KV*L---KLH*QEEGR*-----RLARLLGHPKKLRRRLRHSRGAPIRNEAPLTAPRPSASRARRSNIY--------ANVFGAHSRSL*VHRTL
Gryllus_bimaculatus        YGVMKEIFPTPL*KQ----------H*KSIVKT-TITRKGEGNALADSSDSRRSFGAPTTRSRRAYQKRSVAPRPAPPHATPR----RAHAGVIY--------MQMY-------------
Gryllus_longicercus        ME**KKFLRHPYKRSTKKV**KLQ*HEKAKVTPWQIARTAEGAS-----------GAPTTRSRRAYQKRSVPPRPAPPHATPR----RAHAGVIY--------MQMY-------------
Consensus

Teleogryllus_occipitalis   ------------------------------------------------------------------------------------------------------------------------
Laupala_kohalensis         IY------------YSHIGAWSL-----------AEEEISKGKG-----------------------------------------------------WEL--SRGTGWRGVEGVK----
Acheta_domesticus          IYEQFKKPEPAPASAEPCRAWGLGEAR**KPRRYGQKRVGRRGGGSGGGGGGDGGGGGGGGGGGVAAEAAGGGGDGGGGGGAAAGRRRGGGGTVAVAAWRGWRRRGRRWRRGWGRRRRRR
Gryllus_bimaculatus        -------LEPIPDHYKFIEL*FTSSLKSQSRRPRARSRVERRGG----------------------------------------------------------RRGDERRGGMGRK----
Gryllus_longicercus        -------LEPIPDHYKFIEL*FTSSLKSQSRRPRARSRVERRGG----------------------------------------------------------RRGDERRGGMGRK----
Consensus

Teleogryllus_occipitalis   ------------------------------------------------------------------------------------------------------------------------
Laupala_kohalensis         -*LPVCACACYL---------------------------------------------------LPLCL--------------RAFIPSSEIPFCTRNEL*CEN*E----KLFCL*ILMTW
Acheta_domesticus          QGGDVEAAGRWLWRRGGGGGGGDGGVDGGGGGDGGGGGGGAAAGRRRGGSGAGVAAGTAAGRDAPSLISTAADRASYRVAHYRSGIREFKTRVL---------------RMF--------
Gryllus_bimaculatus        ---------EWV--------GVAVAVEGG--------WGWRRVGE------------------APSLISAATDCASYRVANYRSGIRESKTREYS*KCYPPENAK*QGSTL---------
Gryllus_longicercus        ---------*WV--------GVAVAVAGG---------GGVAAGR------------------GSSFI------------NFRS----HRPRVVSGRELPIGNSRIENARIF--------
Consensus

Teleogryllus_occipitalis   ------------------------------------------------------------------------------------------------------------------------
Laupala_kohalensis         DVSMCSN*KHEIFSWKL**FNGTLK*N-LEVKRSFNGKIVNFRWFFSIDQTLC---------------------IYNSIVQFEA---ILR*K-----LNIKYC*YFI-LINTTIG-----
Acheta_domesticus          -LEMLFS*K-----CKMTRINSAMMES---YRRSHN--------ILAKASQ---------------NQA*I*TKLY*----------VLRVQCWW*KLPIKRL*KSLSLI*LSQQ*YSYM
Gryllus_bimaculatus        -------PW-----WKVTRDSL*RHKTKFKYT*KY------------IQSQKCNVGGGNCQ*KCYKNHYGQFT*VYNVYLARR*KKCISKIN-----------------VHTILG-----
Gryllus_longicercus        -LKMLSS*K-----WKMSRINSAMVES---YKR*L------------IASQ---------------N*V*I*IKLYS----------ISKVQCWWWELPIKML*KPLRPVYLSLQ-----
Consensus

Teleogryllus_occipitalis   ------------------------------------------------------------------------------------------------------------------------
Laupala_kohalensis         ----------------TNKLF**E**QEYFVRLP*---------------GFS-------------------------------------------------MRKSNCCVLVFITFKINF
Acheta_domesticus          KKM*PTFRVDSISCAIDEEQLVNVFVKHNFVNYRAKSLSAWRAGFVIERAGFDPRVGHPGRGFRGFLWS*GEYWDGFKIPRTPPTFHYPT*HLPPSF*SFHTSSSPSPMMMIIIIIIII
Gryllus_bimaculatus        --------------AIDKE*LVNDFAKHYFVTYP*RE----EAKVMKIKNG----QGQ**QAAR-------------------------------------HYMLLPNTDVLYLCAIIIC-
Gryllus_longicercus        --------------CLPSS*MKNVSQKLVYTTFCA-------------------------------------------------------------------LLIRNS**MTLLNIIL*P
Consensus

Teleogryllus_occipitalis   ------------------------------------------------------------------------------------------------------------------------
Laupala_kohalensis         IRCN*KIKNRI*KLLMRKMSSNELAVRLEDL*FPAF*NALKHTNFMNGNS*---------NNSCY----------------------------------KSTNIYANE*DN----TLHNIT
Acheta_domesticus          I------------IITKITSSYHLI-----------------IDYSRQQSC-LNQGM-RP*G*CP----------*FHTFISPE*STTGPRKRETCCVVSSGPYANNNNNF---VNYP*P
Gryllus_bimaculatus        -----------------RWKQLHLTCM*N----------IASIRYTNGSSQDLF*GN*RIQQVCWGPWK*SLLRRV*SVLIV*EQRINMSVKR---------PLMRNMQSLRKQLRYK*H
Gryllus_longicercus        I------------HDAKKPKR*RLK-------------------MGGDSDDKQRGT-----ICY----------------FQVRTYYTSAR---------LL*ADGNN------YTLH
Consensus

Teleogryllus_occipitalis   ------------------------------------------------------------------------------------------------------------------------
Laupala_kohalensis         ADASVTNEINDSMWS*NLLSE*PSS*FGCQLFQLNLIDIVF*ITI-------YSVIWGSL-LCISSW*FYRFR*FIFLSPGI*SSEGPEIITNLMT*YIVVIINIIIIIIGILGLSDI*V
Acheta_domesticus          EEANEMK-VNFKM-------GRGSDGKQNRSWMLCVTPTVWI----------Y*TSVGGV-QGES------------T*NFVTSQG*S----------IVHICLCKA-KCSSTKQQSDK
Gryllus_bimaculatus        ALFTALKAIRISA-------GRAKTAKAGIRKYFSLLRILYA----------FKSDVRAA--------------------KN*LALA-------------V*LQLCRA------------
Gryllus_longicercus        VCKT*L----------------P*DTQMARRKTFSKEAKVYSTCAGDNGSEPYCVECKAC*LCEN-------------KEERCLSKG-------------L**EICRA*RSN*GTNNTH*
Consensus

Teleogryllus_occipitalis   ------------------------------------------------------------------------------------------------------------------------
Laupala_kohalensis         IRTLL----RNI*LF*IIHFYDIFLIKHLTILLATIRQQVCTFVTYFPHDFNALTRTH*LII*TC*YLVVII**HNV*N*IIE---NKAINHQ*TETKYIS*S--------H--------
Acheta_domesticus          *RVT*LGASRK-------------*IQYRNSL*VSIK---------IPPS-------------------------KSQYNLIQ---QHCITY----LQIPSMHE*INSTQHHLSSTPQQP
Gryllus_bimaculatus        -----------------------------E*MMLIMQR-----Y*YITPTY------------------VFLKLHNIPLWLVRCAGQAAITSHLLDLRILCVS---NST-----------
Gryllus_longicercus        LRL----------------------LKPFESVLVEIKH-----*TYI*PTF------------------------NVRNSFIH--------------KII*LS---NS-----------
Consensus

Teleogryllus_occipitalis   ------------------------------------------------------------------------------------------------------------------------
Laupala_kohalensis         -------------------------W*INPDNV*HDQIPHYDFMSVFFF-------------*HGAFIINTFPVQYPSVFFIHR---------------MGFF*AVICFACGFFC-IEVS
Acheta_domesticus          SKNPYNRPSGQ*RKT*R-IQEIYPTYLSNPTNRVRDTEIHNTMSNILICINKKIFTGSLN*I*HDNFLILKSL*NTKSRVALLRVHRTNGRIVL*LQH-----LQCKVSTTPAM*--GRS
Gryllus_bimaculatus        ----------CF*DLLRDISDSIQAWIQ-KSNHVR------GFSNVK---------------DVRHFSILSSRRNSKEE*IIYS-FNNP--------------FQLSLTTNPFFC-----
Gryllus_longicercus        ------------------VSDNLAIIVKGHANKIK------CFRKI*YC-----------CIDPTLFV**VLAVS*QKEIV*LS-HEYPGRKPC*L*NSLSVLFQTRLIYSAIMCSVGLH
Consensus

Teleogryllus_occipitalis   -----------------------------------------------------------------------------RNRN---------------------------------------
Laupala_kohalensis         SMSVN------------------SSFQFF-FSCLFYVTSI--AIFT*YL----------VKHIFYILVLTFFSFITFRNKSTYHHHHHHHS-IN--FSKIYFRKILFYEYTCLPIY----
Acheta_domesticus          KTP-----------------------------------KI-------------------LLHVVVYLCYMGKKSATFSNSGLIQ***TGNTVLN-IFLKI*LECFIFTKIETRVV*SGIV
Gryllus_bimaculatus        -------------------------------------EIKKIEYTIHLREEKS-------------------WL*FHSR*NPRTTAKGVC-LNCLFLQCYPSL**F-------------
Gryllus_longicercus        KTDVKLIGL*LLTILLL*IGTTIVSFQF*EIIVLFIVTSINRAKYVILVGTDFAKNSCTLLHVVAY*CS*IIKWAKFANSGSIQ*IFTGNT-LN-MFLKM**KCIIFNKM----VL**TT
Consensus

Teleogryllus_occipitalis   ------------------------------------------------------------------------------------------------------------------------
Laupala_kohalensis         ----IHSNVTLHRENVTKEKKT*MRNSDYQ-----FI*KKYQQISVLSLKVMLIPRVCCWYRPGILKGHTQK-------NLGPA---LLAV*IC*VCNATFPG--RGKSC----------
Acheta_domesticus          RNPVIHPNIKI*R--ITYQ---LPKSLNFCCKEI-HL*NYY*TL*SYMLTSLCV------*KHSILKS*KNRYLCFTNYNLTHTDVCLLHLGI-------FEGLDRMKRLIEQMTVDKRS
Gryllus_bimaculatus        ----------------P*QKNDFSD*AD*DLQNR---*KTYLS---------------------------AKHVCFQTCQ-RPT----------------------KKC----------
Gryllus_longicercus        *NNVTHLNLKI*R--IPYLLGEL*ESLDFCCKNYIHILKYY*QLCL*TLTILYI------*KRSVFKT*KTRYLHFANCNLTPSDVRSQKLDI-------LKGLDTMKQCIEQMTADKRS
Consensus

Teleogryllus_occipitalis   ------------------------------------------------------------------------------------------------------------------------
Laupala_kohalensis         -SPWFD-------------QS*ANLKLLSYLTDH*KTSTGEYIPIKS*REKMF----YYGAIKETICNF-----------------LLYMSWTSITR-----------NILYVFNLN---
Acheta_domesticus          FYSFYDFIKI*C---------LSSKDLIIA------HDNSSQV-------------YYCKVKN*-----------KL**KSRNFKVMYLCFTE**K-----------NVLKL-------
Gryllus_bimaculatus        ---FYDFTLFQ*KLYLNTDSVYSIQRQIIQMLSKPKESLISLL--KTLNQNVFVHRPF*THSKHLRKKYNGHVHKMKSILPHEYYLQLKPYFSELLFLDQMPRIPNHFNLLKLVSPKELF
Gryllus_longicercus        FY*FYDFIKLQCSY---VKSVVCTRKLIWSLLFHLKQKSHNLV---*MKARIFL*RKFLCVLNNI-------------MI**RSFRVMT*CFTE--------RIKN--NVLILYHYH---
Consensus

Teleogryllus_occipitalis   ------------------------------------------------------------------------------------------------------------------------
Laupala_kohalensis         --KLCYSPYLNDFSSYLTTPS*GI*NKL---------------SFSVKR---FYFLL*KKRSFYHSVWISSQ-----------------------------------VIVWIF*NLNQEL
Acheta_domesticus          ---------L-EMPLHL--------HTL-SNTN-RSSVLLVSGTYSSKKL**FIFMKPGMRRIMMV*GNKLIRGLHYGADID---PPRRLL----------DDAVH-LRKWE--------
Gryllus_bimaculatus        SLLLCI-*YI-TVTCQITT-------TL------L*KLLPTTQQF-------WLHMQIRHQRIYKNT*ISCKRDYDWGVNFSKFQPITSMLFK*QCAKMKLNDAYRNT**WLNTSNST--
Gryllus_longicercus        --YFCI--YL-EILIQLTYCK-SLEHTLP*NFNDLFR*NPICGE*-------WWYKRIS*LEGFIMEQISIYQD-----------AYLMMLFTYANGKL--------TVTWQKV*GSA**
Consensus

Teleogryllus_occipitalis   ------------------------------------------------------------------------------------------------------------------------
Laupala_kohalensis         HINIPIIYFCPKELQVHWH---------------EVMCIVYRNFT*FAILLYYSYMSIGIQEL*KKQLCKI*MNNTKG**IINKE*INYLY*NYIDPRYNPLAG--------IVQHKT--
Acheta_domesticus          -INGYLAEGLRKRLMKS*RLYSDIS*TRETEAPTGKGALQYRRF------------CCG--AV*AKR*VSPS*RHT------------------LSTRYRPTAAPASYGCTLAVNHAQ--
Gryllus_bimaculatus        -----LTYF*CRNNILQQQ---------------NRK--QH*NFVILQG*AIAVHMCLGKAKLFSAKQQSDQKR---------------------------ASKFSYG-----HHASKF
Gryllus_longicercus        NLKGYVVIFPRREKLRRRR---------------EKGALQYRRF------------CSG--AV*ARR*VSPS*RHT------------------LSTRYRPTAAPASYGCTSAVNHAQ--
Consensus

Teleogryllus_occipitalis   ------------------------------------------------------------------------------------------------------------------------
Laupala_kohalensis         ----------------------------------------------------------------------------------TCSEK------------NITKNFV*I-----*INITK
Acheta_domesticus          ----------------LGAVLYRISYAAG-----------S*GCIAPEAPETI---------------------CSTCGLREGNLA---------------ISPCGLLI-----------
Gryllus_bimaculatus        NTEILGRFESKYLRQILNAPLYNDTVQHTYKSPP*TKK*YNAASPIP*AQTAIQTP*PPIFWTIVKNLNDSRDIIS*TYFSNPTNSVRIHTDIVYVIRI*NITQCLIYL------YSSTK
Gryllus_longicercus        ----------------LGAALYGISYGAGREDASHQKR------PKPFAAHAV----------YVKVDSQ*APVIS**SCKKGTFSMKGKTK--*RIKGMQVRECLWF*I*KRNCFTSKL
Consensus

Teleogryllus_occipitalis   -------------------------------------------------------------------------------------------------------------
Laupala_kohalensis         TQIAFIKIFK*N*VLFT----QID*VN--HTIKHRI----SYMGKLIIY----------------------QNKKKYST----------------I*YRKK---------
Acheta_domesticus          ----------------------------------------E-LHEREI*---------HEG*D----------------------------------------------
Gryllus_bimaculatus        TYLLNV*I*STSHLVFS*NFYKTHCTS--HCITSRT------MHKRQNYPIVILDRFIKVGFELGGKHDYN*N*NNYKTITNI*N*N*RFDILYLINF*L--SLYLLLFY
Gryllus_longicercus        TQMYG---SW*SLIQCLKMFRYKCANVLNVV*SKIIPCYELLDSFSLYIIILLSSY---GLE-----------------TNVSNA-----VRKTANFRQKNSYYR----
Consensus
```

**Tocc.2G0004080.1_exon2**

```
Teleogryllus_occipitalis        ------------------------------------------------------------------------------------------------------------------------
Laupala_kohalensis              -------------------------------------------------------------------------------------------------------------------FF--SILYC
Acheta_domesticus               ----------I*F*HKIAI**LILHQEYYNLQIIEK*TKS*EQSLTSRLT*HRICG-----NSKKGNV**HIWDHFSPL*N*----------------TKHINEQTS---------VTIC
Gryllus_bimaculatus             --------------CERLI*L*PCDYHALAARIITQLTSQF--Y*TW*LTVHHLV---*GLLSTPFSVCPPS--------------------------GKEL*QLIP-----------C
Gryllus_longicercus             ACNKSFSA*NLPFFAQHYLRLSPIRRKKVKL*CIMKMN----------LKVHSKV*FCFKLHSTR*HTCI*TF*DIE*VITTNHTFV*VLK*SSYVFIFRTINTLVPAYPLFFI*SFPSC
Consensus

Teleogryllus_occipitalis        ------------------------------------------------------------------------------------------------------------------------
Laupala_kohalensis              G------------------IFFLVLVNY*LSHI*NAM---------------FY------------------------------CVINLVNLSE*NSILFEYLNKSDLCFCYIYL---
Acheta_domesticus               YGQS*DNQGYWLNRTQRALYFYLFLISSKTLHSCQKL---------EQKIIMIF----------LRTICF------------CFERTVLFN*T*LIKNSLLLSSNKNNSVSKFCV*HNAK
Gryllus_bimaculatus             ESSAGATQTAWGTTGQPQ------MTPMRKRHQL------------------------------------------------CWRACVI---------ALRKPASSQATVYDCYVFIKAL
Gryllus_longicercus             *MSLSCNSIK-RSRGLIA------SLPSRRLHELRMVLGASGAMHPHAPRRMIFHTAPPQVEHGLLPRCSHKKRAPPWACI*CLK-CVVN--LGKLTASLRRPRCKTSCTAE------PP
Consensus

Teleogryllus_occipitalis        ------------------------------------------------------------------------------------------------------------------------
Laupala_kohalensis              -----YLNKVFCNIFL*ACCFM-----------------------------------------------------LHYSSQRV-ISGINI-------ILVEVIYLFFINY----------
Acheta_domesticus               FVITW*QSELHYKPYKQL**LV-------SKSNMYNSL*FYIS------------PCHIPRTNICNIIQKTILLELYMTVMFLPIAL*IYALSFELRILIGGPHS*IGSHPHPSSIFSFA
Gryllus_bimaculatus             YVNFSVVSLFQCAS*FTLHVFM----------PFIQSLF*MFGNGILLLF*A------HPHQCV-----------FLYTSQIIPVRGITISLN-*LYVLIHVKYSEFSK**QPPWNHIFI
Gryllus_longicercus             FPDGA*VSRV*EISLRSL*DFIKRFLKPSAR*PLISHLRK*TASSSKRLGRSISAP**SPLISL-----------FSYTTIILRISG-----------FIEINH*SFMEE--------YV
Consensus

Teleogryllus_occipitalis        ------------------------------------------------------------------------------------------------------------------------
Laupala_kohalensis              ---------------------LLTFSVIHSYFTQLFFL*FLNSD*HVRIV*KNCKLCEITIDNAHHFMPMNLEFFR------TEINYWNIYM*FLI-----------------------
Acheta_domesticus               PELPARSS*GQ*KCKMLCRSDYISYCLIIY-*LKSILLNYLNDSNFPRIM---------------YFISY-LQYLHYKNHTGSSRSYCW*VV*LVIIIGHPPESVYCLRMPSLPRKY*KM
Gryllus_bimaculatus             PEI*KR-----------------MYSLAI----NTLFL----------------------------------------------*SY*T*VMDFVLIIGHPPEPVHC----*LHQLYH--
Gryllus_longicercus             PE---------------------TYNTLI----ELVFLNICRSNGNGTISE-------------HYFLFV-------------Q*NTTSLL*NFVTI--------------SLYYLKHKG
Consensus

Teleogryllus_occipitalis        ------------------------------------------------------------------------------------------------------------------------
Laupala_kohalensis              -------------QILKNPYDNL------R*YPN--------TMVKRTFFLQ*EV---KSFYGEA*FVLYSLRRRCEIT*EVI*IWRITKFI*VKNIKNISCYGSPRHV**KITNCFFNS
Acheta_domesticus               ALFVYQ**FCEDYSILDSRNEETINTNV*QTYPSKFINVRYQALK*KSHFS-------I*HSIDS*YVIAN-----*IY*NRTFLSVFVTFTIFENSESST-----------------K*
Gryllus_bimaculatus             -MF*---------KLLSS-----------SSYPPCVSKLI*RTMTFRILTPYL*LPITRSHLCSYLFI----------------IGVIIKVI---------------------------
Gryllus_longicercus             IFFI---------KIFGL-----------SSRPNCVIFALNETVDSR*VFE-----------CKQRFLRMN-----TEV**NHKINRMIAYLP--------------------------
Consensus

Teleogryllus_occipitalis        ------------------------------------------------------------------------------------------------------------------------
Laupala_kohalensis              AVIKHFLSLRFNRDIFTSTSLLM-------------VCQI*KQFQVCLRLV-------------EPR*ATFT------------------------------PSRECGVANLTNSN----
Acheta_domesticus               TCILF*AS--FARFFKKRTA*KNEQFWSTF*KR*GCTCKVLKK*NIRFRLPGYRV*LRKNMKKYQFWRKTYE*LRL*HLQ*IEELIYYIDILQSTTF*YLGAPKRGLWLHNTL*YTSYRN
Gryllus_bimaculatus             -IIKIIVK--FSRSLEEKS-----------------SCRLL*S----IVLPGFSDQHQAN----EYLSSTLT----NRPQTARDLSYLLLALPSP-----------LH*HALHYYT----
Gryllus_longicercus             SFVQYIAS--LCQDL*E----------------------YLISASEHLRV*GYS*RSEDN----EFFK---------------------------------------F*RLNVFK----
Consensus

Teleogryllus_occipitalis        ------------------------------------------------------------------------------------------------------------------------
Laupala_kohalensis              --C*ERGP*IFLCVSF*DSRTVPATYA------------------RY--*HYL*GQYTNLLVLLLNK-----LI----------------------------------------VTVPHS
Acheta_domesticus               NICQLFHNEVATI*TNSCYPA*LH*FI---------KNKNIAL--RSG*FS*RCRLNIP*LVRCIVM---------------------------------SALSRDRLPKQ*S*CEVNYM
Gryllus_bimaculatus             --CRETGRELRTCPGWTGAPLVADTWLGKPPLAGSRRASDLAIERRKTTATIRANTYTTLTVLMT*VVDGPGITNKQSIQRIYLPPNSILT*FRYRV*SHSTLPEAQTIQQQ*KISI-FS
Gryllus_longicercus             --CKE**VSINTI------------------------ASNI--------LKYVCNFYNKNPVILIIL-----LINKECVRFLN*DG*HYFM*F-------TTRPSC*I*Y----ISITFS
Consensus

Teleogryllus_occipitalis        ------------------------------------------------------------------------------------------------------------------------
Laupala_kohalensis              --CFFFFGYIFSV*SYITVNVDW*ASILVE*YFSEIDFGKIDRMMMMMMMIGALIPECNK*EKG*Y*N-IENVFYKILCK-------------------YCDASYIEKTRKKELKT*IHG
Acheta_domesticus               -------------NEPVLLQNGTDFPILVIYFL----------**LFA--------------PDPYSS-------ELPC--IAARSTFYCHILNFSVF*L*NRNIILRI----LKN----
Gryllus_bimaculatus             --IIICGKYIFRNGP*IEAINLWKITLSLIYVNT------IRRKTII*--------------NGATSESRTSAVTSLTCRTMPARVSLR-HGL---------RPLLLRPRS-VLRNN---
Gryllus_longicercus             ETCLMCFQ*IF----IVLIQNLQTLPILLFMSIS------MQRHVTMC--------------TNS*QN-----------------------------------PFLLRLRT--LRG----
Consensus

Teleogryllus_occipitalis        ------------------------------------------------------------------------------------------------------------------------
Laupala_kohalensis              HRGNLNAEEPTGKTNHCSEEAH-----------------SMDKEN-----ARILDRK--------------RIYYKRPVSEEKNTHKIIMGN---------------LIMSNIIGIYLPM
Acheta_domesticus               -------------SKTSS*MAIFGK--IW--------------------SRI*E---NGV*ALL*I*NFIGLFHRKVLSFRKTSLEFI*ST----------------------STYKYS
Gryllus_bimaculatus             ----------TGASSVSRARARAQRPHGWAE----------DKRRAHAR-TRAPEARGGGVEGEQGAT-------PNSAPRRRTRRELRLGRAAFVWLAARGEFLCSVCSYFT-STNLLK
Gryllus_longicercus             ---------------------------*WKLL*IIL*FLKIGKRQL*CRFTR---------EGLLTITG-L*V*HQSYASRRNT*LPSIL-----------DEFGIKLIDCFTISTVFVL
Consensus

Teleogryllus_occipitalis        --------------------------CL--------------------------------------------------------------------------------------------
Laupala_kohalensis              GL*YILGL-CLLMV-----------YCFVFYYLILYVMLLYYDY*ILTRLYN*LMCPCKCIKIMGKICDECTNLLTNCCE*NS*VFYQENVVK-VYNLEQSDIPQ*SSNDLDVGEP*NPN
Acheta_domesticus               SIDKLLEI-CTLTVGEI---FQKNYIC------------VIFEKNIL---------------------LSCTER--------TRIYFDV------------------------------
Gryllus_bimaculatus             AIK*IIRV-VFREIQRL---QSDN*KCL---F*ILIFLNYTYFKKI------VVQLARKY*KMELYFCLSIIDL--------RR**YSE--VKK*YNTKFCRWIR--------------
Gryllus_longicercus             GIRATVKLFLFVRIQQALTRRRE*GRC----------NNIRFF*SILSCSHGLLR*LPNCQKLN*IIK*SCV*K--------NFVH*M*--VK-------CRF-----------------
Consensus                                                  *

Teleogryllus_occipitalis        -----------------------------------------------------------------------------------------------------NI------------------
Laupala_kohalensis              NNNNNINNNNDILRH-----QICY--YFRAF*RLNTWRQENESSETVKSPGRYTQ*TPPNN*INRNLKDDVY*VQLK*LASEL*AWSFR**IL*PH*IINFIRYRGIGSNIMQCIILFIG
Acheta_domesticus               ---NNI*EP-------QSAVPASFS----------------------------------------------T*VQFKRSSTQKPTSLFRDSTVRPVS--IF-----------RA------
Gryllus_bimaculatus             TFDNHIDRN*YILRIKH*NT*ICYMAFYDATLDM*IFV----------------------------LDYLVTRVTSKDLTVTHYL*LLQSRIFRVFTEINL-----------RC------
Gryllus_longicercus             ---NVLSLPALIRMALRAVISACYL-YLNCFFKLCIFL----------------------------IKDLLTDIF---LLCSHTINKLYTRRSRV----HF-----------RCP-----
Consensus

Teleogryllus_occipitalis        ------------------------------------------------------------------------------------------------------------------------
Laupala_kohalensis              VNIS*FVATIILGIPVHKI-----SMF*SILKRWKLQVFQTYSKFV*GHFSH**FSYP----IFYLSVTAYEIYF----EGNK-----D*NAT-VTFPHAKASWKTYKILLLSFLSK-QF
Acheta_domesticus               -----SVASFA----------------ETKFSRDGL----------------------------LYKMVAVHVYIALLAKSNYW**RDDADRTRFNETH*TNRKKM--------------
Gryllus_bimaculatus             -----F*ASFA*--SEKKINAHISSHFEENFESSKVPCY-----------------VPRNINVNLF*LRLYEIHIA*LTESNCWE*RNDANRARFNQNH*ANGKKM*PFYMLCSLTEFNF
Gryllus_longicercus             ---QHTCCTL*L--P*KR-------SFDEPF-------------------------------VYLMEAMFY-IHV----RCNC-------------------------FHRLIIIAQ-RY
Consensus

Teleogryllus_occipitalis        ------------------------------------------------------------------------------------------------------------------------
Laupala_kohalensis              I----------------CSNCCIY*NKILTVFN-------------I*FSPKYC---------F----ELNYRIVNAQRLVNAKEPSEIYNFS---------IKTSFYL*ILL-------
Acheta_domesticus               ---------------YTCNDL*LHIIHLYTAIT-------------ILYSRNVTT*SI*MMD------DLEKILLVVQIFAD*RTLLE--NLSEL***HLTFS*TSIMFFFTNPLLRSTF
Gryllus_bimaculatus             ISS*DNHNY----A*QKCNNLHLYLTNLVSLDHE*S*NDFTGGLYI**FMNSF*KTYLRSIRNF*HYPKPSSLVFT*FHF*IPLSPVK---------------KGSF*FVL*I-------
Gryllus_longicercus             STSVLGSNI*CRAACHHCPRPFLTFISLASSRH--------G*VTK*CLAKSFTNYSLSIAR------KMLCTLISEIHFS-STS*VNIVNLGKLA------VKVSIAFLLAIP------
Consensus

Teleogryllus_occipitalis        ------------------------------------------------------------------------------------------------------------------------
Laupala_kohalensis              --*SSIKLL*FPGKYFMFLIRTH*YIP-SH*NL*TE*FL-------LIFTL------------------------------------------*LVACTE--RDFRGR-------------
Acheta_domesticus               HL*ETASFFIL--NTIMNVIVIKMLLVDDYFKFCLYFFL---------YSFP*KVFSTTLFIE**MKTFDFLIKYH*KV*LKLH*QEEGR*-RLA------RLLGHP-------------
Gryllus_bimaculatus             ----SL*LLS*--KTYLKIIAVNFV*L-----FCLYTFL*GGFFYHLIYGVMKEIFPTPL*KQ------------H*KSIVKTTITRKGEGNALADSSDSRRSFGAP-------------
Gryllus_longicercus             --TTNIALLRL--NIVLFIFRLSFVTL-----*AISCNFPPWQS*SLTFSIFRRITFSGIFSRFRF--------SNSRSVVRDPIRR-----AVGGCGN**RSFPGPPPPPLPPPQPPPR
Consensus

Teleogryllus_occipitalis        ------------------------------------------------------------------------------------------------------------------------
Laupala_kohalensis              YESTETKRKKITRARA----NRKSLHPLHPTPPRSTTQLPTFTLRDFLLR*APCANV-G------------VIYMQMHLKPIPDDFKFIEL*FTSATLHPSDEVSTRRRRRI-*KEIGQT
Acheta_domesticus               ----KKLRRRLRHSRGAPIRNEAPLTAPRPSASRARRS-------------NIYANVFGAHSRSL*VHRTLIYEQFKKPEPAPASAEPCRAWGLG-------EAR**KPRRYGQKRVGRR
Gryllus_bimaculatus             ----TTRSRRAYQKRS---------VAPRPAPPHATPR-------------RAHA---G------------VIYMQMYLEPIPDHYKFIEL*FTSS-----LKSQSRRPRA--RSRVERR
Gryllus_longicercus             LPTISFPYRRAFHHR-----------AYRPGARHGS----------------ALA---GAGSGFLNCS*IKVL*TYNDLEWAPNTFAYILL---------------RRAR-----DAEWR
Consensus

Teleogryllus_occipitalis        ------------------------------------------------------------------------------------------------------------------------
Laupala_kohalensis              GN------------------------------------------EKID------------EGRKTTSRIRKYQKLT*H*RSN*ENVKECHFYAISLQVNTRRDTV*HV*D----------
Acheta_domesticus               GGGSGGGGGGDGGGGGGGGGGVAAEAAGGGGDGGGGGGAAAGRRRGGGGTVAVAAWRGWRRRGRRWR-RGWGR--------RRRRRQGGDVEAAGRWLWRRGGGGGGGDGGVDGGGGGD
Gryllus_bimaculatus             GG------------------------------------------RRGD------------ERRGGMGR-KEW---------------VGVAVAVEGGWGWRRVG----------------
Gryllus_longicercus             GA------------------------------------------ERGG------------EGRFVSDR-R----------------------AASVSW---------------------
Consensus

Teleogryllus_occipitalis        ------------------------------------------------------------------------------------------------------------------------
Laupala_kohalensis              -----------------------------DVN----*ITEIKVAKFY*NALFKFLIRSV----CFY--GIT*F-FITTILNIF--------YELTL---------IILL*YH*LFKLHF-
Acheta_domesticus               GGGGGGGAAAGRRRGGSGAGVAAGTAAGRDAP------SLISTAA--DRASYRV---------AHYRSGIRE--FKTRVLRM----FLEMLFS*KC----------------KMTRINSA
Gryllus_bimaculatus             -----------------------------EAP------SLISAAT--DCASYRV---------ANYRSGIRE--------------SKTREYS*KCYPPENAK*QGSTLPWWKVTRDSL-
Gryllus_longicercus             ------------------------------APPKLLRLSELSAKA--LPSPFRVIVVFTILF*CSFYKGVGEISFITP*IRW*KNPPQRKEYK------KNN*TKFTAIIFK*LFYDN--
Consensus

Teleogryllus_occipitalis        ------------------------------------------------------------------------------------------------------------------------
Laupala_kohalensis              --*KSNNI*NLKVKNFLFLF----------------LL*LECE*-----------LVVSRCVDN*A----------------------RLQVMTSKIN-NVEFNLIFYNN----------
Acheta_domesticus               MMESYRRSHNILAK--------ASQNQA*I*TKLY*VLRVQCWW*KLPIKRL*KSLSLI*LSQQ*Y-SYM-----------KKM*PTFRVDSISCAID-EEQLVNVFVKHNFVNYRAKSL
Gryllus_bimaculatus             --*RHKTKFKYT*K--------------YIQSQKCNVGGGNCQ*---------------KCYKNHYGQFT*VYNVYLARR*KKCISKINVHTILGAID-KE*LVNDFAKHYFVTYP----
Gryllus_longicercus             ---NHKDIHNKN*KEPFFTGESGIQN*NYVKTSE--LGLGKCQ------------------------QFL--------------------------IDLK*VFQKEFMSH*IYNPPVKSF
Consensus

Teleogryllus_occipitalis        ------------------------------------------------------------------------------------------------------------------------
Laupala_kohalensis              -*RQLYNLIK*IK*DLR-----------------------PLMDE---------------FIFCE----ELTTL----RLRFMPSSGDQII----------------IIIIIII*FFNY
Acheta_domesticus               SAWRAGFVIERAGFDPRVGHPGRGFRGFLWS*GEYWDGFKIPRTPPTFHYPT*HLPP-------------------SF*SFHTSSSPSPMMM----------------IIIIIIII----
Gryllus_bimaculatus             -*REEAKVMKIKNGQGQ**QAARHY-------------MLLPNTDV------LYLCA----IIICRWKQLHLTCM*NIASIRYTNGSSQDLF*GN*RIQQVCWGPWK*SLLRRV*S----
Gryllus_longicercus             QDRS*SKLPKLLKYKCKLLHIRYS*-------------L*LPYEDMKLNSVKLHL*EGHIFLPLAQWF*LNLALL---ASFLY----SQQLL---------------------------
Consensus

Teleogryllus_occipitalis        --------------------------------------------------------------------------------R---------------------------------------
Laupala_kohalensis              LLINLILLIYIFLFVL*TRKRYMKINIIS*MCDIRGGFQ-----------------------------------------RRHIL*VNT------------CTHSQSRNNT*WIT*GITD
Acheta_domesticus               ----------IIIITKITSSYHLIIDYSRQQSCLNQGMRP*G*CP*FHTF-----------------ISPE*STTGPRK-RETCCVVSSGPYANNNNNFVN-YP*PEEANEMKVNFKMG-
Gryllus_bimaculatus             ----------VLIV*EQRINMSVKRPLMRNMQSLRKQLR-----YK*HAL---------FTALKAIRISAGRAKTAKAGIRKYFSL----------LRIL--YAFKSDVRAAKN*LALA-
Gryllus_longicercus             -----------------SVNHSMCIS*RRSQNKLTLIFR-----GT*HGVLELSKFS*KWLEMNALIFFSD*AKDA*KE-RRFISV--------NSLNILDCKSYK**VKVRFVDVTLVT
Consensus                                                                                                       *

Teleogryllus_occipitalis        ------------------------------------------------------------------------------------------------------------------------
Laupala_kohalensis              C*SAGWRPLKWTLCELRSRENR*TLCSSFMVRQMFA---------KMHDEFAKI*EISRRQYSKSAEEIYPSYVLV*DSLQQDQNTLNSRTI*PKRDQHIR*VFM*FEFFIY***K*DTA
Acheta_domesticus               RGSDGKQNRSWMLCVTP-------------------------------TVWIY*TSVGGVQG------------------EST*NFVTS--------QG*SIVHICL-----CKAKCSST
Gryllus_bimaculatus             --------V*LQLCRAE*MM---------LIMQRY*YITPTYVFLKLHNIPLWLVRCAG------------------------QAAITS--------HLLDLRILCV--------SNSTC
Gryllus_longicercus             R*S----NTKIHLSTVTS*N---------IM*QIYVF*CFILNMY*LRSIWLSNFRIHR------------------------QNFVLY--------HFLPLEYYHLRRTIIDKQKNSSI
Consensus

Teleogryllus_occipitalis        ------------------------------------------------------------------------------------------------------------------------
Laupala_kohalensis              Y-------SNCSTIMLL*NTNAKLSKFLVV--LKK*SFVNFK*LIN*M*W*IIIIN*N-------ILF*NKHCCSLKILITQFK*TNLFIDHK*IILTEYRIKNHNMSKN*HRK------
Acheta_domesticus               KQQ-----SDK*RVT*L-GASRK*IQYRN------SL*VSIK----------IPPSKSQ------YNLIQQHCIT-------YLQIPSM--HE*INSTQH----HLSSTPQQPSKNPYNR
Gryllus_bimaculatus             F*DLLRDISDSIQAWIQ-KSN-HVRGFSNVKDVRHFSILSSR--------------RNSKEE*IIYSFNNPFQLS-------LTTNPFFCEIKKIEYTIHL-REEKSWL*FHSR*NPRTT
Gryllus_longicercus             FQYFL---ASCITIILN-TYNFQIIKFQN----KHFVIITLQT---------LNFTKN-------HSYNSFNCLQ-------*VSAPKVRTHRA*KFTSR--REPNKSSPAKSKLSPRAA
Consensus

Teleogryllus_occipitalis        ------------------------------------------------------------------------------------------------------------------------
Laupala_kohalensis              ------------------------QQNSKDSS-------------------------NEQITI---------QYHI*TNKVWRQR*GAYSSQNKELTS-SFPAR-----FQMLRR*AFIY
Acheta_domesticus               PSGQ*RKT*RIQEIYPTYLSNP--TNRVRDTE---IHNTMSNIL----ICI------NKKIFTGSLN*I*HDNFLIL--KSL*NTKSRVALLR------------------------VH
Gryllus_bimaculatus             AKGVCLNCLFLQ-CYPSL**FP*QKNDFSD*AD*DLQNR*KTYLSAKHVCFQTCQRPTKKCFY---------DFTLFQ*KLYLNTDSVYSIQRQIIQMLSKPKESLISLLKTLNQNVFVH
Gryllus_longicercus             PGGGVWGCTLFS-LDP-LRPVPLALACVRELA--------------------SCPPPTRAAFVPARARNARRSRVITQNTAWAEEERAEAVAK------THPRR---------------H
Consensus

Teleogryllus_occipitalis        ------------------------------------------------------------------------------------------------------------------------
Laupala_kohalensis              F-------------WDALKC----FLI-ISAVCGI---QNHLSDP--------S**VN------TCYLSDPL-F*EYKWRLRNPPYV*NTFRK-------RSPNQISFLKLAK*------
Acheta_domesticus               R---TNGRIVL*L--QHLQCKVST----TPAM*G------RSKTP-------------------KILLHVVV----YLCYMGKKSATFSNSGL---IQ***TGNTVLNIFL----KI*LE
Gryllus_bimaculatus             RPF*THS--------KHL----------RKKYNG-HVHKMKSILPHEYYLQL------------KPYFSELL----FLDQMPRIPNHFNLLKL-------VSPKELFSLLL----CI*YI
Gryllus_longicercus             RPAS*RGDCGCGSTWTHLDAPGSSFLDYRFPFDGIYMDQR*FNLP*VYYF*LGSISENIFKNIWKIYLSFIVV*FGFLDRLNE*TWYLH*VRIEFGG*VNVLGYELD*LFVSDS*SVNYS
Consensus

Teleogryllus_occipitalis        ------------------------------------------------------------------------------------------------------------------------
Laupala_kohalensis              ------------------IEELIGMFKNPPSRNV-------ISSSK-----------PLI*MKFLVI-----------------LGVML----------------------------LH
Acheta_domesticus               CFIFTKIETRVV*SG----------IVRNP------VIHPNIKI*R--------------------------------------ITYQL------PKSLNFCCKEIHL*NYY*T------
Gryllus_bimaculatus             T-VTCQITTTLL*K----LLPTTQQF----------WLHMQIRHQR--------------IYKNT*ISCKRDYDWGVNFSKFQPITSML---------FK*QCAKMKLNDAYRNT**WLN
Gryllus_longicercus             GH*YCKCSACFCSEGGGGLPPLPGEIRRPPTAG*RGFPQPGISHQRGIAPVGTRSKLPTGLPACVVV*CM*V*G*G*GKEEVGKIARGLCIC*GAAQGFAW-C*SEKPGSTTDQSSRQLD
Consensus

Teleogryllus_occipitalis        ------------------------------------------------------------------------------------------------------------------------
Laupala_kohalensis              VSVLSAW*EL*TPTL*SRLRVNCTP*AHDT----QDLDSLMSQSN*RVLC-------------------------IAYTAW*V-----------------------SMFFPLLL*MRGT
Acheta_domesticus               --------------------L*SYMLTSLC----V*KHSILKS*KNRYLCFTNYN--------------------LTHTDVCL----------------------------LHLG----
Gryllus_bimaculatus             TS------------------NSTLTYF*C------RNNILQQQNR-----------------------------KQH*NFVI--------------------LQG*AIAVHMCLGK-AK
Gryllus_longicercus             LSF--------------RLRLNST*LTR*CG*SCHSRNSNVGPSS*GNYCDINW*RAIYPGLWVGY*FFYECLLDMHHGALVIIIIIIIIIIIIIIIIIMN*LEMICI*FRLVLA----
Consensus

Teleogryllus_occipitalis        ------------------------------------------------------------------------------------------------------------------------
Laupala_kohalensis              SYKVRSE--------ARQPRTTY----------LSPLEKMLKHSLK-ISNQLF-----LIYVNL*KIFEFHKKT*Y--*LLFLIFVVFKIG*I*QLTKTPS*K*LYST------FELVI*
Acheta_domesticus               IFEGLDRMKRLIEQMTVDKRSFYSFYDFIKI*CLSSKDLIIAHDNS---SQVY-------YCKVKN*KL**KSRNF--KVMYLCFTE**KNVLKLLEMP----------------LHLH
Gryllus_bimaculatus             LFSAKQQ--------SDQKR---------------ASKFSYGHHASKFNTEIL---------------GRFESK-YLRQILNAPL--YNDTVQHTYKSPP*TKK*YNAASPIP*AQTAIQ
Gryllus_longicercus             VFSHIDN--------AYSRKS-----PTIVVRFASAVTHTHTHTHTQTHTHIYIYIYIYIYCDVLGDAGTFTARNY*RRVSFS*MGPYEEST*H-------------------------
Consensus

Teleogryllus_occipitalis        ------------------------------------------------------------------------------------------------------------------------
Laupala_kohalensis              R-RFT*FWKPSLRGRFTMTGIFRSRNYISSHT**I*TKR**YLTCNICKNFPLFYI--------FCSTPFTPS*-MILCFYFGV---------------------YMK*FFSKCKLSSE
Acheta_domesticus               T----------------LSNTNRSSVLLVSGTYSSKKL**FIF--------------MKPGMRRIMMV--*GNK-LIRGLHYGADIDPPRRLLDDAVHLRKWEINGYLAEGLRKRLMKS*
Gryllus_bimaculatus             TP*PPIFWT------I-VKNLNDSR-DIIS*TYFSNPTNS------VRIHTDIVYVIRI*NITQCLIYLYSSTKTYL--LNV*I------------------------------------
Gryllus_longicercus             -------WE------I*LENV------VQS*TGFIHKPNQLIT*FIIPIHFALV---PRSGLT*VFIIPWGPNK-YLRCLNYG------------------------------------
Consensus

Teleogryllus_occipitalis        ------------------------------------------------------------------------------------------------------------------------
Laupala_kohalensis              Q-------RNRISKI*G*RQIKVSKAII*LTNIR----P---------YV*NTFRKRRSFQISF---------LKLAK*IVLII*----------------SFHCLQSKLRTTNCDK---
Acheta_domesticus               RLYSDIS*TRETEAPTGKGALQYRRFCCGAV*AKR*VSP------S*RHTLSTRYRPTAAPASYG-------------CTLAVNHAQLGAVLYRISYAAGS*GCIAPEAPETICSTCGL
Gryllus_bimaculatus             -----*STSHLVFS*NFYKTHCTSHCITSRTMHKRQNYP-----------IVILDRFIKVGFELGGKHDYN*N*NNYKTITNI*N*N*RFDILYLINF*L-------------------
Gryllus_longicercus             --------SHRQVSPTFTAAWRSRRLVV*RS-------PRLTAS*IPRAWLLVLRRRTSL*LSF---------------IVLI*LANTR-----PLHVSA-------------------
Consensus

Teleogryllus_occipitalis        --------------------------------
Laupala_kohalensis              -----TLSNIIRYKATFLTVYKEK--------
Acheta_domesticus               REGNLAIS-----PCGLLIELHEREI*HEG*D
Gryllus_bimaculatus             -----SL---------YLLLFY----------
Gryllus_longicercus             ----SVS----PHTVLVLFSLRQIA---
Consensus
```

**Tocc.2G0004080.1_exon3**

```
Teleogryllus_occipitalis      -----------------------------------------------------------------------------------------------CG-----------------------
Laupala_kohalensis            -----------------------------------------FPYKQLRM-----------------------------------L---------LCI**YLITFYHNSWF----------
Acheta_domesticus             I*F*HKIAI**LILHQE--------------------------YYNLQI------------IEK*TKS*EQSL-------TSR------LT*HRICGNSK--------------------
Gryllus_bimaculatus           -------------------------------------ASD*FNYSHVTITHLRHV*SRN*LVNSTKRGN*QCII*YEVY----------YQLHSVCAHHLVRS-YNNSFLARAQPAPRKL
Gryllus_longicercus           ----------YAI*RSENNTRTVCGETDADTCSGRVLAS----YIKTMKDN*REVRRRS--TRSQARGIQEAVSRGDRYTTSRRLRQAAVKVGDTC---------------RCEP*FKQR
Consensus                                                                                                     *

Teleogryllus_occipitalis      ------------------------------------------------------------------------------------------------------------------------
Laupala_kohalensis            ----------------EVLTVNSEMIK*LKR-------SILPVLG-TKFGK--TFFS*KYFRHRDE--YLSIKLWLWKLLSGVTLIF*IFCFSV---LSSA----------YI*KKI---
Acheta_domesticus             ------KGNV**HIW--------------DH--------FSPL*N*TKHINEQTSVTICYGQS*DNQGYW-------LNRTQRALYFYLFLI----SSKT---------LHSCQKLEQK
Gryllus_bimaculatus           ------HGALRVNRR*PQCASDTSCVGA-------------PV------------LSRCGNQRLPKPPFMTV---MFLLKRYM*ILV*FLCFNVRHDSPST----------YLCHSF---
Gryllus_longicercus           RYLLGPQGMMKTHVK-PDLGTKAKCIGIINHVIN*FGL*INPVYDCTTFSN---*ISQCYVDSS*GP-----------IHEKDTLL**FLAVKVPA-SPSTSQYIYIYIYIYMCVCV---
Consensus

Teleogryllus_occipitalis      ------------------------------------------------------------------------------------------------------------------------
Laupala_kohalensis            -------ISCT---------LQNRN---------------------IKSFSSE*RE*SRKYRKEGNFYKYC----KLNIIIALFIFIKYERKYNFCS*RYPSL*I--------YRVNLAS
Acheta_domesticus             IIMIFLRTICF---------CFERT----VLFN*T*L---------IKN------SLLLSSNKNNSVSKFCV*HNAKFVITW*QSELHYK-PYK-------QL**LVSK------SNMYN
Gryllus_bimaculatus           -------RVCFKCLVMVYYYCFKLTRTSVCFFTLHKLYPYEVSQYLLINYTY*FMSSTLNSPNDNNLLGITFLFQKFESVCIRLQ*IHYS--YEAIKRR*WILY*L*GIRLSRFTANYTN
Gryllus_longicercus           -------CVCV-CVCVCVTALAKRT-TIVGLFLLYAL------------------SI*LNTAK-TNLN*IQIISN*FIIIIIIIIII--------------IIIIIIII--------ITN
Consensus

Teleogryllus_occipitalis      -----------------------------------------------------------------GG-----------------------------------------------------
Laupala_kohalensis            KIK*ISAILQAR--------KYYIT----------IFTKEF*LI-------------------------------------VKFNRF*KQQ---------K*E-----------------
Acheta_domesticus             SL*FY---ISPCHIPRTNICNII---------QKTILLELYMTVMFLPIAL*I-YALSFELRILIGGPHS*IGSHPHPSSIFSFAPELPAR---SS*GQ*KCKMLCRSDYISYCLIIY*L
Gryllus_bimaculatus           SIICFKN-YSVRHLIHRV*VN*Y---------RGQ*HSE-FSLLIYNCRSLDLTYVPTFL**E-------------------***R****K***SS--------------------VAVW
Gryllus_longicercus           APWCISNKHS*K--------N*YPTHKPG*MARHQLMSQ*FP*L----------EGPTFEFRE*QDYPHYLV-------S*VEFSRSLKDK---SS--------------------CRLL
Consensus

Teleogryllus_occipitalis      ------------------------------------------------------------------------------------------------------------------------
Laupala_kohalensis            KVINITFFYETQRSFKG*HKSRRADLIFLDCVLASFPAAISTSSSV--------------------ALLLI------SLYM---------------------------------------
Acheta_domesticus             KSILLNYLNDSNF-----------------------PRIMYFISYLQYLHYKNHTGSSRSYCW*VV*LVIIIGHPPESVYCLRMPSLPRKY*KMALFVYQ**FCEDYSILDSRNEETINT
Gryllus_bimaculatus           KRNLVAGCSDR*YSLASPISTRRMST*AVP*QIGHKPRAIFPTSSLHYPHPYTDMHYTTTHA----------GRPVGSLELVPAGPVPRWW-----------------------------
Gryllus_longicercus           *SVVLPGFSDQHQ--ANP--------*AAP*QM-HKPRAIFPTSSLPYPHPYTHMHYTTTHA----------GRPVGSLDLVPTGAMPRWW-----------------------------
Consensus

Teleogryllus_occipitalis      ------------------------------------------------------------------------------------------------------------------------
Laupala_kohalensis            ----KFPA------FKAVTGRT*RLT---------------TLYRQYREPFSCSETLNYQGLEYHELTVCS*LSNEITE-----LVFITLTRHLTLKHVTT*HPVL-----RGTSFKLM-
Acheta_domesticus             NV*QTYPSKFINVRYQAL-------------K*KSHFSI*HSIDS*YVIAN*IY*NRTFLSV----FVTFTIFENSESSTK*TCILF-------------------------*ASFARFF
Gryllus_bimaculatus           ----QIPG-WGNPR*PAVGGR--RISP*SGGRPPPQFEQTHTLHLQY**PE*LTDQES--------------LTNSQSNA----FIF---------------HQIL-F*RNSDTEFNRIQ
Gryllus_longicercus           ----EIPG-WGNPR*PAVGGR--RISPGSGGRPPPPSEQKHALHLQY**PE*LTDQES--------------LTNSQSNS*PNTFTYP-------------PNSILT*CRYQVYSFN---
Consensus

Teleogryllus_occipitalis      ------------------------------------------------------------------------------------------------------------------------
Laupala_kohalensis            -AWRKKLRFATEDS*TFQLTL------QSILPVLGTKFG*GFFS*KYFRHRADS*VATYTLRIRDRL-SNKCLLIMRGHLNGFVFHRRHL**ENT*VHPRNK*MLIC-------VAFGNV
Acheta_domesticus             -----KKRTA*KNEQFWSTF*KR*GCTCKVLKK*NIRFRLPGY-RV*LRKNM----KKYQFWRKTYE-*LRL------------------------*HLQ*IEELIYYIDILQSTTF*YL
Gryllus_bimaculatus           PCQKPKRYNNNKR*VYFPLLYV-----ENTFSEMDPK*KQ*TYGRLHYR*SM*TPSEGKR*SKMEQP-VSRA-------------------------HPQ--------------SPL*LA
Gryllus_longicercus           -LSRNPNYTTIKDKYIFQIFL-------KIFSEMDPN*K**TYGRLNYR*SI*IPSKGKR*SKKELPGASRCVQV--------------------LPHPQ--------------SPL*LA
Consensus

Teleogryllus_occipitalis      ------------------------------------------------------------------------------------------------------------------------
Laupala_kohalensis            RERRMSILYFVNCMLLTSV-AILCLFIYDIEL---------------------------------------LFVHLSYPLN--FVASGVNFLTCCGSLFCILSVLFICDQ*INLFI*IVL
Acheta_domesticus             GAPKRG-LWLHNTL*YTSYRNNICQLFHNEVA--------------------------------------TI*TNSCYPA*LH*FIKNKNIALRSG*F------S*RCRLNIP*LVRCIV
Gryllus_bimaculatus           GRCRRG*VFATASALSSSARAVFCVITRERQAFRARARGHKGRTGGRRTRGEHTHARERQRHGAEGSRENKVQPQTAPPGA----ARGESFDL-AGLL------LFGSRREVNFYARCVL
Gryllus_longicercus           GRCRRG*VFATASALSSSAHAVFCVITRERRAFRARAGTKAARVGGGQEASSRTHASARGT-GRRGSRENKVQPQTPPPGA----ARGESFDL-AGLL------LFGSRREVNFYARCVL
Consensus

Teleogryllus_occipitalis      ------------------------------------------------------------------------------------------------------------------------
Laupala_kohalensis            -LRFLVNNN-------------------------VYFKRVYF-N**L*FTITFN**VT*N----*RNFI--SLIRLKI*TI---------LHLYFIKAL**N------NLSTLYLIFIIN
Acheta_domesticus             -MSALSRDRLPKQ*S*CEVNYMNEPVLLQ-NGTDFPILVIYF----L**LFAPDPYSSELPCIAAR-----STFYCHILNFSVF*L*NRNIILRILK-----------NSKTSS*MAIFG
Gryllus_bimaculatus           TLRALTY*R--------QLNEL*E*FFAKFNGCKVIIKNVYF-----------------------------EF*YF--------------*IIRILKKL*YNSQE---NTEKWNYIFVYQ
Gryllus_longicercus           TLGALTY*R--------QLNEL*EWFFVKFNVCKVIITKCLF*NFII*KLYVFKIIVIQL----ARKY*KMELFFC--------------LSIIVLRR**YSKGKK*YNTKFCRWIRKFD
Consensus

Teleogryllus_occipitalis      ------------------------------------------------------------------------------------------------------------------------
Laupala_kohalensis            K*KIRIT*ILNVCVGHVLVRLYEN*VYSGPVVKNLKPIHNSDISPQHF*NTVFVKFLKFS--RIHHAFLQTFASP*N*NIMSTGFLASVIHIRST*VGANPHFNNQ*F----LK*SIKCC
Acheta_domesticus             K---------------IWSRI*ENGV*ALL*I*N-------------FIGLFHRKVLSFR--KTSLEFI--*STSTYKYSSIDKLLE----ICTLTVG---EIFQKNYICVIFEKNIL--
Gryllus_bimaculatus           ------------------**ICEDD-----------------------------NILKSRN-DTIRNFADGYERLTTISIAINTY*E*STKIRKFATW---HFMMQHLICE-FSCWII**
Gryllus_longicercus           ------------------NHIDRN*-----------------------------YILRIKH*NT*ICYIIFYDVTVDK*IFVLDYL-----VTRVTST---NLTLTHYL*L-LQSRI---
Consensus

Teleogryllus_occipitalis      ------------------------------------------------------------------------------------------------------------------------
Laupala_kohalensis            FGIANEYKC*L--IICGDAGIRP*YHTFNL*YLFSYIFFLFIKQREKYKSIISN*LINN*RIK********FDHLTTA*NEVEA**ALHKI*IRPLKVVSLILF-I*LDYIIDVNCCRIL
Acheta_domesticus             LSCTERTRIYF--DVNNI*EPQS-----AVPASFST*VQFKRSSTQKPTSLFRDSTVRPVSI---------FRASVASFAETKFS--------RDGLLYKMVAVHVYIALLAKSNYW**R
Gryllus_bimaculatus           LGLRQKTSL*L--IICNFCNPEYSEYSLKLICGVSKH--PLLSLRKKLMHTFRAISKK---------------------------------------ILKVLRFHVMCPEISTLICFDCA
Gryllus_longicercus           --FREFTEMNLRSF*ASFA*SEK-----KINAFISSHFQENFESSKTPCYVPRNINVN---------------------------------------LF*LRLYEIHIE*LTESNCWE*R
Consensus

Teleogryllus_occipitalis      ------------------------------------------------------------------------------------------------------------------------
Laupala_kohalensis            N*IQHYLSLMS*PEVLLSYQHIGTLPIIHIPITEEIGTKSSLPS---------------DFKYCSISKNA----------------------------I*ITNDIKARLLV--------S
Acheta_domesticus             DDADR----------------------TRFNETH*TNRKKM--------------------YTCNDL*LH-IIHLYTAITILYSRNVTT*SI-------*MMDDLEKILLVVQIFAD*RT
Gryllus_bimaculatus           FMKYT----------------------LHD*QKVTVGNRGMTPTEQDLIKTIEPMVRKCDPFTCYVA*QNSTSYLHKTITIMRNRNVTTYICT*PTWLAWIMNDLETILLVGYISNDS*T
Gryllus_longicercus           NDANR----------------------ARFNQNH*ANGKKM*PS-----------------YKCSLTEFN-------FISS*GNHNYE*RIC----------NNLHL-----YLSNLGSL
Consensus

Teleogryllus_occipitalis      ------------------------------------------------------------------------------------------------------------------------
Laupala_kohalensis            VHKKYLV*L-------**KIM*FHKNTQT*LKI*I----MHFNKTWLLLF-L*SN*HHLKHVRPCPFVY*LAVKLHRNDTL*RFLSYFVNVT*AFGTCVFLMWFFFLRQFSHFLFVLFPS
Acheta_domesticus             LLENLSEL***HLTFS*TSIMFFFTNPLLRST-------FHL*ETASF--------------------FILNTIM----NVIVIKMLLVDDYFKFCLYFFLYSFP*KVFSTTLFIE**MK
Gryllus_bimaculatus           LFEKPI*GL*ETSDIILNQAHLFLHNSIFE---------FHFPL*KRVLFNLYYEYLYDCYHKKPI*RLLL*I-------LFNYFACILFFEVGFSTTLFME--**KKFFRHPYKSSTKK
Gryllus_longicercus           DHER------S*NDFTGGLYI**LMNSF*KTYLRSIRNC*HFPKPSSLVFT*F*F*IPLSPVKKGSF*FLL*ISL----*LLS*KSYLKIIAVNFV*LFFLYSFL*GGFFYHLIYGVMKE
Consensus

Teleogryllus_occipitalis      ------------------------------------------------------------------------------------------------------------------------
Laupala_kohalensis            IFFSFVL*----------TLRRSGAK--------SRS*I----------------KVL*T*NHQE*A--------------------SNAFAYILLPHWR--------------------
Acheta_domesticus             TFDF-LIKYH*KV*L---KLH*QEEG---------R*RL--------ARLLGHPKKLRRRLRHSRGAPIRNEAPLTAPRPSASRARRSNIYANVFGAHSRSL*VHRTLIYEQFKKPEPAP
Gryllus_bimaculatus           V**K--LQ*HEKAKVTPWQIARTAEG------------------------------ASGRPRHARGAPIRNE---------ASL---------------------------------PAP
Gryllus_longicercus           ISPTPL*KEH*KSIVKT-TITRKGEGNALADSSDSRRSFGGAHDTLAARL-----SETKRPSPPRSAP-RHS---------ASRARRSNIYANVFGAHSRSL*VHRTLIYEQFKKPEPAP
Consensus

Teleogryllus_occipitalis      ---------------------------------------------------------GGGGGGGGGG--RGGGGGGGGSGG-GSGGGGGGGSGGGGGG-------GGGG-----------
Laupala_kohalensis            -------------------------------------MEL-----------------------------SGGGNLEG*RLG-------------------------------VESW----
Acheta_domesticus             ASAEPCRAWGLGEAR---------------------------**KPRRYGQKRVGRRGGGSGGGGGG--DGGGGGGGGGGGVAAEAAGGGGDGGGGGGAAAGRRRGGGGTVAVAAWRGWR
Gryllus_bimaculatus           LRPTPLRV-----ARTPE*YICKCIWSPFQIIISS*NFDLRAV*KARAGARER----GAVSSAGAVGAVMKGAAVWEGKSG--------------------------------*AW----
Gryllus_longicercus           ASAEPCRAPGR-*AR---------------------------**KARRYGKE------------IVG--RRGGGCGGGR-----------------------------------------
Consensus                                                                            *

Teleogryllus_occipitalis      ------------------GG------------GGGGGGGDGG--GGGGGGGGGGGGGSA-----GGGGGGGGSGT---------------------------------------------
Laupala_kohalensis            ------------------NG------------VAWGGGGEVT----------------------SGLRVRVLSSSAL---SP-CFHTFLGNPFLYTQRVIV*KL--REIILFIN-----F
Acheta_domesticus             RRGRRWRRGWGRRRRRRQGGDVEAAGRWLWRRGGGGGGGDGGVDGGGGGDGGGGGGGAAAGRRRGGSGAGVAAGTAAGRDAPSLISTAADRA---SYRVAHYRSGIRE---FKTRVLRMF
Gryllus_bimaculatus           --------------------------RWLWR-VDGGGGGGSGK----------------------------------------LLH*FPQPP---TARLIGSRTTDRE---FEN------
Gryllus_longicercus           ----------------------------------GGGGGPGK-----------------------------------------LLH*FPQPP---TARRIGSRTTDRE---FEN------
Consensus                                                        ****

Teleogryllus_occipitalis      ------------------------------------------------------------------------------------------------------------------------
Laupala_kohalensis            NDL----GC----INVF*----LKT*NIF-----LEIIII*WNSKVKFRGKKKF*WKNCK---------FQMV-L*H*PNAVHLQFDSSIRSNT---*VKIKY*ILLIFYFNKYNNWNK*
Acheta_domesticus             LEMLFS*KCKMTRINSAMMESYRRSHNILAKASQNQA*I*-----TKLY*VLRVQCWW*KLPIKRL*KSLSLI*LSQQ*YSYMKKM*PTFRVDSISCAIDEEQLVNVFVKHNFVNYRAKS
Gryllus_bimaculatus           ----------------------RKRENI------PENVIL-----QKMQNDKDQLCHGGK--------------LQEIAYSVTK--------------------LSLNIHKNIFNPKSAM
Gryllus_longicercus           ----------------------RKRENI------PENVIL-----LKMENVKDQLCHGGK--------------LQEIAYSVTK--------------------LSLNINKTIFNLKSAM
Consensus

Teleogryllus_occipitalis      ------------------------------------------------------------------------------------------------------------------------
Laupala_kohalensis            IVLIR---MIT---------------------------GVFCTSSMRL*HEEK*LLRSSLYYLQNKFHTL*LKDKK*DMKIINEKNVLKRTCCT--FGRLVISSVLECFKTY*FYEREFL
Acheta_domesticus             LSAWRAGFVIERAGFDPRVGHPGRGFRGFLWS*GEYWDGFKIPRTPPTFHYPT*HLPPSF----*SFHTSSSPSPMMMIIIIIIIIIIIITKITSSYHLIIDYSRQQSC-----------
Gryllus_bimaculatus           LVV-----------------------------------GIANKNAIKTI-------------------TASLPKSTMFT*LVDEKNVSRKLMYTPFWAPLIRNS**MTL-----------
Gryllus_longicercus           LVV-----------------------------------GIANKNAIETF-------------------TASLPKFTMFT*LVDEKCISEISVHN-----ILRAIDKE*L-----------
Consensus

Teleogryllus_occipitalis      ------------------------------------------------------------------------------------------------------------------------
Laupala_kohalensis            K**LLQIN*YLR-----Q*IG*YIA*YY----------CRCLCNE*N**FNVVIKSIIGMTKLIVRMPIISV-EPNRHRLLNYDLFSYLGESIVYIFLVILPFPMIHFLVSRYLIFRRP*
Acheta_domesticus             ------LNQGMRP*G*CP*FHTFISPE*STTGPRKRETC-CVVSSGPYANN---------NNNFVNYP*PE--EANEMKVN--------------------------------------F
Gryllus_bimaculatus           ------LN-------------------------------------------------------IIL*PIHDVKKLK**RL----------------------------------------
Gryllus_longicercus           ------VNDFAK-------------------------------------------------HYFVTYP*RE--EAKEMKV----------------------------------------
Consensus

Teleogryllus_occipitalis      ------------------------------------------------------------------------------------------------------------------------
Laupala_kohalensis            NNNKFDDVIYR---CYY*YYYYYYWDFRALRHLGH*NSIAEYLIV---------------LNYTLLRHFLDKTPNYFTR---NNSSASLYI-----------RHIFSP*F*CTYKD----
Acheta_domesticus             KMGRGSDGKQ----------------------------NRSWMLCVTP---------TVWIY*TSVGGVQGEST*NF---VTSQG*SIV--------------HI------C--------
Gryllus_bimaculatus           KMGRDSDDKQRVTICYFQIRTYY----TSARLLYADGSNYTLLVCKT*LP*GTQM---------ARRKTFSKETNVFSRCAGDHGSEAYCVECKAC*LCENKE*I------CLSKGL**E
Gryllus_longicercus           KNGRGQ**Q-------------------AARHYMLLPSTDVLYLCAIIISRWKQLHLTCM*NIASIRYTNGSSKDLF------*GS*SV-------------QHV------C--------
Consensus

Teleogryllus_occipitalis      ------------------------------------------------------------------------------------------------------------------------
Laupala_kohalensis            --------------------------------------------TLINYINVLIFSSHNIIA*R--IKLN-------NRKQSNK--------------------------PS----INRD
Acheta_domesticus             ------------------LCKAKCSSTKQQSDK*RVT-------*LGASRK*IQYR-NSL*VS---IKIP---------PSKSQ-------YNLIQQH-----CITYLQIPSMHE*INST
Gryllus_bimaculatus           ICRA*GSN*GTNNTHYLRLLKPFESVLVELKQQRRES---------GNIFHCFGYCMHSRAMSVQ-LKISWH*PFNYNCAGQSK*C*SCNGINTLPRHTCF*NCTIYH------------
Gryllus_longicercus           ----WGQRK*T-------LLRRV*SLLIV*EQRRKMSVKRSLMRNMQSLKKQLRYK*HALITALKAIRIS---------AGRDK---------TLNLHLTYI*CTKFFY----------T
Consensus

Teleogryllus_occipitalis      ------------------------------------------------------------------------------------------------------------------------
Laupala_kohalensis            QVYIIIPLV------NKSR*CLT*---------SNSPL*F---YECFFLLTRG---VYNKYVS------------CPIS*RFL---------------YPS----NGLLLSSDLFCLWVL
Acheta_domesticus             QHHL----------------SSTPQQPSKNP--YNRP------SGQ*RKT*R----IQEIYPT--------------------------------------------------------Y
Gryllus_bimaculatus           --------------------CGS*DVQGK----LRSPRTCWT*EYCVYQTRR----VSKIY*EIYLILYKHG--------------------------YKSQIM*EGLVM*KMYGILAFY
Gryllus_longicercus           QDYLII*FSF*QFGNYRKRPCEQDKML*KNLILLHRPYSLRLVSACCILTKRNSLTVARIPRTKTVLIVKQSISFIPNSSNILGNYVFRRLA*D*CQTYRPVIVNNPSLVNRHYNCLFPI
Consensus

Teleogryllus_occipitalis      ------------------------------------------------------------------------------------------------------------------------
Laupala_kohalensis            LH*GF--------LYVREFKFSVLFFLSFLCN*HRN------IYIVSCK----------------------------THFLYSSINLFLIY-------YIQE----*EHLSSSSSSSFYQ
Acheta_domesticus             LSNPTNR--------VRDTEIH---------NTMSN--------------ILICINKKIFTGSLN*I*HDNFLILKSL*NTKSRVALLRVHRT--NGRIVL*----LQHLQ-CKVS----
Gryllus_bimaculatus           LHEETQRKNK*FILLIIPFNYHLQPIPFFVK*KKSN---TQFIYA------------KKSPGCDFIVGEIQELQLKGFALIASSFNVIPLYNDFPSKRMISQ----IEQIETCKIAKKHI
Gryllus_longicercus           LRNHS----------IIYSNFH---*PRKVRNLSRNGFC*EFVHIVTCRCILMLMNNKMGKVCKFWINTIN-IHWKHIKHVSE--NVIEMY-------YIQQDGLVVNYMK*CYPS----
Consensus

Teleogryllus_occipitalis      ------------------------------------------------------------------------------------------------------------------------
Laupala_kohalensis            FFQNLF--QKNTILR-----------------------VYLPT---------------------------------NLHSQ*CNFTQRK----------CNQRKKNMNEEQ*--------
Acheta_domesticus             -------------------TTPAM*GRSKTPKILLHVVVYL-----CYMGKKSATFSNSGLIQ***TGNTVLNIFLKI*LECFIFTKI--------------------ETRVV*SGIVRN
Gryllus_bimaculatus           FLPNMSASRHVKGLRKNVSTTSLSSNRSYT*IPIQYT-VFKDKSFKC*ASPRSH*FHYS-----------------RL*IKTFLFTG------------PSRRTVSI*EKSTTAMSTK*N
Gryllus_longicercus           *FKNLT---HSLFIRRIMRITGFLL*KLHTYFKILLAIVFIDT-------------HYS------------------LHLKTFSLQNLKNSLSSLR*L*PHTLRCSLAEIRYS*RS*HNE
Consensus

Teleogryllus_occipitalis      ---------------------------------------------------------------------------------------------LLH*-----------------------
Laupala_kohalensis            -----------LSVYLKEVPAN*C--------IVLKGNVN------------------------TSRMLLVPSWNLKRTHTK--------------KSRARALSSLNLLGLQRHIPGKG-
Acheta_domesticus             PVIHPNIKI*RITYQLPK-SLNFC----CKEIHL*NYY*TL*SYMLTSLCV*KHSILKS*----KNRYLCFTNYNLTHTDV-----------CLLHLGIFEGLDRMKRLIEQMTVDKRSF
Gryllus_bimaculatus           PFYHMN-----IIFS*NHISLNYCF*IKCQEYLIILIY*S-WCLLRN--CSHSYCVFNT*Q*PAK*RLHYCENFCRRRSNFGYTCKSGINVSTKTHRSAAKGITTGA*ISQNSNP-----
Gryllus_longicercus           AMY*TNDGR*AIIL------------------LIL*FY*T-SVFIRKKRCLHS-----------KTHLESTVSFKAKITQFG--------------------------LDESPNI-----
Consensus

Teleogryllus_occipitalis      ------------------------------------------------------------------------------------------------------------------------
Laupala_kohalensis            *KLLTLVRPIVGKLETAFIFDRPLED*YW*IYPY*IVTREN------VLLRRY*RN--------------NL*FFIIHVLDFHNKK----------YSLCF---------*LK*TLLFSI
Acheta_domesticus             YSFYDFIKI*-------------------------CLSSK-------DLIIAHDNS--------------SQVYYCK----VKN*KL**KSRNFKVMYLCFTE**KNVLKLLEMPLHLHT
Gryllus_bimaculatus           -----LLR---------------------------CFSNNNVPK*N*TMPTATRNNG*ILLIPP*PTFNVEITFY--------NNK-----------------TGSNI------------
Gryllus_longicercus           -----FIKKIP-----------------------LCFK*YN------DIVTKF*SN--------------DVVFY*------TNKK-----------------*CSDIVPL---PLLLHI
Consensus

Teleogryllus_occipitalis      ------------------------------------------------------------------------------------------------------------------------
Laupala_kohalensis            FK*LLKLSHNAVLGNIKQIKL---------LRKK----ILLLTVKKTFF-LP*CLDIIASYRM--DFLKFESGITYKYSNNLF-----------LS*R-TPSSLA*SDVH----------
Acheta_domesticus             LSNTNRSSVLLVSGTYSSKKL**FIFMKPGMRR-----IMMV*GNKLIRGLHYGADIDPPRRLLDDAVHL-----RKWEINGY-------LAEGLRKRLMKS*RLYSDIS---*TRETEA
Gryllus_bimaculatus           -KISSSCKVKQSPFTCASAKQNSFLQSNNQTRKELQNSAMVITQVNSIQ--KFWADLNQNTSV-------------KYSMHPYTTTLFNILTNPLHER-------RNNTTQHHLSRKPKQ
Gryllus_longicercus           FRNTNSINVL*VSGTYSSMKL**FISMKPDMRR-----IMVV*ENKLIRGLHYGADIDLPRRLLDDAVHL-----RKWEINGY-------LAEGLRKRLMKS*RLRSDIS---*TRETQA
Consensus

Teleogryllus_occipitalis      ------------------------------------------------------------------------------------------------------------------------
Laupala_kohalensis            -----------------------------------------CLS*FHII-----CNSFILFLH----------VNRNS-----------------RIIKK-----------------TVV
Acheta_domesticus             PTGKGALQYRRFCCGAV*AKR*VSPS*RHTLS-----------TRYRPTAAPASYGCTLAVNHAQ--------LG--------------------AVLYRISYAAGS*GCIAPEAPETIC
Gryllus_bimaculatus           PSKHPNHRSSGQ**KI*TIQEISYPKLTFPIQLIVYVFILILYT*YEY---ET*HNV*YTYIHQQKHIY*MFKFNQHP-------IWFFRKIFIKHIVRRIVL---------------LR
Gryllus_longicercus           PSGKGGSAVQEVLQRGRLSEAVSFPKLT---------------THFKH*-IQAHGGARFLWLH----------LGSKPCSTWGGAVW--------NIIRRGA*-----GCIAPEAPKTIR
Consensus

Teleogryllus_occipitalis      ------------------------------------------------------------------------------------------------------------------------
Laupala_kohalensis            *NMNE*H*RLINN**RINKLPLL-ELY*SQI*PFGWNSAT*NN---------------MLREKYYKKLCLDINKYNKNTN-RFY*DIQIKLSF--IHSD*LS*SHNKTSHFIYGKVNNLP
Acheta_domesticus             STCGLREGN-----LAISPCGLLIELHEREI*HEG*D----------------------------------------------------------------------------------
Gryllus_bimaculatus           EQCT--KGRTIQ*LY*IDLLKLVLSL------------VAN-------MITIRIRTITKLLQTFK---------------------IRIRGSI--SYI*LIS-----NYLYIY-------
Gryllus_longicercus           SSCSLREGR-----LAISPRDLLIELQERDI*HEGKD*IKNKGYAGTRVFMVLNIKT*LLYFKTYTNVWFVVITYSMS*NV*IQVC*RVECSLKQNYT-LL*-----TFRFIFI------
Consensus

Teleogryllus_occipitalis      -------------------------------------
Laupala_kohalensis            KQEKIFHNIV*-------KK-----------------
Acheta_domesticus             -------------------------------------
Gryllus_bimaculatus           --------FYF--------------------------
Gryllus_longicercus           ----IHHNFTFFLRIGDKRK*CCAKNGKF*AEKLLLQ
Consensus
```

**Tocc.2G0008730.1**

```
Teleogryllus_occipitalis        AAGGTATTTTAG-AAAAATTACTCCGTATCCGTCTTCTAAATAACCATTGGAAGTCTATTTATTTCCTACCATTTCGTGAA--------TGTTTATGTTTTTTTTCTTTTTTCCCCTAAA
Acheta_domesticus               AAGCTATTTTAGAAAAAAAAAACTTCTTATCCGTCTCC----------GTGAAAGTCAATTTCTTTCATACAATTTCGTGCATTTGTTTCTGTTTTAATGTTTTTGCTTTTTATCCCTCAA
Consensus                       *** ******** *****  *** * ********* *           ** ***** **** **** *** ******** *        *****   * ***** ******  **** **

Teleogryllus_occipitalis        AACTGTATGGAAACCAAGTATGT--GTGTAAATAGTCGT-GAAATTAGCGTTGCACATATTCAATGTTGTTTTGTGATAACTACGAAAGCTGAAGACAGGAACCTTCAA-----------
Acheta_domesticus               AATTGTTTCTTAACAAAATATTTTGGTGTAAATAGCCATGGAAATTAGCGTTGCACTTATTTAATGCTGTTTTGTGATACCCATGAAAGCTGAAGGCACGAGCCTTGAAAACCTTGAAAC
Consensus                       ** *** *   *** ** *** *  ********** * * **************** **** **** ************ * * *********** ** ** **** **

Teleogryllus_occipitalis        ---CAACTTTCAAATGCGGAGCAAAACGACAACTACCCTTAAAAGTATTTCAGGTTAGAAGAAGTCATTACGTTCCCAGGTTTGGGATGACTTTCTCCATGGTTTCACGGCGGTAAC--A
Acheta_domesticus               CTTGAACCTTGAAATGCGGAGCAAAACGTCAACCATCCTTAAAAGTATTGCAGGAAATAAGACGCCATTTCGTTCCCAGGTTCGCGTTTACGTTCTCCATGGTTTCACGGTGATAACCAA
Consensus                           *** ** ***************** **** * ************* ****  * **** * **** ************ * * * ** ****************** * ****  *

Teleogryllus_occipitalis        TTCTTTTATGAGTGGATAAGAGCCGTATTT-----------TTATAAGAATTTTGTAACGGATTTTTCAGCATCTGCAGTAAATTTTGCAGAAT--TAAAAGAAACTAATAAATTAAAGA
Acheta_domesticus               TACCTTCACGTGGGGATAAGAGCCGTACTTTTTTTTTTACAATTATATAATTATTGTAATTATTTATATAAT-TTGTAATAAATTTGAAGAATTCATACAAAAAACTAATAAATTAAAGA
Consensus                       * * ** * * * ************** **           *****  * * ******    ** *  *   * ** * *******     * *  ** ** ******************

Teleogryllus_occipitalis        AATATTATACATTAATAGGGCTCGAAAATATAGAGAATATAATAAAAATAACATC-GTAATGTCAAATAATTTTTACAATTAATTCGTAGCTGGTAATATCAATATGAAACAAT-TGAAC
Acheta_domesticus               AATATTACATATTAAGAGAGTTCGAAAAGATAGAGAATTGAAAAAAAATAGCACTAGTAATGTTACATAAGTTTTACAGTACATTCGTAGTTGATAATATGAATATGAATCAGTGGGAAC
Consensus                       ******* * ***** ** * ******* *********  ** ******* **   ******* * **** ******* *  ******** ** ****** ******** ** *  ****

Teleogryllus_occipitalis        GTTTTCTCGTACAACGATTAGTTTATCTTTTTGAAAGGGTTAAA---ATCTTTAGAACTGCTAAATGATTTTTATATTTTTTTACACTATATGGATAAGCAAATTGCT-AAAAATGCGAG
Acheta_domesticus               GTTTTCCCGT---ACGATTCAATTATCTTTTTGAGAGCGTTGAAACCATCTTTAGAATTGCCAAATGATATAT---TTTATTTACACTATATGGATAAGCAAATTGTCAAAAAATGCGGG
Consensus                       ****** ***   ******   ************ ** *** **   ********** *** ******* * *   *** **************************   ********* *

Teleogryllus_occipitalis        AGTGAAACTGAATTGTCTAACTTAGTTTTAAAAAGGGCTCCCCTAATACTCTGATAACAGTGTTCAACATTTCGTGTCGAAAGCATAATAGACGAGTGGGTTGCAGTTACACCAACAGTT
Acheta_domesticus               AATGAAACTAAATTGT----CTTAGTTTGAAAACGTGCTCCCCTAAAACTTTGAGAACTGTGTTCAACACTTCGTGTCGAAAGCTTAATAGCAGCATGGGTAGCAGTTACACCAACTGTT
Consensus                       * ******* ******     ******** **** * ********** *** *** *** ********** ************** ******  *  ***** ************** ***

Teleogryllus_occipitalis        CATGCGTCATCAAACGCCGAAGATGGCAACGGACAGCAGACATGGCACGGTCGAATGAGGTGGAGGCTGTCAGGGTCAGTGAAACCTGGAGCCATGGAAAGATACGTAGGCTTCCCCTCT
Acheta_domesticus               CATGCGTCATCAAACGCCGAAGATGGCAACAGACAGCAGACGTGACACGGTCGATTGAGGTGGAGGCTGTCAGGGTCAGTGAAACCTGGACCCATGGAACGATACGGAGGCTTC------
Consensus                       ****************************** ********** ** ********* ********************************** ******** ****** *******

Teleogryllus_occipitalis        GTCCCTCTGTAAATTGCTAGGACAATCTAGCTGACCCCAATTAATTGTTAATTTAACACCTTCACGTCGTTTTCACCTGAACGGCTTTTTTGAGAGCGGTAGGCGCCGGTAAATAACGGT
Acheta_domesticus               --CCCACTGTAAATTGCTAAGACAACCTAGCTG-TCCCATTTAATCGTTAATTTAACACCTTCACGTCTTTTTCACCTCAAGGGTTGTTTTAAGAGCACTAGTCCCTGGCAAATACCAGC
Consensus                         *** ************* ***** *******  **** ***** ********************* ********* ** ** * **** *****  *** * * ** ***** * *

Teleogryllus_occipitalis        GGCGTTCACAACATTCCGTCAGCGCTGGCGGCAACAGAGCGAGAGCGCGAACCAGCACCGGCTG-----------TGCGAGAGTGGGACACGTGGGGTATGGCGCGTGGGGAGCGGCGAG
Acheta_domesticus               TGCGTTCTCTACATTCCAATAGCACTGGCCGCAGCA---CCATCGAGCGAAACAACAATGGCTGCGAACGCCATACGCGAGAGCCCAACA----GGGCATGGGGCGC-----------AG
Consensus                        ****** * *******   *** ***** *** **   * *  * ***** ** **  *****            *******    ***    *** **** ***            **

Teleogryllus_occipitalis        AGTGGGGCGGCGCGTAGGGGGCGGTTTGTAAGTGTGTAGGGGGCGGTTTGTGTGTAGGGGCGGTTTGTGTGTAGGGGCGGTTTGTGTGTAGGGG----CGGTTTGTTTGGGGGCGCGTTA
Acheta_domesticus               AGTTGGGCGGCGCGTGGGGCG---------------------------GCGCGTGGGGGGCGGCGCGTG-GTGGGGGCGGCGAGGGGGCCACAGGCACACGTGTCGCTTGATGATCGGCGG
Consensus                       *** *********** *** *                           * * ** *******   *** ** *******   * *   *   **     **   * * ***   *    *

Teleogryllus_occipitalis        GGGGTCTGCGCGTGGAGGGGGGCCGCG-----TGGAAGGGGCGGCGT---------------------------------------------GGGGGCAAGGGGTCGCGGGGTCGGGG
Acheta_domesticus               AGGCAGGGGAAGTGGCGAGGGCAATGGCGGCTGAAAGGGACCGCACGACCGCGTTGCGTGACGCAATCCACAACAGTTCGCGTTATACAGCGCGGGTGGCGCAACGCGGCGGCGCCGGGG
Consensus                        **    *   **** * ****   *     ** ***** * **                                             ** ***     *  **** * *****

Teleogryllus_occipitalis        GTTCGGGGGGGT-------------CGGGGGGTTCGGGGGGTTTCGGGGGGTTCGGGGGGTCGCGGTTTCGGGGGATCGGGGGGTTCGGGGGGTCGC----------GGTTTCGGGGTTCGG
Acheta_domesticus               CCGCGCTGTGTTTACACCCACTCCCGCGCCGCGCGGGCAGGTCGCCGCGAGCCGAGAGGCCCTCGCCGCGCCAGCCCGGAAAGCCACGCCGCCGCACGCAGAAATGCTTGCACAGCTGGG
Consensus                          **  * * *             **  * *   *  ***  * * *   ** * ** *   *   **   *  ***     *  *  * ***          * ** *   * * **

Teleogryllus_occipitalis        GGGTTCGGGGGGTA--TGGGGGTTTCGGGGGTTGGGGGTTCGGGGGGTATGGGGGTTAGGGGAGTTCGGGGGGGGGGGGGGGGTTCGGGGGGGGTATGGGGGTTGGGGGTTGGGGGGGG
Acheta_domesticus               GGAACCCAGGCGTACCTATGTATTCTTACAGTTAACACC-------------------------------GGCCGAATAGTTGTTCAGCTAAGAGATGCGGTCTGCTGCTTCTGAAGAT
Consensus                       **   *  ** ***  *  *  **      ***                                       **  *    *  **** *    *  *** **  **  * **  *  *

Teleogryllus_occipitalis        GGGG-------------------------GTTGGGGGTGGGGTTCGGGGGGTTCGGGGGTATGGGGGTTGGGGGGTTCGGGG------------------------------------
Acheta_domesticus               GGCGATCACGAAGCCCTTTTGTTCGGAACTGTTTGAATGATATTAAAAGACTTTTGGA--ATGAACTATGAAATGTTTACTGTAAAACAGGATCTACTTAAAATTCTTTTTGACCACTC
Consensus                       ** *                         ***  *  **   **    *  **  **   ***     **    ***     *

Teleogryllus_occipitalis        ----------------------------GGTTTCGGGGGGTT--------------------CGGGGGTATGGGGGTTGGGGGGGTTTGGGGGGGTTCGGGGGGGGTTCGGGGGGGTTC
Acheta_domesticus               TCATCCTATTTGTGCAAATGCAAAACCAAGTTTCAGGGAACTTGCTAAATCACTAGCTTACTACGAACGTTTTGCTCATGTAAAAGCGTTGGAACGTTTAGTGTCGACATGGAAATTTTT
Consensus                                                   ***** ***   *                     **   ** * *    **     *  * **   ***  * *  *    **     **

Teleogryllus_occipitalis        ---------------------------------------GGGGGGGGGGGGGGGTTCGGAGGGTATGGGGGTTAGGGGGTT-----CGGGGGGGGGGTTCGGGGGGTTTGGGGTGTTAGG
Acheta_domesticus               ATACAAATACTCAGGCACCGTTAGAAGAACAAATAAATAGAAGTGAAGGGAAAAAGCATGAGGAATGGAATTTTGAATGTTGCCTCGAACGCGGAAATACTGAAACCTTCGGTTTGCAAC
Consensus                                                              *  * *  ***      *    ** ****   ** *   ***          * **   * * *     ** ** *   *

Teleogryllus_occipitalis        GGAGTTCGGGG--GGTATGGGGTAGGGGGAGTTCGTGGGGGTTCGGGGGGGG--------------------------------TGTCGGGGGTGTCGGGGGTGTCG---------
Acheta_domesticus               TGAGTTACACGTTGGTCTAAGGACAGTTGAAGCCTAAGTGACTAAGAGGAAGGACGATCATCAACTGTTCTGAATGAGAATTCACCACTGCCCCAGGTGCCATGTGTATCGAACATTGTC
Consensus                        *****     *  *** *  **  **  * **      * *  *  * **  **                                ** *   **** *  * ** ***

Teleogryllus_occipitalis        --------------GGGGGTGTCGAGGG-------------------------------------------------------------------------------------------
Acheta_domesticus               AGCGTAGAATTGCTGGAGACTTCAAGGGTAGAATAAGAGACGAACCCTTGCACCTTGTGCTTTCATTCCATTCGCATCATTCAACGGGTAATAAAAAGTAGAACGCAAGCAATCAGTCTT
Consensus                                     ** *   ** ****

Teleogryllus_occipitalis        ---------------------------------------------------------------------------------------------------------TGTCGGGGGGTGAGG
Acheta_domesticus               TCGACATTTCAACAGGCAACTGTTCGGTGAATTAATGCATACATACCATTTGATATGCAACAATGATGGTTTTTACAAGACCGTCACTGTTTATTAGTTTTATTTTGTTGGGAACAGAAG
Consensus                                                                                                                  *** ***    ** *

Teleogryllus_occipitalis        GTCGGGG--------------------------------------------------------GTGTCGGGGGGGGTGTCGGGGAGTTG--TCGGGGGGGTGTCGAGGGGTG-------
Acheta_domesticus               GTCGACGTGGAGTCCTTTTACTACATTCTCCTCAGTCATTTGTTGCAAGTCCTCTTCAATCTCTCTATTCAAAGCAAGACTGATGAGTTGTTTCGAATGAATGCAAAACAGTGTAATAGA
Consensus                       ****  *                                                        * *     *       *  ******  ***   *  **   *   ***

Teleogryllus_occipitalis        -----------------------------------TCGGGGGTTCG-------------------------------------------------------------------------
Acheta_domesticus               AGCTGACTTCGATATTCGTTGAATAGTATGACGCTTCGAGAGTTTGTTTTCAAAATATTCTCCCATTATTTCCTCTTTCCAGGGTCATCGCGCTATGTTCCAACAATTCGCAACGTTTTT
Consensus                                                          *** * *** *

Teleogryllus_occipitalis        ------------------------------------------GGGTGGAGGGT----CGGGGGGATC-----------------------------------------------------
Acheta_domesticus               ACATGAGTATACGACAAACAGTACTAATAATAAGAATGGTATAAAAAAGGAGGATTTTACGCGAGGATTTTATTCTTAAAATCTTGTGTTGATTGTTAAACGCAAATAGTGATTCAAATT
Consensus                                                                  ***** *    ** * ****

Teleogryllus_occipitalis        -----------GGGGGGGCGGGGCGGCGAGCGAGCGTCACCTG-----------------------TTCTGGCATGTGGGGAGGGGC---------------------GCTCCGGTTA
Acheta_domesticus               TAAAATCACAACGAATTGTGATAAAACAATCTAACAATAATTGAGGCGTTCGGGTATAAATAATGGTTTCTGACATATTGGAGGAACTCTATTATTAACCTTCAACAACATTCTTTATTA
Consensus                                   *    * *      * * * * *   *  **                       ***** *** * **  *                             ***

Teleogryllus_occipitalis        CGGCGG-----------
Acheta_domesticus               AGGCGTATATCTAGTTC
Consensus                        ****
```

**Tocc.2G0008730.1_exon1**

```
Teleogryllus_occipitalis                 ------------------------------------------------------------------------------------------------------------------------
Acheta_domesticus                        RKVNVQ*FVRMPLRRPTILLLSLAWITSEKNRLFLS*FIFSGDLFKYSNLFFCMLAGTQLLPVVLVFCKVFTGRKQKTTALFRFILLDLIPDYLTLCERDELVHL*FP*WVNYVKVRQQI
Consensus

Teleogryllus_occipitalis                 ------------------------------------------------------------------------------------------------------------------------
Acheta_domesticus                        *TRVCK*RFHSFVKQRRLFLMVIFY*FSCRM*YLNKMLKLCL*AYYLRPFTKVMKQV*SFHH*SESPLWETSPTFAE*RPQNSSNNLHIPRPLYPAAKVRALVGCFHVTLWIIA*PKGAA
Consensus

Teleogryllus_occipitalis                 ------------------------------------------------------------------------------------------------------------------------
Acheta_domesticus                        AVPVEESL*TKLLTQRHFCATDSFWQMDYSQTVNFHSKLQS*SISTISWVARVC*WRLKPLFYLPSQFA*TLSFFICTLDATAQHFFDILADDVKHKLQ*SVNFICCIF*KFISNKFIRK
Consensus

Teleogryllus_occipitalis                 ------------------------------------------------------------------------------------------------------------------------
Acheta_domesticus                        YFWLLSFNISRFTDLTPTLRNLRTKAKNRWKNLRNETILDI*NCTDELCV*FRDNGLSFPRVYAVIFFKQNTAPPPQKCENNHAKNSRRYSICKRYKR*FPSYFRKKNFLSVSVKVNFFH
Consensus

Teleogryllus_occipitalis                 --------------------------------------------------------------------------------------MTFSM-----------------------------
Acheta_domesticus                        TISCICFCFNVFAFYPSKLFLNKIFWCK*PWKLALHLFNAVL*YP*KLKARALKTLKP*TLKCGAKRQPSLKVLQEIRRHFVPRFAFTFSMVSR**PIPSRGDKSRTFFLQLYNYCNYLY
Consensus                                                                                                                       ****

Teleogryllus_occipitalis                 ------------------------------------------------------------------------------------------------------------------------
Acheta_domesticus                        NFVINLKNSYKKLIN*RNITY*ESSKR*RIEKK*H**CYISFTVHS*LII*I*ISGNVFPYDSIIFLRALKPSLELPNDIFYLHYMDKQIVKKCGNETKLS*FENVLP*NFENCVQHFVS
Consensus

Teleogryllus_occipitalis                 ------------------------------------------------------------------------------------------------------------------------
Acheta_domesticus                        KA**QHG*QLHQLFMRHQTPKMATDSRRDTVD*GGGCQGQ*NLDPWNDTEASPL*IAKTT*LSHLIVNLTPSRLFHLKGCFKSTSPWQIPAAFSTFQ*HWPQHHRAKQQWLRTPYARAQQ
Consensus

Teleogryllus_occipitalis                 ------------------------------------------------------------------------------------------------------------------------
Acheta_domesticus                        GMGRRVGRRVGRRVGAARGGGGEGPQAHVSLDDRRRQGKWRGQWRLKGTARPRCVTQSTTVRVIQRGWRNAAAPGPRCVYTHSRAARAGRREPRGPRRASPESHAAARRNACTAGGTQAY
Consensus

Teleogryllus_occipitalis                 ------------------------------------------------------------------------------------------------------------------------
Acheta_domesticus                        LCILTVNTGRIVVQLRDAVCCF*RWRSRSPFVRNCFE*Y*KTFGMNYEMFTVKQDLLKILFDHSHPICANAKPSFRELAKSLAYYERFAHVKALERLVSTWKFLYKYSGTVRRTNK*K*R
Consensus

Teleogryllus_occipitalis                 ------------------------------------------------------------------------------------------------------------------------
Acheta_domesticus                        EKA*GMEF*MLPRTRKY*NLRFATELHVGLRTVEA*VTKRKDDHQLF*MRIHHCPRCHVYRTLSA*NCWRLQG*NKRRTLAPCAFIPFASFNG**KVERKQSVFRHFNRQLFGELMHTYH
Consensus

Teleogryllus_occipitalis                 ------------------------------------------------------------------------------------------------------------------------
Acheta_domesticus                        LICNNDGFYKTVTVY*FYFVGNRRSTWSPFTTFSSVICCKSSSISLFKARLMSCFE*MQNSVIEADFDIR*IV*RFESLFSKYSPIISSFQGHRAMFQQFATFLHEYTTNSTNNKNGIKK
Consensus

Teleogryllus_occipitalis                 ---------------------------------------------------------------------
Acheta_domesticus                        EDFTRGFYS*NLVLIVKRK**FKFKITTNGDKTI*Q*LRRSGINNGF*HIGGTLLLTFNNILY*GVYLV
Consensus
```

**Tocc.2G0008730.1_exon2**

```
Teleogryllus_occipitalis                 ------------------------------------------------------------------------------------------------------------------------
Acheta_domesticus                        ELDIRLNKECC*RLIIEFLQYVRNHYLYPNASIIVRLFYHNSL*F*I*ITICV*QSTQDFKNKILA*NPPFLYHSYY*YCLSYTHVKTLRIVGT*RDDPGKRK*WENILKTNSRSVILFN
Consensus

Teleogryllus_occipitalis                 ------------------------------------------------------------------------------------------------------------------------
Acheta_domesticus                        EYRSQLLLHCFAFIRNNSSVLL*IERLKRTCNK*LRRM**KDSTSTFCSQQNKTNKQ*RSCKNHHCCISNGMYALIHRTVAC*NVERLIACVLLFITR*MMRME*KHKVQGFVSYSTLEV
Consensus

Teleogryllus_occipitalis                 ------------GSSGGG------------------------------------------------------------------------------------------------------
Acheta_domesticus                        SSNSTLTMFDTHGTWGSGEFSFRTVDDRPSS*SLRLQLSLDQRVTQLQTEGFSISAFEATFKIPFLMLFPFTSIYLFF*RCLSICIKISMSTLNVPTLLHEQNVRSKLVI*QVP*NLVLH
Consensus                                            *  * *

Teleogryllus_occipitalis                 ----------------------------------------GGV-----------------------------------------RGGYGGW---GLGGGG--------------------
Acheta_domesticus                        LHK*DESGQKEF*VDPVLQ*TFHSSFQKSFNIIQNSSEQKGFVIAIFRSSRPHLLAEQLFGRC*L*EYIGTPGFPQLCKHFCVRRRGFPGWRGEGLSARGDLPARRGSGCKHSAAPAPPR
Consensus                                                                        * *                                         * *  **   **   *

Teleogryllus_occipitalis                 --------------------------------------------------------------------------------LGVGFGGFGGMGVG--------------------------
Acheta_domesticus                        CATRAV*RELLWIASRNAVVRSLSAAIALATSPASADHQATRVPVAPRRPHHAPPPRAAPRAAQLCAPCPVGLSRMAFAAIVVSLDGAAASAIGM*RTQLVFARD*CS*NNP*GEKDVKV
Consensus                                                                                                                          *   *      *

Teleogryllus_occipitalis                 ---------------------------GFG-------------------------------------------------GFRGVR-----------------------------------
Acheta_domesticus                        LN*RLNGTARLS*QFTVGKPPYRSMGPGFTDPDSLHLNRPCHVCCLLPSSAFDDA*TVGVTATHAAIKLSTRSVEHSSQSFRGARFQTKTI*FHSRIF*QFAYPYSVNKIYHLAILKMVS
Consensus                                                           **                                                   *** *

Teleogryllus_occipitalis                 ----------------------------------------------------------------------------------GYGGW---------------------------------
Acheta_domesticus                        TLSKR*LNRTGKRSH*FIFILSTTNVL*NLCNITSAIFFQFSIFSNSLNM*YFFNLLVFCMNSSNLLQNYINNYNNYIIVKKKYGSYPHVKVLVITVKPWRT*TRTWERNGVLFPAILLR
Consensus                                                                                                                           **

Teleogryllus_occipitalis                 -----------------------------------------------------------------------------------------------------------GGLGGFGGGS---
Acheta_domesticus                        MVDVLLRISRFKVSRFSRLVPSAFMGITKQH*ISATLISMAIYTKIFC*ETILRDKKQKH*NRNKCTKLYERN*LSRRRIRSFFF*NSLEIIFYISCKLNISANSSHGYSHIFGGGALYF
Consensus                                                                                                                                                  *    ****

Teleogryllus_occipitalis                 -----------------------------------------------GGFGGGG------------------------------------------------------------------
Acheta_domesticus                        V*KKLPHKHAEMRDHCHEIKHRVHLCNFKYPKWSRS*DFSICSSLLFSDFAASG*DL*TEKC*RIITKNIYE*IYWK*TFKICNI*SLHFIVIYV*HHQQGCQRNAELLRLMCK*KMKVF
Consensus                                                                                          *   *

Teleogryllus_occipitalis                 ----------------------------GFGGYGG*------------------------------------------------------------------------------------
Acheta_domesticus                        MQIVTEDKTKV*VSISILELPKILC*SISFEASSGS*RSASSPFARSCQSRRNGVA*AA*STSSLLLGQLPLPSVMLLSTTLHENSQLGRVLWQQDIAVGECANC*NYSVVSTQRKLAKS
Consensus                                                      *    *

Teleogryllus_occipitalis                 ------------------------------------------------------------------------------------------------------------------------
Acheta_domesticus                        PITGIQTSDGSFKLVSSPW*RV*GNMLINIILTFC*DITFYKKINKI*PLKTIYVVLQNCEISTYKLVFKFVV*L*HN*PIRETINARVHLSRKVSNSLE*DLRV*NEIKQSSFVFFL*R
Consensus

Teleogryllus_occipitalis                 ----------------------------------------------------------------------
Acheta_domesticus                        LCKTLVQLVAIVFPLTYKRINLNI*TSLLRK*IMTGRVDFSLTLSMLMKGAR*SVA*VAFLRIIERLLSW
Consensus
```

**Tocc.2G0013240.1**

```
Teleogryllus_occipitalis    TAACATTCCAATGACTAAGATAAAATTCAGTAGGCATCATTTGGATGCTGAAGAATGGCAGCAATTGCAAATCAGCAAGCACGCCATCTTATCTCTACTTTCTTTCTTGGAACGAAAACA
Acheta_domesticus           TAACATTCCAATGACTAAGATAAAATTCAGTAGGCATCATTTGGATGCTGAAGAATGGCAGCAATTGCAAATCAGCAAGCACGCCATCTTATCTCTACTTTCTTTCTTGGAACGAAAACA
Gryllus_bimaculatus         TAACATTCCAATGACTAAGATAAAATTCACTAGGCATCATTTGGATGCTGAAGAATGGCAGCAATTGCAAGTCAGCCAGCACGCCACCTTATCTCTACTTTCTTTCTTGGAACGAAAACA
Consensus                   ***************************** **************************************** ***** ********* *********************************

Teleogryllus_occipitalis    CTTTGTTCTTGCATGGCGTTTCAATGAGAACAGAATAATCACACTAACAACTTCGGCGGAACCCACGTAACCATGTCGCGCAGTCGAAGTACTTAGGAAATCATTAATCATTCCCAAGTT
Acheta_domesticus           CTTTGTTCTTGCATGACGTTTCAATGAGAACAGAATAATCACCCTAACAACTTCAGCGGAACCCACGTAACCATGTCGCGCAGTCCAAGTACTTAGGAAATCATTAATCATTCCCAAGTT
Gryllus_bimaculatus         CTTTGTTCTTGCATGACGTTTCAATGAGAACAGAATAATCACCCTAACAACTTCAGCGGAACCCACGTAAGCATGTCGCGCAGTCAAAGTACTTAGGAAATCGTTAATCATTCCCAAGTT
Consensus                   *************** ************************** *********** *************** ************** **************** *****************

Teleogryllus_occipitalis    CGCATTTGTTCTGTCCATCGATTTGTTCTTCTGTTAAGCGAGCGTGGCATTGATAGCTCGATGCGGCGTTTTAAACTTGCTGTAGAAACTTGCCCTCTCAGATAGGGAAATGCACGTCGA
Acheta_domesticus           TGCATTTGTTCTGTCCATCGATTCGTTCTTCTGTTAAGTGAGCGTGGCATTGATAGCTCGATGCGGCGTTTTAAACTTGCTGCTCGAACTTGTCCTCTCGCGTAGGGAAATGCACGTCCA
Gryllus_bimaculatus         TGCATTTGTTCTGTCCATCGATTCGTTCTTCTGTTAAGCGAGCGTGGCATTGATAGCTCGATGCGGCGTTTTAAACTTGCTGCGAGAACTTGTCCTTTCGCGTAGGAAAATGCACGTCGA
Consensus                    ********************** ************** *******************************************    ****** *** **   **** *********** *

Teleogryllus_occipitalis    CTCCGGAACGTATATTCA-AAAAAGCTTGCCTAGCAGCTCAGTATTTCCTTTCATAAACGTCAACAAGACTCCCGAGGGGCCGTACAAGTTCGCCAGCGCGCACATTAACTCGGACTATC
Acheta_domesticus           GTCCGGAACGTATATTCA-AAAAAGCTTGCCTAGGAGCTCAGTATTTCCTTTCATAAACGTCAACAAGACTCCCGAGGGCCCGTACAAGTACGCCAGCCCTCACATTAACTCGGACTATC
Gryllus_bimaculatus         GTCCGGAACGTATATTCAGAAAAGGCTCGCCTAGGAGCTCAGTATTTCCTTTCATAAACGTCAACAAGACTCCCGTGGGCCCGTACAAGTTCGCCAGCGCTCACATTAACTCGGACTATC
Consensus                    ***************** **** *** ****** **************************************** *** ********** ******* * *******************

Teleogryllus_occipitalis    AACTTATTCATCGCCGACGAGCTCTCTTCCGCGAGCTTCTAATAGATTGAGAACCAATTTTCACGGGAGACACGTTAATCGAAAGTTTACGACGCGACGCGGCGCCGGTGAAGGTGCCGG
Acheta_domesticus           AACTTATTCATCGCCGACGAGCTCTCTTCCGCGAGCTTCTAATAGATTGAGAACCAATTTTCACGGGAGACACGTTAATCGAAAGTTTACGACGCGACGCGGCGCCAGTGAAGGTGCCGG
Gryllus_bimaculatus         AACTTATTCATCGCCGACGAGCTCTCTTCCGCGAGCTTCTAATAGATTGAGAACCAATTTTCACGGGAGACACGTTAATCGAAAGTTTACGACGCGACGCGGCGCTGGCGTAGGTGCCGG
Consensus                   *********************************************************************************************************  * * *********

Teleogryllus_occipitalis    CATCTCGGCGGCAGCAGCGGC----GCAAAGAGGTGGTGGAGGACACGGCGGAAGTGGCAGTGGCGGCGGAGGCGGCGGCGGCAGCGGCGGA-----GGCGGCGGAGGAGGCGGCGGAGC
Acheta_domesticus           CATCTCGGCGGCGACGAAGGGGGGTGGC--GGAGGCGGTGGAGGCTGCGGAGGAGGTGGCACAGGCGAAGGAGGCGGCGGAGATGGCAGC--------GGCGGTGGAGG-----------
Gryllus_bimaculatus         CATCTCGGAAGTGGCGGTGGCGGCGGCAGGGGAGCAGAGGAGAAGGCTGCGGAGGCGGCAACGGCGGAGGAGGCGGCGGGCACAGTGGTGGGGTGGCGGCGCTGGGGGGTGCTGAGGAGG
Consensus                   ********  *   *   **      **   *  *  * ****    * * *** * ****  ****  ***********     *  *         ****  ** **

Teleogryllus_occipitalis    CGGTTGGATCAGTAATGACGGCG--------GGGGTGCGAGGCGGCGGGTGCGGCGG--------------------------------GTTGGCGGGGGCAGCGGGGCGGCGGCGGCGG
Acheta_domesticus           ------------------------------CGGTGAGCGGGGAGACGGCAGGGACGG-AGGAGTGAAATTAGTGGTGGAGGCTGGTGCAGGCAGGAAGAGGGAGGAACAGGTGGTGGAGG
Gryllus_bimaculatus         AGGCTGTGGAGGCGGCAACGGCGGTGGACGCGGAGGACGGGGTGGCGGCTGTGGCGGGGGGTGTGAAAGTGGGGAAGGAGGTTGGTGCAGGCAAGAAAAGAGAGGAAATGGTGGTGGCGG
Consensus                                                  ** *  ** ** * ***  * * ***                                *  ***  *   ** * **    ** ** ** **

Teleogryllus_occipitalis    ---------------------------GGCTGCGGGGGCTGCGG-------------------------------GGGCTCCGGGGGGCGGCGGGGGCGGCGGGGGCGGCGGGGGCGGCG
Acheta_domesticus           CGGCGGCAATG----------------GGGCGAGGGGGCGGTGGCA------------------------------------------GGGGCGGGCTGGCGGAGAGGTGGAGCCGGCG
Gryllus_bimaculatus         CAGAGAGAGTGCCGGCATTTCGGCAGTGGCTGTGGTGGCGGCAGCGTGCGTGCTGAGGAGAGGCTGCAGTCTTGCGGGCACCAATAGTGGGAGGTGGCGGGCAGGTTGGCGACGGTGGCG
Consensus                                              **   * ** *** *  *                                            ** *  *** *   *   ** *  *  ****

Teleogryllus_occipitalis    GGGGCGGCGGGGGCGGCGGGG---------------------------------------------------------------------------------------------------
Acheta_domesticus           AGAACAATGGAGGGGATGATG--TCTTTGCAGGCATCGGCGCTCGTAGTTGACGCTCTTCCCGCCTCGGGCTCCTGGCTCTGATATCGCACTCTCATTGAGTGATCTTTACTGTCTTCAA
Gryllus_bimaculatus         GCGGTGAGAGAGGCGGTGAGGTTTCGTTGCAGGCAGGAAAAAGGGGAAGTTG--------------------------------------------------------------------
Consensus                            * ** *  *  *

Teleogryllus_occipitalis    ----GCGGCGGGGGCGGCGGGGGCGGGGGGGCCG----------------------GCGGGGGCGGGGGGGCGGCGGGGGCGATGGGGGCGACGGGGGCG--------------------
Acheta_domesticus           GTGCGCATCGCAACCGTATTTGAAGTCACGCCTGAATTACAG------------ATATCAGAGAATGAACTGGGAAGAAATTATTGAGAAAATGTGTTAT------------TTGTTGTG
Gryllus_bimaculatus         --ATGGGGCGGGGGAGAAGGAGGAGGGGCGGTGGAGTGAGAGCGGGATGGAAGTTTATGCAGGCAGGAAGAGGGAGGAAGTGGTGGTGGCGGCGGAGAAGGAAGTAAGAGTGACGAAGTG
Consensus                       *   **     *     *  *    *   *                            *   *     **  *      * * *     *

Teleogryllus_occipitalis    ---------------------------------------------------------ACGGGGGCGACGGGGGCGACGGGGGCGACGG--------------------------------
Acheta_domesticus           TTACAGTATATGAATCTCGATTGATCACACTTAATTTTCAAAGGCTTGGCAAGATATCAGAGGAAGAATGAAGTGTCTCAATCATTAG--AAAATCATTACATAAGGAGTAAATGTAGTT
Gryllus_bimaculatus         ATAGTGCGGGTGAAGGTTGGT------------------------------GCAGGCAGGAAGATGGAGGAAGTGGTGGAGGCGGCGGCGCAGATTGCGTGACGCGGCGGCAATGGGGC-
Consensus                                                                     *  *  *   *  * *      *    *

Teleogryllus_occipitalis    ----------------GGGCGACGGGGGCGACGGG-----GGCGACGGGGGCGACGGGGGCGACG-------------------------------------------------------
Acheta_domesticus           GTTTTTCTTTCAGAGCAGGACATTTAGCTTTTAGA---ACGTTTTTTAAGTCAGGGAATATGATATCATTCCCACGACATAATCAAAGTATTCAGAGATTTTTATCATTACAGTTCGTAC
Gryllus_bimaculatus         --------------ACGGGCGGTGGTGGGGGTGGACAGGCGGCGGCGAGAGCAAAGGAGAGGATGTCTTT-----------------GCAGGCGGTGGCGCTTGTAGTTAACGCTCTTCC
Consensus                                     **       *      *      *          *   *     **

Teleogryllus_occipitalis    ------------------------------------------------------------------------------------------------------------------------
Acheta_domesticus           ATTTTTTATTACTTGATTATTACCCGTCTTATTCGTCTCATTTTACTTCCTCGCATACTACTTACGTAGTAGAGGGAAAGTTATGTAATACTGCAGCGAAAGAGCGTATTTTCAGCAGAA
Gryllus_bimaculatus         CTCTTC------------------------------------------------------------------------------------------------------------------
Consensus

Teleogryllus_occipitalis    -----------------------------
Acheta_domesticus           AAAGTATTAAAGGTTCTTGATACC-----
Gryllus_bimaculatus         ----------AGGCTCCTGGCACTGATAT
Consensus
```

**Tocc.2G0013240.1_exon1**

```
Teleogryllus_occipitalis        ------------------------------------------------------------------------------------------------------------------------
Acheta_domesticus               -----Y*NS*ERIRIMLYTEKNNLKNISFQNSFILLVYSNGAKTHMIFYFS------------NHNVAER-----MPVFESCRLTENKMTNYSNMR*TSFFP*IS-HTMK*GTHCQTHAR
Gryllus_bimaculatus             ERTTEIVNNIFQ*RM---TQRTNI*KI-FR--FTTLSYLATVKS*MCYFLFLNRMLLK*FLSLNHVVLRKIK*KIIPI-----------------RNRLHFVRISPHTMK*GTHC*THAR
Consensus

Teleogryllus_occipitalis        ------------------------------------------------------------------------------------------------------------------------
Acheta_domesticus               KIMLCSFLP*ALNRV*NKVHKFTYFSILQK*QTT*WHF*VVFQY*HSND*DKIQ*ASFGC*RMAAIANQQARHLISTFFLGTKTLCSCMTFQ*EQNNHPNNFSGTHVTMSRSPST*EIIN
Gryllus_bimaculatus             KII*CSFLP*ALNRV*NKVDKFTYFSILQK*QST*WHF*AVFQY*HSND*DKIH*ASFGC*RMAAIASQPARHLISTFFLGTKTLCSCMTFQ*EQNNHPNNFSGTHVSMSRSQST*EIVN
Consensus

Teleogryllus_occipitalis        ------------------------MRRFKLA-----------------------------------------------------------------------------------------
Acheta_domesticus               HSQVCICSVHRFVLLLSERGIDSSMRRFKLAARTCPLA*GNARPVRNVYSKKLA*ELSISFHKRQQDSRGPVQVRQPSH*LGLS---------------------TYSSPT--SSLPRAS
Gryllus_bimaculatus             HSQVCICSVHRFVLLLSERGIDSSMRRFKLAARTCPFA*ENARRVRNVYSEK--ARLGAQYFLS*TSTRLP-WARTSSPALTLTRTINLFIADELSSASF**IENQFSRETR*SKVYDAT
Consensus                                               *******

Teleogryllus_occipitalis        ------------------------------------------------------------------------------------------------------------------------
Acheta_domesticus               NRLRTNFHGRHVNRKFTTRRGASEGAGISAATKGVAEAVEA------------------------------------------------------------------AEEVAQ-------
Gryllus_bimaculatus             RRWR---RCRHLGSGGGGGRGAEEKAAEAATAEEAAGTVVGWRRWGVLRRRLWRRQRRWTRRTGWRLWRGV*KWGRRLVQARKERKWWWRQRECRHFGSGCGGGSVRAEERLQSCGHQ*W
Consensus

Teleogryllus_occipitalis        ------------------------------------------------------------------------------------------------------------------------
Acheta_domesticus               --AKEAAEMAAAVEAV-------SGET--AGTEE*N*W--------------------------------WRLVQAGRGRNRWWRRRQWGEGAVAGAG--WRRGGAGENNGGDDVFAGIG
Gryllus_bimaculatus             EVAGRLATVAAVREAVRFRCRQEKGEVDGAGEKEEGRWSESGMEVYAGRKREEVVVAAEKEVRVTK**CG*RLVQAGRWRKWWRRRRRLRDAAAMGHGRWWGWTGGGESK-GEDVFAGGG
Consensus

Teleogryllus_occipitalis        ------------------------------------------------------------------------------------------------------------------------
Acheta_domesticus               ARS*RSSRLGLLALISHSH*VIFTVFKCASQPYLKSRLNYRYQRMNWEEIIEKMCYLLCYSI*ISIDHT*FSKAWQDIRGRMKCLNH*KIIT*GVNVVVFLSEQDI*LLERFLSQGI*YH
Gryllus_bimaculatus             ACS*RSSLFRLLALI---------------------------------------------------------------------------------------------------------
Consensus

Teleogryllus_occipitalis        -----------------------------------------------------------------
Acheta_domesticus               SHDIIKVFRDFYHYSSYIFYYLIITRLIRLILLPRILLT**RESYVILQRKSVFSAEKVLKVLDT
Gryllus_bimaculatus             -----------------------------------------------------------------
Consensus
```

**Tocc.2G0013240.1_exon2**

```
Teleogryllus_occipitalis    ------------------------------------------------------------------------------------------------------------------------
Acheta_domesticus           TTEIVE---NAFE*CFTRRRII*KIFRSKILSYCWFIVTVQKHI*YFIFLIIMLLKECLSLNHAVL------QKIK*QIIQICDRLHFSRKSHIQ*NKALIARLTREK*CYVLFFPKL*I
Gryllus_bimaculatus         *NELLK*LTTYFNNA*HREQTFEKYFVSQL----FHI*QR*NHECAIFFF*I----EC-C*NNSYL*IMSSYAK*NEKLFQYVIDFILSVYLHIQ*NKALIARLTREK*YNVLFFPKL*I
Consensus                   

Teleogryllus_occipitalis    -------------------------------------------------W----------------------------------------------------------------------
Acheta_domesticus           AFETKCTNLHISPSYRNSRLHNGISELFSSTNIPMTKIKFSRHHLDAEEWQQLQISKHAILSLLSFLERKHFVLA*RFNENRIITLTTSAEPT*PCRAVQVLRKSLIIPKFAFVLSIDSF
Gryllus_bimaculatus         AFETKWTNLHISPSYKNSSLHNGTSERFSSTNIPMTKIKFTRHHLDAEEWQQLQVSQHATLSLLSFLERKHFVLA*RFNENRIITLTTSAEPT*ACRAVKVLRKSLIIPKFAFVLSIDSF
Consensus                                                                    *

Teleogryllus_occipitalis    ------------------------------------------------------------------------------------------------------------------------
Acheta_domesticus           FC*VSVALIARCGVLNLLLELVLSRREMHVQSGTYIQKSLPRSSVFPFINVNKTPEGPYK------------YASPHINSDYQLIHRRRALFRELLID*EPIFTGDTLIESLRRDA----
Gryllus_bimaculatus         FC*ASVALIARCGVLNLLRELVLSRRKMHVESGTYIQKRLA*ELSISFHKRQQDSRGPVQVRQRSH*LGLSTYSSP-TSSLPRASNRLRTNFHGRHVNRK--FT-------TRRGAGVGA
Consensus                   

Teleogryllus_occipitalis    --------------RRRRRRQRRR---RRR------------------------------------------------------------------------------------------
Acheta_domesticus           --APVKVPAS----RRRRRGWRRRWRLRRR-----------------------------------------WHRRRRR------------------------------------------
Gryllus_bimaculatus         GISEVAVAAAGEQRRRLRRRQRRR---RRRAQWWGGGAGGC*GGGCGGGNGGGRGGRGGGCGGGCESGEGGWCRQEKRGNGGGGRESAGISAVAVVAAACVLRRGCSLAGTNSGRWRAGW
Consensus                                 ** **  ***   ***

Teleogryllus_occipitalis    --------------------------------------------------RRRSRLDQ*-------------------------------------------------------------
Acheta_domesticus           RRWQR--------------------RWRR*AG------------------RRQGRRSEIS---GGGWCRQEEGGTGGGGGGNG-ARGRW-QGRAGGEVEPARTMEGMMSLQASALVVDAL
Gryllus_bimaculatus         RRWRR*ERR*GFVAGRKKGKLMGRGRRRRGGGVRAGWKFMQAGRGRKWWWRRRRK*E*RSDSAGEGWCRQEDGGSGGGGGADCVTRRQWGTGGGGGGQAAARAKE-RMSLQAVALVVNAL
Consensus                                                                     **

Teleogryllus_occipitalis    ------------------------------------------------------------------------------------------------------------------------
Acheta_domesticus           PASGSWL*YRTLIE*SLLSSSAHRNRI*SHA*ITDIRE*TGKKLLRKCVICCVTVYESRLITLNFQRLGKISEEE*SVSIIRKSLHKE*M*LFFFQSRTFSF*NVF*VREYDIIPTT*SK
Gryllus_bimaculatus         PSSGSWH*Y---------------------------------------------------------------------------------------------------------------
Consensus                   

Teleogryllus_occipitalis    ----------------------------------------------------------
Acheta_domesticus           YSEIFIITVRTFFIT*LLPVLFVSFYFLAYYLRSRGKVM*YCSERAYFQQKKY*RFLI
Gryllus_bimaculatus         ----------------------------------------------------------
Consensus                   
```

**Tocc.2G0020270.1**

```
Teleogryllus_occipitalis      TACATATGTTGGATATTTCCTCGTTGTAAATTTCCTTGACTAAAGCGCATCATCTGTATATTTTTTGAAGA-TTGTGTTGCCTTCTGTACGTCATTGTCAGTATA---GTGACATCTAAT
Gryllus_longicercus           CACAAATGCTAGATATTTCCTCGTTGTACATTTCCTTAAGCAAAATGCATTAGCTGTATATTTTTTGAAGATTTTTTTTGCCTTCTGTACTTCATTGTCAGTCAGTATGTGTCATCCGAT
Consensus                      *** *** * ***************** ******** *  ***  **** * ****************** ** * ************* ***********      *** ****  **

Teleogryllus_occipitalis      TGCCACA----------------------------------ATTTACTTAATCTTCATGAATGTTTCATG-TGTACGAAACTTGTTCGAATAAATAATCAAAAGTCAAATCTCTGAGCC
Gryllus_longicercus           AATTACATTGTTATGGACAATTTTTGTCACCTGTGTTCTCCCTTTTACATTATCTTCATGAATGTTTTATGACGAACAAAACTTGTTCCAATGAATAATCAAATGCCAAATCTCTGAAAC
Consensus                         ***                                   ***** * **************** ***  * ** ********** *** ********** * ***********  *

Teleogryllus_occipitalis      ACTCGAAGGACAATCCCAAGAAACATAATTCGCCTTGACTTAAAAAATATTCCTTATATTACATAAATATTGCATTTAT-GTACATGTGCTT------------GTAGTAGAAATGAGG
Gryllus_longicercus           TC---ACGGACAATTCCAAGAAACAT--TTCGTTTTCACTTAAAAAATATTCCATGCATTATATAATTATTGCACGCATAGTACACGTACATAGTAGTACAATACATAGTAGAAATAAGC
Consensus                      *   * ******* ***********  ****  ** **************** *  **** **** *******   ** ***** ** * *              ********** **

Teleogryllus_occipitalis      GCGCACTACAAAT--------GAAATTTACTTTTTTATATTCCATTGGAAATCAATGTTCTGTATTCACTTATTACTAGGGAATTTTTTTATATGGATAAACGAA---------AGCAAT
Gryllus_longicercus           GC-CAATACAAATTTAAAAAAAAAAATGATTTTCCAAT------TTAAAAAGCAATGTTTTGTATTCACTTATTATTAGGGAATACATTTATGTTTAAAAACGAATTATTTACTGGTAAA
Consensus                     ** ** *******         *** * * ***   **      **  *** ******* *************** ********   ***** *  * *******          * **

Teleogryllus_occipitalis      TACATGTAAAAGCAATTACAAGATACCTATAGCCAGACTTTTTAATTTTTTTTATTTTGAAATTTTTTCGAAAGAATTAAATCTATTTTTACTGTACATATCTTTACGTATTTTAGAATA
Gryllus_longicercus           TACATATTTGTACAGAAAAAGGACATCTGGAGTTTA----------TTTATTTATTTGAAAGGTATTCTGAATGTATTAAATCCATCTTTACTGTACATATTTT----------------
Consensus                     ***** *     **   * * ** * **  **              *** *******  **  * **  *** * ******** ** ************** **

Teleogryllus_occipitalis      AGTACATATTTTACCTATTAAGACAC-TTTTTCACATTTCTCAGCATGGCAGTATTTTGAATGACAAAAAGTAATATTTATACACTATCAACATACGAGTTTGCGATAGACGTTCTTGGA
Gryllus_longicercus           --TAAATATTTTACCTATTATAACACATTTTTCATTTTTCTCAGCTTGACAGTATTTTGAATGACGAAAAGAAATATTTATACAGTATC-ACATACGAGTTTGCAGTACA-GTTCTTGGA
Consensus                       ** ***************  **** ******* ********* ** **************** ***** ************ **** **************  ** * ********

Teleogryllus_occipitalis      ATCGCCTCTCTGAAGC---------------------------------------------------------------------------GTGAAGAA------------
Gryllus_longicercus           ATCGCCTATCTGAAGCGAAGTTCTGCTGCGCTCCTGTGACCTCACCTTCACCGTGAATTGTCAAACTTATCACAATCTTCTGGGAAATTACAAACTCGGAGCGAAGAATGTTGCCGAAAC
Consensus                     ******* ********                                                                           * ******

Teleogryllus_occipitalis      GAAAGAAATTGTGAC-------------------------------------------------------ATTCTTAACAGAGCATTGCCTTTCCACACTCCCTGCTCCCCGCCGACACACAG
Gryllus_longicercus           AGAAGTAATTGCAACATTATGTTTGTTCATGTTTCTTAGAGGCCATGAGAATCTAAATTTGACAGCGATTCTTAACAGAGCATTGTCTTACCGCCCAACCAGCACCCTGCCGAGAAACTG
Consensus                       *** *****  **                                                      ****************** *** ** * *  ** ** *** ***** * ** *

Teleogryllus_occipitalis      AACTCTCTCTACCACGAACGCAGAAATGGAAGTTGTTCTGTTCATGAAATATCGGTCACTTCTGTAAAATGTGTAACTCCTTTAGATTGTAATGGAAGCAAGATGTCGATATGTGGTAA
Gryllus_longicercus           AACTCTCTCTGCCAGGAACGCTGAAATGGGAGTCCTTCTGTTCATAAAATATCGGTCATTCTTGCAAAATGTGTAACTCCTTTGGATTTTTTTGCGAGCAATGTATCGATATGTGGTAA
Consensus                     ********** *** ****** ******* ***  *********** ************ *  ** ******************* ****  *  ***** * *** * **************

Teleogryllus_occipitalis      TTGTTTCAATAAATTTATTTTAATATCAATAAAATGTTGCACGAGGGAT------TTTTGTACCAGATGGTGTTCAAGCTAAGAAAATAATGCCGGAAGTCATTTCGTCAATTTTTTGCA
Gryllus_longicercus           TTGTTTCAATAAATTAATTTCAATAACCATAAATTGTTCATTAGGAGCTGTTTTCTTTTCTATCAGA-AGTTCTCAAGCTAACAAAATAATGCCGG-----------------------
Consensus                     *************** **** **** * ***** ****      * * *      **** ** ****  **  *********  ************

Teleogryllus_occipitalis      GAGGATTTAAATTTGTAGGGTCCTCAATTTTTGGTAGCTTTTGACGGTCGTCATTTTTGGGAGCTTTTGTGGGGTCATCATTTTTGGGATTTTACAAATTTCGAATTTCTTGAGAGATTT
Gryllus_longicercus           --------------------------------------------TGCCAT---------------------------------------------------TTGAGAGTTTT
Consensus                                                                   * ***                                                   ******* ***

Teleogryllus_occipitalis      TAAAATCCATATTGAATTTTAATTTAGAATAAACTTTGTTGCGTCACATTTTCTGTATCCCGTCGTCATTGCTGACACTTTCAAAGTATAAAAAGAAAACAAATAAGAACAGAATGAGAA
Gryllus_longicercus           TAAAAGCCAAATTCAATTTTAATTTATTTGATGCACTATTGCG--ACATTTTCTGTATGCAGACATAATTGCTAACACTTCCAGAGCAGAAAAAAATAGT--------ACTGAATGAGAT
Consensus                     ***** *** *** ************    *  *  * *****  ************ * * * ****** ****** ** * * ***** * *          ** ********

Teleogryllus_occipitalis      AGAGGTAAAGAAGGGAAGAAGTGCAAGTATAAGAAGAGAGCATGGGGAGGA--------------------------------AGAAGTAGAATGTAGGAACGAAAGAAAGGAGATGTA
Gryllus_longicercus           AGAAGTAAAGAGAACAAGAAGTAAGAGTATAAGAGGAGGAAATGCGAAGGAAGAAGAAATATGAGAAAAAGAAAAGATATGAAGAGAAGTAAAATGTAGGGAGGACGGAAAGAAGATATA
Consensus                     *** *******    *******   ********* ***   *** * ****                                   ******* ******** * **  ***** **** **

Teleogryllus_occipitalis      AAC-------------------------------------------------------------------------------------------------------AA
Gryllus_longicercus           AACATTTAAAGAGAAGAAAGCGAGAGGGAAGAAGAAATGAATAGAGAGAAGAACAGAAAAGTGCAGAAGCATGAAGAAGAGGAAAAATGTAATGCAAAATGGAAGAAAAAGCAGAAGAAG
Consensus                     ***                                                                                                       *

Teleogryllus_occipitalis      AAAGAGAAGAAGAAGAAAAAGAAGGCGA--------------------------CAGGGAAAGAAGAGAGTGAATAAAGAGAAGGAATGGAAGACGACGGATAAAGAAAA---------
Gryllus_longicercus           GAGAAGCAGAAGAAGAAAGAGAAGGTGAAGCAGAAGAAGAAAGAAGGTGAAGCAGAAGAAGAAAGAGAAGGTGAAGCAGAAGAAGAAAGAGAAGGCGAAGCAGAAGAAGAAAGAGAAGGC
Consensus                      *  ** *********** ****** **                           **   **  ***  *****  *  ***** **  **** *** * * **  * **

Teleogryllus_occipitalis      ----------------------------AAGAGAAGAAAAAGAAGACAAAATGGAAGAAGCAGTAGCAGCAGCAGCAGAAGAAGAAGAAGAAGCGGCAGCAGAA--------------
Gryllus_longicercus           GAAGCAGAAGAAGAAAGAGAAGGCGAAGCAAAAGAAGAAAGAGAAGGCGAAGCAGAAGAAGAAAGAGAAGGTGAAGCAGAAGAAGAAAGAGAAGGTGAAGCAGAAGAAGAAAGAGAAGGT
Consensus                                                  ** ******** ***** * **    ******* *   ** **  * *************  *****  * *******

Teleogryllus_occipitalis      ------------------------------------------------------------------------------------------------------------------------
Gryllus_longicercus           GAAGCAGAAGAAGAAAGAGAAGGTGAAGCAGAAGAAGAAAGAGAAGGTGAAGCAGAAGAAGAAAGAGAAGGTGAAGCAGAAGAAGAAAGAGAAGGTGAAGCAGAAGAAGAAAGAGAAGGT
Consensus

Teleogryllus_occipitalis      ------------------------------------------------------------------------------------------------------------------------
Gryllus_longicercus           GAAGCAGAAGAAGAAAGAGAAGGTGAAGCAGAAGAAGAAAGAGAAGGTGAAGCAGAAGAAGAAAGAGAAGGTGAAGCAGAAGAAGAAAGAGAAGGTGAAGCAGAAGAAGAAAGAGAAGGT
Consensus

Teleogryllus_occipitalis      ------------------------------------------------------------------------------------------------------------------------
Gryllus_longicercus           GAAGCAGAAGAAGAAAGAGAAGGTGAAGCAGAAGAAGAAAGAGAAGGTGAAGCAGAAGAAGAAAGAGAAGGTGAAGCAGAAGAAGAAAGAGAAGGTGAAGCAGAAGAAGAAAGAGAAGGT
Consensus

Teleogryllus_occipitalis      ------------------------------------------------------------------------------------------------------------------------
Gryllus_longicercus           GAAGCAGAAGAAGAAAGAGAAGGTGAAGCAGAAGAAGAAAGAGAAGGTGAAGCAGAAGAAGAAAGAGAAGGTGAAGCAGAAGAAGAAAGAGAAGGTGAAGCAGAAGAAGAAAGAGAAGGT
Consensus

Teleogryllus_occipitalis      ------------------------------------------------------------------------------------------------------------------------
Gryllus_longicercus           GAAGCAGAAGAAGAAAGAGAAGGTGAAGCAGAAGAAGAAAGAGAAGGTGAAGCAGAAGAAGAAAGAGAAGGTGAAGCAGAAGAAGAAAGAGAAGGTGAAGCAGAAGAAGAAAGAGAAGGT
Consensus

Teleogryllus_occipitalis      ----------------------------------------------------------------------------------AAAAAAGAAAAGAAAAAGAAGAATAAAAAGGAGGAGAAGAA
Gryllus_longicercus           GAAGCAGAAGAAGAAAGAGAAGGTGAAGCAGAAGAAGAAAGAGAAGGTGAAGCAGAAGAAGAAAGAGAAGGTGAAGCAGAAGAAGAAAGAGAAGGTGAAGCAGAAGAAGAAAGAGAAGGT
Consensus                                                                                       ** **  ** ****   **** * ** *** * ******

Teleogryllus_occipitalis      GAAGCAGAAGAAAAAAGCAGAAG--------------------------------------------------------------------------------------------------
Gryllus_longicercus           GAAGCAGAAGAAGAAAGAGAAGGTGAAGCAGAAGAAGAAAGAGATGGAGCTGCAGAAGAAGAAAGAGATGGAGCTGCAGAAGAAGAAAGAGAAGGTGAAGCAGAAGAAGAAAGAGAAGGA
Consensus                     ************ **  *****

Teleogryllus_occipitalis      ---------------AAGAAAACTGAATTAGATGAGAGGTATATCCGAAACGTAACGCGGCGCTCCACTTCCTCGAAACACTTTCGCCGCTTACGGGAGCAGCCGCGCGCGGAGGCGGA
Gryllus_longicercus           GAAGCGGAAGAATAGTAAGGGAAATGAATGGGATGAGAAGTAGCCCCGAAGCGCCAAGCGGCGCTCCACTTCCTCGAAACACTTTCGCCGCTTACGGG---------------------
Consensus                                     ***  ** *****  ******* ***   ***** **  * ****************************************

Teleogryllus_occipitalis      GGCGTTCACCTCGGCGCGGCGCGGAGCGGCGCAAGGTGGGAGACCGTGAGGGACTGCGAGACCTTGACACTCGGACGCCGCGGCGGCGGCGGCGGCCGCCACCACCACCTCCACCTCCAC
Gryllus_longicercus           ------------------------------------------------------------------------------------------------------------------------
Consensus

Teleogryllus_occipitalis      CTCCACCACCACCACCACTCCGGCCAGCGGCCGGCCGCCGCCGCCGCCGCAGACGTCGCGCGATCGCGCACGCGGACGCACGTGCCTCCCGC
Gryllus_longicercus           ---------------------------------------------------------------------------------------------
Consensus
```

**Tocc.2G0020270.1_exon1**

```
Teleogryllus_occipitalis          ------------------------------------------------------------------------------------------------------------------------
Gryllus_longicercus               YR*KRYLSNFFRTKNSRRQLTH*NLVFQ*LNLCNYFFIVLIFLSSTIKIVFI*NIPSLFVSHFKRVICRCMSEMIAMSLISY*SSCQGQKNKTKL*LITASPFFFTIFHLLSLLEINLIH
Consensus

Teleogryllus_occipitalis          ------------------------------------------------------------------------------------------------------------------------
Gryllus_longicercus               ELNRFCLTWHAQKKVQLVKICILKAISKYLLWVTIVSRQKHHYSTEM*M*IMSRLRKRLELQGKFSLMENSNAYSRFTT*LQLQIVASPTLRKNTTSINSRCFQVEPQMLDISSLYISLS
Consensus

Teleogryllus_occipitalis          ------------------------------------------------------------------------------------------------------------------------
Gryllus_longicercus               KMH*LYIF*RFFLPSVLHCQSVCVIR*LHCYGQFLSPVFSLLHYLHECFMTNKTCSNE*SNAKSLKLTDNSKKHFVFT*KIFHALYNYCTHSTRT**YNT**K*APIQI*KKK*FSNLKS
Consensus

Teleogryllus_occipitalis          ------------------------------------------------------------------------------------------------------------------------
Gryllus_longicercus               NVLYSLIIREYIYV*KRIIYW*IHICTEKGHLEFIYLFERYSECIKSIFTVHIFKYFTYYNTFFIFLSLTVF*MTKRNIYTVSHTSLQYSSWNRLSEAKFCCAPVTSPSP*IVKLITIFW
Consensus

Teleogryllus_occipitalis          ------------------------------------------------------------------------------------------------------------------------
Gryllus_longicercus               EITNSERRMLPKQK*LQHYVCSCFLEAMRI*I*QRFLTEHCLTAQPAPCRETELSLPGTLKWESFCS*NIGHSCKM**LLWIFLRAMYRYVVIVSIN*FQ*P*IVH*ELFSFLSEVLKLT
Consensus

Teleogryllus_occipitalis          ------------------------------------------------------------------------------------------------------------------------
Gryllus_longicercus               K*CRCHLRVFKSQIQF*FI*CTIATFSVCRHNC*HFQSRKK*Y*MR*K*REQEVRV*EEEMRRKKKYEKKKRYEEK*NVGRTERRYKHLKRRKREGRRNE*REEQKSAEA*RRGKM*CKM
Consensus

Teleogryllus_occipitalis          ------------------------------------------------------------------------------------------------------------------------
Gryllus_longicercus               EEKAEEGEAEEEREGEAEEERR*SRRRKRR*SRRRKRRRSRRRKRRRSRRRKRRRSKRRKRRRSRRRKRR*SRRRKRR*SRRRKRR*SRRRKRR*SRRRKRR*SRRRKRR*SRRRKRR*S
Consensus

Teleogryllus_occipitalis          ------------------------------------------------------------------------------------------------------------------------
Gryllus_longicercus               RRRKRR*SRRRKRR*SRRRKRR*SRRRKRR*SRRRKRR*SRRRKRR*SRRRKRR*SRRRKRR*SRRRKRR*SRRRKRR*SRRRKRR*SRRRKRR*SRRRKRR*SRRRKRR*SRRRKRR*S
Consensus

Teleogryllus_occipitalis          ------------------------------------------------------------------------------------------------------------------------
Gryllus_longicercus               RRRKRR*SRRRKRR*SRRRKRR*SRRRKRR*SRRRKRR*SRRRKRR*SRRRKRR*SRRRKRR*SRRRKRR*SRRRKRR*SRRRKRR*SRRRKRR*SRRRKRWSCRRRKRWSCRRRKRR*S
Consensus

Teleogryllus_occipitalis          -----------------MR--------------------
Gryllus_longicercus               RRRKRRRSGRIVREMNGMRSSPEAPSGAPLPRNTFAAYG
Consensus                                            **
```

**Tocc.2G0020270.1_exon2**

```
Teleogryllus_occipitalis          ------------------------------------------------------------------------------------------------------------------------
Gryllus_longicercus               V*VKEISLKFLQDKELQTSIDSLESRISIIKFV*LFLHCVNILKQYYKNSVHLKYSFIIRFPFQACNMQMYVGNDCDVLDIVLIKLSRAKKQNKIVANHCKSIFFYNFSFIIFT*N*FDS
Consensus

Teleogryllus_occipitalis          ------------------------------------------------------------------------------------------------------------------------
Gryllus_longicercus               *TE*VLLDMACTKKGSASKDLHLESNFQIFIVGHNSKSSKTSLLHRDVNVNHESSSEATRTSRKILPYGK**CLLKVYDLTSTSNCR*SYFAEKYNFY*FSLLPG*TTNARYFLVVHFLK
Consensus

Teleogryllus_occipitalis          ------------------------------------------------------------------------------------------------------------------------
Gryllus_longicercus               QNALAVYFLKIFFAFCTSLSVSMCHPIITLLWTIFVTCVLPFTLSS*MFYDEQNLFQ*IIKCQISETHGQFQETFRFHLKNIPCII*LLHA*YTYIVVQYIVEISANTNLKKKMIFQFKK
Consensus

Teleogryllus_occipitalis          ------------------------------------------------------------------------------------------------------------------------
Gryllus_longicercus               QCFVFTYY*GIHLCLKTNYLLVNTYLYRKRTSGVYLFI*KVF*MY*IHLYCTYF*IFYLL*HIFHFSQLDSILNDEKKYLYSITYEFAVQFLESPI*SEVLLRSCDLTFTVNCQTYHNLL
Consensus

Teleogryllus_occipitalis          ------------------------------------------------------------------------------------------------------------------------
Gryllus_longicercus               GNYKLGAKNVAETEVIATLCLFMFLRGHENLNLTAILNRALSYRPTSTLPRN*TLSARNAEMGVLLFIKYRSFLQNVITPLDFFASNVSICGNCFNKLISITINCSLGAVFFSIRSSQAN
Consensus

Teleogryllus_occipitalis          ---------------------------------------------------------------------------------------------------------------KKKK-----
Gryllus_longicercus               KIMPVPFESF*KPNSILIYLMHYCDIFCMQT*LLTLPEQKKIVLNEIEVKRTRSKSIRGGNAKEEEI*EKEKI*REVKCREDGKKI*TFKEKKARGKKK*IERRTEKCRSMKKRKNVMQN
Consensus                                                                                                                                           ** *

Teleogryllus_occipitalis          -RKRRIKRRRRRSRRKSRRRKLN*------------------------------------------------------------------------------------------------
Gryllus_longicercus               GRKSRRRRSRRRKRR*SRRRKKVKQKKKEKVKQKKKEKAKQKKKEKAKQKKKEKAKQKKKEKAKQKKKEKVKQKKKEKVKQKKKEKVKQKKKEKVKQKKKEKVKQKKKEKVKQKKKEKVK
Consensus                           ** *  * *** ** *****

Teleogryllus_occipitalis          ------------------------------------------------------------------------------------------------------------------------
Gryllus_longicercus               QKKKEKVKQKKKEKVKQKKKEKVKQKKKEKVKQKKKEKVKQKKKEKVKQKKKEKVKQKKKEKVKQKKKEKVKQKKKEKVKQKKKEKVKQKKKEKVKQKKKEKVKQKKKEKVKQKKKEKVK
Consensus

Teleogryllus_occipitalis          ------------------------------------------------------------------------------------------------------------------------
Gryllus_longicercus               QKKKEKVKQKKKEKVKQKKKEKVKQKKKEKVKQKKKEKVKQKKKEKVKQKKKEKVKQKKKEKVKQKKKEKVKQKKKEKVKQKKKEKVKQKKKEKVKQKKKEMELQKKKEMELQKKKEKVK
Consensus

Teleogryllus_occipitalis          ---------------------------------------
Gryllus_longicercus               QKKKEKEKRKNSKGNEWDEK*PRSAKRRSTSSKHFRRLR
Consensus
```

**Tocc.2G0021750.1**

```
Teleogryllus_occipitalis   CAATGGATTAATCAATGGGCCGTTTCGTACGCTTCAGAG-----CCGTCCACGTTGAGCCGGACGCATCGCCCGC------------------------------------------GG
Acheta_domesticus          TACGGATTTGGCGCAAAGCTTTTTTTCTCCATGCCGAATTGTGAGATTGCCCTTTGAACGCTGTGAATCTGACGCTCCAGTAAACAATGGAGAGGACAAGGACCGAAAACAATTCTCAAG
Gryllus_bimaculatus        TACTGACTTGGTACAAAGATTTTTTTCTCCATGCCAAAT-----GAATTGCCTTTGAACATTGTGTATCTGACGCTCCAGTAAACAATGGATAGGGCAAATACCGAAAACAATTCTCAAG
Gryllus_longicercus        ------------------------------------------------------------------------------------------------------------------------
Consensus
```

[illegible]

**Tocc.2G0021750.1_exon1**

```
Teleogryllus_occipitalis  ------------------------------------------------------------------------------------------------------------------------
Acheta_domesticus         KYCFYIFY*KLYFTFIMNAIDIL*VYFVSAFVKYDHSIIYKHK*LHARTKIIKFFAIF*NILLWNLIIVTN*TK*NV*LYCVLLTIRKESNKNFEGI*VSF*AFYTSSSLSIFFETLYRI
Gryllus_bimaculatus       ------------------------------------------------------FSLCT---LVNLF**HKISK*R*MSDCLL-------------------------------------
Gryllus_longicercus       ------------------------------------------------------FEI-----------------------IL-------------------------------------
Consensus

Teleogryllus_occipitalis  ------------------------------------------------------------------------------------------------------------------------
Acheta_domesticus         *KEGINLQVPVASNLSYLSYKFY*RLTQKIIL*SFNLVPNCIFLCPGIFTQLDILCTSKGTV*CTSSCCRLMFILITQDKYLFN*QVE*HLMKLNILNLEYFHVSSEIIIGV*NTKITIN
Gryllus_bimaculatus       ------------------------------------------------------------------------------------------------------------------------
Gryllus_longicercus       ------------------------------------------------------------------------------------------------------------------------
Consensus

Teleogryllus_occipitalis  ------------------------------------------------------------------------------------------------------------------------
Acheta_domesticus         V*LFTDLAQSFFSPCRIVRLPFERCESDAPVNNGEDKDRKQFSRT*IRMTMLKIQGHIQKVADCKGIDIKNSFCNKKCSRTTQKYFAVLAKDLKHII*DSNKK-------SLTIFRSI--
Gryllus_bimaculatus       -----TWYKDFFLHAK*IA--FEHCVSDAPVNNG*GKYRKQFSRT--QMTMLKVQGHIQKVAKFKGTE-RKKWGNKHCRRTKQKYFAAPTKDLKHTIKDNGKRYELPKIFS*HLYQSIRN
Gryllus_longicercus       -----NFLKRFFGF*TILV--------------------KRFSV-----------------------------------------------VKHKID----------------F*SSNA
Consensus

Teleogryllus_occipitalis  ------------------------------------------------------------------------------------------------------------------------
Acheta_domesticus         YID----LLKC*NDIDF------------------------------FVIKNDSL-LFKRLYSNLSRP*KSESISNVLRQISKL*LNY*PSCIFLHSYNSHTLHTSVIVV-T**SCMVS
Gryllus_bimaculatus       YIEFFKKILWFLNHFT-QTFLGRKT*NRFLKF*RICQRCN*IIATFACFYIQNIV--LIK-YSF*IFYP---------TRYHHQFKQNFE---FFVNVATP*TYGISYS*SPVWRRILVA
Gryllus_longicercus       YVKSVIKLLQLLHVFTFKLFC**NTVSEY------------------FVPQDITINLIEILNFL*MLP---------LQKLTGYP--------IVRVLSGEEF--------LWLTALIG
Consensus

Teleogryllus_occipitalis  ------------------------------------------------------------------------------------------------------------------------
Acheta_domesticus         DFFFQYRFTKLLYSIFVQL*NSHSSDNTAMETVSCKLRC**RIKCNIPVLSSCFQ*RGKVCALVAPLDFFK-TSRMLQQSVKTETYHTRTALPHP---FTILSHAHTRTKLICKGDQTGP
Gryllus_bimaculatus       DGFDR-------------------------------------------------------------*FF*LRPGECRGSVMTNEYLSRAICPSSPLSINFLLHTLT-*SIISLTYRKG-
Gryllus_longicercus       SFFDQ------------------------------------------------------------D----QANAVAVS*QTSIYHERFVPPHPCPFISS*THLHK-V*YHSRTGKGGS
Consensus

Teleogryllus_occipitalis  ------------------------------------------------------------------------------------------------------------------------
Acheta_domesticus         *RSTRLGRCAAGG-DI*RREIPQPGNRDRITCSSGIRTP-----------SRPAM*H-------ALQSRD*CLKQFTHPIY*LNRDSGA-SLGS*VHC--------------TLFFLVG
Gryllus_bimaculatus       ---------------------------------WLRALNELVQFF*LRGSRIAMWQDF*YQLDS*VESSRCLEN--------N-SNGR-SL*SVVFCGFFDQHWVNARA----------
Gryllus_longicercus       GH--QMSQCSSSA*EVLGLEC------GRIFDTSWIATL-----------SPVAIWR-------------RILEA--------GVSNG*FPVVSPINSG*MAEQYPNKRARVCTLYF---
Consensus

Teleogryllus_occipitalis  ------------------------------------------------------------------------------------------------------------------------
Acheta_domesticus         DGDLPSLYGGIQQRLA--CRLAGFYPTRYLPPVVHRPREHAW*CSAC*CKSVCKRRG*DKEEEGKIVRGPW----PVC*DTAPVFAWCLSENPRKTTDQSRRPLEFYSRQQLNLT*LYS*
Gryllus_bimaculatus       ---VP*QTG*GLHAVYPSSSLSYRYPLLTQTVRSHAPHYHT---------------------VGS*ELAPAGVGVPLVGDT-----W-FGKIPTR-----RRRTAPYLIQREKIAATFRE
Gryllus_longicercus       ---LPHPCHIVTHSLH-------------TQRSHAPHYHT---------------------VGS*ELTPAGVGAPLVGDT-----W-FGKIPTR-----RRRAAPYLIQRGKVAATFRE
Consensus

Teleogryllus_occipitalis  ------------------------------------------------------------------------------------------------------------------------
Acheta_domesticus         CRKSCHRRDMNLEPSKTEVMQCHQ-LNALSHLSVTLVRTILTYMMGESFRACPSW-GCVPLDRPRGTNTTRWRIGTESHFLVVEG--HYHFPKNHYNYGTNAQLTHPENKIINQSISGYI
Gryllus_bimaculatus       GNMHCTYETNGL--SR-*RARIHQSINQLPHMPGDQTGT*RSSRLRRCTARGDNWKGKSP*E----------VGVEAEISSNSGARTLTLPP----------IQHAITGLV---TEKKI
Gryllus_longicercus       GNMHCTYGTNGL--SRCSRCLSHQSINQLPYMSENQTGT*RSSRLGRSTARRDNWKGKSP*V----------GGRSQEI---AGEEH*HFQ*----------IQHAIT*LVPLAPAKKI
Consensus

Teleogryllus_occipitalis  ------------------------------------------------------------------------------------------------------------------------
Acheta_domesticus         IQWLVRLMAAHTAPTLLARGACRPVSPCRGRAGAAASGGAARGR------GGGGEGGRTGGRRPRAPAAGA*RVLPPGRL*DGRAPDPGHSLRRLLSAPPDTHSSAMVSLRGAAPALSTV
Gryllus_bimaculatus       SQ---------------------XXXXXXXXXXXXXXXXXXXXXXXXXXXXXXXXXXXXXXXXXXXXXXXXXXXXXXXXXXXXXXXXXXXXXXXXXXASV-APLS*HPPLPW*VRTAPRG---M
Gryllus_longicercus       SQWM------HNP---VAR---TPQSRPHGAGRSAPCGGVAGGRARRPRPGRGRADARAGGGPVRPPPPPPNACFHPRDYKTGASPTPATASVASV-APLN*HPPLPW*VCKAPR-----
Consensus

Teleogryllus_occipitalis  ------------------------------------------------------------------------------------------------------------------------
Acheta_domesticus         RMRCRN-----------------------IDVHHLCISHRFLESHVF*YICFVFDIFIFK*KR*YHNKMQ*CVSD---ISLTFFGTKFG-VEIKIELLKSIQFFK*QFQRKKV---C*I
Gryllus_bimaculatus       RHRCPG-------------------FYGIIDGLHFEFSLHR---------CFVFDIFSFDWKNYQLNKFL*SFSY---IN*TFSRQKLLRSEIKIYVLKSIHFLKYLFK**SIAKYTDI
Gryllus_longicercus       RPRPQGDATPMSRFLWDH*WT*L*IFFASMLCVRYF*F*LEKLPS-----**SLVIIFT-----YQLNMFQTKVP*GVN*N*RFEVHTFF--EISIQMIKYC*IY*YLFS*LNL------
Consensus

Teleogryllus_occipitalis  ------------------------------------------------------------------------------------------------------------------------
Acheta_domesticus         F**GFTVFCKLHPNKHDLRQVNPHV*TLMATIIV*EHQWFGGDILLTSLL*----FPVPVFLEMVAS*SPVVVIT*FSRRNCGVIFIFIF*VFRVLSTYLGFPENICFGICVLTKN--IK
Gryllus_bimaculatus       CLAS*IFYKKL--INATFGQIMSPVCTFLINILIKKHRWFSGDVF*TSIDFEHRCFETCMF----C*RRRFPILQ*SS---------FRS*LKQTVM*F*FFAS*RFWGC*IRIENNIYQ
Gryllus_longicercus       -------L*KL--INATFGQIMSPVCTLLINILIKKT*MVFG*CFLNI----YR-F*TSMFR-YVCSVDDVVSLISSSR-------HFVVR*NKL*CSFDFLLLDVFWGVKYESRTKYIK
Consensus

Teleogryllus_occipitalis  ------------------------------------------------------------------------------------------------------------------------
Acheta_domesticus         KYFRSCFLPVSNN*RIQA-----NKLFFTKIL----------------------------KKVHT-----------------------CLENPHN-----------------------
Gryllus_bimaculatus       K*DFCPFF----NY-IKKLSYLRKTLFFTKIFHVTNNFR*HIY----------------ASVTLEEAWNKCVEFWLIF*TY*V*HVKPILKNVRNFACVTYTYNKYNIIK*FLARPSKG
Gryllus_longicercus       NRIFVRFLTTSKNYRI*GKHYSLPKYFMSQII-CDSTFMHH*L*KKPGTNVLSLG*YFQPTKVGTSS*Y*NMLETSLMLRIFI------ITKT**N---------------NFLARPPKR
Consensus

Teleogryllus_occipitalis  ------------------------------------------------------------------------------------------------------------------------
Acheta_domesticus         ----------IFSSILLKYLKGSGQK*PQ*ICFQRYLWCLRM---------------HNEYLP-------------------------------PHTN-----CNLEV*STHKNVFSGI
Gryllus_bimaculatus       RNMHKIIQISIYKNIFL*IINFS---------YHQFDYVVRAVLSLGTPNSCISLPRNSDWLPTLVKEEVENLINHFMMGRLGYSLTDVHMRLENPHQY-FFISSKIL*KNSHKNSV---
Gryllus_longicercus       RNVHKIIQISIYKNIFL*IINFS---------YHQFDYVVRAVLSLGTPNSCISLPRNSDWLPILVKEEEENLINHFMMDRLGYSLTEVHMHLENPHQYFFFISSKM**KNSQKNSV---
Consensus

Teleogryllus_occipitalis  ------------------------------------------------------------------------------------------------------------------------
Acheta_domesticus         LYMF*NDPEKICLLPVDKDVL---NDTQLAGKQWSRRC-------------ISVSNRQTH-------------ANCTIYMIVCTKI-------------------QMCFQFFRISTKN
Gryllus_bimaculatus       -----NSSCEVLNISVDAS*IPSFRNHQFKSLTYSRSFPQEFLYNVL*GFGKNVSPTSTKML*MSPNVLQNSGPENVYLYRADACKL----------DDLHGYT*TIQVCFYYFKF----
Gryllus_longicercus       -----NSSCEVLIISVDDS*IPSSRNQQFRSLTYSKRFPHEFLYNVL*GFGKNVSPTSTEML*MSLNLLQNSGPEDV*LYRADSCKLAFTWVYVDYSDVLLLFRVSTKL---FF*L----
Consensus

Teleogryllus_occipitalis  ------------------------------------------------------------------------------------------------------------------------
Acheta_domesticus         FQKEL*QKH*KISNHQLTI*ESKKTLVVN*G---SLV*RIFYRNDYFSS-----CCFREC*SRSADE----RVEEG*GGMCEVSGKPGAATARLSFFVSRGFVPSQFKISTNV-------
Gryllus_bimaculatus       -------------------RSNSSFSYDQTLFL*LIEEIWFKRCSIETFIFLLCFFR----EWCSNSLVGRMRRG----GEGRGKAGAAAALSSYFVGREVLQTAVGIAITLVYY*FCE
Gryllus_longicercus       -------------------RPNSSLVSN*G---NLV*KVFYRNVYFSSLLLPGVVFK*--SRWADE-------------EG*GRAAGRPA-----------QRLHDLPISW-------
Consensus

Teleogryllus_occipitalis  ------------------------------------------------------------------------------------------------------------------------
Acheta_domesticus         -------------------SCVSVFTFPYTYGVIYVV*EK**NAARKNGVFWTETNSLPGVDNEKSG*NQ----LIN*EKN------------PLK----------HG---*RIFPEN
Gryllus_bimaculatus       THSIRRRRKKCNNCVCRK*RRCNVNLTEPTD*YLIELVVETVDRY-----IVDYYK*DY*GTDKEASHKEE-----LR*EDS------------PPTTFSQ*VRS-------TTIFPSK
Gryllus_longicercus       ----------------GARYCNRP*V*Q*RWYVISSVKLT-----------------------ASEGEERNVTIVYVENSGV*CQFN*AY*LMPY*IAGRNCRSIYRGLLQMRLLGHR
Consensus

Teleogryllus_occipitalis  ------------------------------------------------------------------------------------------------------------------------
Acheta_domesticus         K----LGKSDVV----FRN--------DKVRGRLHHTHLKNDSIQNNF-------------------------RRTVEVHLTR-----------------------YKYVTNF*GI--
Gryllus_bimaculatus       *RIWTLG*V*IKDY*EFASNV*IHVICVKIETKRLYNFVLQEGILVNISLVTYSVILRI---------------KYIIKVVFAK-----------------------------FLTIST
Gryllus_longicercus       QRSQTLREVEIR----FAAYLVFAI--SKIDPNLSF-----EMTHLNITLSVNKGQLRICK*C*DSSNLCKDRK*TNIQFYFARRYFSEVFPWLLILLF*GSNI**ASR*FLLNVLTISI
Consensus

Teleogryllus_occipitalis  ------------------------------------------------------------------------------------------------------------------------
Acheta_domesticus         --*AANEYHFSYLNMWEM**TKYIC*LSETAFL--------------ICKNV*L-------------*FN*NEHCRD*AS*NLQ--------GRHFKNFL*SKLASKNKKISLLVNRSLV
Gryllus_bimaculatus       LNIACNIFVFNHSNISIYKDSFWIVYVHVSF*FYHIN**DIIRHNN**DKSVNR*LVFQPYCKRKP-AIGFVEIIKTPNQVVISLFGWPIIFGGAGLDN*CALFPSKRGCIGCRLWDSIV
Gryllus_longicercus       LNIACNRFVFNYSNIYIYKDSFWIVYVHVSFWF-------VILTNKKLC*SIIR---FPPLLQKEGWVLGL*KPFKSKSSFSFT-FGLVNYFWRSESG*LMRTVPPQGGCI*CRLWDSIV
Consensus

Teleogryllus_occipitalis  ------------------------------------------------------------------------------------------------------------------------
Acheta_domesticus         SSCVRL--HAKH*CRKCCL---------------------PIRGST*E--I*CPQF------------YL*L-----------PTTHS-LC----------------------------
Gryllus_bimaculatus       -*CLKTVEHTSH*LRE-LFYDQHI*MICQNTRIAKILEIFGSSNISEYEYRANCAVFECY---------*INL--RFSYTCSCTVTTE*LICALGRSS----------WFS*N*KFSTGS
Gryllus_longicercus       -*CLKTVRHTRH*LRKCIFRPTH------------------SYDFSKYE---NCENLRDYWKIRTSVNTSIELIAQFLNTIK*TYVTHTHVCVQ*QQSN*FDLLEVAPGFLENENFRQKA
Consensus

Teleogryllus_occipitalis  ------------------------------------------------------------------------------------------------------------------------
Acheta_domesticus         --------KIFVVAYKT-----*KFLKLIMNHCESF-----LRRSN*LMVP*-----------------------------------ATTAVIRLVSE---------------------
Gryllus_bimaculatus       --------RFHQR*SKTSPHSFVK*LKYT*V*NKCYKCLWHLKCQRLSLFPIKYEGNVKNEVTQVIILISRPGF*SILSFD*VRTCIVITTQPINVMAD-------------------P
Gryllus_longicercus       DCIKDDQKRVHIALSNS-----*ITLKYIMDFTNIYGI---*RVKDYNLLQL-------NMRTMSKMMLHRLSF*YRAPVS*VYLAFIGSERALSLLSNL*TSWLTLQNFL*WKQISKTE
Consensus

Teleogryllus_occipitalis  ------------------------------------------------------------------------------------------------------------------------
Acheta_domesticus         --------------VNVSSRDV-----------------------------------------------------DMFLL------------------------------VGS*DLTP
Gryllus_bimaculatus       SKILCSR-----------NKSVKSKKAAIVSEQVAPESSSECQP-AV*EVLPAN--QGLDVRNSTSVILLY-EISHNIFIVQNVIMFDSIT*PTSFQYSF*IPLAK*CMSSSIG**DPTP
Gryllus_longicercus       NNCHCVRTGRS*EFVRVPTRSVGS---------VAYQSGAQCKKFNICNFVI*NFSQHIHCAKCDYI*LNYMT*KFSIFILNTV---SEVNVPPPQDYEIR----------------RRP
Consensus

Teleogryllus_occipitalis  ------------------------------------------------------------------------------------------------------------------------
Acheta_domesticus         TYMTACRVFL---SLGLVTLQSTAPVWRSSKALSCRLVSPHAK*CSTYVYGCKGSE*GRSKEGIVTLILGQIVRILL-------WH-------------SF---------GIERRNNEQ
Gryllus_bimaculatus       TFM-ACGVFP---SPGLFTHRGTAPAGRSSK------LPTSLPTCIIV*YM*VERE*DRAKEGKTAHDLGPIGRVLLRH*TEDPWKTIDQSTQLHDSR----------KCAGTRTLGHLG
Gryllus_longicercus       TWL-AV-YFPLQASP-----------------------PTGAPP-------------QRGEALSSRPVSPLV**CSR------WR--ENSTEQRKERLRTFLARLVGYCSGIERRIHGK
Consensus

Teleogryllus_occipitalis  ------------------------------------------------------------------------------------------------------------------------
Acheta_domesticus         LPIKTPDP--LNSPPKGD*IRPCNLFNCQKFAWSCTLRPFGQNR**VRDSPGSELISGSPPH*SCKAKVGNTRAHFNGR*NASFAAYTHTNGAW*TSALSPLRDA*V----STE*VCCM-
Gryllus_bimaculatus       LNI-----------------------CR-----------------HQHAGSELISKFPSH--CGTKAGYTREEFNGR-----------------*IHPLYGVHT*MGHGTRTRCAMS
Gryllus_longicercus       LSIRAPDSKILENAPELVR*GPLV*ISAD-----------------ISNVGSELISRFPSH*SCGAKAGYTRAEFNGR-----------------*MHPL*SVHTRMGHGTRTRCAMS
Consensus

Teleogryllus_occipitalis  ------------------------------------------------------------------------------------------------------------------------
Acheta_domesticus         -LLCVSALAPCR*EFFQQKLFVVRTAA*RRKPGTHHNHNRC*KYCP-----------------FTWFPVV-------------*LLLT*MQL-----------------LIGYPNS--
Gryllus_bimaculatus       VVFCLSV----RRRDVCGNMDVLTSLTRR----------SCYR*CTALHI-------------FNYY*SVTHNP**GKGLLHCKVLNLQAI*LVSYYFCSLLPVPFFLFMCIITSP*I--
Gryllus_longicercus       VVFCLSA----RRTDVCGNMDVLTSLM*R----------SCYR*CSALHIIYYYYYYYYYYYYYYY*PVTHIPQ*NKGLLHCKVLNLQAR*LVSYYF-------------LFTSPCTHL
Consensus

Teleogryllus_occipitalis  ------------------------------------------------------------------------------------------------------------------------
Acheta_domesticus         -----AQCGETLLAR-CIILLQNWT---------*NSLVLSIEI*TKKSASSAQA---------------RG*RRAWLTVCVMNCVRDCVRG*LCAWLTVCVANCVRG*RCAWLTVFVTN
Gryllus_bimaculatus       -------FATTF-----VFYICNHI-SGHLT--FASNLRILFHQ*PKTSLRKMYVNFGPSGCWRSD*PRWRAGGRA----------NDVWFAGAVA-----------------------
Gryllus_longicercus       FVHVYFHFTNTFRNHFCVTYM*SHFWSSHICF*FANFISSIVEDFFKKTCMQMLVLLVAGGRFS------RGGARV----------GELMTCGLQALLHVLVA-----LACA--------
Consensus

Teleogryllus_occipitalis  ------------------------------------------------------------------------------------------------------------------------
Acheta_domesticus         CV-----------RVWCGVVRRRCCTCWWRWR------------------------------------------ARVRRRGPPSSAS-AAEDWASAAVSSPSN---PLSLPLTQSA
Gryllus_bimaculatus       --------------RVGGAG----VCVCR---------------------------------GARLAHR---------HRRRRTR------------------------------LRR--
Gryllus_longicercus       CAAAHASLIGVGGGRLGFGGGESQLCACAVEMQLIQLN*IDALFYVARLGP*GPLFRPTTQISLRSSHQLAEKGGISWARNRRFEFSSEASFRLEDLLHA--SKPTNMGQPALLPLRRKA
Consensus

Teleogryllus_occipitalis  ------------------------------------------------------------------------------------------------------------------------
Acheta_domesticus         HE-------------------------------------------------------------------QWKCN**RFCIC-----QLCCLLK*QLS----------IA*PLPSVSCC
Gryllus_bimaculatus       -----R*VSAVRMRSG-----------------------------------NGTDGGLAFVCDIAYGNSGFPWIDNLHQFSAVWKPENE*VLFFGFQTKGN*R*R****LT*N**SGSCC
Gryllus_longicercus       YEGFFRP*NGPRLRSGLNPPTWVQQSSTLTVTPSPPLMEALHLPAILLMEIAAFDGLIIFISFLLFGSQ------KMKTYS-----------FFGFQTKGN*RTR-***LT*N**TGSCC
Consensus

Teleogryllus_occipitalis  ------------------------------------------------------------------------------------------------------------------------
Acheta_domesticus         LK----------------------------------------------------------------------------------------------------------------------
Gryllus_bimaculatus       LETTTDGHFLAFKQLETDDV*AIEYYYFREQYSWK**RL*LLYLECQQLITAKQKMMVY*CVVTAS*RKKNYTWGDT*CSAQVSQ--FS*LGIRLHFTVDSSG----------------I
Gryllus_longicercus       LETTTDGHFLALKLLETDDG*AIEYYYFNKQYSWHD--------------------------NAS---NYYIWNDSD*SPQNQQ*WFTEVWILL---VEKKGTLGATCDVVLSPLDFHV
Consensus

Teleogryllus_occipitalis  ------------------------------------------------------------------------------------------------------------------------
Acheta_domesticus         ---------------ARNRLR**FSGFQTVGN***WLSL*KI*RIFSGFLQFL*LSISPKYVINLHNHCSSKSCTHFQ*LP*G*GI---TKNTI*LPGFKLRNIHVVIVCLISVIASGF
Gryllus_bimaculatus       WV-------------VKAWLRQSFSGFSCL--------------CFSPFLRSLSHAFTDSN--------SSKSYIHFQKRPESFDVGTTINNTIWLLEFKSQKILVER-----MIERGY
Gryllus_longicercus       WVSGFISGSIHQGFGAVKAGLRQSFSGFSGS-------------CFSSFLQSLSHAFTDSH--------SSKSYTHFQKRPEGFDVGTTINNTIWLLEFKFRKILVEL-----MIERGY
Consensus

Teleogryllus_occipitalis  ------------------------------------------------------------------------------------------------------------------------
Acheta_domesticus         LKIISFTWLLFHFSLYHSYGIFIISSVSDN*SILCKIL**RIFTLDVRRTVLK*SGCLIYSSVN---------AVQILACP---------CGIFGEESGFTAGFLESSWFPRIS------
Gryllus_bimaculatus       ----------------SYRFSRITFVSPTSNATYVTSCTDVVINNVHLRLLREP----Y*SGKAVQFILLGIAVQILVCPYDLW*RKRVCGRF------------SWVFTVSLHFIPS
Gryllus_longicercus       ----------------SYRFSRITFVSPTSNATYVTFCTDVVINNVHLRLLREP----Y*SGKAAQFIPLGIAVQILVCPYGL**RKWVCGRF------------SWVFTVPLHFIPS
Consensus

Teleogryllus_occipitalis  ------------------------------------------------------------------------------------------------------------------------
Acheta_domesticus         -FRHCSNNIHIMFLKS---------------HSPKN----PPPCSRY--IQKTSNRHYYEDKMIQFAHGRSAA-*Q*GC*LLLNF--W*TKQISWR*S*DSPISPFFSFYH*FITHIIS
Gryllus_bimaculatus       LLQQNSHHIFVIIYPS---------------KSFTNF*IRPA**R*YGIIVKITSLSLTIRDLL*SR*GIVILLYHGGILGIISFPLHVTSQL---------------LYHP---HYIV
Gryllus_longicercus       LLQKNSYHIYVIIYPSIH*LLDKTSIIKMIWYHSEDNFL-------------IAYYS*SVAKQLGHCNPIVIM---EGFL*IFSFPLHVISQL---------------LYHP---HYIV
Consensus

Teleogryllus_occipitalis  ------------------------------------------------------------------------------------------------------------------------
Acheta_domesticus         S*SSSH*ILIYTESTTHAI*NNFLYK*KDCNMVMMYFCETFLKLHIYCTVSLYY**GRGDKR*YCNFAV*MF*NIS**FYSLTLKFQVRDKV*YNM**ISSERLSSQTK*RKRCKCEDGK
Gryllus_bimaculatus       TIIYSLNLLIYPENIPIFIVKNFYC*WKYQ*HGFGIIL*DIIKCHQ*IVVV--------G*S*YCHVAV*IL*NIN*LLFT*IKILSTR*SVM*HAIIIVEK*QIVKHK-----------
Gryllus_longicercus       TIIYKLNHLIYPGNIPIVMVKNFYCQLKCQ*HGFGIIL*DIIKFHKCIVVV--------**S*YCHVAV*IF*NIN*LLFT*IKIPSTR*SAM*HVIIIFEK*QMMKHK-----------
Consensus

Teleogryllus_occipitalis  ------------------------------------------------------------------------------------------------------------------------
Acheta_domesticus         GKYE***NTKTNNSNIYFKL-VLIQTLNSQRSTL*NNYYISF*NLNRKYSKSHG-ILCLHVHIRCKNAQL*LEFHRR-FHIM------------KYCNL*T*SL*QFLTN-TVHCTADR
Gryllus_bimaculatus       -----------NK*QVVFILNVLKNKR*ISAE*ISE---IMIT------------LCFKI*IVVVSK---IEITRRTFHISCKHGATM*TRISPYYFIL*NTSI--------------
Gryllus_longicercus       -----------NK*QVVFILNVLMNKRGICRENF*NNDYIMF*NLNRSYSKSHGALFIFRANKGQLCA---LEFHRR-FHIM------------KYFNLRT*SLLAISDSITLHCTTDR
Consensus

Teleogryllus_occipitalis  ------------------------------------------------------------------------------------------------------------------------
Acheta_domesticus         KHNMHA*LFSRETWRNVSFNWAI---LFVRF----------CN-----------------------------NIARMSMSNFIIGPM*NIP-----TIQRL-DWHVKIL*TVPSPLAHT
Gryllus_bimaculatus       ---*ERDLF-----------WQF---LTLSHFTAPQIAYTIC------MHNYFLGKHGETFHLIGPFVVFRL*NIAPMSKSNFVIGPM*NIL-----IMHRI-D*YVKVTCEFKI*I---
Gryllus_longicercus       IHNMHA*LFPRETW*NVSFNWAIRCVQTLKH----------CSNEHEQLHNWTYEKY--IYNAWNCLVCYGYL*I*YL-----------NILK*LY*ILNTI*KIFSKLILVFSSLI---
Consensus

Teleogryllus_occipitalis  ------------------------------------------------------------------------------------------------------------------------
Acheta_domesticus         LRLQGF*YNVQ*VPMNNHA*FQS------H-----------SFSFRDFLLRSPVFGII*LS**TPKRMTKISFSLNFQNFR---------KMQRFCK*MECKTK*DY*ISI*SKKRLPNC
Gryllus_bimaculatus       -----F*NNYIKYLILNRKYFQS*F*FFRH*SENIL*QK*LSYDRRFMI*RIQI-----------------YNGEFILFK*LE*LKCYLK*QTGLNVMEI-TGYTMKITCY*RKTKGSR
Gryllus_longicercus       -----*KYTLIKITVI**KI---------HDLKNSDIQRRVYF--------IQVTRTIKV---LP*IANRLEYDGNYWVY------------------------YEDNILLAENKRLPKK
Consensus

Teleogryllus_occipitalis  ------------------------------------------------------------------------------------------------------------------------
Acheta_domesticus         NLSVNYEASYTLLHPRMISHIMAQTL*FLLFYYYYSIV*TTWLSWLKKFVEISFFIVTIKLYCNSK-LKI*SVVLRSLEF*A*NQRR-CIYYNLCALKDEVSTVCVPQVASALAAAAWVW
Gryllus_bimaculatus       RNDV-FQYSFSLL--------------------------------MYLLLSLGFEG*NIDFLCSAGGIGLGGGGLG----LGLGLRGGIGF----------------GGGLGGYG---
Gryllus_longicercus       R----YS*LFIFL--------------------------------MDVFIIL--------------FGL*SAKYR----LSVSRRR--------------------------------
Consensus

Teleogryllus_occipitalis  ------------------------------------------------------------------------------------------------------------------------
Acheta_domesticus         AWACGGEASASAAAWAATGAAWVWATEATAVATEAGMAAATASAAAVWAWAWAESGCWAAAWGGGLGLG-------GGLGLRGGGGGLYGGGI-------GLGKGSYGLGTYGAGFGYYG
Gryllus_bimaculatus       ----GGLGLGGYGGGYGGGYGGGYGIGGGGLGLGLGRIGLLGGGGGL--------------GGGLGLG-----LGGGLGLRGGGGGLYGGGI-------GLGKGAYGLGTYGAGFGYYG
Gryllus_longicercus       -------------------HRPRRRRPGSRLGSAAGRHRLRRRPGRL--------------RGRLGPGRLRRWLRGRLRRR----------LRHRRRRSGLGPGPYRAA---------G
Consensus

Teleogryllus_occipitalis  ------------------------------------------------------------------------------------------------------------------------
Acheta_domesticus         K*ALSGRAAPSKGLSSPSKAP--APAPGSNPRPAKRRFHALMVRFGPVQ---RKGTSPKVFEFRRVDVVEIHDSSFTYDILLFSQN--------------------NYPFFRRFIFGS-
Gryllus_bimaculatus       K*AMSGRAAPPTGLSSPSKAPAPAPAPGSNPRPAKRRFHALMLRFGPIQ---RTGASPKAFEFRRVDVVEIHDSSFTYDILLFSQN--------------------NYPFFRRFTFGS-
Gryllus_longicercus       RRRRLGRRA-------------GPGPGRRLGPARRRRRALRRRHRPGQGRVRPGHVRRGLRLLR-QIGLVRESGPPHGPLLSQQGPGARARAGLEPAAGQEEVPRADGPLRPRPTHGRL
Consensus

Teleogryllus_occipitalis  ------------------------------------------------------------------------------------------------------------------------
Acheta_domesticus         ------------------------------------------FSPKEKIIKSISPHFF---SDKF*VVSCK*RTVCVSVI*IIFVHPKS------------CLTISCNVFPR-------L
Gryllus_bimaculatus       ------------------------------------------FSPKEKSIKSISHHFF---LDKF*VVS*K*RTVCVSVI*IIFVRPKAA*RSAVMYFRA*CMFM-CGILKKYILSCFSL
Gryllus_longicercus       PQGLRVQEGGRGGNTRLVLHLRHTALLAEQLSVLPSVHFWFVFSQGEEH*EHFTSFPFR*ILSGFLKVK---NCVCFSNLN-YFCSPKS------------CLTVSCNVFPR-------L
Consensus

Teleogryllus_occipitalis  ------------------------------------------------------------------------------------------------------------------------
Acheta_domesticus         VHIHVWYIKKIH-----FILFLVDLFAQNPFLHTYTSTHGFCTLIFSGLMQLIE*SF-------LY*ASIWLEEAFSIVNALTFF*NTCKFRIQAFLGINICKCA*FS*K*RTRKITSLF
Gryllus_bimaculatus       LYFH--HILFSH*HIPLFILIFEVVFWHHEIVALLTN------L-------------------------------------------------ASG*NSC----FF**LTQKHICKGL
Gryllus_longicercus       VHVHVWHIKKIH-----FILFLIALFS*HPILPLTLS------LIYFDLRSSILAS*NRPFIDQLCQA*ISF---F*IINSETYLQRTCKFEIKNVSGINNCD--RFNSK*RIRKITS*Y
Consensus

Teleogryllus_occipitalis  ------------------------------------------------------------------------------------------------------------------------
Acheta_domesticus         IIYNLEKSKSLIKINTYSYKINFNI-----------------------------LFFVSII*NLI--LDADFTHLY*MLRTGCGQIQELKRD-----------LRK*I*TSLNV*FT*L
Gryllus_bimaculatus       VSLKLRMFPE*IIL---IDSIQYNVSEK*LPNL*ITIQRTYKD*YLLLQK*T*MFLFFAFIT*NLF--LEAEFTHLFHMF*TGRGQF-ANQN----------------TIY*RVRSS*C
Gryllus_longicercus       INYNSEDL*R*ILI---T*KINVNVSLFCLYNLKLVFR-------------S*VYSLIPHVSNRAWPICESKYD------L*GSGQVNANKTECVIYIT***LLF*KLLTLSLKIKKIYC
Consensus

Teleogryllus_occipitalis  ------------------------------------------------------------------------------------------------------------------------
Acheta_domesticus         DNNPYIPNYSHSV*K*NKYNA*DVMTFPI*NK*LKYLIRKKHFKIMEEI------N----------TYSIL**LII----------FTYWLILPPQFKEKN------QFNM**SELMGD
Gryllus_bimaculatus       QQN*M---------CNLYN----LIIMSILKIINT*FENKKKYIA*E*SLQYNRNN*LLNRK*KLKYWKK*I*ILIHSN***NRI-WGYTHI--GWFCHNNLSKH*Y*KNMKLG--VGD
Gryllus_longicercus       LR---------------------IIIPV*QK*LNY*IENKN*NI--------KRN----------KYEF*FIAI-------NNKIEFGNTHILVG-FCHNNLSKH*Y*KNMKLG--VGD
Consensus

Teleogryllus_occipitalis  ------------------------------------------------------------------------------------------------------------------------
Acheta_domesticus         TVGYKSNTFYRYLSYSNRN*VICSRHFIRFCETNYFGQELTDIAVFWYFQVAALNNHQSSGTNLSFTTADVDLKS-SREVYSRFPWPQRTLFKREAEGVLPFHS-----MIFLLKRSSAK
Gryllus_bimaculatus       TVRYKLSTFYVYLSYSNRN*LIC*DILSDSARLIILGRS-*HIAAVCYFQVVALNNHRAAGTNLAAS---------AADVDLKSPPPARSTLVF-LGRNELCSRGRRRETFLFSARFSSL
Gryllus_longicercus       TVRYKLSTFYVY*SYSNRN*LIC*DILSDSARLILLGRS-*QILPRT--DTFKLWP*TATGQPVPTSPPVQRMST*PPEVYPRFPWPQRTTFEREAGRDVPLER-----TVLLLKHSSAK
Consensus

Teleogryllus_occipitalis  ------------------------------------------------------------------------------------------------------------------------
Acheta_domesticus         I-----MCSFDKN*LKISLLIGTNLLFRSHVISKNVLMLCI---SW-------RLAMMRK*K--SEFYECLTCKPRFTE*NPAPFSTRSITFL---FKLLYFF*LDRSSGDRFFMFFN*Y
Gryllus_bimaculatus       SIPRLKLCALSTKTEKR*VC*SAQTCYLDRMLSPRTF*LYVFHGGWP*L*GNRKVYLTNV*HVNQGFR----------NRTPHSSALEVLHFFLNYY--IYFNYTVLEETD--SLFFN*K
Gryllus_longicercus       S-----MCSFDKN*VKISLLIGSNLLFRSHVISKNVLTF--------------MYFMEVGH---GYEEIEKCILRMFDM*TKVSAIE--HRTVQHWKSCIFS*II--------IFILIR
Consensus

Teleogryllus_occipitalis  ------------------------------------M-----------------------------------------------------------------------------------
Acheta_domesticus         PRLPWSLRGSLY*ET*IQ--THASDMCCR*QKNY--FMIRWKF--LY-----------TLLKNCCD*SANLGKYR*NP--------KLNTLLLYC*KTSNA--RSVIKN*LYKLYTLDNA
Gryllus_bimaculatus       QWLP---LSVMLLYTNKS---ESNTFVGRRVVDIINFMNRCNF---------FINSS*KLLRVRCKL*KT*VKHKNKSLVA*FYA-ITNCSLELIIE-NN*L*RTTIIALC----PLDNA
Gryllus_longicercus       QFLKRQIL*VL*LISMASAVTNATLH**KRIQHVCG-MSRCRYKKLYE*MQIFINSS*KLLRVRCKV*KR*VKHKNKSLVA*FNANATNCSLELIIRI*SAR*R*QYVKLIFQIFPVDCA
Consensus                             *

Teleogryllus_occipitalis  ------------------------------------------------------------------------------------------------------------------------
Acheta_domesticus         GDTEDTSYGNSSVDCAQRLS--VYYGNYRTERPTA-RCVLLTV--*KY--SCD*LFEFI*FVTLF*LT---VFCI---Q*IRLSSIRLLT--RE---------IRFRREEWNSV*QVRGL
Gryllus_bimaculatus       SST*N*SF---------RFSR*IALKDFLLRMATTERSNLLHVAYCTWRTATNRVMNIYM*LII*NYFICHIFLIHYNSVYNKYKCQVFGCLREI*ASEERNGIRFNR*E---VSQ----
Gryllus_longicercus       QRLPT-----------------SYGDY*TKQSTA-RGVLPQTV**TY--ICN*LFEIVLFVTLF*FIIV-VFII---NIIDKYSVAT----RD-*ASEVRNGIRFNR*E---VSQ----
Consensus

Teleogryllus_occipitalis  ------------------------------------------------------------------------------------------------------------------------
Acheta_domesticus         LEINLKVDSFMPKEM*SK**IIDYLVILNMILFYTEMI*LIFDTC*LKK*HIDH-VLLSDN----KCKCTIFSILT---MCVGFLLHNTVQLLNVVQLNVR**PQQHELLVE--------
Gryllus_bimaculatus       -N*I*N*TVLCLRRFGLKNRLYVILFL*T*VCTTQRRMNPDLTTID*TRKTL-----FTELPLEITCKCIHFSKLIPWLFSSA*--HRPIKL----HLDVGS*RQPHKYIFRHC*WCSQR
Gryllus_longicercus       -KLI*N*TEFCLRRSRLKK*IICYFVPLNMILHYAEKIEPRFDNY*LNKENIVHGVVFRESM*MYTFFLK*FCGLV--FFCST----SPIKL----YLNVGS*RQHHKYIFRHC*WSV--
Consensus

Teleogryllus_occipitalis  ------------------------------------------------------------------------------------------------------------------------
Acheta_domesticus         FPVFFLPDRCEKE----NGIILMQIT*I*LILLC**TC*NFEKMSFSFIS*C--------------------------------------------------------------SLQL
Gryllus_bimaculatus       CLVFLLSDCCKNE----KNIIFIKITLIEQT--NLYNCVNEFYQIFKNI------IRIILFHLIVV*RS*NLRDIVGVSNYFLYNLFL----------NASMILKDLKMRHNLLVKSIEI
Gryllus_longicercus       VWAFSYPNVVKKKNSYANNIVLIYIIV-------LMNLIK*WYK*FSFI*LCSRELKICLIW------------SGTQTVFFITYF*MPQCFCMI*N*HTFYLKNR*KTWNL-------
Consensus

Teleogryllus_occipitalis  ------------------------------------------------------------------------------------------------------------------------
Acheta_domesticus         LKFS*YYL---DLKLFETNH*NKLK*TLKADLNDRGGKCIGRLSASR*RVTVVCSLSFSVCVCVCVCVCVCVCVCVVAEGLVRSLATEREAPNVSASSVKVNMCPHEGHIG*DSRNRYLL
Gryllus_bimaculatus       MDFFILYVFDVLLQLIKNSLLEKTIFFIKQ*CKN-----------------SLTILFSFT*CT*IVTMFIM*FWL-----------------------------HSNFISNDQR---IK
Gryllus_longicercus       -PFYTYLMF--YCQLVKLQHKRKLPF*QTIK*KP-----------------LDDIVFIYLVIYLVIHLLIM*LWF-----------------------------HSNFISKDQ------
Consensus

Teleogryllus_occipitalis  ------------------------------------------------------------------------------------------------------------------------
Acheta_domesticus         NMEKATGRKINKNSTLNINFTAILNLIILSVTKNSLRSVVHEIDRIMNR*TT-----SLPTFENLVIPSSGLVYKNN*A*RHQYEITSTIMCSLYAHNMLPHHM----------LET-KT
Gryllus_bimaculatus       GSEKRKRCVYYK---NSMKY---------------------EPVKIKTK-TKFQIV-----IQNLRL----LHFQSNSSSNFNVYMSRRKNCNPYKRNAVEKILGNRMPFHNCKFQTVKN
Gryllus_longicercus       --GKEKYVSY*KTV*NNITYCR-------ETRKN------EDKCKISNRNTKFEIAALLEQLQN--------*FQY-------*YETQKKIVTVTKGRL*KRYLEIKQ*ISNN-----QN
Consensus

Teleogryllus_occipitalis  ------------------------------------------------------------------------------------------------------------------------
Acheta_domesticus         KVMLYVKM----------YSQCNNLYNF*FVCLAYHIIVDVIS----------------------------Y*RLYVLLFCFAHHTS*RN*KIS------------KR*LFIILPIP*T
Gryllus_bimaculatus       NSILLMYQDRKPL---------------YF-----------------------------------------------LWFYLQHVICEGYLISVDV*M*Q*ARRYKQNLF-------S
Gryllus_longicercus       NSILHINQDRKPLLQ*FRF*NCKVVLLYWF*SSIFSILKV*IQLKLVRSNILKEKRLISCVNRDLFAAKLNVFVKMFCIL*FYLQHASCEGYLISVDV*M*Q*TRRYKQNLF-------C
Consensus

Teleogryllus_occipitalis  ------------------------------------------------------------------------------------------------------------------------
Acheta_domesticus         ASLNSSWS*VLQWLKNS*IKTTT------------------FYDKM-----SNFVFN*TWKLRLVIKKKKPLL----------SQVCSESSNKANEVLLRLA*WSAIEFLLYCAETFFIA
Gryllus_bimaculatus       ASFYQN-----HYLIQRLIRETA---TAIQ*LLCLTDTKKWFKFKIKYCVRKGYVSLKTSELR**NKEKAPPLKTTQGTPAKPTKPCFALHNGAH------------SGGSTAVNFFSA-
Gryllus_longicercus       ASF----------LPEPLFESTAYWRNCSTEIIMFDRHKK*FKIKMKYCVRKC*VTLKTSELR**NKEKAPPLKTTQGTPIKPTKPCFALHNGRQ------------QWRLLCGETFSAP
Consensus

Teleogryllus_occipitalis  ------------------------------------------------------------------------------------------------------------------------
Acheta_domesticus         LCKLRK---------------------RRKGILNSR-------------------------------KPR--LWAWHEVLSPVAVGI*THGSA-----FTAPCHSAVYLGLPF*P-GQ--
Gryllus_bimaculatus       L*RLRKRKSRFTVAAKQGSGRGGRF--RRP----SRVVFKRTDPLSQRRVTLQFIWA----SRSDPAK*R---WAQSERVF*ELVCITETINCAQRMWREQAYRTCFVLEMSFH*IGQPV
Gryllus_longicercus       CAGYEKGGADSLCP*SRARGVAGDFVGRREWYLNARIRFHSA-------VSLCSLFGPPVLTRPNDAGPRVSVFFKS*YV*PRP*IVLSECGGNKRI-EPAASSKCLFIE*V--------
Consensus

Teleogryllus_occipitalis  ------------------------------------------------------------------------------------------------------------------------
Acheta_domesticus         -MTLNPE*PCFLRVSMYN---*DDKLSP--------ANVAGTSVSNLLRPRNVFLLNRSTGVSTFFS----VTVMWI*-----------------FLIDI*KEGCGYES*KVVF*FQFN
Gryllus_bimaculatus       WARFSMSL*CGFNFSLM----FKRKVGKGMCILQ*FVNSCSTQIQILFIAFQTLRLD-----FNFFS----FTML*NAAINRCRITDDCS*FLECQLL---------RSFVP*V*CI-FS
Gryllus_longicercus       -NRCRHVFQCHSNVDLISH*CLKGRLGR-VCVFK------SSSLILVQHKFRYLLLSKPCVLISIFSHLQCYEMLQ*I---------DVE*LIE*LFVI---PEASTAKVISSVSVVHFS
Consensus

Teleogryllus_occipitalis  ------------------------------------------------------------------------------------------------------------------------
Acheta_domesticus         T--NSDILY------C--------FPNLAP*IPFF----------*I*KVMKYCNKST-----------SNFDVNTRIANRWY*VCF-----------------SNVCPTMSRYVT--
Gryllus_bimaculatus       SGGGGDVLPLDGEVRCWPPVSQRGFPQ--PRISYHLGSYPPGTTSKVSRHMW*CSACMCKCVYV------RGEDRARKGKERSRAVCCLLLGHCSGIRLVLIVKKPRKTTTQSSW-----
Gryllus_longicercus       SGGGGDVLPLDGEV----PVRQRGFPQ--PGISYQLGSPQPG---QPLRSPDTCGSVVHVCVSVCT*GVR*GQGRGRKHRARSVALC-------------------*GTAPAFAWY*TTE
Consensus

Teleogryllus_occipitalis  ------------------------------------------------------------------------------------------------------------------------
Acheta_domesticus         KVSQTCIA---FVQRVVDWYFCV---------------------RHL------------------------------LYLFFL*VVSY------------------------------
Gryllus_bimaculatus       TLG--------FSSRLADFDLTM*LV*KILPSTGLKP-------QHPHGSDS*YPEKTLPGNRCQILAKTHILGIS*KILLF*VSLFNHVPVSNVSASSEYRIFSKNY*YCTDPMPISTS
Gryllus_longicercus       TIGRRCRPKGLFSSQ*LN---STQLIVKNMPPSGSKPYSTRLLRQNLYTGSCCGPELTQPLVELQTLRRI--------V*LFAV-----------------------------------
Consensus

Teleogryllus_occipitalis  ------------------------------------------------------------------------------------------------------------------------
Acheta_domesticus         ------IWVPPPAGHCPRSDKH*APDWSPDMCGQCRARKVSGCR--CKEG-KGQ**GSKECAQLVARFLKVLIKHSPRIDLRNHEKLSVIDTSPRDCSPYNS*IQPR*LL*IRHSRDSNP
Gryllus_bimaculatus       SCEGFPLRVSPPAGHRHSQDKL*A-----------------*CKWVCM*GEKRQ**GRKDNARPGVRFQKILIQHSPGIELR---KLSVVGAGRRDCSPVST*IQHS*LSKICHLGDAN-
Gryllus_longicercus       ------LRMYPPY-----------------------------------------------------------------------------------------------------------
Consensus

Teleogryllus_occipitalis  ------------------------------------------------------------------------------------------------------------------------
Acheta_domesticus         RTSRSEPLYW----------*LLP*--TNSSACLKQIL*MI*PNSRHCERSFNFSCYYTC----------IIPYLF-----LQFCQQ*FQARLIMWRKSNFH*NPLNFPLQS--------
Gryllus_bimaculatus       HTVTDFP---------VRICTLVPVVALN*PSRLS--------NSRQGEGSFSFSPYYACIPSIFQILSTIVPYAFNNVVYVQFSMKLLKFRKLLF----L*TSPV--PLLK*KIFCT--
Gryllus_longicercus       -----FPNFVNMCSNCV**CGLCPI--FNTAS*LS--------------------------------------------------------EIII----F*ASPI--PLRK*EMFCTDL
Consensus

Teleogryllus_occipitalis  ------------------------------------------------------------------------------------------------------------------------
Acheta_domesticus         VSEVRDCLRICKLSYSTVFFIWYR**IIINAFYLNIQRF----CIIFRTFSIL----LRFFYC*GCQGKIEEYINSLLSTAKRDGRN-----------LEPSCL----------CS*QCL
Gryllus_bimaculatus       LTMLQNQI---K*FYKHVF-------------LTIQRF*ASSC---RTIAIFYKK*TRYFQKCKSKNKFQKRNSSGK*N*TV*AKNFLFEKTLFAKLFQVSCASFTKKLKVFPVKFQG*
Gryllus_longicercus       *TELQNQI---K*FYKHVF-------------ITIQSF*AISC---QTISIFHKK*TRSFQKRKSKNKFQKRNSSGK*N*TV*AKNFLFEKLFLPNYFKSPVLRLQKKLKVFPGKFQG*
Consensus

Teleogryllus_occipitalis  ------------------------------------------------------------------------------------------------------------------------
Acheta_domesticus         NQSLISACAIDNSSA*A*TTLVLP-HCGKTIIRTIILSRQVHRHIDKFLK*G--YLQFRDNTELKSK**NKQSN*TRQ*NI*NVIRRCPPRGTYQQFHRTTKQ*KYK*LVCNTCKFAVHT
Gryllus_bimaculatus       NISRLKIVILGNYYLMMKVMTLIPIKCTMEI*LL*ML-------IYKLFK*NT*FIQYIYNTIFMKN**FRGYSGTWKV-ILNVVK----------------------ILNSSGIIV*K
Gryllus_longicercus       NISRFKIVILGNYYLMMKVMTLIPIKCTMEI*LL*ML-------IYKLFK*NT*FIQYIYNAIFMNN**FRGYSGTWKV-ILNVVK----------------------ILNSSGIIV*K
Consensus

Teleogryllus_occipitalis  ------------------------------------------------------------------------------------------------------------------------
Acheta_domesticus         RKPNGRYSATIPTSTTSKSYATRTSSKIRTTKPNTTTTTSTFSVRCNKQRGGSSR------------IC----PTSTCGTYTNA*IPTPCRRYFRKKQQ*TTINKC--------------
Gryllus_bimaculatus       *INK*K*K*NISESHFSNTL--RLPNSL*ILPQIAPLPVHAFSVDGVDSADGDTSRSE*DTIYCLHALCHVV-STSTSIKEINFTV------KF*KIQ-LL*IKLSWHEILIIVINGMCV
Gryllus_longicercus       *INK*K*K*NISESHFSNTL--RLLNSL*ILPQIAPLPVHAFSVDVVDERFGRWRHFTEREGYNLLPTCALSRRVSTSIKEINFTV------MF*KVQRML*IKHSWHEILLIGINGEL-
Consensus

Teleogryllus_occipitalis  ------------------------------------------------------------------------------------------------------------------------
Acheta_domesticus         ------TSADNKKKR------------IHKSK*L**RKA*NAK**CNK--RPYSNK*ILNLRNN*RTK*QCN*FRTTK*KCKNTSHMH*KCRKHNTPNKRSHTINRKWIHT*NHRRFGK
Gryllus_bimaculatus       NRH*WRNTSDNG*KSTAYLLTPPDKLCCFSKEEYIGNRPLK*EKY*GTKPHKMFFQKIAFTFLKNHK*----NYFVLDNVKLNNITKIIFSNSAIT*CNLNGHTIQQFFSYQKYQQKTCC
Gryllus_longicercus       ------------------------------------RAMK----------------ARNILHSY*L----HQINCDAFKKKNILEIVAYNR--KSADEQSHT--------KCFQKNST
Consensus

Teleogryllus_occipitalis  ------------------------------------------------------------------------------------------------------------------------
Acheta_domesticus         NISKK----------------CN*IPYS*KIL**K*CALLHILYTIRKKLISC------------NKKCTRN-------------------------------------IH*GRDF*
Gryllus_bimaculatus       STSQSLLGNI*WRVSNFALNLKNQQVPAI*E*VK----ELKYLFFLIKSQMLLCFWIPS*FVGT*RNHRKLSRNQSLHVFTSVSVLIGNK*YMC*IC*EYYTS*MYNITTRWAYRGRN--
Gryllus_longicercus       HISQE---------------------PQIK------------FFLIRQ------------------------------------------C*I-------K*YNKT-----------
Consensus

Teleogryllus_occipitalis  ------------------------------------------------------------------------------------------------------------------------
Acheta_domesticus         *IRKPKLSNKK--VTRLTNREQIPFKVVALELANSIE--------GKEIFKL--DKLLHCIVKV--ELRRKPQGPPQCKKCFIH--------------GHTKNYCTMAPKCGYCTQNHW
Gryllus_bimaculatus       -VMDPCLENCKRGA*LFSNVNWFYLELNTIEI*NNILY**YETCCNI*LYEISCN*LQSAIVSCMNEFSESHKG--NTLSLFLHFEII*TYKRLTK*SISFQK*Y-----SCLFGITF*W
Gryllus_longicercus       ---------------------------ILKIRNNIV----------------------------*YKRTH----NTQVLFIP------------------------------------
Consensus

Teleogryllus_occipitalis  ------------------------------------------------------------------------------------------------------------------------
Acheta_domesticus         T--------KDCDLLLSTNPPPPICTLCKGNHMATSDQCKYFPKRKKMQQQSQLNNQYQ-TLQSQNIYE-NKTNSQTYQQQTHINHNPLHTPMTFNNRQNLPTPTYA-----------Q
Gryllus_bimaculatus       IYFIFGIFWRVC-----------------VHV-----CVRVSVRGFILIILHF*KQALG*LNSLSL*SCTKCNS-----------------SLLWKQIT*CADYEGYICFSKVTC*PD
Gryllus_longicercus       ---------------------------------------KIPTKNLLFYIATIIKEYL-----------------------------------MAGKQF--CAESE-----------K
Consensus

Teleogryllus_occipitalis  ------------------------------------------------------------------------------------------------------------------------
Acheta_domesticus         TVSQNNIQS-------NITQILM-QMLP-------TIVNQIIA---TIIPQIQ-------SIVSNQISN------IIRNG*AAQLKIYNTSKLECKWIIK---------------*KKHT
Gryllus_bimaculatus       SFNLSNKHA*YHKFYDKVSYLFLVQVIPLTYYFN*TNVNYFISKNYFIIEII*NGFIYFVSYICKMISSF*T*W*IFKN---------ICEKILGFCICQFIVVMARGKDTRI*L*KPEI
Gryllus_longicercus       SVGTSNMRIG------EITQLFV----------------------FLIK--------------------------------------SQMLLCFWSLQFLCLL---------------
Consensus

Teleogryllus_occipitalis  ------------------------------------------------------------------------------------------------------------------------
Acheta_domesticus         YSLPKKSFNNNCMHNRNTLSTRTNI*N--------TRI*LLQRRQNM-STCSRRSSH----------------HNQKETGPVINFYPKIE*S*TD--------------------CSE-
Gryllus_bimaculatus       LRQPKKVFQ------------RSNIQ*KPLWHFLLTFLLT*QQCGNIVETNSLRKKHISYICFKISFLNVYILVHHKYTSFMVLNTYTTF-LL*TDFYRFLYYCTTNIILFFILLICSSV
Gryllus_longicercus       ----------------------------------------------EINNIRAK------------------------FVKNT-----------------------------------
Consensus

Teleogryllus_occipitalis  ------------------------------------------------------------------------------------------------------------------------
Acheta_domesticus         -NN----------------VRK*ISKYHISI*TT*KTNQYK*IKAN-ITKFRTYHFNG*FK*QTYQLGLQSEHTKWKK-ITKSFNKFKLPNHSTNKSYSLPEKS*RTTRYLRHCHH*KPN
Gryllus_bimaculatus       FNNFY*FLKLSLFCVIYLSIQK---------HLS*WYNYF*LVN*K---NYLSFEF*KL*MKLLCVIGLGALCSVLKKHLMK--NKRNIYSELT----SIP-RSAEVTVHLWQCH-----
Gryllus_longicercus       ---------*PLKCLI---IQPA-------VHTEAAM*KDP*ENCKRVTNFLFFEFSNI*MK*LSVIGLGALCSMLKKHLMK--NERKIYSELT----SIP-RSAEVTVHLWQCH-----
Consensus

Teleogryllus_occipitalis  ------------------------------------------------------------------------------------------------------------------------
Acheta_domesticus         SYPLCSF------FT*IRL*SLSCN----------------------THFRYVL------H*SRIAP*YKYKKHRLDKI---------PNKC**RPSHK---------PIHEKQRRD*F
Gryllus_bimaculatus       ---IIDFQKIFVIFIF*RVHIASCSYIRNCFWRLFFCFNCLCF*LLDITG*RLILSK**TWH**KII----FKKNG--------------------------------------------
Gryllus_longicercus       ---IIEFQKVFVIFIF*RIHIASCSYLRNCFWRLFFCVNCLCF*LLDITG*RLVLSK**TWH**KII----F*KNRIKRKCL*LQR*NRSNNC*LVQGSKIPEFQRILTPFH-------F
Consensus

Teleogryllus_occipitalis  ------------------------------------------------------------------------------------------------------------------------
Acheta_domesticus         CS-*YPH*NYPK-------------------------------------------RSERGKY*S*S*T*IN-------ATNHT--------------------C*HRKINSKETQ---
Gryllus_bimaculatus       ------------------------------------------------------------------------------------------------------------------------
Gryllus_longicercus       CKCTFERSKYKETGLR*IKFRYMCTC*CA*GRRASVRTCKWTDVK*ADVISAYLEVSKCKRGRCGCESSLSMNRRGSRQSASEQT*EYM*NE*ADLKEDALRGRKCKSRRSTSEQT*TLI
Consensus

Teleogryllus_occipitalis  ------------------------------------------------------------------------------------------------------------------------
Acheta_domesticus         ---------NTTAMAAIQKPRF-----KESAK---QTNMPSK-----NNTR*TQNPTI*NHIKK*TLKTTTY-GRLQNKY*KNQQLYLS*RN*IKKLQQQKRSVKY*RNISKK-------
Gryllus_bimaculatus       ------------------------------------------------------------------------------------------------------------------------
Gryllus_longicercus       *YE*ADVKVNVI*ASSR*KWAFVRLSRRESGRTYDQVDMSAKLLSSGTA*RRARKQV*CAVVRNDPLPMSFF*GRFLTTW-----LYSAHCQCF*M*EEHVRAIRTHTIIHARGGIGTRG
Consensus

Teleogryllus_occipitalis  ------------------------------------------------------------------------------------------------------------------------
Acheta_domesticus         ----------LLL*IQLQIITTQSKLLIMK**QFHIQ*NIYITK*NIPNN*TPA*EKISW--NRPNSYHNLQKLNKKTIGYLTAIFNSMLRIGYYPTKWKH----SIIIPIPKP------
Gryllus_bimaculatus       ------------------------------------------------------------------------------------------------------------------------
Gryllus_longicercus       PSSRGSCGSSALATSAIQAGSAEFFFFFFKVASTSRPLTTAIARNGMGTVRMSRYDDDRW*KDKPNS-------------------TMPEAGLEPAVRDRYVTTALTMSAPRPAVSLLF
Consensus

Teleogryllus_occipitalis  ------------------------------------------------------------------------------------------------------------------------
Acheta_domesticus         --------------------------------GK-NSAYTENYRPISLIPIIAK-------------------------VIE--------------------------------KLILKR
Gryllus_bimaculatus       ------------------------------------------------------------------------------------------------------------------------
Gryllus_longicercus       PPALDEVAGGRAGGRVAVRALGDAFSSPAAAEGKGEDAPCSATRPSPLPPSPLKLHLLRDPQRGLLRDSG*EVLNPSH*IIETKYLDTRECIYLLSPYLTLNVKQNCNSI*CGCRIIITS
Consensus

Teleogryllus_occipitalis  ------------------------------------------------------------------------------------------------------------------------
Acheta_domesticus         LLKYINKY-------HTY-----------HISN--------LDLKKNTQR-----HTNF*ESLN-------------*YTKV*TKKAFSSS---------------------FLGRNSS
Gryllus_bimaculatus       ------------------------------------------------------------------------------------------------------------------------
Gryllus_longicercus       ICAYI*IHFICTEITHLYLKIHCTNFTFVHVSNNNVACFVPMKMGKVTRNLCDVIYVLVYNMLNILFVLYSLW*NYLSYTTLL*IEIYQISQGYFNIVKPG*HPID*CLISKLTVGQNIV
Consensus

Teleogryllus_occipitalis  -----
Acheta_domesticus         F----
Gryllus_bimaculatus       -----
Gryllus_longicercus       HKQYG
Consensus
```

**Tocc.2G0021750.1_exon2**

```
Teleogryllus_occipitalis      ------------------------------------------------------------------------------------------------------------------------
Acheta_domesticus             ILLLYFLLKTVFYIYNECNRHFISIFRKCFCEI*SFNNL*A*IIACKNENY*VFCNFLKYIAMEFDHCNKLNEIKCLIILCFIDHPEGIQ*KFRRYLSVLLSFLHEFITFNIF*DTISYL
Gryllus_bimaculatus           ------------------------------------------------------------------------------------------------------------------------
Gryllus_longicercus           ------------------------------------------------------------------------------------------------------------------------
Consensus

Teleogryllus_occipitalis      ------------------------------------------------------------------------------------------------------------------------
Acheta_domesticus             KGRHQSSSACGIQSFLSLLQILLTFNTKNNFVVF*FSTKLYFSLPRNIYTT*YSVYIQRNSLMYFFLLSLDVYFNYAR*IFI*LASRITLDEIKYIKP*IFSCEF*NHYRCIKYQNHDKC
Gryllus_bimaculatus           -------------------------------------------------------------FLYVLLL---IYFN-----------------------------------SIKYQNEDEC
Gryllus_longicercus           ------------------------------------------------------------------------------------------------------------------------
Consensus

Teleogryllus_occipitalis      ------------------------------------------------------------------------------------------------------------------------
Acheta_domesticus             LIVYGFGAKLFFSMPN---CEIAL*TL--*I*RSSKQWRGQGPKTIL---KDI-NQDDDAENPRTHTKGSRLQRYRYKKFVL*QEM-----------Q*NNAKIF--CCTRKRFETHHLR
Gryllus_bimaculatus           LIVY*LGTKIFFSMPNELPLNIVYLTLQ*TMDRANTENNSQGHK*RC*KSKDTYKR*QNSKGPKEKSGETSIVEEQSKSILLHQQKI*NTPLRIMARDMNYQKFFRSICTNQ--------
Gryllus_longicercus           --------------------------------------------------------------------------------------------------------F---------------
Consensus

Teleogryllus_occipitalis      ------------------------------------------------------------------------------------------------------------------------
Acheta_domesticus             **QEISHNFS*HLYRSSKMLK*Y*FFCNKE*FFAV*TTLLKLVSSIKIRIHF*CSKANFKIVIKLLTFLHIFTFIQ*SYITYKCHC-CDLIELYGIRLLFPIQVYKTFVFNFCAVVEFT*
Gryllus_bimaculatus           --LEIILNF---LKR----------------FFGF*TILLKHFSAVKHKIDFLSSNAYVKGVIKLLQLLHVF--------TYKILC**NTVSEYFIPQDITINLNK--ILNFL-------
Gryllus_longicercus           ---EIILNF---LKR----------------FFGF*TILVKRFSVVKHKIDF*SSNAYVKSVIKLLQLLHVF--------TFKLFC**NTVSEYFVPQDITINLIE--ILNFL-------
Consensus

Teleogryllus_occipitalis      ------------------------------------------------------------------------------------------------------------------------
Acheta_domesticus             F*QYCDGNSEL*AKMLVAHKVQYSSTK*LFPITR-------ESMCSCSTTRFF*DQSHAAAECKNGDLSHANCPSSSFYHPLPCTYTD*TNMQGRPDWTLEIDPSGAVRCWGRYLEERNP
Gryllus_bimaculatus           -------------*MLPLHKLTG------YPILRVLSGEEF*WLTALIGSFFD*DQANAVAVS*QTSIYH-----ERFVPPHPCP-----------------------------------
Gryllus_longicercus           -------------*MLPLQKLTG------YPIVRVLSGEEFLWLTALIGSFFDQDQANAVAVS*QTSIYH-----ERFVPPHPCP-----------------------------------
Consensus

Teleogryllus_occipitalis      ------------------------------------------------------------------------------------------------------------------------
Acheta_domesticus             ATR**RPDHMF*WDTDTVTSRNVTCTTITGLMFKAVHASDLLIE*GFRRVIRFLSSLHAFFLGR*RRSPLAVWWDSATTCLPSSGILPNQVSPTSGAPPQGACVVV*CMLVQECV*EERI
Gryllus_bimaculatus           ------------------------------LIFSYTH-------------------LH------------------------------------------------------KV*YHSRT
Gryllus_longicercus           ------------------------------FISS*TH-------------------LH------------------------------------------------------KV*YHSRT
Consensus

Teleogryllus_occipitalis      ------------------------------------------------------------------------------------------------------------------------
Acheta_domesticus             GQGRGRKDRARSVARLLGYCSGIRLVFIGESTENHRSEQTAARILLQTATEFD------LAIQLVSKVLPPSGHEPRTF*DRSNAVSSV-------ECPEPPLRHTCTDYINIHDGREF*
Gryllus_bimaculatus           GKG--------GSGH*MS*CSSSD*EVLG----------------LQCGRIFDTS*IAKLSPVAVWKIILTAGVSDQ*F----SVVSSINTG*MPGQCPNKRAR-VCTLYI---------
Gryllus_longicercus           GKG--------GSGHQMSQCSSSA*EVLG----------------LECGRIFDTSWIATLSPVAIWRRILEAGVSNG*F----PVVSPINSG*MAEQYPNKRAR-VCTLYF---------
Consensus

Teleogryllus_occipitalis      ------------------------------------------------------------------------------------------------------------------------
Acheta_domesticus             SLSELGL---CPAGQTSRY--KHHKMAYRHRI--------------------------------------------------TLFGSGG-----------------SLPF----------
Gryllus_bimaculatus           ------LPPPCPIVTHSLH--KLYALMH-HTI--------------------------------------------------TLLGAKNLPRLGLVSR------*WEIPG----------
Gryllus_longicercus           ------LPHPCHIVTHSLHTQRSHAPHY-HTVGS*ELTPAGVGAPLVGDTWFGKIPTRRRRAAPYLIQRGKVAATFREGNMHCTYGTNGLSR---CSRCLSHQSINQLPYMSENQTGT*R
Consensus

Teleogryllus_occipitalis      ------------------------------------------------------------------------------------------------------------------------
Acheta_domesticus             --------------P*KSLQLRN*CPVDASGKQNN------------------*SINQWLHNPVARTLDGRPHCADAPCAGGVPPCVAVSR--AGGRGGLG------------------R
Gryllus_bimaculatus           --------------SGKS-------PQDDVGQRRI------------------SSSGRRSQLPFERE-----------------TCTALTRLMALAVDALGFISQSTNYHTCREIRRELS
Gryllus_longicercus           SSRLGRSTARRDNWKGKSP*VGGR-SQEIAGEEH*HFQ*IQHAIT*LVPLAPAKKISQWMHNPVARTPQSRPH-----GAGRSAPCGGV----AGG-------------------RARR
Consensus

Teleogryllus_occipitalis      ------------------------------------------------------------------------------------------------------------------------
Acheta_domesticus             GGPGAGRGRGGRTHGRAAAPCARRRRLTRASTWAA-------IRRARAR-PRPQPPS---------------------------------------------------------------
Gryllus_bimaculatus           AHPD*GGARLGEITGKEKARKK*A*RL-RSQVIAGQEH*LFHLYNTQLRD**PKKKSVXXXXXXXXXXXXXXXXXXXXXXXXXXXXXXXXXXXXXXXXXXXXXXXXXXXXXXXXXXXXXX
Gryllus_longicercus           PRPGRG----------------------RADARAG---------GGPVR-PPPPPPNACFHPRDYKTGASPT------------------------------------------------
Consensus

Teleogryllus_occipitalis      ------------------------------------------------------------------------------------------------------------------------
Acheta_domesticus             --------PAV-----------GPSGHTLLCHGESARRCARPLYSKDAVSKH*CSSSMHFTSIFGIPCFLIYMFCVRYFYF*MKKVVS**NAVMRFRYQFNVLRNQVWCG-N*N*AFKVY
Gryllus_bimaculatus           XXXXXXXXPAS-----------HRSVDTLLCHGES----ARPPGGCDTDVQVFMESLMDFTLNFLCTDALCSIFLV-----LIGKITNLTNFSNHFHTSIEHFPDKSYLEVKLKFMF*SP
Gryllus_longicercus           --------PATASVASVAPLN*HPPLPW*VCKAPRR---PRPQGDATPMSRFLWDH*WT*L*IF-----FASMLCVRYF*F*LEKLPS**SLVIIFTYQLNMFQTKVP*GVN*N*RFEVH
Consensus

Teleogryllus_occipitalis      ------------------------------------------------------------------------------------------------------------------------
Acheta_domesticus             TIFQITIPKKK---------GLLNFLIRFYSILQVTSQ*TRSSSG*PSR---------------VNING------DHYSLRTSMVWGRHFVNI----VALISCACFLGDGRVLISSSRHY
Gryllus_bimaculatus           YIF*NIYSNDK---------VLLNILI----FV*LVKSSIKN**TLPSAK*CPLCAHF*SIF*LKNIDGFRVMFSKHLSILNIDV-SRH--------------VCSVDDVVSLFSSSRHF
Gryllus_longicercus           TFFEISIQMIKYC*IY*YLFS*LNLL*K---LINATFGQIMS----------PVCT------LLINI----------LIKKT*MVFG*CFLNIYRF*TSMFRYVCSVDDVVSLISSSRHF
Consensus

Teleogryllus_occipitalis      ------------------------------------------------------------------------------------------------------------------------
Acheta_domesticus             VVF*KKLRCNFYFYFLG-F*GAEYVS--------------------GISG------KYLLWNL------------------------------C-LNKKHQKIFSFLLF-----------
Gryllus_bimaculatus           EVS*NKL*CSFDFLLLDVFGGVKYESRTTYIKNRIFVRFLTTSKNYRI*GKHYSLPKYFMSQIIFDSTFMHQ*L*KKPGTNVLSFG*YFKPIKCSMSSRY*KMLETLLVLRILIINIT**
Gryllus_longicercus           VVR*NKL*CSFDFLLLDVFWGVKYESRTKYIKNRIFVRFLTTSKNYRI*GKHYSLPKYFMSQIICDSTFMHH*L*KKPGTNVLSLG*YFQPTKVGTSS*Y*NMLETSLMLRIFIITKT**
Consensus

Teleogryllus_occipitalis      ------------------------------------------------------------------------------------------------------------------------
Acheta_domesticus             -----------ACIKQLAYSGKQV--------ILHQNIKES------------------------SYVP--------------------------------------------*KPPQYF
Gryllus_bimaculatus           NNFWPVHLRVETCTRLYKYQYIKISFFR*LILVITNSIMWSEQF*V-------------------SELPTLVSRCLGTRIGSLLL*RKK*KI*LIIL*WAG*VIH*PMFTCALKTPTNTF
Gryllus_longicercus           NNFL-----ARPPKRRNVHKIIQISIYKNIFL*IINFSYHQFDYVVRAVLSLGTPNSCISLPRNSDWLPILVKEEEENLINHFMMDR---------LGYSLTEVHM-----HLENPHQYF
Consensus

Teleogryllus_occipitalis      ------------------------------------------------------------------------------------------------------------------------
Acheta_domesticus             LFNSSKIFKRIRTKIASMNLFSEVFMVFEDA**ISP-------PTHKLQFRSL----KYPQKCFLGNSLHVLK*PGKNLFTTSRQRCFKRHPTCGKTMVQKMYKCE*QTDT-----CKLH
Gryllus_bimaculatus           SF-LRKYYKRIHTKIVSIRLVR---------Y*ISPWMPHKYLPSGTTNLKA*----RI-QEVFLRSSFTMFYKDSEKMSLLPVPRCFECHPTCCKTVVQKMYICTGLTHAN*TIYMGIH
Gryllus_longicercus           FFISSKM**K-NSQKNSVNSSCEVLIISVDDS*I-P-------SSRNQQFRSLTYSKRFPHE-FLYNVL*GF---GKNVS-----------PTSTEML*MSLNLLQNSGPED----V*LY
Consensus

Teleogryllus_occipitalis      ------------------------------------------------------------------------------------------------------------------------
Acheta_domesticus             DLHDCMY*NSDVLSVFSHFDQKLSKRVITKTLKNIKPSINYIRV*KDSCS*LRKFG--LKDLL*KRLFF-FLLLPGVLVTF---------G**EGRGGMR--------------------
Gryllus_bimaculatus           RLFRCA-------SIISSFDQTLLS-VTTKLF---------------SCS*LRKFG--LKGVLSKRLFF-FFASSGSGVQIVSLGG*GGVGRAEGRPAQRLHYLPISWGARYCKRP*ELQ
Gryllus_longicercus           RADSCKL--AFTWVYVDYSDVLLLFRVSTKLFF*LRP----------NSSLVSN*GNLV*KVFYRNVYFSSLLLPGVVFK*SRWADEEG*GRAAGRPAQRLHDLPISWGARYCNRP*V*Q
Consensus

Teleogryllus_occipitalis      ------------------------------------------------------------------------------------------------------------------------
Acheta_domesticus             -----------------RNV---------------------------------------------------------------------------------------------------R
Gryllus_bimaculatus           *RWYIISSVKLTASEGEERNVTIVYVENSG-------------------DVMSI*LSLLTDTLLNW---------------------WSKL*IDI*WIITNETIEAQTKKPAIKRS*DEK
Gryllus_longicercus           *RWYVISSVKLTASEGEERNVTIVYVENSGV*CQFN*AY*LMPY*IAGRNCRSIYRGLLQMRLLGHRQRSQTLREVEIRFAAYLVFAISKIDPNLSFEMTHLNITLSVNKGQL------R
Consensus

Teleogryllus_occipitalis      ------------------------------------------------------------------------------------------------------------------------
Acheta_domesticus             GQREARSSN-------------------CKI-------VLFRFTRVRPLPI*N*YQRFVC------------------------------------------------------------
Gryllus_bimaculatus           IRRLPRFRNK*DRPQSFLRNDAFGH*VKCK*RTIKNLQVMFRFM*FV*R*KLNGYTILFCKKVF**IFHWLLILLF*GSNI--**K*FLLNF*QFQPLTSHVT-YLFSITAISLF-----
Gryllus_longicercus           ICK*C*DSSNL-----------------CKDRK*TN--IQFYFAR----------------RYFSEVFPWLLILLF*GSNI**ASR*FLLNVLTISILNIACNRFVFNYSNIYIYKDSFW
Consensus

Teleogryllus_occipitalis      ------------------------------------------------------------------------------------------------------------------------
Acheta_domesticus             ----------*CFYF----------------------------PVYVRSNIRSVGKVMKCCAK-KWGILDGNKFI--------------------AGC**R-----KKWIKPVDKLRKKS
Gryllus_bimaculatus           -IKIHSGLFMYMFHFDFIILTNKT**DIITNKTSLLIDNSFSNPIAKGSLLLGL*KSLKLQIKLLFHFLGGQLFLVELVWIIDAHCSLQKEGASGADCGTRLYNV*KL*NTPVIN*GNYF
Gryllus_longicercus           IVYVH-----VSFWF--VILTNKKLC*SIIR---------FPPLLQKEGWVLGL*KPFK--SKSSFSFTFG---LVNYFWRSESG*LMRTVPPQGGCI*CRLWDSIV*CLKTVRHTRH*L
Consensus

Teleogryllus_occipitalis      ------------------------------------------------------------------------------------------------------------------------
Acheta_domesticus             SKTWLE-----DFS*-K*IRKIRR-------RFPKR*SQGEIAPY--TLKER--FDSK*LSSYS*SSPYAI*ICYKLLGHLSGQ----------------*ISLF----LLKYVGNVIN-
Gryllus_bimaculatus           TTNTFE-----*FV---KIRELRKS*RFLEVRTSVNTSTEQIAQFLNAIK*TYVFHTHVRVQ*QQSN*FALL--EAAPGFLETENFRQEADSIKDDQKRVHIALLNN*NTLKCKINVTNV
Gryllus_longicercus           RKCIFRPTHSYDFSKYENCENLRD---YWKIRTSVNTSIELIAQFLNTIK*TYVTHTHVCVQ*QQSN*FDLL--EVAPGFLENENFRQKADCIKDDQKRVHIALSNS*ITLKYIMDFTNI
Consensus

Teleogryllus_occipitalis      ------------------------------------------------------------------------------------------------------------------------
Acheta_domesticus             ----------------------EIHMLTF*DRVPDL*ECLALI*LE*ALS*LSELKPSRPTLQKLFVVETGQ*KQKNIIASEQIARQFLC----------ASACQTLMQKVLPTN-----
Gryllus_bimaculatus           YGI*SVKDYHFFQLNMRAMSKMKLHRLSF*YRAPVSKVY*ALIESERALSLLRNL*TSWLTLRKFCVVETSQ*NRKKLP---------LCLNRSLLRVRPSANQQCRKCCLPIR-----
Gryllus_longicercus           YGI*RVKDYNLLQLNMRTMSKMMLHRLSF*YRAPVS*VYLAFIGSERALSLLSNL*TSWLTLQNFL*WKQISKTENNCH----------C-------VRTGRS*EFVR--VPTRSVGSVA
Consensus

Teleogryllus_occipitalis      ------------------------------------------------------------------------------------------------------------------------
Acheta_domesticus             -QGL-DVRNLMSTILFVITHHTFIV*NIC-SSLQDLEIFKINYEPL*KLSSAV*LIDGSVSDNCCH*ASQ*SECFIPRRGYVPPR----RFVRSDADLHDGLQGVSLSRSRHPPEHRACL
Gryllus_bimaculatus           -GSM*EIQRL*FCYMRFPTTYSLCKM*LCLTQLHDLQVFNIHSEYR*R------------SDVC------------P-----PP*DDEIRRQPSWLAVYFPLQASSPTGA--PPQRGEAL
Gryllus_longicercus           YQSGAQCKKFNIC--------NFVI*NFS-QHIHCAKCDYI*LNYMT*KFSIFIL--NTVSEVN-----------VP-----PPQDYEIRRRPTWLAVYFPLQASPPTGA--PPQRGEAL
Consensus

Teleogryllus_occipitalis      ------------------------------------------------------------------------------------------------------------------------
Acheta_domesticus             EKL*SSKLPTRIPTCKIV*YVCLWV*GERIGQIKGRNCYTHSWPDC*DIALAFVWH*TKEQRTITDQNTRPLEFSS--ERRLNSTLQSF*LSKICLELYPKAFWSESLV--SQG*PWL*T
Gryllus_bimaculatus           SSRPVSPLV**CSTCK-------W-RENRTEQRKER------------------------LRTIL---ARLVGYCSGIERRIHGKLSIRAPNSMILENAPELVR*GTLV*ISADISMLAL
Gryllus_longicercus           SSRPVSPLV**CSR---------W-RENSTEQRKER------------------------LRTFL---ARLVGYCSGIERRIHGKLSIRAPDSKILENAPELVR*GPLV*ISADISNVGS
Consensus

Teleogryllus_occipitalis      ------------------------------------------------------------------------------------------------------------------------
Acheta_domesticus             NQ----RVPASL---------KLQG*GRKHTSAL*RTLKCVLCGVYTHEWGMVNKRV--KSVTRCLS*YRMSLLHASLCLRIGTLQVRVLSTEIVCSTNCSLKT*TGHASQS**MLKVLS
Gryllus_bimaculatus           N*SASFRLIAGL---------RPDTHGKSLTD-----VKYIRCMVYIHEWGTAHERA--ARCP*C-----------SACLS---------------GVETFVEIWTF*RPSRDARVIVNA
Gryllus_longicercus           ELIS--RFPSH*SCGAKAGYTRAEFNGR*M-----------------HPL*SVHTRMGHGTRTRC----AMSV---VFCLS-------ARRTD-VCG---NMDVLTSLM*RSCYR-*CSA
Consensus

Teleogryllus_occipitalis      ------------------------------------------------------------------------------------------------------------------------
Acheta_domesticus             LHLVSSGLITTNINATFNRIP*FGPVRRDSPGALYHFVTKLDLEQFGS*H*DLD*KECK-------------*RAGAWLTSCVANGVRDELCA*LCAWLTVCVANCVRG*L--CAWLTMC
Gryllus_bimaculatus           LHYIF--LIIISLLPTILSREKASYIVRF*IYKLYNLLVTIFV------HFSLFPSFC----------------------SCVLS-LHHKFSQPLLCFIYVITFLVISHLLLICEFYFI-
Gryllus_longicercus           LHIIYYYYYYYYYYYYYYYY*PVTHIPQ*NKGLLHCKVLNLQAR*LVS-YYFLFTSPCTHLFVHVYFHFTNTFRNH----FCVTYM*SHFWSSHICF*FANFISSIVEDFF---------
Consensus

Teleogryllus_occipitalis      -----------------------------------------ALLHVLVALACACAAAQASLIGIGGGGLGFGG-----------------------------------------------
Acheta_domesticus             VANCVRN*LRACL-----------VWRGA-----------QALLHVLVALACACAAARASLIGIGGGGLGFGGGE*PIKPPLSSPHTVCARAVEM*L-----------------------
Gryllus_bimaculatus           --NSRRLL*EKCM*ILVLLVAGGRISRGGARVGELMTCGLQALLHVLVALACACAAAHASLIGIGGGGLGFGGGE---------SQLCACAVEMEL------------------------
Gryllus_longicercus           --------KKTCMQMLVLLVAGGRFSRGGARVGELMTCGLQALLHVLVALACACAAAHASLIGVGGGRLGFGGGE---------SQLCACAVEMQLIQLN*IDALFYVARLGP*GPLFR
Consensus                                                              **************** ***** *** *****

Teleogryllus_occipitalis      ------------------------------------------------------------------------------------------------------------------------
Acheta_domesticus             --------------------------------------------------------------------------------------------------MKILYLPTLLFIEIAAFDSLAI
Gryllus_bimaculatus           --------------------------------------------------------------------------------------------------MEALHLSAILLMEIAAFHGSII
Gryllus_longicercus           PTTQISLRSSHQLAEKGGISWARNRRFEFSSEASFRLEDLLHASKPTNMGQPALLPLRRKAYEGFFRP*NGPRLRSGLNPPTWVQQSSTLTVTPSPPLMEALHLPAILLMEIAAFDGLII
Consensus

Teleogryllus_occipitalis      ------------------------------------------------------------------------------------------------------------------------
Acheta_domesticus             TVSFLLFESQEST---------------------------------------------------------------QVMIFWLPNSRKLMMAKS--------------------------
Gryllus_bimaculatus           FISFLLFGSQKMNKYYFLAFKQK---------------------ETEDDDDDDD*H-------KTDDRGPAVWKRLQMVIFWLSNSWKLMMSKPSNTTISVNNILGNDNASNYYIWNANN
Gryllus_longicercus           FISFLLFGSQKMKTYSFFGFQTKGN*RTR***LT*N**TGSCCLETT-----TDGHFLALKLLETDD-G*AI----EYYYFNKQYSW------------------HDNASNYYIWNDSD
Consensus

Teleogryllus_occipitalis      ------------------------------------------------------------------------------------------------------------------------
Acheta_domesticus             --LKDLKDFF----------------------------------------------------RVS---PVSLALYFSKICHKSPQSLLIKVMHPFPVITLRLRDHKEYHLITRV*IAEYS
Gryllus_bimaculatus           *LPQNKK*WFTDVWLLLVEEKKTTLGATRDVVLRSLNFHDWVSGFISRSIHQGFGWLKHGCGRVSRGFPVYVSLHFSEV-FRTP--------------LQTPTHQNHTFTSRKDLKALT
Gryllus_longicercus           *SPQNQQ*WFTEVWILLV-EKKGTLGATCDVVLSPLDFHVWVSGFISGSIHQGFGAVKAGL-RQSFS-------GFSGSCFSSF--------------LQSLSH---AFTDSHSSKSYT
Consensus

Teleogryllus_occipitalis      ------------------------------------------------------------------------------------------------------------------------
Acheta_domesticus             CGNCLSNKCNCVRFSQDNFVYLALIP--FFALS---F---------IWYLHY*LCIR*LIHFM*DLVIKNIHLRRSS------------------NCAEMIRLLNLLF------------
Gryllus_bimaculatus           WGP-----------------Q*IIPSGYWNLNPKKF----------LWNV**KGGIR--IGFLE*LSFPLPLMPRMSR-----PVRTL**IMYILDF*ENHIEVGKLFNLFYLVSR----
Gryllus_longicercus           ----------------------------HFQKRPEGFDVGTTINNTIWLLEFK--FR---KILVELMIERGYSYRFSRITFVSPTSNATYVTFCTDVVINNVHL-RLLREPY*SGKAAQF
Consensus

Teleogryllus_occipitalis      ------------------------------------------------------------------------------------------------------------------------
Acheta_domesticus             ------CK---CGSNPG-----LSMRDFW*RKRVHCRIS*VFLVSQNLFSPLF*QYSYHVFKIT*P*KSSTLFQIYPKD**SALLR----R*DDTICT---RSICSVAVRVLAPVEFLVN
Gryllus_bimaculatus           --FKFWCVHMICGEGSGFAAGFLGFSRFLCISFRHCS-SKIHIIS--LSSSIPQNHSL----------TSR*DQHNKDDMAS**R*LPYRLLFVICCEADRAL*SFCI----------
Gryllus_longicercus           IPLGIAVQILVCPYG-------L**RKWVCGRF-----SWVFTVPLHFIPSLLQKNSYHIYVIIYP-------SIH*LLDKTSIIKMIWYHSEDNFLI----AYYS*SV---AKQLGHCN
Consensus

Teleogryllus_occipitalis      ------------------------------------------------------------------------------------------------------------------------
Acheta_domesticus             KAD*LEIILRFSHFPLLLILPLIYYPHHIIII-IFSLNPDIY------RKHHPRHIK*L-----SLQMKRL*HGYDVLL*DIFEIAYILHSILILLVG*R**KVILQFCCLNVLKY--*L
Gryllus_bimaculatus           ----MEGFLELSHFPCMLL-------HNCFIIHIISSRSSIH*IS*YIQKTSPSSL*RTSIVNGSISNMALV*FYETL*NVINEL*W*DSRDIVML--------LFKYCKI*INYY--LL
Gryllus_longicercus           PIVIMEGFL*IFSFPLHVISQLLYHPHYIVTI-IYKLNHLIY----------PGNIPIVMVKNFYCQLKCQ*HGFGIIL*DIIKF----HKCIVVV-------**S*YCHVAV*IF*NIN
Consensus

Teleogryllus_occipitalis      ------------------------------------------------------------------------------------------------------------------------
Acheta_domesticus             IILFSYVKISSTG*SVV*HVI-NIFGKTIFSNKMTKTM*M*RWKR*V*MMMKHKNK*Q*YLF*TCVNSDVEFAEKYIMK*LLH*FLKFKS*VF--EKSRYSLFA-RSYKVQKCAAVTRIS
Gryllus_bimaculatus           ELKF----------*VRDKV*CNMQ*LSLKSNRL*NT---------------KTSNK*CLF*TC*RTNVEFPQSKFLK**LHYVLKFKS*LLVK*KSHGALFIFRASTGQLCELEFHL-
Gryllus_longicercus           *LLFT*IKIPSTR*SAM*HVI------------------IIFEK--*QMMKHKNK*QVVFILNVLMNKRGICRENF*N---NDYIMF*NLNRSYKSKSHGALFIFRANKGQLCALEFHRR
Consensus

Teleogryllus_occipitalis      ------------------------------------------------------------------------------------------------------------------------
Acheta_domesticus             -PTVSYNEILQSMNVISLA-ISD*HSSLHSRSQTQYACI------TIFPGNMEKRFI*LGHSLCQILQQHCPNEHEQFHNWTYVKYTYNPETRLAC*DTLNSTITSSSHPQTSGVLV*CA
Gryllus_bimaculatus           --IISYYEILQSKNVISFGNF*LYHTSLHHRLHTQYACI------TISSGNMVKRFI*LGHSLC------------------------------SDSETLLQ*VRAIS*LDLCEIYL*CI
Gryllus_longicercus           FHIMKYFN-LRT*SLLAISDSITLHCTT-DRIHNMHA*LFPRETW*NVSFNWAIR--------CVQTLKHCSNEHEQLHNWTYEKYIYN-------------------------------
Consensus

Teleogryllus_occipitalis      ------------------------------------------------------------------------------------------------------------------------
Acheta_domesticus             VGTDEQSCIVSKS*FQFPGFFTSITSFWYNLTELIDA*KNDEDLIFVELSEFSKN---AKILQINGM*NQIRLLNFHIKQKKIT*L*LIS*LRSILYITTS*NDLTYYGT----------
Gryllus_bimaculatus           ELIDMLR*LVN---LKFEYFKIIILNT*Y*I----------ENIFKVNFNFFVIN---LKIYFNKNNCHMIEDS*F--KEFRYTT-------ESLFYSSN*NN*SATLNSKQA*M*WKLL
Gryllus_longicercus           ----AWNCLVCYGYL*I*YLN--ILK*LY*ILNTI--*KIFSKLILV-FSSLI*KYTLIKITVI**KIHDLKNSDI---QRRVYF---IQVTRTIKVLP*IANRLEYDGN-----YWVYY
Consensus

Teleogryllus_occipitalis      ------------------------------------------------------------------------------------------------------------------------
Acheta_domesticus             -----------------DAVVFIILLLLFNCLDHMVILA*EVC*NKLLHCNYQIVLQL*T*NLISCT*KLRILSLESKKMHIL*PLRFEG*SIYGLC--PAGGIGLGGGGLGLGLGLRR
Gryllus_bimaculatus           --GIL*R*HVTSGKQKAPEETMFFNIHFLY*C---------------IYYCLWALKGEISTFCVPQAASAWAAVAWVSA----------------WVCGGEASASAAAWAATGAAWVWAA
Gryllus_longicercus           EDNILLAENKRLPKKR--YS*LFIFLMDVFI----------------ILFGL*SAKYRLS----------------VSRRRH--RPRR------------------------------RR
Consensus

Teleogryllus_occipitalis      ------------------------------------------------------------------------------------------------------------------------
Acheta_domesticus             GGIGFGGGLGG-------------YGGGLGLGYGGYGGGYGGGYGGGYGGGYGIGGGGLG-------------------LGLGRIGLLGGG----LGRRPGPG----RRLGP-ARRRRRA
Gryllus_bimaculatus           TAVATGAATGAATASAAAVWAWAWAASGCWAAAAAWAAGWAWAWAEAWACAAAAAGSTAEASAWARA-----------PTAWAHTARASAT----TANRPCPGERPPRRVSPLPARHRRP
Gryllus_longicercus           PGSRLGSAAGRHRLRRRP-----------------------------GRLRGRLG-PGRLRRWLRGRLRRRLRHRRRRSGLGPGPGYRAAGRRRRLGRRAGPG--PGRRLGP-ARRRRRA
Consensus

Teleogryllus_occipitalis      ------------------------------------------------------------------------------------------------------------------------
Acheta_domesticus             LRRRHRPGQGLLRPRHVRRGLRLLRQIGLVRESGPLEGSLLSQQGPG--ARAGLEPAAGQEEVPRAHGPLRP-------------RPTQGHLAQGLRVQEGGRGGNTRLVLHLRHTALLA
Gryllus_bimaculatus           LPRRART-RG--RPR-----------------------------GGS--TR*CSASAPSNARAP----PPRPSSSGGWTWWKYTTRPSPTTYCSSRRTITRSSVGSLLVRFLPRRRALRA
Gryllus_longicercus           LRRRHRPGQGRVRPGHVRRGLRLLRQIGLVRESGPPHGPLLSQQGPGARARAGLEPAAGQEEVPRADGPLRP------------RPTHGRLPQGLRVQEGGRGGNTRLVLHLRHTALLA
Consensus

Teleogryllus_occipitalis      ------------------------------------------------------------------------------------------------------------------------
Acheta_domesticus             EQLSLLPSIYFWFVFSQGEDH*EHFTSFLFR*ILSGFL*VKNCVCFSNLNYFCSPKELLNDQL*CISALSAYSCVVY*KNTFYPVSR*FICTKPF-----FAY--------------LY
Gryllus_bimaculatus           ---------------------FHIISF*IN--FKWFLESKELCVFQ*FKLFLFAQKLLNGQL*CISALSACSCVAY*KNTFYPVSHCFIFIISYSPTDTFPYSF*SSK*YSGIMKSSLY
Gryllus_longicercus           EQLSVLPSVHFWFVFSQGEEH*EHFTSFPFR*ILSGFLKVKNCVCFSNLNYFCSPK----------------SCLTVSCNV-------------------FPR--------------LVH
Consensus

Teleogryllus_occipitalis      ------------------------------------------------------------------------------------------------------------------------
Acheta_domesticus             FHSWLLYFDIFWINAVN*III--SLLS*HLV--------------------RRGILNS*CVNVFLKYL*I*NTGLFGNKYL*MCLIQLKITHQENNFIIYNLQFRE--K*VTKYDKY---
Gryllus_bimaculatus           *PTW---------RQVKIVV----------------------------------FFNN*LRNIFAKDL*V*N*ECFRNK*F*S--IQFNITYQKNNFLIYKLQFRELIKINTY-------
Gryllus_longicercus           VHVW-------HIKKIHFILFLIALFS*HPILPLTLSLIYFDLRSSILAS*NRPFIDQLC-QA*ISFF*IINSETYLQR---TCKFEIKNVSGINNCDRFNSK*RIR-KITS*YINYNSE
Consensus

Teleogryllus_occipitalis      ------------------------------------------------------------------------------------------------------------------------
Acheta_domesticus             -----LQLQNKFQ--YSLL------CFYNLKL--DFGC*LYSFILDVANRVWTNSRIKTRF----KKVNLNLTECVIYIT***PLYSKLFTLSLKIK*I*CLRCNDFSNI-----K*IT*
Gryllus_bimaculatus           ------YFKNKRKC-FSFLPL*LKTCF*KLSLLTYFTC----FEQGVA-----NSRIKIRFIRGSGQVNVNKTECVIYIT***CLF*KLLTLSLKIKKNILLKNNHYS-I-----TEITN
Gryllus_longicercus           DL*R*ILIT*KINVNVSLF------CLYNLKL--VFRS*VYSLIPHVSNRAWPICESKYDL-*GSGQVNANKTECVIYIT***LLF*KLLTLSLKIKKIYCLR------IIPV*QK*LN
Consensus

Teleogryllus_occipitalis      ------------------------------------------------------------------------------------------------------------------------
Acheta_domesticus             VLNQKKTF*NNGRNKYVLYFIVTNN------IHILVDFA------------------TTIQGKKSI*YV-------IIRINGRH----CGV*IEHVL*ILVIF*PELSHLFQTFYQILRD
Gryllus_bimaculatus           Y*IENKN-*NIGRNKYEF*FIAINNKIEFGDTHILVGFV------------------TTI*ANIDIKKI*NSE*ETL*DIN*VRFMCTC-----HILT-------GIDSFVETFYQIQRD
Gryllus_longicercus           Y*IENKN-*NIKRNKYEF*FIAINNKIEFGNTHILVGFCHNNLSKH*Y*KNMKLGVGDTVRYKLSTFYV-------Y*SYSNRN*LI-C----*DILS----------------DSARL
Consensus

Teleogryllus_occipitalis      ------------------------------------------------------------------------------------------------------------------------
Acheta_domesticus             *LFWAGVDRYCRVLVL-----SSCG---PEQPPVIRYQSLLHYSRCRLEILS---------------RG-----------------LLSFSLAATNSVQEGGRGSLTFS*HDFPA*AFLG
Gryllus_bimaculatus           *LFWAGVDILPRSATFKLWP*TTTGQPVPISPPV--------------QQMST*NPP--------PPRG-----------------LLSFSLAATNCVREGGGERRSFSAHDFPP*AFLG
Gryllus_longicercus           ILLGRS*QILPRTDTFKLWP*TATGQPVPTSPPV--------------QRMST-*PPEVYPRFPWPQRTTFEREAGRDVPLERTVLLLKHSSAKSMC---------SFDKN*VKISLLIG
Consensus

Teleogryllus_occipitalis      ------------------------------------------------------------------------------------------------------------------------
Acheta_domesticus             *NYVFFRQKLTKDKFVNRYKPVI*IACYIQERFDVMYLVEVGHD-EEIEK*IL*MFDM*TKVYGIEPRTVQHSKYYIFV*IIIFLLIRQVFGRQVLHVL*LISKA--------SVVITRK
Gryllus_bimaculatus           *NYALFRQKLRKDKFVNRRKPVI*IACYLQERFDFMYFMEVGHDYEEIEKCILRMFDM*TKVSGIEHRTVQHWKSCIFS*IIIFILIIQFLKRQIL--CSLIESNGFRCQ*CYSTPIKAN
Gryllus_longicercus           SNLLFRSHVISKNVLT-----------------FMYFMEVGHGYEEIEKCILRMFDM*TKVSAIEHRTVQHWKSCIFS*IIIFILIRQFLKRQIL*VL*LISMA--------SAVTNAT
Consensus

Teleogryllus_occipitalis      ------------------------------------------------------------------------------------------------------------------------
Acheta_domesticus             PLLRNMNSNTR**HV-LSMTE---KLLYD*MEIFIYPS*ELLRLECKSWKI*VKPKTKYFVAILLKN-----------IQRTKRYQELIVQII-YAR*RWRHGRHI-LRKFLSRLRSKTF
Gryllus_bimaculatus           P--------TRLWDVALSI**---TL*ID--AIFL---*TLLKNCCECGANFEKHK*NTKISHLLPNFMP*PIVL*N*L*RTINYREQ--QL*HYVR*ITLAVRKINLSDFPGRLRSKTS
Gryllus_longicercus           L-----H**KRIQHV-CGMSRCRYKKLYE*MQIFINSS*KLLRVRCKV*KR*VKHKNKSLVA*FNAN------------ATNCSLELIIRI*-SAR*R*QYVKLI-FQIFP------VD
Consensus

Teleogryllus_occipitalis      ------------------------------------------------------------------------------------------------------------------------
Acheta_domesticus             CLLRQLQNGTAYCT-LRTANSV----------KVFL*LII*IYLICHVILINCILYTINTIVKYSVAYERNTIQKRRMEFGLTGKRFIRNKFESRQFYA*RDVV*IIDYRLFGYSKHDS
Gryllus_bimaculatus           CFVWRLLNEAIYCT-WPTARGVLPQTV**TYICN*LFEIILFVTFF*FII----IVFIININVKYSVAYERYKLQKREMEFDLTGKRFLKIEFEIRQCYA*GDLGSKIDYMLFCSSEHEF
Gryllus_longicercus           CAQRLPTSYGDY*TKQSTARGVLPQTV**TYICN*LFEIVLFVTLF*FII----VVFIINIIDKYSVATRD*ASEVR------NGIRFNR*EVSQK---------------LI*N*TEF
Consensus

Teleogryllus_occipitalis      ------------------------------------------------------------------------------------------------------------------------
Acheta_domesticus             VLYRNDLTH--I*HML-IKEVTH*SRT------------------TF-RQ*M*MYNFFYSHYVCWFPSA*HCTI----VECCPVEC*IITPTT*ITSGVS-----------SI-------
Gryllus_bimaculatus           ALRREE*TQ--I*QLL-IKQGKHYSRS------------------CL*RLHVNVYIFLN*FRGCFL---LHNTVQLSSI*MLDHNANLINIYLDIASGVP----------SVAWSFYYP
Gryllus_longicercus           CLRRSRLKK*IICYFVPLNMILHYAEKIEPRFDNY*LNKENIVHGVVFRESM*MYTFFLK*F-CGL--VFFCST-----------SPIKLYLNVGS*RQHHKYIFRHC*WSVVWAFSYP
Consensus

Teleogryllus_occipitalis      ------------------------------------------------------------------------------------------------------------------------
Acheta_domesticus             ---------FLTRSL*KRKRNHSYANNIDLINIIVLMNMLKL*K--NVIFFHFIMQSTALKIFLILSRSQTI-----*N*P------LKQTKVNIESRPE*PRWQVHRPTQRLALKGHGC
Gryllus_bimaculatus           IAVKMKKI*FLLK*H*LNK----------LIYITVLMNFIKFLKILSE*FYFI*LWFRDLKICVILSGSRIIFFITYF*MPQ*Y*KI*K*DTIYL*NR*K--SW---------IFSFYTY
Gryllus_longicercus           NVVKKK--------------NSYANNIVLIYIIVLMNLIK*WY---K*FSFI*LCSRELKICLIWSGTQTVFFITYF*MPQCFCMI*N*HTFYLKNR*K--TW---------NLPFYTY
Consensus

Teleogryllus_occipitalis      ------------------------------------------------------------------------------------------------------------------------
Acheta_domesticus             LFFIFQCVCVCVCVCVCVCVCCCRRSCAFAGHRKRSP------------------QCVSVKCKS*HV-----SP*GTYRIG*SKSIPPEYGKSHRKENK*K*HIEH*FYCNSEFNNI
Gryllus_bimaculatus           LMFYYN*LKILY------------*RKLSFLSNNDVKIP*R-----YCFHLLSVHKL*QCLS--CSSGYILILFQTIKGS-------------KDPRKEKDVY--------------I
Gryllus_longicercus           LMFYCQLVKLQH-----------KRKLPF*QTIK*K-PLDDIVFIYLVIYLVIHLLIM*LW--FHSNFI----------------------SKDQGKEKYVS-------Y*KTV*NNI
Consensus

Teleogryllus_occipitalis      ------------------------------------------------------------------------------------------------------------------------
Acheta_domesticus             V---SYKK*FKECSSRD*SNYE*INNQFANV*KSCNSFIRASVQKQLGMKTSI*NYIHNNVFIICSQHATSSYAG--NKNKSDVV----------------------------CENV--
Gryllus_bimaculatus           I---KTV*NMNQ*KLRQKQNFR----S*YKI*DCC--IFRATPVVTL------------MSI*VAEKIVTRTKGML*KRYLEIE--------------------------------CHFITV
Gryllus_longicercus           TYCRETRKNEDKCKISNR-NTK--------------FEIAALLEQL---QN*FQY*YE-----TQKKIVTVTKGRL*KRYLEIKQ*ISNNQNNSILHINQDRKPLLQ*FRF*NCKVVLL
Consensus

Teleogryllus_occipitalis      ------------------------------------------------------------------------------------------------------------------------
Acheta_domesticus             ----QSV*QFI*LLICLFSI-------SHYCGC----------NFILTSLCLTVLF-----CTPY-------LMKELKDFETLIIYN-STNP----------VNCIP*FILVVSSTMV--
Gryllus_bimaculatus           N-FKQS--KIIPFYLCIKIV-------NHFIFC----------GFISS------------------------MSFVKDISYL*MCRCSSKPD--------VTNKICFLLPFIRTTI*FN
Gryllus_longicercus           YWF*SSIFSILKV*IQLKLVRSNILKEKRLISCVNRDLFAAKLNVFVKMFCIL*FYLQHASCEGYLISVDV*M*Q*TRRYKQNLFCA-SFLPEPLFESTAYWRNCSTEIIMFDRHKK*F-
Consensus

Teleogryllus_occipitalis      ------------------------------------------------------------------------------------------------------------------------
Acheta_domesticus             ------------------KE*LNKNYNIL*QNVKFRF*LNMEIEIGNKEEKAPPLPSLL-RELQ*SK*SPAAPC--IVECNRVSALLRRNLFHSSLQATKKKEGDSQFS*TKALGVARS
Gryllus_bimaculatus           GLFEKLQLQYSNYYV*QIQKNGLNLK*NIV*ERVTF----H*KQAN*DSETKKKPHPSRLHRGLQQSPRSPVSPC--IME-RTVEAPLR*T-FSALCEDYEKGRADSQSPQSKARGVAGD
Gryllus_longicercus           ----KIKMKYC------VRKC*VTLK-------------TSELR**NKE-KAPPL-----KTTQGTPIKPTKPCFALHNGRQQWRLLCGETFSAPCAGYEKGGADSLCP*SRARGVAGD
Consensus

Teleogryllus_occipitalis      ------------------------------------------------------------------------------------------------------------------------
Acheta_domesticus             IVARRGWYLNAWIRFHSALSLCSLFGPPVLTRPNDAEPRVTVFFKS*YV*LRR*IKSSECSGNKRIEPASSSKCLLIE*VNRCKHVFQCHSNVDLISH*YLKGRLW-V*ELKSGLLISVQ
Gryllus_bimaculatus           FVGRREWYLNARIRFHSAVSLCSLFGPPVLTRPNDAGPRVSVFFKS*YV*PRP*IVPSECGGNKRIEPASSSKCLFIE*VNRCGHVFQCHCNVDLISH*CLKGRLGRVCVFYSSLLIPVQ
Gryllus_longicercus           FVGRREWYLNARIRFHSAVSLCSLFGPPVLTRPNDAGPRVSVFFKS*YV*PRP*IVLSECGGNKRIEPAASSKCLFIE*VNRCRHVFQCHSNVDLISH*CLKGRLGRVCVFKSSSLILVQ
Consensus

Teleogryllus_occipitalis      ------------------------------------------------------------------------------------------------------------------------
Acheta_domesticus             HKFRYSLLLSKPCALNSIFLNLESYEILQ*IDVKF*R--KHTYRESLVLSLFFKCLSDHESICYESISNVHSICPESS*LVFLRASPTILIFPLSGFLHLGTSTSGALPQERQALSSGLV
Gryllus_bimaculatus           HKFRYCLLLSKPCALISIFSHLQCCEMLQ*IDVE*LMIVRNSWSANY*GHLFRKC--------------SAFFPPEAAVMFSRW------MVRSGAGHL--LASGGFPNHVSPIT*ALI
Gryllus_longicercus           HKFRY-LLLSKPCVLISIFSHLQCYEMLQ*IDVE*LI---------------------E*LFVIPEASTAKVISSVSVVHFSSGGGGDV-LPLDGEVPV---RQRGFPQPGISYQLGSP
Consensus

Teleogryllus_occipitalis      ------------------------------------------------------------------------------------------------------------------------
Acheta_domesticus             SR-------------HVWPV*STQGKWV*V*RGEGTIMRKQRVRATRSPFFKSIDQAFSTY*SEK-----PRKTVGHRYQPTGLFSIQQLNSTQIIAVNTPQSGLKP*DFPVRTSLLVV
Gryllus_bimaculatus           HPGLPLRSLGICGSVVHVCVSVCT*G--VRIGQGRG---RNDRARSV--AFC*GTAPAFDWC*S*RNHGKLPLRAAGR*DSPRG-----*LIST*LCS*YRKFCHRRDSNLNTRTAVTV-
Gryllus_longicercus           QPGQPLRSPDTCGSVVHVCVSVCT*G--VR*GQGRG---RKHRARSV--ALC*GTAPAFAWY*TTE--------TIGRRCRPKGLFSSQ*LNSTQLIVKNMPPSGSKP--YSTR-----
Consensus

Teleogryllus_occipitalis      ------------------------------------------------------------------------------------------------------------------------
Acheta_domesticus             VALNQLIRLS-------KANTLNDLTELEALRKIF*FFLLLHMYY----PVF-IFTILSTIISSPLNNV----A*IQFSLKPLKFPTPIRF*SERLFA-DM*TKLQYRIFHLVPVMNYYK
Gryllus_bimaculatus           ------------------DTLRKLS-----REIDVKFWQKHTFWGSHKKYC-YFKSLCSTMCQ*AT*VP--QASIEFSPRIIDI-VQIRCRSARQVA---------RVSH--------
Gryllus_longicercus           -----LLRQNLYTGSCCGPELTQPLVELQTLRRIV*LFAVLRMYP----PYFPNFVNMCSNCV**CGLCPIFNTAS*LSEIIIF*ASPIPLRK*EMFCTDL*TELQNQIK*------FYK
Consensus

Teleogryllus_occipitalis      ------------------------------------------------------------------------------------------------------------------------
Acheta_domesticus             RVLSEYSALLHHFPNFFNSPALLLLLGLSRED*RIH**SALNRQAGRT*PRAELLVQLAVLESKSYQCLCDR*FFCIGVDHARFAALRQNYNTNNYIIKTSAPPHRQIL-KVRLSPVS*Q
Gryllus_bimaculatus           ------SVYLH-------------LRGTA--------------TARTSSKPSV-----------SGCVCKE--------------RNDNNEEGKIMRDPESAFRRY*SNIRLVLNYGN
Gryllus_longicercus           ----------HVF-----------------------------------------------------------------------------------------------------------
Consensus

Teleogryllus_occipitalis      ------------------------------------------------------------------------------------------------------------------------
Acheta_domesticus             YRTKK*VIE*TIQLNKTVKYLKCYQKMSP*GDISAVP*NY*TMKVQVTRMQHLQIRSSHTKTKW-----------------*IFSNNTNI------------------------------
Gryllus_bimaculatus           YRS*VPAEGIVLQSAREFNTANC-QK-------------YATLGMQTIQ*QTSPSESVHWYLLWP*INPAGCRTRDRAKDRLVFRRTTHVSPLFSKSCQQLFHMHLIMWYMSNFQ*SFLN
Gryllus_longicercus           ---------ITIQS---F*AISC-QTIS------------------------------------------------------------------------IFH-----------------
Consensus

Teleogryllus_occipitalis      ------------------------------------------------------------------------------------------------------------------------
Acheta_domesticus             -----------------YYIKIICYKNIFKN*NNKTKYYNNNFNILCPVQQTTRGQQQNMSHEHLWNVHKRLNTNTMQKILQEETAVNNNQQVHISR*QKKEDPQK*VTIVTQSVEC*I
Gryllus_bimaculatus           FENYYFFEHPLYHF*SEKYFVR*LCYRTKLSD------FINMFF*LF-------RG---------------------FEPVLVELSQFSTRSKQDIFKSVKAKTSFKSVTRENETE---
Gryllus_longicercus           --------------------------------------------------KK*TRS----------------------------------------FQKRKSKNKFQK---RNSSGK*N*
Consensus

Teleogryllus_occipitalis      ------------------------------------------------------------------------------------------------------------------------
Acheta_domesticus             IVQQKTLL*QI----NTQS*KQLKNQMTM*LI*NHKMKM*KYLPYALEMQET*HS*QKISHN**EMDSHLKPQEIR*KYF*KM*LNTVQLENI--VMKIMCLITHTVYNQKKTY-----
Gryllus_bimaculatus           LCKPRTFFLKKLFLPNYFKSPVLRLQKS*KFF-R*SFKVRTFLD*KL*YWETIT***K**H*FL-*NAQWKYNYCEC*YI----------NY--LSRI-----HDLYN---TYI-IQYL
Gryllus_longicercus           TV*AKNFLFEKLFLPNYFKSPVLRLQKKLKVF-PGKFQG*NISRFKI----------------------------------------VILGNYYLMMKVM------------TLIPIKCT
Consensus

Teleogryllus_occipitalis      ------------------------------------------------------------------------------------------------------------------------
Acheta_domesticus             ----QL**KMYP*YSLRKRFLMN*KA*IIQ---*KGN*IN*QRTNTL*G---------------SCIRTSKFNRRKRN---------I*TR*ITSLYCESRIKKKTPRSTPM--------
Gryllus_bimaculatus           *RINDLEGTVVPG----KLF*TS*KF*ILQELLYKNK*INKNESRTFQNL-------TFLTP*GCLIVCKSSHR*HHCQCTLSLSMVLIRQMETLHGASRIQ----------------
Gryllus_longicercus           MEI*LL*MLIY------KLFK*NT*F--IQ-YIYNAIFMNN**FRGYSGTWKVILNVVKILNSSGIIV*K*INK*K*NISESHFSNTLRLLNSL*ILPQIA-------PLPVHAFSVD
Consensus

Teleogryllus_occipitalis      ------------------------------------------------------------------------------------------------------------------------
Acheta_domesticus             --*KMFHSWTHKKLLHNGPKVWILHTKPLDKRL*SIIINKPTTPNLYSM*RKSY----------------------GYIRSMQILPKKKENATTISTQ*PISNSTISKYL*K*NQLTNIPTTN
Gryllus_bimaculatus           -----FIAYMR-------SVT*CRRRRLLKK*ISLLSFKK-----FNCYR*NFRGTKF*L*SLMGCVLIGINGEIRAIM-----ARNLLHIY*HHQINCAVFQK--------KNILEID
Gryllus_longicercus           VVDERFGRWRH---FTEREGYNLLPTCALSRRVSTSIKEIN-----FTVMF*KVQRML*IKHSWHEILLIGINGELRAMK-----ARNILHSY*LHQINCDAFKK--------KNILEIV
Consensus

Teleogryllus_occipitalis      ------------------------------------------------------------------------------------------------------------------------
Acheta_domesticus             TY*SQSITHTNDI**QTKLTHTHICSNSFTKQYSIKHNTNIDADVTNNS*SNNSNNNSSNSKHSV-KPNIKYNKK--WISS-PT*NL*HF*IGMQMDY*IKETHLFTS*KIIQ*QLHA*Q
Gryllus_bimaculatus           P*NRKSTKEQNHTKCFSKK*HSH---------FSRTTNKIISC*TMLN*II*QK*SSRILQ*HSVI*TDTQYNSS--FHTK-NTNK-----KLAVLHRNRY*GIFNGG*AILR*I*KIS
Gryllus_longicercus           AYNRKSADEQSHTKCFQK-NSTHI-----------SQEPQIKFFLIRQC*IK*YNKTILKIRNNIV*YKRTHNTQVLFIPKIPTKNL-----------------LFYIATIIKEYLMAGK
Consensus

Teleogryllus_occipitalis      ------------------------------------------------------------------------------------------------------------------------
Acheta_domesticus             KHT*YQDKHLKYQDITATEKTEYVNMQQEE*PS*SKRN---WTSH-----*FLSQN*MILN*LQ*K*R*EMNI*ISY-----------QHINHLKNKSI*MN*SKYYKVQNLPF-----
Gryllus_bimaculatus           RYQQYENR-*KNSNICFF**NRKCYFVFGFPPSS*EHNEIIENSHVIRVCMFLLQFLCLLEI--------NNICVKFVKNTIPLKCIILQPAGHIEAEM*WTHV*KIAKEEHNYF-----
Gryllus_longicercus           QFCAESEKSVGTSNMRIGEITQLFVF-------------LIKSQML-LCFWSLQFLCLLEI--------NNIRAKFVKNT*PLKCLIIQPAVHTEAAM*KDP*ENCKRVTNFLFFEFSN
Consensus

Teleogryllus_occipitalis      ------------------------------------------------------------------------------------------------------------------------
Acheta_domesticus             -----*WVI*IA-NISIGAAK*THQMEE---------------NY*KLQQI*TSKS*H-----------QQILLT---TRKIVTHNQIS*TLPSLKT*LIPSMLIL-------YLN----
Gryllus_bimaculatus           -----LTLIGFTLN*TL*KYKTTYYTDNMRPAVIYDCMKYHATNYKAP**VV*TSSLKV---------IKEIL*ACFYTSK*YRHIRD*QNRVSLSKNDTPVCLELLFDGFTSYLE----
Gryllus_longicercus           I*MK*LSVIGLG---ALCSMLKKHLMKNERK------------IYSELTSIPRSAEVTVHLWQCHIIEFQKVFVIFIF*R---IHI-----------------------ASCSYLRNCFW
Consensus

Teleogryllus_occipitalis      ------------------------------------------------------------------------------------------------------------------------
Acheta_domesticus             --*TLITVL*Y-SLQICTTL-------------------------KQNCPI---------V*IQKT*IG--------------*NSKQML-------------ITPF---P*TDS*KTKK
Gryllus_bimaculatus           --FFGVCVCMYVCV*VCVVLF**FCISENRHWDD*TH*VF---EVAQNVTAAYFGNK*RNVQIMKGTFASVK*LASQIHSICLINTRNITNFMIKYHIYF*SKLFP*---RITLTEPM*I
Gryllus_longicercus           RLFFCVNCLCF*LLDITG*R---LVLSK**TWH**K-IIF*KNRIKRKCL*LQR*NRSNNC*LVQG---------SKIPE-------------------FQRILTPFHFCKCTFERSKYK
Consensus

Teleogryllus_occipitalis      ------------------------------------------------------------------------------------------------------------------------
Acheta_domesticus             RLILQLI---------------PSLKLSK-TQRKRQILILIINLN*CNKPYLLTSK-------------N-*FKR-------------NTEYDGNGSNTEAQI*RKC*TN*H--------
Gryllus_bimaculatus           ILFLRII---------------S*LK*FK-TDS----YILLVTY---VK*YLRSKL-------------NGKFSRISARRY*VSVSVNSL**WQGERTQG--------------------
Gryllus_longicercus           ETGLR*IKFRYMCTC*CA*GRRASVRTCKWTDVK*A---------DVISAYLEVSKCKRGRCGCESSLSMNRRGSRQSASEQT*EYM*NE*ADLKEDALRG---RKCKSRRSTSEQT*TL
Consensus

Teleogryllus_occipitalis      ------------------------------------------------------------------------------------------------------------------------
Acheta_domesticus             -----AK*KQH*MNTES----NNIKSH*KINPKDNNLWQVTKQILKEPTIIPELKKLDKKATTTKEK-----------------------------------------C-EILAEHF---
Gryllus_bimaculatus           ------------YDCESLRF*DSLKRFSREATSSENLYGIFY*LFY*P---------DSSAVI*SKQIRFGRNIFLIFALKFLF*MFISSFTISIRLSWF*THTLHFFCKQIFIDFFT-I
Gryllus_longicercus           I*YE*ADVKVNVI*ASSR*KWAFVRLSRRESGRTYDQVDMSAKLLSSGTA*RRARKQV*CAVVRNDP----------LPMSFF*GRFLT--------TWL--YSAHCQCF*M*EEHVRAI
Consensus

Teleogryllus_occipitalis      ------------------------------------------------------------------------------------------------------------------------
Acheta_domesticus             -QET--------------------------------------FALNPTSNNNNTIQIINNEIITVPHPVKYIYHQMKHTKQLNT------------------------------------
Gryllus_bimaculatus           VQQTLFYSS--------------------------------FFLFAPQS------LTIFTDF*NYHYFV*YI-SQYKNTSHNGI------------------------------------
Gryllus_longicercus           RTHTIIHARGGIGTRGPSSRGSCGSSALATSAIQAGSAEFFFFFFKVASTSRPLTTAIARNGMGTVRMSRYDDDRW*KDKPNSTMPEAGLEPAVRDRYVTTALTMSAPRPAVSLLFPPAL
Consensus

Teleogryllus_occipitalis      ------------------------------------------------------------------------------------------------------------------------
Acheta_domesticus             ------------------------------------------------------------CLRKNQLE-----------------*TKFLP*FTKTK*ENYRLLNC--------------
Gryllus_bimaculatus           ------------------------------------------------------------IIFD**IE-----------------KIIYLLNFENFE-*NYFVLLV--------------
Gryllus_longicercus           DEVAGGRAGGRVAVRALGDAFSSPAAAEGKGEDAPCSATRPSPLPPSPLKLHLLRDPQRGLLRDSG*EVLNPSH*IIETKYLDTRECIYLL-------SPYLTLNVKQNCNSI*CGCRII
Consensus

Teleogryllus_occipitalis      ------------------------------------------------------------------------------------------------------------------------
Acheta_domesticus             --------NIQFNVKNRVLSHKMETFYN-------------NSNTKTW*KLSIYRKL*THKPNPYNS*SNRKINT*TTSEIHKQISYIPHFQFGFKEKYSTTHQLLRITELVYKS-----
Gryllus_bimaculatus           --------WVHYVLCLKSI**KTNEIF-----------------TQSLLPFRVLRK-------------*QFICGNVI**IFRK-----YLLFLYFEEY-I*HR-VRI*GIVFGDYFFVL
Gryllus_longicercus           ITSICAYI*IHF------ICTEITHLYLKIHCTNFTFVHVSNNNVACFVPMKMGKV-------------TRNLCDVIYVLVYNMLNIL-FVLYSLW*NY------LSYTTLL*IEIYQIS
Consensus

Teleogryllus_occipitalis      ----------------------------------------
Acheta_domesticus             -----------------------LNKKSIL*QFSWT*LKL
Gryllus_bimaculatus           IAYAFN-CSISRDSVSYFLNSELGINEKLYLKKM------
Gryllus_longicercus           QGY-FNIVKPG*HPID*CLISKLIVGQNIVHKQYG-----
Consensus
```

**Tocc.2G0021750.1_exon3**

Teleogryllus_occipitalis ------------------------------------------------------------------------------------------------------------------------
Acheta_domesticus -KYCFYIFY*KLYFTFIMNAIDIL*VYFVSAFVKYDHSIIYKHK*LHARTK---IIKFFAIF*NILLWNLIIVTN*TK*NV*LYCVLLTIRKESNKNFEGI*VSF*AFYTSSSLSIFFET
Gryllus_bimaculatus FSLCTLV------------------------------NLF**HKISK*R*MSDCLLTWYKDFF--------------------------LHAK*--------------------------
Gryllus_longicercus -------------------------------------------------------LKLY*IF----------------------------*KDS--------------------------
Consensus

Teleogryllus_occipitalis ------------------------------------------------------------------------------------------------------------------------
Acheta_domesticus LYRI*KEGINLQVPVASNLSYLSYKFY*RLTQKIIL*SFNLVPNCIFLCPGIFTQLDILCTSKGTV*CTSSCCRLMFILITQDKYLFN*QVE*HLMKLNILNLEYFHVSSEIIIGV*NTK
Gryllus_bimaculatus ------------------------------------------------------------------------------------------------------------------------
Gryllus_longicercus ------------------------------------------------------------------------------------------------------------------------
Consensus

Teleogryllus_occipitalis ------------------------------------------------------------------------------------------------------------------------
Acheta_domesticus ITINV*LFTDLAQSFFSPCRIVRLPFERCESDAPVNNGEDKDRKQFSRT*IRMTMLKIQGHIQKV----ADCKGIDIKNSFCNKKCSRTTQKYFAVLAKDLKHII*DSNKK-------SL
Gryllus_bimaculatus -----------------------IAFEHCVSDAPVNNG*GKYRKQFSRTQ--MTMLKVQGHIQKV----AKFKGTE-RKKWGNKHCRRTKQKYFAAPTKDLKHTIKDNGKRYELPKIFS*
Gryllus_longicercus -----------------------LVFE-------------PF*SNVSRS*NIK*ISKVLTHMSKV*LNYCNF-------------CMFLHSNYFV--------------NKIQFLNILSH
Consensus

Teleogryllus_occipitalis ------------------------------------------------------------------------------------------------------------------------
Acheta_domesticus TIFRSIYIDLLKC*NDIDFFVIKNDSLLFKRLYSNLSRP*KSESISNVLRQISKL*LNY----*PSCIFLHSYNSHTLHTS---VIVVT**SCMVSDFFFQYRFTKLLYSIFVQL*NSHS
Gryllus_bimaculatus HLYQSI-------RNYIEFF-------------------------KKILWFLNHFTQTFLGRKT*NR--FLKF*RICQRCN----*IIATFACFYIQNIVLIKYSF*IFYP---------
Gryllus_longicercus KISPSI-------*SKF*IFC----------------ECCHSRNLQDILLLESCLEKNSCG*RL*SVVFLIKTRRMPWQCHDKRVSITSDLSLLTLVHLFPLRHTYIKYNI---------
Consensus

Teleogryllus_occipitalis ------------------------------------------------------------------------------------------------------------------------
Acheta_domesticus SDNTAMETVSCKLRC**RIKCNIPVLSSCFQ*RGK-----------VCALVAPL-----------DFFKTSRML-------------QQSVKTE--------TYHTRT-ALPHPFTIL
Gryllus_bimaculatus ------------TRYHHQFKQNFEFFVNVATP*TYGISYS*SPVW-----------RRILVADG--FDR---------*FF*LRPGECRGSVMTN--------EYLSRAICPSSPLSIN
Gryllus_longicercus ------------THVPERVAQGIK*VSAVLLP-------ERS*DWNVAGFLIPAG*LR*VQSLSGGEF*RPESLMGSFPWFLRSTPAEWQSSILTNGLGFARCISFLILVISSPTPYTHS
Consensus

Teleogryllus_occipitalis ------------------------------------------------------------------------------------------------------------------------
Acheta_domesticus SHA-HTRTKLICKGDQTGP*RSTRLGRCAAGGDI*RREIPQPGNRDRITCSSGIRTPSRPAM*HALQSRD*CLKQFTHPIY*LNRDSGASLGS*VHCTLFFLVGDGDLPSLYGGIQQRLA
Gryllus_bimaculatus -FLLHTLT-------*SIISLTYRKGWLRALNELVQF-F*LRGSRIAMWQDF*YQLDS*-----VESSR--CLE---------NNSNGRSL*SVVFCGFF-------------------
Gryllus_longicercus AHMHHTIT-------LSGAKNSPRLG-------LVPL*WEIPGSGKSPQDDVGQRRIS-------------------------SSGGRSPLP------F-------------------
Consensus

Teleogryllus_occipitalis ------------------------------------------------------------------------------------------------------------------------
Acheta_domesticus CRLAGFYPTRYLPPVVHRPREHAW*CSAC*CKSVCKRRG*DKEEEGKIVRGPWPVC*DTAPVFAWCLSENPRKTTDQSRRPLEFYSRQQLNLT*LYS*CRKSCHRRDMNLEPSKTEVMQC
Gryllus_bimaculatus --------------------DQHW----VNARAVP*QTG*--------------------GLHAVYPS---------------------SSLSYRYPLLTQT--VRSHA-----------
Gryllus_longicercus --------------------ERE------TCTALTGLMAL--------------------AVVAVALA---------------------ISQSTNYHTCRKI--RRELS-----------
Consensus

Teleogryllus_occipitalis ------------------------------------------------------------------------------------------------------------------------
Acheta_domesticus HQLNALSHLSVTLVRTILTYMMGESFRACPSWGCVPLDRPRGTNTTRWRIGTESHFLVVEGHYHFPKNHYNY-GTNAQLTHPENKIINQSISGYIIQWLVRLMAAHTAPTLLARGACR--
Gryllus_bimaculatus ------PHY-----HTVGS*ELAPAGVGVPLVGDTWFGKIPTR---RRR---TAPYLIQREKIAATFREGNM----HCTYE---------TNG-----LSR*RA---------RIHQSIN
Gryllus_longicercus ------AH--------------------PDWGGARLGEITGKEKARK*---EADLKK*REKNTDTSNKYNML*HD*CLWHLRKK----SVNGCTIRSLVRPRAGRMEPAAPPRVAES--
Consensus

Teleogryllus_occipitalis ------------------------------------------------------------------------------------------------------------------------
Acheta_domesticus --PVSP------CRG----RAGAAASGGAARGRGGGGEGGRTGGRRPRAPAAGA*R--VLPPGRL*DGRAPDPGH---------------------------------------------
Gryllus_bimaculatus QLPHMPGDQTGT*RSSRLRRCT-------ARGDNWKGKSP*EVGVEAEISSNSGARTLTLPPIQ----------HAITGLVTEKKISQXXXXXXXXXXXXXXXXXXXXXXXXXXXXXXXXX
Gryllus_longicercus ------------------RAG-------GRGGLGRGGGGRTHGRAAAPCARRRRR--------------------LTRASTRATIRR-------------------------------
Consensus

Teleogryllus_occipitalis ------------------------------------------------------------------------------------------------------------------------
Acheta_domesticus -------------------------------------SLRRLLSAPPDTHSSAMVS---------LRGAAPALSTVRMRCRN----IDVHHLCIS-HRFLESHVF*YICFVFDIFIFK*
Gryllus_bimaculatus XXXXXXXXXXXXXXXXXXXXXXXXXXXXXXXXXXXXXXASVAPLS*HPPLPW*VRTAP----------------RGMRHRCPGFYGIIDGLHFEFSLHR----------CFVFDIFSFDW
Gryllus_longicercus -------------------------------------ARARRQPQPPSPASHRSIDTLLCHGESARRRAGHVLRGMRHRCPGFYGIIDGLNFEFSLHR----------CFVFDIFSFDW
Consensus

Teleogryllus_occipitalis ------------------------------------------------------------------------------------------------------------------------
Acheta_domesticus KR*YHNKMQ*CVSDISLTFFGTKF-GVEIKIELLKSIQFFK*QFQRKKVC*I--F**GFTVFCKLHPNKHDLRQVNPHV*TLMATIIV*EHQWFGGDILLTSL----L*FPVPVFLEMVA
Gryllus_bimaculatus KNYQLNKFL*SFSYIN*TFSRQKLLRSEIKIYVLKSIHFLKYLFK**SIAKYTDICLAS*IFYKKLINATFGQIMSPVCTFLINIL-IKKHRWFSGDVF*TSIDFEHRCFE-----TCMF
Gryllus_longicercus KNYHLNEVL*SFSHIN*TCFRRKYLRG*IKINVLKSIHFLKYLFK**SIAKYTDICLAS*IFYKN**TQLSAK-*CPLCAHC*SIF*LKKHRWFSGDVF*TSIDFEHRCFDMYVLLTTLF
Consensus

Teleogryllus_occipitalis ------------------------------------------------------------------------------------------------------------------------
Acheta_domesticus S*S---PVVVIT*FSRRNCGVIFIFIF*V------------------------FRVLSTYLGFPENICF----------GICVL-TKNIKK---------------------------
Gryllus_bimaculatus C*RRRFPILQ*SSFRS*LKQTVM*F*FFAS*RFWGC-*IRIENNIYQK*DFCPFFNYIKKLSYLRKTLFFTK---------IFHV-TNNFR*HIYASVTLEEAWNKCVEFWLIF*TY*V*
Gryllus_longicercus P*S---PVVVIS*FAKTNCDVVLIFCFLT---FFGVLNTNQEQNI-SKIGFLSDF*LHQKIIVFRENIILYQNISCHK*YAIAHLCITNFRR------SLEQM---C*VWANIFNLLRLA
Consensus

Teleogryllus_occipitalis ------------------------------------------------------------------------------------------------------------------------
Acheta_domesticus --------------------------------------------------YFRSCFLPVSN------N*RIQA--------------------------------NKLFFTKILKK----
Gryllus_bimaculatus HVKPILKNVRNFACVTY----TYNKYNIIK*FLARPSKGR-NMHKIIQISIYKNIFL*IINFSYHQFDYVVRAVLSLGTPNSCISLPRNSDWLPTLVKEEVENLINHFMMGR--------
Gryllus_longicercus RQADIKIC*KLRLCYVYL**LKHNKIIFWPVHLSVETCTRLYKYQYIKISFFR*LILVITNLIMWSEQF*VSE------------------LPTLVSRCLGTRIGSLFL*RKKKKI*LI
Consensus

Teleogryllus_occipitalis ------------------------------------------------------------------------------------------------------------------------
Acheta_domesticus --------VHT-----CLENPHNIFSSILLKYLKLGSGQK------*PQ*ICFQRYLWCLRMHNEYLPPHTNCNLEV*S-THKNVFSGILYMF*ND----PEKICLLPVDKDVLNDTQLAG
Gryllus_bimaculatus -LGYSLTDVHM-----RLENPHQYFFISSKIL*KNSHKNSVNSSCEVLNISVDAS*IPSFRNHQF------KSLTYSRSFPQEFLYNVL*GFGKNVSPTSTKM-L*MSPNVLQN----SG
Gryllus_longicercus IL*WTG*VIH*PKFTCTLKTPTNIFSSFLRKCNKRIHK-------KIVSIRLVRY*LSL*MTHKYLPP-GISNLEA*R-IQKDFLMSFFTMFYKD----SEKMSLLPVPRCFE-------
Consensus

Teleogryllus_occipitalis ------------------------------------------------------------------------------------------------------------------------
Acheta_domesticus K-----QWSRRCISVSNR------QTHANCTIYMIVCTKIQMCFQFFRI-STKNFQKEL*QKH*KISNHQLTI*ESKKTLVVN*GSLV*RIFYR-----NDYFSSCCFREC*SRSADERV
Gryllus_bimaculatus PENVYLYRADACKL---------------DDLHGYT*T-IQVCFYYFKFRSNSSFSYD-------------------QTLFL---*LIEEIWFKRCSIETFIFLLCFFREWCSNSLVGRM
Gryllus_longicercus ------CHSTCCKTVVQKMYNCTGQTHAN*PLLGYM*T-IQMCFYYFEFRPNSSFSYD-------------------QTLLL---*VIEEIWFKRCSIETFIFLLCFFREWYSNSHVGRM
Consensus

Teleogryllus_occipitalis ------------------------------------------------------------------------------------------------------------------------
Acheta_domesticus EE-G*GGMCE----------VSGKPGAATAR-----------LSFFV-----------------------------------------------------------------------
Gryllus_bimaculatus RRGGEGRGKAGAAAALSSY-FVGREVLQTA------VGIAITLVYY*FCETHSIRRRRKKCNNCVC-RK*RRCNVNLTEPTD*YLIELVVETVDRYIVDYYK*DY*GTDKEASHKEELR*
Gryllus_longicercus RR-GEGGQREGRRSGCTIFLFRGARGIATGRRYSNNVGMLLVL-------*NSQHRKEKKEM*QLCM*KIAACNVNLTEPTD*CLIELRVETVGRYIGDYYKCDYWDTDKEARH*EKSR-
Consensus

Teleogryllus_occipitalis ------------------------------------------------------------------------------------------------------------------------
Acheta_domesticus -------------SRGFVPSQFKIST------------------NVSCVSV---------------------------------------------------------------------
Gryllus_bimaculatus EDSPPTTFSQ*VRSTTIFPSK*RIWTLG*V*IKDY*EFASNV*IHVICVKIETKRLYNFVLQEGILVNISLVTYSVILRIKYIIKVVFAKF---------------------LTISTLNI
Gryllus_longicercus *DSPPTSFSQ*VRSTPIFLSK*RI*TLR*V*IKDN*EFASNVKIQVICVKIENKLIFNFILQEGILV-------------KYFPGYLFYYFKDQIYNKPQDSFC*MF*QFQSLTSHVTDL
Consensus

Teleogryllus_occipitalis ------------------------------------------------------------------------------------------------------------------------
Acheta_domesticus ----------------FTFPYTYGVIYV-------------------V*EK**N--------AARKNGVFWTETNSLP-----------------GVDNE----------KSG*NQ-----
Gryllus_bimaculatus ----ACNIFVFNHSNISIYKDSFWIVYVHVSF*FYHIN**DIIRHNN**DKSVNR*LVFQPYCKRKPAIGFVEIIKTPNQVVISLFGWPIIFGGAGLDN*CALF-----PSKRGCIGCRL
Gryllus_longicercus FSITAISIFIKIHSGLFMYMFHFGLSF-------------*LIRN------FVNR*FVFHPYCKRKAGYWVCRNPSSPNQVFLSLLGWSIIFGGASLDN*CVLFLHKEVASSADC-GTRL
Consensus

Teleogryllus_occipitalis ------------------------------------------------------------------------------------------------------------------------
Acheta_domesticus ------LIN*EKNPLKHG*RIFPENKLGKSDVVFRNDKVR------GR---LHHTHLKNDSIQNNFRRTVE-----VHLTRYK--------------YVTNF------------------
Gryllus_bimaculatus WDSIV*CLKTVEHTSH*LRELFYDQHI---*MICQNTRIAKILEIFGSSNISEYEYRANCAVFECY*INLR--------FSYTCSCTVTTE*LICALGRSSWFS*N*KFSTGSRFHQR*S
Gryllus_longicercus SN----V*KL*DTPDIN*GNVFFDQHI---RTIFQNTRIAKILEIIGR---FEHQ*IRV*S*LRSF*ILLNEPTLLIHMFVYSDSRVINL---------TSW------------------
Consensus

Teleogryllus_occipitalis ------------------------------------------------------------------------------------------------------------------------
Acheta_domesticus --------*GI------------------------------*AANE---------YHFSYLNMWEM**TK----------------------YIC*LSETAFLICKNV*L*FN*NEHCRD
Gryllus_bimaculatus KTSPHSFVK*LKYT*V*NKCYKCLWHLKCQRLSLFPIKYEGNVKNEVTQV--------------------------IILISRP-----------GF*SILSFD*VRTCIVITTQPINV--
Gryllus_longicercus --------K*L------------LVFLKTKIFDRKPIASK-MIKNEST*LCRIVKLHLSI--*WILQTSMAFKESKIITFSN*I*GQCQK*CYTGYHSDIAPRFLRYI*LLLVQNVHCHY
Consensus

Teleogryllus_occipitalis ------------------------------------------------------------------------------------------------------------------------
Acheta_domesticus *AS*NLQGRHFKNFL*SKLASKNKKISLLVNRSLVSSCVRLHAKH*CRKCCLPIRGST*EI*CPQFYL*LPTTHSL----CKIFVVAYKT*KFLKLIMN--------HCESFLRRSN*LM
Gryllus_bimaculatus ------MADPSKILCSRNKSVKSKKA------AIVSEQVAPESSSECQ---------------PAV*EVLPANQGLDVRNSTSVILLYEISHNIFIVQNVIMFDSIT*PTSF---QYSF*
Gryllus_longicercus *ATYKRHG*PFKIFCSGNRSVKPKII------AIV*EQVAPESSSECQ---------------PEV*EVLPTNQGLNVRNSTSVILLYEISHNIYIVQNVIIFDSIT*PRSF---QYSF*
Consensus

Teleogryllus_occipitalis ------------------------------------------------------------------------------------------------------------------------
Acheta_domesticus VP*ATTAVIRLVSEVNVSSRDVDMFLL-------VGS*DLTPTYMTACRVFLSLGLVTLQSTAPVWRSSKALSCRLVSPHAK*CSTYVYGCK-GSE*GRSKEGIVTLILGQIVRILLWHS
Gryllus_bimaculatus IPLAK-----------------------*CMSSSIG**DPTPTFM-ACGVFPSPGLFTHRGTAPAGRSSKLPT---SLPT--CI--IV*YM*VERE*DRAKEGKTAHDLGPIGRVLLRH*
Gryllus_longicercus TPLAK*-----------------MYLLHRIMRSDAGLHGLRCISL-SRPLHP-PG---HR---PNGEKL*APD---QSPH--LCNSVVGGERIVQSKGRKDCARSWPDW*GIAPALNEGS
Consensus

Teleogryllus_occipitalis ------------------------------------------------------------------------------------------------------------------------
Acheta_domesticus FGIERRNNEQLPIKTPDPLNSP---------PKGD*IRPCNLFNCQKFAWSCTLRPFGQN--R**VRDSPGSELISGSPPH*SCKAKVGNTRAHFNGR*NASFAA-----YTHTNG-AW*
Gryllus_bimaculatus --TED-------------------------PWKTIDQSTQLHDSRKCAGTRTLGHLGLNICR---HQHAGSELISKFPSH--CGTKAGYTREEFNGR*IHPLYGVHT*M-------GHG
Gryllus_longicercus --MENCRSEH-----PTPRFSKMRRNSYARAPWSEYLQTSAM-------------LALN-------*SAGSRLIE----------VAGL-RPDTHGRSLTDVKCIRCRVYIHEWGTAHE
Consensus

Teleogryllus_occipitalis ------------------------------------------------------------------------------------------------------------------------
Acheta_domesticus TSALSPLRDA*VSTE*VCCMLLCVSAL-------APCR*EFFQQKLFVV--RTAA*RRKPGTHH-------------------------NHNRC*KYCPFTWF-PVV*L----LLT-*M
Gryllus_bimaculatus TRTRCAM-----------SVVFCLS---------VRRRDVCGNMDVLTSLTRRSCYR*CTALH---------------------IFNYY*SVTHNP**GKGLLHCKVLNLQAI*LVSYYF
Gryllus_longicercus RAARCPL-----------CSA-CLPGVQTFVEIWTF*RP*C-NARVIVN-----------ALHYILFIIIIIIIIIIIIIIISLLPTYLSKTKASYIVRF*IY-KLDNL----LVT-IF
Consensus

Teleogryllus_occipitalis ------------------------------------------------------------------------------------------------------------------------
Acheta_domesticus QLLIGYPNSAQCGETLLARCIILLQNWT*NSLVLSI--EI*TKKSASSAQARG*RRAWLTVCVMNCVRDCVRG*LCAWLTVCVANCVRG----*RCA------W-----LTVFVTNCVRV
Gryllus_bimaculatus CSLLPVP-------FFLFMCIITSP*IFATTFVFYICNHISGHLTFASN---------LRILFHQ*PKTSLRKMYVNFGP---SGCWRSD*PRWRAGGRANDVWFAGAVARVGGAGVC--
Gryllus_longicercus CSLLPVP-------IFLFMCIFTSPTLFATTFVLHICNHISGHLTFASN---------LRISFHQ*SKTSLRKRVCK--------CWSS--------------W----LLEVGSAEVARG
Consensus

Teleogryllus_occipitalis ------------------------------------------------------------------------------------------------------------------------
Acheta_domesticus WCG---VVRR--RCCTCWWRWRARVRRRGPPSSASAAEDWASAAVSSPSNPLSLPLTQSAHEQWKCN*------------------------------------------------------
Gryllus_bimaculatus -------VCRGARLAHRHRRRRTRLRRR*VSAVRMRSGNGTDGGLA----------FVCDIAYGNSG----------FPWIDNLHQFSAVWKP------------ENE*VL---------
Gryllus_longicercus WVS**RVVCR--RCCTCWWRWRVRVPRRTPRSSASAAEDSASAAVS----------LSCAHAQWKCN*FN*IELMPYFTWRDSGHEARSSVRPRKFRYAPLTN*QRKEVFLGRETGDLNL
Consensus

Teleogryllus_occipitalis ------------------------------------------------------------------------------------------------------------------------
Acheta_domesticus ---------------------------------------------------------------------*RFCICQLCCLLK*QLSIA*PLPSVSCCLKARNRLR**FSGFQTVGN----
Gryllus_bimaculatus -----------------------------------FFGFQTKG--------------------N*R*R------------****LT*N**SGS--CCLETTTDG--HFLAFKQLET----
Gryllus_longicercus AQRRHLGLRIFYMQVNLLTWDSRLYFPSEGRRTKDFFALKTAHGFDRV*TRQLGYNSPVR*PLHHRRR*WRPCICLQFCLWK*RLSMD**SSSVFCCLEARK*KRIRFLAFKQKETEEHD
Consensus

Teleogryllus_occipitalis ------------------------------------------------------------------------------------------------------------------------
Acheta_domesticus ----------------------***WL-------------SL*KI*RI------------------------------------------------------------------FSGFLQFL
Gryllus_bimaculatus -DDV*AIEYYYFREQYSWK**RL*LLYL-ECQQLITAKQK---------------MMVY*CVVTAS*RKKNYTWGDT*CSAQVSQFS*LGIRLHFTVDSSGIWVVKAWLRQSFSGF-SCL
Gryllus_longicercus DDD*HKTDE---RGPAVWKRLQMVIFWL*NCWKLMMAKPSNTIISINNILGTITPLIIIFGMTAINHRKTNNN-GLLRCG-----YC*LKKKVHLGRHAM*CSVPSIFM----SGY-QAS
Consensus

Teleogryllus_occipitalis ------------------------------------------------------------------------------------------------------------------------
Acheta_domesticus *LSISPKYVINLHNHCS--SKSC-----THFQ*LP*G----*GITKNTI*LPGFKLRNIHVVIVCLISVIASGFLKIISFTWLLFHFSLYHSYGIFI--ISSVSDN*SIL------CKIL
Gryllus_bimaculatus CFSPFLRSLSHAFTDSNS-SKSY-----IHFQKRPES-FDVGTTINNTIWLLEFK--SQKILV---ERMIERGY----SYRF------------------SRITFVSP--------TSNA
Gryllus_longicercus YLGRFIRDLGRLKQDCGRVSRGFPVHVSLHFSKVSRTPLQTPTHQNHT--LTSRK--DLKALTWGPQ*IIPSGYWNLNSGKFL---WNL**KGGILIGFLE*LSFPLPLMPRMSRSVRTL
Consensus

Teleogryllus_occipitalis ------------------------------------------------------------------------------------------------------------------------
Acheta_domesticus **RIFTLDVRRTVLK*--------SGCLIYSSV--NAVQILACPCGIF------GEESGFTAGFLESSWFPRISFRHCSNNIHIMFLKSHSPK------NPPPCSRY---IQKTSNRHYY
Gryllus_bimaculatus TYVTSCTDVVINNVHLRLLREPY*SGKAVQFILLGIAVQILVCPYDLW*RKRVCGR---FSWVFTVSLHFIPSLLQQNSHHIFVIIYPSK--SFTNF*IRPA**R*YGIIVKITSLSLTI
Gryllus_longicercus **IMYILDF*ENHIKV---------GKLLNLFL-------LVSRFKFWCVHMVCDEGSGFAAGFLGSSRFLCISFLHCSRKIHIISMSSSIPQFINF*IRPA**R*YGIIVKITSLSLTI
Consensus

Teleogryllus_occipitalis ------------------------------------------------------------------------------------------------------------------------
Acheta_domesticus EDKMIQFAHGRSAA*Q*GC*LLLNFW*TKQISWR*S*DSPISPF-----FSFYH*FITHIISS*SSS---H*ILIYTESTTHAI*NNFLYK*KDC-NMVMMYFCETFLKLHIYCTVSLYY
Gryllus_bimaculatus RD---------LL*SR*GIVILL-------YH-GGILGIISFPLHVTSQLLYHPHYIVTIIYSLNLLIYPENIPIFIVKNFYC*WKYQ*HG-----------FGIIL*DI-IKC------
Gryllus_longicercus RD---------LSRSS*GIVTLL-------*SWRGSCKFSHFPY-----MLFHNCFIIHIISSRSSI---N*IT*YIQETSPSLW*RTSTAN*SVSNMALV*FYETL*NF-INAL**Y--
Consensus

Teleogryllus_occipitalis ------------------------------------------------------------------------------------------------------------------------
Acheta_domesticus **GRGDKR--------*YCNFAV*MF*NI----------------S**FYSLTLKFQV---RDKV*YNM**ISSERLSSQTK*RKRCKCEDGKGKYE***NTKTNNSNIYFKLVLIQTL
Gryllus_bimaculatus ------HQ*IVVVG*S*YCHVAV*IL*NIN*LLFT*IKILSTR*SVM*HAIIIVEK*QIVKHKNK*QVVFILNVL--------------------KNKR*IS------------------
Gryllus_longicercus -----DSRDIVM-------------------LLFKYSKI-------*INYYLLELKFQV---RDKVQCNM**LSL--------------------KSNR**NTKTSNK*YLF*TC**TNV
Consensus

Teleogryllus_occipitalis ------------------------------------------------------------------------------------------------------------------------
Acheta_domesticus NSQRSTL*NNYYISF*NLNRKYSKSHGILCLHVH--------IRCKNAQL*L------------EF--------HRRF---HIMKYCNL*T*SL-*QFLTNTVHCTADRKHN---MHA*L
Gryllus_bimaculatus ------------------AE*ISEIMITLCFKI*IVVVSKIEITRRTFHISCKHGATM*TRISPYY---------------FIL*NTSI*ERDLFWQFLTLS-HFTAPQIAYTICMHNYF
Gryllus_longicercus ----------------EFAEKISEIMITLCFKI*IVV-----IRNHTAHFS-------------YFVQTKGNYVH*NFTGGFIL*NTSI*ERDLFWQFLTLS-HFTAQQIAYTICMHNYF
Consensus

Teleogryllus_occipitalis ------------------------------------------------------------------------------------------------------------------------
Acheta_domesticus FSRETWRNVSFNWAILFVRF-CNNIARMSMSNFIIGPM*NIPTIQRLDWHVKIL*TVPSPLAHTLRLQGF*YNVQ*VPMNNHA*FQSHSFSFRDFLLRSPVFGII*LS**TPKRMTKISF
Gryllus_bimaculatus LGKH---GETFHLIGPFVVFRL*NIAPMSKSNFVIGPM*NILIMHRID*YVKVTCE---------------FKI-*IF*NNYI----------KYLI-----------------LNRKYF
Gryllus_longicercus LAKH---GETFHLIGPFVVFRL*NIAPMSMSNCIIGPMRNISIMHGIAWYVTDICK---------------FNI-*IY*NNYI----------KYLI-----------------LYRKYF
Consensus

Teleogryllus_occipitalis ------------------------------------------------------------------------------------------------------------------------
Acheta_domesticus SLNFQNFRKMQRFCK*MECKTK*DY*ISI*SKKRLPNCNLSVNYEASYTLLHPRMISHIMAQTL*FLLFYYYYSIV*TTWLSWLKKFV---------EISFFIVTIKLYC----------
Gryllus_bimaculatus QS*F*FFRH*SE---------------NIL*QK*L-------SYDRRF------MI*RIQIYNGEFILF---------K*LE*LKCYLK*QTGLNVMEITGYTMKITCY*RKTKGSRRND
Gryllus_longicercus QS*F*FFRH*SE---------------NIL**K*L-------SYDRRS------MI*RILIYNGEFILF---------K*LEQLKCYLE*QTGLNMMEITGYIMKITYY*RKIKGSRRND
Consensus

Teleogryllus_occipitalis -------------------------------------------------------------------GGIGLGGGGLGLGLGLRRGGIGFGGGLGGYGGGLGLG---YGGYSGGYGGGYG
Acheta_domesticus --NSKLKI*SVVLRSLEF*A*NQRRCIYYNLCALKDEVSTVCVPQVASALAAAAWVWAWACGGEASASAAAWAATGAAWVWATEATAVATEAGMAAATASAAAVWAWAWAESGCWAAAWG
Gryllus_bimaculatus VFQYSFSLLMYLLLSLGFEG*N---------------IDFLC----------------------SAGGIGLGGGGLGLGLGLRRGGIGFGGGLGGYGGGLGL-----GGYGGGYGGGYG
Gryllus_longicercus ILNYSFFLWMYLLFSLGFEVRN---------------IDCLC----------------------PAGGIGLGGGGLGLGLGLRRGGIGFGGGLGGYGGGLGL-----GGYGGGYGGGYG
Consensus * * * *

Teleogryllus_occipitalis GGYGIGGGGLGLGLGRIGLLGG--------GLGLGGGLGLRGGGGGLYGGGIGLGKGSYGLGTYGAGFGYYGK*---------------------------------------------
Acheta_domesticus G----------------------------GLGLGGGLGLRGGGGGLYGGGIGLGKGSYGLGTYGAGFGYYGK*ALSGRAAPSKGLSSPSK--APAPAPGSNPRPAKRRFHALMVRFGPV
Gryllus_bimaculatus GGYGIGGGGLGLGLGRIGLLGGGGGLGGGLGLGLGGGLGLRGGGGGLYGGGIGLGKGAYGLGTYGAGFGYYGK*AMSGRAAPPTGLSSPSKAPAPAPAPGSNPRPAKRRFHALMLRFGPI
Gryllus_longicercus GGYGIGGGGLGLGLGRIGLLGGGGGLGGGLGLGLGGGLGLRGGGGGLYGGGIGLGKGAYGLGTYGAGFGYYGK*ASSGRAAPPTGLSSPSKGPAPAPAPGSNPRPAKRRFHALMVRFGPV
Consensus * *************************** ****************

Teleogryllus_occipitalis ------------------------------------------------------------------------------------------------------------------------
Acheta_domesticus QRKGTSPKVFEFRRVDVVEIHDSSFTYDILLFSQNNYPFFRRFIFGSFSPKEKIIKSISPHFFSDKF*VVSCK*RTVCVSVI*IIFVHPKS------------CLTISCNVFPR------
Gryllus_bimaculatus QRTGASPKAFEFRRVDVVEIHDSSFTYDILLFSQNNYPFFRRFTFGSFSPKEKSIKSISHHFFLDKF*VVS*K*RTVCVSVI*IIFVRPKAA*RSAVMYFRA*CMFM-CGILKKYILSCF
Gryllus_longicercus QRTGASPKAFEFRRVDVVEIHDSSFTYDILLFSQNNYPFFRRFTFGSFSPKEKSIKSISPHFLLDKF*VVS*K*RTVCVSVI*IIFVRPKAA*RSAVMYFRA*CMFM-CGILKKYILSCF
Consensus

Teleogryllus_occipitalis ------------------------------------------------------------------------------------------------------------------------
Acheta_domesticus -LVHIHVWYIKKIHFILFLVDLFAQNPFLHTYTSTHGFCTLIFSGLMQLIE*SFLY*ASIWLEEAFSIVNALTFF---*NTCKFRIQAFLGINICKCA*FS*K*RTRKITSLFI--IYNL
Gryllus_bimaculatus SLLYFHHILFSH*HIPLFI---------------------LIFEVVF------------WHHE---IVALLTNLASG*NSCFF*--*LTQKHICK-GLVSLKLRMFPE*ILIDSIQYN
Gryllus_longicercus SLLYFHNILFSH*HFPLFI---------------------LIFEVVS------------WHHE---IVPLLTNFVR-LKLVFFK--*LTQKHICK-GLVSLKLRMFRE*IIVDSIQNN
Consensus

Teleogryllus_occipitalis ------------------------------------------------------------------------------------------------------------------------
Acheta_domesticus EKSKSLIKI-----NTYS------YKINFNILFFVSII*NLILDADFTHLY*MLRTGCGQ------IQELKRD---LRK*I*TSLNV*FT*LDN--NPYIPNYSHSV*K*NKYNA*DVMT
Gryllus_bimaculatus VSEK*LPNL*ITIQRTYKD*YLLLQK*T*MFLFFAFIT*NLFLEAEFTHLFHMF*TGRGQFANQNTIY*RVRSS*CQQN*MCNLYNLIIMSILKIINT*FEN-------KKKYIA*E-*S
Gryllus_longicercus VSGK*LPNI*ITIQRTYKDKYLLLEK*M*MFLFSAFII*NLFLEAEFTHLSHMFRTGRGQFANQNTIYEG------QDK*MPTKLNV*FI*LDN--NFYFENY*LLV*K*KKYIA*E-*S
Consensus

Teleogryllus_occipitalis ------------------------------------------------------------------------------------------------------------------------
Acheta_domesticus FPI*NK*LKYLIRKKHFKIMEEINTYSIL**LII------FTYWLILPPQFK-----EKNQFNM**SELMGDTVGYK------------SNTFYRYLSYSNRN*VICSRHFIRFCETNYF
Gryllus_bimaculatus LQ-YNRNN*LLNRK*KLKYWKK------*I*ILIHSN***NRIWGYTHIGWFCHNNLSKH*Y*KNMKLGVGDTVRYK------------LSTFYVYLSYSNRN*LIC*-DILSDSARLII
Gryllus_longicercus FQ-YNRNN*ITESKIKIKILKEINMNSSL*QLIIKSNLGIHTYWLVFVTTI*ANIDIKKI*NSE*ETL*DIN*VRFMCTSHILTGIDSFVETFYQI----QRD*FFWA-GVDRYCRGLIL
Consensus

Teleogryllus_occipitalis ------------------------------------------------------------------------------------------------------------------------
Acheta_domesticus GQELTDIAVFWYFQVAALNNHQSSGTNLSFTTADVDL------KSS------REVYSRFPWPQRTLFKREAEGVLPFHSMIFLLKRSSAKI----MCSFDKN*LKISLLIGTNLLFRSHV
Gryllus_bimaculatus LGRS*HIAAVCYFQVVALNNHRAAGTNLAASAADVDLKSPPPARSTLVFLGRNELCSR---------GRRRE------TFLFSARFSSLSIPRLKLCALSTKTEKR*VC*SAQTCYLDRM
Gryllus_longicercus SSCGPEQPPGSRYQ----PRHQCSGCRL----------NPP--RSTLVFLGRNELRSR---------GRQGE------TYLLSARFSCLSIPRLKVCALSTKTE*R*VC**VQTCYLDRM
Consensus

Teleogryllus_occipitalis ------------------------------------------------------------------------------------------------------------------------
Acheta_domesticus IS-KNVLMLCI--SWRLAM-MRK--*KSEFYECLTCKPRFTE*NPAPFSTRSITFLFKLLYFF*LDRSSGDRFFMFFN*YPRLPWSLRGSLY-----*ET*IQTHASDMCCR*QKNYFMI
Gryllus_bimaculatus LSPRTF*LYVFHGGWP*L*GNRKVYLTNV*HVNQGFRNRTPHSSALEV----LHFFLN-YYIYFNYTVLEETDSLFFN*KQWLPLSVM-LLYTNKSESNTFVGRRVVDI-----IN-FMN
Gryllus_longicercus LSPRTF*LLCI--SWRLAMVMRK--*KSVFYECLTCKPRFPQ*NTAQFSTGSLAFFLKLLYLF*LDSS*RDRFFKFFN*YQWLPRSLM-LLYTNKSEFNTFVGCRVVDI-----KN-FMN
Consensus

Teleogryllus_occipitalis ------------------------------------------------------------------------------------------------------------------------
Acheta_domesticus RWKFLYT----LLKNCCD*SANLGKYR*NPKLNTLLLYC*KTSNARSVIKN------*LYKLYTLDNAGDTEDTSYGNSSV----DCAQRLSVYYGNYRTERPT----ARCVLLTV*KYS
Gryllus_bimaculatus RCNFFINSS*KLLRVRCKL*KT*VKHKNKSLVA*FYAITNCSLEL--IIENN*L*RTTIIALCPLDNASST*N*SFRFSR*IALKDFLLRMAT------TERSNLLHVAYCT----WRTA
Gryllus_longicercus RCKFL*T----LLRNCCECGAKFEKDK*NTKISHLLPNLMQTQPI--VL*N------*LYAFSPLDNASST*N*SFRFSR*IAHKDFLLRMAT------TERSNLLHVAYC------HKP
Consensus

Teleogryllus_occipitalis ------------------------------------------------------------------------------------------------------------------------
Acheta_domesticus CD*----------------LFEFI*FVTLF*LTVF------CIQ*IRLSSIRLLTREIRFRREEW----NSV*QVRGLLEINLKVDSFMPKE-------M*SK**IIDYLVILNMILFYT
Gryllus_bimaculatus TNRVMNIYM*LII*NYFICHIFLIHYNSVY--NKYKCQVFGCLREI*ASEER----------------NGIRF--NR*EVS---QN*I*N*TVLCLRRFGLKNR-------LYVILF--
Gryllus_longicercus CNE----------------HIYVIDYLKLFYLSRYFDSL**CL**I*LTSIRLL-REIKLQK*EMEFDLTGKRFLKN*FEIR---QSFA*GD-------LGSKNR-------LYVILF--
Consensus

Teleogryllus_occipitalis ------------------------------------------------------------------------------------------------------------------------
Acheta_domesticus EMI*LIFDTC*LKK*H------IDHV-------LLSDNKCKCTIF----SILTMCVGFLLHNTVQLLNVVQLNVR**PQQHELLVEFP--------VFFLPDRCEKENGII---------
Gryllus_bimaculatus ----L*T*VCTTQRRMNPDLTTID*TRKTLFTELPLEITCKCIHF---SKLIPWLFSSA*HRPIKLH----LDVGS*RQPHKYIFRHC*WCSQRCLVFLLSDCCKNEKNIIFIKITLIEQ
Gryllus_longicercus ----L*T*FCTTQRRSNPDLTTID*TRKTLFTELSLENQCKCIHFF*NDSVV--*FSSA--QPVQLSSI*MLD-------------------------------HNANIINIYLDIASG
Consensus

Teleogryllus_occipitalis ------------------------------------------------------------------------------------------------------------------------
Acheta_domesticus ------------------LMQIT*I*LILLC**TC*NF-EKMSFSFIS*CSLQLLKFS*YYLDLKLFETNH----------------------------*NKLK*TLK--------ADL
Gryllus_bimaculatus TNLYNCVNEFYQIF---KNIIRIILFHLIVV*RS*NLRDIVGVSNYFLYNLFLNA------SMILKDLKMRHNLLVKSIEIMDFFILYVFDV-----------LLQLIK---------NS
Gryllus_longicercus A---LCGLSLIQML*KRKILMQITLF*FILLC**I*SND--GINNFLSYNCVLESLKSA*YGLGLKLFSL*H-----IFKCLNAFA*FETDTHFT*KIDRKHGICHFIRI*CFIAN**NF
Consensus

Teleogryllus_occipitalis ------------------------------------------------------------------------------------------------------------------------
Acheta_domesticus NDRGGKCIGRLSASR*RVTVVCSLSFSVCVCVCVCVCVCVCVVAEGLVRSLATEREAPNVSASSVKVNMCPHEGHIG*DSRNRYLLNM---EKATGRKINKNSTLNINFTAILNLIIL
Gryllus_bimaculatus LLEKTIFFIK-Q*CKNSLTIL--FSFT*CT-*IVTMFIM*F-WLHSNFISN---DQRIKG-----SEKRKRC----VYYKNS-MKYE--------PV--KIKTKTKFQI---VIQNLRLL
Gryllus_longicercus NTRESYLFNK-QLSKNLLTIL--FSF-------I*LFI*LFIY*SCNYGSILILFQRIR-------EKKNMC----HIKK----QYEITLHIVEKPV--KMKTNAKFQI---VIQNLRLL
Consensus

Teleogryllus_occipitalis ------------------------------------------------------------------------------------------------------------------------
Acheta_domesticus SVTKNSLRSVVHEIDRIMNR*TTSLPTFENLVIPSSGLVYKNN*A*R--------HQYEITSTIMCSLYAHNMLPHHMLET-------------KTKVMLYVK----------MYSQCN
Gryllus_bimaculatus HFQSNSSSN----FNVYMSRRKNCNP-------------YKRNAVEKILGNRMPFHNCKFQTVK-------------------------------------------NNSILLMYQDRK
Gryllus_longicercus HF*SNSKTN----FNIDMRRKKKL*P-------------LQKDGCRKDTWK----*NSKFQTIKIIPFYI*TKIVNHFYNNSDFKIVRLSYSIGFEAVFLAY*KFKFS*NSLEVIY*KKN
Consensus

Teleogryllus_occipitalis ------------------------------------------------------------------------------------------------------------------------
Acheta_domesticus NLYNF*FVCLAY--HIIVDVISY*RLYVLLFCFAHHTS*RN*KISKR*LFIILPI-P*TASLNSSWS*VLQWLKN-------------S*IKTTTFYDKMSNFVFN*TWKLRLVIKKKKP
Gryllus_bimaculatus PLYFLWF----YLQHVICEGYL---------------------ISVDV*M*Q*AR-RYKQNL-----FSASFYQNHYLIQRLIRETATAIQ*LLCLTDTKKWFKF----KIKYCVRKGYV
Gryllus_longicercus D*Y-----------HV*IEIYLQPS*TCSLKCFAFCSFISSTPVVKDISYL*MCKCSSKQDVTNKICFVLPFYQNHYLNQRPIGE--TAVLKLLCLTDTKNSLKL----K*NIV*ENVKL
Consensus

Teleogryllus_occipitalis ------------------------------------------------------------------------------------------------------------------------
Acheta_domesticus LLSQVCSESSNKANEVLLRLA*W---SAIEFLLYC-------------AETFFIALCKLRKRRKGILNSRKPRLWAWHEVLSPVAVGI*THGSAFTAPCHSAVYLGLPF*PGQMTLNPE*
Gryllus_bimaculatus SLKTSELR**NKEKAPPLKTTQG---TPAKPTKPCFALHNGAHSGGSTAVNFFSAL*RLRKRKSRFTVAAKQGSG--------------RGGRFRRPSRVVFKRTDPLSQRRVTLQ---
Gryllus_longicercus H*KQAN*DSETKKKPHPSRLHRGLQ*SPRSPVSPCIMEGN---SGGSSAVKLFQRPVQATKKEEPIHCAPKAGLGAWREISSAVGSGI*THGSAFTAPCHSAVYLGLPF*PGQMTLGPE*
Consensus

Teleogryllus_occipitalis ------------------------------------------------------------------------------------------------------------------------
Acheta_domesticus PCFLRVS-------MYN*DDKLSPANVAGTSVSNLLRP-------RNVFLLNRSTGVSTFFSVT--VMWI*FLIDI*KEGC-GYES*K--------VVF*FQFNTNSDI---LYCFPNLA
Gryllus_bimaculatus --FIWASRSDPAK*RWAQSERVF*ELVCITETINCAQRMWREQAYRTCFVLEMS------FH*IGQPVWARFSMSL*C-GFNFSLMFKRKVGKGMCILQ*FVNSCSTQIQILFIAFQTLR
Gryllus_longicercus ACFLRVS-------MYNRDHKLCSANVAGTSVSNLLRP-------RNVFSLNRSTGVGTFFSVI--VMWI*FLIDV*KEGWEGYVYLK--------VVR*F*YNINSDI----YCFPNLA
Consensus

Teleogryllus_occipitalis ------------------------------------------------------------------------------------------------------------------------
Acheta_domesticus P*IPFF---*I*KVMKYCNKSTSNFDVN-----------------------------------------------------------------TRIANRWY*VCFSNVCPTM-------SR
Gryllus_bimaculatus LDFNFFSFTML*NA--AINRCRITD---DCS*FLECQLLRSFVP*V*CIFSSGGGGDVLPLDGEVRCWPPVSQRGFPQPRISYHLGSYPPGTTSKVSRHMW*CSACMCKCVYVRGEDRAR
Gryllus_longicercus S*FQFF---LIYNVMKCCNKSMSND*LNDCS*FLKRQLLRSLVP*A*CIF-------------------------------------PPEVAVMFSRWM------------------AR
Consensus

Teleogryllus_occipitalis ------------------------------------------------------------------------------------------------------------------------
Acheta_domesticus Y-----------------------------VTKVSQTCIAFVQRVVDWYFCVR-----------------------------------------HLL------YLFFL*VVSYIW------
Gryllus_bimaculatus KGKERSRAVCCLLLGHCSGIRLVLIVKKPRKTTTQSSWTLGFSSRLADFDLTM*LV*KILPSTGLKPQHPHGSDS*YPEKTLPGNRCQILAKTHILGIS*KILLF*VSLFNHVPVSNVSA
Gryllus_longicercus S-----------------------------------------------------------------------------------------------------------------------
Consensus

Teleogryllus_occipitalis ------------------------------------------------------------------------------------------------------------------------
Acheta_domesticus ------------------------------VPPPAGHCPRSDKH*A-PDWSPDMCGQCRARKVSGCR----CKEGKGQ**GSKEC---AQLVARFLKVLIKHSPRIDLRNHEKLSVIDT
Gryllus_bimaculatus SSEYRIFSKNY*YCTDPMPISTSSCEGFPLRVSPPAGHRHSQDKL*A-----------------*CKWVCM*GE-KRQ**GRKDN---ARPGVRFQKILIQHSPGIELR---KLSVVGA
Gryllus_longicercus -----------------PLDSG---GFPNQVSPIS*ALPSRDNL*GLP----------TRVVV*CMYA*VCVR-KG*DEGKEEDGNIARGLLPFVRALLQHSPGIELR---KPSVVGA
Consensus

Teleogryllus_occipitalis ------------------------------------------------------------------------------------------------------------------------
Acheta_domesticus SPRDCSPYNS*IQPR*LL*IRHSRDSNPRTSRSEPLYW*LLP*TNSSACLKQIL*MI*PNSRHCERSFNFSCYYTCI----------IPYLFLQFCQQ*FQARLIMWRKSNFH*NPLNFP
Gryllus_bimaculatus GRRDCSPVST*IQHS*LSKICHLGDAN-HTVTDFPVRICTLVPVVALN*PSRL-----SNSRQGEGSFSFSPYYACIPSIFQILSTIVPYAFNNVVYVQFSMKLLKFRKLLFL*TSP-VP
Gryllus_longicercus GRRDCSPVSD*IQPS*LSKICHLREAN-HTVRDFSVRISTLVAVVALN*LSRL-----SNSRHCEGSFSCSPYYACIPPIFQILSTCAPIAFNNVAYVQFSIQPLNFRKLLFFEHPL-YH
Consensus

Teleogryllus_occipitalis ------------------------------------------------------------------------------------------------------------------------
Acheta_domesticus LQSVSEVRDCL-RICKLSYST-------VFFIWYR**IIINAFYLNIQRFCIIFRTFSIL-------LRFFYC*GCQGKIEE-----------------YINSLLST------------
Gryllus_bimaculatus LLK----*KIF---CTL---TMLQNQIK*F---------YKHVFLTIQRF*A--SSCRTIAIFYKK*TRYFQKCKSKNKFQKRNSSGK*N*TV*AKNFLFEKTLFAKLFQVSCASFTKKL
Gryllus_longicercus FES----EKCFVRICKLNYRTKLSDFINMFL*LFR---AFEQFLVKLSQFST--RSKQDL----------FKSVKAKTSFKSVTVRENETELCKPRTFFLKNSFCQTISSLLCFVYKKS*
Consensus

Teleogryllus_occipitalis ------------------------------------------------------------------------------------------------------------------------
Acheta_domesticus ----AKRDGRNLEPSCLCS*QCL------------------------------------NQSLISACAIDNSSA*A*TTLVLPHCGK---TIIRTIILSRQVHRHIDKFLK*GYLQFRDN
Gryllus_bimaculatus KVFPVKFQG*NI--SRL----------------------------------KIVILGNYYLMMKVM-----------TLIP--IK---CTMEI*LL*MLIYKL-----------FK*N
Gryllus_longicercus KFFRGSFKVRTFLDSRL*YWETIT***K**H*FL*NAQWKYNYCEC*YINYLSRIHDLYNTYIMQYL*IIND----LEGTVVP--GKLF*TS*KF*ILQELLYK---------------
Consensus

Teleogryllus_occipitalis ------------------------------------------------------------------------------------------------------------------------
Acheta_domesticus TELK----SK**NKQS-----N*TRQ*NI*NVIRRCPPRG---------------TYQQFHRTTKQ*KYK*LVCNTCKFAVHTRKPNGRYSATIPTSTTSKS--YATRTSSKIRTTKPN
Gryllus_bimaculatus T*FIQYIYNTIFMKN**FRGYSGTWKV-ILNVVKILNSSGIIV*K*INK*K*K*NISESHFSNTLRLPNSL*ILPQIAPLPVHAFSVDGVDSADGDTSRSE*DTIYCLHALCHVVSTS--
Gryllus_longicercus --------NK*INKNE-----SRTFQ----NLTFLTP*GCLIVCKSSHR*H--------HCQCTLSL--SMLLMS---------------DSADGDTSRSEKDTIYCLHALCHVECRR--
Consensus

Teleogryllus_occipitalis ------------------------------------------------------------------------------------------------------------------------
Acheta_domesticus TTTTTSTFSVRCNKQ---------RGGSSRICPTSTCGTY-----------TNA--*IPTPCRRYFRKKQQ*TTINKCT----SADNKKKRIHKSK*L**RKA*NAK**CNK--RPYSNK
Gryllus_bimaculatus --TSIKEINFT-----------------VKF*KIQLL-*IKLSWHEILIIVING-----MCVNRH-*WRNTSDNG*KSTAYLLTPPDKLCCFSKEEYIGNRPLK*EKY*GTKPHKMFFQK
Gryllus_longicercus --LLKK*ISLLCFKKFNECYR*NIRG--TKFC**VLMGNYEL*RLEIYCIVTNSTR*IVMLLKRRIYWK*SPIIG-------------------------------KVLMNKATQNVFKK
Consensus

Teleogryllus_occipitalis ------------------------------------------------------------------------------------------------------------------------
Acheta_domesticus *ILNLRNN*RTK*QCN*FRTTK*KCKNTSHMH*KCRKHNTPNKRSHTINRKWIHT*NHRRFGKNISKKCN*IPYS*KIL**K*CALLHILYTIRKKLISCNKKCTRNIH*GRDF**IRKP
Gryllus_bimaculatus IAFTFLKNHK----*NYFVLDNVKLNNITKIIFSNSAIT*CNLNGHTIQQF----------------------FSYQKYQQKTCC-------------STSQSLLGNI*W----------
Gryllus_longicercus IALTFLKNHK----*SSFLLDNVKLNNITK-QFLKSATT*CNINGHTIHKF----------------------FSYQKYQQRTCC-------------STSQPLLRNI*W----------
Consensus

Teleogryllus_occipitalis ------------------------------------------------------------------------------------------------------------------------
Acheta_domesticus KLSNKKVTRLTNREQIPFKVVALELANSIEGK--EIFKLDKLLHCIVKVELRRKPQGPPQCKKCF---------------------IHGHTKNYCTMAPK---CGYCTQNHW--------
Gryllus_bimaculatus --------RVSN--------FALNLKNQQVPAI*E*VKELKYLFFLIK----------SQMLLCFWIPS*FVGT*RNHRKLSRNQSLHVFTSVSVLIGNK*YMC*IC-*EYYTS*MYNIT
Gryllus_longicercus --------RVSN--------FALNLRNQ*VRAI*E*VK*HNYLYF**N----------RKCYFVFGV------------------------------------CSFC--AYWK*IIYVLN
Consensus

Teleogryllus_occipitalis ------------------------------------------------------------------------------------------------------------------------
Acheta_domesticus ----TKDCDLLLSTNPPPPIC--TLCKGNHMATSDQCK--YFPKRKKMQQQSQLNNQY---------------QTLQSQNIYENKTNSQT-----------------------YQQQT
Gryllus_bimaculatus TRWAYRGRNVMD------PCL--ENCKRGA*LFSNVN--WFYLELNTIEI*N--NILY**YETCCNI*LYEISCN*LQSAIVSCMNEFSES-------HK-----GNTLS-------LFL
Gryllus_longicercus L---*RILNLLNA**YSLPCIQKQQCRRTHKKTAKE*QIFYFLNFQTFE*NNFLSLVWVHYVLC------------LKSI**KTNEKFTQSLLPFRVLRK*QFICGNVI**NFRRYLLFL
Consensus

Teleogryllus_occipitalis ------------------------------------------------------------------------------------------------------------------------
Acheta_domesticus HINHNPLHTPMT----------------FNNRQNLPTPTYAQTVSQNNIQSN--ITQILMQMLPTIVNQIIATIIPQIQSIVSNQ-----------ISNIIRNG*AAQLKIYNTSKLEC
Gryllus_bimaculatus HFEII*TYKRLTK*SISFQK*YSCLFGITF*W-------IYFI----FGI-----FWRV---------------CVHVCVRVSVRGF-------ILIILHF*KQALG*LN--SLSL*SC
Gryllus_longicercus YFEESILHRVLI-*EIVFGDYFFVLIVYAFNCSISRDSDSYFLN-SELGINEKLYFEKIELRENAYNFNARIEVTIVNLCKVQKSQSFKEY*PRSIFVSVHLRDQNTKKLD*GESSSGTC
Consensus

Teleogryllus_occipitalis ------------------------------------------------------------------------------------------------------------------------
Acheta_domesticus KWIIK*KKHT--------YSLPKKSFNNNCMHNRNTLSTRTNI*NTRI*LLQ---RRQNMSTCSRRSSHHNQK---------ETGPVINFYPKIE*S*TDCSENNVR------------K
Gryllus_bimaculatus T---KCN----------------------SSLLWKQIT*CAD-YEGYICFSKVTC*PDSFNLSNKHA*YHKFYDKVSYLFLVQVIPLTYYF-----------N*TNVN------------Y
Gryllus_longicercus A---RADVHEEGVRVCGRASGPT*SEQT*SQHTWK*ANVNVDVVDVKVVLV*IDVEVDRVQVSRRRSICRTSE---------QT*KKTHY-----------EGANVKVDAVRVSRRKR*Y
Consensus

Teleogryllus_occipitalis ------------------------------------------------------------------------------------------------------------------------
Acheta_domesticus *ISK-------YHISI*TT*KTNQYK*IKANITKFRTYHFNG*FK*QTYQLGLQ--------SEHTKWKK-------------ITKSFNKFKLPNHSTNKSY-----SLPE---------
Gryllus_bimaculatus FISKN-----YFIIEII*NGFIYFVSYI-----------------------------CKMISSF*T*W*I-FKNICEKILGFCICQFIVVMAR-GKDTRI*L-----------*KPEILR
Gryllus_longicercus STSKQT*K*M*YEQAVVENGRLYG*ADVKVGVRTIK*T*VRSCFQAEQHE-GVRGSRCSVRS*ETTRYL*AFFKVAF*RLGY-IAPTVSVFKC-RKNT*EQYEHTQSSMPEAGLEPVALR
Consensus

Teleogryllus_occipitalis ------------------------------------------------------------------------------------------------------------------------
Acheta_domesticus ------------KS*RTTR------YLRHCH---H*KPN-SYPLCSFFT*IRL*SLSCNTHFRYVLH*SRIAP*YKYKKHRLDKI-----PNKC**RPSHKP----IHEKQRRD------
Gryllus_bimaculatus ---------------QPKKV-----FQRSNI---Q*KPLWHFLLTFLLT*QQCGNIVETNSLR----KKHIS-------------------------------------------------
Gryllus_longicercus AEAAAALQH*RPRPFRPARLSFFFFFLRSHQLLGH*RPL---LLVTVWEL*GCHDMTMIDGER----TNPIQPCPRRDSNPRSEIATSLLP*RC--RPPDRRCRFCFHRRLMKSRAGGRA
Consensus

Teleogryllus_occipitalis ------------------------------------------------------------------------------------------------------------------------
Acheta_domesticus --*FCS*YPH*NYPKRSERGKY*S---------*S*T*INATNHTC*HRKINSKETQNTTAMAAIQKPRFKESAKQTNMPSKNNTR*TQNPTI*NHIKK*TLKTTTYGRLQNKY*KNQQL
Gryllus_bimaculatus --YICFKISFL-----------------------------NVYILVHHKYTS--------------------------------------FMVLNTYTTFLL*TDFYRF---------L
Gryllus_longicercus GGWLCVRWAMLSRAPPLRRGKARTRPVAQPGLPLSPPPPSNSTFFAIHREVSSETLDEKF*IHPIKLLK------------------QNIWILVNVFTYFPPTS---------------
Consensus

Teleogryllus_occipitalis ------------------------------------------------------------------------------------------------------------------------
Acheta_domesticus YLS*RN*IKKLQQQKRSVKY*RNISKKLLL*IQLQIITTQSKLLIMK**QFHIQ*NIYITK*NIPNN*TPA*EKISWNRPNSYHNLQKLNKKTIGYLTAIFNSMLRIGYYPTKW------
Gryllus_bimaculatus YYCTTNIILFFILLICSSVFN----------------------NFY*FLKLSLFCVIYLS---IQ-KHLS-----*W-----YNYF*LVN*KN--YLSFEF*KL*MKLLCVIGLGALCSV
Gryllus_longicercus -H*TLNKI--------ATPFNVDVE***L----------QS-VHISEYILFALK*LIYI*KFIVQILHL-----------------------------FMFQIIMLLVLCQ*RWGKL---
Consensus

Teleogryllus_occipitalis ------------------------------------------------------------------------------------------------------------------------
Acheta_domesticus --KHSIIIPIPKPGKNSAYTENYRPISLIPIIAKVIEKL----ILKR----LLKYINKYHTYHISNLDLKKNTQRHTNF*ESLN*YTKV*TKKAFSSSFLGRNSSF----------
Gryllus_bimaculatus LKKHLM------KNKRNIYSE----LTSIPRSAEVTVHLWQCHIIDFQKIFVIFIF*RVHIASCSYIRNCFWRLFFC-FNCLCF*LLDITG*RLILSK*--*TWH**KIIFKKNG-
Gryllus_longicercus [illegible]PETC-VM*YMYWCIIC------*IYYLYFIHYGKITSLIQLYYELKYTRFLKDISTL*SLVNTPLIDV*Y--RN*Q*DRTLYINNMV
Consensus

**Tocc.2G0023520.1**

```
Teleogryllus_occipitalis       TTTTGGA-----------------------------------------------------------------------------------------------------------------
Tarbinskiellus_portentosus     TCTTCGGTTT---------------------------------------------------------------------TTTAACCTGCGCCTC----------------CAGAGCTGTC
Acheta_domesticus              TTTTGAATTTTGCTCAGAGGCGTCATTTAGGATTGAGGATATTTTACACGCAGCAAATCTACTAACATGGGACAGCCGGCTTTACTTCCTCTCCGAAGCAAGGCGTGCAAGGATTTTTT
Gryllus_bimaculatus            -------------------------------------------------------------------------------TTATATTTCCTCTTT---------------ACAAAATTTTC
Consensus

Teleogryllus_occipitalis       ------------------------------------------------------------------------------------------------------------------------
Tarbinskiellus_portentosus     AGCTTTCACA----CAGGACTTCGTTAAATTCCGACCGCGCACAAAAGG---------------------------------------------------------------GCCCCGG
Acheta_domesticus              CGTCCCTAAAAATCCATGGCTTCGATCGGGTTTGAATCCGCGAACTTGGGTATAGTCCGATACGCTAACTGTTACACCATCGTCGCCATATCTTTCCACCGCTCTCCGGCGCCGCATGAG
Gryllus_bimaculatus            TGTT--------------------------------------------------------------------------------------------------------------------
Consensus

Teleogryllus_occipitalis       -----------------------------------------------------AATTCATATAATTAAGTATCAATAC-----------TAAAAT---TCAAACATTTGTGCAGTTAGAA
Tarbinskiellus_portentosus     GCGAGAAGGACGTGGCGGGGGTCGGGCGGGCG---------------------------------AGCGGGCGGTGC------------------GGCCCCGGGGCGGCGGGGCCGGAG
Acheta_domesticus              GCTGCAGGGCCTTAATGGGGAGTGTGTGCGCCGTAGGTAGGGATGTT-----CGATTCCTGTAA--AAGAATCGATAC-----------TGGAATCGATCCCAGAATCGAAAAGTTGAAA
Gryllus_bimaculatus            -------GATTCTTGTCGAAGTTGCGTACGTTGAAAAAAAATGTTTTTGAGCCGTCTCCCACAG--AAGGAACAGTTTTATCGGCATTATAAAAACGCCACCACCATTTTGAAGTTTGAG
Consensus                                                                          *     *  *                                           *    *

Teleogryllus_occipitalis       TCTGATT-----------------AGAGGAATATATTTCC-----------------------------GTCTGTTGTGTTATTACTGTTAAAATTTTCTGACCTTCT------------
Tarbinskiellus_portentosus     GCAGAGGCGGAGGCG---------------------------------GAGGCGCTCGAGG-------GCGCGTCGAGGGCCGGGCCGTCCTCGCCGCCCCGCGTCCTCGTTAGGTGTG
Acheta_domesticus              TCTATTCTAGAATCGAATCGAAAGGAAAGAATCGATTCCCTAAGGTATCGAATCACTTGAGAATAGGGAGTTTTTGCCATCTCGAACGGTAAGTAATATCGATCTTTTGC-TCATGCGTA
Gryllus_bimaculatus            GTTATCTTGGACT--------------------------------------------------------GCGCGTGATAAAGTGCGCGGCACGAGCTACTTGCCCGCC------TAGGAA
Consensus                                                                                     *    *                            *

Teleogryllus_occipitalis       -------TTTCAAAACAGATTAATACAACT----------------------------------------------------TTTCC-------------------------------
Tarbinskiellus_portentosus     GCT----TTGTGCGGCGCGCGGAGGCGGCGGGGGCGGGGCGGGGCAAGGTGGCGGGCAGGCTGTGAAATTAGGCGCGGGGCGCGTTTCC-------ACGCGGCG---------------
Acheta_domesticus              GTTCAAGTTTCGCATCACTTTAGTGCATCTGAG------CATGCGCAGAAGGTCGATAGTGCTTAACGTTCGAGGTGGGAAAATCTCCCTAATGCGATTCAAGGGAGGAATGGAAACCCG
Gryllus_bimaculatus            GCTAGTTGTTGAAATAACGTGAACTAAATTAA--------------CGAGTGTCTATAGT-----------------TAACAATTCTC--------ACTCGACGGACAGCTGTTAAGAAA
Consensus                             *                                                            *

Teleogryllus_occipitalis       -------------------------------------------------------------GTAATTTATTTATTTATTTAA-------------------------------------
Tarbinskiellus_portentosus     ----------------------------------------------TCCTCGTTGGCGCGAGTATTGATGTTTCAATTTCG-------------------------------GTT
Acheta_domesticus              GATCGAGCAATTTATATTTAATAAAACACAACTAGATGTTAATTAATTAAATTTAATAAAAATAATAATTAACTTATTAATTTAATGAAAATAAATTTAATAAAATATAGATGAAATACT
Gryllus_bimaculatus            AAATAAAAATCTTTTGTTTGTTAAACCTCG--------------------------TAACTTGAAAGTAAATTAATTAATTTATTG----TAAACTTAAGAATACACGA----------
Consensus                                                                             *  *    *  ****

Teleogryllus_occipitalis       -------------------------------------------------------------TGTTTCGA-------CCCTTGTTGGGGTCTTTTTAAAAAT----------------
Tarbinskiellus_portentosus     CGGTGCCGCCGCTGCGGGCGGTGG------CGGGGCGGGGCGGGGGCGCGCGTTCGGAGCGCAGTGGTCCGACAGGAAGCCCAGGCACGCGTGGAGCTCCGACTTCCAAGCGCCGAATCA
Acheta_domesticus              CGGCCCCGTGGCCGAGATTGCTAAGCCTGCTGTGGCGATCTCGAGCCGTAGGTTCGCGGGTTTGTGTCCCGC-------CTCGGGCGTGGGTGTATTTGTGTTT----------------
Gryllus_bimaculatus            ------------------------------------------------------------------------------------------------------------------------
Consensus

Teleogryllus_occipitalis       --------------ATGTATATTGG--------------------GTTCAAAAT-------------------------------------------------------------TGAA
Tarbinskiellus_portentosus     GCGACCAGAAATGTAATGAATGTTGG------------------ACATTTCTAAATAATGATGAAATGAATGCTTCGTTTTTCATCGTCTTTTCCATC---------------------
Acheta_domesticus              --------------GTGTGTGTGTGTGTGTGTGTCGCTTTCTCACCCTTCATAATCATCATCAA--------TTCATCTTCCAT----ATCCCCACGGCTAACAACCCTGGTGTGTAAA
Gryllus_bimaculatus            --------------------------------------------TTACAAAATCAAGATACT--------TTCGATT-----------------------------------------
Consensus                                                                    * *  ***

Teleogryllus_occipitalis       AAAGAATAGGTCAAAATGTTAAAATTATTCGGTTTCTATTAAATATTATTTA----------------------------------------------------------------------
Tarbinskiellus_portentosus     ----------CGAAACGTTCCT---GTTTGTTTTTAATGGTGAGTGATATACGTGCGGTGCGGTGGGTAATGATTCCCCCATTGGTACA-----------------------------
Acheta_domesticus              AATGGACTCAGCAGAA---------ATTCTGATCCCCCCAAACAATGTCTAA--------CGACGAGAAATGCATGTTCGCGTGGTCCATAGAGGCCGAGTCGCGAAAAAAAAATGATGA
Gryllus_bimaculatus            ----------CAAAATATTTAA---GTTTACTTTCAATTGAAAATTG------------------------------------------------------------------------
Consensus                                *  **           **    *

Teleogryllus_occipitalis       --------GAAGTTTTAAAAGTTTGTCAATGTCTTTTTCAAT--CCGGAACACAAGTTACTA----------TGTTTGAAAAAGATCCGTAAGTTTGAAATGTTTAATAAAATAAATTAC
Tarbinskiellus_portentosus     --------GAAGT---AAATACATAAGAGCGTATTCCCCATTGACATGGAAATAATACGTAGATTGGCGTCCTTTTTTGAAGGTTCCAT------------TTTAATAAGGTTATTTTT
Acheta_domesticus              AGTAATACAAAGTTCTAAAAGTGTAACGAAGTCTTT-------------CACAATATACAA----------TATTTGAAAAAGACCGCAACGGTAGAAACGTTGAATAAAGTAAATTAT
Gryllus_bimaculatus            --------GAAATCTGAGAGATATATTATTGTATTT---------AAAGGGATAATTTATTT----------TCTTTAGTAGAG------AAAATGGGAA---TTAATGTAGCTAAATTT
Consensus                               ** *   * *    *      ** **               * **                * ***   *                    * ***      *  *

Teleogryllus_occipitalis       TGAGAAATAAAATCTCTAATTTTGTTTCTGAGAATTTGTTT---ATATTTGTAAAGAGAAAAAT-----------GATAACAGTACAGAAACATTCAT--------------------
Tarbinskiellus_portentosus     AAGAAATAAAAAATCTTAAAATCATTATAGGCTCATTATTT----------TCATTATATAATTTCAAGCCCTGGAAATAACTGCAGAAAAATTGTCCT--------CGTTTTAT-----
Acheta_domesticus              TGAGAAATAGAATGTGCGATTTTGTTTCTGAGTATTTGTTT---AAATTAGTAAAAATAAAAAT-----------GATTATAGTACAAAAACAAATAT--------------------
Gryllus_bimaculatus            GGTGAAATAA-----------TTGTCATCTAGTACTTCTCTTACAAAACAGTAATGAGACAGATA-----TCCAGGGGTAACTACTCACGAGTAATTATAACCTTCATGTTTAATTCGCG
Consensus                          **  *            *  *          ** * *          * *  * * *  *              * *      *  *       *

Teleogryllus_occipitalis       ------------------------GTAATACAATA-TTTTTCAATTTTTTCCCTTGATATTCGTTATCTAATTGTTATCTTCAAGTTGT----------GAATGTGCTATTATTTGTCG
Tarbinskiellus_portentosus     ------------------------CCCTCCTGATTCACTTTTGCTTCTCTTACC--GTATTTATTATCTCATTGTT---------------TAGCATCGAAAAGGGAGATACGAGATAG
Acheta_domesticus              ------------------------CTAAAACAGCATTTTTTTCAATTTTCTTCTACATATTCGTTGTCTAATTGTTATCTTCAAGTTGT----------GTTCGTGCTATTATTTGTGT
Gryllus_bimaculatus            CGCCAGTTTCTAAGGTCGGAGATGCGCGAACTGTTGTATTCTTAGTTACCTTCCCGTGACCTTGTC-----ATGGCTTCCTCCAGATTACCTTAACACAGACGTGCACTATTTATGAAGA
Consensus                                                  * *     *       *       *  *      ** * *                             *    **

Teleogryllus_occipitalis       AACGATTACGTTCTTCTCCGTATATTGTAG----------AAAAAATGTATGTCAATGAAGCAACCCACTGTTTTAAAAGGGGTCGATGCATG--TAGTGTAGGTAATTCTCGGTCTAGA
Tarbinskiellus_portentosus     CAAATTCTACTTTTTGTCTGCGTAAATCGGGCAGTTTTG-GAGCATTGTGTGGCGAGAATAGG------------GAGGGCGGGGGGGGGGGGGGCACGAGGGCTAATTGGGGTGCAGG
Acheta_domesticus              TACGATTATGTTCTACGCTGCATATTCTAG----------AAAAATATACATCGATGTAGCAAGTTATTGTTTTAAAAGGGGTCAAAGCAAGAAAAGTTCTGGTAATCCTCGATCTAGG
Gryllus_bimaculatus            AATGAGTTTGAACCTTGCCGACCAAGATGGCCAGGGTGGTTGAGAGCTTGTACCGATTCAGCG--CTTCTGTTTGAAAGGTCGCGGGCTCGAA---------------TCCCGCCTTGG-
Consensus                       *               * *    *     *              *    *    * *                    * *  *                          *     *

Teleogryllus_occipitalis       CAATTGATAAGCAAGAAGGTCTAGGCTCTTTCTTCC-TCCGTGTGATGACGAGGACGCTTCCTAGTTTT--------TAAATTAGTTGTAGTTGACCAATGACCTAGACTTTGATTTCAC
Tarbinskiellus_portentosus     GGGCGCGGAGACCCGCCGTTGCAACCTGTTGCTGTTGTTGTTGTCGCGGGGCGGCGGCTGGGCCGCTGG------CGGGGGCGGGAGG---------GGGGGCGTTGGCTGAGG-----C
Acheta_domesticus              CAATTGGCGAGCAAAAGGTTCCAGACTCTTTCTTGC-CCTGTGCGATAATGAGAACGCATTCTAGTTTT--------TAAATTAGTAGTACTTGACCGATGACCTTGGCTGTTGTGTCTC
Gryllus_bimaculatus            ---------GCACGGTGGTTTGTCCCTTGTCTCCCATGTATGTGTTGTCGCGGAGGGTTTCGCGTTTTGCTGACCCTAGATAAG------------AGGAGCCCGTGTTAGTGTGTGTT
Consensus                               *     * *     *  *  **        **       * *   *       * *                  *                 *      *

Teleogryllus_occipitalis       TGATAC--TGGAGGATAATTAATAATATTTCCTCTTTCCAACATTTT--------------------------------------------------------------ACAACAGTT
Tarbinskiellus_portentosus     CGAGGCC-CAGGGAGCTGTCGCGGGTGTAACCT------------------------------------------ACCTGCTGATGAGTCGGCGATTGAGGAAGACGAAGCAGTT
Acheta_domesticus              TGTTACCTTCGTGAATTATTTATA----CATCTCTTTCCTACATTTTCTGTTGATTCTTGTGAGACGTATATATGATGGAAAAGATCAATTGA-AGTCGACTCACGCAGCAGGAACAGTT
Gryllus_bimaculatus            AGTGCC--TAGTGGATGGTCGAATATCCCATC----------------------------GGGACGT------------AAAGGTTAACCGACTGTCGGCTGAGGAAAAAAGTTCATTT
Consensus                       *   *    * *     *             *                                                                          ** **

Teleogryllus_occipitalis       TTG----------------------------AGCCTTTAGGCT----------------------------------------------------GGTTGATTCACCTTTCGCCAC
Tarbinskiellus_portentosus     CCC--------------------------------------------------------------------------------------------------------CTGCACCAC
Acheta_domesticus              TTATCAATTTTAGAAAAACGCGACCACCATTTTGCATTTAGGTTATCTTACAGCACACGTGACAAAATACTCCCGCACGAGCTACTTGTCCGCCCGGGAAGCTAGTTATTGTTGCTACAA
Gryllus_bimaculatus            CTTACA--------------------CATGAAGGATTAAGTTTAACTCAGAG----------------------------------GCCCGTCTCG----------------------
Consensus

Teleogryllus_occipitalis       TAAGCAATAATGAGTTAATGA-----------TGGATGATATAA----------------------------------------------------------------------------
Tarbinskiellus_portentosus     TTCGCCGTGATGGATGGCTATATTT--------CGGGCGCCTCGATTT-------ATCTAACAACGCGCCCTACTACGCGGGCGTGTCTGCCTCTT------CTGTTGCGTGCGTCTTTC
Acheta_domesticus              TAATGTGAAATAAATTAACAAATGTGTTACGATTAAAAAATAGAATTTGGTACGAGTGTATCAAC---TCCTATGAGAACGACGCGGCACTGTTTTACACGATTATCTCGAAAAGAGTTT
Gryllus_bimaculatus            TGACCTTTAGTGCATAATTTCGCGTCCTTTCGCGTGGCGCTAAAA-----------------------CTCCCTGAGCCGAAGTGGCACTCTTTTACGCGATTATCTCGGAAAGAGTTT
Consensus                      *         *   *                                 *

Teleogryllus_occipitalis       ------------------------------------------------------------------------------------ATGCTACCGTGTGATACATGGTGGATTC-C
Tarbinskiellus_portentosus     AGAAAGCTCTCGACGCCACGCCGCTTTTCCCTAACGAGTCGAAGTCGTGGCGTTTATAGGCTGTTGTCGCGATTTTACA------------------------AAGGAATTGATT--C
Acheta_domesticus              TGGAAGTGATTAAGAGGAAGC-----------AATCGTTTGAAATCTTTGGAAGTGGTGGTGGAAATCTCGATTTCATCGCAGCATCGCTACGTCTGCGCATGATACGTGGTGGATTTAC
Gryllus_bimaculatus            TGGAAGTGTTTAACACGAAGC-----------AATAGATTGAAGTCCATGAAAGTGGAGGTGAATGCCATCATTTCATCGTAGTATACCAACGCCTGCACTTGATAGGTGATGGATTA-C
Consensus                                                                                                                *     * ****  *

Teleogryllus_occipitalis       CATTGCCTTCTACACCACACTCATAAAGTGAAAACATTC---------GTTTCTTCCTCCATTCGTATTC----------------------ACAACCTGTTGTTC-------------
Tarbinskiellus_portentosus     TTTTACGTTCT----CACACATTTAAGCTGAAACGATTTTGAAATTTAATTTATTTTGCGATTGGTCAAAAATCTTATATAAAGAAAATAACGAAATTTATTGTGTGAGATTTTGAATGA
Acheta_domesticus              TAATGCCTTCCACACCACACTCGTAGAGT-AAAACATTT---------ATGTTTTACTCCACTCGTACT---------------------------CTGTTGTT--------------
Gryllus_bimaculatus            CTTTGCCTTCCACACCACACTCGTAAAGTGAAAACATTC---------GTATTTTCCTCCATTCGTTTC-----------------------ACAACCTGTTGTA--------------
Consensus                         * * ***     *****   **   * ***  ***           * * **   * * * **                              * ****

Teleogryllus_occipitalis       ----CGCGTAAAGAGGTTATCTCCGAAGGAATCGGCACTAGATTCAATTAGCTGTCATTGCTCATCCTCCTGTGTTTTTTATTACTAAAATAATCAATGTGAATTATCTTATGATGAAAT
Tarbinskiellus_portentosus     AAATTCAGTGAATAATGTA-CTCTGTGCGAATATACGCCACTTTACCCTG--------TGGTTTCCACATTTTCCTTCCAGTTATTAAACTTATGA---------ATCTTACAAACAA--
Acheta_domesticus              ----CGCGCAAATGGGTTGTCTCTGCGAGAATCTGCGCTACTCGCAATTAGCTGCCATTGCTCATTCTCTTATATTTGATATTATTAAAACCATC-----GAATTATCTTATGATGAAAT
Gryllus_bimaculatus            -------------------TTTCGCACGAATCTGCGCTACTGAAAATTAACTATCATTGCTAATCCTCTTACATTTGATATTACTAACATCATCGATGTGAATTATCTTATTATGAAAT
Consensus                                         *  *   ****   * * *         *          ** *        *    **    *** ***    **           ******  *  **

Teleogryllus_occipitalis       TGACTTTTATCGAAGTTCAAATACACCACTTGCTTAGAAATGAATAATATATCA--ACACAATTCCGCTTGGAGTGAAAAAAGTGAGCGGAGCGAATAAGTAATGTAAACATCTACCTC-
Tarbinskiellus_portentosus     --------ATCGTACTTCCAACCTGGCATATATGAATGAATTTGCGGCCCGCTACAATATGATTTCTCACCAATCAAGAGGAAAAATATGAACTAATGAATGA---GAAAACACATCCT-
Acheta_domesticus              TGAC-TTTATCGAAGTTCAAATATGTAACCT-CTTAGAAATAAATAGTAAACCA--GCACAATTCCGTATGGCGTCAATGAAGCGAGCAGACTTAATAAATGA---AAAAATCGATCTC-
Gryllus_bimaculatus            TTATTTTTATTTAATTTCAAATACATCACTTACTCAGGAAGCAATAATCCAGC------------------------CGAAGCGAGCGGACAGAATGAATATTGTAAAAACCTACTTTC
Consensus                              **   * *** **      *  *    *  **                                       *   *   **   *** * *      ** *   *

Teleogryllus_occipitalis       ------------------------------------------------------------------------------------------------------------------------
Tarbinskiellus_portentosus     ------------------------------------------------------------------------------------------------------------------------
Acheta_domesticus              ------------------------------------------------------------------------------------------------------------------------
Gryllus_bimaculatus            CCTGGGACATTCCCAATGCGCAGTTATATAATGAACTGGGCATTACACCCATCCCCGTTTACTTCTCCCTCTTATCTGTTCAATGCTATTCCAAAGTATTAAAGATCCCTTCGCAATTTA
Consensus

Teleogryllus_occipitalis       ------------------------------------------------------------------------------------------------------------------------
Tarbinskiellus_portentosus     ------------------------------------------------------------------------------------------------------------------------
Acheta_domesticus              ------------------------------------------------------------------------------------------------------------------------
Gryllus_bimaculatus            CTTTTACATACGTAAGCAAAATCCTCATGGCAAAAGATTAAAGACAAAGTTACCTAGAGAATTATTCAATGCACATTGACATAAATCGATGAGTTTTTTCTCTCTATAAACTACTGTGTA
Consensus

Teleogryllus_occipitalis       ------------------------------------------------------------------------------------------------------------------------
Tarbinskiellus_portentosus     ------------------------------------------------------------------------------------------GGAAGGTATCAATAC---------------
Acheta_domesticus              ------------------------------------------------------------------------------------------------------------------------
Gryllus_bimaculatus            TAACCATTGTATGGTATTATATCTAAAACATAATTTCTGTGAATGATATACCAAATATGTTAATATATACAAATATTATATATAAACTATATCAATGTATTTGTTATGCGTCTGAACAGG
Consensus

Teleogryllus_occipitalis       --------------------------------TCAAAGATGTATTAGTAATGCTTACAATGAGTGTTATATTAGTATATGTAACATTCCTGTAA---------TAAGAACGAATGAAAA
Tarbinskiellus_portentosus     --------------------------------CGAAAGGCCCGTACGTAACCGTGATGAGTAGTCGGCAGCAAACATCTGTAGTAAGTATCTAAGAGGTAACTTAATGAGTAAT-TGCA
Acheta_domesticus              --------------------------------TGAATGAGGTATTTGCAACGTTTTTAAAGAG------------TTTGCAACATTCCCCTAACAA--ACCCCAAGGAAAAATGAGAA
Gryllus_bimaculatus            GTCAACGGCCAAAGCTAGAATAAATTTAAAAAATCAAAGGTGTATTTGTAATGGTTACAAAAAGTATTCTACTAGTACACATACCATTCCCCTAACAGCTGTAGTATGGACCAATGAAAA
Consensus                                                    ** *     *  * **   *    *  **                  *  *       ***          *   *  ***    *

Teleogryllus_occipitalis       GGGTAAAGACGGCCG-ATATAATCAATGACAGT---------CAGCTATTTATTTCC----------------GAAAAATCTGGCCTTTCACTTTCAAAACATCCCTGTTTAGTGAAGG
Tarbinskiellus_portentosus     TATTACA---------ATATATCCTTTGACTGTGTATTAATCTAGCAATCCATCTCCTCAAATAGGACGGTACTATAAATATTAGGACGGTATGTAAATAACAACT---------AAAGG
Acheta_domesticus              CAGTACAGAAGGCCGAATGTAACGAACGACAGT---------GAGCTATTTATTTCCC--------------TCCAAAAAACGGAGTTTCACTTAGAAAACATTCTTGTTTAGTGAGGG
Gryllus_bimaculatus            CAGTAAAGAAGGCCGAATGTAATGAATGACAGT---------TAACTATTTATTTCC----------------ATAAAATCTTGAGTTTCACTTTCAAACCATGCTTGTTTAG-GAGGG
Consensus                         ** *         ** **      *** **           * * **  ** ***                    ***      *       *  *  * * **             * **

Teleogryllus_occipitalis       CTCTTTGAATTTATTTTGCAGCGTCTCGGTCTCTTTCTCAGTTTGGTGGGATGTAAATATGCTCGTGCGCGGAGAGGGTCTGCGAAAACGTCGCTGGCCGCGTATGGAAATGCGGGAAGC
Tarbinskiellus_portentosus     ------AAAATTATTTTTCCCCGGGTGGACCTCCTTC-----------GGACGC--ATAACCTTGTGTCTGAAGAGGGTCTGCGAAAACGTCGCCTCCCGCGCATGGAAATGCGGGAAGC
Acheta_domesticus              CCCTTCAAATTTATTTTCCAGCGTCTCGGTCTCTTTCTTAGTTTCGTGGGATGTAAATATCCTCGTGCGCGCAGAGGGTCCGAGAAAACGGCGCCGGCCGCGTATGGAAATGCGGGAAGC
Gryllus_bimaculatus            CTCTTCAAATTTATTCTCCCGCGTCTTGGTCTCTTTCTTAGTTTGGTGGGATGTAAATTCCTCGTGTGCGGAGAGGGTCT----------GCTGGCCGCGTATGGAAATGCGGGGAGA
Consensus                             ** ***** * *  **  * *  *** ***           *** *   **   ** ***   * ********            **   ***** ************* **

Teleogryllus_occipitalis       GCTCGGTCCGCATATGGAAGTGGC----CTGGCGCTTTGAATCGA---------ACGCGAGCGA----GGATGGGTCGCT---------GGCAAGGGCGGGGGC-------GGGGCGG
Tarbinskiellus_portentosus     CCTACCTCCGCATATGGAAGTGGC----CTGACGTTTTCAATGGGGTGGTGCTGCGCGGGCGAGCGGGGACGGGGCG-----------GGGCGGGGCGAGGCGGGGC---------
Acheta_domesticus              GCTCGGTCCGCATTTGGAAGCGGCCTGGCTGGCGCTTTGAATCGA---------ACGCGGGCGAGCGCGGACGGGGCGCT---------GGCGAGGGCGAGGACGAGGGCGGGGGCGG
Gryllus_bimaculatus            GCTCAGTCCGCATATGGAGGTGGG----CTGGCGCTTTGGATCGG---------GCGCGAGCGA----GGATGGGGCGCTGTGGAGGCGGGGCGGGCGCGGGGCCGG-----CGGAGAG
Consensus                       **   ******* **** * **     *** ** ***  ** *            **** ****    *** *** **             **   * *** **

Teleogryllus_occipitalis       GGGCGGAGAGAAAGACGCGCGCCGCGATGAACTTGTCCCCGTGTGTCGCAATTGATCGGGAGCAGAAGGCCGCGTGTCGCGCTGACAGCACAATGCGGAGCAGAGCAGGCGG--------
Tarbinskiellus_portentosus     ------GGAGGGAGACGAGCGCCGCGATGAACTTGTCCCCGCGTGTCGCAATTGATCGGGAGCAGAAGGCCGCGTGTCGCGCTGACAGCACAATGCGGAGC------------------
Acheta_domesticus              GGGCGGAGAGAAAGACGCGCGCCGCGATGAACTTGTCCCCGCGTGTCGCAATTGATCGGGAGCAGAAGGCCGCGTGTCGCGCTGACAGCACAATGCGGAGCAGTGCAGGCGGCGGCGACG
Gryllus_bimaculatus            GGGCTGGGAGAAAGACGCGCGCCGCGATGAACTTGTCTCCGCGTGTCGCAATTGATCGGGAGCAGAAGGCCGCGTGTCGCGCTGACAGCACAATGCGGAGCAGAGCAGAGCAGAGCAG--
Consensus                             ***  ***** ****************** *** ************************************************************

Teleogryllus_occipitalis       ---------------------------------------------GGTGGCGGTG-GCGGCGGTGGCGGTGGCG----------------GCGGCGGCGGCGGCGGCGGCG------
Tarbinskiellus_portentosus     ---------------------------------------------AGTCGGGGCG-GCGGCGGCGGCGGCGGCG----------------GCGGCGGCGGCGGCGAAGGCG------
Acheta_domesticus              GCGGTGGTGACAGCGGTGAACGCAGTCGCGGAGGCAACGAGGGCCAAGGTGGTGGGG-ATGATGATGGCGGTGGCG----------------GCGGCGGCGGTGGTGAGGGCG------
Gryllus_bimaculatus            --AGCAGAGCAGGCGGCGCGGCGGGCTGCGCGGGCGGCGC-------GGTAGCGGTGAGCGGCGGTGCTGGAAGCGGTTGACGGTAAAGCGGAGCGGTGGCCGGGGGCAGGGCGCGGGTG
Consensus                                                                  ** * ** *   *  *  *  **  ***                  **** *** * **    ****

Teleogryllus_occipitalis       ----------GCGGCGGCGGCGGCGGCGGTGGCGGTGGCGGCG-GTGGCGGTGGCGGCGGCGGTGGCGGTGGCGGCGGCGGCGGTG---------GCGGTGGCGGCGGTGGCGGCGGTGG
Tarbinskiellus_portentosus     ----------GTGACGGCGGCGAG--------------------------------------------------------------------------------------------
Acheta_domesticus              ----------GTGGCGGTGGTGGTGGTGGTAATGGTGGTGGTG-GCGGTGGTGGTGGCGGCGGCGGTGGTGGCGGCAGTGGGGATG---------GCGACGGTGGAAGTGGCGGCGGTGG
Gryllus_bimaculatus            TGTGGGGCGCATGGTGGTGACGTGGTGTGTGGCGGAAGGGGTGCGCTGCGGGCGCGGCGAGCCTGGGGAGCGGGGCTACAGGGGCGGGCCATGGCGCGGCGGTGGACGGATCGGGCGCGG
Consensus                                 *  ** *  *

Teleogryllus_occipitalis       CGGCGGTGGCGGCGGTGGCGGCG------------------------------------GCGGTGGCGGCGGTGGCGGTGGTGGCGGTGGTGGCGGCGGTGGCAGCATCGGTGGTGGTGG
Tarbinskiellus_portentosus     -----------------------------------------------------------GCGGTAGCGACGATGGTGTCGTTGGCGATGGCAGCGTAGGTG-----------GGGAGCCG
Acheta_domesticus              GAATGGCGGCGGTGGTGGGAATG------------------------------------GCGGCGGTGGTGGTGGCGGCGATGGTGGTGGTGGCGGAGGCGGTGG-AGTGGGGGTGGCGG
Gryllus_bimaculatus            GTGGAGTGACAGTTGTACGGACATGACGGAAGGCGTGGCATGCAAAACGGCGATGAGGTGCGGCGGTGGAGGTGGCGGCGGTGGAGGCGGCGGCGGTGGTTTAACCCGTGGCGGAGGCAG
Consensus                                                                                 ****  * *  * *** *  * *** *  **  ***  **              **  *  *

Teleogryllus_occipitalis       CAGTGGCTATGGAGGCGGG-----------TGAGGTGACTGTGGAGGCGCTGATAGGGA----------ATGCAAC--------------------AAGTGTGACAACGGAAAAAGCAAT
Tarbinskiellus_portentosus     CTGTCCACTCCGAATCCGCTC-----CCATTGCATTAAAACCTCAAGCCCTATACGTGCCCTCCT----CCACAATTTCATTTCCGTTTCCCCCTTGAACGTCTC--------------
Acheta_domesticus              CAGTGGCGGCGGGGGCGGGGCGGAGGCTATGGACGTGACGGTGGAGGCGGTGGTGGGGATGACAGGGGGGTGCAACT-----------------AGAAATGTCACAAAGGAGAAAGCAGT
Gryllus_bimaculatus            CGATGGCTGTTGAGGCGGC-----------AGAAGTGGCAGTGGAAGTGATGACAGGAG----------ATAC-----------------------AAGTGTGACAACGCAGAAAGCAGA
Consensus                      *  *       *   * *               *   *         * *   *    *                 *                        *  **  *

Teleogryllus_occipitalis       TGTGGAGATAGAGACGAAGGTCTTGACG----AGAAAGGGTAATTCAAAAG---AAATCCCTTGACAGT------------------------------------------------
Tarbinskiellus_portentosus     ----------------CGCTCTTAGCATTTAGAAGTACGTGGTTCCGCAGTCCCAACCCAATGACAGTGAGACGCTCCGTATCGATTCCCTCAATGTCTTCGTAACTTCCTCAGTCCAT
Acheta_domesticus              GGTGGAGATATAGACAGAAGTCTTGAC------GAAATAGTGGTTCCAGAG---GAGCCCCATGTGGGC------------------------------------------------
Gryllus_bimaculatus            GGTGGAGATAGAGAC--AAGTCTTGGCG----AAAAAGGCTGGTTCCAGAG---AAGTCCCATGACGGC------------------------------------------------
Consensus                                       ****  *       *     *  ***    **     *  **  **    *

Teleogryllus_occipitalis       ---------------------------------------------------------TTGTGGGAAAATGTTCCACTT------------------------------------------
Tarbinskiellus_portentosus     TTTCCTGTTCGTCGTCCTTTGGATAATTTTACTCCGATGTCCAGGTGATTCTGCCATTCAATCACAATTTTCTCCTGTTTTTTTTTTTTGTTTCATATTTATCATTTCTCTCCTCTCCCAC
Acheta_domesticus              ---------------------------------------------------------TTTGGGAATAATAAACCCCCCTT-----------------------------------------
Gryllus_bimaculatus            ---------------------------------------------------------TTTAGGTACAGTGTCCCCCCTTT-----------------------------------------
Consensus                                                                                 **     * * *   *  *

Teleogryllus_occipitalis       ------------------------------------------AAATGTTTAAACAGTCATTAATATCCTTTAAT---------------------------------------------
Tarbinskiellus_portentosus     TTTTCCCGTTCATTTCATTCTTTTTCTTTTTCATTGCTCGCTTAATTTGTTTTACCATTTCTTCTTTCTCTTTGTTTACTTCGTATATCACAATCTAATTTTAATGTCTCACTTCTCCTG
Acheta_domesticus              ----------------------------------------AAAAAATGTTTTAACTCCCAATATTTTTCTTTAAT--------------------------------------------
Gryllus_bimaculatus            ---------------------------------------AAAAATACGTTTAAACTGTCATTAATACCCTTCACT--------------------------------------------
Consensus                                                                  *   **** * *       *  *    ***    *

Teleogryllus_occipitalis       -----------------ATGACAGGAGAATGAGATAGAAACGGCGGAGAGGAAGACGGAAATGTAGACGTGAGCAGAGAAGATGAGAAATGCGAAGGC-----------TGAGGTGA-
Tarbinskiellus_portentosus     ACCTATTCTAATGTTTTAAATTTTAATTAAGGAATTACTGCCGACAAAG-TGCTGACAGGGGTGGAGAGGGGAGGGTGACCGCAAGGAGGCGCCGAAGG--------------------
Acheta_domesticus              -----------------ATGACGGGCGAAGTAGTTAAAAAGGGTGGGGAGGATGACGGGAATGTAGACGTGAGCGCGAATGATAAGATATGCAAAGGC-----------CGAGGCGAA
Gryllus_bimaculatus            -----------------AAGATAGGCGAAGGAGTTAGAA--------------ATGGGAATGTCGACGAGAGCGGAGATGACGAGATACGCGAAGGTGATGCAAATATGGGAGCGGAT
Consensus                                       *          **  *  **                   *  *   **  ** * ***       *    **   **  * *

Teleogryllus_occipitalis       ------------------------------------------------------------------------------------------------------------------------
Tarbinskiellus_portentosus     ----------------------------TGGCGGCGGTGGACGAGAAAGACGAGGA--------------------CGAGGACGAGGACGAGGAC-------------GAGGTGGAGGT
Acheta_domesticus              GTAGGGAAGTCAATTGGCGTGAGGACCATAGAATGGGAGATTAAGAAGAACGAAAAAATGATAGCTTAGGTGCAATTGGGGTTGAATACGGAAAT---------------AATGATGGT
Gryllus_bimaculatus            GTGGAATCGGTAGTTGAGGCTAGGGCCACAGAATCGGAGTTTAAGAAGAATGTAAAAGTAATGATGTATGTGCAGTTGAGGTTGAATGCGGAAGTGATAAGCGATGAGAGAAGTGATGAG
Consensus

Teleogryllus_occipitalis       --------CTATGGGAGCAGATGGGAAATCAGTAATTGAGGCGAGGA------------------------------------------------------------------------
Tarbinskiellus_portentosus     GGAGGTGGAGGTGGAGGTGGAGGTGGAGGTGGAGGTGGAGGTGGAGGTG------------------------------------------------------------------------
Acheta_domesticus              TGTAGAGGGAGTGGAAGTGGACGTCGAGTTAGAGACTGTGACAAAGACGTAAGAGTCGCCTCCGTGGCCGAGATCGCCAAGCCACTAGTGCCGTAGTGATTTCGGACCGAGGAGTCACGG
Gryllus_bimaculatus            TGAAGAAGAAGTGGAAGTGCAGGTCAGGTCAAAGGCTGAGACGAAGACG------------------------------------------------------------------------
Consensus                                ***  *   * *              * *      *

Teleogryllus_occipitalis       ------------------------------------------------------------------------------------------------------------------------
Tarbinskiellus_portentosus     ------------------------------------------------------------------------------------------------------------------------
Acheta_domesticus              GTTCAAGTCCTTCCTCGGGCATGGTTGTATGTGTTCGTCCTTTCACCCCTATCATCACCACACCATTTCATCCTTATACCATAACGTCAAATCCAATAGTAGTCGCTAATGACTTAGTAG
Gryllus_bimaculatus            ------------------------------------------------------------------------------------------------------------------------
Consensus

Teleogryllus_occipitalis       ------------------------------------------------------------------------------------------------------------------------
Tarbinskiellus_portentosus     ------------------------------------GAGGTG------------------------------GAGGTGGAGGTGGAGGTGGAGGTGGAGGTGGAGGT------CACTC
Acheta_domesticus              ATGATGCGAATGCTTAAATAAGAAAGAAAAGACGAATGGGGTGACCTCTAATGGCAATGATGG------GGAAGGTGGTGGAACCGATGGTGGAGACCAAGATGAAGGTATTCTGCTGTC
Gryllus_bimaculatus            --GAGTCGAATG------------------------GGGGTGGCCGCAAGTTGCAATGATGGCGGCGTCGAAAGTGATGGATCTGATGGTGGAGACAAAGATGGAGGTATTCTGCGGTC
Consensus

Teleogryllus_occipitalis       ------------------------------------------------------------------------------------------------------------------------
Tarbinskiellus_portentosus     C-----GAACCCGACCCCGAAACCTGAAGCCCAGTCCTTCCATTGAGCAATA--------------------------------------TCTGTTTATCTTGCGTTTTCTTGCCTGAA
Acheta_domesticus              C-GGACGAACCCGGCTTT---------GTCCACTCCTTCCAGAGGAAAAAGCCTCCGACTTTACGCTAAGTCTCAGGCTCGTTCCACCGCTTTATTAGCGCTGCGCTCTCCCTCGAGGA
Gryllus_bimaculatus            CGGGAGGAACCCGGCTGT---------GTCCACTCTTTCCAGTGGAAAGAGCAGTTGTCTTTATGCTAGGTTTCTGGCTCTTCCCACCGCTTTATTAGCGTTGCGCTCTCCTTCGAGAA
Consensus

Teleogryllus_occipitalis       ------------------------------------------------------------------------------------------------------------------------
Tarbinskiellus_portentosus     TATATAACGC-----------------------------------------------AGAGTGAATATTATTTATTGAGGTGTGGC--------------------------------
Acheta_domesticus              C-TTCTGTGCTCGCCGAGCTTTCCAATTGGTGGGTC-----------------------CACGAGGGTCTTGCCTCCAGATGTTCCACTGTATCTTCTATATTGGCGGGGGCGGCGGCT
Gryllus_bimaculatus            ----CAGTGCTCGCCAAGCTTTCCAATTTGTGGTTCCAGTGAGGTCCAAATGCCAGGGAGCAGGAGAGTCTTGTCTCCAGAGGTTCCGC-------------------------------
Consensus

Teleogryllus_occipitalis       ------------------------------------------------------------------------------------------------------------------------
Tarbinskiellus_portentosus     ------------------------------------------------------------------------------------------------------------------------
Acheta_domesticus              GCGGCGGGGCGGTGGCTGTGGAGTTGATAGTGGAGGTGGTGGAGGCAGTTGTAAGTGGGGGCCAAAGACAAATATGACAACGGAGAAAGCAATGACAGAGATAGAGACAGTCGTCTTGAC
Gryllus_bimaculatus            ------------------------------------------------------------------------------------------------------------------------
Consensus

Teleogryllus_occipitalis       ------------------------------------------------------------------------------------------------------------------------
Tarbinskiellus_portentosus     ------------------------------------------------------------------------------------------------------------------------
Acheta_domesticus              GAAGTGGTGGTTCCAGAGAAGTACCATGAGGACTTAAGGGGTAATGTCCCCCCTTTAAAAAATGTTTTAACTGCCAATATTTTCCTTTAACTCTCTAGTACCCCTTGTTACACATATGTA
Gryllus_bimaculatus            ------------------------------------------------------------------------------------------------------------------------
Consensus

Teleogryllus_occipitalis       ------------------------------------------------------------------------------------------------------------------------
Tarbinskiellus_portentosus     -------------------------------------------GAACGAAATATATTTTTTGCTTGCCACGCTCAATTCAT--------------------------------------
Acheta_domesticus              ACAACGTTTCTGGTCAACTCGTATTTCCAGGTTTTTGCACTAGTTTGCAAAATTTATCTTCTATTTCTCTTTCTCGTATTCCTTTCCATATTTTCCTTTCTGTGTGCCCAACAAGTCAGT
Gryllus_bimaculatus            ------------------------------------------------AAATCTAT-TTCTCTTCTCTCCTCTCGTATTCTTTTCGAAGTTTTCCTTTCTCCTCGCCCAACAAGTCAGT
Consensus

Teleogryllus_occipitalis       ------------------------------------------------------------------------------------------------------------------------
Tarbinskiellus_portentosus     ------------------------------------------------------------CGTATATGTCGTAATCTTTGTT--------------------------------------
Acheta_domesticus              TGGTATTTCTAAAATATTTTCTTCGTATTCTTTCTCGCGTCTTCCTTTGCGGTACTCTGAAACATTTATATTGTTCTTTCAGTT--TCTCACAAATATTTTCTACTTTTTTTAATTTCGT
Gryllus_bimaculatus            T----TGTCTAAAATATTTTCTTGGTATTCTTTCTTGCGTCTTCCTTTGCAAT-TGGTGACACATTTGTATTGTTGTTTTCATTTATCTCAAAAATATTGTTTGTGTTCCTTTATTTCGT
Consensus

Teleogryllus_occipitalis       ------------------------------------------------------------------------------------------------------------------------
Tarbinskiellus_portentosus     ----------------------CATACGTTTC----TGATACCTTCTTGGAGTTGGAATTAAGATGAAGGTGTACAACGCTTTGTGATATCATGAGTCATCA---AAGCCACA-AAAAA
Acheta_domesticus              CAGCTACACACCGATTATTTCCTTGTATATTTTATC---TTATGTTTTCAGCCGTAGAATTATAAC-AAGTTCTGCAGCGATCTTAGTTATCCTGAGTCATCATCTTCACCGCT----AC
Gryllus_bimaculatus            CTGCCACATCCCCATTCGTTTTTTGTATATTTTATCTTGTTACGTTTCTAGCCCTGTAATTATAAC-TAGTTCTGCAGCGATCTGAGTTATCCTGAGTCATCGTCTTAGCCGCTCGAGAA
Consensus

Teleogryllus_occipitalis       ------------------------------------------------------------------------------------------------------------------------
Tarbinskiellus_portentosus     AGAAAGGGAAGTATGAATGAAAGGGTAGATAAATACGAAGTAGCTTGCAAGTTCTCCGTCAGGCGCGTGATGACAAAGTACACGATGCACTCGCCACAAATAAGTTACGGGTCTCCAATC
Acheta_domesticus              AGAAAGAGAAAAATGAATAAGACGATGAGTTAATACAAAACAACTTCCTCCTATGTCTTCACACAC---ATCACAAAGTACACGATGTATACGTCGCAAATAAGATACTAGTCTCCAGTC
Gryllus_bimaculatus            AGAAAGAGAAAAATGAATGAGACGATGAGTTAATACAAAACAACTTCCTGCTATGCCTTCACACAC---ATCACAAAGTACACGATGTGTACGTCGCAAATAAGATACCAGACTCCAGTC
Consensus

Teleogryllus_occipitalis       ------------------------------------------------------------------------------------------------------------------------
Tarbinskiellus_portentosus     CATTTGCAAATAGGGCCCGGGCAATGAATCACTCGGAAGGGGAGCGGTCTCCGCACACGGCCTTC-TGATGAGAGCATTGTCCGTCTTTATCGAGCGTACGTTTACGCGTGCTCACGCCG
Acheta_domesticus              CACTTACAAATAGGGACTCGTCCATGAATCATTCTGA---GAAGTGGTCTCCACATACAATCGTCATGATTACGGTATTGTGCTTCTTTATCGAGCACACTTTTATGCAGGAGATCTGCG
Gryllus_bimaculatus            CACTTACAAATAGGGACTCGTCCATGAATCATTCTGA---GAAGTGGTCTCCACACACAATCGTCATGATTACGGTATTGTCCGCCTTTATCGAGCACACGTTTATGCAGGCGAACTGCG
Consensus

Teleogryllus_occipitalis       ------------------------------------------------------------------------------------------------------------------------
Tarbinskiellus_portentosus     CAAC-------CGATGTCCGCGACCAATCGCAGAGCGCGGGCCCGTT--GCCCGGAGGAGCGGAGCCGCTTTCAAGTGGGCCAATTGGAGCACCTCGAGCCGTCCGCGGGCAGCGCCCGT
Acheta_domesticus              CAACAGGTCCACAATGCGTGAAACCAATCGAAGTGGGAAACGATGGACCGTGCCGTGGCGCGCGCTATGGTCTGGAATGGAGAAAAAGTGGGCAAAAA---------------------
Gryllus_bimaculatus            CAACAGGTCCACAATACGTGAAACCAATCGCACTGGGCAATGACATG-----------GCAGACTATGGTCTGGATTGGGGAAAAAGAGGACAAAAAGTGATCTGCATG----------
Consensus

Teleogryllus_occipitalis       ------------------------------------------------------------------------------------------------------------------------
Tarbinskiellus_portentosus     CACCGCTGCCATCCCCGCGTCTGCCGCCCCCACCCCCACCTACACCCCGCTGCCCCCGTCTCTCGTCCCCGCTTCCCCCGTCCTGCCCCGCCCCCCCCGCCACCGCAACGAGACAGCGTC
Acheta_domesticus              ------------------------------------------------------------------------------------------------------------------------
Gryllus_bimaculatus            -------------------------------------ATCTACGTGTCGCTGCC------------------------------------------------------------------
Consensus

Teleogryllus_occipitalis       ------------------------------------------------------------------------------------------------------------------------
Tarbinskiellus_portentosus     AGCCTCTGTGGTGATGGAGTGGTCCGACCACGACCTTCTGTCCACAGGTGTACGGCGCAAGGATCTGCAAATGCAAGTGGCGGAAGATTCGATTCGATTTTTCGAGTGATCCTTGCATTT
Acheta_domesticus              ------------------------------------------------------------------------------------------------------------------------
Gryllus_bimaculatus            ------------------------------TGATCTCT-----------------------------------CAACTGTCGACCAACTCCATCTTCTTCTT-----------------
Consensus

Teleogryllus_occipitalis       ------------------------------------------------------------------------------------------------------------------------
Tarbinskiellus_portentosus     GGAGTTTGATCGAAAATTCGATGTACGAGTAATCGAAGTTTTTCAATATATATATTTTTATGAAAAAGACGTTCGGTTTGATCGATGGTTTTACATTCGGTTTGTTAAAGAATCGAGTTGCT
Acheta_domesticus              --------ATCATAAATTCTGT----------------TGTGGCAGCAGAGATCTT----------------------------------------------------------------
Gryllus_bimaculatus            ---GTCCCATCGTTGAAATTTT----------------TTTTCCAGTAGACATATTGTAGTATAAGGTGT-----------------------------------GACAGTCGAGCAGTC
Consensus

Teleogryllus_occipitalis       ------------------------------------------------------------------------------------------------------------------------
Tarbinskiellus_portentosus     AGAGGCCAAATGAAGTGGTGGACGATTGATTCGGTTTTTTGAGCGGCGAATTCAGATTTTTTAAATTTATTTAAATTATAACGCGCACACACATGCACACAGGTATTATATATAGTTCAGTT
Acheta_domesticus              ------------------------------------------------------------------------------------------------------------------------
Gryllus_bimaculatus            TGTGCTTAAAT------------------------------------------TTTTTTTTCAGCCATGCAATCTATACTTCA--------------ATATTTTCCAGACATCGATT
Consensus

Teleogryllus_occipitalis       ------------------------------------------------------------------------------------------------------------------------
Tarbinskiellus_portentosus     GACTTTATACTACACGTATGTATTGAAGGCTTATTAGTAAATCTAAACTAACTTAAAAATAAGTACTTTGTAAGAGAAAGGAAACCCATAAGTTAACTCTTTAAAAGTCATTTAAGTCTA
Acheta_domesticus              ------------------------------------------------------------------------------------------------------------------------
Gryllus_bimaculatus            CTTTCTGTCCTGCGAGTA------------------------------------------------------------------CCACAAG-----------------ATTCAATTCAC
Consensus

Teleogryllus_occipitalis       -------------------------------------------
Tarbinskiellus_portentosus     CAAATTTTTTTTTTAAGTTGATATTTATTGGTTATTCTTGTTG
Acheta_domesticus              -------------------------------------------
Gryllus_bimaculatus            CTCCTTATTTCCTAATG--------------------------
Consensus
```

**Tocc.2G0023520.1_exon1**

```
Teleogryllus_occipitalis        ------------------------------------------------------------------------------------------------------------------------
Tarbinskiellus_portentosus      ------------------FFGFLTCAS---------RAVSFHTGLR*IPTAHKRAPGEKDVAGVGRA-----SGRCGPGAAGPEAEAEAEALEGASRAGPSSPP--------------
Acheta_domesticus               LVSPTSGERRYSMGDETGDFEFCSEASFRIEDILHAANLLTWDSRLYFLSEARRA----KDFF-VPKNP----WLRSG---LNPRTWV*SDTLTVT----PSSPYLSTALRRRMRLQGLN
Gryllus_bimaculatus             ---------------------YISS----LQNFLLILVEVAYVEKKCF*AVSHRR----NSFIGIIKTPPPF*SLR--------LSWTARDKVRGT----SYLPA*EASC*N--NVN*IN
Consensus

Teleogryllus_occipitalis        ----------------------------------------------------------------------------------M------------------------------------
Tarbinskiellus_portentosus      -RVLVRCGFVRRAEAAGAGRGKV------AGRL*N*------------------ARGAFPRGGPRWREY*CFN----FGSVPPLRAVAGRGGGARSERSGPTGSPGTRGAPTSKRRISDQ
Acheta_domesticus               GECVR-----RR*GCSIPVKESI-LESIPESKS*NL---------------------F*NRIERKESI------------P*--------GIESLENRE-----FLPSRTV-------
Gryllus_bimaculatus             -ECL*LTILTRRTAVKKK*KSFVC*TS*LESKLINLL*T*EYTITKSRYFRFKIFKFTFN*KLEI*EIYYCI*RDNLFSLVEKM--------GINVAKFGEIIVI*YFSYKTVMRQISRG
Consensus

Teleogryllus_occipitalis        -----------------SF---------------------------------------------------------------------------------------------------
Tarbinskiellus_portentosus      K-CNECWTFLNNDEMNASFFIVFSIRNVPVCF*W*VIYVRCGG**FPHWYRSKYIRAYSPLTWK*YV--------DWRP-----FLKVP-----------F**GYF*EIKNLKIIIGSL
Acheta_domesticus               --SNIDLLLMRSS----SF---ASL*CI*AC----------------------AEGR*CLTFEVGKSP*CDSREEWKPGSSNLYLIKHN*MLIN*I**K**LTY*FNENK---------
Gryllus_bimaculatus             NYSRVIITFMFNSR--ASF---*GRRCANCC----------------------ILSYLPVTL-----------SWLPPD---YLNTD-------------VHYL*RNE---------
Consensus                                        **

Teleogryllus_occipitalis        ------------------------------------------------------------------------------------------------------------------------
Tarbinskiellus_portentosus      FSLYNFK---PWK*LQKNCPRF--------------IPPDS--LLLLLPYLLSHCLASKREIR-----DSKFYFLSA*IGQFWSIVWRE-*GGRGGGGHEG*LGCRGRGDPPLQPVAVV
Acheta_domesticus               FNKI*MKYSAPWPRLLS-------LLWRSRAVGSRVCVPPRA-WVYLCL------CVCV--------------CVSLSHPS*SSSIHLPYPHG*------------------QPWCV-
Gryllus_bimaculatus             FEPCRPR----WPGWLRACTDSALLFERSRAR-----IPPWARWFVPCLP-----CMCCRGGFRVLLTLDKRSPC*CVLVPSGWSNI----PSGR------------------KG*PTV
Consensus

Teleogryllus_occipitalis        ------------------------------------------------------------------------------------------------------------------------
Tarbinskiellus_portentosus      VVAGRRLGRWRGREGGRWLRPRPRELSRV*PTC**VGD*GRRSSSPAPLRRDGWLYFG----RL-----DLSNNAPYYAGVSASSVACVFQKA--------LDATPLFPNESKSWRL*AV
Acheta_domesticus               -----KMDSAE-------ILIPPNNV*RR---EMHVRVVHRG-------------------RV-----AKKNDEVIQSSKSVTKSFTIYN----------I*KRPQR*KR*-------I
Gryllus_bimaculatus             -G*GKKFISYT-------*RIKFNSEARLVTFSA*FRVLSRGAKTP*A---EVALFYAIISERVLEVFNTKQ*IEVHESGGECHHFIVVYQRLHLIGDGLPLPSTPHS*SEN-------I
Consensus

Teleogryllus_occipitalis        ------------------------------------------------------------------------------------------------------------------------
Tarbinskiellus_portentosus      VAILQR--N*FF----YVLTHLS*NDFE---------I*FILRLV---------------------------KN------LI*RK*RNLLCEILNENSVNNVL-------------C
Acheta_domesticus               K*IIEK-*NVRFCF*VFV*ISKNKNDYSTKTNI*NSIFFNFLLHIRCL---IVIFKLCSCYYLCYDYVLRCIF*KNIHRCSKLLF*KGSKQ------EKFW*SSI*AIGEQKVPDSFLPC
Gryllus_bimaculatus             RIFLHSFHNLLY----FARICATEN*LS---------LLILLHLILLTSSM*IILL*NLFLFNFKYIT---------------YSGSNNPAEASGQNEYCKNLLSLGHSQ-------C
Consensus

Teleogryllus_occipitalis        ------------------------------------------------------------------------------------------------------------------------
Tarbinskiellus_portentosus      ANIRHFTLWFPHFPSSY*TYESYKQI-----VLP--TWHI*MNLRPATI*F-------------LTNQEEKY------------ELMNEKTHP-GRYQYRKART*P**VVGS----KHL
Acheta_domesticus               AIMRT--------HSSF*ISST*PMTLAVVSLLP--S*IIYTSLSYIFC*FL*DVYMMEKIN*SRLTQQEQFYQF*KNATTILHLGYLTAHVTKYSRTSYLSAREASYCCYNNVK*INKC
Gryllus_bimaculatus             AVI**--------TGHY----THPRL-----LLPLICSMLFQSIKDPFAIY----FYIRKQNPH----------GKRLKTKLPRELFNAH*HK--SMSFFSL*TTVY---------NHC
Consensus

Teleogryllus_occipitalis        ------------------------------------------------------------------------------------------------------------------------
Tarbinskiellus_portentosus      **VSK-----------------------------------------------R*LNE*LH----------------------------------------------ITIYPLTVY*S
Acheta_domesticus               VTIKK*NLVRVYQLL*ERRGTVLHDYLEKSFGSD*EEAIV*NLWKWWWKSRFHRSIATSAHDTWWIY*CLPHHTRRVKHLCFTPLVLCCSRKWVVSARICATRN*LPLLILLYLILLKPS
Gryllus_bimaculatus             MVL---------------------YLKHNFCE*YTKYV--NIYKYY--------I*T-----------------------------------------------ISMYLLCV*TG
Consensus

Teleogryllus_occipitalis        ------------------------------------------------------------------------------------------------------------------------
Tarbinskiellus_portentosus      --------------------------------------------------------------------SNPSPQIGRYYKY*DGM*ITTKGKLFFPGWTSFGRITLCLKRVCENVASRA
Acheta_domesticus               NYLMMKLTLSKFKYVTS*K*IVNQHNSVWRQ*SEQT**MKKSISE*GICNVFKEFATFP*QTPR-----KNENSTEGR-------M*RTTVSYLFPSKKRSFT*KTFLFSEGPSNLFSSV
Gryllus_bimaculatus             ---------------STAKARINLKN-----------------QRCICNGYKKYSTSTHTIPLTAVVWTNENSKEGR-------M**MTVNYLFP-*NLEFHFQTMLV*EGSSNLFSRV
Consensus

Teleogryllus_occipitalis        ------------------------------------------------------------------------------------------------------------------------
Tarbinskiellus_portentosus      WKCGKPYLRIWKWPDVFNGVVLRAG---------------------------ERGRG----GAGRGGAEGDERRDELVPACRN----------------------*SGAEGRVSR*QH
Acheta_domesticus               ---SVSFLVSW---DV-NILVRAEGPRKRRRPRMEMREALGPHLEAAWLAL*IERGRARTGRWRGRGRGRGRGRRERRAPR*TCPRVSQLIGSRRPRVALTAQCGAVQAAATAVVTAVNA
Gryllus_bimaculatus             ---LVSFLVWW---DV-NFLVCGEGLLAAYGNAGRAQSAYGGGL-ALWIGR--ERGWGAVEAGRARGRRRGAGRKTRAAMNLS------------PRVAIDRE----QKAACRADSTMRS
Consensus

Teleogryllus_occipitalis        ------------------------------------------------------------------------------------------------------------------------
Tarbinskiellus_portentosus      NAEQSGRR-------RRRRRRRRR----------------------------------------------------------------RRR*R----RRGG------SD--------
Acheta_domesticus               VAEATRAKVVGMMMAVAAAAVVRAVAVVVVVMVVVAVVVAAAVVAAVGMATVEVAAVGMAAVVGMAAVVVAAMVV---VAEAV--------EWGWR--QWRRGR---------------
Gryllus_bimaculatus             RAEQSRAE-------QAARRAARA-----------AR*R*AAVLEAVDGKAERWPGAGRGC--------VGRMVVTWCVAEGVRCGRGEPGERGYRGGPWRGGGRIGRGWSDSCTDMTEG
Consensus

Teleogryllus_occipitalis        ------------------------------------------------------------------------------------------------------------------------
Tarbinskiellus_portentosus      ---------DGVVGDGSVGGEPL------------------------------------------------------------STP-------NPLPLH*NLKPYTCPPPQFHFRFPLERLRS*
Acheta_domesticus               ---------GGGYGRDGGGGGG-------------------DDRGVQLEMSQR----RKQWWRYR---QKS*RNSGSR-GAPCGLWE**TPLKKCFNSQYFSL----------I*RAK*L
Gryllus_bimaculatus             VACKTAMRCGGGGGGGGGGGGLTRGGGSDGC*GGRSGSGSDDRRYKCDNAESRGGDRDKSWRKRLVPEKS--HDGFRYSVP--------PLKIRLNCH*YPS----------LR*AKEL
Consensus

Teleogryllus_occipitalis        -------------------------------------------SIRN-----------------------------------------------------------------------
Tarbinskiellus_portentosus      HLEVRGSAVPTQ*Q*DAPYRFPQC-------------------LRNFLSPFSCSSSFG*-------------------------------------FYSDVQVI--LPFNHNFLLFFF
Acheta_domesticus               KRVGRMTGM*T*ARM---IRYAKA----EAK*GSQLA*GP*NGRLRR-TKK**LRCNWG*IRK**WL*REWKWTSS*RL*QRRKSRLRGRDRQATSAVVISDRGVTGSSPSSG------M
Gryllus_bimaculatus             E-----MGMSTRAEM---TRYAKVMQIWERMWNR*LRLGPQNRSLRR-M*K**CMCS*G*MRK**AM-RE----------------------------VMSEEEVE-------------
Consensus                                                          *

Teleogryllus_occipitalis        ------------------------------------------------------------------------------------------------------------------------
Tarbinskiellus_portentosus      LFHIYHFSPLPLFPFISFFF-----------------------FFIARL------------------------------------------ICFTISSFSLFTSYITI*F*CLTSPD--L
Acheta_domesticus               VVCVRPFTPIITTPFHPYTITSNPIVVANDLVDDANA*IRK----KRRMG*PLMAMMGKVVEPMVETKMKVFCCPDEPGFVHSFQRKKPPTLR*VSGSFHRFIS------AALSLEDFCA
Gryllus_bimaculatus             -VQVRSKAETKT--------------------ESNGGGRKLQ*WRRR----------K*WI*WWRQRWRYSAVREEPGCVHSFQWKEQLSLC*VSGSSHRFIS------VALSFEN-SA
Consensus

Teleogryllus_occipitalis        ------------------------------------------------------------------------------------------------------------------------
Tarbinskiellus_portentosus      F*CFKF*L--RNYCRQSAD-----RGGEGR----------------VTARRRRRWRRWTRKTRTRTRTRTRWRWRWRWRWRWRWRWRWRWRWRWRWRWRWRWRWRSLRTRPRNLK-------
Acheta_domesticus               RRAFQLVGP-RGSCLQMFHCIFYIGGGGGCGGAVAVELIVEVVEAVVSGGQRQI*QR--RKQ*QR*RQSS----------------------------*RSGGSREVP*GLKG*CPPFK
Gryllus_bimaculatus             RQAFQFVVPVRSKCQ---------GAGESC--------------------------------LQRFRK-------------------------------------------------
Consensus

Teleogryllus_occipitalis        ------------------------------------------------------------------------------------------------------------------------
Tarbinskiellus_portentosus      ----------------------------P----------SPSI--EQYLFILRFLA*IYNAE*I--LFIEVW-R-TKYIFCLPRSIHRICRNLCSYVSDTFLEL--------------
Acheta_domesticus               KCFNCQYFPLTL*YPLLHICNNVSGQLVFPGFCTSLQNLSSISL--SRIPFHI-FLSVCPTSQLV---FLKYFLRILSRVFLCGTLKHLYCSFSFSQIFSTFFNFVSYTPIISLYILS-Y
Gryllus_bimaculatus             ---------------------------------------SISLLSSRILFEV-FLSPRPTSQFV*NIFLVFF---LASSFAIGDTFVLLFSF-ISKILFVFLYFVCHIPIRFLYILSCY
Consensus

Teleogryllus_occipitalis        -------------------------------------------TSYY-----------------------------------------------------------------------
Tarbinskiellus_portentosus      ----ELR*RCTTLCDIMSHQSHKKRKGSMNER------------VDKYEVACKFSVRRV--------------MTKYTMHS--------------------PQISYGSPIH------
Acheta_domesticus               VFSRRIITSSAAILVILSHHLHRYRKRKMNKT------------MS*YKTTSSYV----------------FTHITKYTMYT--------------------SQIRY*SPVH------
Gryllus_bimaculatus             VSSPVIITSSAAI*VILSHRLSRSRKKEKNE*DDELIQNNFLLCLHTHHKVHDVYVANKIPDSSPLTNRDSSMNHSEKWSPHTIVMITVLSAFIEHTFMQANCATGPQYVKPIALGNDMA
Consensus

Teleogryllus_occipitalis        ------------------------------------------------------------------------------------------------------------------------
Tarbinskiellus_portentosus      ---LQIGPGQ*ITRKGSGLRTRPSDESIVRLYRAYVYACSRRNRCPRPIAERGPVARRSGAAFKWANWSTSSRPRAAPVTAAIPASAAPTPTYTPLPPSLVPASPVLPRPPRHRNETASA
Acheta_domesticus               ---LQIG--------------------------------------------------------------TR---------------------P*IILRSGL-------------
Gryllus_bimaculatus             DYGLDWG-------------------------------------------------------------KRGQKVICMIYVS----------LPDLSTVDQL-------------
Consensus

Teleogryllus_occipitalis        ------------------------------------------------------------------------------------------------------------------------
Tarbinskiellus_portentosus      SVVMEWSDHDLLSTGVRRKDLQMQVAEDSIRFFE*SLHLEFDRKFDVRVIEVFQYIFL*KRRSV*SMV--LHS--VC*RIELLEAK*SGGRLIRFFERRIQIF*IYLNYNAHTHAHRYYI
Acheta_domesticus               --------HI---------------------------------------------QSS*LRYCASLSSTLLCRR--------SAQQV----------------HNA-*NQSKWET
Gryllus_bimaculatus             --------HLLLVP--------------SLKFFFQ*TYCSI--RCDSRA--VCA*IFFSAMQSILQYFPDIDS--FCPA--------STTRF---------------------NSPPYFL
Consensus

Teleogryllus_occipitalis        --------------------------------------------------------
Tarbinskiellus_portentosus      VQLTLYYTYVLKAY**I*TNLKISTL*EKGNP*VNSLKVI*VYKFFF*VDIYWLFLL
Acheta_domesticus               MDRAVARAMVWNG--------------EK----VGKKS*ILLWQQRS----------
Gryllus_bimaculatus             M-------------------------------------------------------
Consensus
```

```
Tocc.2G0023520.1_exon2
Teleogryllus_occipitalis            ------------------------------------------------------------------------------------------------------------------------
Tarbinskiellus_portentosus          --SSVF*PAPPELSAFTQDF------VKFRPRTKGPRARRTWRGSGGRAGGAAPGRRGRR--------------QRRRRR----------RSRARRGPGRPRRPASSLGVALCGARRRR
Acheta_domesticus                   ---------------------------------RFTN*RRKKVFH----------GRRNRRF*ILLRGVI*D*GYFTRSKSTNMGQPALLPLRSKACEGFFRP*KSMASIGF------ESA
Gryllus_bimaculatus                 YISSL------------QNFLLILVEVAYV-------EKKCF*--------AVSHRRNSFI-----GIIKTPPPF*SLRLS-------WTARDKVRGTSYLPA*EAS-------------
Consensus

Teleogryllus_occipitalis            ------------------------------------------------------------------------------------------------------------------------
Tarbinskiellus_portentosus          GRGGARWRAGCEIRRGARFH--------------AAVLVG-------ASIDVSISVRCRRCG-------------------RWR----GGAGARVRSAVVRQEAQARVEL------RL
Acheta_domesticus                   NLGIVRY-ANCYTIVAISFHRSPAPHEAAGP*WGVCAP*VGMFDSC--KRIDTGIDPRIEKLKSI------LESNRKERIDSLRYRIT*E*GVFAISNGK*YRSFAHA*FKF------RI
Gryllus_bimaculatus                 ----------------------------------C*NNVN*INECL*LTILTRRTAVKKK*KSFVC*TS*LESKLINLL*T*EYTITK-----------SRYFRFKIFKFTFN*KLEI
Consensus

Teleogryllus_occipitalis            ---------------------------------------------------------HNAE-----------------------------------------------------------
Tarbinskiellus_portentosus          PSAESATRN----------VMNV---GHF*IMM----K*MLR--FSSSFPSETFLFVFNGE*YT------------------------------------------------CGAVG--
Acheta_domesticus                   TLVHLSMRRRS-------IVLNVR-GGKISLM---------R--FKGGMETRIEQFIFNKTQLDVN*LNLIKIIINLLI**K*I**NIDEILGPVAEIAKPAVAISSRRFAGLCPASGVG
Gryllus_bimaculatus                 *EIYYCI*RDNLFSLVEKMGINVAKFGEIIVI*YFSYKTVMRQISRGNYSRVIITFMFNSR----------------------------------------ASF*GRRCANCCILSYLP
Consensus                                                                                     *

Teleogryllus_occipitalis            ------------------------------------------------------------------------------------------------------------------------
Tarbinskiellus_portentosus          ---------------------------------------------------NDSPIG----------------------------------TEVNT*ERIP-------------------
Acheta_domesticus                   V------------------------------FVFVCVCVCVAFSPFIIIINSSSISPR-------LTTLVCKNG----LSRNSDPPKQCLTTRNACSRGP*----RP----------S
Gryllus_bimaculatus                 VTLSWLPPDYLNTDVHYL*RNEFEPCRPRWPGWLRACTDSALLFER-----SRARIPPWARWFVPCLPCMCCRGGFRVLLTLDKRSPC*CVLVPSGWSNIPSGRKG*PTVG*GKKFISYT
Consensus

Teleogryllus_occipitalis            ------------------------------------------------------------------------------------------------------------------------
Tarbinskiellus_portentosus          --H*HGNNT*IGVLF*RFH-------------FNKVIF----------------KK*KIL------KSL*AHYFHYIISSPGNNCR-KIVLV----------------------------
Acheta_domesticus                   REKK**SNTKF*KCNEVFHNIQYLKKTATVETLNKVNY*EIECAILFLSICLN**K*K*L*Y--KNKYLKQHFFQFSSTYSLSNCYLQVVFV---------------------------
Gryllus_bimaculatus                 *RIKFNSEARLVTFSA*FRVLSRGAKTP*AE------------VALFYAI----ISERVLEVFNTKQ*IEVH-------ESGGECH-HFIVVYQRLHLIGDGLPLPSTPHS*SENIRIFL
Consensus

Teleogryllus_occipitalis            -------------------------------------------------------------QSRRGGGGG--------------------------------------------------
Tarbinskiellus_portentosus          -----LSLLIHFCFSYRIYYLIV*HRKGRYEIANSTFCLRK-----------SGSFGALCGENREGGGGGGTRA----------NWGAGGA---------------------ETRRC--
Acheta_domesticus                   -----LLFVLRLCSTLHILE--------KYTSM*QVIVLK-----------------------------GVKARKVLVILDLGNWRAKSSRLFLALCDNENAF*FLN**YLTDDLGCCV
Gryllus_bimaculatus                 HSFHNLLYFARICATEN*LSLLILLHLILLTSSM*IILL*NLFLFNFKYITYSGS----------------------------NNPAEAS--------------------GQNEYC--
Consensus

Teleogryllus_occipitalis            ---------------------------GGGGG----------------------------------------------------------------------------------------
Tarbinskiellus_portentosus          -------------NLLLLLLSRGGGWAAGGGGRGGVG*GRGPGSCRGCNLPADESAIEEDEAVPLHHFAVMDGYISGASIY---LTTRP-------------------------------
Acheta_domesticus                   SVTFVNYLYISFLHFLLILVRRIYD------GKDQLKSTHAAGTVL--------SILEKRD----HHFAFRLSYSTRDKILPHELLVRPGS*LLLLQ*CEIN*QMCYD*KIEFGTSVSTP
Gryllus_bimaculatus                 ------------KNLLSL-------------GHSQCAVI**TG---------------------HY------------------THP--RLLLPLICSMLFQ-----------SIKDP
Consensus                                                          *

Teleogryllus_occipitalis            ------------------------------------------------------------------------------------------------------------------------
Tarbinskiellus_portentosus          --TTRACLPLLLRASF-----------RKLSTPRRFSLTSRSR-----GVYRLLSRFYKGIDSFTFSHI*AETILKFNLFCDWSKILYKENNEIYCVRF*MKIQ*IMYSVRIYATLPCGF
Acheta_domesticus                   MRTTRHCFTRLSRKEFWK*LRGSN-RLKSLEVVVEISISSQHRYVCA*YVVDLLMPSTPHS*SKTFMFYSTRTLLFAQMGCLCENLRYSQ------------LAAIAHS-----------
Gryllus_bimaculatus                 F-------------AIYFYIRKQNPHGKRLKTKLPRELFNAH*H------------------KSMSFFSL*TTVYNH--CM---VLYLKHN--FCE*YTKYVNIYKYY-----------
Consensus

Teleogryllus_occipitalis            ------------------------------------------------------------------------------------------------------------------------
Tarbinskiellus_portentosus          HIFLPVIKLMNLTNKSYFQPGIYE-*ICG-------PLQYDFSPIKRKNMN**MRKHILEGINTERPVRNRDE*SAANICS--------------KYLRGNLMSNCILQYIL*LCINLAI
Acheta_domesticus                   -LIFDIIKTIEL---SYDEIDFIEVQICNLLEINSKPAQFRMASMKRADL-----------INEKIDL*MRYLQRF*RVCNIPLTN---PKEK*EQYRRPNVTNDS----------ELFI
Gryllus_bimaculatus                 ------I*TISMY------------LLCV-----------*TGSTAKARIN----------LKNQRCICNGYKKYSTSTHTIPLTAVVWTNENSKEGRM**MTVNYLFP*------NL--
Consensus

Teleogryllus_occipitalis            ------------------------------------------------------------------------------------------------------------------------
Tarbinskiellus_portentosus          HLLK*DGTINIRTVCK*QLKENYFSPGGP--------------PSDA*PCV*RGSAKTSPPAHGNAGSPTSAYGSGLTFSMGWCCARASGDGA---GRGEAG---RRETSAAMNLSPRVA
Acheta_domesticus                   SLQKTEFHL-----------ENILV**GPFKFIFQRLGLFLSFVGCKYPRARRGSEKTAPAAYGNAGSARSAFGSGLAGALNR--TRASADGALARARTRAGAGAERKTRAAMNLSPRVA
Gryllus_bimaculatus                 -----EFHF-----------QTMLV*EGSSNLFSRVLVSFLVWWDVNFLVCGEGLL----AAYGNAGRAQSAYGGGLALWIGR--ERGWGAVEAGRARGRRR-GAGRKTRAAMNLSPRVA
Consensus

Teleogryllus_occipitalis            -------------------GGGGGGG--------------------------GGGGGGGGGGG---------------------------------------------------------
Tarbinskiellus_portentosus          IDREQKAACRADSTMRSSRGGGGGGG--------------------------GGGGGEGGDGGEAVATMVSLAMAA*VGSRCPLRIRSHCIKTSSPIRALLHNFISVSPLNVSALSI*KY
Acheta_domesticus                   IDREQKAACRADSTMRSSAGGGDGGGDSGERSRGGN-------EGQGGGDDDGGGGGGGGEGG---------------------------------------------------------
Gryllus_bimaculatus                 IDREQKAACRADSTMRSRAEQSRAEQAARRAARAAR*R*AAVLEAVDGKAERWPGAGRGCVGR----MVVTWCVAE--GVRC-------------------------------------
Consensus                                                                             * * *  *

Teleogryllus_occipitalis            ------------------------------------------------------------------------------------------------------------------------
Tarbinskiellus_portentosus          VVPQSQPNDSETLRIDSLNVFVTSSVHFPVRRPLDNFTPMSR*FCHSITIFSCFFFCFIFIISLLSHFSRSFHSFSFSLLA*FVLPFLLSLCLLRISQSNFNVSLLLTYSNVLNFN*GIT
Acheta_domesticus                   ------------------------------------------------------------------------------------------------------------------------
Gryllus_bimaculatus                 ------------------------------------------------------------------------------------------------------------------------
Consensus

Teleogryllus_occipitalis            -------------------GG---GGG-------------------------------GGGGGGGGGGGG--------GGGGGGGGGGGGGGGG--------GGGGGGGGG--------
Tarbinskiellus_portentosus          ADKVLTGVERGG*PQGGAEGG---GGGRERRGRGRGRG-------------------RGGGGGGGGGGGG--------GGG----GGGGGGGGG--------GGGGGGGGGHS------
Acheta_domesticus                   -------------------GG---GGG--------GNG-------------------GGGGGGGGGGGGGSGDGDGGSGGGGNGGGGGNGGGGG--------GGDGGGGGGGGVGVAAV
Gryllus_bimaculatus                 --------GRGEPGERGYRGGPWRGGGRI------GRGWSDSCTDMTEGVACKTAMRCGGGGGGGGGGGGGLTR-----GGGSDGC*GGRSGSGSDDRRYKCDNAESRGGDRDK------
Consensus                                              **   ***                               ************        ***      **  * *               **

Teleogryllus_occipitalis            ------------------------------------------------------------------------------------------------------------------------
Tarbinskiellus_portentosus          ----------------------------------EPDPET-------*SPVLPLS--NICLSCVFLPEYITQSEYYLLRCG---ERNIFFA-----CHAQFIVYVVIFVHTFLIPSWSWN
Acheta_domesticus                   AAGAGRRLWT*RWRRWWG*QGGATRNVTKEKAVVEI*TEVLTK*WFQRSPMWALGIINPP*KMF*LPIFFFNMTGEVVKKGGEDDGNVDVSANDKICKGRGEVGKSIGVRTI---EWEIK
Gryllus_bimaculatus                 -----------SWRK-------------------RLVPEKSHDGFRYSVPPLKIR-----LNCH*YPS---------LR*AKE-------------------------------LEMGMS
Consensus

Teleogryllus_occipitalis            ------------------------------------------------------------------------------------------------------------------------
Tarbinskiellus_portentosus          *DEGVQRFV-------IS*VIKATKKEREV*MKG*INTK*LASSPSGA**QSTRCTRHK*VTGLQSICK*GPGNESLGRGAVSAHGLLMRALSVFIERTFTRAHAA----TDVRDQS---
Acheta_domesticus                   KNEKMIA*VQLGLNTEIMMVVEGV----EVDVELETVTKT*ESPP------WPRSPSH--------*CRSDFGPRSHGFKSFLGHGCMCSSFHPYHHHTISSLYH------NVKSNSSR*
Gryllus_bimaculatus                 TRAEMTRYAK------VMQIWERM--------------------------WNR*LRL--------------GPQNRSLRRM*K**CMCS*G*MRK**AMREVMSEEEVEVQVRSKA---
Consensus

Teleogryllus_occipitalis            ----------------GSIGGGGSGYGGG*------------------------------------------------------------------------------------------
Tarbinskiellus_portentosus          -------------QSAGPLPGGAEPLSSGPIGAPR----AVRGQ-RPSPLPSPRLPPP-----------PPPTPRC--PR--LSSP----------LPPSCP-----------------
Acheta_domesticus                   *LSR*CECLNKKEKTNGVTSNGNDGEGGGTDGGDQDEGILLSGRTRLCPL----LPEEKASDFTLSLRLVPPLY*RCALPRGLLCSPSFPIGGSTRVLPPDVPLYLLYWRGRRLRRGGGC
Gryllus_bimaculatus                 ---------ETKTESNG--------------GGRK----------------------------------------------------------------------LQ*WRRRK-------
Consensus                                           *

Teleogryllus_occipitalis            ------------------------------------------------------------------------------------------------------------------------
Tarbinskiellus_portentosus          -------------APPATATRQRQ---------------------------------PLW*WSG----PTTTFCPQ-VYGARICKC-------KWRKIRFDFS---SDPCIWSLIENSM
Acheta_domesticus                   GVDSGGGGGSCKWGPKTNMTTEKAMTEIETVVLTKWWFQRSTMRT*GVMSPL*KMF*LPIFSFNSL--VPLVTHM*QRFWSTRISRFL-----H*FAKFIFYFSFSYSFPYFPFCVPNKS
Gryllus_bimaculatus                 ------------------------------*WI*WWRQR-------------------WRYSAVREEPGCVHSFQ--WKEQLSLC*VSGSSHRFISVALSFENS-ARQAFQFVVPVRS
Consensus

Teleogryllus_occipitalis            ------------------------------------------------------------------------------------------------------------------------
Tarbinskiellus_portentosus          ------------------YE*SKFFNIYFYEK-DVRFDRWFYIRFVKESSC*RPNEVVDD*FGFLSGEFRFFKFI*IITRTHMHTG--II*FS*LYTTRMY*RLIS------------
Acheta_domesticus                   ------------------VGISKIFSSYSFSRLPLRYSETFIL-----------------FFQFLTNIFYFF*F------RQLHTDYFLVYFILCFQP*NYNKFCSD-----LSYPESS
Gryllus_bimaculatus                 KCQGAGESCLQRFRKSISLLSSRILFEVFLSPRPTSQFV*NIFLVFFLASSFAIGDTFVL-LFSFISKILFVFLY------FVCHIPIRFLYILSCYVSSPVIITSSAAI*VILSHRLSR
Consensus

Teleogryllus_occipitalis            ------------------------------------------------------------------------------------------------------------------------
Tarbinskiellus_portentosus          -----KSK-------LT*K*VLCKRKETHKL-------------TL*KSFKSTN-----------------FFFKLIFIGYSC------------------------------------
Acheta_domesticus                   SSPLQKEKNE*DDELIQNNFLLCLHTH-HKVHDVYVANKILVSSPLTNRDSSMNHSEKWSPHTIVMITVLCFFIEHTFMQEICATGPQCVKPIEVGNDGPCRGARYGLEWRKSGQKII--
Gryllus_bimaculatus                 SR--KKEKNE*DDELIQNNFLLCLHTH-HKVHDVYVANKIPDSSPLTNRDSSMNHSEKWSPHTIVMITVLSAFIEHTFMQANCATGPQYVKPIALGND----MADYGLDWGKRGQKVICM
Consensus

Teleogryllus_occipitalis            ------------------------------------------------------------------------------
Tarbinskiellus_portentosus          ------------------------------------------------------------------------------
Acheta_domesticus                   ------------------------------------NSVVAAEI----------------------------------
Gryllus_bimaculatus                 IYVSLPDLSTVDQLHLLLVPSLKEFFQ*TYGSIRCDSRAVCA*IFFSAMQSILQYFPDIDSFCPASTTRFNSPPYFLM
Consensus
```

**Tocc.3G0002060.1**

```
Teleogryllus_occipitalis    AAAGGACCAAAAGCATT-------------------------------------TTTGTTTTATTAAATGAATTTC----------------------------------------ATGA
Acheta_domesticus           ------------------------------------------------------------------------------------------------------------------------
Gryllus_longicercus         AGCGCATCAAGACCCTCATTGTGACTCAAAGTTTTCGGGAATAATGTTTTTAAATTTATTTATTTCAGTTAATTTTTCGGTAAGCCTGATTTAACTGGGATTTATGGGAAACTGAAACAA
Consensus

Teleogryllus_occipitalis    ATTAGTATTTGCTTTTAGGTAAT-------------------------------------------------------------------------------------------------
Acheta_domesticus           ------------------------------------------------------------------------------------------------------------------------
Gryllus_longicercus         GCGAGTCTCTAATTATAGAGAATCAATGCTACTCTACTCTAGAGACTATACACGTCACAGATTCACACACCACACATAGGGGAACATGCACTCTCCCACAACTAGACACAAGCCGTCAGT
Consensus

Teleogryllus_occipitalis    ------------------------------------------------------------------------------------------------------------------------
Acheta_domesticus           ------------------------------------------------------------------------------------------------------------------------
Gryllus_longicercus         CACCCGGAGGAACACACACGCAAGACCGCGTACACATTCAGACACACACACACACACACACACACACACACACACTAACAGCTACATTATTACAACAACACATAGACGACACACTACC
Consensus

Teleogryllus_occipitalis    ------------------------------------------------------------------------------------------------------------------------
Acheta_domesticus           ------------------------------------------------------------------------------------------------------------------------
Gryllus_longicercus         ACTCACACGGGCCGCACAATATTCACGGATAGACACACACGAGGGCTCGCGCACGCATACACAAACGCACGCGCGCACACACACACACACACACACACACAAACACTCCACACACAAG
Consensus

Teleogryllus_occipitalis    ------------------------------------------------------------------------------------------------------------------------
Acheta_domesticus           ------------------------------------------------------------------------------------------------------------------------
Gryllus_longicercus         ACCTGACGTAGAAGCTGACGTATAGCGAAACACCATATCGCGTTGCTTGGCGTGGCCTCGAATTACGTAACTTCTCCCCACATCCTAAAGAGCCCAGGTGCAAGCCCAAAACATTTGGTG
Consensus

Teleogryllus_occipitalis    ----------------------------------------ATTTTTAACGATCTTTATGT-----TCTTATTTACTTATATTCAGAAAATGGCC--------------------------
Acheta_domesticus           ------------------------------------------------------------------------------------------------------------------------
Gryllus_longicercus         GCCATCGTATTGTGACCGTCTTCAGGGATGAAATGTCGCAATATTTGACACTCCCTTTGTCACACCGTTTTCCATGCATGTTAAATAAATAACACGTTGGGCGTGTTATATACAGCCATT
Consensus

Teleogryllus_occipitalis    -----------------------TAAACAGCGTCCGAAAGTGTGTATAT-----------------------------------------------------------------------
Acheta_domesticus           ------------------AATGATAAATGCCGTTAAAGGCTTTGCG--------------------------------------------------------------------------
Gryllus_longicercus         CATTTCGCGGATAGAGTCATCGGTGACCACCGTTAGAATTTTGGCATTTCAGCCCAAAAGAACACAACGGATGAACTTGAATTTCATAAAAATCGATGTTTTATAAAGGGGGCTTTTAAT
Consensus                                          * *    ***   *   *  *

Teleogryllus_occipitalis    -------------------------------------------------------------------------------------------------GTAACAGGGTCTT----------
Acheta_domesticus           -----------------GCATGTTACGA---------------------------------------------------------------------GTGACAGGGC-------------
Gryllus_longicercus         ATTTGCTTATTTTATTTGTATAGTACAGTATACACATTCAACTGCTTACCGAATCAAGAATCGGAAAAAGTCAATGCACTCTTTGATACAACACTTCGTGAAGGGGCATTTATATTCACA
Consensus                                                                                                                    ** *  ***

Teleogryllus_occipitalis    -------------------------------------------------------------CCATGTGCCGTGATTTTCCAGACTTCAATTTGATTCAAGGCCCAATTATATGGATTTTT
Acheta_domesticus           ------------------------------------------------------------------------------------------------------------------------
Gryllus_longicercus         GAGAGATTTGATTTCGTAATTCCACAGAAAGAAACACATACTCTCGGAACATTTAAAACAGTCTTGCGCAACCAATTTTTTTATTGAAATGTAACTTAAA----AATAATACCAATATTT
Consensus

Teleogryllus_occipitalis    GATTGA----------------------------ATATGCGCTTATAAAATC------------------------ATAAGTTCAATTTATTGTTA------------------------
Acheta_domesticus           ----------------------------------------------------------------------------ATAAGCGCCGCCTGCCGCGT------------------------
Gryllus_longicercus         AACAGATTCCGATCTATGATTAGTAGCTTTCTTCGTACACTCAAATAAAATAATAAACAGAATTCATTCTAAATAAATACATTTCATTTGCCGCATTAGTTTATTTTAATATTGCAAATA
Consensus                                                                               ***         *   *

Teleogryllus_occipitalis    --------TCTTTAATGATATATTGAAATTATT-------------------GAGCTGAGCGGAAGAGCAAT---------------------------GAGAGCAGGGCCGTCAGGG--
Acheta_domesticus           ----------------------------------------------------AGGACAAGTCAGTAAACTA-------------------------------------------------
Gryllus_longicercus         TCAAACGATTGTTAATATTAAATTAACCACATTAGAGAAATTTTACGAAAACAAAATAATCCAATATACTATATTCATTTCTTTTGAACCGGATGATCGGAAATCTAGTACGTCAGGGTT
Consensus                                                                     *         * *

Teleogryllus_occipitalis    -----------GTGGTTGTGTGG-------------------------------------CGAAAGCGGTCAAAGGAGCGACACCGTTAGAGAGAAGATTTAAATTAGTAGATTGGATAA
Acheta_domesticus           ------------------------------------------------------------------------------------------------------------------------
Gryllus_longicercus         AGTGTAACGCAACGGTTGTATAATATATTTCAAATTTCAGAATGTTTCATGTTAATACTCTGAGTACAGTGATGAGTATAATAATGCTCAATAGAATATTAAAATGAGAATATGAAACGA
Consensus

Teleogryllus_occipitalis    GCAGTTTGCTCA-------------------GAGGAAAACATTACAA-------------------------------------------------------TTATAATTCGCAT-----
Acheta_domesticus           -------------------------------CGCGCAAACCTGACAG--------------------------------------------------------CGTGATGTTCAC-----
Gryllus_longicercus         GTTAATTGTGTGTACAACGCTCATGTTAAACCGCGCTAATGTGACAAACGCATCTAGCCATCTTTTATAATGCATGGACAATACCGCTTGATTTTAATGGGTTTATGACATTCACATATG
Consensus                                                     *  **  * ***                                                           * *    **

Teleogryllus_occipitalis    ------------------------------------------------------------------------------------------------------------------------
Acheta_domesticus           ------------------------------------------------------------------------------------------------------------------------
Gryllus_longicercus         CATTGATTTATCTTTGATACGCTGTTTAACACCATTAAAGTTTAACTCATTTTCCTGACGAATAATTTTATACGACATGGTATCATTCGAAAAAAGAATTAATATTCTTTATGAAATATT
Consensus

Teleogryllus_occipitalis    -------------------------------------------------------TAAAGAACGTAA----------------------------------------------TTATAAT
Acheta_domesticus           ------------------------------------------------------------------------------------------------------------------------
Gryllus_longicercus         CATACAATTTTTCACGCAAGTCATAATTTGAAGACGACTTTATGAATTGTTAATGTAAAAATTATAAACCTCATTGTTCATTCGTTCTCGAGCCATGTCTTCACAAATTTTCCGTGTAAT
Consensus

Teleogryllus_occipitalis    TTATCTGCAAGTCCACCGGCAGCGGCGG-----------------------------------------------------------------GACAGCGCTGAGGAGA-----------
Acheta_domesticus           ------------------------------------------------------------------------------------------------------------------------
Gryllus_longicercus         TCTTTTGCAATTACATTAATTATACTGGTGTAAGATTTGTAATTATTCTTTATAACTGAAATCCAATTTCTTCGTTGTAGGGACATTAATTTTAAAAACGTTTGAGAGAGTAAATAAAAA
Consensus

Teleogryllus_occipitalis    ------------------------------------------------------------------------------------------------------------------GGCGTA
Acheta_domesticus           ------------------------------------------------------------------------------------------------------------------------
Gryllus_longicercus         TCTTGAAAACAAAGTATTTGTGGAATTCAACGTACCTAGAATATTAATCACTTTGGAAATTGATACATTTGCAGCACACGCACACCAAAACTGTTTCTAAAGAATCTTAAACCTGATGTG
Consensus

Teleogryllus_occipitalis    GTCTACGGGGGTGGATGTGACA--------------------------------------------------------------------------------------------------
Acheta_domesticus           ------------------------------------------------------------------------------------------------------------------------
Gryllus_longicercus         TTCCATCGAAACATTTGTAGCACGCACGTTCGACAAATTTCATTCGAGAGCCCTTGCCAAACCACAGAGCACTTCTCTCCCCTTCTCATTCATTCTTTATCATCTTGCGGGACATTTATG
Consensus

Teleogryllus_occipitalis    ------------------------------------------------------------------------------------------------------------------------
Acheta_domesticus           ------------------------------------------------------------------------------------------------------------------------
Gryllus_longicercus         TCGAAATAATTTTCCAGAACGTGTGGTAATTAATAACTGTCCAAATAGGGTGTCGCCGGCACCCCCCGGTCTCCGATTGAACCGGTGCCAAACCCCACCGTCTGCTGAGTCTCCCCGTTT
Consensus

Teleogryllus_occipitalis    -----------------------------GCGTGGGGGTGGCGAGGAAGGC-CAGGGCAACGAGGTCGGTGGATGCGGCAG--------------------------------------G
Acheta_domesticus           ------------------------------------------------------------------------------------------------------------------------
Gryllus_longicercus         TCCTTTCCCTTCTCAGAGTTCGGGCCGGGGGGCGGGGGTGGGGGAGAGGGTGCTGGAGGGGGGGAGGCGGGGATTGCGGGAAACCACGCCTAATCCCTGCGGGTCCCGGAATCCGGAGTCG
Consensus

Teleogryllus_occipitalis    GAAGAAGGTATCGGCGGGGCGG----------------------------------------------CCGCGGCCGTGGCGGAGC------GGAGGGGAGGGCGGCACATACGACGGGG
Acheta_domesticus           ----CAAGTCTCC-----------------------------------------------------------------------------------------------------------
Gryllus_longicercus         AACACAAGTCTTCCCGGGGTGGGCTTTAAGCCGCTGCACTTCGTCACTGTCGCTTGACAAACACGCAATCAGAGCGGCGGCGAAGCGGGCGGGGAGAGTGGGGCTGGGGATGGGGGGGG
Consensus                        * ** *

Teleogryllus_occipitalis    GTGGCGAGG--------GTGGCAGGGTAGTGTGTGG-----TATGGTGGTGG--------------------------------------------------------------------
Acheta_domesticus           -----------------------------------------CATGATGATGA--------------------------------------------------------------------
Gryllus_longicercus         GCGCCGAGGCGAAGGACCCAACAGGACGGCGCACAAGAAGTCATAGTGATGGGACGAAGGAAAGTGGAGAATCGAAGCGACGCATGTAATGCCTGTTAGATGTTACCAGGGCAAGAAAGA
Consensus                                                             **  ** **

Teleogryllus_occipitalis    ------------------------------------------------------------------------------------------------------------------------
Acheta_domesticus           ------------------------------------------------------------------------------------------------------------------------
Gryllus_longicercus         TAACAATAAAATAATATGAAAGTAGTAAAGAAAAGGCGGGAGGCAAAAGAGTTAGCAATTATATTAGTCACACATGTTATTGCAGGCAAAATGAAAGAAGAGACCAAAGAGGTCTAGGAG
Consensus

Teleogryllus_occipitalis    ------------------------------------------------------------------------------------------------------------------------
Acheta_domesticus           ------------------------------------------------------------------------------------------------------------------------
Gryllus_longicercus         TCATAGTTAAGACGGCAGACGGTAACTACAGCAGAGAATGGGTCAGAAATTCTAAAGCCAGAACCGAAGAAACGTTTGTGAGAAAAATAATGTTGGCAAGCGCAGAAAAAGCAAATGAAA
Consensus

Teleogryllus_occipitalis    ------------------------------------------------------------------------------------------------------------------------
Acheta_domesticus           ------------------------------------------------------------------------------------------------------------------------
Gryllus_longicercus         TTGTGAAAGACAATACACAATCCGTGGAGAAGATGGGCGTTGTATGTATGTATGTATATATATATATATATATATATATATATATATATATAGAGAGAGAGAGAGAGAGAGAGAGAGA
Consensus

Teleogryllus_occipitalis    ---------------------GGATGGCGAAGGTAGCATTG-------------------------------------------------------------------------------
Acheta_domesticus           --------------------------GCGGATCTACTGCCGCGATCTCATGGAAGCGTTCT-----------------------------------------------------------
Gryllus_longicercus         GGAATAAACCGACGCAGTGGCAGAGTGCGAGTGAACCATTGTCATATAAAGACAGTGTTCTAATAAGATGAGATAATGCCTTCGCAGAAGCGGAAACGTCAAGAGCCATATGAAATATCG
Consensus                                             ***     *     *

Teleogryllus_occipitalis    ---------------------ACGGTGGGGAT---TGCAAAGGTGACGGGGAT---------------------GGCGGGGCGGCACAGGCGGCGGTTG-----------GTGACAAGGA
Acheta_domesticus           ---------------------ATGTAGCGAAT----------CTGACGGG--------------------------------------CGCGATGGTTC---------------------
Gryllus_longicercus         CACCTTGACACAAAGGGACAGATGCAGTGAATAATAGTAATGCTGGCGAAGCTATGCCAGCCATAATCAGATCAGGATGAGGTGCTCCAGCGATGGTGCACATGTCTCCAATATTCAGCA
Consensus                                        * *  * * **           ** **                                         ***  ***

Teleogryllus_occipitalis    TGGCGTCGATTGTTGGACGGCGG--------CGGGGGCAGTC---------------------------------GGGGTGGCGGGGGAAAGGTGCCGGGTT--TGATTGATTGATTGAT
Acheta_domesticus           --CCAACAATATATAGACAGCTG--------------CCACTCGCTCCCGACCTGGTGGCAGTGGCCCACTGCGAGGCGTGGTGGA------------------AGGTTGATTGATTGAT
Gryllus_longicercus         GACCAACCCTTTGCAGACATCTGACAGCCACCGGCGTCAACCCGCTTGCATCCCTCAAGAACCGTCCCCTTTCGGGGGGCAGCCGGTACAGACTCTTAAGATCTTGATTGATTGATTGAT
Consensus                      *  *  *     ***  * *              *                                     ** *  *  *                    * *************

Teleogryllus_occipitalis    TTTTTCGTGTTTCCTGGCGCATCAACTGCTTTGGTCATTAGTCCCGTTAGTGTAGTGCATGTTCTTTTCTCGGGAGATGTAGGTGATC-TTCCACCACTGGATGGCGAGAT------CCG
Acheta_domesticus           TGATTCGTGTTTCCGGGCGCGTCAACTACTTAGGTCATTAGTCCCGTAAGTGTAGTGCGTGT--TTTTCTCGGGAGGTGGCGATGATCTTTTCCCCGCTGGATGGCGAGATCTGGCGCCG
Gryllus_longicercus         TGTTTAGTGATTCCGTGCACGTCAACTACCAGGGTCA--AGTCCCGTAAGTGTAATGCATGT-GTTTGCTCGGGAGGTGGCGGTGATCTTTCCCCCACTGTATGGCGAGATCTGGCGCCG
Consensus                   *  ** *** ****  ** * ****** *   *****  ******** ****** *** ***  *** ******** **  * ***** ** * ** *** **********      ***

Teleogryllus_occipitalis    ACCTACCGTCGGGTAGTGAGGTTTCCCCC-TAACCAGGACTCTCCCACCAGCGGAGCACCGCCCAAGCCGGGACAAGCTCTAAGCTCCCGACCGATCTCCCGGAATATGTGGTAGTGTAG
Acheta_domesticus           ACCTACCGTCGACTAGCGGGGTTTTCCCCTCAACCAGGTATCTCCCACCAGCGGGGCACCGCCCCAGCCGGGACAAGCTGTAAGCTCCCGACCGGTCTCCCGGCATGTGTGGTAGTGTAG
Gryllus_longicercus         ACCTACCGTCGGTTAGAGGGATTTTCCC--CAATCAGGTATCTCCCACCAGCGGGGCACCGCCCCAGCCGGGACAAGCTCTATGCTCCCGATCAATCTCCCGGCATGTGAGGTTGTGTAG
Consensus                   ***********  *** * * *** ***   ** ****  ************** ********* ************** ** ******** *  ******** ** ** *** ******

Teleogryllus_occipitalis    TGCATGTTAGTGTAAGTGTGTGTGTAAGGGTAGACGATAGGACAAGGAAGAGGAAGGACAGATATCGCGTGGTCCGTGGCCCGTTTGTTAGGGTACTGCTCCGGCATTCGCCTGGTGTTT
Acheta_domesticus           TGCATGTTAGTGTAAGTGTGTGTGTAAGAGGAGAGGATAGGACAAGGAAGAGGAAGGAAAGATCGCGCGCGGTCCGTGGCCCGTTTGTTAGGGTACTGCTCCGGCATTCGCCTGGTGTTG
Gryllus_longicercus         AGCATGTTAGTGTAAGTGTGTGTGTAAGGGGTGAGGATAGGACAAGGAAGAGGAAGGTCAGACCGTGCGCGGTCTGTGGCCC-TTTGGTACGGTACTGCTCCGGCATTTGCATGGTGCTG
Consensus                    *************************** *  ** **********************  ***    *** **** ******* **** ** **************** ** ***** *

Teleogryllus_occipitalis    ATCGGAGAAACCACGG-AAACTACCGATCAGTGCAGCTGACCTCTAGAATTCTCCTCCAGACAGCTACTGAATTCGACCTAGCAGAAAAGCTAATATCGAAAATCCTGCCGCCGTCAGGA
Acheta_domesticus           ATCGGAGAAACCACGGAAAACTACCGATCAGAGCAGCCGGCCGCTAGAATTCTCCTCCAGACAGCGACTGAATTCGACCTAGCTGACAAGCTAGTGTCGAAAATCCTGCCGCCGTCGGGA
Gryllus_longicercus         ATCAGAGAAACCACGGAAAACTAGCGATCAGAGCAGTCGGCCGGTAGAATTCATTTCCAGAGAGCGACTGAATTCGACCCAGCTGACAAGCTAGTGTCGAAAATCCTGCCACAGACGGGA
Consensus                   *** ************ ****** ******* ****  * **  ********   ****** *** ************* *** ** ****** * ************** * * * ***

Teleogryllus_occipitalis    CTCGAACCTAGTACCTCCCGGTCAGAGGCGCTGC------------------------------------CGCTCCCGGT-------------GGAAAGGTGCCGGGGATGGCGCAGGCA
Acheta_domesticus           CTCGAACCTAGGACCTCCCGGTCAGAGGGACTGCAGTGACATCAGGTTGATGCACTGAGCCATCTCTCCCGGTGCCTGGT----------------------------------------
Gryllus_longicercus         CTCGAACCTGGGACCTCCCGGTCACAGCGACTGT--------------------CTGAGTCATCTCTCCAGGCTCCCGACTGGTCTTCCCACACGAATGGT----AGTATAGTGTAAATG
Consensus                   ********* * ************ **   ***                                      *  ** *

Teleogryllus_occipitalis    ACGGTTGGTGGCGGGGGTGATAAGGGTGGCGTGGATGGCGGGGGTGACAA---------------GGGTGGCGTGGATGGTGGGGGCCG---------------------GAATGGAGGA
Acheta_domesticus           -----------------GGAAGGGGATGGTGTCGATGGAGAGGGGG-------------GTCCTTGAGTGGC-TATGAGTCTTAGGCCC------------------------GAGACAA
Gryllus_longicercus         TGTGTT---------CGGGAAGAGTATAG-GACACAGAAGAGGGGAAAAATCGCACGCTGTCCTTGACTCGTGTTTAAGGCGTTGCTCCGGCATTCGCCTGATATTGATCGGAGAAACTA
Consensus                                     **   *  * * *     *  * ***                     *  * *  *    *     *  *                            *  *

Teleogryllus_occipitalis    GTGACAGGGGCGGCGCGGAGCGGAG-------GCTTCGAGGTGGGGGGA---------------------------------------GCGGGGTGGCGGGGGGCACAGGGGGCGAGGCG
Acheta_domesticus           TGGGCGGGGGCA--------------------CGCGCGAGGCGCGTGAA-----------------------------------------------------------GGCGCGGCGGCG
Gryllus_longicercus         TGGACCAACGCAGCCTGCCGCTAGAATTCTCCTCCAGAAAGCGGATGAATCCGACCCACCCATCCAGCTAGTATCCACCACCGTCACCGTCTGAAATTGAACCGCGCCCGAGGAGAGGTG
Consensus                     * *    **                           * * *   * *                                                            * *  * ** *

Teleogryllus_occipitalis    GGGCGGGGCTGGGCGGGCGGGGCTGGGCGGGGCTGGGCGGGGC--TGGGCGGGGCTGGGCGGGGCTGGGCGGGGCTGGGCGGGGCTGGGCGGGCGGCGGGGCTGGGCGG-------GGCT
Acheta_domesticus           CTTTGAAGGAGCGTGCGACCGCGCTA-------CAACTCACAC-----AGTGGGCAGGGCGCGCGACGGCACCACTGCTTAGAAGAAGGTGCAAAAATAAGCCGAACAT--------GCC
Gryllus_longicercus         GCTCAGGGCATCATGCGATGTCACTAGTGTCTCTAACCGAGACGTCCTAGTTTCAAGTCCGGAGGACGGCAGAATT-TTCGACAATAGGTACATGGCTAGGTCGAATTCAGTCTCTAGCC
Consensus                          *      * *                         *             *  **      ***     *           **           *  *             **

Teleogryllus_occipitalis    GGGCGGGGCTGGGCGGGGCTGGGCGGGCT--------------------------------------GGGCGGGGCTGGGCGGGGCTGGGCGGGGCTGGGCGGGGCTCGGGCTG------
Acheta_domesticus           GAAAGAAGCTCTGAAGTGCC---------------------------------------CACCCCGCGAAAGGGGTGGGTCGGCGGCGCTGGGTACAGGCAAAGTACC------------
Gryllus_longicercus         AGAAGGGAATCCTAGCGGCCAGCTGCACTGATCAGTAGTTTTTGGTGGTTTTTCCGATAAACCCCAGGTGAATGACGAAGCAGTACCCTAAGAAAGAGGCTAGGGACTGAGCGGTCTTTC
Consensus                       *    *       **                                                *     *      * *        *     **    *

Teleogryllus_occipitalis    ------------------------------------------GGCGGGGCTGGGCGGGGCTGGGCGGGGCTGGGCGGGGCTGGGCGGGGCTGGGCGAGGCGGGGCGGGGCTGGGCG----
Acheta_domesticus           ----------------------------------------------------CGCGGCGGCAGAAGAT------------------CAACTGCCCGCTCGGGGGAGGGG-----------
Gryllus_longicercus         CTTCCTCTTTCTTGTCCCATCCTCTCCCCTTACTTACACTAACACTTAGTTGCACTACATTGGGAGATACTGATTAGGAAAAACTCCACTAGCTCACCATCGAGAGGGGGAAGGGAACCG
Consensus                                                                         *       *  *                         *  *      * * ****

Teleogryllus_occipitalis    -----------------------------AGGATGGGCGGGGAAAGCGGGCGGCGAGGGTGGTGAGGATGATGGGGTGGCGGGGGCGGCGTAAAGAGTGAAAGGGTAAAGATAAGGTG--
Acheta_domesticus           -----------------------------TACGCTTACGGG-----CAGGCGGCCGCGGC------------------------GCGCGACAAAA-------------------------
Gryllus_longicercus         ACACCTCCCACGATAAAAAAAGCTTGCGCTACACTTACGGGGCTTACGGGTTACCGCGCCT-----AATCATCAATCAATGGAATCGCGGTGAAAGCTGCAAAGATGCAGTAGCAATGCT
Consensus                                                        ****     * **   *   *                           **     **

Teleogryllus_occipitalis    -------------GAAGGTGGATGCTGTGAAGTGGAGGGAAAGTACAGGCTGCAGAGGTAGAGAGAAGCG-----------GCGTGT-----TTGCGAGGCGACAGCGAACGCGCCGGG-
Acheta_domesticus           ----------------------TGCGGCCGACCCTAC-------------------------------------------------------GACCGCAGCGGCGGCGGGCGCCTTTGA-
Gryllus_longicercus         GTCCCTGACTTCCGACAGCGGTCGTGGTCGACCTAGCTGACAAGCTAGTGTCGAAAATCGGTAGGGCGCGCTTGTTTGTGTGCGTGTGTGTGTGTCGCAGTGGTGGTGTGTGCTGCGGGC
Consensus                                          *  *   *                                                                **  * *   * *   **    *

Teleogryllus_occipitalis    -------------AATGTCATCAAGAGGTCTTCGCGTGGTCCCGCAGCTGGAGAAGGGGGATTTTTGGCATCGGTAGTGC----------------------------GTCGACGTC---
Acheta_domesticus           -----TGATACAAAAGGTCGTGCAATACCTTT----CGTCTTGCCGGCTACCGCGAAAGAGTGTCTGAGAGCCCCAACAC----------------------------------------
Gryllus_longicercus         AGAGCCGGCGCGAGAAGGCAGGCGAGGCCCGTTTGGTGTCTTAGCCGCCTTCGCCCGAGAACTTCTCTTGTTCTCGGCGCCCGGCCCGCGACCCTTCCGGGCTTCGATTTCGACGCCTGG
Consensus                                 * * *            *     *      * **    *     *    * *             *

Teleogryllus_occipitalis    -------------------------AGTGCAAGGAACGT-------------------------------------------------------AGTGGGAGCGGCGGCAACGAAATGTC
Acheta_domesticus           ---------------------------------------------------------AGGTCGCTTCGACTATCT-------------------CGAAAAAAC-----------------
Gryllus_longicercus         CGGCTCCGGCTTTCATCCACTCGGGACTCCGAGGAGCGTCCAATGGTCTGGCCCACGAGGCCCCACCAACTTTCTTTTCTCTCCAATGCGAAACCGAAAGAACAACGGTAAAGGCGTGTC
Consensus                                                                                                                  *    * *

Teleogryllus_occipitalis    AATGACGG-----------CGCGGTGTCGGTA-----------------------------------------------------------------------------GTGAAGAAGGA
Acheta_domesticus           -------------------TTCATTTTTTGTACATAAATTTCCTAA-------------------------------GTTTTGACTGTTCAAAAACATCTTAAACTTTTTTA--------
Gryllus_longicercus         ATTTACGAAAACCACAGCATGCGATTTCGGAATGTTGAACTTTCAATCTGGTATTAATTAATTGGCATATGACGTTTTTTCTATGTATTCAAAAACGTTGTTAATTCATTTGAACATGAG
Consensus                                        *  * *  * *                                                                              *

Teleogryllus_occipitalis    GATGGAAGGGAAGG-----------------------AAAGTTGATACTCGATCC------------------------------TTCATGAGAAG--------CGAAATACTGTACGAG
Acheta_domesticus           ------------------------------------------TTGTACATGGTTT-------------------------------------------------ACAAATGTCGGCCA--
Gryllus_longicercus         GTGAAAAGAGAAGGTTCACAAAATGGCATTTTATGGCACAATCCGTACATAGCTTTTTTGTACTGCTCCGGCATTCGCCTGGGCGTTTGTCGGAAAAAAGACGAAAAACTACCGATCAGA
Consensus                                                                ***                                                          ** *   *  *

Teleogryllus_occipitalis    TTAGACGATGG------------------------------------------TGACATTATGATGAATAAAT-----------------------------------------------
Acheta_domesticus           -----------------------------------------------------------------------------------------------TTTTTCTTTCGCGTCGTGC------
Gryllus_longicercus         ACAGCTGGTCGCTAGCATGCCCTTCCGGCCCGCGACTGAATTCGACTCAGCCATGTAACTATTGTGGAAAATCCTCCCCATCCGACCTTCCCGTGTCTTTCCCGCACGTCGTTCCACCCG
Consensus

Teleogryllus_occipitalis    -----GCAGTGAGTCGCTGTGAAGGTGGGATTGTGA-----------------------------------------------------ATAAGAGTCGGAGCACGTAAATGCGTATGGC
Acheta_domesticus           --------GCCATTTTTTGCGTTTGCAGGTATT---------------------------------------TACTTTTCTATTCAGCTCTAGATGTTCTAGCATTCTAGAATA-AAAAA
Gryllus_longicercus         TCCTGCTAGTCAGTCACCCTGCCCGCACGTACTCAACGCTTACTGTCCGTCCTCACTGCCCACCTCACCACCCACTCTTCTTCCGCGCAGTCGAAATTCCAACATC-------------G
Consensus                           *  * *      *   *   *                                                             *     *  * **

Teleogryllus_occipitalis    ATGTTGGGAATGGCATGTTAATCCGAGGGAGAAATGAAGAAATGAAATGAGCGAAATATATGG---------------------------------------------GATATAGTGCA-
Acheta_domesticus           GCATTATTAATTCAGTAGCC-TCTGTTCTACAAATCGAAATATCATTTGCACATCTTCTGT----------GTTTTTATTTGGGA-----------------------AATATATTGCC-
Gryllus_longicercus         ACATTGTCAACTACATAACTACCTATCATATAAGTTTCGAAAATAATTGAGGTTCTTTTATAAGTGTATAAAGCTCTCGTTCAAGAAAATTGCCTATGCAACTTGTTTAAAATATTGCTT
Consensus                      **  **     *      *      * ** *    * *  *  **       * * *                                               * *** ***

Teleogryllus_occipitalis    -----------GATAAGGTTGAGGGACATTAG------------------AACGGGGTG---ATGTGGAATAGGGGATGAGGTCAGGAGACACAGAGAGGATGAAC------------AC
Acheta_domesticus           -----------ACCAAATCTGTGGAACGTGAATGAACGAAATTCGATTTTCAGAGAGTA-------------------------------------------------------------
Gryllus_longicercus         GATTGTCTTTGTGCAAATTTGAAAACAACTAACAAGACGATGCTTGTTTTCAAAGAATGTACTTATGGAATAATACATAACATTAACAACTACTCGTATAATAAACTATTTTATTTGAAG
Consensus                                 **   **         *                    *  *  *

Teleogryllus_occipitalis    CAGAACGGGGGGAGTTAAGGATGGCGAGAGTTGAGG----ACGGCGGGAGACCACAGGGGTTCGGGCATAGCGTAATGATATACCTCCTCAGTCTATGGTTCG--GGGGAAGTTGAGAAT
Acheta_domesticus           ----------ATTCTCAAAGAAGAATAGAATCAGAG------CCTTAATTTTCCTATACTTTTAAACGTAAAAGTATAGTTACTGTTTTTTGTAAACTATCAA---AATCAAACTAGAAT
Gryllus_longicercus         TAATCTAAGTAATCTAAAAATCGACTAGATTCAAAGGGTACCCCTTAAATACCCAACATATTTTGCCAAGGAAATA---------TTCCCTCCAAAATACCCAATGAATTAATTTAAGCC
Consensus                                 * **    *   *** *    *                *  *    **    *        *         *         *             *    *

Teleogryllus_occipitalis    GGGAGACAAAGACGGGGAAGGGAAGAACGGGAAATGAGAAGACGGAATGG------------------------AAAGTCTCTGGATGTGAGGCCGGGAGGAAGCGCGGTGGCCGACGGC
Acheta_domesticus           AAACTATGTCGACTGTGTAAGAAAGAGTGAGCGTTTCATGGTGAAA----------------------------TAGATATTGTAAAGA--GCTCAAAAACAAGCTT-------------
Gryllus_longicercus         AGATTATGT------AGTAGGAAATTGGTTTCATTTCGAAGTGACAATCTAATTATCTAAACCGAAACACGATTTAAATCTCATAATG-----TAAATAGGAAACCTAATGGC-----AC
Consensus                        *          * * * **          *     *    *                             * * *    * *          *  ** *

Teleogryllus_occipitalis    CCGCTGCGAGAGAGAGCGGCAACGTCGTGCGGGGGAGTGAGCAACGTTCGCCCAGAGCAGCTGATGCAAATGGCT--------------GGCTGGCGCAAGGACACGGCGCGCCGCGGCG
Acheta_domesticus           ---TCACATGAATATTGCAAGTTCTCTCGC-------TGAAAAAAGACTATTTCACGAAGCCAGGTAAGTCGTCTCTGATTAAT------------------------------------
Gryllus_longicercus         GAACTACGTAAATAAGACATATT-----------------GAAATATCTGCCGACAGAAATTATCGAAGTCATTTATCGTAAATTTTTGGGCTGAAGCAAACAAATG-------------
Consensus                         *   *                               **            * *        *      *

Teleogryllus_occipitalis    GCGGCGGCGGCGGCGGCGGCGGCAGCGGCGGCGTCTGGGGAGGTGGCGAGGGGTTTGCGTGTGGCGCCGCCGA---------CGCCCGGCGCCCGCGGGGGGGAGGGGGGCGGGGGAGGG
Acheta_domesticus           ------------------------------------------------------------AAAATACCTATAA----------AATATCAGCGGTTGAAGGTATTGGAAAACCTACTAA
Gryllus_longicercus         --------------------------------------------------------TTAAACAACACTCCTAATTCGAGTTTTCCGTGCCAATTGTTGGAGATAACCATACACGGATTAG
Consensus                                                                                     *     *                *    *  *  *  *

Teleogryllus_occipitalis    GCAGGGTGGAGCGGGGGACGGCCTCGACGGGGCGAGGAAGGGCGGGGCGGGGCGGAGCGGGGCAGTGGAGGGGAGGGGAGGGGCGGGGCGGGGCCGTGCTGGGCGGGGCGGGGTCAGAG
Acheta_domesticus           ACTACGTTGTGTTTATGACTGAAATCTGTATAAATG--------------------------------------------------------------TATCCGTTTCG-ATAGTTACA-
Gryllus_longicercus         ACCATGTAAGGTACTACATAATTATCTGCATAACTGA------------------------------------------------------------TCTCCAGTTTCGTGTAGTTATAC
Consensus                    *   **   *      *      **                                                                             *    *    ** * *

Teleogryllus_occipitalis    CGAGCTAGGAGGCCGCGACCTTCCGCCTGGCGCCGCCCGAGCGCTGCCAACACGCCGTCCCTCCTGTCTGTCCCAGCCCCAGACGCAAGGCGGGTGCCTTCCTACCTCTCCGACATCGGC
Acheta_domesticus           -------------------------------ATTAAAATATTACTTATTATTGGTTTTATAATTTGT---CCCGCTTTTTCCCAACAAATCACGCTTTTTTTTCACTA------------
Gryllus_longicercus         -------------------CTTGCATTTGACATTGAATAACGTGATATTACAGATGTAATATCTTATGTGCCAAATTTTCACAAACTTATTACACACTATTTTAATTATTTGTCTTAGTT
Consensus                                                          *         *              * *    *             *             *  *   *

Teleogryllus_occipitalis    GACGATTCACTCGCTGCGAACATT--------------TGCATGCGGTTCTCCTGTCTATCCTCGCTCT---CTCATCCCAAGGGGATCGTCCAATGTGGAATT-----------TCTGC
Acheta_domesticus           -----------------------------------------------------AGATTATTAAAAATCACGACTTGGAGGAAAAAAATCTGGCAACGCTGAATGCAGATATAACAGGTAT
Gryllus_longicercus         GTACAGTCATGCAATAATTTCATTATTATTCCATATAACATAGATAATTGACGAGATTGTTACTGAT-----TTCGAAGTTAAGACATTATATAAACCATAACCGAGCAACTACATACAA
Consensus                                                                         *  * *      *      *       *    **     **     **

Teleogryllus_occipitalis    GCATTCCAGTCTAAATTGAGTTCTGCAGCCCCTTTAGCGACATTTGATCGACTCCTGAATAGACTGCCGTTGAAAGCAAATGTGTCTGTCGAATATGTAACAACGCTAATTTAATTTTGT
Acheta_domesticus           ATAAACTCGTTCATGTTT-----------------------------------------------------------------------ACATAAGGTAATACCTCAAAT----GTTTCA
Gryllus_longicercus         ATATACTCGCGCAAAAATAG---TGTTGCTAATTCAGCAACAGAAGATGTATT-----AGGGAATTTTAATGAAGAGAAAGATAATGAAAGAGAATGTGAGAATAGAAAT----CTTTCT
Consensus                     *  *  *   *                                                                              *  * ** * *     ***     ***

Teleogryllus_occipitalis    TAACAATTG---CAATATATAGTACAAGAGAAAATGGTGAGAGATTATAC-TGAATTATTGGTGTGTAATTTGAATTTTAC-----------------ATTATGCTGATTAAT-------
Acheta_domesticus           AAATAATTG---TGGTATTTTTATCAATGGT------TTGAAAATTGTGG----------------TGACCCGGCATTTGT---------------------------------------
Gryllus_longicercus         AAATAGACAAACAAATATTTATTTCAAGATTTTCCCATTGAAAATTCTGGTTTTATCATGAAAATATGAATCAGAATTTATAGTATAAATTCATAACGATTACGCTCAGTCATATACTAT
Consensus                    ** *          *** *    ***          *   * *** *                  * *       ***

Teleogryllus_occipitalis    ------------------------------------------------------TCTCTGAAGGTCTGCGATAAGCTTCC-TGAGAAAATTTAACTTTCTTTATT---------------
Acheta_domesticus           --------------------------------------------------ATTCTCTTTGAAAACGTTCA-----------AGAAAAGACACAAA-------------------------
Gryllus_longicercus         TTAGGAAAGTAATTGATTTACGTTATATCGCGCAATTATTTTTCTATAAAATTATCTCTGAATGAGTGCACTAAAATAATGAGTGAGGAAATAAACTTACTAATTCAATACAGTAATGCT
Consensus                                                                         *** ****    * *             *  *  *   **

Teleogryllus_occipitalis    -ATTTCATCGAATT----------------------ATGTCGAATATAAAACAACT----------------------------TCGCATCATGATTTTATTCTGTATAAATAAA-----
Acheta_domesticus           -TACACATCACATTTGAA---------------------------------------------------------------------TATCAAAATGTATTTGTGT--------------
Gryllus_longicercus         TTGCTCATCGTATCTGTGAAGGATAGCCGCAGGAATATGCCGAGGGGAAAATGTTTATTTTCAGAATATTTTATTTCCTGTTGCTCTTACGAAAACATAATTATGTAAAGAGGGGAACAC
Consensus                        ****  **                                                                           *  *  *  * ** ***

Teleogryllus_occipitalis    -------------------------------------------------------------TAACAAATGTTTATTTAATACACCAACAGACT-------TTTGCTTAGAAAT--AAGTT
Acheta_domesticus           -------------------------------------------------------------TATTGATTGTAATGTAATAACGTAAACAAAAA---------------------------
Gryllus_longicercus         CATCTTTACATTTACCTCAGAGAGACCAGAGGAACAGATATTATATATTGTATAATTGATGCATCGTTTGAAAAATAAAAACACCAGGAAATAATTCATCTCTACCCAAAATTTCAACCC
Consensus                                                                                 *     **     * *  **   *  * *

Teleogryllus_occipitalis    TGAGCATTCTTCTCACTCAAATTTCCT--------------------------------GTTTCTTTTAAAAATAAAGGT----------------------------------------
Acheta_domesticus           ------------------------------------------------------------------------------------------------------------------------
Gryllus_longicercus         CAAACATTTTTTTCTCCAAAAAATCCCCTAGCAGCGGACAAAATCTGCGAAAGCGCCGAGTTTATCCTCAGGGTTAAACCATCCGATCGCGTCCGCCGCCAGAGCGTTTGACGGGGAACG
Consensus

Teleogryllus_occipitalis    -----------------------TTGTG---GACTCTAGATATGTT---------------------------------------------------------------TTTAGTTGCTC
Acheta_domesticus           -----------------------TTGTATACAATTTGAGGTAAATA------------------------------------------------------------------TGTCGCAC
Gryllus_longicercus         CGGAATATCCAGCGAGGGAATCCTTGCACGCGAAACCAAACAAACAGCGCGAGCCGCGCAGGGGAACTCCCCGAAAAGCGCCGCGCCGCGTGGAGCTGTGCGGCGTTCGGCTGTTTGCGC
Consensus                                          ***      *    *   *                                                                        * ** *

Teleogryllus_occipitalis    AT---------------------------------
Acheta_domesticus           ATATATCTATATAGACTGAGAAGCT----------
Gryllus_longicercus         ACGCGTCTGAAACGGAAGAGCAGCGCACCTGGCGC
Consensus                   *
```

**Tocc.3G0002060.1_exon1**

```
Teleogryllus_occipitalis      ------------------------------------------------------------------------------------------------------------------------
Acheta_domesticus             --------------------------------------------------MINA------------------------------------------------------------------
Gryllus_longicercus           MATSSASRPSL*LKVFGNNVFKFIYFS*FFGKPDLTGIYGKLKQASL*L*RINATLL*RLYTSQIHTPHIGEHALSHN*TQAVSHPEEHTRKTAYTFRHTHTHTHTHTHTNSYIITTTHR
Consensus

Teleogryllus_occipitalis      ------------------------------------------------------------------------------------------------------------------------
Acheta_domesticus             ----------------VKGFAACYE*Q---------------------------------------------------------------------------------------------
Gryllus_longicercus           RHTTTHTGRTIFTDRHTRGLAHAYTNARAHTHTHTHTHKHSTHKT*RRS*RIAKHHIALLGVASNYVTSPHILKSPGASPKHLVAIVL*PSSGMKCRNI*HSLCHTVFHAC*INNTLGVL
Consensus

Teleogryllus_occipitalis      ------------------------------------------------------------------------------------------------------------------------
Acheta_domesticus             ------------------GISAA-------------------------------------------------------------------------------------------------
Gryllus_longicercus           YTAIHFADRVIGDHR*NFGISAQKNTTDELEFHKNRCFIKGAFNICLFYLYSTVYTFNCLPNQESEKVNALFDTTLREGAFIFTERFDFVIPQKETHTLGTFKTVLRNQFFY*NVT*K*Y
Consensus

Teleogryllus_occipitalis      ------------------------------------------------------------------------------------------------------------------------
Acheta_domesticus             --------------------------------CRVG-------------QVSKL------------------------------------------------------------------
Gryllus_longicercus           QYLTDSDL*LVAFFVHSNKIINRIHSK*IHFICRISLF*YCKYQTIVNIKLTTLEKFYENKIIQYTIFISFEPDDRKSSTSGLV*RNGCIIYFKFQNVSC*YSEYSDEYNNAQ*NIKMRI
Consensus

Teleogryllus_occipitalis      ------------------------------------------------------------------------------------------------------------------------
Acheta_domesticus             -------------RANLT------------------------------------------------------------------------------------------------------
Gryllus_longicercus           *NELIVCTTLMLNRANVTNASSHLL*CMDNTA*F*WVYDIHICIDLSLIRCLTPLKFNSFS*RIILYDMVSFEKRINILYEIFIQFFTQVII*RRLYELLM*KL*TSLFIRSRAMSSQIF
Consensus

Teleogryllus_occipitalis      ------------------------------------------------------------------------------------------------------------------------
Acheta_domesticus             -------------A*C--------------------------------------------------------------------------------------------------------
Gryllus_longicercus           RVILLQLH*LYWCKICNYSL*LKSNFFVVGTLILKTFERVNKNLENKVFVEFNVPRILITLEIDTFAAHAHQNCF*RILNLMCSIETFVARTFDKFHSRALAKPQSTSLPFSFILYHLAG
Consensus

Teleogryllus_occipitalis      ------------------------------------------------------------------------------------------------------------------------
Acheta_domesticus             ---------------------------------------SPSLP***AD-----------------------LLP-----RSHGSVL-------CSES----------------------
Gryllus_longicercus           HLCRNNFPERVVINNCPNRVSPAPPGLRLNRCQTPPSAESPRFPFPSQSSGRGAGVGERVLEGGGGDCGKPRLIPAGPGIRSRTQVFPGWALSRCTSSLSLDKHAIRAAAKRAGRVGLGM
Consensus

Teleogryllus_occipitalis      ------------------------------------------------------------------------------------------------------------------------
Acheta_domesticus             -DGRDGS----------------QQYIDSCHSLPT-WWQ---------WATARRGGRL--------------------------------------------------------------
Gryllus_longicercus           GGGRRGEGPNRTAHKKS**WDEGKWRIEATHVMPVRCYQGKKDNNKII*K**RKGGRQKS*QLY*SHMLLQAK*KKRPKRSRSHS*DGRR*LQQRMGQKF*SQNRRNVCEKNNVGKRRKS
Consensus

Teleogryllus_occipitalis      ---------------------------------------------------------------------MR-------------------------------------------------
Acheta_domesticus             -----------------------------------------------------------------ID*LIRVSGRV------------------------NYLGH*S--------------
Gryllus_longicercus           K*NCERQYTIRGEDGRCMYVCIYIYIYIYIYIYIERERERERERE*TDAVAECE*TIVI*RQCSNKMR*CLRRSGNVKSHMKYRTLTQRDRCSE***CWRSYASHNQIRMRCSSDGAHVSN
Consensus                                                                            *

Teleogryllus_occipitalis      ------------------------------------------------------------------------------------------------------------------------
Acheta_domesticus             -------------RKCSAC--------------------------------------------FSREVAMIFSPLDGEIWRRPTVD*RGFPLNQVSPTSGAPPQPGQAVSSRPVSRHVW*
Gryllus_longicercus           IQQTNPLQTSDSHRRQPACIPQEPSPFGGQPVQTLKILID*LIV**FRARQLPGSSPVSVMHVFAREVAVIFPPLYGEIWRRPTVG*RDFPQSGISHQRGTAP----AGTSSMLPINLPA
Consensus

Teleogryllus_occipitalis      ------------------------------------------------------------------------------------------------------------------------
Acheta_domesticus             C----SAC*CKCVCKRR---G*DKEEEGKIARGPWPVC*GTAPAFAWC*SEKPRKTTDQSSRPLEFSSRQRLNST*LTS*CRKSCRRRDSNLGPPGQRDCSDIRLMH*AISPGAWWKGMV
Gryllus_longicercus           CEVV*SMLV*VCV*GVRIGQGRGRSDRAR-SVALW---YGTAPAFAWC*SEKPRKTSDQSSRPVEFISRERLNSTQLTS*CRKSCHRRDSNLGPPGHSDC----LSHLSRLPTGLPTRMV
Consensus

Teleogryllus_occipitalis      ------------------------------------------------------------------------------------------------------------------------
Acheta_domesticus             -----------SMERG-----VLE---------WL*VLG--------------------------------------PRQ--------------------------------WAG---AR
Gryllus_longicercus           V*CKCVFGKSIGHRRGEKSHAVLDSCLRRCSGIRLILIGETMDQRSLPLEFSSRKRMNPTHPSS*YPPPSPSEIEPRPRRGGSGHHAMSLVSLTETS*FQVRRTAEFSTIGTWLGRIQSL
Consensus

Teleogryllus_occipitalis      ------------------------------------------------------------------------------------------------------------------------
Acheta_domesticus             ARRVKAR------------RRFEGACDRATTHTVGRARDGT------------TA*KKVQK*AEHAE---RSSEVPTPRKG---------------WVGGAGYR--QSTRG---------
Gryllus_longicercus           ARRES*RPAALISSFWWFFR*TPGE*RSSTL----RKRLGTERSFLPLSCPILSPYLH*HLVALHWEILIRKNSTSSPSRGGREPTPPTIKKACATLTGLTGYRA*SSINGIAVKAAKMQ
Consensus

Teleogryllus_occipitalis      ------------------------------------------------------------------------------------------------------------------------
Acheta_domesticus             -----------------------GRRS------TARSGEGYAYGQ-----AAAARDKMRP--------TLRPQR-------RR--APLMIQKVVQYLSS--------------CRLPRKS
Gryllus_longicercus           *QCCP*LPTAVVVDLADKLVSKIGRARLFVCVCVCRSG-GVCCGQSRREKAGEARLVS*PPSPENFSCSRRPARDPSGLRFRRLAAPAFIHSGLRGASNGLAHEAPPTFFSLQCETERTT
Consensus

Teleogryllus_occipitalis      ------------------------------------------------------------------------------------------------------------------------
Acheta_domesticus             V-----*EPQHRSLRLSRKNFIFCT*IS*VLTVQKHLKLFYC------------------TWFTNVGHFSFASCAIFCV-------------------------------------CRYL
Gryllus_longicercus           VKACHLRKPQH---AISE-----C*TFNLVL-INWHMTFFLCIQKRC*FI*T*GEKRRFTKW-----HFMAQSVHSFFVLLRHSPGRLSEKRRKTTDQNSWSLACPSGPRLNSTQPCNYC
Consensus

Teleogryllus_occipitalis      ------------------------------------------------------------------------------------------------------------------------
Acheta_domesticus             -------------------------------------------LFYSALDV-------------------------------------------------LAF*NKKALLIQ--------
Gryllus_longicercus           GKSSPSDLPVSFPHVVPPVLLVSHPARTYSTLTVRPHCPPHHPLFFRAVEIPTSTLSTT*LPII*VSKIIEVLL*VYKALVQENCLCNLFKILLDCLCANLKTTNKTMLVFKECTYGIIH
Consensus

Teleogryllus_occipitalis      ------------------------------------------------------------------------------------------------------------------------
Acheta_domesticus             -------*PLFY-----KSKYHL---------HIFCV------FIWEIYCHQICGT*M-----------NEIRFSESNSQRRIESEP*FSYTF--------KRKSIVTVFCKLSKSN*NK
Gryllus_longicercus           NINNYSYNKLFYLK*SK*SKNRLDSKGTP*IPNIFCQGNIPSKIPNELI*ARLCSRKLVSFRSDNLII*TETRFKSHNVNRKPNGTNYVNKTY*NICRQKLSKSFIVNFWAEANKC*TTL
Consensus

Teleogryllus_occipitalis      ------------------------------------------------------------------------------------------------------------------------
Acheta_domesticus             LCRLCKKE-----------------------------*AFHGE---IDIVKSSKTSFH------MNIASSLAEKR----LFHEAR---*VVSD-------------------**NTYKIS
Gryllus_longicercus           LIRVFRANCWR*PYTD*TM*GTT*LSA*LISSFV*LYLAFDIE*RDITDVISYVPNFHKLITHYFNYLS*LYSHAIISLLFHIT*IIDEIVTDFEVKTLYKP*PSNYIQIYSRKNSVANS
Consensus

Teleogryllus_occipitalis      ------------------------------------------------------------------------------------------------------------------------
Acheta_domesticus             AVEGI------GKPTKLRCVYD*NLYK---------------------CIRFDSYN----*NITYY----WFYNLSRFF----------------------------------------P
Gryllus_longicercus           ATEDVLGNFNEEKDNEREC-ENRNLSK*TNKYLFQDFPIENSGFIMKI*IRIYSINS*RLRSVIYYLGK*LIYVISRNYFSIKLSLNECTKIMSEEINLLIQYSNALLIVSVKDSRRNMP
Consensus

Teleogryllus_occipitalis      ------------------------------------------------------------------------------------------------------------------------
Acheta_domesticus             TNHAFFSLRLLKITTWRKKIWQR*MQI*QVYKLVHVYIR*YLKCFK---IIVVFLSMV*KLW*PGICIL---FENVQEKTQIHITFEYQ-------------------NVFVLLIVM**R
Gryllus_longicercus           RGKCLFSEYFISCCSYENIIM*RGEH--------HLYI--YLRETRGTDIIY--------------CIIDASFEK*KHQEIIHLYPKFQPQTFFSPKNPLAADKICESAEFILRVKPSDR
Consensus

Teleogryllus_occipitalis      -------------------------------------------------------------
Acheta_domesticus             KQKLYTI*G---------------KYVAHISI*TEK-------------------------
Gryllus_longicercus           VRRQSV*RGTRNIQRGNPCTRNQTNSASRAGELPEKRRAAWSCAAFGCLRTRLKRKSSAPG
Consensus
```

**Tocc.3G0002060.1_exon2**

```
Teleogryllus_occipitalis     -----------------------------------------------------------------------------------------------------------------------
Acheta_domesticus            -------------------------MINAVKGF----------------AACY------------------------------------------------------------------
Gryllus_longicercus          GYL*RIKTLIVTQSFRE*CF*IYLFQLIFR*A*FNWDLWETETSESLIIENQCYSTLETIHVTDSHTTHRGTCTLPQLDTSRQSPGGTHTQDRVHIQTHTHTHTHTHTH*QLHYYNNT*T
Consensus

Teleogryllus_occipitalis     --------------------------------------------------------------------------------GARR---------------------------------
Acheta_domesticus            --------------------------------------------------------------E*QGIS-------------AACR----------------------------VGQV-
Gryllus_longicercus          THYHSHGPHNIHG*THTRARARIHKRTRAHTHTHTHTQTLHTQDLT*KLTYSETPYRVAWRGLELRNFSPHPKEPRCKPKTFGGHRIVTVFRDEMSQYLTLPLSHRFPCMLNK*HVGRVI
Consensus                                                                                        *

Teleogryllus_occipitalis     -----------------------------------------------------------------------------------------------------------------------
Acheta_domesticus            -----------------------------------------------------------------------------------------------------------------------
Gryllus_longicercus          YSHSFRG*SHR*PPLEFWHFSPKEHNG*T*IS*KSMFYKGGF*YLLILFV*YSIHIQLLTESRIGKSQCTL*YNTS*RGIYIHREI*FRNSTERNTYSRNI*NSLAQPIFLLKCNLKIIP
Consensus

Teleogryllus_occipitalis     -----------------------------------------------------------------------------------------------------------------------
Acheta_domesticus            -----------------------------------------------------------------------------------------------------------------------
Gryllus_longicercus          IFNRFRSMISSFLRTLK*NNKQNSF*INTFHLPH*FILILQISNDC*Y*INHIREILRKQNNPIYYIHFF*TG*SEI*YVRVSVTQRLYNIFQISECFMLIL*VQ**V**CSIEY*NENM
Consensus

Teleogryllus_occipitalis     -----------------------------------------------------------------------------------------------------------------------
Acheta_domesticus            -----------------------------------------------------------------------------------------------------------------------
Gryllus_longicercus          KRVNCVYNAHVKPR*CDKRI*PSFIMHGQYRLILMGL*HSHMH*FIFDTLFNTIKV*LIFLTNNFIRHGIIRKKN*YSL*NIHTIFHASHNLKTTL*IVNVKIINLIVHSFSSHVFTNFP
Consensus

Teleogryllus_occipitalis     -----------------------------------------------------------------------------------------------------------------------
Acheta_domesticus            -------------------------------------SKLRANLTA*C-------------------SPSL-----------------------------------P***ADL-----
Gryllus_longicercus          CNSFAITLIILV*DL*LFFITEIQFLRCRDINFKNV*ESK*KS*KQSICGIQRT*NINHFGN*YICSTRTPKLFLKNLKPDVFHRNICSTHVRQISFESPCQTTEHFSPLLIHSLSSCGT
Consensus

Teleogryllus_occipitalis     ---------------------------------------------------GGAGRAGLGGAGRG-----------------------------------------------------
Acheta_domesticus            ----------------------LPRS----------------------------------------------------------------------------HGSVLCSES---DG
Gryllus_longicercus          FMSK*FSRTCGN**LSK*GVAGTPRSPIEPVPNPTVC*VSPFSFPFSEFGPGGGGGGEGAGGGRRGLRETTPNPCGSRNPESNTSLPGVGFKPLHFVTVA*QTRNQSGGEAGGESGAGDG
Consensus

Teleogryllus_occipitalis     -----------------------------------------------------------------------------------------------------------------------
Acheta_domesticus            ---------RDGSQQYI-------------DSCHSLP----------------------------------------------------------------------------------
Gryllus_longicercus          GGAPRRRTQQDGAQEVIVMGRRKVENRSDACNAC*MLPGQER*Q*NNMKVVKKRREAKELAIILVTHVIAGKMKEETKEV*ES*LRRQTVTTAENGSEILKPEPKKRL*EK*CWQAQKKQ
Consensus

Teleogryllus_occipitalis     -------------WA--------------------------------------------------------------------------------------------------------
Acheta_domesticus            ---------TWWQWA----------------------------TARRGGR---------------------------------------------------------------------
Gryllus_longicercus          MKL*KTIHNPWRRWALYVCMYIYIYIYIYIYIYRERERERERGINRRSGRVRVNHCHIKTVF**DEIMPSQKRKRQEPYEISHLDTKGQMQ*IIVMLAKLCQP*SDQDEVLQRWCTCLQY
Consensus                                  **

Teleogryllus_occipitalis     ---------------------------------------------------------------------------GLGG----------------------------------------
Acheta_domesticus            ---------------------------------------LID*LI---RVSGRVNYLGH*SRKCSACFSREVAMIFSPLDGE----IWR--RPTV-D*RGFPLNQVSPTSG---------
Gryllus_longicercus          SADQPFADI*QPPASTRLHPSRTVPFRGAAGTDS*DLD*LIDCLVIPCTSTTRVK-----SRKCNAC-------VCSGGGGDLSPTVWRDLAPTYRRLEGFSPIRYLPPAGHRPSRDKLY
Consensus                                                                                   *

Teleogryllus_occipitalis     -----------------------------------------------------------------------------------------------------------------------
Acheta_domesticus            APPQ-PGQ-------AVS------SRPVSRHVW*CSAC-----*CKCVCK---------------------------------------------------------------------
Gryllus_longicercus          APDQSPGM*GCVEHVSVSVCVRGEDRTRKRKVRPCAVCGPLVRYCSGICMVLIRETTEN*RSEQSAGRIHFQRATEFDPADKLVSKILPQTGLEPGTSRSQRLSESSLQAPDWSSHTNGS
Consensus

Teleogryllus_occipitalis     -----------------------------------------------------------------------------------------------------------------------
Acheta_domesticus            ------RRG*DKEEEGKIARGPW---------------------------------------------PVC*GTAPAFAW---------------------------------------
Gryllus_longicercus          IV*MCVREEYRTQKRGKIARCP*LVFKALLRHSPDIDRRNYGPTQPAARILLQKADESDPPIQLVSTTVTV*N*TAPEERWLRASCDVTSVSNRDVLVSSPEDGRIFDNRYMARSNSVSS
Consensus

Teleogryllus_occipitalis     -----------------------------------------------------------------------------------------------------------------------
Acheta_domesticus            --------C*SE--------KPRKTTDQSSR---------PLEFSSRQ------RLNST*L-------------------TS*CRKSCR-----------------------------
Gryllus_longicercus          QKGILAASCTDQ*FLVVFPINPR*MTKQYPKKEARD*AVFPSSFLSHPLPLLTLTLSCTTLGDTD*EKLH*LTIERGKGTDTSHDKKSLRYTYGAYGLPRLIINQWNRGESCKDAVAMLS
Consensus

Teleogryllus_occipitalis     ----AGRG-------------------------WAGL------GGRRG---------------------------------------------WAGLGG--------------------
Acheta_domesticus            -RRDSNLGPPGQRDCSDIR---LMH*AISPGAWWKGM-----VSMERGVLEWL*VL----------GPR-----------------------QWAGARA---------RRVKARRRFEG-
Gryllus_longicercus          LTSDSGRGRPS*QASVENR*GALVCVRVCVSQWWCVLRAEPAREGRRGPFGVLAAFARELLLFSAPGPRPFRASISTPGGSGFHPLGTPRSVQWSGPRGPTNFLFSPMR----NRKNNGK
Consensus                           *                         *            **                                            * *

Teleogryllus_occipitalis     -----------------------------------------------------------------------------------------------------------------------
Acheta_domesticus            --------ACDRAT-----------------------THTVGRARDGTTA*KKVQK*AEHAERSSEV--PTPRKGW---------------VGGAGYRQSTR------------------
Gryllus_longicercus          GVSFTKTTACDFGMLNFQSGIN*LAYDVFSMYSKTLLIHLNMR*KE------KVHKMAFYGTIRT*LFCTAPAFAWAFVGKKTKNYRSEQLVASMPFRPATEFDSAM*LLWKILPIRPSR
Consensus

Teleogryllus_occipitalis     -----------------------------------------------------------------------------------------------------------------------
Acheta_domesticus            --GGRRSTARSGEG------YAYGQAAAARDKMRPTLRPQRRRA--------------------------------PLMIQKVVQYLSSCR---------------------------
Gryllus_longicercus          VFPARRSTRPASQSPCPHVLNAYCPSSLPTSP--PTLLPRSRNSNIDIVNYITTYHISFENN*GSFISV*SSRSRKLPMQLV*NIA*LSLCKFENN*QDDACFQRMYLWNNT*H*QLLV*
Consensus

Teleogryllus_occipitalis     -----------------------------------------------------------------------------------------------------------------------
Acheta_domesticus            --------------------------LPRK-SV*EPQHRSLR-----------------LSRKNFIF-------------------------------------------------CT
Gryllus_longicercus          *TILFEVI*VI*KSTRFKGYPLNTQHILPRKYSLQNTQ*INLSQIM**EIGFISK*QSNYLNRNTI*IS*CK*ET*WHELRK*DILKYLPTEIIEVIYRKFLG*SKQMLNNTPNSSFPCQ
Consensus

Teleogryllus_occipitalis     -----------------------------------------------------------------------------------------------------------------------
Acheta_domesticus            *IS*VLTVQKHLKLFYCTWFTNVGHFSFASCAIFCVCRYLLFYSALDVLAF*NKKALLIQ*PLFYKSKYHLHIFCVFIWEIYCHQICGT*MNEIRFSESNSQRRIESEP*FSYTFKRKSI
Gryllus_longicercus          LLEITIHGLDHVRYYIIICITDL-QF---RVVIPCI*H*IT*YYRCNILCAK-----------FSQTYYTL--F*LFVLVV---QSCNNFI-IIPYNIDN*RD--------CY*FRS*DI
Consensus

Teleogryllus_occipitalis     -----------------------------------------------------------------------------------------------------------------------
Acheta_domesticus            VTVFCKLSKSN--*NKLC-----RLCKKE*AFH----------------GEIDIVKSSKTSF---HMNIASSLAEKRLFHEAR*VVSD**NTYKISAVEGIGKPTKLRCVYD*NLYKCIR
Gryllus_longicercus          I*TITEQLHTNILAQK*CC*FSNRRCIREF**RER**KRM*E*KSF*IDKQIFISRFSH*KFWFYHENMNQNL*YKFI-------------TITLSHIL-FRKVIDLRYIAQLFFYKIIS
Consensus

Teleogryllus_occipitalis     -----------------------------------------------------------------------------------------------------------------------
Acheta_domesticus            FDSYN*N---------ITYYWFYNLSRFFPTNHA---------FFSLRLLKITTWRKK------IWQR*MQI*QVYKLVHVYIR*YLKCFKIIVVFLSMV*KLW*PGICILFENVQEKTQ
Gryllus_longicercus          E*VH*NNE*GNKLTNSIQ*CFAHRICEG*PQEYAEGKMFIFRIFYFLLLLRKHNYVKRGTPSLHLPQRDQR--------NRYYILYN*C---------IV*KIKTPG------NNSSLPK
Consensus

Teleogryllus_occipitalis     -------------------------------------------------------------------------------------------------------
Acheta_domesticus            I---HITFEY------QNVFVLLIVM**RKQKLY----TI*G-------------------------------------------KYVAHISI*TEK----
Gryllus_longicercus          ISTPNIFFSKKSPSSGQNL-------RKRRVYPQG*TIRSRPPPERLTGNAEYPARESLHAKPNKQREPRRGTPRKAPRRVELCGVRLFAHASETEEQRTWR
Consensus
```

**Tocc.3G0002060.1_exon3**

```
Teleogryllus_occipitalis         ------------------------------------------------------------------------------------------------------------------------
Acheta_domesticus                ------------------------------------------------------------------------------------------------------------------------
Gryllus_longicercus              GYL*RIKTLIVTQSFRE*CF*IYLFQLIFR*A*FNWDLWETETSESLIIENQCYSTLETIHVTDSHTTHRGTCTLPQLDTSRQSPGGTHTQDRVHIQTHTHTHTHTHTH*QLHYYNNT*T
Consensus

Teleogryllus_occipitalis         ------------------------------------------------------------------------------------------------------------------------
Acheta_domesticus                ------------------------------------------------------------------------------------------------------------------------
Gryllus_longicercus              THYHSHGPHNIHG*THTRARARIHKRTRAHTHTHTHTQTLHTQDLT*KLTYSETPYRVAWRGLELRNFSPHPKEPRCKPKTFGGHRIVTVFRDEMSQYLTLPLSHRFPCMLNK*HVGRVI
Consensus

Teleogryllus_occipitalis         ------------------------------------------------------------------------------------------------------------------------
Acheta_domesticus                ------------------------------------------------------------------------------------------------------------------------
Gryllus_longicercus              YSHSFRG*SHR*PPLEFWHFSPKEHNG*T*IS*KSMFYKGGF*YLLILFV*YSIHIQLLTESRIGKSQCTL*YNTS*RGIYIHREI*FRNSTERNTYSRNI*NSLAQPIFLLKCNLKIIP
Consensus

Teleogryllus_occipitalis         ------------------------------------------------------------------------------------------------------------------------
Acheta_domesticus                -------------------------------------------ND---------------------------------------------------------------------------
Gryllus_longicercus              IFNRFRSMISSFLRTLK*NNKQNSF*INTFHLPH*FILILQISNDC*Y*INHIREILRKQNNPIYYIHFF*TG*SEI*YVRVSVTQRLYNIFQISECFMLIL*VQ**V**CSIEY*NENM
Consensus

Teleogryllus_occipitalis         ------------------------------------------------------------------------------------------------------------------------
Acheta_domesticus                ------------------------------------------------------------------------------------------------------------------------
Gryllus_longicercus              KRVNCVYNAHVKPR*CDKRI*PSFIMHGQYRLILMGL*HSHMH*FIFDTLFNTIKV*LIFLTNNFIRHGIIRKKN*YSL*NIHTIFHASHNLKTTL*IVNVKIINLIVHSFSSHVFTNFP
Consensus

Teleogryllus_occipitalis         ------------------------------------------------------------------------------------------------------------------------
Acheta_domesticus                --------------------------KCR----------------*RLCGMLRV------------------------------------------------------------------
Gryllus_longicercus              CNSFAITLIILV*DL*LFFITEIQFLRCRDINFKNV*ESK*KS*KQSICGIQRT*NINHFGN*YICSTRTPKLFLKNLKPDVFHRNICSTHVRQISFESPCQTTEHFSPLLIHSLSSCGT
Consensus

Teleogryllus_occipitalis         ----------------------------------------------------GGGGGAGWSGGR----------------------------------------------------ASTG
Acheta_domesticus                ------------------------------------------------------------TGHKRRL-----------------------------------------------------
Gryllus_longicercus              FMSK*FSRTCGN**LSK*GVAGTPRSPIEPVPNPTVC*VSPFSFPFSEFGPGGGGGGEGAGGGRRGLRETTPNPCGSRNPESNTSLPGVGFKPLHFVTVA*QTRNQSGGEAGGESGAGDG
Consensus                                                                      *

Teleogryllus_occipitalis         RG----------------------------------------------------------------------------------------------------------------------
Acheta_domesticus                ---PRRTSQ--------------------------------------------------------------------------------*TTRKPDSVMFTKSP---------------M
Gryllus_longicercus              GGAPRRRTQQDGAQEVIVMGRRKVENRSDACNAC*MLPGQER*Q*NNMKVVKKRREAKELAIILVTHVIAGKMKEETKEV*ES*LRRQTVTTAENGSEILKPEPKKRL*EK*CWQAQKKQ
Consensus

Teleogryllus_occipitalis         ----------------------------------------------RAGR----------------------------------------------------------------------
Acheta_domesticus                MMSGSTAAISWKRSM*------------------------------RI*RARWFPTIYRQLPLAPDLVAVGHC--EAWWKVD*LIDSCFFRARQ-------------------LLRSL---
Gryllus_longicercus              MKL*KTIHNPWRRWALYVCMYIYIYIYIYIYIYRERERERERGINRRSGRVR-----------------VNHCHIKTVF**DEIMPSQKRKRQEPYEISHLDTKGQMQ*IIVMLAKLCQP
Consensus                                                                      *  *

Teleogryllus_occipitalis         ----------------------------------------------------------------------------------GGAG----------------------------------
Acheta_domesticus                ----------------------------------------VP--------------*V*CVFF------------------SGGGDDLFPAGWRDLAPTYRRLAGFSP---QPGISHQRG
Gryllus_longicercus              *SDQDEVLQRWCTCLQYSADQPFADI*QPPASTRLHPSRTVPFRGAAGTDS*DLD*LIDCLVIPCTSTTRVKSRKCNACVCSGGGGDLSPTVWRDLAPTYRRLEGFSPIRYLPPAGHR--
Consensus                                                                                          **

Teleogryllus_occipitalis         ------------------------------------------------------------------------------------------------------------------------
Acheta_domesticus                TAPAGTSCKLPT---GLPACVVV*CMLV*VCV*EERIGQGRGRKDRARSVA---RLLGYCSGIRLVLIGETTENYRSEQPAARILLQTATEFDLADKLVSKILPPSGLEPRTSRSEGLQ*
Gryllus_longicercus              --PSRDKLYAPDQSPGM*GC--VEHVSVSVCVR----GEDRTRKRKVRPCAVCGPLVRYCSGICMVLIRETTEN*RSEQSAGRIHFQRATEFDPADKLVSKILPQTGLEPGTSRSQRLSE
Consensus

Teleogryllus_occipitalis         ------------------------------------------------------------------------------------------------------------------------
Acheta_domesticus                HQVDA---LSHLSRCLVEGDGVDGEGGP*VAMSLRPETMGGGTREAREGA---------AAL*RSVRPRYNSHSGQGARRHHCLEEGAKISRTCRKKL*SAHPA-----------KGVG
Gryllus_longicercus              SSLQAPDWSSHTN------------GSIV*MCVREE-----YRTQKRGKIARCP*LVFKALLR--------HSPDIDRRNYGPTQPA--ARILLQKADESDPPIQLVSTTVTV*N*TAP
Consensus

Teleogryllus_occipitalis         ------------------------------------------------------------------------------------------------------------------------
Acheta_domesticus                RRRWVQA---------------------------------------------------------------KYPRRQK----------INCPL---------------------------G
Gryllus_longicercus              EERWLRASCDVTSVSNRDVLVSSPEDGRIFDNRYMARSNSVSSQKGILAASCTDQ*FLVVFPINPR*MTKQYPKKEARD*AVFPSSFLSHPLPLLTLTLSCTTLGDTD*EKLH*LTIERG
Consensus

Teleogryllus_occipitalis         ---------------------------QWRGGE---------------GRG------------------GAVL-----------------------------------------------
Acheta_domesticus                GGV------------------------RLRAG----------------GRGARQNAADPTTAAAA----GAFD-----------------DTKGRAIPF-VLPATAKECL----------
Gryllus_longicercus              KGTDTSHDKKSLRYTYGAYGLPRLIINQWNRGESCKDAVAMLSLTSDSGRGR------PS*QASVENR*GALVCVRVCVSQWWCVLRAEPAREGRRGPFGVLAAFARELLLFSAPGPRPF
Consensus                                                        *                ***                  **

Teleogryllus_occipitalis         -------GGA--------------------------------------------------------------------------------------------------------------
Acheta_domesticus                --------------------------RAPTQ---------------VASTISKKLHFLYINF------LSFDC-SKTS*TFLLYMVYKCRP-----FFFRVVRH--FLRLQVFTFLFSSR
Gryllus_longicercus              RASISTPGGSGFHPLGTPRSVQWSGPRGPTNFLFSPMRNRKNNGKGVSFTKTTACDFGMLNFQSGIN*LAYDVFSMYSKTLLIHLNMR*KEKVHKMAFYGTIRT*LFCTAPAFAWAFVGK
Consensus

Teleogryllus_occipitalis         ------------------------------------------------------------------------------------------------------------------------
Acheta_domesticus                CSSILE*KSIINSVA------------SVLQIEISFA-------------------HLLCFYLGNIL---PPNLW------NVNERNSIFRE*FSKKN-----------RIRAL------
Gryllus_longicercus              KTKNYRSEQLVASMPFRPATEFDSAM*LLWKILPIRPSRVFPARRSTRPASQPSPCPHVLNAYCPSSLPTSPPTLLPRSRNSNIDIVNYITTYHISFENN*GSFISV*SSRSRKLPMQLV*
Consensus

Teleogryllus_occipitalis         ------------------------------------------------------------------------------------------------------------------------
Acheta_domesticus                ----------------------IFLYF*T*KYSYCFL*TI---------------------------KIKLE*TMSTV*ERVSVSW*NRYCKELKNKLSHEYCKFSR*KKTISRSQVSRL
Gryllus_longicercus              NIA*LSLCKFENN*QDDACFQRMYLWNNT*H*QLLV**TILFEVI*VI*KSTRFKGYPLNTQHILPRKYSLQNTQ*INLSQIM**EIGFISK*QSNYLNRNTI*IS*CK*ET*WHELRK*
Consensus

Teleogryllus_occipitalis         ------------------------------------------------------------------------------------------------------------------------
Acheta_domesticus                *LIKYL------------------------------------------------------------------------*NISG*RYWKTY*TTLCL*LKSV-----------------*M
Gryllus_longicercus              DILKYLPTEIIEVIYRKFLG*SKQMLNNTPNSSFPCQLLEITIHGLDHVRYYIIICITDLQFRVVIPCI*H*IT*YYRCNILCAKFSQTYYTLF*LFVLVVQSCNNFIIIPYNIDN*RDC
Consensus

Teleogryllus_occipitalis         ------------------------------------------------------------------------------------------------------------------------
Acheta_domesticus                YPFR*LQLKYYL-------LLVL*FVPLFPNK------------------------------SRFFFTKIIKNHDLEEKNLAT--LNADITGI*TRSCLH-KVIPQMF-----------Q
Gryllus_longicercus              Y*FRS*DII*TITEQLHTNILAQK*CC*FSNRRCIREF**RER**KRM*E*KSF*IDKQIFISRFSH*KFWFYHENMNQNL*YKFITITLSHILFRKVIDLRYIAQLFFYKIISE*VH*N
Consensus

Teleogryllus_occipitalis         ------------------------------------------------------------------------------------------------------------------------
Acheta_domesticus                NNCG-----------------------------IF-INGLKIVVTRHLY------SL*KRSRKDTNTHHI*ISKCICVIDCNV---------IT*TKIVY---------NLR--------
Gryllus_longicercus              NE*GNKLTNSIQ*CFAHRICEG*PQEYAEGKMFIFRIFYFLLLLRKHNYVKRGTPSLHLPQRDQRNRYYILYN*CI-V*KIKTPGNNSSLPKISTPNIFFSKKSPSSGQNLRKRRVYPQG
Consensus

Teleogryllus_occipitalis         -------------------------------------------------------------------
Acheta_domesticus                ----------------------------------------------*ICRTYIYID*EA--------
Gryllus_longicercus              *TIRSRPPPERLTGNAEYPARESLHAKPNKQREPRRGTPRKAPRRVELCGVRLFAHASETEEQRTWR
Consensus
```

**Tocc.3G0002060.1_exon4**

```
Teleogryllus_occipitalis      ------------------------------------------------------------------------------------------------------------------------
Acheta_domesticus             ------------------------SFSVYI---DICATY---------LPQIVYNFCLRYYI-------------------------------------TINNTNTF*YSNVMCI-----
Gryllus_longicercus           MATSSASRPSL*LKVFGNNVFKFIYFS*FFGKPDLTGIYGKLKQASL*L*RINATLL*RLYTSQIHTPHIGEHALSHN*TQAVSHPEEHTRKTAYTFRHTHTHTHTHTHTNSYIITTHR
Consensus

Teleogryllus_occipitalis      ------------------------------------------------------------------------------------------------------------------------
Acheta_domesticus             ---------CVFS------------------------------*TFSKRIQMPGHH--------NFQT---IDKNTTIILKHLRYYLM--------------*T*TSLYTC---------
Gryllus_longicercus           RHTTTHTGRTIFTDRHTRGLAHAYTNARAHTHTHTHTHKHSTHKT*RRS*RIAKHHIALLGVASNYVTSPHILKSPGASPKHLVAIVL*PSSGMKCRNI*HSLCHTVFHAC*INNTLGVL
Consensus

Teleogryllus_occipitalis      ------------------------------------------------------------------------------------------------------------------------
Acheta_domesticus             YICIQ------------------------------RCQI--------FFL--QVVIFNNLSEKKA*FVGKKRDKL*NQ**VIF-----*L*LSKRIH---------------------LY
Gryllus_longicercus           YTAIHFADRVIGDHR*NFGISAQKNTTDELEFHKNRCFIKGAFNICLFYLYSTVYTFNCLPNQESEKVNALFDTTLREGAFIFTERFDFVIPQKETHTLGTFKTVLRNQFFY*NVT*K*Y
Consensus

Teleogryllus_occipitalis      ------------------------------------------------------------------------------------------------------------------------
Acheta_domesticus             RFQS*TQRSLVGFPIPSTADIL*VFY*SETTYLAS*NSLFS-ARELAIFM*KLV------------FELFTISISP*NAHSF------------------LHSRHSLF*FDFDSLQKTV-
Gryllus_longicercus           QYLTDSDL*LVAFFVHSNKIINRI--HSK*IHFICRISLF*YCKYQTIVNIKLTTLEKFYENKIIQYTIF-ISFEPDDRKSSTSGLV*RNGCIIYFKFQNVSC*YSEYSDEYNNAQ*NIK
Consensus

Teleogryllus_occipitalis      ------------------------------------------------------------------------------------------------------------------------
Acheta_domesticus             -----------TILL-RLKV*E--------------------------------------N*GSDSILL*ELLS-ENRISFIHVPQIWWQYISQI-------------------------
Gryllus_longicercus           MRI*NELIVCTTLMLNRANVTNASSHLL*CMDNTA*F*WVYDIHICIDLSLIRCLTPLKFNSFS*RIILYDMVSFEKRINILY--EIFIQFFTQVII*RRLYELLM*KL*TSLFIRSRAM
Consensus

Teleogryllus_occipitalis      ------------------------------------------------------------------------------------------------------------------------
Acheta_domesticus             KTQ---------------KMCK*YFDL*NRGY*I----------------NNAFLF*N------------------------------------ART-----SRAE*KSKYL--------
Gryllus_longicercus           SSQIFRVILLQLH*LYWCKICNYSL*LKSNFFVVGTLILKTFERVNKNLENKVFVEFNVPRILITLEIDTFAAHAHQNCF*RILNLMCSIETFVARTFDKFHSRALAKPQSTSLPFSFIL
Consensus

Teleogryllus_occipitalis      ------------------------------------------------------------------------------------------------------------------------
Acheta_domesticus             ---------------------------------------------------------------------------------------QTQ------------------------------
Gryllus_longicercus           YHLAGHLCRNNFPERVVINNCPNRVSPAPPGLRLNRCQTPPSAESPRFPFPSQSSGRGAGVGERVLEGGGGDCGKPRLIPAGPGIRSRTQVFPGWALSRCTSSLSLDKHAIRAAAKRAGR
Consensus

Teleogryllus_occipitalis      ---------------------------GKW*-----------------------------------------------------------------------------------------
Acheta_domesticus             ---------------KMAHDAKE-----KWPTFVNHVQ*KSLRCF-----------------*TVKT*EIYVQKM------------------------------KFFRDSRSDLCWGS-
Gryllus_longicercus           VGLGMGGGRRGEGPNRTAHKKS**WDEGKWRIEATHVM--PVRCYQGKKDNNKII*K**RKGGRQKS*QLY*SHMLLQAK*KKRPKRSRSHS*DGRR*LQQRMGQKF*SQNRRNVCEKNN
Consensus                                                 **

Teleogryllus_occipitalis      ------------------------------------------------------------------------------------------------------------------------
Acheta_domesticus             -------------QTLFRGSRQDERYCTTFCI-----------IKGARRR----------------------------CGRRVGRILS--------------------------------
Gryllus_longicercus           VGKRRKSK*NCERQYTIRG--EDGR-CMYVCIYIYIYIYIYIYIERERERERE*TDAVAECE*TIVI*RQCSNKMR*CLRRSGNVKSHMKYRTLTQRDRCSE***CWRSYASHNQIRMR
Consensus

Teleogryllus_occipitalis      ------------------------------------------------------------------------------------------------------------------------
Acheta_domesticus             -----------------------RAAAACP*AYPSP---------ERAVDLL--------------PPRVLCLYPA------PPTH--------------PFRGV---------GTSEL-
Gryllus_longicercus           CSSDGAHVSNIQQTNPLQTSDSHRRQPACIPQEPSPFGGQPVQTLKILID*LIV**FRARQLPGSSPVSVMHVFAREVAVIFPPLYGEIWRRPTVG*RDFPQSGISHQRGTAPAGTSSML
Consensus

Teleogryllus_occipitalis      ------------------------------------------------------------------------------------------------------------------------
Acheta_domesticus             ---LSACS-------------------------AYFCTFF*AVVPSRA----------------------------------------------------------------LPTV*VV-
Gryllus_longicercus           PINLPACEVV*SMLV*VCV*GVRIGQGRGRSDRARSVALWYGTAPAFAWC*SEKPRKTSDQSSRPVEFISRERLNSTQLTS*CRKSCHRRDSNLGPPGHSDCLSHLSRLPTGLPTRMVV*
Consensus

Teleogryllus_occipitalis      ------------------------------------------------------------------------------------------------------------------------
Acheta_domesticus             --------------ARSHA-------------------------------------------------PSK-------------------------------------------------
Gryllus_longicercus           CKCVFGKSIGHRRGEKSHAVLDSCLRRCSGIRLILIGETMDQRSLPLEFSSRKRMNPTHPSS*YPPPSPSEIEPRPRRGGSGHHAMSLVSLTETS*FQVRRTAEFSTIGTWLGRIQSLAR
Consensus

Teleogryllus_occipitalis      ------------------------------------------------------------------------------------------------------------------------
Acheta_domesticus             -----------------------------RRRAFTRLARAPAHC--LGPKTHSH---------------------SRTPL---------------------SIDTIPFHQAPGEMAQCIN
Gryllus_longicercus           RES*RPAALISSFWWFFR*TPGE*RSSTLRKRLGTERSFLPLSCPILSPYLH*HLVALHWEILIRKNSTSSPSRGGREPTPPTIKKACATLTGLTGYRA*SSINGIAVKAAKMQ*QCCP*
Consensus

Teleogryllus_occipitalis      ------------------------------------------------------------------------------------------------------------------------
Acheta_domesticus             LMS---------------------------------------------LQSL*PGGPRFESRRR--QD-----FR---------------------------------------------
Gryllus_longicercus           LPTAVVVDLADKLVSKIGRARLFVCVCVCRSGGVCCGQSRREKAGEARLVS*PPSPENFSCSRRPARDPSGLRFRRLAAPAFIHSGLRGASNGLAHEAPPTFFSLQCETERTTVKACHLR
Consensus

Teleogryllus_occipitalis      ------------------------------------------------------------------------------------------------------------------------
Acheta_domesticus             ---H*LVS*VEFS----------------RC---------------------------LEENSSGRLL*SVVFRGFSDQ-----------------------------------------
Gryllus_longicercus           KPQHAISEC*TFNLVLINWHMTFFLCIQKRC*FI*T*GEKRRFTKWHFMAQSVHSFFVLLRHSPGRL--SEKRRKTTDQNSWSLACPSGPRLNSTQPCNYCGKSSPSDLPVSFPHVVPPV
Consensus

Teleogryllus_occipitalis      ------------------------------------------------------------------------------------------------------------------------
Acheta_domesticus             ----HQANAGAV----P*QTGHGP---RAI-FPSSSLS-------------------YPLLLHTH---------LH*HALHYHTCRETGRELTACPGWG---------------------
Gryllus_longicercus           LLVSHPARTYSTLTVRPHCPPHHPLFFRAVEIPTSTLSTT*LPII*VSKIIEVLL*VYKALVQENCLCNLFKILLDCLCANLKTTNKTMLVFKECT-YGIIHNINNYSYNKLFYLK*SK*
Consensus

Teleogryllus_occipitalis      ------------------------------------------------------------------------------------------------------------------------
Acheta_domesticus             --------GAPLVGDTWLRGK-----PR----*STVGRRQISPSSGEKIIATS--------------------------REK--------------------------------------
Gryllus_longicercus           SKNRLDSKGTP*IPNIFCQGNIPSKIPNELI*ARLCSRKLVSFRSDNLII*TETRFKSHNVNRKPNGTNYVNKTY*NICRQKLSKSFIVNFWAEANKC*TTLLIRVFRANCWR*PYTD*T
Consensus

Teleogryllus_occipitalis      ------------------------------------------------------------------------------------------------------------------------
Acheta_domesticus             -------------------------------------------------------HAL-----HL--------RD**PK*LTRP----------------------------ETRINQ--
Gryllus_longicercus           M*GTT*LSA*LISSFV*LYLAFDIE*RDITDVISYVPNFHKLITHYFNYLS*LYSHAIISLLFHIT*IIDEIVTDFEVKTLYKP*PSNYIQIYSRKNSVANSATEDVLGNFNEEKDNERE
Consensus

Teleogryllus_occipitalis      ------------------------------------------------------------------------------------------------------------------------
Acheta_domesticus             ----------------------------------SIN-----------------------LPPRLAVAHCHQVGSE-----WQLS-----------------------------------
Gryllus_longicercus           CENRNLSK*TNKYLFQDFPIENSGFIMKI*IRIYSINS*RLRSVIYYLGK*LIYVISRNYFSIKLSLNECTKIMSEEINLLIQYSNALLIVSVKDSRRNMPRGKCLFSEYFISCCSYENI
Consensus

Teleogryllus_occipitalis      ------------------------------------------------------------------------------------------------------------------------
Acheta_domesticus             ---------------------IYC--------------------------WEPSRP---------SDSLHRTLP*DRGSRSAHHHGRLGEHHAVRFARSLLTCPTRQAALMPCHS*HAAK
Gryllus_longicercus           IM*RGEHHLYIYLRETRGTDIIYCIIDASFEK*KHQEIIHLYPKFQPQTFFSPKNPLAADKICESAEFILRVKPSDRVRRQSV*RGTRNIQRGNPCTRNQTNSASR-AGELP-EKRRAAW
Consensus

Teleogryllus_occipitalis      --------------------
Acheta_domesticus             PLTAFII-------------
Gryllus_longicercus           SCAAFGCLRTRLKRKSSAPG
Consensus
```

**Tocc.3G0015530.1**

```
Teleogryllus_occipitalis    TTTCGGGTTATCCACGAGAATAAAGTTTCTTGAAGAGTGACTTGTGTTGCTGGTTGCTGATGCGCCTGTTTACTCTCATCACCATATTCTGGGA-ACTTCTAT----------------
Acheta_domesticus           TTCCGGGTTATTCACGACGGTAACGTTTGTTGAAGAGTGGCTTGTATTGCGGGTTGCATGCCCGCCTGTTTTCTCTCTTCCCCAAATTCTGCTATATTTTTATTCTATCTTTTCTTTCCC
Consensus                   ** ******** *****   *** **** ********** ***** **** ******     ********* ***** ** *** ******  * * ** ***

Teleogryllus_occipitalis    ---------------------TTTTTTCGTCTTCAGTCTATTTGATGCAGGCGTGG-----------------------------------------------------------------
Acheta_domesticus           GTATTCTGCTGTGTGCTTCTATTTTTTTCGTCTTCATACCCTTT-ATGCAGGAGTGGGAGTGGCTCAGTCCCTTCTCAAGTGTAGTTACACACTTACGGAGTGCTGTGTGATGGTTTCCTA
Consensus                                        **************  *  *** ******* ****

Teleogryllus_occipitalis    ------------------------------------------------------------------------------------------------------------------------
Acheta_domesticus           TAAATGATACAACATCCTTATATTTCCTTTGCGAATTATGTATAGAGAGAACGTTTACAATGGGGCCGCCTGCTAGTCAGCCCGTGAATAAAGTTTTTTCGGTATGATTATTACCAAAGC
Consensus

Teleogryllus_occipitalis    -----------------------------------------------------------------------------------------------GGGTCCTCAGACTCTCATCTCGTAAT
Acheta_domesticus           ATGCTGAACTTTTAATTGATGAATGTCTCCATGGTTTCCAAATTGGAATTCAAGCCTCTCGACCAATCACATTCCTCGAATTTGGGAAATCCACGAGTCCTCCGTCACTCGTGTCGTAAT
Consensus                                                                                                  * ****** * * *** * *******

Teleogryllus_occipitalis    TATACTTGTAAATAATTGATACATACGTACGATAATTG-AAAACAAAATATTCTCAATGTTACTGAAATCCCCATTTTTGAATTTCAAAAATATGAAGAAAGTTTTAAATTAAGAAATTT
Acheta_domesticus           TATACCTACAAATAATAGGTACCTCTGTACGATTATTGAAAAAATATACAATTCTCAACCTCACTGAAACCCCCAATTTTTGAATTTCAAA------ATGATAGTTTTCAATTAAGAATTTT
Consensus                   ***** *  ******* * *** *  ******* **** **** * *  ********  * ******* *** ****************      * ** ****** ********* ***

Teleogryllus_occipitalis    TCAAACAAACGTTATGTGATGTGTTCGTAATCA---AAGACAAGTTACAGAAGTGAAGTTTTCGATTGGGTTCATTGCGAGCCACTGTGCGTTGGCGGCGCGGCGCGGCGCGGCGGGGCG
Acheta_domesticus           TCAATCGTACGATCTGTGATGAGTTTCTATTCAATCACGGCAAGTTCCAGAAGTGACATTTTCGATTGCGTTCATTGTGAGCCACTGTTCGCTGGCGGCGCGGCGCGGCGCGGCGGGGCG
Consensus                   **** *  *** * ******* ***  ** ***   * * ****** *********  ********** ******** ********** ** ****************************

Teleogryllus_occipitalis    GGGCGCTGGACCGCTCGCTTGCTCGCCCAGATGCCAGCGAGCGCCGATCGATGCACGCCCGCGGCCTCCCCCGCGCCGCCGCGCCGCGGCAACCGTTACTAAGGCGGCGGAGGCACGGCC
Acheta_domesticus           GGGCGCTGGACCGCTC----GCTCGCCCAGATGCCGGCGGGCGCCGATCGATGCACGCCCGCGGCCGCCCCCGCACCGCCGCGCCGCGGCAACCGTTACTAAGGCGGCGGAGGCACGGCC
Consensus                   ****************    *************** *** ************************** ******* ********************************************

Teleogryllus_occipitalis    TGGGCCGGGCGCGCGCCCGCCACCGCGCCTGCCCCCGCCGCCGCGCCCGTCCCCGCCCCCGCGTCCGCCCCGCCCCCGCGCCCGCCCCCGCGCCCCCCCCCGCGCCCGCCCCGCGCCCGC
Acheta_domesticus           TGGGCCGGGCCCG-----------------------------------------------------------------------------------------------------------
Consensus                   ********** **

Teleogryllus_occipitalis    CCGCGCCGCCCCCCGCCGCCCCCCGCGCCCGCCCCCGCGCCCGCCCCCGCGCCCGCCCCCGCGCCCGCGCCCGCCCCCGCGCCCGCCCCCGCGCCCGCCCCCGCGCCCGCCCCCGCGCCC
Acheta_domesticus           -------------------------------------------TCCCGCGCCCGCCCCCGCGCTCGCGCTC-------------------------------------------------
Consensus                                                             ******************* ***** *

Teleogryllus_occipitalis    GCCCCCGCGCCCGCCACCGCCACCGCGCCCGCCACCGCGCCCGCCACCGCGCCCGCCCCCGCCCCCGCGCCCGCCTCCGCGCCCGCCACCGCGCCCGCCACCGCGCCCGCCACCGCGCCC
Acheta_domesticus           ------------------------------------------------------------------------------------------------------------------------
Consensus

Teleogryllus_occipitalis    GCCACCGCGCCCGCCACCGCGCCCGCCACCGCGCCCGCCACCGCGCCCGCCACCGCGCCCGCCCCCGAGCAGGCACCCCGTTTGCGCAGGATCCTTCCTTCCCTCCCGCTCGGTCAACGC
Acheta_domesticus           -----------------------------------------------------------------------GCACCCCGTTTGCGCAGGATCCTTCCTTCCCTCCCGCTCGGTCAACGC
Consensus                                                                                          ************************************************

Teleogryllus_occipitalis    CATTGACTCTAACAAGCCCTGCCCCGGCCCGTGTCTCTGCCATTCATGCCCAGTCGCGCCGCACCGCGCCGCGCTGCCTTTTCAGTTCCTCGTCCGCTTGGATTTATCATCGACCCCGCC
Acheta_domesticus           CATTGACTCTAACAAGCCCTGCCCTGGCCCGTGTCTCTGCCATTCATGCCCAG----------CCGCGCCGCGCTGCCTTTTCAGCTCCTCGTCCGCTTGGATTCGTCATCGACCCCGCC
Consensus                   ************************ ****************************          ********************** ******************  **************

Teleogryllus_occipitalis    TCCGCTGCTCCTGTTTTCCC----------TCCGATATCATCTTCGTCTGTTTCTG--CCGTTTTTTTCATTCCTCCTTCTTTCGTCGCTAG--TTGTCACTACGATCGCTTCTCCCCTT
Acheta_domesticus           TCCGCTGCTTCTGTTTTTCCCTCCGGCCGGTCCGATATCATCTTCGTCTGTTTCTGCCCCTTTTTTTTCATTCCTCCCTCTTCCGTCGCTCGTTTTGTCACTACTGTTGCTTCTCCCCTC
Consensus                   ********* ******* **          **************************  ** **************** **** ******* *  **********  * ***********

Teleogryllus_occipitalis    GCTCGCAGCCCCGT------------------------CAGCCGCTTCCTCCATCGTCGCAGAGCGCGCTCCTCCCGCGGCAGCGCTCCCTCGCGCAGCGGCCG----------------
Acheta_domesticus           GCTCTCGGCCCCGTCCGCCGCTGTGCTCCTCGTCGCAGCCGCCGCTTCCGCCGTCGTCGCAGAGCGCGCTCCTCTCGCCGCAGCGCTCCCTTGCGCAGCTGCCAGCTCCTCAACGCCCTC
Consensus                   **** * *******                        * ********* ** ******************** *** ************ ******* ***

Teleogryllus_occipitalis    ------------------------------------------------------------------------------------------------------------------------
Acheta_domesticus           GCGCGCTCGCTTCCGACCATCCTGCCCCTGCCCCGCTCCTGCCCCGCTCCTGCCCCGCCTCGCGCCCGCCACCGCCTTCCCCGCCCCTCACTGCGTTTGCCCTCCCCAGACAGCGCTCCA
Consensus

Teleogryllus_occipitalis    ------------------------------------------------------------------------------------------------------------------------
Acheta_domesticus           CCCCCTGCGCCGGGGACAGCCAGCCGCTCACGCTGGATGCGCTGATGGGACCCTCTGATGTGGAGACGACGTACTAGAAATGGTACACATCCAAAGGAGGATTTCCATGCACATGCATGC
Consensus

Teleogryllus_occipitalis    ------------------------------------------------------------------------------------------------------------------------
Acheta_domesticus           GCGTGAGAAATTTCAACACTTGTTAATGAAAAATGTTTCTATTTTTTGTTGATACGTTATTGCTTGTAATTGCCTACGATACGGAGTTCGATATACACTGCATAACGTATCTGGATTGAG
Consensus

Teleogryllus_occipitalis    ------------------------------------------------------------------------------------------------------------------------
Acheta_domesticus           AGGATCGTGGTTACGAGGAAAACAGTTGAATGGTGCAACATGTTTTCATACAGAGGGAATTATGGAAGGGAAATGGGAAAAGACAAACAGTCAAGTTAACATCTACTTAAGTAAATATGA
Consensus

Teleogryllus_occipitalis    -----------------------------------------------------------------------------------------------------------------------
Acheta_domesticus           ATTATTAAAAGTACATAAAAATAAAATCGTGTATGTGCCACGATCGTCAGAACCCTTAAACGGTCCTTGGTGTATAAATAACTATTATTGTCCCATATCACTTACGATGAACAATGTGA
Consensus
```

**Tocc.3G0015530.1_exon1**

```
Teleogryllus_occipitalis                      --------------------------------------------------------------------------------------------------------------
Acheta_domesticus                             ILDTYAVLEFFSRHYIHLCLFHDHFLKLSSLKHF*YTLNPRFQSK*RVFPLKIRRFSAQHYRTFSS*RQESKNSESGTQLSTFHTTNALERGDIQTCFIWLIMIQTLRALLGKRMRF*SP
Consensus

Teleogryllus_occipitalis                      --------------------------------------------------------------------------------------------------------------
Acheta_domesticus                             RNDHFRVIHDGNVC*RVACIAGCMPACFLSSPNSAIFLFYLFFPVFCCVLLFFRLHTLYAGVGVAQSLLKCSYTLTECCVMVSYK*YNILIFPLRIMYRENVYNGAAC*SARE*SFFGMI
Consensus

Teleogryllus_occipitalis                      --------------------------------------------------------------------------------------------------------------
Acheta_domesticus                             ITKAC*TFN**MSPWFPNWNSSLSTNHIPRIWEIHESSVTRVVIIPTNNRYLCTIIEKYTILNLTETPIFEFQNDSFQLRIFQSYDL**VSIQSRQVPEVTFSIAFIVSHCSLAARRGAA
Consensus

Teleogryllus_occipitalis                      --------------------------------------------------------------------------------------------------------------
Acheta_domesticus                             GRGAGPLARPDAGGRRSMHARGRPRTAAPRQPLLRRRRHGLGRARPAPAPALALAPRLRRILPSLPLGQRH*L*QALPWPVSLPFMPSRAALPFQLLVRLDSSSTPPPLLLFFPPAGPIS
Consensus

Teleogryllus_occipitalis                      --------------------------------------------------------------------------------------------------------------
Acheta_domesticus                             SSSVSAPFFSFLPLPSLVLSLLLLLPSLSAPSAAVLLVAAAASAVVAERAPLAAALPCAAASSSTPSRARFRPSCPCPAPAPLLPRLAPATAFPAPHCVCPPQTALHPLRRGQPAAHAGC
Consensus

Teleogryllus_occipitalis                      --------------------------------------------------------------------------------------------------------------
Acheta_domesticus                             ADGTL*CGDDVLEMVHIQRRISMHMHAREKFQHLLMKNVSIFC*YVIACNCLRYGVRYTLHNVSGLRGSWLRGKQLNGATCFHTEGIMEGKWEKTNSQVNIYLSKYELLKVHKNKIVYVP
Consensus

Teleogryllus_occipitalis                      ----------------------M---
Acheta_domesticus                             RSSEPLNGPWCINNYYCPISLTMNNV
Consensus                                                           *
```

**Tocc.3G0015530.1_exon2**

```
Teleogryllus_occipitalis      -----------------------------------------------------------------------------------------CGG----------------------------
Acheta_domesticus             FLILMQS*NFLVDITFTFVYFTIIF*SYLP*NIFNTL*ILDFKVNKEFFR*KFAVFRRNITELSLRKGKKVKIVKVEPSFRRFIQPMLWRGGIFKHVLFG*L*YRHFVPFWENE*DFNHP
Consensus                                                                                               **

Teleogryllus_occipitalis      ------------------------------------------------------------------------------------------------------------------------
Acheta_domesticus             GMIISGLFTTVTFVEEWLVLRVACPPVFSLPQILLYFYSIFSFPYSAVCFYFFVFIPFMQEWEWLSPFSSVVTHLRSAV*WFPINDTTSLYFLCELCIERTFTMGPPASQPVNKVFSV*L
Consensus

Teleogryllus_occipitalis      -------------------------------------------------------------------------------------------------------------------GTAWA
Acheta_domesticus             LPKHAELLIDECLHGFQIGIQASRPITFLEFGKSTSPPSLVS*LYLQIIGTSVRLLKNIQFSTSLKPQFLNFKMIVFN*EFFNRTICDEFLFNHGKFQK*HFRLRSL*ATVRWRRGAARR
Consensus                                                                                                                        * *

Teleogryllus_occipitalis      G----RAPATAPAPAAAPVPAPASAPP------------------PRPPPR------------------------PPPRP----------------------------------------
Acheta_domesticus             GGALDRSLAQMPAGADRCTPAAAPAPPRRGNRY*GGGGTAWAGPVPRPPPRSRSHPVCAGSFLPSRSVNAIDSNKPCPGPCLCHSCPAAPRCLFSSSSAWIRHRPRLRCFCFSLRPVRYH
Consensus                     *    *  *  ** *    ** * ***                  ******                        * * *

Teleogryllus_occipitalis      -------------------------PRARPRRPPP------PPAPAPAPAPAPAPAPAPAPAPAPA-PAPAPAPAPAPAPATATAPATAPATAPAPAPAPASAPATAPATAPATAPATAP
Acheta_domesticus             LRLFLPLFFHSSLFRRSFCHYCCFSPRSRPR--PPLCSSSQPPLPPSSQSALLSPQRSLAQLPAPQRPRALASDHPAPAP-------LLPRSCPASRPPPPSPPLTAFALPRQRSTPCAG
Consensus                                              ** ***  **      ** *         *    *  ***  *       *****         *   **  * * * * ** *         *

Teleogryllus_occipitalis      ATAPAT-----APATAPATAPA--------PEQAPRLRR-----------ILPSLPL-------------------------------------------GQRH*---------------
Acheta_domesticus             DSQPLTLDALMGPSDVETTY*KWYTSKGGFPCTCMRVRNFNTC**KMFLFFVDTLLLVIAYDTEFDIHCITYLD*EDRGYEENS*MVQHVFIQRELWKGNGKRQTVKLTST*VNMNY*KY
Consensus                        * *      *     *           *    * *                * *                                           * *

Teleogryllus_occipitalis      -----------------------------------
Acheta_domesticus             IKIKSCMCHDRQNP*TVLGV*ITIIVRYHLR*TM*
Consensus
```

**Tocc.4G0008930.1**

```
Teleogryllus_occipitalis    ACATCGGAGCTTGAAATCTGTAGGAGTGTGGGTGACCTGGCTC---------TGAGGAAATACCAAACATAGCGTA-----AGAGCACAGCGAAGT--GGAAGCCATGGGAAATGCGAAA
Acheta_domesticus           ACTTGGATGCTTGGAATCTGTAGGGGTGAGGGTGCCACGGCACAAAAAGGATTGAGGAAATACCAAAGATAACGT----------------CGAAC--AAGAGCCATGAGAAATGCGAAA
Gryllus_longicercus         AAATCGTAGCTGGCGATCTGTAGCGGTGTAAGTGCCACGGCACAAAAAGGATTGAGGAAATGCCGAAGATAACGCAGAACGAGAGCACAGCCGTGTGGGGGGGGGTGAGAAACGCGAAA
Consensus                   *  * *  *** *  ********  ***   *** *  *** *         ********* ** ** *** **                         *   ** **** ******

Teleogryllus_occipitalis    GAGAGATGTGA--ATGGAGGGGTGCGAATGGGCGCGACGCGAGGCAAGAGACAGACGGGAACGATGCCATTGCCCGCGGTCGCCGCGATCGGCTCTCTTCCACTCCTCGCCCCGCCCCCC
Acheta_domesticus           GAGAGATGTGA--ATGCTGCGGAGAGAGTGGGCGCGACGCGAGGCGAGAGGCAGGCGAGAACGATGCCATTCCCCGCGGCCGCTGCGATCGGCTCTCTTCCACTTCGCTCCTGCACCCCC
Gryllus_longicercus         GAGAGACGTGACGCGAGTGCGGTGAGAACGGG-----AGTGAGGCAAGCGACTGACGGGAACGATGCCATTCCGCGATGCCGCCGCGATCGGCTCTCTTCCCCTCCCCCCCGCCCACCCT
Consensus                   ****** ****       * ** * **  ***      * ***** ** * * * ** ************* * **  * *** **************** ** * * **     ***

Teleogryllus_occipitalis    TCGCACACGCACAC-TCCCATGCTTCTGTTCGGCACTGCGCGGTAAATGGGCGCGTGAGATACGAGCCACGTGTCGCAGATGTGCATTGTGGGCCGACAGCGACCTGCGACCGCGTCAAG
Acheta_domesticus           AAACACACATACACACTTCAGGCTTCTGTTCGGTAGCGTGCGGTGCGTGGACGCGTGAGATACGAGTCACGTGTCGCAGATGTGCATTGTGGGCCGACAGCGACCTGCGACCGCGTCAAG
Gryllus_longicercus         CCCCCCACGCAC---TTCCAGGCTTCTGTTCGGCAGCGCGCGGTGCGTGGGCGCGTGAGATACGAGCCACGTGTCGCAGATGTGCGTTGTGGACCGACAGCGACCTGCGACCGCGTCAAG
Consensus                      * ***  **      ** ************ *  * *****   *** *************** ***************** ****** *************************

Teleogryllus_occipitalis    CATTCGCCCGGCCGCCCCGGCCGCCCCCGCCGCCCCCGCCGCCCCCGCCGCCCCCGCCGCCCCCGCCGCCCCCGCCGCCCCCGCCGCCCCCGCCGCCCCCGCCGCCCCCGCCGCCCCCGC
Acheta_domesticus           CACTCGCCCGGACGCCCCAGCAGCCCCCGCC-TCCCAGCCGTCCCCGCCGCCCCCGCCGCACCTGCCATGCCCGCCGCTCCCACCGTTC---CGCCCCCCGCCGCCCCCGCC-CCCCCGC
Gryllus_longicercus         CACTCGCCCGGCCG----------------------------------------------------------------------------------------------------------
Consensus                   ** ******** **

Teleogryllus_occipitalis    CGCCCCCGCCGCCCCCGC-CGCCCCCGCCGCCCCCGCCCCCGCCGCCCCCGCCG-CCCCCGCCGCCCCCGCCG-CCCCCGCCGCCCCCGCCGCCCCGCCGCCCCGCCGCCCCCGCCGCCC
Acheta_domesticus           CAACCCTGCAGCCCCCACTCACCCCCGCCGCCCCCA-----GCCACTCTCGCCGCCCCCCACCACCCCCGCCGCCCCCCACCACCCCCGCCGCC------CACCACCACCTCCGCCGCCC
Gryllus_longicercus         ------------------------------------------------------CCCCCGCCGCCCCCGCCG-CCCCCACTGCCCCCGCCGCC-------------------------
Consensus                                                                         ***** ** ********* ***** *  ***********

Teleogryllus_occipitalis    CCGCCGCCCCCCCGCCCCCGCCGCCCCGCCGCCCCCGCCGCCCCCGCCGCCCCCGCCGCCCCCGCCGCCCCCGCCGCCCCCGCCGCCCCCGCCGCCCCCGCCGCCCCGTCGCCCCCGAAC
Acheta_domesticus           CTACCGCTC---------------------CCACCACCTCCCCCACCGCTCCCACCACCT------CCCCCACCGCTCCCACCACCTCACCACCCTGCCCGTCTCCGTCTCCCCCGAAC
Gryllus_longicercus         ------------------------------------------------------------------------------------------------------------------CCGGAAC
Consensus                                                                                                                                     ** ****

Teleogryllus_occipitalis    ATAAGGTGCGCGCGTGATGGACGCTCGTGAATGGCGGTGGGCAGACAGCCGCCGTCTGGGGCC-----------TGCGGTGATAAATATGCCTGGACAGCTGCGAGGGGATCTGGGAGTG
Acheta_domesticus           TCAAGGTGCGCGTGTGATGGACGCTCGTGAACGGCGACGGGCAGGCAGCTGCCGTCTGGGGTCGCCGCTGC-GGGGCGGTGATAAATATGCCTGGGCTGTGGCGAGGGGATCCGGGAGTG
Gryllus_longicercus         GTAAGGTGCGCCCGTGATGGACGCTCGTGAATGGCGGCGGGCAGGCAGCTGCCGTCTGGGCTCCCCACTGCTCGGGCGGTGATAGATATGCCTGATCTGTGGCGAGGGGATCCGGGTGTG
Consensus                     *********  ****************** ****  ****** **** **********  *            ********* *********  * *  *********** *** ***

Teleogryllus_occipitalis    TTGTAAACCGGTATTGCGCGGTCTTTGAGAGAGCGATAAGTTCGGATTTGAAAATGAAACATCATTTCTTACGACCAAAAGTATCGTTGTTTTTTGTTTTGTTTTTTTGATGTAATGATG
Acheta_domesticus           TTGTAAACCGGTATTGCGCGGTCTTTGACAGCGCGATAAGTTTGGATTTGAAAATGAAACACCTTTTCTTATGATCGAAAGCA-----------GCTTGTTCTTGTTGATACCATGATG
Gryllus_longicercus         TTGTAAACCGGTGTTGCTCGGTCTTTGAAAAAGCGATAAGTTAGGATTTGGAAACGAGTCACCTTTTCTTATGATCAAAAGGA-----------GCACTTTTGTTTTGTTACAATGATG
Consensus                   ************ **** ********** *  ********** ******* *** **  ** * ******* ** * **** *            *     *  * *** *   ******

Teleogryllus_occipitalis    CATTATAGATTTTGTCCAGAAGGGAATGTATTCTTGGTCTTTTGGCTTTCTAACTGGTTTAGGGCCATGAAAGCT---------------------------------------------
Acheta_domesticus           CATTATAGATTTCGTCCA-CATAGAATGTGTTCTTGGTGTCTTTGGTTTCTAACTGGTTT-GGGCCATAAAAGTTC----AAGCTCTCTGGACATTCTTGAACAGCTCCAACTGTGTGTG
Gryllus_longicercus         CATTATAGATTTCATCCA-GGTGGAATGTATTCTTGATCTCTTGGCTTTTCAAGTGGTTTGGGGCCATAAAAGTTCAATCAATCTCTCTGGATATTTGTGAACAGTTTGTGCTGTGTG--
Consensus                   ************  ****     ****** ****** * * ** * ***  ** ****** ******* **** *

Teleogryllus_occipitalis    ------------------------------------------------------------------------------------------------------------------------
Acheta_domesticus           TATTGGTCGGAACAAAAAAATTGAAGAGCATTAAATTTTCAACAGTCTCAAATTAAAATGAGCATTAAATTTCCCACAGGTGTTCAAAA-TGCACTTGTTGCTTAAATGCGTGTATCAAA
Gryllus_longicercus         --TTGTTAAGAACAAAAAAA-TAAGGAGC-------------------------GAATTGAGCATCAAATATCCCAAAGTGGTTGAAAATTCCATATTTTGCCTAAATACGCATATTAGA
Consensus

Teleogryllus_occipitalis    ------------------------------------------------------------------------------------------------------------------------
Acheta_domesticus           AACTGTTAAGTAAACCGAAACAG-CAAATAAGTAAATATCACTGGTCTTCTGTATAATCAAATCAAATAAATTATTTATTGATATTTATAAAATTTAAAATACTAGGAAAACAAGTAAAC
Gryllus_longicercus         AACTGTTCAGAAAACAGTAACAGATAAATAAGTAAATATCAATAATCTTCTCTATAATCAAGTAAATTAATTATCTTTTTAATGTTTA--------AAACTATTAGCAAAATAAGTAACC
Consensus

Teleogryllus_occipitalis    ------------------------------------------------------------------------------------------------------------------------
Acheta_domesticus           GGACCACATTCAAATGTTAATTCATCTGTTAATTTTTAAACCTTTACATTTAGA-----------------AATGTATGTTCTCT-------TAAAATTATCATAAAC------------
Gryllus_longicercus         GG-CCACATTCAAATG-TAATTCGTATTACAATTTTTATACCTTTCCTCTTATAATAAACATAATCTTAATAGGGTTTGTTTTTCGAAAGTGTAATACTATTATTAGTAAGCTTTAGACT
Consensus

Teleogryllus_occipitalis    ------------------------------
Acheta_domesticus           ------------------------------
Gryllus_longicercus         TACGGTTCACGTTTGCAAATTTTCTATGCT
Consensus
```

**Tocc.4G0008930.1_exon1**

```
Teleogryllus_occipitalis                ------------------------------------------------------------------------------------------------------------------------
Acheta_domesticus                       KTLKMQNKYILWFDF*LIEQF*F*SFPRISVRILELRNN-------FNFRIIY*LFLLIT------I*FS*HTKSIT-----ASYFTIVLPE*LNCE*LQRICPIAKIRMSLRMKL*KSR
Gryllus_longicercus                     HTESIRPYFWLKLVFRRTKE-------EITKHI*HSNNKTKNT*QIL*KKCVFLFDL*LTQKIHNSIFFYYYCT*IT*L*MIAAY----LP---NCKD-PNIAEMDDLRWRKMTNDNTKG
Consensus                               

Teleogryllus_occipitalis                ------------------------------------------------------------------------------------------------------------------------
Acheta_domesticus                       QKR--DEMELTMT-LNKDSHTIQESIEVVLSESAKNLDAWNL*G*GCHGTKRIEEIPKITSNKSH------EKCEREM*MLRREWARR--EARGRRERCHSPRPLRSALFHFAPA--PPN
Gryllus_longicercus                     QNRVNDDFKKGVTLLNRGS----------MWYCQKKIVAGDL*RCKCHGTKRIEEMPKITQNESTAVWGGGEKRERET---*RECGENGSEASD*RERCHSAMPPRSALFPSPPPTLPPR
Consensus                               

Teleogryllus_occipitalis                -------------------------MCIVGRQRPATASSIRPAAPAAPAAPAAPAAPAAPAAPAAPAAPAAPAAPAAPAAPAAPAAPAAPAPAAPAAPAAPAAPAAPAAPPPRRPRRPRR
Acheta_domesticus                       THTHFRLLFGSVRCVDA*DTSHVSQMCIVGRQRPATASSTRPDAPAAPA-----SQPSPPPPPHLPCPPLPPFRPPPPPPPRQPCSPHSP-PPPPATLAAPHHPRRPPPPPP--------
Gryllus_longicercus                     TS---RLLFGSARCVGA*DTSHVSQMCVVDRQRPATASSTRPAAPAAPA---------------------------------APTAPAAP-------------------------ERKVR
Consensus                                                        ** * ********* ** ******                                  *  *  *

Teleogryllus_occipitalis                P-----------PAPAAPPPPPPPPPPPPPPPPPPPPPPPPPPPR---------------------------------------------------------------------------
Acheta_domesticus                       PTTT-------SAAPTAPTTSPTAPTTSPTAPTTSPPCPSPSPPNSRCACDGRS*TATGRQLPSGVAAAGR**ICLGCGEGIRECCKPVLRGL*QRDKFGFENETPFLMIESSLFLLIP*
Gryllus_longicercus                     P*WTLVNGGGQAAAVWAP----------------------------HCSGGDRYA*SVAR----------------GSGCVVNRCCSVFEKAI------S*DLETSHLFL*SKGALLF--
Consensus                               *            *  **

Teleogryllus_occipitalis                ---------------------------------------------------------------------------------------RPRT*----------------------------
Acheta_domesticus                       CIIDFVHIECVLGVF--GF*LVW--AIKVQALWTFLNSSNCVCIGRNKKIEEH*IFNSLKLK*ALNFPQVFKMHLL-LKCVYQKLLSKPKQQISKYHWSSV*SNQINYLLIFIKFKILGK
Gryllus_longicercus                     CYNDAL*ISSRWNVFLISWLFKWFGAIKVQSISLDICEQFVLCVVKNKKIRS---------ELSIKYPKVVENSIFCLNTHIRNCSENSNR*ISKYQ*SSL*SSK---LIIFLMFKTISK
Consensus                               

Teleogryllus_occipitalis                -------------------------------------------------------
Acheta_domesticus                       QVNGPHSNVNSSVNF*TF---------------------TFR---NVCSLKIIIN
Gryllus_longicercus                     ISNRPHSNVIRITIFIPFLL**T*S**GLFFESVILLLVSFRLTVHVCKFSM---
Consensus
```

**Tocc.4G0010540.1**

```
Teleogryllus_occipitalis  CAAAAAATTTAAA-----------------------------------------------TAAAATTGAAAACGTTCATTCTTTGTGAGAAAATTAGACGAACTGAAGAACAAAAACAAA
Acheta_domesticus         CAAAAACTGTTCAATTGTTTTTCTTTAAATTTTGTGTTCCAATAAAGCGAAATTAACGACTAATATTGCACGAGTACAATCGTCATTAAAACTTCAAAATCACTAAAATTCCAAATTTTT
Consensus                 ****** * ** **                                              *** **** *   ** ** ** *  * * **  * * *   *** **   * ***

Teleogryllus_occipitalis  GTAACCCTTCTGTCAAGGTTTTGATAAAGGAA----------TATAAA------------------------------------------------------------------------
Acheta_domesticus         TCCACCCTCCTACAACGGTATTGCTAATATAAGAGGTTTCTTTATAAATCAAGCTGTTCCATTATATTTGAGCTGTAAATCCATTAGTATTTCCATACTATTTTATCTATTTGTGATTTT
Consensus                    ***** **   * *** *** ***   **          ******

Teleogryllus_occipitalis  ----------------------------------------------GTTAGGAAAATATAATGTATGACACCTTAAAAACAAATA-----------------------------------
Acheta_domesticus         TACATAATTTTCAGTTACATCGTTTGTGTTTTTAATGTGCTGAAGTGGTAGGAGGATGGAAGGTGCAATATTATAATCTTAGCTAATTTCTTGTAATATTGGAGGAGTTAAGTCTTACTT
Consensus                                                               * *****  **  ** **   * *   ***    *  **

Teleogryllus_occipitalis  ---CAAAATCAAGACTTCATAGTTAAAGGACTTTTTTCGATA------------------------------------------------------------------GGGGGTTGCTAC
Acheta_domesticus         TAGCACAACTAGGTTTTCTGTGTGACAGCACTCTCGCAAATAAATGATCTGAATTCTTTGTAAGCATTTTGTATTTCATTCGTATAATTTCGCTCTCTTCCCATTTTCGGGCATTACTTC
Consensus                    ** **  * *  ***   ** * ** *** *     ***                                                                  ***  ** ** *

Teleogryllus_occipitalis  AA------------------------TATTGACGAGGACGTAA----GCCAATATCCGAAGCCTAAATGA--------------TCAGATGAATTAAT-----------CTGTTCCAA--
Acheta_domesticus         AACTTTTTGGGGAAGAAATGTAATTTTGCTGAGAGGTACATAAAACCGGTGAAATTATCGACATAAATGAAAGTGCTGTTTTTCTTAGAAAAATTAGTAAGGATGTCAATAATTTCAATG
Consensus                 **                        *  ***   * ** ***    *   * **      * *******              * ***  ***** *              ** ***

Teleogryllus_occipitalis  -------CAAATATTGAAAATATT--------ATTTCTTTTACTTTTTA----------------------------TGCAGTTCTTTGTTTCAT-------------------------
Acheta_domesticus         AAATAACCAAATACAAAAAATAGTTTTCTCAAATTCTGTTTACTTGTAAAAACAATGTTAAACTGTACTTGACAACTTGTCGTTTTTCATTTTATGCCTCAGTCTCACAACTGGGTCTAA
Consensus                        ******   ****** *        ***   ******* * *                            **  *** **  *** **

Teleogryllus_occipitalis  -----------TAGACTTCGACTGTACGTCATTTTTTCGTTT------------------------------------------------------------------------------
Acheta_domesticus         TGTCACTACTCCATCTTTCGAACGAACAACAACTTCCCATTTGGAGGGTCATCTCAACCACTGCCGTCACCGCCGCTTCTTCGCACTTTCCTAGAGCAGCCATCCATCTTCGGGCCGCAC
Consensus                             *   *****  * **  **  **  * ***

Teleogryllus_occipitalis  ----------CTTCCAGGACTTCG------------------------------------------------------------------------------------------------
Acheta_domesticus         GTTTCTCGCGTTTCCAAAGCTCCGGAGGCAAACTTGTCGTGAGAGCGCGCTAAAATGTATCGCCAGAGCTTCTCGCAACATCCTCATGTAAGCTTACAAATTTAAAAAACAAACAAAACG
Consensus                            *****   ** **

Teleogryllus_occipitalis  ------------------------------------------------------------------------------------------------------------------------
Acheta_domesticus         CGAGAGCATAGAATAAAGTGAAATTGGATAGCACATTTCGATTCCGAGAATCGAAGCAGCCGACAATGTTAAGCGAATTCAGTACAATGGAGAACTTGTGATGTTAGTGTGTTTACATGG
Consensus

Teleogryllus_occipitalis  ------------------------------------------------------------------------------------------------------------------------
Acheta_domesticus         AATATAGAGTGATATAGTGTGTTTGAAACTTTGAAGGGTTTTTCGTTGCACCGAAGTGTTCTTCTTGCCTTCTGAAGGGGAGACTGCTCTTCTTATTTGGATCGTTGAAAGTGTTTGTGA
Consensus

Teleogryllus_occipitalis  -----------------------------------------------------TTTCTTTGAAATTAAGAAAGTACTCATACAATCTATTGCATCTTTTAACG------ATTGCGTTTTC
Acheta_domesticus         ATGTAGGAAAACATATGTGACATGCTAAAAGTATATTTCACATACATTTCGCATAATTTTGGGATAAAAAATGCATCATAATTGCCGGTTATATCTTTTTACGTGGCTTATTTCTTCCTC
Consensus                                                                      *   ****  ** ** ** * *     *    *  **  ******* ***      *** * *  **

Teleogryllus_occipitalis  TG-----AATTTACA---------------------------------GTTTTTATTGC-------------------------------------AAACCTGAATAATTCTGAAAC---
Acheta_domesticus         TATCCATAATTAACACAATTCAATTCATTGTTGACAAAATATTGGGGGGTCTTTAATGCTGCTAGCTTTAATTTTATCAACAAGAAAAATTACTTGAAAAATGAAAATCTTCGAAACAAT
Consensus                 *      **** ***                                 ** **** ***                                     ***  **** *  *  *****

Teleogryllus_occipitalis  -----CTATACAATTCGAAAGAAGAT-------GTTGGACCCAAAGGATTTCATCTTAGACAGGTGTTTGTGTGTAATAAATAGATAACAATAGAGCAGAGTAGTTACTGCGTTGGTGTG
Acheta_domesticus         TTAATATTTGTAGTATTAAAATAGATCACAGTAGCTTACCTGTATCGTTTTCTTTCAACATAAGAAATTGGGTGCACATTTTTGAGAATTATGAGACTTCTTAATTATTGCAATGGAATG
Consensus                       * *  * *   ***  ****       * *   *   *  * **** *   * * * *   *** *** *     * ** **  **    *    ** *** ***  ***  **

Teleogryllus_occipitalis  CAATAGGTAGAGAGATATTGTAATAGAGAGAAACTAATCGTCGTGTTGGAAGTTCAAGGTCCAT-AAAAATCCAAATTAAAATAAACGAAGAAAGTAATTTAAATAAATAACTAATTGAG
Acheta_domesticus         CAACAG-----GAGACATTGCAATAGAA------TACTGGTCGCGTTGGAGAT--AACGTCTTTGAGAAATCCAAGTTA----GAACGAAGAAAATAATTTAAATAAATAACTAATTGAA
Consensus                 *** **     **** **** ******      ** * **** ******  *  ** ***  * * ******** ***     ********** **********************

Teleogryllus_occipitalis  CAGAAGTCAGAGTTCTTCGTCCCTCCCTTTTCTTGCAATGCAAGAAGCGCATAGCAGCAGTCTCTGCCAGCTGAACCACCCTTTAAAAGTTCGAAATCCCATAAAAAAGTCGAAGTATCA
Acheta_domesticus         CAGCAGTCAGGGTACTTTGTCCC-----TTTTTTATAATACAAGATGCGCACAGGAGCGTCCTCTGCCAGCTGAACCACTCCTTAAAAGTTCGAAATACCATAAAAACGTCGAAGTATCA
Consensus                 *** ****** ** *** *****     *** **  *** ***** ***** ** ***   ***************** * *************** ********* ***********

Teleogryllus_occipitalis  GAACTGTCATTGAATATTCCGCGCGCAGTCGTCAGAAAA-AACGACCACTTCTCCGATCGTTTGACGCTTTATTAAAAGAGCGGCCGCCCGTGCAGAACGCGCTGCC-------------
Acheta_domesticus         GAACCGTCATTGAATATTCCGCGCGCAGTCGTCAGAAAAGAACGGCTACTTCTCCGATCGTTTGACGCTTTATTAAAAGAGCGGTCGCGAGCAGAGAACGTGCCGCCCTGGCGGCGTTAA
Consensus                 **** ********************************** **** * ************************************* ***  *   ****** ** ***

Teleogryllus_occipitalis  GCACAGCCCGCGACGCGCGGCGGCGCTGCGATCGCCCGTAACCTCGCGCCGGGCGCGCCCCGCCCCGCCCTCGCTCTCGTTTATGGCGGCGCGGGGAGCTTTCACAGAGTTATGGAAATG
Acheta_domesticus         GCGCAGCCCGCGACGC----------------------------------------------CCCCGCCCCTGC-CCCGTTTATGGGAGCGCGGAGAGCTTTCGCAGAGTTATGGAAATG
Consensus                 ** *************                                              ********  ** * *********  ****** ******** ***************

Teleogryllus_occipitalis  CGTTTGAGGATTGTAGCCCCGCGTCAAGTGGCTGCGAGGCAAAGGTGGTGGCTGTGTGCGC----------CGGAAGGGAG---------------------------------GAGGCG
Acheta_domesticus         CGTTTGAGGATTGTAGTCCCGCGCCAAGTGGCTGC---------GTGGTGGCTGCGTGCGCGGGGTGTGAACGGAAGGGGGCCGGTGGCGGGGCCAGTCACGCGGGCGCGGGCAGGAGCG
Consensus                 **************** ****** ***********         ********** ******          ******** *                                 *  ***

Teleogryllus_occipitalis  CGGGAGGAGGAGGAGGAGGA---------------------------------------------------------------------------------------------GGAGGCG
Acheta_domesticus         GGGGTGAAGGTGGAGGCGAAGTTATGGGCGGAGGCGGAGACGGGTTATGCAGCGGGGGCGGCAGCGGGGGCGGCAGCGGGGGCGGCAGCGGGGGCGGCAGCGGGGGCGGCAGCGGGGGCG
Consensus                  *** * *** ***** * *                                                                                            ** ****

Teleogryllus_occipitalis  CGGGAGGAGGAGGAGGAGGAGGAGGAGGAGCGGGAGGAGGAGGAGGAGGAGGAGGAGGAGGC-GCGGGAGGAGGAGGAGGAGGAGGAGGAGGAGGAGGAGCGGGGA-------GGAGGAG
Acheta_domesticus         GCAGCGGGGGCGGCAGCGGGGGCGGCAGCGGGGGCGGCAGCGGGGGCGGCAGCGGGGGCGGCAGCGGGGGCGGCAGCGGGGGCGGCAGCGGGGGCGGCAGCGGGGGCGGCAGCGGGGGCG
Consensus                    * ** ** **  * ** ** **  * * *** **  * ** ** **  * ** ** *** ***** *  * **  ** *  * **  ** *  * *******        ** ** *

Teleogryllus_occipitalis  GAGGAGGAGGAGGAGGAGGAGGAGTGGGAGGAGGAGGAGGAGGAGGAGGAGGAGGAGGAGGAGGAGGAGGGGGAGGAGGAGGAGGAGGGGG--------------------------
Acheta_domesticus         GCAGCGGGGGCGGCAGCGGGGGCGGCAGCGGGGGCGGCAGCGGGGGCGGCAGCGGGGGCGGCAGCGGGGGCGGCAGCGGGGGCGGCAGCGGGGGCGGCAGCTGAGGCGGCAGCGGGGCGG
Consensus                 *  * ** ** **  * ** ** *   * ** ** **  * ** ** **  * ** ** **  * ***** **  * ** ** **  * *****

Teleogryllus_occipitalis  -------------------------------------------------------------GGAGGAGGAGGAGGAGGAGGAGGAGGGGGGGGAGGAGGAGGAGGAGGAGGCGGGAGGAG
Acheta_domesticus         CAGCGGGGGCGGCAGCGGGGGCGGCAGCGGGGGCGGCAGCGGGGGCGGCAGCGGGGGCGGCAGCGGGGGCGGCAGCGGGGGCGGCAGCGGGGGCGGCAGCGGGGGCGGCAGCGGGTGGTG
Consensus                                                                  * ** ** **  * ** ** **  * ***** **  * ** ** **  ***** ** *

Teleogryllus_occipitalis  GAGGAGGAGGAGGAGGAGGAGGAGGCGCCAGGAGGAGGAGGAGGAGGAGGAGGAGAGGAGGAGGC----------------------------------GCGGAGGAGGAGGAGGAGGTG
Acheta_domesticus         GCAGTGGGGGCGGCAGCGGGGGCGGCAGCGGGGGCGGCAGCGGGGGCGGCAGCGGGGGCGGCAGCGGGGGCGGCAGCGGGGGCGGCAGCGGGGGCGGCAGCGGGGGCGGCAGCGGGGGTG
Consensus                 *  * ** ** **  * ** **** ***  ** *  * **  ** *  * **  * ** **  **                                  **** ** **  * ** ****

Teleogryllus_occipitalis  GGAGGAGG-------------------------------------------GCGCGGGAGGAGGAGGAGGAGGTGGGAGGAGGAGGCGCGGGAG-----------------GAGGAGGAG
Acheta_domesticus         GCAGTGGGGGCGGCAGCGGGGGCGGCAGTGGGGGCGGCAGCGGGGGCGGCAGCGGGGGAGGCAGCGGGGGCGGCAGCGGGGGCGGCAGCGGGGGCGGCAGCGGGGGCGGCAGCGGGGGCG
Consensus                 * **  **                                           *** ******  * ** ** **  *  ** *  *  ***** *                 * ** ** *

Teleogryllus_occipitalis  GAGGAGGAGGCGCGAGGAGGAGGAGGAGGAGGAGGAGGAGGCGAGGAGGAGGAGGAGGAGGAGGAGGAGGAGGAGAAGGAGGAGGAGGAGGAGGAGGAGGAGGAGGAGGAGGAGGAGGAG
Acheta_domesticus         GCAGCGGGGGCGGCAGCGGGGGCGGCAGCGGGGGCGGCAGCGGGGGCGGCAGCGGGGGCGGCAGCGGGGGCGGCAGCGGGGGCGGCAGCGGGGGCGGCAGCGGGGGCGGCAGCGGGGGTG
Consensus                 *  * ** ****  **  ** *  * **  ** *  * **  * ** **  * ** ** **  * ** ** **    ** ** **  * ** ** **  * ** ** **  * ** ** *

Teleogryllus_occipitalis  GAGGAGGAGGAGGATGAGGAGGAGGAGGAGGAGGATGAGGAGGAGGAGGAGGAGGAGGAGGAGGAGGAGGAGGAGGAGAGG--AGGAGGTGGGATGAGGACGTGAGGAGGAGGAGGA
Acheta_domesticus         GCAGCGGGGGCGGCAGCGGGGGCGGCAGCGGGGGCGGCAGCGGGGGCGGCAGCGGGGGCGGCAGCGGGGGCGGCAGCGGGGGTGGCAGCGGGGGGCGGCAGCGGGGGC
Consensus                 *  * ** ** **  * ** ** **  * ** ** **  * ** ** **  * ** ** **  * ** ** **  * * **   * **  *** **    ** * ** **  * ** **

Teleogryllus_occipitalis  GGCGGGGGAGGAGGAGGAGGAGGAG-----------------------------------------------------------------------------------------------
Acheta_domesticus         GGCAGCGGGGGCGGCAGCGGGGGCGGCAGCGGGGGCGGCAGCGGGGGCGGCAGCGGGGCGGCAGCGGGGGCGGCAGCGGGGGTGGCAGTGGGGGCGGCAGTGGGGGCGGCAGTGGGGGCG
Consensus                 *** * ** ** **  * ** ** *

Teleogryllus_occipitalis  -----------------GAGGAGGAGGAGGAGGAGGAGGAGGAGGAGGAGGAGGAGGAGGAGGAGGAGGAGATGAAAGGAGGAGGAGGAGGAGGAGGAGGAGGAGGAGGAGGAGGAGGAG
Acheta_domesticus         GCAGCGGGGGCGGCAGCGGGGAGGCAGCGGGGGAGGCAGCGGGGGCGGTAGCGGGGGTGGCAGTGGGGGCGGCAGCGGGGGGAGGAGGCGGGAGTGGTGGCGGGGGCGGAGGCGGGGCCA
Consensus                                  * *****  * ** *****  * ** ** **  * ** ** **  * ** ** *      ** ******** **  * ** ** ** ** ***** ** *

Teleogryllus_occipitalis  GAGGAGGAGG-------------------------AGGAGGAGGGGGGAGGAGGAGGAGGAGGATGGAGGAGGAGGAGGAGGAGG--------------------------------
Acheta_domesticus         GAGGAGGAGGTGCTGCGGAGAGCGGAAGTGGGGGCGGGAGCGGAGGCGAAGGCGGGGACGGGGACGTGGGTGGAGGTGGGGGCGGGGGGCGGCAGCTCGAGCGGTGGCGGAAGCGGGGCC
Consensus                 **********                         ****  * ** ** *  * *** * *** *  ** ***** ** ** ** **

Teleogryllus_occipitalis  -AGGAGGAG---------------------------------------------------------------------------------------------------------------
Acheta_domesticus         AGGGGGGAGGTGCTGCGGAGGGCGGAGGCGGGTGTCAGAACGCCGGCAGCAACGATGTTCGGTTGAAAGGTGAGAATGGCTGTGACGGCGGCGGGGAGGCGTCGGCGGCATCGGTGAAAG
Consensus                   ** ****

Teleogryllus_occipitalis  ------------------------------------------------------------------------------------------------------------------------
Acheta_domesticus         CCAAGCGAGGGCGACTGACTGTGAAGGTACGCTGGCAAAGGTGAATGAATAAAAGGATAAGGCAACAAAGGAATTGCTTCTGCACACGCGTGGCTGGATGCTGACGGCAGTGACGTTAGT
Consensus

Teleogryllus_occipitalis  ---------------------------------------------------------
Acheta_domesticus         GACGATTACTGTAATTTGACAAGTGCGTCAGCGTTATCGATGCTGTCGTTAGAGGAC
Consensus
```

**Tocc.4G0010540.1_exon1**

```
Teleogryllus_occipitalis        ------------------------------------------------------------------------------------------------------------------------
Acheta_domesticus               FQTE*FSLNF**NCETEKSKILKNCSIVFL*ILCSNKAKLTTNIARVQSSLKLQNH*NSKFFPPSYNGIANIRGFFINQAVPLYLSCKSISISILFYLFVIFT*FSVTSFVFLMC*SGRR
Consensus                       

Teleogryllus_occipitalis        ------------------------------------------------------------------------------------------------------------------------
Acheta_domesticus               MEGAIL*S*LISCNIGGVKSYFSTTRFSV*QHSRK*MI*ILCKHFVFHSYNFALFPFSGITSTFWGRNVILLRGT*NR*NYRHK*KCCFS*KN**GCQ*FQ*NNQIQKIVFSNSVYL*KQ
Consensus                       

Teleogryllus_occipitalis        -------MQFF-------------------------------------------------------------------------------------------------------------
Acheta_domesticus               C*TVLDNLSFFILCLSLTTGSNVTTPSFERTTTSHLEGHLNHCRHRRFFALS*SSHPSSGRTFLAFPKLRRQTCRESALKCIARASRNILM*AYKFKKQTKRESIE*SEIG*HISIPRIE
Consensus                                **

Teleogryllus_occipitalis        ------------------------------------------------------------------------------------------------------------------------
Acheta_domesticus               AADNVKRIQYNGELVMLVCLHGI*SDIVCLKL*RVFRCTEVFFLPSEGETALLIWIVESVCECRKTYVTC*KYISHTFRIILG*KMHHNCRLYLFTWLISSSIHN*HNSIHC*QNIGGSL
Consensus                       

Teleogryllus_occipitalis        ------------------------------------------------------------------------------------------------------------------------
Acheta_domesticus               MLLALILSTRKIT*KMKIFETI*YL*Y*NRSQ*LTCIVFFQHKKLGAHF*EL*DFLIIAMECNRRHCNRILVALEITSLRNPS*NEENNLNK*LIEQQSGYFVPFL*YKMRTGASSAS*T
Consensus                       

Teleogryllus_occipitalis        ------------------------------------------------------------------------------------------------------------------------
Acheta_domesticus               TP*KFEIP*KRRSIRTVIEYSARSRQKRTATSPIV*RFIKRAVASRERAALAALSAARDAPAPAPFMGARRAFAELWKCV*GL*SRAKWLRGGCVRGV*TEGGRWRGQSRGRGQERG*RW
Consensus                       

Teleogryllus_occipitalis        ------------------------------------------------------------------------------------------------------------------------
Acheta_domesticus               RRSYGRRRRRVMQRGRQRGRQRGRQRGRQRGRQRGRQRGRQRGRQRGRQRGRQRGRQRGRQRGRQRGRQRGRQRGRQRGRQRGRQRGRQRGRQRGRQRGRQRGRQRGRQLRRQRGGSGGG
Consensus                       

Teleogryllus_occipitalis        ------------------------------------------------------------------------------------------------------------------------
Acheta_domesticus               SGGGSGGGSGGGSGGGSGGGSGGGSGGGSGGGSGWWQWGRQRGRQRGRQRGRQRGRQRGRQRGRQRGRQRGRQRGWQWGRQRGRQWGRQRGRQRGRQRGRQRGRQRGRQRGRQRGRQRGR
Consensus                       

Teleogryllus_occipitalis        ------------------------------------------------------------------------------------------------------------------------
Acheta_domesticus               QRGRQRGRQRGRQRGRQRGRQRGRQRGRQRGRQRGRQRGWQRGRQRGRQRGRQRGRQRGRQRGRQRGRQRGWQRGAAAGAAAGAAAGAAAGAAAGAAAGRQRGRQRGWQWGRQWGRQWGRQRGR
Consensus                       

Teleogryllus_occipitalis        ------------------------------------------------------------------------------------------------------------------------
Acheta_domesticus               QRGGSGGGSGGGSGGGSGGGSGGRRREWWRGRRRGQRRRCCGERKWGRERRRRRGRGRGWRWGRGAAARAVAEAGPGGRCCGGRRRVSERRQQRCSVER*EWL*RRRGGVGGIGESQARA
Consensus                       

Teleogryllus_occipitalis        ------------------------------------------------------
Acheta_domesticus               TDCEGTLAKVNE*KDKATKELLLHTRGWMLTAVTLVTITVI*QVRQRYRCCR*R
Consensus                       
```

**Tocc.4G0010540.1_exon2**

```
Teleogryllus_occipitalis        ------------------------------------------------------------------------------------------------------------------------
Acheta_domesticus               FKLSSFRLIFSEIVKLKRVRFSKTVQLFFFKFCVPIKRN*RLILHEYNRH*NFKITKIPNFFHPPTTVLLI*EVSL*IKLFHYI*AVNPLVFPYYFIYL*FLHNFQLHRLCF*CAEVVGG
Consensus

Teleogryllus_occipitalis        ------------------------------------------------------------------------------------------------------------------------
Acheta_domesticus               WKVQYYNLS*FLVILEELSLTLAQLGFLCDSTLANK*SEFFVSILYFIRIISLSSHFRALLQLFGEEM*FC*EVHKTGEIIDINESAVFLRKISKDVNNFNEITKYKK*FSQILFTCKNN
Consensus

Teleogryllus_occipitalis        ------------------------------------------------------------------------------------------------------------------------
Acheta_domesticus               VKLYLTTCRFSFYASVSQLGLMSLLHLSNEQQLPIWRVISTTAVTAASSHFPRAAIHLRAARFSRFQSSGGKLVVRAR*NVSPELLATSSCKLTNLKNKQNARA*NKVKLDSTFRFRESK
Consensus

Teleogryllus_occipitalis        ------------------------------------------------------------------------------------------------------------------------
Acheta_domesticus               QPTMLSEFSTMENL*C*CVYMEYRVI*CV*NFEGFFVAPKCSSCLLKGRLLFLFGSLKVFVNVGKHM*HAKSIFHIHFA*FWDKKCIIIAGYIFLRGLFLPLSIINTIQFIVDKILGGL*
Consensus

Teleogryllus_occipitalis        ------------------------------------------------------------------------------------------------------------------------
Acheta_domesticus               CC*L*FYQQEKLLEK*KSSKQFNICSIKIDHSSLPVSFSFNIRNWVHIFENYETS*LLQWNATGDIAIEYWSRWR*RL*EIQVRTKKII*INN*LNSSQGTLSLFYNTRCAQERPLPAEP
Consensus

Teleogryllus_occipitalis        ------------------------------------------------------------------------------------------------------------------------
Acheta_domesticus               LLKSSKYHKNVEVSEPSLNIPRAVVRKERLLLRSFDALLKERSRAENVPPWRR*AQPATPPPLPRLWERGELSQSYGNAFEDCSPAPSGCVVAACAGCERKGAGGGASHAGAGRSGGEGG
Consensus

Teleogryllus_occipitalis        ------------------------------------------------------------------------------------------------------------------------
Acheta_domesticus               GEVMGGGGDGLCSGGGSGGGSGGGSGGGSGGGSGGGSGGGSGGGSGGGSGGGSGGGSGGGSGGGSGGGSGGGSGGGSGGGSGGGSGGGSGGGSGGGSGGGSGGGSGGGS*GGSGAAAGAA
Consensus

Teleogryllus_occipitalis        ------------------------------------------------------------------------------------------------------------------------
Acheta_domesticus               AGAAAGAAAGAAAGAAAGAAAGAAAGAAAGAAAGGGSGGGSGGGSGGGSGGGSGGGSGGGSGGGSGGGSGGGSGGGSGGGSGGGSGGGSGGGSGGGSGGGSGGGSGGGSGGGSGGGSGGG
Consensus

Teleogryllus_occipitalis        --------------------------------------------------------VGGGGAGGGGGGGGARRR-------------RRRRRRRGGGGGGGGGGEGGGGGG---------
Acheta_domesticus               SGGGSGGGSGGGSGGGSGGGSGGGSGGGSGGGSGGGSGGGSGGGSGGGSGGGSGGGSGGGSGGGSGGGSGGRQRGRQRGRQRGRQRGRQRGRQRGGSGGGSGGGSGGGSGGGSGGGSGGG
Consensus                                                                                ***  ** *** * * *             * * * *** *** *** *** **

Teleogryllus_occipitalis        ---------------------GGGGGGGGGGGGG*-------------------------------------------------------------------------------------
Acheta_domesticus               SGEAAGEAAGAVAGVAVGAAAGGGGGSGGGGGGGARGGGAAESGSGGGSGGEGGDGDVGGGGGGGRQLERWRKRGQGGGAAEGGGGCQNAGSNDVRLKGENGCDGGGEASAASVKAKRGR
Consensus                                            ***** *******

Teleogryllus_occipitalis        ------------------------------------------------------
Acheta_domesticus               LTVKVRWQR*MNKRIRQQRNCFCTRVAGC*RQ*R**RLL*FDKCVSVIDAVVRG
Consensus
```

**Tocc.4G0011610.1**

```
Teleogryllus_occipitalis        CGGCGTGCTCACCAGATAGCATTACTGT--------------------------------------------------------------------------------------------
Acheta_domesticus               TTTAGTG------AAGTGCAATTATAGTGACAAGCGAATAGCAGGTAA-----------------------------------------AGAAATACTTGGGCATTCA------------
Gryllus_bimaculatus             TGTTATGTTCA--AAATACTCTTACAGATA-----------------------------------------------------------AGAGATGTAAAACTGTGTGAAATTAC-----
Gryllus_longicercus             TGTTGTGGGATCCAGATGCGATTACATTTAGAGGTGAATGAGAAGCGAGTTTGGAAGAGTAAAATTGCTAAGCGAAAGACAGAATACGCAAGGACGTTGAATCATTTAAAATGGCAGGTA
Consensus                            **      *  *    ***

Teleogryllus_occipitalis        --------------------------------------------------------------------------GCGT------------------CTTGGCTGTCGTCTTCTGTGAATA
Acheta_domesticus               ----TGCCACCA-------------------------------------------AGGCCAATCTCTCAGGCA---------------------------GTTTCTGTCTTCCATTTCCA
Gryllus_bimaculatus             ----TGCTGATA-------------------------------------TGTGAAAAAGAAACACATCTAACTTGCGTAACTATACATATTATTTCCCTGAATGTTTTCTTCCTGGTTTT
Gryllus_longicercus             GGGTTGCCAACAACAAGGATGGAAAACGAGGGAGCTGCGATTGGAGACGGTCGAGAAGCGAACATGTGGAGCAGGTGT------------------CTCCGCTGGTGTTGGCAGATTGCA
Consensus                                                                                                             *    *   *

Teleogryllus_occipitalis        AGATAGTATGCA-------------------------------------------TTGTGTTAGAATATTGTTTAATTA-------------CAAACACTTGAAGGAAGCGTGGCAACGT
Acheta_domesticus               CTCCAACATAAAGCTCACGACGGCCAGAGTTG-----------------------CCGCTTTAGTGTGCTCTGT-----------TATCTCCCGATCCTTCATTTCCAGTGGCGCAACAG
Gryllus_bimaculatus             CAAATGTACGTA----------------------TAAA-----------------TTTTCTGAAGATATTTTTT---------------TTATGTATTCCACTAATCAACAAAAAAATAA
Gryllus_longicercus             GGCGGGTATTAAGGCCGCGTGCGCCGGAGGAGGGTAGAGGGCGAGTGCAGGAACTCCGCCTCGGGGAGAGATGCGGCGATGGGGGCAGCCGGCGCGCCCGTATGGCGAGTGGGGTGGGAG
Consensus                              *   *                                                *          *                                   *

Teleogryllus_occipitalis        CAGATTACGACAAGG---------------------CTAAAACCAATGTGTACTCGTTTGTACATTTCGCAAATT---------------------------------------------
Acheta_domesticus               AAATTTAAAATATCGTAGGCCATTCATTTCGGCTTTTGAAAACCTCGGGAAAAATGTTCGTACATCAATCAAACTACACAGTGCTTCTATCTCGCCATTTTTCACTTTGCATCTCAAACT
Gryllus_bimaculatus             TAATTTCTGGTTACGT----------------CATAAACATAATTTACTTGAGTTGTAAAAATATTATGCAAGCT---------------------------------------------
Gryllus_longicercus             AAAGCTGCGGC-------GTCTGTCTTCCCCGCCTGCGCCGGGTTAATGAAACTCGAGCAAATATTAAACGCACC--------------TCCCGCGCTGCTCCCTTTTCCTTTCCGCCTG
Consensus                        *   *                                             *   *     * **     *

Teleogryllus_occipitalis        ------------------------AACTTTCTGCATGTTCTT------------------------TGTAAAATAAA------------GAAAACGGAGAAATAATTTTAT---------
Acheta_domesticus               CACATTCCT--------ATGATGAAACTTTTTGTTCTCTTTT-----TACTCCGACG-----TATTTTTAAAATAAATGCT--------------GGAAAATCAATATTGT-------CC
Gryllus_bimaculatus             -----------------ATATCAAACTTTGGCGGTCATTCTTGAGGACATTTCAATGAGCTATATTTTTGAAATCGATTTTTAACCTGCTATAACAAAGTAGTTGTATTCTCGTAACTAT
Gryllus_longicercus             CGCGCCCCTCCCGAGCGGAGAGGGAATTCTCGGGCCGCCCGTA--CGTCTTCCCAAG-----ATATTTTAAGGTGGACATTTACGGAAGAAGAGAGGATAACACGTGTTGTC------CC
Consensus                                               *  *    *        *                        * * *  *  *                    *  *    * ** *

Teleogryllus_occipitalis        ---------------------------------------------------------------GTATGAGGTCACC------------GGAAAGTAG---GTGCCTATGCGGTAACT---
Acheta_domesticus               AAAATGCTGGACCAA----------------------CGATTGAACAATCAGTTTCATAGAATTTCTAAAGTGATA------------TAAAATTA--------TCTTCCGATACTT---
Gryllus_bimaculatus             GAAGTACTATACAGGTTAAAATTAGACATTTTTTTCTAGAGTGGGTAAGAAAATTCAAGGCGTTCCTGGAATAGCATGGTGAATGATGGAGAAGTGCTACCCATATTTGGGGCAGCC---
Gryllus_longicercus             ATAATGCCACACTGATTTAAA--------GGTCTTGCTAATGGATTTGATCGTTTAATCTTATTTCTACCGTAA---------------GGAAGTA--------TTTTAGGACAACTAGT
Consensus                                                                                         *    *                   ** *            *  *  *

Teleogryllus_occipitalis        --------------------------------------------TTCACTACAAAATGA------------------------------TAAACGGCAAAGATAATTGC-GATGGCGACT
Acheta_domesticus               -------------------------------TGATGCTGAAC---TTCCTACGAAATGGA-----------------------TATCGCAATACGAGAAAAAGCAATAC-AACGGAGTTT
Gryllus_bimaculatus             --------CATTGAGCGGCGCGGAGAGAGAAAGGTGATGGAG--TAGGTCGCGGGGTGGCGGCCCGTGCGTCATGCGAGGGCCTTGGGGAGAGGGGCGCAAGGCGGCGGCGGTGGCAGAC
Gryllus_longicercus             GGCCTGGCAATTAGTTTGCTGTAGGCAAAAATGCTGCTGAACAAAATCCCACAAAATGG--------------------------TAGCATGGAGCCAAAGAGTAGCATGGACAGTGTTC
Consensus                                                                  *    **                                     *    *              *

Teleogryllus_occipitalis        TGTAA-----ATATAATTAAAT--------------------------------AATTAAACTTTAGATCGGATTAATTTCGTTGTCTAGTGATCTGTTGCGGAGGTTTTAATTCCCGTG
Acheta_domesticus               TGAAGCTTTCAGACAGTCAGGT-CTGTTCGAAAGTGCATTGCCATTATTCCTTTTTCAAAATGTTATATCATGCTACT---------------TCTACTCAGGAACTTTCGAATCAAATA
Gryllus_bimaculatus             AGCGC----GGGGAGGCCGGGT------------TGCACCACACTCGGCTCCAGACGGCGTTCTCCCGGCGGGTAACTGTTTCTGCAGCATTCTCCGTC----TCTCCTTTACTTTGACG
Gryllus_longicercus             AGCAA----CAAACAATTTACTATTGTAAGAAAGTATAACG---------------AATAAAATGTTTTA-----------------------------------ACTTCACAAATTCAGCA
Consensus                        *                   *                                          *                                               * *

Teleogryllus_occipitalis        TCTTTGG--------------------------------------------AAAAACAATCA------TAACAACATAATGAATTTCATCTATCAGTCTACTGGGT-CCACTTCCAAATT
Acheta_domesticus               ACGCGGGTGAAAAGTAAAGTATTGAATATCGAAAAAGAGAGAGGGTTAAGAAGAAATCACTTCCGATGTAGAAAGAAATTGAAATTTAATTATC--TTAGCCGA-----GATTTCAAATG
Gryllus_bimaculatus             TCTTCCTCGCAATGTA-----------------------------CTGATAAGATGGTATGCATTATGTTGGAATATTGTTAATTACAAATTTTGGACAGAGGCGTGGCAATGACAGATT
Gryllus_longicercus             ACTCTGGCACGAAGTAGGTGAAACAAACTAAAGAAAGAG------------AAGAACCACTT----------------GGCAACTTCGATTATTAGTTCGACGA-----ACTTTCGGAAG
Consensus                        *                                                 *      *                     ** *     * *         *        *  *  *

Teleogryllus_occipitalis        C---------------------------------------------------------------------------GTGATTGGTTGCCCTACCGAC-----------------------
Acheta_domesticus               TAGGTTACTAGTAAAACTGAATTTGTAATATTTTACTTGTGTTTGTACAATTCTCTCGCTTCCAAAGAAAATAACATTGATCAATTATATCAATGTCGTATCGTTTTCCCTAAAACCTGA
Gryllus_bimaculatus             A-------CGACTAGGCTGTAACTCATGTTTTTTTTTTTT---------------------------------------GTTTTTATATAAATACAAATTCTCATTTCAGAAATTC---
Gryllus_longicercus             C---------TCAAAATTAAATATTTTCCCGTTTAACTGT---------------------------------------ATCATTTATACGAATG-------------------------
Consensus                                                                                       *   **     *

Teleogryllus_occipitalis        ------------------------------------------------------------------------------------------------------------------------
Acheta_domesticus               TGAACTGGAAGCTAAGAAGATAGCAAAGAGAATTGCAGGCAGCATGAATATTGGTATTATTTTGACATCGAACT-----------------TTATTACCACGACGATGGCTTCCTCGGTT
Gryllus_bimaculatus             -ACTCTGCGTGTTAGTTTTAAAACAAAGAAAAATGCTGGGGACATACATTAT----TTATGTGAGCATGAAAATGGTTGATCGTCAGAAAATAGTAACCATTGCGTTATCTTTCCTAGAA
Gryllus_longicercus             --------------------TAATAAAAGTATTAATAAAAACAATAAATATTGTTATAACTTAATTTTGAAATTA--TTATTATT-----ATTATTATTATTATTATTATTATTATTATT
Consensus

Teleogryllus_occipitalis        -------------------------------------------------------------------------------------------------AGCCGCTTCCGGTCACCTTGCCC
Acheta_domesticus               TATGTACCTACGCATACAAAGATATATTATGTAGCCATAATACAGAGTGTAGG--AGTAAACGTTTTGAGACTATTAAATACCGTGACCAGC-----ATCCATTTTTGTTTATTTAGCTT
Gryllus_bimaculatus             AAT-------TATAAACGCAGAGAT-------------------GGGTGTAGACTAACGAATATAATTACATAATTAAAATTCA-GATAAATGTTGTACCGTCTTCTGGGGATTTAAATG
Gryllus_longicercus             AATATTATTACATGAACATAGAGCCAGTTCTACCACTTATTGTAAAATGTG-----------ATTTTTTAATTATTAAATTATAATATAAATA---AATGTATTTCTGTTAAATTAAC--
Consensus                                                                                                        *     **  *   *  *

Teleogryllus_occipitalis        GTCTCTGCATCCGTTACAGC----------------------------------------------------------------------------------------------------
Acheta_domesticus               AATACGGTATCAATTAAAACAAACCAGCTATCATATGCCAGTTGCCTCTAGAGTTCTTCGATAGTATTATACGAGTAATACAAATGAATTAAATTTAAAATCCCATGGTTCCCAAATGTG
Gryllus_bimaculatus             TCTTCGGGAAGGGGCATAATGA--------------------------------------------------------------------------------------------------
Gryllus_longicercus             ---------------------------------------------------------------------------------------ACTGAATTTGGGG----AGCGTTCCGAGATTT-
Consensus

Teleogryllus_occipitalis        ------------------------------------------------------------------------------------------------------------------------
Acheta_domesticus               TTCCTAAGAAGTTTCGGAAATCA------GAGGTAAGGTAGATTTGCACACAAGTTGAGAGAAGATCTGGACGAACACGGTGCCGCGTGCCCACCAGCCCGGATTATCCCGCGCCGAAAC
Gryllus_bimaculatus             ----CAAGAAGTTTCATCTGTCAAC------------TGGGCCCACTTCCAAATTCGTGGTTGATTT-----------------------------------------CCCTACCATAGC
Gryllus_longicercus             -------GAAATTTTATAAATTAAGCCCTGGAGTAGTGTATACGGATACACA----GTAAACAGATGT-------TATCATGCCAAGAATATCAAAGACGTTGTTCTTCCATAACGCAAC
Consensus

Teleogryllus_occipitalis        -------------------------------------------------------------------------------------------------------------------CGTCT
Acheta_domesticus               AAGGAATACAGGCGAGACGGCGCGGTGGAGCTGGCGACGGGAGGCGGGGGCGAAGGTGAGGTATCTGGGTCACGCCTCTACGGCTGACTGGTCTCCGCCGCCGCCGCCGCCGCCGCC
Gryllus_bimaculatus             C-----------------------------------------------------------GGTTCCGCCGCTTCCTCTATCAGCTCGCCCGTCTCCGCATCCGTTCCAGC-----CATAT
Gryllus_longicercus             A------------------------------TGTGAATAGCAAATATTTAATGGAGAGCGGCCTCTTCCTTATAACTGTGCTGTTGCATGACAT---------------------CGGTT
Consensus                                                                                                                          *

Teleogryllus_occipitalis        CCTTCGCCAATAGCACCGCAGC----------------------------------------------------CGTCTTCGACAACCCC-----GCAATCATAGACTCCGCCT------
Acheta_domesticus               GCGTCGCCTCCAATGCCGCGGCGCCCGCGAGGGTCAATCACGTCGACGACCCCGAGTCACCCGAAACGCGCCCGCGCCTTGCGCAACGCTCGCCGTCACCCAAGGGACTCCCTCGTGGAG
Gryllus_bimaculatus             CTTTCGCTATTGCCACCGTAGC----------------------------------------------------CGTCTTTGGCGACCCC-----GTAATCATAACTTCGGC--------
Gryllus_longicercus             GTGTCGTAAGTAGTGAGGCACTGAGTGTGAGAACTGA---------------CGAACTAACATAATCAAATATTTGTATTTTTCAAATCTAGT--TTATTTACAGTTTTTCT--------
Consensus                          ***           *                                                          *  **   * *  *         *   *

Teleogryllus_occipitalis        ------------------------------CGGTTGCATTAGGGAGTTTGTACACCCTCGAACGTTAA----------------AGCACTATCAATCTTCCGTGCATGCGCAGTTCAAGT
Acheta_domesticus               AGGAGACGCCGATGACAACTGTGCGGGCGGCCGCGACATTAGGGAGTTTTTGCACCCTCGAACGTTATAACAATCCAAAGCGCTAGCACTATCGATCTTCTGCGCATGTGCAGTTCAAGT
Gryllus_bimaculatus             ---------------------------------------------------ACGGTGTCGATCACT-----------------TGTCACTGCCGTTCCCCCTATTGCTACCAACACA---
Gryllus_longicercus             -GGAAATATGTATAGAAAT--TCAGAATTGTAGGGATGTTAGGGAGTTTTTGCTCTATCGAACGTTAAAATAATCGAA------AGTACTATCGATCTTCTGTGCATGCGCAGTTCAAGT
Consensus                                                                           *    **** *  *                   ***  *  **  *          **   **

Teleogryllus_occipitalis        TTCGCATCACGTTTGTGCATCTGCGCATGCGCAGAAGATCGATATTGCAATCGATTGCTGCCAGCTGAGGCGTGCAAAAAC-----TCAAAGAGTTTGTGCACGCTTTACAGGACCGCAC
Acheta_domesticus               TTCGCATCACTTTTGTGCATCTGCGCATGCGCAGAAGATCGAGATTGCAGCCGATTGCTGCTCGCTAAGGCGTAAAAAGCTCTCTATTAGAGAGTATTTGCACCCTCGAATGGTAAGCAC
Gryllus_bimaculatus             -----ATCACCGTCACGCACCTAC----------CACGGCGGTCACGTTACCAGCTGCCGCTCCCCATCAC----------------------------------------------CGC
Gryllus_longicercus             TTCGCATCACTTTTGTGCATCTGCACATGCGCAGAAGATCGTGATTGCTGTCGATTGCTGCCCGCTAAGGC-------------------------------------------------
Consensus                            *****  *   *** ** *           *   **     *    *   *** **   *     *

Teleogryllus_occipitalis        TATCGATCTTCTGCGAATGCGGAGATGCACCAACGTGATGCGAAACTTTTTAACTGTGTATGCGC---AGATCGA-GTCCTTTAACGTTGGACGGTGCAGAACCTCTCCCTAAT--CACT
Acheta_domesticus               TATCGATCTTCTGCGCATGCGCAGATGCACTAAAGTGATGCGAAACTTGC--ACTGCGCATGAGCAGAAGATCGATAGTGCTTACCGTTCGAGGGTGCAAAAACTCCTTATTCCCGCACT
Gryllus_bimaculatus             TGCCGCTCCCC--------------------------------------------------------ATCACCGCTGCTGCTCCCCATCT----CCGCTGCCGCTCCCCATCACCGCTGC
Gryllus_longicercus             ----------------------------------------------------------------------------------------------GTGCAAAAACTCCCTATTGATGCATT
Consensus                                                                                                         **     ***    *     *

Teleogryllus_occipitalis        TGT-----------------------------------------------------CACTGTCTTCATCTCGCATCACCGCC-------------GACGCCCCCCATCAGCACTGTG-GC
Acheta_domesticus               GGTTCGCAAAGATGAGAAGCACGGGAGATCAGACGATATCAAAGAAGGAGCTTCTGCACTTTTTTACCGCTCATTCGTGCGCAGCCACAGCCTGGAATGTTTTCTGGCCGAGCTAGA-AA
Gryllus_bimaculatus             CGCTCCCCAT----------------------------------------------CACCGCTGCCGCTCCCCATCACCGCT-------------GCCGCTCCCCATCACCGCTGTGTGC
Gryllus_longicercus             AGC-----------------------------------------------------AACTTTTCTGGAGTTAATTCATTGAC-------------GATTCATTATATGTCACTTAGC-AA
Consensus                        *                                                       **                **                                      *

Teleogryllus_occipitalis        CTCTTCGCCACCTGCGCTAGCACTG----CGGTCTTCGTCGCCGCC-----------------GCCTTTCCACGCGGTCGCAAGGT----AGCTCAAGGAAGTCGTGT------------
Acheta_domesticus               ACTTACTTTCGCTTTTCTAAGTTAAGGTCAGATTTCTTCACTTGCCTATTGAAAACATTATTAGTCTTTACAGAATCATAACAGGTCACCGGTTTCAGAAAATCACGTCAACGATGATGA
Gryllus_bimaculatus             CTCTTCGCCACCTGCGCAGCCACTG----CGATCTCCGCCATCGTC-----------------GCCTTTCCACGCGGTCGCAAGGT----AGCTCAAGGAAGTCGCGGCGGCG-------
Gryllus_longicercus             GCTTTCTTTGTTTGCACGATGTTCA----CAATTCTCATACTT---------------------CCTTTGTGTATTTACGCCAGGT-----------GGAAACCCCACTGACAAAGCCCT
Consensus                          * *      *   *               *                                ****             ****           * **  *

Teleogryllus_occipitalis        ------------------------------------------------------------------------------------------------------------------------
Acheta_domesticus               AATAAATATAGAAACCAGTTTTAGTAATAAACGACCGACGAGGAATTAACAAACTCTGTTAATTAAAACGCGGTGCTAATGCGAAAAAGATTCACAGAGCTTGAGTTTGAAAATACACTC
Gryllus_bimaculatus             ------------------------------------------------------------------------------------------------------------------------
Gryllus_longicercus             CGTTAAGATAACAGC----------------------GCAAGCAGCTGACGGATGTTG-AGATTAGAAGGCCGT-------------CGACCGACTGAGCGCTAGACTATCCGTCCAGTC
Consensus

Teleogryllus_occipitalis        -----------------------CGGGCGACGGCTGCG----GCAC---------------------------GGCGCGACTAGGTGCAGCGGGGCGG----------------------
Acheta_domesticus               CAGCAGCGAGTTTATTCATCATTCAAATGAAAGCCGCA----ACGCAAAAGCTAACGA---------------AGCGCGATGAAGTCCAACGGATCAGATGAAGTCCGATCTAAAAAAGA
Gryllus_bimaculatus             -----------------------GCGGCGGCGGCGGCG----GCGGCGGCGGCGGCGG--------------------------------------------------------------
Gryllus_longicercus             GCGCGGTG---------------CTGACGACAGGCGCGCGGTGTGCAAACGCTGGCGACTGATTCAGCTTGCGAGTGCG-------CTGACGCGGTAGGCGCAGT---------------
Consensus                                                   *   *  **

Teleogryllus_occipitalis        --------------------------------CGCGGGGAGAGTGTTTCGCCGAAGCATGCATG-----------------------------ACTCACGCAGCGCCGCGCGCAGAC---
Acheta_domesticus               TCAAAGCCTGCTTACGACGGAGCGCCGGGTTGCACCACGGAGGTCCTGCTTCGAGGCCCGGGGGCGGCGCGAGCGGCGTGGGCGCCGGCGCCGGCGGAGGCAGGGGCGCGAAGGAGGCCA
Gryllus_bimaculatus             --------------------------------CGCGGGGGCAGAGTTTCACCGAAGCATG---------------------------------ACTCACGCAGCGCCGCGCGCAGAC---
Gryllus_longicercus             TCTTGGCGTGCTTGCGAAAGGCGTCCGTG------GGCGCGCGTTCTTCCGCGGATCGGCAGTGCACAGTGATCGTTGTGA--------------------AGTGTCGAGGAGAAGA---
Consensus                                                              *   *   * *  **   *                                            ** * ** *

Teleogryllus_occipitalis        ----------TCGCACCGCCGCCGCACGCCCAGCCCCAGCTTCTGCCTCTGCCTCTGCCTCTGCCCCTGCCCCTGCCCCTGCCCCTGCCCCTGCCCCTGCCCCTGCCCCTGCCCCTGCC
Acheta_domesticus               GCGACGTCGCCTCCGAGCGCCGCGCCGCGCCGGGACATGCAGTTTGGGTTCAAGGCGGC-----------------------CCACCGCTCCCCCTCCCCCCCCCCACGCGCCGCCGCT
Gryllus_bimaculatus             ----------TCGCACCGCCGCCGGACGCGCAGCCCAAGCACAAGCCCAAGCCCCAGCCTCTGCCGCAGCCTCGG-----------------------------CCCCGGCCGCAGCAGCC
Gryllus_longicercus             ----------TCCAAGTG----------TGAGATTTGTTGTT-----------------------------------------------------TTTTACTTCCCAGCTTGCAGCATCT
Consensus                                  **  *  *              *                                                                      *      **  *  *

Teleogryllus_occipitalis        C-----------------------------------------CTGCCCCTGCCCCTGCCCCTGCCTCTGCCCCTGCCTCTGCCCCTGCCTCTGCCCCTGCCCCTGCCCCTGCCTCTGCCC
Acheta_domesticus               CAGCTGCAGCGCGGTGACGACGGCGG----------------CAGGGCACAGGCAGGGCAGCGCCGCCGCCGCCGGGGCAATGGGCCTCTGACGCACTGCACC-GCCAATACTTTCACTC
Gryllus_bimaculatus             AGAGCGAGGAATTCCATGAACGACGCTGACCCAAAAAACAGAGTGGTTTAGTTGGAAGTTACGCCATTCACGCTACCTCATTCCCATTCTCGGTAC-----TCTAACAATAATTCTGCCC
Gryllus_longicercus             CAAATTGGACGTATTGGTGAAAACAGTCATC--AGTTTCGTTTTGCATCAGGATGTGATATAGCTTTTGTTCTTCAGTTAATGGTCATTTAGGTAA-----------AGTACTTTATTTT
Consensus                                                                    *                 **                          *                   *   *

Teleogryllus_occipitalis        CTGCCTCTG---------------------------------------------------CCCCTGCCTCTGCCCCTGCCTCTGCCCCT----------------------GCCTCTGCC
Acheta_domesticus               GCGCTGCCT---------------------------------------------------ATCCCTCCCT---------------TCCCCGAGATACGATGACGCGCGCGGACCCCAGGG
Gryllus_bimaculatus             GGGCGGTTTCAAAAAAGAACTGTGCTGCGAAAGGCGTAAGGAGAATGTTCGGAAATTCGGATTCATCGCTTGGAAATCTTTCCCATTTCCGTAA------------------------
Gryllus_longicercus             ATGCTGGTT---------------------------------------------------GGCCCTTATTGACGTATGCTTGCGACTTTTGAGTTACGGTAGTTATTACCGGTTTC----
Consensus                         **                                                            *

Teleogryllus_occipitalis        CCTGCCTCTGCTCCTGCCTCTGCCCC------------TGCCTCTGCCCCTGCCTCTGCCCCTGCCTCTGCCCCTGCCTCTGCCCATGCCTCTGCCCATGCCTCTGCCCCTGCCTCTGC
Acheta_domesticus               AACTCTTCTTTTTGTATGGTTTCTCC------------TTGCCCCCTATACGGCAATTTATTCAACTCTGCCCCTGCCAGGTATCAGC----TGTTGATGGTTTCATCC--AACTGGGA
Gryllus_bimaculatus             --ATTTTATTTTATTTTATTTATTGCGAGCAATTTTTATTTTCCCTTTTTACATTAATGAATTCAA---------------AGATCACTAACATCTGCGTAACTCCATTTATAGTGGAGT
Gryllus_longicercus             --TTCTTTTTTTATTAGGGTT------------------------GTTATGTTTCTGTGCAATAAATTTAGTCAGTAACAGTAATCAG--------------------------------
Consensus                             * *  *  *     *                                    *                            **

Teleogryllus_occipitalis        CCCTG-----CCTCTGCCCCTGCCTCTGCCCCTGCCTCTGCCCCTGCCCCTGCC------------------------------------------------------------------
Acheta_domesticus               GCTTA-----TATGTCTTTACCGTGACGTCACTGGTGACGTTCAAGCTCCCGATTCGATGAAACCGTTAACAATAAATC-------------------------------GCAGTATAGA
Gryllus_bimaculatus             AATTGCTTAATTTTTCTCTCACGTACATCCGTTGTAGATTTTTGGTGATTTCTTTTAACGGGATAAGTGAATGTTAAATTCTTCAACCCATTCGATACAACTCGAGAGGAGTGATATACT
Gryllus_longicercus             -----------ATCTATTGTCAGTGAATTCAATAACAATAAT--------------AATAAAAACAGTAACAACTAAT----------------------------------GAAATAGA
Consensus                                   * *               *  *

Teleogryllus_occipitalis        ---------------------------------CCTGCCTCTGCCCCTGCCCCTGCCCCTGCCCCTGCCCCTGCCCCTGC-----------------------------CCCTGCCCCTG
Acheta_domesticus               GGAGACCGGTAT-GTTTGAAAATCGAATGGTGACGACCTTTTGCAAGTAAATTCATT----------------------------------------ATTATCGAATCATCTTATTATTA
Gryllus_bimaculatus             GATGGCAGACTT-GCTGAATTGCTAGTCGCCAGGTTGTCTTATTCAATGTTTTCCTT--------------------------TCATGCAGCTCAACTCTGTGAGAGGCCACTATATTTA
Gryllus_longicercus             AGATGTTAAAATAGTGTTACAAATTTGTACTGACGTGTCTTTAAAAATATTTTCGTTGGTTTTCTTTTACTTGAATTTACTGTTTATGTA----AACGTAATCGAAGTTCGTTGTATTCA
Consensus                                                               *       *                                                              *

Teleogryllus_occipitalis        CCCCTGCCCCTGCCTCTGC----------------------------------CCCTGCCCCTGCCGCAGCCGCAGC-------------------------------------------
Acheta_domesticus               CCGC-------------------------------------------------ATCTTCCTCTGCTGAAGTAGTAGA-------------------------------TAAGTAGAGAAT
Gryllus_bimaculatus             TTTTTATTTTCCGATTCAAATTTATCGAGGCGGGTGAGGGGCGCGGTTATCGAATCTTCAATTTTTCAATTTAAAGC---TCCGGATTGGATAACAGTGACTCCCATCCAATCCAAGAAG
Gryllus_longicercus             CTGTTGACCGTATGTATTTACTTATTGGGTTTAACG---------------TGCCCTTCAACTGCTGAAGTCATTGAGACCCGTGATTG------------------TTGATCTGAGTAA
Consensus                                                                              ** *   *     *      *

Teleogryllus_occipitalis        ------------------------------------------------------------------------------------------------------------------------
Acheta_domesticus               TCATGCACAA-----------------------------------------------------------CGTGAAGC-------------------------------------------
Gryllus_bimaculatus             TGGAGAGAAGCCTCAATAGGATATTCCAATTCTCTTGACTACGTTTTACAACACTAATATAATCGCGCGTGTGAACTAAAGAAAATTCATTTGTATTATTCTGCTTTCAGGAAGAGGGGA
Gryllus_longicercus             TTGAGAGCGATTATGA----------------------------------------------------CTGTGAAGTAAAAG--------------------------------------
Consensus

Teleogryllus_occipitalis        -----------------CGCATCCGCAGCCG---------CAGCCGCCAGAGCGAGGCATTCCGTGAACGCAGCTGA-------------------------------------------
Acheta_domesticus               ---TCCTTAGCGTATACCACGTTTGCGCCTCTGCG-----TAAGCGCCCACTCGA-----TCTCGAAATGAAGCAGAGCCTGCT------------------------------------
Gryllus_bimaculatus             ATATGAGCAATGTTTGCCCCTTTTCTAGGATTGATTTTAGCAGTTACTAGCATAA-CCTTCGTTAAAACGACTTAAAAACAAACGCAATGAATGACCATCGAGCAAGAACCTTATATCAG
Gryllus_longicercus             --ATATTTAAGGACTGTCAGATTTGGAAGTGAGAGAGCTCCGAATACCTACCTAAAGCCCTCTCTGACCAGCACAGAGTTCATTTAGAT-------------------------------
Consensus                                        *   *                        *       *           *         *

Teleogryllus_occipitalis        -----CCCAAAAAACTGAGTGGTTTAGGTGGAA------------GTTACGCCATTCACGCTACCTCATTCCCATTC-----------TCAGTACTCTTACAATAATTCTGTT-----
Acheta_domesticus               ------------ATCACGTCGAATTTAGCGGGC------------AACTTATATCACACAGTAATCCATCACTC--------------------------------------GGTCGTAG
Gryllus_bimaculatus             AGAGAGCAAAAAAGCTCTGTGAAATAAAAGAAAAACAATTTCCTCTATTCTGTAATTCGCAAAAATCCTTTCCTT------------CATTAATGCGTATCT------------------
Gryllus_longicercus             ---TACCTAATGATCGCGGTGGAAAGCGAATGC------------GTTAACTTTTACATCATAATCCCCCACCCTCCCAATGTTAGACATTGGTGATTGTCTAACCTTTGTAGTCGT--
Consensus                                   * *     *                                    *     *   *    *

Teleogryllus_occipitalis        TGGACGGGTCCAAAAAGGAACTGTGGTGCGAGAAGGGTAAAGAAAATG-----TTCCGTA---AATTCATATTCACAGCATGAAAATCTTTCCCATGTTGCTTATTTCATTGTTTTATTT
Acheta_domesticus               TAAATCGATTTAGTTCCCAACTGTGGCGACAGAGTGGGCTGCTAAATGTGCATCCTCATG----ATGCAATATCAAGTTGGTGGCATCCATCTGCGAATATAAATATCTGTATATATGAA
Gryllus_bimaculatus             CGAACGCTATCAAGGTAAGGATGAAGCACTAAAGGTTTCCGGCCGTGGG----TTGCGTA---ACTCCGCCCTGTGGGCACCGGAAGTCTCTCGCGCATGCATGTGCATTT---------
Gryllus_longicercus             TGAATACATGCGAGACCCAATTGTATTGTCA----------------------TCTCATGTTATCTGTGAATTCATAACGTTGACACTCATTTCCAGGAACTTGTATTCCTGGTTAATAA
Consensus                          *                 **        *                         * *       *       *             *                   *     *

Teleogryllus_occipitalis        ATTTATTGTGGACAAATTTTGT-----------------------------TTTACATTTCACATTCATGAATTTCACTAAAGTCTGGTTAATATCCT--------------------TT
Acheta_domesticus               AT------TAGTTATAATTAA---------------------------------------TATAATAATGATAATAATAATACTAATAATAATATTCTATAATATAATTGTATATTCATA
Gryllus_bimaculatus             -------CTGGGCCAGCTGCGCCGCCGCCGCCGCCGCCGCTGTCGCATCCGCTGGCAATTCGCGACAGGAA-----GCAAGGCTCTCAGATGCGTCCTCTCCCCCTCCTCTTTGTGCGCC
Gryllus_longicercus             ATT----CTAGGAATATTTTGC-----------------------------GAGGCATTTTGTGCTATCGACTTTACTACAGTTCAGAGTCGTATTCCGTAATTACGCATCCGAAGCATG
Consensus                               * *      *                                                     *             *          * *

Teleogryllus_occipitalis        TTAGTAGGAGGATATTT----------------------------------------------------------------------------------------------------CTGAT
Acheta_domesticus               TCAGTGA------------------------------------------------------------------------ACAAGGGCAGCTGTACTG--------------------AAC
Gryllus_bimaculatus             TTAGTGTTGGCGCGCTG-----------CCTTTGAGCGATCCGATCCCACTCACCCCTGCACCGACTCCCCTCTCGCACTCCGTCCCGCTACGCCGCTCCTTGACGTTCCCCATCTGTC
Gryllus_longicercus             TTGGTGACGTGTGATTTGGGTGCTCATCAGGTTTCAGTAATTTG-------CAGCTCCTAATACGACGTGTTTTCTGTGGCAATGCCAGACGTCCTGT----TGGTGTTCCAC-CTTGGC
Consensus                       *  **

Teleogryllus_occipitalis        CCTCTCGAGT--------------------------------------------------------------------------------------------------------------
Acheta_domesticus               CAATGCGAAT--------------------------CAAGAGAATGGGGCAGTATTGAA-----ATTTCGCGCTCACTTTG---------------TCGCGAGAGTTCGAGACTACATCT
Gryllus_bimaculatus             CCACCCCCACGTTGCTCGGCTCGTCTGTTGTGTTC-CGCCCTGCCAGGCCTTTACTCTTTCCGCCCCTCCCACTTTGTTCCGCTCCATTGAACTATGTTCCACCGTACCCTGCTCTCCCC
Gryllus_longicercus             CGCCGCGAATGTCGGCCGACCCGCGTCCATTGTTCCCACGGCTGTGGGCGCCTGCTGTAC-GGCTCTTGGTGCGCGGTTCGAGTCCGGCCAGC-AACTATTAGCGCTAGATACGGCGTTC
Consensus                       *    *

Teleogryllus_occipitalis        -------ATTTCCATTGTAG----------------------------------------------------------------------------------------------------
Acheta_domesticus               AATATTGATTTCGATGATAGCATCGTAAATGCCAC-------------------------------------------------------------------------------------
Gryllus_bimaculatus             CACTCCCACTCCTACTCTACTCTACTGCCTTGCCC-------TGACTTGCCCTGACTTGCCCTGACTTGCCCTGACTTGCCCTGACTTGCCCTGACTTGCCCTGCCCTACTTCCACCTGT
Gryllus_longicercus             AACTCCGGCCGCGGCCGTGGCCGCGTGCGCCCCGCGGTGGAGCGGGTGCCGCTG------------TTGCTAGGTTCTCCCCGCCCATGCCCCATACTATCCCGCCCGGCC---------
Consensus                                  *     *

Teleogryllus_occipitalis        ------------------------------------------------------------------------------------------------------------------------
Acheta_domesticus               ------------------------------------------------------------------------------------------------------------------------
Gryllus_bimaculatus             CCCGTTCCCGTTTCTTCACCTTGCCGATCCTTCAGCATTTCCCCGCTTCGTAAGCCCATAAATTTCGATGTTCTCATCGCAGGGCAATCACACCTTTCTCACTATTACACAATTTCCCTC
Gryllus_longicercus             ------------------------------------------------------------------------------------------------------------------------
Consensus

Teleogryllus_occipitalis        -------------------------------
Acheta_domesticus               -------------------------------
Gryllus_bimaculatus             TCGCCTTATCTCCCTTCACTCTCCTGAGGTT
Gryllus_longicercus             -------------------------------
Consensus
```

**Tocc.4G0011610.1_exon1**

```
Teleogryllus_occipitalis      ------------------------------------------------------------------------------------------------------------------------
Acheta_domesticus             --------RSTIL*VPDPTFHYLVKCNYSDKRIAGKEILGHSCHQGQSLRQFLS------------SISTPT*SSRRPELPL*CALLSPDPSFPVAQQKFKIS*AIHFGF*KPR-----
Gryllus_bimaculatus           TPCTKLYNSSNVSACPFE*IDILLLC------------------RNK*INKY*N------------ELKITFLHS---SVY*YCV-----PSF---KRMSNINCEIKLNWTCFK*NMYK
Gryllus_longicercus           ---------------------SLPGC------LAAR--------RGAAQRRGGSGGGCAGAGADGKAEVKSRVRSDGRGARGGKCTCA---GSN---KRRWTICCGIQM-----------
Consensus

Teleogryllus_occipitalis      ------------------------------------------------------------------------------------------------------------------------
Acheta_domesticus             ---EKCSYINQTTQCFYLAIFHFASQTH-----IPMMKLFVLFLLRRIFKINAGKS---ILSKMLDQRLNNQF---HRISKVI*NYLPIL*C*TSYEMDIAIREKAIQRSFEAFRQSGLF
Gryllus_bimaculatus           MLCSKYSYR*EM*NCVKLLLI-CEKETHLTCVTIHIISLNVFFL---VFKCTYKFSEDIFFMYSTNQQKNNNFWLRHK-----HNLLEL*KYYASY-----IK---LWRSFLRTFQ*AIF
Gryllus_longicercus           -------------------------RLHLE-----------------------------------------------------------------------VNEKRVWKSKIAKRKTE-Y
Consensus

Teleogryllus_occipitalis      ------------------------------------------------------------------------------------------------------------------------
Acheta_domesticus             ESALPLFLFQN---VISCYFYSG---TFESNNAGEK*S------------------------IEYRKRERVKK-----------KSL---------------------------------
Gryllus_bimaculatus           LKSIFNLL*QSSCILVTMKYYTG*N*TFFS-RVGKKIQGVPGIAW*MMEK----CYPYLGQPIERRGERKVME*VAGWRPVRHARALGRGAQGGGGGRQRGEAGLHHTRLQTAFSRRVTV
Gryllus_longicercus           ARTLNHLKWQV-----------GLPTTRME-NEGAAIG--DGREANMWSRCLRWCWQIAGGY*GRVRRRRVEG------ECRN-SASGRDAAMGAAGAPVWRVGWE-------------
Consensus

Teleogryllus_occipitalis      ------------------------------------------------------------------------------------------------------------------------
Acheta_domesticus             -------PM---*KEIEI*LS*PR-------------------------------------FQM*VTS---------KTEFVIFYLCLY--------------NSLASKENNIDQLYQC
Gryllus_bimaculatus           SAAFSVSPL---L*RL------PRNVLIRWYAL----------------CWNIVNYKFWTEAWQ*QITT---------RL*LMFFFFVFYI-------------NTNSHFRNSL-----C
Gryllus_longicercus           KAAASVFPACAGLMKLEQILNAPPALLPFPFRLRAPPERRGNSRAARTSSQDILR---WTFTEEERITRVVP*CHTDLKVLLMDLIV*SYFYRKEVF*DN*WPGN*FAVGKNAAEQN---
Consensus

Teleogryllus_occipitalis      ----------------------------------M---------------------------------------------------------------------------------------
Acheta_domesticus             RIVFPKT**TGS*EDSKENCRQHEYWYYFDI--EL----------------YYHDDGFLGLCTYAYKDILCSHN---TECRSKRFET-IKYRDQHPFLF-------------I*LNTVSI
Gryllus_bimaculatus           VLVLKQRKMLGT------------YIIYVSM--KMVDRQKIVTIALSFLENYKRRDG----------------------CRLTNIIT*LKFR---*MLY-------------RLLGI---
Gryllus_longicercus           ----PTKW*HGAKE*HGQCSATNNLLL*ESITNKMF*LHKFSNSGTK*VKQTKEREEPLGNFDY*FDELSEAQN*IFSRLTVSFIRM**KY**KQ*ILL*LNFEIIIIIIIIIIIIINI
Consensus

Teleogryllus_occipitalis      ------------------------------------------------------------------------------------------------------------------------
Acheta_domesticus             KTNQLSYASCL*SSSIVLYE*YK*IK-------FKIPWFPNVFLRSFG--------NQR*GR---FAHKLREDLD---------------EHGA---ACPPARIIPRRNKEYRRDGAVEL
Gryllus_bimaculatus           ---*MSSGRGIMTRSFIC----------------QLGPLPNSWLISLP*PVPPLPLSARPS-----PHPFQPYLSLLPP*PSLATP*S*LRHGVDHLSLPFPLLLPTQSPSRTYHG----
Gryllus_longicercus           IT*T*SQFYHLL*NVIF*LLNYNINKCISVKLTLNLGSVPRFEIL*IK------PWSSVYGYTVNRCYHAKNIKDVVLP*RNM*IANI*WRAAS---SL*LCCCMTSVVS*VVRH*V*EL
Consensus

Teleogryllus_occipitalis      ------------------------------------------------------------------------------------------------------------------------
Acheta_domesticus             ATGGGG----------------EGEVSGSRLYG*L-----------------VSAAAAAAAAASP---PMP-----------RRPRGSITSTTPSHPKRARALRNARRHPRDSLVERRRR
Gryllus_bimaculatus           -----------------------GHVTSCRSPSPLPL--------------PITAAAPHLRCRSPSPLPLPITA-AAPHHRCRSPSPLCASSPPAQPLRS---------PPSS-------
Gryllus_longicercus           TN*HNQIFVFFKSSLFTVFLEICIEIQNCRDVREFLLYRTLK*SKVLSIFCACAVQVSHHFCASAHAQKIVIAVDCCPLRRAKTPY*CISNFSGVNSLTIHYMSL----SKLSLFARCSQ
Consensus

Teleogryllus_occipitalis      ------------------------------------------------------------------------------------------------------------------------
Acheta_domesticus             *Q--LC-----GRPRH*GVFAPSNVITIQSASTIDLLRMCSSSFASLLCICACAEDRDCS----RLLLAKA*KALY*RVFA--PSNGKHYRSSAH-------AQMH*SDAKLALRMSRRS
Gryllus_bimaculatus           -------------PFHAVAR*LKEVAAAAAAAAAAAAAARGQSFTEA*LTQRRAQTRTAA---GRAAQAQAQAPASAAASA--PAAAARARNS-----------MNDADPK------NRV
Gryllus_longicercus           FSYFLCVFTPGGNPTDKALVK----ITAQAADGC*D*KAVDRLSARLSVQSRGADDRRAVCKRWRLIQLASALTR*AQFLACLRKASVGARSSADRQCTVIVVKCRGEDPS------VRF
Consensus

Teleogryllus_occipitalis      ------------------------------------------------------------------------------------------------------------------------
Acheta_domesticus             IVLTVRGCKNSLFPHWFAKMRSTGD-QTISKK---------------------ELLHFFTAHSCAATAWNVFWPS*KTYFRFSKLR-----SDFFTCLLKTLLVFTES*QVTGFRKS--
Gryllus_bimaculatus           V*LEV---TPFTLPHSHSRYSNNNSARAVSKKNCAAKGVRRMF---GNSDSSLGNLSHFRKFYFILFIASNFYFPFLH**IQRS-LT-----SA*LHL*WSNCLIFLSRTSVVDFW*FLL
Gryllus_longicercus           VVFYFPACSISNWTYW*KQ-------SSVSFC------IRM*YSFCSSVNGHLGKVLYFM-----------LVGPY*RMLATFE-LR*LLPVSSFF-------------IRVVMF-----
Consensus

Teleogryllus_occipitalis      ------------------------------------------------------------------------------------------------------------------------
Acheta_domesticus             RQR**NKYRNQF***TTDEELTNSVN*NAVLMRKRFTELEF----------------ENTLQQRV------YSSFK*KP---------QRKS*RSAM--KSNGSDEV-------------
Gryllus_bimaculatus           TG*VNVKFFNPF-DTTREE*YTDGRLAELLVARLSYSMFSFHAAQLCERPLYLFLFSDSNLSRRVRGAVIESSIFQFKAPDWITVTPIQSKKWREASIGYSNSLDYVLQH*YNRACELKK
Gryllus_longicercus           -------LCNKF----SQ*Q*SD------------------------------LLSVNSITIIIK-----------------TV--------------TTNEIEDV------------K
Consensus

Teleogryllus_occipitalis      ------------------------------------------------------------------------------------------------------------------------
Acheta_domesticus             ---------------------------------RSKKDQSLLTTERRVAPRRSCFEARGRRERRGRRR------------------RRRQGRE--------GGQRRRLRAPR--RAGT
Gryllus_bimaculatus           IHLYYSAFRKRGI*AMFAPFLGLILAVTSITFVKTT*KQTQ*MTIEQEPYIRES------KKAL*NKRKTISSIL*FAKILSFINAYLERYQGKDEALKVSGRGLRNSALWAPEVSRACM
Gryllus_longicercus           IVLQICTDVSLKIFSLVFFYLNLLFM*T*SKFVVFTVDRMYLLIGFNVPFNC*SH*D---------------P*LLI*VIESDYDC-----EVKDI*------GLSDLEVRELRI-----
Consensus

Teleogryllus_occipitalis      ------------------------------------------------------------------------------------------------------------------------
Acheta_domesticus             CSLGSRRPTAPPPPPPRAAAQLQRGDDGGRAQAGQRRRR--RGNGPLTHCTANTFTRAAYPSLPRDTMTRADPRELFFLYGFSLPPIRQFIQLCPCQVSAVDGFIQLGA----YMSLP*R
Gryllus_bimaculatus           CISG---PAAPPPPPPPLSHPLAIR----DRKQGSQMR--------PLPL----LFVRLSVGALP-----LSDP--------IPLTP-------APTPLS---------------HSVPLR
Gryllus_longicercus           -------PT*SPL*PAQSSFRLPN----DR--GGKRMR*LLHHNPPPSQC---------------*TLVIV*PL-------*SLNT-------CETQLY---CHLMLSVNS*R*HSFPGT
Consensus

Teleogryllus_occipitalis      ------------------------------------------------------------------------------------------------------------------------
Acheta_domesticus             HW*RSSSRFDETVNNKSQYRGDRYV*KSNGDDLLQVNSLLSNHLIITASSSAEVVDK*RIHAQREAP*RIPRLRL------CVSAHSISK*SRACYHVEFSGQLISHSNPSLGRSK----
Gryllus_bimaculatus           --------------------------RSLTFPICPTPTLLGSSVVFRPAR----------------PLLFPPLPL------CSAPLNYVPPYPALPH--------SHSYSTL--------
Gryllus_longicercus           --------------------------------CIPG**ILG---IFCEAFCA--IDFTTV------QSRIP*LRIRSMLVTC--------------------DLGAHQVSVICSS*YDVF
Consensus

Teleogryllus_occipitalis      ------------------------------------------------------------------------------------------------------------------------
Acheta_domesticus             SI*FPTVATEWAA----------KCASS*CNIKLVASIC-------EYKYLYI*N*L*LI******Y***YSII*LYIHISEQGQLY*TNANQENGAVLKFRAH------------FVAR
Gryllus_bimaculatus           -LPCPDLP-*LAL----------TCP----DLP*LALTCPALLPPVPFPFLHLADP---------------SAFPRF--VSP*ISMFSSQGNHTFLTITQFPSR------------LISL
Gryllus_longicercus           SVAMPDVLLVFHLGRRECRPTRVHCS-HGCGRLLYGSWCAVRVRP-------------------------ATISARY---------------------GVQLRPRPWPRAPRGGAGAAVAR
Consensus

Teleogryllus_occipitalis      -----------------
Acheta_domesticus             VRDYI*Y*FR**HRKCH
Gryllus_bimaculatus           HSPEV------------
Gryllus_longicercus           FSPPMPHTIPPG-----
Consensus
```

**Tocc.4G0011610.1_exon2**

```
Teleogryllus_occipitalis    ------------------------------------------------------------------------------------------------------------------------
Acheta_domesticus           -----GIYDA------------------IIE--INIRCSLELSRQSEREISI-LPHS-------------------LDSHW--FST---AALVH*YEYTIIL*NIIISIIIIIIILIITN
Gryllus_bimaculatus         -TPCTKLYNSSNVSA---------CPFE*ID--ILLLCRNK*INKY*NELKITFLHSSVY*YCVPSFKRMSNINCEIKLNWTCFK*NMYKMLCSKYSYR*EM*NCVKLLLI---------
Gryllus_longicercus         SLP--GCLAARRGAAQRRGGSGGGCAGAGADGKAEVKSRVRSDGRGARGGKCTCAGS-------------------NKRRW---------------------------------------
Consensus

Teleogryllus_occipitalis    ------------------------------------------------------------------------------------------------------------------------
Acheta_domesticus           FIYTDIYIRRWMPPT*YC--IMRMHI*QPTLSPQLGTKSIYYDRVMDYCVI*VAR*IRRDSRLCFISRSSGR-------LRRGANVVYAKELHVVHEFSTYLLLQQRKMR***DDSI---
Gryllus_bimaculatus         -----------------C--EKETHLTCVTI--HIISLNVFF---------LVFKCTYKFSEDIFFMYSTNQQKNNNFWLRHKHNL-----LEL*KYYASYIKLWRSFLRTFQ*AIFLKS
Gryllus_longicercus         --------------TICCGIQMRLHL-------EVNEKRVWKSKI----------------------------------AKRKTE--YAR------------------------------
Consensus

Teleogryllus_occipitalis    --------------------------------------------------------------------------------------------------RRAQTR----------------
Acheta_domesticus           IMNLLAKGRHHSIFKHTGLLYTAIYC*RFHRIGSLNVTSDVTVKTYKLPVG*NHQQLIPGRGRVE*IAV*GARRNHTKRRVPWG--PRASSYLGEGRDRQRE*K------YWRCSASEAH
Gryllus_bimaculatus         IFNLL*QS---SCILVTMKYYTG*N*TFFSRVGK------------KI-------QGVPG--------------------IAW*MMEKCYPYLGQPIERRGERKVME*VAGWR-------
Gryllus_longicercus         TLNHL-------------KWQVGLPTTRMENEGA------------AI-----------GDGR---------EANMWSRCLRWC-WQIAGGY*GRVRRRRVEGE----------------
Consensus                                                                                                                 *

Teleogryllus_occipitalis    ---------------------------------------------------------------------------------TAAARPAPASASASASA----------------------
Acheta_domesticus           CPGGGGAALPVPCRRRHRAAAERRRVGGGEGERW--------AALNPNCMSRRGAALGG---------------------DVAGLLRAPASAGAGAHA--ARAAPGPRSRTSVVQPGAPS
Gryllus_bimaculatus         ---------PV----RHARALGRGAQGGGGGR-------------------QRGEA--GLHHT---------------RLQTAFSRRVTVSAAFSVSPLL*RL---PRNV----------
Gryllus_longicercus         ------------C--RNSASGRDAAMGAAGAPVWRVGWEKAAASVFPACA--------GLMKLEQILNAPPALLPFPFRLRAPPERRGNSRAARTSSQDILRWTFTEEERITRVVP*CHT
Consensus                                                                                                          *

Teleogryllus_occipitalis    ------------------------------------------------------------------------------------------------------------------------
Acheta_domesticus           *AGFDLF*IGLHLI---------------------------------RW--------------------------------------------------------------TSSRFV-SF
Gryllus_bimaculatus         ------------LI---------------------------------RWY----------------ALCWNIVNYKFWTEAWQ*QITTRL--------*LMFFFFVF-YINTNSHFRNSL
Gryllus_longicercus         ----DLKVLLMDLIV*SYFYRKEVF*DN*WPGN*FAVGKNAAEQNPTKW*HGAKE*HGQCSATNNLLL*ESITNKMF*LHKFSNSGTK*VKQTKEREEPLGNFDY*FDELSEAQN*IFSR
Consensus

Teleogryllus_occipitalis    ------------------------------------------------------------------------------------------------------------------------
Acheta_domesticus           CVAAFI*M------MNKLAAGVYFQTQAL*IFFALAPRFN*QSLLIPRRSFIT----KTGFYIYFIIVDVIF*NR*PVMIL*RLIMFSIGK*RNLTLT*K--SESKF-SSSARKHSRLWL
Gryllus_bimaculatus         CV---LVL------KQRKMLGTY-------IIYVSMKMVDRQKIVTIALSFLENYKRRDGCRLTNIIT*LKFR-----*MLYRLLGI*MSSGRGIM-------TRSF-ICQLGPLPNSWL
Gryllus_longicercus         LTVSFIRM**KY**KQ*ILL*LNFEIIIIIIII----------IIIIIINIIT----*T*SQFYHLL*NVIF*----------LLNYNINKCISVKLTLNLGSVPRFEIL*IKPWSSVYG
Consensus

Teleogryllus_occipitalis    -------------PAPAPA------------PAPAP--------APAP------------------------------------------------------------------------
Acheta_domesticus           RTNER------*KSAEAPSLI----------SSDLP---CFSSLRTSAGIRSFCTLER*A--------------------------------------------------LSIFCSCAVQ
Gryllus_bimaculatus         IS----------LP*PVPPLPLSA------RPSPHPFQPYLSLLPP*PSLATP*S*LRHG--------------------------------------------------VD-------H
Gryllus_longicercus         YTVNRCYHAKNIKDVVLP*RNM*IANI*WRAASSL*LCCCMTSVVS*V--------VRH*V*ELTN*HNQIFVFFKSSLFTVFLEICIEIQNCRDVREFLLYRTLK*SKVLSIFCACAVQ
Consensus                                    *

Teleogryllus_occipitalis    ---------------------------APAPAPAPAPASAP--------ASAPASAPAPAPASAP---------------ASAPASAPASAPASAPASAPASAP----------ASAPAS
Acheta_domesticus           VSHHFSASAHAQKIDSAYHSR--VQILSNRELFTP*RAAIGCN-LDLLRMRRCTKVMRNLNCTCAEDR*C*RFGLL*RSRVQKLPNVAAARTVVIGVSSPRGSPLGDGERCARRGRVSGD
Gryllus_bimaculatus         LSLPFPLLLPTQSPSRTYHGGHVTSCRSPSPLPLPITAAAPH-----LRCRSPSPLPLPITAAAPH------------HRCRSPSPLCASSPPAQPLRSPPSSPFHAVAR*LKEVAAAAA
Gryllus_longicercus         VSHHFCASAHAQKIVIA------VDCCPLRRAKTPY*CISNFSGVNSLTIHYMSLSKLSLFARCSQ----FSYFLC--------------------VFTPGGNP---TDKALVKITAQAA
Consensus                                                     *                                                                *   *

Teleogryllus_occipitalis    APASAPASAH-----------------------ASAH-----------ASA---PASAPASAPASA------------------------------------------------------
Acheta_domesticus           SGSST*LTLAGAAALEATRRRRRRRRRRPVSRRGVTQIPHLRPRL--PSPA---PPRRLACIPCFGAG*SGLVGTRHRVRPDLLSTCVQIYLTSDFRNFLGTHLGTMGF*I*FICITRII
Gryllus_bimaculatus         AAAAAAAAARGQSFTEA*LTQRRAQTRTAAGRAAQAQ-----------AQA---PASAAASAPAAAA----------RARNSMND--------ADPKN-------------------RVV
Gryllus_longicercus         DGC*D*KAVDRLSA------RLSVQSRGADDRRAVCK----RWRLIQLASALTR*AQFLACLRKASVG----------ARSSADRQC------------------------------TVI
Consensus                                                                     *        *

Teleogryllus_occipitalis    ------------------------------------------------------------------------------------------------------------------------
Acheta_domesticus           LSKNSRGNWHMIAGLF*LIPY*AK*TKMDAGHGI**SQNVYSYTLYYGYIIYLCMRRYINRGSHRRGNKVRCQNN------TNIHAACNSLCYLLSFQFIRF*GKRYDIDIIDQCYFL--
Gryllus_bimaculatus         *LE------------------------------------VTPFTLPHSH------SRYSNNNSARAVSKKNCAAKGVRRMFGNSDSSLGNLSHFRKFYFILFIASNF--------YFP--
Gryllus_longicercus         VVK--------------------------------------------------------------------CRGEDP------------------SVRFVVF-------------YFPAC
Consensus

Teleogryllus_occipitalis    ------------------------------------------------------------------------------------------------------------------------
Acheta_domesticus           ----WKRENCTN--TSKILQIQFY**PTFEIS-AKIIKF-------QFLSTSEVISS*P-SLFFD----------------------IQY--FTFH-------PRYLIRKFLSRSSMI*H
Gryllus_bimaculatus         ----FLH**IQRSLTSA*LHL*WSNCLIFLSR-TSVVDF------W*FLLTG*VNVK-----FFNPFDTTREE*YTDGRLAELLVARLSYSMFSFHAAQLCERPLYL---FLFSDSNLSR
Gryllus_longicercus         SISNWTYW*KQSS-VSFCIRM*YSFCSSVNGHLGKVLYFMLVGPY*RMLATFELR*LLPVSSFF-----------------------IRV-------------VMFLCNKF---------
Consensus

Teleogryllus_occipitalis    ------------------------------------------------------------------------------------------------------------------------
Acheta_domesticus           FEKGIM----AMHFRTD--LTV*KLQ------------NSVVLLFLVLRYPFRRKFSIK---------------------------------------------VSEDNFI-------SL
Gryllus_bimaculatus         RVRGAVIESSIFQFKAPDWITVTPIQSKKWREASIGYSNS---LDYVLQH*YNRACELKKIHLYYSAFRKRGI*AMFAPFLGLILAVTSITFVKTT*KQTQ*MTIEQEPYIRE--SKKAL
Gryllus_longicercus         -----------SQ*Q*SDLLSV----------------NSITII-------------IKTV---------------------------------TTNE------IEDVKIVLQICTDVSL
Consensus

Teleogryllus_occipitalis    ----------------------------------------------------------------------PASAPAPAPASAP------------APAP---------------------
Acheta_domesticus           *KFYETDCSIVGPAFWTILIFQHLF-*KYV--GVKREQKVSS*ECEFEMQSEKWRDRSTV*FD*CTNIF-------------PEVFKSR------NEWPTIF*I----------------
Gryllus_bimaculatus         *NKRKTISSIL--*FAKILSFINAYLERYQ--GKDEALKVSGRGLR---NSALWAPEVS---RACMCISGPAAPPPPPPLSHPLAIRDRKQGSQMRPLPLLFVRL---------------
Gryllus_longicercus         KIF-----SLV--FFYLNLLFM*T*-SKFVVFTVDRMYLLIGFNVPFNC*S--------------------------------------------H*DP*LLI*VIESDYDCEVKDI*GL
Consensus                                                                                                                 *

Teleogryllus_occipitalis    -------APAPAPAP-APAPAPAS------------------APAPAAAAAASAAAAARA------------------------------RHS---------------------------
Acheta_domesticus           ----------------SVAPLEMK-----DREIT--------------------------------------------------------EHTKA-------------------------
Gryllus_bimaculatus         ---SVGALPLSDPIPLTPAPTPLSHSVPLRRSLTF-----PICPTPTLLGSSVVFRPARPLLFPPLPLCSAPLNYVPPYPAL------PHSHSYSTLLPCPDLP*LALTC-PDLP*LALT
Gryllus_longicercus         SDLEVRELRIPT*SPL*PAQSSFR--LPNDRGGKRMR*LLHHNPPPSQC*TLVIV*PL*S-----LNTCETQL-----YCHLMLSVNS*R*HSFPG------------TCIPG**ILGIF
Consensus                                     *                                                                        *

Teleogryllus_occipitalis    --------------------------------------VNAADPKN*-------------------------------------------------------------------------
Acheta_domesticus           --ATLAVVSFMLEWKWKTETA*E-------------IGLGGMNAQVF------LYLLFA-----------------------CHYNCTSL--------------NSEKLGQVL-------
Gryllus_bimaculatus         CPALLPPVPFP--------------------------FLHLADPSAFPRFVSP*ISMFS-----------------------SQGNHTFL--------------TITQFPSRL-------
Gryllus_longicercus         C-EAFCAIDFT--------TVQSRIP*LRIRSMLVTCDLGAHQVSVI---CSS*YDVFSVAMPDVLLVFHLGRRECRPTRVHCSHGCGRLLYGSWCAVRVRPATISARYGVQLRPRPWPR
Consensus

Teleogryllus_occipitalis    ------------------------
Acheta_domesticus           ---------IESYF----------
Gryllus_bimaculatus         ---------ISLHSPEV-------
Gryllus_longicercus         APRGGAGAAVARFSPPMPHTIPPG
Consensus
```

**Tocc.4G0012180.1**

```
Teleogryllus_occipitalis    CAGCTCGACGCTGTGCGCGGACGGGACCGGACGAGAGGGCCACAATTCACGCGGCCGACAGCTCCGCAGGCGGGGACATTTTTAC------CGGCCGGCCGCGTCCACGGGCCGTGGGCC
Acheta_domesticus           CAGCTCTACGCTGTGCGCGGACGGGACCGGACGAGAGGGCCACAATTCACGCGGCAGACAGCTCCGCAGGCGGGGACATTTTTAC------CGGCCGGCCGCGTCCACGGGCCGTGGGCC
Gryllus_longicercus         CAGCTCGACGCTGTGCGCGGACGGGACCGGACGAGAGGGCCACAATTCACGCGGCGGACAGCTCCGCAGGCGGGGACATTTTTACTGGCCGCGGCCGGCCGCGACCACGGGCCGTGGGCT
Consensus                   ****** ************************************************ *****************************      ************ ***************

Teleogryllus_occipitalis    TCATCCGGTCTCATTAAGATCCAGCGCGAGGGGGTGATAGCGGCGGCGGAGGTGGAGGTAGCGGTGGCAGTGGCTTTGCCGGTGGTGGCGGAGGGGAAGGTGGCGGCGGCGATGGTGGTG
Acheta_domesticus           TCATCCGGTCTCATTAAGATCCAGCTCGAGGGGGTGATA---GCGGTAGTGGTGAGTGTGAC------------------------------------------------------------
Gryllus_longicercus         TCATCCGGTCTCATTAAGATCCAGCGCGAGGGGGTGAAAGAGGCGGCGGAGGTGGGCGTTTC------------------------------------------------------------
Consensus                   ************************* *********** *   ****  * ****   **  *

Teleogryllus_occipitalis    GTAGTGGTGGAGGCGAAGGTGGTGGTGGCGGTGGTGGTGGTGGCGGTGGCGGCGCTGGCGACACCGATGGCGGTGCTGGTGGTGGTGGTGGTGGTGGTGGTGGTGGTGGTGGTGGTGGTG
Acheta_domesticus           ------------ACGAAGGAGGCGGTGGTGGTGGTGGTGGTGGTGGTGGTGG-----------------------TGGCGGTGGCGGTGGCGGTGGCGGTGGCGGTGGCGGTGGCGGTG
Gryllus_longicercus         ------------ACGAAGGCGGTGGTGGCGGTGATGGTGGTGGCGGTGGCGG-----------------------TGGCGGTGGCGGTGGCGGTGGCGGTGGCGGTGGCGGTGGCGGTG
Consensus                                ****** ** ***** **** ********* ***** **                       *** ***** ***** ***** ***** ***** ***** ****

Teleogryllus_occipitalis    GTGGTGGTGCAGGTGGTGGTGGTGGTGGTGGTGGTGGTGGCGGCGGCGGCGGAGGCAATGGTGGTGGCGGCAATGGTGGTGGCGGTGGTGGAGGCGAAGGTGGCGGTGGCTACAGTG
Acheta_domesticus           GCGGTGGCGGTGGCGGTGGCGGTGGCGGTGGCGGTGGCGGTGGCGGTGGCGGTGGCGGGGGCGGGGGCGGGGGCGGGGGCGGGGGCGGGGGCGGGGGCGGGGGCGG---GGGCG
Gryllus_longicercus         GGGGTGGCGGCGGTGGCGGTGGCGGTGGAGGTGGCGGCGGTGGCGGTGGC---------------------------------------------------------------------
Consensus                   * ***** *  ** ** ** ** ** ** ** ** ** ** ******** ***

Teleogryllus_occipitalis    GTGGTGGAGACGAAGGTGGAGGTGGCTGCGGTGGTGGTGGAGACGAAGGTGGCGGTGGCGGGGGCGCGGGGGTGGTAGCGGGGGGGTAGGGGGGTGGTAGCGGTGGTGGCGGCGGCGATG
Acheta_domesticus           GGGGCGGGGGCGGGGGCGGGGGCGGGGGCGGCGGCGGTGGCGGCGGTGGCGGCGGGGGCGGCGG-----------------------GGGCGGCGGTGGCGGCGGTGGCGGCGGTGGCG
Gryllus_longicercus         -------------------------GGCGGCGGCGGTGGCGGCGGTGGTGGCGGTGGCGGC--------------------------GGCGGCGGTGGCGGCGGTGGCGGCGGTGGCG
Consensus                                            **** ** ***** * **  ** ***** *****                           ** ** *** **** *********** *  *

Teleogryllus_occipitalis    GAGGTGGCGGAAGTGGCGGCGGTAGCGGCGGTGGTGGCGGCGGCGTGGCGGCGGTGGCAGCGGAGGGGGCGGCGGCGGCGGCGGCGGTGGCGGCGTTAGCGGTAGCGGCCTCTCCCG
Acheta_domesticus           GCGGTGGCGGCGGTGGCGGCGGTGGCGGCGGTGGCGGC----------------------------GGTGGCGGCGGTGGCGGCGGTGGCGGCGGCGTTAGCGGTAGCGGCCTCTCCCG
Gryllus_longicercus         GCGGTGGCGGCGGTGGCGGCGGTGGCGGCGGTGGCGGT----------------------------GGCGGCGGTGGCGGTGGCGGCGACGGCGGCGTTAGCGGTAGCGGCCTCTCCCG
Consensus                   * ********  *********** ********** **                             ** ***** ** ** ***** *  ****************************

Teleogryllus_occipitalis    CCCCAAAAAATGGCGCTATTTCCGTGCAGGCGCTGAGCGAATCTCCAGAAATATCCGGCAGCGACGGCGGAGGCGGAGGTGGC-GGCTGTGGCGGCG--GACGGTGGCGGCGCCGCGGCA
Acheta_domesticus           CCCCAAAAAATGGCGCTATTTCCGTGCAGGCGCTGAGCGAATCTCCAGAAATATCCGGCAGGGACGGCGGAGGCGGTGGCAGCGAGCGGCGGCTGCGGCGGCGGCGGCGCCGCGGCA
Gryllus_longicercus         CCCCAAAAAATGGCGCTATTTCCGTGCAGGCGCTGAGCGAATCTCCAGAAATATCCGGCAGCGACGGCTGCGGCGGAGGC----GGCGGCGGCCGCG--GACGGCGGCGGCGCCGCGGCA
Consensus                   ************************************************************* ****** * ***** **      ** * *** ***  * *** ***************

Teleogryllus_occipitalis    TTCCAGGCGGTGTGTGATCGCGGGCAGCCCCAATTACGGCTCGCTGACGGCGCGCGCCGGAGGCGGCCACTTCGCGCAGGGTGATGGGCTTCTTCGCTTGAGGAGCGACGTCGTGGGTCT
Acheta_domesticus           TTCCAGGCGGTGTGTGATCGCGGGCAGCCCCAATTACGGCTCGCTGACGGCGCGCGGCGGAGGCGGCCACTTCGCTCAGGGTGATGGGCTTCTTCGCTCCAGGGGCTACATCATGGGTCT
Gryllus_longicercus         TTCCAGGCGGTGTGTGATCGCGGGCAGCCCCAATTACGGCTCGCGGACGGCGCGCGCCGGAGGCGGCCTCTCCGCGCAGGGTGATGGGCTTCTTCGCGCGAGGGGCGACGTCGGGGGCGT
Consensus                   ******************************************** *********** *********** ** *** ********************   *** ** ** **  ***  *

Teleogryllus_occipitalis    AGCTTGGAGCCCCTGCGGAACACCATTATTTTCTGGAGTTATGAATATTTTTT--------CTCTTCTGCTAATAGTAGTACAACCCAATTCGAACTTATACAAGATGTTCATGTCCTCA
Acheta_domesticus           CGCTTGGAGCCCCTGCGGGACACCATTATTTGCTGCATTTCTGAATATATTTTTCCCCTTATTCTTCTGC-AGTTGGATTCGAGCTCGTGTAGAA-TGTTGCC-----------TTGTCA
Gryllus_longicercus         CGCTTGGAGACCCTGGGGCACACCTTTATTTCTTGGGTTTCTGAATGTTCTTTTCCCCCTTTTCTTCTGCTAATGGCACTGCAGCTCAATTGGAGCTTTTGCA-----------TTTTTG
Consensus                    ******** ***** ** ***** ******  **   ** ***** *  ***         ******** * * * * *  * * *   * **  *  * *            *  *

Teleogryllus_occipitalis    CAGAACTCTCTTGAGAATTTGTAGGA----CCACATGGAGAA------------------------------------------------------------------------------
Acheta_domesticus           TAGAACTGTCTTGAGAATTTGTCGGGTGTTCCACATGGAGAATGTGTATCATCAAATCGTCTGAGAAGGTTGAAGGATTGTGCTTGGTTTATCTCCGGTCTAATGTGCTATGGCATCCGC
Gryllus_longicercus         CAGAACTAACTTAGGAATATGTGGGGTGTGCGACCTGGAGAAGGGGAATACCCAA----CCTTAGAAGGTTGAAGGATTGCGCCAGGTCCATCTTT-GTCTAACGTACTATGGCCTCTGA
Consensus                    ******  ***  **** *** **     * ** *******

Teleogryllus_occipitalis    ------------GAAGGAAACAAGCCAA---------------------------------------------------------------------------------------CAG
Acheta_domesticus           CAGGCCCTGATCGGAGAAATCGAC---------CCGTCAGAGCTGCGGGGATTTAGAATTCCAGGTAGAAAATTAGTCGGATCCAGCTATTTATCTCGTGTCGAACAACAGTAATCGTAG
Gryllus_longicercus         CAGGTGCTGATGGAAGGAAACGACGAAAACAAACCATCAGAGTTGCCGAGATTTCGAATTAT-CGTAGAAGATTCGACGGATCCAGCTATTTAGCTCATCTCGAA-----------TAG
Consensus                               * ** ** * *                                                                                           **

Teleogryllus_occipitalis    CGAGTG---------------------------------------------------------------------------------CCTTAGAACGATG-------------------
Acheta_domesticus           GGACTG--GAACCTATGATCCCAATGTATTTGAACGTTTTCTTGCTAATTCGCTACAATGTAAGCTGGACAAAAC--TTTGTTTTTTTCTGAAAACTACGCTCATTAAATATTCACGTCA
Gryllus_longicercus         GGATTGTAGAACTTATGATCACGGCGTATTTGTAGCTCTTTCTGAGGATTTGCTACAATACAGGTTACAGAAAACTGTATGTTCTTTTCTAAAAACGACGCTCATTAAATATCCATGCCA
Consensus                    ** **                                                                                 ** * *** * *

Teleogryllus_occipitalis    -------------------------------------------------------------------------AAGCACTGC------------------------------------
Acheta_domesticus           AAAAAATTAATCATGTGTATTGTCTATATGACTCACCCTTTTCAGGACTGTCGTCAACGTCATCAGTGATCATCAGTCACTTCTGAGACCAATTGCCTGATTTGAGTAGTGTGATGGACA
Gryllus_longicercus         AAAATTTCGACTATATATAATATGTTTATAAGACTCACCCAGCAGGACTGACGTTATCGTCATCAGCGACTA----TCGCTGCGGAGACCAACTCAGTGATTGGAGTAGTGTGATTGACG
Consensus                                                                                             * ** *

Teleogryllus_occipitalis    ------------------------------------------------------------------------------------------------------------------------
Acheta_domesticus           GAAAAGTGGAGATGTTACCACATATCCGCATACATATAATAACTATT-----------AGTTATTACTCTACCGAGAGTTTTAAGGACCTCCCTTTGAACACGTTGAACGTTACC-AAAA
Gryllus_longicercus         GAAAAATGGAGATATTACAACATATCAGCA---ATATAATAGTTTTTTTCTTTCAAAAAATTATAACTCTACAGAGAGTTTTCGTGACCTATCTTTAAATAC--TGAATATTTTTGAGAA
Consensus

Teleogryllus_occipitalis    ---------------------------------------------------------------
Acheta_domesticus           TTACATGTTTATAAAATTCAGGCAATACCTAATACTATCAAGTCAAAATGTGATTTTATTCAA
Gryllus_longicercus         AAATACATTGATAAAATTCAGTGAATACCGAACACTTGCAGGTCAAAATATGATTCGAATGAT
Consensus
```

**Tocc.4G0012180.1_exon1**

```
Teleogryllus_occipitalis    -------------------MA---------------------------------------------------------------------------------------------------
Acheta_domesticus           LNKITF*LDSIRYCLNFINM*FW*RSTCSKGGP*NSR*SNN**LLYVCGYVVTSPLFCPSHYSNQAI-GLRSD**SL--------MTLTTVLKRVSHIDNTHD*FF*REYLMSVVFRKKQ
Gryllus_longicercus         -----YFL*NLITLLKYIGML------------------------------------------SQTIQGVISDSTELGRFKLVSKIMLFSLLNDFEVVDGS-------SR*N*LILMKDN
Consensus                                       *

Teleogryllus_occipitalis    ------------------------------------------------------------------------------------------------------------------------
Acheta_domesticus           SFVQLTL*RISKKTFKYIGIIGSSPYDYCCSTRDK*LDPTNF--------------LPGILNPRSSDGSISPIRAWRMP*HIRPEI------------------NQAQSF--NLLRRFDD
Gryllus_longicercus         VLFFVNKPRVQEK-FRTIHLY----VLYCYFIRDFRRKTVKFGFFTSLNGNAKMIKLNKKY*PGTSYALIVP*KVLMFLI*IVKFILP*SY*LYNIATLSFRKGNH*ESFDRNLEQHY--
Consensus

Teleogryllus_occipitalis    ------------------------------------------------------------------------------------------------------------------------
Acheta_domesticus           THSPCGTPDKFSRQFYDKATFYTSSNPTAEE*GEKYIQKCSK*WCPAGAPSETHDVAPGAKKPITLSEVAASAARR-------QRAVIGAARDH------TPPGMPRRRRRRRSRRSLPP
Gryllus_longicercus         ----C-TPSPRIR----------FTNPRPSENTE*HT-------------AETFRRHR*SRR----HQITGQTARRCARTGPDERATIHAADSSAGGDIFTGRGRPRPRAVGFIRSH*DP
Consensus

Teleogryllus_occipitalis    ------------------------------------------------------------------------------------------------------------------------
Acheta_domesticus           PPPSLPDISGDSLSACTEIAPFFGAGEAATANAAATAATAATAATAATAATAATAATAATAAPAAPAATAATAAAPAPAPAPAPAPAPAPAPAPAPAPAPATATATATATATATA
Gryllus_longicercus         AR------GGER-----------GGGGGRFTKAVVAVMVVAVAVAVAVAVAVAVAVAVGVAAVAVAVEVAAVAVAAAAVAAVVAVA---------------AAAVAAVAAVAAVAAVAAV
Consensus

Teleogryllus_occipitalis    ------------------------------------------------------------------------------------------------------------------------
Acheta_domesticus           TATATATATATATATATATATTTTTTTTTTASFVSHSP----------------------LPLSPPRAGS**DRMRP-TARGRGRPVKMSPP-------------AELS--AA*IVAL--
Gryllus_longicercus         AAVAVAAVAVAATAALAVAASPAPKNGAISVQALSESPEISGSDGCGGGGGGRGRRRRRGIP------GGV*SRAAPITARGRRAPEAASPRRVMGFFARGATSGASLGDPGAHLYFLGF
Consensus

Teleogryllus_occipitalis    ------------------------------------------------------------------------------------------------------------------------
Acheta_domesticus           --------SSGPVRAQRRAVLPVI**RLLYLCL*D--VSAVCHSVFSDCLDS*ILFLAKV-CCFKFQLDSKCYSQ**IL-------------------------AVFYS**L*CRINYFT
Gryllus_longicercus         *MFFSPFSSANGTAAQ---------LELLHFCRTNLGICGVCDLEKGNTQP*KVEGLRQVHLCLTY------YGL*QVLMEGNDENKPSELPRFRIIVEDSTDPAI*LISNRDCR-----
Consensus

Teleogryllus_occipitalis    ---------------------------------------------------------------------------VLVV-----------------------------------------
Acheta_domesticus           IYKRYINAC*ETIEHGLSGYIMI*FCHMSVSIQRVKNNMDSEFCSF----------------------SPEISDKIIKVI*MNC--------------FKFYQHLRFIHKK*NIV--FHE
Gryllus_longicercus         -----------TYDHGV---------FVALSEDLLQYRLQKTVCSFLKTTLIKYPCQKFRLYIICL*DSPSRTD-VIVISDYRCGDQLSDWSSVIDGKMEILQHIS------NIIVFFFQ
Consensus

Teleogryllus_occipitalis    ------------------------------------------------------------------------
Acheta_domesticus           N*LVLATGSV*IFWFLNII*K*KYHNSGYWLGTANL*TMCYSTLYCA*GKILKFPSVE----IKFPRKSNCH
Gryllus_longicercus         KIITLQRVFVTYL*ILNIFEK---------------------NTLI----KFSEYPTLAGQNMIRM-------
Consensus
```

**Tocc.4G0012180.1_exon2**

```
Teleogryllus_occipitalis        ------------------------------------------------------------------------------------------------------------------------
Acheta_domesticus               K*QFDFLGNLISTLGNFSIFPQAQYKVL*HIVQRLAVPSQYPEL*YFHF---*MMFK--NQNIHTDPVASTN--*FS*KTIFYFL*INRRCW*NLKQFIYITLIIL--------------
Gryllus_longicercus             ----YFL*NLITLLKYIGMLSQT--------IQ--GVISDSTELGRFKLVSKIMLFSLLNDFEVVDGSSR*N*LILMKDNVLFFVNKPRV----QEKFRTIHLYVLYCYFIRDFRRKTVK
Consensus                       

Teleogryllus_occipitalis        ------------------------------------------------------------------------------------------------------------------------
Acheta_domesticus               ----SEISGEKLQNSESILFLTLWIETLIWQNQIII*PDNPCSIVS*QALMYLL*IVK*FILH*SY*L*NTARI-YH*E*HFES-N*NLKQHTFAKN---KIYESRQSENTE*HTAETSH
Gryllus_longicercus             FGFFTSLNGNA---------------KMIKLNKKY*PGTSYALIVP*KVLMFLI*IVK-FILP*SY*LYNIATLSFRKGNH*ESFDRNLEQHYCTPSPRIRFTNPRPSENTE*HTAETFR
Consensus                       

Teleogryllus_occipitalis        ----------------------------------------------------------------------------AMVVAAMVVAVV--------------------------------
Acheta_domesticus               RHR*SRRHQITGKTALRCARTGPDERATIHAADSSAGGDIFT--GRPRPRAVGLIRSH*DPARGGDSGSGECDTKEAVVVVVVVVVVAVAVAVAVAVAVAVAVAVAVAVAVAVAVAVAVA
Gryllus_longicercus             RHR*SRRHQITGQTARRCARTGPDERATIHAADSSAGGDIFTGRGRPRPRAVGFIRSH*DPARGGERGGGGGRFTKA-VVAVMVVAVAVAVAVAVAVAVAVAVGVA-AVAVAVEVA-AVA
Consensus                                                                                                   * **   ** *

Teleogryllus_occipitalis        -----------------------------EAKVA--------------------------------------------------------------------------------------
Acheta_domesticus               VAGAGAGAGAGAGAGAGAGAGAGAGAGAGAAAVAAVAAGAAGAAVAAVAAVAAVAAVAAVAAVAAVAAVAAALAVAASPAPKNGAISVQALSESPEISGRD--GGGGGSERRLRRRR
Gryllus_longicercus             VA---------------------------AAAVAAVVAVAA-AAVAAVAAVAAVAAVAAVAAVAVAAVAVAATAALAVAASPAPKNGAISVQALSESPEISGSDGCGGGGGG----RGRR
Consensus                                                      * **

Teleogryllus_occipitalis        ------------------------------------------------------------------------------------------------------------------------
Acheta_domesticus               RRRGIPGGV*SRAAPITAR*RRAAEAATSLRVMGFFAPGATSWVSLGAPAGHHYLLHF*IYFSPYSSAVG----FELV----*NVALS*NC-LENLSGVPHGECVSSNRLRRLKDCAWFI
Gryllus_longicercus             RRRGIPGGV*SRAAPITARGRRAPEAASPRRVMGFFARGATSGASLGDPGAHLYFLGF*MFFSPFSSANGTAAQLELLHFCRTNLGICGVCDLEKGNTQP----*KVEGLRQVHLC----
Consensus                       

Teleogryllus_occipitalis        ------------------------------------------------------------------------------------------------------------------------
Acheta_domesticus               SGLMCYGIRQALI---GEIDPSELRGFRIPGRKLVGSSYLSRVEQQ*S*GLEPMIPMYLN---------VFL-----LIRYNVSWTKLCFFLKTTLIKYSRQKN*SCVLSI*LTLFRTVV
Gryllus_longicercus             --LTYYGL*QVLMEGNDENKPSELPRFRI----IVEDS------------TDPAI*LISNRDCRTYDHGVFVALSEDLLQYRLQKT-VCSFLKTTLIKYPCQKFRLYIICL*DSPSRTDV
Consensus                       

Teleogryllus_occipitalis        -------------------------------------------------------------------------------------------
Acheta_domesticus               NVISDHQSLLRPIA*FE*CDGQKS-------GDVTTYPHTYN------NY*LLLYREF*GPPFEHVERYQNYMFIKFRQYLILSSQNVILF
Gryllus_longicercus             IVISDYR-----------CGDQLSDWSSVIDGKMEILQHISNIIVFFFQKIITLQRVFV-TYL*ILNIFEKNTLIKFSEYPTLAGQNMIRM
Consensus                       
```

**Tocc.4G0012180.1_exon3**

```
Teleogryllus_occipitalis      ------------------------------------------------------------------------------------------------------------------------
Acheta_domesticus             K*QFDFLGNLISTLGNFSIFPQAQYKVL*HIVQRLAVPSQYPEL*YFHF---*MMFK--NQNIHTDPVASTN--*FS*KTIFYFL*INRRCW*NLKQFIYITLIIL-------------
Gryllus_longicercus           ----YFL*NLITLLKYIGMLSQT--------IQ--GVISDSTELGRFKLVSKIMLFSLLNDFEVVDGSSR*N*LILMKDNVLFFVNKPRV----QEKFRTIHLYVLYCYFIRDFRRKTVK
Consensus

Teleogryllus_occipitalis      ------------------------------------------------------------------------------------------------------------------------
Acheta_domesticus             ----SEISGEKLQNSESILFLTLWIETLIWQNQIII*PDNPCSIVS*QALMYLL*IVK*FILH*SY*L*NTARI-YH*E*HFES-N*NLKQHTFAKN---KIYESRQSENTE*HTAETSH
Gryllus_longicercus           FGFFTSLNGNA---------------KMIKLNKKY*PGTSYALIVP*KVLMFLI*IVK-FILP*SY*LYNIATLSFRKGNH*ESFDRNLEQHYCTPSPRIRFTNPRPSENTE*HTAETFR
Consensus

Teleogryllus_occipitalis      ------------------------------------------------------------------------------------------------------------------------
Acheta_domesticus             RHR*SRRHQITGKTALRCARTGPDERATIHAADSSAGGDIFT--GRPRPRAVGLIRSH*DPARGGDSGSGECDTKEAVVVVVVVVVVAVAVAVAVAVAVAVAVAVAVAVAVAVAVAVA
Gryllus_longicercus           RHR*SRRHQITGQTARRCARTGPDERATIHAADSSAGGDIFTGRGRPRPRAVGFIRSH*DPARGGERGGGGGRFTKA-VVAVMVVAVAVAVAVAVAVAVAVAVGVA-AVAVAVEVA-AVA
Consensus

Teleogryllus_occipitalis      ----------GGGSGGGGGDG-GGGSGGGS-----------------------------------------------------------------------------GGGGG--------
Acheta_domesticus             VAGAGAGAGAGAGAGAGAGAGAGAGAGAGAAAVAAVAAGAAGAAVAAVAAVAAVAAVAAVAAVAAVAAVAAVAAALAVAASPAPKNGAISVQALSESPEISGRD---GGGGGSERRLRRR
Gryllus_longicercus           VA---------------------------AAAVAAVVAVAA-AAVAAVAAVAAVAAVAAVAAVAVAAVAVAATAALAVAASPAPKNGAISVQALSESPEISGSDGCGGGGGG-----RGR
Consensus                                                                                                                                ***** 

Teleogryllus_occipitalis      ------------------------------------------------------------------------------------------------------------------------
Acheta_domesticus             RRRRGIPGGV*SRAAPITAR*RRAAEAATSLRVMGFFAPGATSWVSLGAPAGHHYLLHF*IYFSPYSSAVG----FELV----*NVALS*NC-LENLSGVPHGECVSSNRLRRLKDCAWF
Gryllus_longicercus           RRRRGIPGGV*SRAAPITARGRRAPEAASPRRVMGFFARGATSGASLGDPGAHLYFLGF*MFFSPFSSANGTAAQLELLHFCRTNLGICGVCDLEKGNTQP----*KVEGLRQVHLC---
Consensus

Teleogryllus_occipitalis      ------------------------------------------------------------------------------------------------------------------------
Acheta_domesticus             ISGLMCYGIRQALI---GEIDPSELRGFRIPGRKLVGSSYLSRVEQQ*S*GLEPMIPMYLN---------VFL-----LIRYNVSWTKLCFFLKTTLIKYSRQKN*SCVLSI*LTLFRTV
Gryllus_longicercus           ---LTYYGL*QVLMEGNDENKPSELPRFRI----IVEDS-----------TDPAI*LISNRDCRTYDHGVFVALSEDLLQYRLQKT-VCSFLKTTLIKYPCQKFRLYIICL*DSPSRTD
Consensus

Teleogryllus_occipitalis      --------------------------------------------------------------------------------------------
Acheta_domesticus             VNVISDHQSLLRPIA*FE*CDGQKS-------GDVTTYPHTYN------NY*LLLYREF*GPPFEHVERYQNYMFIKFRQYLILSSQNVILF
Gryllus_longicercus           VIVISDYR-----------CGDQLSDWSSVIDGKMEILQHISNIIVFFFQKIITLQRVFV-TYL*ILNIFEKNTLIKFSEYPTLAGQNMIRM
Consensus
```

**Tocc.4G0012180.1_exon4**

```
Teleogryllus_occipitalis          -------------------------------------------------------------------------------------------------------------------------
Acheta_domesticus                 K*QFDFLGNLISTLGNFSIFPQAQYKVL*HIVQRLAVPSQYPEL*YFHF---*MMFK--NQNIHTDPVASTN--*FS*KTIFYFL*INRRCW*NLKQFIYITLIIL-------------
Gryllus_longicercus               ----YFL*NLITLLKYIGMLSQT--------IQ--GVISDSTELGRFKLVSKIMLFSLLNDFEVVDGSSR*N*LILMKDNVLFFVNKPRV----QEKFRTIHLYVLYCYFIRDFRRKTVK
Consensus

Teleogryllus_occipitalis          -------------------------------------------------------------------------------------------------------------------------
Acheta_domesticus                 ----SEISGEKLQNSESILFLTLWIETLIWQNQIII*PDNPCSIVS*QALMYLL*IVK*FILH*SY*L*NTARI-YH*E*HFES-N*NLKQHTFAKN---KIYESRQSENTE*HTAETSH
Gryllus_longicercus               FGFFTSLNGNA---------------KMIKLNKKY*PGTSYALIVP*KVLMFLI*IVK-FILP*SY*LYNIATLSFRKGNH*ESFDRNLEQHYCTPSPRIRFTNPRPSENTE*HTAETFR
Consensus

Teleogryllus_occipitalis          -------------------------------------------------------------------------------------------------------------------------
Acheta_domesticus                 RHR*SRRHQITGKTALRCARTGPDERATIHAADSSAGGDIFT--GRPRPRAVGLIRSH*DPARGGDSGSGECDTKEAVVVVVVVVVVAVAVAVAVAVAVAVAVAVAVAVAVAVAVAVAVA
Gryllus_longicercus               RHR*SRRHQITGQTARRCARTGPDERATIHAADSSAGGDIFTGRGRPRPRAVGFIRSH*DPARGGERGGGGGRFTKA-VVAVMVVAVAVAVAVAVAVAVAVAVGVA-AVAVAVEVA-AVA
Consensus

Teleogryllus_occipitalis          ---------------------------------------------------------------------------------------------------EISGSDGGGGGGG-CGGGRWRR
Acheta_domesticus                 VAGAGAGAGAGAGAGAGAGAGAGAGAGAGAGAAAVAAVAAGAAGAAVAAVAAVAAVAAVAAVAAVAAVAAVAAVAAALAVAASPAPKNGAISVQALSESPEISGRDGGGGGSERRLRRRRRR
Gryllus_longicercus               VA--------------------------AAAVAAVVAVAA-AAVAAVAAVAAVAAVAAVAAVAVAAVAVAATAALAVAASPAPKNGAISVQALSESPEISGSDGCGGGGG--GRGRRRR
Consensus                                                                                                                             **** ** ***        * **

Teleogryllus_occipitalis          RGIPGGV*-----------------------------------------------------------------------------------------------------------------
Acheta_domesticus                 RGIPGGV*SRAAPITAR*RRAAEAATSLRVMGFFAPGATSWVSLGAPAGHHYLLHF*IYFSPYSSAVG----FELV----*NVALS*NC-LENLSGVPHGECVSSNRLRRLKDCAWFISG
Gryllus_longicercus               RGIPGGV*SRAAPITARGRRAPEAASPRRVMGFFARGATSGASLGDPGAHLYFLGF*MFFSPFSSANGTAAQLELLHFCRTNLGICGVCDLEKGNTQP----*KVEGLRQVHLC------
Consensus                         ********

Teleogryllus_occipitalis          -------------------------------------------------------------------------------------------------------------------------
Acheta_domesticus                 LMCYGIRQALI---GEIDPSELRGFRIPGRKLVGSSYLSRVEQQ*S*GLEPMIPMYLN---------VFL-----LIRYNVSWTKLCFFLKTTLIKYSRQKN*SCVLSI*LTLFRTVVNV
Gryllus_longicercus               LTYYGL*QVLMEGNDENKPSELPRFRI----IVEDS------------TDPAI*LISNRDCRTYDHGVFVALSEDLLQYRLQKT-VCSFLKTTLIKYPCQKFRLYIICL*DSPSRTDVIV
Consensus

Teleogryllus_occipitalis          -----------------------------------------------------------------------------------------
Acheta_domesticus                 ISDHQSLLRPIA*FE*CDGQKS-------GDVTTYPHTYN------NY*LLLYREF*GPPFEHVERYQNYMFIKFRQYLILSSQNVILF
Gryllus_longicercus               ISDYR-----------CGDQLSDWSSVIDGKMEILQHISNIIVFFFQKIITLQRVFV-TYL*ILNIFEKNTLIKFSEYPTLAGQNMIRM
Consensus
```

**Tocc.4G0012190.1**

```
Teleogryllus_occipitalis      ACACATTGAAACGGAGTCACGACGTTTGCTAATTAGTGTTACGATATGCTTCATATATTGAACGCGTTTCATGTAAGAGGAGGAAATGCGAAGGCAGGATATGCGCTCCAGCTTGATGAT
Acheta_domesticus             ACTCACCAAAATGAAGTCGCGACGTGTGCTAATTAGTA-TAAAGTGTGCTTAATATATTTGATATGTTTCATGCGCGAGGAGAAAATACGAGGGTAGGATATGCGCTCCAGTTTCACGGA
Consensus                     ** **   *** * **** ****** ***********  **   * ***** *******  *   ********   ****** **** *** ** **************** ** * *

Teleogryllus_occipitalis      GCTGGGTAAAATAATTTAAATACTTAACCAATCTCTCATATCCATTTAAACACATTTGGAAAGTTCACGACATTGTACTCTCCCGGA-TTTTTATGTTTAATCTATTCTGTTCATATACT
Acheta_domesticus             GCTGGGCAAAATAA-TTGAGGAATTAACCACTGTCTCATGTCCATTTAGACAAATTTGGAATGTTCAAGTTATTGGTATCTCCCGCATTTTTTAAGTTTAATCTACTCTGCCCCTAAACT
Consensus                     ****** ******* ** *  * ******* * ****** ******** *** ******** ***** *  ****   ******* * ****** ********** ****  * ** ***

Teleogryllus_occipitalis      AATTATCACAAACTTACAATGTTGTTTAGTTAACCATACCAACACATAGTACGAATACTATATGAATTTTATAGGTATAAATTGCGCCA-------TGAAAGACTCAGTAAGGCTTAACT
Acheta_domesticus             AACTCTCCGAAAAGTTCAGCGGC-TGTAGTTTAACGTATCAACACGTA-CATAA-----ATATGAATTCTGTAGGTATTATGTGCCCCATATACACTATAACACACAGCATGGCTTATCT
Consensus                     ** * **  ***  * **  *   * ***** * * ** ****** **  *  *     ********* * ******* *  *** ***       *  ** ** *** * ****** **

Teleogryllus_occipitalis      TGAATTT--TATTATTAATAAAGAGTGTAGAGAAATTTTAATAAGTGAATCTTCGTCAAGAACAGTTTTTTTGGAATTCTCAGGAAAAGAC--TTCAAAAACTGCTGCGTAAGTAAAAGA
Acheta_domesticus             TGAATTTGATAACATTAAGAAACAGTGCAGCGAAA-TTGAATAAGTGAATCTTCTTCAAGAACAGCTTTCTTGGAATTGTAAGGAAAATACTGTTCCAAACCTGCTGCGTGAGTAAA--A
Consensus                     *******  **  ***** *** **** ** **** ** *************** ********** *** ******** * ******* **  *** *** ********* ******  *

Teleogryllus_occipitalis      CAAAAATTTTTCATTTGAATTTTCCAGGCATTTTTATTGGT---------ATCCTGGAAACTAAGTAATCTTTCTGAAATGCCAATGTCTTTGGAGAGGAAAATATTAGATTAAATCAGT
Acheta_domesticus             TGATAATTGTTCATTTGACTTTTCCAGGCATTTTTATTGGTAGACATTCCATCCTAGTAA-AAAGTAATCTTAATGAAATATGAATATTTTTGGAGGGGAAAATATCTGATAAAATCAGT
Consensus                       * **** ********* *********************         ***** * **  **********  ******    *** * ******* *********  *** ********

Teleogryllus_occipitalis      TAGGTGCACTCTCACATTCTTTTGACTTAATTAAAATTAAATTTCCATTGT-AAATAAAATTGATAAAAGCTA--TTCTTATGTGTAACTATTCTAGGTCCAAGATAACGAAAGAATATC
Acheta_domesticus             TATGTGCAGTCTTACAATGTTTTGAAATAAATAAAATTACACATTCACTGCAAAACTATTCTGATAATAGCTGACATCTTGTAGTTAACTATTCTATGCGCAAGGTAACAAAAAGATACC
Consensus                     ** ***** *** *** * ******  *** ******** *  * ** **   ***  *   ****** ****    **** *   *********** *  **** **** ***  *** *

Teleogryllus_occipitalis      TCAATAGGCCACACCTTTTAGCAGTGAACTACTTGTTGATCCCAAGGAGGCTTTCGAAAC----AGTGAGAAGAAAACTCCAACCTGAAAAAACAGA-----TGTTTTCCTAAGTACCAA
Acheta_domesticus             ACAATTTGCCTTACCTTTAAACAGTAAACCACTTGTTGATCTCATGGAAACTTGCAATACAGTGAGTGAGAAGAAAACTCGAACCTGAAACAACAGAGAATTGATTTCAATGAGTTCGGA
Consensus                      ****  ***  ****** * **** *** *********** ** ***  *** * * **    **************** ********* ******       ***   * *** *  *

Teleogryllus_occipitalis      ACATAAATGTGTCACAATAAGCTAACGAATACCAAAATGTATGACTATGATGTACTTCATCTGATCAAACGGAACAC-TAATTGTAAT--AAATTGTGATTATAGAAAGCACTTAAATAT
Acheta_domesticus             ATATAAATGTGTCATGATAAGGTAACGAAT-TAGAAATTTATGGCTAAAATGTTC---------TCATTCGAAACACATAATTGTTATCAATATTTGGCGTATAGAAATCACTTATATAT
Consensus                     * ************  ***** ********    **** **** ***  **** *         ***  ** ***** ******* **  * ***  *  ******** ****** ****

Teleogryllus_occipitalis      AATATTAATTTATGTTGGATCTAATACACAAGTCATTGGAGCGAATAATTTTTCAGCACTATATAATGCGAAATAAATCAAATAATTAAATTCTCCAGCGACGCTGATGAATATTGCGAG
Acheta_domesticus             AATATAAA------TTGAAAATAATATAGACGTCATTGCATTGAATAATTTTTCAGCCCTATATAAATCGAATTAATCACAATAATTTAATTATCCAGTGACGACAATGAACATTGCTTG
Consensus                     ***** **      *** *  ***** * * ******* *  *************** ********  **** ***    ******* **** ***** ****   ***** *****  *

Teleogryllus_occipitalis      TAAATACCCACAACAGCTGGTTTACTTCA-AAACACCTTGTTACGTCGTTGCAGCGCTAAGTGGTCTGCCAGTCTGTTGAGCGAGCGAGCAAGTGTGCTATAATGAGGCTGCGGTGCATG
Acheta_domesticus             TTAAGGCCCATAACAGCTGGTTTACTTCAGAAACATCTTGTTACGTCGTTGCAGCGCTAAGTGGTCTGGCAGTTTGTTCAGGGAGCGA--------GCTATAATGAGGCTGTGGTGCATG
Consensus                     * **  **** ****************** ***** ******************************** **** **** ** ******        *************** ********

Teleogryllus_occipitalis      CAGGCCAAGGGGGTGGCAAAACGCCAGCATCAACGGACAGGCGCCGAACTCAGCTCCCGCTACAGGTTCCGCACCGACATCGTTTTGATCTCGCTCGCCCCTCTCGCCCGTTCCC-----
Acheta_domesticus             CAGGCCAAGGGGGTGGCAAAGCGCCAGCATCAACGGACAGGCGCCGAACCCCGCTCCCGTTACAGGTACCACACCGACTTCGTTCTGATCTTGCCCGCTCCCCTGGCCCGTTCCCGTTCG
Consensus                     ******************* *************************** * ******* ******* ** ******* ***** ****** ** *** ** ** **********

Teleogryllus_occipitalis      -------------GTCCTCATTTCAGGCTGCTCCCGCTCATGCCCACGCTCACATGCCTGAGCACGGCCGCGCTGACGACTGCAATCACAACCGTGTTCACGACAGCGCTCACAACCCC
Acheta_domesticus             CATCCATAGTTCCTGTTCCCGTTTCAGCCTGCTCCCGCTCATGCCACAGCTCACGTGCCTAATCATGGCCGCGCTCACGGCCGCGCTCACGGCCGCGC---------------------
Consensus                                  ** * * ****** ****************   ****** ***** * ** ********* *** * **  ****  *** *

Teleogryllus_occipitalis      GCTGTAGATTGTAATCACGACCACGTTCACGACCGCGATCACCACCGTCGTCACGACCGCAATCACGATCGCAACCACGACTGCAATCACAATCGCCATCACGATCACAATCATGATGGC
Acheta_domesticus             -------------TCACGACCGCGCTCACGGCCGCGATCACGACTGCAATCACTTCCGCAAT---GATCGCAATGACCGCAGCAATTATTCTCGCAATGACGATAGCAATGACGACAGT
Consensus                                  ******** ** ***** ********** ** *   ****  *******   ********  **  * ***** *   **** ** *****  **** * **  *

Teleogryllus_occipitalis      GATCGCGATCATAATCGCGCGCACGACCGCGATCGCACCGGAATCATGGTCACAATCACAATCACCTCCACAATCACGACTGC----------GCTCACGACTGCGATCACCACCTCATC
Acheta_domesticus             ATTCACTA------------------------AGCTGCGGATTTTTGCCTATTTGCCTATTTTTTCCTTCAATTTAAAATGCAGTATTTTAGGTTAATTACTCTTTTTGAGGTATTAGT
Consensus                       ** * *                        **  **** *  **   *    *  * *     *  ****    * ***          * * *  ***    *       * *

Teleogryllus_occipitalis      ACGACTGCAATCACGATCACAATCACGTCTGCGATCACAATCACGATCACAATCACGATCACGATCACAATCACGATCACAGTCACAATCACGGCCGCAATCGCGATCACAATCACGTCT
Acheta_domesticus             AGTACTGTATTT---------------------------------TAAAAAACCTAGTGCT--TCAAAATGAATATGTAACTTAAAAT---------------------------TCT
Consensus                     *  **** * *                                  * * ** *    *     *** *** *  **   * * * ***                           ***

Teleogryllus_occipitalis      GCGATCACGACCGCAATCACGATCACAATCACGATCACGATCACAATCACGACCACAATCACTATCACGATCACGATCACAATCACGACTGCAATCACGATCACAATCACGTCTGCGATC
Acheta_domesticus             GCGTTTTTAATTGAAG----------------GATAAAAATAGACAAAGAGGCAAAAATCCGCAGCTTAGT---------AATCACTACGGCAATGACGATTGCAATGACGACTGCAATC
Consensus                     *** *    *  * *                 *** *  **    *    * * * ****   * *    *         ****** ** ***** *****  **** *** **** ***

Teleogryllus_occipitalis      ACGACCGCCATCACGATCACAATCACGATCACAATCACGATCACAATCACGATCACAATCACGATCACAATCACGATCACAATCACGATCACAATCACGATCACGATCACAATCACGATC
Acheta_domesticus             ACTACCGCAATGACGATATCAGTGACGACCGCAATCACTACCGCAATGACGATCGCAATGGCGACCGCGAACACGACCACGACCGCGCGCACGACCGCGACCACGATCTCGACTATGCT-
Consensus                     ** ***** ** *****  ** * **** ******* * * **** ****** ****  *** * * * ***** *** * * **  *** * * *** ******* * *  * * *

Teleogryllus_occipitalis      ACGATCACAATCACGGCCACAATCACGATCACAATCACGATCACGATCACAATCACGATCACGATCACAATCACGGCCGCAATCACGATCACAATCACGTCTGCGATCACGACAGCCATC
Acheta_domesticus             ------------------------------------------------------------------------------------------------------------------------
Consensus

Teleogryllus_occipitalis      ACGATCACAATCACGATCGAGATCACAATCGCGCGCACGACCGCAATCACGATCACAATCACCACCACAATCATGACTAAGCCCATGATTGTGTTCACGACCGCTCTTACAACCGCAATC
Acheta_domesticus             ------------------------------------------------------------------------------------------------------------------------
Consensus

Teleogryllus_occipitalis      ACGACCGCTATTCTCTCTCCGTTGGTACCTCTTGTTTCGAAATGCGTTCCAAATCTTTACGTTTAGCCTATTCCATCTACCATTCCCGGCCGCGCTCTTCCGGTCCCTTTACTGTTCTGC
Acheta_domesticus             ----CCGTTGGCATCTCGCGCGTGGCCCCTCTTCTTCCGAAATGCGTCCCAACTCTGTACTTTTAGCCTATTCCATCTACCATTCGGACCCGCTGTCCTCCGATCCCGTTCCTGTACAGC
Consensus                         *** *    **** *   ***  ****** ** ********** **** *** *** ***********************    ****  ** **** **** ** **** * **

Teleogryllus_occipitalis      CTTTGATA-TTTTCCGTTACCTCTCTCGCGACGTTGCCAACACTGCTCCCGCCCCACGCCCCTCTCCGCTCTTCAACCAGGCAAACTACTAGTAGGAACTCGACTTCATCGATTGTTTGC
Acheta_domesticus             CTTTGATACCTTTCCATTACCTCTCCCACGCCGTTCCCAACACTGCTTCCCCCCGACGCCCCTCTCCGCCCTTTCACCAGGCAAACTACTCGTAGGAACTCGACTTCATCGATTGTTTGC
Consensus                     ********  ***** ********* * ** **** *********** ** *** ************** ***  ************** *****************************

Teleogryllus_occipitalis      CCTATTTAGTCAATGCGATTT---------------------------------------------------------------------------------------------------
Acheta_domesticus             CCTATTTAGTCAATGAGATTTATTCAGTAGCTCCGTGTACTGGCGGAACAGAGTTACGAGAAGTTCCCTCTGCGGTTGCTGGAGTACTGGTTCATGTCGCTGGAGTTGGGAAGTGGGGCC
Consensus                     *************** *****

Teleogryllus_occipitalis      ------------------------------------------------------------------------------------------------------------------------
Acheta_domesticus             GAACCCGATTTATACCTGCTTGCCAGAGTATCTGAGAAAGGGGATACCAGACAATAGGGGCAGAATATCGAAACATGGTCATTTACTTTCAGCCATGGATGCTTTTGAATACACGCGTGT
Consensus

Teleogryllus_occipitalis      ------------------------------------------------------------------------------------------------------------------------
Acheta_domesticus             TATCGAAGTATGGTTGCTGTTAGTTTGAAAACAGATTTCGCTTTCAGTTTGAGTGCCCAGTAGTGATTCGTCTATTTTTTGGAAGACCAAAAAAATTATGTATCATTATATACTAAGCAT
Consensus

Teleogryllus_occipitalis      ------------------------------------------------------------------------------------------------------------------------
Acheta_domesticus             TGGCGTTAAAGAGATCTCTTCTTTTCTTCTTCTTCGACGGTTTCTACCCCTAGGATTTTTCCGTTTTCGATTTCAATATTTTTCGGTCCTTCCATTGACCTTCTTCTAGTCCTTTCTCAT
Consensus

Teleogryllus_occipitalis      ------------------------------------------------------------------------------------------------------------------------
Acheta_domesticus             CCATTACTTCCTGTATACAGGCCTTCCAGGTCAAAACAGGTCTGCCTCTTCTACGCCTTCTTTCAAGTATCCAGTTTAAAATTTCATTTGGCCATCTCTCATTTTGCATCGTCTGGACAT
Consensus

Teleogryllus_occipitalis      ------------------------------------------------------------------------------------------------------------------------
Acheta_domesticus             GTCCGTACCATTGAAGAGCTTTATTTTCTAGTCTTCTGGTTACGGTTGTTTGAACCCCAGGTCTTTCCCATATCACTTCGTTTCTTATCTTGTCCAGCCTTGTAATCTGCAAAGATCTTC
Consensus

Teleogryllus_occipitalis      ------------------------------------------------------------------------------------------------------------------------
Acheta_domesticus             TCATAAAGTCGAGCTCTACTGACCGTATTCTGTCTCTATGGCGTTTTGTTAAAGGCCTCATTTCTACTCCATATAGTATAATATTTTCTACTACACTGTGGAATATTCTTCTCTTTGTGT
Consensus

Teleogryllus_occipitalis      ------------------------------------------------------------------------------------------------------------------------
Acheta_domesticus             CTTGTCGTAACACATTATTCCATATAAGGCCACGTAGAGCTTTGATTGCATTTCTGCCCATTGCGAGTCTAGTGGTAACATCTTCATCAACTTTCCCATCGTCAGTAATTATAGTCCCCA
Consensus

Teleogryllus_occipitalis      ------------------------------------------------------------------------------------------------------------------------
Acheta_domesticus             GATATTTAAATTTTGAACAGTTAGTTATTTCGTCATCACCAATTTTTATATTACAGGTATCTCCTCCAATGCATAAATATTCGATCTGTTTAAGGTTAATTTTAAGTCCCCACTTTTCTT
Consensus

Teleogryllus_occipitalis      ------------------------------------------------------------------------------------------------------------------------
Acheta_domesticus             TTAATTTCCTAAGCATATACTCCATATCTTCCCGATCTGATGCCATAATGACCTGATCATCAGCAAAAAGTAGAGAATGTAAGCACCTATCATTTATGACAACTCCCATTTGAGAACATT
Consensus

Teleogryllus_occipitalis      ------------------------------------------------------------------------------------------------------------------------
Acheta_domesticus             TCTTGTACCAAGTTTCCAAAACTTGTTCCATATATTATATTAAAGAGGAGTGGCAATAAACTACATCCTTGTTTTAACCCTTTGTTAGTTTTAATACTTTGGGTAATTTTCCTCCCCATT
Consensus

Teleogryllus_occipitalis      ------------------------------------------------------------------------------------------------------------------------
Acheta_domesticus             TAATACACGCTTCAGTATTCTGGTACATTTTTTGAACCAATAAATCCATTTTTCTGGAACATTCAGTCTTCTCATTGCTGACCACATTTTACATCTGGGAACACTGTCATAGGCCTTATC
Consensus

Teleogryllus_occipitalis      ---------------------------------------------
Acheta_domesticus             CAGGTAAATAAATATCATATTTGTGTCTATATTGCATATTCCATTT
Consensus
```

**Tocc.4G0012190.1_exon1**

```
Teleogryllus_occipitalis      --------------------------------------MNFIG-----------------------------------------------------------------------------
Acheta_domesticus             MEYAI*TQI*YLFTWIRPMTVFPDVKCGQQ*ED*MFQKNGFIGSKNVPEY*SVY*MGRKITQSIKTNKGLKQGCSLLPLLFNIIYGTSFGNLVQEMFSNGSCHK**VLTFSTFC**SGHY
Consensus                                                             ***

Teleogryllus_occipitalis      ------------------------------------------------------------------------------------------------------------------------
Acheta_domesticus             GIRSGRYGVYA*EIKRKVGT*N*P*TDRIFMHWRRYL*YKNW**RNN*LFKI*ISGDYNY*RWES**RCYH*TRNGQKCNQSSTWPYME*CVTTRHKEKNIPQCSRKYYTIWSRNEAFNK
Consensus

Teleogryllus_occipitalis      ------------------------------------------------------------------------------------------------------------------------
Acheta_domesticus             TP*RQNTVSRARLYEKIFADYKAGQDKKRSDMGKTWGSNNRNQKTRK*SSSMVRTCPDDAK*EMAK*NFKLDT*KKA*KRQTCFDLEGLYTGSNG*ERTRRRSMEGPKNIEIENGKILGV
Consensus

Teleogryllus_occipitalis      ------------------------------------------------------------------------------------------------------------------------
Acheta_domesticus             ETVEEEEKKRSL*RQCLVYNDT*FFWSSKK*TNHYWALKLKAKSVFKLTATILR*HACIQKHPWLKVNDHVSIFCPYCLVSPFSDTLASRYKSGSAPLPNSSDMNQYSSNRRGNFS*LCS
Consensus

Teleogryllus_occipitalis      ------------------------------------------------------------------------------------------------------------------------
Acheta_domesticus             ASTRSY*INLID*IGQTIDEVEFLRVVCLVKGRRGASGGSSVGNGVGEVMERYQRLYRNGIGGQRVRMVDGIG*KYRVGTHFGRRGATREMPTEHSRDRGRGRARGRGRVRGRHCDRHCG
Consensus

Teleogryllus_occipitalis      ------------------------------------------------------------------------------------------------------------------------
Acheta_domesticus             SDCGRH*YRHCGSDCSRHCNRHCRSDY*AADFCLFVYFYPSIKNAEF*VTYSF*STRFFKIQYY*YLKKSN*PKILHFKLKEKIGK*AKIRSLVNTVVIAIVIARIIAAVIAIIAEVIAV
Consensus

Teleogryllus_occipitalis      ------------------------INCAMKDS----------------------------------------------------------------------------------------
Acheta_domesticus             VIAAVSAVVSAAVSAAVSAAMIRHVSCGMSGSRLKREQELWMRTGTGQGSGQDQNEVGVVPVTGAGFGACPLMLALCHPLGLHAPQPHYSSLPEQTARPLSAATT*QDVSEVNQLLWALT
Consensus                                                * *  *

Teleogryllus_occipitalis      ------------------------------------------------------------------------------------------------------------------------
Acheta_domesticus             SNVHCRHWIIKLL*LIRFI*G*KIIQCNDVYIIFNLYYI*VISIRQILITIMCFE*EHFSHKFLIRYLIMTHLYSELIEINSLLFQVRVFFSLTVLQVSMRSTSGLLFKGKANCGIFLLP
Consensus

Teleogryllus_occipitalis      ------------------------------------------------------------------------------------------------------------------------
Acheta_domesticus             CA*NS*LQDVSYYQNSFAVNV*FYLFQNIVRLHITDFIRYFPLQKYSYFIKITFY*DGMSTNKNAWKSQMNNYHFTHAAGLEQYFPYNSKKAVLEEDSLIQFRCTVS*CYQIQDKPCCVL
Consensus

Teleogryllus_occipitalis      ------------------------------------------------------------------------------------------------------------------------
Acheta_domesticus             *CIWGT*YLQNSYLCTC*YVKLQPLNFSES*FRGRVD*T*KMREIPIT*TFQICLNGHETVVNSSIILPSSVKLERISYPRIFSSRMKHIKYIKHTLY*LAHVATSFW*VLHRREKHCSL
Consensus

Teleogryllus_occipitalis      ------------------------------------------------------------------------------------------------------------------------
Acheta_domesticus             T*FRFPFSFNS*KQGRSII*STLLVFYSHFLCLSFVKQFVKLHEMKTQFRKN*WCYRMAQTFPY*ILFPWVFASYSELPQTLFCQLSSITAKSHFFVIFLIFRFPIGRTVTLILEY*RTQ
Consensus

Teleogryllus_occipitalis      ------------------------------------------------------------------------------------------------------------------------
Acheta_domesticus             KPSHQWQFSFKNYVPFKRKWRFSATKNAKFEAHFHRKVLSFQNTSFEVI*STYTHKYNHLYKLLVMCILKM*HSFQTNYNYVIFEKTKNN*K*TLRYTKECN*HYPLQKFEENNCSHI*E
Consensus

Teleogryllus_occipitalis      ------------------------------------------------------------------------------------------------------------------------
Acheta_domesticus             NGV*AFLKMRNFELLFT*NGCVFEKPYLTFFKAHVLVNIIM*TNRR*CKYYRREKYVKKFTFL*FLKKTNQ*SNMTFM*SDRVHLPLFLLKKVKFLKFKEKIWSRF*KNLVFIIFLNFHI
Consensus

Teleogryllus_occipitalis      ------------------------------------------------------------------------------------------------------------------------
Acheta_domesticus             LHAISHENY*NFENHLHDLYERAQYAEQILFEEKINIFNFVAGERRRKKFKFSNFSKIRSYPRKSDICHVNKCI*LYSYKKFKEKIWSCF*EKFENPRGFI*STPKF*RIVFKISELRLQ
Consensus

Teleogryllus_occipitalis      ------------------------------------------------------------------------------------------------------------------------
Acheta_domesticus             SGRSGNIHFDRNRTYGLGDSSVHRRTDGRTDGRTDGRCNVRLCQHFFCRNLSLQLL*LLFFLVVTLQFFFILVFISVFFVQCTSILYLLTLYK*ISHISLEHVLRSILSLYLINISVPNL
Consensus

Teleogryllus_occipitalis      ----
Acheta_domesticus             IH**
Consensus
```

**Tocc.4G0012190.1_exon2**

```
Teleogryllus_occipitalis                 ------------------------------------------------------------------------------------------------------------------------
Acheta_domesticus                        Y**MRLGTDILMRYRERIERKTCSKEMCEIYLYKVRRYSIEVHCTKKTEIKTKIKKN*SVTTKKKSSYSNCNERFLQKKCWQSLTLHRPSVRPSVRPSVRL*TLLSPKP*VRFLSKWIFP
Consensus

Teleogryllus_occipitalis                 ------------------------------------------------------------------------------------------------------------------------
Acheta_domesticus                        DRPDCSRSSEILKTILQNLGVLYMKPRGFSNFSQKQLQIFSLNFLYE*S*IHLLTWHMSDFRG*LRIFEKFENLNFFRRLSPATKLKILIFSSNKICSAY*ALS*RSCKWFSKF**FSWD
Consensus

Teleogryllus_occipitalis                 ------------------------------------------------------------------------------------IVITTT------------------------------
Acheta_domesticus                        IACNMWKFKKIIKTRFFQKRLQIFSLNFKNLTFLSRNSGKCTLSDYIKVILLYWLVFFKNYRNVNFFTYFSRR*YLH*RRFVHIIIFTSTCALKNVK*GFSKTQPFYVKSSSKFRIFRNA
Consensus                                                                                                    *  * *

Teleogryllus_occipitalis                 ------------------------------------------------------------------------------------------------------------------------
Acheta_domesticus                        QTPFSHIWLQLFSSNFCNG*C*LHSLV*RKVYFQLFFVFSKIT*L*FV*KLCHIFNMHITSNL*R*LYL*VYVLQITSNEVF*KLKTFL*K*ASNFAFLVALKRHFLLNGT*FLKENCH*
Consensus

Teleogryllus_occipitalis                 ------------------------------------------------------------------------------------------------------------------------
Acheta_domesticus                        *LGFCVL*YSKINVTVLPIGNRKIKKMTKK*LLAVIDDSWQKRVCGNSE*LAKTQGNKI*YGKVCAIL*HH*FFLNWVFIS*SFTNCLTKERHKKCE*KTSRVL*IIDLPCFYELKENGN
Consensus

Teleogryllus_occipitalis                 ------------------------------------------------------------------------------------------------------------------------
Acheta_domesticus                        RN*VKLQCFSLLCKTHQNEVATCAN*YKVCLIYLICFMREEKIRG*DMRSSFTELGKIIEELTTVSCPFRQIWNVQVIGISRIF*V*STLPLN*LSEKFSGCSLTYQHVHKYEFCRYYVP
Consensus

Teleogryllus_occipitalis                 ------------------------------------------------------------------------------------------------------------------------
Acheta_domesticus                        HIHYNTQHGLS*I**H*ETVQRN*ISESSSRTAFLEL*GKYCSKPAA*VK**LFI*LFQAFLLVDIPS**KVILMKYEYFWRGKYLIKSVMCSLTMF*NK*NYTFTAKLF***LTSCS*L
Consensus

Teleogryllus_occipitalis                 ------------------------------------------------------------------------------------------------------------------------
Acheta_domesticus                        FYAQGNKKIPQFALPLNSKPLVDLMETCNTVSEKKTRT*NNRELISMSSEYKCVMIR*RIRNLWLKCSHSKHIIVINIWRIEITYI*YKLKII*TSLH*IIFQPYINRINHNNLIIQ*RQ
Consensus

Teleogryllus_occipitalis                 ----------------------------------------------------------------------------------------------------------FTTAITTVVTTAIT
Acheta_domesticus                        *TLLVKAHNSWFTSETSCYVVAALSGLAVCSGSEL**GCGACRPRGWQSASINGQAPNPAPVTGTTPTSF*SCPLPWPVPVRIHSSCSRFSLLPLMPQLTCLIMAALTAALTAALTTALT
Consensus                                                                                                                       * * *   *** *

Teleogryllus_occipitalis                 IATTTAITIAITITIMMAIAIIIARTTAIA------------------------------------------------------------------------------------------
Acheta_domesticus                        AAITTAITSAM---IAMTAAIILAMTIAMTTVFTKLRIFAYLPIFSFNLKCSILG*LLFLRY**YCILKNLVLQNEYVT*NSAFLIEG*K*TKRQKSAA**SLRQ*RLQ*RLQSLPQ*RY
Consensus                                 * ***** *    * *  *** * * *

Teleogryllus_occipitalis                 ---PESWSQ--SQSPPQSRLRSRLRSPPHH---------------DC---------------------------------------------------------NHDHNHVC--------
Acheta_domesticus                        Q*RPQSLPQ*RSQWRPRTRPRPRARPRPRSRLCSVGISRVAPLLPKCVPTLYF*PIPSTIRTRCPPIPFLYSL*YLSITSPTPFPTLLPPDAPLRPFTRQTTRRNSTSSIVCPI*SMRFI
Consensus                                   * *  *  **  *  * * * *  *                  *                                                         *     **

Teleogryllus_occipitalis                 ----------------------------------------------DHNHDHNHDHDHNHDHSH----------------------------------------------NHGRNRD---
Acheta_domesticus                        Q*LRVLAEQSYEKFPLRLLEYWFMSLELGSGAEPDLYLLARVSEKGDTRQ*GQNIETWSFTFSHGCF*IHACYRSMVAVSLKTDFAFSLSAQ**FVYFLEDQKNYVSLYTKHWR*RDLFF
Consensus                                                                      *               **                                               * * **

Teleogryllus_occipitalis                 ------------------------------------------HNHVC-----------------DHDRNHDHN-----------------------------------------------
Acheta_domesticus                        SSSSTVSTPRIFPFSISIFFGPSIDLLLVLSHPLLPVYRPSRSKQVCLFYAFFQVSSLKFHLAISHFASSGHVRTIEELYFLVFWLRLFEPQVFPISLRFLSCPAL*SAKIFS*SRALLT
Consensus                                                                     **                  *     *

Teleogryllus_occipitalis                 ---------------HDHDHNHDH-----------NHYHDHDHN----HDC----NHDHNHVCDHD----------------RHHDHNH-------------------------------
Acheta_domesticus                        VFCLYGVLLKASFLLHIV*YFLLHCGIFFSLCLVVTHYSI*GHVEL*LHFCPLRV*W*HLHQLSHRQ*L*SPDI*ILNS*LFRHHQFLYYRYLLQCINIRSV*G*F*VPTFLLIS*AYTP
Consensus                                                *       *            **    *     * *       * *   *                 ***

Teleogryllus_occipitalis                 ----------DHNHD-HNHDHNH-------------------------DHNHDH-------------------------------------NHDHNHDHDHNHDHDHNHGHNHDHNHDHD
Acheta_domesticus                        YLPDLMP**PDHQQKVENVSTYHL*QLPFENISCTKFPKLVPYIILKRSGNKLHPCFNPLLVLILWVIFLPI*YTLQYSGTFFEPINPFFWNIQSSHC*PHFTSGNTVIGLIQVNKY---
Consensus                                           **     *    *                           *  *                                     *    *   *        *

Teleogryllus_occipitalis                 HNHDHDHNHGRNHDHNHVCDHDSHHDHNHDRDHNRAHDRNHDHNHHHNHD*
Acheta_domesticus                        ----------------HICYYIAYSI-------------------------
Consensus                                                * *
```

**Tocc.5G0008020.1**

```
Teleogryllus_occipitalis  TGTATATTTATAATATAATTTAATAAATATTCTC--------------------------------------------------------------------------------------
Acheta_domesticus         TGCCCGTTTGTTCGTTTACTGGTCCGGCATTCGCCTGCTGTTGATCGCAGAAGCTGCGGAAAACTACTGATCAGAGGAGCCGGCCGCTATAA---TGCTCCAGTCAGTGGCTGAAGTCGT
Gryllus_longicercus       ---------GTTAGGGTACTGTTCCGACCCTAAT-----------------------GGAAAACTACTGACCATAGCAACCGGCGGTTAGAATTGTTCTCTAGACA--GACCGAATTCGA
Consensus                           *      * *          *

Teleogryllus_occipitalis  ----------------------------------------------------------------------GAGGTACCGGAACGCC--CTACCGACGCCTTCCGCCAT------------
Acheta_domesticus         CCTGCCTATGTGGTTAGTGTTGAAAATCGTGCCACA--GTCGGGACTCGAACCCTAAGACCACAAGGTCAGAGGCACTGCAGTGACATCAGTTGATGCCCTGAGCCACCT----------
Gryllus_longicercus       CCTAGCGTTGTTACTAGTAACGAAAATCCTGCCGCGCCGTCGAGGCTCGAACCTC-GGACCTCGCAGTCAGAGACATCGCAGTGATATCAGTTAATGCCCTGAGCCACCTCTTCCGGTTA
Consensus                                                                                 ***  *  * *  *    *     * *** *  ****

Teleogryllus_occipitalis  --------------------------------------------------AAATCAAGCCCTGC--------------------------------------------------------
Acheta_domesticus         -------------------------------GAGTGTGAGCGAGGCAAAGAAAGGAAGCGGTGCGGC-----------------------------------------------------
Gryllus_longicercus       CTTTAAACAAATTAAATTAAAAATTCAATTGAATTATAAGTGAAGAGCA-ACACCCAATGACGCAGCAACGGGTTCTGCGATGCACCGGCCGCCGAGCCAGGTCGGCCGCCGCCCGAGAC
Consensus                                                                   * *   *     **

Teleogryllus_occipitalis  ------------------------------------------------------------------------------------------------------------------------
Acheta_domesticus         -------------------------------------------------------------------------------------------------CTGGCAACTGTTCA---------
Gryllus_longicercus       AGGTGCGCGACGAAGGCCGAACGCGTCGCGTAGCTCTGGGCTCTGCGTCGTTACGAAATGCTTATATCCCACCACGCAGCACGCGATATCTGCGCTGCTGGCGTCCGCCCACGCCGGTGA
Consensus

Teleogryllus_occipitalis  ------------------------------------------------------------------------------------------------------------------------
Acheta_domesticus         ----AATGCAGGCTGCAGTGGAAGTTACGGCAATGTAGC---------------------------------------------------------------------------------
Gryllus_longicercus       CTACGGCGACGGCTGCAATCGGGCGAAAGGTGATGGAACACACCAGGCGAACATTGATTGATTGATTAAGATTACGGGTGCAACGGCTGCTACAGTCATTATTCTCGTAAGTTTAGTGTC
Consensus

Teleogryllus_occipitalis  ------------------------------------------------------------------------------------------------------------------------
Acheta_domesticus         ------------------------------------------------------------------------------------------------------------------------
Gryllus_longicercus       TGTTTTTATAGCGGGAGATGGCAGTAACCTTCCCCCGCTGGATGGTGAGATCTGTCGCCAACCTACGTCGGCTTTGGGGGATTTTCCCTAACCAGGTATCTCCCAACAGCAGGGCACCGC
Consensus

Teleogryllus_occipitalis  ------------------------------------------------------------------------------------------------------------------------
Acheta_domesticus         ---------------------------------------------AGAGATGGAGGAGACTTGGAGGT----------------------------------------------------
Gryllus_longicercus       TCCAATCCGGACAAGTTCTAAACTACCAACTGATTTCGCGGCTCGAGAGGTAGAGTAGTGTTGTAAGTTCAAGAGCGGGTAAGAAAAGATGAGGATATGACAAGGAAGAGGAAGGAAATG
Consensus

Teleogryllus_occipitalis  ------------------------------------------------------------------------------------------------------------------------
Acheta_domesticus         -----------------------------AGAAAACTTGCGAAGAGCAAAAAAACAGACGATGAATAAG---------------------------------------------------
Gryllus_longicercus       TTGCGCGCGGAGAAACGGCAGAAAAACGGAGAAAAACGGCAGAAATCGGAGAAACGGCAGAAAAATCCGCTCAGAGTAGCAAGCCGCTGGAATTCTTCTCCAGACAGCGGCTGAACACGG
Consensus

Teleogryllus_occipitalis  ---------------------------------------------------------------------GTAGTTTAGAGTGATGTTGTACGATGCCT----------------------
Acheta_domesticus         -------------------------------------------------------------------AAGTAGATTAGAGAGATGTGTTGCGATGTCT----------------------
Gryllus_longicercus       CCTAGCTATCTAGTTTGTGTCAAAAGTTCTGACACCGTCGGGACTCGAACCTTAGGACCTCACGGTCAGATACTCTGTAGTGACACCAGTTGATGCCCTGAGCCACCTAAGCCGGCACGA
Consensus                                                                                 **   *  ** **        **** *

Teleogryllus_occipitalis  -------------------------GACGTAGGTTCTACGCGAATGAGAAGAGCAGAATTTAAGAATAAAGAACAACCAAAACAAAATAAAGAACGGATTATTTTCTAGAAAAC------
Acheta_domesticus         -------------------------GATGTAGGTTCTAGGCGAAAAAGAAGAACAGAATTTAAGAATAAAGAAGAACCTAAACACAATCAACAAAGGATTATTTTCTGGAAAACCTGTAG
Gryllus_longicercus       AGAACGGAAGCGGACCGACCTGGCGACTGTAGGTTCTA-GCAAAAGAGAAGAGCAGAATATAACATTTAAGGAGAACCTAAACACAGTAAACAAAGTATTATTTTCTGGAAAACCAATAG
Consensus                                               ********** ** **  ****** ****** *** * * *** * **** ***** * * ** ** * ********** ******

Teleogryllus_occipitalis  -------AAATGAGTCGTGCCGTGAAAAAA---GAGAACCAAAGATTCTATCAGGTGGACATGATTGAA-----------ACAGATATATAGTT---GAAAAAAACAAGGTAATAGTGAC
Acheta_domesticus         TTTTAGTGAAGCAGTTGGGTCGTGAAAAAAA-TGAGGACACAAGATTATATCAGAAAGATATAATTAAAACTGATACACTACAGATACATAGTT--CGAGAAAAACAAGGAGATGGTGCC
Gryllus_longicercus       -TTTAGCGAAGAAGTTGAGCCGTGAAAAAAAATGAGGACACAAGATTACATCAGGAAGACATGATGCAA-----------ACACATATATAGTTGAAAAAAAAAACAAGGAAAATGTGAC
Consensus                         **  *** * * **********   *** **  ******  *****   ** ** **  **           *** *** ******    * **********  *  *** *

Teleogryllus_occipitalis  AAATGAAAACGGACACGCTGTGTAAAGATTGAGAGGAATGTGAAAGTCAGACAAATAAAATGTTGAAAAGTTCTGAGGTATGGCAAAGAGAAGGAACAAATATAAAAACAAGGAGGAAAC
Acheta_domesticus         AAATGGAAACAGAAACGCTGTGTAAAGAAAGAGAGAAACGTGAAAGTTAACCAAATAAAATGTTAAAAATTTCTAGAGTATGGGAGCGAGAAGTGACAAAAATCAAAGCAAGCAG--AAT
Gryllus_longicercus       A-----ACGCAGACACGCTATATAAAGAATGAGAGGAATGTGAAAGTTAGCCAAATGAATTGTTGAATAGTTCTAGGGGATGGTAGCGAGAAGCGACAA-----AAAACAAAAAGTGGAA
Consensus                 *     *  * ** ***** * ******  ***** ** ******** *  ***** ** **** ** * ****   * **** *  ******  ****     *** ***  **   *

Teleogryllus_occipitalis  ACAAGTACAAGCGAAGCGCAAGGAGGAAACACGTTAGAG---------------------------------------------------------------------------------
Acheta_domesticus         GAAGGTGCAAGCGAAGTGCACGGAGGCATCGCGTTAGGGCAGGGCTTTCCAACTGGCGGCCCGCGGGCCTCATGCGGCCCGCCGGATCTTTTGTGCGGCCCGCCACTGCTTTGAACAATT
Gryllus_longicercus       GTAGGTATCAGCGAAAAGCACGGAGGAAACGTGTTAGAA---------------------------------------------------------------------------------
Consensus                   * **   ******  *** ***** * *  *****

Teleogryllus_occipitalis  ----------------------------------------------------------------------------------------AAATTATAACTAGACA----------------
Acheta_domesticus         TTATTGTACCAACGAAATACAATGTTCATTCAGTTATTCTATTACGCATTTTTATCATTGTATAAGTAATTATAAATGTGAAATATGTGAATTATAAATGGAAAATATGAAAAATTGATT
Gryllus_longicercus       ----------------------------------------------------------------------------------------GAAATGTAAG----------------------
Consensus                                                                                          ** * ***

Teleogryllus_occipitalis  -----------------------------------------------------------------------------------------------CGAAACCAAGGCGGA----------
Acheta_domesticus         ATAAATATGAAATATTTGAATTATAAATGAAAAATATGACCAATTAATTATAAATGTCAAATATGTGCATTAAAAACATTATGTTGAATATAAGTCGAATCTAATATGTATTTGCTTCAT
Gryllus_longicercus       -----------------------------------------------------------------------------------------------CGAAACCAAGGCGGA----------
Consensus                                                                                                **** * **   * *

Teleogryllus_occipitalis  ------------------------------------------------------------------------------------------------------------------------
Acheta_domesticus         TATAATGTCAGTAGTTTGGGTAAAAACGATTTAAAATGACGGTGCGGCCAGCGGCTTAACTAGAATGTGATATTTGGCCTATAAGCCCTTCTAAGTTACAGAGTCCTGCGTTAGAGGAAT
Gryllus_longicercus       ------------------------------------------------------------------------------------------------------------------------
Consensus

Teleogryllus_occipitalis  ----------------GACAAGAAAATCACGGTG---------AGATTGGAAGAGCCTAAAGCTTCAAGTCCAA-------GAGGTGGGAGGCGGCGACTGGAATAAAACGGGGCCGGCG
Acheta_domesticus         TATGAGTAAACGAAACGGCGAGAAAATCACGGTAAGGTAAGCCAGACCGAAGGAGCGCGAAGCAGCAAGTCCGAAAGGCGGGAGGTGGGAGGAGGCAACTGGAAGGAAACGGGGCCAGCG
Gryllus_longicercus       ----------------GACGAGAAAATCATGGTGAGATAAGCCAAACTGAAGGAGCACGAAGCTGCAAGTCCCA-------AAGGTGGGAGGCA--AGCCGGAATAAAACGGGGCCAGCG
Consensus                                  * * ********* ***          * *  * * ****   ****  ******* *        **********      * ****  ********** ***

Teleogryllus_occipitalis  GGAGGACAATACGCACGCGAACAGAGTGGCAAGAGCAAGAGCGGGCATTCCTGCAGAC--GCGCGGGCGGGAGCGCGGGCGGCGACATAACGGGCGCATCCTTGCG--------------
Acheta_domesticus         GGAGGACAATGCGCAGGCGCGCAGATTGGCAAGAGC----GCGGGCATTCCTGCGGA--------------------GGCGGCGACGTAACGGGCGCATCTCCGC---------------
Gryllus_longicercus       GGATGACAATACGCACGCGAACAGATTGGCAAGAGC----GCGGGCATTCCTGCGGAGGCGCGCGGGCGGGAGCGCAGACAGCGACATAACGGGCGCATCTCTGCGGCGGAGACAAGCGG
Consensus                 *** ****** **** ***  **** **********    ************** **                    * * ***** *************   **

Teleogryllus_occipitalis  ----GCGGAGGCGAGCGGAGGCGGCGAAACAGCGCGACTTCACGCACGCATCTGCATGCGCATCTGCATACCCGTGCGCCTGCACGGACGCAGTCGGGGGTCCGCAAGCCTCCG------
Acheta_domesticus         -------------AGCGGAGGCTGCGAAACCGCGCCGCTTCACGCGCGCATCTGCATGCGCATCTGCATACCCGTGCACCCGCACGCACGCGGTCGGGGGTCCGGCGGCCGCCGCGCCGC
Gryllus_longicercus       GGCGGCGGAGGCGAGCGGAGGCGGCGAAACAGTACGGCTTCACGCACGCATCTGCATGCGCATCTGCATACCCGTGCGCCTGCACGTACACAGTCGGGGGTCCGGCGGCCGCCGCGCCGC
Consensus                              ********* ******* *  *  ******** ******************************* ** ***** ** * ************   *** ***

Teleogryllus_occipitalis  --------------------------------------------CCACCACCTCCATCTTCCTCCAAGACCACGCCTCAGCCTCCGCCATCCTCGCCGTCTCTGGCATCGCCATCGCATC
Acheta_domesticus         CTCCGTCACCACCAGCGCCACCACCACCACCACCACCACCACCACCACCACCATCTTCCCCAAAGA-----CCTCAGCCTCCGCCGTCGCAATAGCCTTCGCCACCGCCACCGCCTC
Gryllus_longicercus       CTCAAGCTCAGG--------------------------------CCACCACCACCGTCTTCCTTTAACACCCTGCCTCA-----------------------------------------
Consensus                                                             ******** ** ******   ** *     *****

Teleogryllus_occipitalis  TAGAATCGCCATCGCCTGCGCCATCCCCATCGCCTCTGGCGGCGCCATGGTCCTCTGGCATCGCCATCGCCTCTGGCTTTGCCATCGCCTCTGGCTTTGCCATCGCCTCTGGCTTTGCCA
Acheta_domesticus         CGCCTCCGTCAAAGCCTCTGCTTCCGCCAACGCCTCTGCCTCCGCCAACGCCT-------CCACCACCGCCTCCGCCTCTGCCACCGCCTCCGCCATCGCCACCGCCTCCGCCATCGCCA
Gryllus_longicercus       --------------------------CCAACGCCTCTGCCACCGACAACCTTCTGC-------------CCCCCACATCCTCCTCCATCGATGCCGACGCCGACGCCTCCGCCTTCGCCA
Consensus                                           *** ******** *  ** **                      ** *    *   **  *  *   * *   ***  ****** * * * ****

Teleogryllus_occipitalis  TCGCCTCTGGCTTTGCCATCGCCTCTGGCTTCGCCATCGCCTCTAGCTTCGCCATCGCCTCTGGCTTCGCCATCGCCTCTAGCTTCGCCATCGCCTCTAGCTTCGCCATCGCCTCTAGCT
Acheta_domesticus         CCGCCTCCGCCAACGCCTCCGCCAACGCCTCCGCCAACGCCTC------CGCCAACGCCTCCGCCAACGCCTCCGCCAACGCCTCCGCCAACGCCTCTGCCTCCGCCTAGGCTTCTGCCT
Gryllus_longicercus       ACGCC-------ACGCCACCG----------------------------CGCCAACGCCCCCGCCAACGCC-------CCGCCTCTGCAAACGCCCCCGCCTCCGGCAACGCCTCCGCCT
Consensus                  ****         ***  **                            ***** **** * * *  ****           **  ** * **** *   ** ** *   ** **   **

Teleogryllus_occipitalis  TCGCCATCGCCTCTAGCTTCGCCATCGCCT------CTAGCTTCGCCATCGCCTCTAGCTTCGCCATCGCCTC------TAGCTTCGCCATCGCCTCT------AGCTTCGCCATCGCCT
Acheta_domesticus         CCGCCAACGCCTCTGCCTCCGCCACCCCTTCCGTTACTACCTCTGTCATCGCCTCCACCTTCGCCAACGCCTCCTCCAACGCCTCCGCCAACGCCTCTGCCAACGCCTCCGCCAACGCCT
Gryllus_longicercus       CCGGCAACGCCTC------------------------------------CGCCTCCGCCTCCGCCAGCGCATCCGTCA-GCGCTCCGTCATCGGCTAT-----CACCTCCCCCATTCCTT
Consensus                  ** ** ******                                    ******   ** ***** *** **         **  ** ** ** ** *       ** * ***   * *

Teleogryllus_occipitalis  CTAGCTTCGCCATCGCCTCTAGCTTCGCCATCGCCTCTAGCTTCGCCATCGCCTCTAGCTTCGCCATCGCCTCTGGCATCCCCTT------CGCCTCCGCCATCGCCTCCGCCCACCCCT
Acheta_domesticus         CCGCCAACGCTTTCGCCAACACCTCCGCCAACGCCTCCGCCAACGCCTCCGCCAACGCCTCCGCCAACGCCTCCGCCAACGCCTCCGCCAACGCCTCCGCCAACGCCTCCGCCAACGCCT
Gryllus_longicercus       CCACCTACTGCTCCGCCAATTCCAATGCATCCACCT--------GCCCCCTCTTTCTCTTTCTTAATTTCTCTTTCTCTCTCCTTCTCTCGTTCACTCTCTCCCTCTCTCGTTCACTCTC
Consensus                 *   *  *     ****     *   **   * ***        ***  * *       * *   *   *         * ***         *   * *   * *   **   ** *

Teleogryllus_occipitalis  C------CGCCTTCGTGAACGCCTCCGGC--------------TAGCTTTGCCATCGCCTCTAGCTTCACCATCGCCTCTAGTTTTGCCATCGCCTCTGGCATTGCC-------------
Acheta_domesticus         CCGCCAACGCCTCCGCCAACGCCTCCGCCAACGCCTCTGCCAACGCCTCTGCCAACGCCTCTGCCTCCGCCTAGGTCTCTGCCTCCGCCAACGCCTCTGCCTCCGCCACCCCTTCCGTTA
Gryllus_longicercus       TCC----CTCTCTCGTTCACTCTCTC-----------------CCTCTCTCGTTCACTCTCTCCCTCTCTCTTTTTCTCTCCCTCTCTCTTTTTCTCTCCCTCTCTC-------------
Consensus                        * *   **   ** *   *                    ** *        ****  **    *     ****   *    *     ****  *     *

Teleogryllus_occipitalis  ----------CATCCCCTCTGGCATCCCCTTCGCCTCCGCCATCGCCTCCGCCCA---------------------CTCCTCCGCCTTCGCGAACGTCTCGG------------------
Acheta_domesticus         CTGCCTCTGTCACCGCCTCCGCCAACGCCTCTGCCTCCGCCAACGCTTCCACCAATGCATCCGCAAATGCCTCTGCCTCCTCCGCCTTCGCCATCGCCCTGACTCCACCATCGCCATCGC
Gryllus_longicercus       -----TTTTTCTCTCCCTCTCTCTTTTTCTCTCCCTCTCTCTTTCTCTCTTCCTCT--------------------CTCTTTCTCTCTCTCCCTCCCCCTAT------------------
Consensus                           *    ****   *     **   ****   *      **  **                       *** * * *  ** *   *  *

Teleogryllus_occipitalis  --------------------------------GCTCCGCCATTGCCCAAGTGTCCACTTCCTCTCTCTCCATCTCCACTGCCTC-----CACCGTTAGTACCACTATTACCAGCACCTGC
Acheta_domesticus         CTTCCCCATCCTCTCCGCCACCGTCTCCACCCACACCGCCGTTGCCAATGCATCCACCT-TTCTCTCTCCAATTCCATCGCCCCATCGTTACCACCACCTCCACCACCACCATCACCTTC
Gryllus_longicercus       ------------------------------------------------------------ATCTCTATCTATCTC--------------CACCGATACCACCACCATTACCATCACCTTC
Consensus                                                                              ***** ** *  **               ***   *   **** *  **** ***** *

Teleogryllus_occipitalis  GCCGCCGCTGTCTGCTCCGCCGCCTTCTCGGAAACCCAGGTCCGTTTCGCGCGAGGATGATGAAGCGTCCC-------------------------------------------------
Acheta_domesticus         GCCGGCGCTGTCTGCTGCGCCGCCTTCTCGGAAACCCAGCTCCGTTTCGCGCGAGGATGATGGAGCATCCGGCTGCGCTGGCAGACGGGCGCGCGGGCAGTCGGCTTGTCGCGGGCGCCG
Gryllus_longicercus       GCCGCCACGGTCTGCCCAGCCGCCTTCTTGGAAACCCAGCTCCGTTTCGCGCGAGGATGATGAAGCATCCCGCTGCACTGGCAGACGGGCGC----GCAGTCTGCTTTTCGCGA-----G
Consensus                 **** * * ******   ********** ********** ********************** *** ***

Teleogryllus_occipitalis  ------------------------------------------------------------------------------------------------------------------------
Acheta_domesticus         CGCCGGGCTTTCCAGAAATCTTAGCCGCAAAGGCGAGCGATATCGGTGCGAAATAAAACACGAAATAGCGGAGTGGCCGACGGGTTTGGCAACGGCTCGGCACACGAGGCCGTGTAAGGA
Gryllus_longicercus       TGCCGGGATTTCCAGAAATCTTAGCCGCAAAGGCGAGCGATATCGGTGCGAAATAAAACACGAAATAGCGGAGTGGCCGACGAGTTTGGCAATGACTCGGCACACGAGGCCGTGTAAGGA
Consensus

Teleogryllus_occipitalis  ------------------------------------------------------------------------------------------------------------------------
Acheta_domesticus         GTGCGCGCTGGAACAAGTTATTTTCAATATTTCGTGTTCTCTGGCTGCAGGAGGACGCTGTGAAAAATTCTTATTACAGCGGTTAACAGATTTTTAAGAGGGGAAAGTATAAACGAATGG
Gryllus_longicercus       GTGCGCGCTGGAACAAGTTATTTTCAATATTTCGTGTCGTCTGGCAGCAGAAGGACGCTGTGAAAAATTCTTATTACAACGGTCAACAGATTTTTAAGAGGGGAATGTATAAACCAATGG
Consensus

Teleogryllus_occipitalis  ------------------------------------------------------------------------------------------------------------------------
Acheta_domesticus         TAATGTACAATGTGGGA-------------------------------------------------------------------------------------------------------
Gryllus_longicercus       TAATGTACAATGTGGGACGAATGTGTTTGCAAATTAGGCTTTCCATGCAGACAAAATATGCAAAATGTTTCCAAGCTCGTTTAGTCACAAATATGTTAATTTGGAACGTGTATTACTGTT
Consensus

Teleogryllus_occipitalis  ------------------------------------------------------------------------------------------------------------------------
Acheta_domesticus         ------------------------------------------------------------------------------------------------------------------------
Gryllus_longicercus       TTTATGGATACTAATTTGAGTTTCAGAAATGCCTGTCCAGGAACTGACAAATAATTTCCTCATAGAATGTGATCTATGAATCAGTGTATAGTTCATTGCACTTGTTGCATCTTTGATGGG
Consensus

Teleogryllus_occipitalis  ------------------------------------------------------------------------------------------------------------------------
Acheta_domesticus         ------------------------------------------------------------------------------------------------------------------------
Gryllus_longicercus       CTCTCATGATAGTCTCAATAATCTGCTCAGTATAGGACCCTTTGCAAGAGATCTTCAGTTATACCTAAAGCACTAATGATTATTCATACAACATAACATATTGATTCATGTAGCTAAGAC
Consensus

Teleogryllus_occipitalis  --
Acheta_domesticus         --
Gryllus_longicercus       AG
Consensus
```

**Tocc.5G0008020.1_exon1**

```
Teleogryllus_occipitalis    ----------------------------------------------------------------------------------------------------------------------
Acheta_domesticus           --------------------------------------RSRKRKKRS-----------RAGLCPFVRLLVRH----------------SPAVDRRS--------CGKLLIRGAGRYN
Gryllus_longicercus         CLSYMNQYVMLYE*SLVL*V*LKISCKGSYTEQIIETIMRAHQRCNKCNELYTDS*ITFYEEIICQF---LDRHF*NSN*YP*KQ*YTFQINIFVTKRAWKHFAYFVCMESLI-------
Consensus

Teleogryllus_occipitalis    ----------------------------------------------------------------------------------------------------------------------
Acheta_domesticus           APVSG*SRPAYVVS-----------------------------VENRATVGTRTLRPQGQRHC--------------------SDIS*CPEPPECERGKERKR-------CGLATVQM
Gryllus_longicercus         ---CKHIRPTLYITIGLYIPLLKIC*PL**EFFTASFCCQTTRNIENNLFQRALLTRPRVPSHCQTRRPLRYFVFYFAPISLAFAAKISGNPG----TREKQTARPSASAAGCFIILARN
Consensus

Teleogryllus_occipitalis    ---------------------------------------------------------------------------------------------------------MKTDTL--
Acheta_domesticus           QAAVEVTAM*QRWRR--------------LGGRKLAKSKKTDDE*EVD*RDVLRCLM*VLGEKEEQNLRIKKNLNTINKGLFSGKPVVLVKQLGRE---KNEDTRLYQKDIIKTDTLQI
Gryllus_longicercus         GAGFPRRRLGRPWRRR*W*WWWYRWR*IEI*GEGERERERKRERERERKR-----------ERERKRERERK----------------------RERERKRERERVNER--GRE*TREG
Consensus                                                                                                                            *

Teleogryllus_occipitalis    ----------------------------------------------------------------------------------------------------------------------
Acheta_domesticus           HSSRKTRRWCQMETETLCKEREKRES*PNKMLKISRVWEREVTKIKASRMKVQAKCTEASR*GRAFQLAARGPHAARRIFCAARHCFEQFYCTNEIQCSFSYSITHFYHCISNYKCEICE
Gryllus_longicercus         ESER--------------ERE------------RVNERRREKEKLRKRK------------------RGQVDA--------------------------------------LELAE
Consensus

Teleogryllus_occipitalis    ----------------------------------------------------------------------------------------------------------------------
Acheta_domesticus           L*MENMKN*L*I*NI*IINEKYDQLIINVKYVH*KHYVEYKSNLICICFIIMSVVWVKTI*NDGAASGLTRM*YLAYKPF*VTESCVRGIMSKRNGEKITVR*ARPKEREAASPKGGRWE
Gryllus_longicercus         Q*VEGMG-------------------------------------------------EVIADDGA---------------LTDA-----LAEAEAEALPEAEALPE--AGAFAEAGRWR
Consensus

Teleogryllus_occipitalis    ----------------------------------------------------------------------------------------------------------------------
Acheta_domesticus           EATGRKRG------------QREDNAQARRLARARAFLRRRRRNGRISAAEAAKPRRFTRASACASAYPCTRT------HAVGGPAAAAPPPSPPAPPPPPPPPPPP----PPSSPKTS
Gryllus_longicercus         ---GRWRGGVALAKAEASASASMEEDVGGRRLSVAEALVRQGVKGRRWWWPELEAARRPPDPRLCT----CRRTGMQMRMQMRA*SRTVSPPPLASAAPLVSAAEMRPLCRCLRSRPRAS
Consensus

Teleogryllus_occipitalis    ----------------------------------------------------------------------------------------------------------------------
Acheta_domesticus           ASAVA---------------IAFATATASASVKASASAN-------------ASASANASTTASASATASAIATASAIATASANASANASANASA------------------NASAN
Gryllus_longicercus         AGMPALLPICSRAYCHPAGPVLFRLASHLWDLQLRAPSVWLISP*FSRLRLGFAYISSNTFPPCFSLIPTSTFCFLSLLATIP*NYSTIHLANFHIPLILYIACLRCHIFLVFFFNYICV
Consensus

Teleogryllus_occipitalis    ----------------------------------------------------------------------------------------------------------------------
Acheta_domesticus           ASANASANASASA*ASA--------------------SAN----ASASATPSVTTSVIAST-----FANASSNASANASANASANASANAFANTSANASANASANASANASANASANAS
Gryllus_longicercus         CIMSS*CNLVSSFFFTAQLLR*TIGFPENNTLFTVFRFSLNVIFCSSLLLEPTVARSVRFRSSCRLRWLRASTGVTTEYLTVRS*GSSPDGVRTFDTN*IARPCSAAVWRRIPAACYSER
Consensus

Teleogryllus_occipitalis    ----------------------------------------------------------------------------------------------------------------------
Acheta_domesticus           A--------------------------------------NASANASANASANASANASANASA----------NASASA*V-----SASANASASATPSVTASVTASANASASANAST--
Gryllus_longicercus         IFLPFLRFLPFFSVFLPFLRAQHFLPLPCHILIFSYPLLNLQHYSTSRAAKSVGSLELVRIGAVPCCWEIPG*GKSPKADVGWRQISPSSGGRLLPSPAIKTDTKLTRIMTVAAVAPVIL
Consensus

Teleogryllus_occipitalis    ----------------------------------------------------------------------------------------------------------------------
Acheta_domesticus           -NASANASASSAFAIALTPPSP----SPSPSSPPPSPPTPPLPMHPPFSLQFHRPIVTTTSTTTITFAGAVCCAA----FSETQLRFARG*WSIRLRWQTG----ARAVGLSRAP-----
Gryllus_longicercus         IN-------QSMFAWCV--PSPFARLQPSP*SP--------------------------AWADASSADIACCVVGYKHFVTTQSPELRDAFGLRRAPVSGGGRPGSAAGASQNPLLRHW
Consensus

Teleogryllus_occipitalis    ----------------------------------------------------------------------------------------------------------------------
Acheta_domesticus           ------------------RRAFQKS*PQRRAISV--RNKTRNSGVADGFGN--------GSAHEAV*GVRAGTSYF-----QY-------FVFSGCRRTL*KILITAVNRFLRGESINE
Gryllus_longicercus         VLLFTYNSIEFLI*FV*SNRKRWLRALTDITAMSLTARSEVRASTARQDFRY**QR*VEFGLSREQF*PPVAMVSSFPLGSEQYPNSRDLSFLFGALSSPLAYTQLRL------------
Consensus

Teleogryllus_occipitalis    ------
Acheta_domesticus           W*CTMW
Gryllus_longicercus         ------
Consensus
```

**Tocc.5G0008020.1_exon2**

```
Teleogryllus_occipitalis        ------------------------------------------------------------------------------------------------------------------------
Acheta_domesticus               --------GQGRG--RKDRARAFARLFVYWSGI--RLLLIAEAAENY*SEEPAAIMLQSVAEVVLPMWLVLKIVPQSGLEP*DHKVRGTAVTSVDALSHL-----------SVS------
Gryllus_longicercus             KRN*V*ARGEDKAPKRKERSRELG----YCSDPNGKLLTIATGG*NCSLDRPNS---------T*RCY**RKSCRAVEARTSDLAVRDIAVISVNALSHLFRLL*TN*IKNSIEL*VKSN
Consensus

Teleogryllus_occipitalis        ------------------------------------------------------------------------------------------------------------------------
Acheta_domesticus               -----------------------------EAKKGSGAAWQLFKCRLQWKLRQ------------------CSRDGGD----LEVENLRRAKKQTMNKK*IREMCCDV*CRF*AKKKNRI-
Gryllus_longicercus             TQ*RSNGFCDAPAAEPGRPPPETGARRRPNASRSSGLC-VVTKC--LYPTTQHAISALLASAHAGDYGDGCNRAKGDGTHQANID*LIKITGATAATVIILVSLVSV---FIAGDGSNLP
Consensus

Teleogryllus_occipitalis        ------------------------------------------------------------------------------------------------------------------------
Acheta_domesticus               ---*E*RRT*TQSTKDYFLENL*F**SSWVVK---KMRTQDYIRKI*L--KLIHYRYIVREKQGDGAKWKQKRCVKK-ERNVKVNQIKC*KFLEYGSEK*QKSKQAE*RCKRSARRHRVR
Gryllus_longicercus             PLDGEICRQPTSALGDFP*PGISQQQGTAPIRTSSKLPTDFAAREVE*CCKFKSG*EKMRI*QGRGRKCCARRNGRKTEKNGRNRR---------NGRKIRSE*QAAGILLQTAAEH---
Consensus

Teleogryllus_occipitalis        ------------------------------------------------------------------------------------------------------------------------
Acheta_domesticus               AGLSNWRPAGLMRPAG----------SFVRPATALNNFI----------------------VPTKYNVHSVILLRIFIIV*VIINVKYVNYKWKI*KIDYKYEIFEL*MKNMTN*L*MSN
Gryllus_longicercus             -GLAI*FVSKVLTPSGLEP*DLTVRYSVVTPVDALSHLSRHEERKRTDLATVGSSKREEQNITFKENLNTVN--KVLFSGKPIV*RRS*AVK-----------------KNEDTRLHQED
Consensus

Teleogryllus_occipitalis        -------------------------------------------------------------------------SSPSPLASPS-------------------------------------
Acheta_domesticus               MCIKNIMLN------------ISRI*YV--FASL*CQ*FG*KRFKMTVRPAA*LECDI-------------WPISPSKLQSPALEEL*VNETAR----------KSR*GKPDRRSAKQQV
Gryllus_longicercus             MMQTHI*LKKKTRKM*QRRHAI*RMRGM*KLAK*IVE*F*G----MVARSDKKQKVEVGISEKHGGNVLEEM*AKPRRRRENH-GEISQTEGARSCKSQRWEASRNKTGPAG*QYAREQI
Consensus                                                                                                  *

Teleogryllus_occipitalis        ------------------------------------------------------------------------------------------------------------------------
Acheta_domesticus               RKAGGGRRQLEGNGASGRTMRRRADWQ-ERGHSCGGGDVTGASPQRRLRNRAASR--AHLHAHLHTRAPARTRSGVRRPPRRLRHHQRHHHHHHHHHHHHHLPQRPQPPPSQ*PSPPPPP
Gryllus_longicercus             GKSAGIPAE-----ARGRERRQRHNGRISAAETSGAAEASGGGETVRLHARICMRICIPVRLHVHSRG-----SGGRRAASS-------------SGHHHRLPLTPCLTNASATD-----
Consensus

Teleogryllus_occipitalis        -----PLASPSPLASPSPLASPSPLASPS------------------------------------------------------------PLASPSPLASPSPLASPSPLASP--------
Acheta_domesticus               PPPSKPLLPPTPLPPPTPPPPPPPLPPPPPSPPPPPSPPPPPTPPPTPPPTPPPTPPPTPPPTPPPTPLPPPRLLPPPTPLPPPPLPLLPLSSPPPSPTPPPTPPPTPLPTPPPTPPPTL
Gryllus_longicercus             -----NLLPPTS-SSIDADADASAFANAT-----------------------------------PPRQRPRQR----------------PASANAP-ASGNASASGNASASA--------
Consensus                             *  *                                                                               *     *

Teleogryllus_occipitalis        SPLASPSPPPSPPPTPPPS*----------------------------------------------------------------------------------------------------
Acheta_domesticus               SPTPPPTPPPTPPPTPPPTPPPTPPPTPPPTPPPTPPPTPPPTPPPTPL--------PTPLPTPLPPPRSLPPPTPLPPPPLPL---------------LPLSPPPPTPLPPPTLPPMHP
Gryllus_longicercus             SASASVSAPSSAITSPIPST----------------YCSANSNASTCPLFLFLNFSFSLLLSFTLSLSRSLSPSL---VHSLPLSFTLSLSLFLSLSLFLSLS----LFLSLSLFLSLSL
Consensus                       *       *      * *

Teleogryllus_occipitalis        ------------------------------------------------------------------------------------------------------------------------
Acheta_domesticus               QMPLPPPPSPSP*L--------HHRHRL------------PHPLRHRLHPHR------RCQCIHLSLSNSIAPSLPPPPPPPSPSPALSAAPPSRKPSSVSREDDGASGC----AGRRAR
Gryllus_longicercus             SLFLSLSLSPSPYISIYLHRYHHHYHHLRRHGLPSRLLGNPAPFRARMMKHPAALADGRAVCF------SRVPGF----------PEILAAKAS---------DIGAK*NTK*RSGRRVW
Consensus

Teleogryllus_occipitalis        ------------------------------------------------------------------------------------------------------------------------
Acheta_domesticus               GQSACRGR----------------------------------RAGLSR-----NLSRKGERYRCEIKHEIA-------------------------EWPTGLATARHTRPCKECALEQV-
Gryllus_longicercus             Q*LGTRGRVRSARWNKLFSIFRVVWQQKDAVKNSYYNGQQIFKRGMYKPMVMYNVGRMCLQIRLSMQTKYAKCFQARLVTNMLIWNVYYCFYGY*FEFQKCLSRN*QIISS*NVIYESVY
Consensus

Teleogryllus_occipitalis        ----------------------------------------------------------
Acheta_domesticus               -------------IFNISCSLAAGGRCEKFLLQRLTDF*EGKV*TNGNVQCG------
Gryllus_longicercus             SSLHLLHL*WALMIVSIICSV*DP-------LQEIFSYT*STNDYSYNITY*FM*LRQ
Consensus
```

**Tocc.5G0008120.1**

```
Teleogryllus_occipitalis        AAACACCAACACCCAAA----------------------------------------------------------------------------------------------------------
Tarbinskiellus_portentosus      AAATATGGCAGTGTAGA-----------AGAATTAAAATGAAAAAATAGAGATGTC--------ATTGGTGACGGAATTACCACTTACGGGGTCGCGCCGGGGTGTTCCGACACCCCAA
Acheta_domesticus               AATTTTAATCACTGTAA---------------------------------------------------ATTATTGTATTTTTATCTTCCAGAGGCTTTAAGATCGCCATTTTATTGAAA
Gryllus_bimaculatus             AAATCTAAAAATTTGAACACGTCTGT----AAGTCTAAATTCGA---CGTAATTTTC-AATCATAGCCAATTATCGTATTATTATCGTTTAGAGACTTTAAGACCATTACTGCATTGAAT
Gryllus_longicercus             ATACACAAACACACAAATACACCCATGCCCGAGGCGGGATTCGAACCCGCGATCTTCTGGTCCGAGACCACTCCGGCACTACTGCC--TTAGCGATCTCGG---------CCATCTGAT
Consensus                       *               *

Teleogryllus_occipitalis        ------------------------------------------------------------------------------------------------------------------------
Tarbinskiellus_portentosus      GAAATTCCA----------------------------------------------------------------------------------CCCTGTTATCGATATGTTTTTTTTT
Acheta_domesticus               TAACTTTCAGAAATAGTTCTTTGCACTTTCTCTTGCTTAACATTGTTCTTA----------------------------------------------AATTAATGTATTTTCTGTTT
Gryllus_bimaculatus             CGACTTG----AATAAAAGTGTTCACCTTCTGTTGGTTATTATTATTATTATTATTATTATTATTATTATTATTATTATTATTA------TTATTATTATTGTTAACGTATTTTCTGTTT
Gryllus_longicercus             CGGCATT----ATTATTATTATTATTATTATTATTATTATTATTATTATTATTATTATTATTATTATTATTATTATTATTATTGTTGTTGTTATTATTATTAACGTATTTTCTGTTT
Consensus

Teleogryllus_occipitalis        -----------------------------------------------------------------------------ATATGGTGAGAAAAGTA---------------------
Tarbinskiellus_portentosus      TTT--------------------------------------------------------------------------TTTTGAAAAAAAAGTTAATATCTACGACTTCTTTAGGGT
Acheta_domesticus               TTCGGAGATCATTTTATTTTATATTCTTAGCCTATTCTTCGCCGTT----------------------------------------------------------------------
Gryllus_bimaculatus             TCCTGAGATCATGTCATTGTATTTTATTAGCCCATACTTATTTATTTGACGTCAAAAACAGTAAAAAAACTTGTGTGACTTGCTATGGATAGAAGTGTATTCTGTGCATGTTTTTCGAAAT
Gryllus_longicercus             TCCTGTGATCATTTCATTGTATTTTATTAGCCCTTACTTATTTATTTGACGTCAAAAACAGTTAAAAACTTGTGTGACATGCTATGGATAGAAGTGTATTCTGTGCATGTTTTTCGAAAT
Consensus

Teleogryllus_occipitalis        ------------------------------------------------------------------AAAGGAAATTAATCACCGGAGTATTTTTTTGCCGT--------------
Tarbinskiellus_portentosus      CTACATAATAATGCCTTATAGTTTGAGTTAATTATTCCAGTAGTCCTTAGAGTTGATTATTCCAGAACAGGAAAGAAATTGA---------------------------AGAATCTTT
Acheta_domesticus               ----GAAGTAAAACCTGGGATTC---------ACGTTAAACACCTCTTAAAAATAAATGTTTAA------ACTGTAGGTTGATTTCCA-------TTGCCACTACACTTCCAAAATTTGT
Gryllus_bimaculatus             TCTGTAAGTAAAATCTGAGATTCA----CTATACATGAAACAACTCACAACAATAAATATTTAAATGTTTAAACTAAATTGATTGCTAGTAAACTCCATTGCCATATTTTCAAAACTCGT
Gryllus_longicercus             TCTGTAAGTAAAATCTGAGATTCA----CTATACATGCAACAACTCACAACAAAAAATATTTAAATGTTTAAACTAAATTGATTGCCAGTAAACTCCATTGCCACATTTTCAAAACTTGT
Consensus                                                                                                *   ** *

Teleogryllus_occipitalis        ------------------------------------------------------------------------------------------------------------------------
Tarbinskiellus_portentosus      CAGGAT------------------------------------------------------AAGGGAATATTTGCTCTTTTTTTT----------------------TACATTTCAAC
Acheta_domesticus               TAAAA---GTAAGAATTATGAGTTTGGCACATGAATTTTAACACATTACACATTAAGCTTACGAGTGAAATAGTAGTTTTCGTTTTTATGAACAAAAATGTAAATGATATAATTTATGAT
Gryllus_bimaculatus             TACAATTTGTTAGAATTATGAGTTTGCCACATTCATTCTGA-GCATTACACAAGAAGCTGAGGAGTGAAATAGCGGTTTCCGTTTTTATGACCAAAAATGCTAGTGATATAATTTATAAT
Gryllus_longicercus             TACAATTTGTTAGAATTATGAGTTTGCCACATTCATTCTGA-GCATTACACAATACGCTGAGGAGTGAAATACCGGTTTCCGTTTTTATGACCAAAA-----AGTGATATAATTTATGAT
Consensus

Teleogryllus_occipitalis        ------------------------------------------------------------------------------------------------------------------------
Tarbinskiellus_portentosus      AAAATGATATTTTTAAGAATTC-ATAATATTTTTGAATGTTCTTTCGATCATGCGGG---------------ACTAGTAATGCAAAAT----------TTATTTTATATTATATTACAT
Acheta_domesticus               GTATTGATATTTTTGCACACT--AACATATTTTGACTGCCTCTTTTCTTTGCGTCTAGACGAAATTTAGTGACGTTGCCAAAGCACAACGTCAATCTAGTTACCGTGGATTTTCATACAC
Gryllus_bimaculatus             CTATTGATATTTTTGCATACTTAATCATGTTTTGACTGCCTCTCTCCTCTACGTCTA---------------GTTGCCGAAGCAATACGTCA----CGTTGCTGTGTATTTTCATACA-
Gryllus_longicercus             CTATTGATATTTTTGCATACTTTATTATGTTCTGACTGCCTCTCTCCTCCGCTTCTA---------------GTTGCCGAAGCAATACGTCA----CGTTGCCGTGTATTTTCATACA-
Consensus

Teleogryllus_occipitalis        ------------------------------------------------------------------------------------------------------------------------
Tarbinskiellus_portentosus      AATTAATTTTTAAATATGTTAATTTATTGGTCGCCTGGGATGGCTGTAA------------GAAATAAATCGAATTA-------------------------------------------
Acheta_domesticus               ------CTCTTGGAAATGGCCAATTATAAATACCATACTCTGAATATAATTTCAAGTTTGAAACACAAATAAAAGATTAGTGCTTCAATGAAAATCAAGGACAGAATATACGTCCTCTGT
Gryllus_bimaculatus             --------GTTAGAAATGA-AAGTTATAAACACCATAAACTGAATTTAATTCCAAGTTCCGAACAGTAATCAAACTT-------------------------------------------
Gryllus_longicercus             --------CTTAGAAATGG-AAGTTATAAACACCATAAACTGAATTTAATTCCGAGTTCCGAACAGTAATAAAACTT-------------------------------------------
Consensus

Teleogryllus_occipitalis        ------------------------------------------------------------------------------------------------------------------------
Tarbinskiellus_portentosus      ------------------------------------------------------------------------------------------------------------------------
Acheta_domesticus               TTCCAGAAAATATCCCCCTATTTTCTGCCCATGGCCAATTTTCCTTCCTTAGAACGACTTCGATAAGGAATTTATTTTGTAAATCATGAATTTTCAAAAGACAGAAAAATTATGGTAGCC
Gryllus_bimaculatus             ------------------------------------------------------------------------------------------CTTTCAAAATCCATAA-------------
Gryllus_longicercus             ------------------------------------------------------------------------------------------CTTTCAAAATCCAGAA-------------
Consensus

Teleogryllus_occipitalis        ----------------------------------------------------------------AATTGTGTCTTATATTCGCTCAA---TACAGGAGGTTGCAACAAAAGCACCTGGG
Tarbinskiellus_portentosus      ------TACTGGCGGTAGAATGAT----------------------------TATTTTT-----TGTGGCGCATTTTGCTCGTTCAAGTTTCTACGCGAATTTAAGAAACGAAATTAAG
Acheta_domesticus               TCTCTGTGCTTGCGACGTGTAAATAAAAGATAGGAGGAAATTAATCAGCGGAGTATTTTTGCCGTAATTGCATCTTCTATTCATTAAA---AACATGAGATTACAATAAAAGAGCTTAGG
Gryllus_bimaculatus             ---------TTGTAATATAAAA----------------------------------------GAAATTGTATCTTCTATTCATTCAA---TATATGGGGCTTTAACAAAAGCACTTGGG
Gryllus_longicercus             ---------TTGTAATCTAAAA----------------------------------------GAAATTGTATCTTCTATTCATTCAA---TATATGGGGTTTTAACAAAAGCACCTGGG
Consensus                                                                           * *    ** *  **  * **      * * *  *  ** *** *    *  *

Teleogryllus_occipitalis        ATTGACGGCGAGAGAAAATGGCGTACAGTAACGATTCAAATTTTAGAGGGA------------GATGTCTTTTCT--TTGATGCTTCAAGTAAATGCAGAGCCGAACATTGAATGACCT
Tarbinskiellus_portentosus      A------------AGAAATTACGAACAATAACGGGGAAAATGTACGTAGGTGACATGTGCATCCGAAATGGCTACTAACGGGTTTGAGAACAAGAAACTTAAAGAGAGAACCAATGTAAT
Acheta_domesticus               ATTGACTGTGGGGGGAAATGGCTCACGATAATGGTTCAAATTTTAGAGAGAGAGAGGAAGAGATGGAGTATTTTCT--TGGATGCATAAAGTAAACGGTGAGAC-GATATTGAATGTGTC
Gryllus_bimaculatus             ATTGACTGCGGGAGGAAATAGCTCACAGTTACAATTCAAATTTTAGAAGGA------------GGTGTAATTTTT--TGGATTCATAAAGTAAATGGTAGAACAGACATTGATTAAACC
Gryllus_longicercus             ATTGACTGCGGGAGGAAATAGATCACAGTTACGGTTCAAATTTT---AGGA------------GGTGTAATTTTT--TGGATTCATAAAGTAAATGCTAGAATAGACATTGATTAAACC
Consensus                       *          ****     **  * *      **** *      *          *   *   *  *    * *     **  * *         * *   * *

Teleogryllus_occipitalis        G------------------------TAGT------------TACAGTAGAAACCT-----------------------------------------------------------------
Tarbinskiellus_portentosus      C---------GTAAACAATGTAAACGTAAAGAATGTAAATACATACAGTGCGAATGTAAAGAACGTAAACTTATCGAAAGAATGCAAATCAAGTCCGTTTCAGAGTTAATACGCTGAAAT
Acheta_domesticus               T-GTTATTTCCTAAATGATTTATCATTAAT----AAACAAACATACAGTAGAATTCT-----------------------------------------------------------------
Gryllus_bimaculatus             TGTTTTTTACCTAAATGTTTTATCAGTAAT----AAACAAATATACTGCTGAATCCT-----------------------------------------------------------------
Gryllus_longicercus             T-TTTTTTACCTAAATGTTTTATCAGTAATAAACAAACAAATATACAGTTGAATCCT-----------------------------------------------------------------
Consensus                                                 **               *** *    *   *

Teleogryllus_occipitalis        -TTCAGTTATAATTTACTTAGCTCA----GCAAGTTCTAAGAAGTTTTCTACCACTTATAATGAATAAT-----ATTGTTGGAACTTACAACGCACAGACACTTACACGTACATGTGAGA
Tarbinskiellus_portentosus      ATTTAAATATTATTACTAAAATTCATTTGAAAAATTAAATTCAAATTTGAGGCAAATATCGAAAATAGTTAACGTTTGTTAATGTTTAATGTGA------------ACATTTGAGTGATG
Acheta_domesticus               -TTCAGTTAGTATCATCATAGAATA----GCAAGTCCAATGAAGTTTTCTACCACGTATAATGTAAAA--------TGTTGGAACCTACAATGC-------------------------G
Gryllus_bimaculatus             -TTCAGTTA---GTCTTGTAGTTCA----GCAAGTACTATGAAGTTTGCTACTACATATAATTAATAAT-----ATTGTTGGAACTTCCAACG--------------------------
Gryllus_longicercus             -TTCAGTTACAGTCTTCGTAGTTCA----GCAAGTACTATGAAGTTTGCTACTACATATAATTAATAAT-----ATTGTTGGAACTTCCAACGC-------------------------
Consensus                        ** *  **          *    *      ** *   *   *  **      *  ***     * *         ****      *     *

Teleogryllus_occipitalis        GCGCGCGCGCACACACACACATACACACACACACATACACACACACACACACACACACACCATA-----------ATCGTTTGAACCTACTAAAAACACTTTTATTGAAGAATAT-
Tarbinskiellus_portentosus      AGATCATTGTACGGA---------------AGACAATCAGAGACAAAAACACACGTTTTAAAAATTTGAAAAATATGCTTTTTTATTACCCTCCCTGAAATCTGCATGTGAAATTATATC
Acheta_domesticus               GTGCGCGCGCACGCACGCACGCACGTG---------ACACGTACACACACGCGCACGTTAATCTT---------------ACTCTTGAATCTACTAAAAATACCATTAATAAATAATAT-
Gryllus_bimaculatus             ---------------------------------CACACACGCGCACACACACATACACTAATATT---------------TTATTGAGACCTACCAAAAATACTATTATTAAATAATTT-
Gryllus_longicercus             ----------ACGCACGCACACACACACACACACACACACACACACACACATACACTAATATT---------------TTATTGAGACCTACTAAAAATACTATTATTAAATAATTT-
Consensus                                                            **    ** * ** *        *                        ** *   ***      *    **  ** *

Teleogryllus_occipitalis        -------------------TATCTTCCCGTAAATATG--------------AAACTGCATTCATTCGTAGGCTCTCTGAAGCCTAACACC-----GCATCCCCGCAGA--ATATTTTTC
Tarbinskiellus_portentosus      ACACTTTTTGCGTGTCCAATATCTTTCCTATATTACGTAAACGAGCAGATAAAAAATGCCATCATTCTTACATTCTTTTGCGCATAACGCCATTGAACCTCTCCGTAGA--ACATAAATC
Acheta_domesticus               -------------------TATCTTCTCTTCAATATG--------------AAACTGCATTCATTCGTAGGTTATTTGACGCCTACCACC-----ACACCCACGCAGAATATATTTTTC
Gryllus_bimaculatus             -------------------TATCTTTTCGTGAATCTG--------------AAACTGCATTCATTTTTAAGCTATTTGACTCCTATCACC-----ACATCTCTGCAGA--ATATTTTTC
Gryllus_longicercus             -------------------TATCTTTTCGTGAATCTG--------------AAACTGCATTCATTTTTAAGCTATTCGACGCCTATCACC-----ACATCTCTGCAGA--ATATTTTTC
Consensus                                          ******  *   * *  *              *** ***  *****  **   * *      * ** * **      *  *   * ***  * **   **

Teleogryllus_occipitalis        GAGCCTATCTCCACTCAATCGC-AGGCTCGGAGCGATAGCCGCCGCGAACGTCCGCCAAGTCCCGCGCTCTGATGCGCCCCTGGAGTTTACTGCTGCTTATAAAACGCAATACGGCGAAC
Tarbinskiellus_portentosus      AAAACCCTCTCCACTCAATCACAAAGCTCGAAATGATATCCGCGGTGGATGTCCACGGAATTACGCGCTCTGATGCGCCCCGGCAGTTTACTGCTGCTTGTAAAACGCAACACGGCGAAC
Acheta_domesticus               GAACCTATCTCCACTCAATCGCAAAGCTCCAAACGACATCCGCCGCGAACGTCCGCGAAGTTGCGCGCTCTGATGCGCCCCTCGAGTTTACTGCTGCTTGTAAAATGCAATACGGCGAAC
Gryllus_bimaculatus             GAACCGATCTCCCCTCAATCACTAAGCTGCAAACGATATCCGCCGCGAACGTCCACGAAGTTGCGCGGTGTGATGCAGCCCTGGAGTTTACTGCTCCTTGTAAAACGCAATACGGCGCGC
Gryllus_longicercus             GAACCTATCTCCACTCAATCGCAAAGCTTCAAACGATATCCGCCGCGAACGTCCGCGAAGTTGCGCGCTGTGATGCGGCCCTGGAGTTTACTGCTCCTTGTAAAACGCAATACGGCGCAC
Consensus                        *  *  ***** ******* * * ***   *  ** * **** * * * **** *  * *  **** * ******  ***   *********** *** ***** **** ******  *

Teleogryllus_occipitalis        ACTAAATCCCGACACGAAACGTCGCTCCCGCGCATTTCCGTCATTGAGTCCTCTCGTCGCAGACG------CCGCGCACGCACTC-------------CACAGCGGTCGCCGCTCCGCG
Tarbinskiellus_portentosus      ACTAAATCCCGACACGAAACGTCGCTCCGGTGCATTTGCCTTATTGAGTTGGCGCTGCCGAGACG------TTACGGACGCGCGCTGCCAGGTCGTCGTAGCAACGACGGCGGCGACGGG
Acheta_domesticus               ACTAAATCCCGACACGAAACGTCGCTCCGGCGCATTTCCCTTATTGAGTCCTCGCGGCGGAGGCGCCGCCCCCGCGCACGCACTC-------------CAGAGCGGACGCCGTGCCGTG
Gryllus_bimaculatus             ACTAAATCCCGACACGAAACGTCGCTCTGGTGCATTTCCCATATTGAGTCCCCGCGTCGGAGGCG------CCGCACACGCACTCGA-------GGCGGCGCAGCG-------------
Gryllus_longicercus             ACTAAATCCCGACACGAAACGTCGCTCTGGTGCATTTCCCATATTGAGTCCCCGCGTCGGAGGCG------CCGCACACGCACTCGA-------GGCGGCACAGCG-------------
Consensus                       **************************  * ****** *   *******   * *  *  ** **         *  **** * *                * **

Teleogryllus_occipitalis        ---CCCGCGGGACGCGCGCTCCCAGGCCGGCGGCAGGTG----------------------GCGGTGGAGGTGGCAGTG----GCGGTGGCGGTG-----------------------
Tarbinskiellus_portentosus      GCAGCCACTGGGCGCGTGCGTGTGCGTGTGTGCGCAGCCG-------------------CTCGGCCAAAGGTCAGGGTT---CGCGGTG----------------------------
Acheta_domesticus               --CCGCACCGCGCACTCACTCCAGGGCGGACGCCGTGCCGTGCCGCACCGCGCACGCACTCTAGGGCGGACGTCGTGCCGTGCCGCACCGCGTACGCACT-------------------
Gryllus_bimaculatus             ----CCGCGGTACGCGCGTCCAAAGGCCGCCGAGGCGGG----------------------CGGGGGAAGACGGTAGCT----GCGACGCGGGCGTGCGACGCGGGCGGCCAGTGGGCG
Gryllus_longicercus             ----CCGCGGTACGCGCGTCCAGAGGCCGTGGAGGCGG------------------------GAGGGAAGGCGGTAGC---------------------------------------
Consensus                           * * *  * *          *     *    *                            *

Teleogryllus_occipitalis        ---------------------GCGGCGGTGACGGCG-----------------------------GTGGCGGCGGCGGCGGCGGCGGCGGGGGTG------GCGGCGGTGGCGGC
Tarbinskiellus_portentosus      ---------------------AAGCGGCGGGCGCCGCGCTGCGCCGGACCGTGCG----------GTCACTGCTGTGTTG--GGAGACTGCGGACGCG------GTGCGAATGGTGGG
Acheta_domesticus               ---------------------CTAGGGCGGACGCCGTGCCGTGCCGCACCGCGCACTCACTCCAGGGCGGACGCCGTGCCG--TGCCGCACCGCGCACGCAC---TCTAGGGCGGACGT
Gryllus_bimaculatus             TGCGGCGGTTGCTAGCTGGGGCGCGGCGCTGGCGGCGTACGGAGTAGGGGCGCGCG-------GGTTGCGGAGGCGGAGGCGA-GGCGGAGGCGGAGGCGGGCGGTGCGGTGGCGGTGGC
Gryllus_longicercus             ------------------------------------------------------------------------GGCGGCGGCGG-GGCGGCAGCGGG----------GCGGCGGCGACGGC
Consensus                                                                                         ** * *  *   *      **                  *   *

Teleogryllus_occipitalis        GGTGGCG-------------------------------------------GCGGTGGTGGTGGCGGTGGC-----------------------------------GGCGGCG-------
Tarbinskiellus_portentosus      GGTTGGA---------------------------------------------------ATGAGGGTGGGGGC-----------------------------------GGCAGTG-------
Acheta_domesticus               CGTGCCG------TGCCGCACCGCGT------------------------ACGCACTCTAGGGCGGACGC-----------------------------------CGTGCCGTGCCGCA
Gryllus_bimaculatus             GGTTGCGTGGCGGTGCGGCGTGGCGTGGCAATGGCGGGGTGGGCAAACGTCGCGCTGGCGGCGGTGGGAACGAGCGGTAGTGGGGCGGCCGATGCTAGTGTGACTGAGCGGCGGATCGCG
Gryllus_longicercus             GGTGGCG----------------------------------------------------GGTGGTGGTAAC-----------------------------------AGCGGCG-------
Consensus                        **                                                       ** **    *                                    *    *

Teleogryllus_occipitalis        -------------------------------------------GCGGCGGCGGCG----------GCGGCGGCGGCGGCGGTGGT--GGCGGTGGCGGTGGCGGCGGTGGCGGCGGCGG
Tarbinskiellus_portentosus      -------------------------------------GTTAGGGTGGTGGGAGTG-------CGGTGACGGGATGGTGGTTGTTGTT-ACTGGTGTTGG--------------------
Acheta_domesticus               CCGCGCACGCACTC------------------------CAGGGCGGACGCCGTG-------CCATGCCGCACCGCGGACGCACTCT-------------------------------
Gryllus_bimaculatus             TCGTCAGCGCGTCCGGTGTGTGGCGGAGCCGGTGGCAGACGAGGTGGCGGCGGGGAGAGGGGCGGTGTGGCAATGGTGACGGTAGATGGCGGATGCCAGTTGCGGAGGTTTGCAATTGGG
Gryllus_longicercus             ------------------------------------------------------------------GTGGCAATAGTGACGGTAGCT-GCGGATGCCAGTTGCGGAGGTTGCAATTGCAG
Consensus                                                                                          *  *       *

Teleogryllus_occipitalis        CGGTGGCGGTGGCGGCGGTGGCGGCGGTG--------------------------------------------GCGGCGGCGGCGGCGGCG--------------GTGGCGGCGG
Tarbinskiellus_portentosus      -----------GTGGTGGTGGTGGTGGTG--------------------------------------------GTGGTGGTGGTGGTGGTG--------------GTGGTAGT--
Acheta_domesticus               -----------AGGGCGGACGCCGTGCCG-------------------------TGCCGCACCGCGCACGCACTCCAGGGCGGACGCCGTGCCG-------------TGCCGCACC--
Gryllus_bimaculatus             TTGT-GTGTTCGGGGGGGAGGTTGTGATG-GAGCGTTGTTTTAGGACTGGCATGCTGGTGCTCGTGGTGGCGGCGTCGCGGCGGCGGCGGCGGTGTGGTGTGGTGGTGGTGTGGTGGTAG
Gryllus_longicercus             TTGTGGTGTTCGGGGGTGGGTTGTTGGTGAGACCGTTGTTTTGG---------ATCGGTGATTAAGATGGACATGCGGTGGTTGTGGTGGTGGTG--------------GTGGTGGT--
Consensus                                     **  *       *  *                                                **  *  *  * *  *               *  *

Teleogryllus_occipitalis        TG----------------------------------------------------------------------------------------------------------------------
Tarbinskiellus_portentosus      ---------------------------------------------------------------------------------------------TGTTGGACCATTGAGAATAA
Acheta_domesticus               -------------------------------------------GCGCACGCACTCTAGGTCGG------------------------------------------------------A
Gryllus_bimaculatus             TGTGGTTGTGGGTATGTAGATGTTGGTGAGGTGAAGAAAGTGCGTGCGTGCGTAGTGGGTAGTGTCTGTGTGTGTGTGTGTGTGTGTGTGTGTGTGTGTGTGTGATTGTTGTGGATTG
Gryllus_longicercus             ----------GGTATGTGGATGCGGGTGAGATGAGTAGTG---GTGCGTGAGTGGCGGGTAGT----------------------TGTGTGTGTGTGTGTGTGATTGTTGTGGATTG
Consensus

Teleogryllus_occipitalis        -GTGGTGGCGGTGGCGGTGGCG-----GTGGCGGTGGCGGTGGCG--GCGGTGGCGGTGGCGGTGGCGTGCG----------------------------GTGGCGGTGGTGGCGGTG-
Tarbinskiellus_portentosus      GGTGGAGGTAATAATGCACTTATAAAAGAGCAGGTGGGAACAGATTGCTTATTCAAACAATAGCGACATGTTTG---------------------GTTCACTTGGAAAAGGATGAAATG-
Acheta_domesticus               CGCCGTGCCGT-----GCCGCAC---CGCGCACGCGCT----------CCAGGGCGGACGCCGTGCCGTGCCGC-----ACCGCGCACGCAC---------TCCAGGGCGGACGCCGTG-
Gryllus_bimaculatus             CGTAGTGTGGCTGGTGGTTGTG-----GTGTGTGTGGTGATAAC------GTGTGGGTGATGGTGACATGTGGTTCATAGTTGTGTATTGGTGATGGTGATGTATGGGTGGTGGCGGTGT
Gryllus_longicercus             CGTAGTGTGGCTGGTGGTAGTGG---TGTGTGTGTGGTAATAAC------GTGTGGGTGGTGGTCACATGTGGGTCGTAGTGGTGTATGGATGATTGTGATGTGTGGGTGGTGGCGATGT
Consensus                        *  * *                    * *   * *                     *   * **                                  **  *   **

Teleogryllus_occipitalis        ----------------------------------------GCGGGGGCGGTGGC-------------------------------------------------------------GGTGG
Tarbinskiellus_portentosus      ATGTCAAAGGGAGAAGTA----------------------TTATAAACAGTGGCATT----------------------------------------------------------GATAA
Acheta_domesticus               ------------------------CCGTGCCGCACCGCGCACGCACTCTAGGGC-------------------------------------------------------------GGACG
Gryllus_bimaculatus             ATGTTCGGTGGGGAAGAAAATGAGTAAGTGGTGGTGTAGGCGTGTGAGTGGTGGTAATGGTATCGACGGCGTAAGGATGTAGTGGAATAGGTTGTTGTGTTGGCGTGTAAGTGGAGGTAA
Gryllus_longicercus             GTGTCAGGTGGGGAAGAAAATGAATAGGTGGTGGTGTAGGCGTGTGAGTGGT-------------------------------------------------------------GGTAA
Consensus                                                                                                                                   *

Teleogryllus_occipitalis        CGGTACGG-----------------------------------TGTGGTTGC------------------GGCGGCGGTGACGGCGGTGG-CGGCGGTGGCGGCGGTGGCG-------
Tarbinskiellus_portentosus      TCATTTCG-------------------------------CTATTATCATCGG-----AATAATGAATGTGTGAATGTGGTAATAACGCTGACTGGTAAAAACGACAATTGCAATTTTTG
Acheta_domesticus               CCGTGCCGTGCCGCACCGCGCACGCGGGACGCCG----------CGCCGCCGC---------------------------------------------------ACACAGGGGTATTTTG
Gryllus_bimaculatus             TGGTATGAGCGTGTGTTGTGTATAACTGTTGTTGTTGTTGTTGTTGTTGTTGTTGGTTGTTGTTGGTTGT-TGGTGGTGTTGGTGGTGTTGG-TGGTGTTGGTAGTGTTAGCGTGTTTTG
Gryllus_longicercus             TGGTATGAGCGTGTGTTGTGTATGACTGTTGTTGTTGTTGTTGTTGTTGTTGT-----------------------TGTTGTTGTTGTTGG-TGGTGGTGGTGGTGTTAGCGTGTTTTG
Consensus                          *                                            *                                                      *

Teleogryllus_occipitalis        -------------------------------------------------------------------------------------------------------------GCGGTGC
Tarbinskiellus_portentosus      ----------------------------------------------------------------------------AACCATTATT-----CTTTTGGTTTGTAATTTCACTCGTAGGGC
Acheta_domesticus               TAAGAAAAGCTGGCCAA--------------------------AAGTCGAAAAAAAAAATTTTAACATACGAGAAAACTAACACTATTTCCTTTAAGTTGAATATATATTACGTCATAC
Gryllus_bimaculatus             TGATGTAGACGGATGAAGGTGGTGATATGTGAATAGGGACGGAGAGGTGCGATGAAGAAACGATGCTGCAGACTGATAGTGATGTT----------AAGGAGGCGTTAATCATGTGATGT
Gryllus_longicercus             TGATGTAGATGGATGAAGGTGGTGATATGTGAATAGGGACGGAGAGGTGAGATGAAGAAACGATGCTGCAGAGTGATAGTGATGTT----------AAGGAGGCGTTCATCATGTGATGT
Consensus                                                                                                                               *

Teleogryllus_occipitalis        GGCGGCAGTG--GCGGCGGTAATGG---------------CGGTGGCGGTGGTGCCGGCGGTGGCAGCGGTGGTGGTGGTGGCGGTGGAGGCGGAGGCGGCGGTGAGAACGGCGGCTGCC
Tarbinskiellus_portentosus      GGCATAAGTG--GTATTTATCGAATGTAGATC-----------------------------TGAATGTAGTGCTGATAATGGTGAGAGTGGCAAAGGTGGA------------------
Acheta_domesticus               GACAGTTAT---CTGCTAGTCTCAGGCTAGTA--------TCTTCGTTGTAGCATCAGTCTAAAGTGAGTCGTTGTACGCGCTGCTAGAATTTG--------------------CTACC
Gryllus_bimaculatus             GGTGATGATGGAGTGGTGGTAGAGGGTCAGTAGTCTGGCAGAGTGGCATTGGTAGCGCGGCAAAGAGTGGTGGTGATGGCGGTGGCAGGATCGGAGGCGGC------------------
Gryllus_longicercus             GGTGATGATGGAGTGGTGGTAGAGGGTCAGTAGTCTGGCAGAGTGGCATTGGTAGCGCGGCAAAGAGTGGTGGTGATGGCTGAGGCAGGATCGGAGGCGGC------------------
Consensus                       *       *          *                                           *    * **       *   *

Teleogryllus_occipitalis        GTTGATGCAGATGCCAATTGGGGGGGGGG--------------------------------TGGAAGGTGA--------------------GTTAAGAA---ATGGGGGTTCGATGA
Tarbinskiellus_portentosus      --TGATTATAATGTTATTTTGTTGACCGGTAT------------------------------CGTAAAATGACTTCGAATACTTAACAACATTAGCTAGCG--CGCACTCGTGTGCATG
Acheta_domesticus               A-----------GCCGTTCAATACAAAGGGATCCTTTATTCGCTTTGTGAGTGCATTCAAATTCGTGAAGTGT------------ATTAATTCGGCACGGATTTAGAAGTTCCAGGCAT
Gryllus_bimaculatus             ------------GCCAGTTGGGACGGAGAG--------------------------------CAAAAGATGG-------------CGACGACGGCAAGGA---AGACATGCGCGGTGG
Gryllus_longicercus             ------------GCCAGTTGGGACGGAGGG--------------------------------CAAAAGATGGCGACGA-----CGACGACGACGGCAAGGA---AGGCATGCGCGGTGG
Consensus                                    *    *         *                                 *  **                        *              *

Teleogryllus_occipitalis        -GATTGT-----------------------------------------GGTGGTGGCTGTGGCGGTGGTGGTG------------------------TATGGATGGTGGTGTCGTGGG
Tarbinskiellus_portentosus      TGTTTGTACATTTGTTTATTTTTTCTCTCTGGAAAGCATTAGTTCAAATGGTGCTTTTTATTAGGTAAAAGATGATCTGCATTTCTTATTTCCCTTTACCGTATTACATCGTTTCCTGAT
Acheta_domesticus               ATACTGT---------------------------GTTTAAAATTAAAAACTGGCCCATTTAACATTATTATTTA-----------------------CATAATTTAAGAATTTTTGGC
Gryllus_bimaculatus             GGACTGCG------------------------CGGTGGGGACCAGGGGGTAGA-----------GGCGGGGA----------------------CTGCGGATGACGGCTCCATCGC
Gryllus_longicercus             TGCCTGT-------------------------------GGGGCCAGGGGGTAGAGGCGGGGACGACGGCGGGGA----------------------CTGCGGATGACGGCTCCATCGC
Consensus                           **                                               *                                                           *

Teleogryllus_occipitalis        TAG----------TGATGTGTGTG------------------GGGGGTGGTGTGTGGGGGGAT--------------------GCGCCGTTGGT-----------------------
Tarbinskiellus_portentosus      CGAATTTTTCCATCTTTATATTCATTATCTTCAAA-----ATTGCTGCTGGAATGTGGTGGCTTCCCTGTTACGTAAATCGATTTGCACCTCCGAC-----CGAACACACAAAACGCAAG
Acheta_domesticus               CAG----------CTTTACGTTCACTATACCCGTATGTGCACCGGTGGGAGGCCATGGAGGCGG-------------AAGGGAATTCGGCGATGGT-----------------------
Gryllus_bimaculatus             TGG----------GGCAAAGGTCA------------------GGGTTCGCGGCATAGAGGC---------------------TGGTCGGTGGCCGCT-GGAGTATCGCGGTCGCAGA
Gryllus_longicercus             TGG----------GGCAAAGGTCA------------------GGGTTCGCGGCATAGAGGCGT-------------GGAGGGCTGCGTCGGTGGCCGCTGGGAGTATCGCGGTCGCAGA
Consensus                                                                *            * * **                         *   *

Teleogryllus_occipitalis        ------------------------------------------------------------------------------------------------------------------------
Tarbinskiellus_portentosus      CGT-------------------------------------------------GTTAACAATATGCTGAATGA--------------------AAGATGCTCTAACAAACGGGT
Acheta_domesticus               -----------------------------------------------------------------------------------------------------------GAGACCGAATT
Gryllus_bimaculatus             ------------------------------------------------------------------------------------------------------------------------
Gryllus_longicercus             GGTCGGGTTGGGACGCAGACACGGTTATGGTGGCGTTATTGACAATAAGGGTTGTAGCGGCTAGGAATTCATTGGGGGAACAATAGATGGATATTCACCAAAGCACTTCAAAAATGAAGT
Consensus

Teleogryllus_occipitalis        ------------------------------------------------------------------------------------------------------------------------
Tarbinskiellus_portentosus      TTCGAATAATTAAAAGTTT--------------------TCACAAATTATTTATATTGAGA-------------------------------------CGGCACAGATAATATTATC
Acheta_domesticus               TTTATGTAATTACATATTTTTTTGTCTTTCTACTTACCGTTCTCAAAATTTTAATCTGACGAATCGTTGCCGTCCTACCACTATATCACATTTTTCCGTTACATATTTGGTAATATATTA
Gryllus_bimaculatus             ------------------------------------------------------------------------------------------------------------------------
Gryllus_longicercus             GGAATGTACGTAGGTATGCATTTGGTTTATTATTTGTCAGTT----AATATTATTATTATTATTATTATTATTATTATTATTAT-----------------TATTATTATTATTATTATT
Consensus

Teleogryllus_occipitalis        -------------------------------------------------------------------------
Tarbinskiellus_portentosus      TGTGTGCGTCTCAAGAAACA----GAGTTTCTGCAGCAGAGCTAGAAAGCGGCGCAGGAATGCACGCCCGTTG
Acheta_domesticus               CATATTTATATTGAAATATACCTTCATCCTTCACTTC-------------------GTAATACGTTGTCGTG-
Gryllus_bimaculatus             -------------------------------------------------------------------------
Gryllus_longicercus             ATTATTACTATTATTAT-TACTATTAGTTTGCAC---------------------------------------
Consensus
```

**Tocc.5G0008120.1_exon1**

```
Teleogryllus_occipitalis        -------------------------------------------------------------------------------------------------------------------------
Tarbinskiellus_portentosus      --------------------------------------------------FPTDFK---------------------SNVYEYEEVEDRVTNI*LRNMAV*KN*NEKIE----------
Acheta_domesticus               -----------------------------------------------------NFN--------------------------HCKLLYFYLPEALRSPFY*NNFQK*FF----ALSLA*
Gryllus_bimaculatus             SA*MLHSYI*YLLCLTIRY*AFISKFYIMELSILKMYA*I*KFEHVCKSKFDVIFN--------------------------HSQLSYYYRLETLRPLLH*IDLNKSVH----------
Gryllus_longicercus             -------------------------------------------------KLDPVFNPKVVSPGGMGDEIMI***WG*KDNTYRHTQTHKYTH---ARGGIRTRDLLVRDHSGTTALAISA
Consensus

Teleogryllus_occipitalis        -------------------------------------------------------------------------------------------------------------------------
Tarbinskiellus_portentosus      --------------------------------MSLVTELPLTGSRRGVPTPQEIPPC-------YRYVFF--FFFEKKVNIYDFFRVYI----IMPYSLS*LFQ*SLELIIPEQERN*-R
Acheta_domesticus               HCS*---------------------------INVFSVF-----RRSF----------YFIFLAYSSPLK*NLGFTLNTS*K*MFKL*V---DFHCHYTSKIC*-K*EL*VWHMNFNTLH
Gryllus_bimaculatus             ---------LLLVIIIIIIIIIIIIIIIIIIIVNVFSVF-------------LRSC-HCILLAHTYLF-------DVKNSKKLV*LAM-DRSVFCACFSKFCK*NLRFTIHETTHNN--
Gryllus_longicercus             I*SALLLLLLLLLLLLLLLLLLLLLLLLLLLLLLLLLTYF-------------LFSCDHFIVFY*PLLIY--LTSKTVKNLCDMLWIEVYSVHVFRNSVSKI*D---SLYMQQLTTKNI*
Consensus

Teleogryllus_occipitalis        -------------------------------------------------------------------------------------------------------------------------
Tarbinskiellus_portentosus      IFQ---------------------------------------DKGIFALFFYISTK*--YF*EFIIFLNVLSIMRD**CKIYF-ILYYIINF*I-------------C*FIG-RLGWL
Acheta_domesticus               I--------------KLTSEIV--------------------------VFVFMNKNVNDIIYDVLIFLHT---------NIF*LPLFFASRRNL--VTLPKHNVNLVTVDFHTPLG--
Gryllus_bimaculatus             ----KYLNV*TKLIASKLHCHIFKTRYNLLEL*VCHIHSEHYTRS*GVK*RFPFL*PKM--LVI*FIIY------------*YFCILNHVLTASLLYV*LPKQYVTLLCIFIQLEMKVI
Gryllus_longicercus             MFKLN*LPVNSIATFSKLVTIC*NYEFATFILSITQYAEE*NT-------GFRFYDQK----VI*FMIY------------*YFCILYYVLTASLLRF*LPKQYVTLPCIFIHLEMEVI
Consensus

Teleogryllus_occipitalis        -------------------------------------------------------------------------------------------------------------------------
Tarbinskiellus_portentosus      *EINRIILAVE*LFFVAHFARSSFY---------------ANLRNEIKKK---LRTITGKMY-VGDMCIRNGY*RV*EQET*RENQCNRKQCKRKECKYI----QCECKERKLIERMQIK
Acheta_domesticus               ----------NGQL*IP-YSEYNFKFETQIKD*CFNENQGQNIR-PLFPENIPLFSAHGQFSFLRTTSIRNLF----------------------CKS*IFKRQKNYGSLSVLATCK*K
Gryllus_bimaculatus             NTIN*I------*FQVPNSNQTSFKI-----------HNCNIK-EIVSS---IHSIYGAL----------------------------------TKAL----GIDCGRK*LTVTIQIL
Gryllus_longicercus             NTIN*I------*FRVPNSNKTSFKI-----------QNCNLK-EIVSS---IHSIYGVL----------------------------------TKAP----GIDCGRK*ITVTVQIL
Consensus

Teleogryllus_occipitalis        -----------------------------------------------------------------------------------------------------MNNI-----------
Tarbinskiellus_portentosus      ---------------------------------------------------------------------------------SVSE-LIR*NI*ILLLKFI*KIKFKFEANIENS*
Acheta_domesticus               IGGN*SAEYFCRNCIFYSLKT*DYNKRA*D*LWGEMAHDNGSNFREREEEMEYFLGCIK*TVRRY*MCLLFPK*FIINKQTYSRILSVSIIIE*QVQ*SFLPRIM*NV-GTYNA------
Gryllus_bimaculatus             EGG---------VIFWIHKV---NGRTD---------------------------ID*T------CF-LPKCFISNKQIYC*ILSVSL-VVQQVL*SLLLHIINNIVGTSNA------
Gryllus_longicercus             -GG---------VIFWIHKV---NARID---------------------------ID*T------FFYLNVLSVINKQIYS*ILSVTVFVVQQVL*SLLLHIINNIVGTSNARTHTHT
Consensus

Teleogryllus_occipitalis        -------------------------------------------------------------------------------------------------------------------------
Tarbinskiellus_portentosus      RLLMFNVNI*VMRSLYGRQSETKTHVLK----------------I-------*KICFFITLPEICM*NYITLFACPISFLYYVNEQIKNAI------ILTFFCA*RH*TSP*NINQNPLH
Acheta_domesticus               -----------VRA-HARTHVTRTHTRTLILLLNLLKIPLINNIIFSSI*NCIHS*VI*RLPPHPRRIYFSNLS-PLN------RKAPNDIRRERPRSCAL*CAPRVYCCL*NAIRRTLN
Gryllus_bimaculatus             --------------------HTRTHTYTNILLR-----PTKNTII-------K*FYLFVNLKLHSFLSYLT----PIT------------------------------TSLQNIFRTDLP
Gryllus_longicercus             ------------------HTHTHTHTYTNILLR-----PTKNTII-------K*FYLFVNLKLHSFLSYST----PIT------------------------------TSLQNIFRTYLH
Consensus

Teleogryllus_occipitalis        -------------------------------------------------------------------------------------------------------------------------
Tarbinskiellus_portentosus      SITKLEMIS----------AVDVHGI--------------------------TRSDAPRQFTAACKTQH----GEH*IPTRNVAPVHLPY*VGAAETLRTRAARSS*QRRRRRGSHWAR-
Acheta_domesticus               PDTKRRSGAFPLLSPRGGGAAPAHALQSGRRAVPHRALTPGRTPCRAAPRTHSRAD-----VVPCRTAYAL*GGRRAVPHRALT----PGRTPCRAAPRTHSRADVVPCRTAYAL*GGRR
Gryllus_bimaculatus             SITKLQTIS----------AANVHEV--------------------------ARCDAALEFTAPCKTQY----GAH*IPTRNVALVHFPY*VPASEAPHTHSRRRSAAVRASKGRRGGRG
Gryllus_longicercus             SIAKLQTIS----------AANVREV--------------------------ARCDAALEFTAPCKTQY----GAH*IPTRNVALVHFPY*VPASEAPHTHSRRHSAAVRASRGRGGGRE
Consensus

Teleogryllus_occipitalis        -------------------------------------------------------------------------------------------------------------------------
Tarbinskiellus_portentosus      ---------------AC-ACVRSRSAKG---QGSR----*SGGRRAAPD-------RAVTAVL--GDCGRGANGGGWNE---GG------------GGSG*GGGSAVTGW----------
Acheta_domesticus               AVPHRAR---TPGRTPCHAAPRTHSRAD--AVPCRTAHALQGGRRAVPH-RAR-------------TLGRTPCRAAPRTRSRADAVPCRTAH---------------------------
Gryllus_bimaculatus             KTVAATRACDAGGQWACGGC*LGRGAGGVRSRGARVA-EAEARRRRRRAVRWRWRLRGGAAWRGNGGVGKRRAGGGGNER*WGGRC*CD*AADRVVSASGVWRSRWQTRWRRGEGRCGNG
Gryllus_longicercus             G--------------------GSGGGG-----------AAAGRRRR---RWRVVVTAAVAIV---TVAADASCGGCN-------CSC-----------GV------RGWVVGETVVLDR
Consensus

Teleogryllus_occipitalis        -------------------------------------------------------------------------------------------------------------------------
Tarbinskiellus_portentosus      ---WLLLL-------VLGGG--------GGGGGGG-----GGGGGSCW-----------------------------------------------------TIENKVEVIMHL*KSRWEQIAY
Acheta_domesticus               ---------------ALQGGRRAVP-----------HRARTLGRTPCR------------AAPRTRDAAPPHTGV-----FC-----------------------------KKSWPKVEK
Gryllus_bimaculatus             DGRWRMPVAEVCNWVVCSGGRL*WSVVLGLACWCSWWRRRGGGGGVVWWWCGGSVVVGM*MLVR*RKCVRA*WVVSVCVCVCVCVCVCVIVVDCVVWLVVV----VCVVITCG*W*HVVH
Gryllus_longicercus             *LRW-----------TCGG--------------CG-----GGGGGGMW--------------MRVR*VVVREWRVVVCVCVC----------DCCGLRSVAGGSGVCVVITCGWWSHVGR
Consensus

Teleogryllus_occipitalis        -------------------------------------------------------------------------------------------------------------------------
Tarbinskiellus_portentosus      SN--NSDMFGSLGKG*NDVKGRSIINSGIDNHFAIIIG-------------IMNV*-------MW**R*LVKTTIAIFEPLFFWFVISLVGRHKWYLSNVDLNVVLIMVRVAKVDDYNVI
Acheta_domesticus               KN-------------FNIR---------EN*HYFL*V-EYI---------------------LRHTTVIC*SQASI-------FVVASV*------------------------SESLY
Gryllus_bimaculatus             SCVLVMVMYGWW----------RCMFGGEENE*VVV*ACEWW*WYRRRKDVVE*VVVLACKWR*WYERVLCITVVVV---------VVVVVG-CCWLL--VVLVVLVVLVVLA-----CFV
Gryllus_longicercus             SGVWMIVMCGWW----------RCVSGGEENE*VVV*ACEWW---------------------*WYERVLCMTVVVV---------VVVVV----------------------------V
Consensus

Teleogryllus_occipitalis        -------------------------------------------------------------------------------------------------------------------------
Tarbinskiellus_portentosus      LLTGIVK*LRI------------------------LNNIS*RALVCMCLYICLFFLSGKH*FKWCFLLGKR*SAFLISLYRITSFPD-----------------RI---------FPSLYS
Acheta_domesticus               ALLEFATSRSI---------------QRDPLFAL*VHSNS*SVLIRHG------FRSSRHILCLKLKTGP----FNIIIYII*EFLASFTFTIPV-------------------------
Gryllus_bimaculatus             M*TDEGGDM*IGTERCD----------EETM--LQTDSDVKEALIM*C---------GDDGVVVEGQ*SGRVA--LVARQRVVVMAVAGSEAAPVGTESKRWRRRQGRHARW--------
Gryllus_longicercus             VVVGGGGGVSV---FCDVDG*RW*YVNRDGE--VR*RNDAAE***C*G---------GVHHVMW**WSGGRGS--VVWQSGIGSAAKSGGDG*G----------RIGGGASWDGGQKMAT
Consensus

Teleogryllus_occipitalis        -------------------------------------------------------------------------------------------------------------------------
Tarbinskiellus_portentosus      LSSKLLLECGGFPVT*IDLHLRPNTQNASVLTIC*MKDALTNGFRIIKSFHKLFILRRHR*YYLCASQET--EFLQQS--------------------------------*KAAQECTPV
Acheta_domesticus               --------CA--PVG----GHGGGREFGDGET----------------EF-------LCNYIFFCLS-----TYRSQN--------------------------------FNLTNRCRPT
Gryllus_bimaculatus             -------------------GLRGGDQGVEAGT---ADDGSIAGAKV-----------RVRGIE---AGRWPLEYRGRR------------------------------------------
Gryllus_longicercus             TTTTARKACAVVPV---GPGGRGGDDGGDCG*---RLHRWGKGQGS-----------RHRGVEGCVGGRW--EYRGRRGRVGTQTRLWWRY*Q*GL*RLGIHWGNNRWIFTKALQK*SGM
Consensus

Teleogryllus_occipitalis        -----------------------------------------
Tarbinskiellus_portentosus      -----------------------------------------
Acheta_domesticus               TISHFSVTYL---------VIYYIFILKYTFILHFVIRCR--
Gryllus_bimaculatus             -----------------------------------------
Gryllus_longicercus             YYGMHLVYYLSVNIIIIIIIIIIIIIIIIIIITIIIITISLH
Consensus
```

**Tocc.5G0008120.1_exon2**

```
Teleogryllus_occipitalis        --------------------------------------------------------------------------------------------------------------------------
Tarbinskiellus_portentosus      ---------------------FPTDFK----------SNVYEYEEVE----DRVTNI*L----------------------RNMAV*KN*NEKIEMSLVTELPLTGSRRGVPTPQEIP-
Acheta_domesticus               -----------IILITVNYCIFI-------------------FQRL*----DRHFI-----------EI-----------TFRNSSLH--------------------------------
Gryllus_bimaculatus             SA*MLHSYI*YLLCLTIRY*AFISKFYIMELSILKMYA*I*KFEHVCKSKFDVIFN--------------------------HSQLSYYYRLETLRPLLH*IDLNKSVH----------
Gryllus_longicercus             -------------------------------------------------KLDPVFNPKVVSPGGMGDEIMI***WG*KDNTYRHTQTHKYTH---ARGGIRTRDLLVRDHSGTTALAISA
Consensus

Teleogryllus_occipitalis        -----------------------------------------------------------------------------VAVAV--------------------------------------
Tarbinskiellus_portentosus      -----------------------------------------PC-------YRYVFFFF---------FEKKVNIYDFFRVYI----IMPYSLS*LFQ*SLELIIPE-QERN*RIFQ----
Acheta_domesticus               --------FLLLNIVLKLM------------------YFLFFGDHFILYS*PILRR*SKTWDSR*TPLKNKCLNCRLISIAT--------TLPKFVK-----------SKN---------
Gryllus_bimaculatus             ---------LLLVIIIIIIIIIIIIIIIIIIIVNVFSVFLRSC-HCILLAHTYLF--------------DVKNSKKLV*LAM-DRSVFCACFSKFCK*NLRFTIHETTHNN------KYL
Gryllus_longicercus             I*SALLLLLLLLLLLLLLLLLLLLLLLLLLLLLLLLLTYFLFSCDHFIVFY*PLLIYLT---------SKTVKNLCDMLWIEVYSVHVFRNSVSKI*D---SLYMQQLTTKNI*MFKLN*L
Consensus

Teleogryllus_occipitalis        --------------------------------------------------------------------------------------------------------------------------
Tarbinskiellus_portentosus      -------------------------------------DKGIFALFFYISTK*YF*EFIIFLNVLSIMRD**CKIYF-ILYY---IINF*I----------------C*FIG---------
Acheta_domesticus               -----------------YEFGT*ILT-------HYTLSLRVK**FSFL*TKM*MI*FMMY-------------*YFCTLTYFDCLFSLRLDEI**RCQSTTSI*LPWIFIHLLEMANYKY
Gryllus_bimaculatus             NV*TKLIASKLHCHIFKTRYNLLEL*VCHIHSEHYTRS*GVK*RFPFL*PKMLVI*FIIY-------------*YFCILNH---VLTASLLYV*LPKQYVT---LLCIFIQL--------
Gryllus_longicercus             PVNSIATFSKLVTIC*NYEFATFILSITQYAEE*NT-------GFRFYDQK--VI*FMIY-------------*YFCILYY---VLTASLLRF*LPKQYVT---LPCIFIHL--------
Consensus

Teleogryllus_occipitalis        ---------------------------------------------------------------AVAAVTAVAAAA---------------------------------------------
Tarbinskiellus_portentosus      -----------------RLGWL*EINRIILAVE*LFFVAHFARSSFY---ANLR------NEIKKKLRTITGKMYVGDMCIRNGY*RV*EQET*RENQCNRKQC-----------KRKE
Acheta_domesticus               HTLNIISSLKHK*KISASMKIKDRIYVLCFQKISPYFLPMANFPSLE----RLR*GIYFVNHEFSKDRKIMVASL-----CLR----RVNK----R*EEINQRSIFAVIASSIH*KHEIT
Gryllus_bimaculatus             -----------------EMKVINTIN*I------*FQVPNSNQTSFKIHNCNIK--------EIVSSIHSIYGAL---------------------------------------------
Gryllus_longicercus             -----------------EMEVINTIN*I------*FRVPNSNKTSFKIQNCNLK--------EIVSSIHSIYGVL---------------------------------------------
Consensus

Teleogryllus_occipitalis        AAAAGV-----------------------------------------------------------AAVAAVAAVVV------------------------------------------
Tarbinskiellus_portentosus      CKYIQCECKERKLIERMQIK----------------------------------------------SVSE-LIR*NI*ILLLKFI*KIKFKFEANIENS*RLLMFNVNI*VMRSLYG
Acheta_domesticus               IKELRIDCGGKWLTIMVQILERERKRWSIFLDA*SKR*DDIECVCYFLNDLSLINKHTVEFFQLVSS*NSKSNEVFYHV*CKML---------------------------EPTMRCA--
Gryllus_bimaculatus             TKALGIDCGRK*LTVTIQILEGGVIFWIHKVNGRT----DID*TCF-LPKCFISNK------QIYC*ILSVSL-VVQQVL*SLLLHIINNIVGTSNA-----------------------
Gryllus_longicercus             TKAPGIDCGRK*ITVTVQIL-GGVIFWIHKVNARI----DID*TFFYLNVLSVINK------QIYS*ILSVTVFVVQQVL*SLLLHIINNIVGTSNARTHTHT-----------------
Consensus

Teleogryllus_occipitalis        ---------------------------------------------------------------------------------------AVAAAAAAAAAAAVVAVAVAAV----------A
Tarbinskiellus_portentosus      RQSETKT-HVLK-----------I*KICFFITLPEICM*NYITLFACPISFLYYVNEQIKNAIILTFFCA*RH*TSP*NINQNPLHSITKLEMISAVDVHGITRSDAPRQFTAACKTQHG
Acheta_domesticus               -RTHART*HVHTRA--R*SYS*IY*KYH**IIL---SSLQYET------------------AFIRRLFDAYHH-THAEYIFRTYLHSIAKLQTTSAANVREVARSDAPLEFTAACKMQYG
Gryllus_bimaculatus             ---HTRT-HTYTNILLRPTKNTIIK*FYLFVNLKLHSFLSYLT----PIT-----------------------TSLQNIFRTDLPSITKLQTISAANVHEVARCDAALEFTAPCKTQYG
Gryllus_longicercus             -HTHTHT-HTYTNILLRPTKNTIIK*FYLFVNLKLHSFLSYST----PIT-----------------------TSLQNIFRTYLHSIAKLQTISAANVREVARCDAALEFTAPCKTQYG
Consensus                                                                                                                      *          *

Teleogryllus_occipitalis        AAAVAVAAVAAV-------AAAA----------AAAVAA-----------------------------------------------VVVAVA------------------------VA
Tarbinskiellus_portentosus      EH*IPTRNVAPVHLPY*VGAAET--------LRTRAARSS*QRRRRRGSHWA--RAC---------ACVRSRSAKGQGSR-----------*SGGRRAAP-------------------DR
Acheta_domesticus               EH*IPTRNVAPAHFPY*VLAAEAPPPRTHSRADAVPCRTAHSLQGGRRAVPH--RARTLGRT----SC---RAAPRTHSRADAVPCRTAHSLQGGRRAVPH-RARTLGRTSCRAAPRTHS
Gryllus_bimaculatus             AH*IPTRNVALVHFPY*VPASEAP--HTHSRRRSAAVRASKGRRGGRGKTVAATRACDAGGQWACGGC*LGRGAGGVRSRG-----ARVAEAEARRRRRRAVRWRW-----------RLR
Gryllus_longicercus             AH*IPTRNVALVHFPY*VPASEAP--HTHSRRHSAAVRASRGRGGGREG---------------------GSGGGG---------------AAAGRRRR---RWRV-----------VVT
Consensus                              **         *

Teleogryllus_occipitalis        VAVAVA---AVAVAVACGGGG-----------------------GG-----------------------------------GGG------------------------GGGGGTVW-
Tarbinskiellus_portentosus      AVTAVL--GDCGRGANGGGWNE---G---G-----------GGSG*GGGSAVTGW------------WLLLL-------VLGGG------------GGGGGGG-----GGGGGSCW-
Acheta_domesticus               RADAVP--CRTAHALQGGRRAMPHRGRTLGRTPC-RAAPRTHSRADAVPCRTAHAL*VGRRAVPHR--------------ARAPGRTPCRAAPRTHSRADAVPCRTAHAL*GGRRAVPH
Gryllus_bimaculatus             GGAAWRGNGGVGKRRAGGGGNER*WG---GRC*CD*AADRVVSASGVWRSRWQTRWRRGEGRCGNGDGRWRMPVAEVCNWVVCSGGRL*WSVV-----LGLACWCSWWRRRGGGGGVVWW
Gryllus_longicercus             AAVAIV---TVAADASCGGCN----------CSC-----------GV------RGWVVGETVVLDR*LRW-----------TCGG-------------------CG-----GGGGGGMW-
Consensus                           *            *                                                                                          **

Teleogryllus_occipitalis        -------------LRRR*-----------------------------------------------------------------------------------------------------
Tarbinskiellus_portentosus      ------------TIENKV-------------------------------------EVIMHL*KSRWEQIAYSN--NSDMFGSLGKG*NDVKGRSIINSGIDNHFAIIIG-----------
Acheta_domesticus               RARG--------TPRRRT-------------------------------------------------------------------QGYFVRKAGQKSKKKILTY--------EKT
Gryllus_bimaculatus             WCGGSVVVGM*MLVR*RKCVRA*WVVSVCVCVCVCVCVIVVDCVVWLVVV----VCVVITCG*W*HVVHSCVLVMVMYGWW----------RCMFGGEENE*VVV*ACEWW*WYRRRK
Gryllus_longicercus             -------------MRVR*VVVREWRVVVCVCVC----------DCCGLRSVAGGSGVCVVITCGWWSHVGRSGVWMIVMCGWW----------RCVSGGEENE*VVV*ACEWW-------
Consensus

Teleogryllus_occipitalis        --------------------------------------------------------------------------------------------------------------------------
Tarbinskiellus_portentosus      --IMNV*-------MW**R*LVKTTIAIFEPLFFWFVISLVGRHKWYLSNVDLNVVLIMVRVAKVDDYNVILLTGIVK*LRI----------------------LNNIS*RALVCMCLYI
Acheta_domesticus               NTISFKLNI-----YYVIRQLSASLRLV-SSL*HQSKVSRCTRC*NLLPAVQYKGILYSLCEC----IQIREVY*FGTDLEVPGIYC-----------------V*N*KLAHLTLLF---
Gryllus_bimaculatus             DVVE*VVVLACKWR*WYERVLCITVVVV--------VVVVVG-CCWLL--VVLVVLVVLVVLA-----CFVM*TDEGGDM*IGTERCD----------EETMLQTDSDVKEALIM*C---
Gryllus_longicercus             --------------*WYERVLCMTVVVV--------VVVVV----------------------------VVVVGGGGGVSV---FCDVDG*RW*YVNRDGEVR*RNDAAE***C*G---
Consensus

Teleogryllus_occipitalis        --------------------------------------------------------------------------------------------------------------------------
Tarbinskiellus_portentosus      CLFFLSGKH*FKWCFLLGKR*SAFLISLYRITS-----------------FPDRI---------FPSLYSLSSKLLLECGGFPVT*IDLHLRPNTQNASVLTIC*MKDALTNGFRIIKSFH
Acheta_domesticus               --------------------------------------------T*FKNF---------W---PALRSL----------YPY----------------------------------------
Gryllus_bimaculatus             ------GDDGVVVEGQ*SGRVA--LVARQRVVVMAVAGSEAAPVGTESKRWRRRQGRHARW--------------------------GLRGGDQGVEAGT---ADDGSIAGAKV-----
Gryllus_longicercus             ------GVHHVMW**WSGGRGS--VVWQSGIGSAAKSGGDG*G----------RIGGGASWDGGQKMATTTTTARKACAVVPV---GPGGRGGDDGGDCG*---RLHRWGKGQGS-----
Consensus

Teleogryllus_occipitalis        --------------------------------------------------------------------------------------------------------------------------
Tarbinskiellus_portentosus      KLFILRRHR*YYLCASQET---EFLQQS---------------------------------*KAAQECTPV-------------------------------------------------------
Acheta_domesticus               ---------------VHRWEAMEAEGNSAMVRPN-----------------------------------------------FYVITYFFVFLLTVLKILI*RIVAVLPLYHIFPLHIW*YITYL
Gryllus_bimaculatus             ------RVRGIE---AGRWP-LEYRGRR---------------------------------------------------------------------------------------------
Gryllus_longicercus             ------RHRGVEGCVGGRW---EYRGRRGRVGTQTRLWWRY*Q*GL*RLGIHWGNNRWIFTKALQK*SGMYVGMHLVYYLSVNIIIIIIIIIIIIIIIIIIIIITIIITISLH--------
Consensus

Teleogryllus_occipitalis        ---------------
Tarbinskiellus_portentosus      ---------------
Acheta_domesticus               Y*NIPSSFTS*YVVV
Gryllus_bimaculatus             ---------------
Gryllus_longicercus             ---------------
Consensus
```

**Tocc.5G0008290.1**

```
Teleogryllus_occipitalis      TCAATTGTATGAATATGATTCAAGTTCTTCAATTA------------------------------ATTTATTCTGGAAGATTGAAAACGTACATTCCAAATACATCCGAAATTAATATTA
Acheta_domesticus             TCAACTGTACGAAGAAGACTAAAATACTCCATTTACCAATACCACACCTAAAATATCAATTGGTTGTTTATTCTGGAAGTTTGAAAACGACCATCGAAAAAACATTCAAAATCAATATTA
Gryllus_bimaculatus           TCAATTGTATGAAGATGATTAAAATTCTTTATCAGTCG---------------------------ATTTATTCTGAAAGTTTGAAAACGAACATTCCAAAAACATACGAAATAAATTTTA
Gryllus_longicercus           TCAATTGTATCAAGATGATTAAAATTCTTCATCAGTTG---------------------------ATTTATTCTGAAAGTTTGAAAACGAACATTC------CATACGAAACGAATATTA
Consensus                     **** ****  ** * ** * ** * **  *                                   ********* *** *********  ***         *** * ***  *** ***

Teleogryllus_occipitalis      CCAATA------------------------------------------------------------------------------------------------------------------
Acheta_domesticus             CCGGAA------------------------------------------------------------------------------------------------------------------
Gryllus_bimaculatus           CCGAGATGCAATATGTGATCAAAAGTATCCGGACATGTCCAAAATGATGTGATGGTGTGGTTGTGTTTTTCATGGACAGGGCTTGCACCCCTTGTTTTGTGTTGCACACTCACGGCACAG
Gryllus_longicercus           CCGAGATACACTATGTGATCAAAAGTATCCGGAAACGTCCAAAATGGCGTTATGGTGTGGCAGTGTTTTTCATGGAGAGAGCTTGCACCCCTTGCTTTGTGCGGCACTATCACAACTCAC
Consensus                     **   *

Teleogryllus_occipitalis      ------------------------------------------------------------------------------------------------------------------------
Acheta_domesticus             ------------------------------------------------------------------------------------------------------------------------
Gryllus_bimaculatus           GCCTACATTGAAATTTTGAACAC---CTTGCTTTCCACTGTTGAAGAGCAATTCGGGGATGGCGA-------TTTCAATTCAATTGAGCCCCTGTTCATACTAAACGGCCTGTGGCGGAG
Gryllus_longicercus           GCCTACGTTCGTGTTTTAAACACCTTCTTGCTTTCCTCTGTTGAAGAGCAATTCGGGGATGGCGATTGCCTCTTTCAACACGATTGAGCACCTGTTCAAACTACACGACCCGTGGCAGAG
Consensus

Teleogryllus_occipitalis      ------------------------------------------------------------------------------------------------------------------------
Acheta_domesticus             ------------------------------------------------------------------------------------------------------------------------
Gryllus_bimaculatus           TGATTCCACGACAATAACATCCCTGTAATGGATTGTCCTGCACAGAGTCTTGACTTCAATCCTATCGAACACCTTTGGGATGTTTTGGAACGCCGAATTCTTGCCAGGCCTCACCGACCT
Gryllus_longicercus           TGGTTCCACAATAATAACATCCCTGTAATGGATTGTCTTGTACAGAGTTCTGACTAGAATCCTATCGAACACCTTTGGGATGTTTTGGAACGCTGATTTCATGCCAGGCCTCATTGACCT
Consensus

Teleogryllus_occipitalis      ------------------------------------------------------------------------------------------------------------------------
Acheta_domesticus             ------------------------------------------------------------------------------------------------------------------------
Gryllus_bimaculatus           ACAACGATACCTTCCCTTCAGTGTAGCACTCCGTGAACAAAGGGCTGCCATTGCCCAAGGAACCTTCCAGCACCTGATTG--AGTATGCCTGCGAGAGTGGAAACTGTCATTATGGCTAA
Gryllus_longicercus           TCAACGATACC-TCTCTTCAGTGCAGCACTCCGTTAAGAAAGGGCTGCCATTGCCCAAGAAACCTTCCAGCACCTGATTGAAAGTATGTCTGCCGGAGTGGAAACTGTCATCATGGCTAA
Consensus

Teleogryllus_occipitalis      ----------------------------------------------------------------------------------------------------------TAAGCGAAGTTTGACA
Acheta_domesticus             ----------------------------------------------------------------------------------------------------------TATTCGAAGTTTGACA
Gryllus_bimaculatus           GAATGCTCCAACACCCTATTGAATTCCAGAATTACCAATGGAGAGCGCCACGAACTGGAAAGTCATTTTCAGCCACGTGCCCGGATACTTTTGATCTCATAGTGTATTCGAAGTTTGGCA
Gryllus_longicercus           GAGTGGGCCAACATCCTATGGAATTCCAGAATTACCAATGGCGAGCGCCACGAACTGGAAAGTCATTTTCAGCCACGTGTCCGGATACTTTTGATCTCATAGTGTATTCGAAGTTTGGCA
Consensus                                                                                                                         **  ********* **

Teleogryllus_occipitalis      GAACATCAATCTAAATATCGGAAAAGAAAAA---TTCCTGGATGAAATTACGTTAAT--AAGTAGTTGAAAGTAAATATTCTCATAATGTTCTAATAGAGATAATGAAATCGTAGCATTG
Acheta_domesticus             GAACATCAATCCAAATATCAGTAAAGACAGATGGTTTGTGAATCAAATTACGTTAAT--AAATATTCTAAAA---------------ATTCTAATAGAAGTGATGAATTGAAAATAGAA
Gryllus_bimaculatus           GAACATCACTCTAAATGTCGGTAAAGAAAGATG-GTTGTAGATGAAATCACATTAATGGAAGTGCTGGAAAGTATATATTCTGAAAATTCATCAATAGTGATGATGAAATCGTAACATTG
Gryllus_longicercus           GAACATCAGTCTAAATGTCGGTAAAGAAAGATG-GTTGTAGATGAAATCACATTAATGGAAGTGCTGGAAAGTATATATTCTGAAAATTCATCAATAGTGATGATGAAATCGTAACATTG
Consensus                     ******** ** **** ** * ***** * *    *  *  ** **** ** *****  ** *  *  ***                      *****   * ***** *   *  *

Teleogryllus_occipitalis      GTG---------------------------------------------------------------------------------------------------------------------
Acheta_domesticus             GTGATGATCATTAACTTCTTCTTCTTCAAGCATTGACCGTAGATCCGTTGCGTACTCTTTGATCTGACACCTTGTTTTCGATCGTCCAACACACCGATTTCCTTTAGCATGGTATTGTGT
Gryllus_bimaculatus           GTT---------------------------------------------------------------------------------------------------------------------
Gryllus_longicercus           GTT---------------------------------------------------------------------------------------------------------------------
Consensus                     **

Teleogryllus_occipitalis      ------------------------------------------------------------------AATTCGTTCATTTCATTTCA-------------------------GTAAAGAA
Acheta_domesticus             GACTTCTTTTGATGTTCTGGAATTTTCCATTCTGTACGTCGTTTGCCTATACGCTCCTGTTTCTTTTATTTTTTTTTCTTCATTTTAATTAAATATTCCTAACTGATTTTGCGTAAAGAA
Gryllus_bimaculatus           ------------------------------------------------------------------TCATCCGTCCCTTTAATTGA-------------------------GTAAAGAA
Gryllus_longicercus           ------------------------------------------------------------------TCATCCGTCCCTTTAATTGA-------------------------GTAAAGAA
Consensus                                                                                            *   *   ** * ** *                        ********

Teleogryllus_occipitalis      GTTTAGAAGCAAAATAAACAGAAT---ATAAAACAAGGAAATATAATGGAAAGTACTTGGGTGCTGATTAATACGGTTATCATTAGTA-TACATCACATTACGGCTCGATGCTAGTGGCT
Acheta_domesticus             ATTAATATGCAAAATAAAAAGGGTTTAAAAAAACAATGAAATATAATGGAAAGT-CTTCGGTGTTGATTAATATGTGTATCGTTTACAGCACATCACATCATGGTTAAGTGTTATTGGCT
Gryllus_bimaculatus           GTTTGTAAGCGAAATAAATAGAAT---GAACAACAATGAAATATAATGAAAAGT-ATTTGGTGGTGATTAATATGAGGAC-----------------------------------------
Gryllus_longicercus           GTTTATAAGCGAAATAAATAGAAT---GAACAACAATGAAATATAATGGAAAGT-ATTTGGTGGTGATTAATATGAGTAC-----------------------------------------
Consensus                      **   * ** ******* **  *     * ***** *********** *****  ** **** ********* *   *

Teleogryllus_occipitalis      ATTGAAAGTACCTATGCTTTAAGTTCGGTAAAATCGATGTAAACGTTAGATTATTCCGCCGCGCTGCCTTTAGTATCGCAAGATCCGTAAAAGTTTTAAGCAATTACAGAGCCACATCGT
Acheta_domesticus             A---AAGGTACCGATAC--TAGGTTC------------GTAAAAATTACTTTATTCCGCCGGGATCCCTCTAATACCGCAAAATCCGTAAACGTTTCAAGCCATTTCAGCGCTA----GC
Gryllus_bimaculatus           -------------------TAGGTTC------------------------------CGCCGAGATCCCGCAAGTATCGCAAAATTCGCAAACGTTTCAAGCAATTACAGAGGTACACAGC
Gryllus_longicercus           -------------------TAGGTTC------------------------------CGCCGAGATCCCTCAAGTGTCGCAAAATCCGTAAACGTTTCAAGCAATTACAGAGGTACACAGC
Consensus                                        ** ****                                 ***** * * **   * *  ***** ** ** *** **** **** *** *** *  *    *

Teleogryllus_occipitalis      CACGCATGCAGGCCATTCGACAGCCACA-------------CGGCACGCGGCACGCG---------------------------------------------------------------
Acheta_domesticus             CACGCATGCAGGCCATTCGACAGCCACATGGCA--------CCGGCACGCGGCACGCA---------------------------------------------------------------
Gryllus_bimaculatus           CACGCATGCAGGCCATTCGACAGCCACATGGCACGCGGCACGCGGCACGCGGCACGCG---------------------------------------------------------------
Gryllus_longicercus           CACGCATGCAGGCCATTCGACAGCCACATGGCACGCGGCACGCGGCACGCGGCACGCGGCACGCGGCACGCGGCACGCGGCACGCGGCACGCGGCACGCGGCACGCGGCACGCGGCACGC
Consensus                     ****************************              ***************

Teleogryllus_occipitalis      -GTACGTCGGCCGCAA------------------------CGGCGGAGGCGCGTGGGCGGGTGACGCAGATGCCGACGACGCCCCACTTGGGCGGCACAGAAGCGAGCGCGGGCGGGCG
Acheta_domesticus             -GCACGTCGGCCACAACGGCGG---------TGGCGGCGGGCGGCGGCGGCGCGTGGGCGGGTGACGCAGATGCCGACGACGCCCCACTTGGGCGGCACAGAAGCGAGCGCGGGCGGCTG
Gryllus_bimaculatus           -GCACGTCGGCCACAA---------------CGGCGGCGG-CGGCGGCGGCGCGTGGGCGGGTGACGCAGATGCCGACGACGCCCCACTTGGGCGGCACAGAAGCGGGCGCGGGCAGCCG
Gryllus_longicercus           AGCACGTCGGCCACAACGGCGGCGGTGGCGGCGGCGGCGG-CGGCGGCGGCGCGTGGGCGGGTGACGCAGATGCCGACGACGCCCCACTTGGGCGGCACAGAAGCGGGCGCGGGCGGCCG
Consensus                      * ********* ***                          ****** ********************************************************** ******** *  *

Teleogryllus_occipitalis      GGCGGGCGGGCGGGCGGGGGCGCCACACGCCTCCGGAGCAGGTCCCGACCGGCGCGACCCGGCGGCGGCGGCA---ACGGGCGCGCCAAATGTGGTGGTGGTGGTAGAGGGGCCTGCGGA
Acheta_domesticus             GGCGAGCGGGCGGGCGGGGGCGCCACACGCCTCCGGAGCAGGTCCCGACCAGCGCGACCCGGCGGCGGCGGCTACGGGCAAGGCGAATGTGGAGGCGGTG---GAAGGGCCTGGGGA
Gryllus_bimaculatus           GGCGGTCGGGCAGGCGGGGGCGCCACACGCCTCGGGAGCAGGTCCCGACCGGCGCGACCCGGCGGAGGCA---ACGGCCACGGCAAACCTGGCGGAGGTA---GAGGGGCCTGA---
Gryllus_longicercus           GG----CGGGCAGGCGGGGGCGCCACACGCCTCGGGAGCAGGTCCCGACCGGCGCGACCCGGCGGAGGCA---ACGGCCACGGCAAACGT-GCGGAGGTA---GAGGGGCCTGAGGA
Consensus                     **    ***** ********************* *************** **************** ***    **** *  * * **  * * ** ***    ** *******

Teleogryllus_occipitalis      AGCGGGGGCGGTGGTACTGGGAGCGTTAAAGGTGGAGATGCAGTTCAAGGTGGCTGCAGGGGCGTGAGGGGGAGGATGATGGTGGCAGGC-AGGCGGTGGGTGAAGGTGGTGGAGGTGGC
Acheta_domesticus             GGCGGGGGCGGT-----TGGTAGCGTTAAAGGTGGAG-TGCAGATGAAGGCGGC----------GGAGAAGGGGAGTGATGAAGGTAGGTGAGGCGGTGG-------------------
Gryllus_bimaculatus           ---GGTGGTGTTGGAGTTGCGAGCGTTAAAGATGGAGATGCAGATGAAGGTGTCAGCGGGGG--AAGAGGGGGGGATGATGAAGGAAGG--GGGCGGTGGG------------------
Gryllus_longicercus           GGCGGGGGTATTGGGGTTGGGAGCGTTAAAGATGGAGATGCAGATGAAGGTGTCAGCGGGGG--AAGAGGGGGGGATGATGAAGGCAGG--GGGCGGTGGG------------------
Consensus                        ** **   *     **  ********** ***** ***** * **** * *              ** *  *****  ** ***   ********

Teleogryllus_occipitalis      GTAGGCGGAGGCGGAGTGGGAGTGGGTGGAGGAGGTGGAGGGTTTGGAGGAGACGGAGGGGTGGATAAGACAGAGGAGAACGAGGAGGCGGAGGTAACGGAGGAGGCGGATGTGACGGAG
Acheta_domesticus             --------------------------------------------------------------------------------------------------------------------GGAA
Gryllus_bimaculatus           --------------------------------------------------------------------------------------------------------------------GAAG
Gryllus_longicercus           --------------------------------------------------------------------------------------------------------------------GAAG
Consensus                                                                                                                         * *

Teleogryllus_occipitalis      GGGATGGAGGTGACGGCGGAGGCTATGGGGGCGGATTAGGTGGAGGTGGCGGAGGTGATGGAGAAGGCGGGGAGGACGGAGGGGACGGATGAGGCGGAGGAGGAAACGGAGGCGGAGGGG
Acheta_domesticus             GGGGAGGAGATGGGGACGGAGGTGACGGCGGTGGAGGAGGTGGAGAAGGCGGAAGAGACGGAGGAGGCGAGGAGGA-----------------GGAGGTGGCGAAGGAGGCGGAGGGG
Gryllus_bimaculatus           GTGGTGGTGACGGCGGTGGAGGTAACGACCACGGCAAACGTGGCGGAGGGAGAGGGGCCTGCTGAGGCGGGGGCGTTGTG----------------------------------------
Gryllus_longicercus           GTGGTGGTGACGGCGGCGGAGGCAACGGCCACGGCAAACGTGGCGGAGGTAGAGGGGCCTGCTGAGGCGGGGGCGTTGTG----------------------------------------
Consensus                     * *  ** *  *  *  *****  * *     **   * **** *  **  ** * *   *   ***** **  *

Teleogryllus_occipitalis      ACAGAGGGTACGGAGGAGGTGGAGGGGACGGAGGAGGCGAAAGGCACAGAGGAGGCGGAGGGGATGGAGGAGGCGGAGGGAACGGAGGAAGCGGTGGGGACGAAGGAGGCGAAGGGGACG
Acheta_domesticus             ACGTTGGGGGCGGAGGGGGCGGGGGGGGCGGAGGGGGCGGGGGGC----GGGAGGCGGAGGGGGCGGAGGGGGCGGAGGGG--CGGAGGGGCGGAGGGGGCGGAGGGGGCGGAGGGGGCG
Gryllus_bimaculatus           --GTGGAGAGCGTTAAAGGTGGAGATGAAGGTGGCGGCGGGCG------CGA----------------AAGGCGGGGGGC-----------------ATTGGGGAAGGTGGTGGTGACG
Gryllus_longicercus           --GTTGGGAGCGTTAAAGGTGGAGATGAAGGTGGAGGCGGGAG------CGAAGGCGGGGGGGGGATGAAGGCGGGGGGC-----------------AGTGGGGAAGGTGGTGGTGACG
Consensus                          * *  **     ** ** *  *  ** ** ****   *       *                 ***** ***                 *  *  ** *  ** * **

Teleogryllus_occipitalis      GAGGAGGCGGAGCGGGTGCAGGAGAAGGAGGAGACAGAGGGTATGGAGGAAGTGGATGTGACGGAATAGTCGGAGGTGACGGAGGAAGCGCAGGTGACGGAGGAGGCGGAGGGGTCGGAG
Acheta_domesticus             GAGGAGGCAAA---------------------------------------------------------------------------------------GGAGGCGGAGGGGGAGCTGGAG
Gryllus_bimaculatus           GCGGAGGTAAC---------------------------------------------------------------------------------------GGTAGCGGTGGAGGAGGCGGAG
Gryllus_longicercus           GCGGAGGTAAC---------------------------------------------------------------------------------------GGTAGCGGTGGAGGAGGCGGAG
Consensus                     * *****                                                                                           **  * ** ** ** *  ****

Teleogryllus_occipitalis      GAGGCGGACGGTACCGAGCAGGCAGAGGAGGCGGAGGAGGTGGAAGAAGTGGAGGGAGCGGAGGAAGCGGAGGAAGACCTGGAGGCGGCGCGGTGGCGCGGCGGCGCGGCGGCGGCAGCG
Acheta_domesticus             GAGACAG------------CGGCAGCGGGGGCGGCAACGGTG---------------------------------------------------------------ACTGCGGCGGCGGCG
Gryllus_bimaculatus           GAGATGG-----------------------------------------------------------------------------------------------------------CGGTG
Gryllus_longicercus           GAGAT-G-----------------------------------------------------------------------------------------------------------CGGTG
Consensus                     ***   *                                                                                                            * * *

Teleogryllus_occipitalis      GCGACGGTGACTGCAACGGCGGCGTCGAAGAATAGATCGTCCATCACGAGCGGGCGGCGAAGGTGGTGGCAGCGGTCGCCGCGT-GGAGCGCCACTTTCCAGCGTCTGCTCCTGCGCCGC
Acheta_domesticus             GCGACGGTGACTGCGACGGCGGCGTTGGAGAATAGATCGTCCATCACGCGCGGGCGCCGAAGGTGGTGGCATCGGTCGCCGCGTAGGAGCGCCACTTTCCAGCGTCTGCTTCTGAGCCGC
Gryllus_bimaculatus           GCAACGGTGACTGCAACGGCGGCGTCGAAGAATAGATCGTCCATCACGCGCGGGCGGCGAAGGTGGTGGCAGCGGTCGCCGCGT-GGAGCGCCACTTTCCAGTGTCTGCTCCTGCGCCGC
Gryllus_longicercus           GCAACGGTGACTGCAACGGCGGCGTCGGAGAATAGATCGTCCATCACGCGCGGGCGGCGAAGGTGGTGGCAGCGGTCGCCGCGT-GGTGCGCCACTTTCCAGTGTCTGCTCCTGCGCCGC
Consensus                     ** *********** ********** * ******************** ******* ************** *********** ** ************** ******* *** *****

Teleogryllus_occipitalis      ACGCGAATCTCCGTTACTCACCATAACACCCAGTTTTGAGCATCAAACGCTGAGTTCTTTTTAC-------------------TTTTTTCGTTACATCTCGACC----------------
Acheta_domesticus             CTGCGAATCTCCGTTACTCTCCGCAACACCAAG-TTTGAGTCTCAAACGCTGAGTCCTTTAAATCTTTTAAGTCTTTGTTATGTTTTTTACTTACATCTTGACC----------------
Gryllus_bimaculatus           ATGCGAATCTCTGTTACTCTACGTAACACCCAATTTTGAGCCTCAAACGCTGAGTCCTTTAAGT-------------------TTTTACTTTCTCACACTGACTAACGTAAGTAGAGAAA
Gryllus_longicercus           ATGCGAATCTCCGTTTCTCTACGTAACACCCAATTTTGAGCCTCAAACGCTGAGTCCTTTAAGT-------------------TTTTACTTTCTCACACTGACTAACGTAAGTAGAGAAA
Consensus                       ********* *** ***  *  ****** *  ******  ************* ****                       ****    *  **    ***

Teleogryllus_occipitalis      ---------TCGTA---CATCTCTCTCGGATATCTTGAGTAAACACATTTTATTTTTCATTTGTTTATTTTTTG----------------------------------------------
Acheta_domesticus             ---------TAGTGTTTCATCTCTCACGGATATCTTAAGTAGG---ATTTTTATTTTGCTTTGTTTAATTTTTGTTTGCTTTGTTTTTTCTGTTTGTGCGTGTTTTATGTTTGTTTCTTG
Gryllus_bimaculatus           GTATTGTAATCGTGCACCGGACACCGGCGAACGATTGAGTAGACACTTTGTATATTTTTATTTTTATATTTTTGGGGGACATTGTTTTTGTGTTTGTTTCCTTTTT-TGC---CATCTGT
Gryllus_longicercus           GTATTGTAATCGTGCACCGGACACCGGCGAACGATTGAGTAGACACTTTGTACATTTTTATTTTTATCTCTTTGGGGGGCATTGTTTTTGTGTTTGTTTCCTTTTTGTGC---GATCTGT
Consensus                              * **    *      *   **    ** ****      ** *   ***   ** **   * ****

Teleogryllus_occipitalis      --------CTTTGTTTTGAGAAA-GTTTTAAACATCAATCAAAAGCACCGTCT-------------------------------------------------------------------
Acheta_domesticus             TTTTTGACTTTTGTTTTGGGAAATATTTAAAACACCACACAAAAACGCCTTCTTCGGCATTCTGTACCCTTGGCTCTTCATGATTTGAAATAGAAATTTCGAAACATTTTCATTACAAAA
Gryllus_bimaculatus           TTTAGGACATTTGTTTTAAGAA--GTTTTAAATATCAATCAAAAACGACTTCTTCGACATTCTGTATCTTCGTCTCTCGA-----------------TTTCATACACTTTCATTGCACAA
Gryllus_longicercus           TTTAGGACATTTGTTTTAAGAA--GTTTTAAATATCAATCAAAAACGACTTCTTCGACATTCTGTATCTTCGTCTCTCGA-----------------TTTCATCCACTTTCATTGCACAA
Consensus                              ********  ***   *** *** * **  ***** *  * ***

Teleogryllus_occipitalis      ------------------------------------------------------------------------------------------------------------------------
Acheta_domesticus             GGCGAGTCATTTTGCCCGTAAAAGTTTCAACACTT-----------------AGTATTTTG---CAAACACATCTCATGTAATGTCAGGTGATACACTTAGTTACTGGAGATAAATTTTT
Gryllus_bimaculatus           GGCGAGCCATTTTGACCGTAAAACAATCAGCACTTTGGTGTAATGCACAATAAGTTGTTTGGTTAAAACACATCTTGTAGAATTCGAGGTGATG-ACGTAGTAACTAGTGATAAGATTTT
Gryllus_longicercus           GGCGAGTCATTTTGACCGTAAAAAAATCAGCACTTTGGTGTAAAGCACAATAAGTTGTTTGGTTCAAACACATCTTGTCGAATTCGAGGTGATG-ACCTAGTAACTAGTGATAAGTTTTT
Consensus

Teleogryllus_occipitalis      ------------------------------------------------------------------------------------------------------------------------
Acheta_domesticus             AAGGAATTCGATTGTCTTT-ATTGTCAAGAAGCTGGAGATTTTTCTGAGTCAAATTGGTAAGCATGCCGATAAATAACGTCTAAGAGAAATTCTACATTG-TTTGATAATAATGTCTTCA
Gryllus_bimaculatus           AACGAATTCGATTGTCTTTAATTGCCATGAATTC--AGTTTCTTCTAAGTGGAATTTATAGGTACGCCTATAAATAAAGCTAAAGGGAAATTCTAGATTGTTTTGATAGTAATTATTTCA
Gryllus_longicercus           AACGAATTCGATTGTCTTTAAGTGCCATGAATTT--AGTTTCTTC-------GTTTTATAAGTACGCCTATAAATAACGCTAAAGGGAAATTCTAGATTGTTTTGATAGTAATTATTTCA
Consensus

Teleogryllus_occipitalis      ------------------------------------------------------------------------------------------------------------------------
Acheta_domesticus             AACGGGAATTTAAAAACTTAGAAAATTGGATTGGGAACGACAAAAGACCTAAATATAAAATCCTAGAGCTACGCTACTTCGACAACCTCCTCTGCGACTAAA----AGTAACTGCAGAAG
Gryllus_bimaculatus           AATGGGAATTATAAGACTTACAGAATTGGTGTGTTAACAACAAAAGATCT--GTATGAAATCCGAGAGCGACG-ATTTTCGACAACCTCTTCTGCGATTAAAATTCAGGAACTGCAGATG
Gryllus_longicercus           GATGGTAATTATAAGACTTACAGAATTGGTGTGTTAACAACAAAAGATCT--GTATGGAATCCGAGAGCGACG-AATTTCGACAACCTCTTCTGCGACTAAAATTCAGTAACTGCAGATG
Consensus

Teleogryllus_occipitalis      ------------------------------------------------------------------------------------------------------------------------
Acheta_domesticus             AGTATTTTCAGGAGCG-CAAGGACTTATCACGTCAGCACCGACGCTTCGTGTCTCTCAGAACTGTTCACATTGCCAACAGGAGACTGGTTGCAAGGAAATTTGGAAGTTACACTAAAACC
Gryllus_bimaculatus           AGTATTTCCAGATGTG-GAAGGACTTATCACATC-----------------TCTCACTGGATTG----------CGATAGGATACTGTCTGCAATGAAATTTTGAAGTTACATTAAAATT
Gryllus_longicercus           AGTATTTCCAGATGTGCAAAGGACTTATCACATC-----------------TCTCACTGGATTG----------CGATAGGATACTGTCTGCAATGAAATTTTGAAGTTACATTAAAATC
Consensus

Teleogryllus_occipitalis      ------------------------------------------------------------------------------------------------------------------------
Acheta_domesticus             TTCCACTTCTTCA-ATTTTATGTCTTAAAGGAAAAGAATGTACTGGTAATACACTGAACTCTCATTAACGCGAATTTCAAGGGACCGAAATAATCCGAGTTCGTGTTATAAAAATTCGTA
Gryllus_bimaculatus           TTTGGTTTCATCACTTTTTAGGCCTAAATAGTAAAGAATTTATT----------------------------------------------------------------------------
Gryllus_longicercus           TTTGGCTTCATCATTTTTTAGGCCTAAATAGTAAAGAATTTATT----------------------------------------------------------------------------
Consensus

Teleogryllus_occipitalis      ------------------------------------------------------------------------------------------------------------------------
Acheta_domesticus             TTAACCGAGTATTTTGTATATCCATATTACCGTAATCCCTTGCCAGGACCGAAGTTTTTTCGTTTAACGGCGAAATTCGTGTTAACGAGCGAATTCGGGGGTAACGACGGATTACTATAT
Gryllus_bimaculatus           -------------------------------------------------------------------------------------------------------------GTTTGATAATA
Gryllus_longicercus           -------------------------------------------------------------------------------------------------------------GTTTGATAATA
Consensus

Teleogryllus_occipitalis      ------------------------------------------------------------------------------------------------------------------------
Acheta_domesticus             GTAAAAAAGGAGCAAGATGATACTGATTTTTTGTGATATAGCACATCATGTATACCAGAAAAGAAATCGTAGCTTAAATCAATTTGTTAAATTCCTGTACCATGATCATAAACTTCAAAC
Gryllus_bimaculatus           GTAAAAGAGGAGCAAGATTATAATTATTTTCTTTAG-ATGACTCATTTTGTATACCTGAAAGCAATTCGTATCTT------TTGTTCTAAATCCCTGT------TTCATAAGCTTCAAAC
Gryllus_longicercus           GTAAAAGAGGAGCAAGATTATAATTATTTTCTTTAG-ATGGCTCATTTTGTATACCTGAAAGCAATTCGTATCTT------TTGTGCTAAATCCCTGT------TTCATAAGCTTCAAAC
Consensus

Teleogryllus_occipitalis      ------------------------------------------------------------------------------------------------------------------------
Acheta_domesticus             TGAGAATGT---GTATTTTATTATTT-TTTTTTAATTATGCTT--------------------TATTTAAAAGGTCGCA-----------------------------------------
Gryllus_bimaculatus           TGAGGAAT----ATTTTTTATTGTTTATTTTTTAATTTAATTTTCGAACCTTCTGTCAATTACATATTGAGGAGGTTGCATCGTTGGTTTGAGTTCTCGACTGAGAAACTCTGACAGGTT
Gryllus_longicercus           TGAGGAATTTTATTTTTTTATTGTTTATTTTTTTATTTCATTTTCGAACCTTCTGTCAATTGCATATTGAGGAGGTTGCATCGTTGGTTTGAATTCTCGATTGAGGAACTCTGACAGGTT
Consensus

Teleogryllus_occipitalis      ------------------------------------------------------------------------------------------------------------------------
Acheta_domesticus             ------TGAACTACTAGGTCATTAGCGACCA----CATTCAAAATACAGTGTATACAATGTACA------TGAGAGCATACAACAAAATTTAAGGTTAAAGCACAAATACATAGTTTGTC
Gryllus_bimaculatus           ATCATCTGAACGAAGACTTTGTTCACGCCTGATACCACACAAAAAGCGATTTCTTTGCCTCAAAGAATCTTGAGAGAATAATATAATAATTATTGTAACA----------ATTGTTCTTG
Gryllus_longicercus           ATCATCTGAACGAAGAGTTCGTTCACGCCTGATACCACACAAAAAGCGATTTATTTGCCTCAAAGACTATTGAGAGAATAATATAATAATTATTGTAACA----------ATTGTT----
Consensus

Teleogryllus_occipitalis      -----------
Acheta_domesticus             AA---------
Gryllus_bimaculatus           ACTTCGTTAGT
Gryllus_longicercus           -----------
Consensus
```

**Tocc.5G0008290.1_exon1**

```
Teleogryllus_occipitalis     ------------------------------------------------------------------------------------------------------------------------
Acheta_domesticus            DICSK*ILFEYKIRRFSK-------LIFIK*I*NRTIDIVK*IVL----------RI*VY---YGRNL*LYPIKNFN*D--KNLF-SIG---KKCQDN--------------------I
Gryllus_bimaculatus          ------LEPSFK-KRFLRNADACLVSIF*PNFKAEE*EWIRKFKQHSCMHEKV--QLTCHLNEFSTILS*Y*YLDMKSRLFQSLYLSMQFKMQ*C*INWLQYLSVLWKDILLNFQYILYI
Gryllus_longicercus          --RSKSIE-E*R-KRF-----------------NDGEF-QTFKM-VCLNSNMS*KITVKMN-VTQNLD*NLIIEGKSR-------------K*CFDSKLHFIFILTK---LHH*Y----
Consensus

Teleogryllus_occipitalis     --------------------------------------------------ME-------------------------------------------------------------------
Acheta_domesticus            SKGSSPDGC----------------RIFYTSDTVYVEF---------RRSLEFNSSCS--DR*FSVVSPINIRRMEEYYRKK*AVGRTRSFLPLPCPIFTSHLHTLTCNTLPHTSEDA*E
Gryllus_bimaculatus          FK**INV*CIKLIY*IIYLYSHQIPSLAQIYSSIYSSFIQTSGIHIQRAGMKLLNVCDHFSKTF*W*N----------YL*MSIV*R*LKF-------FISRF-----------------
Gryllus_longicercus          ----------KFIHQFIIFYS----DIGYSYTTCWD---------------EIADVCDHFSRTF*W*N----------YL*MSIVSR*LKF-------FIS*F-----------------
Consensus

Teleogryllus_occipitalis     ------------------------------------------------------------------------------------------------------------------------
Acheta_domesticus            LVNNLKRAT--HAEI*NHRHIL-R*NMHST---NGTNDLTSSQKHQLI*SYSLNIYSP-----NLSNH*YKFISSSNLSLVSISAEFLDRHLELV*D---I*SV*PLN---------VCD
Gryllus_bimaculatus          ILKV*KRTFQKHT-----K*ILPRCNM*SKVSGHVQNDV------MVWLCFSWTGLAPLVLCCTLTAQAYIEILNT-L-LSTVEEQFGDGD---------FNSIEPLF---------ILN
Gryllus_longicercus          ILKV*KRTF--HT-----KRILPRYTM*SKVSGNVQNGV------MVWQCFSWRELAPLALCGTITTHAYVRVLNTFL-LSSVEEQFGDGDCLFQHD*APVQTTRPVAEWFHNNNIPVMD
Consensus

Teleogryllus_occipitalis     ------------------------------------------------------------------------------------------------------------------------
Acheta_domesticus            DIPHFSKN-------VLLDENIYK--SQLYEED*NTPFTNTTPKISIGCLFWKFENDHRKNIQNQYYRNIRSLTEHQSKYQ*RQMVCES----------NYVNKYSKNSNRSDELK---
Gryllus_bimaculatus          GLWRSDSTTITSL*WIVLHRVLTSILSNTFGMFWNAEFLPGLTDLQRYLPFSVALREQRAAIAQGTFQH---LIE---------YACESGNCHYG*ECSNTLLNSRITNGERH-ELESHF
Gryllus_longicercus          CLVQSSD*NPIEHLWDVLER-----------*FHARPH*PSTIPL-----FSAALR*ERAAIAQETFQH---LIESMSAGVETVIMAKSGPTSYGIP---ELPMASATNWK---------
Consensus

Teleogryllus_occipitalis     ------------------------------------------------------------------------------------------------------------------------
Acheta_domesticus            -------IEVMIINFFFFKH*P*IRC----------VLFDLTPCFRSSNTPISFSMVLCDFF*CSGIFHSVRRLPIRSCFF--YFFFFILI------------KYS*LILRKEINMQNKK
Gryllus_bimaculatus          QPRARILLIS*CIRSLAEHHSK---CR*RKMVVDEITLMEVLESIYSEN---------------------SSIVMMKS*HW---------------------FHPSL*LSKEVCKRNK*
Gryllus_longicercus          -------VIFSHVSGYF*SHS----------------VFEVWQNI-SLNV------------------GKERWL*MKSH*WKCWKVYILKIHQ****NRNIGFIRPFN*VKKFISEINRM
Consensus

Teleogryllus_occipitalis     ------------------------------------------------------------------------------------------------------------------------
Acheta_domesticus            GLKKQ*NIMESLRC*LICVSFTAHHIMVKCYWLKVPILGS*KLLYSAGIPLIPQNP*TFQAISALATHAGHSTATWH-------------RHAARSTSATT-----AVAAGGGGAW----
Gryllus_bimaculatus          N-EQQ*NIMKSIWW*LI*G-----------------------LGSAEIPQVSQNSQTFQ-----AITEVHS----HACRPFDSHMARGTRHAARGTSATT-----AAAAAARGRVTQMP
Gryllus_longicercus          N-NNE------I*WKVFGGD---------*YEY*VP----PRSLKCRKIRKRFKQLQRYT-----ATHAGHSTATWHAARG-TRHAARGTRHAARGTRHAARGTRHAARRPQRRRWRRRR
Consensus

Teleogryllus_occipitalis     ------------------------------------------------------------------------------------------------------------------------
Acheta_domesticus            -----AGDADADDAPLGRHRSERGRLGERAGGGATRLRSRSRPAR--PGGGG-GYGQ---------------------------GECGGGGRAWGGGGGW*R*RWSADEGGGEGE**R*
Gryllus_bimaculatus          TTPHLGGTE----------AGAGSRAVGQAGAPH---ASGAGPDRRDPAAEATATANLAEVEGPEV--VLELRALKMEMQMKVSAGEEGGMMKEGGGGEG--------------------
Gryllus_longicercus          RRRRVGG*RRCRRRPTW-AAQKRARAAGRAGGGATRLGSRSRPAR--PGGGGNGHGKRAEVEGPEEAGVLGLGALKMEMQMKVSAGEEGGMMKAGGGGEG--------------------
Consensus

Teleogryllus_occipitalis     ------------------------------------------------------------------------------------------------------------------------
Acheta_domesticus            VRRWGRGGDGDGGDGGG--GGGEGGRDGGGEEEEVAKEAEGTLGAEGAGGAEGAGG-----------GRRRGR----RGRRGGGAEGAEGAEGAEEAKEAEGELEETAAAGAATVTAAAA
Gryllus_bimaculatus          ------GGDGGGGNDHGKRGGGRGAC*GGG---VVVESVKG--GDEGGGG---------------------------RERR--------GALGKVVVTA-----EVTVA--VEEAEEMAV
Gryllus_longicercus          ------GGDGGGGNGHGKRGGGRGAC*GGG---VVVGSVKG--GDEGGGGSEGGGGDEGGGQWGRWW*RRR*R*RWRRRRRCGGNGDCNGGVGE*IV-----------------HHARAA
Consensus

Teleogryllus_occipitalis     ------------------------------------------------------------------------------------------------------------------------
Acheta_domesticus            ATVTATAALENRSSITRGRRRWWHRSPRR--------SATFQRLLLSRLRISVTLRNTKFESQTLSPLNLLSLCYVFYL-------------------HLDLVFHLSRIS*VGFLFCFV*
Gryllus_bimaculatus          ATVTATAASKNRSSITRGRRRWWQRSPRGAPLSSVCSCAACESLLL-------------YVTPNFEPQTLSPLSFYFLTLTNVSRESIVIVHRTPAND*VDT------------------
Gryllus_longicercus          KVVAAVAA--------------W--------------CATFQCLLLRRMRISVSLRNTQF*ASNAE-------SFKFLLSHTD*RK*RKYCNRAP-----DTGERLSR-----------H
Consensus

Teleogryllus_occipitalis     ------------------------------------------------------------------------------------------------------------------------
Acheta_domesticus            FLFALFFLFVRVLCLFLV--FDFCF--------------------GKYLKHHTKTPSSAFCTLGSS*FEIEISKHFHYKRRVILPVKVSTLSILQ------THLM--------------
Gryllus_bimaculatus          -LYIFIFIFLGDIVFVFVSFFAICFRTFVLRS----------------FKYQSKTTSSTFCIFVSRFHTLSLHKASHFDRKTISTLV*CTISCLVK-----THLV--------------
Gryllus_longicercus          FVHFYFYLFGGHC-------FCVCF--LFVRSVLGHLF*EVLNINQKRLLRHSVSSSLDFIHF-------------HCTRRVILTVKKSALWCKAQ*VVWFKHILSNSR**PSN***VFN
Consensus

Teleogryllus_occipitalis     ------------------------------------------------------------------------------------------------------------------------
Acheta_domesticus            --*CQVIHLVTGDKFLRNSIVFIVKKLEIFLSQIGKHADK*RLREILHCLIIMSSNGNLK-----T*KIGLGTTKDLN-IKS*SYATSTTSSATK--------SNCRRVFSGAQGLITSA
Gryllus_bimaculatus          EFEVMT**LV-----IR----F*RIRLS--LIAMNSVSSKWNL----*VRL*IKLKGNSRLF***LFQMGIIRLTELV-C*QQKICMKSESDDFRQPLLRLKFRNCR*VFPDVEGLITSL
Gryllus_longicercus          EFDCL*VP*I*---FLR----FIS-------TPINNAKGKF*I-----VLIVIISDGNYK-----TYRIGVLTTKDLYGIRERRISTTSSATKIQ--*LQMSISRC----------AKDL
Consensus

Teleogryllus_occipitalis     ------------------------------------------------------------------------------------------------------------------------
Acheta_domesticus            PTLRVSQNCSHCQQETGCKEIW---------KLH*NLPLLQFYVLKEKNVLVIH*TLINANFKGPK*SEFVL*KFVLTEYFVYPYYRNPLPGPKFFRLTAKFVLTSEFGGNDGLLYVKKE
Gryllus_bimaculatus          TGLR*D---------TVCNEIL---------KLH*NFWFHHFLGLNSKEFIV**---------**KRSKIIII--------------------FFR*LILY------------------
Gryllus_longicercus          SHLSLD-----------CDRILSAMKF*SYIKI---FGFIIF*A*IVKNLLFDN---------SKRGARL*L-----------------------FSLDGSF------------------
Consensus

Teleogryllus_occipitalis     ---------------------------------------------------------------------------------------------------------------------
Acheta_domesticus            QDDTDFL*YSTSCIPEKKS*LKSIC*IPVP*S*TSN*ECVFYYFFLIMLYLKGRMNY*V------------------------ISDHIQNTVY---TMYMRAYNKI*G*STNT*FV
Gryllus_bimaculatus          ------------T*KQFVSFVLNPCFISF--KLRN-----IFYCLFFNLIFEPSVNYILRRLHRWFEFSTEKL*QVII*TKTLFTPDTTQKAISLPQRILRE*YNNYCNNCS*LR*-
Gryllus_longicercus          -----------CIPFSNSYLL--C*IPV--S*ASN*GILFFYCLFFYIFIFEPSVNCILRRLHRWFEFSIEEL*QVII*TKSSFTPDTTQKAIYLPQRLLRE*YNNYCNNC------
Consensus
```

**Tocc.5G0008290.1_exon2**

```
Teleogryllus_occipitalis                 ------------------------------------------------------------------------------------------------------------------------
Acheta_domesticus                        -------------DICSK*ILFE-YKI------------RRF-------------------------SKLIFIK*I*NR-----------------------TIDIVK*IVLRI--*VY-
Gryllus_bimaculatus                      LEPSFKKRFLRNADACLVSIF*PNFKAEE*EWI------RKFKQHSCMHEKVQLTCHL---NEFSTILS*Y*YLDMKSRLFQSLYLSMQFKMQ*C*INWLQYLSVLWKDILLNFQYILYI
Gryllus_longicercus                      -----DQNQLKNKER---DLMMENFKL--LKWFV*IQTCRKR*P*K*M*LKISIKT*LLRGSRESNVSTLNFILSLFSR-----------------------NSIISTNLFINL------
Consensus

Teleogryllus_occipitalis                 ------------------------------------------------------------------------------------------------------------------------
Acheta_domesticus                        YGRNL*LYPIKNFN*DKNLFS--IGKKCQDNISKGSSPDGCRIFYTSDTVYVE------------FRRSL-------------EFNSSCSDR*FSVVSPINIRRMEEYYR-----KK*--
Gryllus_bimaculatus                      FK**INV*CIKLIY*IIYLYSHQIPSLAQIYSSIYSS------FIQTSGIHIQRAGMKLLNVCDHFSKTF---------*W*NYL*MSIV*R*LKFFISRFILK--V*KRTF---QKHTK
Gryllus_longicercus                      -----------------------------------SS------FIQTSGIRTQRAGMKLLTCATIFQEPFNGEIIYECQLYQDD*NSSSVDLF*KFENEHSIRN--EYYRDTLCDQKY--
Consensus

Teleogryllus_occipitalis                 ------------------------------------------------------------------------------------------------------------------------
Acheta_domesticus                        -----------AVGRT------------RSFLPL-PCPIFTSHLHTLTCNT-----LPHTSEDA*ELVNNLKRATHAEI*NHRHILR*NMHSTNGTNDLTSSQKHQLI*SYSLNIYSPNL
Gryllus_bimaculatus                      *ILPRCNM*SKVSGHVQNDVMVWLCFSWTGLAPLVLCCTLTAQAYIEILNT-----LLSTVEEQFGD---------GDFNSIEPLFILNGLWRSDSTTITS-----L*WIVLHRVLTSIL
Gryllus_longicercus                      ---PETSKMALWCGSV---------FHGESLHPL-LCAALSQLTPTFVF*TPSCFPLLKS-NSGMAI---------ASFNTIEHLFKLHDPWQSGSTIITS-----L*WIVLYRVLTRIL
Consensus

Teleogryllus_occipitalis                 ------------------------------------------------------------------------------------------------------------------------
Acheta_domesticus                        SNH*YKFISSSNLSLVSISAEFLDRHLELV*DI*SV*------------PLN----VCDDIPHFSKNVLLDENIYKSQLYE----ED*NTPFTNTTPKISIGCLFWKF----------EN
Gryllus_bimaculatus                      SNTFGMF----------WNAEFLPGLTDLQRYL----------------PFS----VALREQRAAIAQGTFQHLIEY------ACESGNCHYG*ECSNT---LLNSRI----------TN
Gryllus_longicercus                      SNTFGMF----------WNADFMPGLIDLQRYLSSVQHSVKKGLPLPKKPSST*LKVCLPEWKLSSWLRVGQHPMEFQNYQWRAPRTGK-SFSATCPDTFDLIVYSKFGRTSV*MSVKKD
Consensus

Teleogryllus_occipitalis                 ------------------------------------------------------------------------------------------------------------------------
Acheta_domesticus                        DHRKNIQNQ---------YYRNIRSLTEHQSKYQ*RQMVCESNYVNKYSKNSNRSDELKIEVMIINFFFFKH*P*IRCVLFDLTPCFRSSNTPISFSMVLCDFF*CSGIFHSVRRLPIRS
Gryllus_bimaculatus                      GERHELESHFQPRARILLIS*CIRSLAEHHSKCR*RKMVVDE-----------------------------------ITLMEVLESIYSENSSIV-------MMKS*HWFHPSL*LSKEV
Gryllus_longicercus                      GCR---*NHINGSAGKYIF*KFINSDDEIVT-------LVSS-----------------------------------VPLIE-*RSL*AK*IE*T-------TMKYNGKYLVVINMS---
Consensus

Teleogryllus_occipitalis                 ------------------------------------------------------------------------------------------------------------------------
Acheta_domesticus                        CFFYFFFFILIKYS*LILRKEINMQNKKGLKKQ*NIMESLRC*LICVSFTAHHIMVKCYWLKVPILGS*KLLYSAGIPLIPQNP*----TFQAISALAT--HAGHS----TATWHRHAAR
Gryllus_bimaculatus                      C-----------------------KRNK*NEQQ*NIMKSI------------------WW*LI*GLG------SAEIPQVSQNSQ----TFQAITEVHS--HACRPFDSHMARGTRHAAR
Gryllus_longicercus                      --------------------------TRFRRDPSSVAKSV--------------------------------------NVSSNYRGTQPRMQAIRQPHGTRHAARG-TRHAARGTRHAAR
Consensus

Teleogryllus_occipitalis                 --------------------------------------------------------------------------------RDGGDGGGYG------------------------------
Acheta_domesticus                        STSATT---AVAAG----------------GGG------AWAGDADADDAPLGRHRSERGRLGERAGGGATRLRSRSRPARPGG-GGGYGQGECGGGGRAWG-----------GGGGW*R
Gryllus_bimaculatus                      GTSATTAAAAAARGRVTQMPTTPHLGGTEAGAG--------------------------SRAVGQAG-------------APHASGAGPDRRDP--------------------------
Gryllus_longicercus                      GTR------HAARG-------TQHVGHNGGGGGGGGGGGAWAGDADADDAPLGRHRSGRGRPGGQAG-------------APHASGAGPDRRDPAAEATATANVRR*RGLRRRGYWGWER
Consensus                                                                                                                     * *

Teleogryllus_occipitalis                 -----------------------------------------------------------------------------------GGLGGGGG-----------------------------
Acheta_domesticus                        *RW--------------------------SAD----------EGGGE-------------------GE**R*VRRWGRGGDGDGGDGGGGGGEGGRDGGGEEEEVAKEAEGTLGAEGAGG
Gryllus_bimaculatus                      -----------------------------AAEATATANLAEVEGPEV--VLELRALKMEMQMKVSAG-EEGGMMKEGGGGEG-GGDGGGGN-DHGKRGGGRG---AC*GGGVVVESVKGG
Gryllus_longicercus                      *RWRCR*RCQRGKRGG**RQGAVGKVVVTAAEATATANVAEVEGPAEAGALWLGALKVE--MKVEAGAKAGGGMKAGGSGEG-GGDGGG----NGSGGGG------------------GG
Consensus                                                                                                                   ** ***

Teleogryllus_occipitalis                 ------------------------------------------------------------------------------------------------------------------------
Acheta_domesticus                        AEGAGGGRRRGRRGRRGGGAEGAEGAEGAEEAKEAEGELEETAAAGAATVTAAAAATVTATAALENRSSITRGRRRWWHRSPRRS--------ATFQRLLLSRLRISVTLRNTKFESQTL
Gryllus_bimaculatus                      DEGGGGRERRGALGKVVVTAEVTVAVEEAEE---------------------MAVATVTATAASKNRSSITRGRRRWWQRSPRGAPLSSVCSCAACESLLL-----YVT---PNFEPQTL
Gryllus_longicercus                      D----------------------------------------------------AVATVTATAASENRSSITRGRRRWWQRSPRGAPLSSVCSCAACESPFL-----YVT---PNFEPQTL
Consensus

Teleogryllus_occipitalis                 ------------------------------------------------------------------------------------------------------------------------
Acheta_domesticus                        SPLNLLSLCYVFYLHLDLVFHLSRIS*V-------------GFLFCFV*FLFALFFLFVRVLC-LFLVFDFCFGKYLK-----------HHTKTPSSAFCTLGSS*FEIEISKHFHYKRR
Gryllus_bimaculatus                      SPLSF------YFLTLT---NVSRESIVIVHRTPAND*VDTLYIFIFI-FLGDIVFVFVSFFA-------ICFRTFVLRSFK-------YQSKTTSSTFCIFVSRFHTLSLHKASHFDRK
Gryllus_longicercus                      SPLSF------YFLTLT---NVSRESIVIVHRTPAND*VDTLYIFIFI-SLGGIVFVFVSFLCDLF--*DICFKKF*I-SIKNDFFDILYLRLSISST---FIAQGESF*P*KNQHFGVK
Consensus

Teleogryllus_occipitalis                 ------------------------------------------------------------------------------------------------------------------------
Acheta_domesticus                        VILPVKVSTLS-ILQTHLM*CQVIHLVTGDKFLRNSIVFIVKKLEIFLSQIGKHADK*RLREILHCLIIMSSNGNL---KT*KIGLGTTKDLNIKS*SYATSTTSSATK--------SNC
Gryllus_bimaculatus                      TISTLV*CTISCLVKTH---------LVEFEVMT**LVIRF*RIR--LSLIAMNSVSSKWNL*VRL*IKLKGNSRLF***LFQMGIIRLTELVC*QQKICMKSESDDFRQPLLRLKFRNC
Gryllus_longicercus                      H-NKLFGSNTSCRIRGDD-------LVTSDKFLTNSIVFKCHEF-------------SFFVL*VRL*ITLKGNSRLF***LFQMVIIRLTELVC*QQKICMESESDEFRQPLLRLKFSNC
Consensus

Teleogryllus_occipitalis                 ------------------------------------------------------------------------------------------------------------------------
Acheta_domesticus                        RRVFSGAQ------------------------GLITSAPTLRVSQNCSHCQQETGCKEIWKLH*N------LPLLQFYVL------KEKNVLVIH*TLINANFKGPK*SEFVL*KFVLTE
Gryllus_bimaculatus                      R*VFPDVE------------------------GLITSLTGLR*DT--------V-CNEILKLH*NFWFHHFLGLNSKEFIV****KRSKIIIIFFR*LILYT*K-----QFVS--FVLNP
Gryllus_longicercus                      R*VFPDVQRTYHISHWIAIGYCLQ*NFEVTLKSLASSFFRPK**R--------IYCLIIVKEEQD--YNYFL*MAHFVYL--------KAIRIFCAKSLFHKLQT---EEFYF--FIVY-
Consensus

Teleogryllus_occipitalis                 ------------------------------------------------------------------------------------------------------------------------
Acheta_domesticus                        YFVYPYYRNPLPGPKFFRLTAKFV--------LTSEFGGNDGLLYVKKEQDDTDFL*YSTS------CIPEKKS*LKSIC*IPVP*S*TSN*ECVFYYFFLIMLYLKGRMNY*VISDHIQ
Gryllus_bimaculatus                      CFI-----------SF-KLRNIFY---CLFFNLIFEPSVNYILRRLHRW------FEFSTE-KL*QVII*TKTLFT---------------------------------------PDTTQ
Gryllus_longicercus                      FFI-----------SFSNLLSIAY*GGCIV-------GLNSRLRNSDR---------LSSERRVRSRLIPHKKRFIC-------------------------------------LKDY*E
Consensus

Teleogryllus_occipitalis                 ---------------------------
Acheta_domesticus                        NTV---YTMYMRAYNKI*G*STNT*FV
Gryllus_bimaculatus                      KAISLPQRILRE*YNNYCNNCS*LR*-
Gryllus_longicercus                      NNIIIIVTIV-----------------
Consensus
```

**Tocc.5G0008290.1_exon3**

```
Teleogryllus_occipitalis  ------------------------------------------------------------------------------------------------------------------------
Acheta_domesticus         ---------------------------------------------------DICSK-----------*ILFEYKIRRFSK---LIFIK*I-----*NRTIDIVK*IV-------------
Gryllus_bimaculatus       RALF*EEISEERRCMFGVHFLTKFQG*RIRMDSQVQTTFMHARKSSINMPFE*I*H-------DIKLILIFGYEIATFPK---FIFIYAI*NAIMLNKLASVFKCIVEGHIVEFPIYIIH
Gryllus_longicercus       RS---KSIEE*RK---------RFN------DGEFQTFKMVCLNS--NMS*KITVKMNVTQNLD*NLIIEGKSRK*CFDSKLHFIFI--------LTKLHH*YK----------------
Consensus

Teleogryllus_occipitalis  ------------------------------------------------------------------------------------------------------------------------
Acheta_domesticus         ---LRI*VYYG----RNL*LY---PIKNFN*DKNLFSI--------GKKCQDNISKGSSPDGCRIFYTS----------------DTVY-VEFRRSLEFNSS-----CSDR*FSVVSPIN
Gryllus_bimaculatus       I*IINKCVMY*THILNNLSLFSPNSIISTNLFINIFII-YSDIGYSYTTSWDEIA-----*RM*PFFKNVLMVKLFMNVNCMKMIKILYQSIYSESLKTNIP--------KTY------E
Gryllus_longicercus       -------------------------------FIHQFIIFYSDIGYSYTTCWDEIA-----DVCDHFSRT------F*W*NYL*M------SIVSR*LKFFIS*FILKV*KRTF------H
Consensus

Teleogryllus_occipitalis  ------------------------------------------------------------------------------------------------------------------------
Acheta_domesticus         IRRMEEYYRKK*-AV-------------GRTRSFLPLP-CPIFTSHLHTLTCNTLPHTSEDA*ELVNNLKRATHAEI*NHRHILR*NMHSTNGTNDLTSSQKHQLI*SYSLNIYSPNLSN
Gryllus_bimaculatus       INFTEMQYVIKSIRTCPK*CDGVVVF--FMDRACTP---CFVLHTHG------TGLH*NFEHLAF----------------HC*R----AIRG-------------WRFQFN*APVHTKR
Gryllus_longicercus       TKRILPRYTM*S-KVSGNVQNGVMVWQCFSWRELAPLALCGTITTHAYVRVLNTFLLSSVEE-QFGD-------------GDCL------------------------FQHD*APVQTTR
Consensus

Teleogryllus_occipitalis  ------------------------------------------------------------------------------------------------------------------------
Acheta_domesticus         H*YKFISSSNLSLVSISAEFLDRHLELV*DI*SV*PLNVCDDI--------PHFSKNVLLDENIYKSQLYEED*NTPFTNTTPKISIGCLFWKFENDH-------------RKNIQNQYY
Gryllus_bimaculatus       PVAE*FHDNNIPVMDCPAQSLDFN-----------PIEHLWDVLERRILARPHRPTTI------------------PSLQCSTP*TKGC--------HCPRNLPAPD*VCLREWKLSLWL
Gryllus_longicercus       PVAEWFHNNNIPVMDCLVQSSD*N-----------PIEHLWDVLER*FHARPH*PSTI------------------PLFSAA----------------------------LR*ERAAIAQ
Consensus

Teleogryllus_occipitalis  ------------------------------------------------------------------------------------------------------------------------
Acheta_domesticus         RNIRSLTEHQSKYQ*R--------QMVC----ESNYVNKYSKNSNRSDELKIEVMIINFFFFKH*P*IRCVLFDLTPCFRSSNTPISFSMVLCDFF*CSGIF--HSV-------------
Gryllus_bimaculatus       RMLQHPIEFQN-YQWRAPRTGKSFSATCPDTFDLIVYSKFGRTS-----L*MSV--------------------------KKD-GCR*NHI-------NG----------------SAGK
Gryllus_longicercus       ETFQHLIE--------------SMSAGV----ETVIMAKSGPTSYGIPELPMAS--------------------------ATNWKVIFSHV-------SGYF*SHSVFEVWQNISLNVGK
Consensus

Teleogryllus_occipitalis  ------------------------------------------------------------------------------------------------------------------------
Acheta_domesticus         -RRLPIRS----CFFYFFF-------------FI--------LIKYS*LILRKEINMQNKKGLKKQ*NI-MESLRC*LICVSFTAHHIMVKCYWLKVPILGS*KLLYSAGIPLIPQNP*T
Gryllus_bimaculatus       ----------------YIF-----------*KFINSDDEIVTLVSSVPLIE*RSL*AK*IE*TTMKYN--EKYLVVINMRTRFR------------------------------------
Gryllus_longicercus       ERWL*MKSH*WKCWKVYILKIHQ****NRNIGFIRPFN*VKKFISEINRMNNNEI*WKVFGGD*YEY*VPPRSLKCRKIRKRFK------------------------------------
Consensus

Teleogryllus_occipitalis  ------------------------------------------------------------------------------------------------------------------------
Acheta_domesticus         FQAISALATHAGHSTATWH-------------------RHAARSTSATTAVAA-------------GGG------GAWAGDADADDAPLGRHRSERGRLGERAGGGATRLRSRSRPARPG
Gryllus_bimaculatus       -RDPASIAKFANVSSNY----RGTQPRMQAIRQ-PHGTRHAARGTRHVGHNGG-------------GGG------GAWAGDADADDAPLGRHRSGRGQPGGRAGGGATRLGSRSRPARPG
Gryllus_longicercus       -QLQRYTATHAGHSTATWHAARGTRHAARGTRHAARGTRHAARGTRHAARRPQRRRWRRRRRRRRVGG*RRCRRRPTWAAQK-------------RARAAGRAGGGATRLGSRSRPARPG
Consensus

Teleogryllus_occipitalis  ----GY----------------------------------------------GGGGG---------DGGGERH---------RGG-------GGDGG-GGGNGGSGGDEGG---------
Acheta_domesticus         GGG-GYGQ----------------------------GE------CGGGGRAWGGGGGW*R*RWSADEGGGEGE**R*VRRWGRGG-------DGDGGDGGGGGGEGGRDGGGE-------
Gryllus_bimaculatus       GGGNGHGK------P----------------------------------------------------GGGRGA---------*GGVGVASVKDGDADEGVSGGRGGDDEGRGRWGRWW*R
Gryllus_longicercus       GGGNGHGKRAEVEGPEEAGVLGLGALKMEMQMKVSAGEEGGMMKAGGGGEGGGDGGG---GNGHGKRGGGRGAC-------*GGGVVVGSVKGGDEGGGGSEG-GGGDEGGGQWGRWW*R
Consensus                     *                                                              ***             **        **   *   *  *   *

Teleogryllus_occipitalis  -------------------------EGDGGG-----------------------------------GAGAGEG-----------------------------------------------
Acheta_domesticus         -----EEEVAKEAEGTLGAEGAGGAEGAGGG-----RRRGRRGRRG----------------GGAEGAEGAEGAEEAKEAEGELEETAA--AGAATVTAAAAATVTATAALENRSSITRG
Gryllus_bimaculatus       RWR*RPRQTWRRERGLLR-------RGRCGGER*RWR*RWRRARKAGGIGEGGGDGGGNGSGGGGGGDGGGNG----DCNGGVEE*IVHHARAAKVVAAVAA------------------
Gryllus_longicercus       R---------------------------------R*R*RWRRRRRC-----------------------GGNG----DCNGGVGE*IVHHARAAKVVAAVAA------------------
Consensus                                                                                         *

Teleogryllus_occipitalis  ------------------------------------------------------------------------------------------------------------------------
Acheta_domesticus         RRRWWHRSPRRSATFQRLLLSRLRISVTLRNTKFESQTLSPLN-LLSLCYVFYLHLD---LVFHLSRIS*VGFLFC--FV*FLFALF--------FLFVRVLCLFLVFDFCFGKY-----
Gryllus_bimaculatus       ----W------SATFQCLLLRRMRISVTLRNTQF*ASNAESFKFLLS-------HTD*RK*RKYCNRAPDTGERLSRHFVYFYFYIFGGHCFCVCFLFC-----HLF*DICFKKF*ISIK
Gryllus_longicercus       ----W------CATFQCLLLRRMRISVSLRNTQF*ASNAESFKFLLS-------HTD*RK*RKYCNRAPDTGERLSRHFVHFYFYLFGGHCFCVCFLFVRSVLGHLF*EV----LNINQK
Consensus

Teleogryllus_occipitalis  -------------------------------------------------------------------GDR--------------------------------------------------
Acheta_domesticus         -----LKHHTKTPSSAFCTLGSS*FEIEISKHFHYKRRVILPVKVSTLSILQTHLM*C--QVIHLVTGDKFLRNS--------IVFI-----------VKKLEIFLSQIGKHADK*RLRE
Gryllus_bimaculatus       NDFFDILYLRLSISYTFIAQGEP-F*P*NNQHFGVMHNKLFG*NTS-----------CRIRGDDVVTSDKILTNS--------IVFNCHEFSFF*V------EFIGTPINKAKGKF*I--
Gryllus_longicercus       R----LLRHSVSSSLDFI-------------HFHCTRRVILTVKKSAL--------WC--KAQ*VVWFKHILSNSR**PSN***VFN--EFDCL*VP*I*FLRFISTPINNAKGKF*I--
Consensus

Teleogryllus_occipitalis  ------------------------------------------------------------------------------------------------------------------------
Acheta_domesticus         ILHCLIIMSSNGNLKT*KIGLGTTKDL-NIKS*SYATSTTSSATK--------SNCRRVFSGAQGLITSAPTLRVSQNCSH------CQQETGCKEIWKLH*NLPLLQFYVLKEKNVLVI
Gryllus_bimaculatus       ---VLIVIISNGNYKTYRIGVLTTKDLYEIRERRF--STTSSAIKIQELQMSISRCGRTY-----------------HISHWIAIGYCLQ------------------------------
Gryllus_longicercus       ---VLIVIISDGNYKTYRIGVLTTKDLYGIRERRI--STTSSATKIQ*LQMSISRCAKD-------------------LSH---------------------------------------
Consensus

Teleogryllus_occipitalis  ------------------------------------------------------------------------------------------------------------------------
Acheta_domesticus         H*TLINANFKGPK*SEFVL*KFVLTEYFVYPYYRNPLPGPKFFRLTAKFVLTSEFGGND----GLLYVKKEQDDTDFL*YST-----------------SCIPEKKS*LKSIC*IPVP*S
Gryllus_bimaculatus       ----------------------------------------*NFEVTLKFLVSSLFRPK**RIYCLIIVKEEQDYNYFL*MTHFVYLKAIRI--------FC---SKS-------------
Gryllus_longicercus       ------------------------------------------LSLDCDRILSAMKF*SYIKIFGFIIF*A*IVKNLLFDNSK----RGARL*LFSLDGSFCIPESNSYL--LC*IPV--S
Consensus

Teleogryllus_occipitalis  -------------------------------------------------------------------------------------------------------------
Acheta_domesticus         *TSN*ECVFY------YFFLIMLYLKGRMNY*VISDH------IQNTVYTMYMRAYNKI*G*-----------STNT*FV-----------------------------
Gryllus_bimaculatus       -------LFHKLQTEEYFLLFIF*FNFRTFC---QLH------IEEVASLV*VLD*ETLTGYHLNEDFVHA*YHTKSDFFASKNLERII**LL*QLFLTSL--------
Gryllus_longicercus       *ASN*GILFF------YCLFFYFIFEPSVNGILRRLHRWFEFSIEEL*QVII---------------------*TKSSFTPDTTQKAI---YLPQRLLRE*YNNYCNNC
Consensus
```

**Tocc.5G0008290.1_exon4**

```
Teleogryllus_occipitalis    ------------------------------------------------------------------------------------------------------------------------
Acheta_domesticus           -------------DICSK*I--------LFEY-----------------KIRRF---------------------SKL-------------------------------------I
Gryllus_bimaculatus         LEPSFKKRFLRNADACLVSIF*PNFKAEE*EWIRKFKQHSCMHEKVQLTCHLNEFSTILS*Y*YL---------DMKSRLFQSLYLSMQFKMQ*C*INWLQYLSVLWKDILLNFQYILYI
Gryllus_longicercus         -----------------IKI-N*RIKKEI**W-----------------RISNF*NGLSEFKHVVKDNRKNECNSKSRL--------------------------------------
Consensus

Teleogryllus_occipitalis    ------------------------------------------------------------------------------------------------------------------------
Acheta_domesticus           FIK*I*NRTIDIVK*IVLRI*VYYGRNL*LYPIKNFN*DKNLFS-----------IGKKCQDNISKGSSPDGCRIFYTSDTV-YVEFRRSLEFNSSCSDR*FSVVSPINIRRMEEYYRK
Gryllus_bimaculatus         FK**INV*CIKLIY*I---IYLYSHQIPSLAQIYS-----SIYSSFIQTSGIHIQRAGMKLLNVCDHFS-----KTF*W*NYL*MSIV*R*LKFFIS-------------RFILK-V*K
Gryllus_longicercus         -KPNY*GEVAKVMFRL*TSFYLYSHETPSLVQIYS-----SIYHLLFRHRVFVHNVLG*NC*RVRPFFKNLLMVKLFMNVNCIKMIKILHQLIYSESLKTNIPYETNITEIHYVIKSIRK
Consensus

Teleogryllus_occipitalis    ------------------------------------------------------------------------------------------------------------------------
Acheta_domesticus           K*AVGRTRSFLPLP---CPIFTS-HLHT--LTCNTLPHT----------SEDA*ELVNNLKRATHAEI*NHRHILR*NMHS---TNGT--NDLTS--SQKHQLI*SYSLNIYSPNLSNH
Gryllus_bimaculatus         RTFQKHTK*I--LPRCNM*SKVSGHVQNDVMVWLCFSWTGLAPLVLCCTLTAQAYIEILNTLLSTVEEQFGDGDF--NSIEPLFILNGLWRSDSTTITSL*WIVLHRVLTSIL-------
Gryllus_longicercus         RPKWRYGVAVFFMERACTPCFVR-HYHNSRLR-SCFKHL----LAFLC*RAIRGWRLPLSTRLSTCSNYTTRGRVVPQ**HP---CNGL---------SCTEF*LESYRTPL--------
Consensus

Teleogryllus_occipitalis    ------------------------------------------------------------------------------------------------------------------------
Acheta_domesticus           *YKFISSSNLSLVSISAEF------LDRHLELV*DI*SV*PLNVCDDIPHFSK--------------NVLLDENIYKSQLYEED*NTPFTNTTPKISI----GCLFWKFENDHRKNIQN
Gryllus_bimaculatus         -----SNTFGMFWN--AEFLPGLTDLQRYLPFSVALREQRAAIAQGTFQHLIEYACESGNCHYG*ECSNTLLNSRITNGERHELE-----SHFQPRARILLIS*C--------------
Gryllus_longicercus         ------GCFGTLISCQASLTFNDTSLQCSTPLRKGCHCPRNLPAPD-----*KYVCRSGNCHHG*EWANILWNSRITNGERHELE-----SHFQPRVRILLIS*C--------------
Consensus

Teleogryllus_occipitalis    ------------------------------------------------------------------------------------------------------------------------
Acheta_domesticus           QYYRNIRSLTEHQSKYQ*RQMVCESNYVNKYSKNSNRSDELKIEVMIINFFFFKH*P*IRCVLFDLTPCFRSSNTPISFSMVLCDFF*CSGIFHSVRRLPIRSCFFYFFFFILIKYS*LI
Gryllus_bimaculatus         -----IRSLAEHHSKCR*RKMVV---------------DEI-------------------TLMEVLESIYSENSSIVM--------------------MKS*HWF--------HPSL*
Gryllus_longicercus         -----IRSLAEHQSKCR*RKMVV---------------DEI-------------------TLMEVLESIYSENSSIVM--------------------MKS*HWF--------HPSL*
Consensus

Teleogryllus_occipitalis    ------------------------------------------------------------------------------------------------------------------------
Acheta_domesticus           LRKEINMQNKKGLKKQ*NIMESLRC*LICVSFTAHHIMVKCYWLKVPILGS*KLLYSAGIPLIPQNP*TFQAISALATHA----GHSTATWHRHAARS--------------------T
Gryllus_bimaculatus         LSKEVCKRNK*N-EQQ*NIMKSI------------------WW*LI*GLG------SAEIPQVSQNSQTFQAITEVHSHACRPFDSHMARGTRHAARG--------------------T
Gryllus_longicercus         LSKEVYKRNK*N-EQQ*NIMESI------------------WW*LI*VLG------SAEIPQVSQNP*TFQAITEVHSHACRPFDSHMARGTRHAARGTRHAARGTRHAARGTRHAARST
Consensus

Teleogryllus_occipitalis    --------------------------------------------------------------------GRRR-GRRR----------RTVPSRQ------------------RRRR---
Acheta_domesticus           SATTAVAAGGGGAWAGDADADDAPLGRHRSERGRLGERAGGGATRLRSRSRPARPGGGGGYGQGECGGGGRAWGGGG-------GW*R*RWSADEGGGE--GE*--------*R*VR---
Gryllus_bimaculatus         SATT---------------------------------AAAAAARGRVTQMPTTPHLGG----TEAGAGSRAVGQAGAPHASGAGPDRRDPAAEATATANLAEVEGPEVVLELRALKMEM
Gryllus_longicercus         SATTAAVA----------------------------AAAAAAARGRVTQMPTTPHLGG----TEAGAGGRA-GRRG-----------RHTPREQVPTGA--------------TRRRRQ--
Consensus                                                                                       * *  *             *                         *

Teleogryllus_occipitalis    -------------------------------------------RWKKWRERRKR---RKTW-------------------------RRRGGAAAR-----------------------
Acheta_domesticus           --RWGRGGDGDGGDGG--------GGGGEGGRDGGGEEEEVA----------KE--------AEGTLGAEGAGG------------AEGAGGGRRRGRRGRRGGGAEGAEGAEGAEEAKE
Gryllus_bimaculatus         QMKVSAGEEG---------GMMKEGGGGEGGGDGGGGNDHG------------KRGGGRGAC*GGGVVVESVKGG------------DEGGGGRERRGALGKVVVTAEVTVAVEEAEEM--
Gryllus_longicercus         RPRQTCGGRGA*GGGGIGVGSVKDGDADEGVSGGRGGDDEGRGRWGRWW*RRRRQRPRQTWRR*RGLLRRGRCGWER*RWR*RWRRERRRGGG*RRGAVGKVVVTAEVTVAVEEAEEM--
Consensus                                                                                                     **   *

Teleogryllus_occipitalis    ------------------------------------RRQRRR*-----------------------------------------------------------------------------
Acheta_domesticus           AEGELEETAAAGAATVTAAAAATVTATAALENRSSITRGRRRW-------------WHRSPRR---SATFQRLLLSRLRISVTLRNTKFESQTLSPLNLLS--LCYVFYLHLDLVFHLSR
Gryllus_bimaculatus         -------------------AVATVTATAASKNRSSITRGRRRW-------------WQRSPRGAPLSSVCSCAACESLLLYVT---PNFEPQTLSPL---------SFYFL-TLT-NVSR
Gryllus_longicercus         ------------------------------RWQR*LQRRRRRIDRPSRAGGEGGGSGRRVVRHFPVSAPAPHA-------------NLRFST*HPILSLKR*VL*VFTFSH*LT-*VEK
Consensus                                                    * ***

Teleogryllus_occipitalis    ------------------------------------------------------------------------------------------------------------------------
Acheta_domesticus           -------------IS*VGFLFCFV*FLFALFFLFVRVLCLFLVFDFCFGKYL----KHHTKTPSSAFCTLGSS*FEIEISKHFHYKRRVILPVKVSTLSIL-QTHLM*CQVIHLVTGDK
Gryllus_bimaculatus         ESIVIVHRTPAND*VD-TLYIFIFIFL-GDIVF------VFVSFFAICFRTFVLRSFKYQSKTTSSTFCIFVSRFHTLSLHKASHFDRKTISTLV*CTISCLVKTHLVEFEVM-------
Gryllus_longicercus         ---VL*SCTGHRRTIE*TLCTFLFLSLWGALFL------CLFPFCAICFRTFVLRSFKYQSKTTSSTFCIFVSRFHPLSLHKASHFDRKKISTLV*STISCLVQTHLVEFEVM-------
Consensus

Teleogryllus_occipitalis    ------------------------------------------------------------------------------------------------------------------------
Acheta_domesticus           FLRNSIVFIVKKLEIFLSQIG------------KHADK*RLREILHCLIIMSSNGNLKT*KIGLGTTKDLNIKS*SYATSTTSSATKS--------NCRRVFSGAQGLITSAPTLRVSQN
Gryllus_bimaculatus         ----T**LVIRF*RIRLSLIAMNSVSSKWNL*VRL*IKLKGNSRL--F-------***LFQMGIIRLTELVC*QQKICMKSESDDFRQPLLRLKFRNCR*VFPDVEGLITSLTGLR----
Gryllus_longicercus         ----T**LVISF*RIRLSLSAMNLVSSFY----KYAYK*R*REILDCF-------DSNYFRW*L*DLQNWCVNNKRSVWNPRATNFDNLFCD*NSVTADEYFQMCKGLITSLTGLR----
Consensus

Teleogryllus_occipitalis    ------------------------------------------------------------------------------------------------------------------------
Acheta_domesticus           CSHCQQETGCKEIWKLH*NLPLLQFYVLKEKNVLVIH*TLINANFKGPK*SEFVL*KFVLTEYFVYPYYRNPLPGPKFFRLTAKFVLTSEFGGNDGLLYVKKEQDDTDFL*YSTSCIPEK
Gryllus_bimaculatus         -----*DTVCNEILKLH*NFWFHHFLGLNSKEFIV---------****KRSKIIII--------------------FFR*L-------------ILYT*KQ-----FVSFVLN-----
Gryllus_longicercus         -----*DTVCNEILKLH*NLWLHHFLGLNSKEFIV---------****KRSKIIII--------------------FFRWL-------------ILYT*KQ-----FVSFVLN-----
Consensus

Teleogryllus_occipitalis    ------------------------------------------------------------------------------------------------------------------------
Acheta_domesticus           KS*LKSIC*IPVP*S*TSN*ECVFYYFFLIMLYLKGRMNY-----------------------------------------------*VISDHIQNTV---YTMYMRAYNKI*G
Gryllus_bimaculatus         ------------PCFISFKLRNIFYCLFFNLIF-EPSVNYILRRLHRWFEFST--------------------------EKL*QVII*TKT-LFTPDTTQKAISLPQRILRE*YNNYCN
Gryllus_longicercus         ------------PCFISFKLRNFIFLLFIFL-----------------FHFRTFCQLHIEEVASLV*ILD*GTLTGYHLNEEFVHA*YHTKSDLFASKTIERII---**LL*QL------
Consensus

Teleogryllus_occipitalis    --------
Acheta_domesticus           *STNT*FV
Gryllus_bimaculatus         NCS*LR*-
Gryllus_longicercus         [illegible]
Consensus
```

**Tocc.6G0002000.1**

```
Teleogryllus_occipitalis    GAAAAAAATATCGACGATTA----------------------------------------------------------------------------------------------------
Acheta_domesticus           ---------GCCCGCTAGTA---------------------------TTCACTTTGTGA--------------------GCTGATTTGAAGTTAACTAT---------------------
Gryllus_bimaculatus         AAGAAATGTACTCACCGTCACTTTATCAACGACCTCATTTACATTCTTTTACTTCGCGAAACAGCACGACCCACATTTGGCTGTTACTGAGTTAATTGTACAGAGTGGGAACCGACTAAG
Gryllus_longicercus         TGAAATTGAACTTACTTGTA-------------------CACCTCTTTCAACTCTATTAATTGGGACTAATAACAATTGTGCATTTTTAACACAGTGGG-----------------TAAA
Consensus                                 *    *

Teleogryllus_occipitalis    --------------------------------------------------------GGATGAAAGAAAATATCCAGA---------CCCACATCCAACATG------------------C
Acheta_domesticus           ----------------------------------------------------------------GAATTTTTGTCGAAAATCC---------TCCACCA---CCAGGGT----------G
Gryllus_bimaculatus         CTACACGCATACACACAGAGANNNNNNNNNNNNNNNNNNNNNNNNNNNNNNNNNNNNTAAAGAGAGAGAGTGTTTGTAATTAATACAATTTGGTATGCAATA---CCAACGTCAAAATTAAAA
Gryllus_longicercus         TTGCA-----------------------------------------------------------GCAAATTTGCAGAAAAAAT----TAATATTCAAAATAGCCGAATTCAACATTGA-G
Consensus                                                                                   *    * *                    * **  *

Teleogryllus_occipitalis    CAAGTACCTTACATACCTTTATACCT------------------------------------------------------------------------------------------TACC
Acheta_domesticus           CGATCTCTCTAACCTCCC---------------------------------------------TATTCTAAACGCTTCAA----------------TGGCAACAGTCGTCAGCCTGAGTC
Gryllus_bimaculatus         CATTTTCTTTATATACCCGTATTTC-------------------------------------GTTTTTAGAACTCCTCAAAACTGCTTTGTCGGTTCTGCGTCACTTGTACATTT-AATT
Gryllus_longicercus         TGCCTTTCTTAGTTGCTTCCGCACTTAGTTGTATTGTAGTATGCTACAGATGGCTAAAAACATTATTCTAAGAACTTCAATATTATCTATGTCTATTTGCAAGAGTTTTAAATCGATATT
Consensus                            **    *

Teleogryllus_occipitalis    ACCGAGCATAAC---------------------------------------------ACACAATCAATGCATCCAAATTAAAGTAGGGT-------------------------------
Acheta_domesticus           ATCATACCCAGTTACGTGGTTTGCTTATTGAATTTATTTAT----------ATTTTGGAAAAATTCGCAAAGTTCAGTTGATGTTCAACTTTGCATTTGTATT-------TACGGGGTTA
Gryllus_bimaculatus         ACAGTTCTCAGTTACTTGTACATTTAATTGCAGTTCCTTGTTGCGACACAAACGGCTAAACGATTCATAGATTCATATCATATTATTAGTTTATGATCATATTA-----ATTCGGAAAAT
Gryllus_longicercus         AATTTATTCAAT--------------------------------------AGAGCTAAAATGATGCATGTATTAGAAATAAAATAAGATATAATTTTCGTCGCAACAGAATTTGCGAATG
Consensus                   *        *                                                 *  **      *            *

Teleogryllus_occipitalis    --------------------------------------------------------------------------------------TTTCATGACAGATTTTA-----------------
Acheta_domesticus           CAATACAGCTTATTCTAATTTAATACTAACACTA-------------------AACTTCTTCTGACGAGGGGACTCACAGTACACTTCAATTGATTTATTCTATATTGCTGAGTATATTC
Gryllus_bimaculatus         CATTAGATCAAATTCAAAATGAAAACAAATGGTATTTCATATTATTTGTTGCTGATGCCTTCACACAATGAGATTGAC--------TAAAATGAAAAATCTCATATT--------TTACC
Gryllus_longicercus         CTCAGCAGTTGGTGCCCACTAAATGTAGTAGGGG------------GACGC-GAGAATTTCTGAGCTTTGGATCGGGAGTGT---TCGAAGGCCAGGTTTGACCTTAAGTGGC-TCGCA
Consensus                                                                                         *     *     *   *

Teleogryllus_occipitalis    --AGCCATATTTAGGCAATGAG--------------------------------------------------------------------------------------------------
Acheta_domesticus           AAAAACACATGTTAAGGATATC-TATTTCTTATTCGAATTTGACGTTCTGAATATTTACCCAGCTTGAGATTTCTGCAATGGCTGATACAAGTGGAGGTTTGAGTAAAAGTTATT-----
Gryllus_bimaculatus         AAAAATAAACGTAAATAATATTGTCATTGAAATGCAAATTACGCCACTACAGTATGTAGTTAGTTT----------------------CTAGTGTAGTATTCGGTCAAGGATACTCCATC
Gryllus_longicercus         ACAAATTTAGTTAAGCAATGATTTATTATTAGTTGGAAT--AATATTTCAAATGTAAATATGAATTAAAC------------------CCTGAGGATGGCTCTTTAAAATTAACT-----
Consensus                     *     *  *     **

Teleogryllus_occipitalis    -------------ACTTTACGGACGTTTTACAGTTTTGAA--------------------------------------------------------------------------------
Acheta_domesticus           ---------AGCG-ATATTATCAATACGGTAAGGGTTTGGTGTTCAATTCAATAAGTGGTTTGATGTTTAATT--CTGTGTTTAGTTGCTTGTTGGATA---------------------
Gryllus_bimaculatus         ACAGTGCGCAGCGTGTTTCATGAAAGCAGTGAAGATT-----TTTACCCAAATGAAAGCATAGTTATCCCATCAACCACGCGGAGATATATGTTGATTACATCAGTATCCTGGCCAATCA
Gryllus_longicercus         ---GAATGGTACATTTTCTAGCGTTCTGTTAGAAACTT----TTTCAGGGACTCAAAAGATTATAAACTTATTGAATGTGTTAAGTTACATTCTG------------TCAGTTCAATCC
Consensus                                      *         *      *

Teleogryllus_occipitalis    ----------------------------------------------------GAGAACCCCCACGTCG--TAGCCCAGGT----------------------------------------
Acheta_domesticus           -------------------------------------------------------------TAAACCAGAGGGCTTGGGC--------GATGGGTCTTAAGAATTGGTTTTGGGTGATTT
Gryllus_bimaculatus         ACACAACTTGACTAGTGCCGGCGCTACATATATGTAGACCGGGCGATCGCCCGGGGACCCGCAAGTCAAGGGGCCTGAATATGGAACGGATGAGAACAAGGAATTGTTTTCGGGAGAATG
Gryllus_longicercus         AGGCACGTCAAATCGCGCCCGCACGAGCTACTTTCCGCTTGG----------GAAGCCCGATGTTCCAACGAGCACGGGC----------------------------------------
Consensus                                                                                     *     **

Teleogryllus_occipitalis    ------------------------------------------------------------------------------------------------------------------------
Acheta_domesticus           GTAATTT---ATTAATATTATTTGTCCATACCGGGTAGGCATATAGGAGGAGGAGGAAGAGTTTGATGATG-------------GACTTGTATATAATGACATTGTTTAAAGAG-----
Gryllus_bimaculatus         ATACATTTCAGCACATTTCAGGCACTGAACTTGACTGCACATTTGCATTGGGTAACTGACGTTCCGTTATGTTCGAAGTAACCACCATTCACATAACTTCACATCAGTGAGAAAGCAGCA
Gryllus_longicercus         ---AACTTCCCCTCACTCCGCACTCATTTCCCCTTAAAATACAAAAAGTGATGTACAAGCGCAAGCTGACG-------AGGCCCGTGGCTGTGAATAGTGACTTCATACGAAAAT-----
Consensus

Teleogryllus_occipitalis    -----------------------------------------------------------CAGAAC--------------------------AATTCGAGGTTGGTTTTCCGTACGAAGCT
Acheta_domesticus           ------------------AGTTAAGTAGTTTTATTG-----------------------AAGAGC---------------------AGTTTCACTTGAATTTTGATTTCGGCATGCTGAT
Gryllus_bimaculatus         ATACTTTTCCGGGCGTTTGGTAAAACAGTTTCATTGCCACTGAAAAAGGACTTAAAAGTGAGAGCCACAACCTGCGACCTCACAAAGGCGCCAAGTGGACCAGGATCGTGACATTCCAAT
Gryllus_longicercus         -----------------ATGTTTGCAGGTTTTGTTTCTACCAAATAACC----------AAGAAC------------------------------TGTATCTTGTTTACTATTTGATGTT
Consensus                                                                               *** *                               *      * *             *

Teleogryllus_occipitalis    ATTTT-------------------------------------------------------------------------------------------------------------------
Acheta_domesticus           CTTTT---------------------GTTATCGATGTGAGTTTTCCACTTGAGAGTGCCCTGAAATTGGGAGTCCTCATTATTTCACTGCCTTATTAATTGGAGTCCACGGAATGATACT
Gryllus_bimaculatus         TTTTTCAACATATTTAGGAATAAAGCGCGATCGGTTCAAATGAAAATGCGGAAAGTTACCAAGAAAT---AATTGTCAATATCTCCATAACGAGGTTTCTTAAGAAAGTGTAAAGGCACA
Gryllus_longicercus         TATTT-----TCTTCCTTAATT-----------------------------AAATTCTCCCTTAAATAAACATCTTTGCTA------------AGTGACAGAAATATGTTTAAAGGGCGA
Consensus                     ***

Teleogryllus_occipitalis    ------------------------------------------------------------------------------------------------------------------------
Acheta_domesticus           GTCATTTATTTATTTTTGTTTTATTG----------------GTTGGGGGTTTCGTGG-----------------------TGTTGAAGTTGGTAGACGTGCCCGAGCTATCCACTTGAA
Gryllus_bimaculatus         TTTGTTTACTATTTTACATTATGTCGGTGTTTTTTTCGCCTCGTTGGAACATCTGTAGAATCAGTTTTTATTCTATGATATTCTTAAAATTTAAAGTTAAATTTTTGTTCTATTATCGGG
Gryllus_longicercus         ATTGTTTCCTTGTTTTCGTTTTGCATGCTGTTATTATATC-------------------------------ACTCTGTTACTTATTGTATTTAGAAAATAATGTGAAATAAATTAATGTC

Teleogryllus_occipitalis    ------------------------------------------------------------------------------------------------------------------------
Acheta_domesticus           TTGGCCTGTCGGGCTGCAGCGCACAGCGACGCTGCCTAC---------TCCGCTTGCGACGAAGTAACCGCAGGATGAACCTTCGCTCGGTTCACGATCGCGCCAGGGCTCACTGCCTCC
Gryllus_bimaculatus         ACAAAATATCGAGAAAACGCGAAAATATGAACGGCTTGCGGGATTTTGCCCACTAGCGTGGCAGC-ACCCCAGCAAAACAATCTGTAAGTTCCACAAAAGTATGGAAGAGTAGTGCACAT
Gryllus_longicercus         TCTATACAAACAATTCTCGCTTACAGATGGACAGC------GCTTTTGTCTGTT--------------------------ATTTGTCTGTTAAACCTTCTAACCTGAACGTAGAATGTTA

Teleogryllus_occipitalis    ------------------------------------------------------------------------------------------------------------------------
Acheta_domesticus           CCGCTCAGAAG---GCGTGGAGGGGAGGAGGGGGTGATAAATAA------------------CACCTTTCCACTCCAGT---------CCGCGCTGATACTCGGTC--TCTGAACCTGAA
Gryllus_bimaculatus         GAGCGTAAAAGTTTGTACGTAAGGGGTGAGGATAGAACAAGGAAGAGGAAGGAAAGACGGTGCGCAGTTCCGGGCCCGTTTGTTAATACTGCTCCGACCTTCGTCTGGTGTTGACCGGAG
Gryllus_longicercus         ATGCTGTTATA-TCACTCTTAAGTAAAAAGAATAACAAAATCAAGA------------------------------------------TAATTTTGATAAATTGTT--------TTAAAG

Teleogryllus_occipitalis    ------------------------------------------------------------------------------------------------------------------------
Acheta_domesticus           AAATGGCGCGCGACGGGCCGCTGGAACGGCAGCCAATCGACTGAATCTCT--------------------------GTACTCCGTGTAATTGTGCGAATATATTTTTACCAACGACAGGC
Gryllus_bimaculatus         AAACCACGAAAAACTACCGATCAGAGCAGCGGCTGCTAGA--------------------------------------ATTCTGCGCCAGTCTGCAGCTGAATTCGTCCTAGCTAC----
Gryllus_longicercus         AAATTACTCTCAATTGTGAATTTGAAATCTGACCAATGTATTATTTCCTTTCAATGGATATGTCTTTGCTTCAAAGAGAACATGGGAATTAGTGTAGCTCAATTTCGCGCAATAACTGTC

Teleogryllus_occipitalis    ------------------------------------------------------------------------------------------------------------------------
Acheta_domesticus           CACCGACTGCAAGCTAGCAATCTCTTTGCAAACAGATCATCGCTGAGAGGAACTGGCGGACGA------------------TTTTTAATTCCACAGCAATTATATAAATGGAAATTT---
Gryllus_bimaculatus         ---------CTAGCTAGTG-----TCAAAATCCTGCCACCACCGGGA-----------------------------------TTCAAACCCTA---GAACCTCCCGGTCAGAGGCTC---
Gryllus_longicercus         AACTTACTCCTTGTAAAGAACG--TCATAGAGCAGGTATCCATTGATCTGGCCTCTCCGAAGAGATTTGTACAACTCTTTCTTTTAAATCTCGCGCCAATTTCTCGGGACGCAGTCCCCG

Teleogryllus_occipitalis    ------------------------------------------------------------------------------------------------------------------------
Acheta_domesticus           -----TCTAGCCGTTTCACGAGATACACAGTAACTCTGAAAAATGTCAATAATAATAATAAAATTGAAAACAAGAAAAGGAAAGCTGAGGTTCATACATAAGATTTCAATTAGCAAAAA-
Gryllus_bimaculatus         -----TTCAGTGATATCAGTTTATGCCCTGAGCC-------ACCTTCGCCGGTAATAATTAA--TGGGAGTTGTGAAAACAAAACAGAACACAACGCACACATTTTTATTTTTGTAATAT
Gryllus_longicercus         CGAGATCTAATAAGCTTGTTACCTTGCCTGCGACCTTGA-----CTTGCCTGCTTTAATCGC--TCGTATTCTGTAACAGCGCACATGATGCAAAGTACATAATTAAGAATAACGCAAAG

Teleogryllus_occipitalis    ------------------------------------------------------------------------------------------------------------------------
Acheta_domesticus           -------ATATGAAACGCGAATACATTGTCCTAGTAAACAACGCATGTTGTTACTTTTTATTACCATTTTTAAAGTATACTTGAAGCTTTCTACACACACACACACACACACACACGC
Gryllus_bimaculatus         AATCATGATGTAAAAAGTGTGTATATTTTTCTAATA-----------------------------GTTTTGCAAATAGTCTCGA--------------TCCCGCATATCCATAAAATTCC
Gryllus_longicercus         GATAATAAACAAAAACTCAAGTGTTAATTTGTAATC-----------------------------GTTTTCAGAAGAG--------------------------------AGAAATTTGT

Teleogryllus_occipitalis    -------------------------------------------------TACGTTTAAAGCAATTATTT---------------------------------------------------
Acheta_domesticus           ATATTATATTTTTCCTGGAAGAGATGAAAAAATATGTCATAATTTTTTATACATCACAATAAATAATGTTTTACTCATATGTAGCAGAGTGAAAGTAACAGGGAATAATAAGAGAAATAA
Gryllus_bimaculatus         ACAAAATGTTTTTCCTAGACAAG------------------------TGTACCTTAAAGTTAATTAATTTAGAACGATA------AGAATGGCAGTGCTAGGAGCT--------------
Gryllus_longicercus         ATGGAATTATTCGGCTGGTGTAGTTGAAAGGAT----------------CACATAATTACAATTTTTTTTAAACT---------------------------------------------
Consensus                                                     ** *       * *    *

Teleogryllus_occipitalis    ------------------------------------------------------------------------------------------------------------------------
Acheta_domesticus           CAAATTCCTATGCAGAAGCAATGAGCCCCTTGGAACTAAAAAACATTCTATTTCGTAGTTTCACAATTCAAGGAACACGCTGATATCATTC---------ATGGGAACGAGTACTTATTA
Gryllus_bimaculatus         CGAGTGTCTAAACTAATTTGTTAAAAACATTGCTCCTCAGGGGCTTCATATTTCTGTTGGTCCTAACAGAGAAAATATGAAGCTTTAGCCCTGTAAGTTAACAGGAAAATTTACCTGCAA
Gryllus_longicercus         CGGGGGTTTCTGCAGAAATGATG-------------------------------------------------------------------------------------------------

Teleogryllus_occipitalis    ----------------------------------------------------------------------------------------------AAATGTGAGGAATATTC---------
Acheta_domesticus           ----------TTCAACTGCCTAAAAATTGAAATTCTTTAATGCTTGTCCAAAACATGTTACACTGAACTGTCCCCAATGGTCACTAATAGTGCCAACAAAAATTGACCTTTTTAGGATTA
Gryllus_bimaculatus         AGGATTGACTTCCAACAAACAAAAATCTGGAAAACCGGAATCCTTG------------------------------------------------AAAAATATTTGGTTTTT-----AATA
Gryllus_longicercus         ---------------------------TGGAACCCCATAATTCTAATACAACGTGAGA------------------------------------AAAAAGTAATGATCTTC-----CTCA
Consensus                                                                                                 **            **

Teleogryllus_occipitalis    -----------------------------------------ACATGAGTCTATCTACACACATTT-------------------------------------------------------
Acheta_domesticus           AGGAGGCCGCGTCATCAAATTTGTTCGATTACAGTTAGCATACATAAAATCGTAAACATACAAACATAATTGTTACGATGTA----------------TTACGTACTCCTCGTAAACTC
Gryllus_bimaculatus         AGTAACCTACTGCATTAAA------------------------ATTAATCTATAAACATTCAAAC-----------------------------------TACATAATTTTAGGAG----
Gryllus_longicercus         AACAGAATTTATTAACTAA-CTATTCGTCAG-----------TTACAACATTACACAGGCAAACA----TGTTTCGAGGCATTTTACTCCAGTGTCTTCAACAGGATATTAAGATAAAC
Consensus                                                               *       *  ***  **

Teleogryllus_occipitalis    ----TATTCA--------------------------------------------------------------------------------------------------------------
Acheta_domesticus           TGTTTATTTACTTTAAACAACAAAATACAATTTAAATAATGGATCTAGGCCCAACTTCTCGGCTATCTCAAACTCCCTACTGAC-------TATCAAT--CATGCTCAAACTCATCAACT
Gryllus_bimaculatus         ----TATAAGATACAAGCCACACAAAATCATGTAGTAAATGAATACAAAGGTAAGTACT-AATTGTTTCAAAATTCACAATGGTACAATTATATCAATAACCTAGTGGAACTAATCTGCG
Gryllus_longicercus         ----TATAAAATATAAAT--CATAAAATTGTAAAACTAATAAATAACTAAAACACTATT-AAAAACCTTACAAAATATAATTATT----TTACCCAAATAACATGGTCAACATGATTT---
Consensus                       ***

Teleogryllus_occipitalis    ----------------------CCATTGTAACAACCGCGTC-------------------------------------------------------------------------------
Acheta_domesticus           CATTTTAGGTGCTCACCCAAC-CAATTGCATAAGCAGACT----------ACAGACAGACTGGATGACGCATTTCAATATCAGGCAGAGTCTTTAAAGCAGCGCCGTCCGACAAAATAGT
Gryllus_bimaculatus         CATTATCAGCACTAATC-------ATCGTATTAAC-------------------------------------------------------------------------------------
Gryllus_longicercus         --TTTGTGTTTCTCACGCGATGCCATTTTGTGCATTGTCTTCCGGGGGAAGCAGGGGGGAGGGAGGGCGGTCTATAAGCGGAGTTGCAGTCCTTACAATTAAGCTGATTGAAAGGACAGC
Consensus                                           **

Teleogryllus_occipitalis    ------------------------------------------------------------------------------------------------------------------------
Acheta_domesticus           GAAAGGCCTACAGGTCGGTTTGACGCATGTGAAAAC-------------------GCCAGAAACA-----------CACCCGAGAATGCTGCGAAAGGGCAGTTGAGGCTGCGGCTACAG
Gryllus_bimaculatus         ------------------TATTGCAAATTTCGAAAA-------------------TTCAGAA--------------------------------------AGTTCAACTTGCG-------
Gryllus_longicercus         AA---------------TTCAGCTTATATTGGAAAAACTGGTAAGAAATTCTCTTTCGGAATCAAAGAAAATTCTCGTGCATGGATATTACAAAAAGAACACTCCACCTTTGCCTCTCA

Teleogryllus_occipitalis    ------------------------------------------------------TGTCCGGGCTTTGTCCTTG--------------------TACCACTCGCAGCGATCTGCAACGCGC
Acheta_domesticus           CATAAGTGCTGAAAGAGCTTTTGAA-----TAACAGGTCT-------------TTCTGAGCTTTTGTCAT----AAGTAGGGACCGGATTTCTATGATTTTGCATATTTTCTCCATTGC
Gryllus_bimaculatus         ---------------------------------CAGTTTT--------------TTACCATACTTTGCTCT----ATCCGAGGCAAAAATTAATAATATTTGAGGTCTTCTATAAATT--
Gryllus_longicercus         TCT--------------TCTTGAAAATATCAACATCTTTGAGCACTATACAAATTTTAAGATTTTACATTTGGAAAACAAAGGACCAAAATTGAACATTTTAGAAAATTTTGAAATT--
Consensus                                                                         *        ***    *                       *  * *        * *

Teleogryllus_occipitalis    ------------------------------------------------------------------------------------------------------------------------
Acheta_domesticus           TTTAAGTCATAGCTAAAAGTAATTTCTTGGCATTTGGTTATAATTGGATGCTTTGTAGATTAATTTGTTAGTGCATATTTGATAGAT------------TTTTCCTGTTGGAGCATATT
Gryllus_bimaculatus         ----------------AAAAATGTTCTTAGGACT--------------------------------------------------------------------------------------
Gryllus_longicercus         ------TTAAAACTAGTAAAAAATCCT-----------AAAATCGACTGCCTCAACGACCAAACTA---------ATTTGATATACAGCCCTCTGCTAAGTATCCTAATAGACCGCCCT

Teleogryllus_occipitalis    ------CACCGGC-----CCGAATTGAAGCGAGCCACGCCCCACGGTGGGAGA----------CGCGGTCTTGCGGCGGCGGTGGTGTAGCGGTTAGCGTATCGGACTGCTAAACCCAA
Acheta_domesticus           CATTTGCGGCGGC---------------------------------------------------GGTGGTGTTGCGGCGGCGGTGGTGTAGTGGTTAGCGTAACGGACTGCTATACCCAA
Gryllus_bimaculatus         ---------------------------------------------------------------------TCTTGCGGCGGCGGTGGTGTAACGGTTAGCGTACTGGACTGTTGTACCCAA
Gryllus_longicercus         CCCCCGGACCTGCTTCCACCGGAGTACAATGCACAAAATGACACCGTATGAGAAAAACACCATGAATCATGTGGCGGCGGCGATGGTGTAACGGTTAGCGTACTGGACTGTTGTACCCAA
Consensus                                                                                        * * ********* *******  **********  ****** *  ******

Teleogryllus_occipitalis    GTTCGCGGGTTCAAACCCGATCGAAGCCGTGGA-TTTTTAAGGGCG---AAATCCTTCATACGCCTTCCTTCGGAGGGGAAGTAAAGCCGGCTGTCCCATG-TAGTAGATTTAC-TGCAT
Acheta_domesticus           GTTCGCGGGTTCAAACCCGATCGAAGCCGTGGA--TTTTAGGGGCGAAAAAATCCTTCGTACGCCTTCCTTCGGAGGGGAAGTAAAGCCGGCTGTCCCATGTTAGTAGATTTACTTGCAT
Gryllus_bimaculatus         GTTGGCGGGTTCAAACCCGATCGAAGCCGTGGATTTTTTAAGGGCGAAAAAATCCTTCATACGCCTTCCTTCGGAGGGGAAGTAAAGCCGGCTGTCCCATGTTCGTAGATTTACTTGCAT
Gryllus_longicercus         GTTGGCGGGTTCAAACCCGATCGAAGCCGTGGA-TTTTTAAGGGCGAAAAAATCCTTCATACGCCTTCCTTCGGAGGGGAAGTAAAGCCGGCTGTCCCATGTTCGTAGATTTACTTGCAT
Consensus                   *** *****************************  ***** *****   ********* ****************************************** * ********** *****

Teleogryllus_occipitalis    GTAAAAGATCCTCAAGCCTAAATGATGCCTATGGGCAAAATTCAAATCGCCTGTTTCGTCGCCCAAGTAATACCTTCTTTCTCGTAT---TGAAAGGAGTGTAACGAAATTTGTGTAGTC
Acheta_domesticus           GTAAAAGATCCTCAAGCCTAAATGACGCCTCTGAGCTAAATTCAAATCGCCGGTTTCGTCGCCCAGTAATACCTTCTTTCCTTAGTTGGTGAAAGGAGTGTAACAAAATTTGCGTGGTC
Gryllus_bimaculatus         GTAAAAGATCCTCAAGCCTAAATGACGCCTCTGAGCTAAATTCAAATCGCCTGTTTC-TCGCCCAAGAAATACCTTCTTTCTCTGCTATATGAAAGGAGCGTAGCGAAATTTGCGTGGTC
Gryllus_longicercus         GTAAAAGATCCTCAAGCCTAAATGACGCCTCTGAGCTAAATTCAAATCGCCTGTTTC-TCGCCCAAGAAATACCTTCTTTCTCTGCTAGATGAAAGGAGCGTAGCGAAATTTGCGTGGTC
Consensus                   ************************* **** ** ** ************** ***** ****** ** *************      *    ********* *** * ******* ** ***

Teleogryllus_occipitalis    AGGCGGAAGAGCAGGCCTCATGGCCTGAGTCTCGCCTCGTAAAATAAGGCAAATTCAATTCAATTCTGTACTTCAGAAGTGCGCACCTACTGGCCTATAGTAACCTCACTGAACTTTATG
Acheta_domesticus           AGACGGAAGAGCAGGCCTCATGGCCTGAGTCTCGCCACGTAAAATAAGGCA-ATTCAATTCAATTCATTGAATTATGGGAGCATATTTCGCATATTTCGTTG----AAATAGAGAAAATA
Gryllus_bimaculatus         AGACGGAAGAGCGGGCCTGATGGCCCGAGTCTCGCCACGTAAAATAAGGCATATTCAATTCAATTCA----------------------------------------------------A
Gryllus_longicercus         AGACGGAAGAGCGGGCCTGATGGCCCGAGTCTCGCCACGTAAAATAAGGCAAATTCAATTCAATTCA-----------ATCATGTTGACCATGTT----------ACTTGGGAAAATA
Consensus                   ** ********* ***** ****** ********** ************** **************

Teleogryllus_occipitalis    TCCATA------------------------------------------------------------ACGCCTCTTCTAACATCCAAATGCACTCCAAATGAGTGAACAAGTATTGAATCC
Acheta_domesticus           TTCATGGTTTCGTAGAATTTATTTGTGAGCGTTCATTGTCGGGAAAGCTTGTCACAATTTTCATTTTCCCCCTTTCTTTTCTATAGACG---ATAAAATACTTGTGCAAATGAAGGATAT
Gryllus_bimaculatus         TTCATG----------------------------------------------------------------------TAGTGTATGGACG---GACGGACGGACGTGCCGACCGA------
Gryllus_longicercus         ATTATA----------------------------------------------------------------TTTTGTAAGTTTTTTAATA---GTGGTGTAATTGTACCAAAGTAAAGTCT
Consensus                      **                                                                           *    *                *  *

Teleogryllus_occipitalis    GGGTACATCAACTATTCAGATCATTAGTCATGTAGTTAACTCTATGATACATACCGGGAGGTGGCGGGTTCCTCCCTGTCTGG--------------------------AACGTAAAGTC
Acheta_domesticus           TTCTTGATATATTTTTTAAACCAGGCAGAAAAAGT--------------------------------------------------------------------------GAAAGAAGACC
Gryllus_bimaculatus         ----------------CGGACCGGCAACAATATTT---------------------------------------------------------------------------TAACAAAAAT
Gryllus_longicercus         AGCCTACTAAATTGTTTCTAATAGTAACAAATATTATTTATTACTCATAGATA---AGAACTTTAGTTTTATTATTTTTTTGGCTCGGAATTTCTTTAGGAGGTAAGCAAGAAGAAGAAA
Consensus                                      *         *                                                                                    **

Teleogryllus_occipitalis    CGCTGCCGACTTACACGGCAGTTTGTCATGTCTTGGTATCTCACAC-------------AGCGACACACCGTCCATGTAGAGAAAACTCTA-----------------------------
Acheta_domesticus           TATAATCAAGGTACTTCACTAAAGTAAATATTTACTTCAATTATCTCTCCCCAGAAGTAACTTAGTACATTATGTTTGCA----------------------------------------
Gryllus_bimaculatus         TATTATGAACGTACT--------GGAAATACATTCAGTGTTCATCT-------------TTTGGCACATTGTGTATACGA----------------------------------------
Gryllus_longicercus         ATTATCCAATACGTTTTAATATACAAAATGTTTTCAATAACAACTTACCTTGTTAATGGAATTAATACAAAACGTATGGAGATTAACATCGACGACTTCTGGTAGGGGTACGCTAGTTAC
Consensus                           *                  **   *         *                       ***       *

Teleogryllus_occipitalis    -------AGCTCATTACTGGATTCCCAACATATTTAGTAGTTTAGCCCATGTACGTTTGTTTGTGTGTACGAAGAAAGGACAACAACGAAAAAGTAAAGATTGAGCC--AAAGCAATGGG
Acheta_domesticus           -------TATTATTTTTTTTATTCCAAGCATGCTT----------CTTCTGAATT-----------TTATCAAGAAAGGTTTATACTGAAAAT---------------GAAATAGTTAA
Gryllus_bimaculatus         ------TCATTCATTTTTGTTTTCT----------------------------------------------------------------AAT-----------------ACATCAGACGG
Gryllus_longicercus         ATTATATTACCTGCTTTTAAACTCAAAACATATGTGAGTTGTTCACCCGCACATG----------GTGCAATGTCATGTGTACATTGTCCATAT---------TCCTAGCCGTGATTAA
Consensus                                 *  *    **                                                                   *

Teleogryllus_occipitalis    CCGCCAAACACACTTCAAAT-------TTCGACAAGTTAGTTGACACCGTGTGACAAAATGCTGGCAAATTTTGACTCTATGCGTTTTCCATGATAACGTCTTGCACGGTAGAGTAGG--
Acheta_domesticus           TTTTAATTGAATATTAAAGTGCATATTTCAGAATTGTTAGGTCATA-------------AATGCATAATTTAAAAAGCATATTTCAA-------GATATTTAGGGCATAAAAA------
Gryllus_bimaculatus         TATTGGATGAAACCTAAAAC----------------------------------------------------------------------------------------------------
Gryllus_longicercus         TATTTAAAGAATTCTTTAAT-TATCTTTTATATATTTAATGTGACATTTTTT-----CCTGTTATATATTTTTAATTTGATATTTTA---------ATATTTTAGCCTTAAGAATGAGAA
Consensus                            *    *  *

Teleogryllus_occipitalis    -GAAAAAGTCTGCGCGTCCGCCGCGGTGAATGGTGCGTGCGAG----------------------------AAAGAGCCAAAAAGCAGAGGGATCGAAGTGCTCGAGCTGTGTCTTTACG
Acheta_domesticus           -----------TCCGCCCCCTAGTAATAAGAG---------------------------------------AATATTCTGAAAACCACAATAA------------------GTCTGTTAG
Gryllus_bimaculatus         ----------------------------------------------------------------------------------------------------------------TATGTTAA
Gryllus_longicercus         CGAATAGAGTTTTGAAACGTTGGCAGTAACAGGTTCATATTACATAACTTCACTCTCTGTGCCCGACGTTTAGAGTTCTGGGAGTAATAT----------------------TGTAACAG
Consensus                                                                                                                   * *

Teleogryllus_occipitalis    CTGAATGCGTAATAACCGATCTGAATAAGAAAAATGGGGCTTTGGCCGGCTAAAGGCGTAGTTTGTCGGGCTGTGGAGAGGTCGGCCGCGGAGGTCTGAGACCGCTCGTAAA--------
Acheta_domesticus           CACACTTTGCAAAGAATGATTAAATTTTGAAGAA-------------GACATAGTGT----TTCCTGGGCGTAAAGAAATCGTATTCCTTCCCGTT------------------------
Gryllus_bimaculatus         AACAATTTCTAATAAACTATT---------------------------------------------------------------------------------------------------
Gryllus_longicercus         TACATCCATCCACGCCCTATTAAACGGGTAAAAA------------GGCAGAATACGCGATGTGGGCGCTTGAGGAAAATCCGCCCTTCGCCGCCATCGCCCCCGCGTCAGCTCCCGCC
Consensus                      *       *      **

Teleogryllus_occipitalis    ------------TCGGCAGTAAATCCCCCGG-------------CGCGGGCTCAGGAGCGAAGGACTGGCGGTCGCGGTTGGTGGCGGTGGTGCTGGCGGTGGTGGTGGCGGAGGCGG
Acheta_domesticus           -----------------------CTCTCTCGG---------------------------------------------------------------------------------TACTCGT
Gryllus_bimaculatus         ------------------------------------------------------------------------------------------------------------------------
Gryllus_longicercus         CTAAGAGTTTCCTTTTGTCGTTACTTTCTCGGAAACGTTGTCCATTCACCATCTTAGG---------CTCGTGTTCACCACTTAAACCGCT----------------CGGCCCAGTCGT

Teleogryllus_occipitalis    AGGCGGAGGCGGAGGCGGAG---------------------GCGGAGGCGGTGGCGGTGGTGGAGGCGGTGGCGGTGGCGGTGGTGCTGGCGGTGGTGGTGGCGGAGGCGGAGGCGGAGG
Acheta_domesticus           GAGCGTGAGTGAAATTTACG-----------------------------------------------------------------------------CGCCAACCACGCTGGTTCACGGA
Gryllus_bimaculatus         ---------------------------------------------------------------------------------------------------TTCATTACGCACCTACAAAAA
Gryllus_longicercus         TACCGGATGCAGAACTTAAGAGTTTCCCAGAAAGAAATTAAACAAAAATTGAACTGTCGGAAATCTAGCCAAATATTTTCAACGGGCGGGCGCCGATACGTGATTGGGCAAAGTTAGGGC
Consensus                                                                                                              **

Teleogryllus_occipitalis    CGGAGGCGGAGGCGGAGGCGGAGGCGGAGGTGGAGGTGGAGGCGGTGGCGGAGGCGATGGCGGAGGCGGTGGCGGT------------GGCGGTGGCGGTGGCGGTGGCGGTGGCGGTGG
Acheta_domesticus           GGATTCCACGACCACGG----------------AGAGGGGCGTCTGGCAGGAGGACCAGCGTGACACCCAGCAACCTCCGACTATTCTGCGATTTTTCTAG------GAAGGACACTCA
Gryllus_bimaculatus         AAATTTCAAAACTATA------------------------------------------------------------------------TCCAATTCTC-------------AAACTTTTA
Gryllus_longicercus         TTATTTCGAAACCACTCGTTCAAATAGAAGGAACGGAGGGCTACCTCCTCGGTGGATGAGTGGGAAGCAGAA-----------GTTCGGTGGTAAGTTTAGCTGAGATGGAGATCCGTA
Consensus                         *                                                                                      *

Teleogryllus_occipitalis    CGGTGGCGGTGGGCGGTGGCGGGTGGCGGTGGCG-------------------------------------------------------------------------GTGGCGGTGGGTG
Acheta_domesticus           ACATAACTCAAATCCCA---------------------TCTTCGCGATGGAATATGTCACGGCATTTCGCATCTTTCTCAGTCGTTCCGCTAGATTTCGCACGATCGCGACCGTATGAA
Gryllus_bimaculatus         TCAGGGTTTATATCCC-----------------------CTCAGCAATACAATGGATTTCTGGATCTCTTTTTAATTTAAGAAATGCTA--GGATTTCTTATTTTCTTCTCTGTTTGAA
Gryllus_longicercus         TTGAGACAAAAATCACCGACGGGCAAGAATAGAGCAACGTGTTAGTTACACCATGTGTTATGGGATT--ATTGTTTTCTTAATTAGCAACTGGATTTATCCTTTTCTTGGTTATCAGGG
Consensus                                *                                                                                                    *  *

Teleogryllus_occipitalis    GCGGTGGCGGCGGTGGCGATGGCGATGGCGATGGCGATGGCGGGGCGCG-----------GCCGTG---------------GCGGTGGCGGTGGCGGCGGTGG--------------
Acheta_domesticus           GCAGTGCTTGC---GATCATGCGACTTCTACCGGCAAGACCGAGAAAAG---ACGCGAAATGCCGTG---------------ACATATTCCATCGCGAAGATGGGATGTGAGTTATCTT
Gryllus_bimaculatus         AAAG-------------------GTTAGTTTAGCAAAACTAAG-------------AATATTAGA---------------ACATATACTA--------------------------
Gryllus_longicercus         CCCA------------AAAATGTATTTTTCTGTCGAAAATAGGACGTATTTTCGCTCAATATTACATAACTTTCTCAATACTACGTATATTACTGCA---------------------
Consensus                                            *        * *      *                                        *

Teleogryllus_occipitalis    ------CGGCGGTGGCGGCGCCGGGTGGCGGCGGT-------GGCGGCGGTGGCGGCGGTGCGGCG-----ATGGCGGCGAT----------------------------------------
Acheta_domesticus           GAGAGTCCTTCCTAGAAAAATCGCAGTATAGCGAAATAGCTCGGGTGAATTTAGCCGCTGTCTGGAGCACAATTGGCTCAGATCGATTTCAGAACTTG--AGATGGCTGCTTCAAACCGG
Gryllus_bimaculatus         --------------AAATATCACATAGAAATAAA--------GATGTATTTA----------------------------------------------------------------------
Gryllus_longicercus         --------------TACAAACGTACGTATATGAG--------ATGGTATTAAGTGGTTCTCCAGAA------TGGCTCAGAT--ATTCCGGAATTTGTTTTCTATTTCCTTTGATCTGG
Consensus                                        *                        *

Teleogryllus_occipitalis    -----------------------------------------GGCGGCGGTGGCGGCC---------------------------GGTGGCGGCGGTGGCGGCGTGGCGGCAGCGACGTCT
Acheta_domesticus           TTTTCCGGTTCATTCAAGGGAAATGCCTATTGAGTTCCCAAAGAAACTGCACGTGCCCGATTTTTATTCCTTTTCTTAGTCCAAGGCAGCGTCGTTAGCCAAATTGCGTAACTAAGAGTT
Gryllus_bimaculatus         --------------------------------------------------------------------------------------------------------TGCATTACTCACCATT
Gryllus_longicercus         GCTAGCTATTCTTTCTTGCTGCATACTCTTGCCAGCTGCCATCTATCAACGCCAACCCCGTTTTTTTCTTTGTTTTTTT-----------------------------------------TT
Consensus                                                                                                                          *

Teleogryllus_occipitalis    TCCGGCGCTGGCGTGCGTGGCGGCTGGCGGGCTGCGGGTGG-----CAGCGGTG------------------------------------------------------GCAGCGGTGGCG
Acheta_domesticus           TTTATGTTTGATATGAGCTAAAGTTCATTTACTTCCAATGAGAATCCTAGACGCAAAAAAGTATTTCACGTAAATGAACGATGAATGTA------------------AGATCGCTGGCA
Gryllus_bimaculatus         TCTATTTTCACTGTCAGC-AGGTTTCATTTGGATGGGATGA-----TGCATCTGAAAAAATATGTAATGT-----ATTAATGAATATGCTTTTTTTACTCCCATTGTACCTTACTCTCA
Gryllus_longicercus         TTAATTTTTTTTTTAATCTCAATTTTTCAGTTTTCTCATAG-----TAGAGCTGAAATATTATTTTAGGAATATTACAAAACAGTAAACCCCAGTTATCA-------ATGGCAATTTTC
Consensus                   *            *          *             *                                                                            *

Teleogryllus_occipitalis    GCGGTGGCGGCGTGGCGGCGGTGGCGGCGGTGGCGGCGGTGG--CGGCGGTGGCGGCGGTGGCGGCGGTGGCGGCGGTGGCGGCGGTGGCGGCGGTGGCGGCGGTGGCGGCGGTGGCGGC
Acheta_domesticus           CTTGTTAATACACAGATCAACTGTGATAAAGTTGAATGACCGACAGCACGTGGGAAAGTGGACTATCAATTCCA----------------------------------------------
Gryllus_bimaculatus         CTTGATGGCACTC------ATTACGATAAGATTTCATTATGG------------------------------------------------------------------------------
Gryllus_longicercus         ATTGAAAAAATGTGGAGCCGTTTGAAGAAAATAGCGTTTAAG--CGCTACTGGCGAAGTGGATAATAAAGCTC-----------------------------------------------
Consensus                      *                 *                   *

Teleogryllus_occipitalis    GGGGGCGGCGGTGGCGGCGGTGGCGGCGGTGGCGGCGGTGGCGGCGGTGGCGGCGGTGGCGGCGGTGGCGGCGGTGGCGGCGGTGGCGGCGGTGGCGGCGGTGGCGGATGCGTCGGCGTC
Acheta_domesticus           -------------------------------------------------------------------------------------GACACGAACACAATGCTCTCTTGCCTTTAAACGTC
Gryllus_bimaculatus         ----------------------------------------------------------------------------------------------------------------TACACGTT
Gryllus_longicercus         --------------------------------------------------------------------------------------------------------------------CTTT
Consensus                                                                                                                       * *

Teleogryllus_occipitalis    AGCGTCGGCGTCGGCATCAGCGTCAATG--------------------------------------TTGAATGGATTTG--------------GGAGTGCGCAGGAGAGCAGGAATGGAG
Acheta_domesticus           TCAGAGGAGTCTAACGTTTGTTTGAAGGGCTGCTTATCTCGTGCATTGTTC-----CCGTCAATTATTGATCCAACTTGTTTTCGACGTAGCCCATATTCTTTCCAAAGATGTAATAAGG
Gryllus_bimaculatus         CTTGACCAATCGCAGATTAGTTTACAAA-----------------------------------------------------------------CAAATTATTGCCAATTAATCATTGAGA
Gryllus_longicercus         TCACAGAAAACTATCGATATTTTGAAAA----CCTTCATTAGGCGTCATTTGAATAACATTAATTCATAAATGGAATAGAGTTTA--------CAAATTGATAGAAGAGAACATATTACA
Consensus                                         *  *                                                                       *       *         *

Teleogryllus_occipitalis    GATATAGTAG-------------------GTTGCAGGACCTTATGAAAGTTGCGAAATTATATTTT------------------------------------------------------
Acheta_domesticus           ACTTCAATAACCAAAAGCCTTAAATTTATATTCTTATTATTAAGCAAGAAACCTAAAAATATGTACAAAAAACATAACACTAAAGAACAATGAAATT----TCACGCAACACACTGTAAT
Gryllus_bimaculatus         ATCGTTATAG----------TAAATGATTAGTTTCATAACTTCGTAAAGTAATTAGTCTTATTT-----------------------------------------------CACTTTGAT
Gryllus_longicercus         GGCGTGATAG---------------------------ATCTTTCAAGAAAATTATATTTGTATATTAAAAAAAAAAGAATTCATCGAAATGAACTTTACATCATATTAAATATCGCAAA
Consensus                          **                             *       *            * * *

Teleogryllus_occipitalis    -------------------------------------------------------TTTTATGTTTCAAGTTAGAATTTTTA-------------------AAAATGAAA-----------
Acheta_domesticus           CATGGATTTCTATACTCAGAATACGGAAATGAATATCAATTCTGTGGCCACGGGGTAAAAGCTCTAAAGTAAAAATCTT----AGTTTGA----------AGAAAACAA--------GGG
Gryllus_bimaculatus         CCTATGTGACTA-------------------------------------GCGCAGTGATAGTGGTAGAGGAAACATTT------GTCTGACGTTAAGATCAGAACGCAATTCATTTTGTG
Gryllus_longicercus         TAGCGCGTTCCGTTCGAACAGTTTGAAGAACACTTT------------------TGGAGTCGTTTGTGAGAAAATCTTTCAAGATCTTTCATTAAGAGAATGGTGCAG---GTCTCATA
Consensus                                                                          *        *   *  *   ** *                     *      *

Teleogryllus_occipitalis    ------------------------GGTTTTCTGGGGATATTGATGTCATTCATGTAGTTTAT--------------GGGGAATATTGATTTAGAAAGAGAAAG-----------------
Acheta_domesticus           CTAGATTACTGAGCCAACTGTT----TTTCCTAAGCTTA---ACGCTATTTTCTAACTTTTTTTTCTCGTTAAGGCGGCAATATTTGAATT------AGGTCGTTTTATTACAATAAAAT
Gryllus_bimaculatus         TTATATGCTGCTGCCGTCTGTC----TGTCCTTAACAAT---ATGCCAATTCTTTAGTTT---------------AAAGAGAATTGAATCA----GAGGTTGTGTTGACACCTTCAGAA
Gryllus_longicercus         TTCATTGATATCCCCGATTTTTCGAATTTTTTTAAAAAA---ATGTGGAAATACAAAATTGT----------AGATAGAGATTTCTCGATTATACAAAGACTGTTGTA-CACAATTAGGT
Consensus                                             * *  *          * *          *  **                    *    *   *       **   *

Teleogryllus_occipitalis    ----AGAGAGAAGAAGAGGGAAACCGAGATGACAGATGCTA-------------------------CACGATGACACGC-----TTAAAAACAGCGCAC--------------------
Acheta_domesticus           AT----AGATACTTAACTTGAAGTTTTCGTAA-----ACTCAACTTTCGCGAAAACTGATTTTACCCCGTGACCACAGAA-----TTAAAGCCATCACGTACAGTTTTCCTTATTTCCTG
Gryllus_bimaculatus         AT----AAAAGAAAAAGTGAAAAT-------------------------------------------GGTAACCTCAGAA-----CTA------------------------TTTCCACA
Gryllus_longicercus         ATTTAAAAAAAGGCATGTACAATTTTTTGTAAAAGTTAGTAGTTTTGCGTAATAAATGCTTGAAATTTACAACAACAAAAAATTCTCAAAGCAGCTCC-------TAATTTTTTTCTTTA
Consensus                         * *     *     **                                                 *   **          *

Teleogryllus_occipitalis    -----------------------AATTTGTAGAAAT-----------------------------------------------------GCAGAA------------AAATACTAATGA
Acheta_domesticus           AACTTTCTGTTTAGAATGTTACGCGCCCGTCGTTAATTGCGCTGCTTTTCCCGGTTTACATGGTGCTTGGCGTTGCTGCAATGTCATACACAAAACCACAGCACTACTGTGTAATAATAA
Gryllus_bimaculatus         AATTTTCTACT-----------CTCTCGTAG-----------------------------------------------------------------------------GTGGACTGATAA
Gryllus_longicercus         AATTTTCTCCA-----------CGTATGTAATAAG-------------------------------------------------------------------------GGATGTTAAGAA
Consensus                                               **                                                                                    * *  *

Teleogryllus_occipitalis    GTGTAA------------------------------------------------------------------------------------------------------------------
Acheta_domesticus           ACATGA-GTCACACTTTTGTTTATATTTCCCTTCCAAGGGTTCGAGAAGTGTGTACTGGTAGGGACTGCGTAGCCAACGTCAGGCTG---TAGGCTGAATTCACCGCAGTAATATTCCGC
Gryllus_bimaculatus         ACCAAA-GCCAC------------------------------------CTGCGACCGGGTAGG------ATAGCTGTCCGCTAGGTGACCTTGGCTGAATCCTGCATGTCAACTGCC---
Gryllus_longicercus         ATACGGCGGCAGAATCTTTTTTGGGTAT-TATTCAATTATTTCGAAAAGTACA-------------------------------------------AAATCAAATATAGAGATTTTCCTC

Teleogryllus_occipitalis    ------ATAAAGCGATGAAATGTGA----------------------------------------------TACAGAAGAAACAAACAGTAAGAAAATCGATA-----------------
Acheta_domesticus           TTAATGTCGAGAGTGGTGTACATAACCAGTTCTTTTTGTGTCTTGTCTTCATAACTAAGAAGCTTTAAAACTGTGTGATTCTTCAAATGTTTCTAAGTTGATATCATTCGTGACAGTGGT
Gryllus_bimaculatus         ------ACGAAACGATTAAATATAA--------------------------------------TTTTAACTTATAAAAGTCACCATAAGTCGTAAAATTGATAGTGTCTG----------
Gryllus_longicercus         T-----CAAAGACTGTTGTACAAAA----------------TTATTTATGTAGAAAAGGTCATGTACAATT-------TTTTGAAAAACTACTAGTTTTGCGTAATTTGCGCTTAAAAC
Consensus                            *         *    *                                              *            *         *  *

Teleogryllus_occipitalis    -----------------------------------------------GTCAAT------------------------CAGATGAGAGAGATCACTA------------------------
Acheta_domesticus           TT---CTAATTAACCCCGTAAGTATGTAAATTCATCACGTCTCTTTTGTTACTTTTGTGGACGTGCACTATGAAATCCTAGAGAAAGGAACTCACATAGCGCCGACATTG----------
Gryllus_bimaculatus         -----------------------------------------------------------------------------ACGGAGAGAGTGTCTGAC-------------G-----------
Gryllus_longicercus         TTACAATAACAAACCTTCT-----CACAAACTGCTCCAATTTCTT---TAAATTTTCTCCACATACAGTAAGAAAT-ATGGTGCTAGTCTCTTTT-----TCTAGTATTGCTCGTTTTTT
Consensus                                                                                                     *  **

Teleogryllus_occipitalis    ----------GAGAGAATTGATGAACAAACGGGTACATGAAGACATTTTATAGAAAAG--------AATGAAAGGCAATTTCGTGTATA-------TGCTTACAAGAGAGGGAAA----
Acheta_domesticus           ----------AAAAAAGCTTCCGCATTGTAT-TTAAATTTAAA---CTTGGAAATCAGGACAAACAACGGGCACCCCGTTTAAATGATACTCGAAATATTTACATGAATGGTTGTCGGG
Gryllus_bimaculatus         ----------GAGAGAGTGTCTG--------------------------------------------ACGGAGAGAGTGTCTGACGGAGA---------------GAGTGNNNNNNNNNN
Gryllus_longicercus         TCGAAGGTTGCAAAAGAATTTCCAAACTACA----AAATTGAAATATTTGG--------------CCACAGAAAATGTTTGCTGTAGT-----AAATTTTAAATAAATAA--------
Consensus                               * * *                                                        *     * *     *                   *

Teleogryllus_occipitalis    ------------------------------------------------------------------------------------------------------------------------
Acheta_domesticus           TTTACTCTTCCATGCTAT------------------------------------------------------------------------------------------------------
Gryllus_bimaculatus         NNNNNNNNNNNNNNNNNNNNNNNNNNNNNNNNNNNNNNNNNNNNNNNNNNNNNNNNNNNNNNNNNNNNNNNNNNNNNNNNNNNNNNNNNNNNNNNNNNNNNNNNNNNNNNNNNNNNNNNN
Gryllus_longicercus         ------------------------------------------------------------------------------------------------------------------------
Consensus

Teleogryllus_occipitalis    ------------------------------------------------------------------------------------------------------------------------
Acheta_domesticus           ------------------------------------------------------------------------------------------------------------------------
Gryllus_bimaculatus         NNNNNNNNNNNNNNNNNNNNNNNNNNNNNNNNNNNNNNNNNNNNNNNNNNNNNNNNNNNNNNNNNNNNNNNNNNNNNNNNNNNNNNNNNNNNNNNNNNNNNNNNNNNNNNNNNNNNNNNN
Gryllus_longicercus         ------------------------------------------------------------------------------------------------------------------------
Consensus

Teleogryllus_occipitalis    ------------------------------------------------------------------------------------------------------------------------
Acheta_domesticus           ------------------------------------------------------------------------------------------------------------------------
Gryllus_bimaculatus         NNNNNNNNNNNNNNNNNNNNNNNNNNNNNNNNNNNNNNNNNNNNNNNNNNNNNNNNNNNNNNNNNNNNNNNNNNNNNNNNNNNNNNNNNNNNNNNNNNNNNNNNNNNNNNNNNNNNNNNN
Gryllus_longicercus         ------------------------------------------------------------------------------------------------------------------------
Consensus

Teleogryllus_occipitalis    ------------------------------------------------------------------------------------------------------------------------
Acheta_domesticus           ------------------------------------------------------------------------------------------------------------------------
Gryllus_bimaculatus         NNNNNNNNNNNNNNNNNNNNNNNNNNNNNNNNNNNNNNNNNNNNNNNNNNNNNNNNNNNNNNNNNNNNNNNNNNNNNNNNNNNNNNNNNNNNNNNNNNNNNNNNNNNNNNNNNNNNNNNN
Gryllus_longicercus         ------------------------------------------------------------------------------------------------------------------------
Consensus

Teleogryllus_occipitalis    ------------------------------------------------------------------------------------------------------------------------
Acheta_domesticus           ------------------------------------------------------------------------------------------------------------------------
Gryllus_bimaculatus         NNNNNNNNNNNNNNNNNNNNNNNNNNNNNNNNNNNNNNNNNNNNNNNNNNNNNNNNNNNNNNNNNNNNNNNNNNNNNNNNNNNNNNNNNNNNNNNNNNNNNNNNNNNNNNNNNNNNNNNN
Gryllus_longicercus         ------------------------------------------------------------------------------------------------------------------------
Consensus

Teleogryllus_occipitalis    ------------------------------------------------------------------------------------------------------------------------
Acheta_domesticus           ------------------------------------------------------------------------------------------------------------------------
Gryllus_bimaculatus         NNNNNNNNNNNNNNNNNNNNNNNNNNNNNNNNNNNNNNNNNNNNNNNNNNNNNNNNNNNNNNNNNNNNNNNNNNNNNNNNNNNNNNNNNNNNNNNNNNNNNNNNNNNNNNNNNNNNNNNN
Gryllus_longicercus         ------------------------------------------------------------------------------------------------------------------------
Consensus

Teleogryllus_occipitalis    ------------------------------------------------------------------------------------------------------------------------
Acheta_domesticus           ------------------------------------------------------------------------------------------------------------------------
Gryllus_bimaculatus         NNNNNNNNNNNNNNNNNNNNNNNNNNNNNNNNNNNNNNNNNNNNNNNNNNNNNNNNNNNNNNNNNNNNNNNNNNNNNNNNNNNNNNNNNNNNNNNNNNNNNNNNNNNNNNNNNNNNNNNN
Gryllus_longicercus         ------------------------------------------------------------------------------------------------------------------------
Consensus

Teleogryllus_occipitalis    ------------------------------------------------------------------------------------------------------------------------
Acheta_domesticus           ------------------------------------------------------------------------------------------------------------------------
Gryllus_bimaculatus         NNNNNNNNNNNNNNNNNNNNNNNNNNNNNNNNNNNNNNNNNNNNNNNNNNNNNNNNNNNNNNNNNNNNNNNNNNNNNNNNNNNNNNNNNNNNNNNNNNNNNNNNNNNNNNNNNNNNNNNN
Gryllus_longicercus         ------------------------------------------------------------------------------------------------------------------------
Consensus

Teleogryllus_occipitalis    ------------------------------------------------------------------------------------------------------------------------
Acheta_domesticus           ------------------------------------------------------------------------------------------------------------------------
Gryllus_bimaculatus         NNNNNNNNNNNNNNNNNNNNNNNNNNNNNNNNNNNNNNNNNNNNNNNNNNNNNNNNNNNNNNNNNNNNNNNNNNNNNNNNNNNNNNNNNNNNNNNNNNNNNNNNNNNNNNNNNNNNNNNN
Gryllus_longicercus         ------------------------------------------------------------------------------------------------------------------------
Consensus

Teleogryllus_occipitalis    ------------------------------------------------------------------------------------------------------------------------
Acheta_domesticus           ------------------------------------------------------------------------------------------------------------------------
Gryllus_bimaculatus         NNNNNNNNNNNNNNNNNNNNNNNNNNNNNNNNNNNNNNNNNNNNNNNNNNNNNNNNNNNNNNNNNNNNNNNNNNNNNNNNNNNNNNNNNNNNNNNNNNNNNNNNNNNNNNNNNNNNNNNN
Gryllus_longicercus         ------------------------------------------------------------------------------------------------------------------------
Consensus

Teleogryllus_occipitalis    ------------------------------------------------------------------------------------------------------------------------
Acheta_domesticus           ------------------------------------------------------------------------------------------------------------------------
Gryllus_bimaculatus         NNNNNNNNNNNNNNNNNNNNNNNNNNNNNNNNNNNNNNNNNNNNNNNNNNNNNNNNNNNNNNNNNNNNNNNNNNNNNNNNNNNNNNNNNNNNNNNNNNNNNNNNNNNNNNNNNNNNNNNN
Gryllus_longicercus         ------------------------------------------------------------------------------------------------------------------------
Consensus

Teleogryllus_occipitalis    -----------------------
Acheta_domesticus           -----------------------
Gryllus_bimaculatus         NNNNNNNNNNNNNNNNNNNNNGG
Gryllus_longicercus         -----------------------
Consensus
```

**Tocc.6G0002000.1_exon1**

```
Teleogryllus_occipitalis     ------------------------------------------------------------------------------------------------------------------------
Acheta_domesticus            PASIHFVS*FEVNYEFLSKILHHQGAISLTSLF*TLQWQQSSA*-----VIIP----------SYVVCLLNLFIFWKNSQSSVDVQLCICIYGVTIQLILI*Y*H*TSSDEGTHSTLQLI
Gryllus_bimaculatus          ----------GKREQKIKKCTHRH---FINDLIYILLLRETAR---------P----------TFGCY*VNC-TEWEPTK----------LHAYTQRXXXXXXXXXXXX*RERVFVINTIW
Gryllus_longicercus          -------------------MH*HF-------------QTTRFS*FRRIVFPFPLLLYK*N*TYLYTSFNS-INWD**Q--------LCIFNTVGKL-----------------QQIC
Consensus

Teleogryllus_occipitalis     ------------------------------------------------------------------------------------------------------------------------
Acheta_domesticus            YSILLSIFKNTC*GYLFLIRI*RSEYLPSLRFLQWLIQVEV*VKVISDIINTVRVWCS-IQ*VV*C-------LILCLVA---CWI*TRGLGRWVLRIGFG*FVIY*YYLSIPGRHI---
Gryllus_bimaculatus          YAIPTSKLKH------FL-------YIPVFRF-----------------*NSSKLLC---RFCVTCTFNYSSQLLVHLIAVPCCDTNG*TIHR---------FISYY*FMIILIRKIIRS
Gryllus_longicercus          RKN*YSK*PNS-----TL-----SAFLSCFRT---------------------*LYCSMLQMAKNII------LRTSILSMSICKS---------------------FKSILIYSI---
Consensus

Teleogryllus_occipitalis     ------------------------------------------------------------------------------------------------------------------------
Acheta_domesticus            GGGGRV**WTCI**HCLKRVK-------*FY*RA----VSLEF*FRHADL-LLSM*VFHLRVP*NWESSLFHCLINWSPRNDTVI------YLFLFYWLGVSWC*SW*TCPSYPLELACR
Gryllus_bimaculatus          NSK*KQMVFHIICC*CLHTMRLTKMKNLIFYQK*T*IILSLKCKLRHYSM*LVSSVVFGQGYSITVRSVFHESSEDFYPNESIVIPSTTRRYMLI---TSVSWPIN--TT*LVPALHICR
Gryllus_longicercus          ----------------------ELK*-CMY*K*NKI*FSSQQNLR-----MLSS--------------------WCPLN--VVGGRENF*ALDRECSKARFDLKWLATNLVKQ*FIIS
Consensus

Teleogryllus_occipitalis     ------------------------------------------------------------------------------------------------------------------------
Acheta_domesticus            AAAHSDAAYSACDEVTAG*TFARFTIAPGLTASPLRRR----------------------GGEEGVI--------------NNTFP---------LQSALILGL*T*KMARDGPLERQ
Gryllus_bimaculatus          PGDRPGTRKSR----------------GLNMERMRTRNCFRENDTFQHISGTELDCTFALGN*RSVMFEVTTIHITSHQ*ESSNTFPG-------VW*NSFI---ATEKGLKSESHNLR
Gryllus_longicercus          WNNISNVNMN*-----------------TLRMAL*N*LN---------------------GTFSSVLLE--------------TFSGTQKIINLLNVLSYILS-VQSRHVKSRPHEL-
Consensus

Teleogryllus_occipitalis     ------------------------------------------------------------------------------------------------------------------------
Acheta_domesticus            PID*ISVLRVIVRIYF----------------YQRQATDCKLAISLQTDHR*EELADDF*FHSNYINGNFLAVSRDTQ*L*KMSII--IIKLKTRKGKLRFIHKISISKKYET-----
Gryllus_bimaculatus          PHKGAKWTRIVTFQFFQHI*E*SAIGSNENAESYQEIIVNISITRFLKKV*RHICL----LF---YIMSVFFSPRWNICRISFYSMI--FLKFKV---KFLFYYRDKISRKRENMNGLRD
Gryllus_longicercus          -----------------LSAWEARCSNEH---GQLPLTPHSFPLKIQKVMYKRKLTRPVAVNSDFI-------RKYVCRFCFYQITKNCILFTI---*CLFSSLIKFSLK*TSLLSDRN
Consensus

Teleogryllus_occipitalis     ------------------------MSLSTH-----------------------------------------------------------------------------------------
Acheta_domesticus            ----RIHCPSKQRMLLLFITIFKVYLKLSTHTHTHTH-------------------------THIIFFLEEMKKYVIIFYTSQ-----------*IMFYSYVAE*K*---QGIIR----
Gryllus_bimaculatus          FAH*RGSTPAKQS-----VSSTKV-WKSSAHERKSLYVRGEDRTRKRKERRCAVPGPFVNTAPTFVWC*PEKPRK-----TTDQSSGC*NSAPVCS*IRPSYLASVK------------
Gryllus_longicercus          MFKGRIVSLFSFCMLLLYHSVTYCI*KIM*NKLMSLY---KQFSLTDGQRFCL----------LFVC*TF*PERRMLMLLYHS*VKRITKSR*F**IVLKKLLSIVNLKSDQCIISFQWI
Consensus

Teleogryllus_occipitalis     ------------------------------------------------------------------------------------------------------------------------
Acheta_domesticus            ----------EITNSYAEAMSPLELKNILFR-SFTIQGTR*YHSWERVLIIQLPKN*NSLMLVQNMLH*TVPNGH-------**CQQKLTF------------LGLRRPRHQI-CSITVS
Gryllus_bimaculatus          -------------NPATTGIQTLEPP-----------GQRLFSD-----ISLCPEPPSPVIINGSCENKTEHNAHIFIFVI*S*CKKCVYFSNS-----FANSLDPAYP*NSTKCFS*TS
Gryllus_longicercus          CLCFKENMGISVAQFRAITVNLLLVKNVIEQVSIDLASPKRFVQ--LFLLNLAPISRDAV--------PARSNKLVTLPATLTCLL*SLVFCNSAHDAKYIIKNNAKDNKQKLKC*FVIV
Consensus

Teleogryllus_occipitalis     ------------------------------------------------------------------------------------------------------------------------
Acheta_domesticus            I-------------HKIVNIQT*LLR-----------CITYSS*TLFIYFKQ-----------------------------QNTI*IMDL-GPTSR-------------LSQTPY*LS
Gryllus_bimaculatus          VP*S*LI*NDKNG-SARSSSV*TNLLK-----------TLLLRGFIFLLVLTEKI*S-FSPVS--------------------*QENLPAKD*-LPTNKNLEN-RNP*KIFGF**VTYCIK
Gryllus_longicercus          FRREKFVWNYSAGVVERIT*LQFFLNSGVSAEMMWNPIILIQREKKVMIFLKQNLLTNYSSVTTLHRQTCFEAFYSSVFNRILR*TIKYKS*NCKTNK*LKHY*KPYKI*LFTQITWST*
Consensus

Teleogryllus_occipitalis     ------------------------------------------------------------------------------------------------------------------------
Acheta_domesticus            IML---KLINSF*VLTQPIA*ADYRQTG*RISISGRVFKAAPSDKIVKGLQVGLTHVKTPETHPRMLRKGS*GCGYSISAERAFE*QVFLSFCHK*GP-DFYDFAYFLHCFKS*LKVISW
Gryllus_bimaculatus          INL*TFKLHNFRSIRYKPHKIM**MNT----KVSTNCF------KIHNGTIISIT-------------------*WN*SAH----------YQH*S--------SY*LLQISKIQKVQLA
Gryllus_longicercus          FFV---FLTRCHFVHCLPGE-AGGREGGL*AELQSLQL------S*LKGQQFSLYWKNW*EILFRNQRKFSCMDITKRTLHLCLSSS--*KYQHL*ALYKF*DFTF------GKQRTKIE
Consensus

Teleogryllus_occipitalis     ------------------------------------------------------------------------------------------------------------------------
Acheta_domesticus            HLVIIGCFVD*FVSAYL--------------------------IDFSCWS---------IFI----CGGGGVAAAVV*WLA*RTAIP-KFAG-SNPIEAVDFRGEKILRTPSFGGEVKP
Gryllus_bimaculatus          QFFTILCSIRGKN**YLR-----------------------SSIN*KCS*---------DFLRRRWCNG*RTGLLYPSWRVQTRSKPWIF*GRKNPSYAFLRRGS-----------KA
Gryllus_longicercus          HFRKF*NFKTSKKS*NRLPQRPN*FDIQPSAKYPNRPPSPGPASTGVQCTK*HRMRKTP*IMWRRRWCNG*RTGLLYPSWRVQTRSKPWIFKG-----------EKILHTPSFGGEVKP
Consensus

Teleogryllus_occipitalis     ------------------------------------------------------------------------------------------------------------------------
Acheta_domesticus            AVPC**IYLHVKDPQA--*MTPLS*IQIAGFVAPVIPSFLSW*KECNKICVVRRKSRPHG----LSLAT*NKA--IQFNSLN----YGSIFRIFR-------*NRENIHGFVEFICERSL
Gryllus_bimaculatus          GCPMFVDLLACKRSSSLNDASELNSNRL--FLAQEIPSFSAI*KERSEICVVRRKSGPDG----PSLAT*NKAYSIQFNSCS----VWTDGRTCRPTDRQQYFNKKLL*TYWKYIQCSSF
Gryllus_longicercus          AVPCS*IYLHVKDPQA--*MTPLS*IQIACFSPKKY--LLSLLDERS----VAKFAWSDGRAGLMARVSPRKIRQIQFNSIMLTMLLGKIIIFCK------FFNSGVI------VPK*SL
Consensus

Teleogryllus_occipitalis     ------------------------------------------------------------------------------------------------------------------------
Acheta_domesticus            SGKLV-----------TIFIFPLSFL*T-IKYLCK*RIFLDIF------FKPGRK-SERRPIIKVL----------H*SKYLLQL-------------------------SLP-RSNLV
Gryllus_bimaculatus          GTLCI-----------RSFIFVF*YIR---RYWMKPKTMLKQFLINYFHYAPTKKISKLYPILKLLSG---FISPQQYNGFL----------------------------------------
Gryllus_longicercus          AY*IVSNSNKYYLLLIDKNFSFIIFLARNFFRR*ARRRKLSNTF-----*YTKCFQ-*QLTLLMELIQNVWRLTSTTSGRGTLVTLYYLLLNSKHM*VVHPHMVQCHVYIVHIPSRD*YL
Consensus

Teleogryllus_occipitalis     ----ILFTI---------------------------------------------------------------------------------------------------------------
Acheta_domesticus            HYVCILFFLFQA----CFF*IL-------------------------SRKVYTEN-------EIVNF--N*ILK--CIFQNC*VI-----------NA*FKKH----ISRYLGHKNPPPSN
Gryllus_bimaculatus          ----DLFLI*EMLGFLIFFSV*-------------------------KRLV*QN*-------EY*NIY*NIT*K*RCIYALLTI------------SIFT------VSRF--HLDGMMHL
Gryllus_longicercus          KNSLIIFYIFNV----TFFPVIYF*FDILIF*P*E*ERIEF*NVGSNRFILHNFTLCARRLEFWE*YCNST----SIHALLNG*KGRIRDVGA*GKSALRRHRPRVSSRPKSFLLSLLSR
Consensus                          *

Teleogryllus_occipitalis     ------------------------------------------------------------------------------------------------------------------------
Acheta_domesticus            KRI-------------------------F*KPQ*VC*HTLQRMIKF*RRHSV-----------SWA*RNR----IPSRSLSVLVSVSEIYAPTTLVHGGFHDHGEGRLAGGPA*HPATSD
Gryllus_bimaculatus          KK-----------------------------------YVMY**ICFFYSHC----------------------------------------------TLLSLDGTH------------YDKISL
Gryllus_longicercus          KRCPFTILGSCSPLKPLGPVVTGCRT*EFPRKKLNKN*TVGNLAKYFQRAGADT*LGKVRAYFETTRSNRRNGGLPPRWMS----------------------GKQKFGGK---FS*DGD
Consensus

Teleogryllus_occipitalis     ------------------------------------------------------------------------------------------------------------------------
Acheta_domesticus            YSAIFLGRTLNITQIPSSRWNMSRHFASFSVVPLDFARSRPYEAVLAIMRLLPARPRKDAKCR--DIF---------HREDGM*VILRVLPRKIAV*RNSSGEFSRCLEHNWLRS------
Gryllus_bimaculatus          WYTFLTNRRLVYKQIIAN--------*SLRIVIVN-----------------------------D*F---------HNFVK*LVLFHFDPM*LA-------------Q**W*RKH----
Gryllus_longicercus          PY*DKNHRRARIEQRVSY--TMCYGIIVFLI*QLDLSFS------------WLSGPKNVFFCRK*DVFSLNIT*LSQYYVYYCIQTYVYEMVLS------G------SPEWLRYSGICFL
Consensus

Teleogryllus_occipitalis     ------------------------------------------------------------------------------------------------------------------------
Acheta_domesticus            --------------ISELEMAASNRFSGSFKGNAY*VPKETARARFLFLFLVQGSVVS-QIA*LRVFMFDMS*SSFTSNENPRRKKVFHVNER*M*DRWHLLIHRSTVIKLNDRQ---HV
Gryllus_bimaculatus          --------------LSDVKI---------------------RTQFILCYMLLPSVCP*QYA-----------NSLV*RE---------LNQR-------LC*H------LQK*K---KK
Gryllus_longicercus          FPLIWASYSFLLHTLASCHLSTPTPF----------------FSLFFF*FFF*SQFFSFLIVELKYYF-----RNITKQ*TPV------INGN---------FH*KNVEPFEENSV*ALL
Consensus

Teleogryllus_occipitalis     ------------------------------------------------------------------------------------------------------------------------
Acheta_domesticus            GKWTI------NSRHEHNALLPLNVSEESNVCLKGCLSRALFPSIIDPTCFRRSPYSFQRCNKDFNNQKP*IYILIIKQET*KYVQKT*H*RTMKFHATHCNHGFLYSEYGNEYQFCGHG
Gryllus_bimaculatus          *KW*P-----QNYFHKFSTLS*VD**TKATCDRVG*LS-------------AR*PWLNPACQ-------------------------------------------------------------
Gryllus_longicercus          AKWIIKLLFTENYRYFEN------------------LH*ASF--------------------------E*H*FINGIEFTN**KRTYYRRDRSFKKII----FVY*KKKNSSK*TLHH
Consensus

Teleogryllus_occipitalis     ---------------------------------------------------------------------------------VTT--------------------------------
Acheta_domesticus            VKALK*KS*FEENKG*ITEPTVFPKLNAIF*LFFLVKAAIFELGRFITIKYRYLT*SFR---------KLNFREN*FY-PVTTEL---KPSRTVFLIS*TFCLECYAPVVNCAAFPGLHG
Gryllus_bimaculatus          ---------------------LPRND*I--*F*LIK-----------------------------------------VTI-------SRKIDSV*RRECL-----------------
Gryllus_longicercus          IKYRK*RVPFEQ-----FEEHFWSRL*ENL-SRSFIK-------RMVQVSYSLISPIFRIFLKKCGNTKL*IEISRLYKDCCTQLGI*KKACTIFCKS**FCV------IN---------
Consensus

Teleogryllus_occipitalis     ------------------------------------------------------------------------------------------------------------------------
Acheta_domesticus            AWRCCNVIHKTTALLCNNKHESHFCLYFPSKGSRSV--YW*GLRSQRQAVG*IHRSNIPLNVESGVHNQFFLCLVFITKKL*NCVILQMFLS*YHS*QWFLINPVSM*IHHVSFVTFVDV
Gryllus_bimaculatus          ---------------------------------------------TERVSDG----------ESV*RREXXXXXXXXXXXXXXXXXXXXXXXXXXXXXXXXXXXXXXXXXXXXXXXXXXX
Gryllus_longicercus          ---A*NLQQQKILKAAPN----FFL*IFSTYVIRDVKKY-----GGRIFFG--YYSIISKSTKSNI--EIFLSKTVVQNYLCRKGHVQFFEKLLVLRNLRLKLTITNLLTNCSNFFKFSP
Consensus

Teleogryllus_occipitalis     ------------------------------------------------------------------------------------------------------------------------
Acheta_domesticus            HYEILEKGTHIAPTLKKASALYLNLNLEIRTNNGHPV*MILEIFT*MVVGFTLPCY---------------------------------------------------------------
Gryllus_bimaculatus          XXXXXXXXXXXXXXXXXXXXXXXXXXXXXXXXXXXXXXXXXXXXXXXXXXXXXXXXXXXXXXXXXXXXXXXXXXXXXXXXXXXXXXXXXXXXXXXXXXXXXXXXXXXXXXXXXXXXXXXX
Gryllus_longicercus          HTVRNMVLVSFSSIARFFRRLQKNFQTTKLKYFGHRKCLL**ILNK*-------------------------------------------------------------------------
Consensus

Teleogryllus_occipitalis     ------------------------------------------------------------------------------------------------------------------------
Acheta_domesticus            ------------------------------------------------------------------------------------------------------------------------
Gryllus_bimaculatus          XXXXXXXXXXXXXXXXXXXXXXXXXXXXXXXXXXXXXXXXXXXXXXXXXXXXXXXXXXXXXXXXXXXXXXXXXXXXXXXXXXXXXXXXXXXXXXXXXXXXXXXXXXXXXXXXXXXXXXXX
Gryllus_longicercus          ------------------------------------------------------------------------------------------------------------------------
Consensus

Teleogryllus_occipitalis     ------------------------------------------------------------------------------------------------------------------------
Acheta_domesticus            ------------------------------------------------------------------------------------------------------------------------
Gryllus_bimaculatus          XXXXXXXXXXXXXXXXXXXXXXXXXXXXXXXXXXXXXXXXXXXXXXXXXXXXXXXXXXXXXXXXXXXXXXXXXXXXXXXXXXXXXXXXXXXXXXXXXXXXXXXXXXXXXXXXXXXXXXXX
Gryllus_longicercus          ------------------------------------------------------------------------------------------------------------------------
Consensus

Teleogryllus_occipitalis     ----------------------------------------
Acheta_domesticus            ----------------------------------------
Gryllus_bimaculatus          XXXXXXXXXXXXXXXXXXXXXXXXXXXXXXXXXXXXXXXX
Gryllus_longicercus          ----------------------------------------
Consensus
```

**Tocc.6G0002000.1_exon2**

```
Teleogryllus_occipitalis  -----------------------------------------------------------------------------------------------------------AEGLAVAV-----
Acheta_domesticus         --------AR*YSLCELI*S*L*IFVENPPPPGCDLSNLPILNASMAT-------------------------VVSLSHHTQLRGLLIEFIYILEKFAKFS*C-----STLHLYL---RG
Gryllus_bimaculatus       -ESASKK*RNVLTVTLSTTSFTFFYFAKQHDPHLAVTELIVQSGNRLSYTHTHRXXXXXXXXXXXXKERECL*LIQFGMQYQRQN*NIFFIYPYFVFRTPQNCFVGSASLVHLI------
Gryllus_longicercus       PCIDIFRLRDFRNFDALYSLFRYCYTNEIELTCTPLSTLLIGTNNNCAFL------------------TQWVNCSKFAEKINIQNSRIQH*VPFLV-----------ASALSCIVVCYRW
Consensus

Teleogryllus_occipitalis  --------------------------------------------------GGGGAGGGGGG---------------GGGGGGG-------------------------------------
Acheta_domesticus         YNTAYSNLILTLNFF*RGDSQYTSID-----------------------------------------------------------------------LFYIAEYIQKHMLRISISY----
Gryllus_bimaculatus       --TVLSYLYI*LQFLVATQTAKRFIDSYH----IISL*SY*F----------------------------------GKSLD---QIQNENKWYF--ILFVADAFTQ*D*LK*KISYFTKN
Gryllus_longicercus       LKTLF*ELQYYLCLFARVLNRY*FIQ*S*NDACIRNKIRYNFRRNRICECSAVGAH*M**GDARISELWIGSVRRPGLTLSGSQQI*LSNDLLLVGIIFQM*I*IKP*GWLFKIN*MV--
Consensus

Teleogryllus_occipitalis  -----------------------------------------------------------------------------------------------------------GGGGG--------
Acheta_domesticus         -------SNLTF-----------------------------*IFTQLEISAMADTSGGLSKSY--------------------*RYYQY--------------------GKGLVFN----
Gryllus_bimaculatus       KRK*YCH*NANYATTVCS*FLV*YSVKDTPSQCAACFMKAVKIFTQMKA*LSHQPRGDIC*LHQ---------YPGQSTQLD*CRRYIYVDRAIARGPASQGA*IWNG*EQGIVF-----
Gryllus_longicercus       -------HFLAF----C*KLFQGLKRL*TY*MC*-------VTFCQFNPGTSNRARTSYFPLGKPDVPTSTGNFPSLRTHFP-LKYKK*CTSAS*RGPWL*IVTSYENMFAGFVSTK*PR
Consensus                                                                                                                                *

Teleogryllus_occipitalis  ------------------------------------------------------------------------------------------------------------------------
Acheta_domesticus         SISGLMFNSVFSCLLDINQRAWAMGLKNWFWV----ICNLLILFVHTG*AYRRRRKSLMMDLYIMTLFKES*VVLLKSSFT*ILISAC-----------*SFVIDVSFPLESALKLGVLI
Gryllus_bimaculatus       ---GRMIH--FSTFQALNLTA------HLHWVTDVPLCSK*PPFT---------------*LHI----SEKAAILFR-----------------------AFGKTVSLPLKKDLKVRATT
Gryllus_longicercus       TVSCLLFDVYF--LP*LNSPL----NKHLC*VTE--ICLKGELFP----CFR-----FACCYYITLL-----LIVFR---K*CEIN*CLYTNNSRLQMDSAFVCYLSVK-PSNLNVEC*C
Consensus

Teleogryllus_occipitalis  -----------------------------------------------G------------------------------------------------------------------------
Acheta_domesticus         ISLPY*LESTE*YCHLFIFVLLVGGF--VVLKLVDVPELST*IG-----LSGCSAQR----------RCLLRLRRS-------------NRRMNLRSV-------HDRARAHC--LPAQK
Gryllus_bimaculatus       CDLT-KAPSGPGS*HSNFFNIFRNKARSVQMKMRKVTKK*LSISP*RGFLRKCKGTFVYY---FTLCRCFFRLVGTSVESV-FIL*YS*NLKLNFCSIIGTKYRENAKI*TACGILPTSV
Gryllus_longicercus       CYIT--LK*KE*QNQDNFDKLF*RNY--SQL*I*NLTNVLFPFN---GYVFASKRTWELV*LNFAQ*LSTYSL*RTS*SRYPLIWPLRRDLYNSFF*I-SRQFLGTQSPRDLISLLPCLR
Consensus

Teleogryllus_occipitalis  ---GGGG---------------------------------GGGGGGAGG-------------------------------------------GGGG------------------------
Acheta_domesticus         AWRGGGGDK*HLSTP---------------VRADTRSLNLKNGARRAAGTAANRLNLCTPCNCANIFLPTTGHRLQASNLFANRSS----LRGTGGRF---------LIPQQLYK-----
Gryllus_bimaculatus       A----APQQNNL*VPQK-YGRVVHM----SVKVCT*GVRIEQGRGRKDG-AQFRARLLI------LLRPSSGVDRRNHEKLPIRAAAARILRQSAAEFVLAT*LVSKILPPPGFKP*NLP
Gryllus_longicercus       P---------*LACFNRSYSVTAHMMQST*LRITQRIIN-KNSSVNL*SFSEER-------NLYGIIRLV*LKGSHNYNFF*TRGF----LQK*CG------------TP*F*YNVRK--
Consensus

Teleogryllus_occipitalis  ------------------------------------------------------------------------------------------------------------GGGGGG------
Acheta_domesticus         WKFSSRFTRYTVT-LKNVNNNNKIENKKRKAEVHT*DFN*QKI*NANTLS**TTHVVTFYYHF*SILEAFYTHTHTHTHAYYIFPGRDEKICHNFLYITINNVLLICSRVKVTGNNKRNN
Gryllus_bimaculatus       VRGSSVISVYALSHLRR**LMGVVKTKQNTT--HTFLF------------L*YNHDVKSVYIFLIVLQ-IVSIPHIHKIPQNVFPRQ--------VYLKVN*FRTI--RMAVLG--ARVS
Gryllus_longicercus       K**SSSNRIY*LT-IRQLQHYTGKHVSRHFTPVSSTGY-*DKL-------*NINHKIVKLINN*NTIK--------NLTKYNYLPK*HGQ--HDFLCFSRDAILCIVFRGKQGG-GRAVY
Consensus                                                                                                                                   *

Teleogryllus_occipitalis  ------------------------------------------------------------------------------------------------------------------------
Acheta_domesticus         KFLCRSNEPLGTKKHSIS*FHNSRNTLISFMGTSTYYSTA*KLKFFNACPKHVTLNCPQWSLIVPTKIDLFRIKEAASSNLFDYS*HT*NRKHTNI----IVTMYYVLLVNSVYLL*TTK
Gryllus_bimaculatus       KLIC--------*KHCSSG--------------ASYFCWS*QRKY------------EALAL*VNRKIYLQRI---------DF-------QQTKIWKTGILEKYLV-------------
Gryllus_longicercus       KRSC---SPYN*AD*KDSN--------------SAYIGKTGK-KF-------------------------------------SFGI----KENSRAW---ILQKEHS-------------
Consensus

Teleogryllus_occipitalis  ------------------------------------------------------------------------------------------------------------------------
Acheta_domesticus         YNLNNGSRPNFSAISNSLLTINHAQTHQLILG----AHPTNCISRLQTDWMTHFNIRQSL*SS--AVRQNSERPTGRFDACENARNTPENAAKGQLRLRLQHKC*KSF*ITGLSELLS*V
Gryllus_bimaculatus       ----------FNK*PTALKLI-YKHSNYIILGV*DTSHTKSCSK*IQ---------R*VLIVSKFTMVQLYQ*PSG-----TNLRIISTN-----------HR-----------------
Gryllus_longicercus       ---------TFASHLLENINIFEHYTNFKIL------HLENKGPKLN-------------ILENFEILKLVKNPK------IDCLNDQTN------------------------------
Consensus

Teleogryllus_occipitalis  GGG-------------------------------------------------------------GGGGGGGGG------------GGGGGDGGGG-------------------------
Acheta_domesticus         GTGFL*FCIFSPLL*VIAKSNFLAFGYNWMLCRLIC*CIFDRFFLLEHIHLRR-----------RWCCGGG-GVVVSVTDCYTQVRGFKPDRSRGF*GRKNPSYAFLRRGSKAGCPMLVD
Gryllus_bimaculatus       ----INYCKFRKFRKFNLRS-------------------FLPYFALSEAKINNI*GLL*IKNVLRTSCGGG-GVTVSVLDCCTQVGGFKPDRSRGFF-----------KGEKI-------
Gryllus_longicercus       -------LIYSPLLSILIDR-------------------PPPDLLPPEYNAQN--DTV*EKH--HESCGGGDGVTVSVLDCCTQVGGFKPDRSRGFL-----------RAKKS-------
Consensus                                                                                             *** *             *   *   *

Teleogryllus_occipitalis  ------------------------------------------------------------------------------------------------------------------------
Acheta_domesticus         LLACKRSSSLNDASELNSNRRF---------------------RRPSNTFFP*LVKGV*QNLRGQTEEQASWPESRHVK*G--NSIQ------------FI----------------ELW
Gryllus_bimaculatus       ------------------------------------------------------------------------------------------------------------------------
Gryllus_longicercus       ---FIRLPSEGK*SRLSHVRRFTCM*KILKPK*RL*AKFKSPVSRPRNTFFLC*MKGA*RNLRGQTEERA*WPESRHVK*GKFNSIQSC*PCYLGK*LYFVSFLIVV*LYQSKV*PTKLF
Consensus

Teleogryllus_occipitalis  ---------------------------------------------------------------------------------------------------GG-------------------
Acheta_domesticus         EHISHISLK*RKYSWFRRIYL*AFIVGKAC----HNFHFPPFFSIDD---------------------KILVQMKDIS*YI--F*TRQ--KK*KKTYNQGTSLK-------*IFTSIISP
Gryllus_bimaculatus       ------------------------------------LHTPSF---------------------------------------------------------GGEVKPAVPC-S*IYLHVKDP
Gryllus_longicercus       LIVTNIIY----YS*IRTLVLLFFWLGISLGGKQEEENYPIRFNIQNVFNNNLPC*WN*YKTYGD*HRRLLVGVR*LH-YITCF*TQNICELFTRTWC-------NVMCTLSIFLAVINI
Consensus

Teleogryllus_occipitalis  ------------------------------------------------------------------------------------------------------------------------
Acheta_domesticus         QK*LSTLC-LHIIFFIPSMLLLNFIKKGLY*K*N------------S*F*LNIKVHISELLGHKCII----------*KAYFKIFRA*KSAP***ENILKTTISLLAHFAKND*-ILKKT
Gryllus_bimaculatus       QA*MTPLS*IQIACFSPKKYLLSL----LYERS------------------------------------------------------------------------VAKFAWSD-------
Gryllus_longicercus       *RIL-*LSFIYLM*HFFLLYIFNLIF*YFSLKNENE*SFETLAVTGSYYITSLSVPDV*SSGSNIVTVHPSTPY*TGKKAEYAMWALEENPP-------------FAAIAPASAPALRVS
Consensus

Teleogryllus_occipitalis  -----------------------------------------------------------------------------------GGGGGGGGG----------------------------
Acheta_domesticus         *C--FLG-------------VKKSY---------SFPFSLGTR----ERE*NLRANHAGSRR------IPRP-----------RRGASGRRTS-VTPSNLRLFCDFSRKDTQHNSNPI--
Gryllus_bimaculatus       -------------------------------------------------------------------------------------GRAGLMAR-VSPRKI--------------------
Gryllus_longicercus       FCRYFLGNVVHSPS*ARVHHLNRSAQSLPDAELKSFPERN*TKIELSEI*PNIFNGRAPIRDWAKLGLISKPLVQIEGTEGYLLGG*VGSRSSVVSLAEMEIRIETKITDGQE*SNVLVT
Consensus                                                                                                               *  *

Teleogryllus_occipitalis  ---------------------------------------------------------------GGGGGR-----------W---------------------------------------
Acheta_domesticus         ---------------------------FAMEYVTAFRIFLSRSARFRTIATV*SSACDHATSTGKTEKRREMP*HIPSRRWDVSYLESPS*KNRSIAK*LG*I*PLSGAQLAQIDFRT*D
Gryllus_bimaculatus       -----------------------------------------RHIQFNSIHVVY----------GRTDGR--------------------------------------------ADRRTGN
Gryllus_longicercus       PCVMGLLFS*FSNWIYPFLGYQGPKMYFSVENRTYF-----RSILHNFLNTTY------ITAYKRTYMR-----------W---Y*---------------------------VVLQNGS
Consensus                                                                                             *

Teleogryllus_occipitalis  ------------------------------------------------------------------------------------------------------------------------
Acheta_domesticus         GCFKPVFRFI-QGKCLLSSQRNCTCPIFIP-FLSPRQRR*PNCVTKSFYV*YELKFIYFQ*ES*TQKSISRK*TMNVRSLALVNTQINCDKVE*PTARGKVDYQFQTRTQCSLAFKRLRG
Gryllus_bimaculatus       -----------------------------------------NILTKNYY----------------------ERTGNTFSVHLL-------------AHCVYDHSF---------------
Gryllus_longicercus       DIPEFVFYFL*SGLAILS------CCILLPAAIYQRQPRFFLCFFFNFF--FNLNFSVFS-----------**S*NIILGILQNSKPQL-------SMAIFIEKMWSRLK-KIAFKRY--
Consensus

Teleogryllus_occipitalis  ------------------------------------------------------------------------------------------------------------------------
Acheta_domesticus         V*RLFEGLLISCIVPVNY*SNLFST*PIFFPKM**GLQ*PKALNLYSYY*ARNLKICTKNITLK----------NNEIS-----------RNTL*SWISILRIRK----*ISILWPRGKS
Gryllus_bimaculatus       ----------------------------LFSNTSDGI-------------G*NLKLC*NNF**TIFITHLQKKFQNYIQFSNFYQGLYPLSNTMDFWIS-F*FKKC*DFLFSSLFEKG*F
Gryllus_longicercus       -----------------WRSG**SS---FSQKTIDIL---------KTFIRRHL----NNINS*ME*S-LQIDRREHIT------GVIDLSRKLYLYIKKKRIHR--NELYIIL------
Consensus

Teleogryllus_occipitalis  ------------------------------------------------------------------------------------------------------------------------
Acheta_domesticus         SKVKILV*RKQGLDY*ANCFS-----------------*A*RYFLTFFSR*GGNI*IRSFYYNKI*ILNLKFS*TQLSRKLILPRDHRIKAITYSFPY----------FLNFLF------
Gryllus_bimaculatus       SKTKNIRTYTKISHRNKDVF--------------------MHYSPFLFSLSAGFIWMG*CI*KNM*CINEYAFFTP-----IVPYSHLMALITIRFHYGTRS*PIAD*FTNKLLPI-NH*
Gryllus_longicercus       ----NIANSAFRSNSLKNTFGVVCEKIFQDLSLREWCRSHIH*YPRFFE----------FF*KNVEIQNCR*RFLDYTK--TVVHN*VFKKRHVQFFVKVSS------FA**MLEIYNNK
Consensus

Teleogryllus_occipitalis  ---------------------------------RVAVAV---------------------------------------------------------------------------------
Acheta_domesticus         ----RMLRARR*LRCFSRFTW--------------CLALLQCHTQNHSTTV***T*VTLLFIFPFQGFEKCVLVGTA*PTSGCR------LNSPQ*YSA*CREWCT*P------------
Gryllus_bimaculatus       ESL**MISFITS*SN*SYFTLILCD*RSDSGRGNICLTL---RSERNSFCVICCCRLSVLNNMPIL*FKEN*IRGCVDTFRNKRKSENGNLRTISTNFLLSRRWTDKPKPPATG*DSCPL
Gryllus_longicercus       KFSKQLLIF------FFKFSP------------RM**GMLR-NTAAESFLGI------------IQLFRK--VQNQI*RFSSQR------L-----------------------------
Consensus

Teleogryllus_occipitalis  --------------------------------------AVGG----------------------------------------------------------GGG-----------------
Acheta_domesticus         -----VLFVSCLHN*EALK----LCDSSNVSKLISFVTVVSN*PRKYVNSSRLFCYFCGRAL*NPRERNSHSADIEKS-------------FRIVFKFKLGNQDKQR---APRLNDTRNI
Gryllus_bimaculatus       GDLG*ILHVNC---HETIKYNFNL*KSP*VVKLI-----VSD----------------------------------------------------------GESV*RRECLTERVSDGESX
Gryllus_longicercus       --LYKIIYV------EKVMYNF-----------------LKN---------Y*FCVICA*NLQ--*QTFSQTAPISLNFLHIQ*EIWC*SLFLVLLVFFEG---------CKRISKLQN*
Consensus                                                                                                                           *

Teleogryllus_occipitalis  ------------------------------------------------------------------------------------------------------------------------
Acheta_domesticus         YMNGCRVYSSML------------------------------------------------------------------------------------------------------------
Gryllus_bimaculatus       XXXXXXXXXXXXXXXXXXXXXXXXXXXXXXXXXXXXXXXXXXXXXXXXXXXXXXXXXXXXXXXXXXXXXXXXXXXXXXXXXXXXXXXXXXXXXXXXXXXXXXXXXXXXXXXXXXXXXXXX
Gryllus_longicercus       NILATENVCCSKF*IN--------------------------------------------------------------------------------------------------------
Consensus

Teleogryllus_occipitalis  ------------------------------------------------------------------------------------------------------------------------
Acheta_domesticus         ------------------------------------------------------------------------------------------------------------------------
Gryllus_bimaculatus       XXXXXXXXXXXXXXXXXXXXXXXXXXXXXXXXXXXXXXXXXXXXXXXXXXXXXXXXXXXXXXXXXXXXXXXXXXXXXXXXXXXXXXXXXXXXXXXXXXXXXXXXXXXXXXXXXXXXXXXX
Gryllus_longicercus       ------------------------------------------------------------------------------------------------------------------------
Consensus

Teleogryllus_occipitalis  ------------------------------------------------------------------------------------------------------------------------
Acheta_domesticus         ------------------------------------------------------------------------------------------------------------------------
Gryllus_bimaculatus       XXXXXXXXXXXXXXXXXXXXXXXXXXXXXXXXXXXXXXXXXXXXXXXXXXXXXXXXXXXXXXXXXXXXXXXXXXXXXXXXXXXXXXXXXXXXXXXXXXXXXXXXXXXXXXXXXXXXXXXX
Gryllus_longicercus       ------------------------------------------------------------------------------------------------------------------------
Consensus

Teleogryllus_occipitalis  ------------------------------------------------------------------------------------------
Acheta_domesticus         ------------------------------------------------------------------------------------------
Gryllus_bimaculatus       XXXXXXXXXXXXXXXXXXXXXXXXXXXXXXXXXXXXXXXXXXXXXXXXXXXXXXXXXXXXXXXXXXXXXXXXXXXXXXXXXXXXXXXXXX
Gryllus_longicercus       ------------------------------------------------------------------------------------------
Consensus
```

**Tocc.6G0002000.1_exon3**

```
Teleogryllus_occipitalis    ------------------------------------------------------------------------------------------------------------------------
Acheta_domesticus           -----------------PL--VFTL*ADLKLTMNF------CRKSSTTRVRSL*PPYSKRFNGNSRQP-----ESSYPVT--WFAY*IYLYFGKIRKVQLMFNFAFVFTGLQYSLF*FNT
Gryllus_bimaculatus         GKREQKIKKCTHRHFINDLIYILLLRETAR--PTFGCY*VNCTEWEPTKLHA----YTQRXXXXXXXXXXX*RERVFVINTIWYAI----------------PTSKLKHFLYIPVFRF*N
Gryllus_longicercus         -----------------PCIDIFRLR-DFR---NFDALYSLFRYCYTNEIELTCTPLSTLLIGTNN------------------------------------NCAFLTQWVNCSKFAEKI
Consensus

Teleogryllus_occipitalis    ------------------------------------------------------------------------------------------------------------------------
Acheta_domesticus           NTK---------LLLTRGLTVHFN---------*FILYC*VYSKTHVKDI-YFLFEF-DVLNIY-----PA*DFCNG*YKWRFE*KLLAILSIR*GFGVQFNKW---FDV*FCV*LLVGY
Gryllus_bimaculatus         SSK---------LLCRFCVTCTFNYSSQLLVHLIAVPCCDTNG*TIHRFISYY*FMI--------------------------------IL-IRKIIRSNSK*KQMVFHIICC*CL----
Gryllus_longicercus         NIQNSRIQH*VPFLVASALSC--------------IVVCYRWLKTLF*ELQYYLCLFARVLNRY*FIQ*S*NDAC-----------------IRNKIRYNFRR------NRICECSAVGA
Consensus

Teleogryllus_occipitalis    ------------------------------------------------------------------------------------------------------------------------
Acheta_domesticus           KPEGLGDGS-*ELVLGDL*FI-------NIICPYRVGI*EEEEEFDDGLVYNDIV*RELSSFIEEQFHLNFDFGMLIFCYRCEFST*ECPEIGSPHYFTALLIGVHGMILSFIYFCFIGW
Gryllus_bimaculatus         ----------HTMRLTKMKNLIFYQK*T*IILSLKCKLRHYSM*LVSSVVFGQGYSITVRSVFHES---SEDF------YPNE----------------SIVIPS--TTRRYMLITSVSW
Gryllus_longicercus         H*M**GDARISELWIGSVR---------------RPGL-------------------TLSG--SQQI*LSNDL---------------------------LLVGI--IFQM*I*IKP*GW
Consensus

Teleogryllus_occipitalis    ------------------------------------------------------------------------------------------------------------------------
Acheta_domesticus           GFRGVEVGRRARAIHLNWPV-----------GLQRTATLPTPLATK*PQDEPSLGS---------RSR---------------------------------------------QGSLPPR
Gryllus_bimaculatus         PIN------------TT*LVPALHIC--------------------RPGDRPGTRKSRGLNMERMRTRNCFRENDTFQHISGTELDCTFALGN*RSVMFEVTTIHITSHQ*ESSNTFP--
Gryllus_longicercus         LFK------------IN*MVHFLAFC*KLFQGLKRL*TY*MC*VT-FCQFNPGTSN---------RAR---------------------------------------------TSYFP--
Consensus

Teleogryllus_occipitalis    ------------------------------------------------------------------------------------------------------------------------
Acheta_domesticus           SEGVEGRRG**ITPFHSSPR*Y-SVSEPEKWRATGRWNGSQSTESLYSV*LCEYIFTNDRPP----TAS*QSLCKQIIAERNWRTIFNSTAII*---MEIF*PFHEIHSNSEKCQ****N
Gryllus_bimaculatus         --------GVW*NSFIATEKGLKSESHNLRPHKGAKWTRIVTFQFFQHI*E*SAIGSNEN------AESYQEIIVNISITRFLKKV*RHICLLFYIMSVFFSPRWNI------CR-----
Gryllus_longicercus         ------------------------LGKPDVPTSTGNFPSLRTHFPLKYKK*CTS--AS*RGPWL*IVTSYENMFAGFVSTK*PRTV---SCLLF---DVYFLP*LNSPLNKHLC*VTEIC
Consensus

Teleogryllus_occipitalis    ------------------------------------------------------------------------------------------------------------------------
Acheta_domesticus           *KQE-----KES*GSYIRFQLA---KNMKREYIVLVNNA------------------------CCYFLLP---------FLK-YT*SFLHTHTHTHTRILY------FSWKR*KNMS*FF
Gryllus_bimaculatus         ----------------ISFYSMIFLK-FKVKFLFYYRDK-----------ISRKRENMN------------------------GLRDFAH*RGSTPAK-------QSVSSTKVWKSS---
Gryllus_longicercus         LKGELFPCFRFACCYYITLLLIVFRK*CEIN*CLYTNNSRLQMDSAFVCYLSVKPSNLNVEC*CCYITLK*KE*QNQDNFDKLF*RNYSQL*I*NLTNVLFPFNGYVFASKRTWELV*LN
Consensus

Teleogryllus_occipitalis    ------------------------------------------------------------------------------------------------------------------------
Acheta_domesticus           IHHNK*CFTHM*QSESNRE**EK*QIPMQKQ*APWN*KTFYFVVSQF--------KEHADIIHG--NEYLLFNCLKIEIL*CLSKT---------------------CYTELSPMVTNSA
Gryllus_bimaculatus         -AHERKSLYVRGEDRTRKRKERRCAVP----------GPFVNTAPTFVWC*PEKPRKTTDQSSGC*NSAPVCS*IRPSYLASVKNPATTGIQTLEPPGQRLFSDISLCPEPPSPVIINGS
Gryllus_longicercus         FAQ*LSTYSL*RTS*SRYPLI----WPLRRDL----YNSFF*ISRQFL-----GTQSPRDLIS-------LLPCLR-P*LACFNRS----------------------YSVTAHMMQST*
Consensus

Teleogryllus_occipitalis    ------------------------------------------------------------------------------------------------------------------------
Acheta_domesticus           --------NKN*PF*D*GGRVIKFVRLQLAYIKS*TYKHNCYDVLRTPRKLCLFTLNNKIQFK*WI*AQLLGYLKLPTDYQSC--------------SNSSTHFRCSPNQLHKQT-----
Gryllus_bimaculatus         CE------NKT--------------------------EHNAHIFIFVI*S*C----------------------------KKC-----------VYFSNS--------------------
Gryllus_longicercus         LRITQRIINKNSSVN----------------L*SFSEERNLYGIIR------LV*LKGSHNYNFF*TR---GFLQ-----K*CGTP*F*YNVRKK**SSSNRIY*LTIRQLQHYTGKHVS
Consensus

Teleogryllus_occipitalis    ------------------------------------------------------------------------------------------------------------------------
Acheta_domesticus           -----------TDRLDDAFQYQAESLKQRRPTK**KAYRSV*R---------------------M*KRQKHTRE--------------------CCERAVEAAATA*V------------
Gryllus_bimaculatus         ----------FANSLDPAY--------P*NSTK------------------CFS*TSVP*S*LI*NDKNGSARSSSV*TNLLKTLLLRGFIFLLVLTEKI*SFSPVS*QENLPA------
Gryllus_longicercus         RHFTPVSSTGY*DKL*NINHKIVKLINN*NTIKNLTKYNYLPK*HGQHDFLCFSRDAI--LCIVFRGKQGGGRA--------------------VYKRSCSPYN*AD*KDSNSAYIGKTG
Consensus

Teleogryllus_occipitalis    -----------------------------------------------------------------------------------------------------------EFLK---------
Acheta_domesticus           ------LKELLNNRS--------F*AFVISRDRISMILHIF-SIALSHS*K*FLGIWL*LDAL*IN---------------------------LLVHI**IFPVGAYSFAAAVV------
Gryllus_bimaculatus         -KD*LPTNKNLENRNP*KIFGF**VTYCIKINL*TFKLHNFRSI----RYKPHKIM**MNTKVSTNCFKIHNGTIISIT*WN*SAHYQH*SSY*LLQISKIQKVQLAQFFTILCSIRGKN
Gryllus_longicercus         KKFSFGIKE--NSR-----------AWILQKEHSTFASHLLENINIFEHYTNFKILHLENKGPKLN---------------------------ILEN---------FEILKLVK------
Consensus

Teleogryllus_occipitalis    ------------------------------------------------------------------------------------------------------------------------
Acheta_domesticus           ----------------LRRRWCSG*RNGLLY-----------------PSSRVQT----------------------------------RSKPWILGAKKSFVRLPSEGK*SRLSHVSRF
Gryllus_bimaculatus         **YLRSSIN*KCS*DFLRRRWCNG*RTGLLY-----------------PSWRVQT----------------------------------RSKPWIF*GRKN----PSYAFLRRGS-----
Gryllus_longicercus         ----------------NPKIDCLNDQTNLIYSPLLSILIDRPPPDLLPPEYNAQNDTV*EKHHESCGGGDGVTVSVLDCCTQVGGFKPDRSRGF-LRAKKSFIRLPSEGK*SRLSHVRRF
Consensus

Teleogryllus_occipitalis    --------------------------------MKG-------------------------------------------------------------------------------------
Acheta_domesticus           TCM*KILKPK*RL*AKFKSPVSSPQ--*YLLSLVG----------ERSVTKFAWSDGRAGLMA*VSPRKIRQFNSIH*IMGAYFA---YFVEI-------------EKIFMVS*NLFVSV
Gryllus_bimaculatus         ---------------KAGCP-------MFVDLLAC----------KRS----------------SSLNDASELNS-------------------------------NRLFLA--------
Gryllus_longicercus         TCM*KILKPK*RL*AKFKSPVSRPRNTFFLC*MKGA*RNLRGQTEERA----*WPESRH--------VK*GKFNSIQSC*PCYLGK*LYFVSFLIVV*LYQSKV*PTKLFLIVTNI----
Consensus

Teleogryllus_occipitalis    ------------------------------------------------------------------------------------------------------------------------
Acheta_domesticus           HCRESLSQFSFSPFLFYRR*NTCAN------EGYFLIYF---LNQAEKVKEDL*SRYFTKVNIYFNYLSPEVT*YIMFA--------------YYFFYSKHASSEFYQERFILKMK*LIL
Gryllus_bimaculatus         ---QEIPSFSA---I*KERSEICVVRRKSGPDGPSLAT*-----------NKAYSIQFNSCSVWTDGRTCRPTDRQQYF--NKKLL*T-----YWKYIQCSSFGTLCIRSF---------
Gryllus_longicercus         --------------IYYS*IRTLV-----------LLFFWLGISLGGKQEEENYPIRFNIQNVFNNNLPC*WN*YKTYGD*HRRLLVGVR*LHYITCF*TQNICELFTRTWCNVMCTLSI
Consensus

Teleogryllus_occipitalis    ------------------------------------------------------------------------------------------------------------------------
Acheta_domesticus           IEY*SAYFRIVR-S*MHNLKSIFQDI*GIKIRPLVIREYS-ENHNKSVSTLCKE*LNFEEDIVFPGRKEIVFLPVLSRYS*A*VKFTRQPRWFTEDSTTTERGVWQEDQRDTQQPP----
Gryllus_bimaculatus         ----------------------------IFVF*YIRRYW-------------------------------------------------------------------------MKPK----
Gryllus_longicercus         FLAVINI*RIL*LSFIYLM*HFFL----LYIFNLIF*YFSLKNENE---------*SFET-LAVTGSYYITSLSVPDV*SSGSNIVTVHPSTPY*TGKKAEYAMWALE----ENPPFAAI
Consensus

Teleogryllus_occipitalis    ------------------------------------------------------------------------------------------------------------------------
Acheta_domesticus           ----------TILRFF*EGHST*LKSHLRDGICHGISHLSQS-----------------------------FR*ISHDRDRMKQCLRSCDFYRQDREK---------TRNAVTYSIAKMG
Gryllus_bimaculatus         ----------TMLKQFL------------------INYFHYA--------------------------PT-------------------------------------KKISKLYPILKL-
Gryllus_longicercus         APASAPALRVSFCRYFL-GNVVHSPS*AR------VHHLNRSAQSLPDAELKSFPERN*TKIELSEI*PNIFNGRAPIRDWAKLGLISKPLVQIEGTEGYLLGG*VGSRSSVV-SLAEM-
Consensus

Teleogryllus_occipitalis    ------------------------------------------------------------------------------------------------------------------------
Acheta_domesticus           CELS*ESFLEKSQYSEIARVNLAAVWSTIGSDRFQNLRWLL--------QTGFPVHSREMP--IEFPKKLH----------------------------VPDF-YSFS*SKAASLAKLRN
Gryllus_bimaculatus         ----LSGFISPQQY-----------------NGFLDLFLI*EMLGF---LIFFSV*KRLV*QN*EY*NIY*N------IT*K*R------------------------------------
Gryllus_longicercus         -EIRIETKITDGQ--E*SNVLVTPCVMGLLFS*FSN--WIYPFLGYQGPKMYFSVENRTY-----FRSILHNFLNTTYITAYKRTYMRWY*VVLQNGSDIPEFVFYFL*SGLAILS----
Consensus

Teleogryllus_occipitalis    ------------------------------------------------------------------------------------------------------------------------
Acheta_domesticus           *EFLCLI*AKVHLLPMRILDAK------------------------KYFT*MN-------DECKIAGTC*YTD-QL**S*MTDSTWESGLSIPDTNTMLSCL*T--SQRSL----TFV*R
Gryllus_bimaculatus         ----CIYA----LLTISIFTVS-RFHLDGMMHLKKYVMY**ICFFYSHCTLLS-----------LDGTHYDKI----------SLWYTFL----------------TNRRL---------
Gryllus_longicercus         ----CCI-----LLPAAIYQRQPRFFL---------------CFFFNFFFNLNFSVFS**S*NIILGILQNSKPQLSMAIFIEKMWSRLKKIAFKRYWRSG**SSFSQKTIDILKTFIRR
Consensus

Teleogryllus_occipitalis    ------------------------------------------------------------------------------------------------------------------------
Acheta_domesticus           AAYLVHCSRQLLIQL--------VFDVAHILSKDVIRTSITKSLKFIFLLLSKKPKNMYKKHNTKEQ*NFTQHTVI-----MDFYTQN-TEMNINSVATG*KL*SKNLSLKKTRARLLSQ
Gryllus_bimaculatus         ----------------------------------VYKQIIAN*SLRIVIV-----ND*FH--------NFVK*LVLFHFDPM*LAQ**W*RKHLSDV--------------KIRTQFIL-
Gryllus_longicercus         HLNNINS*ME*SLQIDRREHITGVIDLSRKLYLYIKKKRIHRNELYIILNIA---NSAFRSNSLKNTFGVVCEKIF-----QDLSLREWCRSHIH*YPRFFEFF*KNVEIQNCR*RFLD-
Consensus

Teleogryllus_occipitalis    -------------------------------------------------------------------------------FLGILMSFM*-------------------------------
Acheta_domesticus           LFFLSLTLFSNFFFSLRRQYLN*VVLLQ*NIDT*LEVFVNSTFAKTDFTP*PQN*SHHVQFSLFPELS---V*NVTRPSLIALL------------FP----VYMVLGVAAMSYT--KPQ
Gryllus_bimaculatus         CYMLLPSVC-------P*QYANSLV*RELNQRLC*HLQK*KKK*K----W*PQN--YFHKFSTLS*VD---**TKATCDRVG*LSA---------R*P-----------------WLNPA
Gryllus_longicercus         ---YTKTVVHN*VF--KKRHVQFFVKVSSFA**MLEIYNNKKFSK-------QLLIFFFKFS--PRM**GMLRNTAAESFLGIIQLFRKVQNQI*RFSSQRLLYKIIYVEKVMYNFLKNY
Consensus

Teleogryllus_occipitalis    ------------------------------------------------------------------------------------------------------------------------
Acheta_domesticus           HYCVIINMSHTFVYISLPRVREVCTGRDCVANVRL*A-EFTAVI---FRLMSRVVYITSSFCVLSS*LRSFKTV*FFKCF*VDIIRDSGF*LTP*VC--KFITSLLLLLWTCTMKS*RKE
Gryllus_bimaculatus         --CQL------------PRND*I*-----------------------F*LIKVTI------------SRKIDSV*RREC------------LTERVS--------------------DGE
Gryllus_longicercus         *FCVI-----------------------CA*NLQ*QTFSQTAPISLNFLHIQ*EIWC*SLFLVL--------LVFFEGCKRISKLQN*NILATENVCCSKF*IN----------------
Consensus

Teleogryllus_occipitalis    ------------------------------------------------------------------------------------------------------------------------
Acheta_domesticus           LT*RRH*KKLPHCI*I*TWKSGQTTGTPFK*YSKYLHEWLSGLLFHA-------------------------------------------------------------------------
Gryllus_bimaculatus         SV*RREXXXXXXXXXXXXXXXXXXXXXXXXXXXXXXXXXXXXXXXXXXXXXXXXXXXXXXXXXXXXXXXXXXXXXXXXXXXXXXXXXXXXXXXXXXXXXXXXXXXXXXXXXXXXXXXX
Gryllus_longicercus         ------------------------------------------------------------------------------------------------------------------------
Consensus

Teleogryllus_occipitalis    ------------------------------------------------------------------------------------------------------------------------
Acheta_domesticus           ------------------------------------------------------------------------------------------------------------------------
Gryllus_bimaculatus         XXXXXXXXXXXXXXXXXXXXXXXXXXXXXXXXXXXXXXXXXXXXXXXXXXXXXXXXXXXXXXXXXXXXXXXXXXXXXXXXXXXXXXXXXXXXXXXXXXXXXXXXXXXXXXXXXXXXXXXX
Gryllus_longicercus         ------------------------------------------------------------------------------------------------------------------------
Consensus

Teleogryllus_occipitalis    ------------------------------------------------------------------------------------------------------------------------
Acheta_domesticus           ------------------------------------------------------------------------------------------------------------------------
Gryllus_bimaculatus         XXXXXXXXXXXXXXXXXXXXXXXXXXXXXXXXXXXXXXXXXXXXXXXXXXXXXXXXXXXXXXXXXXXXXXXXXXXXXXXXXXXXXXXXXXXXXXXXXXXXXXXXXXXXXXXXXXXXXXXX
Gryllus_longicercus         ------------------------------------------------------------------------------------------------------------------------
Consensus

Teleogryllus_occipitalis    -------------------------------------------------------------------------------------------------
Acheta_domesticus           -------------------------------------------------------------------------------------------------
Gryllus_bimaculatus         XXXXXXXXXXXXXXXXXXXXXXXXXXXXXXXXXXXXXXXXXXXXXXXXXXXXXXXXXXXXXXXXXXXXXXXXXXXXXXXXXXXXXXXXXXXXXXXXX
Gryllus_longicercus         -------------------------------------------------------------------------------------------------
Consensus
```

**Tocc.6G0004470.1**

```
Teleogryllus_occipitalis        CTTCTATTCCACACCCGAACTG--------CCATCCTTACATCCGTACTTTGTCCTACATCTAGGCACATATATGCGAACAAG------CACAACTCGCCCTAACTTGCATCCCGCAAAC
Acheta_domesticus               CTTCTCTCCCCCATCCAAATCGTCACGGAACCATCCTTACGTCAGTACTTTGTCCTACTTCACCGCACATATAAGCGAACGAGCAGTCGCAAAACTCGCCCTCACTTGCATCCTGCGAGC
Gryllus_longicercus             CTTCTCATCCACACCCGAACCGTCAAGGAACCATCTTCACATCACTACTTTGTCCTACTTCACCGCACAGATATGCGAACTAGGAGTTACACAACTCGCCCGAACTTGCATCCCGCAAAC
Consensus                       *****    ** ** ** **  *        ***** * ** **  ************* **    ***** *** ****** **       ** *********  ********** ** * *

Teleogryllus_occipitalis        ATCTTCCCGTTTCCGATATGCGCCAGACCACCTCCCGCATCACCACCGCAGCCGCTGTGTCGTCACGCCGTCTGAACATCTCTCTCAGCTCACCCGGGGGCCCACCACGCTGCTCGCCAC
Acheta_domesticus               ATCTTCCCGTTTCCGATATGCACCAGAACATCTCTCGC-TCGCCACCGCAGCCACCGTGTCGCCGCGCCGTCTGAACATCTCTCCCAGCTTACTCGTGGAACGGCCAAGTTGCTCTCCTC
Gryllus_longicercus             ACCATCCCGTTTCCGATATGCGCCACAACATCTCCCGCATCACCGCCGCAGCCACCGTGTCGTCCCGCCGTCTGAACATCTCTCCCAGCCCACCCAGAGAGCGACCAAGCTGCTCGCCTC
Consensus                       * * ***************** *** * ** *** *** ** ** ******** * ****** * ****************** ****   ** *    *   *   *** * ***** ** *

Teleogryllus_occipitalis        GTCCTCTTCTCT--ACCTCCTACCGTCGCCCTCGCCCCGGCGC---GCGGCAGCAGAAGTAAAAACGTATAGGGTGACGATGCGCCGCGCGCGGAACGGTCTCATACAGCGGCGCGCCGG
Acheta_domesticus               TACCTCCCACCGTGTCCTCCCACCGTTGCTCTCACCCCGGCGC---TCGGCAGCAGAAGTAAAAACCTATAGGGTGACGATGCGCCGCACGCGGAACGGTCTCATACAGCGG-----CGG
Gryllus_longicercus             GTCCTC--------TCCTCACACCGTCGCCCTCGCCCCGGCGCGCGGCGGCAGCAGAAGTAAAAACGTATAGGGTGACGATGCGCCGCGTGCGGAACGGTCTCATACAGCGGCGCGCCGG
Consensus                         ****          ****  ***** ** *** *********    ****************** *********************  *********************      ***

Teleogryllus_occipitalis        CGGCGGCGGCGGCGGGGTCGGTGGGGGCGGCGGAGGCGGCGGGGGCGGCGGGGGCGGCGGGGGCGGCGGGGGCGGCGGGGGCGGCGGGGGCGGCGGGGGCGGCGGGGGCGGCGGGGGCGG
Acheta_domesticus               C-----------------------------------------------------------------------------------------------------------------------
Gryllus_longicercus             C-----------------------------------------------------------------------------------------------------------------------
Consensus                       *

Teleogryllus_occipitalis        CGGGGGCGGCGGGGGCGGCGGGGGCGGCGGGGGCGGCGGGGGCGGCGGGGGCGGCGGGGGCGGCGGGGGCGGCGGGGGCGGCGGGGGCGGCGGGGGCGGCGGGGGCGGCGGGGGCGGCGG
Acheta_domesticus               ------------------------------------------------GGCGGTGGGGGCGGCGCGGGCGGCGGGAGTGA---------------------------------------
Gryllus_longicercus             ------------------------------------------------GGCGGCGGGGGCGGCGGGGGCGGCGGGGGCGGCGGGGGCGGCGGGGGCGGCGGGGGCGGCGGGGGCGGCGG
Consensus                                                                       ***** ********** ********** * *

Teleogryllus_occipitalis        GGGCGGCGGGGGCGGCGGGGGCGAGCGCGCGCGCGCTATTCGTATATTTGCGGAAGTGTGTCGCGTCTGAAACAGAGCGCGTTGTCGGTGCGCCCTGATTGTTATGCAGCGGAGACGCCG
Acheta_domesticus               ----------------------GCGCGCGCGCGCGCTGTTCGTATATTTGCGGAAGTGTGTCGCGTCTGAAACAGAGCGCGTTGTCGGCGCGCCCTGATTGTTATGCAGCGGAGACGCCG
Gryllus_longicercus             GGGCGGCGGGGGCAGCGGGAGTGAGCGCGCGCGCGCTGTTCGTATATTTGCGGAAGTGTGTCGCGTCTGAAACAGAGCGCGTTGTCGGTGCGCCCTGATTGTTATGCAGCGGAGACGCCG
Consensus                                             * ************* ************************************************** *******************************

Teleogryllus_occipitalis        TGGAAATGTCGCCGTCGCGGGACGGGGCGTGGCAAGACCGGC---------------------------------------------------CGGCCGGGCCGGGGAAG---------
Acheta_domesticus               TGGAAATGTCGCCGGGGCGGAGCGAGGCGGGGC-TGGTTGGTACCGGACGGTG----------------------TGAGGTGAGGCGGGGT--GAGATGGGCCGGGGTGACACGAGGGA
Gryllus_longicercus             TGGAAATGTCGCCAGGGTGGGGAGGGGCCGGGCAGGGGAGATGCCGGGCAGTGGAGGCGCGGGGGCGAGTAGGGGCCTGGGCGGGGCGGGGCTGGAACTGGGCCGGGGGGAGGAGCGGGG
Consensus                       *************   * **   * ***  ***  *   *                                                     *********

Teleogryllus_occipitalis        ------------------------------GGCAGGGTGATGAGGGGCG------------------------GGGCGGAGA-------------------------------------
Acheta_domesticus               TGAGATGG---------------------GGCGGGGTGAGATGGGGCGTGGTGAGATGGGGCGGGGTGAGATGGGGCGGGGTGAGATGGGGCGGGGTGAGATGGCGGGGAAGGAGCGG
Gryllus_longicercus             CAGGGTGGCTGGGGCGGTTCGCGGCAAAGCGGCGGTGTGATGAGGGGGTG------GACGGGGCTGGGAACGGCGGGGCAGGGT-AGGCAGTGCGAGGCGGGGTGGAGAGGAGCAGGGCGG
Consensus                                             *** * ****    **** *                         **** * *

Teleogryllus_occipitalis        ----------------------------------------ACGAGATGGAGGGAGACTAGGCAGTGTAAAGCGAGTGGGGGGGAAGGAACGGGTCTGGCAGGAGACGGTGCGAGATGGTA
Acheta_domesticus               AGAGAGGTGGGGCGAGGCGGGGCACAGCGGGTGAAATTGGGCGGGGCGGGGAGAGACTGGGCAGGGCAGAGCGA--GGCGGCGACGGAACGCGTCTGGCCGGGAACGACGCGAGATGGTA
Gryllus_longicercus             GGCTGTATGGGGCGTAGCGGCACAGGACGGGAATGAAAAGGCAGGGCGGGGGGAGACCGGGCAGAGCACAGCGC--CGTGAGAACGGAGCGGGTCTAGCCAGGGAAGATGCGGGATGGTA
Consensus                                                         *  *  ** * *****  ***** * * ****     * *   * *** ** **** **  *   * *  *** *******

Teleogryllus_occipitalis        TGGACAGGGTTGGGCTGGGTGGTGCAAGGGAGAGAGATGGGCAGAGGAGGGGTGCGGAAGGCTGGGGAGAGTTGGGGCATTGT-------------------------------------
Acheta_domesticus               TGGAGAGGGTTGGGCAGGGCGG-GGGAGGGAGAGAGCTGGGCCGAGGCGGCGTGCAAAAGACTGTGGAGGGGTGGGATGAAGCGGGTTTGGGGCGGAGTAACGGCAAGAATAGCCGCACT
Gryllus_longicercus             TGAACTGGGCGGGGT----------GATGGAGAGAGCTGGACAGTGGAGG-GCGGGGAGCGGCGGGGTGAGGTGGGGTGAGGTGGGTGTGGGGCGGAGTAACGGCGAGAAGAACCGCTCT
Consensus                       ** *  ***  ***             * ******** *** * * ** ** * *   *     * ** * * ****      *

Teleogryllus_occipitalis        ------------------------------------------------------------------------------------------------------------------------
Acheta_domesticus               TCACGACTTTCTCTGAAACGCTA-----------CGTAAAGAGGACTGTTTTTGAATTATTTGTGTAGTTGCCGTGTGGCGTACGTGTGATGTAGAAGTGAAGGAGTGTTTATGGCACA
Gryllus_longicercus             CCACGACTTGCTTTGCGACGCTACGTTGGCCGCGTCGTAAAGGGGTCTGTGTTTGAATTATTTGTGCAGGTACCGTGTGGCGTGCGTGCGATGGAGAAGTGTACGAGTGTGTATGTCACA
Consensus

Teleogryllus_occipitalis        ----------------------------------------------------------------
Acheta_domesticus               TCTGGAATCCCAATGTGCTCTTCTACACGTGATGCCTTGAGGAAACAAATCCTTTCTCTATTTG
Gryllus_longicercus             TCTGGAATCCCAATGCGCTTT-------------------------------------------
Consensus
```

**Tocc.6G0004470.1_exon1**

```
Teleogryllus_occipitalis                 ------------------------------------------------------------------------------------------------------------------------
Acheta_domesticus                        HPPLLPPHQLRLGCRYSRVAELPP*PQLLLSSPQASLALPLGSGVSLHPPDHASCLLHFCLVPQCSQFPT--LTVGP*LLYDVTIASLIPVRMPSLNKLLLFPPLLSPIQIVT--EPSLR
Gryllus_longicercus                      ---LIP---LR-----PRSDPVPTPPL----HPGYSLYL---HGSTCYPP----------LCPLSQVFPTILLTTSPVSF--TSV*SLPWSRITT--------PTVSPLFAGTTMSPSRR
Consensus                                

Teleogryllus_occipitalis                 ------------------------------------------------------------------------------------------------------------------------
Acheta_domesticus                        QYFVLL-HRTYKRTSSRKTRPHLHP---ASIFPFPICTRT-------------SLARHRSHRVAAPSEHLSQLTRGTAKLLSSTSHRVLP------------PLL----SPRR-SAAEVK
Gryllus_longicercus                      PFLP*VPHYFSGQAIAVSTSSHPHPNRQGTIFTSLLCPTSPHRYAN*ELHNSPELASRKHHPVSDMRHNISRIT--AAATVSSRRLNISPSPPRERPSCSPRPLLTPSPSPRRAAAAEVK
Consensus                                

Teleogryllus_occipitalis                 -----MRRARNGLIQRRAGGGGGGVGGGGGGGGGGGGGGGGGGGGGGGGGGGGGGGGGGGGGGGGGGGGGGGGGGGGGGGGGGGGGGGGGGGGGGGGGGGERARAIRIFAEVCR
Acheta_domesticus                        TYRVTMRRTRNGLIQRRRRWGRRGRRE*A------------------------------------------------------------------------------RARAVRIFAEVCR
Gryllus_longicercus                      TYRVTMRRVRNGLIQRRAGGGGGGGGGGGG--------------------------------------------------------GGGGGGGGGGGGGGGGGSGSERARAVRIFAEVCR
Consensus                                     *** ********   *  *                                                                                   **** ********

Teleogryllus_occipitalis                 V*----------------------------------------------------------------------------------------------------------------------
Acheta_domesticus                        V*NRARCRRALIVMQRRRRGNVAGAERGGAGWYRTV*GEAG*DGPG*HEG*DGAG*D-----------GAW*DGA----G*DGAG*DGAG*DGRGRSGERWGEAGHSG*NWAGRGETGQ-
Gryllus_longicercus                      V*NRARCRCALIVMQRRRRGNVARVGRGRAGEM------PGSGGAGASRGLGGAGLELGRGEERGRVAGAVRGKAAV**GVDGAGNGGAG*AVRG--GVERSRAGLYG---A*RHRTGMK
Consensus                                **

Teleogryllus_occipitalis                 ------------------------------------------------------------------------------------------------------------------------
Acheta_domesticus                        -------GRARRRRNASGRERREMVWRGLG----RAGEGESWAEAACKRLWRGGMKRV-WGGV-----------TARIAALHDFL*NAT*RG----LFLNYLCSCRVAYV*CRSEGVFMA
Gryllus_longicercus                      RQGGGRPGRAQRRENGAGLAREDAGWYELGGVMERAGQ------------WRAGSGGVRWGEVGVGRSNGEKNRSPRL-ALRRYVGRVV-KGSVFELFVQVPCGVRAM------------
Consensus                                

Teleogryllus_occipitalis                 ----------------------
Acheta_domesticus                        HLESQCALLHVMP*GNKSFLYL
Gryllus_longicercus                      ---EKCTSVYVTSGIPMRF---
Consensus
```

**Tocc.6G0004530.1**

```
Teleogryllus_occipitalis    TAGGCGCCGCCTTAATTGCGGCGTGCGCGCTGCGGCGGCGCCCGGCACGGCACCGCAGCTTGCGGACGGCGGCAGGCGGAGGGCGCGCGCCGGTCGAAGC-CCCGCTCTGGCCACGTGCG
Gryllus_bimaculatus         CAA--GGCGCCGTTATTGCGGTCGGCCC-GTGCGGCGTCGCCCGGCG-GGCACCGCAGCTT----AGGCCGGCAGCGGAGGCGCGCCCG--GTCGCAAGCGCGCGCTCTGGTCACGCTCG
Gryllus_longicercus         TAGGCGCCGCCTTAATTGCGGTCGGCCCGGTGCGGCGTCGCCCGGCGCGGCACCGCAGCTT----AGGCCGGCAGGCGGAGGGCGCGCA--GGTCGAAGCACGCGCTCTGGCCACGTGCG
Consensus                    *   * **** * *******    ** *   ******* ********   ************     * * ******  *  * **** *    *     **** * ******** ****  **

Teleogryllus_occipitalis    GGGAGGGATCCGGTCGTTAGCGGCTTCA-GCTCCGGTCGATGCCGCGTGAAAGCG-AGCGGATAAATAAGAAGACTGCTGCAACTATTTTTCTGAACTTTGAACACGAAGGGG-GTACTA
Gryllus_bimaculatus         GGGAT-GAGTCGTTCGTTAGCGGTAACGCGTGTCG---GATGCCTCGTGAAAACGGGGCGGATAAATAAGAAGATTGCTGCAACTAATTTTTCGATTTTAAACAACGCAAGCG-GTATTG
Gryllus_longicercus         GGGATGGAGTCGGTCGTTAGCGGGTACA-GTGCTGGTCGATGCCTCGTGAATACG------------AGCGGATTGCTGCAAGTATTTTCCAGATCTTTAAACACGCAAGGGAGTATTG
Consensus                   ****  **  ** **********   *  *    *   ****** ******  **              **  ** ******** ** ***    **  **   *  *** * * * *** *

Teleogryllus_occipitalis    AC-GGTAAGGCAGAAACAACGTTCAATCTGGTTCTTCTATTA-CGTTGTGAA-TACTTGCGGGTTTAATGTTCAAGAAAATTAATGATGG-CTACAAAATATTGAATAGTTAAATTA----
Gryllus_bimaculatus         TTTAGTGCAGCAGAAACAACC-ACAATCACTGTTCTTTTATACCCTTGTGAATTTCTTGCGGGTTTAGTATCTACGAGTGTGAATAACCTCCACAACCAAAAAACCAGTAAAATAATCAA
Gryllus_longicercus         TT------AGCAGAAACAACCTACAATCACGTTCTTTTATTACCATTGTGAA-TTCTTGCGGGTTTAGTATCTATGAGTGTGAAAA----CTGCCACACAAAAAAAAGTTAAATAA----
Consensus                           **********    *****     *  * *  ** * ******* * *********** * *   * **    * **       *  * *    *    *   *** **** *

Teleogryllus_occipitalis    ---TTAGACAGTTTTATAAAAAGCTCGAGAAACAAAATAACTGCTTGCTACTCTTCTAAAGGTTAATTTAATTTCTAGTCTGCAGGTTAAATACGTAT-----CCCGTCGTAAAAAATGT
Gryllus_bimaculatus         AGGCAAGGTAACAGTACATATAGCTCGG-AAGCAAAATAACTAC-TTTCACGGTCGTAAAGATTAA-TTAGTTTCTATATTGT-GATTAAACACGTATATCG-TTCGGTAAGACAGCTGC
Gryllus_longicercus         ---CAAGGTAATAGTACATATAGCTCGGCAAGCAAAATAACTACTTTTTACGCTCGTAAAGATTAATTTAGTTTCTATTCAGCGGATTAAACACGTATATCGTCTTCGGGAAGACAGCTGC
Consensus                         **  *     ** * * ******  ** ********** * *    **   *  ***** **** *** ******     *   * ***** ******         **      * *  **

Teleogryllus_occipitalis    AAATCAATCCATTTTAAATGTTAATAAAGTAAA--AAATTCTCCCAACCGTTCAGTGTAATGGGGTGTGAAGTATAGCTC-------------------------------------
Gryllus_bimaculatus         CAATCAATCCATAGTATTTGGTGATAAAATCGAAGACATTGTTGCAACGTTC---GATTAAGGCAGTGACAGTACAGTTCCTTTTTTTTTTTGTTTTTTGTTTTTGTGTTTTTTTTTTTT
Gryllus_longicercus         CAATCAATCCATGGTAAATGTTGATAAAATCGA--ACCTTCTCACAACCTTTCAGTATTAAGGTGGGTGAACTACAGTTC-------------------------------------
Consensus                    ***********  **   ** * ***** *  *  *  *  ** *  ****  *        * * **        * ** ** **

Teleogryllus_occipitalis    ----------------------------------------------------------------------------------------------------------------TCGCTA
Gryllus_bimaculatus         GTTGTTTTTTTTTTGGTTTTTTTTTTTTTTTTTGTTTTTTTTTTTCTTTTTTTTTATACTATTTCCCCACTGCGACAGCTCTTTCGTTCGCCTTGATTATCTGCGTCCATCCCTTTCGCAT
Gryllus_longicercus         -------------------------------------------------------TAGCTATTGTCCACTGTGACAGCTC-TTCGTTCGCC-TGATTATCTG-GTCCATCCCTTTCCCAT
Consensus                                                                                                                                       ** *

Teleogryllus_occipitalis    CTACTTTGCTCTC------------------------------------------------------------TTTTTCGTTATTTTGTCATTTTACATCACACAT--TCTTCTTGT
Gryllus_bimaculatus         TTACTGCCTTTCCAATGTTACGTTTTGTTGGTCGGAAGTTCTATTGATCAAACGGCGTTAATTCATCTCTCTTCCTTTTTTTGCTCTTTCGATA-TTTTCATGACATATACCCTCTTTGC
Gryllus_longicercus         TTACTTACCTTTCCAATATAAGTTTCGTTGGTCGGGAAGTTCTTGATC-AACGGCGTTAATTCATCTCTCTTCC-TTTTTTGCTCTTTCGATATTTTTCATGACATAT-TCCTCTTTGC
Consensus                    ****    *  *                                                            ***** * * *** *  * *** *** *** **   **   ***

Teleogryllus_occipitalis    TCAATTAAGAATACGTTG---------------------CTAATGAGAGTGTCATATTCAGTGTTAT--------------------TATTATTATTATTATTATTATTATTATTATT
Gryllus_bimaculatus         TTATTTAAGAATACCTAG-TAGTAGTGAGTTAATAACCTACGAATAAACAGTTTATGTTCTTTATTATTATTATTTTATTATTATTATTATATAATATTATTATTATTATTATTATTATT
Gryllus_longicercus         TTATTTAAGAATACCTAGTAGTAGTAGTTAGTTAATAACGTACGAATAACAGTTTCATGTTCTTTATTAT--------------------TATTATTATTATTATTATTATTATTATTATT
Consensus                   * * ********** * *                          * *** *      * ** ***  * ****                    ******************************

Teleogryllus_occipitalis    ATT---------------------------------------------------------------------------------------------------------------------
Gryllus_bimaculatus         ATTTCTTACATTATTCACAGTATCTACTGCGGATGTCAACCACACACAGTATATGACCAGGTGTGTGGTGCTTTACCTGAAGTTCAGATGGCTTCTGACATACGCATTGTATTTCAAATT
Gryllus_longicercus         ATT---------------------------------------------------------------------------------------------------------------------
Consensus                   ***

Teleogryllus_occipitalis    --------------------ATTATTATTAT----------------------ATTTACGTGACTAAAGTCT-----TCACTCTCGT-AACTTCATTTAATTCTACAATATAAAA--G
Gryllus_bimaculatus         ATTATTTATATTATATTATTAATTATTATTATTATTATTATTATTATTATTATTTACGTTTTAAGCGAGTATCTTCTAAAATCTCTCTCGTGGACTTAATTTACGTTTACAATACAAACAAA
Gryllus_longicercus         --------------------ATTATTATTATTATTATTATTATTATTATT-TTTACGTTTTAAGTGAGTAGTCTTCTAAAATCTCTCTCGT-GACTTAATTTACGTTTACAATACAAAT-A
Consensus                                       ***********                    **** * ** **     *        ** *******  **** *****  * ******* ***

Teleogryllus_occipitalis    CTCTACGAATTTAAAATAGGCAGGTTAAAAATACCATTTTAAA--CGACACATTTTCATAGGCCGGAGTTGGCTGTCACAGAATGTCACCGCAATCGTAAATCCGGGCACGGGTACAAAT
Gryllus_bimaculatus         AATTACGAATTTAAAATAGGCAGGTTAAAATATTATTTTAAACCCGACATGC-TTCATAGGCCGAGTATGGCTGTCACACTTGTCACCCGCAATCGTAAATCCGGCCTTGGGTCCAAAT
Gryllus_longicercus         AATTACGAATTTAAAATAGGCAGGTTAAAAA---TATTTTAAA--CGACACGCTTTCATAGGCCGGAGTTGGCTGTCACAGCTTGTCACCGCAATCGTAAATCCGGCCTTGGGTCCAAAT
Consensus                      ****************************    ********  *****    ******** **    ***********         ****************** *  **** *****

Teleogryllus_occipitalis    TAAGACCAGCTTTTCATAGCCGCCGTGGCATGATTCACCTGCACCACCACA-CCTCCTCCCCCGTCCCTACCACCATCTCTACTACCGCCTCCACTCC-----ACTGCCGCCATATTATC
Gryllus_bimaculatus         T-AGAGCAGCTTTTCATAGCCTCCGTGGCATGATTCACTGCGTCTTCAACAGCCTTCTCTTCCTCCTCCATCACCTCCTCC-------CCTCCACCCCTGCTGACCGCCGTCTCACCACC
Gryllus_longicercus         TAAGAGCAGCTTTTCATAGCCGCCGTGGCATGATTCACCTGGGTCTTCACAGCCTCCTCCTCCTCCATCACCT-------------------------------------------------
Consensus                   * *** *************** *******************         *** *** ***   **   *  *  * ****

Teleogryllus_occipitalis    GCGCGCTTACAGCACCACCACCACCGTCCCAA--TCACCGCCACCATCACCACCACCGCCTCCGCTGCTGCCGCCCCAGCCGCCTCGACCACCACCATCGCCACAGTCTCCGTCATTACTA
Gryllus_bimaculatus         TCAGCTTGCCAT----CAATCATCGCCCGCGTCCCACTGCTGCCCCCACTATGACCACCAACCCCTCACCACCACACCACCACCAACCACTA-------------------------
Gryllus_longicercus         ---------------------------------CTACCGCTGCCAT-------AGCGTCGTCGCGTCCACCACCTTA---------------------------------------------
Consensus                                                   ** **   **           * *   *   * *   ** **  *

Teleogryllus_occipitalis    CTTCTACTACACCACCCCCACTGCCACCACCACCACCACTACCACCACCACTATTACCACCACCACCACCACCACCAC----------------CACCACCACCACCACCACCACC
Gryllus_bimaculatus         --------------CACCACCACCACTACCAACACCACCACTACCACCACCACTACTACCACCA--ACCACTACGACCACACCAAACTACTACACCACACTACCACCACCACCACTACC
Gryllus_longicercus         ------------------GTCGCCATCATCAACGCTGCCGTTCCCACCGCCGCT---------------------------------------------------------GCCACCACTACC
Consensus                                      ***  * ** * *   **   * ***** ** **                                                     ******* ***

Teleogryllus_occipitalis    ACCACCACCACCACCACCACTACCACTACTACGAGCACCACTACCACCACCACCA--CCACCACCACCACCACCACCACCACTACCACGAGCACCATCACCATCACCATCACCATCA
Gryllus_bimaculatus         ACCACCACCACCACAACTACCGCCACCACCACCACCACCACCACCTTCTACCCCACTCTACCACCACTACTACCA-CACCACTACTACCACCACCACTACTATCACCACCACTAACT
Gryllus_longicercus         ACCACCACCAACAACAACAACAACAACAACTCCACGAACTCCACCGCCGTCT-------------------------------------------------------------------
Consensus                   ********* **   *  * *    * **    **** * * ****  **  *

Teleogryllus_occipitalis    AAACCACCACCACCACCACCACCACCACAACCACCACCACCACCACCACCACCACCACTACTACTACCACTACCACTAGCACCAGCACCACCACCACAACAATTTAAACAATATCA
Gryllus_bimaculatus         ACCACCACCCTACCAACACCACCACCACCACCACCACCTACCCCTACTCCACCACCACCACCACCACCACCACTACCACCGTCTCCACCGCCTCCG------------------
Gryllus_longicercus         ---------CTGTAACCGTCACCTCCAGCACCATCGCCGCCGCTGCGCCACCACCACCGCCACCACCACCGCTACC-------CCCCTCCGTCAC----------------------
Consensus                            *    * *  **** *** *** * * *   ** *  *   *********** * ** ** *** * ***        **  * **  * *

Teleogryllus_occipitalis    CTCTCTTCGCTGTCGCTATTGCCACCGCCACTACCATCTTCGATGCCATCACAACTACCACCACCAATACCACTATCACCCATCGTCACCCGTTGCCGCGCCTCATTTTCGTGTTTGAG
Gryllus_bimaculatus         --CACCCCTTCATCACTACCGCCATCGCCTC-------------------------------------------------------------------------------------
Gryllus_longicercus         ----CCCCTTCATCACTAGCGCCATCGCCTC-------------------------------------------------------------------------------------
Consensus                       *   *     ** ***   **** **** *

Teleogryllus_occipitalis    CAGGGTGGAGAGTCGGTGCGGCGGGTGTTGCAGAGGTCGAAAGGCGCCAGGAATACGCGGCGAGTGGGCGGCGGGCGGCGCGCTTTGAAGCCGACCTCCCGCGAATGCACCGCCCGACCT
Gryllus_bimaculatus         ------------------------TGTTGGAGCAGCCGAAAGGCGCCAGGAATACGCGGCGAGTGGGCGGCGGGCGGCGCGCTTTGAAGTCGACATCCCGCGGATGTACTGCTCTGGCT
Gryllus_longicercus         ------------------------TGTTGGAGAGGCCGAAAGGCGCCAGGAATACGCGGCGAGTGGGCGGCGGGCGGCGCGCTTTGAAGTCGACCTCCCGCGGATGTACCGCTCTGACG
Consensus                                           ***** **  * ********************************************************** **** ******* *** ** ** *   *

Teleogryllus_occipitalis    AC--------------------------------TCCGAACGCGCTTGCTGGCA-------------------------------------------------------------------
Gryllus_bimaculatus         GCGCACGTCTTCCCATTCCTGCCCTCCACCCGTATTCTGGACCG----CGTTGGCATCCCC-GACGTTTTAATTGGTCGGTTGCAAGACAACTGTTCAGTCTGCAAGTATCAACGTTAAT
Gryllus_longicercus         GCGCACGTCTTCCCATTCCTGCCCTCCACCCGTATTCGGACCGCGCTCGTTGGCATCCCCGGACGTTTTAATTGGTCGGTTGCAAGACAACTGTTCAGCCTGCAAGTATCAACGTTAAT
Consensus                    *                                ** **      * *****

Teleogryllus_occipitalis    -------------------------------------------------------------------------------------------------------------------
Gryllus_bimaculatus         AAATAACATTACTAAAATATTTATCGTCGAATGGTGCTACTAGGGCCAAGGTATACAAAATTAAGACAAGCGTGTAATTTGCATTGCATAGGCGCCTTGCAATGCCAATCCGGCACCCGA
Gryllus_longicercus         AAATAACATAACTAAAATATTTATTGTTGAATGGTGCAACTAGGGCCAAGGTATACAAAATTAAGGCAAGCGTATCATTTGCATTGCATAGGCGCCTTGCAATGCCAATCCGGCACCCGA
Consensus

Teleogryllus_occipitalis    -------------------------------------------------------------------------------------------------------------------
Gryllus_bimaculatus         AGAACAGCAATCTGGGAAGGGCAATGAGGGGGGAGGGCTATATTCGGCGTATTTCAGAATAATATATTTGAAGGAAAATTGTTTGCTTTTAATAAATGATACCTGTATTTAGCAGTATAT
Gryllus_longicercus         AGGACAGCAATCTGGGAAGGGCATTGAGGGGGGAGGGCTATAAACTGCGTATTTCAGAATAATACATTTGAAGGACAATTGTTGTACTTAATATATGATACGTGTATTTAGCAGTATAT
Consensus

Teleogryllus_occipitalis    -------------------------------------------------------------------------------------------------------------------
Gryllus_bimaculatus         TACAATAATTTTCTGCATTTTTACTTCATAAGATTTTGAATAGCCGAAAAGATTATGATGATATCGCATTGTCCACTGTACATTTGCTGAAAATGCATTGTAACCTTAACGTAGCGTAAA
Gryllus_longicercus         TACAATAATTTTCTGCATTTTTACTTCATAAGATTTTGAATATCCGAAAATATTATAATG-CTCGCATTGTCCACTGTACATTTGC------CGCATTGTAACCTTAACGTAGCGTAAA
Consensus

Teleogryllus_occipitalis    -------------------------------------------------------------------------------------------------------------------
Gryllus_bimaculatus         CATAGAGTACACGTGCACACGAAGTGCCAGCTTTATGTGTTGACGAAGTGCCAGTTATGTGCACACGAAGTGCCAGTGTTCATGGTGTTTTCTCCCGTAATTTTTGCGGCACACCCACTT
Gryllus_longicercus         CATAGAGTGCGCGTGCACACGATGTGCCAGTTTTAC---------------------------------------GTGTTCATGGTGTTTTCTCCCGTAATTTTCGCGGCACACCCACTT
Consensus

Teleogryllus_occipitalis    -------------------------------------------------------------------------------------------------------------------
Gryllus_bimaculatus         TATTCCACGGCACACTGCTGTCCACGACACACCACTGCTGAGAGCCATTATCGACATGCTTTCCGTACTCCCACAACATAAAATTTCCATTCCAACCAGCTCTTCGTTTGAGCCGGCGCT
Gryllus_longicercus         TATTCCACGGCACACTGCTGTCCACGGCACACCACTGCTGAGAGCCATTGTCGACATGCTTTCCGTACTCTCACAATATTAAATTTCCATTCCAACCAGCTCTTCGTTTGAGCCGGCGCC
Consensus

Teleogryllus_occipitalis    -------------------------------------------------------------------------------------------------------------------
Gryllus_bimaculatus         TTTCTGCTCTATCCTGCATTTGTCTCGCGTCAAACTGCACCTTCTCCGTTCCTCCGATCCCACCTCTGGGTCCTCCCACTCAAAACCTATTCTCATGCTATCCCATTTGGCCATTTATT
Gryllus_longicercus         TTTCCGCTCTGTCCTGCATTTGTCTCGCGTCAACTCGCACCTTCTCCGCTCCACCCCATCCCACCTCTGGGTCCTCCCACTCACACCCTGCCCTCATGCTGT-CCATTTGGCCATTAATT
Consensus

Teleogryllus_occipitalis    -------------------------------------------------------------------------------------------------------------------
Gryllus_bimaculatus         CGGCACTACTCCCACTTCTTGCGACATGATCCCATACCGTAACATTTCGAGTGTTTTTTCTGGCTTTCATTTCC----------TCCTTTCCTTCCTTCCTTCCGGTGACATGTATCTCA
Gryllus_longicercus         CGG---------TACTTCTTACAACACGATCTCATATCGTACCATTTCGAGTGTTTTTTCTGGCTTTCATTCCCATTCCCTTTTTCCATTTCTTCCTCCCTTCCGGTGACATGTTTCTCA
Consensus

Teleogryllus_occipitalis    -------------------------------------------------------------------------------------------------------------------
Gryllus_bimaculatus         ACCTGACCGCGCGCGGCGATGGTGTAACGGTTAGCGTATCGGA--CTGTTTTACCAAAGTTGGCGGTTTCAAATCCAGGATGGATGAACTAGACAATTCCCAACAATTTAGACCATTTTT
Gryllus_longicercus         AAATCACCGCGCGCGGCGATGATGTAATGGTTAGCGTATCGGACCCTGTTATGCCCAAGTTGGCAGGTTTAAACCCAGGACGGATGAACTAGACAATTCCCAACAATTCAGACCATTTTT
Consensus

Teleogryllus_occipitalis    -------------------------------------------------------------------------------------------------------------------
Gryllus_bimaculatus         -GTGTACAAACACGATCAAAGCCTTGAATTGTTAAGGGCGAAAAAATTCTTCGAAAGGGAAGTAAAGCCGGCTGTCCCATGTTAGTAGATTTACTTGCATGTAAAGATCCTCAAACCTA
Gryllus_longicercus         CGTGTTCAAACTCGATCTAAGCCGTAGATTTTTAAGGGCAA------------------------------------------------------------------------------
Consensus

Teleogryllus_occipitalis    ---------------------------------------------------------------------------
Gryllus_bimaculatus         AATGACGCCTCTGAGCTAAATTCAAATCGCCTGTTTCTCGTCCATGAAATACTTGCCTTTTCTGATAGTTGGTGAAA
Gryllus_longicercus         ---------------------------------------------------------------------------
Consensus
```

**Tocc.6G0004530.1_exon1**

```
Teleogryllus_occipitalis     ------------------------------------------------------------------------------------------------------------------------
Gryllus_bimaculatus          ----------------------------TRLPLV-----SRYPPPPSWPP*AFHRL---HDPC------QTATCVLKPIAVISPPP---------------------PPRTATLAT---
Gryllus_longicercus          ATTATSVISVASITSVASVTPVTPVTLITPVALVAHVIPSSMPPPSSQPPLSKLTLIASIGPCCLLVAITTVATLASPTYLLIPPSPNKLSRFLCPDRPVYLQCHAVRPPLLATLCTPPS
Consensus

Teleogryllus_occipitalis     ------------------------------------------------------------------------------------------------------------------------
Gryllus_bimaculatus          -----------FIHSIVSTPP----------------LASTDSTRPSLFPPHPT*LNLTHAFPPHSPIPSRPPVL*RSSLRNPDRRAQG-AVIAVGPCGVARRAPQLRP--------AAE
Gryllus_longicercus          APPLPNCRMRP*AHAVHITPPTQDAPSPPHPVVSNITVASTDSTAPSLFPTSPPTNST*PTVPATFSVRTRSPVL*RSSLRSPDWACLGAALIAVGP--VRRRPARHRSLGRQAEGAQVE
Consensus

Teleogryllus_occipitalis     ------------------------------------------------------------------------------------------------------------------------
Gryllus_bimaculatus          ARPVASARSGHARG*VVR*R*RVSDAS*KRGG*IRRLLQLIFRF*TTQAVLFSAAETTTITVLLYPCEFLAGLVSTSVNNLHNQKTSKIIKGKVTVHIARKQNN-YFHGRKD*LVSIL*L
Gryllus_longicercus          ARALATCGDGVG-------R*RVQCWSMPR--EYERIAASIFQIF-KHAREYC*QKQPTITFFYYHCEFLR-V*YL*V*KLPHKKK---LNNKVIVHIARQAK*LLFTLVKINLVSIQRI
Consensus

Teleogryllus_occipitalis     ---------------------------------------------------------------------------------------MFKKI--------------------------
Gryllus_bimaculatus          -----------NTYIVR*DSCQSIHSIW**NRRHCCNVRLRQ*QYSSFFFLFFVFVFFFFVVFFLVFFFFLFFFFFFFILFPHC--DSSFVRLDYLRPSLSHLLPFQCYVL----LVGSS
Gryllus_longicercus          KHVYRFGKTAANQSMVNVDKIEPSHNLSV----------LRWVNYSS---------------------------------SYCPL*QLFVRL------IIWSIPFPFTYLSNISFVGRE
Consensus                                                                                                         *

Teleogryllus_occipitalis     ------------------------------------------------------------------------------------------------------------------------
Gryllus_bimaculatus          ID-QTALIHLSSFFCSFDIFMTYTLFAYLRIPSSSELITYE*TVYVLYYYFIIII*YYYYYYYYYYYFLHYSQYLLRMSTTHSI*PGVWCFT*SSDGF*HTHCISNYYLYYIINYYYYYY
Gryllus_longicercus          VS*STALIHLSSFFALSIFFMTYSSLLI*EYLVSS*LITYE*QFHVLYYY------YYYYYYYYYYYY---------------------------------------------YYYYYY
Consensus

Teleogryllus_occipitalis     ------------------------------------------------------------------------------------------------------------------------
Gryllus_bimaculatus          YYYLRFKRVSSKNLSRGLNLRLQYKQKLRI*NRQVKNIILNPTCFIGRVWLSHLSPAIVNPALGPN*SSFS*PPWHDSLRLQQPSLPPPSPPPLHPC*PPSHHL-SLPSIIARVPLLPPL
Gryllus_longicercus          YFYVLSE*SSKISLVT*FTFTIQINYEFKIGRLK----IF*TTRF-------H-RPELAVTACHRNRKS----GLGSKLRAAFHSRRGMIHLGLHSLLLLLHHLYRCHSVVASTTLVAII
Consensus

Teleogryllus_occipitalis     ------------------------------------------------------------------------------------------------------------------------
Gryllus_bimaculatus          *PPTPPPPHHHQPLHHHHYQHHHYHHHYYHQPLRPHQTTTPHYHHHHYHHHHHNYRHHHHHHHHLLPHSTTTTTTPLLPPPLLSPPLTTTTLPTPPPPPPLPLLPPPPPPPPPPLPPSPP
Gryllus_longicercus          NAAVPTAA-------------------------ATTTTTTNNNNNN-------------------NSTNSTA--------VSVTVTSSTIAAAAATTTATTTATPPPSPPSSLAPSPL
Consensus

Teleogryllus_occipitalis     ------------------------------------------------------------------------------------------------------------------------
Gryllus_bimaculatus          PPHPFITTAIASVGAAERRQEYAASGRRAARFEVDIPRMYCSGCARLP------IPALHPYSGPRWHPRRFNW---SVARQLFSLQVSTLIN-NITKIFIVEWCY*GQ-GIQN*DKR---
Gryllus_longicercus          LERP--------KGARNTRR--VGGGRRAL-------KSTSRGCTALTAHVFPFLPSTRIPDRARWHPRTF*LVGCKTTVQPASINVNK*HN*NI-------YC*MVQLGPRYTKLRQAY
Consensus

Teleogryllus_occipitalis     -------------------------------------------------------ND---------------------------------------------------------------
Gryllus_bimaculatus          -VICIA*APCNANPAPEEQQSG-------KGNEGGGLYSAYFRIIYLKENCLLLINDTCI*Q-YITIIFCIFTS*DFE*PKRL**YRIVHCTFAENAL--------*P*RSVNIEYTCT-
Gryllus_longicercus          HLHCIGALQC---------QSGTRRTAIWEGH*GG-------RAI----NCVFQ-NNTFEGQLFVLNI*YVYLAVYY---NNFLHFYFIRF*ISENIIMLALSTVHLPHCNLNVA*T*SA
Consensus                                                                   *

Teleogryllus_occipitalis     ------------------------------------------------------------------------------------------------------------------------
Gryllus_bimaculatus          RSASFMC*RSASYVHTKCQCSWCFLP*FLRHTHFIPRHTAVHDTPLLRA--IIDMLSV-LP-QHKISIPTSSSFEPALFCSILHLSRVKLHLLR-------------------SSRSH--
Gryllus_longicercus          RAHDVPVLR----VHG----------VFSRNFRGTP--TLFHGTLLSTAHHC*EPLSTCFPYSHNIKFP----FQPA-----LRLSR-RLSALSCICLASTCTFSAPPHPTSGSSHSHPA
Consensus

Teleogryllus_occipitalis     ------------------------------------------------------------------------------------------------------------------------
Gryllus_bimaculatus          --LWVLPLKTY--SHAIPFGHLFGTTPTSCDMIPYRNISSVFSGFHFLLSF-LPSGDMYLNLTARGDGVTVSVS---DCFTKVGGFKSRMDELDNSQQFRPFLCTNTIKALNC*GRKNSS
Gryllus_longicercus          LMLSIWPLIRYFLQH---------------DLISYH-----FECF-FWLSFPFP-------------FSISSSLPVTCFS-----KSPRAAMM*WLAYRTLLC--------------PS
Consensus

Teleogryllus_occipitalis     ------------------------------------------------
Gryllus_bimaculatus          KGK*SRLSHVSRFTCM*KILKPK*RL*AKFKSPVSRP*NTCLF**LVK
Gryllus_longicercus          WQV*TQDG*TRQFPTIQTIFRVQTR---------SKP---*IFKG---
Consensus
```

**Tocc.6G0004530.1_exon2**

```
Teleogryllus_occipitalis        ---------------------------------------------------------SP-----SLL--LL----------LHHPHCH--------------------------HHHHHY
Gryllus_bimaculatus             --QPVYL*CHAIRPPRPGHPE------------------------HSTVYTTLAKLPP-----ASLSPLL----------SYHPHLHPGP------------PPSPPSSIPSSQHHRW
Gryllus_longicercus             LLPPPPSSASPPSPPLPPSPQSPQSP*LLPSPSSPTSSLLQCLLHPHSHLCPS*HSLPPLAPAASLLLSLLLPL*PPLLTCLFHPHQTSCPVSSVQIDPSTSSVTPSAPPSWPPCAPHRL
Consensus                                                                                 *       *  **            ***                               *

Teleogryllus_occipitalis        ------------------------------------------------------------------------------------------------------------------------
Gryllus_bimaculatus             --------------------------------------PPPILHGRLSFLHIPRNLT*PTRSRHILR-----------YPADPQSSRDPRFETPTGVLKA-PLLRSARAASPGGHRSLGR
Gryllus_longicercus             RHPCQTAACALKPMLSISPLPPRTPPRPLIQSSPTSPLPPPILPRRLSF-----------RPHHPLTPPDPPFPPPSPYARGPQSSRDPRFEAPTGRA*APP*LRSARCGVARRGTAA*A
Consensus

Teleogryllus_occipitalis        ------------------------------------------------------------------------------------------------------------------------
Gryllus_bimaculatus             QRRRARSQARAL-VTLGDESFVSGNACRMPRENGADK*EDCCN*FFDFKQRKRYCLVQQKQPQSLFFYTLVNFLRV*YLRV*ITSTTKKPVK*SKAR*QYI*LGSKITTFTVVKIN*FLY
Gryllus_longicercus             GRRRARRSKHALWPRAGMESVVSGYSAGRCLVNTSGLLQ-----VFSRSLNTQGSIVSRNNLQSRSFITIVNSCGFSIYECENCHTKKS*I----TR**YI*LG---------KQNNYFL
Consensus

Teleogryllus_occipitalis        ------------------------------------------------------------------------------------------------------------------------
Gryllus_bimaculatus             CD*TRISFGKTAANQSIVFGDKI----------EDIVATFD-*GSDSTVPFFFCFLFLCFFFLLFFFWFFFFFCFFFSFFLYYFPTAT-ALSFALIIC-VHPFRIYCLSNVTFCWSEVLL
Gryllus_longicercus             RS*RLI*FLFSGLNTYIASGRQLPINPW*MLIKSNLLTTFQY*GG*TTV-------------------LAIVHCDSSSFA*LSGPSLSHLLTFPI*VSLVGKF----------------L
Consensus

Teleogryllus_occipitalis        ---------------------------------------------------------------------HHHY----------------------------------------------
Gryllus_bimaculatus             IKRR*FISLPFFALSIFS*HIPSLLI*EYLVVVS**PTNKQFMFFIIILLLLYNIIIIIIIIIIISYIIHSIYCGCQPHTVYDQVCGALPEVQMASDIRIVFQIIIYIILLIIIIIIIII
Gryllus_longicercus             DQRR*FISLPFLLFRYFS*HIPLCLFKNT*LVVS**RTNNSFMFFIII------IIIIIIIIII--------------------------------------------IIIIIIIF
Consensus

Teleogryllus_occipitalis        ----------------------------------------------YHHHHHH--------------------------------HH----------------------H---HHHHH
Gryllus_bimaculatus             IYVLSEYLLKISLVDLIYVYNTNKNYEFKIGRLKILF*TRHAS*VEYGCHTCHPQS*IRPWVQIRAAFHSLRGMIHCVFNSLLFLLHHLLPSTPADRRLTTSACHQSSPASHCCPHYDHQ
Gryllus_longicercus             TF*VSSLLKSLS*LNLRLQYK*ITNLK--------------*AG*KYFKRHAF---IGRSW---------LSQLVTAIVNPAL---------GPN*EQLFIAAV----------------
Consensus                                                                     *

Teleogryllus_occipitalis        HHHHHHHHYHY--------------HEHHYHHHHHHHHHH----------------------------------------------HHHYHEHHHHHHH----HQNHHHHHHHHHNHHHHH
Gryllus_bimaculatus             PLHHHTTTNHYTTTTTNTTTTTTTTTTNHYDHTKLLHHTT--------------TTTTTTTTTTTATTTTTTTTFYPTLPPLLPHHYYHHHYYHHH*LPPPYQHHHHHHHHYPYSHHHHH
Gryllus_longicercus             ------A*FTWVFTASSSSSITSTAAIASSRPPP*SPSSTLPFPPPLPPLPPPPTTTTTTTPRTPPPSL*PSPPAPSPPLPPPPPP------------PPPLP--------PLRHPLHH
Consensus                                                                                                                          *  **

Teleogryllus_occipitalis        HHHHYYYHYH*-------------------------------------------------------------------------------------------------------------
Gryllus_bimaculatus             HHHHHYHRLHRLRTPSSLPPSPLLEQPKGARNTRRVGGGRRALKSTSRGCTAL----AAHVFPFLPSTRILDRVGIPDVLIGRLQDNCSVCK--YQR**ITLLKYLSSNGATRAKVYKI-
Gryllus_longicercus             *RHRLCWRGRK---------AP------GIRGEWAAGG---AL*SRPPADVPL*RRTSSHSCP--PPVFRTALVGIP----GRF--NWSVARQLFSLQVSTLINNIT-------KIFIVE
Consensus                          *

Teleogryllus_occipitalis        ------------------------------------------------------------------------------------------------------------------------
Gryllus_bimaculatus             ------------KTSV*FALHRRLAMPIRHPKNSNLGRAMRGEG----YIRRISE*YI*RKIVCF**MIPVFSSILQ*FSAFLLHKILNSRKDYDDIALSTVHLLKMHCNLNVA--*T*S
Gryllus_longicercus             WCN*GQGIQN*GKRIICIA*APCNANPA--PEGQQSGKGIEGGGL*TAYFR-----IIHLKDNCLYL---IYDTCI*QYITIIFCIFTS*DFEYPKI------L*CSHCPLYICRIVTLT
Consensus

Teleogryllus_occipitalis        ------------------------------------------------------------------------------------------------------------------------
Gryllus_bimaculatus             TRAHEVPALCVDEVPVMCTRSASVHGVFSRNFCGTPTLFHGTLLSTTHHC*EPLSTCFPYSHNIKFPFQPALRLSRRFSALSCICLASNCTF----SVPPDPTSGSSHSKPILMLSHLAI
Gryllus_longicercus             *RKHRV------RVHTMC--QFYVFMVFS------PVIFAAHPLYSTAHCCPRHTTA--ESH-----------CRHAFRTLTILNFHSNQLFV*AGAFPLCPAFVSRQPAP---------
Consensus

Teleogryllus_occipitalis        ------------------------------------------------------------------------------------------------------------------------
Gryllus_bimaculatus             YSALLPL--LAT*SHTVTFRVFFLAFISSFPSFLPVTCIST*PRAAMV*RLAYRTVLPKLAVSNPGWMN*TIPNNLDHFCVQTRSKP*IVKGEKILRKGSKAGCPMLVDLLACK-RSSNL
Gryllus_longicercus             -SPLHPIPPLGPPTHT----------------LPSCCPFGH*FGTSYNTISYRTISSVFSGFHSHSLFPFLPPFR*HVSQNHRARR*C--------NG*RIG-PCYAQVGRFKPRTDEL
Consensus

Teleogryllus_occipitalis        ---------------------------
Gryllus_bimaculatus             NDASELNSNRLFLVHEILAFSDSW*--
Gryllus_longicercus             DNSQQF---RPFFVFK-LDLSRRFLRA
Consensus
```

**Tocc.6G0011210.1**

```
Teleogryllus_occipitalis      GGTGGATT-------------------------------------------------------------GTCTTTTCTGAACATCAATTCCAGGGGAATGCT-AAGGAATGCCGT
Acheta_domesticus             GCTGAGTTGGGCATCTGGATATCAATGCCTGGAATGAGAATTCTAAAGGCCACTTTCTCTGATCGGGTATTCACGTCTATTGTGAAGATCATTTCCATGGGAATGCTGAAGGAATACCGT
Consensus                     * **  **                                                             **** ** **** **** ***** ********* ******* ****

Teleogryllus_occipitalis      AAAAATAG---GCTATCCTTATCCTATTTAACTCTC-TTTTTAAATGTTGCTCCACTATGGAAATAATACGTCCATTACAAATGCACAACAAATACAAGCCCAGTTATACGTTATACGGG
Acheta_domesticus             AAAAATAGGCAGCCCTTCTTATCCTATTTTACTCCCATCTTTTAAGTGTTGTGCAACTACGGAAACAATA-GATCGTGACAAATTCACAAC----ACAAGCTCAGTTATACGTTA------
Consensus                     ********   **  * *********** **** * * ***** *****  * **** ***** **** *  * * ****** ******    ****** ************

Teleogryllus_occipitalis      GCTCTTTACAAATGATGGAGATGATATCACTCGCCCACAGT-GGGCGACCTACGGTCACGTGATTAGGGCTCTCTCCCTCCACCTCTGGGTAGCGTGTTCAAATCCCGGTAAAGCTGGTG
Acheta_domesticus             ------TACAAATAATGGAGATGATATCACTCGCCCATATTGGGGCGACCCACGGTGAAGTGAACAGGACGATTACTCTCCACTTCTAGGCAGCGGGTTCAGATCTCGGTAACGCTGGTG
Consensus                           ******* ******************* * * ******* ***** * ****  *** *  *  * ****** *** ** **** ***** *** ****** *******

Teleogryllus_occipitalis      AAATTTGTGACTA----TGC-GGTTCGCGGTAGGTTTCTCGAGCTTCTCCCGGTTTACCGACCACTCATTCCGTATTCGTTCCGTCATTATACAGCACCTACATCATC-----------
Acheta_domesticus             GAATTTGTGATTGACACTGCGGGATTGCGGCAGGTTTCTCGACTTTCTCCCATTTTACCGGCCACTCATTACACTATGGTTACGTCATTATATCGCAACTACATCATCCCTGGCGGCTTC
Consensus                      ********* *     *** ** * **** ***********  *******  ******* ******* ********* *    * *** **********  *** **********

Teleogryllus_occipitalis      -----------------------------------------------------------------------------------------------------------------------
Acheta_domesticus             GCTCATTGGTCAGCGTGCTCGCTTTCCGTGACGCGACCCCGGGTTCGATTCCCGGTCGGGGTAAAAGTATAGTGTGCGTTTTACTGGCAATGTCCCAGTATATGTTGCTGTGTGACGATG
Consensus

Teleogryllus_occipitalis      -----------------------------------------------------------------------------------------------------------------------
Acheta_domesticus             AGAAAATTCACTTCTCAGGTTGAGAAGATCAGATTAAGGAGTGGGATGTCTTCGTGTAAACCACACGAACCCCCTTCACCCCCAAGGCCGCGAAGCCGGGACTACGCTGGTAGGCCCAAG
Consensus

Teleogryllus_occipitalis      ----------------------------------------ATTATCAGCACCGAATATCAATACC----ATCTTAGGCATGACGAAAAGTATAGATACTCAGGATCATGAAGATGAA
Acheta_domesticus             CCCGCACACAGGCTGTTGCGCCAGGTACGTACATACATCATTATCATCATCACCTGTCTATCAAAACCATTTATCTCACGTATGTCGAAAAGAATAGGTACTCAGGATCATTAAGATAAA
Consensus                                                             ** **** **** *  ****** ***    **** * * *** ******* **** ************* ***** **

Teleogryllus_occipitalis      TACACTTATCTTTT--------------------------------------------------------------------------------------ATCTCCTTTAA---------
Acheta_domesticus             TACACTTCTCTCTTTGCCCCAGACATACGAGGTCTGTCTAAAAAGTATCCGACCTTTGGCCAGAAAAAATATTTCGAATACTTGACAGATTTGGGACCCTAATCCCCTTCAAATAGGCCCC
Consensus                     ******* *** **                                                                                      *** **** **

Teleogryllus_occipitalis      -----------------------------------------------------------------------------------------------------------------------
Acheta_domesticus             TTGTGCTTCCACACACTTAGCCCACCGATCCTGCCACTGCTGGAAACACCTTTGGAAGTCTTCTTTTGGAATGGTGTTCAGCTCCGTCGTCGCGTTCCGCATTATTTCTTCTTTACTCTC
Consensus

Teleogryllus_occipitalis      -----------------------------------------------------------------------------------------------------------------------
Acheta_domesticus             AAAACGGGATCCTTTCAGTGACATCTTCAATTTTGGAAACAACCAGAGAAGCTAAGAAAGTGTGGATCAATTGGGATGAATGTGCAGGGGCGTTGTCATGATGCAAATGCCAGTTTTTCG
Consensus

Teleogryllus_occipitalis      -----------------------------------------------------------------------------------------------------------------------
Acheta_domesticus             CCGTCCACATGTCTGGTCTTTTGCGCCGAATTGCGTCACGGAGTCTCCGGAGAACATCTCCCGCACTGTCAATCGGCGATCTGCCATCACCAAATTTCGCACCCTTTCAACAACAGCTGC
Consensus

Teleogryllus_occipitalis      -----------------------------------------------------------------------------------------------------------------------
Acheta_domesticus             ATTGCGAGCAGTTTGGGGCCTGGCGCCACTCTTCTCACTCTCGGCTGATGTTCGGCCATCCTTGAATCGGTTGAACCACTGCTTAATTTGTGTTACACCATTGCATCTTCTCCAAACAC
Consensus

Teleogryllus_occipitalis      -----------------------------------------------------------------------------------------------------------------------
Acheta_domesticus             CTGCTGAATCTTACGAATTGTTTGGCTTTGAGAATCACCTAGCTTTTGACAGAATTTGATGCAGTATCTTTGCTCAATTCGTTCAGTCATCTTGCGAGAAAACTAAATCCGACGAACACT
Consensus

Teleogryllus_occipitalis      -----------------------------------------------------------------------------------------------------------------------
Acheta_domesticus             ATGAACAGCACCTTACTTAGCGACCACCAGCCAGCGACTGGCACCAGCGAACGCGGTGAAAAAATTCAAGCATGCGCATTAAGGATCCTCCTTCCATCGTGCACAGTGGCGCCGCCTTAG
Consensus

Teleogryllus_occipitalis      ---------------------------------------------------TCATACATTATTTATTTGAGATGGTTTAGTTAATACTATAATATTTTGATAACTTTGTATGAGATGTA
Acheta_domesticus             AATCACTACTTTGCGCGGAAAAAATTAAGGTCGGATACTTTTTAGACAGGCCTCGTACATTATCCATTTGAGTAGGTTTAGTTAATAC-----TTCTTTGATA------------TTGAA
Consensus                                                                      ** ********  *******  **************     *  *******              ** *

Teleogryllus_occipitalis      TAATTGTTAACACAATATTTATTTTGTACGTGTTTTGTGAATCTACGAGAGTTTTGACTTGTGTGTGTAAAAATCATGGTTGCTAGAGGGGAGGAAGAGAGGTAAAATTTGATTATAGG
Acheta_domesticus             TAATTATCAGTTCAATATTTATTTTGTACCTATTTTGTTAATCTAGGA-------GACTTATATAAATAAAAATCATGGTTGTTAGAGGTGAGGAGAAGAGGTGGAA--TGTTTAGGAG
Consensus                     ***** * *   ***************** * ****** ****** **       ***** * *   ***************** ****** ****** ******  ** ***   *

Teleogryllus_occipitalis      TCTTGTATTCGTTGAGCGGGGAGTAGTTCACCGTTATGGAGAGGCACAGATACTTTTCAGAATAACATAAGAAATGTTAATACATATGTACCGTGTGTACTATTTGGGTGCTAGTGCAAG
Acheta_domesticus             TCCTGAATACTCTGAGAGGAGAGTCATTCACCGTAACGTGAAAATACAGAAACTTTCCCGAATAACTTGAGTAATGTTATTAGATGTATGTCTTGTATACTATTTGGACGCTAGTGCCAG
Consensus                     ** ** ** *  **** ** ****  ******** * *   *   ***** ***** * ******* * ** ******* ** ** *  * **  * ** **********  ******** **

Teleogryllus_occipitalis      CATACTTTGAGCTCGAAAGTTATTAGTTAATGAGAGACAGTAAGTTGG-TACAGAATTCCTTTGAATAAGTTGGATAATGTCCTTCAGTAG------AACTGTTTATAAATAAAAAATGG
Acheta_domesticus             -AAAGTTTGAGCTTGAAATTTAATAGTTAAAGTGAGATAGTAAGTTGGACTACAGAATTCCTTTGAAAAAGTTGGATTTTGCCCTGAGGTAGCACGCAAGCTGCTTATAAAT-AAAAATGG
Consensus                      * * ******** **** *** ******* * **** **********  *************** *********  ** ***    ****      * *** ******** ********

Teleogryllus_occipitalis      TTCTGCT-GGTTGAACGAACTACGTTGAGTACATTACGGAAGAAAGACGACCTTCACGCTACTAATTTTTTTTTGGCTGCGGCAAGACACACGAGCATTTATTTTCTTATTGGTGAAAGT
Acheta_domesticus             TTCTGTTGGGTTGAACGAACTACGTTGAGTACATTACTGAAGTAAGATGACCTTCACGCTACCA----TTTTTTGGCAGAGGCAACACACACGAACATATATTTTTCTTATTGGTGAAAGT
Consensus                     ***** * ******************************** **** ** ********** ** *    ********* * ***** ******* *** ********* ********************

Teleogryllus_occipitalis      AACTATCGATTTTCTGAAACATGTCGGAAAGTGCCGAGTTGACTTGACTTTGTTTGGAGACCTAACACTTCACACCTCCCTAAT-ATTCCCCATCGCTCTTGATCCAATATGATCATCGC
Acheta_domesticus             AACAGTCAATTTTCTGAAACATGACGGAAATGTGCTGAGTTGACTTGAAGTTGTTTGGAGAC--ATCACTATCACACCTCCCTAATAATTCCCCATTGGCGCATGATCCATCATGATAATCGC
Consensus                     ***  ** *************** ***** **** **** ************  ************  * ****  ************ * ******** ** * *******  ***** ****

Teleogryllus_occipitalis      CCATCGAATAATGTGCAAAGCTGGCTG--ACACTACAGCAGCACACAAAGAAAGACAGAAAAAAGATAAATAAAGAAAAGGAAACAAAGGAAAGAAAGAGAAAGAAAAAAGAGAAAGAG
Acheta_domesticus             CCATCGAATGATGTACAAAGCTGGCTAACACACTATAGCAG----------------------------------------------------------------------------
Consensus                     ********* **** ***********   ****** *****

Teleogryllus_occipitalis      AAAGAAAGAAAGAAAAAAGGGAAAAAAAGAAAGAAAGAAAAAAGGAAAGAGAAAAGAAAAAGAAAAAAGAGAAAGAAAAGAGAAAGAAAAAGAGAAAGAGAAAGAAAGAAAGAAAAAAG
Acheta_domesticus             -----------------------------------------------------------------------------------------------------------------------
Consensus

Teleogryllus_occipitalis      AGAAAGAAAGAAAGAGAAAGAAAGAAAGAAAGAAAAAAGAGCAAAAGAAAGAAAGAAAAAGAAAGAGAAAGAGGGAGAAAGAGAAAGAAAGAAGAAAGAAAGAAAGAGAAAGAAAGAAAG
Acheta_domesticus             -----------------------------------------------------------------------------------------------------------------------
Consensus

Teleogryllus_occipitalis      AGAAAGAAAGAAAGAAAAGAGAAAGAAAGAAAGAGAAAGAAAGAGAGAGAAAGAAAGAAAGAGAAAGAAAGAGAAAGAAAGAGAAAGAAAGAAAGAAAGAGAAAGAAAGAGAAAGAAAGA
Acheta_domesticus             -----------------------------------------------------------------------------------------------------------------------
Consensus

Teleogryllus_occipitalis      AAGAAAGAGAAAGAAAGAGAAAGAAAGAAAGAAAGAGAAAGAAAGAAAGAAAGAAAGAAAGAGAAAGAAAGAAAGAAAGAAAGAAAGAAAGAGAAAGAAAGAGAAAGAAAGAGAAAGAAA
Acheta_domesticus             -----------------------------------------------------------------------------------------------------------------------
Consensus

Teleogryllus_occipitalis      GAAAGAGAAAGAAAGAGAAAGAAAGAGAAAGAAAGAAAGAAAGAAAAAAAAGAGAAAAAAAGAGAAAAAGAGAAAGAAAGAAAGAAAGAAAGAAAGAAAGAGAAAGAAAGAAAGAAAGAGAA
Acheta_domesticus             -----------------------------------------------------AAACAAAGAAAGAAAGAAAGAAAGAAAGAAAGAAAGAAAGAA--------------------------
Consensus                                                                          *** * ***** *  ****************************

Teleogryllus_occipitalis      AGAAAGAGAAAGAAAGAAAGAAAGAGAAAGAAAGAAAGAGAAAGAAAGAAAGAGAAAGAAAGAGAAAGAAAGAAAGAGACAGAACTTATACTTTTAGGCTAATTAATTCTCTATC
Acheta_domesticus             --------------------------------------------------------------------------------AGAAATTCGTAGTTTCAAGCTAATTAATGCTCTCCT
Consensus                                                                                                      ***** *   ** *** * ********* ****

Teleogryllus_occipitalis      ACGAAATTTGGGCTCTCGTGGTTTTCTCAAAAGTCAATGTTAAACCAGTT-------------------CACTCGAGCTTAAAAG-TATGAGCCATTTTTC-TTGTTTTCTGCTGCAGTG
Acheta_domesticus             A-AAATTTTAGGTTCTTGCAATTTTCTCAAAAATCAATGTTCAGCCAGATTTAAAATAAAGCAAGTACGCACTCTAACATAAAAATTATGAGCCGTCTTTCTTTGTGTTCTGCTGTAGTG
Consensus                     *  ***** ** *** *   ************ ******** * **** *                   *****  * * *****  ******** * ****  ***** ********* ****

Teleogryllus_occipitalis      TATCACCTAGCTTTAGAGACATCCCCGAACGCGTCTGGTGTGAGCACGCGGAAATCAGGCAAACGCGCGAACCGGGATG-TAGTGACATCGAAGAGAGTCGCCC-TGACACGCCATCACT
Acheta_domesticus             TGTCAGCTAGCTTTATAGA-ATCTCCGAATGCGTTTGGCGTGAGCATGCGGAAATCAG--------GCCAACCGGGTTGTTAGTGACATGGAAGAGAGTCGCCCTTGACACGTCATCACT
Consensus                     * *** ********* ***  *** ***** **** *** ******* ********** *         ** ******* ** *********  ******** ************** ******* *******

Teleogryllus_occipitalis      TCCAGGCGAAAGCCTTTTTCAGCACCTGAGAACACATTTCACCCCGCCCACATCTCTATACGTTGCAAAA---------------------------------------------------
Acheta_domesticus             TCCAGGCGAAAGCCGTTTTCAGCACCTGAGAACACATTTCACCCCGTCCACATCTCCATACCTTGCAAAACACAGCCGTAAGACATTAATGAGGGGCAGAGCCACACGGACGGTACGCT
Consensus                     ************** ****************************** ********* ****  ********

Teleogryllus_occipitalis      -----------------------------------------------------------------------------------------------------------------------
Acheta_domesticus             GATGCGACAAGGTATTAGATTCGCTGCCATATTTGTCGATGCGAGAGGCCAGCGTGGAGATGTTTACTACGTCTTCACGTCGAAGGTGAAAGAGCCTTTTAAAAAACGTGTGTGCGGTGT
Consensus

Teleogryllus_occipitalis      -----------------------------------------------------------------------------------------------------------------------
Acheta_domesticus             CATGTGACAAACTCCAGGGGTGTCAATTTTACTCCAGTAATTCCGGACATTTTTTCCGCATAGAAAATTCAAGTCTGCCTCATGGGGTGGTTACTTTGTTTTTAGTTTCTCAAAATACTT
Consensus

Teleogryllus_occipitalis      -----------------------------------------------------------------------------------------------------------------------
Acheta_domesticus             GTTAGGTCTTGAAATATGTCGTATATTTAACGACTGGATTTCTACGATGTTCGTGATGGTGGAATGATTTCATTATACATAGCCAGTTATAAAACGTCAAACACTGAACGCTTTGATGCG
Consensus

Teleogryllus_occipitalis      -----------------------------------------------------------------------------------------------------------------------
Acheta_domesticus             TCCGTTGGATTTTTAATATTTTTACTATTTCAGATTTAAATAATGAGTAATATTGTAACAGTTAATATTTAATATATAATTTATCATAAACATTTAAATATTTTTAGGCGAAATAATTGT
Consensus

Teleogryllus_occipitalis      -----------------------------------------------------------------------------------------------------------------------
Acheta_domesticus             TTTTATTTAATATACAGTTCCAATATTTTCCATGTTAATCTTATGTTTAGAATAGTGCATTACGAAAGAAAGACAGCAAAATACAAATAATAGAAAAAACACATCGCTTAGTATGATAA
Consensus

Teleogryllus_occipitalis      -----------------------------------------------------------------------------------------------------------------------
Acheta_domesticus             AGAACACTACATAAGATTTGCTAAGGTACCATAAAATGTGAGGTTCAAAAATATCGGTTAATCCAACTGAGATGTGATATCAACTAAAATATCATTCACGAGATAAACGGCCTATGATAA
Consensus

Teleogryllus_occipitalis      -----------------------------------------------------------------------------------------------------------------------
Acheta_domesticus             TGGAGCCATGATAATGGAATTATCAATATTTACTTCGATTATCGATAAATCAGTTCCACAGTTAAGAGTGACGAGACAACAAAAATATTGTCAGACTTCCGTAACATTGCATCTTCTTCA
Consensus

Teleogryllus_occipitalis      -----------------------------------------------------------------------------------------------------------------------
Acheta_domesticus             GCACTTCCTTCAGTCGTTAGGTTTTCAGGCAGTCCAACAGAAACCTGTCATAATTCAAAGAGGACTGTTCCCGGCACATCAAATAAACGATCCGAGAAAAAAAATCGAAATGATGCATCC
Consensus

Teleogryllus_occipitalis      -----------------------------------------------------------------------------------------------------------------------
Acheta_domesticus             TCTATATAAACATTTCAGGGGAACTGGTCGGCGAGTGGCCGAACCGGCGCGGCGCGGCGCGACGCGGCTGGGGCGTTCCAGGGAGGGCGCTCACCCACGGCCCGAATCGCTCTGCCCGCC
Consensus

Teleogryllus_occipitalis      -----------------------------------------------------------------------------------------------------------------------
Acheta_domesticus             GCGCCGCGCCGGGGCCACTCACGGGGAGCGGAACCACAGAAACCACCGCGCGGAGACATTCGGTTGGCGAGGGGGTTATGAGCTCATAGGGCAAAGGTGAATTTAAGCCAGTGGCGGTTC
Consensus

Teleogryllus_occipitalis      -----------------------------------------------------------------------------------------------------------------------
Acheta_domesticus             GTGGGACATTTTGTTGGCGGTTGGATACAACTGACGTAGTCAAACAAATTCCAGCAGTTAATGCAAACTTAACCCCAACATAAACAAATCGATTAACGCGGCCTGCAGCAGCAAAATTC
Consensus

Teleogryllus_occipitalis      -----------------------------------------------------------------------------------------------------------------------
Acheta_domesticus             TGCTGTACAAAATGATTTTCCAAAAGTAGCAAGCACGCTGCTAAATAACAACAAATAAAATACTAAAGTTACAATAAAATACTAACCTACTATATACTATAAATACTATACACGAAAA
Consensus

Teleogryllus_occipitalis      -----------------------------------------------------------------------------------------------------------------------
Acheta_domesticus             TATTATAATGACTACTTAAATTAAATAGTTAAAAGAAACAGACTTAAGAAATACTGTTAAAATATAAATGGGTGAAAACCAGAACGGTATAAGTGACATTTTATAAATTGTGAGAAAAT
Consensus

Teleogryllus_occipitalis      --------------------------------
Acheta_domesticus             GTGTTGAGTTTAGGCTATATTCAGTTGCAATG
Consensus
```

**Tocc.6G0011210.1_exon1**

```
Teleogryllus_occipitalis    ------------------------------------------------------------------------------------------------------------------------
Acheta_domesticus           H*EQLLLLLYVSTPDTKYYAGASSFNYLSIFFHIFVPINSSPAQPIKYQNFNNLKY*KSEV*NTYS*VGHLDINAWNENSKGHFL*SGIHVYCEDHFHGNAEGIP*K*AALLILFYSHL*
Consensus

Teleogryllus_occipitalis    ------------------------------------------------------------------------------------------------------------------------
Acheta_domesticus           VLCNYGNNRS*QIHNTSSVIRYTNNGDDITRPYWGDPR*SEQDDYSPLLGSGFRSR*RWWNL*LTLRDCGRFLDFLPFYRPLITLWLRHYIATTSSLAASLIGQRARFP*RDPGFDSRSG
Consensus

Teleogryllus_occipitalis    ------------------------------------------------------------------------------------------------------------------------
Acheta_domesticus           *KYSVRFTGNVPVYVAV*R*ENSLLRLRRSD*GVGCLRVNHTNPLHPQGREAGTTLVGPSPHTGCCARYVHTSLSSSPVYQNHLSHVCRKE*VLRIIKINTLLSCPRHTRSV*KVSDLWP
Consensus

Teleogryllus_occipitalis    ------------------------------------------------------------------------------------------------------------------------
Acheta_domesticus           EKIFRILDRFGTLIPFK*APCASTHLAHRSCHCWKHLWKSSFGMVFSSVVAFRIISSLLSKRDPFSDIFNFGNNQRS*ESVDQLG*MCRGVVMMQMPVFRRPHVWSFAPNCVTESPENIS
Consensus

Teleogryllus_occipitalis    ------------------------------------------------------------------------------------------------------------------------
Acheta_domesticus           RTVNRRSAITKFRTLSTTAALRAVWGLAPLFSLSADVRPSLNRLNHCLICVTPIASSPNTC*ILRIVWL*ESPSF*QNLMQYLCSIRSVILREN*IRRTL*TAPYLATTSQRLAPANAVK
Consensus

Teleogryllus_occipitalis    ------------------------------------------------------------------------------------------------------------------------
Acheta_domesticus           KFKHAH*GSSFHRAQWRRLRITTLRGKN*GRILFRQASYIIHLSRFS*YFFDIE*LSVQYLFCTYFVNLGDLYK*KSMVVRGEEKRWNV*ES*IL*EESHSP*RENTETFPNNLSNVIRC
Consensus

Teleogryllus_occipitalis    ------------------------------------------------------------------------------------------------------------------------
Acheta_domesticus           MSCILFGR*CQKV*A*NLIVKVR**VGHRIPLKKLDFALR*HASCL*IKMVLLG*TNYVEYITEVR*PSRYHFLAEATHTNIYFLIGESNSQFSET*RNVLS*LEVVWRHHYTPP**FPI
Consensus

Teleogryllus_occipitalis    ------------------------------------------------------------------------------------------------------------------------
Acheta_domesticus           AHDPS**SPIE*CTKLANTL*QKQRKKERKKERKKERNS*FQAN*CSPKILGSCNFLKNQCSARFKIKQVRTLT*KL*AVFLCVLL*CVS*LYRISECVWREHAEIRPTGLLVTWKRVAL
Consensus

Teleogryllus_occipitalis    ------------------------------------------------------------------------------------------------------------------------
Acheta_domesticus           DTSSLPGESRFQHLRTHFTPSTSPYLAKTQP*DINEGQSHTDGTLMRQGIRFAAIFVDARGQRGDVYYVFTSKVKEPFKKRVCGVM*QTPGVSILLQ*FRTFFPHRKFKSASWGGYFVFS
Consensus

Teleogryllus_occipitalis    ------------------------------------------------------------------------------------------------------------------------
Acheta_domesticus           FSKYLLGLEICRIFNDWISTMFVMVE*FHYT*PVIKRQTLNALMRPLDF*YFYYFRFK**VIL*QLIFNI*FIINI*IFLGEIIVFI*YTVPIFSMLILLFRIVHYERKTAKYK**KKHI
Consensus

Teleogryllus_occipitalis    ------------------------------------------------------------------------------------------------------------------------
Acheta_domesticus           A*YDKEHYIRFAKVP*NVRFKNIG*SN*DVISTKISFTR*TAYDNGAMIMELSIFTSIIDKSVPQLRVTRQQKYCQTSVTLHLLQHFLQSLGFQAVQQKPVIIQRGLFPAHQINDPRKKI
Consensus

Teleogryllus_occipitalis    ------------------------------------------------------------------------------------------------------------------------
Acheta_domesticus           EMMHPLYKHFRGTGRRVAEPARRGATRLGRSREGAHPRPESLCPPRRAGATHGERNHRNHRAETFGWRGGYELIGQR*I*ASGGSWDILLAVGYN*RSQTNSSS*CKLNPNINKSINAAC
Consensus

Teleogryllus_occipitalis    ------------------------------------------------------------------------------------M----------
Acheta_domesticus           SSKNSAVQNDFPKSSKHAAK*QQIKY*SYNKILTYYIL*NTIHENIIMTT*IK*LKRNRLKKYC*NING*KPERYK*HFINCEKMC*V*AIFQLQ
Consensus                                                                                                       *
```

**Tocc.6G0011210.1_exon2**

```
Teleogryllus_occipitalis      ------------------------------------------------------------------------------------------------------------------------
Acheta_domesticus             H*EQLLLLLYVSTPDTKYYAGASSFNYLSIFFHIFVPINSSPAQPIKYQNFNNLKY*KSEV*NTYS*VGHLDINAWNENSKGHFL*SGIHVYCEDHFHGNAEGIP*K*AALLILFYSHL*
Consensus                     

Teleogryllus_occipitalis      ---------------------------------------------------------RW----LTLQ-----------------------------------------------------
Acheta_domesticus             VLCNYGNNRS*QIHNTSSVIRYTNNGDDITRPYWGDPR*SEQDDYSPLLGSGFRSR*RWWNL*LTLRDCGRFLDFLPFYRPLITLWLRHYIATTSSLAASLIGQRARFP*RDPGFDSRSG
Consensus                                                                              **    ***

Teleogryllus_occipitalis      ----------------------------------------------------------------------------------------------------------QHTKKDRK------
Acheta_domesticus             *KYSVRFTGNVPVYVAV*R*ENSLLRLRRSD*GVGCLRVNHTNPLHPQGREAGTTLVGPSPHTGCCARYVHTSLSSSPVYQNHLSHVCRKE*VLRIIKINTLLSCPRHTRSV*KVSDLWP
Consensus                                                                                                                **    *

Teleogryllus_occipitalis      -KI---------------------------------------------------------------------NKEKETK----------------------------------ERKEKEK
Acheta_domesticus             EKIFRILDRFGTLIPFK*APCASTHLAHRSCHCWKHLWKSSFGMVFSSVVAFRIISSLLSKRDPFSDIFNFGNNQRS*ESVDQLG*MCRGVVMMQMPVFRRPHVWSFAPNCVTESPENIS
Consensus                      **                                                                     *                                        *  *

Teleogryllus_occipitalis      RERERKKE----------------------------------------------------------------------------------KREKKKER---------------------K
Acheta_domesticus             RTVNRRSAITKFRTLSTTAALRAVWGLAPLFSLSADVRPSLNRLNHCLICVTPIASSPNTC*ILRIVWL*ESPSF*QNLMQYLCSIRSVILREN*IRRTL*TAPYLATTSQRLAPANAVK
Consensus                     *   *                                                                                      **    *                     *

Teleogryllus_occipitalis      KRK--------RKEKEKRERKEKEKEKEKER-------------------------------------------------KKKEK-----------ERKRKKERKKKE------------
Acheta_domesticus             KFKHAH*GSSFHRAQWRRLRITTLRGKN*GRILFRQASYIIHLSRFS*YFFDIE*LSVQYLFCTYFVNLGDLYK*KSMVVRGEEKRWNV*ES*IL*EESHSP*RENTETFPNNLSNVIRC
Consensus                     * *              * *      *   *                                                    **           *      *   *

Teleogryllus_occipitalis      ----------QK--------KERKRKRKR------------------------------EKEKERRKKER--------------------ERKKEKERK---------------------
Acheta_domesticus             MSCILFGR*CQKV*A*NLIVKVR**VGHRIPLKKLDFALR*HASCL*IKMVLLG*TNYVEYITEVR*PSRYHFLAEATHTNIYFLIGESNSQFSET*RNVLS*LEVVWRHHYTPP**FPI
Consensus                               **        * *     *                              *   * *   *                        *  *

Teleogryllus_occipitalis      ---------------------KRERKKEKERERKKERERK-----------------RKKEKERKKEKE-------------------------------RERKKER-------------
Acheta_domesticus             AHDPS**SPIE*CTKLANTL*QKQRKKERKKERKKERNS*FQAN*CSPKILGSCNFLKNQCSARFKIKQVRTLT*KL*AVFLCVLL*CVS*LYRISECVWREHAEIRPTGLLVTWKRVAL
Consensus                                             ****   ******                          * * *                                **    *

Teleogryllus_occipitalis      -------ERKRKKER--------------------------------------------------------KRKKERKKER---------------------------------------
Acheta_domesticus             DTSSLPGESRFQHLRTHFTPSTSPYLAKTQP*DINEGQSHTDGTLMRQGIRFAAIFVDARGQRGDVYYVFTSKVKEPFKKRVCGVM*QTPGVSILLQ*FRTFFPHRKFKSASWGGYFVFS
Consensus                            *      *                                                           **  * *

Teleogryllus_occipitalis      ----------------------------------------------------------------------------------------------------------ERKKERKK------
Acheta_domesticus             FSKYLLGLEICRIFNDWISTMFVMVE*FHYT*PVIKRQTLNALMRPLDF*YFYYFRFK**VIL*QLIFNI*FIINI*IFLGEIIVFI*YTVPIFSMLILLFRIVHYERKTAKYK**KKHI
Consensus                                                                                                                     ***    *

Teleogryllus_occipitalis      ---EKE---------------------------------------------------------------RERK-------------------------------------------RKK-
Acheta_domesticus             A*YDKEHYIRFAKVP*NVRFKNIG*SN*DVISTKISFTR*TAYDNGAMIMELSIFTSIIDKSVPQLRVTRQQKYCQTSVTLHLLQHFLQSLGFQAVQQKPVIIQRGLFPAHQINDPRKKI
Consensus                         **                                                               *  *                                           ***

Teleogryllus_occipitalis      ------------------ERKRKKEKERERKKERKKEKKEK----KRKKERKKERKRKKERKR---------------------------------------------------------
Acheta_domesticus             EMMHPLYKHFRGTGRRVAEPARRGATRLGRSREGAHPRPESLCPPRRAGATHGERNHRNHRAETFGWRGGYELIGQR*I*ASGGSWDILLAVGYN*RSQTNSSS*CKLNPNINKSINAAC
Consensus                                       *  *       *  *      *      *      **     *

Teleogryllus_occipitalis      ------------KKEKERKKEKERKR-----------------------------KKERERK-------KEKERKR------QKLILLG*-----
Acheta_domesticus             SSKNSAVQNDFPKSSKHAAK*QQIKY*SYNKILTYYIL*NTIHENIIMTT*IK*LKRNRLKKYC*NING*KPERYK*HFINCEKMC*V*AIFQLQ
Consensus                                 *  *   *    *                              *  *  *           **        *
```

**Tocc.6G0014840.1**

```
Teleogryllus_occipitalis      ATATTCCGTCCTGAAAATCTTATCTTGAATGCAATGAGAGGCACTCATGACT--------ACGTTCATAATAGATGTTATTAGTT--------GTATGTTTCGT---------------
Tarbinskiellus_portentosus    ATGCGAGATTGGGAAAG------------AGTGACGAGATGTGGCTATGAGGCATCCAGAGCGACCAAGCAAGACAGGAGAAGTCTGGAAATAACAGGTCAAGT------AATTGAGGAA
Acheta_domesticus             ATATTCCGTCGTGAAAATCCGCTCTTAAATACAATGAGAGGCCCTCGTGAGT--------CTAT------TAGATGTTACCAGTCTGTAATGAATATATTTCATCCTGGGGATTCAGAAT
Gryllus_bimaculatus           GTACTCCGTCTTGAAAATCCTGTCTTGAATAGCACGAGAGGCCTTCATGAATG------TATGTTCTCTTAAGATGATAGTAGTCTGTTATG---------------------------
Gryllus_longicercus           CTACTCCGTCTTGAAAGTCCTATCTTGAATATAATGAGAGGCCCTCATGAGT---------TGTTCTC--------ATGTTAGTCTGTAATG---------------------------
Consensus                      *      *   ****                 * **** *      ***                               ***

Teleogryllus_occipitalis      -------------------------------------------------------------------GATGTACTAGTAATATTATTT--------------------------------
Tarbinskiellus_portentosus    GAAAGAAAAA-------------------TGTGGGGAAACGGAAAT----------------TATAGTAATTTCTAGTAACACCATTT-------------TTATCGCTGATTGATATC-
Acheta_domesticus             GAAAGAAATATCCCAATATCTTCTGTCTTTGTGTTTTAATAAAATTTCTTTGTCCAGTTTTATTTTAGACAGTCTAATACTATCATTTGAAAACCAACAAGTTAGTAAAGGGACATATTT
Gryllus_bimaculatus           ------------------------------------------------------------------------------AATATTATTT--------------------------------
Gryllus_longicercus           ------------------------------------------------------------------------------AATATTATTT--------------------------------
Consensus                                                                                   *  *  ****

Teleogryllus_occipitalis      ---TTCTGGGGATTTA----------------------------------------------------------------------GGATGAAAGAAATAATTGCAATTTGA-TTTCGGC
Tarbinskiellus_portentosus    ---TTCAAAGTCTTCG-------------------------------TCGTTGTCATCTCATTACGGCCA---------------------------------GCAGCTTCG---TTGAC
Acheta_domesticus             TAAACTTAAAAATTTATTGAAATGTGTTTTTTATGCATGTTGATTCATTATTCTCTTTGAATCACGACTATCATTACTATCTATTTTAATGAAAAAAATAGTTGCAATTTCA-TTTTTGT
Gryllus_bimaculatus           ---TTCGTAGGATTCA----------------------------------------------------------------------GGTTGAATGAAATAATTGCAATATGA-TTTCGGC
Gryllus_longicercus           ---TATG-AGGATTCG----------------------------------------------------------------------GGTTGAATGAAATAATTGCAATATGAGTTAGGGT
Consensus                                 **                                                                                         ***   *

Teleogryllus_occipitalis      GTGAGATGGCTAGTTTCCTTTCTGTGAAATGCATAGCGAATTTGTTATGACA---CAACTTCGATCATAGATCTAG--ACTAGTGTAATGATTATG---------------------GCC
Tarbinskiellus_portentosus    TTCATATCTCCGTCTCGTTGCCAAT---------GGCGAGTTTGACGTAGTAGTCAATTCACAGTCACAGATCCAGAATCGAAATCAATAATGATATAAAAGTGTACTCTCACCGACACC
Acheta_domesticus             GTGAGATGGCCATTTTCCTACCAGTTAAATACGCGTCGAATGTGCTGTGACA---CACCTCTGATCGTAGATCTAGTCTCTAAGGTTATGATTATA---------------------GCT
Gryllus_bimaculatus           GTGTTATGGCCATTTTCCTACCAGTTAAAT-AGGGGCGAATTTGTTATGACA---CACCTTTGATTATAGATCCAG--------------------------------------------
Gryllus_longicercus           GTGTTAC-GCCATTTTCCTACCAGTTAAATAAGGGGCGAATTTGTTATGATA---CACCTTTGATTATTGATCCAG--------------------------------------------
Consensus                      *   *   *    *   *  *  *           ***  * **    *   *   *       *    **** **

Teleogryllus_occipitalis      TTGACTGTCACTCTGGCAACGTGGGCCCAGCTCTCTATTTTGAAACCTGCTATGGACATCTTATCTTCCCCAACC------------------------------GTATTGCCATTCATG
Tarbinskiellus_portentosus    TTCATTCTTCCGCTAATCAAACCAATTC--TCTGTCCTTTCTAATTCCGCTTCACACATTTCTCCCCCTTCCGCATTTTGAATCCTTGATGTTTCCAGAATCCTCCTGTCTTCCTTCGTC
Acheta_domesticus             TTCTCTGTCA--CTGGCAACGTGGGCC---------GTTTTGAAACTTGCTATGGACATCTTATCGTCCTCAGC--------------------------------GCTCCCCATTCATG
Gryllus_bimaculatus           ---------ACTCTGTCAACGTGGGCCG----AGCTGTTTCGAAACTTACTATGGACATCTTATCTTCTCCAACG------------------------------TTCTTGCCATTCATG
Gryllus_longicercus           ---------ACTCTGTCAACGTGGGCCGAGCTAGCTGTTTCGAAACTTGCTATGGACATCTTATCTTCTCCAACG------------------------------CTCTTGCCATTCATG
Consensus                                 **    *                  ***  **     **    **** *   *  *  *  *                                  *   * *** *

Teleogryllus_occipitalis      TG----TTTTTTTACGTAGTCGCCATGTTCGTCCCA--------GCTGCGCGTCTGTCTTCACCTCCGCGT---------------CGGCCGTCGCCCTCGCCTCCGCCCACGCGGCGT-
Tarbinskiellus_portentosus    GC-TCCTGTTGCCTTATAGCCGC--TCCTCGTCACCCTTTCCCAATCTCGCGTTCGTTTCCTCGTCTTTCG-ACCCTGCGCCGCTCCCACCAGCAGCGGCACCTCCTCCCACTCACAGTC
Acheta_domesticus             CGTTTTTTTTTTTATGTAGTCGCCATCTTCGTCACT--------GCCGCGCGTGTGTCTTCACGTCCGCCTCCTTCTCGGCCTGCGCGGCCGTCGCCCTCGCCACCGCCCACGCCGCGT-
Gryllus_bimaculatus           TT-TTTTT----TATATAGTCGCCATCTTCGTCACT--------GCCGCGCGTCTGTCTTCACCTCCGTGT------CGGCCGGCGCGGCCGTCGCCCTCGCCACCGCCCACACAGCGT-
Gryllus_longicercus           TT-TTTTTTTTATATATAGTCGCCATCTTCGTCACT--------GCCGCGCGTCTGTCTTCACCTCCGTGT------CGGCCGGCGCGGCCGTCGCCCTCGCCACCGCCCACACAGCGT-
Consensus                           *         *** ***  *  ***** *            *****  ** * * * **                    *  **  *  *  * ** ** ***** *   **

Teleogryllus_occipitalis      CGCCACCGCCTCCGTTCGCCTTCCACTCCGCTATCTCGACCTGCCG---------------CCGCCGCCGCCGCCGCATACGTGATGCAATGCACCTGGCCGCCGGCCACTGGCTGTGCA
Tarbinskiellus_portentosus    CGCTACCGACA----CCGCCTTCCACTCCGCTATCTCGACCTGCCAACGCCTCCGCCGGCCTGGCCGCCGTCGCCGCATACGTGATGCAATGCACCTGGCCGCCGGCCACTGGCTGTGCA
Acheta_domesticus             CGCCACCGCCAGCGCCCGCCTTCGACTCCGCTATCTCGACCTGCCGCTG----------CCCCGCCGCCGTCTCCGCATACGTGATGCAATGCACCTGGCCGCCGGCCACTGGCTGCGCA
Gryllus_bimaculatus           CGCCACGGACACTGCCCGCCTTCCACTCCGCTATCTCGACCCGCCGCCGCCGCCGCCGCCGCCGACTCCGTCTCCGCATACGTGATGCAATGCACCTGGCCGCCGGCCACTGGCTGTGCA
Gryllus_longicercus           CGCCACGGACACTGCCCGCCTTCCACTCCGCTATCTCGACCCGCCGCCGCCGCCGCCGCCGCCGACTCCGTCTCCGCATACGTGATGCAATGCACCTGGCCGCCGGCCACTGGCTGTGCA
Consensus                     *** ** * *      ******* ***************** ***                  * * *** * ******************************************* ***

Teleogryllus_occipitalis      GGGGCGCGG--------TGTGCGCCGCTCCGCACGCGGCGGCGGCGGCGGGGGCAGGTCCGTGACGTCACGAAATATGACGCCGGGGGCAGCCGTCCGCCGCGCCGCGCCGCGCCGCGCT
Tarbinskiellus_portentosus    GAGGCGCGGTGCGGTGCGGTGCGCTGATCCGCGG-------CGGCGGCGGGGGCTGGTCCGTGACGTCACGAAATATGACGCCGGGGGCAG------------------------------
Acheta_domesticus             GGGGCGCGG--------TGTGCACCGCTCCGCAC---GCGGCGGCGGCGGGGGCAGGTCCGTGACGTCACGAAATATGACGCCAGGGGCAGCCGTCCGCCACGCCGCGCC----------
Gryllus_bimaculatus           GAGGCGCGGT---GTACTGTGCACCGCTTCGCAC--------GGCGGCGGGGGCAGGTCCGTGACGTCACGAAATATGACGCCGGGGGCAGCCGGCCGCCGCGCCGCGCCGCGCC-----
Gryllus_longicercus           GAGGCGCGGT---GTACTGTGCACCGCCTCGCAC--------GGCGGCGGGGGCAGGTCCGTGACGTCACGAAATATGACGCCGGGGGCAGCCGGCCGCCGCGCCGCGCCGCGCC-----
Consensus                     * *******         **** * *   ***          ************ **************************** *******

Teleogryllus_occipitalis      GTGCCGGTTGAATATCCGCGTAAT-AGCCGTGGGCGGGCCCGCC-----CCGGCCCAGCCAGCGCGGCGATCAGGCGGTGGCGGTGGCGGCGGCGGCGGCGGCGGCGGCGGGGGCTG
Tarbinskiellus_portentosus    CCGCCGGTTGAATATCCGCGTAATGGGCCACGGGCGAGCTGGG-------------------CGCGGCGATCACACGCCGACCA------------------------------------
Acheta_domesticus             GTGCCGGTTGAATATCCGCGTAAT-GGCCGTGGGCGGGCCCGGCCCGGTCCGGCCCAGCCAGCGCAGCGATCAGGCGGC-----------------------------------------
Gryllus_bimaculatus           GCGCCGGTTGAATATCCGTGTAAT-GGCCGTGGGCGGGCCCGC------CTGCCCCAGCCAACGCGGCGATCAGGCGGC-----------------------------------------
Gryllus_longicercus           GCGCCGGTTGAATATCCGCGTAAT-GGCCGTGGGCGGGCCCGC------CTGCCCCAGCCAGCGCGGCGATCAGGCGGC-----------------------------------------
Consensus                       **************** *****  ***  ***** **  *                    *** *******  **

Teleogryllus_occipitalis      AGGCGGCGGGCGGAGGTGGAGGTGGAGGTGGAGGTGGAGGCGGAGGTGGAGGCGGAGGCGGAGGCGGAGGCGGAGGCGGAGGCGGAGGCGGAGGCGGAGGCGGAGGCGGAGGCGGAGGCG
Tarbinskiellus_portentosus    ------------------------------------------------------------------------------------------------------------------------
Acheta_domesticus             ------------------------------------------------------------------------------------------------------------------------
Gryllus_bimaculatus           ------------------------------------------------------------------------------------------------------------------------
Gryllus_longicercus           ------------------------------------------------------------------------------------------------------------------------
Consensus

Teleogryllus_occipitalis      GCGGCGGAGGCGGAGGCGGCGGCGGAGGCGGAGGCGGCGGCGATGGCGGCGGCGGCGGCGGAGGCGGCGGCGGAGGCGGAGGCGGAGGCGGTGGCGGCGGCGGCGGTGGCGGCGGCGGAG
Tarbinskiellus_portentosus    ------------------------------------------------------------------------------------------------------------------------
Acheta_domesticus             ------------------------GCAGCGCCGCTGGTGGCGGTGGCGGCGGCGGCGGCGGCGGCGGCGGCGGCGGCGGCGGCGGCGACGGCGGCGGCGACGGCGGCGGCGGTGGCG
Gryllus_bimaculatus           -------------------------------------------------------------------------------------------------------------GGCGTCC
Gryllus_longicercus           -------------------------------------------------------------------------------------------------------------GGCGGCG
Consensus

Teleogryllus_occipitalis      GCGGCGACGTCCAGCAGC-CTGTACCCCACCGCGACCGGAGCGCGAACCGGCAGTGAACCGTTTAACCCAATGAAACTCCTGTTCTTCCCGGTTGGGAAGCTGGTGCAGTGGAACCGAGA
Tarbinskiellus_portentosus    --------ACCCGGCTCC-GCGACCTGCTCCGCGATCAGGTCCCGAACCGGAACTGAACCGTTT-ACGCGAAGATCTTGATATCTTCCCGTTTCTGGGGCCCAAAGTATTTTGATCATGA
Acheta_domesticus             GTGGTGGCGTCCGTCACCGCCGTGCCACACCGAGACCGCAGCGCGAACCGGCACTGAACCGTTTAACCCAATGAAACTCCTGTTCTTCCCGGTTGGGAAACTGGTGCAGTG-----GAGA
Gryllus_bimaculatus           GGCGCCTGATCCGTCCGCGCTGTGCCCCACCGGGGCCGGAGCGCAAACCGGCACTGAACCGTTTAACCCAATGAAAATCTTCTTCTTCCCGGTTGGGAAACTGGTGCAGTGGAACCGGGA
Gryllus_longicercus           GCGGC---GTCCGGCAGCGCTGTGTCCCACCGGGGCCGGAGCGCGAACCGGCACTGAACCGTTTAACCCAATGAAAA---TCTTCTTCCCGGTTGGGAAACTGGTGCAGTGGAACCGGGA
Consensus                               **  *  *   *     * *** *  *    * * ****** * ********** ** * * **      * *  * **   *  **   *    * * *         **

Teleogryllus_occipitalis      CGATTATCCGACTCGGAAGTGGGAAACAATTTAACGGAAATAAAATGTTTTCGTGTATCGAGGTTGGGACCAATGGTGAAGCA--AAAGGAAATCAAACCTCGAGTTATCAATAATTTCG
Tarbinskiellus_portentosus    C-------------------------TCAAGTTCACGTGATCACAGTTTGT----------------GTAGTATTGTTGGCGTCGTATAGGGAGGAAAGACT---GTTATCAATGTTGTGG
Acheta_domesticus             CGATTATTCGAATCCAAAGAGTGAAACAATTTCACAGAAATAAAATAATTTCTCGTGTCGAGATTGATGATGTT--------------GAAATCAAAACTGAAGTTATCAATAATCAAA
Gryllus_bimaculatus           CGATCATTCGAAACGCAAGCGGGAAACGATTTAACGGAAATAAAATATTTTCTCATGTCGATAT----GATATTGGTAAAGC-------GTATAAAAAATATATGTTACCAAAGATTTCT
Gryllus_longicercus           CGATCATTCGAATCGCAAGCGGGAAACGATTTAACGGATATAAAATATTTTCTCATGTCGATAT----GATATTGGTAAAGC-------G--AAAAAAATCAATGTT-----AGATTTCG
Consensus                     *                         * * ** **      * * *   *                       *               *     **      ***        *

Teleogryllus_occipitalis      -ACACAGATGAACTAACGAGCTAAAGT--AATTGATAAATTC--------ATGAATTGGAGTTAAAAAATAATTCGT-------------GATTTTTCAAATAATTAGTATCATTGCAGT
Tarbinskiellus_portentosus    -GGAAGATACAGTTGGCAAACCGTACTTTGAGCTCAGAATTCCGGTCGT-GCGGACCGGAACT---GAACCGTTTGC-------------CTGCCTTTCTGAAATCGTTCCATTTGAGGG
Acheta_domesticus             CTCATAAATAAGGTGACAAGTTAAAAT--AATCACTAAGTTTGGACGACGATGAATTAGGGTTACCAAAGAATTTGTCACGAAAGTTTAAGGGTTTTCTAATAATTAGAATTATTGCAGG
Gryllus_bimaculatus           -ACTCAGATGAACAGACGAGTTAAAG-----GCAATAAATTC--------ATGAATAAGAGTTATCAAAGATTTTGT--CGATCGAAGAATAGTTTTTAAAGAATTAGTATTATTACAGT
Gryllus_longicercus           -ACTCGGATGAACAGACGAGTTAAAG-----TCAATAAATTC---------CGAATTAGAGTTATCAAAGATTTGGT--CGATCGAAGAGTAGTTTTC-AAGAATTAGTATTATTACAGT
Consensus                               *     * *     *            * **           * *   *   *    **   ** *                   **     ***        **   *

Teleogryllus_occipitalis      AATAGAGTTATC------------------------------------------------------------------------------------------------------------
Tarbinskiellus_portentosus    GGTCACATTGCC-----------TTGATCACAACTCCAAAGTTCGTACATTTGTGTGGTATTAT---TTTTGTGTCCAACG---------------------------------------
Acheta_domesticus             AATCGAGTTATCGAAGAATTCGAGTTATCAGAATTTTAAAGTTTGAAACGTTCCATTTTATTGTGAGTTATAATCCGAACGAAAGTCCAGAATCTGAACGATAAGTTACTACGTAGTACT
Gryllus_bimaculatus           AATCGCGTTATCAA-----------------GATTTT--CGTTTGAAATGTTCCATT----------TTATAATGCGAACGAAATTTCATATTATAAACGATAGGCTATGACAAAATTTT
Gryllus_longicercus           AATGGCGTTATCAAT-------------CGCGATTTT--GGTTTGAAATGTTCCATT----------TTATAATCCAAACGAAATTTCATATTAAAAACGATAGGCTATGACTAAAATT
Consensus                       *    **  *

Teleogryllus_occipitalis      ------------------------------------------------------------------------------------------------------------------------
Tarbinskiellus_portentosus    -----------------------------------------------------GTGACGACTGTTGCCA-TTGTTGTTGCAAAGATAAGGTTGGTATTGGCAACAGTTTTTTTGTTCATG
Acheta_domesticus             CAGAAACGGGTATCTAGATCCAAATTTTTACGTTGAGTAATAATTTATGATTTGTGACCACACAAACCTGCAGTATGAACCACAGAAACTTTGGCATATGAAATGACACGTTTGTTTACA
Gryllus_bimaculatus           AATGAAGTG-----------------------------AATTACTAAACAATTGTGACCACTCACACCT---GTATGAACCATTGATACGCTAGCACATGGTCTGATACGTTTG-TAACA
Gryllus_longicercus           AATGAAGTG-----------------------------AATTACTAAAGAATTGTGACCACTCAAACTT---GTATGAACGACGGATACTCCAGCACATACTGTGATACGTTTG-TTACA
Consensus

Teleogryllus_occipitalis      ----------------------------------------------------CAGATTTATCC---------------------------------------------------------
Tarbinskiellus_portentosus    ACTGGAGAGCCCAGATTTCTTTACTTTATACAAATATTTCTCGA--------AAAATTTGATGAGTAACTGGACAGACACATCTGCTTGGCACGTGCCTGTCACA---TGTCTGAACGGG
Acheta_domesticus             GATTT--AAGTCAGATAAACTGGCTTCAATCGAACGTTCTGGAA-AATAAGTCGGATTCGTTCG-GGGCCAG-CAGACAC-TGGGAGCAGCACGTGCCTGTCACACGCTGTCTGCGCGGG
Gryllus_bimaculatus           GATTTCAGAGTCACATAACGTGATTTAAATGGAACCTTTTGGAACAAAATATGGGATTCGTTGAAGGGCTAG-CAGACAC-CGGGCGCAGCACGTGCCTGTCACCCGGCGTCTGCGCGGG
Gryllus_longicercus           GATTTCAGAGTCACATGAAGTGATTTAAATGGAACGTTCTGGAACAAAATATGGGATTCGTTGAAGGGCTAG-CAGCCAC-CGGGCGCAGCACGTGCCTGTCACCCGGCGTCTGCGCGGG
Consensus                                                            ***

Teleogryllus_occipitalis      ------------------------------------------------------------------------------------------------------------------------
Tarbinskiellus_portentosus    GCTTGATTCACGATGCGCCCGGGGCTTCCAGCCTCTGCGGTTGCGAAGAACTCCAAGGTGCGGAGATTCCTGGAA--------CGCCTGGGTTTCGTCTCGTCAGATCAGTGGGCT----
Acheta_domesticus             CCCTGACTC-----------ACGGTTTCCGGCCTCTGCGGTTGCGAGGAACTCCAAGGTGCAGGGCTTCTCGGAACGGGGGGTCGCCTGCGCTTCGTCTGCTCGGATCACTGCATCACTC
Gryllus_bimaculatus           CCCTGACTCACGTTGCGCAGACGGTTTCCGGCCTTTGCGGTTGCGACGAACTCCGAGGTGCGGAGGTTCTCGGAACGGGGGGTCGCCTGGGCTTCGTCTGGGCGGATCACTGCATCACTC
Gryllus_longicercus           CCGTGACTCACGTTGCGCAGACGGTTTCCGGCCTTTGCGGTTGCGACGAACTCCGAGGTGCGGAGGTTCTCGGAACGGGGGGTCGCCTGGGCTTCGTCTGGGCGGATCACTGCATCACTC
Consensus

Teleogryllus_occipitalis      ------------------------------------------------------------------------------------------------------------------------
Tarbinskiellus_portentosus    -CTCCGCGATAGGAAGCTCGATGGCTATGAATAATACAGACAGGAAGTGGAGAAATAGATTCCTCGAATTTCAACGCAAGTGCGATGAAAGAAAAATTTGAAGTTGGGAGGAAGG-AGGA
Acheta_domesticus             GCCCGCCGCGGGCACGCGGGACGCCTATGAATAATACAGACGGGAAGTGAAGGAATTGATTCCCCGAATTTCGACGGAAGTGTGATG---AAAAAATTAGAAGTTGGAGGAAGAAAAAAA
Gryllus_bimaculatus           GCCCGGCGCAGGCATGCCGAACGGCTATGAATAATACAGGCGGCAAGGGAAGAAATTGATTCCCCGAATTTCGACGGAAGCGTGATG---GAAGAATTAGAAGTTGGAGGAAGAA-GAAA
Gryllus_longicercus           GCCCGGCGCAGGCACGCCGAACGGCTATGAATAATACAGGCGGCAAGGGAAGAAATTGATTCCCGGAATTTCGACGGAAGTGTGACG---GAAGAATTAAAAGTTGGAGGAAGAA--AAA
Consensus

Teleogryllus_occipitalis      ------------------------------------------------------------------------------------------------------------------------
Tarbinskiellus_portentosus    ATATAATTACCAATATCAGGATTCCGACAGCGTAGTGGAAATT-TTATGTATGAAAATGTTAATTA-ACGATCATCAAATATATTAATAATTCTGTGTATTATATATATATATATATATA
Acheta_domesticus             ATGTAGATTTCTTT----GGAATTCGA-AGCATTAGGGAAATA-TTTTTAAAAAATATGTTAAT----TATTGATCAAGGGAAGTGATAATTCTGAATTTCGTTGTAACTTGAAT-----
Gryllus_bimaculatus           ATGTAGATTTCTGT----GGAGTCCGATGGTATTAGAGAAATATTTTTTTTTAAATATGTGGATTATATATCCATTACGGGAAGTGATAATTCTAAGTTTCGTTGAAACGTGTTT-----
Gryllus_longicercus           ATTTAGATTTCTGT----GGAGTCCGGTGGTATTAGGGAAATA-TTTTTTTTAAATATCTGGGTGATATAGCGATTATGGGAAGTGATAATTCTAAGTTTCGTTGAAACGTGTTT-----
Consensus

Teleogryllus_occipitalis      ------------------------------------------------------------------------------------------------------------------------
Tarbinskiellus_portentosus    TATGTATGTGTGATTAATATAAGAAAATCTATAAAATATAATTAAATCTCTTAAATTCAAATAATAATTATAGACGTACCGATAAATTATTT-----TAGAAAAGTAAATGTGTGAAGAG
Acheta_domesticus             --------------------------------------TACTTCAGACTTTCCACTGAAAGTAAGAAT-------ATATTGGAATATTA-------------------------------
Gryllus_bimaculatus           --------------------------------------TAATTCTGACTATCGAATGATAGTAATAAT-------ATTTAGGAATGTAATTTATTTGCAGGTGCGACAAAGAGTGAGAAG
Gryllus_longicercus           --------------------------------------TAATTCTGGCAATCCAATGATAGTAATAAT-------ATTTAGGAATGTAATTTATTGGCAGATGCGACAAACAGTGAGAAG
Consensus

Teleogryllus_occipitalis      ------------------------------------
Tarbinskiellus_portentosus    AGATAATAC---------------------------
Acheta_domesticus             ------------------------------------
Gryllus_bimaculatus           GGAT--------------------------------
Gryllus_longicercus           GGATAGTGTAGGTCAATGCAAGGCTTGAGGAGAGTG
Consensus
```

**Tocc.6G0014840.1_exon1**

```
Teleogryllus_occipitalis            ------------------------------------------------------------------------------------------------------------------------
Tarbinskiellus_portentosus          IISLHTFTFLK*FIGTSIIII*I*EI-*L-YFIDFLILITHTYIYIYIYNTQNY*YI**SLINIFIH---KISTTLS------ES*YW*LYSSFLPTSNFS-----FIALALKFEES---
Acheta_domesticus                   --------------------------------------------------------------------SLSLTSSYSCLFPSSKDHVLIQLTTLMPHRRIR-----SAQTSVLF------
Gryllus_bimaculatus                 -----------------MKYFKIKETFPL**IINFL--------------TQNFHTRCSNKIEHF*LNSTKIK*TKSDL----KSVLYCKLISFAL-RRFRL*FLQSLSFSISFSPSHTP
Gryllus_longicercus                 ---------------------KIKEIFPL-YIINLL--------------TLNLHT*CSNKIGHF*HNSTKVK*TKSDL----KSDVFCKLISFAP-RRFRLYFLQSLSFSISFSPSNTP
Consensus

Teleogryllus_occipitalis            ------------------------------------------------------------------------------------------------------------------------
Tarbinskiellus_portentosus          ------ISPLP-----------------------------------------------VC---IIHSHRASYRGEPTDLTRRNPG-------------------------VP--------
Acheta_domesticus                   -----LIPPLC-------LSLSLS------------------------------LRLALC----LRGAPAARGMREAECKQEQEG---------------ARRSAPCGELVPDAAAAASS
Gryllus_bimaculatus                 SLTH-FLSPTP-------LTISLSSYSQSSEDRIIKQLPTERPVPHRRPSSSNPLSVRLCRAQRLRGAPASRGMREAECKQEHDGAA*RTLRRRRRRRLRARDGTGDG-----G-EGNRG
Gryllus_longicercus                 PLSHLFTHPLPQSEPPHTLPLFLLSVFRRSRHKAIAYVSAAPKDPFRIDVRPLPIPLSVC---VVRSASAERRPR-AACA-----------RRNANKSTTARRSAPCG-----GCGGDFA
Consensus

Teleogryllus_occipitalis            ------------------------------------------------------------------------------------------------------------------------
Tarbinskiellus_portentosus          ----------------GISAP------------W-----------SSSQPQRLEAPGAS*IK----PRSDM*QARA--KQMCL-------------SSYSSNFSRNICIK*RN-------
Acheta_domesticus                   LAMALAMAVKEIGTRIAISA--GVIVRGEMCGCWERKTA*AVVMVTAGPPQP---PKAASRF-----RELFYALIADYSCCC*DCVDILFAVGINGTRFR---RRYSVVKIRS*IQ*EA-
Gryllus_bimaculatus                 LESDLGGRGS---ARREVRLEGEEGGVGDDGGRW*R*PC------GYGSPQGL-LPGLGSLLNVH--R*LFSLL----PVFCR----------LPVCWWN---KRYSVLRLENPVLNSTR
Gryllus_longicercus                 LAMALATAAKAIGARRAISV--GEGVRGERCG-WRERKA------AWAMTEG---GGDGDRLATAAPRSL--------------------------TWFGISFKRSSVIILADASILST-
Consensus

Teleogryllus_occipitalis            ------------------------------------------------------------------------------------------------------------------------
Tarbinskiellus_portentosus          -----------------LGSPV------MNKKTVANTNLI----------------------------------------------FATTMATVVTV--GHKNNTTQMYELWSCDQGNVT
Acheta_domesticus                   -LVSLLDVTSL**IYFILGI--------QNERNIPISSVFVF**NFFVQFYFRQSNTII*KPTS**RDIF*T*KFIEMCFLCMLIHYSL*ITTIITIYFNEKNSCNFIFV*----DGHF-
Gryllus_bimaculatus                 GLHECM----------------------FS*DDSSLL*IL-----FFVGFRL------------------------------------------------NEIIAI*FRRV----MAIFL
Gryllus_longicercus                 ---SCLLVE*T-----VLATPS*KSYLEYNERPS*VVLMLVCNEYYFMRIRV------------------------------------------------E*NNCNMS*GV----LRHF-
Consensus

Teleogryllus_occipitalis            ---------------------------------------------MDILSSPT-------------------------------------------------------------------
Tarbinskiellus_portentosus          PSNGTISERQANGSVPVRTTGILSS-----------------KYGLPTVSSPQH**QSFLPIRRQQYYTNCDHVNLSHDQNTLGPRNGKISRSSRKRFSSGSGPDRGAGRGAGLVGV*SP
Acheta_domesticus                   PTS*IRVE-----CAVTHL*S*I*SLRL*L*LSLSLATWAVLKLAMDILSSSA------LPIH-AFFFLCSRHL-----------RHCRACVFTSASFSACAA-----------VALATA
Gryllus_bimaculatus                 PVK*GR--------ICYDTPLIIDPDSVNVGRAVS-------KLTMDILSSPT-----FLPF---MFFLYSRHL-----------RHCRASVFTSV--SAGAA-----------VALATA
Gryllus_longicercus                 PTS*IRGE-----FVMIHL*LLIQTLSTWAELAVS-------KLAMDILSSPT-----LLPF---MFFFY----------------------------------------------I*SP
Consensus                                                                            **

Teleogryllus_occipitalis            ------------------------------------------------------------------------------------------------------------------------
Tarbinskiellus_portentosus          RPARPWPITRIFNRRLPPASYFVTSRTS--PRRRRGSAHRTAPRL---------------------CTASGRRPGALHHVCGDGGQAGGGVGRSR*RSGRRC----------R*RTVSGR
Acheta_domesticus                   HAASPPPA---------PAFDSAISTCR---------------------------CPAAVSAYVMQCTWP---P-AT------------------GCAGARCAPLRTRRRRGQVRDVTKY
Gryllus_bimaculatus                 HTASPRTL---------PAFHSAISTRRRRRRRRLRLRIRDAMHL---------------AAGHWLC-----------------------------RGAVYCAPLRTAAGAGP*RHEI*R
Gryllus_longicercus                 SSSLPRV-----------CLHLRVGRRGRRPRHRPHSVATDTARLPLRYLDPPPPPPPPPTPSPHT*CNAPGRRPLAV-----------------QRRGVLC----TASHGGGGRSVTSR
Consensus

Teleogryllus_occipitalis            ------------------------------------------------------------------------------------------------------------------------
Tarbinskiellus_portentosus          RCRCWWERRRVER-------RGNER---EIGKG*RG--------------------AA-----------------IRQQERRRK--------------------TGGFWKHQGFKMRKGE
Acheta_domesticus                   DARGSRPPRRAVPVEYPRNGRGRAR-PGPAQPAQRS-GGAAPLVAVAAAAAAAAAAAAAAAATAAATAAAVAVVASVTAVPHRDRSANRH*TV*PNETPVLPGWETGAV-----------
Gryllus_bimaculatus                 RGQPAAAPRRAAPVEYPCNGRGRARLP---QPTRRSGGGVRRLIRPRCAPPGPERKPA-----------------LNRLTQ*KSSSSRLGNWCSG---------TGT-------------
Gryllus_longicercus                 NMTPGAAGRRAAP--------------------RRA--G*ISA*WPWAGPPAPA-SAA-----------------IRRRRRRRPAA-----LCP----------TGA-------------
Consensus

Teleogryllus_occipitalis            ------------------------------------------------------------------------------------------------------------------------
Tarbinskiellus_portentosus          KCVKRN*KGQRIGLISGRMKVSVRVHFYIIIDFDSGSVT----VN*LLRQTRHWQRDGDMKSTKLLAVMR*QRR--------RL*RYQSAIKMVLLEITIISV------SPHFSFFLNYL
Acheta_domesticus                   ETIIRIQRVKQF----HRNKIISRVEI------DDVEIK----TEVINNQT---HK*GD----KLK*SLSLDDD------ELGLPKNLSRKFKGFLIIRIIAGIELSKNSSYQNFKV*NV
Gryllus_bimaculatus                 --IIRNASGKRF----NGNKIFSHVDMIL-------------VKRIKNI--C-YQ----------RFLLR*TDELKAINS*IRVIKDF--------------------------------
Gryllus_longicercus                 --GARTGT-EPF----NPMKIFFPVGKLVQWNRDDHSNRKRETI*RI*NIFSCRYDIGKAKKINVRFRLG*TDELKSINSELELSKIWSIE-E*FSRISIITVMALSI-----AILV*NV
Consensus

Teleogryllus_occipitalis            ------------------------------------------------------------------------------------------------------------------------
Tarbinskiellus_portentosus          TCY--------------------------------------------FQ----------------------------------TSPVLLGRSGCLI------ATSR--HSF-PI-SHSLP
Acheta_domesticus                   PFYCEL*SERKSRI*TISYYVVLRNGYLDPNFYVE**FMICDHTNLQYEPQKLWHMK*HVCLQI*VR*TGFNRTFWKISRIR-SGPADTGSSTCLSHAVCAGPDSR--------------
Gryllus_bimaculatus                 ----------------------------------------VDRRIV-FKELVLLQ*SRYQDFRLKCSIL*CERNFIL*TIGY-DKILMK*ITKQL*PLTPV*TIDTLAHG-----LIRL*
Gryllus_longicercus                 PFY---------NPNEISY---*KR*AMTKN**SE-LLKNCDHSNL-YERRILQHIL*YVCYRFQSHMK*FKWNVLEQNMGFVEGLAATGRSTCLSPGVCAGRDSRCADGFRPLRLRRTP
Consensus

Teleogryllus_occipitalis            ------------------------------------------------------------------------------------------------------------------------
Tarbinskiellus_portentosus          RLSNLRR-------------RRLHDLHRESETAAARMRRRPRHRRRD-------------RRTSPRMAARAVAGRVPISLAATARPRRHRSIRYPLAARCATRVALLLFACRPSHAARDR
Acheta_domesticus                   ------------------------------------------------FPASAVARNSKVQGFSERGVACASSARITASLAR----RGH-----------AGRL*IIQTGSEGIDSPNFD
Gryllus_bimaculatus                 QISESHNVI*MEPFGTKYGIR*RASRHRAQHVPVTRRLRGP*LTLRRRFPAFAVATNSEVRRFSERGVAWASSGRITASLAR----RRH-----------AERL*IIQAAREEIDSPNFD
Gryllus_longicercus                 RCGGSRN-------GGSPGLRLGGSLH---HSP--------------------------------------GAGTPNGYE*Y----RRQ-----------GKKL--IPGISTEV*RKN*K
Consensus

Teleogryllus_occipitalis            -------------------------------------------------------------------------------------------------------------
Tarbinskiellus_portentosus          RSAGERE--------GGRRRR-----------------WNL---------------------------RLGTEDVNL------QMDRMDNWRREFNMPSEFIRC-----
Acheta_domesticus                   GSVMKK------LEVGGRKKM*ISLEFEALGK-YF*KIC*L----------LIKGSDNSEFRCNLNYFRLSTESKNILEY-----------------------------
Gryllus_bimaculatus                 GSVMEE------LEVGGRRKCRFLWSPMVLEKYFFLNMWIIYPLREVIILSFVETCFNSDYRMIVIIFR------NVIYLQVRQRVRRD--------------------
Gryllus_longicercus                 ---LEEEKI*ISVESGGIREI------------FFLNIWVI*RLWEVIILSFVETCFNSGNPMIVIIFR------NVIYWQMRQTVRRDS----------VGQCKA*GE
Consensus
```

**Tocc.6G0014840.1_exon2**

```
Teleogryllus_occipitalis            ---------------------------------------------------------------------------AVAVAAAAAAAA-------------------------------
Tarbinskiellus_portentosus          ------------------SIE*IRSAY*IPF-----ASCPSYPSVS*RL-----------LSRVGDSISSSGPPLSL----------------------------------------
Acheta_domesticus                   ----------------------ISQSDLLLF-----LSLSIF--*RSRLNTIDYINAAPKNPFRTNIRTLSNPPPLSVSVSLSASCAV-------------------------------
Gryllus_bimaculatus                 --PFSLFVAPANKLHS*ILLLSFDSQN*NTFQRNLELSLPVMDI*STYL-------------KKNISLI---PSDSTEIYIF------FFLQLLI-------LPSRF-----------
Gryllus_longicercus                 HSPQALH*PTLSLLTVCRICQ*ITFLNIITI-----IGLPELKHVSTKLRIITSHNRYITQIFKKNISLI---PPDSTEI*IFSSSNF*FFRHTSVEIPGINFFPCRLYYS*PFGVPAPG
Consensus

Teleogryllus_occipitalis            -------------------------------------------------------AGAEAAGGG--------GGGGG-GGG------------GG---------------GGGGGGGG
Tarbinskiellus_portentosus          -------PRSAD-----------PARHARGG------MQTRARRR-A*RT-SRRAGTGCCG--VFAGERWRPRKSGLGRRRRGQPCAEKYGGRGGGDAAADAFELLPSPIRGAGHGGGD
Acheta_domesticus                   ------PPRSAG-----------RARHARGG------MQTRARRRAA*RT-LRRAGAGCCGGGVVAS---DGTGDG-GEGNRDPHC-----DLGGSDRARRDVRLLGEE-DGVGGGDGD
Gryllus_bimaculatus                 -------RRNSGNQFLPLPPVLFIAVRHACAGRVMQ*SAQTKPRRPPVPRTSAPRSSSQPQRPETVCAT*VRARADA-G*QAR------------------------------------
Gryllus_longicercus                 E*CSDPPRRSPGDPPFREPPHL--GVRRNRKGRKPSAQRESRPAQTPGDRHVLRPVAASPSTNPIFCS-------------------------------------------------
Consensus

Teleogryllus_occipitalis            GGGG-----GGGGG---------------------------------------------------------------------------------------------------------
Tarbinskiellus_portentosus          DAGSKDEEVNARLG--------------------KSDE---------------MWL*GI---------QSDQARQEKSGNN------------RSSN*GRKKNVGKRKL**FLVTPF
Acheta_domesticus                   RGAT-----AAS*GRFPVS*TLLRVNR*LFL-----------LLLGLCRHPVC-CWNKR-------YSFSS*IFRRENPLLNTMRGPRESIRCYQSVMNIFHPGDSE*KK---------
Gryllus_bimaculatus                 ----------AAPG----VC*PFNESHILFQKVPFKSRYVTLKSVTNVSDHVLAYQWFIQV*VVTIV**FTSLKF---------CHSLSFII*NFVRIIKWNISNEN------------
Gryllus_longicercus                 --------------------RTFHLNHFM-------------*L*NL*QTYHSMCWSIR----------RSYKFEWSQFFSNSLH*FLVIAYRF*YEISFGL*NGTFQTKI--------
Consensus

Teleogryllus_occipitalis            -----------------------------------------------------------------------------------------------------------------------
Tarbinskiellus_portentosus          LSLIDIFKVFVVVISLRPAASLTSYLRLVANGEFDVVVNSQSQIQNRNQ**YKSVLSPTPS----------FFR*SNQFSVLSNSASH--------------ISPPSAF*I--------
Acheta_domesticus                   --YPNIFCLCVLIKFLCPV------------LF*T--------V*YYHLKTNKLV--KGHILNLKIY*NVFFMHVDSLFSLNHDY-HYYLF**KK*LQFHFCVRWPFSYQLNTRRMCCD
Gryllus_bimaculatus                 --LDNAITVIILI--L*KL------------FFDR--------QNL**LLFMNLLPLTRLFI*VEIFGNIYF-----LYALPISYRHEKIFYFR------------------------
Gryllus_longicercus                 -AIDNAITVIILI--LENYSSI--------DQIFDN--------SNSEFIDFNSSVHPSRNLTLI------------FFALPISYRHEKIFYIR------------------------
Consensus

Teleogryllus_occipitalis            -----------------------------------------------------------------------------------------------------------------------
Tarbinskiellus_portentosus          -----LDVSRILL----------------------------SSF--VAPVAL-------*PLLVTLSQSRVRFLVFRPCAAPTSSGTSSHSQSATDTAFHSAISTCQRLR-------RPG
Acheta_domesticus                   TPLIVDLVSKVMIIAFSVTGNVGRFETCYGHLIVLSAPHSCVFFFM*SPSSSLPRVCLHVRLLLGLRGR-------RPRHRPRRVATAS----------------ARLRLRYLDLPLPR
Gryllus_bimaculatus                 -----*IVSRL-----------RFE--------*SSRFHCTSF----PTGK-------KKIFIGLNGS-------VPVCAPAPVG---------------------------------
Gryllus_longicercus                 -----*IVSRL-----------RFE--------*SSRFHCTSF----PTG--------KKIFIGLNGS-------VPVRAPAPVG---------------------------------
Consensus

Teleogryllus_occipitalis            --------------------------------GGG------------------------------------------------------------------------------------
Tarbinskiellus_portentosus          RRRRIRDAMHLAAGH-WLCRGAVRCG--ALIRGGGGGWSVTSRNMTPG---AAAG*ISA*WATGELGAAITRRPTRLRDLLRD--QVPNRN*TVYAKILISSRFWGPKYFDHDSSSRDHS
Acheta_domesticus                   RRLRIRDAMHLAAGH-WLRRGAVCTAPHAAAAGAGP*RHEI*RQGQPSATPRRAG*ISA*WPW--AGPA--RSGPASAAIRRR-------------------------------------
Gryllus_bimaculatus                 ---------HSADGSGAGRRRLI-----AALAGAGGPAHGHYTDIQPA---RR------------GAA--RRPAAPGVIFRDVTDLPP-----------------PPCEAVHSTPR---
Gryllus_longicercus                 ---------HSAAGR-RRRRRLI-----AALAGAGGPAHGHYADIQPA---RR------------GAA--RRPAAPGVIFRDVTDLPP-----------------PPCEAVHSTPR---
Consensus                                                           * *

Teleogryllus_occipitalis            -----GGGGGG-----GGGGGGGGGGGGDGGGGGGGGGGG-------------------------------------------------------------------------------
Tarbinskiellus_portentosus          LCSIVGVV*GG--------------------------------KTVINVVGKIQLANR-------------------TLSSE----------------------FRSCGPELNRLPAF
Acheta_domesticus                   --SAAGGGGGG-----GGGGGGGGGGGGDGGGDGGGGGGGGGVRHRRATP--------------RPQREPALNRLTQ*NSCSSRLGNWCSGDDYS-------NPK--SETISQK*NNFS--
Gryllus_bimaculatus                 LCTASGRRPGALHHVCGDGVGGGGGGGGSR*RSG----------RRAVSVA---TLCGRWRGRRPRRPTRR*RQTRG---SDEDGDYIKKT*MARTLEKIRCP**VSKQLGPR*QSLD--
Gryllus_longicercus                 LCTASGRRPGALHHVCGDGVGGGGGGGGSR*RSG----------RRAVSVA---TLCGRWRGRRPRRPTRR*RQTRG---SDEDGDYI*KKNMN------------GKSVGEDKMSIASF
Consensus                                *   *

Teleogryllus_occipitalis            -----------------------------------------------------------------------------------------------------------------------
Tarbinskiellus_portentosus          --LKSFH-----LRGSHCLDHNSKVRTFV--W-------------------------YYFCVQR*RLLPLLLQR*GWYWQQFFCS*LESPDFFTLYKYFSK------------------
Acheta_domesticus                   ------------CRD**C*NQN*S------------------------------YQ*SNS*IR*---QVKIITKFGRR*IR---------VTKEFVTKV*GFSNN*NYCRNRVIEEF
Gryllus_bimaculatus                 ------------L*SKVCHNKFAP-I*LVGKWP*HAEIILQLFHST*ILRKIIFITDYYHLK---RT--YIHEGLSCYSRQDFQDGVPSTVYSTSRQEVDKILAAA--------RIISDE
Gryllus_longicercus                 ETASSAHVDRVWINNQRCIITNSPLI*LVGKWRNTP*LILQLFHSTRILIK------*YSLQTNMRT---THEGLSLYSR*DFQDGVASTVHSTSRQEVDKILASA--------RIITDE
Consensus

Teleogryllus_occipitalis            -----------------------------------------------------------------------------------------------------------------------
Tarbinskiellus_portentosus          -------------------NLMSNWTDTSAWHV----------------PVTCL-NGA*F-----------------TMR--------------------------------------P
Acheta_domesticus                   ELSEF*SLKRSILL*VIIRTKVQNLND---------KLLRSTQKRV---------SRSKFLR*---VIIYDL*PHKPAV*--------TTETLAYEMTRLFTDLSQIN------------
Gryllus_bimaculatus                 RLKE-------------IPNQVRDLGDCRSHTVTVTTALRHRPRRLPLPPAAPLSAHS--------LSHRDRSPGPDCLRRHRQCHREREVAAAAAAARCATPRRRALVCIPPRACRARP
Gryllus_longicercus                 RLKE-------------IPNQVRDLGAAVA-------------KRSPSPPPSVI-AHAAFLSLQPHLSPRTPSPTEIALR--------APIAFAAVASAIASAKS------PPQPPQG--
Consensus

Teleogryllus_occipitalis            ----------------------------------------------------------------------GGGGG-----GGGGGGGGG---------------------------GDVQ
Tarbinskiellus_portentosus          GLPASA-------VAKNSKVRRFLERLGFVSSDQWALRDRKLDGYE*YRQEVEK*IPRISTQVR*---KKNLKLG-----------------------------------------GRRN
Acheta_domesticus                   WLQSNV-------LENKSD---------SFGASR----------------HWEQHVP-VTRCLRGP*LTVSGLCGCEELQGAGLLGTGGRLRFVCSDHCITRPPRARGTPMNNT-DGK*R
Gryllus_bimaculatus                 ALRGGA--------AHDTDGRRGGWKRTDVDAER----------------VVP*AIA-L*CDLRT---TESKKRG--RL*EGSD*GSG*----------VTVCVRERMI*RKKVIEGTTA
Gryllus_longicercus                 ALRRAVVLLFAFRLAHAARGRRSAEALRTTQTERGIGRGRTSMRNGSFGAALT*AIA-L*RDLRK---TESKKRG--RV*GGSD*GSG*----------VKRCERGGVFEGENEIEKESD
Consensus                                                                                     *

Teleogryllus_occipitalis            QPVPHRDRSA-NRQ*--------------------------------------------------------------------------------------------------
Tarbinskiellus_portentosus          IITNIRIPTA-*WKFYV*KC*LTI--IKYINNSVYYIYIY--------IYMYV*LI*ENL*NIIKSLKFK**L*TYR*IILEK*MCEER*-Y-----------------------
Acheta_domesticus                   N*FPEFRRKC-DEKIRSWRK----------KKNVDFFGIR----------------------------------SIREIFLKNMLIIDQGK**F*ISL*LELLQTFH*K*EYIGIL
Gryllus_bimaculatus                 GTASVRKKLACSKEPT*GRIWST*SS*N*VRNVQFYLNIA--CGN----FVS------------ESL*F-----TTREMFL----------*F*NISC-----------------
Gryllus_longicercus                 *RKYSRNRLG-AKEISLQKT------SDLRSDLVYLTFVELC*KCPILFEH------------HVCKF-----SVRRFII----------YKGNISLIL----------------
Consensus
```

**Tocc.7G0009600.1**

```
Teleogryllus_occipitalis        AACGCAACGAATTAATTGAAATAATTTCAAGCAATTTTCCATTCTTCATTAATTTCTCATGTTCCTAGTCCTTGACCTTCACATCCCCATGAGAAATTTCGCCATCAATAACCATCAACT
Acheta_domesticus               AAAGCAATAAATAATTTGAAATAATTTCAAGCAATTTTCCTGTCTTTATTTATTTGGCATATTCCTGATCCTCGATCTTCACATCCCCTTGAGAAATTTCGCAATCAATAACCATCAACT
Consensus                       ** ****  *** * *************************  **** *** ****  *** *****  **** ** ************ ************* *****************

Teleogryllus_occipitalis        TTGCTCTTTCGTTGAGATTAGGTGAGCCAGATGCGATATGGTAATTTCCGATCAAATGATACTTGAATGCAAGCGTCGACAAAACATTACACAATGGTCGACAAAACACATGTTAAAGCC
Acheta_domesticus               TTGCTCTTTCGTTGGGATTAGGTGAGCCTGACGCGATAGGGTAATTTCCGATCAATTGATACTCGAATGCAAGCGTCGACAAAACATTACACAATGGTCGACAAAACACACGTTAAAGCC
Consensus                       ************** ************* ** ****** **************** ******* ********************************************** *********

Teleogryllus_occipitalis        AGGACACTGGTGCTTAAAAGTAGTAATGCGAGAGAACAGTAATGCGAGTTTCTAACGACCATGAAGGCCAAACTGTGAAATAGGGACACACTGTCGACGAACACTCGCACAGATAAATCT
Acheta_domesticus               AGGACGCTGATGCTTAAAAGTAGTAATGCGACTGACCACTAATGCGAGTTTCTAACGACCATGAAGGCCAAACTGTGAAATAGGGACACCCTGTCGACGAACACTCGCACAGATAAATCT
Consensus                       ***** *** *********************  ** ** ************************************************** ******************************

Teleogryllus_occipitalis        GAACGGCATAAATCAAGAAGTCGGATTGCGATACAAGGTTCCAGAATCATACGCTCTAAAATACTAGAAGAATTTTCTAGTCGCGCCACGTGGCTTGGCACGCGACTGGTGTAAAGGCGT
Acheta_domesticus               GAGCGGCATAAATCAAGCAGTCCGATTGCGATACAAGGTTCCAGAATCATACGCTCTAAAACACTAGAAGAATTCTCTA--TGCGCCACATGGCTTGGCACGCGTCT--------AGGGT
Consensus                       ** ************** **** ************************************** ************ ****   ******* ************** **         * **

Teleogryllus_occipitalis        GATGGAGACGGTGGTAGCGGAGATAGCGGCCGCGGTGGAGGCGGCGACGACGGGAATGGCGTCTCTGGTGGTGTTGACGACGATGGCGATGGTGGTGGCGGTGATGGCGGCGGCGGGCAG
Acheta_domesticus               GATGGACACGGTCGTGGCGGAGGCAGAGGC-----------------------------------------------------GGCGATGGTGATGACGGCGGTGGC------------
Consensus                       ****** ***** ** ******  ** ***                                                     ********** ** *** * ****

Teleogryllus_occipitalis        AGGTTTTCTGGTGGCGTCGGTTGTGACGCTGGAGGCGGTGGCGGCAGTGGTGACAGTAACAGAGGCGGCAGCAGCGTTAGTGGCGGTGGCGGTGGTGGCGGTGGTGGCGGTGGTGGCGGT
Acheta_domesticus               -------------------------ACGATGGGGGCGGTGGTGGAGGTGGTGACGGTAGAGTAGGCGACAGCAAC--------------------------------------------
Consensus                                                *** *** ******** **  ******** ***    ***** ***** *

Teleogryllus_occipitalis        GGTGGCGGTGGTGGCGGTGGCGGTGGCGGTGGCGGCGGCGGTGGCGGTGGCGGCGGTGGCGGTGGCGGTGGCGGTGGCGGTGGCGGTGGCAGTGGTGGCGGTGGCGGTGGCGGTGGCGGC
Acheta_domesticus               ------------------------------------------------------------------------------------------------------------------------
Consensus

Teleogryllus_occipitalis        GGTGGTGGCGGTGGCGGTGGCGGTGGCGGTGGCGGTGGTGGTGGTGGCGGTGGCGGTGGTGGCGGTGGCGGTGGTGGTGGCGGTGGTGGCGGTGGCGGTGGCGGTGGCGGTGGTGGCGGT
Acheta_domesticus               ------------------------------------------------------------------------------------------------------------------------
Consensus

Teleogryllus_occipitalis        GGCGGTGGCGGTGGCGGTGGCGGTGGCGGTGGCGGTGGCGGTGGCGGCGGTGGTGGTGGTGGTGGTGGCGGTGGTGGCGGTGGCGGTGGCGGTGGCGGTGGCGGTGGCGGTGGCGGTGGT
Acheta_domesticus               ---------------------------------------------------------------------------------------------------------------------GGC
Consensus                                                                                                                                             **

Teleogryllus_occipitalis        GGCGGTGGCGGCGTCGGCGGCGGTGGCCGCAGGCGAGGGCGGACTCGCTTTGGTAGCTGACCCGAGGCGGCGGAGGAAGGG-----GCTGTGATCTAGTGATCATATAGGCGGCGATGTT
Acheta_domesticus               GGCGGCGAGAGCGAGGGCGGGGGCAGCCGCAGGCGAGGGCGGACTCGCTTTGGTCGCCGGCCCCACGCGGCGGAGACAGGGGGGCAGCAGTGGACTAGTGATCACG-------------
Consensus                       ***** *   ***  ***** **  *************************** ** * *** * *********  ****     ** ***  **********

Teleogryllus_occipitalis        GCTGTTGGTTGCGATGGAGGAAGTGGTAGTGGCGACAA--CTAAGTTTCTGGAAATAAGTATCTAGGGGACCGAGTTGATTATATTGCCGTTGGCGTAGATAAAACAATCAAGAAACATA
Acheta_domesticus               ---------GTCAATGGGGGAAGTGATGGAGGCAACAAACCACAGTTTCTGGAAATCAATATCTCGGGGATCGAGCTGATTATGTTGCTGTCGGCGTAAGTTAAAC-------GAACATA
Consensus                                  * **** ******* * * *** ****  *  ************* * ***** ***** **** ******* **** ** ******  * ****        ******

Teleogryllus_occipitalis        TCATTTACATTTAAGGAAAATGTTTCTTTTGTGAATTTT-TTATTTTTTCATTAATATTACATTAAAGTGTAATAAACATGGTACACAGACTTACCATAAAATTTTAGTTCACGAAACAT
Acheta_domesticus               TCATGTACATTTGGGGAAATTG--TCTTTAGTGATTTTTGTCGTCCATTCATTAATATTAGATTAATTTGTAATACACATGGTAAATAGACATCCCAT---ATTTCAGCTCAAGAAACAT
Consensus                       **** *******  ***** **  ***** **** **** *  *   ************* *****  ******* ******** * **** * **** ****   **** ** *** *******

Teleogryllus_occipitalis        CTTCGCTT----------------------------------------------------------------------------------------------------------------
Acheta_domesticus               ATTAGCTTGTGGGATTGGATGGTTTCAAACAGAACAGCCTTATTCTAATAATTTAATTTTAATTTGACATGTTAATTATTCTAATGAGGTGTTTTACACTTTTACATATTTGTTATCGTT
Consensus                        ** ****

Teleogryllus_occipitalis        ------------------------------------------------------------------------------------------------------------------------
Acheta_domesticus               TCTTCATTTTCTACGGAAAAGGTTAATTCCAAGAGAAGCAGAAGAAAAATAAGAGAGGGAAATGACAAAAAAGTGGGAGAAGGATTGGGAAAAACACAAAATAAAAAAGACGAATTTACA
Consensus

Teleogryllus_occipitalis        ------------------------------------------------------------------------------------------------------------------------
Acheta_domesticus               TTCTCCATTAAAAATTACACTTCGTACCAATTTACCAATTCTCTTTCGACATGTAACAGCTGATGATAATGGTAATGACTGATAGAAGGATGATGGATGTATCACAATGGCAGACAGGAG
Consensus

Teleogryllus_occipitalis        ------------------------------------------------------------------------------------------------------------------------
Acheta_domesticus               AGAAGTTCAAGAAAATCTACTGCGAACTCTCTTGGATCCACACCAAGACGCACCAGCAAGACGAAAACGACACTGGAAACAAAATATGGTTTGTCGGAGGTGACACGGCCGTTAGTCACA
Consensus

Teleogryllus_occipitalis        ------------------------------------------------------------------------------------------------------------------------
Acheta_domesticus               ACGATTGCAAATCCTTTTCCTCGGTGACTCGCGTCACCACCCACCGCGCGGGCGGGCGTGGAAGTGACGGAGCAGCGCGGCGCCGCGGAGCGGGAGGGGGAGGAGGGTGGGGTGGGTGGG
Consensus

Teleogryllus_occipitalis        -----------------------------------------------------------------------------------------------------
Acheta_domesticus               TCAGGAGGGGTGGGGAATGGCGGGGGAGGGGGCAGGGCAGGGAAGGGCGGGACGGGGAGGGGTGAGGCGGGACGTCGAGGGGTGGAGGGAAGAAGCGGGGCG
Consensus
```

**Tocc.7G0009600.1_exon1**

```
Teleogryllus_occipitalis          ------------------------------------------------------------------------------------------------------------------------
Acheta_domesticus                 C**IKICILLLQN*MHKTSHTFLCLKSFNIRILFKISLLKQIIHLQHIITYRLNSLTLIY*KKILLLQITHL*LPT*SQIISTLIPQIQTLLQNLLPPLLMGINNNRNINITLLNWNSKH
Consensus

Teleogryllus_occipitalis          ------------------------------------------------------------------------------------------------------------------------
Acheta_domesticus                 *RDNKIFLITF*LFIILT*LVFLKLILTLKKNDI*KVITFIEVIRFLIMHLEE*Q*KSRILYIINKHFFIFLTSSKLSSKKRV*TS*ECNEKCLDPLQRVKFHDDRAFFQEMYTPPK*FK
Consensus

Teleogryllus_occipitalis          ------------------------------------------------------------------------------------------------------------------------
Acheta_domesticus                 IHLIKLQHLA*KVDARMVSSSHGSLKLLSQELS*KLR*KEKAINNLK*FQAIFLSLFIWHIPDPRSSHPLEKFRNQ*PSTLLFRWD*VSLTR*GNFRSIDTRMQASTKHYTMVDKTHVKA
Consensus

Teleogryllus_occipitalis          --------------------MK--------------------------------------------------------------------------------------------------
Acheta_domesticus                 RTLMLKSSNATDH*CEFLTTMKAKL*NRDTLSTNTRTDKSERHKSSSPIAIQGSRIIRSKTLEEFSMRHMAWHASRVMDTVVAEAEAAMVMTAVARWGRWWRW*R*SRRQQRRRRERGRG
Consensus                                             **

Teleogryllus_occipitalis          ------------------------------------------------------------------------------------------------------------------------
Acheta_domesticus                 QPQARADSLWSPAPRGGDRGAAVD**SRSMGEVMEATNHSFWKSISRGSS*LCCCRRKLNEHIMYIWGNCL**FLSSIH*Y*INL*YTW*IDIPYFSSRNILACGIGWFQTEQPYSNNLI
Consensus

Teleogryllus_occipitalis          ------------------------------------------------------------------------------------------------------------------------
Acheta_domesticus                 LI*HVNYSNEVFYTFTYLLSFLHFLRKRLIPREAEEK*EREMTKKWEKDWEKHKIKKTNLHSPLKITLRTNLPILFRHVTADDNGND**KDDGCITMADRREVQENLLRTLLDPHQDAPA
Consensus

Teleogryllus_occipitalis          -----------------------------------------------------------------------------------------------
Acheta_domesticus                 RRKRHWKQNMVCRR*HGR*SQRLQILFLGDSRHHPPRGRAWK*RSSAAPRSGRGRRVGWVGQEGWGMAGQGAGQGRAGRGGVRRDVEGWEGRSGA
Consensus
```

**Tocc.7G0009600.1_exon2**

```
Teleogryllus_occipitalis    ------------------------------------------------------------------------------------------------------------------------
Acheta_domesticus           VNK*KFAFSFSKIKCIKLPTLSSVSNLSI*EFYSKSPF*NKSYISNI*SHIV*ILSL*YIKRKSFYFKLHIFNYQHSHKLFLHLSRKYRLYYKTFFLHF*WALITTEI*I*HY*TGTLNT
Consensus                   

Teleogryllus_occipitalis    ------------------------------------------------------------------------------RFSG-----------GVGCDAGGGGGSG---------DSNRG
Acheta_domesticus           NATTKYF*SLFNFS*Y*HSLYF*NSS*P*RKTIFKRL*HL*K*YDFSSCIWRSSNKNQEFCISSTNIFSYF*HHLNYHRRSGYRQVKNVMRNVWIPCNA*NSMMIGHFFKKCTPRQSNLK
Consensus                                                                                                             * **              * *      *          **

Teleogryllus_occipitalis    GSSVS-------------------------------------------------------------------------------------------------------------------
Acheta_domesticus           FTSLSYNTWRKKSTLAWFLLLMVL*NF*VKNSRRS*DEKRKQ*II*NNFKQFSCLYLFGIFLILDLHIPLRNFAINNHQLCSFVGIR*A*RDRVISDQLILECKRRQNITQWSTKHTLKP
Consensus                     * *

Teleogryllus_occipitalis    ----------------------------------------------------------------------------------------------GGGGGGGGGGGGGGGGGGGGGGGGGG
Acheta_domesticus           GR*CLKVVMRLTTNASF*RP*RPNCEIGTPCRRTLAQINLSGINQAVRLRYKVPESYALKH*KNSLCATWLGTRLG*WTRSWRRQRRRW**RRWHDGGGGGGGDGRVGDSNGGGESEGGG
Consensus                                                                                                                   ******* *  *   ***   ***

Teleogryllus_occipitalis    GGGGGGGGGGGGGGGGGSGGGGGGGGGGGGG-------------------------------------------------------------------------GGGG------------
Acheta_domesticus           SRRRGRTRFGRRPHAAETGGQQWTSDHGQWGK*WRQQTTVSGNQYLGDRADYVAVGVS*TNISCTFGEIVFSDFCRPFINIRLICNTHGK*TSHISAQETY*LVGLDGFKQNSLILII*F
Consensus                       *    *        **       *  *                                                                         *  *

Teleogryllus_occipitalis    -------------------------GGGGGGGGGGGGGGGGGGGGGGGGG----------------------------------------------------------------------
Acheta_domesticus           *FDMLIILMRCFTLLHICYRFFIFYGKG*FQEKQKKNKRGK*QKSGRRIGKNTK*KRRIYILH*KLHFVPIYQFSFDM*QLMIMVMTDRRMMDVSQWQTGEKFKKIYCELSWIHTKTHQQ
Consensus                                            * *           *     *   *

Teleogryllus_occipitalis    ------------GGG---------------------GGGGGGGGGGGGGGGGGGGGGGGGGGGGGGGGGGGGGGGGGGGGGGGGVGGGGRRRGRTRFGS*
Acheta_domesticus           DENDTGNKIWFVGGDTAVSHNDCKSESSVTRVTTHRAGGRGSDGAARRRGAGGGGGWGGWVRRGGEWRGRGQGREGRDGEG*GGTSRGGREEAG------
Consensus                               **                       ** *  *     * ***** **    **   * * *  *  * * **   ***
```

**Tocc.7G0010080.1**

```
Teleogryllus_occipitalis      GCGTCAACTGCGAAGGCCATTAGTCCTGCAACTGTAGTGCGTATTTTTGCTCGAGA---------CGTGGCGGTGATCTTCCCTCG--------------------------CTGGGTGG
Acheta_domesticus             AATTCATATTTGAATAGAAATAGTTATATATTTATAATATAATGTAATAATTATTCCCGAGGTATCGGATCGTCGCGCTGGCATCGAGCCCTATCTTAGATACACGAGGTCCTTGCGGAG
Gryllus_bimaculatus           ----------TAAAATTAATTAGCCAAAGGTCAGCCGCGCGCTGCACCGCGTAGCC---------CCCGTCGCTGCGTCCCCGCCGC-----------------CAGGGCTTCCGAGAAT
Consensus                                 **    * ***                                          *    **  *      *  **                            * *

Teleogryllus_occipitalis      CGAGACCCGGAC------------------------CCGATCTACCGCCA--GCTAGCGGG-----------------------------------------------------------
Acheta_domesticus             AGACGTGCGATCAACATTGAACAGAACTATATCTATTCTACACGACGTTATCGCTAAGGAGAGGGTAAATGCGGATTACCTGCTATCTTGGAATTACCCATTTGTTTGTATGGAGATATA
Gryllus_bimaculatus           CGACGTTCAGCC------------------------CCGTTTCGCCGCGAGCGCTAACGG------------------------------------------------------------
Consensus                      **    *   *                         *       **  *  ****  *

Teleogryllus_occipitalis      --------------------------------GTTTTCCCCAACCAATAT----------------------------------------------------------------------
Acheta_domesticus             TTTGTGACACATAATGAGTGACCACAGAAAATGACTTTTCGAGTCAGTTCACTGAGACTAAAGCAGGCCTTGTAGTTCCGTTCGTATCCAATGAGAATGTCCACCAAGATCTATCAGTAG
Gryllus_bimaculatus           --------------------------------------CTCGGCCAGCGCGCCG------------------------------------------------------------------
Consensus                                                                 **

Teleogryllus_occipitalis      ---------------------------------------------------------------------------------------CTCCCACCAGGGCGGCACCGCCCAGCCGGGACA
Acheta_domesticus             TAGCGCATTCATCCATATTTTCTGTCACAGAGTCTCTACTCAGCAGAAAACTGAAGCCTCGAAACAGTCGATTTTCGTTCCCAACTTTCCCCCCAAACCCAGTCCCGCCCACTGTGCGCC
Gryllus_bimaculatus           ------------------------------------------------------------------------------CCGCCATCACCGCCGCCAGCGCAGCCGCGCCCCGTTTTTGCA
Consensus                                                                                               ** * *   * *   *****        *

Teleogryllus_occipitalis      AGCTCTA-------------------------------AGCTCCCGACTG----------------------------------------------------------------------
Acheta_domesticus             CGGGCGG---AGACCTCCCTGCACTTCAAAGGAGCGC-TGCCCCGGCCGGCGCCCCTCTGCCCGCCGAGATGGTGGCTTGTAGTGTGACCTTTCGAACATGATCAGGGTCTTCCGGGAGA
Gryllus_bimaculatus           TAATCCGCGGAGTCGCCCTGGTACCTCGCAGCCTCGCCTGCCGTTGCCAG----------------------------------------------------------------------
Consensus                         *                                  **    * * *

Teleogryllus_occipitalis      -GCCTACCGGCATGT---------------------------------------------------------------------------------------------------------
Acheta_domesticus             CGTTTGTGGGCCCGCGCGGAGCGGTGCGAGCGCGGAGGTCTTCCACTTAGGCGAGGGGACGGGACGGGACGGGAGGGACGGGACGGGACGGGACGGGAGGAGGGCTCCGCCCAGAAGCGA
Gryllus_bimaculatus           -GCTCGCGAGCCCGTGCCCG---------------------------------------------GAGACAATAGGAGCGG---------------------------------------
Consensus                      *       **  *

Teleogryllus_occipitalis      ------------------------------------------------------------------------------------------------GTTAAGTTCCAGAACAT-------
Acheta_domesticus             CTCCTCGCCCGGCGCGGCCGCCGCCCAAGATGCGCTCGCACCGCGATCTGTCTCCGTCCCGTTACATCGGCCGCCGCCGCCGCGCGCGACACGCCAAACGCATTACGGGGCCCTCGCCGC
Gryllus_bimaculatus           --------------------------------------------------------------------------CGTAATCGTACCCGCCAGGCCGGTCACGTTCCTGGACAC-------
Consensus                                                                                                           ** * *  *

Teleogryllus_occipitalis      -------------------------------------------------------------------------------------------------GGCATGAGTTTTGTGGCCCTTCT
Acheta_domesticus             GAACGAGCTGCCGATTGTGGATTCTGACAAAACGTCGAGATACGCTGCTTCCGAAAGCGTTCGAGGAGCTTTTTGAAGTGAAACTTCCCGTCCGTGGGGCGACGACGCTTTGCCTCTGCG
Gryllus_bimaculatus           -------------------------------------------------------------------------------------------------GGCACTGGTTTTGCGGCTTTTCT
Consensus                                                                                                      ***        *  * *  * *

Teleogryllus_occipitalis      TCAATTTTG------------------------GGAACA-TTTGCAATGCGTTACGTTTATTTACAGGTGTCTAAG--------------------------------------------
Acheta_domesticus             TCAAATCTATTTCAGTGTGATTGACTGAGATACGGAGTGTCTCACAATAAATGTTACAAACCCTCCGGAGGCTAAGTACCTTTAAACAGTTGAAGCGTTCTAAAAACAGTCAGGTATACA
Gryllus_bimaculatus           TCAATATTG------------------------AGAACATTTTGCAATGCATTTTGTTTGTTTGCAGGAATCT-----------------------------------------------
Consensus                     ****   *                          **     *  ****   *            * **   **

Teleogryllus_occipitalis      -----------AATAAATGTTACCAATGT-------------------------------------------------------------------------------------------
Acheta_domesticus             GGATGTCTTGTAATAAATGTTACGAATTTCTAGGGCAAATAGAGGAGTCGAAAATGATCACTTTTTGTCAATGAACATATAGTGGGAAATTGTTTATGGCGGAGTTTATCAAATGACGGA
Gryllus_bimaculatus           ----------------ATGTTACAAACTT-------------------------------------------------------------------------------------------
Consensus                                     ******* **  *

Teleogryllus_occipitalis      -------------------------------------------------------------------------------------------ACTCACGCAA-------------------
Acheta_domesticus             GTTACATGTTGCATTTGCCCCTGGTTTTAGGCAACCTGGTATAAACAGATTAGAAATTTCCCTGACTGCGTCAGAACGTGACTTGCCGTGCACGTGCGTAAGGCGTTGCATAAAAATCAG
Gryllus_bimaculatus           -------------------------------------------------------------------------------------------ACGCACTTAA-------------------
Consensus                                                                                                **   *  **

Teleogryllus_occipitalis      --------------------TAAGACCGTTACATACAATTTGCCACTGTATGTTCAAAGTAAAAAAGTGTTCCTTCCCGACCCCTCTACGTGCCCTAGAAGTTTGTAACATTTATTATA
Acheta_domesticus             TGTGGGTAGAAAAGATATGTGGAATTCGGTTATATGACATTTGCCACTACATGTTCATTGTCAAAAAGTTATCATTTTCGATCCCTACACATGTTCTAGAAGTTTGTAACATTTATTATA
Gryllus_bimaculatus           --------------------TAAGTCCGTAACGTGCAATTTGCCACTGTACGTTCATAGGCAAAAAGCGTTCCTCGCT-ACCCCTCTAC-TGCCCCACACGTTTGTGATATTTATTCTA
Consensus                                           **  * ** *  *   **********  * *****  *  ******   ** *     * ****  ** **  * * * ****** * ******* **

Teleogryllus_occipitalis      AGATGTA-----------------------------------------------------------------------------------------------------------------
Acheta_domesticus             AGATATT-----------------------CTATATACTGGT----------TTTAGACTTTC---------------------------------------------------------
Gryllus_bimaculatus           CGATATGAATTAGTGAACGGTAGAAGTTTCTCATATTCTAATAATTTAATACGTTATAATTTCTTGAGAAAATTTTTTAGGTCAAAATCACTGTAATGTCGTGGTGCTCTGTTGAATCAT
Consensus                      *** *

Teleogryllus_occipitalis      ------------------------------------------------------------------------------------------------------------------------
Acheta_domesticus             ------------------------------------------------------------------AACTAGTTACAGGTTTTAAAT---------------------------------
Gryllus_bimaculatus           TAAAACCACGGCATTTCAATAGTTCAATACAACTACGAAAATGTATTGACACTGAAATATAGCCAAAAATAATCAGTTTTTTTAAATGTATAAAATGTCACTTATACCGTGATGGCTTTC
Consensus

Teleogryllus_occipitalis      -------------------TTTATTTATTGGAGTTTAGAGGCACATAAACTACGGAGGTCATTCAGACTCATTGATGAATCGCTGATACCGATAAAGACCATGAGCTGACAGAA------
Acheta_domesticus             ------------------TCTCCTGTATT---TTTTATAAATAGGTCGACTACATAAAAATTATGAATATATTTGTAAA--AGAAATATTGGTCCTTACAACATACCCAAAGAATTTCCC
Gryllus_bimaculatus           ACCCCTTCGTATATAGTATTTTTTGTATTGGAGTTTAGAGGCACGTTAACTAGGAAGGTCACTCAGGCTCATTGGTGAGTAGATGATAGCGATAATGTCGATGAGCTGAATAAA------
Consensus                                         *  * ****    **** *   *  *  ****   *              *** * *        *** * *      * *   *  *    **

Teleogryllus_occipitalis      -------------AATGGATAAATGGAGGAATCAGAGGTAGGGGAGATGGTACAACCC--------AGTGAAAACCTACCGGG-------------ATCTCACCGTCTGCTTCACAAATT
Acheta_domesticus             AATGCTTTGCCTCGTTTAATAAAT-ATAAAATGAAAAATGGTGAAAATGGTACGACTTGTAGGACAAGAAAATATCTGGAGAGGTACGACATCAATGTTTCAACGCGTACCACATGGACT
Gryllus_bimaculatus           -------------AATGGAGAAAT---AGAATCAGAGGCAGGAGAGATGGTACAATCC--------TGAGAAATCCTA------------------------------------------
Consensus                                    *  * ****     *** * *    *   * ******* *            *  **   **

Teleogryllus_occipitalis      CCTCGTGAAGTGATTCAATCCGGGCTCTAGTGGTGGAAGGTGTGTATGCATTAAGTCACGAGTCTCCCTCCCCCC--------------------------CCCCGCCCTAAACACACAC
Acheta_domesticus             CCAACTATGAAGAATTAA------------------GAGAACCAAATGAAGAGGAAAATGACTGTCCTTATCCTTATGAACGTTTTGTTCATAGTCTGCCATTATGCCTTAGGTATTGTA
Gryllus_bimaculatus           --------------------------------------------GATGCATTAAGACACGATCCTCC---------------------------------------CCCCACATACCCAC
Consensus                                                                  *** *       * **   ***                                       **  *   *

Teleogryllus_occipitalis      ATCATAATGCTGTAATGGAATGATCATCAAATAAATTATTTTCGTTGTCAGCATCCTTTTTTATCAAA--------GAATTCGAAGTCTACAGCAATGAATAAAAGTAGTATTAATAGAA
Acheta_domesticus             GATGTTAGTCAATGATTAACCAAACATAGACTATGTTGAAATTTCTGTATGCTGTTTCTTTGCCCAAACACGCTTTGAATTCG---------------------------------ATAA
Gryllus_bimaculatus           ATCACAATGCTGCAATAAACGTATCATCAGATACATTGTTTTTATTGTCTGTGTCCT------TCATA--------GAATACGAAGTGTACAGCAAAGAATAAAAGTATTAGTAG-ATAG
Consensus                           *  *    **  *   *  ***     **  **    *  ***   *      *       ** *        **** **                                 * *

Teleogryllus_occipitalis      GGATTTTTTTGCCTTTTTCACTAGCACATAACTTCTC-----TTTAAAACGCATTGTTTTGCTTATTATTTAAGTTAAACAAAAACC--ACTTTCAGACGTTTCTTTTTCACTGTCATTT
Acheta_domesticus             TGGTCCTTTA------TTAACCAATAAATACAATCTCATCTTGTACAAGTGCAT------------TATCCAAGGAAATTTAATTCA--ATTTTCAGTCAT---ATTTTCAATGGAA---
Gryllus_bimaculatus           GAATTTTTTAACCTATTTCACTAACATACAACTTCTCGCAGATATCAAATGCATTGTTTTGTCGAATATTCAAGTTAAAAGAAATCCTTATTTTCAGACGTTTCCTTTTCAAAGGCATTC
Consensus                        *  ***       ** ** *  * * *   ****         **  ****            ***  ***  **   **   *   * ****** * *    ******  *  *

Teleogryllus_occipitalis      CTCCTCCAATAATA--ATACACACAATGTCTAGCATGCACC---CACTCTTAATATTTCAATGTGCTACTACACGTCACAGAGAAGGGGTGCATTTTTTTTTATCCTGAGAAGGTCATCA
Acheta_domesticus             -------AATGAAATGTTACAGATGCTACCTAGTA---------------------------------CTGCGTTAGATAAAATACGACCGCATTTC-------------AACATCATCC
Gryllus_bimaculatus           CTCTTCTAAATATATTATATATACAGCGTCTAGCATACGTCTAGCACTTCGAATGTTTGGATGTAGTTCTGTACGCCACAATGAAGCAG-GTATTTTTTTGGTTCTT----AGGTCATCA
Consensus                            **  * *   ** * *      **** *                                 **       * *    *     * ****               *  *****

Teleogryllus_occipitalis      GCTGTCCAGCCCTCCCCCAAGTACGTGTCCGAATGGGAGTGTTTTTAAGATTCTGCTCTTCCAGAGTTTTGAGCGCTGTTGGGCGACGGAGGGCTGTGTGGGCCGACGGTGGCGCAACAG
Acheta_domesticus             GCGACTC--TTGTCGTGGTAATTTATGTGCAGGT-----CGTTCGAACGAGCCCGTTGTCACTCTG----AAACGTCGTTTAGCGACATTCAGTA----TGACAAGTGATGCGGCAACGA
Gryllus_bimaculatus           GC-------CCTCCCAAGAAGTGCATGTCTGGAGGGGAGTGTTTTTAAGAATCTGCTCTCTCTTAG---------TGTTGGGCGGCGGAGGGCC-----GACCGACGGTGGCGCAACAG
Consensus                     **           *     * *   ***            ***   * **   * * * *  *   *           ***  *** *     *       * *    * **  *****

Teleogryllus_occipitalis      GTCCCCGCCGGCGCCCCCGCGCCCGCCCCCGCCCCCGCCCCCGCGCCCGCCCCCGCGCCCGCCCCCGCGCCCGCCCCCGCCCCCGCGCCCGCCCCCGCGCCCGCCCCCGCGCCCGCCCCC
Acheta_domesticus             CATTACGACTTCA-----------------------------------------------------------------------------------------------------------
Gryllus_bimaculatus           GTTCCCGCCCCCGCCCCCGCCGGCGGCCCCAGCCCCGC----------------------------------------------------------------------------------
Consensus                          ** *  *

Teleogryllus_occipitalis      GCCCCCGCGCCCGCCCCCGCGCCCGCCCCCGCGCCCGCCCCCGCGCC-CGGGCCGAGCCGGGCCGCTCAATGCCTGCGCCGGAAGTGCTTGAAATAGCCGCGTTACGGGCACATCGAATC
Acheta_domesticus             ----------------AAGGATCAGAGGTGTTAACGACCGACTCGTC------------AGGGCGCTTAG----AGCATCGGAAACAACGGAGACTTCCAC-------------------
Gryllus_bimaculatus           ---------------------------------ACCGCCGCCGCGTCGCGGGTCGTGCCGGGCCGCTCAATGCCTGCGCCGGAAGTGCTTGAAATAGCCGCGTTACGGGCACATCGAATC
Consensus                                                       *  **  * ** *             ** **** *      **  *****      ** *   ** *

Teleogryllus_occipitalis      TCGCGCTCGCCGCCATGCCATGGCGCTGCGTTCCAGGGCTA---------------CTTCGCACTGCCGCTCTACTC-------------------------------------------
Acheta_domesticus             ---------------TGGAGCGACGCCATGCATGAAAG----------------------------------------------------------------------------------
Gryllus_bimaculatus           GTGCGCCCGCTGCCCTGCCACGGCGCTACGTACCAGAGCTGCTTCGCTCCGCGCCCCTTCGCACGGCCGTTTTACTCAGCGGGGAATATTGTAGCGTCACGTTGGCTGTCCTGCGTGGGG
Consensus                                    **    * ***   *    *  *

Teleogryllus_occipitalis      ------------------------------------------------------------------------------------------------------------------------
Acheta_domesticus             --------------------------------------------------AATAGGAGATGCAGGATA----------------------------------------------------
Gryllus_bimaculatus           GGAGAACGCGGAAACCTCTCATATCTGAGAACTATAGCTAGCAGCCATGGAGTTGGGGGTGACAGATATTTTTAGAGCCAGGTAGACACTTGCTTGTATTGCAGTATGTCACGTGGGGTT
Consensus

Teleogryllus_occipitalis      ---------------------GTCAACT--------------------------------------------------------------------------------------------
Acheta_domesticus             ---------AGAATGAATGTGGTTG-------------------------------------------------------------TGGAGGCGGTGTGGTAGAGGGCACTGGCATGGCA
Gryllus_bimaculatus           AAGCCGGCGAGTATAAATTGGGCCGACTCGCTCATGCCACCGGTTTTTTTTTCTTGTAAATCTCGCGCTTTGCGCAAAAAATCACGTCGAGGCTGGTGTGTATTCGTGCATTGCACGGGTG
Consensus                                          *

Teleogryllus_occipitalis      ------------------------------------------------------------------------------------------------------------------------
Acheta_domesticus             ACGACAGCAGCAAAGGTTGCGGCGGAG-------------------------------------------------------GGTTGTGAATGGCAATGAAGGG-------AGGCGGAGA
Gryllus_bimaculatus           ACAAAGGTCACGAGTGGCGCGGTAGGAGCGTGCACTTGAGGCAGGACATTCAGGTACATGCGGTGTATCCACACGATGGTACGATCGCGGCTGGCGGTCGGGAGCTACTGTAAAGTGTGT
Consensus

Teleogryllus_occipitalis      --------------------------------------------------------------------------------------AATATTTACTTATTATACAGAGTGATCAAGAGAC
Acheta_domesticus             ATGGGGC--------------------------------------------------------------------------------------------------GGATGACTAAGAGA-
Gryllus_bimaculatus           ATGGGGTAGTCCATACATTTTTAAATTTTGTAAATAAATCTGTATATACTTTTAAATTGTCTGTGTGCTTAATTCCACGATGCCACAATATTAATTTATTATGCAGGGTGATCAAAAGGT
Consensus                                                                                                              *  ***  ** **

Teleogryllus_occipitalis      ACTTTGCCTACGAGCGGGTGGCATTTTT-TCGAAAGGTTAATGTATTTG---TTTGCCTAAAGTCAAATTTACGAAGCTGAGATCGGTGTTGTAGTAGGGGTTATTACGAAACA------
Acheta_domesticus             ------------GGTGAGTGGTAGGGGTAAAAGAAGGTAGCGGGGACTGGATCATGGTTATGGGAAGGTATGGAAAGTAGAGAAAGATAATGAAGGAGGTCGGACGTCGAAGGTGACGAT
Gryllus_bimaculatus           ACTTTTCCTACGAGCGAGTGCTATTTTTCTCGAGACGTTAATGTATTTG---TTTGTCTAAAGTCACATTTACGAAGCCGAGGTCGGTCTTGTAATAAGAGTGATTACGAAACAA-----
Consensus                                  * * ***  *    *      * **     *    **      **  **  *  *  * *   ***  ***   * *  ** *  * *    *   ****

Teleogryllus_occipitalis      ------------ATTTCTAATAATGAACGATCT-------CTAATAATTTCTAAT-----------------------------------------------------------------
Acheta_domesticus             GGCGATGGGTTCAGGTCGAGGAATACAGGAGGGAAAAT--ATAATGAAGTTCGGC-----AATAAGAATAAGAAAAGA----------------------------------ACTAACGA
Gryllus_bimaculatus           -----------GAGTTCTAGTAACGCATGGCTTACTATTAACAACAATTTTCAGTTTCGTTATGTAAGTAATATAAAAGCACATCTATGTCCTCGTATCTATCACTATCAATATCATCAG
Consensus                                 *  ** *  **   * *             **  *  *

Teleogryllus_occipitalis      --------------------AATAATCGAGTTCTAATAATGAAT----------------------------------------------------------------------------
Acheta_domesticus             CACGCCAGACGTCTGTGGAAAAAGTTCAGGACGGAAGAATGCGC-----------------------GAAGTGAGTGAGAACGTGAA---------------------------------
Gryllus_bimaculatus           TATATCACATTCGCAGAGAGAATAGTCAAATATAAACAAAAAATATAGTTCATTGGAATTTTTATGGGAAATGGGTAAAACTTTGAATGATATTGGTAGTGTAACATCTTTTACAACAAG
Consensus                                         **   **       ** **

Teleogryllus_occipitalis      ------------------------------------------------------------------------------------------------------------------------
Acheta_domesticus             --------TATTGATTCCGAGCGAGTGAACTACTTAGGTCGTAATCCCGT----------------------------------------------------------------AGGTGT
Gryllus_bimaculatus           GGATTAGTTATCAAACTCGGCCTAACGACTTTATTTGATAATTCTTCAATCTCTGTTTTTTAATTTGTTGCGTCTTTGCAACACGTCTCTTAACCCGTTTTCGGCACGGGCAGATTGCAT
Consensus

Teleogryllus_occipitalis      ------------------------------------------------------------------------------------------------------------------------
Acheta_domesticus             AGTGCATGCCATATCCCTGAAGT----------------------------------------GTCACTTCCCACTGAAACGTAAAGCCCCTGAAACGTAAGGACTT---ACCCGTA---
Gryllus_bimaculatus           AATACGTACCGAATATCTGGAGTACTCCGATACGTTTTAGCCTTTTTGTCTCGTTACGTCGCAGTCCTTTCTTCCCTTCGGTCAAAGGATCGACGCCGTTAGTACTCGGGATTCGCAGAT
Consensus

Teleogryllus_occipitalis      ---------------------------------------------------------------------------GGCT-----------------------------------------
Acheta_domesticus             ---------------------------------------------------------------------------GGCTTTGCGGGGATTTTTCTCCTCAAGGTATCTCTCTCAATGA--
Gryllus_bimaculatus           CGCCATCCGTCCAGGAATCCGGGGTGCCTCGAGTTGCTAACTTCCTTGACTTCACGGAGCATAGTTAAGCAATTCGGCTAACGGCGCAGGTTTTGTAATAAGATATCTCACTATACCAAC
Consensus                                                                                                ****

Teleogryllus_occipitalis      --------------------------------------------------------
Acheta_domesticus             --------AGCGCCGCCACAAGT---------------------------------
Gryllus_bimaculatus           GTCCTAATATTATCGTCAAGTGTAATATTTTATTTAATTTTATTCCATTCCTTTTA
Consensus
```

**Tocc.7G0010080.1_exon1**

```
Teleogryllus_occipitalis      ------------------------------------------------------------------------------------------------------------------------
Acheta_domesticus             FLSFRSRNAPLINSYLNRNSYI----------------FII*CNNYSRG----------------IGSSRWHRALS*IHEVLAERRAINIEQNYIYSTRRYR*GEGK--CGLPAILEL--
Gryllus_bimaculatus           -----KIN*PKVSRALHRVAPVAASPPPGLPRIDVQPRFAASANGSASAPPPSPPPAQPRPVFA*SAESPWYLAASPAVARLASPCPETIGAA*SYPPGRSRSWTRHWFCGFSSILRTFC
Consensus

Teleogryllus_occipitalis      ----------------------------------------MLPMYS--------------------------------------------------------------------------
Acheta_domesticus             -PICLYGDIFVTHNE*PQKMTFRVSSLRLKQAL*FR----SYPMRMST-----KIYQ**RIHP------------YFLSQ---------SLYSAEN*SLETVDFRS----------QLS
Gryllus_bimaculatus           NAFCLFAGIYVTN--------LRT**VRNVQFATVRS*AKSVPRYPSTAPHVCDIYSTI*ISER*KFLIF**FNTL*FLEKIF*VKITVMSWCSVE--SLKPRHFNSSIQLRKCIDTEI*
Consensus                                                               *

Teleogryllus_occipitalis      ------------------------------------------------------------------------------------------------------------------------
Acheta_domesticus             PQTQSRPLCARAETSLHFKGALPRPAPLCPPRWWLVV*PFEHDQGLPGD-VCGPARSGASAEVFHLGEGTGRDGRDGT-GRDGR----RAPP----------------------------
Gryllus_bimaculatus           PKIISFFKCIKCH--LYRDGF----HPFVYSIFCIGV*RHVN*EGHSGSLVSR**R*CR*AE*KWRNRIRGR--RDGTILRNPRCIKTRSSPHIPTSQCCNKRIIRYIVFIVCVLHRIRS
Consensus

Teleogryllus_occipitalis      ------------------------------------------------------------------------------------------------------------------------
Acheta_domesticus             --------------------------------------------------------------------------------------RSDSSPGAAAAQDALAPRSVSVPLHR--------
Gryllus_bimaculatus           VQQRIKVLVDRNFLTYFTNIQLLADIKCIVLSNIQVKRNPYFQTFPFQRHSSSKYIIYTASSIRLALRMFGCSSVRHNEAGIFLVLRSSALPRSACLEGSVFKNLLSLSVGRRRADRRWR
Consensus

Teleogryllus_occipitalis      ------------------------------------------------------------------------------------------------------------------------
Acheta_domesticus             ----PPPPRATRQTHYGALAANELPI---VDSDKTSRYAASESVRGAF*SETSRPWGDDALPLRQIYFSVID*DTEC-LTINVTNPPEAKYL*TVEAF*KQS---GIQDVL**MLRISRA
Gryllus_bimaculatus           NRFPPPPPPAAPAPHRRRVAGRAGPLNACAGSA*NSRVTGTSNRAPAALPRRYVPELLRSAPLRT---AVLLSGEYCSVTLAVLRGGRTRKPLISENYS*QPWSWG*QIFLEP-------
Consensus

Teleogryllus_occipitalis      ------------------------------------------------------------------------------------------------------------------------
Acheta_domesticus             NRGVENDHFLSMN-I*WEIVYGGVYQMTELHVAFAPGFRQPGINRLEISLTASERDLPCTCVRR-CIKI-SVGRKDM-WNSVI*HLPLHVHCQKVIIFDPYTCS--RSL*HLL*DILYTG
Gryllus_bimaculatus           -----GRHLLVLQYVTWG--*AGEYKLGRL--AHATGF-----------FSCKSRALRKKSRRGWCVFVHCTGDKGHEWRGRSVHLRQDIQVHAVYPHDGTIAAGGRELL*SVYGVVHTF
Consensus

Teleogryllus_occipitalis      ------------------------------------------------------------------------------------------------------RNKT--------------
Acheta_domesticus             FRLSTSYRF*ILLYFL*IGRLHKNYEYICKRNIGPYNIPKEFPNALPRLINIK*KMVKMVRLVGQENIWRGTTSMFQRVPHGLQL*RIKRTK*---------RGK*LSLSL*T------F
Gryllus_bimaculatus           LNFVNKSVYTFKLSVCLIPRCHN--------------------------INLLCRVIK--RYFSYERV------LFFSRR*CICLSKVTFTKPRSVL**E*LRNKSSSNAWLTINNNFQF
Consensus                                                                                                                           * *

Teleogryllus_occipitalis      ------------------------------------------------------------------------------------------------------------------------
Acheta_domesticus             CS*SAIMP*VL*MLVND*PNIDYVEISVCCFFAQTRFEFDN-----GPLLTNKYNLILY-KCII---QGNLIQFSVIFSMENEMLQMLPSTA-LDKIRPHFNIIRDSCRGNLCAGRSNEP
Gryllus_bimaculatus           RYVSNIKAHL-------CPRIYHYQYHQYITFAERIVKYKQKI*FIGIFMGNG*NFE*YW*CNIFYNKGLVIKLGLTTLFDNSSISVF*FVASLQHV--------------------S*P
Consensus

Teleogryllus_occipitalis      ------------------------------------------------------------------------------------------------------------------------
Acheta_domesticus             VVTLKRRLATFSMTSDAATTLR-----LQRIRGVNDRLVRALRASETTETSTGATPCMKE*EMQDKNECGCGGGVVEGTGMATTAAKVAAEGCEWQ*REAENGADD*ER*VVGVKEGSGD
Gryllus_bimaculatus           VFGTGRLHNTYRISG----VLRYVLAFLSRYVAV----LSSLRSKDRRR*YSG---------FADRHPSRNPGCLELLTSLTS-------------------------------------
Consensus

Teleogryllus_occipitalis      ------------------------------------------------------------------------------------------------------------------------
Acheta_domesticus             WIMVMGRYGK*RKIMKEVGRRR*RWRWVQVEEYRRENIMKFGNKNKKRTNDTPDVCGKSSGRKNARSE*EREY*FRASELLRS*SRRCSACHIPEVSLPTET*SP*NVRTYP*ALRGFFS
Gryllus_bimaculatus           -----------RSIVKQFG*RR-----------------RFCNKISHYTN----------------------------VLILSSS----------------------VIFYLILFHSF--
Consensus

Teleogryllus_occipitalis      ------------
Acheta_domesticus             SRYLSQ*SAATS
Gryllus_bimaculatus           ------------
Consensus
```

**Tocc.7G0010080.1_exon2**

```
Teleogryllus_occipitalis                 -----------------------------VPAG---------------------------------------------------------------------------------------
Acheta_domesticus                        FLSFRSRNAPLINSYLNRNSYIFII*CNNYSRGIGSSRWHRALS*IHEVLAERRAINIEQNYIYSTRRYR*GEGKCGLPAILELPICLYGDIFVTHNE*PQKMTFRVSSLRLKQAL*FRS
Gryllus_bimaculatus                      --------------------------*N*LAKG---------------------------------------------------------------------------------------
Consensus                                                                *

Teleogryllus_occipitalis                 ---------------------------------------------------------------APAPAP---------------------------------------------------
Acheta_domesticus                        YPMRMSTKIYQ**RIHPYFLSQSLYSAEN*SLETVDFRSQLSPQTQSRPLCARAETSLHFKGALPRPAPLCPPRWWLVV*PFEHDQGLPGDVCGPARSGASAEVFHLGEGTGRDGRDGTG
Gryllus_bimaculatus                      ---------------------------------------------------------------QPRAAPRSPRR------------------CVPA------------------ARASEN
Consensus                                                                                                         *  **

Teleogryllus_occipitalis                 -----AP-----------APAPAPAPAPAP--------------------APAPAPA-----------PA--------------------------------------------------
Acheta_domesticus                        RDGRRAPPRSDSS-----PGAAAAQDALAP-----RSVS-VPL-------HRPPPPRATRQTHYGAL-AANELPIVDSDKTSRYAASESVRGAF*SETSRPWGDDALPLRQIYFSVID*D
Gryllus_bimaculatus                      R--RSAPFRRER*RLGQRAAAITAASAAAPRFCIIRGVALVPRSLACRCQAREPVPGDNRS---GVIVPARPVTFLDT---------------------------ALVLR-LFFN-----
Consensus                                     **             *     * **                       * *             *

Teleogryllus_occipitalis                 ------------------------------------------------------------------------------------------------------------------------
Acheta_domesticus                        TECLTINVTNPPEAKYL*TVEAF*KQSGIQDVL**MLRISRANRGVENDHFLSMNI*WEIVYGGVYQMTELHVAFAPGFRQPGINRLEISLTASERDLPCTCVRRCIKISVGRKDMWNSV
Gryllus_bimaculatus                      ----------------------------IENILQCIL-------------FVCRNL--------CYKLTHL-----------------IS--------P*RAICHCTFIGKKRSSL----
Consensus

Teleogryllus_occipitalis                 -----------------PAPAPA-------------------------------------------------------------------------------------------------
Acheta_domesticus                        I*HLPLHVHCQKVIIFDPYTCSRSL*HLL*DILYTGFRLSTSYRF*ILLYFL-----*IGRLHKNYEYIC-------------------KRNIGPYN----------IP-----------
Gryllus_bimaculatus                      -----------------PLYCPTRL*YLFYDMN**TVEVS---HILII*YVIIS*ENFLGQNHCNVVVLC*IIKTTAFQ*FNTTTKMY*H*NIAKNNQFF*MYKMSLIP*WLSPLRI*YF
Consensus                                                 *

Teleogryllus_occipitalis                 ------------------------------------------------------------------------------------------------------------------------
Acheta_domesticus                        -------KEFPNALPRLIN------IK*KMVKMVRLVGQENIWRGT----------------TSMFQRVPHGLQ----L*RIKRTK*RGK*LSLS------------------L*TFCS*
Gryllus_bimaculatus                      LYWSLEAR*LGRSL-RLIGE*MIAIMSMS*IKMEK*NQRQERWYNPEKS*MH*DTILPPHTHITMLQ*TYHQIHCFYCLCPS*NTKCTAKNKSISR*EFFNLFH*HTTSRRYQMHCFVEY
Consensus

Teleogryllus_occipitalis                 ----------------------------------------------------------------------------------------------------PAPAPAPAPAPAPAPAPGPS
Acheta_domesticus                        SAIMP*VL*MLVND*P-------NIDYVEISVC------------CFFAQTRFEFDNGPLLTNKYNLILYKCIIQGNLI---QFSVIFSMENE--MLQMLPSTA-LDKIRPHFNIIRDSC
Gryllus_bimaculatus                      SS*KKS----LFSDVSFSKAFLF*IYYI-YSV*HTSSTSNVWM*FCTPQ*SRYFFGS*VISPPK------KCMSGGECF*ESALS*CWAAEGRPTVAQQVPAPAPAGGPSPAPPPRRGSC
Consensus                                                                                                                    *  *      *

Teleogryllus_occipitalis                 RAAQCLRRKCLK*-----------------------------------------------------------------------------------------------------------
Acheta_domesticus                        RGNLCAGR--------------SNEPVVTLKRRLATFSMTSDAATTLRLQRIRGVNDRLVRALRASETTETS------------------------------------------------
Gryllus_bimaculatus                      RAAQCLRRKCLK*PRYGHIESCARCPATALRTRAASLRAPSHGRFTQ-----RGIL*RHVGCPAWGENAETSHI*EL*LAAMELGVTDIFRAR*TLACIAVCHVGLSRRV*IGPTRSCHR
Consensus                                *   *  *

Teleogryllus_occipitalis                 ------------------------------------------------------------------------------------------------------------------------
Acheta_domesticus                        -------------TGATPCMKE*EMQDKNECGCGGGVVEGTGMATTAAKVAAEGCEWQ*REAENGADD*ER*VVGVKEGS---------------------------------GDWIMVMG
Gryllus_bimaculatus                      FFFL*ISRFAQKITSRLVCIRALHG*QRSRVAR*ERALEAGHSGTCGVST-----RWYDRGWRSGA------TVKCVWGSPYIFKFCK*ICIYF*IVCVLNSTMPQY*FIMQGDQKVLFL
Consensus

Teleogryllus_occipitalis                 ------------------------------------------------------------------------------------------------------------------------
Acheta_domesticus                        RYGK-----------------------*RKIMKEVGRRR*RWRWVQ------------------------VEEYRREN----------------------------IMKFGNKNKKRTND
Gryllus_bimaculatus                      RASAIFLETLMYLFV*SHIYEAEVGLVIRVITKQEF**RMAYY*QQFSVSLCK*YKSTSMSSYLSLSISSVYHIRRENSQI*TKNIVHWNFYGKWVKL*MILVV*HLLQQGISYQTRPND
Consensus

Teleogryllus_occipitalis                 ------------------------------------------------------------------------------------------------------------------------
Acheta_domesticus                        TPD--------VCGKSSGRKNARSE*EREY*FRASELLRS*SRRCSACHIPEV-SLPTET*SP*NVRTYP*ALRGFFSS--------------------------RYL--SQ*SAATS--
Gryllus_bimaculatus                      FI**FFNLCFLICCVFATRLLTR--------FRHGQI---------A*YVPNIWSTPIRFSLFVSLRRSP-----FFPSVKGSTPLVLGIRRSPSVQESGVPRVANFLDFTEHS*AIRLT
Consensus

Teleogryllus_occipitalis                 -------------------------------
Acheta_domesticus                        -------------------------------
Gryllus_bimaculatus                      AQVL**DISLYQRPNIIVKCNILFNFIPFLL
Consensus
```

**Tocc.8G0000430.1**

```
Teleogryllus_occipitalis       TGAGACCCTTGAGTAAATGAGTAGTTTAAATGAGAGTATGGAATAATTAGGCCTTGT------------------GTTTCTGAATGGGATCGTGT------------------------
Acheta_domesticus              TGAGCTTCGTCGAACATTAAGAAGCACGGGCTATAATTTGAGCCTTTAGCACGCTGTTCATGACCGTAAAGTAACCGCTTTTACTAGAGACTGCCTAAAAGTATTTTCTTTTTGGCGTAT
Consensus                      ****   * *     * * ** **        * * * **      *    *  ***                  * ** *    * **  *  *

Teleogryllus_occipitalis       -------------TTGTTGGGCTGCAGCATGGAGTATTAGTGAATAAAATTTGAAATTTTTATAG-----TAGATGCGTTAAATGTTCTGGTGGTGACGTATGGTGGGCT---------
Acheta_domesticus              AGCTCTACAATCCACTGTCAGGTCAGGACTTGGAGCGTACTGCAGTAAAA--CGGAGTCCTTAGGGCGAATTATATATATGAATTATTTT-ATATTAAGTTACGTCAGGCTGACCAATAA
Consensus                                   ***  **      * *****  *     * *****    * * *  ***  *      ** **   * ** * ** *  *  * *  ** *    ****

Teleogryllus_occipitalis       -----------------------------------CGTTCAATCTCATAATGTTTTATTGTAGACCAGATGCGTGGTATGTTCATGTGGTGATGTAACGTAGAGACGTGTTACAGGCTC
Acheta_domesticus              CGTTGAATTATATTATTCAATTAATATCACAGACAGTGCTTAATAGTGCAATACGTTTTTAAAATATATTTGAGGGTTCTATCAAATTAGAAATTTCATTTAATTACACAGTGCAA----
Consensus                                                        * * ***     ***    ** **  *     *  ** * * * * *  *  * *  ** * *  **    **     * **

Teleogryllus_occipitalis       GCTCAACCTCTTAAACTGGTATTGTAGACCAGA--------------------------TGCATTGTATGTTCATGTGGTGCT-----------------------------------
Acheta_domesticus              ---TTGCTTTTAGAGATGACACTATAAAATTGAGAGTGCAATTATGACTATTTTCGCTTTTATATTAGATATTAATACTGTTTTTGATATTCACAAGGGTTTTCTGGTTAGGTAAGTCAC
Consensus                            * * *  *  **  * * ** *   **                          *  ***  ** ** **   **  *

Teleogryllus_occipitalis       --------GTGGCGTGTTTCAGGCTTGCTCAACCTC-----------------------ATAATGTAGTATTGTAGACCAGATA---------------------------------
Acheta_domesticus              ATGCGAAAACAGTACAATTTATGTCGGCACCGTTTCAACAACTCTATTCTGGATCACAAAATAATATGGCATTGCATAAGGAACAATAGAGAAACAGCGATAGCGAAGATGAAGAACTTG
Consensus                                *     ** * *   ** *    **                        ***** * * **** * *    * *

Teleogryllus_occipitalis       --CGTTGCATGTTCA----------------------TGTGGTGAC---------------------------------------------------GTGACGTGTTGCAGGCTCG
Acheta_domesticus              ATCATTGGTTTTCCGCCCAGATAATATTACTTCATAATTGTAATGACATTTCTTATTTTTACCGAGTTCTTGAATTGATAAGTAAGGAAAATATATCGTTTATTACTTGGTACAGATCCA
Consensus                        * ***  * * *                        ***  ****                                                  * ** ** * ***   *

Teleogryllus_occipitalis       -------------------------------CTCAACCTCATAATGT-------------GTTATTGTAGAC---------------------CAGCTGCGTTGCATGTTCATGTGG
Acheta_domesticus              AGACGTAAACAAAAAAATCGTAAAAAGCATGCCTCAATATTTAAATGTTTATTCGAGCGAAATTTGTTGCTGGTGCTGAATATAGATGCGTGCATATAGATGTGACACAGGGTTTAATGG
Consensus                                                      *****  *   *****               ** ***  *                        ** ** *   ** * *    ***

Teleogryllus_occipitalis       TGACGTGACGTGTTGC------------------------------------------------------------AGGCTCGCTCAACCTCATAATGTGTTATTGTAGACCAGCTGCG
Acheta_domesticus              TGGCATGGCACCCCATGGTTTAAGGGTGGGGTGTTAGGATATAACGTAGGCGGAAGAAACAAGAGAGAAGGGTGGAAGGGCGCGGCCAGCTAGGCGAGCGGCGACGCTCCACCGGCCCCA
Consensus                      ** * ** *                                                                   *** **  ** *      *   *  *   *  *** **  *

Teleogryllus_occipitalis       TT--------------GCATGTTCATGT-------------------------------------------------------------------------------GGTGACGTGAC
Acheta_domesticus              TTCGCTTCAAGCAGTCGACGTGTGCAGATCGGAGGCAGCAGCCGGGTGGGACGCGGACCGCGTGTGCGGAGGCTAGGGCCAGCAATGTACTGTGACACAGTGAGACTCAAGTGACGTGTC
Consensus                      **                * *** **  *                                                                               ******** *

Teleogryllus_occipitalis       GTGTTGCAGGCTCGCTCAACCTCATAAAGTAGTATT-------------------------------------------------------GTAGACCAGATGCGTTGCA---------
Acheta_domesticus              GTGTAGGAATTCTGTGGTGCTTCATACGATCGGATTCATTCCATAATTAAGGAGAGGGTTTGGTAGCCGTAGAATCAACATTGCTACTAGTAGTGGGATCTATATATTACATTCATTTAG
Consensus                      **** * *     *     * *****   * * ***                                                        ** *      **   ** **

Teleogryllus_occipitalis       ----------------TGTTCATGTGGTGA------------------CGTGACGTGTTGCAGGCTCGCTCAACCTCATAATGTGTTATTGTAGACCAGTGCGT---------TCATG
Acheta_domesticus              CTATTATTCTTAAGTTTTATTTTGGTGGTGGATTTATTTATCAATAAACTAGCAATAAGTTAAACTCTCTCTCGTCT-----GTGTATTATTGCAACT---TGCATGACGCCACACTAGA
Consensus                                      * **   ******                      *  *    ***  *  *** ***  *       *** ****** *       *** *           *

Teleogryllus_occipitalis       TTCATGTGGTGACGTGACGTGTTGCAGGCTCGCTCAACCTCATAAGTTGTA--TGTAGACCAGTGATTGTATGTTCATGTGGTGACTGTTGTCGTCCATAAGTATTGTAGCAGTTGATGT
Acheta_domesticus              TGCGTGGTATGTGAGGATGTATTGCA-GCTCACTTAACCTCATAATGAGGAATTGTAGCAGATCTGTTGCATGTTGTGGTGGTGAC----------------------------------
Consensus                      * * **    **     ** ** ***** **** ** **********    * *   *****    *     *** *****    ********

Teleogryllus_occipitalis       TCAGGCGTGACGTGTTGCAGGCTCGCTCAACTTCATGCCGCGCACGCTGCGGCTGCTGCGCGACGGCGGCGCGGAGTTCCGGCGCGCCTACGTCACCACGCCCATGTGCTGCCCGTCGCG
Acheta_domesticus              ---GCGGTGATGTGTTGCAGGCTCGCTCAACTTCATGCCGCGCACGCTGCGGCTGCTGCGCGACGGCGGCGCGGAGTTCCGGCGCGCCTACGTCACCACGCCCATGTGCTGCCCGTCGCG
Consensus                         *  **** *****************************************************************************************************

Teleogryllus_occipitalis       CTCGTCGCTGCTCACCGGCATGTACGTGCACAACCACCACGTCTTCACCAACAACGAGAACTGCTCCTCGGCCGCGTGGCAGGCGCACCACGAGACGCGCTCCTTCGCCACCTACCTCAG
Acheta_domesticus              CTCCTCGCTGCTCACCGGCATGTACGTGCACAACCACCACGTCTTCACCAACAACGAGAACTGCTCGTCGGCCGCGTGGCAGGCGCACCACGAGACGCGCTCCTTCGCCACGTACCTCAG
Consensus                      *** ******************************************************************* ***************************************** ********

Teleogryllus_occipitalis       CAACGCCGGCTACCGCACCGGTCAGTACCAGTA------------------------------------------------------------------------------------------
Acheta_domesticus              CAACGCCGGCTACCGCACCGGTCAGTACCACCAAATACTATAGAGATAATGCTATTAGTTCATTTCAGTAGCACCATTACACTTCGAGTAATTGCAGACCAAACTTGTAACTGAAAGTTT
Consensus                      ******************************  *

Teleogryllus_occipitalis       -----------------------------------------------CCAGTAGCAGAACTAGCGGCCG------------------------------CTCCCCGACCCCAACCCC
Acheta_domesticus              TGCCGGGAGTATGCAACAATGAATTTGTTTGCAGCTGCCCGTAGCATCCTTGGTGGTAGAAACAGTGATCGTAGTGAACATGGTAGGTGATTATGGTAATGCTTTTCTTGTCACATGCCT
Consensus                                                                      ** * ****  ** *  **                              *  *    * **  **

Teleogryllus_occipitalis       GCCCCAACCCCAACACCAACCCCC-----------GCCCCCGCCCCCGCCCCCGCCCCAGCCCCAACCCCAACCCCGAACCCGACCCCA-----------ACCCAACCCAACCCAACCCC
Acheta_domesticus              ATCCCGCTTTTAATACGGGTTATCCCAGAATATTGGGGGCTGACTATCGTTTGGCCTCAATCGCCATTAGAA---TGAACTAAAAACCAATAAATTAGCCATCTAATCCATCAGATCTGT
Consensus                        ***      ** **       *           *   * * *         *** **  * * *    **    ****   *   ***            * * ** *** *   * *

Teleogryllus_occipitalis       AACCCCAACCACAACCACAGCCCCAGCCCCA-------------GCCCCAATCCCAATCCCAATCCCAATCCCAACCCCAACCCCAACCCCAAC-------------------------
Acheta_domesticus              ATATCTGATCAGACTTGTATATCCAACAGCAATGAAAAAATGAATGTGTTAAATCTACTCCTCTCACAATGCCCAACTCAAAGTGTAGCAGTTACAGTGTGTTTGCCGTTACAGCCTGCT
Consensus                      *   *  * ** *     *   *** *  **              *    **  * * ***     * *  ****** * **    * *    **

Teleogryllus_occipitalis       ---------------------------------------------------------------------------------------------------------------------------
Acheta_domesticus              ACTGTTTTACGTCTGGTAAACACATATTGTATCCTCATGAAAATATTTTTAAAATATATGTAACATATATTTTCTCTGTACATGTTTAATGACACCACTTTTCAACACACGATAGTATGT
Consensus

Teleogryllus_occipitalis       -----------------------------------------------------------------------------------------------CCCAACCCCGACCCCGGC
Acheta_domesticus              ACCGCAATGAAAAATATTGGCTTGCGGCTGTGAGTCTAGCATGGCACGACTGTATTTATTGAACGTCGTTGATTAGTCCCCATGTGTGTACATCGTGCATCACGTAACCTTCACTTCTGC
Consensus                                                                                                                        *  ****   **  * **

Teleogryllus_occipitalis       CCCAACCCAAACCCCAACCCCAACCCCAACCCCAACCCCA---------ACCCCAACCCCAACCCCAACCGCAACCCCAATCCCTAACCCC-----------------TAACCTCTTAAC
Acheta_domesticus              ACCTCCTTGAA---TAGCGCGAAAGGCGATTTCATCTCCACAAGGCAAGTGCTCGGTTTTGATTTGTGAGGCGAGCGCTTTTCCGATCGCCGTATAACGTGACGAAGGCAACATCTTGCA
Consensus                       **  *   **    * * * **   * *   ** * ***           * *       *         ** * * *  * ** * * **                 *** ****

Teleogryllus_occipitalis       TCTTAACTTACCCCAAT-----------------CCTAACCCCTAACCC-------------TTACCCTTAACC---------ACTAACCTCTAACCCTAACCCCTAACCCTTAAAC
Acheta_domesticus              AATCACCTGCCACCGGCGCGGCCTTGAGTGATTTGTCTGAGCTCCAGTTCGGCGCGAGGAGGCACTCGCCTTGACTGACGGGTTTGGCGAGCCGCGAAAATATCCTCCCGGGCGTTGAAG
Consensus                        * * **  * **                     ** * * * *   *               *   **** **             * * ** * **       * **    * ** **

Teleogryllus_occipitalis       CCCACCCCGACCCCAACCCA-----------ACCCCAAC-----------------------------------------------------------------------CCCAGCCC
Acheta_domesticus              TCTCCGCGTGTCCCGGCCGACGCGACCGGGCGCTGCAACTGGTTTTATTAGAGCAGTGACTCGCAGATGGCTGCTGCCGGGCTTGTGCACAGGTGGGCAGGAATGTGGGTGGCCCGGCGC
Consensus                       *  * *    ***  ** *             *   ****                                                                         *** ** *

Teleogryllus_occipitalis       GGCTCGCGA-----------------GGGAGGTTGGGTTGGGAGG--------------------------------------CTCGATTTATGCCTGGCTCGCGGGTCGGGT----
Acheta_domesticus              GACTGGCGATCCGTGGGGATCGTGCCCGGGCGAGGGGGATTTGGAGGGCGTTGTAATCCGCTGGGGCCGCCTTTGATCCCGCAGCCCCTGGGCGCCCGCCCAGGCAGTCGGTGCGGCGGCT
Consensus                      * ** ****                  *** *    ** ** *****                                       ** *       ***  *    *   ***   **

Teleogryllus_occipitalis       ------------------CCGTTTTTTGTTG-----------TTTTTTTGTTCGATTTTCTGTTTTTT--------------GTTTTCATTTTTTTCTTAGTGATTCCGGTCGCATC
Acheta_domesticus              GAAACACCAAAGCGTCCCATCGGTTTTTATTACGGTTTCAACATTTATTTTATTTTAATCGTACCTTATCTTATGCAATTTTACAAGTTAATATTTGTTCGTATGACGTTCTGACACACC
Consensus                                         ** ***** **              ** **** ***  **  *    ** * *                ***    **** *** **      * * * * ** *

Teleogryllus_occipitalis       AGCAGCTAA--------------------GGTCATTCCTGTAAGTGTAGTGCATGTTTTTCTCGAACCGTAGGGGTGATCTTCCCCCGCTGGGTGG----------------------
Acheta_domesticus              ACGAGTCAGAGAAAACCGGTGTGAACAATCGAGCACCGAAGGAAATAAGGAAGAAGATCTTCCTGAATAAT-------ATATCCATATAATTGATTGTAATAACAACAACAATAATAATA
Consensus                      *  **  *                    *  **      * ** *    *    * * * ***  ***    *       ** * *       * * * *

Teleogryllus_occipitalis       ----------------
Acheta_domesticus              ATAATAATAACAACAAC
Consensus
```

**Tocc.8G0000430.1_exon1**

```
Teleogryllus_occipitalis          ------------------------------------------------------------------------------------------------------------------------
Acheta_domesticus                 HSQLYRNIELRRTLRSTGYNLSL*HAVHDRKVTAFTRDCLKVFSFWRIALQSTVRSGLGAYCSKTESLGRIIYMNYFILSYVRLTNNVELYYSINITDSA**CNTFLKYI*GFYQIRNFI
Consensus

Teleogryllus_occipitalis          ------------------------------------------------------------------------------------------------------------------------
Acheta_domesticus                 *LHSAIAFRDDTIKLRVQL*LFSLLY*ILILFLIFTRVFWLGKSHAKTVQFMSAPFQQLYSGSQNNMALHKEQ*RNSDSEDEELDHWFSAQIILLHNCNDISYFYRVLELISKENISFIT
Consensus

Teleogryllus_occipitalis          ------------------------------------------------------------------------------------------------------------------------
Acheta_domesticus                 WYRSKT*TKKS*KACLNI*MFIRAKFVAGAEYRCVHIDVTQGLMVAWHPMV*GWGVRI*RRRKKQERRVEGRGQLGERRRSTGPIRFKQSTCADRRQQPGGTRTACAEARASNVL*HSET
Consensus

Teleogryllus_occipitalis          --------MW-----------------------------------------------------------------------------------------------CCG------------
Acheta_domesticus                 QVTCRVGILWCFIRSDSFHN*GEGLVAVESTLLLVVGSIYYIHLAIILKFYFGGGFIYQ*TSNKLNSLSSVYYCNLHDATLDAWYVRMYCSSLNLIMRNCSRSVACCGGDAVMCCRLAQL
Consensus                                  *                                                                                               ***

Teleogryllus_occipitalis          ------------------------------------------------------------------------------------------------------------------------
Acheta_domesticus                 HAAHAAAAARRRRGVPARLRHHAHVLPVALLAAHRHVRAQPPRLHQQRELLVGRVAGAPRDALLRHVPQQRRLPHRSVPPNTIEIMLLVHFSSTITLRVIADQTCN*KFCREYATMNLFA
Consensus

Teleogryllus_occipitalis          ------------------------------------------------------------------------------------------------------------------------
Acheta_domesticus                 AARSILGGRNSDRSEHGR*LW*CFSCHMPIPLLIRVIPEYWGLTIVWPQSPLE*TKNQ*ISHLIHQICISDQTCISNSNEKMNVLNLLLSQCPTQSVAVTVCLPLQPATVLRLVNTYCIL
Consensus

Teleogryllus_occipitalis          ------------------------------------------------------------------------------------------------------------------------
Acheta_domesticus                 MKIFLKYM*HIFSLYMFNDTTFQHTIVCTAMKNIGLRL*V*HGTTVFIERR*LVPMCVHRASRNLHFCTSLNSAKGDFISTRQVLGFDL*GERFSDRRIT*RRQHLANHLPPARP*VICL
Consensus

Teleogryllus_occipitalis          ------------------------------------------------------------------------------------------------------------------------
Acheta_domesticus                 SSSSARGGTRLD*RVWRAAKISSRALKSPRVPADATGRCNWFY*SSDSQMAAAGLVHRWAGMWVARRDWRSVGIVPGRGDLEGVVIRWGRL*SRSPWAPAQAVGAAAETPKRPIGFYYGF
Consensus

Teleogryllus_occipitalis          ----------------------------------------------------------------------
Acheta_domesticus                 NIYFILIVPYLMQEYKLIFVRMTE*HTTSQRKPV*TIEHRRK*GRRSS*IIYPYN*L**QQQ******QQ
Consensus
```

**Tocc.8G0000430.1_exon2**

```
Teleogryllus_occipitalis                 -------------------------------------------------------------------------------VTSGRSPTPT-------------------------------
Acheta_domesticus                        IHSCIGTLSFVEH*EARAII*AFSTLFMTVK*PLLLETA*KYFLFGV*LYNPLSGQDLERTAVKRSP*GELYI*IILY*VTSG*PITLNYIIQLISQTVLNSAIRF*NIFEGSIKLEISF
Consensus                                                                                                               ****   *

Teleogryllus_occipitalis                 ----------------------------------------------------------------------------------------PPQ-----------------------------
Acheta_domesticus                        NYTVQLLLEMTL*N*ECNYDYFRFYIRY*YCF*YSQGFSG*VSHMRKQYNLCRHRFNNSILDHKIIWHCIRNNRETAIAKMKNLIIGFPPR*YYFIIVMTFLIFTEFLN**VRKIYRLLL
Consensus                                                                                                                        **

Teleogryllus_occipitalis                 ---PQ----------------------------------------HQPP-------------------------------PPPP------------------------------------
Acheta_domesticus                        GTDPRRKQKNRKKHASIFKCLFERNLLLVLNIDACI*M*HRV*WWHGTPWFKGGVLGYNVGGRNKREGWKGAAS*ASGDAPPAPFASSSRRVQIGGSSRVGRGPRVRRLGPAMYCDTVRL
Consensus                                   *                                         *  *                               ** *

Teleogryllus_occipitalis                 --------------------------PPPQ------------------------------------------------PQP---------------------------------------
Acheta_domesticus                        K*RVV*EFCGASYDRIHSIIKERVW*P*NQHCY**WDLYITFI*LLFLSFILVVDLFINKLAIS*TLSRLCIIATCMTPH*MRGM*GCIAAHLTS**GIVADLLHVVVVTR*CVAGSLNF
Consensus                                                          *  *                                                *

Teleogryllus_occipitalis                 QPR-----------------TRPQPNPTQPQ-----------------------------------------------------------------------------------------
Acheta_domesticus                        MPRTLRLLRDGGAEFRRAYVTTPMCCPSRSSLLTGMYVHNHHVFTNNENCSSAAWQAHHETRSFATYLSNAGYRTGQYHQIL*R*CY*FISVAPLHFE*LQTKLVTESFAGSMQQ*ICLQ
Consensus                                 **                 * *   *

Teleogryllus_occipitalis                 ---------------------------------------------------------------------------PQPQPQ---------------------------------------
Acheta_domesticus                        LPVASLVVETVIVVNMVGDYGNAFLVTCLSRF*YGLSQNIGG*LSFGLNRH*NELKTNKLAI*SIRSVYLIRLVYPTAMKK*MC*IYSSHNAQLKV*QLQCVCRYSLLLFYVW*THIVSS
Consensus                                                                                                                   *

Teleogryllus_occipitalis                 ----------------------------PQPQ-------------------------------------PQSQSQSQSQPQ---------------------------------------
Acheta_domesticus                        *KYF*NICNIYFLCTCLMTPLFNTR*YVPQ*KILACGCESSMARLYLLNVVD*SPCVYIVHHVTFTSAPP*IARKAISSPQGKCSVLICEASAFPIAV*RDEGNILQITCHRRGLE*FV*
Consensus                                                            **                                       *       * **

Teleogryllus_occipitalis                 -P--------------QPQPQP---------RP-RP-------------------------------------------------------QPKPQPQPQPQPQPQPQPQPQP-------
Acheta_domesticus                        APVRREEALALTDGFGEPRKYPPGR*SLRVSRPTRPGAATGFIRAVTRRWLLPGLCTGGQECGWPGATGDPWGSCPGEGIWRAL*SAGAAFDPAAPGRPPRQSVRRLKHQSVPSVFITVS
Consensus                                  *               *   *         ** **                                                        *     *  *       *  *

Teleogryllus_occipitalis                 -----------------------------------QSLTPNLLTLNLPQS*-------------------
Acheta_domesticus                        TFILLF*SYLILCNETS*YLFV*RSDTPRVRENRCEQSSTEGNKEEDLPE*YIHIIDCNNNNNNNNNNNNN
Consensus                                                                   ** *       **
```

**Tocc.8G0002370.1**

```
Teleogryllus_occipitalis      CACGGGAAGCTA-------------------------------------------------CCGATTAGAGTAGCTCAAAATTCTTCTCCATACAG------------------------
Gryllus_bimaculatus           TGCGGAAGGCTACAGTATGTTGGCCGAACAGTCAGCCACCTCGACAGAGACGCGCCTCGTCCCGTATAGCATACTAAATAATTCCACAAAATATAGATTCTAACTTCACAACGTGTTAGA
Gryllus_longicercus           ----------------------GCCGAAACGTCAGCCACCTCGACAGAGACGCGCCTCGTCCCGAATAGCATACTAAATAATTCCAGAAAATATGGATTCTAACTTCACAACGTGTTAGA
Consensus                                                                                  ***  ***  **    * *****      ***  *

Teleogryllus_occipitalis      ------------------------------------------------------------------CGACTCGATTCGATCACGCTGACAAGCAGGTGTCAAAGATCCTGCCACCGTCGG
Gryllus_bimaculatus           GTACGTACGCGCGCGTACTACGCGCTTAGGGGTTACAAAAGAGTTTTGTTCGTGTTCGAGTGCCTCCCATCCAACCGACTTGCAATG-AATATAGTCTTCCAAGTTTGCACGTCCAGATG
Gryllus_longicercus           GTACGCACGCGCGCGTACTACGCGCTTAGGGGTTACAAACGAGTTTTGTTCTTCTTTGAGTGCCTCCCATTCAACCGTATTGCAATGAAATATAGTCTTCCAAGTTTGCACGTCCGGATG
Consensus                                                                                       * *  * *     *  *  **  *   **   ** *** *    *  **    *

Teleogryllus_occipitalis      GACTCCAGGAT-------------------------------CTCCCTGAAC--------------------------------------------------------------------
Gryllus_bimaculatus           AGCTCTCGCATATTTTCACCTCCGTCTAACTGCATCACCTTATTCTCTGGACATTGTTCGCTACAAGAGAAAGGGAAAACATACGCTTGAATGTAGACTGAGGCTATTAGTTGGCTACTT
Gryllus_longicercus           AGTTCTCGCATATTTTCACCTCGATCGAACTGCATCACTTTATTCTCTCGACATTGTTCGCTACAAGAGAAAGGGAAAACATACGCTTGCATATAGACTGA-ACTATTAATTTACTACTT
Consensus                        **  * **                                ** **  **

Teleogryllus_occipitalis      ---------------------------------------------------------CAGGACCTCCCAGCCAGAGATACTGAAGTAAC----------------------ATCAGGTGA
Gryllus_bimaculatus           ACGGAAGAAACTTTCTTGCGGCATAGTTTACCCCTCTTTGTTGTTACATTGCTCGGGCCATGCCTTCTGGGAAGGGAAGCGGGTAAGACGGACGGCTGGCCGGTGGATAGAATCCGGCGA
Gryllus_longicercus           CCGGAAGAAACTTTCTTGCGGCATAGTTTACCCCTCTTTGTTGTTCCATAGCTCGGGCCATGCCTTCTGGGAAGGAAGGCGGGTGAGACGGATGGTTGGCCGGGGGATGCAATCCGGTGA
Consensus                                                                              *    *** *  *  **  *  * *     **                    *** ** **

Teleogryllus_occipitalis      TG---------------------------------------------------------------------------------------------------------TCCAGAGTCACCT
Gryllus_bimaculatus           TGTTCTAAGGACAACCGTGAGCCTGCTTCCCAATGAAAATCATGTTGAGTCGGAAGAGACAAAAATCGAGGAGCGGAGGTGCAGTCAGGCGGCCACTACGATCGCCCTTCGGAGTCATTC
Gryllus_longicercus           TGTTCTAAGGACAACCGTGAGCCTGTTTACCAACGCAAATTATGTTGAGCCGGAAGAGACAAAAATCGAGGAGCGGAGGTGCATTCAGGCGGCCACTGAGATTGCCCTCCGGAGTTACTC
Consensus                     **                                                                                                         * * **** *

Teleogryllus_occipitalis      CTA-CCGGTCGGA---------------------TATTTTGATTTTTAGGGGTAATTGATATTAA---------------------------------------------------ACCT
Gryllus_bimaculatus           ATG-AAAATCGGACCTATGACATCTTTACAAGTGTATTTTACTCTCTTGTTACAAGAAATTTAAAGAAAAAATTTGGAGCCGTTTTCAAAAAAATTCAAAAATGTCCAGGAGATTCATGT
Gryllus_longicercus           ATGAAAAATCGGATCTATGCCATCTTTACAAGTGTAATTTACTCTCTTGTTACAAGAAATTTAAAGAAAAAATTTGGAGCCGTTTTCAATAAATTTCCCAAATGTCCAGGAGATTCGTGT
Consensus                      *      *****                     ** ***  * * * *    **   ** * **                                                   *

Teleogryllus_occipitalis      CAAATGTTCTAAAATTTTG--------CTCCAACAGCT--------------------------------------------------------------TGGTTTAAAT--TTGTTGTC
Gryllus_bimaculatus           CAAATTCTCCCAAATTTTAAAGAATG-TTTTAACAATCTCAAAACATCGACTCGAGCTAGATGGTCAGGAGTTTCTTCAATTTGTAAAAAATACTACGTATGGTTTAGGTGCCTGTGGTG
Gryllus_longicercus           CAAGTACTCCCAAATTTTAAAGAATGTTTTTAACAATCTCAAAGCATCGACTCGAGCTAGATGGACAGGAGTTTCTTCAATTTTAAAAAAATATTACGTATGGTTTAGATGCCTGTGGTG
Consensus                     *** *  **  *******          *  ****                                                                 *******  *   *** **

Teleogryllus_occipitalis      T-------------------CGGATAGAAAAAAAG----------AAACTAAAAAAATGACGTATGTTT-----------------------------------------TATTTTTGTT
Gryllus_bimaculatus           TTCATACACGGGCAGACGGACGGACAGATAGAT------------GGACGGAGGGTGAGAAGTACATT----AAAAAAACGTTTTTGGTATCAGGGAACCCCAAAACGTGTATTTCCGTT
Gryllus_longicercus           TTCATACACAGACAGACGGACGGACAGATAGATAGACGGACGGACGGACGGTGAGAAGTACATTTTAAAAAACAACGTTTTTAGTATCAGG----------ACGTGTATTTCCGTT
Consensus                     *                   **** *** * *              **  *      ** ***  **                                           *****  ***

Teleogryllus_occipitalis      GAATTTTC----TTATTTTTCGAAA--ACTACAAAAGTTTC-----------AAATTTATTCGACGTCTTCCTATTACTATAAAATTGAATACAGAAATTG-------------------
Gryllus_bimaculatus           GAAATTTCGGAATCAATTTTCCACAGCATTGCATAACTTTCCCTATATTACGTAAATTACTTCACATATACGTATATACACGGGTAAGAAAAAATGCATTGAGAGCTATAATGCTGGCCA
Gryllus_longicercus           GAAATCTCGGAATCAATTTTCCACAGCATTGCATAACTTTCCCTATATTACGTAAATTACTCCACATATACGTATATATACGGGTAAGAAAAAATGTATTGAGAGCTATAATGCTGGCCA
Consensus                     *** * **    * * ***** * *  * * ** ** ****            ** *** *  ** * * * ***    *       *** * *   ****

Teleogryllus_occipitalis      ----------------TTTTGAAATGAAATTTGTTCTACGCATATAGGCATTTAATG----AATGCATCTTTTTTTAAATATTAGTAGTTTTAAGTGCAGTTTTTTAGTACAATACGCAC
Gryllus_bimaculatus           TAACATGGCAGGGAGCCTTTGGGTCTGAGTACAACCTGCCCAAATCGGCGGGTAAAACCAATTCGCATC-------------CACTAAGTGAAGGGGCCTTATTTAGGGACCATGCGAAA
Gryllus_longicercus           TAACATGGCAGGGAGCCTTTGGGTCTGAGCACAACCTGCCCAAATCGGCGCGTAAAACCACATCGCATC-------------CACTAAGTGAAGGGGCCTTATTTAGGGACAATGCGAAA
Consensus                                     ****      *       ** * ** ** ***   ***         *****              * **  *  * * **  * ***  * ** ** ** *

Teleogryllus_occipitalis      AATAT------TAAAAACATTTTTTCTCAAAAAGGCTCCAA-----------------------------------------ACTTTTGCACTGTGAAAG---------------GTG
Gryllus_bimaculatus           TAAATATCTGGTGAAATAACTATTTAGTTGAAAAGGTCCAAACATGGAACGGATAATGTATTGTAATGCTGCGGCTAAAACGGACTTTTTCGTTAG-AAAACTCGTAAAGTGGGGCCCTG
Gryllus_longicercus           TAAACTTCTGGTGAAATAACTATTTAGTTGAAAAGATCCAAACATGGAATGGATAATGTATTGTAATGCTGTGGCAAAAACGGACTTTTTCGATAGAAAAACTCGTAAAGTGGGGCCCTG
Consensus                      * *       * ***  * * ***     *** * *****                                          ****** *  *   ***                **

Teleogryllus_occipitalis      ATGTACACTACATTGTTTTTTATCCTTTCGATTTTTCAGAAATTACGCAAATT---------TATTGTTTTGAATTGCCTCTCATT---------------------------------
Gryllus_bimaculatus           ATATCGAAAATGTCGATAAATCTTGTTTCAATATCTTGTAAATTGTACTATCTGTATCATAGTGTAGCCGTACTCTACGTTTTATTGGTCCACGTCACCAAAAAATGGGCTTACGGCTAT
Gryllus_longicercus           ATATCGGAAATGTCGATAAATCTTGTTTCAATATCTTGTAAATTGTACTACCTGTATCATAGTGTAGCCGTAATCCACGTTTTATTGGTCCATGTCACAAAAAAATAGGCTTACGGCTTT
Consensus                     ** *     *  * * *   * *  **** ** * *   *****   * *  *         * * *   *      * * * ***

Teleogryllus_occipitalis      -------------------------TCGTTAACATTAACACTGTGG----------------------------------------------GCGG--------------------
Gryllus_bimaculatus           ACGTCTGTCCGCCTTTCTGTGAAAATCGCTAGCATCTTTACCGTAGATATAATTTTCAACAAAATTGTTATTTCTCAATCGATCGGCCAAATGTCACGGTTCTCTTGAATTTGAAACGGA
Gryllus_longicercus           ACGTCTGTCCACCTTTCTGTGAAAATCGCTAGCATCTAAACCGTAGATATAATTTTCAACAAAATTGTTATTCCTCAATCGTTCGGCCAAATGCCACGG-TCTCTGGAATTTTAAATGGA
Consensus                                              *** ** ***    ** ** *                                           ***

Teleogryllus_occipitalis      ---TTTTATATATTATTCATT-------------TGCAAATAAGAGT-GTTTTAATTATATCTAATAAAATGTCTCAAAAACTTAATTTTTTTCTGTAAAAATGTGCTGAAAAGTGAGCA
Gryllus_bimaculatus           AATATTTTTATTTGATCGATTAAGTAAGTGATAGAGTAAGTATGAGC-TTTTAAATTATTTATAATGAAAATTCTCAAAATCAAACTATTTTTCGGTGAAAATGAGATGAAAATAGCTCA
Gryllus_longicercus           AATGTTTTGATTTTATTGATTAAGTAAGTGATTGAGTAAGTATGAGCGTTTTAAATTATTTCTAATGAAATTTCTCAAAAACAAACTTTTTTT---------TTCGAGGAAAATAGCGAA
Consensus                         ***  ** * **  ***              * ** ** ***   *** ****** * **** ***  ******** *  * * *****         *  *  *****  *   *

Teleogryllus_occipitalis      TTTTCGGAAACACTTAAACGACACTTTCTGGAGAACTGTTCCAAATTT------------TTCCATGAAAATTTGCATGAAGGACAAGATTTTACGCTTCTCTAATATTATTAACGTCAC
Gryllus_bimaculatus           TTTTCGGAAAAGCTCAAACAATACTTTCGCGAGAAGGGTTTCATATTTTCTGAACCCAAATCCAACGAACATTTGCATGTATGTACAGATTTTACTCTTCCCTAATAGTCTTA-------
Gryllus_longicercus           TTTTCGGAAAAGCTCAAACGATACTTTCTCGAGAAGGGTTTCATATTTTCTGAATCCAAATCCAACGAACATTTGCATGTATGTACAGATTTTACTCTTCCCTAATATTCTTA-------
Consensus                     **********  ** **** * ******  *****  *** ** ****            * * * *** ********* * *   ********* **** ****** * ***

Teleogryllus_occipitalis      GTCACATTTCAGTTCTACTATTAGGATACCAAACAACTTATAAA-ATGTGAAAAAAAAGTATCTTA-TAGGCACCAACAGATGGCAACTGGGAAGAGGGTGGGGCAAAACCGAAAAGCTA
Gryllus_bimaculatus           GTCACATTTCAATTCTTCCATTAGG-----------------------------------------------------------------------------------------------
Gryllus_longicercus           GTCACATTTCAATTCTTCCATTAGGGTACCGAACAACTGAGAAATATGTAAAAAGAAAACAATGTATTTGAGTCCCAAAAATGGCAGCTGGGAAGAGGGTGGAGCAAAATCGAAAAGCTA
Consensus                     *********** **** * ******

Teleogryllus_occipitalis      GCTCAGACTAATGAGAACAGGAAATTTGCAAGTACGGGAATTTCTATGTCATTTTGTAGCAT-----------------TATTTCCTTTCTTGATTGTGTGTGTATGATGTGTGTGTGT
Gryllus_bimaculatus           -------GTAATGAGAACAGAAAATTAAGTAGTACCAGA------AAGTCATTTACTAGCACCAATAGTT----------TACTTCCTTACATGACTGTGTGTGTGTG-TGTGTGTGTGT
Gryllus_longicercus           TCCCAGAGTAATGAGAACAGGAAATTAACTAATACCAGA------AAGTCATTTAGTAGCACCACTAGTTTACTTGGGTCTACTTCCTTCCTTGATTGTGTGTGTGTG-TGTG-------
Consensus                             ************ *****    * ***  **      * *******  *****                 ** ****** * *** ********* ** ****

Teleogryllus_occipitalis      GTGTGCGTGTGTGTGCGTGTGCCTGTGTGTGCGCGCGTGCGTATGGTAAACAAGAACGCAATTAAATTACTCACAGAAAGGAAAAATAATTTAGAGTAGTCAACACGCTGTTATACGGTG
Gryllus_bimaculatus           GTGCGCGTGTGTGCGCGCGTGTGTGCTTGTGTATGCGTGTGTGTAGTAAATGAGACCGCAAAGAAATTACTCACAGAAAGGAAAGATAATTTAGAGTAGTCAACACTCTGTTAAATGGTG
Gryllus_longicercus           ------------GGGGGGGCGTGTGCGTGTATATGCGTGTGTGTGGTAAATGAGACCGCAAAGAAATTACTCACACAAAGGAAAGATAATTTAGAGTAGTCAACACTCTGTTATATGGTG
Consensus                                 * * * * *  **  ***    ***** ** * *****  *** *****  ************ ******** ******************** ****** * ****

Teleogryllus_occipitalis      GCTGAAAGGAAACGAAAA---AACTCAATGTATACAGGTATTGGAGTAATCTCATTTCCTTTGCGAGTTCATTCATTCCGCGGTTACCGACGGTGTGGAGGCGTTCGACGATAGCGAGAA
Gryllus_bimaculatus           GCTGAAGGAAAA--AAAAAATAAATAATTCTACCCAAATATTCGAGTTATCGAAGTCCCTTTGCGGGTTCATTCATTCCGCTGTGAATGATGGTGTGAAGGCGATCGTAGGTAGCGAGAG
Gryllus_longicercus           TCTGAAGGAAAATGAAAAACAAAAACATTCTACCCAAACATTCGAGTTATCGAAGTCCCTTTGCGAGTTCATTCATTCCGCTGTGTCCGATGGTGTGAAGGCGATCGTAGGTAGCGAGAG
Consensus                      ***** * ***  ****   **   * * **  **   *** **** ***  * * ******** ************** **    ** ****** ***** ***  * ********

Teleogryllus_occipitalis      CAGACCCACTTGGGCAGTCACTACCAGAGGGATGGAGGGGGGGGG-------------------GTGATCGGCGAAGCGCGTCGCGTGAGGTATCTGCGAGCCGCTTATCGTCCACAAA
Gryllus_bimaculatus           CAGA-ACACTTGGCCAGCCGCAACCAGAGCGAGGGTGGGTGGGGGTGGGGGGTACAAGGTTGGACGCGGACAGAAAAGCGCGCCGCGTGAGGTATCTGCGAGCCGCTTATCGTCCGCAAA
Gryllus_longicercus           CAGA-ACACTTGGTCAGCCGCCACCAGAGCGAGTGTAGGTGGGGGTGGGGGGTACAAGGTTGGACGCGGACAGAAAAGCGCGCCGCGTGAGGTATCTGCGAGCCGCTTATCGTCCGCAAA
Consensus                     ****  ******* *** * * ******* **  *  ** *****                    * *  * *  ******* ******************************** ****

Teleogryllus_occipitalis      CACGCGATACGGCGCCGCGGCTCAGACACCCATTCCATCGTCTCCGCCGGCGCGCCGCGGAGACGATAGGACGAGTGTCTGAGGTCGGCGCAAAGCGA-GGGACATCGGTAAACGCGGTC
Gryllus_bimaculatus           CACGCGATACGGCGCCG---------------------------------------GCTGAGCCGATAGGAGAGGGGGCTGCGGACGGCGCAAACCGAGGGGACATCGGTG--------C
Gryllus_longicercus           CACGCGATACGGCGCCG---------------------------------------GCGGAGGCGATAGGAGAGGTGGCTGAGGACGGCGCAAAACGAGGGGACATCGGTG--------A
Consensus                     *****************                                       ** *** ********   * * *** ** ********* *** ***********

Teleogryllus_occipitalis      ATGTCACTAGGGATAGCGGAGATAAAGATGGAAATAAAGATTGACAAAGCGGCGGTGGTGATGGTG--------------------------------------------------AATG
Gryllus_bimaculatus           ATGCCATTAGGAATAGCCAAGACAAAGAAGGAAATAAAGATTGGCAAAGCGGCGGCGAC-------------------------------------------------------------
Gryllus_longicercus           ATGCCATTAGGAATAGCCAAGATAAAGAAGGAAATAAAGATTGGCAAAGCGGCGGCGGCGGCGGCGGCGGCGGCGGCGGCGGCGGCGGCGGCGGCGGCGGCGGCGGCGGCGGCGGCGGCG
Consensus                     *** ** **** *****  *** ***** ************** *********** *

Teleogryllus_occipitalis      AAGGTGGGTATAAAGTCGAAGGTGACGAAAGAAGTGGGAATTG-----------------------------------------------------------------------------
Gryllus_bimaculatus           ------------------------------------------------------------------------------------------------------------------------
Gryllus_longicercus           GCGGCGGCGGCGGCGGCGGTGGTGGTGGTGGTGGTGGTGGTGGTGGTGTTGGTGGTGTTGGTGGTGTTGGTGGTGTTGGTGGTGTTGGTGGTGTTGGTGGTGTTGGTGGTGGTTGTGGTG
Consensus

Teleogryllus_occipitalis      --------------------------------------------GGGGTCGTGGAAAAGGTGGTGAGCGAGGAGGAGTGGATAGAGAGGGTGGAAGAATATGGAAAGGGTATGAAGAAGA
Gryllus_bimaculatus           --------------------------------------------GGCATCGTGAAAAAGAT------GGAGAAGAAATGAATATAGAGAGTGGAAGAAAGTG--AAGGGTATGTAGGAGA
Gryllus_longicercus           GTTGTGGTGGTTGTGGTGGTTGTGGTGGTTGTGGTGGTGGTGGTGGCATCGTGGAAAAGATGGTGAAGGAGAAGGAGTGAATATAGAGAGTGGAAGAATGTGGAAAGGGTATGTAGGACA
Consensus                                                                 **  ***** ***** *       *** ** * ** *** **** *********  **  ********* ** * *

Teleogryllus_occipitalis      GTGCCGGAGA-TAGTGGAGGAGAGTGTGGAGCAAAGTGAGGACGGAGGGTAGAGAAGAGGATATCGATAGG----------------------GTGGAGAGAGTTTGTAGAGGAGTCGA
Gryllus_bimaculatus           GTGCGGAGAGTTAGTGGAGGAGAGTGTGGAGGAAAGTGA--TGAGAGGGTAGAAAAGAGGGAAATGGAAAGGAAAGAGAAGTATAC--------TGGAAAAGAGTGCATAGAGAGGTTGG
Gryllus_longicercus           GTGCGGAAGGTTAGTGGAGGAGAGTGTGGATGAAAGTGAGGAGAGAGGGTAGAAAAGAGGGTATTGGGAAGGGAAGAGAAGTATACTGGGAGGGTGGAAAAGAGTGCATAGAGAGGTTGG
Consensus                     **** *     *******************  *******     ********* ******  *  *  * *                        * * *****   *****  ** *

Teleogryllus_occipitalis      AGGAGAATGTGAAGAGGGGTGGAGGAGAGCATGGTTGGAGGAAAGAGTAGGGACAGGATGAAGGAGAGTGTGAATAAAAG-ATGGAGGATTGTGATGAAAATAGTTGGTGGATGAAACAG
Gryllus_bimaculatus           AGGTGAATGTGAAGAGGAGTGGAGGAGAGCGTGTTTGGAGG--ACAATAGGGAGAGGGTGGGGGAGAGTGTGGAAAGAGGAAAAAAGGACAGTGATGAAAATATT---TGGAGGAAACAG
Gryllus_longicercus           AGGTGAATGTGAAGAGGGGTGGAGTAGAGCGTGTTTGGAGG--ACAATAGGGAGAGGGTGGAGGAGAGTGTGGAAAGAGGGAAAAAGGATAGTGATGAAAATAGT---TGGAGGAAAC-G
Consensus                     *** ************* ****** ***** ** *******  * * ****** *** **  ********** * * * * *   ****  ************ *   **** ***** *

Teleogryllus_occipitalis      TGGCCGGTGGAGACGGTGTAGTAGGCAAAGCTCCACAAATTTTGTCCTGGCGATGAATTTGTCGGGTGGCAGCGATAGTTGGAAAGAGTCAAAGAGATAGTGCTGATGGAAGAAGAGTGG
Gryllus_bimaculatus           TGGGCGAAGGAGACAATGTAGCTGGCAAAACTGAACAAATTTTGCTTTCGCGATGAATTTGTCAGGTGGCAGCGGCAGCTGAAGGGAGAGAAAGAGATAGTGCTAGTGGTCGAAGAGCGG
Gryllus_longicercus           TGGGCGAAGGAGACAATGTAGCAGGCAAAACGGCACAAATTTTGCTTTGGCGATGAATTTGTCAGGTGGCAGCGGCAGCTGAA--GAGAGAGAGAGATAGTGCTTGTGATCGAAGAGCGG
Consensus                     *** **  ******  *****  ****** *   **********   * ************** **********  ** ** *  ***  * ************  **   ****** **

Teleogryllus_occipitalis      TAGAGATAGTGGTGAAGGCGGAAGCAACTTCGATCCTACAGTTGTTGTGGCGTGGGTGTTGCCATAACTATAAGAGGAAGGATTGGCAGTATGTAGAGTATGGGGATTTCCCCTCAGTGG
Gryllus_bimaculatus           TAGAGATAGTGGCGAATGCGGAAGCAACATCGATCGTACAGTTGTTGTGGCGTGGGTGTTGCCATAACTATAAGAGGAAGGTTGGCGGCAATATATACCGTGCGGGTTTCCCCTCGGCGA
Gryllus_longicercus           TAGAGATGGTGGTGAATGCGGAAGCAACTTCGATCGCACAGTTGTTGTGGCGTGGGTGTTGCCATAACTATAAGAGGAAGGTTGGTGGCAATATATACCGTGCGGGTTTTCCCTCAGCGA
Consensus                     ******* **** *** *********** ******  ******************************************* * *     ** ** *   ** ** *** ***** * *

Teleogryllus_occipitalis      CAC-TTGCATAAGAGCGTCGACAACGGAGGAGGAGGCGGCGGCGGCGGCGGCG---------------GTGGCGGCAGAGGCAGAGGCGGCGGCGGCGG----------AGGTGGAGGC
Gryllus_bimaculatus           CACTTTGCGT-----------------ATAGGGATGAGACAACGATGTCGATGCTTTCAAAACATCGATGTTTTTGTAGATACATATATGTTTTAAACAT--------------CGATGT
Gryllus_longicercus           CAC-TTGCGT-----------------ATAGGGATGCGACATCCATGTCGATGTTTTCAAAACATCGATGTTTTTGTAGATACATGGATGTTTTAAACATCGACGTGTTAACACCGATGT
Consensus                     *** **** *                *   *** * * *  *   * **  *               **    * ***  **     *       *                 ** *

Teleogryllus_occipitalis      GGCGGCGACGGCGGCG------------------GCGGCGGCGGAGGCGGAGGTGGAGGCGGCGGCGGAGGAGGAGGAGGCGGCGGAGACGGCGGCGTGGCGAAGGTGCAGC-----
Gryllus_bimaculatus           TTTTCTAACGACGATGTTTCTTCTTTTCCGATGTTCTGTTTCTGAATGAAAAATTCCCATGACCACAATACAGTAAAAGTTAAAATATGAAATAATAACTGATAGTAAACACTGCTCTTC
Gryllus_longicercus           TTTTATGACATCGATTTTTTTGCTTTTCCGATGTTCTGTTTCTGAATGAAAAATTCCCATGCTCACAATACAGTAAAAGTTAAAATATGAAATAATACCTGATAGTAAATACTGTTGTTC
Consensus                            **  **                     * *   * **     *  *      *    *     ** *  **         ** *      *     * *    * *

Teleogryllus_occipitalis      -------GGAGGCGGCGGA-----------------------GGCGGCGGCGGCGGCAGCGGGGGTGGCGGCGGCGGCGGTG------------------------------ACGGAGGC
Gryllus_bimaculatus           ATGTGTTGGAACCGAAGGAACGCTGCTTGTACTACCCTCCTGGACCACACTAATGTCGCCCGATATGGCCTCCATACCCTTGTATTTCCTCTTCTTCATATTCAATTCGCACATTGAGAT
Gryllus_longicercus           ATGTGTTGGAACCGAAGGAACGCTGTTTGTACTACCCTCCTGGACCACACTAATGTCGCCCGATATGACCTCCATGCCCTTGTATTTCCTCTTCTTCATATTCATTTCGCACATTGAGAT
Consensus                            ***  **  ***                      * *  *      * *  * *   ** *  *     *  **                           *  ***

Teleogryllus_occipitalis      AGAGGCGGGAGCGGAGGCGGAGGCGGAG---------------------GCGGTGGTGGAGGCGGCGGCGGAGG----------------------TGGCGGTGGCGAAGGTGGCGGCGG
Gryllus_bimaculatus           CTGGCCAACAAATACGCTGTGGGCATAGGGTCTGCCCCTCCTTCCCGTAGCCGTATTCGGCGAATTGATAAACACGCATACTGTCAAATTAAATTTTAATGCACATGAATTTAATGTTAG
Gryllus_longicercus           GTGGCCAACAAGTACGCTGTGGGCATAGGGTCTGCCTCTCCTTCCCGTAGCCTTATTCGGCGAATTGATAAACACGAATACTGTCAAATTAAATTTTAATGCACATGAATTTAATGTTAG
Consensus                        * *   *     *  *  ***  **                     **  *  * * *    *    *                           *   *     ***  *   *   *

Teleogryllus_occipitalis      AGGTGGCGGCGGCGGCGGCGG--------CGGCGGCGGCGGTGG----CGGTGGCGGTGGCGGTGGCGGCGGCG------------GCGGCGGCGGTGGCGGTGGCGGCGG-CGGCGGCG
Gryllus_bimaculatus           AGGTTACTACTCCAGTCTCGAAACATGTTGAAAAACATCGATGGTTTTTAGTCGCTGTATTTTTATCATCGATGCTTTAGTTTTGAACATCGATGTTTGCTAACTCGATGATTTGTTATG
Gryllus_longicercus           AGGTTACTCCTCCAGTCTGGAAACATGTTGAAAAACATCGATGGTTTTTAGTCGCTGTATTTTTAACATCGATGCTTTGGTTTTGAACATCGATGTTTGCTAACTCGATGATTTGTTATG
Consensus                     ****  *  *  * *    *               *  ** ***      ** ** **     *  *  **  *            *  **  * * **     **  *    *    *

Teleogryllus_occipitalis      GCGGCGGCGGCGGAGGCGGCGGAGGAGGAGGCGGCAGAGGC-GGCGGCGGCTGAGACCGGGCATCGCCGCAGCTGCGACCGCGACCCCCCGCTTGATCCCGATCCCATCGGCGCGTGGCC
Gryllus_bimaculatus           GATGTCGCATCACTACTTGCGTAAGAGCCGC-----------GGCACCGGC--------GGCGTCGCCGCAGCTGCGGCCGCGACCCTCCGCTTGATCCCGAT-CCAGCGGCGCGTGGCC
Gryllus_longicercus           GATGTCGCATCACTACTTGCGTAAGAGCGGCGGACAGCGGGGGGCGGAGGC--------GGCATCGCCGCAGCTGCGGCCGCGACCCTCCGCTTGATCCCGAT-CCAGTGGCGCGTGGCC
Consensus                     *  *  **  *       *** * ***  *             ***   ***        *** ************** ********* ************** ***  **********

Teleogryllus_occipitalis      TCTGAAAGCGCGCGGCAAGGCCACTGCTTGACCGGCTTCCGAAACTCCGCCTTTTA----------------------------------------------------------------
Gryllus_bimaculatus           TCTGGACGCGCCCGGCAAGGCCACTGCTTGAATGTCTCCCTAAACTCCGCCTTCTACTTGTCTTGGAAGAACATGCGAAACAGTGGCTTCGAATGTTAAGTTCGAAGTTGTTACGTTCGG
Gryllus_longicercus           TCTGGACGCGCCCGGCAAGGCCGCTGCTTGAATGCCTCCCTAAACTCCGCCTTCCACTTGTCTTGGGATAACATTCGAAACAGTGGTTTCGAATGTTAAGTTCGAAGTTGTTTCGTTCGG
Consensus                     **** * **** ********** ********  * ** ** ************  *

Teleogryllus_occipitalis      ------------------------------------------------------------------------------------------------------------------------
Gryllus_bimaculatus           AGATGTAACGCTTAGTGTTTCCAACTGTTAAGTACTGGGGGAACAAAGGTGATCAAAACCATGGATTTCCATGAAACACATGG-AAAAAATCCTTCGTACGCCTCCCTCCGGAGGAGAAG
Gryllus_longicercus           AGATGTAACGCTTAGCGTTTCGAACTGTTAAGTTCGGGGGGTACAAACACGATCGAAACCATGGATTTCCAAGGAAGACAAGGAAAAAAATCCTTCGTACGCCTCCCTCCGGAGGAGAAG
Consensus

Teleogryllus_occipitalis      ------------------------------------------------------------------------------------------------------------------------
Gryllus_bimaculatus           TAAAGCCAGCTGTCCCATGTTATAGATTTCCTGCGTGTAAAAGATTCTCAAGTCTAAAAGACACCTCCGAGGGAAATTAAAATCGCCCGTTTCATCACCCGGGTAATACGACCTGTGTCT
Gryllus_longicercus           TATAGCCAGCTGTTCCATGTTATAGATTTACTGCGTGTAAAAGATTCTCAAGCCTATAAGACACCTACGAGGGAAATTAAAATCGCCCGTTTCGTCACCCGAGTGATACGACCTCTCTTT
Consensus

Teleogryllus_occipitalis      -----------------------------------------------------------------------------------------------------------------AGCGAGA
Gryllus_bimaculatus           GCCATACGGTGAAAGGAGCGCAGCTAAATTTGCGTGGTCAGACGGAAGAGCAGGCCTCATGACGTAACCCTCGCCACAGAAAATATGGCATAATTGAAGTGAATTGAACAATGAATAACA
Gryllus_longicercus           GC------------------------------------CAGACGGAAAAGCAGACCTCACGACGAGAGCCTCGCCACAGAAAATAAGGCATAATTGAAGTGAATTGAACAATGAATAACA
Consensus                                                                                                                      *   * *

Teleogryllus_occipitalis      AACCCACTAGGAGCTCCCGGCTC----GGCTCGGCCGACAGCCCGGCGAGGCGCGG----------------------------------------------------------------
Gryllus_bimaculatus           ATCTCAATGAAGGCTTACAATTTTTAGTTTTTAGTTGACAAATCTAAGAGTTTTAGGTAATTGCGACTTAGACTATAACAAGTTCTAACACATACTAACATTCTGAAATAATAAGTGATT
Gryllus_longicercus           ACCTCAATGAAGGATAACAATT-----CATTTAGTTGACAAGACTAAGAGTTTTAGATAATTGCGACTTAGACTATAACAAGTTCTAACATATACTAACATTCTGAAATAACAAGAACTT
Consensus                     * * ** *    * *  *   *        *  *  ****   *   ***      *

Teleogryllus_occipitalis      ------------------------------------------------------------------------------------------------------------------------
Gryllus_bimaculatus           CCTACTTTTCACTTCAGCTTCATCAGTATTGTAAAAGAAAGCGCACTGTGAGCCAGTATCAATTATTTGACCCTTATTTTGATTGAATAAAAAAACTGCTAACCGATTAAGACAAAATCT
Gryllus_longicercus           CCTAGTTTTCACTTCAACTTCATCAGTATTGTAAAGGAAAGCGCACTGTGAGCCAGTATCAATAACTTGGCCCTTATTTTGTTTG-ATCAAAAAACTGCTAACCATTTAAGACAGAATCT
Consensus

Teleogryllus_occipitalis      ------------------------------------------------------------------------------------------------------------------------
Gryllus_bimaculatus           TAAATGTTCTATAAATATTTGTGCCCCAGCTGCGGCCACGTCTCTTCAATATTCTTGAACTAGATTAGAATGGTGTCGCCGACGGGTATTTTCATGTCAAAAACAATTTTTTGTGCTTTT
Gryllus_longicercus           TAATTGTTCTATAAATATGTTTGCCCCAGCTGCGTCCGCGTCTCTTCAATATTCTTGAGCTAGATTAGAATGGCGTCGCGGAAGGGTATTTTCATGTCAAAAACTATTATTTGTGCTTTT
Consensus

Teleogryllus_occipitalis      -------------------------------------------TGCCGCTCCGCTCGGGCCTGCCAG-----------------------------------------------------
Gryllus_bimaculatus           TAAGTTCATAATAAAAA-ACATTTTAAATATATTCCAGTTGCTCGTCGTCCAATTCGGGCATACAAAGTTACATGAAATAGTATCGATTAAAATACTCGTTTAAGCTTTAGTTTCATCAC
Gryllus_longicercus           TAAGTTCATAATAAAAACTCATTTTAAATATATTACAGTTGCTCGTCGTCCAATTCGGGCATACAAAGTTACATGGAAAAGTATCGATTAAAATGCTCTTTTAAACTTTTGTTTCATCAC
Consensus                                                                 * **  *   ****** * * *

Teleogryllus_occipitalis      ------------------------------------------------------------------------------------------------------------------------
Gryllus_bimaculatus           AAAATTACGTCTAATTAACGTCAAACTGATCAATCAAGTCCAAATGATTCGTGTGAGGCCCGGAAGGAGAGAAGTAATTCAGGGTTTCTCCAGTGTTCTCGTAGTTGCGTTGAAAACGCA
Gryllus_longicercus           AAAATTACGTTTAATCAACGTCATACTGATCAATGAAGTCCAAATGATTCGTGTAAGGTCCGGAAGGAGAGAAGTAATTCAGGGTTTCTCCAGTGTTCTCGTAGGTACGTTGAAAACGCG
Consensus

Teleogryllus_occipitalis      ------------------------------------------------------------------------------------------------------------------------
Gryllus_bimaculatus           AAAAGCTCGTGTTTTTATAGAATTTTTCCACAATACCGCGGAGGGATTCAGCTAAGAATTGAAGGGCTCGGTAGAACAAGAGGGTATTAGGT-------------------TATTAGCAT
Gryllus_longicercus           AAATGCGCGTGTTTTTATAGAATTCTTCCACAATACCCTGGAGGTAATCAGCTAAGAATTGAAGGGCTCAGTAGAACAAGGGGGGTATTCGACATTGTGGTGAAATTGGGTATATTAGCAT
Consensus

Teleogryllus_occipitalis      ------------------------------------------------------------------------------------------------------------------------
Gryllus_bimaculatus           TATACAAAACTTTAAATATTAGTCTT---------TTAGTGTAATATGTTAAATCCAGTATGGACGTGTATGCAAATGTACTGTCTTTAAATGTTTAAAAATGTACTGATTTATATGGTG
Gryllus_longicercus           TATACAAAACTTCAAATATTATTCTTACTTAAATATTAATATGTTAC-TTAAATCCTGCATGAACTTATATGCAAGTGTACTGACTTTAAAAGTTTAAAGATGTACTGACTTATATGGAG
Consensus

Teleogryllus_occipitalis      ---------------------------------------------------------------------------------------------TCACCCATTACGGACGGCCCGGC----
Gryllus_bimaculatus           GAAGTTTAGGTAAAATCTGGGTAATTAGGAAGTTTAGGTAGGTTATCTGAGAACTATGCCAATGTGGGATAGCCTATTCTACTACTGATCTCTTCAATTGTCACAATCGACTCGTCTTGT
Gryllus_longicercus           GAAGTTTAGGTAAAATGTGGGTAATTAGGAAGTTTAGGCAAGTTATCTGAGAACTATGCCAATGTGGGATAGCCTCTTGTTCTACTGGTCCCTTCAATTGTCACAATCGACTCGTCTTGT
Consensus                                                                                                    ***    * **   ** * ** *

Teleogryllus_occipitalis      --------------------------------------------------------ACGGATCGGAACTG--------------------------------------------------
Gryllus_bimaculatus           CGAGAGGGATCGGCTGAGCAATGTTTCGCCTCTGTAGGTGCCGTACGTTACCTAAAACACATTTGAGTTA---CAAAATTACACAAATACGTTTTTTAGAGAAATGTTTTAGAATTTTCT
Gryllus_longicercus           CGAGAGGAATCGACTGAGCAATTGATCGCCTCTGTAGGTACCGTACGTTACCTAAAACACATATGAGTTGCTTCAAAATGACAAAAATACGCTTTTTAGAGAAATGTTTTAGAATTTTCT
Consensus                                                                             **  **  **  *

Teleogryllus_occipitalis      ---------------------------------CACGAGACGC--------------CACTTGTTCCAGTAGCGATG-------------------------------------------
Gryllus_bimaculatus           CAATGAATTCGAAAAAAATAATCAATTCTCCATCACGAGGTGCGTCAAATGGCATGTCATTTGTCTCGATT--GATGGACCGGAAGTGGTCTGTTAGTGGTGGAAGAACGGAGTAGGTTA
Gryllus_longicercus           CAATGAATTCGCAAAGAATAATCAGTTCTCCACCACGAGGCGCGTCAAATGGCATCTCATTTGTCTCGATTGAGACAAATGAGTCTGGGTCTGTTAGTGGCGGAAGAACGGAGCAGGTTA
Consensus                                                      ******  **               ** ****  *  *   **

Teleogryllus_occipitalis      ------------------------------------------------------------------------------------------------------------------------
Gryllus_bimaculatus           CTCTACTACTATACAAATAATACGGTTAAGTCCCTTAAATTGCGGTCAATTTTTACGGTGCTTTTTTTTATTTTTCGAAAACATAAGCCGAGTATAATAAGTAATATTCTAATCGCTTGT
Gryllus_longicercus           CTCGACTACTATACAAGTAATACGGTTAAGTC----------------------------------------------------------------------------------------
Consensus

Teleogryllus_occipitalis      -----------------------------------------------------------------------------------AAGAGTTTAA---------------------------
Gryllus_bimaculatus           GAAATGTACAGTGCTAATAACGGGTTAACTATACAGATTAATTTAAAGAAATAAAGAAAAACATCTTATTATTTTTAAACTGTAAAAGTTCAATTTGTGGCCAGTTAA-AAATAAATTAA
Gryllus_longicercus           ---------------------------------------------------------------TCTTATTATTTTGAAAGTGTAAAAGTTCAATTTGTGGCCATTTAACAAATAAATTAA
Consensus                                                                                        ** **** **

Teleogryllus_occipitalis      -------------------------TACTTTCGATGGAG-GAAT----------------------------------------------------------------------------
Gryllus_bimaculatus           GTAAATAAAAGTGTTCTTCCAAAGTTAATTTAAATATAA-AAACAAGATAAAATATGAGACTGCTGCAATATCATCGCAGGACGG-----------------------------------
Gryllus_longicercus           GTAAATAGAAGTCCTCTTTCAAAGTTAATTTAAATATAACAAACAAGATAAAATATGGAACTGTTGCAATATCCTCGCAGAACTGCCTGACCTCGTGGCCGTATTAACGACAGTGGATGC
Consensus                                              ** ***  **  *   **

Teleogryllus_occipitalis      ------------------------------------------------------------------------------------------------------------------------
Gryllus_bimaculatus           ------------------------------------------------------------------------------------------------------------------------
Gryllus_longicercus           GACCGGATGTGAATGCGGTCGGGACAAGCAAGAGCGGAACAAGACGCGACGGGACAAGACGGGGACGGATACGACGGGACGGGCAGGACAAGATGGGGCGGTACGAGACGAGGCGGCACA
Consensus

Teleogryllus_occipitalis      ----------------------
Gryllus_bimaculatus           ----------------------
Gryllus_longicercus           AGACGAAGCGGGACCCGACAGG
Consensus
```

**Tocc.8G0002370.1_exon1**

```
Teleogryllus_occipitalis      ------------------------------------------------------------------------------------------------------------------------
Gryllus_bimaculatus           GGRADRGTDARTDGRSDGRTLGRTDGRTDGETFAFN*FFSINDPQYIIMLPRPEIVLSRPYGTSRVSVKLADVSANYYGLPQKELLEAVGSAVRSAEGYSMLAEQSATSTETRLVPYSIL
Gryllus_longicercus           ------------------------------------------------------------------------------------------------------AETSATSTETRLVPNSIL
Consensus

Teleogryllus_occipitalis      ------------------------------------------------------------------------------------------------------------------------
Gryllus_bimaculatus           NNSTKYRF*LHNVLEYVRARTTRLGVTKEFCSCSSASHPTDLQ*I*SSKFARPDELSHIFTS---V*LHHL----ILWTLFATREREN------IRLNVD*GY*LATYGRNFLAA*FTPL
Gryllus_longicercus           NNSRKYGF*LHNVLEYARARTTRLGVTNEFCSSLSASHSTVLQ---------*NIVFQVCTSG*VLAYFHLDRTASLYSLDIVRYKRKGKHTLAYRLN----Y*FTTSGRNFLAA*FTPL
Consensus

Teleogryllus_occipitalis      ------------------------------------------------------------------------------------------------------------------------
Gryllus_bimaculatus           CCYIARAMPSGKGSG*DGRLAGG*NPAMF*GQP*ACFPMKIMLSRKRQKSRSGGAVRRPLRSPFGVIHENRTYDIFTSVFYSLVTRNLKKKFGAVFKKIQKCPGDSCQILP-NFKECFNN
Gryllus_longicercus           CCSIARAMPSGKEGG*DGWLAGGCNPVMF*GQP*ACLPTQIMLSRKRQKSRSGGAFRRPLRLPSGVTHEK---------------------------------SDLCHLYKCNLLSCYKK
Consensus

Teleogryllus_occipitalis      ------------------------------------------------------------------------------------------------------------------------
Gryllus_bimaculatus           LKTSTRARWSGVSSICKKY-------------------------YVWFR---------CLWCSYTGRRTDR*MDGG*E----VH*KNVFGIREPQNVYFR*NFGINFPQHCITFPILRKL
Gryllus_longicercus           FKEKIWSRFQ*ISQMSRRFVSSTPKF*RMFLTISKHRLELDGQEFLQF*KNITYGLDACGVHTQTDGRTDR*TDGRTDGEKYILKNNVFSIRT------------------CISVEISESI
Consensus

Teleogryllus_occipitalis      ------------------------------------------------------------------------------------------------------------------------
Gryllus_bimaculatus           LH-------IYV-YTRVRKNALRAIMLAITWQGAFGSEYNLP----------KSAGKTNSHPLSEGALFRDHAK*ISGE----------ITI*LKRSKHGTDNVL*C--CG*NGLFR*KT
Gryllus_longicercus           FHSIA*LSLYYVNYS--------------TYTYIYG*EKMY*EL*CWP*HGREPLGLSTTCPNRRVKPHRIH--*VKGPYLGTMRNKLLVK*LFS*KDPNMEWIMYCNAVAKTDFFDRKT
Consensus

Teleogryllus_occipitalis      ------------------------------------------------------------------------------------------------------------------------
Gryllus_bimaculatus           RKVGP*YRKCR*ILFQYLVNCTICIIV*PYSTFYWSTSPKNGLTAIRLSAFL*KSLASLP*I*FSTKLLFLNRSAKCHGSLEFETEIFLF-DRLSK**SKYELLNYL**KFSKSNYFSVK
Gryllus_longicercus           RKVGP*YRKCR*ILFQYLVNCTTCIIV*P*STFYWSMSQKNRLTALRLSTFL*KSLASKP*I*FSTKLLFLNRSAKCHGLWNFKWKCFDFID*VSD*VS-MSVLNYF**NFSKTNFF---
Consensus

Teleogryllus_occipitalis      ------------------------------------------------------------------------------------------------------------------------
Gryllus_bimaculatus           MR*K*LIFGKAQTILSREGFHIF*TQIQRTFACMYRFYSSLIVLVTFQFFH*G----------------------------------NENRKLSSTRKSFTSTNSLLPYMTVCVCVCVCA
Gryllus_longicercus           FRGK*RIFGKAQTILSREGFHIF*IQIQRTFACMYRFYSSLIFLVTFQFFH*GTEQLRNM*KENNVFESQKWQLGRGWSKIEKLSQSNENRKLTNTRKSFSSTTSLLGSTSFLDCVCVCG
Consensus

Teleogryllus_occipitalis      ------------------------------------------------------------------------------------------------------------------------
Gryllus_bimaculatus           RVCARVCLCMRVCSK*DRKEITHRKER*FRVVNTLLNGG*RKKKINN-----STQIFELSKS---LCGFIHSAVNDGVKAIVGSESRTLGQPQPE--RGWVGVGGTRLD-----------
Gryllus_longicercus           ---GGVCVYMRVCGK*DRKEITHTKER*FRVVNTLLYGV*RKMKNKNILPKHSSYRSPFASSFIPLCPMV*R------RS*VARAEHLVSRHQSECRWGWGVQGWTRTEKRAA*GICEPL
Consensus

Teleogryllus_occipitalis      ------------------------------------------------------------------------------------------------------------------------
Gryllus_bimaculatus           -----ADRKARR--VRYLRAAYRPQT-----RDTAPAE---------------------------------------------------------------PIGEGAADGANRGDIGACH
Gryllus_longicercus           IVRKHAIRRRRRR*ERWLRTAQNEGTSVNAIRNSQDKEGNKDWQSGGGGGGGGGGGGGGGGGGGGGGGGGGGGGGGGGGGGGGGGVGGVGGVGGVGGVGGVGGGCGGCGGCGGCGGCG
Consensus

Teleogryllus_occipitalis      ------------------------------------------------------------------------------------------------------------------------
Gryllus_bimaculatus           *E*PRQRRK*RLAKRRRRHREKDGEEMNIESGRK*RVCRRVRRVSGGECGGK**EGRKEGNGKEREVYWKRVHREVGGECEEEWRRACLEDNRERVGESVERGKKDSDENIWRKQWAKET
Gryllus_longicercus           GGGGIVEKMVKEKE*I*RVEECGKGM*DSAEG*WRRVWMKVRR-----------EGRKEGIGKGREVYW-----EGGKECIERLEVNVKRGGVERVWRTIGRG--------WRRVWKEGK
Consensus

Teleogryllus_occipitalis      ------------------------------------------------------------------------------------------------------------------------
Gryllus_bimaculatus           M*LAKL---------NKFCFRDEFVRWQRQLKGEKEIVLVVEER*R*WRMRKQHRSYSCCGVGVAITIRGRLAAIYTVRVSPRRH-----------FAYRD-------------ETTMSM
Gryllus_longicercus           RIVMKIVGGNVGEGDNVAGKTAQILLWR*IC----QVAAAAEERER*CL*SKSGRDGGECGSNFDRT-------VVVAWVLP*L*EEGWWQYIPCGFSLSDTCV*GCDIHVDVFKTSM-F
Consensus

Teleogryllus_occipitalis      ------------------------------------------------------------------------------------------------------------------------
Gryllus_bimaculatus           LSKHRCFCRYIYVLNIDVFLTTMFLLFRCSVSE*KIPMTTIQ*KLKYEIITDSKHCSSCVGTEGTLLVLPSWTTLMSPDMASIPLYFLFFIFNSHIEIWPTNTLWA*GLPLLPVAVFGEL
Gryllus_longicercus           L*IHGCF-KHRRV-NTDVFMTSIFLLFRCSVSE*KIPMLTIQ*KLKYEIIPDSKYCCSCVGTEGTLFVLPSWTTLMSPDMTSMPLYFLFFIFISHIEMWPTSTLWA*GLPLLPVALFGEL
Consensus

Teleogryllus_occipitalis      ------------------------------------------------------------------------------------------------------------------------
Gryllus_bimaculatus           INTHTVKLNFNAHEFNVRGYYSSLETC*KTSMVFSRCIFIIDALVLNIDVC*LDDLLWMSHHYLRKSRGTGGVAAAAAATLRLIPIQRRVASG----------------RARQGHCLNVS
Gryllus_longicercus           INTNTVKLNFNAHEFNVRGYSSSLETC*KTSMVFSRCIFNIDALVLNIDVC*LDDLLWMSHHYLRKSGGQRG-AEAASPQLRPRPSA*SRSSGAWPLDAPGKAAA*MPP*TPPSTCLGIT
Consensus

Teleogryllus_occipitalis      ------------------------------------------------------------------------------------------------------------------------
Gryllus_bimaculatus           LNSAFYLSWKNMRNSGFEC--------*VRSCYVRRCNA*CFQLLSTGGTKVIKTMDFHETHGKNPSYASLRRRSKASCPML*ISCV*KILKSKRHLRGKLKSPVSSPG*YDLCLPYGER
Gryllus_longicercus           FETV-------VSNVKFEVVSFGDVTLSVSNC*VRGVQTRSKPWISK------------EDKEKNPSYASLRRRSIASCSML*IYCV*KILKPIRHLRGKLKSPVSSPE*YDLSLP----
Consensus

Teleogryllus_occipitalis      ------------------------------------------------------------------------MTY---------------------------------------------
Gryllus_bimaculatus           SAAKFAWSDGRAGLMT*PSPQKIWHN*SELNNE*QSQ*RLTIF------SF*LTNLRVLGNCDLDYNKF*HILTF*NNK*FLLFTSASSVL*KKA----HCEPVSII-----------*P
Gryllus_longicercus           --------DGKADLTTRASPQKIRHN*SELNNE*QPQ*RITIHLVDKTKSFR*LRLRL-----------*QVLTYTN------------ILK*QELPSFHFNFISIVKESAL*ASINNLA
Consensus                                                                                              *

Teleogryllus_occipitalis      ------------------------------------------------------------------------------------------------------------------------
Gryllus_bimaculatus           LF*LNKKTANRLRQNLKCSINICAPAAATSLQYS*TRLEWCRRRVFSCQKQFFVLFKFIIKNILNIFQLLVVQFGHTKLHEIVSIKILV*ALVSSQNYV*LTSN*SIK------------
Gryllus_longicercus           LILFDQKTANHLRQNLNCSINMFAPAASASLQYS*ARLEWRRGRVFSCQKLLFVLFKFIIK---THFKYITVARRPIRAYKVTWKSID*NALL---NFCFITKLRLINVILINEVQMIRV
Consensus

Teleogryllus_occipitalis      ------------------------------------------------------------------------------------------------------------------------
Gryllus_bimaculatus           ---SK*FV*GPEGEK*FRVSPVFS*LR*KRKKLVFL*NFS------TIPRRDSAKN*RAR*NKRV--------LGY*HYTKL*ILVF*C-NMLNPVWT-----CM--------QMYCL*M
Gryllus_longicercus           RSGRREVIQG--------FSSVLVGTLKTRNARVFIEFFHNTLEVIS*ELKGSVEQGGIRHCGEIGYISIIQNFKYYSYLNINMLLKSCMNLYASVLTLKV*RCTDLYGGSLGKMWVIRK
Consensus

Teleogryllus_occipitalis      ------------------------------------------------------------------------------------------------------------------------
Gryllus_bimaculatus           FKNVLIYMVEV*VKSG*LGSLGRLSENYANVG*PILLLISSIVTIDSSCREGSAEQCFASVGAVRYLKHI*VTKLHKYVF*RNVLE-FSQ*IRKK*SILHHEVRQMACHLSRLMDRKWSV
Gryllus_longicercus           FRQVI*ELCQCGIASCSTGPF--------NCHNRLVLSRGID*AIDRLCRYRTLPK-----------THMSCFKMTKIRFLEKCFRIFSMNSQRIISSPPRGASNGISFVSIETNESGSV
Consensus

Teleogryllus_occipitalis      ------------------------------------------------------------------------------------------------------------------------
Gryllus_bimaculatus           SGGRTE*VTLLLYK*YG*VP*IAVNFYGAFFYFSKT*AEYNK*YSNRL*NVQC**RVNYTD*FKEIKKNILLFLNCKSSICGQLKIN*VNKSVLPKLI*I*K-----------------Q
Gryllus_longicercus           SGGRTEQVTRLLYK*YG*VS----------YYFESVKVQF----------------------------------------VAI*QIN*VNRSPLSKLI*I*QTR*NMELLQYPRRTA*PR
Consensus

Teleogryllus_occipitalis      -------------------------------------------------------
Gryllus_bimaculatus           DKI*DC-CNIIAGR-----------------------------------------
Gryllus_longicercus           GRINDSGCDRM*MRSGQARAEQDATGQDGDGYDGTGRTRWGGTRRGGTRRSGTRQ
Consensus
```

**Tocc.8G0002370.1_exon2**

```
Teleogryllus_occipitalis    ------------------------------------------------------------------------------------------------------------------------
Gryllus_bimaculatus         GGRADRGTDARTDGRSDGRTLGRTDGRTDGETFAFN*FFSINDPQYIIMLPRPEIVLSRPYGTSRVSVKLADVSANYYGLPQKELLEAVGSAVRSAEGYSMLAEQSATSTETRLVPYSIL
Gryllus_longicercus         ------------------------------------------------------------------------------------------------------AETSATSTETRLVPNSIL
Consensus

Teleogryllus_occipitalis    ------------------------------------------------------------------------------------------------------------------------
Gryllus_bimaculatus         NNSTKYRF*LHNVLEYVRARTTRLGVTKEFCSCSSASHPTDLQ*I*SSKFARPDELSHIFTS---V*LHHL----ILWTLFATREREN------IRLNVD*GY*LATYGRNFLAA*FTPL
Gryllus_longicercus         NNSRKYGF*LHNVLEYARARTTRLGVTNEFCSSLSASHSTVLQ---------*NIVFQVCTSG*VLAYFHLDRTASLYSLDIVRYKRKGKHTLAYRLN----Y*FTTSGRNFLAA*FTPL
Consensus

Teleogryllus_occipitalis    ------------------------------------------------------------------------------------------------------------------------
Gryllus_bimaculatus         CCYIARAMPSGKGSG*DGRLAGG*NPAMF*GQP*ACFPMKIMLSRKRQKSRSGGAVRRPLRSPFGVIHENRTYDIFTSVFYSLVTRNLKKKFGAVFKKIQKCPGDSCQILP-NFKECFNN
Gryllus_longicercus         CCSIARAMPSGKEGG*DGWLAGGCNPVMF*GQP*ACLPTQIMLSRKRQKSRSGGAFRRPLRLPSGVTHEK--------------------------------SDLCHLYKCNLLSCYKK
Consensus

Teleogryllus_occipitalis    ------------------------------------------------------------------------------------------------------------------------
Gryllus_bimaculatus         LKTSTRARWSGVSSICKKY------------------------YVWFR---------CLWCSYTGRRTDR*MDGG*E----VH*KNVFGIREPQNVYFR*NFGINFPQHCITFPILRKL
Gryllus_longicercus         FKEKIWSRFQ*ISQMSRRFVSSTPKF*RMFLTISKHRLELDGQEFLQF*KNITYGLDACGVHTQTDGRTDR*TDGRTDGEKYILKNNVFSIRT----------------CISVEISESI
Consensus

Teleogryllus_occipitalis    ------------------------------------------------------------------------------------------------------------------------
Gryllus_bimaculatus         LH-------IYV-YTRVRKNALRAIMLAITWQGAFGSEYNLP----------KSAGKTNSHPLSEGALFRDHAK*ISGE----------ITI*LKRSKHGTDNVL*C--CG*NGLFR*KT
Gryllus_longicercus         FHSIA*LSLYYVNYS-------------TYTYIYG*EKMY*EL*CWP*HGREPLGLSTTCPNRRVKPHRIH--*VKGPYLGTMRNKLLVK*LFS*KDPNMEWIMYCNAVAKTDFFDRKT
Consensus

Teleogryllus_occipitalis    ------------------------------------------------------------------------------------------------------------------------
Gryllus_bimaculatus         RKVGP*YRKCR*ILFQYLVNCTICIIV*PYSTFYWSTSPKNGLTAIRLSAFL*KSLASLP*I*FSTKLLFLNRSAKCHGSLEFETEIFLF-DRLSK**SKYELLNYL**KFSKSNYFSVK
Gryllus_longicercus         RKVGP*YRKCR*ILFQYLVNCTTCIIV*P*STFYWSMSQKNRLTALRLSTFL*KSLASKP*I*FSTKLLFLNRSAKCHGLWNFKWKCFDFID*VSD*VS-MSVLNYF**NFSKTNFF---
Consensus

Teleogryllus_occipitalis    ------------------------------------------------------------------------------------------------------------------------
Gryllus_bimaculatus         MR*K*LIFGKAQTILSREGFHIF*TQIQRTFACMYRFYSSLIVLVTFQFFH*G---------------------------------NENRKLSSTRKSFTSTNSLLPYMTVCVCVCVCA
Gryllus_longicercus         FRGK*RIFGKAQTILSREGFHIF*IQIQRTFACMYRFYSSLIFLVTFQFFH*GTEQLRNM*KENNVFESQKWQLGRGWSKIEKLSQSNENRKLTNTRKSFSSTTSLLGSTSFLDCVCVCG
Consensus

Teleogryllus_occipitalis    ----------------------------------------------------------------------------------------------------------RVE-----------
Gryllus_bimaculatus         RVCARVCLCMRVCSK*DRKEITHRKER*FRVVNTLLNGG*RKKKINN-----STQIFELSKS---LCGFIHSAVNDGVKAIVGSESRTLGQPQPE--RGWVGVGGTRLD----------
Gryllus_longicercus         ---GGVCVYMRVCGK*DRKEITHTKER*FRVVNTLLYGV*RKMKNKNILPKHSSYRSPFASSFIPLCPMV*R------RS*VARAEHLVSRHQSECRWGWGVQGWTRTEKRAA*GICEPL
Consensus                                                                                                                             *

Teleogryllus_occipitalis    ------------------------------------------------------------------------------------------------------------------------
Gryllus_bimaculatus         -----ADRKARR--VRYLRAAYRPQT-----RDTAPAE----------------------------------------------------------------PIGEGAADGANRGDIGACH
Gryllus_longicercus         IVRKHAIRRRRRR*ERWLRTAQNEGTSVNAIRNSQDKEGNKDWQSGGGGGGGGGGGGGGGGGGGGGGGGGGGGGGGGGGGGGGGGGGGVGGVGGVGGVGGVGGVGGGCGGCGGCGGCGCG
Consensus

Teleogryllus_occipitalis    --------------------------------------------------------------------------EYG------------------------KG-----------------
Gryllus_bimaculatus         *E*PRQRRK*RLAKRRRRHREKDGEEMNIESGRK*RVCRRVRRVSGGECGGK**EGRKEGNGKEREVYWKRVHREVGGECEEEWRRACLEDNRERVGESVERGKKDSDENIWRKQWAKET
Gryllus_longicercus         GGGGIVEKMVKEKE*I*RVEECGKGM*DSAEG*WRRVWMKVRR----------EGRKEGIGKGREVYW-----EGGKECIERLEVNVKRGGVERVWRTIGRG--------WRRVWKEGK
Consensus                                                                             * *                         *

Teleogryllus_occipitalis    ------------------------------------------------------------------------------------------------------------------------
Gryllus_bimaculatus         M*LAKL---------NKFCFRDEFVRWQRQLKGEKEIVLVVEER*R*WRMRKQHRSYSCCGVGVAITIRGRLAAIYTVRVSPRRH-----------FAYRD-------------ETTMSM
Gryllus_longicercus         RIVMKIVGGNVGEGDNVAGKTAQILLWR*IC----QVAAAAEERER*CL*SKSGRDGGECGSNFDRT-------VVVAWVLP*L*EEGWWQYIPCGFSLSDTCV*GCDIHVDVFKTSM-F
Consensus

Teleogryllus_occipitalis    ------------------------------------------------------------------------------------------------------------------------
Gryllus_bimaculatus         LSKHRCFCRYIYVLNIDVFLTTMFLLFRCSVSE*KIPMTTIQ*KLKYEIITDSKHCSSCVGTEGTLLVLPSWTTLMSPDMASIPLYFLFFIFNSHIEIWPTNTLWA*GLPLLPVAVFGEL
Gryllus_longicercus         L*IHGCF-KHRRV-NTDVFMTSIFLLFRCSVSE*KIPMLTIQ*KLKYEIIPDSKYCCSCVGTEGTLFVLPSWTTLMSPDMTSMPLYFLFFIFISHIEMWPTSTLWA*GLPLLPVALFGEL
Consensus

Teleogryllus_occipitalis    ---------------------------------------------------------------MKKSAG---------------------------------------------------
Gryllus_bimaculatus         INTHTVKLNFNAHEFNVRGYYSSLETC*KTSMVFSRCIFIIDALVLNIDVC*LDDLLWMSHHYLRKSRGTGGVAAAAAATLRLIPIQRRVASG----------------RARQGHCLNVS
Gryllus_longicercus         INTNTVKLNFNAHEFNVRGYSSSLETC*KTSMVFSRCIFNIDALVLNIDVC*LDDLLWMSHHYLRKSGGQRG-AEAASPQLRPRPSA*SRSSGAWPLDAPGKAAA*MPP*TPPSTCLGIT
Consensus                                                                     ** *

Teleogryllus_occipitalis    ------------------------------------------------------------------------------------------------------------------------
Gryllus_bimaculatus         LNSAFYLSWKNMRNSGFEC--------*VRSCYVRRCNA*CFQLLSTGGTKVIKTMDFHETHGKNPSYASLRRRSKASCPML*ISCV*KILKSKRHLRGKLKSPVSSPG*YDLCLPYGER
Gryllus_longicercus         FETV-------VSNVKFEVVSFGDVTLSVSNC*VRGVQTRSKPWISK------------EDKEKNPSYASLRRRSIASCSML*IYCV*KILKPIRHLRGKLKSPVSSPE*YDLSLP----
Consensus

Teleogryllus_occipitalis    --------D---------------------------------------------------------------------------------------------------------------
Gryllus_bimaculatus         SAAKFAWSDGRAGLMT*PSPQKIWHN*SELNNE*QSQ*RLTIF------SF*LTNLRVLGNCDLDYNKF*HILTF*NNK*FLLFTSASSVL*KKA----HCEPVSII-----------*P
Gryllus_longicercus         --------DGKADLTTRASPQKIRHN*SELNNE*QPQ*RITIHLVDKTKSFR*LRLRL-----------*QVLTYTN------------ILK*QELPSFHFNFISIVKESAL*ASINNLA
Consensus                           *

Teleogryllus_occipitalis    ------------------------------------------------------------------------------------------------------------------------
Gryllus_bimaculatus         LF*LNKKTANRLRQNLKCSINICAPAAATSLQYS*TRLEWCRRRVFSCQKQFFVLFKFIIKNILNIFQLLVVQFGHTKLHEIVSIKILV*ALVSSQNYV*LTSN*SIK-----------
Gryllus_longicercus         LILFDQKTANHLRQNLNCSINMFAPAASASLQYS*ARLEWRRGRVFSCQKLLFVLFKFIIK---THFKYITVARRPIRAYKVTWKSID*NALL---NFCFITKLRLINVILINEVQMIRV
Consensus

Teleogryllus_occipitalis    ------------------------------------------------------------------------------------------------------------------------
Gryllus_bimaculatus         ---SK*FV*GPEGEK*FRVSPVFS*LR*KRKKLVFL*NFS------TIPRRDSAKN*RAR*NKRV--------LGY*HYTKL*ILVF*C-NMLNPVWT-----CM--------QMYCL*M
Gryllus_longicercus         RSGRREVIQG--------FSSVLVGTLKTRNARVFIEFFHNTLEVIS*ELKGSVEQGGIRHCGEIGYISIIQNFKYYSYLNINMLLKSCMNLYASVLTLKV*RCTDLYGGSLGKMWVIRK
Consensus

Teleogryllus_occipitalis    ------------------------------------------------------------------------------------------------------------------------
Gryllus_bimaculatus         FKNVLIYMVEV*VKSG*LGSLGRLSENYANVG*PILLLISSIVTIDSSCREGSAEQCFASVGAVRYLKHI*VTKLHKYVF*RNVLE-FSQ*IRKK*SILHHEVRQMACHLSRLMDRKWSV
Gryllus_longicercus         FRQVI*ELCQCGIASCSTGPF--------NCHNRLVLSRGID*AIDRLCRYRTLPK----------THMSCFKMTKIRFLEKCFRIFSMNSQRIISSPPRGASNGISFVSIETNESGSV
Consensus

Teleogryllus_occipitalis    SGG---------------------------------------------------------------------------------------------------------------------
Gryllus_bimaculatus         SGGRTE*VTLLLYK*YG*VP*IAVNFYGAFFYFSKT*AEYNK*YSNRL*NVQC**RVNYTD*FKEIKKNILLFLNCKSSICGQLKIN*VNKSVLPKLI*I*K----------------Q
Gryllus_longicercus         SGGRTEQVTRLLYK*YG*VS----------YYFESVKVQF---------------------------------------VAI*QIN*VNRSPLSKLI*I*QTR*NMELLQYPRRTA*PR
Consensus                   ***

Teleogryllus_occipitalis    -------------------------------------------------------
Gryllus_bimaculatus         DKI*DC-CNIIAGR-----------------------------------------
Gryllus_longicercus         GRINDSGCDRM*MRSGQARAEQDATGQDGDGYDGTGRTRWGGTRRGGTRRSGTRQ
Consensus
```

**Tocc.8G0002370.1_exon3**

```
Teleogryllus_occipitalis    ------------------------------------------------------------------------------------------------------------------------
Gryllus_bimaculatus         GTGGQGDGRSDGRTLGRTDARTDGRTDGRRDFCLQLIFQHKRPTVYNYASEAGDCTI*AVWNESRLSQVG*RFC*LLWPSSKRTAGGSRLSSTLCGRLQYVGRTVSHLDRDAPRPV*HTK
Gryllus_longicercus         ---AETSATSTETRLVPNSILNNSRKYG---F*LHNVLEYAR---------------------ARTTRLG-------------------VTNEFCSSL-----SASH-------------
Consensus

Teleogryllus_occipitalis    ------------------------------------------------------------------------------------------------------------------------
Gryllus_bimaculatus         *FHKI*ILTSQRVRVRTRAYYALRGYKRVLFVFECLPSNRLAMNIVFQVCTSR*ALAYFHLRLTASPYSLDIVRYKRKGKHTLECRLRLLVGYLRKKLSCGIVYPSLLLHCS-GHAFWEG
Gryllus_longicercus         -------------------------------------STVLQ*NIVFQVCTSG*VLAYFHLDRTASLYSLDIVRYKRKGKHTLAYRLNY*FTTSGRNFLAA*FTP---LCCSIARAMPSG
Consensus

Teleogryllus_occipitalis    -------DGGGGGG----------------------------------------------------------------------------------------------------------
Gryllus_bimaculatus         KRVRRTAGRWIESG-----------DVLRTTVSLLPNENHVESEETKIEERRCSQAATTIALR-----SHS*KSDL*HLYKCILLSCYKKFKEKIWSRFQKNSKMSRRFMSNSPKF*RMF
Gryllus_longicercus         K--EGG*DGWLAGGCNPVMF*GQP*ACLPTQIML-----------SRKRQKSRSGGAFRRPLRLPSGVTHE-KSDLCHLYKCNLLSCYKKFKEKIWSRFQ*ISQMSRRFVSSTPKF*RMF
Consensus                                 *

Teleogryllus_occipitalis    ------------------------------------------------------------------------------------------------------------------------
Gryllus_bimaculatus         ----*QSQNIDSS*MVRSFFNL*KILRMV*VPVVFIHGQTDGQ----IDGRRVRSTLKKRFWYQGTPKRVFPLKFRNQFS-----------TALHNFP----YIT*ITSHIRIYTGKKKC
Gryllus_longicercus         LTISKHRLELDGQEFLQF*KNITYGLDACGV-----HTQTDGRTDR*TDGR--------------TDGEKYILK-NNVFSIRTCISVEISESIFHSIA*LSLYYVNYSTYTYIYG*EKMY
Consensus

Teleogryllus_occipitalis    ------------------------------------------------------------------------------------------------------------------------
Gryllus_bimaculatus         IESYNAGHNMAGSLWV----------------*VQPAQIGG*NQFASTK*RGLI*GPCEINIW*NNYLVEKV-------QTWNG*CIVMLRLKRTFSLENS*SGALISKMSINLVS--IS
Gryllus_longicercus         *EL---------*CWP*HGREPLGLSTTCPNRRVKPHRI------------H*VKGP-YLGTMRNKLLVK*LFS*KDPNMEWIMYCNAVAKTD-FFDRKTRKVGP*YRKCR*ILFQYLVN
Consensus

Teleogryllus_occipitalis    ------------------------------------------------------------------------------------------------------------------------
Gryllus_bimaculatus         CKLYYLYHSVAVLYVLLVHVTKKWAYG----YTSVRLSV----KIAS----IFTVDIIFNKIVISQSIGQMSRFS*I*NGNI----FI*SIK*VIE*V*AFKLFIMKILKIKLFFGENEM
Gryllus_longicercus         CTTCIIV*P*STFY---------WSMSQKNRLTALRLSTFL*KSLASKP*I*FSTKLLF--------LNRSAKCHGLWNFKWKCFDFID*VSD*VS---------MSVLN---YF**NFS
Consensus

Teleogryllus_occipitalis    ------------------------------------------------------------------------------------------------------------------------
Gryllus_bimaculatus         KIAHFRKSSNNTFARRVS---------YFLNPNPTNICMY--VQILLF-------------------PNSLSHISILPLG**EQKIK*Y------QKVIY*HQ*F---------TSLHDC
Gryllus_longicercus         KTNFFFRGK*RIFGKAQTILSREGFHIF*IQIQRTFACMYRFYSSLIFLVTFQFFH*GTEQLRNM*KENNVFESQKWQLGRGWSKIEKLSQSNENRKLTNTRKSFSSTTSLLGSTSFLDC
Consensus

Teleogryllus_occipitalis    ----GGG-----------------------------------------------------------------------------------------------------------------
Gryllus_bimaculatus         VCVCVCV-CACVRAC--------------------VLVYACV**MR-----PQRN---------------------YSQKGKII*SSQHSVKW------WLKEKKNK*FYPNIRVIEVP
Gryllus_longicercus         VCVCGGGVCVYMRVCGK*DRKEITHTKER*FRVVNTLLYGV*RKMKNKNILPKHSSYRSPFASSFIPLCPMV*RRS*VARAEHLVSRHQSECRWGWGVQGWTRTEKRAA*G-----ICEP
Consensus

Teleogryllus_occipitalis    -------------------------------------------SGGGGGGGDGGRGGSGGGGGGGGGGGGGGGGGEGGGGGGGGGGGGGGGG---------GGGGGGGGGGGGGGG
Gryllus_bimaculatus         L--RVHSF-----RCE*WC-----EG-------DRR*REQNT-WPAA--------TRARVGGGGGYKVGRGQKSAPREVSASRLSSANTRYGAG*ADRR-----GGCGRRKPRGHRCMPL
Gryllus_longicercus         LIVRKHAIRRRRRR*ERWLRTAQNEGTSVNAIRNSQDKEGNKDWQSGGGGGGG----GGGGGGGGGGGGGGG-----GGGGGGGGGGGGGGGGVGGVGGVGGVGGVGGVGGVGGCGGC
Consensus                                                                   *****   * *                   * *          ** *     *

Teleogryllus_occipitalis    GGGGGGGGGGGGGG----------------------------------------------------------------------------------------------------------
Gryllus_bimaculatus         G-------------IAKTKKEIKIGKAAATAS*KRWRRNEYREWKKVKGM*ESAES*WRRVWRKVMR-----G*KRGK---WKGKRSILEK---SA*RG-----WR*M*RGVEESVFGG
Gryllus_longicercus         GGCGGCGGCGGGGGIVEKMVKE---------------KE*I*RVEECGKGM*DSAEG*WRRVWMKVRREGRKEGIGKGREVYWEGGKECIERLEVNVKRGGVERVWRTIGRGWR------
Consensus                   *

Teleogryllus_occipitalis    ---------------------------------------------------------------GRGGGG*------------------------------------------------
Gryllus_bimaculatus         Q*GEGGGECGKRKKGQ**KYLEETVGEGDNVAGKTEQILLSR*ICQVAAAAEGRER---DSASGRRAVEIVANAEATSIVQLL---------WR--GCCHNYKRKVGGNIYRAGFPSATL
Gryllus_longicercus         -------RVWKEGKRIVMKIVGGNVGEGDNVAGKTAQILLWR*ICQVAAAAEERER*CL*SKSGRDGGECGSNFDRTVVVAWVLP*L*EEGWWQYIPC---------------GFSLSDT
Consensus                                                                  **

Teleogryllus_occipitalis    ------------------------------------------------------------------------------------------------------------------------
Gryllus_bimaculatus         CV*G*DNDVDAFKTSMFL*IHICFKHRCFSNDDV---SSFPMF-CF*MKNSHDHNTVKVKI*NN------------N***TLLFM-CWNRRNAACTTLL--DHTNVARYGLHTLVFPLLH
Gryllus_longicercus         CV*GCDIHVDVFKTSMFL*IHGCFKHRRV-NTDVFMTSIFLLFRCSVSE*KIPMLTIQ*KLKYEIIPDSKYCCSCVGTEGTLFVLPSW-------TTLMSPDMTSMPLYF-------LFF
Consensus

Teleogryllus_occipitalis    ------------------------------------------------------------------------------------------------------------------------
Gryllus_bimaculatus         IQFAH*DL---ANKYAVGIGSAPPSRSRIRRIDKHAYCQIKF*CT*I*C*RLLLQSRNMLKNIDGF*S--------LYFYHRCFSFEHRCLLTR*FVMDVASLLA*EPRHRRRRRSCGR-
Gryllus_longicercus         IFISHIEMWPTSTLWA*GLPLLP----------------VALFGELINTNTVKLNFNAHEFNVRGYSSSLETC*KTSMVFSRCI-FNIDALVLNIDVC*LDDLLWMS--HHYLRKSGGQR
Consensus

Teleogryllus_occipitalis    ------------------------------------------------------------------------------------------------------------------------
Gryllus_bimaculatus         -----DPPLDPDPAA----RGLW-----------TRPARPLLECLPKLRLLLVLEEHAKQWLRMLSSKLLRSEM*RLVFPTVKY----WGN----------KGDQNHGFP*NTWKKSFVR
Gryllus_longicercus         GAEAASPQLRPRPSA*SRSSGAWPLDAPGKAAA*MPP*TPPSTCLG-----ITFET---------------------VVSNVKFEVVSFGDVTLSVSNC*VRGVQTRSKP---W------
Consensus

Teleogryllus_occipitalis    ------------------------------------------------------------------------------------------------------------------------
Gryllus_bimaculatus         LPPEEK*SQLSHVIDFLRVKDSQV*KTPPREIKIAR-------FITRVIRPVSAIR*KERS*ICVVRRKSRP--HDVTLAT---------------ENMA*LK*IEQ---*ITISMKAYN
Gryllus_longicercus         ISKEDKEKNPSY--------------ASLRRRSIASCSML*IYCV*KILKPIRHLRGKLKSPV------SSPE*YDLSLPDGKADLTTRASPQKIRHN*SELNNE*QPQ*RITIH-----
Consensus

Teleogryllus_occipitalis    ------------------------------------------------------------------------------------------------------------------------
Gryllus_bimaculatus         F*FLVDKSKSFR*LRLRL*QVLTHTNILK**VIPTFHFSFISIVKESAL*ASINYLTLILIE*KNC*PIKTKS*MFYKYLCPSCGHVSSIFLN*IRMVSPTGIFMS--KTIFCAF*VHNK
Gryllus_longicercus         ---LVDKTKSFR*LRLRL*QVLTYTNILK*QELPSFHFNFISIVKESAL*ASINNLALILFDQKTANHLRQNLNCSINMFAPAAS--ASLQYS*ARLEWRRGRVFSCQKLLFVLFKFIIK
Consensus

Teleogryllus_occipitalis    ------------------------------------------------------------------------------------------------------------------------
Gryllus_bimaculatus         KHFKYIPVARRPIRAYKVT*NSID*NTRLSFSFITKLRLINVKLINQVQMIRVRPGRREVIQGFSSVLVVALKTQKARVFIEFFHNTAEGFS*ELKGSVEQEGIRL------------LA
Gryllus_longicercus         THFKYITVARRPIRAYKVTWKSID*NALLNFCFITKLRLINVILINEVQMIRVRSGRREVIQGFSSVLVGTLKTRNARVFIEFFHNTLEVIS*ELKGSVEQGGIRHCGEIGYISIIQNFK
Consensus

Teleogryllus_occipitalis    ------------------------------------------------------------------------------------------------------------------------
Gryllus_bimaculatus         LYKTLNISLLV*YVKSSMDVYANVLSLNV*KCTDLYGGSLGKIWVIRKFR*VI*ELCQCGIAYSTTDLFNCHNRLVLSRGIG*AMFRLCRCRTLPKTHLS-YKITQIRFLEKCFRIFSMN
Gryllus_longicercus         YYSYLNINML---LKSCMNLYASVLTLKV*RCTDLYGGSLGKMWVIRKFRQVI*ELCQCGIASCSTGPFNCHNRLVLSRGID*AIDRLCRYRTLPKTHMSCFKMTKIRFLEKCFRIFSMN
Consensus

Teleogryllus_occipitalis    ------------------------------------------------------------------------------------------------------------------------
Gryllus_bimaculatus         SKKIINSPSRGASNGMSFVSIDGPEVVC*WWKNGVGYSTTIQIIRLSPLNCGQFLRCFFLFFENISRV**VIF*SLVKCTVLITG*LYRLI*RNKEKHLIIFKL*KFNLWPVKNKLSK*K
Gryllus_longicercus         SQRIISSPPRGASNGISFVSIETNE------SGSVSGGRTEQVTRLLYK*YG----*VSYYFESV-----------------------------KVQFVAI*QIN*VNRSPL--------
Consensus

Teleogryllus_occipitalis    ----------------------------------------------------------------------------
Gryllus_bimaculatus         CSSKVNLNIKTR*NMRLLQYHRRT----------------------------------------------------
Gryllus_longicercus         --SKLI*I*QTR*NMELLQYPRRIA*PRGRINDSGCDRM*MRSGQARAEQDATGQDGDGYDGTGRTRWGGTRRGGTRRSGTRQ
Consensus
```

**Tocc.8G0009300.1**

```
Teleogryllus_occipitalis    GGCAGCGGC---------------------------------------------------------------------------------------------------------------
Gryllus_bimaculatus         GGCATTGGCCACAGAGAAGCGGATAGATGTTTAGGCATCAACCGATTTGCAGTATCAAGGACTTTCTAAGCATCGGTTCTGACTACGCAACGTGTTAGAGGAAGTGATTCGCTGCAAATA
Consensus                   ****  ***

Teleogryllus_occipitalis    ------------------------------------------------------------------------------------------------------------------------
Gryllus_bimaculatus         TGAATCAAGATTTACTCCCTTTTTGGTTACCAGGAGTAGCCAACAAGAGTTTTTCTGTCGAAAAATGGCCAATTTCCAGCGCACGATTACAAAACTGTGTTTTCATGGGAATTAAGGCGA
Consensus

Teleogryllus_occipitalis    ------------------------------------------------------------------------------------------------------------------------
Gryllus_bimaculatus         GAAATTAAAAAAAAGCCTGCTCAACGAACGGATGTCTTGCCTGATAAAAATATAAAATTACACTTTGATTTATAATCAGATTATTATTAATTAATATTTTCACTAAAATTACTGAATATG
Consensus

Teleogryllus_occipitalis    -------------------------------------------------------------------------------------------GGTGTTAATGGCGCGAAGCGA--------
Gryllus_bimaculatus         CATATTGATGGAGAATCCTCTTCAATAATTATTTAATAGCAAATTTAAAAAAAAGTCGATATACCACATATTTTTGTAGAATTTCTGATTTGATATTGCAGACGCCATGTGATGTTTAAT
Consensus                                                                                              * * **   * *** * * **

Teleogryllus_occipitalis    ---------------------------------------------------------------------------GGCAGCCCAT------------------------------------
Gryllus_bimaculatus         CCATTGCTTCTCCTTAATATTTATGAAATGTAATTATGCCGACAATTAGAGTTAGTGGTATACATTATATTTGTAGAAGCCCATTCTTAATCCCGCTTATTTCACATTGACGTGTCTCAT
Consensus                                                                                    * *******

Teleogryllus_occipitalis    ------------------------------------------------------------------------------------------------------------------------
Gryllus_bimaculatus         TTAACTTTTTCTGAAATTTGAGAATAGCACATTATTAAAAGGATTTTTTGTTGAGAATCAATGTATCAGCTAATGAATAATTAAGTTTTTGAGTTCACTCGCTATAATTTCCTGCCAACA
Consensus

Teleogryllus_occipitalis    ------------------------------------------------------------------------------------------------------------------------
Gryllus_bimaculatus         ATTTTACCGTTCATCACACAACCAACTCTTCTCTCCGCGGACGCTACGCTAGAGTTAATTTCGCCACTGTGATGAAAATGTGATGGGGGTGCAAATGCGGAAAGCGACACCAAGTGTGGA
Consensus

Teleogryllus_occipitalis    --------------------------------CTGCGGCGTCGGGAGAAGAGGGGGCCACGGCGAGCGTTGCAGCCGCCGC---------------ACCGCGACG-------------
Gryllus_bimaculatus         GGAACGTTGCAGGTAATGGCGAGTGGTAGGTGTCGCGTCGCGGGGAACAGAAACGGCGGCGGCTACGGCGGCAGCGGCGGCGGCGGTGGCGGCAGCGGCGGCGGCGGTGGCGGTGGCGGT
Consensus                                      *** **  ****  ***   ***  **** *  *  ***** ** **                *  *** **

Teleogryllus_occipitalis    ------------------------------------------------------------------------------------------------------------------------
Gryllus_bimaculatus         GGCGGTGGCNNNNNNNNNNNNNNNNNNNNNNNNNNNNNNNNNNNNNNNNNNNNNNNNNNNNNNNNNNNNNNNNNNNNNNNNNNNNNNNNNNNNNNNNNNNNNNNNNNNNNNNNNNNNNNN
Consensus

Teleogryllus_occipitalis    ------------------------------------------------------------------------------------------------------------------------
Gryllus_bimaculatus         NNNNNNNNNNNNNNNNNNNNNNNNNNNNNNNNNNNNNNNNNNNNNNNNNNNNNNNNNNNNNNNNNNNNNNNNNNNNNNNNNNNNNNNNNNNNNNNNNNNNNNNNNNNNNNNNNNNNNNNN
Consensus

Teleogryllus_occipitalis    -----------------------------------------------------GGCGGGCGAGTTCAGTGAAGTTGGCTGGAGCTGTTGCTATAAGCGGCGCAAGTAGTTGCAGTGACGG
Gryllus_bimaculatus         NNNNNNNNNNNNNNNNNNNNNNNNNNNNNNNNNNNNNNNNNNNNNNNNNNNNNGAGGCGGTCGAGTTGGTTGAAGTTGGCAGGAGCCGTTGCTACAAGCGGCGCAAGTAGGCGCAATGAGGC
Consensus                                                                   ***** ******   ********** ***** ******* **************  *** *** *

Teleogryllus_occipitalis    GCTCGCTCGTGTAAGGACCAGTGGCGAATCCTTAACATTAATTTAAGCACACCGCACTTTATGGAAGCTACCTCTACCGTTGTTTAACTGGCTCTCGGTAGAAAAACAGAATTATTTTGC
Gryllus_bimaculatus         GC----------------------------------------------------------------------------------------------------------------------
Consensus                   **

Teleogryllus_occipitalis    TTCGCTCGTATAAGAATGTCACGAGCATCGCAGTTGCTCGTCAACAACAGAAACAGGACTCGTAAGATACGTCT----------------------------------------------
Gryllus_bimaculatus         -TCGCTCGTATAAGAACGACACGAGCCTCACCGCTGCTCGACAACAGGAAAAACACCTCTCATAAGATACGTTTGTACGTAACGTACGTACATCACGTACGTACGTTACGTACATCAAAC
Consensus                    *************** * ******* ** * * ****** *****  * *****   *** ********** *

Teleogryllus_occipitalis    ---------GTTTAATTTAAAACATCATTAACTCGTTGGGTAAGGTTAAGATTCATCCAGATATTTATAAAATGTCACTTAATTTTAAATATTTCGTAGCAGGCAACAGGTGTTTACTAC
Gryllus_bimaculatus         GTAATGTACGTTTAATGTGGAATGCCATTACTTCGTTGGGAAAGGTTAAGGTAGCTCCAGACATTTATAAAATATAAATTAA-CTTAAATTTTTTTCGCATGCAACAGGTGTTTGATAG
Consensus                            ******* *  **   *****  ********* ********* *   ****** *********** * * ****  ****** ***  * *** ************  **

Teleogryllus_occipitalis    ATTTATTCAAATGCAGCACATAACCATCGCGGATCTCTAGAGTAAAATTTTTAAATGGTTTTGTTCATGATCTGTTT-TTAATGATGGTAGAATTACAGCTGAGAGATGAGACGAATTAA
Gryllus_bimaculatus         AACTATTCAAATACAGCCCCAAAACATCTAGGACTTGTTAAGTTAATTTATTAAATGATTTTGTATGTTAACTGCTCATTAATGGTAGTAGAGTT-----------------------A
Consensus                   *  ********* **** *  ** ****  ***  * *  *** ** ** ******* ******   * * *** *  ****** * ***** **                       *

Teleogryllus_occipitalis    TCAGGTTAATTATTTAAAAAAACGTGTTAAACAGGTTATCGTGTTAAAGAACGTGAACGCAGCAGAGCTGACAAATAGA----------GATAATTACCCAGAAAACGGAAAATAAACAA
Gryllus_bimaculatus         TTAGACTAATTAT---AAAAAAATGGGCAAACAGGTTTTTGTGTAAAATAACGTCAACGCAGCAGAGCTGG-AAACAGATCCTTCGTGGACTAATACCCTCGAAGACGGAACAAAAATAA
Consensus                   * **  *******   ******   *  ********* * **** *** ***** ***************  *** ***          ****  **  *** ****** * *** **

Teleogryllus_occipitalis    ATAAATTAGTGGCGAGAAAAAAGAGATAACTAAGGAGAAAGAAAAAAGGGAAACCGAGAAGTATAAGAAGAAAAACAACAAGAGAACGGCAGGAAAAAACACTGATGTGAAGAATCGTAA
Gryllus_bimaculatus         AGAAATTATTAGTGTGAAAAAAGTGATAACTGATGGGAAAGAATACGAGGAACCC-------------AGAGGTATGACAAGGTACCTAGAAGTAGGATGATTGAAATTAGGA----TGG
Consensus                   * ****** * * * ******** ******* * * ******* *   **** **             ***   *  *****  * *   * * *  *  * ***  * * **    *

Teleogryllus_occipitalis    TATCAGTAGGAAAGATAGAAACGAAGGAATAAGGCAAAAAGGCTGAGGAGGAAGAAACGAGGTACGCTGGGAGAGCTGAAATCTCAGTAAGAAAGAATAGGCGAGAAATATCGGTCATGA
Gryllus_bimaculatus         TAGTAGTAGGGAGGATTGAAACACAGGTATACTGCAGAAAGTCTGGGGAAGAGGAAACGGGATAGAATGGGTGTGAGGCACC-----TAAGACAGAAACGCCGAGAAAGATCGGTCATAA
Consensus                   **  ****** * *** *****  *** ***  **** **** *** *** ** ****** * **   **** * *  * *       ***** ****  * ******* ********* *

Teleogryllus_occipitalis    CTACAATGAAAATTACAGCAAAGGAAGTAGAAATCTGCGCGACTATTCCGCCTCTTGCTTGGAAGTAAAGTGGGGCGCATTAGAAGTTTCGC-CGCGAGGCG---------TCGTCGGAG
Gryllus_bimaculatus         TCATAATGAAAAGTACAGCAAAGGCAGCAGAAATCTTTGCGAGCATTCCGCCTCTTCCTTGGAAGTGCAGTGGGGCGCATTAGAAGTATGGTGCGCGAGGCGGCTGTCGCTTCCTCGGAG
Consensus                     * ******** *********** ** ********  ****  ************ *********  ****************** * *  *********         ** ******

Teleogryllus_occipitalis    TCAGCCTGGCCCCATCAGTGTGCACATCAGCGGCGTCAACGACCGCAGTGGCGGCGCGCGTTATCCCTACTCCCCTCCTTGGGGGTCGATAGGTGCCGTCGCTCGCGCCGCCCGGGCAAT
Gryllus_bimaculatus         TCAG-CTGACCCCATCAGCGCGCACATCAGCGGCGTCAACGACCGCAGTGGCGGCGCGTTGTGTCCCTCCTTCCCCCCTCGGGGGTCGATAGGCGCCGCCGCCCG--------------
Consensus                   **** *** ********* * *************************************   * ***** ** *** *** ************ **** *** **

Teleogryllus_occipitalis    CGGACTGTGTGCGGCGGGCGGCGGCGGGCGGGTGGTGGCGGGCGGTGTGGTGGTGGCAGGGTGGGCGGGGGTGGGTGGAGGCGGGCGTGCGGCGGTGGGCGGGCGGCGGCGGGTGGGCGG
Gryllus_bimaculatus         --------------------------------GGCGACGCGCGAT---------------------------------------------------------------------------
Consensus                                                   ** * ** *** *

Teleogryllus_occipitalis    GCGGCGGCGGGTGGGCGGGCGGCGGCGGGGGGGCGGGCGGCGGCGGGCGGGCGGGCGGCGGCGGGGTGCGTGGTGGCGGCGGGTGGGCGGGCGGCGGCGGCTGGTGGGGGGGCGGCGGCG
Gryllus_bimaculatus         ------------------------------------------------------------------------------------------------------------------------
Consensus

Teleogryllus_occipitalis    GGGGCGGGCGGCGGGGGTGGGCGGGGCGGCGGGGGGTGAGCGGGCGGCGGCGGGTTGGTGTGGATGGGCGGGCGGCGGCGGGTGGGCGGAGTGGGCGGACGGTCGGCGGGTGGGGCGGGT
Gryllus_bimaculatus         ------------------------------------------------------------------------------------------------------------------------
Consensus

Teleogryllus_occipitalis    GGGCGGGTGGGCGGGCGGGCGGGCGAGCGAGCGAGAGGGCGAGCGGGCGAGCGGTCGGGAGGGCGAATGGCGGGCGACGGTGCGGGCGGGCGGTGGCGGCGGGCGGGCAGGCGGCGGGCG
Gryllus_bimaculatus         ------------------------------------------------------------------------------------------------------------------------
Consensus

Teleogryllus_occipitalis    TGGGTGGGTTGTGGGTGGGCGACAGGCTGCGCTGCGAGGAGAGGGCGATGAGGGCGAGAGGGCGAGGGGGCGAGAGGGCGAGAGGGCGAGTGGGGAGAGAGGGGGCGAGGGGAGAGGGCG
Gryllus_bimaculatus         ------------------------------------------------------------------------------------------------------------------------
Consensus

Teleogryllus_occipitalis    AGAGGGCGAGAGGGCGAGCGGGGGGGCGGGCGGGCGGGCGGGCGGACGGGCGGGAGGGTCGGCGGGCGGCGGCGTGTGGGCGGGCGGCGGCAGCGGCGGCGGCGGCCCCGCAACCTGACA
Gryllus_bimaculatus         ------------------------------------------------------------------------------------------CGGCGGCGGCGCCAGCCCCGCAACCTGACA
Consensus                                                                                                             * ********* * ****************

Teleogryllus_occipitalis    CCATGCCGCGCCGCGGGCCGCGTAAACAAATATGGGCAGCGGTGCGGCGCCGCCAGGCGGGCAATCAATACCCACGCGAGCGGACGCCCGCGCTCATTCTCTCCCCC-------------
Gryllus_bimaculatus         CCGCGCCGCGCCGCGGGCCGCGTAAACAAATACGGGCAGCGGTGCGGCGCCGCCTGGCGGGCAATCAATACCCACGCGAGCGGACGCCCGCGCTCATTCCCTCTCCCGTCCATCTCCCGT
Consensus                   **   *************************** ********************* ******************************************** *** ***

Teleogryllus_occipitalis    ------------------------------------------------------------------------------------------------------------------------
Gryllus_bimaculatus         CCCTTTCCGTCCCTCTCCTCTCCCTCTCCCGTCCCTTTCCATCCCTCTCCCGTCCCGTCCCTCTCCCGTCCCTTTCCGTCTCTGTCCCGTCCCTTTCCCTCGTCCCATCCCTCTCTCTCC
Consensus

Teleogryllus_occipitalis    ------------------------------------------------------------------------------------------------------------------------
Gryllus_bimaculatus         CTTTCACTCGTCCCGTCCCGTCCTGTCCCGTCTCGTTAGGTCCCGTAATGTCCCGTCCCGTCCCTCTCCCTCTCCCTCTCCCTCTCCTGTCTCGTCCCGTATCGTCCCGTCCTGTCTCGT
Consensus

Teleogryllus_occipitalis    ------------------------------------------------------------------------------------------------------------------------
Gryllus_bimaculatus         CCCGTCCCGTTAGGTCCCGTCCCGACCCTCTCCCGTCCCTCACCCTCTCCCTTTTCCTCTTCCCGTCCCGTCTTGTTAGGTCCCGTACCGTCCCGTCTCGTCTCGTCCCATCCCATTAGG
Consensus

Teleogryllus_occipitalis    -------------------------------------------------CAGCCGCCCTCCCGGCGCGTCC-------------------------------------------------
Gryllus_bimaculatus         TCCCTCCCGACCCTCTCCCGTCCCTCTCCCTCGTCCCGTCCCGTCACTTCAAGTACCGTCCCGTCCCGTCCCTGTCCCGTCCCTATCTGTCCCTTTCTGTCCCTCTCCTGTCCCTCTCCC
Consensus                                                                    **    ** ***** * *****

Teleogryllus_occipitalis    ------------------------------------------------------------------------------------------------------------------------
Gryllus_bimaculatus         GTCCCTTCCCGTCTCTCTCCCGTCCTGGCCCTCTCCCATCCCTTTTCCTCGTCCTGTCCCTCTCTCTCTCTTTCCCTCGTTCCGTCCCGTCCCGTCCTGTCTCGTTAGGTCCCGTCCCTC
Consensus

Teleogryllus_occipitalis    ------------------------------------------------------------------------------------------------------------------------
Gryllus_bimaculatus         TCCCTCTCCCTCTCCCTCTCCCTCTCCCTCTCCCTCTCCGTCCCGTCTCGTCCCGTCCTGTCTCGTCTCGTTCCGTCCCGTTTTGTCCCGTCCCGTTATGTCCCGTCCCGTTATGTCCCG
Consensus

Teleogryllus_occipitalis    ---------------------------------------------------------------------------TCCGAGATGGCAGGCAAGCGCCGGAGGCTGACAAAGGAGCTTGTGC
Gryllus_bimaculatus         CCCCGTTATGTTCCGTCCCGTTATGTTCCGTCCCGACCCTCTCTCGTCCCTCTCCGTCCCGTCCCGTCCCTCAGTTCGTGGTAGCAGCGAAGCGCCGGAGGCTGACAAAGGAGCTCCTGT
Consensus                                                                                      * ** * * ****  **************************  **

Teleogryllus_occipitalis    TAATCCATTGGGGGCAGGTGCCCGCCCCGGCGCCGCGTCGCGTCGCGCAGCCCGATCCGCGCCCCGCTGACCTCGAGAGGCGCCGCCGGCGCCCGCCTCAATGGCACCGGCGCCGCC---
Gryllus_bimaculatus         TAATCCATTGGAGACAGGT------------GCCGCGCCGCGCCGCGCCGCGCGTTCCGTGCCCCGCTGACCTCGAGAGCGCCCGCCGGCGACCGCCTCAATGGCACCGGCGCCGCTGTG
Consensus                   *********** * *****            ****** **** ***** ** ** **** *******************   ********* ***********************

Teleogryllus_occipitalis    ----GATATGAGCGGGGGCGGCGGCGGAGGTGATGTCGGCTGCGGTGGGGAGAGCAGCGTCGGCGTAGGCGTGGGTTCCGGTGGCGGGGGA--GATGGTGAAAGCGGGAGCGGCGGCGGA
Gryllus_bimaculatus         GGCGGACGCGGCGGGCGGCGGCGGCGGATGTAATATCGGTTGCCGCGGTGTGAGTGGCGTGGGCGTTGGCGTAGGTTGAGGTGGACGGCACCGAGGGTGAAGGCGGAGAGAGCGACGGA
Consensus                       **   *   ** ************ ** ** **** *** * ** * ***  **** ***** ***** ****  *****  *** *  ** ****** ****     *** ****

Teleogryllus_occipitalis    GGCGAAAGCCACGAAGGCGGAGACACTGGAGGAGACGTCTGGGAGGGGTTTTGGTGACGG---------------------------TGGTTTTGGTGCCTGCGACTTCAGGATGTCGA
Gryllus_bimaculatus         GGCGGTGACGGCGGAGGCTGTGACGGCCGAGGCGGTGACGGCCGAGG----CGGTGACGGCCGAGGCAGTGACGGCCGAGGCGGTGATCGGTGAGAGTGCCGGTGATTGCTGGACGAC-A
Consensus                   ****    *  ** **** * ***    **** *  * * *    **     ********                               ***    ***** * ** * * *** * * *

Teleogryllus_occipitalis    GGACTGTGGGCGA--GGAGGAAACGGATGGCGGACGTGGGAGTGGTAATGGGTTG---------------ATTGATTGATGTTTCCGGGTAAGTCAACTGCTACGGTCATTAGTCCGGC
Gryllus_bimaculatus         GTACGATTGGTGACGGGGGAAATGGGAAGAAGGTCCTGGTATAGGCAATGGGTTGGTTGTTTGGTGGGTTAGTTGATTGATGTTTCTGAGCACGTTATCAGCTAC------------GT
Consensus                   * **  * ** **  ** * **  *** *  ** * *** *  ** *********                *************** * * * ** * * *****            *

Teleogryllus_occipitalis    TAGTTGAATGAATGTTTTGTCACGGGAGGTGGCGG-ATAACCTCTCCCACTGGA------------------------------------------------------------------
Gryllus_bimaculatus         TAGTGCAGTGAATATTTTGTTTCGGGAGGTGGCGGTGTACCTCCTCGCACTGGATGGGGAGAACCGACGCGCGTATGTCGGCTTGCAGGGCATTTCCCTCTCGAGAATTTGCTTGAGGGG
Consensus                   ****  * ***** ******  *************  ** *  *** *******

Teleogryllus_occipitalis    ------------------------------------------------------------------------------------------------------------------------
Gryllus_bimaculatus         CACTTACGCAGCCAGTTTAAAGTTTGCCGTTCAAGGTGGGCGAGATCGTTAAGGCAGTAGTTCCGTAGCGATCTCGGACCGAAGGATCGCAGGTTCGAGTTCCGCTTCGGGCATGCTTGT
Consensus

Teleogryllus_occipitalis    ATTGACGCTCG-------------------------------------------------------------------------------------------------------------
Gryllus_bimaculatus         ATTTGTGCTCGTCCTTTCGCCCCCTTTCATCACTATAGCATGCCATCTTCACAGTTACCAAACCGTTAAATGCAATACTGGTCGCTAATAACCTAGATGATGAAGATAATGCGACCTTTA
Consensus                   ***   *****

Teleogryllus_occipitalis    -----------------------------GTCGGAAGCATAGAGCTTGTCCCG-------------------------------------------------------GTTGAGGCGGTGCC
Gryllus_bimaculatus         ATAACAAAAACGTTCTAAGCTTCCACGTCGTCAGAAGCACAGCATTTATACAGAGAATTCTCCTCGAGATAGACGCTGAATTCGACCTAGCTAGTATATAGCCAATGTCGGAAGTGCGCC
Consensus                                                *** ****** **   ** * * *                                                     ** *     * ***

Teleogryllus_occipitalis    CC----------------------------------------------------------------------------------------------------------------------
Gryllus_bimaculatus         CCCATCTGGTCTCGAACCCTAGAACCTCCCAATCAGAGTCGCAGAGTCGACAATTGTCGTCGGCAATGTGAGCTCGCAACTAAAGGGGCAGCCGGTGTGGCAATGTAGCTGTGGAAAAGG
Consensus                   **

Teleogryllus_occipitalis    ------------------GCTGGTAGGAGATTCCTGGTTG--------------------------------------------------------------------------------
Gryllus_bimaculatus         AGAAAACGCCATGCGGTGGCAGGTGCGAGAGGCTTGGGAGCAGTGACGGGTGCACCGCCTCCGTCGCCACAGGCGAAGGAGTGGGGGTGTCTGCGGCGGCGCGGGCGGTGGGCGGTGGGC
Consensus                                     ** ***  ****  * ***  *

Teleogryllus_occipitalis    --------------AGGAAAACCACGCTAGCCGACGTATAACACCATCAAGCGGGGTATGATCATCACCGCCACCTCGCGATGAAA-----------------------------------
Gryllus_bimaculatus         GTTGGAAGGTGCGCGGGGAAGGAGCAGCGAGAGGCGTCGTGCGTCAGCAAGCACAATGCTGCCATTGTGGGTTCGCCTGAATGATATCTTTAAGTGCAAACGCTTTTCGTGTTGTTGTTG
Consensus                                 ** **    *       * ***    *  ** *****    *     ***     *   *  *   **** *

Teleogryllus_occipitalis    ------------------------------------------------------------------------------------------------------------------------
Gryllus_bimaculatus         GGTTGTAGTGGGAGGGGGGAGGGGAATGAAATTCCCCTCGACAAGAACCCCAGGTCAAGTTATTTCTGTTCAGTATTCTTTGTTTGGGATTGTTCAAGCTTTCGATTTTTTGTGCTTTTT
Consensus

Teleogryllus_occipitalis    -------TACATGCACTAC-----------------------------------------------------------------------------------------------------
Gryllus_bimaculatus         TTTCGTGTACATGCATCATTCATGCTATTTTAATAACATTACGAGTCTTTTATGTGAATTAAAATTTTGTTGATATCTTTTGCTTCTCACTCCTCACTTCTCATGACAACATAGTTTTGC
Consensus                          ********  *

Teleogryllus_occipitalis    ---------------ACTTGCGGGACGAATGACCTCTGTAGTTGACGTG-
Gryllus_bimaculatus         AAAAAAAGTCATAAAACATATGAAAAGTATCTCGTTTGAAATTAATTCGT
Consensus                                  ** *  *  * * **  * * ** * ** *   *
```

**Tocc.8G0009300.1_exon1**

```
Teleogryllus_occipitalis     ------------------------------------------------------------------------------------------------------------------------
Gryllus_bimaculatus          IFTNFSENIN*L*SDYKSKCNFIFLSGKARLKNQISVSNSQGLQCV*IEWRQFWRKSAAGFMCLRFSCGIGHREADRCLGINRFAVSRTF*ASVLTTQRVRGSDSLQI*IKIYSLFGYQE
Consensus

Teleogryllus_occipitalis     ------------------------------------------------------------------------------------------------------------------------
Gryllus_bimaculatus          *PTRVFLSKNGQFPAHDYKTVFSWELRREIKKKPAQRTDVLPDKNIKLHFDL*SDYY*LIFSLKLLNMHIDGESSSIII**QI*KKVDIPHIFVEFLI*YCRRHVMFNPLLLLNIYEM*L
Consensus

Teleogryllus_occipitalis     ------------------------------------------------------------------------------------------------------------------------
Gryllus_bimaculatus          CRQLELVVYIIFVEAHS*SRLFHIDVSHLTFSEI*E*HIIKRIFC*ESMYQLMNN*VFEFTRYNFLPTILPFITQPTLLSADATLELISPL**KCDGGANAESDTKCGGTLQVMASGRCR
Consensus

Teleogryllus_occipitalis     ------------------------------------------------------------------------------------------------------------------------
Gryllus_bimaculatus          VAGNRNGGGYGGSGGGGGGSGGGGGGGGGGGXXXXXXXXXXXXXXXXXXXXXXXXXXXXXXXXXXXXXXXXXXXXXXXXXXXXXXXXXXXXXXXXXXXXXXXXXXXXXXXXXXXXXXX
Consensus

Teleogryllus_occipitalis     ------------------------------------------------------------------------------------------------------------------------
Gryllus_bimaculatus          XXXXXEAVELVEVGRSRCYKRRK*AQ*GARSYKNDTSLTAARQQEKHLS*DTFVRNVRTSRTYVTYIKRNVRLMWNAITSLGKVKVAPDIYKI*INLNFFSHATGV**NYSNTAPKHLGL
Consensus

Teleogryllus_occipitalis     ------------------------------------------------------------------------------------------------------------------------
Gryllus_bimaculatus          VKLIY*MILYVNCSLMVVELLD*L*KNGQTGFCVK*RQRSRAGNRSFVD*YPRRRNKNKEIISVKKVITDGKEYEEPRGMTRYLEVG*LKLGW***GGLKHRYTAESLGKRKRDRMGVRH
Consensus

Teleogryllus_occipitalis     ------------------------------------------------------------------------------------------------------------------------
Gryllus_bimaculatus          LRQKRRERSVIIIMKSTAKAAEIFASIPPLPWKCSGAH*KYGARGGCRFLGVS*PHQRAHQRRQRPQWRRVVSLLPPSGVDRRRRPGDARSAAAPAPQPDTAPRRGPRKQIRAAVRRRLA
Consensus

Teleogryllus_occipitalis     ----------------------------------------------------------------------------MSR-----------------------------------------
Gryllus_bimaculatus          GNQYPRERTPALIPSPVHLPSLSVPLLSLSRPFPSLSRPVPLPSLSVSVPSLSLVPSLSLPFTRPVPSCPVSLGPVMSRPVPLPLPLPLLSRPVSSRPVSSRPVRSRPDPLPSLTLSLFL
Consensus                                                                                                ***

Teleogryllus_occipitalis     ------------------------------------------------------------------------------------------------------------------------
Gryllus_bimaculatus          FPSRLVRSRTVPSRLVPSH*VPPDPLPSLSLVPSRHFKYRPVPSLSRPYLSLSVPLLSLSRPFPSLSRPGPLPSLFLVLSLSLSFPRSVPSRPVSLGPVPLPLPLPLPLPLPLRPVSSRP
Consensus

Teleogryllus_occipitalis     ------------------------------------------------------------------------------------------------------------------------
Gryllus_bimaculatus          VSSRSVPFCPVPLCPVPLCPAPLCSVPLCSVPTLSRPSPSRPVPQFVVAAKRRRLTKELLLIHWRQVPRRAAPRVPCPADLESARRRPPQWHRRRCGRTRRAAAADVISVAAV*VAWALA
Consensus

Teleogryllus_occipitalis     ------------------------------------------------------------------------------------------------------------------------
Gryllus_bimaculatus          *VEVDGHRG*RRRERRRR*RRRL*RPRR*RPRR*RPRQ*RPRR*SVRVPVIAGRQYDW*RGKWEEGPGIGNGLVVWWVS*LMFLSTLSATLVQ*IFCFGRWRCTSSHWMGRTDARMSACR
Consensus

Teleogryllus_occipitalis     ------------------------------------------------------------------------------------------------------------------------
Gryllus_bimaculatus          AFPSREFA*GALTQPV*SLPFKVGEIVKAVVP*RSRTEGSQVRVPLRACLYLCSSFRPLSSL*HAIFTVTKPLNAILVANNLDDEDNATFNNKNVLSFHVVRSTAFIQRILLEIDAEFDL
Consensus

Teleogryllus_occipitalis     ------------------------------------------------------------------------------------------------------------------------
Gryllus_bimaculatus          ASI*PMSEVRPHLVSNPRTSQSESQSRQLSSAM*ARN*RGSRCGNVAVEKEKTPCGGRCERLGSSDGCTASVATGEGVGVSAAARAVGGGRWKVRGEGAARGVVRQQAQCCHCGFA*MIS
Consensus

Teleogryllus_occipitalis     -----------------------------------------------------------------------------------------------------------
Gryllus_bimaculatus          LSANAFRVVVGL*WEGGGE*NSPRQEPQVKLELFSILCLGLFKLSIFCAFFSCTCIIHAILITLRVFYVN*NFVDIFCFSLLTSHDNIVLQKKS*NI*KVSRLKLIR
Consensus
```

**Tocc.8G0009300.1_exon2**

```
Teleogryllus_occipitalis                 ------------------------------------------------------------------------------------------------------------------------
Gryllus_bimaculatus                      IFTNFSENIN*L*SDYKSKCNFIFLSGKARLKNQISVSNSQGLQCV*IEWRQFWRKSAAGFMCLRFSCGIGHREADRCLGINRFAVSRTF*ASVLTTQRVRGSDSLQI*IKIYSLFGYQE
Consensus

Teleogryllus_occipitalis                 ------------------------------------------------------------------------------------------------------------------------
Gryllus_bimaculatus                      *PTRVFLSKNGQFPAHDYKTVFSWELRREIKKKPAQRTDVLPDKNIKLHFDL*SDYY*LIFSLKLLNMHIDGESSSIII**QI*KKVDIPHIFVEFLI*YCRRHVMFNPLLLLNIYEM*L
Consensus

Teleogryllus_occipitalis                 ------------------------------------------------------------------------------------------------------------------------
Gryllus_bimaculatus                      CRQLELVVYIIFVEAHS*SRLFHIDVSHLTFSEI*E*HIIKRIFC*ESMYQLMNN*VFEFTRYNFLPTILPFITQPTLLSADATLELISPL**KCDGGANAESDTKCGGTLQVMASGRCR
Consensus

Teleogryllus_occipitalis                 -AGVRRWAGGGGWAGGGGWAGGGGGAGGGG------------------------------------------------------------------------------------------
Gryllus_bimaculatus                      VAGNRNGGGYGGSGGGGGGSGGGGGGGGGGXXXXXXXXXXXXXXXXXXXXXXXXXXXXXXXXXXXXXXXXXXXXXXXXXXXXXXXXXXXXXXXXXXXXXXXXXXXXXXXXXXXXXXXXXX
Consensus                                 ** *   * **  ****  ***** ****

Teleogryllus_occipitalis                 ------------------------------------------------------------------------------------------------------------------------
Gryllus_bimaculatus                      XXXXXEAVELVEVGRSRCYKRRK*AQ*GARSYKNDTSLTAARQQEKHLS*DTFVRNVRTSRTYVTYIKRNVRLMWNAITSLGKVKVAPDIYKI*INLNFFSHATGV**NYSNTAPKHLGL
Consensus

Teleogryllus_occipitalis                 ----------------------------------------RAG-----------------------------------------------------GG----------------------
Gryllus_bimaculatus                      VKLIY*MILYVNCSLMVVELLD*L*KNGQTGFCVK*RQRSRAGNRSFVD*YPRRRNKNKEIISVKKVITDGKEYEEPRGMTRYLEVG*LKLGW***GGLKHRYTAESLGKRKRDRMGVRH
Consensus                                                                         ***                                                     **

Teleogryllus_occipitalis                 -----------------------------------------GVRGG--------------------------------------------------------------------------
Gryllus_bimaculatus                      LRQKRRERSVIIIMKSTAKAAEIFASIPPLPWKCSGAH*KYGARGGCRFLGVS*PHQRAHQRRQRPQWRRVVSLLPPSGVDRRRRRPGDARSAAAPAPQPDTAPRRGPRKQIRAAVRRRLA
Consensus                                                                          * ***

Teleogryllus_occipitalis                 ------------------------------------------------------------------------------------------------------------------------
Gryllus_bimaculatus                      GNQYPRERTPALIPSPVHLPSLSVPLLSLSRPFPSLSRPVPLPSLSVSVPSLSLVPSLSLPFTRPVPSCPVSLGPVMSRPVPLPLPLPLLSRPVSSRPVSSRPVRSRPDPLPSLTLSLFL
Consensus

Teleogryllus_occipitalis                 ------------------------------------------------------------------------------------------------------------------------
Gryllus_bimaculatus                      FPSRLVRSRTVPSRLVPSH*VPPDPLPSLSLVPSRHFKYRPVPSLSRPYLSLSVPLLSLSRPFPSLSRPGPLPSLFLVLSLSLSFPRSVPSRPVSLGPVPLPLPLPLPLPLPLRPVSSRP
Consensus

Teleogryllus_occipitalis                 ------------------------------------------------------------------------------------------------------------------------
Gryllus_bimaculatus                      VSSRSVPFCPVPLCPVPLCPAPLCSVPLCSVPTLSRPSPSRPVPQFVVAAKRRRLTKELLLIHWRQVPRRAAPRVPCPADLESARRRPPQWHRRRCGRTRRAAAADVISVAAV*VAWALA
Consensus

Teleogryllus_occipitalis                 ------------------------------------------------------------GGWAGGGG-------WW-------------------------------------------
Gryllus_bimaculatus                      *VEVDGHRG*RRRERRRR*RRRL*RPRR*RPRR*RPRQ*RPRR*SVRVPVIAGRQYDW*RGKWEEGPGIGNGLVVWWVS*LMFLSTLSATLVQ*IFCFGRWRCTSSHWMGRTDARMSACR
Consensus                                                                                                    * *  * *       **

Teleogryllus_occipitalis                 ------------------------------------------------------------------------------------------------------------------------
Gryllus_bimaculatus                      AFPSREFA*GALTQPV*SLPFKVGEIVKAVVP*RSRTEGSQVRVPLRACLYLCSSFRPLSSL*HAIFTVTKPLNAILVANNLDDEDNATFNNKNVLSFHVVRSTAFIQRILLEIDAEFDL
Consensus

Teleogryllus_occipitalis                 -------------------------------------------------------GG----------------------------GGGGGR---RGWAGRRGV-----------------
Gryllus_bimaculatus                      ASI*PMSEVRPHLVSNPRTSQSESQSRQLSSAM*ARN*RGSRCGNVAVEKEKTPCGGRCERLGSSDGCTASVATGEGVGVSAAARAVGGGRWKVRGEGAARGVVRQQAQCCHCGFA*MIS
Consensus                                                                                                **                              ****   **    ***

Teleogryllus_occipitalis                 ---------------------SGRR---------------------------------------RV-----------------------------------------
Gryllus_bimaculatus                      LSANAFRVVVGL*WEGGGE*NSPRQEPQVKLFLFSILCLGLFKLSIFCAFFSCTCIIHAILITLRVFYVN*NFVDIFCFSLLTSHDNIVLQKKS*NI*KVSRLKLIR
Consensus                                                      * *                                     **
```

**Tocc.8G0009300.1_exon3**

```
Teleogryllus_occipitalis                  ------------------------------------------------------------------------------------------------------------------------
Gryllus_bimaculatus                       YSQILVKILINYSLIINQSVILYFYQARHASKTKYPFQTRRVFSVFR*SGDNFGENRLQVLCACGLVAALATEKRIDV*ASTDLQYQGLSKHRF*LRNVLEEVIRCKYESRFTPFLVTRS
Consensus

Teleogryllus_occipitalis                  ------------------------------------------------------------------------------------------------------------------------
Gryllus_bimaculatus                       SQQEFFCRKMANFQRTITKLCFHGN*GEKLKKSLLNERMSCLIKI*NYTLIYNQIIIN*YFH*NY*ICILMENPLQ*LFNSKFKKKSIYHIFL*NF*FDIADAM*CLIHCFSLIFMKCNY
Consensus

Teleogryllus_occipitalis                  ------------------------------------------------------------------------------------------------------------------------
Gryllus_bimaculatus                       ADN*S*WYTLYL*KPILNPAYFTLTCLI*LFLKFENSTLLKGFFVENQCIS**IIKFLSSLAIISCQQFYRSSHNQLFSPRTLR*S*FRHCDENVMGVQMRKATPSVEERCR*WRVVGVA
Consensus

Teleogryllus_occipitalis                  ------------------------------------------------------------------------------------------------------------------------
Gryllus_bimaculatus                       SRGTETAAATAAAAAAVAAAAAAVAVAVAVAXXXXXXXXXXXXXXXXXXXXXXXXXXXXXXXXXXXXXXXXXXXXXXXXXXXXXXXXXXXXXXXXXXXXXXXXXXXXXXXXXXXXXXXXXX
Consensus

Teleogryllus_occipitalis                  ------------------------------------------------------------------------------------------------------------------------
Gryllus_bimaculatus                       XXXXXRRSSWLKLAGAVATSGASRRNEALARIRTTRASPLLDNRKNTSHKIRLYVTYVHHVRTLRTSNVMYV*CGMPLLRWERLR*LQTFIKYKLT*IFFRMQQVFDRTIQIQPQNI*DL
Consensus

Teleogryllus_occipitalis                  ------------------------------------------------------------------------------------------------------------------------
Gryllus_bimaculatus                       LS*FIK*FCMLTAH*W**SY*TNYKKMGKQVFV*NNVNAAELETDPSWTNTLEDGTKIKKLLV*KK**LMGKNTRNPEV*QGT*K*DD*N*DGSSRED*NTGILQKVWGRGNGIEWV*GT
Consensus

Teleogryllus_occipitalis                  ------------------------------------------------------------------------------------------------------------------------
Gryllus_bimaculatus                       *DRNAEKDRS*S**KVQQRQQKSLRAFRLFLGSAVGRIRSMVREAAVASSESADPISAHISGVNDRSGGALCPSFPPRGSIGAAARATRDRRRRQPRNLTPRRAAGRVNKYGQRCGAAWR
Consensus

Teleogryllus_occipitalis                  ------------------------------------------------------------------------------------------------------------------------
Gryllus_bimaculatus                       AINTHASGRPRSFPLPSISRPFPSLSSPSPVPFHPSPVPSLSRPFPSLSRPFPSSHPSLSLSLVPSRPVPSR*VP*CPVPSLSLSLSLSCLVPYRPVLSRPVPLGPVPTLSRPSPSPFSS
Consensus

Teleogryllus_occipitalis                  ------------------------------------------------------------------------------------------------------------------------
Gryllus_bimaculatus                       SRPVLLGPVPSRLVSSHPIRSLPTLSRPSPSSRPVTSSTVPSRPCPVPICPFLSLSCPSPVPSRLSPVLALSHPFSSSCPSLSLSLVPSRPVLSR*VPSLSLSLSLSLSLSLSVPSRPVL
Consensus

Teleogryllus_occipitalis                  ------------------------------------------------------------------------------------------------------------------------
Gryllus_bimaculatus                       SRLVPSRFVPSRYVPSRYVPPRYVPSRYVPSRPSLVPLRPVPSLSSW*QRSAGG*QRSSC*SIGDRCRAAPRRAFRAPLTSRAPAGDRLNGTGAAVGGRGGRRRRM*YRLPRCEWRGRWR
Consensus

Teleogryllus_occipitalis                  -----------------------------------------------------------GGNG---------------------------------------------------------
Gryllus_bimaculatus                       RLRWTGTEGEGGESDGGGDGGGCDGRGGDGRGGDGRGSDGRGGDR*ECR*LLDDSTIGDGGNGKKVLV*AMGWLFGGLVD*CF*ARYQLR*CSEYFVSGGGGVPPRTGWGEPTRVCRLAG
Consensus                                                                                            ****

Teleogryllus_occipitalis                  ------------------------------------------------------------------------------------------------------------------------
Gryllus_bimaculatus                       HFPLENLLEGHLRSQFKVCRSRWARSLRQ*FRSDLGPKDRRFEFRFGHACICARPFAPFHHYSMPSSQLPNR*MQYWSLIT*MMKIMRPLITKTF*ASTSSEAQHLYREFSSR*TLNST*
Consensus

Teleogryllus_occipitalis                  ------------W-RT-------------------------------W----------------------------EW*-----------------------------------------
Gryllus_bimaculatus                       LVYSQCRKCAPIWSRTLEPPNQSRRVDNCRRQCELATKGAAGVAM*LWKRRKRHAVAGARGLGAVTGAPPPSPQAKEWGCLRRRGRWAVGVGRCAGKEQREASCVSKHNAAIVGSPE*YL
Consensus                                              * **                               *                            **

Teleogryllus_occipitalis                  ----------------------------------------------------------------------------------------------------------
Gryllus_bimaculatus                       *VQTLFVLLLGCSGRGEGNEIPLDKNPRSSYECSVFFVWDCSSFRFFVLFFRVHASFMLF**HYESFM*IKILLISFASHSSLLMTT*FCKKSHKTYEKYLV*N*F
Consensus
```

**Tocc.9G0000100.1**

```
Teleogryllus_occipitalis      T--GTTGGTGTTGTTCACAAACAGATGGACACATGTTTTGAAAAAG------AAAACACAAACACTTTT---TTGGTATCAGGGGAGCCAAAAATTT-------------GAATTTCTAT
Tarbinskiellus_portentosus    TAGATTGCAATGAACTTCCAATTTAATAAAAAATGTTGTGAACAAATATGGTTATCCTAAGACGCCCCTCTATTATCACCAATT-----------------------------TTTGCAA
Acheta_domesticus             T---TAGTAGGGATCATCACATGCAAATTCACAGATTGAAGACAGATTTAACACTACATAAACACATCTCCACCTTCACCAGTTGAAAAGAAAGCTCTTACGTATATATACCATTTATAA
Gryllus_longicercus           T---TTAAAATCAGTGACACATGTCAGTGGTCAAGATTA--GCTGCTCTAACAATTAACAGATGCATTT-----TTTACTCAATGAAAAACAATTC-------------TAGTTAACAG
Consensus                     *   *            *  *         *   *                   * *  *   *         *                         **   *

Teleogryllus_occipitalis      GAAATCTGAAAATCAATTTTCTGCTACATTACATAACTTTCTCTTTTATACTA--CATAT-------------------AATAGTCTGTACACATGTACATATTTGAGAAAGTC-----
Tarbinskiellus_portentosus    CAGATGTGAGAT------------------------------------------CTTAT-------------------------------TGTTTATATAAATTAA------------
Acheta_domesticus             CAAGCCTGGTGTTCAGATTTACGTCAT-TAGGACCAGGATTTGTGTAATTATA--CATTT--CTTTATTTACAAGGTATAAACTTTTTGT-CATGTTTATAAATTAGGCTTTTTG-----
Gryllus_longicercus           GAATTTCAGGATGGATGATAGCAGCAT-TCCGACAGGGTTCTCTGTTGGCAAAATCATTTAACTTTATTTAATTGACAAGTACACTCTGTCCATTCTGACAGACCAGGAAACATAATTAA
Consensus                      *                                                      * * *                                          * * *

Teleogryllus_occipitalis      -AAAGAGTTGAAA------------------------AGAAAGCAAATGAAACTGGCTTGTGT-----------------------------------------GTGTCTCACC-----
Tarbinskiellus_portentosus    -AAATAG------------------------------AACAAGTAAATGCCATAGATATGTGG-ATGATATTT------------------------------------------------
Acheta_domesticus             -AAGTGATTTGAAATCTATTTGATTATCCATATTTGTCCACATTCAAATATGTAGAACATATAGCCTGATATTTAAGGAAAACTCATATTTCATTCCAAATTTCAATATTTCACTTGCAA
Gryllus_longicercus           CAAATAAATAAAAGCATTTTTGTCAAATAATTTTTACTAATAGGAAACTATAAAAACTATGAGTAATTATACTTTCAAGGAACTC----------------------TCTCTCCTTTAT
Consensus                      **                                      *    ** *           *

Teleogryllus_occipitalis      ------------------------------------------------TGCTGCTGCTGGGCCAA-------------------------CTGCTGCTGTGCCCGCTGGGAGCCTGCTG
Tarbinskiellus_portentosus    ------------------------------------------------TTTGTACTGTGAACAAA-------------AATGAGGTTATTAATATAATTAATGTTTTCAA----------
Acheta_domesticus             TATGTACCATATTGTACCCTATCAAAACCATTCTTGGAAGAATATAT-TTTACCTTTTAGGCCAATTTAAAGTGTTACATCAGGTTTACAAGTAGATCTGGATATTTTGGAAGGTT--CA
Gryllus_longicercus           TCAGCACC--AGTGTACTTTGTCCTGATCAATATCATCAATGAATGTGTTCACATGTTAAAAGAATTATTTAAATCTCACTAAGTTTAAAACATTTACTAAGATTTTTGG----------
Consensus                                                                *        *     **                           *

Teleogryllus_occipitalis      AGAAGGACTCCCATACAAT-----------------------------------------------------------------------------GAGGCCACAAGCCCTCGGGGT
Tarbinskiellus_portentosus    ---------CAATTACCAC-------CATAAACTCTAATTTACAAACAAAATAGAAATAAATAA----------------------------------------ATTAGTTTCCTGAAT
Acheta_domesticus             AGAGTGACTCCATTACTACTTCTGATGTTGAGCTTAATTTTAGCAAGTACAAAGCAAGATATAAAAGGGGCAGTTTCACATTTGAACATTTAAAACATGTAACCCACTGGTTTAT-AAAT
Gryllus_longicercus           --------TTCATTATTAC--------------------TAAAAATTCAAAATAAATAAATTATATTGTTGACTTTACATT------------------ACATAATAAATTATGAGAT
Consensus                                 **  *                                                                                   *            *

Teleogryllus_occipitalis      GGGCAGGGCACTGCTGC-------------------------------------------------------------------TGCTGCTGCTGCTGCTGGTGC------------
Tarbinskiellus_portentosus    GTACAGGATATTAACAA----------------------------------------------------------------------CAATAATATTACTAAAAC------------
Acheta_domesticus             TTTCAGAATAGGATTAAAAAACTTGCTTTCAGCAACAAAAAATGTTCAGGATATTTTTGTACATATATGCTCAAATTTTGTTAGATAGCTAATATTAAAATTAATTTCCTTTAC------
Gryllus_longicercus           ACACAGAAAAGTAATAAAATATT-----------------AAGAGTCAAGACTT-----------------------------------CGATTTTAGAATTAAATTTTTTTATTTATAC
Consensus                        ***   *                                                                                 *  *     *

Teleogryllus_occipitalis      -------------------------------------CACTGCTGCCTTTGCTCTTATGGCTGCTGCCATTG--------CTACCCCCTCCACTCGACT--------GCTGTTGCTGCTG
Tarbinskiellus_portentosus    --TGATTGGTTCAGAAAACCGACACACTCAGGTGATGTATTATATTTCCTA-----------------------------TCAAATCATCCATTACATGAAAAAAGAGCTGTGGTGTTTA
Acheta_domesticus             --TGTTCTATTTATGAAACTTAAAAATGTGATTAACACATTATTTTCACTATTTTTATGTTTTCAATTGTTAGCCCATATTTACACCTTATTCTCAACA---------------TTTTTA
Gryllus_longicercus           TGTACTTTATT--------------TTTATTTCACTTTCCATTTTGTCTATTCTGATAACTTTTAAAATTTCTCCTGATTTACATTACTTTTTCCAT---------------------
Consensus                                                                *                                  *           *  *

Teleogryllus_occipitalis      CTCT---------------------------------------------TGTTGCTGTCGCTGTTGTTGTTGTTGTTGCTGCTGCTGCTGCTGTTG-----------------------
Tarbinskiellus_portentosus    ATCT-CATAAAGTCATTTTATTATCAGTACAAAAGAACATAAAAAGAC------ATTTTGATTGATAATGTTGATATCCCTTAC-----------------CCATAAATAAATATACAAA
Acheta_domesticus             GTCCATACATAGGTTTACATTTTTCCATATAA---------------------ATCCTAGCCCTGGATATTAAGACTTCCTGCCCTGTTCTTACTG----CTATTATCAAGTCTGCGCT
Gryllus_longicercus           ---------TAATTATATGATTTTCTATTCAAAAAAAGAAATAAAATTGAAGTTATCTAAGAACTGAAAAGGAATATGGTATATCATTTCCTTACTAAAATCTTTCCTTAAAAATATATA
Consensus

Teleogryllus_occipitalis      -----------------------------------------------------------------------------------------------------------------------
Tarbinskiellus_portentosus    AAGGAATTAA-----------------------TAAACAAAAATATAATCATCATCATGAA----------------------------------------------------------
Acheta_domesticus             AATCATCCCA------GTGTGGA---GTCAAAGCAACATAATAGAATTGTCAGAGTGCTG-----------TTTTTTTTTTTACTCAACATTTTATTGCAATACCTGTCTATGTAAAAAA
Gryllus_longicercus           AATAAATAAAAATGACATGTAGATAGATAAAAGTCAATTAACAGGAATTACATATTCCTGAAAAGCACGAAATGACTTTTTTAGCTATTAATTTACTAGCAATTTTGTCAATTTCATAAG
Consensus

Teleogryllus_occipitalis      -----------------------------------------------------------------------------------------------------------------------
Tarbinskiellus_portentosus    -------------------------------------------CTGCAGGATGGTAGGTCTGTTGCCTGTTCCT-GTTAAAACAGTAGATCTGGTGGTTATTCCTC------------
Acheta_domesticus             TGTGTTGCAACAATAAAAGATTGACTGTTCTATTCCTAAGTCACCTGCAGTATGTGCATTAAATCTCTTATTTCTTGGAAAAACCCAGAAGATGGACAGCACTGGTTCGTTAGCAGACAG
Gryllus_longicercus           TTCATTCCAA------------------------------------------------AAATCGCTTGTACCT----GGAACTGGCAAGGCGCAGGGCATCG---------------
Consensus

Teleogryllus_occipitalis      ------TTGTTGTTGTTGCTGCTGCTGCTGTTGTTGTTGCTGCAGCTGCATACGTTGA--------ATGTGCTGGACCTATGAATT---------------------------------
Tarbinskiellus_portentosus    ------CAGATGTTCCTGTGCAATCCATC-----TTTCCTTGGGTCTTCTTATATCTC-------------TTCGCCCTATGGGTT---------------TATATTTTAAGA------
Acheta_domesticus             TTTATATTCAAACCCTTATGTCAGCTTTCAATGATATCTCTGCATTTACCAATGTTGT------------GTCAGTCTTTCATATTAGACACATGATTAGATTGTAGTTTGAAATAAATC
Gryllus_longicercus           ------TGAAAGTCCTTGCG--------------AATCTGACGCTGCCACATGTTGTTGGGGAGGGAGCGTTGAGTCTGTGGATT-GGCTCGCGGTT--------GTTGGAAAAGGGGA
Consensus                                    *                                *   *  *                         *       **

Teleogryllus_occipitalis      --TTGCTT--------------------------------------TTCTTTCACACTATTCATC-------------------------AACAAA-----------------------
Tarbinskiellus_portentosus    --CTGCTTTAG------------------GAAATCCTTCCACTTGCTTTCTTCACTGCACATGTG---------------------------------------------AAAGCCATCTC
Acheta_domesticus             CACTCATTCAGTCATTCAAGTTGTAGACCTATGTCCACCTATATT-TCTCTTCAGTTCACTAATG--TGAAATTCTTGGAT---------------------GTGAGAAAGGTCAATTT
Gryllus_longicercus           ATCCCATTTAG-------------AGCTGAGGTGCTGCCAT-TG-TCCCCTCTGGGCACTTGTGGCTGGGACTGTTGGTTAGAGGGCTGCAACAAAAAAACAGCTGAAAAGGGAAAATG
Consensus                           **                                  *    **      *    *

Teleogryllus_occipitalis      -----------------------------------------------------------------------------------------AACCGAGTTGT-------------------
Tarbinskiellus_portentosus    CTCTGGTTCTCTCTAATGTCATCCACTATATTTTTTGCATTGAGTTTTTCT-----GATATCTGTGTTTCTTTGTCAATCTAGG------CAGGTAATTCC------------------
Acheta_domesticus             CCCAGATTCTGAACTA-----------------GACACAACAA-------------AAAATAAAGACACCTTTTTTGATTTATA------AAGAAAATTTT--------ATTTGTTCACA
Gryllus_longicercus           CACAATGTATGTACTATTATGTTTACTAACTCCATCACAATAACTTTTTGCTGGATATTGTATTTACATTTCTTATGATTAATACTTTGGAAGAAAATTTTCACAAAGGATCTG-ACGGA
Consensus                                                                                                        *    * **

Teleogryllus_occipitalis      ------------------------------------------------------------------------------------------------------AATAAGGTTTGAAA--
Tarbinskiellus_portentosus    ------------------------------------------------------------------------------------------------------TACAAGATGCTGAA--
Acheta_domesticus             GGTTGAAATTATCGATAAGCTCTGATCACATATTTTATTGTAATATCAATACAATGTAAAATGTATGAAATACAAATATCATTTATGATTTTTTGCAAAAAGAAAATAAAATACAATAAT
Gryllus_longicercus           GATGGACATTGAGAAT--------------TTTTAAAATATAATAAGAATTTATTAGCAAAGGAATGGATAACGCATTTTA---ATGAGTTTTTAT-CATAGCATACAAAATATAACA--
Consensus                                                                                                              * **  *     *

Teleogryllus_occipitalis      -----------------------------------------------------------------------------------------------------------------------
Tarbinskiellus_portentosus    ----------------------------------------------------------------------AAAATCTCATTTGTGTGGCGTCTAACCTTT-----------------GAC
Acheta_domesticus             GAAGTTTAACTAAATTTGGCAACATCAACATTTCCACTTCTCTCAGCCA--------------------ATACTTTTACTTATGGAGTTTACAACCCAGAAGATTCACA-------ATA
Gryllus_longicercus           -------------------CCATATCAACAAGTTTACTATTGTACTCTACGAGTGTACAAGTTTGCTATTGTACTCTAATCTATGGTATTAATTATACATACATATTATAAGAGAACAAA
Consensus

Teleogryllus_occipitalis      --------------------------------------------------------------------------------------------------------GCTACCA---ACATT
Tarbinskiellus_portentosus    TCTCTCTGTTTTAGGGTCCACATTTCTGATCTATACATTAACATTGGCTTAGTAGTTATGTATT-----------TTTAGTTT--------------GGTCTCAGCTGTCATATCTCTT
Acheta_domesticus             TCTATTTCTCCTATTGTTTGTATTTTTATTTTATTTATTTATTTTTGTTTTGTTTTTTTGCATCCAGCTTGTGTATTCAACTTTATAGAGTGAAAGCTGGACTGAGCCTCAAAATACATT
Gryllus_longicercus           TTTATAAGCAAAACTGTTTGTCTCATGTCTTAGAAAATTAATTTTAGACCAATAC--------------------TTCAGATTTTTATGGTAAGGAC--AACTGTTTTGCAA---ACATT
Consensus                                                                                                                         *       **

Teleogryllus_occipitalis      TTAAATAT---------------------------------------------------------------------------------------------------------------
Tarbinskiellus_portentosus    CCAAAG-----------------------------------------------------------------------------------------------------------------
Acheta_domesticus             CAAAAATT------CAAAAAGAAATCTTAAAAAAATGTGTTCAATGGCACATCAAAAAATCCG--------------------------------------------------------
Gryllus_longicercus           CAAAGAATATGTGCCAATAATAAATTTTTCAGAATTGTCTTTATAAGTGCATAAGGAAAATAACATTGTGAAATCAGCAATGCAATGTTGCCTAGATTGTAGGCAGCAAAGCTAACATCA
Consensus                       **

Teleogryllus_occipitalis      ----GAATAAATTCTGAAGTGGAAAGTGTGATTTGGAGTAATTAC--------------------------------------------------------------------------
Tarbinskiellus_portentosus    --------TTTTTCTTAATTATTCCACTTACCCCATTGTATTTTT-------------------------------------------GGATTTTTCTCTCTTGATCTGAATTTTGGCC
Acheta_domesticus             --------TTTATGTGAATTGCAGTATTTTTTTTATAATTTTTATTTT-----------------------------------------CTTCAATGCCTTTTGTACTTCATTTGCTG
Gryllus_longicercus           TAAAAAGTTATTCCTGAATTACAATATCTACTTGGAAACTATTACACTGTGATACAAATATCTGTGATATGACACAAAATCTTAATGGAAGGTTCACCCACCTCTGTCTAATTTATATCA
Consensus                                * ** *         *              **

Teleogryllus_occipitalis      -----------------------------------------------------------------------------------------------------------------------
Tarbinskiellus_portentosus    GATGTTTCCTAGATAAATAAA--------------------------------------------------------------------------------------------------
Acheta_domesticus             TGTGATCAGTGGAACGGGGAGGAAGGGGCGATGCATCTGGGGGCCCGTGGCTGACTGGAGGCAGGGGCAGAGGCGGAGTCAGGGTCGGGGTCGGGGGCAGGGGCGAGCAGGGGTCTGCCT
Gryllus_longicercus           TATCTTTTATGAAACAAAAATGTTCAAAGAATGT-------------------------------------GCACAACAAAGATCAAAATTAAACACAG--------------------
Consensus

Teleogryllus_occipitalis      -----------------------------------------------------------------------------------------------------------------------
Tarbinskiellus_portentosus    ---------------------------------------------------------------------------------------TTTTGTTACTAGTTCTATTATTT---------
Acheta_domesticus             CTGTAGCGAGCACTCACCGCAGAGGGCGGCCTGGGGAAGGCGATGAGCAGTGGCGTGGTGCCCACTGACTGGAACAGCTGCAGAGGGCCCCTGTTCTCCGCTGCGTCATCCACAGGCGTC
Gryllus_longicercus           -------------TAAGTGTTGAAGTAGACATGAGAAA-------------------------------------------------TATTTTCTTGTGATTCTTGAATCACATACGCC
Consensus

Teleogryllus_occipitalis      ------------------------------------------------------------------------------------------------------------CAACGAAAA
Tarbinskiellus_portentosus    -------------------------------------------------------------------CCCCATCTATAATTATAACTATTTTGATTCT----------------------
Acheta_domesticus             TGCCCATGGGGCTGCAGACTCCTGCATGTGGACAAATAGCCAATGTGGATTGGAAGGAAAACAACAGCCCGGTACATAACTTAAACTACGCTAACTGTGTATACTGTTTATTAATACAAA
Gryllus_longicercus           CA-CAATGATGCATCACTACTCAACATGTTGTTCATTAGGTAATGCGTATTGTATCAAAAAATCAAATCTAAAATGTATCCAAA---------------------------TATCACACA
Consensus

Teleogryllus_occipitalis      TAAT------------------------------------------AATGTTTGAAAA-------------------------------------------------------------
Tarbinskiellus_portentosus    --------TCTCATTTCTTTGCCTTGGTATGCCATAGATTTTGTCTTATTTGTAGAAA-----------------------TTCTCACGTCATATGTTAGGGCTATTTTATGATGTT--
Acheta_domesticus             GGGTACGGTATTAACTTTGTTGATGAATATGAAATAACCTTAAAATAATATTTGAAGAGACCAGCAGAATTGTTCTTTCTATTTCAAATGAAATAATTCAGTTTTACCTGCATACATTCA
Gryllus_longicercus           GAAT--------------------GGTGAGATATAAGATGGAATCATTACATGCAAA-----------------------TTCACAGATTGTAGTTTATGACTACATAAACACAT---
Consensus                                                                *   *  * *

Teleogryllus_occipitalis      -----------------------------------------------------------------------------------------------------------------------
Tarbinskiellus_portentosus    ---------------------------------------------------------------------GGTTTACTGCTCTTTGT---------------------------------
Acheta_domesticus             TTGCTGTGACCTTTGTTTCATCATAGCAATTTTATGGAATGTACAGATTGGGTACTGTCAATCCCCACAGAGTTTATTAATGTACATTAATTGGAGATGGGGACCTGTCAATTTAAATA
Gryllus_longicercus           ---------------CTCCACCTTCACCAGCTGGGAAGAAAAAAAAAAAAAAGC-----ATCCTAGCAGAGCTAGTATGTGTATGT------------------------AATTACATA
Consensus

Teleogryllus_occipitalis      -------------------------------------------TATAATTGTTGCTACATGAGTATGAAAC------------------------------------------------
Tarbinskiellus_portentosus    ---------------------AAAACATCCTCTGATTCCATTGTTAGGATCATGTTGTAAGCACAGAGAAT------------------------------------------------
Acheta_domesticus             TTTTATAGCAATACAGAAATAAAAATAATATTAAATCAGAGTGTACAAATACT-TTGTATTCATTTATAATGAAGCCAGCTAGTTAAACCATATTATGAGCACAACTTAGCTCAACTAAA
Gryllus_longicercus           TTTCTTTACTCATCAGTTATCAAATTTTTGTCATGTT------TATGAATAATGATGAAGTGATTTGAAATCAATTTGATTATCCATATTTGTCCATATGCAAATATGTG----------
Consensus                                                           *     *  *  *  *         **

Teleogryllus_occipitalis      --------TCTTTGTTCTTACATCACCAGTA-------------------------------------------ATAGTTATTTTGGT----------------------AATGGTTAC
Tarbinskiellus_portentosus    --------TGTTTGGACATTTTGTCTT-TTGTTTAAGTCTATTCCAGTTTTCA----------------------CTATTTGTTTCCAC-------------------------------
Acheta_domesticus             CTATAGTTCGTCAGGGCTCAAGTTCCCAATAAATGAGCAGGGCCCGAGTTTCAGTTAGATGGGGGCCCAAGGCTCTCAAAGATTTGGGGTCCCTAGGTAACTGTCTGGAAAAATGACTAT
Gryllus_longicercus           -----------TAGTGCACATATTCCC-CTATTTAAGGAAAACTCATATTTCA----------------------TTACAAATTTGAACTTTTTACGTG--------------CAATAT
Consensus                                  *  *          *                                                *    ***

Teleogryllus_occipitalis      --------------------------TTTTTTCAATTGGTAAGTA-----------------------------------------------TGAGTAGAAATGACCAAACTTCAAAACTTA
Tarbinskiellus_portentosus    ---------------------------TATTTTAACATGTGATTG-------------------------------------GCATAATGTTGAATAGGCATGGTGACAGTCCACACCCCT
Acheta_domesticus             GTCATATTTCATATTGTCATAAAAATTTATTCTGAAAAGTTAATA---------TAAAAATAGTATGTAAGCAAATGAAATGTAAAAAGATTATTAG---AGGGAAGGGATTAAAATCTC
Gryllus_longicercus           GTACTATATTGTAGCCTTTCAAAAG--CATTCTTGGAAGAAAATATTTTGTCTTTTAGTGCAACAAAGGGGTAAATCAGATTTTCAAGATTTGAGAGATCAGGGGGTAAATCAGATTTTC
Consensus                                          **       *  * *                                                  *    **    *           * *

Teleogryllus_occipitalis      GT---------------------------------------------------------------------------------------------------------------------
Tarbinskiellus_portentosus    GTCTCAGTCCTCTGGTTGTTCTAGAAGCCTCTGTGTTGCTCCCGTCATTCATTCTTA---------CATTAATTGGAGAGTTTGCATATGGATTC------------------------
Acheta_domesticus             AT--TAGGGTTCCGATAATTTTATA------TTATTTGCTTTTTCAACTTCCCGTTAGAGAACAAGAATTCATTAGAGAGATTGCTTGATGATTTTTTAGCATCTGTGGGGCCTGTTAAT
Gryllus_longicercus           A----AGATCTGGGTTAACCTAAAA------TATATTGCAAGTTCA----------------------------AGATGGACTCCGTTACTACTTCTGAGGTT----------------
Consensus

Teleogryllus_occipitalis      -----------------------------------------------------------------------------------------------------------------------
Tarbinskiellus_portentosus    ------------------TGGATTCCTTTTCTTATGTCT--------------------------------------------------------------GTGGGTATCTCTTCATATT
Acheta_domesticus             CATGGGGGCCCAAGGCACAAGCTTTGTCAGCCTGTGCCCAAAGCCGACCCTGCTAATTAGCAGAAACATAGAGTAAAGAGTCTATGGAAGTAATAGTGAAAACATATTTCTCAGCCAAGT
Gryllus_longicercus           ------------------GAGCTTCATTTTAACAAGTTCAAAGC----------AATATTAAGAGATAAAAGGGATAGTTTC-------------------ATGTGTGTTTTCACTGCTT
Consensus

Teleogryllus_occipitalis      ----TGTTACC-------------------------TATATT------------------------------
Tarbinskiellus_portentosus    CCAGTATTTCC-------------------------CACATTTTGCT-------------------------
Acheta_domesticus             TAAATATTATTGTTACAGAATCACCAAATATATCAATCACATCAGGATAACTGGCAGTGAATTTGTTGACTCAAG
Gryllus_longicercus           TACATATGGCC-------------------------------AATTATCAGAAAAGGATTAGAAAACT----
Consensus                         * *
```

**Tocc.9G0000100.1_exon1**

```
Teleogryllus_occipitalis      ------------------------------------------------------------------------------------------------------------------------
Tarbinskiellus_portentosus    ----------------------------------------------RNCNQHYRLQ*TS------N--------LIKNV-----------VNKYGYPK-------TPL--YYHQFLQQ
Acheta_domesticus             LESTNSLPVILM*LIYLVIL*Q*YLTWLRNMFSLLLP*TLYSMFLLISRVGFGHRLTKLVPWAPMINRPHRC*KIIKQSL**ILVL*REVEKANNIKLSEP**DFNPFPLIIFLHFICLH
Gryllus_longicercus           -------------------------------------Y**QAQIFKISDTCQWSRLAALT-----IN---RC---IFYSM-----------KNNSS*QE-----------FQDG**QH
Consensus

Teleogryllus_occipitalis      ------------------------------------------------------------------------------------------------------------------------
Tarbinskiellus_portentosus    M*DLIVYIN--------------------*K*NK*M-P*ICG*-----YFLYCEQK*GY*YN*C-----FQQLPP*T----LIYKQNRNK*ISF----LNVQDIN-------------
Acheta_domesticus             TIFILTFQNKF-L*QYEI*HSHFSRQLPRDPKSLRALGPHLTETRALLIYWELEP*RTIV*LS*VVLIIWFN*LASL*MNTKYLY----TLI*YYFYFCIAIKYLN*QVPHLQLMY----
Gryllus_longicercus           SDRVLCWQNHLTLFN*QVHSVHSDRPGNIINK*IKAF---LSNN-----FY**ETIKTMSNYT-------FKELS-------LLY----SAPVYF----VLINIINECVHMLKELFKSH*
Consensus

Teleogryllus_occipitalis      ------------------------------------------------------------------------------------------------------------------------
Tarbinskiellus_portentosus    NNNITKTDWFRKPTHSGDVLYFL-------------SNHPLHEKRAVVFNLIK--SFYYQ---YKRT*KDILIDNVDI------PYP*INIQKGINKQKYNHHH---ELQDGRSVA----
Acheta_domesticus             INKLCGD*QYPIC-----------TFHKIAMMKQRSQQ*MYAGKTELFHLK*KEQFCWS-LQILF*GYFIFINKVNT------VPFV-LINSIHS*R-SLSYVP-----GCCFP----
Gryllus_longicercus           V*NIY*DFWFIITKNSK*INYIVDFTLHNKL*DTQKSNKILRV-KTSILELNF---FIYTVLYFYFTFHFVYSDNF*NFS*FTLLFPLI-I*FSIQKKK*N*SYLRTEKEYGISFPY*NL
Consensus

Teleogryllus_occipitalis      ------------------------------------------------------------------------------------------------------------------------
Tarbinskiellus_portentosus    ---------------------------------------------------------------------------CSC*NSRSG----------------------------------
Acheta_domesticus             SNPHWLFVHMQESAAPWADACG*RSGEQGPSAAVPVSGHHATAHRLPQAALCGECSLQRQTPARPCPRPRP*LRLCPCLQSATGPQMHRP------FLPVPLI---------TQQMKYKR
Gryllus_longicercus           SLKIYK*IKMT---------------------------------------CR*IKVN*Q------------------------ELHIPEKHEMTFLAINLLAILSIS*VHSKNRLYLE
Consensus

Teleogryllus_occipitalis      -----------------------------------------------------MRPQ------------------------------------ALG---------------------
Tarbinskiellus_portentosus    -------------GYSSRCSCAI-HLSLGLL--------------------ISLRPMG--------------------------LYF----KTALGNPSTCF---------------
Acheta_domesticus             H*RK*KL*KKYCNSHKRIF*CAIEHIFLRFLFEFLNVF*GSVQLSL-YKVE-----------YTSWMQKNKTKINK*NKNTNNRRNRY---CESSGL*TP*VKVLAERSGNVDVAKFS*T
Gryllus_longicercus           LARRRAS*KSL-----RI*RC---HMLLGRER*VCGLARGCWKRGIPFRAEVLPLSPLGTCGWDCWLEGCNKKTAEKGKCT-----MYVLLC----LLTPSQ*LFAGYC----------I
Consensus                                                                                                                     *

Teleogryllus_occipitalis      ------------------------------------------------------------------------------------------------------------------------
Tarbinskiellus_portentosus    ---------------------LHC-------------------------------------TCESHLLWFSLMS---STIFF--ALSFS---------DICVSLSI*AGNSYKMLKKSH
Acheta_domesticus             SLLYFIFFLQKIINDICISYILHCIDITIKYV------------IRAYR*FQPVNK*NF-LYKSKKVSLFFVVSSSESGKLTFLTSKNFTLVN*REI*VDIGLQLE*LNEWIYFKLQSNH
Gryllus_longicercus           YISYD*YFGRKFSQRI*RRWTLRIFKI**EFISKGMDNAF**VFIIAYKI*HHINKFTIVLYECTSLLLY--------SNLWY-----------------*LYIHIIREQIYKQNCLSH
Consensus

Teleogryllus_occipitalis      --------------------------------------------------------------------------------VGRALL----------------------LLLL------
Tarbinskiellus_portentosus    L---------------------------------------------------------CGV*PLTLS-----------VLGSTFL----------------------IYTLTLA**L
Acheta_domesticus             V------SNMKD*HNIGKCRDIIES*HKGLNINCLLTNQCCPSSGFFQ-----------EIRDLMHILQVT*E*NSQSFIVATHFL---HRQVLQ*NVE*KKKQHSDNSIMLL*LHTGM
Gryllus_longicercus           VLEN*F*TNTSDFYGKDNCFANIQ--------------RICANNKFFRIVFISA*GK*HCEISNAMLPRL*AAKLTS*KVIPELQYLLGNYYTVIQISVI*HK-------ILMEGSPTSV
Consensus                                                                                           *

Teleogryllus_occipitalis      ---------------------------------------------------------------------------------------------LLV------------------------
Tarbinskiellus_portentosus    CIFSLVSA--------------------------------------------------------------------VISLPKFFLII------------------------
Acheta_domesticus             ISADLIIAVRTGQEVLISRARIYMEKCKPMYGLKMLRIRCKYGLTIENIKIVKIMC*SHF*VS*IEQ*RKLILILAI*QNLSIYVQKYPEHFLLLKASFLILF*KFINQWVT--------
Gryllus_longicercus           *FISYLL*NKNVQRMCTTKIKIKHSKC---------------------------------------------------------*SRHEKYFLVILESHT---PTMMHHYSTCCSLGNAY
Consensus                                                                                                 *

Teleogryllus_occipitalis      ---------PLLPLLLWLL-------------------PLLP-----------------------------------PPLD------------------------------------
Tarbinskiellus_portentosus    ---------PLTPLYFWIFLS*S------------EFWPMFP-----------------------------------R*INFVTSSIIS--------PSIIITILILLISLPWYAIDFV
Acheta_domesticus             CFKCSNVKLPLLYLALYLLKLSS---TSEVVMESLLNL-PKYPDLLVNLM*HFKLA*KVKYILPRMVLIGYNMVHIASEILKFGMKYEFS-----------LNIRLYVLHI*MW---TN
Gryllus_longicercus           CIKKSNLKCIQISHRMVRYKMESLHANSQIVVYDYINTSPPSP---AGKKKKKKAS*QS*YVY----VITYFFTH---QLSNFCHVYE***SDLKSI*LSIFVHMQICVVHIFPYLRKTH
Consensus                                                            *  *

Teleogryllus_occipitalis      -------------CCCCCSCCC--------------RCCCCCCCCCCCCCCCCC-------------------CCCC--------------------CCS---------------
Tarbinskiellus_portentosus    LFVEILTSYVRAIL*CWFTALCKTSSDSIVRIML*AQRIVWTFCLLFKSIPVFTICFHYFN-----------M*LA*C*IGMVTVHTP-VSVLWLF*KPLCCSRHSFLH*LESLHMDSG
Acheta_domesticus             MDNQIDFKSLQKA*FINMTKSLYLVNKEMYNYT----------------------------NPGPNDVN------LNTRLVINGIYT*ELSFQLVKVEMCLCSVKSVFN----------
Gryllus_longicercus           ISLQI*TFYVQYVLYCSLSKAFLEENILSFSATKG*IR-----------------FSRFERSGGKSDFQDLG*PKIYCKFKMDSVTTSEVELHFNKFKAILRDKRDSF-----------
Consensus

Teleogryllus_occipitalis      ----CIR*-------------
Tarbinskiellus_portentosus    FLFLCLWVSLHIPVFPTFC--
Acheta_domesticus             -L*ICM**SLLIISP------
Gryllus_longicercus           ---MCVFTALHMANYQKRIRK
Consensus                         *
```

**Tocc.9G0009060.1**

```
Teleogryllus_occipitalis        TCACCCTTTTCCGAAACATCAAACATTCTGATCGAACTAAAGCGAAACCGTCTCCAGAATAATAAACGCCACCCACAGCAAATTTCGAGCGCAAAATAAAGG-AAGAATCGCGCCACAAA
Acheta_domesticus               TCATCCTTTTCCGAAACATCGAACATTCTGATGGACCCAAAGCGAAACCGTCCCTGGAATAATAAACGCCTCCCACAGCAAATCTCGAGCGCGAAAATAAAGTAAAAATCACGCCACAAA
Gryllus_bimaculatus             TCATCCTTTTCCGAAACATCAAACATTCTGATCGAACCGAAGCAAAACCGTCTCGAGAATAATAAACGCCACCCACAGCAAATTTCGAGCACCAAATAAAGG-AAAAATCGCGCCACAAA
Gryllus_longicercus             TCATCCTTTTCCGAAACATCAAACATTCTGATCGAACCAAAGCAAAACCGTCTCGAGAATAATAAACGCCACCCACAGCAAATTTCGAGCACCAAATAAAGGAAAAAATCGCGCCACAAA
Consensus                       *** **************** *********** ** *  **** ******** *  ************** ************ ****** * ***  ** * ** **** *********

Teleogryllus_occipitalis        CGAGCC--GGGAGAAGGGCGACTTCTTCACACAAAACTTCTTGCTCC-TTTGAGTCACTCTACATACATCGGT-GCCCTCGACGGAAGCTCTCCTGCGAGAAGGATTCGTTCC-CCAATT
Acheta_domesticus               CGACCCCGGGAGAAAGGGCGACTTCTTCACACAAAACTTCTTGCTCC-TTTGAGTCA--CTACATACATCGGTGGCTCTCGACGGAAGCTCTCCTGCGAGAAGGATTCGTTCT-CCAATT
Gryllus_bimaculatus             CGACCC--GGGAGAAGGGCGACTTCTTCACATAAAACTTCTTGCTCCTTTTGAGTCACTCTACATACATCGGT-GCCCTCGACGGAAGCTCTCCTGCGAGAAGGATTCGTTCTCCCAATT
Gryllus_longicercus             CGACCC--GGGAGAAGGGCGACTTCTTCACACAAAACTTCTTGCTTCTTTTGAGTCACTCTACATACATCGGT-GCCCTCGACGGAAGCTCTCCTGCGAGAAGGATTCGTTCTCCCAATT
Consensus                       *** **  **   ****************** ************* * *********  ************** ** ***********************************  ******

Teleogryllus_occipitalis        CGTTCCAAATCATTTAGCGCGGAGTTCCCGGCAGAAACCACATACTGTGCGACACTCGACACAGGAATGCTTCGAAGGTCTGGCTTGGCTGCACACTCGGCAGAGCGCGGCGGAGGCGGC
Acheta_domesticus               CGTTCCAAATCATTTAGCGCGGCGTTCCCGGCAGAAACCACATACTGTGCGACACTCGACACA-GAATGCTTCGAAGGTCTGGCTTGGCTGCACACTCGGCAGAGCGCGGCGG-------
Gryllus_bimaculatus             CGTTCCAAATCATTTAGCGCGCAGTTCCCGGCAGAAACCACATACTGTGCGACACTCGACACAGGAATGCTTCGAAGGTCTGGCTTGGCTGCACACTCGGCAGAGCGCG----------
Gryllus_longicercus             CGTTCCAAATCATTTAGCGCGGAGTTCCCGGCAGAAACCACATACTGTGCGACACTCGACACAGGAATGCTTCGAAGGTCTGGCTTGGCTGCACACTCGGTAGAGCGCG----------
Consensus                       *********************  **************************************** ************************************ ********

Teleogryllus_occipitalis        GGTGGTGGCGGCGGCAGCTGTGGCAGCGGACGCGGAGGCGGTGGAGATGGTGGCGGAGGCGGAGGCGGAGGCGGAGGCGGAGGCGGAGGCGGAGGCGGAGGCGGAGGCGGTGGCGGAGGT
Acheta_domesticus               ------------------------------------------------------------------------------------------------------------------------
Gryllus_bimaculatus             ------------------------------------------------------------------------------------------------------------------------
Gryllus_longicercus             ------------------------------------------------------------------------------------------------------------------------
Consensus

Teleogryllus_occipitalis        GGTGGCGGTGGCGGTGGTGGAGGCGGTGGCGGCGGCGGTGGTGGCGGTGGCGGTGGCGGTGGCGGTGGCGGTGGCGGTGGTGGAGGCGGTGGCGGAGGCGGCGGCGGAGGCGGAGGCGGA
Acheta_domesticus               ------------------------------------------------------------------------------------------------------------------------
Gryllus_bimaculatus             ------------------------------------------------------------------------------------------------------------------------
Gryllus_longicercus             ------------------------------------------------------------------------------------------------------------------------
Consensus

Teleogryllus_occipitalis        GGCGGTGGCGGAGGCGGTGGCGGTGGCGGTGGCGGAGGCGGTGGCGGCGGCGGCGGCGGCGGCGGCG-GCGGCG--GCGGCGGCGGCGGTGGCGGCGGCGGTGGCGGTGGCGGAGGTGGT
Acheta_domesticus               -----------------------------------------CGCGGCGGCGGCGGAGGCGGCGGCG-GCGGCG--GCGGCGGCGGCGGGGGCG-------------------------
Gryllus_bimaculatus             ------------------------------------------GCGGCGGCGGCGGCGGCGGCGGGGGGCGGCGGCGGGGCGGCGGCGGGGGCGGC-----------------------
Gryllus_longicercus             ------------------------------------------GAGGCGGCGGCGGGGGCGGCGGCA---------GGGGCGGCGGCGGGGGC--------------------------
Consensus                                                                 * *********** ********           * *********** ***

Teleogryllus_occipitalis        GGCGGAGGCGGTGGCGGAGGCGGGGGCGGTGGCGGCGGCGCCGGTGGCGGCGGTGGCGTCGGTGGCGGTGGCGGCGGTGGCGGCGGAGGCGGTGGCTACGGTGGTGGCGGAGGCGGAGGC
Acheta_domesticus               ------------------------------------------------------------------------------------------------------------------------
Gryllus_bimaculatus             ------------------------------------------------------------------------------------------------------------------------
Gryllus_longicercus             ------------------------------------------------------------------------------------------------------------------------
Consensus

Teleogryllus_occipitalis        GGAGGCGGAGGTGGTGGCGGAGGCGGAGGCGGAGGCGGAGGCGGAGGCGGAGGCGGAGGCGGAGGCGGAGGCGGAGGCGGAGGCGGAGGCGGAGGCGGAGGCGGTGGCGGTGGCGGTGGC
Acheta_domesticus               ------------------------------------------------------------------------------------------------------------------------
Gryllus_bimaculatus             ------------------------------------------------------------------------------------------------------------------------
Gryllus_longicercus             ------------------------------------------------------------------------------------------------------------------------
Consensus

Teleogryllus_occipitalis        GGTGGCGGTCGCGGCGGTGGCGGTGGCGGCGGCGGTGGCAGCGGCGGCGGCCTCAGTCGTTGATGGCGACGCGGTGACGTTGCAGGATTCACCCACGGAGTTCTGCCCAGCCATTGGATT
Acheta_domesticus               -------------------------------------GGGGCGGCGGTGGCCTCAGTCGTTGATGGCGACGCGGTGACGTTGCAGGATTCACCCACGGAG-TCGGCCCAGCCATTGGA-T
Gryllus_bimaculatus             -GCGGCGGGCGCGG---------------------CGGGGCGGCGGCGACCTCAGTCGTTGATGGCGACGCGGTGACGATGCAGGATTCACCCACGGAGTTCTGCCCTGCCATTGGATT
Gryllus_longicercus             ---------------------------------------GGCGGCGGTGGCCTCAGTCGTTGATGGCGACGCGGTGACGATGCAGGATTCACCCACGGAGTTCTGCCCTGCCATTGGATT
Consensus                                                             ******* * ***************************** ******************** ** **** ********* *

Teleogryllus_occipitalis        GCGCGCGGGATTGACAAGGTCTCGTGGTTTCCCTCCGTGCCGCAAACAGTCATGATTCGAAGACTGGAGCAGGGCCCATTACAATGGTGCATCTGGAGCACAGAGCACCTTTCCGCGGTC
Acheta_domesticus               GCCCGCGGGATTGACAAGGTCTCGAGGTTT-CCTCCGTGCCGTGAGCCGCCATGATTCGAAGTCTGGAGCAGGGCCT-TTACAATGGTGCACCTGGAGCACAGAGCACCTTTTCGCGGCC
Gryllus_bimaculatus             GTACGCGGGATTGACAAGGTCTCGTGGTTTCCCTCCTTGCCGCGAGCAGCCATGATTCGAAGACTGGAACAGGGCCCATTACAATGGTGCATCGGGGGCACGGAACACCTTTCCGCCGCC
Gryllus_longicercus             GTGCGCGGGATTGACAAGGTCTCGTGGTTTCCCTCCTTGCCGCGAGCAGCCATGATTCGAAGACTGGAACAGGGCCCATTACAATGGTGCATCGGGGGCACAGAACACCTTTCCGCGGCA
Consensus                       *  ******************** ***** ***** *****  * * * ************ ***** *******  ************* * ** **** ** ******* *** *

Teleogryllus_occipitalis        GGAGACACCGGTCGGAGCGAAAGCAATAGCGACCGAGTTGACGAAGGAGGTGGGGGCATGATACCG----------------------------------------------------
Acheta_domesticus               GGAGATACCGCTCGGAGCGGCAGCAATAGCGACCGAGTTGACGACGGAGGTGTGGTCATGATACCGGAATGAATTATCNNNNNNNNNNNNNNNNNNNNNNNNNNNNNNNNNNNNNNNNNN
Gryllus_bimaculatus             GGAGATATCCGTCGGAGGGAAAGCAGTAGCGACCGAGTTGACGACGGAAGTGGGGGCTTGATACCG----------------------------------------------------
Gryllus_longicercus             TGAGATATACGTCGGAGGGAAAGCAGTAGCAACCGAGTTGACGACGGAAATGGGGGCTTGATACCG----------------------------------------------------
Consensus                        **** *    ****** *  **** **** ************* ***  ** ** * ********

Teleogryllus_occipitalis        ----------------------------------------------------------TAATGAATTATGGTGGGTGGTGTAGCAGAGG-----------------------------T
Acheta_domesticus               NNNNNNNNNNNNNNNNNNNNNNNNNNNNNNNNNNNNNNNNNNNNNNNNNNNNNNNNNNTTGTCAGCTGCCATCTGTCGCCTCTTAAAGTACATTTTTTCGTCATTTTCTTAATTTTCCGT
Gryllus_bimaculatus             ----------------------------------------------------------TCCTTAGTTAACGTTTGTGGTGTAGCAGATT-----------------------------T
Gryllus_longicercus             ----------------------------------------------------------TCATTAGTTATCGTTGGTGGCGTAGCAGATT-----------------------------T
Consensus                                                                                 *  * *  *    *  ** *  *   * *                             *

Teleogryllus_occipitalis        TTTGCAATGGACGATAT-------------------------------------------------------------------------------------------------------
Acheta_domesticus               TTCCTAATGGTAGAACAGAAATATTAAATTT-------------AGAATATTACAGAATACTAAAATCTCGTCCTAAATGGAA----------------ATTTTCTTTGAAAAATTTG
Gryllus_bimaculatus             TACGTAATGGAAGATGTAATGTATTACATTTCGATGGTTTCTGCCAAAATCTCGTAAATGTTTATAATATTAGGTGAAATGAAACGAAAATCACGCATATTATTCATTTTAGAGTATTAC
Gryllus_longicercus             TTTGTAATGGAAGATGTAATGTATTACATTTTGATAGTTTCTGCCAAAATCTCATAAATGTTTATAATATTAGGTGAAATGAAACGAAAATAATGCATTTTATTCATTTTAGAGTATTAC
Consensus                       *    *****  **

Teleogryllus_occipitalis        ------------------------------------------------------------------------------------------------------------------------
Acheta_domesticus               TTACCGTTTTTAAAAATCTGGAGTTTTAGTGCTACTAAAAATGCGAAATTTTAAGCTCATTCT---------------------------------------------TACAGAAAATT
Gryllus_bimaculatus             TTTCTTTTTGCAAAAAATGGGATATTTATTTGTTCTACACGAGT---ATTACCAGCCCATCACAGCCCATTACAGGCCTTACCAGCCCATTACAATGGTGCATCGGGGGCACAGAACACC
Gryllus_longicercus             TTTCTTTCTGC-AAAAATGGAATATTTATTTGTTCTACACGAGT---ATTACCAGCCCATCACAGCCCATTACAGGCCTTACCAGCCCATTACAATGGTGCATCAGGAGAACAGAACACC
Consensus

Teleogryllus_occipitalis        ------------------------------------------------------------------------------------------------------------------------
Acheta_domesticus               TTTGAGCTTTAGAAAAACTTCATTTGAAGTCATTTAAAGCAGT----------TTAACTACAAAAACAATAGT-------------------------------------GCTTCCAAATT
Gryllus_bimaculatus             TTTCCGTGGCCGGAGATATCCGTCGGATG-----GAAAGCAGTAGCGACCGAGTTGACGACGGAAATGGGGGCTTGATACCGTCCTTAGTTAACGTTTGTGGTGTAGCAGATTTTACGTA
Gryllus_longicercus             TTTCCGTGGCCTGAGATGTACGTCGGAGA-----GAAAGCAGTAGCGACCGAGTTGACGACGGATGTGGGGGCTTGATACCGTCCTTAGTTAACGTTGGTGGTGTAGCAGCTTTTTCTTA
Consensus

Teleogryllus_occipitalis        -------------TGTGTTAC---------------------------------------------------------------------------------------------------
Acheta_domesticus               GTGGGAAATG--------TGCATATTAACGGTTTATTAACGGAGACCTTTCAACAAAATTACATTTGTGTAATTTTTGGGAAAAAAATATACTCCAAATTGAACTTTACGTCATAATAAA
Gryllus_bimaculatus             ATGGAAGATGTTATGTATTACATTTCGATAGTT--TCCCCCAACATCTC---------GTAAATATTTATAATTTCAGGTGAAATAAAACG---------------------AAAATAAT
Gryllus_longicercus             ATGGAAGATCTTATATATTACATTTCGATGGTT--TCAGCCAAAATCTT---------GTACATTTTTATAGTTTTAGATGAAATACAACA---------------------AAAATAAT
Consensus                                         * *

Teleogryllus_occipitalis        ------------------------------------------------------------------------------------------------------------------------
Acheta_domesticus               GAGTGTAACTAGTGTTAGCAATTCGAAAAGTTTGAAGAGAGAATTTGGATGCAATTT----TCTGAAAATGTCGTTTGAGCGTTTCTGAAAATGTGGA------ATTTTGGATACGA---
Gryllus_bimaculatus             GCATTTAATTCATTTTAG-AGTGTTACTTTCTTCCTGCGAAAATGTGATAATTATTTGTTCTACACAAGTATTACCAGCCCATCCCTAATATTACGGAGCGCTCATCTCAAATGCAGGGA
Gryllus_longicercus             GTATTTAATTAATTTTAG-AATATTACTTTCTTTTAGCGAGAATGGGGTATTTATTTGTCCTACACGAGTATTACCAG------CCTAATATTACAGAGCGCTCATCTCAAATGCAGGGA
Consensus

Teleogryllus_occipitalis        ------------------------------------------------------------------------------------------------------------------------
Acheta_domesticus               ----------CTGGCAAACTTTCTTTGAAATTATATAAAGCACGTGCACTCTACTTCTT------CTTCTTCTTCTTCTTCATTGCCAAGCGTTGGGAACCATTG--------------
Gryllus_bimaculatus             TTCCACGGTTGTGCCAAGTTGCTCCTTACCCTATCACGAGCACGACCGGTCGAGCGTCTAACGAGTCACTCGTTTTTTCCATGGGTCAAAGGGTCAATGCCGATGAGAAGAGTAATCCAT
Gryllus_longicercus             TTCCACAGTTGTGCCAAGTTGCTCCTTACCCTATCACGAGCACGACCGGTCGAGCATCTAACAAGTCACTCGTTTTTCCT-TGGGTCGAAGGTTCAATGTCGATGAGAAGAGCAATGCAT
Consensus

Teleogryllus_occipitalis        -------------------------------------------------------
Acheta_domesticus               ---GCCCGTTCCGAGATTCAGAACTGCTCTGACCTCTTCATTCTATA--------
Gryllus_bimaculatus             TCTGGTGATTCCGACGTCC------------------------------------
Gryllus_longicercus             CCTGGTGGTTCCGACGCCCCTTGCTGCTGTATACGTATTTTCATTTGCATGTAAG
Consensus
```

**Tocc.9G0009060.1_exon1**

```
Teleogryllus_occipitalis        ------------------------------------------------------------------------------------------------------------------------
Acheta_domesticus               ----TRTNNRNY-LQELKTDTIFSK*SFRFSNNLV*NVYFTS*IGPTTKYNPLSV--CL*KRQEGRYNV--YLMQNVNSP*HPTHSP*HKET-LCL*KLCL*K---VYNVSLHHSQ----
Gryllus_bimaculatus             ----TGSVNWYYVIKIIL*NCVF---------RLI*RKIFISLVNLVNFSIVISVL*VYTCRNVKKEKQCRFLIAERQLTVSYTHSPKHTVT-LCLRSL*GLQCFLTANSSFVL*SNSEN
Gryllus_longicercus             VRTDIISSK*YY-------RTVFCE---*FRIKLLFRLLTWATTPLVSTQCIL----V*TSR--KRYNVIVFLMQN-------VNSPYHTHTHLNIK*LFVFESFEVHNGSLLYSQ----
Consensus

Teleogryllus_occipitalis        ------------------------------------------------------------------------------------------------------------------------
Acheta_domesticus               ---RFILIAIEVQEQVTGLVYLDVFRHCRLYFK*YTILNASL*YNYRIYLI---HIIKCLPDGISSNTY--LHICKLNN-----------------*YLCLKKFQNCSR*S**FTLLHNS
Gryllus_bimaculatus             K**NFLS*IVSQLST*L*IIY---------DIESVFLI*LQVQVNTLNKVLVRRYFI*SL----------LTHFANLTTSICV*KSFKIAQDDTDYIYFCTIRFKCMEAC*LQLQCL---
Gryllus_longicercus             ---QFILIAIKVQEQITELLYLELFRYCRLDCKSYAMLKASF*YNYRFKLI---HLIKYLSDVISSSTY*LTHFANLTTSICV*KSFKIVQDDTDYIYFCRIRFKYMEAC*LQLQCL---
Consensus

Teleogryllus_occipitalis        ------------------------------------------------------------------------------------------------------------------------
Acheta_domesticus               F*GHGSTNDDAVS-FRELHEKVQRNYDAPRVTACITFF-FFSSFSETSNI----LMDPKRNRPWNNKRLPQQISSAKIK*KSRHKRPREKGR-----------LLHTKLLAPLSHYIHRW
Gryllus_bimaculatus             -----ST**LSVQGFSELHEKVQRK*EC---HSCYRMYHIF--FILFRNIKHSDRTEAKPSR--E**TPPTANFEHQIKEKSRHKRPGRRATSSHKTSCSF*VTLHTSV--P--------
Gryllus_longicercus             -----STY*LSVQGFSELHEKVQRMYEC---RSCYRM*HIFSSFSETSNI----LIEPKQNRLENNKRHPQQISSTK*RKKSRHKRPGRRATSSHKTSCFF*VTLHTSV--P--------
Consensus

Teleogryllus_occipitalis        --------------------------------------------------MLRRSGLAAHSAERGGGGGGGGGSCGSGRGGGGDGGGGGGGGGGGGGGGGGGGGGGGGGGGGGGGGGGG
Acheta_domesticus               LSTEALLREGFVLQFVPNHLARRSRQKPHTVRHSTQNASKVWLGCTL--GRARRRG--------GGGGGGGGG-----------------------------------------------
Gryllus_bimaculatus             -STEALLREGFVLPI-------RSKS------FSAQFPAETTYCATLDTGMLRRSGLAAHSAERGGGGGGGGGR----------------------------------------------
Gryllus_longicercus             -STEALLREGFVLPI-------RSKS------FSAEFPAETTYCATLDTGMLRRSGLAAHSVERGGGGGGGG------------------------------------------------
Consensus                                                                           ** *        ********

Teleogryllus_occipitalis        GGGGGGGGGGGGGGGGGGGGGGGGGGGGGGGGGGGGGGGGGGGGGGGGGGGGGGGGGGGGGGGGGGGGGGGGGGGGGGGGAGGGGGVGGGG-----------------------GGGGGG
Acheta_domesticus               ------------------------------------------------------------GGGGGGGGGGLSR*WRRGDVAGFTHGVGPAI-----------------------GCPRD*
Gryllus_bimaculatus             -----------------------------------------------------------RRGGGGGGAAGAAGR-RRPQSLMATR*RCRIH---------------PRSSALPLDCTRD*
Gryllus_longicercus             ------------------------------------------------------------RGGGGGGGGGLSR*WRRGDDAGFTHGVLPCHWIVRGIDKVSWFPSLPRAAMI----RRLE
Consensus                                                                                    ******  *

Teleogryllus_occipitalis        GGY-----GGGGGGGGGGGGGGGGGGGGGGG---------GGGGGGGGGGGGG--------GG---------------------------------------------------------
Acheta_domesticus               QGL-------------------------------------EVSS----------VP*A--------------------------------------------AMIR--------------
Gryllus_bimaculatus             QGLVVSLLAASSHDSKTGTGPITMVHRGHGTPFRRRRYPSEGKQ*R---------PS*RRKWGLDTVLS*RLWCSRFYVM---------------EDVMYYISMVSAKIS*MFIILGEMK
Gryllus_longicercus             QGPLQWCIGGTEHLSAA*D-------------IRRR----ESSSNRVDDGNGGLIPSLVIVGGVADFL*WKM*CITF**FLPKSHKCL*Y*VK*NENNAFY-SF*S--------------
Consensus                        *

Teleogryllus_occipitalis        -----------------------------------------GGGGGGGGGGGGR--------------------------GGGG--------GGGGG-----------------------
Acheta_domesticus               --------------------------------SLEQGL---YNGAPGAQSTFSR-----PEIPLGAAAIATELTTE------------------VWS*YRNELSXXXXXXXXXXXXXXXX
Gryllus_bimaculatus             RKSRILFILEYYFLFAKNGIFICSTRVLPAHHSPLQALPAHYNGASGAQNTFPW-----PEISVGWKAVATELTTE------------------MGA*YRP*LT----------------
Gryllus_longicercus             ----ITFFLQ------KWNIYLFYTSIT----SPSQPI-------TGLTSPLQWCIRRTEHLSVA*DVRRRESSSDRVDDGCGGLIPSLVNVGGVAAFS*WKIL----------------
Consensus                                                                     *

Teleogryllus_occipitalis        ------------------------------------------------------------------------------------------------------------------------
Acheta_domesticus               XXXXXXXXXXXXXXXXXLSAAICRLLKY---------IFSSFS*--FSVS*W*NRNIKFRILQNTKISS*M--EIFFEKFVTVFKNLEF*--CY*KCEILSSFLQKIFEL*KNFI*SHLK
Gryllus_bimaculatus             -----------FVV*QILRNGRCYVLHFDSFPQHLVNIYNFR*N-------------------KTKIMHLI----HFRVLLSSCENVIII--CSTQ-----------------VLPAHP*
Gryllus_longicercus             -----------YITFRWFQPKSCTFL*F-*MKYNKNNVFN*F*NITFF*REWG------IYLSYTSITSLILQSAHLK-----CRDSTVVPSCS--------------------LPYHEH
Consensus

Teleogryllus_occipitalis        ------SGGGLSR*----------------------------------------------------------------------------------------------------------
Acheta_domesticus               QFNY--KNNSASKLWEMC--ILTVY*RRPFNKITFV*FLGKKYTPN*TLRHNKECN*C*QFEKFEERIWMQFSENVV*AFLKMWNFGYDWQTFFEII*STCTLLLLLLLLHCQALGTIGP
Gryllus_bimaculatus             YYGALISNAGIPRLCQVAPYPITSTTGRASNE-SLVFSMGQRVNA------DEKSNPFW*FRR---------------------------------------------------------
Gryllus_longicercus             ------DRSSI*QVTR--------------------FSLGRRFNV------DEKSNASWWFRRP---------------------------------------LLLYTYFH---------
Consensus

Teleogryllus_occipitalis        -------------
Acheta_domesticus               FRDSELL*PLHSI
Gryllus_bimaculatus             -------------
Gryllus_longicercus             ---------LHV-
Consensus
```

**Tocc.10G0002420.1**

```
Teleogryllus_occipitalis      CCACTCTAAATTTCGTATTATTCCTGAGCCCCTGCCATCTTGGAATATATATTTTTTAAATCACTCTCTGAATGTAATATCTGAACAAGTCTGTGAACTACGGTCATTACCTACTGAACT
Acheta_domesticus             TCGCACTAAATTGCATATTGTTCCTGA-CCCGCACCATCTTGG--CATATATTTTTTAAATCACTCTCTGGA--GAGTGTCTGACTAAGTCACTCATCTACGGTCCTTGCCCATTGAATC
Gryllus_bimaculatus           CTGCACTAAATTTCG---ATGTTTTGAGCCCCCACGATCTTG---TAGAATTTTTTTAAACGACTCTCTGGATGGAATGTCTGAATAAGCCTGTGATCTACGGTCCCTGTGCATTGAACC
Gryllus_longicercus           TCGCACTAAATTTCG---TTGTTCTGAGCCCCCACGATCTTG---TAGAATTTTTTTAAACCACTCTCTGGATGGAATGTCTGAATAAGCCTGTGATCTACGGTCCTTGTGCATTGAACC
Consensus                       * ******* *      *  *** ***  * ******   *  *********  ******** *   * * *****  *** *   * * ********  *    * ****

Teleogryllus_occipitalis      ACTTCGAGTTGAATTTGTATGCAATAGCAAAGAAATAAAATGCATCTATTTATTTACTTATAGCAAAAGAAATATTGTATAAAATAAAAAAAAACATTTTAAAAATGAATTTTTAGAT-
Acheta_domesticus             ACTCCGAGCTGAATTTGTATGCAACAGCAAAGAAGAAA-------TTTTATTTTTTTGTGGAAAATGAAATATTATACATCAAACATTGCAAACACATTTTAAAAATGTATTTTTAGATT
Gryllus_bimaculatus           ACTGCGAGTTGAATTTGTATGTAATAGCAAAGAAGAAATGGA-----ATTTATTTACGTACAGAAAAGGAAAGATTGGAAAAAACACAATAAACACATTTAAAAACGTTAAC------TT
Gryllus_longicercus           ACTGCGAGTTGAATTTGTATGCAATAGCAAAGAAGAAATGGA-----ATTTATTTACGTACAGAAAAGGAAAGATTGGAAAAAACACAATAAACACCTTTAAAAACGTTAACTTC-GATT
Consensus                     *** **** *********** ** ******* **         *******   *  * *** ***** ***  *  *** *     *** ** *** ***    *         *

Teleogryllus_occipitalis      ----------------GGTATCTCCAAGATCGCGAATTAATCCTGAATCATCAATCCAACGCATAAATATTTCATAATGGAACGTAATAAACTTATAAATGTTACAGATATGACTA-
Acheta_domesticus             TACATGTTA-------ATGTGTCTCCAAGAATCGTGAACTGACCCTGAGTCATCAGTCCAATGTATAAACGAGTGACAGTAGGTAATAATATACTTATAAATGTTACCAAATGTGAGTAA
Gryllus_bimaculatus           CACAGGTTAATGATACTCGTATATCCAAGAATCGTGAACTGACCCTGAATTATCAGTCACCGTGCACAAACGAGTGGTAGTAGGGCATAATAAACTTATAAATGTTATCAAATGTAAATAA
Gryllus_longicercus           CACAGGTTAATGATACTCGTATATCCAAGAATCGTGAACTGACCCTGAATGATCAGTCACCGTGCACAAACGAGGGGTAGTAGGGCATAATAAACTTAT---------AAATGTAAATAA
Consensus                         * * ********** *** * * **** * *  *  ***         * * *     ***** *****             * * * * **

Teleogryllus_occipitalis      -----ACATTTGGCTCTC-TTTTGGTTTTCAATGTAAATCGGAAAACAACATTTTTTTTCCCAGAGAGTTATTTTCAAATTATTAATTATTATTTTACAGAAACATTAATAGTCATGTA
Acheta_domesticus             TGCACACATTTGACTCTT-TTTTGGTTTTGAAGGCAAATCGGAAAACAAC----TTTATTTCCAGAAAATTATTTTCAAATTATTAATTATTATTTATCAG-ACCATTATTGATCATA-A
Gryllus_bimaculatus           TACACACATTTGACTCTTATTTTGGTTTTGAACATAAATCGGGAAAGTAC----CTTGTTTCCAGGCAATTATCTTCAAATTTTTAATTATTATTTACCAG-AACATT-----TCATATA
Gryllus_longicercus           TACACACATTTGACTCTGTTTGGTTTTGAACATAAACCGGGAAAGTAC----CTTGTTTCCAGACAATTATCATCAAATTGTTAATTATTATTTACCAG-AACATT-----TTATATA
Consensus                          ******* ****  ********* **    *** ***  *****  **     ** * ***** * ****  ******* *** *********  *** * ****     * **

Teleogryllus_occipitalis      TTGTGAGTAATGACTATTGAATAAACTCTGTCAATTTTCTTTTGGTTAAATA-TATAATTTTGCATTACGAAACTATTTTCTTCCAAAGTGCAAAGGGCAAGTTTACCACAACCATAT
Acheta_domesticus             TTGTGAGTAATGACTATTGAATAAACTTTGT--TTCCTTCTGTATAAAAAA--TACAATTTTGCATTAAGAAACAATTTGCTTCCAAAGTGCAATGGGCAAGTTGACCACAACCGTAG
Gryllus_bimaculatus           TTATGAGTAATAACAACTAAATATACTGTGTCTCTTTCTTTTAGAAAAAAAAAGTATAATTTTCGTTACGAAACGATTTGCTTCCGAAGTGCAATGGGCATGTTTACCTCTAGCGTAT
Gryllus_longicercus           TTATGAGTAATAACCACTTAATATACTTGTGTCTCTTTCCTTTTAGATAAAAAAGTATAATTTTCCTTACGAAACGATTTGCTTCCGAAGGGCAATGGACATGTTTACCTCAAGCGTAT
Consensus                     ** ******** ** * * **** ** *   * ***  ***  **    *** * *  *  ******   * ** * ***** ***** ****** ** ** ** ***  * * * **

Teleogryllus_occipitalis      AACATGTCTGTCATTCAATGTATCGTGTATTTTCCGGCGTCAATCTTATTGCTGTTACGGCAATCAAACAGTTTGCAAAATCTTACTTGAGCCCCTAAAAATTATGATTCCACTCCACTA
Acheta_domesticus             AACATGTCTGTCATGCAATAGATCTTGTATTTTCGTGCATCAATCTTATTGCTGTGACGGCAATCAAACAGTTTGCAAACACTCACCTGCG-CCCTAAAAATTATGATTCCACACCACTA
Gryllus_bimaculatus           AACATGCCTGTCATATATTATATATCTGTATTTTCGTGCATCAATTTTACTGCTGTTATGGTGATTAAACAGTTTGCAAATCTTACTTGCGCCCGTAAAAATTATGATTCCACTCTACTA
Gryllus_longicercus           AACATGTCTGTCATATAATATATCTTATATTTTCGTGCATCAATCTTACTACTGTTATGGTGATTAAACAGTTTGCAAATCTTACTTGCGCGCGTAAAAATTATGATTCCACTCTACTA
Consensus                     ****** *******  * *   *** * *********  ** ** **** ****   ** ** **  *** ****  ********** **  * *  ******************  ****

Teleogryllus_occipitalis      ACAAACGTCGACGGCCCAGGCGTCCCTTGTAAATATTACCGCCGACTTGTACGCACCGCTGTTCTCAGCGCTCAGGCAAAATAGACGGATTATGCTTCCCACATTTTTTAGCCTTTCCTC
Acheta_domesticus             ACAAACGTCTGT-GCCCAGGCGGCCCTCGTAAATATTTCCGCCGACGAGTGCGC---------CCTCGCGCTCGAGCAAAATAGACGGATTATGCTTCCCACG-GTTTTGGTCTTTCCTC
Gryllus_bimaculatus           ACACACGTCTATTGCCCAGGCGTCCCTTGTAAATATTACCGCTGACTTGTGCGCACCGCTGTTCTCCGCACTCAGGCAAAATACACGAATTATGCTTCCCACGATTTTTTGTACTTCCTC
Gryllus_longicercus           ACA------------CAGGCGTCCCTTGTAAATATTACCGCTGACTTGTGCGGACCGCTGTTCTCCGCGCTCAGGCAAAATACACGGATTATGCTTCCCACGATTTTTTAGTCTTTCCTC
Consensus                     ***             ****** **** ********* ***** *** **  **          ** ***  ******** *** ******** ******    **** *    ******

Teleogryllus_occipitalis      TCTAAACACGATCCGAATATAAAAAAGAACTC-TTCAAAAGATCGCAAATTCATCTTAAA--GAAAAAACCGCTTTTTTCCTTATTTCATTCTCATGCAAATTGCCGATTGAACCTTGCTG
Acheta_domesticus             TCTAAACACGACCCGAATATAAAAAGGAACTC-TTCAAAAGATCGCAAATTCATCTTAAAGGGAAAAAACGCTTTTTTCCTTATTTCATTCTTATGCAAATTACCGATTGAACGTTGCTG
Gryllus_bimaculatus           TCTTAACACGACCAGAATGTAAAAAGAACTCTTTCAAAAGATCGCAAATTCATCTTACA--GAAAAAC---TTATTCCTTATTTCATTCTTATGCAAATTACCGATTGAACCTTGCTG
Gryllus_longicercus           TCTTAACACGACCAGAATGTAAAAAGAACTC-TTCAAAAGATCGCAAATTCATCTTACA--GAAAAAC---TTCTTCCTTATTTCATTCTTATGCAAATTACCGATTGAACCTTGCTG
Consensus                     *** ******* **** * **** ** ***** *************************** *  ********   ** ***************** ********** ********** ******

Teleogryllus_occipitalis      CCTGTTTATAATTGCGTTAAGACAACCGCAAAAGAGTTTTCTTTCAATTGCGAGAATTTAATAGGTTATTCTGAGCCGCCTCTGCAAATTGGCGAC----GCCTATCTTTTTTAGCCAC
Acheta_domesticus             CCTGTTTATAATTGCGTTAAGACAACCGCCAAAGAGTTTTCTTTCAATTGCGAGAATTTAATAGGTTATTCTGAGCCACCTCTGCAAATTGGCGACCCGGCCTATCTATCTTTTCGAGCCAC
Gryllus_bimaculatus           CCTGTTTATAATTGCGTTAAGACAACCGCAAAAGAGTTTTCTTTCAATTGCGAGATATTTAATAGGTTATTCGAGCCGCTCTGCAAATTGGCGAC----GCCTATCTTTTT-AGCCAC
Gryllus_longicercus           CCTGTTTATAATTGCGTTAAGACAACCGCAAAAGAGTTTTCTTTCAATTGCGAGATATTTAATAGGTTATTTGAGCCGCTTCTGCAAATTGGCGAC----GCCTATCTTTTT-AGCCAC
Consensus                     ***************************** ************************* ***************  ***** * ****************       *********  ******

Teleogryllus_occipitalis      TTTTCCGCACCCGAGCAAAGAAAATGGCGGGATCTGGTTTTTCTCCGGGCCCGCCCGTGGCGCCAGAGAAAACATTGACACAATCCCCGGCCGCCCCGCAATTACGCTCCCTGTTATTCC
Acheta_domesticus             TTTTCCGCACCCGAGCAAAGAAAATGGCGGGATCTGGTTTTTCTGTGGGCCCGCCCGTGGCGCCAGAGAAAACATTGACACAATCCCCGGCCGCCCCGCAATTACGCTCCCTGTTATTCC
Gryllus_bimaculatus           TTTTCCGCACCCGAGCAAAGAAAATGGC-GGATCTGGTTTTTGTCCGGGCCCACCCGCGGCGCCAGAGAAAACATTGACACAATCCCCGCCGCCCCCGCAATTACGCTCCCTGTTATTCC
Gryllus_longicercus           TTTTCCGCACCCGAGCAAAGAAAATGGCGGGATGTGGTTTTTCTCCGGGCCCACCCGCGGCGCCAGAGAAAACATTGACACAATCCCCG-CCGCCCCGCAATTACGCTCCCTGTTATTCC
Consensus                     **************************** **** ******** *  ****** **** *************************************** *************************

Teleogryllus_occipitalis      ATTGTTTCTCGCACTGGCGCCGCTCCGCGGCGGCGA------------------------GGCGGTGGAGGCGGTGG------AGGCGGTGGAGGCGGTGGAGGAGGC-------AAG
Acheta_domesticus             ATTGTTTCTCGCACTGGCGCCGCTCCGCGGCGGTGGAGGCGGTGAAGGTGGCCGCGGAGGTGGCGGTGGAGACGGCGGTGGAGACGGCGGTGGAGACGGCGGTGGAGACGGCGGTGGAG
Gryllus_bimaculatus           ATTGTTTCTCGCACTGCCGCCGCTCCGCGGCGGA------------------------GGTGGTGCAGGCCGTGG------AGGC-------------------------------------
Gryllus_longicercus           ATTGTTTCTGGCACTGCCGCGCGCTCCGCGGCGGA------------------------GGTGGTGCAGGCCGTGG------AGGCAGTGGACGC---------------------------
Consensus                     ********* ****** *** *************                          ** **** ** * * **        ***

Teleogryllus_occipitalis      GCGGAGGTGGTGGAGGAAGTGGAGGAGGCGGAGGAGGCGGAGGAGGTGAAGGAGGCAGAGGAAGTGGAGGAGATGGAGGCGGTGGAGGCGGTGGAGGAGGTGGAGGCGGTGGAGGAGGTG
Acheta_domesticus             ACGGCGGTGGAGACGGCGGTGGAGACGGCGGTGGAGACGGCGGTGGAGACGGCGGTGGAGACGGCGGTGGAGACGGCGGTGGAGACGGCGGTGGAGACGGCGGTGGAGACGGCGGTGGAG
Gryllus_bimaculatus           -------------------------------------------------------------------------------------------------GGTAGAAGCAGTGGGGGCAGTGGAGGTGGTG
Gryllus_longicercus           -------------------------------------------------------------------------CTGCAGGTGGAGGTGGTGGAGGAAGTAGAGGCGGTGGAGGCAATGGAGGCGGTG
Consensus                                                                                                                 ** **    * **  *     **  * *  *  *

Teleogryllus_occipitalis      AAAAGATGGAGGCGGTGGAGGAGGCGAAGGAGGTGTGGACGGTGGAGGAGGGCAAAGGA------GGTTGAGGCAGTAGAGGAAACGGATGAGGTGGAGGCAGGGGAGGAGGCGAAGGA-
Acheta_domesticus             ACGGCGGTGGAGACGGCGGTGGAGACGGCGGTGGAGACGGCGGTGGAGACGGCGGTGGAGACGGCGGTGGAGACGGCGGTGGAGACGGCGGTGGAGACGGCGGTGGAGACGGCGGTGGAG
Gryllus_bimaculatus           GAGGAAGTAGAGGCGGTGGAGG-------------------------------------------------------------------------------------------------
Gryllus_longicercus           GAGACGATGGAGCGGTGGAGG--------------------------------------------------------------------------------------------------
Consensus                          * *** *** ** **

Teleogryllus_occipitalis      -----GGTGGAGACGGTGGAGGAGATGG---------AGGAGATGGAGGAGGCGGAGGAGGAAGTGGAGGCGGAGGAGGCGATAGAGTAGGCGATGGA-------------------------
Acheta_domesticus             ACGGCGGTGGAGACGGCGGTGGAGACGGCGGTGGAGACGGCGGTGGAGACGGCGGTGGAGACGGCGGTGGAGACGGCGGTGGAGACGGCGGTGGAGACGGCGGTGGAGACGGCGGTGGAG
Gryllus_bimaculatus           -----------------------------------------------------------------------------------------------------------------------
Gryllus_longicercus           -----------------------------------------------------------------------------------------------------------------------
Consensus

Teleogryllus_occipitalis      ---------------------------------GGAGACGTGGGAGGTGGAGGCGGTGGAATAGGCGATAGAGGACGCGGAGAGAATGGAGGAGGTGGTGGAGCCTACGATGGAGAGGT
Acheta_domesticus             GACGGCGGTGGAGGCGGCGGTGGAGACGGCGGTGGAGGCGGCGGTGGAGGCGGCGGTGGAGGCGGCGGTGGAGGCGGCGGTGGAGGCGGCGGAGGCGGCGGAGGCGATGGAGGCGGTGGAG
Gryllus_bimaculatus           ------------------------------------------------------------------------------------------------------------CGGCGATGGAGAGAT
Gryllus_longicercus           ------------------------------------------------------------------------------------------------------------CGGCGATGGAGAGAT
Consensus                                                                                                                               ********** *

Teleogryllus_occipitalis      TGAATGATGTGTGGAAGCAGAAGGAGGCGTTGGAGGCAAGTTCAGTACCGGTGGGGCAAGGGAGATAGGGTGCGGAGAC--TAGACGAAAAACAAG---------GGCTGTGATAAAGG
Acheta_domesticus             TAAATGATGTGTGGTGGCAGGAGA---GTCGGAGGCGAGTGCTGCGCAGGTCAGGCAGTGTAGATGAGGTGCGGAGACTATAGACGAAAAACAA-CGGTTTT---GGCAGTGGTAA---
Gryllus_bimaculatus           TCGACTATGTGTGGAGGGAGAGGGGGAGTTGGAGGCAAGAACAGTTCCGGTGAAGCAGGCGAAGGTGGGGTGCGGAGAC--TAGACAAATAACAG----------GGTTGTGGAATAGG
Gryllus_longicercus           TCGACTATGTGTGGAGGGAGAGGGGGAGTTGGGGGCAAGAACAGTTCCGGTGGAGCAGGCGAAGGTGGGGTGCGGAGAC--TAGACAAATAACAAGCAGGAATGGAATAGGTTGTGTAATAGG
Consensus                     *  *  ********  * ***    * ** ***   ** ***   ** ***  ** ** *   ***   *** * ** ************   ***** ****            **  ***  *

Teleogryllus_occipitalis      GGCGGTGACGGGATAGTCATTGAATTTGACATGGCGTGGCTATGGTG-------------------------------------------------------------------------
Acheta_domesticus             --------GGGATCGTCACTGAATTTTGCATGACGTTGCGAAGGAGGCAGTGAGGAGGCAGTTGCCTGTGAAGAGTGGATCAGTAGACGTGGATGGTAGCCGAGTGACAGATACCGAAGCGGTA
Gryllus_bimaculatus           GACTGTGACGGGATCGTCATTGAATTTGATATGACGTTGCGATGGAGGTGGAACGTCCTGCTATGAAGAGGCAGATCA-----------------------------GACGTGGGA
Gryllus_longicercus           GACTGTGAC-GGATCGTCATTGAATTTTGC-----------GAGGAGGTGGAACGTCCTGCTATGGAGAGGCAGATCA-----------------------------GATGTGAGA
Consensus                               **** **** *******                  ** *

Teleogryllus_occipitalis      -----------------------------------------------------------------------------------------------------------------------
Acheta_domesticus             GAGGAACAAAGTGGTGGTGTAAACGGGGAGGCGTGGAGGAAGCCGAACAAGAAGGGGCAGGGTAGTGATAGTGTGAGACAAAGAAGTGATTGGGTGACTTGTGTGTGGCAGGAAGATGAAT
Gryllus_bimaculatus           GAGGA--AAACTGCGGTGCCGACGGGGAGGCGTGGAAAGGGTCGAACGAGTAAGGACGGGGTCG-------------AAAGGGTGGTACAGGTTGTAGGGGGAGGGCGGCGGAAATGGC
Gryllus_longicercus           GA-------------------------CGTGGAAAGGGTCGAACGAGTAGGGACGGGGTAGTGTCAGTGT----TAAGGTGAGGTGTGAGTGGCAGTGGGGAGGCGG------GGGC
Consensus

Teleogryllus_occipitalis      -----------------------------------------------------------------------------------------------------------------------
Acheta_domesticus             CGTAGGCAGGAAAGGAGACGGTGGAGGAAAATGTTCGCAGGATCTTGGGGACAGGGC-----------AGGGAGACGACGGAGAAACGACAAGGCGGTGGTTA----CCGTCACACCGT
Gryllus_bimaculatus           AACGAGGGGGAGGAGCAGCGGTGAAG-----------AGGAGGATGACGAAGGGGC-----------GAAGAGATGCTTGAGAACTGGTGCGGAGGAGCAGACAAGCAGTAGCGCAAG
Gryllus_longicercus           AGCGGGGCGAATGTGCTGTGATGGCG-----------ACGGGGTCGAAAGGGTGGTACAGGTTGTAGGAGGGGAGGGGGTGGAGCAGCGGTGAAGAGGAGCAGACAAGCAGTAACGCCAT
Consensus

Teleogryllus_occipitalis      ------------ATGGTGGCTTGTTAAG---------------------------------------------------------------------------------------------
Acheta_domesticus             TGCGACGACGGTGTGGTGGCTAGTCGAGTGGTAGGGATTGAGATTGGAGGAGGGGCGGAGATGCTGG--------------------------------------------AGCTGGA---
Gryllus_bimaculatus           GGCAGGGGCC--CTGGGCGATGGCCAGA--GTAGGACCTGGCGGAGGAGGGTCAGTGGAGGCGATGGGTCAATTACGTTCGAAGAAGCGGGTTGACACATTGGAAGGGACAGCGAGATGA
Gryllus_longicercus           GGCAGGGGCC--CTGGGCGATGGCCAGAGTGTAGGACTTGGTGGAGGAGGGTCAGTGGA--------------------------------------------------AGCGAGATGA
Consensus                                ***  * * *

Teleogryllus_occipitalis      -----------------------------------------------------------------------------------------------------------------------
Acheta_domesticus             --------------------------------------------------------------GAGGCAATGAGGGGGTGGAGCGGCGG------------------------------TGAT
Gryllus_bimaculatus           CAGCGGACAGTCCAAGGCG----------------------GACTTACTCCATCGATGCAGGGAACGAGGGGTCTGACGGAAGGAGCACGGGAAGAGGTGATTCGATTGATTGCTGAT
Gryllus_longicercus           CAGCGCTGAGTCCAAGGCGGAGCGCCTCGCGTTCTGCGTTCGTGACTTACT-CATCGATGTAGGGAGCGA-GGGTCTGACGGAAGCAGCGCGGGAAGAGGTGATTGATGGATTGCTGAT
Consensus

Teleogryllus_occipitalis      ---------------------------------------------------------CTCTGTAATGTAGCGGT-----------------------------------------------
Acheta_domesticus             TCCGGCGTGTCAATTATGGACGTCATTAGTCCGTAAGCATAGTACGTGTTTTTACCACCGGGAGGTGGCGGTGATCTTTT----CCCCGCTGGATGGCGAGATCCGGCG-CGACCTAC
Gryllus_bimaculatus           TCCGGGTGCGTCAACTGTTAAGCCCATT-------------------------------------------GCGGTGGTCT------CCCTCGTTGGACGGCGAGAATCAGCATGTTGCCTGT
Gryllus_longicercus           TCCGGGTACGTCAGCTGTTGAGCCCATT-------------------------------------------GCGGCGGTCTCCCCCCGCCCCCTGGACGGCGAGATCTGAATGTTGCCTGT
Consensus                                                                                                   ****

Teleogryllus_occipitalis      -----------------------------------------------------------------------------------------------------------------------
Acheta_domesticus             CGTTGACTAGCGGGGTTTTCCCCTCAACCAGGTAGCTC-------------------CCACCAGCGG--------------------------------------------------------
Gryllus_bimaculatus           TGGCGGTC-GCAGGATCTCTCGTTACACAGAATGGTTCAAGGCCTAATGTGGACCTCAG-ACACGTGCGTCATGATGCAATAAACCTTGAGTGATTTTAGGTGTTTGCAATATCATAGGG
Gryllus_longicercus           TGGCGGTC-GCAGGATCTCTCGTTACACAGAATGGTTCAAGGCCTAATGTGGACCTCAGAACACGTGCGTCATGATGCAATAAACCGTGAGTGATTTTAGGTTTGCAATATCATAGGG
Consensus

Teleogryllus_occipitalis      -----------------------------------------------------------------------------------------------------------------------
Acheta_domesticus             ----------GGCACCTCTCCAGCCACTGTAAGCTCCC--------------------------------------------------GATCGGTCTCCCGGCATGTGGTACTGTAGAA
Gryllus_bimaculatus           GAGACCAGGCTACATTTATACGAACATTATGAACTGTCGTGCATTTGTGTTTTAAATTTCTCAACAGTTATTAAAACGGTTATTCAAGAACCATTCTGTCGGTTTACTGTTAATTGTATTT
Gryllus_longicercus           GAGACCAGGCTACATTTATACGAACATTATGAACTGTCGTGCATTTGTGTTTTAAATTTCTCAACAGTTATTAAAACGGTTATTCAAGAACCATTCTGTCGGTGTACTGTTAATTGTATTT
Consensus

Teleogryllus_occipitalis      -----------------------------------------------------------------------------------------------------------------------
Acheta_domesticus             TATATTAGTGTAAATGTGTGTGTAAGGGGGTGAGT-----ATAGGACAAGGAAGAGGAAGGAAAGT---------------TCACG---------------------------------T
Gryllus_bimaculatus           TATATT----TAATCGCGTGAAGAAGGAATCGGCTCAGAGACGGAGCGATGGTGCGGAGGAAGAGGGGC---------------TGACGAGGAGGCGCCGCTCCACTGCACTGGAAACGGGCT
Gryllus_longicercus           TATATT----TAATCGCGTGAAAAAGGAATCGGCTCAGAAACGGAGCGATGAGGTGGAAGAGGGGCAACGAAGATGAGGCTGACGAGGAAGGCGGCGTCCACTGCATTGGAAACGGGCT
Consensus

Teleogryllus_occipitalis      -----------------------------------------------------------------------------------------------------------------------
Acheta_domesticus             GCGCTCC-------------------------------------------------------------GTAACCCGTTTCTT------AGGGTACTGCTCC-------
Gryllus_bimaculatus           GCGCTTCCAGCTACCTCCATCGCCTTTTATTAACTATGTTAATTACTCTTCCCTGGCCCCGCCGGAGCCAAGTCTTTGTGGTAATCTGTTTTTCTTTGGGAGGGCGGCGGTTTCCTCTCT
Gryllus_longicercus           GCGCTTCCAGCTACCTCCATCGCCTTTTATTAACTATGTTAATTACTCTTCCCTGGCCCCGCCGGAGCCAAGTCTTTGTGGTAATCTGTTTTTCTTTGGGAGGGCGGCGGTTTCCTCTCT
Consensus

Teleogryllus_occipitalis      -----------------------------------------------------------------------------------------------------------------------
Acheta_domesticus             -----------GGCGTTCA-----------CATGGATTGGAAAACCACAGAAAACTGC---------------------------------------------------------------
Gryllus_bimaculatus           TTTCCACCAGGGGCGCCCTTGGTGGGCCATTCGCGGACTTCAAAAAGAGGGAGGACNNNNNNNNNNNNNNNNNNNNNNNNNNNNNNNNNNNNNNNNNNNNNNNNNNNNNNNNNNNNNNNN
Gryllus_longicercus           TTTCCACCAGGGGCGCCCTTGGTGGGCCATTCGCGGACTTCAAAAAGAGGGAGGAAGACAGCTCCGCAGTGACGGGGGGCGGCGAGGACGGCCGCGGGTGGAGGATGGCGAGAGCGGTGGCG
Consensus

Teleogryllus_occipitalis      -----------------------------------------------------------------------------------------------------------------------
Acheta_domesticus             --------------------------------------------------------------------------CGATGAGG--------------GCAGCCGGCTGCCAGAATTCTCCTCT
Gryllus_bimaculatus           NNNNNNNNNNNNNNNNNNNNNNNNNNNNNNNNNNNNNNN--------------------------------GGGTGGGGTGGGGGAGTGGGGTTGCGACCAACCCTCAGGCCCCCTACGC
Gryllus_longicercus           GGGGCGGCGTAGACGGTGGCAGGGACGGCAGCGGCGGCGGGGACGGCGGCGGGGACGGAGGGAGCGGCCGTGGGCGGGGTGGGGAAGGGGGGTTGCGCCCAACCCTCAGGCCCCCTACGC
Consensus

Teleogryllus_occipitalis      ------------------------------------------------------------------AGATAAGATGTG-------------------------------------------
Acheta_domesticus             AGAGA-----------------------------------GCGTCGGAATTCGACCAAGCAGACAAGCTGGAGTCGAAAATCTCGCCGC-------------------------------
Gryllus_bimaculatus           AGGCGTGTTATGTTCCGCCATTGTTTCCACTCTATTCATTTCGTGTTAACTTTTGTCCG---AGCGATCCTGTGGACGGAAGCCTTGTCTCCCGCCTCTGCCCGCCATGCTTTATGACCT
Gryllus_longicercus           AGGCGTTTTATGTTCCGCCATTGTTTCCAGTCTATTCATTTCGTGTTAACTTTTGTCCG---AGCGATCCTGTGGACGGAAGCCTTGTCTCCCGCCTCTGCCCGCCATGCTTGATGACCT
Consensus                                                                                        **  *   **

Teleogryllus_occipitalis      --------------------------------------------------------------------------------
Acheta_domesticus             --CGTCGGGACTCGAACC------------------------------CAGGACCTTCCG--------------------
Gryllus_bimaculatus           AACATTGTAGTTCAATTTTCTCTTTAACGCGTTTCGAAAAAAGGAGAGAAGTCCCTCTGTGTCCCGTGTGTTTGAGTTTC
Gryllus_longicercus           AACATTGTAGTTCAATT
Consensus
```

**Tocc.10G0002420.1_exon1**

```
Teleogryllus_occipitalis      -----------------------------------------------------------------------------------------------------------------------
Acheta_domesticus             DLKHSTTFMFRSMNKILFKITLKY*SFRKLPTIHPDAFATSVQK---CAP*VRLSSELNRILRAGFHIECNPTTLICNEFSERCHQRNVFR----------IHLYIRAHVV*R-TTLFER
Gryllus_bimaculatus           --------------------ECKIVRYKQV--*APD*IICPVQDFTLCA------------------------------F*KLSFATNFRRGVISAMYLG--YIYTSACTFYNITTLFER
Gryllus_longicercus           ---------------------------------LNHMLCRISH---CL------------------------------RSKNSHLQRIFGEVSSAQCI*ATFIYLRARSIT*LPCLKDL
Consensus

Teleogryllus_occipitalis      -----------------------------------------------------------------------------------------------------------------------
Acheta_domesticus             L--------HSG*D*TPNR----------------IVIV*QPQRVTNCLPLSSTKIIA*IVITR*NHLNNYSN--RRN------AHCNIFTRSK--------MIIKTKTRSGVLIAQR-Y
Gryllus_bimaculatus           L------------NSAPNR--IAATMESYDNPNASPIVYHYPQLK*S--PKSSSRGIF*TTI-----------------PTGAMFAATFSRTA--------MIIKTKTRSSVLIAQR-Y
Gryllus_longicercus           ILHLIGLQPQWNHTSTPTRHQLFTIILN*NNP-----INHQHEV--------SSKQLFQQAPYSLQHFHEQQ*LLKRKLDPAC**HNATS**TERHPYRLVWAVMRLAARGMR*RA*REF
Consensus

Teleogryllus_occipitalis      -----------------------------------------------------------------------------------------------------------------------
Acheta_domesticus             QLINRATSRSFGLGRNAIGCSGNEITCLARISRAA-----PIKRVAHFQGSRNSFGMAGPSGR*KSQNRITQTNK-*TVAER*LPERKFARKKGNGGFKVRASDHTNRRSKLLSNFAYFE
Gryllus_bimaculatus           *LINRATSISFGLGRNEIGCSGNEITSLARISRAAPLNEWPISRAAGIHLS*LV--------RVTGESRKTASHKQINNHRGRITSRTEIYE-GKGKWRI*SA---SKRPHKLTVKAVIK
Gryllus_longicercus           QEPRRLTSGPFP-GQPEFICHD--------------WSE*PVKVAKPHH-----------------------TNK*ITIAVG*LPERKCTRERGNGGFKVRASDHTNRRSKLLSSFASFE
Consensus

Teleogryllus_occipitalis      -----------------------------------------------------------------------------------------------------------------------
Acheta_domesticus             NYLCSALQYVAFR*FL*EYFTEKKLNKVKIKTRS*DIKSFINGNSLTDR---------------------GIGEGR-----------------PARKC------------------TIS
Gryllus_bimaculatus           FYCFY---LIFIRIFHR-----KQFNIVNVKTSF*NIKCFINGNSLGIEKIQAW-SLELYTINWSIFGVLIGIIEGKKCLRSPRT**SSFELDVSRKTCQVSAA*YVQ*SNLRKYISVIS
Gryllus_longicercus           TYV*YS-ALLLLRNFYKNIL-QKTIQYC*CENKFLKHKMLYKRKFLGNSKNSGVKSTTIHNKWWPIFGVLIGIIEGKKYLRSPRT**SSSELDFPRKACQVSAA*YVQ*SNLRKYISVIS
Consensus

Teleogryllus_occipitalis      -----------------------------------------------------------------------------------------------------------------------
Acheta_domesticus             KNKTTRKSSSFK*WKCFY*GSGKYI*KHVVFAVRPHCNLC*RQSRVWLDDSQIALNCILFLTRTILAYIF*ITLW------------------RVSD*-------VTH-----------
Gryllus_bimaculatus           KNKTTRK-SCFKCCKCSY*ESGKQN*KHVASTVSPDCNLR*RQSRGDLVDSKAALNFDVLSPHDLVEF-F*TTLWMECLNKPVIYGP--------CA---------LNH---------CE
Gryllus_longicercus           KIK----------------------PHESHRVSNVVNVLTEKWKTKL-KTRGVCCFS*LQS-------------ALKAKPRVFG*LTSRTKFRCSEPPRSCRIFLNHSLDGMSE*ACD
Consensus

Teleogryllus_occipitalis      ------------------------------------------------------------------------------------M-----------------------------
Acheta_domesticus             LRSLPIESLRAEFVCNSKRR-NFIFLWKMKYYIKHCKHILK-----------MYF*IYMLMCLQES*TDPESSVQCINE*Q*VIIYL*MLPNVSNAHI*LF------FG----FEGKSEN
Gryllus_bimaculatus           LN--------------------LYVIAKKKWNLFTYRKGKIGKNTINTFKNVNFTG**YSYIQES*TDPELSVPCTNEW**GIINL*MLSNVNNTHI*LLFWF*T*IGKVPCFQAIIFK
Gryllus_longicercus           LRSLCIEPLRVEFVCNSKEEMEFIYVQKRKDWKKHNKHL*K---------R*LRFTG**YSYIQES*TDPE*SVPCTNEG**GIINL*M---------*IIHTFDSCFG----FEHKPGK
Consensus                                                                                                         *

Teleogryllus_occipitalis      -----------------------------------------------------------------------------------------------------------------------
Acheta_domesticus             NFISRKLFSNYYLLFIRPLLIIIVSND-Y*INF-----VSFCIKNTILH*ETICFPKCNGQVDHNRR----------TCL--SCNRSCIFVHQSYCCDGNQTVCKHSPAP*KL*FHTTNK
Gryllus_bimaculatus           FLI---------IIYQNISYIMSNNN*IYCVSFLLEKKV*FFVTK-RFASRSAM-----GMFTSSV*HACHILYLVFSCI----NFTAVMVIKQF----AKSYLR----P*KL*FHSTNT
Gryllus_longicercus           YLVSRQLSSNC*LLFTRTFYIL*VITT*YTLS------LSF*IKKYNFSLRNDLLP--EGQWT---------------CLPQAYNMSVI*YILYF----RASIL--------LLLW*LNS
Consensus

Teleogryllus_occipitalis      -----------------------------------------------------------------------------------------------------------------------
Acheta_domesticus             RLCPGGP---------RKYF---------------------------RRRVRPRARAK*TDYASHGFGLSSLNTTRI*KGTLQKIANSS*REKTLFSLFHSYANYRLNVAAC----L*L
Gryllus_bimaculatus           RLLPRRPL*ILPLTCAHR------------------CSPHSGKIHE---LCFPRF-----------FVLPLLTRPECKNELFQKIANSSYRKNLFLISFLC----KLPIEPCCLFIIAL
Gryllus_longicercus           ---------LQNLTCARKNYDSTLLTQASLVNITADLCGPLFSALRQNTRIMPPTI-----------FSLSSLNTTRM*KRTLQKIANSSYRKNFFLISFLC----KLPIEPCCLFIIAL
Consensus

Teleogryllus_occipitalis      -----------------------------------------------------------------------------------------------------------------------
Acheta_domesticus             R*DNRQRVFFQLREI**VILSHLCKLATRIYLFEPLFRTRAKKMAGSGFSVGP-----------PVAPEKTLT------QSPAAPQLRSLLFHCFSHWRPLRGGGGGEGGRGGGGGDGGG
Gryllus_bimaculatus           R--QPQKSFLSIARYLIGYFEPLLQIGD-AYLFSHFSAPEQRKWR-IWFLSGPTRGARENIDTIPARPAITLPVIPLFLALPPAPRRR----WC-RPWRR*KQWGQWRWWRK*RRWRR--
Gryllus_longicercus           R--QPQKSFLSIARYLIGYFEPLLQIGD-AYLFSHFSAPEQRKWRDVVFLRAHPRRQRKH*HN-PRRPAITLPVIPLFLALPRAPRRR----WC-RPWRQWTPAGGGGGGSRGGGGNG--
Consensus

Teleogryllus_occipitalis      -----------------------------------------------------------------------------------------------------------------------
Acheta_domesticus             DGGGDGGGDGGGDGGGDGGGDGGGDGGGDGGGDGGGDGGGDGGGDGGGDGGGDGGGDGGGDGGGDGGGDGGGDGGGDGGGDGGGDGGGDGGGDGGGDGGGDGGGDGGGDGGGDGGGDGGG
Gryllus_bimaculatus           ----------------------------------------------------------------------------------------------------------------RWRDSTM
Gryllus_longicercus           ----------------------------------------------------------------------------------------------------------------RRGDDGA
Consensus

Teleogryllus_occipitalis      -----------------------------------------------------------------------------------------------------------------------
Acheta_domesticus             DGGGDGGWR--------RR--WRRRWRRRWETAVEAAVEAAVEAAVEAAVEAAVEAAEAMERLNDVWVAGGVGGECCAGRAV*MGCGDYRRKTRQW*GIVTEFCMTLRRRQ*TPL*RSGS
Gryllus_bimaculatus           CGREEGELEARTVPVKQRR--WGAET----------------------------------RQITGLW-------------------------NRDCDGIVIEFCMTLRWRWNVLL*R-GR
Gryllus_longicercus           GGGGDGEIR---LCVGGRRGSWGQEQF-RWSS-----------------EGGVRRLDK*QEWNRLC------------------------NRDCDGSSLNFAR----RWNVLLWR-GR
Consensus

Teleogryllus_occipitalis      -----------------------------------------------------------------------------------------------------------------------
Acheta_domesticus             VDV-DGSRV---------TDTEAVEEQMVV*TGRRGGSRTRRGRVVIV*DKEVIG*LVCGRKMNRRQERRRWRKMFAGSWGQGRETTEKRQ----GGGYRHTVATTVWWLVEW*GLR---
Gryllus_bimaculatus           SD---------VGEENCGADGEAWKGSNE*GRGRKGGT-GCRGRAAEMATR-------------------------GRSSGEEEDDEGAKRCLRTGAEEQTSSSARAGALG---------
Gryllus_longicercus           SDVRDVERVERVGTG*CQC*GEV-----*VAVGRRGQ----RGECAVMAT--------------------------GSKGWYRL*EGGG-----GAAVKRSRQAVTPWQGPWAMARV*D
Consensus

Teleogryllus_occipitalis      -----------------------------------------------------------------------------------------------------------------------
Acheta_domesticus             -LEEGR-RCWSWR----------------------------------------GNEGVERR*FRACQLWTSLVP*A*YVFFHREVAVIFSPLDGEIRRRPTVD*RGFPLNQVAPTSGAP
Gryllus_bimaculatus           ---DGQSRTWRRRVSGSDGSITFEEAG*HIGRDSEMTADSPR-----RTYSIDAGNEGSDGRSTGR--------GDS--IDC*FRVRQLLSPL----RWSPPWTARISML----PVGGRR
Gryllus_longicercus           LVEEGQ---WK--------------------RDDSAESKAERLAFCVRDLLIDVGSEGLTEAAREEVI*W---IADSGYVSC*AHCGGL--P-------PPPWTARI*ML----PVGGRR
Consensus

Teleogryllus_occipitalis      -----------------------------------------------------------------------------------------------------------------------
Acheta_domesticus             L-------QPL*APDRSPGM---------------------CGTVEYISVNVC------VRG*V*DK-----------------------------------------------------
Gryllus_bimaculatus           ISRYTEWFKA*CGPQT--------------------RAS*CNKP*VI-LGVC--------NIIGETRLHLYKH*LSCICVLNSQQLLKRLFKNHSVGLLLIVFYI*SREEGIGSETERC
Gryllus_longicercus           ISRYTEWFKA*CGPQNTCGMMQ*TVSDFRFLQYHRGDQATFIRTL*TV-VHLCFKFSTVIKAVIQEP---------FCRCTVNC-----------------ILYLIA*KRNRLRNGAMRW
Consensus

Teleogryllus_occipitalis      -----E-----------------------------------------------------------------------------------------------------------------
Acheta_domesticus             -----EEEGKFTC------------------------------AP*P-----------------------------------------VS*GTAPAFTW--------IGKT--------
Gryllus_bimaculatus           GGRGADEEGGVHCTGNGLRFQLP------PSPFINYVNYSSLAPPEP----SLCGNLFFF------------------------GRAAVSSLFHQGRPWWAIRGLQKRGRTXXXXXXXX
Gryllus_longicercus           -KRGNEDEA-----GRGRRRPLHWKRAALPATSIAFY*LC*LLFPGPAGAKSLW*SVFLWEGGGFLSFPPGAPVVGHSRTSKKREDSSAVTGGGEDGRGWRMARAVAGAA*T--------
Consensus

Teleogryllus_occipitalis      -----------------------------------------------------------------------------------------------------------------
Acheta_domesticus             -------------------TENCR*G--------QPAA---RILL*RAS--------EFDQADKLESKILPPSGLEPRTF---------------------------------
Gryllus_bimaculatus           XXXXXXXXXXXXXXXXXXXXXXXXXGGVGEWGCDQPSGPLRRRVMFRHCFHSIHFVLTFVRA------ILWTEALSPASARHAL*PNICSSIFSLTRFEKRREVPLCPVCLSF
Gryllus_longicercus           -----VAGTAAAAGTAAGTEGAAVGGVGERGCAQPSGPLRRRFMFRHCFQSIHFVLTFVRA------ILWTEALSPASARHAS*PNICSSI--------------------
Consensus
```

**Tocc.10G0002420.1_exon2**

```
Teleogryllus_occipitalis     ----------------------------------------------------------------------------------------------------------------------------------------------------------
Acheta_domesticus            FKTFHHIHVSFYE*NFIQNHVKILKLPKTPYYPS*CVCDQCAKMRAVSTAELRTKSHIARRISHRVQSNNSHLQRIFGEVSSTQRI*DTFIYPRARCITNYLV*KTSFWMRLNT**DCNR
Gryllus_bimaculatus          ---------------------------------------SAKLCAISKSELRTKSYALCRISHCARSKNSHSQRIFGEVSSAQCI*ATFIHLRARSIT*LPCLKDLILHLIGLPPQWNH
Gryllus_longicercus          ----------------------------------------------------LNHMLCRISHCLRSKNSHLQRIFGEVSSAQCI*ATFIYLRARSIT*LPCLKDLILHLIGLQPQWNH
Consensus

Teleogryllus_occipitalis     ----------------------------------------------------------------------------------------------------------------------------------------------------------
Acheta_domesticus            MTTPTRHQLFAVILN*NNRINRHHEVESSKQLFQQAQRALQHFHEKQNDY*NE-NPIRRADSTTLPANKQSDIQIVWFGPQCDWLLGE*DNVFSANFKGRA--------H*TSGPFPGQP
Gryllus_bimaculatus          TTTPTRHQLFTIILN*NNPLNHHHEV-SSKQLFQQAPCLLQHFHEQQ*LLKRKLDPAC**HNATS**TERHPYRLVW----------AVMRLAARGMR*RA*REFQEPRRLTSGPFPGQP
Gryllus_longicercus          TSTPTRHQLFTIILN*NNPINHQHEV-SSKQLFQQAPYSLQHFHEQQ*LLKRKLDPAC**HNATS**TERHPYRLVW----------AVMRLAARGMR*RA*REFQEPRRLTSGPFPGQP
Consensus

Teleogryllus_occipitalis     ----------------------------------------------------------------------------------------------------------------------------------------------------------
Acheta_domesticus            EFIWHGRSEWPVKVAKPHHTNK*IN-RRGTITSRTEICQEKGK--WRI*SASK*SHESAVKTVVEFCLF*KLPM*CLTVCCV*VVSIRIF--------------------YRKKVK*SE
Gryllus_bimaculatus          EFICHDWSE*PVKVAKPHHTNK*ITIAVG*LPER-KYTREKGNGGFKVRANDHTN*RS--KLLSSFTAFT*F----L*EYFTENNSIL*M*KQVSET*NAL*MEILWELKKYRRGV*NYT
Gryllus_longicercus          EFICHDWSE*PVKVAKPHHTNK*ITIAVG*LPER-KCTRERGNGGFKVRASDHTNRRS--KLLSSFASF---------ETYV*YSALLLL---------------------RNFYKNIL
Consensus

Teleogryllus_occipitalis     ----------------------------------------------------------------------------------------------------------------------------------------------------------
Acheta_domesticus            NK--------NKILGHKKLYKWKFPD*QRNRGGASST*MH-------------------------------------------------------------YIKKQNHAKIIVFQMMEM
Gryllus_bimaculatus          Q*IGPSLEC**GL*RERNVYEVHELNDQ-------------------VLN*TFLGKHVRFRQPNTY-----------------------NDRTYGNT*A*YLKIKPHEN-RVSNVVNV
Gryllus_longicercus          QKTIQYC*CENKFLKHKMLYKRKFLGNSKN-SGVKSTTIHNKWWPIFGVLIGIIEGKKY-LRSPRT**SSSELDFPRKACQVSAA*YVQ*SNLRKYISVIS---KIKPHESHRVSNVVNV
Consensus

Teleogryllus_occipitalis     ----------------------------------------------------------------------------------------------------------------------------------------------------------
Acheta_domesticus            LLLR-KRKI----HL-----------KTRCIRCPSSL*S-VLKAKQSVVG*LTNRTKLHIVPDPHHLGIYFLNHSLE--SV*LSHSSTVLAH*ITPS*ICMQQQKKKF-YFFVENEILHQ
Gryllus_bimaculatus          LTEKVENKTENTWHLLFLLIVICAEGKAEGIWLTQKLH*ISMF*APTIL*NFFKR-------------------LSGWNV*ISL*STVPVH*TTAS*ICM**QRRNGIYLRTEKERLEK
Gryllus_longicercus          LTEKWKTKL----------------KTRGVCCFS*LQS-ALKAKPRVFG*LTSRTKFRCSEPPRSCRI-FLNHSLDGMSE*ACDLRSLCIEPLRVEFVCNSKEEMEFIYVQKRKDW--K
Consensus

Teleogryllus_occipitalis     ----------------------------------------------------------------------------------------------------------------------------------------------------------
Acheta_domesticus            TLQTHF------------KNVFLDLH-------VNVSPRIVN*P*VISPMYKRVTVGNNILINVTKCE*CTHLTLFWF*RQIGKQLYFQK------IIFKLLFIIYQTIIDHNCE**LL
Gryllus_bimaculatus          TQ*THLKTLTSQVNDTRISKNRELTLN---YQYRAQTSG---SRA**TYKCYQ------------M*IIHTFDSYFGFEHKSGKYLVSRQLSSNF*LLFTRTFHIL*VITTK-----YT
Gryllus_longicercus          KHNKHL------------*KR*LRFTG**YSY-IQES*---TDPE*SVPCTNEG**G---IINL*M*IIHTFDSCFGFEHKPGKYLVSRQLSSNC*LLFTRTFYIL*VITT*-----YT
Consensus

Teleogryllus_occipitalis     ----------------------------------------------------------------------------------------------------------------------------------------------------------
Acheta_domesticus            NKLCFLLYKKYNFALRNNLLPKVQWAS*PQP*NMSVMQ*ILYFRASILLL*RQSNSLQTLTCALK----IMIPHH*QTSVPRRPS*IFPPTSAPSRSSKIDGLCLPRFWSFLSKHDPNIK
Gryllus_bimaculatus          VSLSF*K-KKYNFSLRNDLLPEVQWACLPPLAYNMPVIYYILYFRASILLLLW*LNSLQNLTCARKNYDSTLLTHVYCPGVPCKYYR*LVRTAVLRTQAKYTNYASHDFSYFLS*HDQNVK
Gryllus_longicercus          LSLSF*I-KKYNFSLRNDLLPEGQWTCLPQAYNMSVI*YILYFRASILLLLW*LNSLQNLTCARKNYDSTLL----------------------TQASLVNITADLCGPLFSALRQNTR
Consensus

Teleogryllus_occipitalis     ----------------------------------------------------------------------------------------------------------------------------------------------------------
Acheta_domesticus            R-----------NSSKDRKFILKGKNA-----FFLISFLCKLPIERCCLFIIALRQPPKSFLSIARNLIGYSEPPLQIGD-------------P------YLSFRATFPHPSKENGGI
Gryllus_bimaculatus          T---NSFKRSQIHLTEKTYSLFHSYAN--YRLNL-----------AACL---*LR*DNRKRV-----------FFQLRDI**VISSRFCKLATP--------IFLATFPHPSKENGGS
Gryllus_longicercus          IMPPTIFSLSSLNTTRM*KRTLQKIANSSYRKNFFLISFLCKLPIEPCCLFIIALRQPQKSFLSIARYLIGYFEPLLQIGD-----AYLFSHFSAPEQRKWRDVVFLRAHPRRQRKH*HN
Consensus

Teleogryllus_occipitalis     ----------------------------------------------------------------------------------------------------------------------------------------------------------
Acheta_domesticus            WFFCGPARGARENIDTIPGRPAITLPVIPLFLALAPAPRRWRR*R--------WPRRWRWRRRWRRRWRRRWRRRWRRRWRRRWRRRWRRRWRRRWRRRWRRRWRRRWRRRWRRRWRRRW
Gryllus_bimaculatus          GFCPGP-----------PAAPEKTLT------QSPPAPQL----RSLLFHCFSHCRPLR-------------------------------------------------------------
Gryllus_longicercus          ----------------PRRPAITLPVIPLFLALPRAPRR----R--------WCRPWR------------------------------------------------------------
Consensus

Teleogryllus_occipitalis     ---------------------------------------------------------------------------------RGGGGGGGGG------------------------
Acheta_domesticus            RRRWRRRWRRRWRRRWRRRWRRRWRRRWRRRWRRRWRRRWRRRWRRRWRRRWRRRWRRRWRRRWRRRVETAVETAVEAAVGDGGGGGGGGGGGGGGGGGGGGGGG----GGGGDGE
Gryllus_bimaculatus          ---------------------------------------------------------------------------------GGGGAGRGGGRSSGGSGGGGGGSRG----GGGGDGE
Gryllus_longicercus          ---------------------------------------------------------------------------------QWTPAGGGGGGGSRG--GGGNGRRGDDGAGGGGDGE
Consensus                                                                                                              * ***

Teleogryllus_occipitalis     ----------------------------------------------------------------------------------------------------------------------------------------------------------
Acheta_domesticus            VK*CVGGRR-SRRRVLRRSGSVDGVRRL*TKNKAVVRDRH*ILHDVAKEAVNPPVKKWISRRGW*PSDRYR-----SGRGTN-GGVNGEA-----------WRKPNKKGQGSDSVRQRSD
Gryllus_bimaculatus          IRLCVGGRRGSWRQEQFR*SSEGGVRRL--------------------------------------DK*Q--------GCGIGTVTGSSLNF--------------------V*RCDGG
Gryllus_longicercus          IRLCVGGRRGSWGQEQFRWSSEGGVRRL-------------------------------------DK*QEWNRLCNRDC-----DGSSLNFARRWNVLLWRGRSDVRDVERVERVGTG
Consensus

Teleogryllus_occipitalis     ----------------------------------------------------------------------------------------------------------------------------------------------------------
Acheta_domesticus            WVTCVWQEDES*AGKETVEENVRRILGTGQGDDGETTRRWLPSHRCDDGVVASRVVGIEIGGGAEMLELERQ*GGGAAVIPGV---------------SIMDVISPVSIVRVFSPGGGG
Gryllus_bimaculatus          GTSCYEEADQTWERKTAVPTGRR-------G-KGRTSK---------DGV--ERVVQV-VGGGRRKWQR----GGGAAVKRRMTKGRRDA*ELVRRSRQAVAQGQGPWAMARVG-PGGGG
Gryllus_longicercus          *CQC*GE--------V*VAVGRR-------GQRGECA-------------------V-MATGSKGWYRL*EGGGGAAVK--------------RSRQAVTPWQGPWAMARV------*
Consensus

Teleogryllus_occipitalis     ----------------------------------------------------------------------------------------------------------------------------------------------------------
Acheta_domesticus            DLFPAG-----WRDPAATYR*-------LAGFSPQ-PGS-----------------------SHQRGTSPA---------TVSSRSVS----------RHV------------------
Gryllus_bimaculatus          SVEAMGQLRSKKRVDTLEGTA-R*QRTVQGGLTPSMQGTRGLTEGAREEVIRLIADSGCVNC*AHCGGLPPGRRESACCLLAVAGSLVTQNGSRPNVDLRHVRHDAINLE*F*VFAIS*G
Gryllus_longicercus          DLVEEGQW---KRDDSAESKAERLAFCVRDLLIDV--GSEGLTEAAREEVI*WIADSGYVSC*AHCGGLPPPPWTARI*MLPVGGRRIS----------RYT-------EWFKA*CGPQN
Consensus

Teleogryllus_occipitalis     ----------------------------------------------------------------------------------------------------------------------------------------------------------
Acheta_domesticus            ---------------------------WYCR----IY----------------*CKCV--CKGVSIG-------------------QGRGRKVHVRSVTRFLGYCSG--VH------
Gryllus_bimaculatus          RPGYIYTNINCPAFVF*ILNSY*NG----YSRTILSVY-----------------------C*LYFIF---NRVKKESAQ-RRSDAEEEGLTRR---AASTALETGCASSYLHRLLLTM
Gryllus_longicercus          TCGMMQ*TVSDFRFL-----QYHRGDQATFIRTL*TVVHLCFKFSTVIKAVIQEPFCRCTVNCILYLIA*KRNRLRNGAMRWKRGNEDEAGRGRR---RP-----------LH------
Consensus

Teleogryllus_occipitalis     ----------------------------------------------------------------------------------------------------------------------------------------------------------
Acheta_domesticus            ---MDWKNH------------------RKLPM-RAAGCQNSPL---------------------------------------ESVGIR------------------------------
Gryllus_bimaculatus          LITLPW-----PRRSQVFVVICFSLGGRRFPLFSTRGALGGPFADFKKEGGXXXXXXXXXXXXXXXXXXXXXXXXXXXXXXXXGVGW-------------GSGVATNPQAPYAGVLCS
Gryllus_longicercus          -----WKRAALPATSIAFY*LC*LL----FP--GPAGAKSLW*SVFLWEGGGFLSFPPGAPVVGHSRTSKKREDSSAVTGGGEDGRGWRMARAVAGAA*TVAGTAAAAGTAAGTEGAAVG
Consensus

Teleogryllus_occipitalis     ---------------------------------------------------------------------
Acheta_domesticus            -------------PSRQAG--------------------------------VENPAAVGTRTQDLP------
Gryllus_bimaculatus          AIVSTLFISC*LLSERSCGRKPCLPPLPAMLY-----DLTFVVQFS-----L*RVSKKGEKSLCVP--CV*V
Gryllus_longicercus          [illegible]GFRGCA-QPSGPLPRRFMFRHCFQSIHFVLTFVRAILWTEALSPASARHAS*PNICSSI
Consensus
```

**Tocc.10G0002420.1_exon3**

```
Teleogryllus_occipitalis                 ------------------------------------------------------------------------------------------------------------------------
Acheta_domesticus                        DLKHSTTFMFRSMNKILFKITLKY*SFRKLPTIHPDAFA-TSVQKCAP*VRLSSELNRILRAGFHIECNPTTLICNEFSERCHQRNVFRIHLYIRAHVV*RTTLFER--LHSG*D*TPNR
Gryllus_bimaculatus                      --------------------SAKLCAISK-SELRTKSYALCRISHCAR--SKNSHSQRIFGEVSSAQCI*ATFI------------------HLRARSIT*LPCLKDLILH---------
Gryllus_longicercus                      -----KSYALQ---------DFTLFAF*KL------SFA-TNF-------RRGVISAMYLGYIYISAC---TFY-----------NI--------------TTLFER--LNSA----PNR
Consensus

Teleogryllus_occipitalis                 ------------------------------------------------------------------------------------------------------------------------
Acheta_domesticus                        IVIV*QPQRVTNCLPLSSTKIIA*IVITR*NHLNNY---SNR-------RNAHCNI--FTRSKMIIKTK---------TRSGVLIAQRYQLINRATSRSFGLGRNAIGCSGNEITCLARI
Gryllus_bimaculatus                      -LIGLPPQWNHTTTP--TRHQLFTIILN*NNPLNHHHEVSSKQLF----QQAPCLLQHFHEQQ*LLKRKLDPAC**HNATS**TERHPYRLV-------WAVMRLA--------------
Gryllus_longicercus                      IAATMESYVNPNASP----------IVYHYPQLK*SHKSSARGIF*TTIPTGAILAATFSRTAMIIKTK---------TRSSVLIAQRY*LINRATSISFGLGRNEIGCSGNEITSLARI
Consensus

Teleogryllus_occipitalis                 ------------------------------------------------------------------------------------------------------------------------
Acheta_domesticus                        SRAAPIKRVAHFQGSRNSFGMAGP--------SGR*K------SQNRIT-----QTNK-*TVAER*LPERKFARKKGNGGFKVRASDHTNRRS--KLLSNFAYFENY---LCSALQYVAF
Gryllus_bimaculatus                      ARGMR*RA*REFQEPRRL--TSGP-------FPGQPEFICHDWSE*PVKVAKPHHTNK*ITIAVG*LPERKYTREKGNGGFKVRANDHTN*RS--KLLSSFT----------------AF
Gryllus_longicercus                      SRAAPLN---EWPISR----AAGIHLS*LVRVTGESRKTA---SHKQIN----NHRGR-IT--------SRTEMYQGKGKWRI*SASKRSH*STVKAVIKFCFF*NLRII*CLT----AF
Consensus

Teleogryllus_occipitalis                 ------------------------------------------------------------------------------------------------------------------------
Acheta_domesticus                        R*FL*EYFTEKK----LNKVKIKTRS-------*DIKSFINGNSLTDRGI-------------------------------------GEG---RPARKCT--ISK----NKTTRKSSSFK
Gryllus_bimaculatus                      T*FL*EYFTENNSIL*M*KQVSET*NAL*MEILWELKKYRRGV*NYTQ*I-------------------------------------G------PSLEC**GL*R----ERNVYEVHELN
Gryllus_longicercus                      T*FL*EYFTENNSILLM*KQVSET*NAL*TEIPW**QKFRREVYNYTQ*MVAHLWSANRDYRGKEISTKSTNLMIKF*TRLS*ESMSGFGSLIRTMIELT-*IHKRDI*N*TTRKSSRFK
Consensus

Teleogryllus_occipitalis                 ------------------------------------------------------------------------------------------------------------------------
Acheta_domesticus                        *WKCFY*GSGKYI*KHVVF-------------------AVRPHCN-----------------------------LC--*RQSRVWLDDSQIALNCI-LFLTRTILAYIF-*ITLWRVSD*
Gryllus_bimaculatus                      DQVL----N*TFLGKHVRFRQPNTYNDRTYGNT*A*YLKIKPHENRVSNVVNVLTEKVENKTENTWHLLFLLIVICAEGKAEGIWLTQK---LH*ISMF*APTIL*NFFKRLSGWNV--*
Gryllus_longicercus                      CCKCSY-------------------------------------------------*EVENKIENTWCLLFLLIAICAEGKAEGIWLTHK---SH*ISLF*APTIL*NFFKPLSGWNV--*
Consensus

Teleogryllus_occipitalis                 ------------------------------------------------------------------------------------------------------------------------
Acheta_domesticus                        VTHLRSLPIESLRAEFVC-NSKRRNFIFLWKMKYYIK-----HCKHI--------------LKMYF*IYMLMCLQES*TDPESSVQCINE*Q*VIIYL*MLPNVSNAHI*LFFGFEGKSE
Gryllus_bimaculatus                      ISL*STVPVH*TTAS*ICM**QRRNGIYLRTEKERLEKTQ*THLKTLTSQVNDTRISKNRELTLNYQYRA----QTSGSRA**TYKC---YQM*IIHT----------FDSYFGFEHKSG
Gryllus_longicercus                      ISL*STVLVH*TTAS*ICMQ*QRRNGIYLRTEKERLEKTQ*TPLKTLT--------SIHRLMILVYPRIV------N*P*MISTV-----HKRGVVGHNKLINVNNTHI*LLFWF*T*TG
Consensus

Teleogryllus_occipitalis                 ------------------------------------------------------------------------------------------------------------------------
Acheta_domesticus                        NNFISRKLFSNYYLLFIRPLLII------IVSND---Y*INF------VSFCIKNTIL-H*ETICFPKC-NGQVDH----------------NRRTCLSCNRSCIFVHQSYCCDGNQTVC
Gryllus_bimaculatus                      KYLVSRQLSSNF*LLFTRTFHIL-----*VITTK---YTVSLSF*KKKYNFSLRNDLLPEVQWACLPLAYNMPVIYYILYFRASILLLLW*LNSLQNLTCAR------KNY----DSTLL
Gryllus_longicercus                      K-------VPCFQTIIIKLLIIIYQNILYIMSNNHLIYFVSF-LLDKKV*FFLTKRFASRRAMDMFTSS-V*HVCH-IIY------LIFSCINLTTVMVIKQ---FA-KSYL--------
Consensus

Teleogryllus_occipitalis                 ------------------------------------------------------------------------------------------------------------------------
Acheta_domesticus                        KHSPAP*KL*FHTTNKRLCPGGPRKYFRRRVRP---RARAK*TDYASHGFG-LSSLNTTRI*KGTLQKIANSS*REKTL--FSLFHSYANYRLNVAACL*LR*DNRQRVFFQLREI**VI
Gryllus_bimaculatus                      THV--------------YCPGVPCKYYR*LVRTAVLRTQAKYTNYASHDFSYFLS*HDQNVKTNSFKR-SQIHLTEKT---YSLFHSYANYRLNLAACL*LR*DNRKRVFFQLRDI**VI
Gryllus_longicercus                      ----RA*KL*FHSTNT----GVPCKYYR*LVRTAVLRAQAKYTDYASHDF*SFLS*HDQNVKTNSSKD-RKFILQKKLLPYFILMQITD*TLLPVYNCV---KTTAKEFSFN--------
Consensus

Teleogryllus_occipitalis                 ------------------------------------------------------------------------------------------------------------------------
Acheta_domesticus                        LSHLCKLATRIYL----------FEPLFRTRAKKMAGSGFSVGPPVAPEKTLTQSPA----APQLRSLLFHCFS--------------------------HWRPLR--------------
Gryllus_bimaculatus                      SSRFCKLATPIFL------------ATFPHPSKENGGSGFCPGPPAAPEKTLTQSPP----APQLRSLLFHCFS--------------------------HCRPLR--------------
Gryllus_longicercus                      ----CEIFNRLF*AASANWRRLSF*PLFRTRAKKMAGCGFSPGPPAAPEKTLTQSPPPRNYAP--------CYSIVSGTAARSAAEVVQAVEAVDACRWRWWRK*RRWRQWKAWRRWSRW
Consensus

Teleogryllus_occipitalis                 ----------GGGGGE-GGRGSGGDGGGGG--GGGGGSG-------------------------------------------------------------------------GGG-----
Acheta_domesticus                        ----------GGGGGE-GGRGGGGGDGGGDGGGDGGGDG------GGDGGG---------DGGGDGG----------------------GDGGGDGGGDGGG---------DGGG-----
Gryllus_bimaculatus                      ---------GGGGAGRGGGRSSGGSGGGGGSRGGGGGDGEIRLCVGGRRGSWRQEQ---FR*SSEGGVRRL----------------DK*QGCGIGTVTGSSLNFV*R--CDGGGTSCYE
Gryllus_longicercus                      RRRWRDSTMCGREEGELGART---------------------VPVEQRRWGAETRQITGME*VV**GL*RIVIEFCEEVERPAMERQIRCERRGKGRTSRDGVVSVLR*GVSGSG-----
Consensus                                          *   *  * *                                                                                            * *

Teleogryllus_occipitalis                 ----------------------EKDG----------------------------------------------------------------------------------------------
Acheta_domesticus                        ------------DGGGDGGGDGGGDG-----GGDGGGDGGGDGGGDGGGDGGGDGGGDGGGDGGGDGGGDGGGDGGGDGGGDGGGDGGGDGGGDGGGDGGGDGGWRRRWRRRWRRRWETA
Gryllus_bimaculatus                      EADQTWERKTAVPTGRRGKGRTSKDGVERVVQVVGGG----------------------------------------------------------------------------RRKWQRG
Gryllus_longicercus                      ------------EAGAAGRMCCDGDGVERVVQVVGG------------------------------------------------------------------------------RGWSSG
Consensus                                                        **

Teleogryllus_occipitalis                 -----------------------------------------GGGGG--------------------------------------------------------------------------
Acheta_domesticus                        VEAAVEAAVEAAVEAAVEAAVEAAEAMERLNDVWVA-----GGVGG-ECCAGRAV*MGCGDY-------RRKTRQ----W*GIVTEFCMTLRRRQ-----*TPL*RSGSVDVDGSRVTDT
Gryllus_bimaculatus                      GGAAVKRRMTKGRRDA*ELVRRSRQAVAQGQGPWAMARVGPGGGGSVE---------AMGQL-------RSKKR---------VDTLEGTAR*QRTVQGGLTP-------SMQGTRGLTE
Gryllus_longicercus                      EEEQTSSNAMAG-------------ALGDGQS------VGLGGGGSVEAR*QR*VQGGAPRVLRS*LTHRCRERGSDGSSAGRGDLMDC*FRVRQL----LSPLRRSPPAPLDGENLN--
Consensus                                                                         ** *

Teleogryllus_occipitalis                 ------------------------------------------------------------------------------------------------------------------------
Acheta_domesticus                        EAVEE--QMVV*TG------RRGG-SRTRRGRVVIV*DKEVIG*LVCGRKMNRRQERRRWRK---------MFAGSWGQGRETTEKRQG---------------------GGYRHTVATT
Gryllus_bimaculatus                      GAREEVIRLIADSGCVNC*AHCGGLPPGRRESACCLL--AVAGSLVTQNGSRPNVDLRHVRHDAINLE*F*VFAIS*G--------RPGYIYTNI-NCPAFVF*ILNSY*NGYSRTILSV
Gryllus_longicercus                      ---------------VACW-RSQDLSLHR----------MVQGLMWTS---------EHVRHDAINRE*F*VFAIS*G--------RPGYIYTNIMNCRAFVF*IFNSY*SGYSRTILSV
Consensus

Teleogryllus_occipitalis                 ------------------------------------------------------------------------------------------------------------------------
Acheta_domesticus                        VWWL-VEW*GLRLEEGRRCWSWRGN---EGVERR*FRACQLWTSLVP*A*YVFFHREVAVIFSPLDGEIRRRP------TVD*RGFPLNQVAPTSGAPLQPL*APDRSPGMCGTVEYISV
Gryllus_bimaculatus                      YC*LYFIFNRVKKESAQR----RSDAEEEGLTRRAAS-----TALETGCASSYLHRLLLTMLITLPWPRRSQVFVVICFSLGGRRFPLFSTRGALGGPFA--------------------
Gryllus_longicercus                      YC*LYFIFNRVKKESAQK----RSDEVEEGQRR*GWTRKAASTALETGCASSYLHRLLLTMLITLPWPRRSQVFVVICFSLGGRRFPLFSTRGARGGPFA--------------------
Consensus

Teleogryllus_occipitalis                 -------------EGG--------------------------------------------------------------------------------------------------------
Acheta_domesticus                        NVCVRG*V*DKEEEGK-------------------FTCAP*PVS*GTAPAFTWIGKTTENCR*GQPAARILL*R--ASEFDQADKLESK-------------------------------
Gryllus_bimaculatus                      ---------DFKKEGGXXXXXXXXXXXXXXXXXXXXXXXXXXXXXXXXXGVGWGSGVATNPQ--APYAGVLCSAIVSTLFISC*LLSERSCG----------------------------
Gryllus_longicercus                      ---------DFKKEGRQLR--------------------------------------SDGGRRGRP------------------RVEDGESGGGGGVDGGRDGSGGGDGGGDGGSGRGRG
Consensus                                             **

Teleogryllus_occipitalis                 -----------------------------------------------------------------
Acheta_domesticus                        --------ILP-------PSGLEPRTF--------------------------------------
Gryllus_bimaculatus                      -----RKPCLPP-----LPAMLYDLTFVVQF--------SL*RVSKKGEKSLCVPCV*V------
Gryllus_longicercus                      GGKGLRPTLRPPTQAFYVPPLFPVYSFRVNFCRPSDPVDGSL--VSR-----LCPPCFMT*HL*FN
Consensus
```

**Tocc.10G0002420.1_exon4**

```
Teleogryllus_occipitalis      -------------------------------------------RKR---------------------------------------------------------------------
Acheta_domesticus             I*NIPPHSCFVL*IKFYSKSR*NIKASENSLLSILMRLRPVCKNARRKYG*APN*IAYCAQDFTSSAIQQLSFATNFRRGVINATYLG--YIYISARTLYNELPCLKDFILDEIKHLIG-
Gryllus_bimaculatus           ---------------------------------------ECKIVRYKQV*APD*IICPVQDFTLCAF*KLSFATNFRRGVISAMYLG--YIYTSACTFYN-ITTLFERL---------
Gryllus_longicercus           --------------------------------------------------LNHMLCRISH---CLRSKNSHLQRIFGEVSSAQCI*ATFIYLRARSIT*-LPCLKDLIL----HLIGL
Consensus

Teleogryllus_occipitalis      ---------------------------------------------------------------------------------------------------------MR----------
Acheta_domesticus             -----------------L*SYDNPNASPIVCRYPQLK*SHKSSSRGRII*TTIPTGATRTATFSREAK*LLKRKPDPAC**HNATS**TERHPDRLVWAAMRLAARGMR*RV*REFQGP
Gryllus_bimaculatus           ------NSAPNR--IAATMESYDNPNASPIVYHYPQLK*SPKSSSRG-IF*TTI----------------------PTGAMFAATFSRTA--------MIIKTKTRSSVLIAQR-Y*LI
Gryllus_longicercus           QPQWNHTSTPTRHQLFTIILN*NNP-----INHQHEV------SSKQ-LFQQAPY-----SLQHFHEQQ*LLKRKLDPAC**HNATS**TERHPYRLVWAVMRLAARGMR*RA*REFQEP
Consensus

Teleogryllus_occipitalis      -----------------------------------------------------------------------------------------------------------------------
Acheta_domesticus             R------------------------------PLNEWPISRAAGIHLAWPVRVAGESRKTASHKQINK-----PSRNDNFPNGNLPG----------KREMEDLKCEQVITRIGGQNCCR
Gryllus_bimaculatus           NRATSISFGLGRNEIGCSGNEITSLARISRAAPLNEWPISRAAGIHLS*LVRVTGESRKTASHKQINNHRGRITSRTEIYE-GK--GKWRI*SA---SKRPHKLTVKAVI----------
Gryllus_longicercus           RRLTSGPFP-GQPEFICHD-------------WSE*PVKVAKPHH--------------TNK*ITIAVG*LPERKCTRERGN--GGFKVRASDHTNRRSKLLSSFASF----------
Consensus

Teleogryllus_occipitalis      -----------------------------------------------------------------------------------------------------------------------
Acheta_domesticus             ILLILKTTYVVPYSMLRLGSFYKNILQKKS*I----------------------------------------------K*K*KQDPRT-*KAL*MEIP*LTEE*GRG--VQHVN----
Gryllus_bimaculatus           ------KFYCFY--LIFIRIFHR----KQFNIVNVKTSF*NIKCFINGNSLGIEKIQAW-SLELYTINWSIFGVLIGIIEGKKCLRSPRT**SSFELDVSRKTCQVSAA*YVQ*SNLRKY
Gryllus_longicercus           ------ETYV*YSALLLLRNFYKNILQKTIQYC*CENKFLKHKMLYKRKFLGNSKNSGVKSTTIHNKWWPIFGVLIGIIEGKKYLRSPRT**SSSELDFPRKACQVSAA*YVQ*SNLRKY
Consensus

Teleogryllus_occipitalis      -----------------------------------------------------------------------------------------------------------------------
Acheta_domesticus             -ALYQKTK---------------------PRENHRLSNDGNAFTEEAENTSENTLYSLSVLIVICAEGKAECGWMTHKSH*IAYCS*PAPSWHIFFKSLSGECLTKSLIYGP-------
Gryllus_bimaculatus           ISVISKNKTTRKSCFKCCKCSY*ESGKQN*KHVASTVSPDCNLR*RQSRGDL-------------VDSKAALNFDVLSPHDLV---------EFF*TTLWMECLNKPVIYGP-------
Gryllus_longicercus           ISVISKIK---------------------PHESHRVSNVVNVLTEKWKTKL--------------KTRGVCCFS*LQS---------------------ALKAKPRVFG*LTSRTKF
Consensus

Teleogryllus_occipitalis      -----------------------------------------------------------------------------------------------------------------------
Acheta_domesticus             -CP---------LNH---------SELN--------------------LYATAKEEI-LFFCGK*NITSNIANTF*KCIFRFTC*CVSKNRELTLSHQSNV*TSDSR**YTYKCYQ---
Gryllus_bimaculatus           -CA---------LNH---------CELN--------------------LYVIAKKKWNLFTYRKGKIGKNTINTF------------KNVNFTG**YSYIQES*TDPELSVPCTNEW*
Gryllus_longicercus           RCSEPPRSCRIFLNHSLDGMSE*ACDLRSLCIEPLRVEFVCNSKEEMEFIYVQKRKDWKKHNKHL*K---------------------R*LRFTG**YSYIQES*TDPE*SVPCTNEG*
Consensus

Teleogryllus_occipitalis      -----------------------------------------------------------------------------------------------------------------------
Acheta_domesticus             -------M---------*VMHTFDSFLVLKANRKTTLFPENYFQIII-YYLSDH----Y*S*L*VMTIE*TLFPSV----*KIQFCIKK-QFASQSAM---GKLTTTVEHVCHAIDLVFS
Gryllus_bimaculatus           *GIINL*MLSNVNNTHI*LLFWF*T*IG-----KVPCFQAIIFKFLI---------IIYQNISYIMSNNN*IYCVSFLLEKKV*FFVTK-RFASRSAM---GMFTSSV*HACHILYLVFS
Gryllus_longicercus           *GIINL*M---------*IIHTFDSCFG---------FEHKPGKYLVSRQLSSNC*LLFTRTFYIL*VITT*YTLS------LSF*IKKYNFSLRNDLLPEGQWT--------------
Consensus

Teleogryllus_occipitalis      -----------------------------------------------------------------------------------------------------------------------
Acheta_domesticus             CI----NLIAVTAIKQFANTHLRP--------------------------KNYDSTPLTNVCAQAALVNISADECAL---ALEQNRRIMPPTVLVFPL*TRPEYKKELFKRSQIHLKGK
Gryllus_bimaculatus           CI----NFTAVMVIKQFAKSYLRP*KL*FHSTNTRLLPRRPL*ILPLTCAHR----------------------CSPHSGKIHE---LCFPRFFVLPLLTRPECKNELFQKIANSSYRK
Gryllus_longicercus           CLPQAYNMSVI*YILYFRASIL----LLLW*LNS---------LQNLTCARKNYDSTLLT----QASLVNITADLCGPLFSALRQNTRIMPPTIFSLSSLNTTRM*KRTLQKIANSSYRK
Consensus

Teleogryllus_occipitalis      ---------------------------------------------------------------------------------------WRQ-----------------------------
Acheta_domesticus             KRFFPYFILMQITD*TLLPVYNC------VKTTAKEF----SFNCEKFNRLF*ATSANWRPVSIFSSHFSAPEQRKWRDLVFLWARPWRQRKH*HN-PRPPRNYAPCYSIVSRTGARSAA
Gryllus_bimaculatus           NLFLISFLCK-------LPIEPCCLFIIALRQPQKSFLSIARYLIGYFEPLLQIGDA------YLFSHFSAPEQRKWR-IWFLSGPTRGARENIDTIPARPAITLPVIPLF---------
Gryllus_longicercus           NFFLISFLCK-------LPIEPCCLFIIALRQPQKSFLSIARYLIGYFEPLLQIGDA------YLFSHFSAPEQRKWRDVVFLRAHPRRQRKH*HN-PRRPAITLPVIPLF---------
Consensus

Teleogryllus_occipitalis      -----------------------------------------------------------------------------------------------------------------------
Acheta_domesticus             VEAVKVAAEVAVETAVETAVETAVETAVETAVETAVETAVETAVETAVETAVETAVETAVETAVETAVETAVETAVETAVETAVETAVETAVETAVETAVETAVETAVETAVETAVETAV
Gryllus_bimaculatus           ---------LALPPAP-------------------------------------------------------------------------------------------------------
Gryllus_longicercus           ---------LALPRAP-------------------------------------------------------------------------------------------------------
Consensus

Teleogryllus_occipitalis      -----------------------------------GRR------RRRW-RRW-RRW------RRWRRRRKWR---------------------------------------------------
Acheta_domesticus             ETAVETAVETAVETAVETAVETAVETAGGDGGGDGGGGGGGRRRWRRRWRRRWRRRWRRRWR-----RRWRRRRRRWRG*M----MCGWQE-ESEASAAQVGQCR--WGAET-----------ID
Gryllus_bimaculatus           --------------------------------------------RRRWCRPW-RR*KQWGQWRWWRK*RRWRRRWRDSTMCGREEGELEARTVPVKQRR--WGAET------------
Gryllus_longicercus           --------------------------------------------RRRWCRPW-RQWTPAGGGGGGSRGGGGNGRRGDDGAGGGGDGEIR---LCVGGRRGSWGQEQFRWSSEGGVRRLD
Consensus                                                                   **** * * *

Teleogryllus_occipitalis      -----------------------------------------------------------------------------------------------------------------------
Acheta_domesticus             E-KQGSG------KGSSLNFA*RCEGGSKPPCEEVDQ*TWMVAE*QIPKR*RNKWWCKRGGVEEAEQEGAG***CETKK*LGDLCVAGR*IVGRKGD------------GGGKCSQDLG
Gryllus_bimaculatus           --RQITGLWNRDCDGIVIEFCMTL-----------RWRWNV----LL*RGRSD------------VGEENCGADGEAWKG----SNE*GRGRKGGTGCRGRAAEMATRGRSSGEEEDD
Gryllus_longicercus           K*QEWNRLCNRDCDGSSLNFAR---------------RWNV----LLWRGRSDV----RDVERVERVGTG*CQC*GEV---------*VAVGRRGQ---RGECAVMAT--GSKGWYRL*
Consensus

Teleogryllus_occipitalis      ----RRRR*--------------------------------------------------------------------------------------------------------------
Acheta_domesticus             DRAGRRRRRNDKAVVTVTPLRRRCGG----------*SSGRD*DWRR------------------GGDAGAGEAMR-----------GWSG------------GDSGRVNYGRH*SRK
Gryllus_bimaculatus           EGAKRCLRTGAEEQTSSSARAGALG-----------DGQSRTWRRRVSGSDGSITFEEAG*HIGRDSEMTADSPR-----RTYSIDAGNEGSDGRSTGR-----GDS--IDC*FRVRQL
Gryllus_longicercus           EGGG-----GAAVKRSRQAVTPWQGPWAMARV*DLVEEGQ---WK-------------------RDDSAESKAERLAFCVRDLLIDVGSEGLTEAAREEVI*WIADSGYVSC*AHCGGL
Consensus

Teleogryllus_occipitalis      -----------------------------------------------------------------------------------------------------------------------
Acheta_domesticus             HSTCFFTGRWR*SFPRWMARSGGDLPLTSGVFPSTR*LPPAGHLSSH---------------------------------------------C-------KLPIGLP---------A
Gryllus_bimaculatus           LSPL----RW---SPPWTAR-------------ISMLPVGGRRISRYTEWFKA*CGPQT--------------------RAS*CNKP*VILGVC--------NIIGETRLHLYKH*LS
Gryllus_longicercus           --P----------PPPWTAR-------------I*MLPVGGRRISRYTEWFKA*CGPQNTCGMMQ*TVSDFRFLQYHRGDQATFIRTL*TVVHLCFKFSTVIKAVIQEP---------F
Consensus

Teleogryllus_occipitalis      -----------------------------------------------------------------------------------------------------------------------
Acheta_domesticus             CVVL*N-----------------ILV*MC-----V*GGEYRTRKR---KES-----SRALRNPFL--------------------------------------------------------
Gryllus_bimaculatus           CICVLNSQQLLKRLFKNHSVGLLLIVFYI*SREEGIGSETERCGGRGADEEGGVHCTGNGLRFQLP------PSPFINYVNYSSLAPPEP----SLCGNLFFF----------------
Gryllus_longicercus           CRCTVNC-----------------ILYLIA*KRNRLRNGAMRW-KRGNEDEA-----GRGRRRPLHWKRAALPATSIAFY*LC*LLFPGPAGAKSLW*SVFLWEGGGFLSFPPGAPVVGH
Consensus

Teleogryllus_occipitalis      -----------------------------------------------------------------------------------------------------------------------
Acheta_domesticus             -RVLLRR-----------------SHGLEKPQKT------------------------------------ADEGSRLPEFSSRER---RNS----------TKQTSWSRK---SCRRRD
Gryllus_bimaculatus           --------GRAAVSSLFHQGRPWWAIRGLQKRGRTXXXXXXXXXXXXXXXXXXXXXXXXXXXXXXXXXXXGGVGEWGCDQPSGPLRRRVMFRHCFHSIHFVLTFVRAILWTEALSPASARHA
Gryllus_longicercus           SRTSKKREDSSAVTGGGEDGRGWRMARAVAGAA*T-------------VAGTAAAAGTAAGTEGAAVGGVGERGCAQPSGPLRRRFMFRHCFQSIHFVLTFVRAILWTEALSPASARHA
Consensus

Teleogryllus_occipitalis      -----------------------------
Acheta_domesticus             SNPGPS-----------------------
Gryllus_bimaculatus           L*PNICSSIFSLTRFEKRREVPLCPVCLSF
Gryllus_longicercus           S*PNICSSI--------------------
Consensus
```

**Tocc.11G0001970.1**

```
Teleogryllus_occipitalis      AATATTAGTTTAAAAACGCTGATGTTTAAGTGGAATAACTCCTCACAAAAACTCTTTGAATTCACTTGTATGTTCGAGCGTTACATTTTAACAATACAAGTGAAAATATAGTTGAAGTCT
Gryllus_bimaculatus           AATATTAGTTGAA-------TATGTTAATATGAAATAACTTCTCACAAAAATTCTTTGAATCCACTTGTATGTTCGAGCGTTACATTC---------------AGATCTATACAAGGAAA
Gryllus_longicercus           AATATTAGTTGAA-------TATGTTAATATGAAATAACTTCTCACAAAAATTCTTTGAATCCACTTGTATGTTCGAGCGTTACACTC---------------AGATCTATTCAAGGAAA
Consensus                     ********** **        ***** *  ** ******* ********** ********* *********************** *               * ** **    * *

Teleogryllus_occipitalis      TACTAAACCATACTAGATTCGAATATAACGTTTTATTTTTCTTGTTCAAACATATTTAATTCTCATTA--AGTGTAAATCACAACTACATCTTCATACTGTTCTACCGGTGAATTATAAT
Gryllus_bimaculatus           TGTAAAACAATTCGGGCTTCGAAGATTACATTTTTATTTTCTTGTTCCAACATATCTAATTATCATTA--AGAGAGATTCACAAACAAATCTTCACAC--TTCTGCC-GTGAATTATAAT
Gryllus_longicercus           TGTAAAACAATACGGGCTTCGAAGATTACATTTTTATTTTCTTGTTCCAACATATCTCATTATCATTAAGAGAGAGAGAATCACAAACAAATCTTCACAC--TTCTGCC-GTGAATTATAAT
Consensus                     *   **** ** *  * ****** ** ** ****  *********** ******* * *** ******  ** *  * ******  * ******* **  **** ** ***********

Teleogryllus_occipitalis      ATGATTCGATCTATTTTTCAGCTTAACTGATCGCCTTCCCTGAACTTCTATTCATTGACGAAATGCCCCGTCGTGGACACCATGGACACCATCACCACCATGGTGGTCACCATGGTCACC
Gryllus_bimaculatus           GTGATTCGATCTATTTTTCAGCATAACTGATCGTCTTTGCTGAACTTATATTCTTTGACGAAATGCCCCGTCCTGGACATCGTGGACACCATCATCACCATGGCCATCATGGTCACCACC
Gryllus_longicercus           GTGATTCGATCTATTTTTCAGCGTAACTGATCGTCTTTGCTGAACTTATATTCTTTGACGAAATGCCCCGTCCTGGACATCGTGGACACCATCATCACCATGGCCATCATGGTCACCACC
Consensus                      ******************** ********** ***  ******** ***** ***************** ****** * ************ ********   ***   *   ****

Teleogryllus_occipitalis      ACCACCATCACCGGCCGCCGCCGCCACCGCATCACGGTCACCATCACCACCACCGACCAGTGCTCTGCTGTACGATCATTTAGATGACGTCAGAAACTGTGACTTGGGA-----------
Gryllus_bimaculatus           ACCACCATCACCGACCGCCGCCTCCACCGCATCACGGTCACCATCACCACCACCGGCCAGTGCTCTGCTGTACGATCATTTAGATGACGTCAGCAACTGTGACTTGGGAAGTT--AGTGT
Gryllus_longicercus           ACCACCATCACCGACCGCCGCTTCCACCGCATCACGGTCACCATCACCACCACCGGCCAGTGCTCTGCTGTACGATCATTTAGATGACGTCAGCAACTGTGACTTGGGAAGTTTAAGTGT
Consensus                     ************ *******  ******************************** ************************************* ***************

Teleogryllus_occipitalis      --AGAAGAAGTGTTCTTCCTCACTACGAGGATAAAGAATTGCACCGAATACCAAAGAATCAATTCTTCACATTTTTATTTCTCGTATCATCTCTGGAACCGTACTTTTTCGAGTTGCA--
Gryllus_bimaculatus           TCAGAAGATCTACTCATCTTCCCTACGAGGGTAAAGGCCTGCACCAAATACCAAAGAAACGATTCTTCACATTTTTACTGCTCATATCATCTCTGGAACCGCACGTTTCCGAGGTGAATG
Gryllus_longicercus           TCAGAAGATCTACTCATCCTCCCTAAGAGGGTAAAGGCCTGCACCGAATACCAAAGAAACGATTCTTCACATTTTTACTTCTCATATCTTCTCTGGAACCGCACGTTTTCGAGGTGAATG
Consensus                       ******  *  ** ** ** *** **** *****   ****** ************ * **************** * *** **** ************ ** *** **** ** *

Teleogryllus_occipitalis      -CAGATGGCGTTTGACGTGGGAGAGGCAGCCTTAGATATGGCCCTTGCGCGACATGTGTGCGCCGAGAAGATAAACTGTGCAAATTGTATTGAACGCAAAGTGGAGTGTCAAGATCAAAG
Gryllus_bimaculatus           ACAGATGGCGTTTGACGTGGGAGAGGCAGCCTTAGATATGGCGCCTGAGCGACATGTGTGCTCCGAGAAGATAAACTGTGCAAACTGTATTGAACGCTAGGTGGAGTGTCAAGATCAAAG
Gryllus_longicercus           GCAGATGGCGTTTGACGTGGGAGAGGCAGCCTTAGATATGGCGCCTGAGCGACATGTGTGCTCCGAGAAGATAAACTGTGCAAACTGTATTGAACGCTAGGTGGAGTGTCAAGTTCAAAG
Consensus                      ***************************************** * ** ************ ********************* ************ * ************* ******

Teleogryllus_occipitalis      CCACTGGCTCGAGTGTTTATG---------------------------------------------------------------------------------------------------
Gryllus_bimaculatus           CCACTGACTCGAGTGATCAGGATATTGTAGTGCTATTTGCATGTCATTCATATTCCACAATTTTTTTAAACTTAACACAAACACGAAAATATTTATTCCTAAGCTGTGTCTGTCATTATG
Gryllus_longicercus           CCACTGACTCGAGTGATCAGGATATTGTAGTGATATTTGCATGTCATTCATATTCCACAATTCTTCTAAACTTAACACAAACACGAAAATATTTATTCCTAAGCTGTGTCTGTCATTATG
Consensus                     ****** ******** * * *

Teleogryllus_occipitalis      ------------------------------------------------------------------------------------------------------------------------
Gryllus_bimaculatus           TTAAAGATTACATTTTCCATAAAGATTTACGTCTGATATGTAATACTAAATATAATAATAAAACAAAATAATTATAAAAATAATCATATATTATTATCAGCAATGTTTATACTTCTGTGT
Gryllus_longicercus           TTAAAGATTACATTTTCCATAAAGATTTACGTCTGATATGTAATACTAAATATAATAATAAAACAAAATAATTATAAAAATAATCATATATTATTATCGGCAATGTTTATACTTCTGTGT
Consensus

Teleogryllus_occipitalis      --------------------------------TTTATTTTAGTGCTATTC----------------------------------------------------------------------
Gryllus_bimaculatus           TCTATAAATAAAATATTGTATGTTTGAAAAGTTTTATTTTCGCATTATTTATGTGACATTTAGAGCATCTAAGATAAACCCTCATTCATATGTTATCGTAATTTCCTAGATTTGTGTATA
Gryllus_longicercus           TCTATAAATAAAATATTGTATGTTTGAAAAGTTTTATTTTCGCATTATTTATGTGACATTTAGAGCATCTAAGATAAATTCTCATTCATATGTTATCGTAATTTCCTACATTTGTG-ATA
Consensus                                                     ******** *   ****

Teleogryllus_occipitalis      ------------------------------------------------------------------------------------------------------------------------
Gryllus_bimaculatus           GTACAAAACCGAAATCACTTAAATATTTCTACATGAGAAGACTGTACTT-----------------TTTATTAGTTTCGTTGTAATTTATTTGGCAATTTCGATAGCATTATGATTATGT
Gryllus_longicercus           GTACAAACCCGAAATCAATTAAATATTTCTACATGAGAAGTTTCTAATTGAGAAATGTGAAAAAAATGTATTTCGTGCGACAAAATGAAAGCGGCAG--TGAAGAGAATGAGGCTTGTCG
Consensus

Teleogryllus_occipitalis      -------------------------------------------------------------------------------------
Gryllus_bimaculatus           TACGA--CAATTTCAAATATGTAGCATTAGAATCTCATGAAGTTTTGATTAGCTTCCAAATTTAACTCGTTTTGATTGAGTTCA
Gryllus_longicercus           CAAAAGCTAGTCAAGAATAGAGAGATTAGAAAACGAATTAGGGCGAGA---------AAGTCTGTGCCATTTTATAG-------
Consensus
```

**Tocc.11G0001970.1_exon1**

```
Teleogryllus_occipitalis                 ------------------------------------------------------------------------------------------------------------------------
Gryllus_bimaculatus                      -GDNFYFRLALKNLLLSELTGSVLK--------RLNHILSCWSEFIFMLAPV*DNC---------RTMFLY*SSF----YF*KTSKFT*VL*Y*LNMLI*NNFSQKFFESTCMFERYIQI
Gryllus_longicercus                      TGDNLDFRLARKN-LLSELTGSFLND*ITFSLVEVN-LYTC*PQ----CKTIVDQCFCIRVHFTSRRQVNYLSTLILVEYVNMK*LLTKIL*IHLYVRALHSDLFKEM*NNTGFEDYI--
Consensus

Teleogryllus_occipitalis                 ---------------------------------------------------------------MPRRGHHGHHHHHGGHHGHHHHHRPPPPPHHGHHHHHRPVLCCTII*----------
Gryllus_bimaculatus                      YTRKCKTIRASKITFLFSCSNISNYH*ERFTNKSSHFCREL*CDSIYFSA*LIVFAELIFFDEMPRPGHRGHHHHHGHHGHHHHHHRPPPPPHHGHHHHHRPVLCCTII*MTSATVTWEV
Gryllus_longicercus                      --------------FIFLFQHISLSLRERITNKSSHFCREL*CDSIYFSA*LIVFAELIFFDEMPRPGHRGHHHHHGHHGHHHHHHRPPLPPHHGHHHHHRPVLCCTII*MTSATVTWEV
Consensus                                                                                               *** ** ******* *  ******** ********************

Teleogryllus_occipitalis                 ------------------------------------------------------------------------------------------------------------------------
Gryllus_bimaculatus                      SVQKIYSSSLRG*RPAPNTKETILH------IFT---------AHIISGTARFRGE*QMAFDVGEAALDMAPERHVCSEKINCANCIE-R*VECQDQSH*LE*SGYCSAICMSFIFHNF-
Gryllus_longicercus                      *VFR------RSTHPP*EGKG--LHRIPKKRFFTFLLLISSLEPHVFEVNGRWRLTWERQP*IWRLSDMCAPRR*TVQTVLNARWSVKFKATDSSDQD-----------IVVIFACHSYS
Consensus

Teleogryllus_occipitalis                 ------------------------------------------------------------------------------------------------------------------------
Gryllus_bimaculatus                      -FKLN-TNTKIFIPKLCLSLC*RLHFP*RFTSDM*Y*I***NKIIIKIIIYYYQQCLYFCVL*IKYCMFEKFYFRIIYVTFRASK---INPHSYVIV----------IS*ICV*YKTEIT
Gryllus_longicercus                      TILLNLTQTRKYLFLSCV-------------------------------------CHY-----VKDYIFHK-DLRLICNTKYNNKTK*L*K*SYIIIGNVYTSVFYK*NIVCL--KSFIF
Consensus

Teleogryllus_occipitalis                 ------------------------------------------------------------------------------------------
Gryllus_bimaculatus                      *IFL----HEKTVL------FISFVVI----YLAISI----------------AL*LCYDNFKYVALESHEVL-----ISFQI*LVLIEF
Gryllus_longicercus                      ALFM*HLEHLR*ILIHMLS*FPTFVIVQTRNQLNIST*EVSN*EM*KKCISCDKMKAAVKRMRLVAKASQE*RD*KTN*GEKVCAIL*--
Consensus
```

**Tocc.11G0007880.1**

```
Teleogryllus_occipitalis     TCGA------------AAGTAAGGAACATGTCCCTAACACAATTACGGGTCGAATAAAGAATCTCATCCTTGTTTTGAATTCGGTGAAAAAGACTACTCCTACTTTTCTATAAATATAC
Acheta_domesticus            TCGAACGCTTACCGTATAAGTAAGGAACATGCCCACAACATAACAACTAATCGAATAAACAATCTC-CCCTTGTTTTGAATTCGGTTAAAAA--TACCCCTACTTTCCTGTAAATATTC
Consensus                    ****            ************** **  **** **  **   ********* ******  ***************** *****   *** ******** ** ******* *

Teleogryllus_occipitalis     ATCTTTCGACGACGAAAGAGTTGACACAAAATAAATGCCATAAGCGGTTGAAGTGATTTTCGAGAGAAAAAAATTCTAATATCATTCGGAAAATCTTCCTCGAGCTGTTGGAATTTTCTGA
Acheta_domesticus            ATCTTTCGAAAACGAAAGAGATGACAATAAATAACTGCGAAAAGTAGTGGAAGTGATTTTAGAGCAAAAAAATTCTAAAATTACCCAGAAAATCTGACTGGAAATGTTGGAATTTTCTGG
Consensus                    *********  ********* *****  ****** *** * ***  ** *********** ***  ************ ** *  * ********  ** **  ***************

Teleogryllus_occipitalis     TACTGGAGATTTTTACAATATTTCTGTTTCTGTGGGTTCT---------------GTGGTGGTCGTTAAGACAATCACGCGCTAATGGATGTCTCTAATTGCAGCGATCTCTCAGACTG
Acheta_domesticus            TACTGCAGATATTTACAATATTTCTCTTTCTTTCGGTTCTCTCCATCTTTGTCACAGTGGTAGTCGTTAAGACAATCACGCACTAATGGATGTCTCTAATTCCAGCGATCTCTCAGGCTC
Consensus                    ***** **** ************** ***** * ******               ***** ****************** ******************* ************** **

Teleogryllus_occipitalis     GGGGTTCCGCTCCACCGA------------------TCCGCACTTGGCGCGCCCGGGGCCCGTCGCAGCCG-CCTGCGAGTTGGGTCGCCGGCGCGCTCCCAGGGGTCCCGTGCCCCGC
Acheta_domesticus            GGGGTTCCGCTCCACCGAGCGCGCGGTCCTCGGCGCGACCGCACTTCGAGCGCCCGGGGCTCGTCGCAGCCGCCCTGCGAGTTCGGGCGCCGGC-CGCTCCCAGGGGTCCCGTGCCCCGC
Consensus                    ******************                  ******** * *********** *********** ********** ** ******* ************************

Teleogryllus_occipitalis     GGCGGCGGAAGCGGCGACGACCACGCCGACGACGCTGGTGGCGGCGGCAGCGGCGGTGGCAACGGGGCGGTGGCGGTGGGACGGTGGTAGAAGCGGTGGCAGCGGCGACGGCGGCGGGAC
Acheta_domesticus            GGCGGCGGAAGCGGCGACGGCGACGGCGACGCCAACGACGGCGGCGGCGGTGGCAGCGGCGGCGG---------------TCACGGCAGAAGCGG-GGCGGTGGCGGTGGTGGCGAGGT
Consensus                    ****************** * *** ***** *   *  ********* * *** * ***  ***                   ** ******** *** * ****  ** **** *

Teleogryllus_occipitalis     GGTGGTAAAAGCGGCGGTATTGGTGGTGGTGGTGGTGGTGGCGGCAGTA-------GCGGGCCGAGGGTGGCGGTGGCAACGG-------------------GGCGGTGGCGGCAG
Acheta_domesticus            GGTGACGATAGTGGCGGTGGTGGCGAGGCGGGTAGCGGTGGTGACGGGACGATGGCAGTGTGGGGAGGTGGTGGCAGCGGCGGCGGTAGTGGTCTTGATGGCGGTCGGTAGTGGTGGTAG
Consensus                    ****   * ** ******  *** *  *  *** * ******* *** *  *       * ***  *  ****** * ***  ***                   **  **** ** **

Teleogryllus_occipitalis     GACGGTGGTAGAAGCGGTGGTGA-GGTAGTGGTGGTGGCGGCGGTGGCAGTAGCGGGCCGATGGCTGAGGCGGTGGCGTGGCAGAGGCAGTGGTAGTGGTGATGGTGGCGGTCGGTAGAG
Acheta_domesticus            --TGGCCGTGGAGACGGCGGTGGCGGTGACAGCGGCGGCGGTGACAGCAGTAGCGGGGCGGTGGCGGTGGTGGCGAGGTGGTGACGATAGTGGCGGTGGTGGC-------------GAG
Consensus                       **  ** **  *** ****  ***    * ** ***** *   *********** ** **** * ** ** *  ****    *  *****  ******                ***

Teleogryllus_occipitalis     GTGGTAGTGGTGGTGGTGGTAGTGGTCGGGGTGGTGTGGTAGGTCGGGGTGGTGGTGGTGGTCGGGGTGGTGGTGGTGGTGGTGGTGGTCGGGGTGGTGGTGGTGGTGGTGGTGGGGTGG
Acheta_domesticus            GCGGTAGCGGTGGTGACGGGA-----------CGATGGCAGTGTGGGGAGGTGGTGGCAGCGGCGGCGGTAGTGGTCTTGATGGCGGTC--GGTAGTGGTGGTAGTGGCCGTGGAGACG
Consensus                    * ***** *******  ** *               *** **   **** ********  *  * ** *** *****  ** *** ****  *** ******** ****  **** *  *

Teleogryllus_occipitalis     GTGGTGGTGGTGGTGGTGGTGGTGGTGGTGGTAGTGGTGGTGGTGGTGGTGGT--GGTGGTGGTGGTGGTGGGGGTGGTGGTG----GGTGGTGGTGGTGGTGTGGTAGTGGTG
Acheta_domesticus            GCGGTGGCGGTGGCAGCGGCGGCGGTGACAGCAGTAGCGG-GGCGATGGCGGTGGTGGCGAGGTGGTGACGGTGGTGGCGGCGGTGGCGGGGCGGTGGCGGTGGTGACGAGGCGATGGCA
Consensus                    * ***** *****  * ** ** ****   *  **** ** ** * *** ********   *******  ******** ** ***** *    ***** *******  * **   ***

Teleogryllus_occipitalis     GT-------GGTGGTGGTGGTGGTGGTGGTGGTGGTGGTGG----TGGTGGTGGTGGTGGTGGTGGTGGCGCTGGTGGTGGTGGTGGTGGTGGTGG-CCATCGCCGTCCCC
Acheta_domesticus            GTGTGACGAGGTGGTGGCAGCGGCGGCGGTAGTGGTGTTGATGGCAGTCGGTAGTGGTGGGAGTGGCCGTGGAGGCGGCGGTGGCGGTGGCGACGGCGCTGGCGGACCATCGCTGACACC
Consensus                    **       ********  * ** ** *** ****** ** ***     *** *******  ****  **** ****  ***** ***** *  ** * *** ** ******* * * **

Teleogryllus_occipitalis     GATGACCTGCAGGACGTCGCTCCTGCTCGGGCCGGATTTTGGGGGACTGTCCTACTCTCTCTCACACACACACACACATACGCGCGCTCGAGGAGAGACACACAGAGGAACTTTTCTG
Acheta_domesticus            GATGTCCTGGAGGA-GTCGGTCCTGCACGGGCCGGATTTTGGTGGGCTGTCCTACTCTCT----------CGCACACATACACGCGCGCTGGAGGAGACACACACAGAAGAACTTCTGTG
Consensus                    **** **** **** **** ****  ****** ************** ** ************          * ****** * ********* ******* ********* ****** * **

Teleogryllus_occipitalis     GGAAAATCGCTACGGCAGACG----TTTGACATCTCGTTCCATGTTTCTCACTTCCTCTGTCCTCATCGATATTTTGGATGATATCCAGAATGAGAACATGGGCCAGGCTGATATTGCCA
Acheta_domesticus            GGAAAATCGCTACGGCGAAGAACGTTGTGACACCTTCTTCCATGTTTCTCACTTCCTGTCTCCTCATCGATATTTTGGACGATATCCTGAATGAGATCACGGTCCAGGCTGATATTTGCA
Consensus                    ****************  *       * ***** **  ******************* * ****************** ******* ******** ** ** ************  **

Teleogryllus_occipitalis     --------GGCTAGTTTCCTGTCTAACAGAGAGGATGTCAAGAGCA--AAAAACATTCAATTAAACTTTAGAGTTGATC----------------------------------------
Acheta_domesticus            GGCTACATGGCTAGTTTCCTGTCCAGTAGAGAGGAAGTAGGGAGCAATAAAAACATCCTTGAGGAATTTAGGATTGATCAAAAGATGATGATAATTAATACCATAAAAGGTGATTTTTGT
Consensus                            *************** *  ******** **    *****  ******** *      * *****  ******

Teleogryllus_occipitalis     ------------------------------------------------------------------------------------------------------------------------
Acheta_domesticus            GTGAAAGGTCAAACTACATATTTCATTTTTATGTGTCTTCTTTGAAAACTTTGCATAGTATATTAAACAGTCAAGGCTATACATCCTTAAAACTGTGAAAGGCAAAGCAACAAAACATTG
Consensus

Teleogryllus_occipitalis     ------------------------------------------------------------------------------------------------------------------------
Acheta_domesticus            AAGTTGCTGATGCTGCAAATTTGTTTACTGACAGTCGTGGAAACGATGACGACTGACACCATACGAGTTTTCAAATGTCGGAGATTCCAGACAGCCTGTTGTGGAATTGAGTATAACAAT
Consensus

Teleogryllus_occipitalis     ------------------------------------------------------------------------------------------------------------------------
Acheta_domesticus            GAGTTCAATTTGTGTTCGAATGTCTTATTATTATCCTCAACAACAGGAAAAATTCGAATAGATAACCAGAGACAGATAACTGATTTAAAAGATTTATATATTCCTTTAAGAAAATTTCAG
Consensus

Teleogryllus_occipitalis     ------------------------------------------------------------------------------------------------------------------------
Acheta_domesticus            GCACTTGTTAGACGATTACCTTATCATAAAAGTTATAAATGTATTTAATTTTTATTCACTTTAATGCACAAACATTAATGGACAATAAAAATATTGCACAATTAATTATTTAGCAGAAAT
Consensus

Teleogryllus_occipitalis     ------------------------------------------------------------------------------------------------------------------------
Acheta_domesticus            ATTAGCAGCTCTCTTCTTAAACGTACAGCTATATACTAAAGATATAAAAATCAAGATGGTCAATGGATGACAAGATATGCTCATCAACGGATATTAATATACTGATTCATGTACTTTCCT
Consensus

Teleogryllus_occipitalis     ------------------------------------------------------------------------------------------------------------------------
Acheta_domesticus            AGTAGTGAGAAATCTCACAAAATGTCATTGAGAAATGCTCTGAAAGTATTTGTCATAACGAAACAACGATTCTTCACTTTGTTTGGCCACATTTATGACGCATACAACAACGGAAGTGTT
Consensus

Teleogryllus_occipitalis     -------------
Acheta_domesticus            TCGAAAATATTTG
Consensus
```

**Tocc.11G0007880.1_exon1**

```
Teleogryllus_occipitalis     ------------------------------------------------------------------------------------------------------------------------
Acheta_domesticus            FIPLSVVGIFRREECGGGDISSPDSMIQCQSTRSLAVFLSLDISHQQGTSARTSSCV*ERFKDKAMNRKDRSRAIRSPFVRVLLRH*CRVGRRNRVKLPIRAAGLKS*DSPPLS*AVQ*H
Consensus

Teleogryllus_occipitalis     ------------------------------------------------------------------------------------------------------------------------
Acheta_domesticus            QLMP*TT*AGFKCGVLADCLPVLLYFGAWRDHVNNKVIQGHTTVENKALWGSLQS*KIIVIPTKFRLIN*LHFNLCCT*KSETFNFT*HYIFRVVTSTGVVKKKLVCSLEIKKNPTSKSV
Consensus

Teleogryllus_occipitalis     ------------------------------------------------------------------------------------------------------------MSLI--------
Acheta_domesticus            ERLPYK*GTCPQHNN*SNKQSPLVLNSVKKYPYFPVNIHLSKTKEMTINNCEK*WK*F*SKKILKLPRKSDWKCWNFLVLQIFTIFLFLSVLSIFVTVVVVKTITH*WMSLIPAISQARG
Consensus                                                                                                                                ****

Teleogryllus_occipitalis     ------------------------------------------------------------------------------------------------------------------------
Acheta_domesticus            SAPPSARSSARPHFERPGLVAAALRVRAPAAPRGPVPRGGGSGDGDGDANDGGGGGSGGGHGRSGAVAVVARW*R*WRWWRGG*RW*RDDGSVGRWWQRRR*WS*WRSVVVVVAVETAVA
Consensus

Teleogryllus_occipitalis     ------------------------------------------------------------------------------------------------------------------------
Acheta_domesticus            VTAAAVTAVAGRWRWWRGGDDSGGGGEAVAVVTGRWQCGEVVAAAAVVVLMAVGSGGSGRGDGGGGGSGGGDSSSGAMAVVARW*RWWRRWRGGGGGDEAMAV*RGGGSGGGSGVDGSR*
Consensus

Teleogryllus_occipitalis     ------------------------------------------------------------------------------------------------------------------------
Acheta_domesticus            WWEWPWRRRWRWRRRWRTIADTDVLEESVLHGPDFGGLSYSLAHIHARWRRHTQKNFCGKIATAKNVVTPSSMFLTSCLLIDILDDILNEITVQADICRLHG*FPVQ*RGSREQ*KHP*G
Consensus

Teleogryllus_occipitalis     ------------------------------------------------------------------------------------------------------------------------
Acheta_domesticus            I*D*SKDDDN*YHKR*FLCERSNYIFHFYVSSLKTLHSILNSQGYTSLKL*KAKQQNIEVADAANLFTDSRGNDDD*HHTSFQMSEIPDSLLWN*V*Q*VQFVFECLIIILNNRKNSNR*
Consensus

Teleogryllus_occipitalis     ------------------------------------------------------------------------------------------------------------------------
Acheta_domesticus            PETDN*FKRFIYSFKKISGTC*TITLS*KL*MYLIFIHFNAQTLMDNKNIAQLII*QKY*QLSS*TYSYILKI*KSRWSMDDKICSSTDINILIHVLS***EISQNVIEKCSESICHNET
Consensus

Teleogryllus_occipitalis     ----------------------
Acheta_domesticus            TILHFVWPHL*RIQQRKCFQNI
Consensus
```

**Tocc.11G0007880.1_exon2**

```
Teleogryllus_occipitalis      -------------AVVVVM---------------------------------------------------------------------VAVGR---------------------------
Acheta_domesticus             LFP*A*WVFFVGRSVVVVISPRRIA*SSANLHVH*LFFSL*ISPTNKAPQPEQARVCKKGLRTKL*IERIDRARSVAHLLGYYSGISVVLVGEIA*NYRSEQPVSNHRTPLPCLKQCSDI
Consensus                                   ****                                                                      * **

Teleogryllus_occipitalis      -------------------------------------------------GG----------------------------------------SG---------------------------
Acheta_domesticus             S*CPEPPKPVLNVEYWQIVCRFSCTLARGEIM*IIKLSKATRR*RIKLCGGLYKVRKLL*FQQNLD*LINYILIYVVPKSQKPLTLLDITFSGW*LLQVWLKRN*SVL*K*KKIQHQNLS
Consensus                                                                        **                                        **

Teleogryllus_occipitalis      -----------------------------------------------------GGGSGRG-----------GVVG---------------------------------------------
Acheta_domesticus             NAYRISKEHAHNITTNRINNLPLF*IRLKNTPTFL*IFIFRKRKR*Q*ITAKSSGSDFRAKKF*NYPENLTGNVGIFWYCRYLQYFSFFRFSPSLSQW*SLRQSRTNGCL*FQRSLRLGV
Consensus                                                                             *   *            * **

Teleogryllus_occipitalis      ----------------RGGG-------------------------------------------GGRGGGGGGGRGGGGGGGGGVGGGG-----GGGGGGGGSGGGGG-------------
Acheta_domesticus             PLHRARGPRRDRTSSARGSSQPPCEFGRRPLPGVPCPAAAEAATATATPTTAAAVAAAAVTAEAGRWRWWRGGDDSGGGGEAGSGGDGTMAVWGGGGSGGGSGLDGGR*WW*WPWRRRWR
Consensus                                     **                                              **     **   ****  * ** *     **** *****  **

Teleogryllus_occipitalis      ----------GGGGGG------------------GGGGGGWW-------------------------------------WWCGSGGGGGGGGG-------------------------GG
Acheta_domesticus             *QRRR*QQ*RGGGGGGEVVTIVAVVARR*RW*RDDGSVGRWWQRRR*WS*WRSVVVVVAVETAVAVAAAAVTAVAGRWRWWRGGDGGGGGGGAVAVVTRRWQCDEVVAAAAVVVLMAVGS
Consensus                               ******                   *  * **                                     ** *  *******                          *

Teleogryllus_occipitalis      GGGGGGGGGGGGAGGGG------------------------------GGGGGHRRPR*--------------------------------------------------------------
Acheta_domesticus             GGSGRGGGGGGGDGAGGPSLTPMSWRSRSCTGRILVGCPTLSHTYTRAGGDTHRRTSVGKSLRRRTL*HLLPCFSLPVSSSIFWTIS*MRSRSRLIFAGYMASFLSSREEVGSNKNILEE
Consensus                     ** * ******* * **                               **  ***

Teleogryllus_occipitalis      ------------------------------------------------------------------------------------------------------------------------
Acheta_domesticus             FRIDQKMMIINTIKGDFCVKGQTTYFIFMCLL*KLCIVY*TVKAIHP*NCERQSNKTLKLLMLQICLLTVVETMTTDTIRVFKCRRFQTACCGIEYNNEFNLCSNVLLLSSTTGKIRIDN
Consensus

Teleogryllus_occipitalis      ------------------------------------------------------------------------------------------------------------------------
Acheta_domesticus             QRQITDLKDLYIPLRKFQALVRRLPYHKSYKCI*FLFTLMHKH*WTIKILHN*LFSRNISSSLLKRTAIY*RYKNQDGQWMTRYAHQRILIY*FMYFPSSEKSHKMSLRNALKVFVITKQ
Consensus

Teleogryllus_occipitalis      ----------------------
Acheta_domesticus             RFFTLFGHIYDAYNNGSVSKIF
Consensus
```

**Tocc.12G0010430.1**

```
Teleogryllus_occipitalis    ATAACACCTT---------------------------CAAATGCGAC-----------------------------------------------------------------TAAATGCA
Gryllus_longicercus         GTACCATCCTAAGGGCGAGAAAGGAAAATGAATTACACAAATGCCTCTTTCTGTCTTTCCTATTGTAAGCTATACTATTGTCGGAGATATTCCCAATCCCTCATATATATCATTAATACC
Consensus                    ** ** * *                           *******  *                                                                 * *** *

Teleogryllus_occipitalis    ATAATTCTTCTATG----------------------------------------------------------------------------------------------------------
Gryllus_longicercus         CTAATCCTTCAGTGGCGGTTTCTCAGGGGAGGAACGTTCATCTGACCTTTTCTTCTGAACGTAATTACCAAATAAAATATTTACTTTGAAGTTTAAGGGAAAAAGGTACGTTTTATTTTA
Consensus                    **** ****  **

Teleogryllus_occipitalis    -------------------TTCGATAGATTTCTGACCGTAAAAA----------------------------------------------------------------------------
Gryllus_longicercus         CTGCTATAAGGTAGATTATTTCGATGGTATTTCGATGGCGAAAACGAAGTTACTAGTTATGGAAATTAATCGATTAATTGCTTTGAGAGGAATCTCATTGTTGACGCTGCGAAAATAGCT
Consensus                                      ****** *  **  **  *  ****

Teleogryllus_occipitalis    ------------------------------------------------------------------------------------------------------------------------
Gryllus_longicercus         GAATGAGGATGGCTGGGAGAGCATAGCATTGTTGATGAAAATTAAAATGACAATGGAACATCGCTTCTTCTATCGCAGTAGTGCTTTCGTCAGAAAAGAATATTCTGGAAAATAAATGAC
Consensus

Teleogryllus_occipitalis    -------------------------------------------------------TCACACTTAC-------------------------------------------------------
Gryllus_longicercus         ACACAATTTACAAGTATTGTTACGTGGCGAAAGAAAACAAAGAGTATTGAAATTGTCACAGTTACCTCTGCCCACGAAAACTGTCCTTCCTTACGAATTCCGTGCCTTTAGCTACGTTTT
Consensus                                                                  ***** ****

Teleogryllus_occipitalis    ------------------------------------------------------------------------------------------------------------------------
Gryllus_longicercus         TCTATTACATACATTTCCTGGTACACTCTGGGTAGAACCTTGAAGACTTTCCAAAACGCCTTTTTACGAAGCGTTTCGCACTGCTTGTCGTGACAAGGCGATCGGCCGTCTGATATAGAA
Consensus

Teleogryllus_occipitalis    ------------------------------------------------------------------------------------------------------------------------
Gryllus_longicercus         ATTCTCAGCATTTCGACCGTGTAATTTTCGCGGGAGTGGAGAATCTTTCCCCGGGAGCTGGTGCGATCTTTCGGTGGCGCTCCGCACTAATAAGACAAGTTGAAATCGAATTCAAGGAAC
Consensus

Teleogryllus_occipitalis    --------------------------------------------------------------------------------AAAGCATTGTAATCAAAA----------------------
Gryllus_longicercus         TTGCAAAAAGAAATGAATGTTTTCGACACCAAAGAAGAAGAAATTCGCGCGGGTTTTGGGAGAAAATTAAGACTATCTCGAAGGCGCCACGTTTAAAATGTATGGTCTTTTCCTTTGAAA
Consensus                                                                                   ** **       * ****

Teleogryllus_occipitalis    --------------------------GGCAATGTAGCCCGGGCG----------------------------------------ATCCTTCCTTCCC-----------------------
Gryllus_longicercus         AGCCTTCAAAGAACATCAAACGCCCTGTTAACGAAGCCAAGGCGCCAAATTCGAAGTAATTTGGCCGGCAATGTGTCAACACTCATCCTTTCTTCCTCTCAGGTGTAAATAGTCAATATC
Consensus                                             *  ** * ****  ****                                        ****** *****

Teleogryllus_occipitalis    ------------------------------------------------------------------------------------------------------------------------
Gryllus_longicercus         TCCCAGCATTCCTGCGGCCAGAAGACAAAATCTTTTATAATGCATCTACGCAATAGGAAGGGAGAATATCCTCCCTTGAACTGCCATCTTTTGGCTTCTATCTGAACCAGACAAACGCGT
Consensus

Teleogryllus_occipitalis    ------------------------------------------------------------------------------------------------------------------------
Gryllus_longicercus         CACGGGCGGAGAGGGATCGCACTTCCGCAGGCAGAGCTCGAATGTGCTCTTACAATGTTGGCTGACGACAGCAACGAGGCCGAACTGATCTTTCTTATTCCATTGCAATTCGACTATTTA
Consensus

Teleogryllus_occipitalis    ------------------------------------------------------------------------------------------------------------------------
Gryllus_longicercus         ATCACATTTGTTTTATGTTGGGTTTGAAGCAGATTACGCGCCTGATTTCTATTCTACGGCTAAGAATCGTGCACCAGCCGCCACTGCACTGGTAAATTCTTGGCTATCGGAGAGACGCCG
Consensus

Teleogryllus_occipitalis    --------------------------------------------TTTCCTTAGGGCACAA------------------------------------------------------------
Gryllus_longicercus         CGTCGTTGGCTGACAGGCCGACGTTTCGTCTGACATGTCGCGGGCTTCCTCAGGGCGAAACGTCGGCCTCTCAGCCAAGGACGCGGCGTTTCCGGACAGCCAAGAATTTATGGATTCGTG
Consensus                                                    ***** *****  **

Teleogryllus_occipitalis    ------------------------------------------------------------------------------------------------------------------------
Gryllus_longicercus         CCGCGGAAGCTTACGTGTGCACCATTGCACTGGATTGATCGTTCATTGCCACCTGCTCCTACCGGCGCAGCGTATGTGGCCTTCTCTTAACCAGTGTAGGTTAGGTTGGTACTCGTGTCG
Consensus

Teleogryllus_occipitalis    -----------------------------------------------------------------------CCCATCGAGCCCATCTTCATCTCAAAGATGGAGAAAACCTCCGGCA---
Gryllus_longicercus         CTCGGTCTTGCCCTTAGCACTCACGACCCCAAAACCCATCACAGCATCAGCTGGGAGAGGACCTGCACAATCACATCAACGCTTCCTCCAATTCTTAGGATGAGAGCGCCTTTACCATAT
Consensus                                                                          * **** *  *   ** **  **  **   ****   ***    **

Teleogryllus_occipitalis    ------------------------------------------------------------------------------------------------------------------------
Gryllus_longicercus         AGGTTTTTCTTCATTAAGTGAACATCGACCAGCACACTGTACACAATCTTATTCATTCGGCTGCCACTTTCAAAACAATTCAACAGTGGGGAATATTTTGATACTACTACACAGAAGAAA
Consensus

Teleogryllus_occipitalis    -------------------------------------------------------------------------------------------------AGTGCCAA---------------
Gryllus_longicercus         TTGGCTTCTGGAAAGATTATGTACGCACTTAATAGAAAATGAACGCAAAACGAAATCAAGGAAAGAAGACAAAGCGAAGGAACTAATTTGTCACATCAGTGTCAAGTTAACTTACGAAGT
Consensus                                                                                                    **** ***

Teleogryllus_occipitalis    ------------------------------------------------------------------------------------------------------------------------
Gryllus_longicercus         GGTGAAAGAAAATATTTGGTCTACAAACCACCCAATATGAATACTTCCTCCACAGAATACAGTTATGGTTTTACTTAATTTTCAGCCAAAACATAACACCGTGACGAAGTGAAATGATTT
Consensus

Teleogryllus_occipitalis    -----------------------------------------------------------------------------------------------------------CAGATTCGAG---
Gryllus_longicercus         TTTTCTTCTACATTCGATAGATTTCTGATCGTACAAATGACAGATCTACGAAACACTTGACCGTTCGTTCCCCATTCTTAGGGCCAAATCCATAATTTCTTGGGGGTCATGTTCAAAGAT
Consensus                                                                                                              **  *** *

Teleogryllus_occipitalis    --------ATGTCACGAACAGCTGCAAATTCCTCCACATAACATAATGTCTCAGATCTTGCACAAACACGTAACCGAATACATACATAAATAAAAGCACTAAATGCA-CGCTTTTATCCC
Gryllus_longicercus         AAAATGTCATGTCTCGGACAGCTGCGAATTCCTCGA---------------CAGATTATGTCCTGAAGTACAACCCAATACATACATAAATACAAGCACTAATTAAACCGTTTTTATGTC
Consensus                           ***** ** ******** ******** *               *****  **  *  *     **** **************** ********* *  * ** ******  *

Teleogryllus_occipitalis    TTACGAAATTGTATGGTATTTATCGGAAGTACCAAATAACAGAAAATGTATTTGAGTACGAGCA---TAGGTTCAGAATAGGTTGAAATCTAGGAATTTCGTACTGATTAATGAAAAGTT
Gryllus_longicercus         ATTTGGAATTGTTTGCTATTTCTCGGAAGATCCAAATACTAAAATATTTAATCCAGTAAGAGCAAACTAGGTTCAGAATAAGTCAAAATCTTGGAACTTCGTGCTGATGAAGAAGGTGGG
Consensus                    *  * ****** ** ***** *******  *******  * ** ** ** *  **** *****   ************ **  ****** **** ***** ***** **  *   *

Teleogryllus_occipitalis    GATTAACGTCTATTTCGCAACAAATTTCTGGGACTTCGTCTGGTTTTCCAGTTCTTC-----CTCCTACTGTGAC--TATTTGTTGGATTTGAACAAAACATAAAAATTTTACATTAAAA
Gryllus_longicercus         CAC-----TTTTTTCTGCAATACCTTTCTCTGACTTCGTCTCGTTTTCCAGGTTTTCCAGCACTTCGACTGTGACAATATTTGCTTCAGCAAAGCAATATATCACAA-TTTACATTTAAA
Consensus                    *      * * **  **** *  *****  ********** ********* * ***     ** * ********  ****** *  *    * *** * ** * ** ******** ***

Teleogryllus_occipitalis    TAATACATCAGATTTCAGCAATAGCAAG------GTGACTATTCAGCTCCGATTTCATCGTGAAAAGGACCCAACGAATAAAAAG--------------------CTATGAATCCA----
Gryllus_longicercus         TAATACACGATAGTTAAGTAATACCAGGGCAGCTGTAACTATTCAGCTCTGATTTTACTGTTAAAAGGAATAAAAGGAACGAAAGGTTCACAGTGCCGATATTATTTACGGAGCAACGTC
Consensus                   *******  * * ** ** **** ** *      ** ************ ***** *  ** *******   ** * *   ****                   ** * * * *

Teleogryllus_occipitalis    ----TCTACGAGTGTGATT-------------------------------------------------GCGCGCG---------------------CAGAGGCCGTTGGCTCCTCGGGCG
Gryllus_longicercus         GGACTTCATGAATGTGGTCTCCAAAACGAATCATTCTTCCGCCACGCCGGGACTCTCCCACTTGCACAGTGAGCGTTTCGCGTTCATCGGAGGCATCGGAGGCCGTTGGCTCCCCCGGCG
Consensus                       *  * ** **** *                                                 * * ***                     * ************** * ****

Teleogryllus_occipitalis    ACCGCCGCGCCGCG---CGCCTCCGCCCGGGATCAAACCGGCGACCTTGGGGGCTAGTATGGATCT----------GCTAGGCCGCATGGGGCGCTGCGTTGTGCAGGTCTGATGTAGAC
Gryllus_longicercus         AAAAAAACGGCGCGCGCCGCCGCCGCCCGGGATCAAACCGGCGACCTTGGGGGCTAGTATGGATCTGCGGCTGAGAGCTAGGCCGCATGGGGCGCTGCGTTGTGCAGGTCCGATGTAGAC
Consensus                   *      ** ****   **** ********************************************          ********************************* *********

Teleogryllus_occipitalis    GCACCCCAGCCCCGCTCCCCGCGCCCCGCCCCGCC------CGCCGCCCGCGCGTGCCAGACCGGCGGGATGACGTCAGAACCGGCCTGTTACCATGACGACGTGCTGCTGCCGCCGCCG
Gryllus_longicercus         GCATCCCGGCCCCG--CGCCGCGCCCCGCCCCGCCCCCACGCCCCGCCCGCCCGCGCCAGACCGGCGGGATGACGTCAGAACCGGCCTGTTACCATGACGACGTGCTGCTGCCGCCGCCG
Consensus                   *** *** ******  * *****************      * ******** ** *****************************************************************

Teleogryllus_occipitalis    CCCGAAGGCCGAGGTGGAGGGGAGGCGCAAGGTGACGCATGCGCGCTGGGCCG-CCGCGCAGAGCCGGAGGGAGG------------GGGAGGGGAAAGAGGTCTTTGGCCCAGGAGTC
Gryllus_longicercus         CCCGAGGAGGGCGGGCGGGCGGGCGGGCGAGGTGACGCATGCGCGCTGGGCCGCCCGCGCAGCGCGGGGGAGGGGGCAGGGGCCAGGAGGGGGGAGAGAGAGGTCTTTGGCCCAGGAGGC
Consensus                   ***** *   * **  * * **  * ** *********************** ******** ** ** * * **             *** ** ** ******************* *

Teleogryllus_occipitalis    GCGCATCGCCGCTGCGCTGTGTACAAATACCCCGTCCTCCTCCTCCTCCTCCTCCGCACGGAGAGGGCGCAGACCCCGCACGCACTCGCTCACAAGGCGGCCGCCACGCACACACCA
Gryllus_longicercus         GCGCATCGCCGCTGCGCTGTGTACAAATACCCCGCC-----------------------------------------------CGCACTCGCTCACGAG-----------AAACAGAAAA
Consensus                   ********************************** *                                               ************* **           *** *  *

Teleogryllus_occipitalis    AGAAACAGGAGGGGAAATAGGCAGAAATAGAAGGTAAAGGAAAATACGACCAAACAGAAAATAGCACGAAATACAAAATAGAA------------AGAAAAATATCAAAAGGAGA-AATG
Gryllus_longicercus         GGAAACACGCGAAGAAACAGA-AGGAACGGGAGATACAGGAAGAAACGG-GACACGAAAAATCAGAAAAAACAGGAGGAAGAACTAGAACCGAACAAAAAAATTTCGAAATGACATGATT
Consensus                    ****** * *  **** **  ** **  * ** ** ***** * ***    * **  *****   *  *** *  *   ****           * ****** ** *** ** *  **

Teleogryllus_occipitalis    TGTGACCTGTGTCTATAAGACTAGTATACATTCTAAGAAG-TATCATGCTGTACTAACAGTCAACTTAAACTAGTGGTGCTGCAACTCAACATATTTTCTCGCTTTATCTCTAACGAGAA
Gryllus_longicercus         TGTGACCAGCATCTACAAGACTAGAATACAATTCAAGACGTTGCCATGCTTTGGTAACGGACTACTTAAACTAGTGGTGCTACAGCTCCACACATTTTATCGGATTATTTGTACTGGCTC
Consensus                   ******* *  **** ******** ***** *  **** * *  ****** *  **** * * ***************** ** *** *** ***** ***  **** * **  *

Teleogryllus_occipitalis    GACGACAATGAAAGACGAC----------------AATGAAAGAA-------------------GAAGAAGACGAAGAAGAAGA------------------AGAAGACGACAATGAAA-
Gryllus_longicercus         CGCAATATTATACTACTACCTGAATCTGTAATTCTAATCAGCAAATTAGTATTATTCCGCACCCAAGCAAGTCCATACAGTACACTCGCCGTGCGCTGTCCCATTACACGACACAGACAC
Consensus                     * * * *  *  ** **              *** *   **                 *  *** * *   ** * *                *  * ******  ** *

Teleogryllus_occipitalis    ----------------------------------GACGACAATGAAAGAAGAAGAAGACGAAGAAGAAGA------CGACAATGAAAGACGACAATGAAA----------------GAAG
Gryllus_longicercus         TCCGTACTCAATTGCGCTCAAGCAGAAACGATGTGGCGAGAATGTAAAAAGTAACCAATAAACACCCAGAATCTCTCAACAGT--AACTCAATAATGCCATGCACACTGTGGTTTTAACA
Consensus                                                     * *** **** ** *** *    *  ** *   ***      * *** *  **  * * ****  *               *

Teleogryllus_occipitalis    AAGACGAAGAAGAAGAAGAAGAAGACGACAATGAAAGAAGACGA------CAATGAAAGAAGAAGAAGACGAAGAAGAAGAAGAAGACGAAAGAAGAAGACGACGAAAGAAAACGAAAGA
Gryllus_longicercus         TAATCGGTGGTAAAGAAAAAGAGGACGAAGGAGAAAGAAGACGACGAAAGCGACGAAGGAGAAAGAAGACGACGAAAGCGACGAAGGAGAAAGAAGACGACGA-AAGCGACGAAGGAGAA
Consensus                    *  **  *   ***** **** *****    ************      * * *** **  ********** ***   ** ****  ********* *****  *  **  * * *  *

Teleogryllus_occipitalis    AGAAGACGACGAAA-----GAAAACGAAAGAAGAAGACGACGAAAGAAGAAGAAGAAGAAGAAGAAGAAGACGAAGAAGAAGACGAAAGAAGAAGAAGACGAAGAAGAAGAAGAAGAAG-
Gryllus_longicercus         AGAAGACGACGAAAGCGACGAAGGAGAAAGAAGAAGACGA---------AAGCGACGAAGGAGAAAGAAGACGACGAAAGCGACGAAGGAGAAAGAAGACGACGAAAGCGACGAAGGAGA
Consensus                   **************     ***   ***************         ***      * **  ********** ***   ****** **  ********** ***   ** **** **

Teleogryllus_occipitalis    AAGACGAAGACGAAGACGAAGAAGAAGACGAAGAAGACGAAGAAGACGAAGAAGAAG-AAGAAGAAGAAGAAGACGAAGAAGAAGAAGACGAAGAAGAAGAAGAAGACGACGAAGAAGAA
Gryllus_longicercus         AAGAAGACGACGAAAGCGACGAAGGAGA--AAGAAGACGACGAAAGCGACGAAGGAGAAAGAAGACGACGAAAGCGACGAAGGAGA-----AAGAAGACGACGAAAGCGACGAAGGAGA-
Consensus                   **** ** ******  *** **** ***  ********** ***  *** **** ** ******* ** ***  *** **** ***     ******* ** ***  ******** ***

Teleogryllus_occipitalis    GAAGAAGGCGACGAAGACGAAGAAG----AAGAAGAAGACGAAGAAGAAGAAGAAGACGAAGAAGACCAAGAAGAAGAAGAA-GAAGACCAAGAAGAAGACCAAGAAGAAGACCAAGAAG
Gryllus_longicercus         -AAGAAGACGACGAAAGCGACGAAGGAGAAAGAAGACGACGAAAGCGACGAAGGAGA--AAGAAGACTAAAACTACGCAGAACGAAGACGAAAAAGAAGACCAACTGAAAAATCAAAAGG
Consensus                    ****** *******  *** ****    ******* ******   ** **** ***  ******** ** *  * * **** ****** ** ***********    ** * *** * *

Teleogryllus_occipitalis    AAGAAGAAGACCAAGAAGAAGAAGAAGAAGAAGACGAAGAAGACGAAGACCAAGAAGAAGACGAAGACCAAGAAGAA-------------------------------------------
Gryllus_longicercus         GGAAAATGAATTAGGACGAAAATTAAGAGACCTGAAATGAGTAT-----CTGATGAGGGATGGAGGGGCAAGGGGAGGCATGTGATGTGATTGGGCGCAGCTTATCTGCGCGGGTGGACC
Consensus                      **    *  * ** *** *  ****        * **  *       *  *  **     ** *  ****  **

Teleogryllus_occipitalis    ------------------------------GACGAAGACCAAGAAGAAGAAGAAGACCAAGAAGACGAAGAACAAGAAGAC---------------------------------------
Gryllus_longicercus         TTACCCAGCACCCCTCCCCCTTCCCCCGCCGCCGAACAACACACAACATTACAAACACAGTGAAATGAACATATCGAGGACACGTCTCTACGAACTGTGCTCTCAAGGCAGCAATTCGCC
Consensus                                                 * **** * **   *  *  * **   **   * * *** *    ** ***

Teleogryllus_occipitalis    -----------------------------------------GAAGAAGAAGAA--------GACCAAGACGAAGACCAAGAAGAAGAAGAAGACGAA-----------------------
Gryllus_longicercus         TTCCCCGCCTTCTCGAGCACTTCTTGCCATTTCACCGCTCTAGAAAAAAACAATGTTCGCCAACTAACGTGTTGCGCAAAAATAATGAGTTGGCAAACTAACTACCAACGGGCTCCGCGA
Consensus                                                                * ** ** **         ** **   *  *  *** ** **  **  * * **

Teleogryllus_occipitalis    -------GACCAAGAAGAAGA---------CCAAGAAGAAGACC----------------------------AAGACCAAGAAGAAGACCAAGACCAAGAAGAAGACG------------
Gryllus_longicercus         ATGATTTGCGCAGGCAGCAGAGTTGAGGCTCCGAAACGAAGAACTACCGCTTTTCCATCTTCCTTCTCCAAGGAAATAAACAATCGGCCGTACCCATGAATTAGTTATCGTTTAAACTG
Consensus                          *  ** * ** ***         ** * * ***** *                           * *  ** **   * **    *** ***  **

Teleogryllus_occipitalis    ------------AAGACCAAGAA--------------------------------------GAAGACGAAGAC-----------------------------------------------
Gryllus_longicercus         TATTAATTGTACAAGACTAGATAAGCACGTTATAAATATTGCAATCTCTATTCCGCTTCTGGGATAAGAAGTCTCTCGTTGACATACAGTTTTATTGATCCGTTCAGTGAATGTTCTTAC
Consensus                               ***** *   *                                      * * * **** *

Teleogryllus_occipitalis    -----------------------------------------------------------CAAGAAGAAGACGAAGACCAAGAAGAAGACGAAGACCAAGAAGAAGACGAAGACCAA----
Gryllus_longicercus         GACCCTTCGCTTTTCTCCCTTCCACGCGTCACGTGCAGCTTTCACAGTTTTAAACGTTATAAGTAAATTACGCTAATTA--AAAAAGTCCGTGTATTAAGAGAGGACAAACTGCAATTGC
Consensus                                                                               *** * *  ***   *  *  ** *** *   *    *  *** *** **   ***

Teleogryllus_occipitalis    -----GAAGAAGACGAAGACCAAGAAGAAGACGAAGACCAA----------------------GAAGAAGACGAA----GACCAAGAAGAAGACGAAGACCAAGA---------------
Gryllus_longicercus         TCAGGGTAAAACCTAAATTTCGGCGAACAAATACAAATCAATCTATTCATTTCAAACTCTGCGAAATAATATCAAAGTGAACTTAATTACACACAGTTGTCAAAATAGCGTTTCTTGTGT
Consensus                        * * **    **   *    *  * *   * * ***                      ** ** *  **     **  *     * **      *** *

Teleogryllus_occipitalis    --------------------------------------------AGAAGACGAAGACCAAGAAGAA-----------------------------GACGAAGAAGACCAAGAAGAAGACG
Gryllus_longicercus         GATCTCTAACGCATTATTCGTTATAATTTGTCTCGACTGTTCTTAATAAACACAGGCCATTAAGAGTAGGTTACCGTTCATCTTTGTTTGGTTGCGATAAGTTGGTACAACGGTACGACG
Consensus                                                               *  * **  ** ***  ****                             **  *    *  ***    * ****

Teleogryllus_occipitalis    AAGAAGAAGAAG-------------------------------------------------------------------------ACGAAGAAGACCAAG--------------------
Gryllus_longicercus         AAAAGTAATTAACGGCAATCTCGTCATCCATCGAATTCTCCGACGCCCGGCCACGTCTAAAACTTCACACACTACCAATCACTTTCAAAGAAAATCACGCGCAAACCTTAACCTCTGAC
Consensus                   ** *  **  *                                                                           * ***** * ** *

Teleogryllus_occipitalis    -------------------------------------AAGAAGACGAAGAAGAAGAA------------------------------------------------------------GAC
Gryllus_longicercus         TCTCGGGTAGCGCTTACATCTTTCCCCTCCCTTTGTCGAGAGGCCTGCGAGTAATTACCTCCAATATTTTTCTTCTTGGTATATTAAAATAATTGGCGTCTTTCAACATTCAATACCAAT
Consensus                                                         *** * *   **  **  *                                                             *

Teleogryllus_occipitalis    GAAGAAGAAGAAGACGAAGAAGAAGAAGACGAAGAAGAAGAAGACGAAGAAGAAGA--------AGACGAAGAAGAAGAAGACGAA----------------GACGAAGAAGAAGACG--
Gryllus_longicercus         GCGGGCGGGCGGGCGGGCGGGCGGGCGGGCGGGCGGGCGGGCGGCCAGGACGTCAAAGCCATCTAGACGTAGAACCGAGATGCGAACTGGTTGCATTGGCACGCCGAAAGACCGGACGTT
Consensus                   *  *  *     *  *  *     *  * **     *  *  * * * ** *   *        ***** ****     *  ****                * ****  *   ****

Teleogryllus_occipitalis    --------------------------------------------AAGACGAAGAAGAAGACGAAGACGAAGAAGAAGATG--------------------AAGACGAAGAAGAAGATGAA
Gryllus_longicercus         TTAAATTAGCGAAACTTAATCCGATTTTTCGCGATTTTTCACTAAATACTGAATATAAATTTTTGAAATAAAACTAGATACCACCGACCCCATCCACCTCAGAATGACACGTCATTTGTG
Consensus                                                               ** **  *  * **      **   * **  ****                    *  * **      *  **

Teleogryllus_occipitalis    GACGAAGAAGACGAAGAAGATGAAGACGAAGAAGATGAAGACGAAGAAGATGAAGACGAAGAAGAAGACGAAGAAGATGAAGACGAAGATGAAGACGAAGAAGATGAAGACGAAGAAGA-
Gryllus_longicercus         AACGTAACACACTCTGAAACTACGCACGAAATCGTGTATTAATAATAATAATAATAATAATAATAATAATAATAAGAAGAAGAAGAAGAAGAAGAAGAAGAAGAAGAAAACAAAGGGGAA
Consensus                    *** *  * **   ***  *    *****   *    *   *   ** ** *   ** *   ** ** ** *   ** **** ***** ***** ***** ******** *** ** ***  **

Teleogryllus_occipitalis    --------------------------------------------------------TGAAGACGAAGAAGATGAAGACGAAGAAGATGAAGACGAAGAAGATGAAGACGAAGAAGATGAA
Gryllus_longicercus         CAGCCTTCGCTCGTAAGATCGCTGGATAACCTCAACACTATAACCTATTGATCTATTAGTATCAAAGGCTACGGTCATTAGCCCCGTAAGTTTAGTGCATATTTTATTGCAGGAGGTGGC
Consensus                                                                           *     * ***   * *   *  *      * *       * * **      * ** ** **

Teleogryllus_occipitalis    GACGA------------AGAAGATGAAG-------------------------------------------------------ACGAAGAAGATGAAGACGAAGAAGATGAAGACGAAGA
Gryllus_longicercus         GGTGATCTTCTCCGGCTGGATGACGAAATTCAGCGCCGACCTATCCGTCGACCAACGGGGGTTTCCCCACCCAGATATTTCCCACCAGAGGGGCAATGTCCCGGCAGAGACAAGCTCTAA
Consensus                   *  **            ** ** ***                                                         ** *    *   * * *   * ***   *  *    *

Teleogryllus_occipitalis    AGAT---------------------GAAGACGAAGAAGATGAAGACGAAGAAGATGAAGACGAAGAAGATGAAGACAATGAAAGAAGACGAAGGACGAAGATGACGAAGGACGAAGGCTA
Gryllus_longicercus         GGTTCCGACTGGTTCTCCGGCATGGGGAGTGGCGTGGACTGCATGTAAGGAAGGAAGGAAGGAAGGAAGGAGGGAAGGAGGGAAGGAAGGAAGGAGGGAAGGAGGGAAGGAAGGAA
Consensus                    * *                     * **  *       ** *    * ****      * **** *    * **       **   * ****** * **    * ** *** * * *  *

Teleogryllus_occipitalis    AGGAACCGTATCGGATGGAGGTGATGGGGAGGGCAGGGGAGGCGTGCGCAGTGAATGGGTCCATTACAAATGCAGTGACATGCAGACGCCGAGGGCGCCGCTCCACCTGCTTCTTTACTA
Gryllus_longicercus         GGAA--------GGAAGGAGGGAAGGAGGGAAGGAAGGAAGGAAGGAAGGAAG---------------------------------------------------------------
Consensus                    * *        *** *****  * * **   * * ** ***   *      * * *

Teleogryllus_occipitalis    CGCCCGCCTCTCCGCCACTTAGTGCCTTTTCACAACATCTCTTCACAAACGAACCACGGGTTCAGGCAACAAATCATGTGTGCCAATGAATTAGATTTTTAAATTGTAAAATTCTAAAA
Gryllus_longicercus         ------------------------------------------------------------------------------------------------------------------------
Consensus

Teleogryllus_occipitalis    GACTGGATATTCTTGTAACAACTACTGCAAGCTCTATTCAGCTTCTGGGATAATGGTCTGT
Gryllus_longicercus         -------------------------------------------------------------
Consensus
```

**Tocc.12G0010430.1_exon1**

```
Teleogryllus_occipitalis      -------------------------------------------------------------------------------------------------------MH---------------
Gryllus_longicercus           LPSFLPSFLPSFPPSFLPSFPPSLLPSFPPSLLPSFLPSFLTCSPRHSPCRRTSRNLRACLCRDIAPLVGNIWVGKPPLVDG*VGAEFRHPAGEDHRHLLQ*NMH*TYGANDRSL*Y**I
Consensus                                                                                                                                  **

Teleogryllus_occipitalis      ------------------------------------------------------------------------------------------------------------------------
Gryllus_longicercus           NRL*C*GYPAILRAKAVPLCFLLLLLLLLLLLIIIIIIIIIIINTRFRA*FQSVLRSQMTCHSEVDGVGGI*FYFKNLYSVFSEKSRKIGLSFANLKRPVFRRANATSSHLGSTSRWL*R
Consensus

Teleogryllus_occipitalis      ------------------------------------------------------------------------------------------------------------------------
Gryllus_longicercus           PGRPPARPPARPPARPHWY*MLKDANYFNIPRRKILEVITRRPLDKGRGKM*ALPESQRLRFARDFL*K*LVVCEVFRRGRASENSMDDEIAVNYFSSYRCTNLSQPNKDER*PTLNGLC
Consensus

Teleogryllus_occipitalis      ------------------------------------------------------------------------------------------------------------------------
Gryllus_longicercus           LLRTVETNYNE*CVRDHTRNAILTTVCN*VHFDIISQSLK*ID*FVFVRRNLGFTLSNCSLSSLNTRTFLISVIYL*RLKL*KLHVTRGREKSEGS*EHSLNGSIKLYVNERLLIPEAE*
Consensus

Teleogryllus_occipitalis      ------------------------------------------------------------------------------------------------------------------------
Gryllus_longicercus           RLQYL*RAYLVLYN*YSLNDN*FMGTADLFISLEKEDGKAVVLRFGASTLLPAQIIRGARW*LVCQLIIFAQHVSWRTLFFSRAVKWQEVLEKAGKANCCLESTVRRDVSSICSFHCVCN
Consensus

Teleogryllus_occipitalis      ------------------------------------------------------------------------------------------------------------------------
Gryllus_longicercus           VVCCSAAGEGGGVLGKVHPRR*AAPNHITCLPLPLHPSSDTHFRSLNFRPNSFSPFDFSVGLLFRLRSA*F*SSFSFVAFVVFFLLRRFRRLLSPSSLSSSSFSFVAFVVFFLLRRFRRL
Consensus

Teleogryllus_occipitalis      ------------------------------------------------------------------------------------------------------------------------
Gryllus_longicercus           LSPSSLSSSSFSFVAFVVFFLLRRFRRLLSPSSLSSSSFSFVAFVVFFLLRRFRRLLSPSSLSSSSFSFVAFVVFFLLRPLFLYHRLC*NHSVHGIIELLLRDSGCLLVTFYILATSFLL
Consensus

Teleogryllus_occipitalis      ------------------------------------------------------------------------------------------------------------------------
Gryllus_longicercus           ERN*VRSVCVV*WDSARRVYCMDLLGCGIILIC*LELQIQVVV*YCGASTNNPIKCVEL*HH*FK*SVTKAWQRLELYSSLVDAGHKSCHFEIFLFGSSSSSCFF*FFVSRFFLYLPFLL
Consensus

Teleogryllus_occipitalis      ------------------------------------------------------------------------------------------------------------------------
Gryllus_longicercus           FLRVFPFLFLVSECGRGICTQRSGDARLLGQRPLSPPSWPLPPPPRCAGGPARMRHLARPPARPPRAAAAAAARRHGNRPVLTSSRRSGAGGRGVGAGRGAARGRDASTSDLHNAAPHAA*
Consensus

Teleogryllus_occipitalis      --------------------------------------------------------------------------------AFIPY-----------------------------------
Gryllus_longicercus           LSAADPY*PPRSPV*SRAAAARAVFFAGGANGLRCLR*TRNAHCASGRVPAWRKNDSFWRPHS*SPTLLRK*YRHCEPFVPFIPFNSKIRAE*LQLPWYYLTIVYYLNVNCDILLC*SKY
Consensus                                                                                                                ***

Teleogryllus_occipitalis      ------------------------------------------------------------------------------------------------------------------------
Gryllus_longicercus           CHSRSAGKPGKRDEVRERYCRKKCPPSSSARSSKILTYSEPSLLLLD*IF*YLDLPRNSKQFQMT*KRFN*CLYLCMYWVVLQDIICRGIRSCPRHDILSLNMTPKKLWIWP*EWGTNGQ
Consensus

Teleogryllus_occipitalis      ------------------------------------------------------------------------------------------------------------------------
Gryllus_longicercus           VFRRSVICTIRNLSNVEEKNHFTSSRCYVLAEN*VKP*LYSVEEVFILGGL*TKYFLSPLRKLT*H*CDKLVPSLCLLSLISFCVHFLLSAYIIFPEANFFCVVVSKYSPLLNCFESGSR
Consensus

Teleogryllus_occipitalis      ------------------------------------------------------------------------------------------------------------------------
Gryllus_longicercus           MNKIVYSVLVDVHLMKKNLYGKGALILRIGGSVDVIVQVLSQLML*WVLGS*VLRARPSDTSTNLTYTG*EKATYAAPVGAGGNERSIQCNGAHVSFRGTNP*ILGCPETPRPWLRGRRF
Consensus

Teleogryllus_occipitalis      ------------------------------------------------------------------------------------------------------------------------
Gryllus_longicercus           ALRKPATCQTKRRPVSQRRGVSPIAKNLPVQWRLVHDS*P*NRNQARNLLQTQHKTNVIK*SNCNGIRKISSASLLSSANIVRAHSSSACGSAIPLRP*RVCLVQIEAKRWQFKGGYSPF
Consensus

Teleogryllus_occipitalis      ------------------EI----------------------------------------------------------------------------------------------------
Gryllus_longicercus           LLRRCIIKDFVFWPQECWEILTIYT*EEERMSVDTLPAKLLRIWRLGFVNRAFDVL*RLFKGKDHTF*TWRLRDSLNFLPKPARISSSLVSKTFISFCKFLEFDFNLSY*CGAPPKDRTS
Consensus                                         **

Teleogryllus_occipitalis      ------------------------------------------------------------------------------------------------------------------------
Gryllus_longicercus           SRGKILHSRENYTVEMLRISISDGRSPCHDKQCETLRKKAFWKVFKVLPRVYQEMYVIEKRS*RHGIRKEGQFSWAEVTVTISILFVFFRHVTILVNCVSFIFQNILF*RKHYCDRRSDV
Consensus

Teleogryllus_occipitalis      ------------------------------------------------------------------------------------------------------------------------
Gryllus_longicercus           PLSF*FSSTMLCSPSHPHSAIFAASTMRFLSKQLID*FP*LVTSFSPSKYHRNNLPYSSKIKRTFFP*TSK*IFYLVITFRRKGQMNVPPLRNRH*RIRVLMIYMRDWEYLRQ*YSLQ*E
Consensus

Teleogryllus_occipitalis      ------------------------------------------------
Gryllus_longicercus           RQKEAFV*FIFLSRP*DGTPLEKKFM*SCATKGRVFPKKNSYC*TLVT
Consensus
```

**Tocc.12G0010430.1_exon2**

```
Teleogryllus_occipitalis        KKKTKKKK------KTKE-------------EDDERKRKKKTTK---------------------------------------------------------------------------
Gryllus_longicercus             --VTRV*Q*EFFFGKTRPFVAHDYINFFSNGVPS*GRERKMNYTNASFCLSYCKLYYCRRYSQSLIYIINTLILQWRFLRGGTFI*PFLLNVITK*NIYFEV*GKKVRFILLL*GRLFRW
Consensus                          *          **                    * **   *

Teleogryllus_occipitalis        ----ENE---------------------------------------------------RRRR---------------RKKKKKKKKKTKKKTKE----------------------
Gryllus_longicercus             YFDGENEVTSYGN*SINCFERNLIVDAAKIAE*GWLGEHSIVDEN*NDNGTSLLLSQ*CFRQKRIFWKINDTQFTSIVTWRKKTKSIEIVTVTSAHENCPSLRIPCL*LRFSITYISWYT
Consensus                           ***                                                     *  *                *** *     *     *

Teleogryllus_occipitalis        ----------------------EEDEEEEEE-----------------------------EDEDEDEEEDEEDEEDEEEEEEEE---------------------------------
Gryllus_longicercus             LGRTLKTFQNAFLRSVSHCLS*QGDRPSDIEILSISTV*FSREWRIFPRELVRSFGGAPH**DKLKSNSRNLQKEMNVFDTKEEEIRAGFGRKLRLSRRRHV*NVWSFPLKSLQRTSNAL
Consensus                                               *     *                               *           *       ***

Teleogryllus_occipitalis        ------------------------------------------------DEEEE--------------------------DEEEEE-------DDEEEEE--------------------
Gryllus_longicercus             LTKPRRQIRSNLAGNVSTLILSSSQV*IVNISQHSCGQKTKSFIMHLRNRKGEYPPLNCHLLASI*TRQTRHGRRGIALPQAELECALTMLADDSNEAELIFLIPLQFDYLITFVLCWV*
Consensus                                                           *                             * *       **  * *

Teleogryllus_occipitalis        ----------GDED---------------------------------------------------------------------------------------------------------
Gryllus_longicercus             SRLRA*FLFYG*ESCTSRHCTGKFLAIGETPRRWLTGRRFV*HVAGFLRAKRRPLSQGRGVSGQPRIYGFVPRKLTCAPLHWIDRSLPPAPTGAAYVAFS*PV*VRLVLVSLGLALSTHD
Consensus                                 * *

Teleogryllus_occipitalis        -----------------------------------------------------EEEEEDEEEEEDEEDQEE----------EEEDQEEDQEEDQEE----------------------
Gryllus_longicercus             PKTHHSISWERTCTITSTLPPILRMRAPLPYRFFFIK*TSTSTLYTILFIRLPLSKQFNSGEYFDTTTQKKLASGKIMYALNRK*TQNEIKERRQSEGTNLSHQCQVNLRSGERKYLVYK
Consensus                                                                    *  *   *                 * *  *  * *

Teleogryllus_occipitalis        -----------------------EEDQEE------------------------------------------------------------------------------------------
Gryllus_longicercus             PPNMNTSSTEYSYGFT*FSAKT*HRDEVK*FFSSTFDRFLIVQMTDLRNT*PFVPHS*GQIHNFLGVMFKDKMSCLGQLRIPRQIMS*STTQYIHKYKH*LNRFYVIWNCLLFLGRSKY*
Consensus                                        *

Teleogryllus_occipitalis        -----------------EEEEDEED----------------------------------------------------------------------------------------------
Gryllus_longicercus             NI*SSKSKLGSE*VKILELRADEEGGHFFLQYLSLTSSRFPGFPALRL*QYLLQQSNISQFTFK*YTIVK*YQGSCNYSALILLLKGIKGTKGSQCRYYLRSNVGLHECGLQNESFFRHA
Consensus                                *   ***

Teleogryllus_occipitalis        ---------------------------------EDQEED------------------------------------------EDQEED-------------------------------
Gryllus_longicercus             GTLPLAQ*AFRVHRRHRRPLAPPAKKTARAAAARDQTGDLGG*YGSAAES*AAWGAALCRSDVDASRPRAAPRPAPTPRPPAPDRRDDVRTGLLP*RRAAAAAARGGRAGGRAR*RMRAG
Consensus                                         **  *                                            *   *

Teleogryllus_occipitalis        ---------EDQEEEE------------------------DQEDEEQEDEEEEDQDEDQEEEEDEDQEEDQEEDQDQEEDQDQEEDEDQEE---------------------------
Gryllus_longicercus             PPAQRGGGGRGQEGGERGLWPRRRASPLRCVQIPRPHSLTRNRKGNTRRNRRNGRYRKKRDTKNQKKQEEELEPNKKISK*HDL*PASTRLEYNSRRCHALVTDYLN*WCYSSTHFIGLF
Consensus                                  **  *                                                   ***  *         *        *

Teleogryllus_occipitalis        ------------DEDQE---------------------------------EDEDQEEDEDQEEDED-------------------------QEEDEDQEEDEDQEEDEDQEEDEDQE
Gryllus_longicercus             VLAPQYYTTT*ICNSNQQISIIPHPSKSIQYTRRALSHYTTQTLRTQLRSSRNDVARM*KVTNKHPESLNSNSIMPCTLWF*HNRW*RKRGRRRKKTTKATKEKEDDESDEGERRRRKRR
Consensus                                       *                                    *            *                                   * ** * **

Teleogryllus_occipitalis        EDEDQEEDEEDQEEDEEEEDEEDQEEDEEEEDEEEEDEEEEDEEEEDEEEEDEEEEDEDEEEDEDEEEDEDEEEDEDEEEDEDEEDEEDEDEEDEDEEDEDEEEDEEDEDEDEDEEDEDE
Gryllus_longicercus             RRKKTTKATKEKEEDESDEGERRRRKRRRRKKTTKATKEKED-DESDEGERRRRKRRRRKKTTKATKEKEDDESDEGERRRRKRRRRKKTTKATKEKED*NYAERRRKRRPTEKSKGENE
Consensus                                   ****  * *                 * **  * ** *                 * ** * ** *                   **    *        *    * *

Teleogryllus_occipitalis        EDEDEED----EDE------------------------------EDEDEEDEDEEDEDEEDE-----------------------------------------------------
Gryllus_longicercus             LGRKLRDLK*VSDEGWRGKGRHVM*LGAAYLRGWTLPSTPPPSPAAEQHTTLQTQ*NEHIEDTSLRTVLSRQQFAFPAFSSTSCHFTALEKNNVRQLTCCAKIMSWQTNYQRAPRMICAG
Consensus                                   *     **                                           *  **

Teleogryllus_occipitalis        -------------------------------------------------------------------DEEDEDEEDED----------------------------EEDE---------
Gryllus_longicercus             SRVEAPKRRTTAFPSSFSKEINKSAVPMN*LSFKLY*LYKTR*ARYKYCNLYSASGIRSLSLTYSFIDPFSECSYDPSLFSLPRVTCSFHSFKRYK*ITLIKKVRVLREDKLQLLRVKPK
Consensus                                                                                      *   *   *                                **

Teleogryllus_occipitalis        -------------------------------------------------------------------------DEEDEDEEDEDEEDED-----------------------------
Gryllus_longicercus             FRRTNTNQSIHFKLCEIISK*T*LHTVVKIAFLV*SLTHYSL*FVSTVLNKHRPLRVGYRSSLFGCDKLVQRYDEK*LTAISSSIEFSDARPRLKTSHTTNHFQRKSRANLNL*LSGSAY
Consensus                                                                                        **          *  *

Teleogryllus_occipitalis        -----------------------------------------------------------------------------------------------------------------------
Gryllus_longicercus             IFPLPLSRGLRVITSNIFLLGILK*LASFNIQYQCGRAGGRAGGRAGGRPGRQSHLDVEPRCELVALARRKTGRFKLAKLNPIFRDFSLNTEYKFLK*N*IPPTPSTSE*HVICERNTL*
Consensus

Teleogryllus_occipitalis        -------------------NERRRRTKMTKDEG*-------------------------------------------------------------------------------------
Gryllus_longicercus             NYARNRVLIIIIIIIIIIRRRRRRRRRRRKQRGTAFARKIAG*PQHYNLLIY*YQRLRSLAP*V*CIFYCRRWR*SSPAG*RNSAPTYPSTNGGFPTQIFPTRGAMSRQRQALRFRLVL
Consensus                                             ****    *  *

Teleogryllus_occipitalis        --------------------------------------------------
Gryllus_longicercus             RHGEWRGLHVRKEGRKEGRREGGKEGRREGGKEGRKEGGKEGRKEGRKEG
Consensus
```

**Tocc.13G0004800.1**

```
Teleogryllus_occipitalis        TTCTTTTTTGCCACAATTATCGATTCTTTGGTTATTGTGCCACTGCATTTAATCCTTAGGAACAACTAATGGTGTAGAAATGCTGGTCCAAGCTAGTCAGAAGAATGCCGGATCGCCCTT
Acheta_domesticus               ----TTTTTTCAGCAATTAACGATTCTTCGGCTATTGTGCCACCGCATTCAATCCTTAGGGACAAGGAATGACGTGGAAAGACGGGT----GCAAAACAGAAGAATGCTTGATCGTCCTT
Gryllus_longicercus             TTCTTTTTTGCAGCAATTAACGATTCTTCGGCTATTATGCCACCGCATTCTATCCTTAGCGACAAAGAATAGCGTAG-AATGATGGTGCAAGCAAGACAGTGGAATGCTGGATCGTCCTT
Consensus                           ***** *  ****** ******** ** **** ****** *****  ********  ****  ***   ** * **    ***    ** *  ***  ******  ***** ****

Teleogryllus_occipitalis        T-GATGTGTATCGTCTTTCATTAGGGTCAGTATCGGGCGCTTAATAAAAGAGCACGTTGCCGCATATAATGAAAATATGCATCATTAATTTCGCTGGTATATTTGCAACCGAAGTACGTG
Acheta_domesticus               TTGACGTGTAACGTCTTTCATTT--GTCATTATCGGTCGCTTATTCAAACAGCACGTTATTGCA---GATGAAAATAAGTATCATTAATTTCGCTGATATATTTGCAACTGTAGTACTTG
Gryllus_longicercus             ----------TTGGCGTATTTTTGAGTCAGTTGCGGTCGCTCACTAAAACAGCATTTTAATGCATATAATGGAAGTATGTATCTCTAATTTCGCTGATATGTTTGCAACCGTAGTTCCTG
Consensus                                 * * *   **   **** *  *** **** * * *** ****  **   ***     *** ** ** * ***  *********** *** ******** * *** * **

Teleogryllus_occipitalis        CGAATCGTATCGTTTGATCCTTGCTGTTGTATTTAGAAAAGAAGAAAT-AACTCGATCTCGCATGAAAATAAAATTACTCTCAAATATCATTGTGCGGTTATAGTTT--TTCTAAAAAAT
Acheta_domesticus               CGAACCGTAGCGTTTGATCCGTCCTCTTGTGTTTAAAAAAGTATAATT-AACTCGATCTCTCATGAAAATAAAATTCCTCTCAAATACCATGGTTCGGTTATAGTTTTGTTCTAAAAAAT
Gryllus_longicercus             CAAACCGTATCGTTTGGTCCTTGCTGTTGGGTTTAGAAAAGAGGAAATAAACTCGATCTCGCATGGCAATAAAATTCCTCTCATGTATCATTGTGCGGTTATAGTAT--TTTCTAAATAA
Consensus                       * ** **** ****** *** * ** ***  **** *****   ** * *********** ****  ********* ******  ** *** ** ********** *  **   *** *

Teleogryllus_occipitalis        TGGCAAATGTTACCCTGTACTTTAATACCTATCCCATTCCTATCGGCAATTTACTAATATGAATTAAATGCTCAAAATATAGCGAATTACTACTGCTCAATTCCACTGTAAAAAATCAAC
Acheta_domesticus               GGTCTAATGTTAC--------------TCTATCCTATCCCTAT---CTAATTACTAATAGTGAATAAATACACAAAATATAGTGAATTCCTACTACCCAATTACACCGTAAAAAAT-TAT
Gryllus_longicercus             TATCAAATGTTACGCTGTACTTTAATACCTATCCCAACCTTATCGTCAAATTACTAATATAGAATAAATACACAAAATACAGCGAATTACTACTGCCCAATTACACTGTAAAAAGT-AAC
Consensus                          * ********              ****** *  * ***   * * *********   * ***** * ******* ** ***** ***** * ***** *** ******* *  *

Teleogryllus_occipitalis        AGCACTACCTGGAATATAAGTTCAGTTTGTAAGTTTTGTGAGGCAGGAAAAAAATGAATTTGCGTATTTGTTTTCA-TTCGTTTCCCAACGCTGTCGCGTCTGCAGCACTCGTACTAATA
Acheta_domesticus               CGCAGTACCCGTAATGTAATTTCAGTTAGTAAGGTTTGTGAGGCGGGAAATAA-TTAATTTGCGTATTTGTTTTTATTTCATTTCCTAAC-CTCACGCGTCTGTAGCATTCCTATTAATA
Gryllus_longicercus             TGCACTACCTGTAATTTAAGTTCAGCTTGTTAGTTTTGTAAGGCCAGAAAAAA---AATTTGCGTGTTTGTTTTCATTTCATTTTCCTAAGCACACGGATCTGCTACATTCGCACTAATA
Consensus                        *** **** * *** *** ***** * ** ** ***** ****  **** **   ********* ******** * *** *** *  *  *   **  ****   ** **  * *****

Teleogryllus_occipitalis        TTCTACAAAGGCTGCGTCCACGGTACCGTCTACTCGAATTTTTTTCTGTTTCTCATAAATGTGTT----------TCTGTACATGACC-------CATTTATATGTTCTATGTACCGCTT
Acheta_domesticus               TTCTACAACGTCTGCATTCACGATACCGCCTAATCGACTTTTTTTCTTTTCCTCATAAATGTGTT----------TTTATACATCTCCCCATCATCATTCGTATATTCTATGTACAGGTT
Gryllus_longicercus             TTCTACAACGGCTGCGTTCACGATATAGCCTATACGAATTTTTTTCTGTTCCT-ACAAATGTGTTTTTACTGTACTTTATACATCACC-------AATTCGTATATTCTTTATTCCGCTT
Consensus                       ******** * **** * **** **  * ***  *** ********* ** ** * *********          * * *****  **        ***  *** **** * * * * **

Teleogryllus_occipitalis        ACGTCTAGTGGTATAGTTCTTTTTACAGCCGTTTTTCAATATCTTTAAAAATGATTTGTAAAATAACTGCAATCCGTAACCAGATGGAGGGAC--AACGTGATGTACTAATT-------A
Acheta_domesticus               ACATCTGGAATTATAGTTCGTTTTATAGCC-TTTTTCAATAACTTTAAAAAGGATTTGTGAAATAAGTGCAATCTGTGACCAGATAGAGGAGCAGAACGTGAAATAATAATT-----ACT
Gryllus_longicercus             ACGTCTAATATTATATTTCGTTTTATAGCCGTTTTCAAAAAACTTTAAAAGGGATTTGTAAAATAACTGCACTCCTTAACCAGATCTAGAGACAGAATGTGTTATACTTATTATTTTACA
Consensus                       ** ***     **** *** ***** **** ****  ** * ********  ******* ****** **** **  * *******  **   *  ** ***   ** * ***

Teleogryllus_occipitalis        ATACGTTGTGAATAAAATAAATACTTTAATTACGAACAACAATGAAACCTCGTGTATTAGCGAATGCTAGGTTTTTAAAATTTTTGATTCTGTGTAATAAGGAAAGTGCGATCTCAAAAT
Acheta_domesticus               TTACATAGTGAATTAAATAAATACTTAAATTATG---AATAAAGGAAACTCGGATACTAGTAAATACTACATTTTT--TATTTTTGTTTCTGTGTAATAATAAAAATGCATTCTGAGAAT
Gryllus_longicercus             TTACATTGTGAATTAAGTAAAAAGTTTAATTATG---AACAATGAAACATAGACTA-TAGTAAATGCTATTTTTTTTAAAAAAATTGTTTCTTTGTAATAAGGAAACAGCGATCTCAAAAT
Consensus                        *** * ****** ** **** * ** ***** *   ** ** * **  * *  ** ***  *** ***  *****   *   *** **** ********  ***  **  *** * ***

Teleogryllus_occipitalis        AATTCCTTCTGGTCCGTCAACTGAATAACTGAAATGAATCGTAGTCGCGCACAATCCGATAGTTTTGTTATGTTGCCAGAAGGTAACCGGAAACTT--TAATGTGTAACTTTTTA-TTTT
Acheta_domesticus               GATTCCTTCTCCTTT-CCAAATTAATAACTGAAGACAATCGAAGTCACGCACAGTCTGATAGTTTTGTTATTTTGTTAAAAGGTAATCGGAAATTT--TAATGCCTAACTCTTTATTTTT
Gryllus_longicercus             GATTCCTTCAGCTCTGTCAACTGAATAACTGAAATGAATCGAAGTCACACTCGGTCCGATAGTTTTGTTATATTGTTAGAAGGTAACCGGAAGCTCTATAATGCGCCACTCTTTA-TTTT
Consensus                        ********   *    *** * **********   ***** **** * * *  ** ************** ***  * ******* *****  *   *****    *** **** ****

Teleogryllus_occipitalis        TGATGAAAGCTTGACCATGCTTCTTAACTCTGCAAATTGATTTCTACTCTAAATTTGTTTCAAATACAAACAAAATCAGCATCTAATGC-----------AAGTGCGTTCACTTC-----
Acheta_domesticus               TGATGAACCCTTGAACATGCTTCTTAACTCCGCAAAGTAATTTCTACTCTAAATTTGTTTCAAATACAAACTAAATCAGAATCAAATACTGTAAGTGCAAATAACTGTTCAGTTCTTCCT
Gryllus_longicercus             TGATGAACGCTTGAACTTGATTCTTTACTCCTCAAATTGATTTCTACTCTAAATTTGTGTCAAATACAAACAAAATTAGCATCAAATACTGAAACTGC-AAGGTCTGTGCAGTTC-----
Consensus                       *******  ***** * ** ***** ****  **** * ****************** ************ **** ** *** *            *     ** ** ***

Teleogryllus_occipitalis        ------TTAACAAATTTCCAACAATTT---------------------------------------------------------------------------------------------
Acheta_domesticus               TCAATGTTAACAAATTTCCAACATTTTCCTGTAATAAGTGCCGACACCAGGCATGGACTGACCGGACGATTGCCCGCGGCCCTGCAAGTCATGGGGCACGATGATGGGACTGATGGAGGA
Gryllus_longicercus             ------TTAACAAATTTCTAACGATTT---------------------------------------------------------------------------------------------
Consensus                             ************ ***  ***

Teleogryllus_occipitalis        ------------------------------------------------------------------------------------------------------------------------
Acheta_domesticus               AAAATTAGTAAAAAATGGAAAAAGAATTTACAACTCGAGTAAATCGTTATCATACTTCTAATTTTATAAGAGTATGTCGCGATACAGTAGTCTGATATTTCATAAAAGAAAGTATTAAAA
Gryllus_longicercus             ------------------------------------------------------------------------------------------------------------------------
Consensus

Teleogryllus_occipitalis        ------------------------------------------------------------------------------------------------------------------------
Acheta_domesticus               AACTAGCTAAACAACAAAATTTTTTAGCGTTTTAAGCTCACCGTAAAAATTGTATTATTTGATCACTAAATACTTTTATTTAAATCTATTTTTAAATATATTTACAAAAAGTTTTTCTAA
Gryllus_longicercus             ------------------------------------------------------------------------------------------------------------------------
Consensus

Teleogryllus_occipitalis        ------------------------------------------------------------------------------------------------------------------------
Acheta_domesticus               CTTAAACATTATAATTTTTGTTATTTTTTGAAATAAATAAGGCATAAAGGTTCTTAATACGACTTTCACATTTTGGGGGGGGGGGCGGGCTTCGTTTCAGCATTGCCCCTAGCCCAGAAT
Gryllus_longicercus             ------------------------------------------------------------------------------------------------------------------------
Consensus

Teleogryllus_occipitalis        ------------------CCAATGATTTCAGATGTCAAAAATCTTATGGGGGCAAAGAGGTTGATGGTTCGCCATTTTTAAACCGAAGATAATTACATGCGCGCCGAAGTTAGTAAACCG
Acheta_domesticus               TTTCTATCTCCAACACTGCCAAAGATTTCAGATGTCAAGAATCTTAT-GGATAAAAGAGGTTGATGGTTCGCCATTTTTAAACTGAAGCTAATTATATGCGTACCGAAGTTAGTAAACCG
Gryllus_longicercus             ------------------CTAGTGATTTCAAATGTCAAGAGTCTTGT-GGAGTAAAGAGGTTGATGGTTCGCCATTTTTAAACCGAAGATAATTACATGCGCGCAGAACTTAGTAAACCG
Consensus                                         * *  ******* ******* * **** * **   ****************************** **** ****** *****  * *** ***********

Teleogryllus_occipitalis        TCCAATAAACCTCTCGAGTTGCTTGTAAGATTCCACGAGAACAATCAAGCATACGATAATTCATTCCAGAAGACTTCCTCATTTGCACAGCAGTCTCCCCCTGCTGCCGCCAGACGCAGA
Acheta_domesticus               TCCAATAAACCTCTCGAGTTCCTTGTAAGATTCCACGAGAACAATCAAGCACACGATAATTCATTCCAGAAGACTTGCTCATTTGCACAGTAGTC-TCCCCCGCTGCCGCCAGGC-----
Gryllus_longicercus             TCCAATAAACCTCTTGAGTTCGTTGTAAGATTCCACGAGAACAATCAAGCATACGATAATTCATTCCAGAAGACTTCCTCATTTGCACAGTAGTC-CCTCCTGCTGCCGCCAGACGCGGG
Consensus                       ************** *****  ***************************** *********************** ************ ****  * ** *********** *

Teleogryllus_occipitalis        GGCGG---GGGCGCGCGCGCCTCACCCTCAGGTTAAAGATGATCCGGCCACCCTGCGCCGGACGGGGTAGGGGTGAAGAATTTACTTTGGACGCAGCGGGGCGGGGCAGAGCGGAGCAGG
Acheta_domesticus               GGCGACGCGGGGGTGCGCGCTGCAGCCTCAGGTTAAAGATGATCCGGAAG-------------------GGGGTGAGGAATTTACTTTGGGCGCAGCGGGGC------------------
Gryllus_longicercus             GGCGACGCGGGGGCGCGCGCTTCAGCCTCAGGTTAAAGATGATCCGGCCGACCTGCGCCGGAAG-----GGGGTGAGGAATTTACTTTGGGCGCAGCGGAGC------------------
Consensus                       ****    *** * ******  ** **********************                      ******* ************* ******** **

Teleogryllus_occipitalis        CGGGCAGGGCAGGGGCGAGGAGGGGCGGGGTGGGGGGCAGGAGCAGGGCAGGGGGGGGGGGGAGGGCAGGAGCAGGGGCAGGAGCAGGGCAGGAGCAGGAGCAGGGCGCAGGAGCAGGGC
Acheta_domesticus               -----------GCGCCAGGCCGGGGCAGGGC-----------------------------------------------------------------------------------------
Gryllus_longicercus             -----------GGGGCGGGAAGGGGCGGGGCAGGGAG-----------------------------------------------------------------------------------
Consensus                                  * * *  *  ***** ***

Teleogryllus_occipitalis        AGGGGCAGGAGGGAGGACAGGGGAGCAGGGGGCAGGAGCAGGAGCAGGAGCAGGGGCAGGGGCTGGGGTAGGAGCAGGGGCAGGAGCAGGAGCAGGGGCAGGGGCAGGAGCAGGAGCAGG
Acheta_domesticus               ------------------------------------------------------------------------------------------------------------------------
Gryllus_longicercus             ------------------------------------------------------------------------------------------------------------------------
Consensus

Teleogryllus_occipitalis        GGCATGAGCAGGGGCAGGGGCGGGAACAGGGGGGGCAGGGGCAGCACATTGGGGGGGCGCGGGCTCCATGAGGCATGAGGCTATCCGTTTTGGTAACGGC-----------------GGC
Acheta_domesticus               ------------------------------------AGGGGCAGCCCAGTGGGGCGGCGCGGGCTCCGTGAGGCATGAGGCTATCCGTCTTGGTAACGGC--------------GGCGGC
Gryllus_longicercus             ------------------------------------AGAGGCAGCCCAGTGGGGCGGCGCGGGCTCCGTGGGGCATGAGGCCATCCGTCTTGGTAACGGCGGGGGCGGGGGCGGGGTGGC
Consensus                                                           ** ****** ** ***** ************ ** ********** ****** ***********                 ***

Teleogryllus_occipitalis        GGAGGGAGGCGCACAGTGGGCCGAAAGA----GTCGGGGGGGAACGACAACAATGTAAGTAAACATTTCCTCAAAATGCGTTTTATTATGGT----------------------------
Acheta_domesticus               GGCGGGCGGCGCACAGTGGGCCGAAAGG----GGCGCGTGGGAACAACAA-AATGGAAGGAAACATTTCC-TCAAATGTGCATTTTTCTTTTGGAAAACAGAAAAATTTTCAAGTATATA
Gryllus_longicercus             CGAGGGCGGCGCAAAGTGGGCCGAGACAACATGCAGCATGGGAACAACAA-AATGTAAGTAAACATTTCC-AAAAATGTGCATTTTTACGGT----------------------------
Consensus                        * *** ****** ********** *      *  *   ****** **** **** *** **********   ***** *  ** **    *

Teleogryllus_occipitalis        ----------------------------------------------AAACAGGAAAAGGTGAATTTTTAGATGTCGATTTTTTTGCACATAATAAAACAAGTTTTCGGATGCAGTCATTT
Acheta_domesticus               ATATTAATAACCATTACCGAACCAATTACCGTTACAGATTACTACAAAACAGGAAAAGTTGCGTAATTAAATGTCGATTCTTTTGCACATATTGAAACAAGTTTTCAGATGCAGTAATTT
Gryllus_longicercus             ----------------------------------------------AGACAGGAAAAAATGAAGTTTTAGATGTCG------------------GATCAAGTTTTCGAATGCAGCCATTT
Consensus                                                                     * *********  **     *** ******                  * *********  ******  ****

Teleogryllus_occipitalis        GCAGTTTATGAATCGACACAAAGCGAAATTCCTT--------------------------------------------------------------------------------------
Acheta_domesticus               GAAGTTTATGAATCGATACGAAGCCTAATTCGTTTAAACAAATCTAAACATTTCGTAAGAAGCGGCAGAGTATTGGACATTAAGAAAAGGCGTAAAAGTAGCGAAAATCATTCTGAAGGT
Gryllus_longicercus             GATTTTTATAAATCCATACGAAGCCAAATTCTTTGAAA-ATACTACAGCATTTCGTACGAAGAGGCG-------------TAAGAAAAGG-GTAAATGTAGCGAAAATCATTCTCTGTGT
Consensus                       *   ***** **** * ** ****  ***** **

Teleogryllus_occipitalis        ---------------CTTAAAAA-------------------------------------------------------------------------------------------------
Acheta_domesticus               AAAACGAAAATATTGTTACAGAAGACATTGAATGGTGATAATCCGTTTCAATAAA-----------AAATTATAAATAAAATAAATGTAATGA---AATGTATTTACCTGGGGCTATTTC
Gryllus_longicercus             GAAATGAAAAGATTGTTTAAAGAGACATTGAAACGTCCTGTGCCATTTGAACAAATATTATAAATGTAATTGATAATGAAGTGTGAATAATGAATGAGTGCATTTAACTGGATCAATTTC
Consensus                                       *  *  *

Teleogryllus_occipitalis        ------------------------------------------------------------------------------------------------------------------------
Acheta_domesticus               --TTTGTATGTTTTATTCATTTTATGATTTCAGTATTTTTTTAACTTTGAAATTTGCCTTTGCTTTAGAA------------TGTTTGCCTGCAAATTGGCCTCATACGTGCCTTTATCG
Gryllus_longicercus             ATTCTGAAGGTTTTATT----------GTTCAGTATTTTTTT--CTTTAAAATTTCCCTTTGCCTTTGAATGAGCCGCTTCCTGCGTGACTGCAAATGAGCTTCACATTTGCGTTTATCA
Consensus

Teleogryllus_occipitalis        ------------------------------------------------------------------------------------------------------------------------
Acheta_domesticus               GATTGTATGTGCGTGTTTAAAATATGCT--------------------------------------------------------------------------------------------
Gryllus_longicercus             GATTATACGTGCGTGTTTAAAATATGCTAAGGGCGGTTTGCCGATCCGGTGGCCGAGATCGCTAAGGCAGTAGTGCCGTAGCGATCTCGTACCCAAAGATCGCGGGTTCGAGTCCCGCCT
Consensus

Teleogryllus_occipitalis        ------------------------------------------------------------------------------------------------------------------------
Acheta_domesticus               ------------------------------------------------------------------------------------------------------------------------
Gryllus_longicercus             CGGGCATGGTTGTATTGTGTTCGTCCTTTCACCCCCTATCATCATCATCATCATATCAGGTTAGGCAACGGGAAACCATCCTGAATAATTTATCCCAGGAAAACCCGTTATGGGTCACCT
Consensus

Teleogryllus_occipitalis        ------------------------------------------------------------------------------------------------------------------------
Acheta_domesticus               ------------------------------------ATGCGCACTTTTGTGCCGTGTCATTTCATTGGGTTGTGCAACTCGGACAATGGATATCTTAT-TGACAAATTCGCGGTCAAACA
Gryllus_longicercus             AACTTCCGTATATTGAAGATAGGAACAAAGAAGAAGAAGGGCGGTTTGGTGCCGTGTCACTCTACTGTGTTGTGCAACCCGCACAATGGGTACCTTATATGACAAATTCGTGGT-GAACC
Consensus

Teleogryllus_occipitalis        ------------------------------------------------------------------------------------------------------------------------
Acheta_domesticus               AACTCGTCGTGTAACTAGGTCTTTTCATTTATCCAATAAAAAAATGCTTGACCCGAACATTCTAGAAAGTAGGCTGACGGAGCGCTTTCGGCCCACTGTCCGGGGCAGGGCGGGCGATCT
Gryllus_longicercus             AATTCGTTGTGTAACTTCGCCTTTTCATTTATCCATT--AAAAATGACTGACCCGAAC-TTGTAGAAAATAGGCTTCGGGTGCCCTTTCGGCCCACTGCCCCGCGCGGGGTAAGCGCTCC
Consensus

Teleogryllus_occipitalis        ------------------------------------------------------------------------------------------------------------------------
Acheta_domesticus               GCTCGCGGAGGGCGCGTGGC----------------------------------------------------------------------------------------------------
Gryllus_longicercus             TCTCGCGGAGGGCGCGGGGCGGGCGGGCGGCGTTATGGCGGCGCATTTGTCATGGAAATCAGATGAGGGCTGCGGGAGCGGCGACTCGCACCTGATGTGGATGGCGTTAAAGAAGAGCGA
Consensus

Teleogryllus_occipitalis        ------------------------------------------------------------------------------------------------------------------------
Acheta_domesticus               ----------------------------------------------------------------------------------------GCAGGGGGGGAGTGGGGCGGG-----------
Gryllus_longicercus             GCGGCTGCGGCGGGGGCGTTCCCCAGGGGCCGCACCGCGTTTGCGCCACGGCGCTGGGCGGCCGCGGGGCTGTGAGGCCTTGCGCCGTGCGGGGGGGGGGGGGGGGGGGGGGGCGCGACG
Consensus

Teleogryllus_occipitalis        ------------------------------------------------------------------------------------------------------------------------
Acheta_domesticus               ------------------------------------------------------------------------------------------------------------------------
Gryllus_longicercus             ACCATGCTCAAATTGTTCCCGGGCCGGCTCTTTCAGGTGTGTCAAAGAGCACGGCGCCTTGAATCTCTCTCTTGTTAATTATTTGTGACATTACCATCGGGGGCAGGGACAGAGAGTTTG
Consensus

Teleogryllus_occipitalis        ------------------------------------------------------------------------------------------------------------------------
Acheta_domesticus               ------------------------------------------------------------------------------------------------------------------------
Gryllus_longicercus             CTGTGTTTTTAATGAACAAATGCTGAAGGTTTATAGAAGATCATGGTTAAAAAGCTTACTTGCTTATTAAAATAGTTAATAAAACTTTATAATATTGGTTACTTATGTCGTACTTTTTCT
Consensus

Teleogryllus_occipitalis        ------------------------------------------------------------------------------------------------------------------------
Acheta_domesticus               ------------------------------------------------------------------------------------------------------------------------
Gryllus_longicercus             CGTATCTTTAATATTTCCTCATCAATTCTTTTTTTTTTACAGGGCAATAAAATACCAAAAAAATCAGCTATAAAAGGTTCTAATTGGATGCGAAAGTAAATATTCCTATTTAATTATTAT
Consensus

Teleogryllus_occipitalis        -------------------------------------------------------------------------------------------------
Acheta_domesticus               -------------------------------------------------------------------------------------------------
Gryllus_longicercus             GTGTGGTGATGAAGGATTGCTTTTCATATTTAGCGCGATAAAAGCCTCGTTTTAGTGCTCTAAATTATAAATATATTAATAACTTATTCGTTTTTTT
Consensus
```

**Tocc.13G0004800.1_exon1**

```
Teleogryllus_occipitalis        ------------------------------------------------------------------------------------------------------------------------
Acheta_domesticus               --------------------------------------------VCSLE------DTI*KKLIPGIKTRASPSLDPSTQRAN*NFFHVVDFYFSLPQFVLKFNVFFLLFLILLYVSRSGH
Gryllus_longicercus             RENVEAESSAHGLGLQ*GGKKPT*IRRHAVSSLVFFYNLH*EHFICS*NVDSGN*DN-------SQYSYSSPSFDPSTLRPH*NLFDVVYFYFSLPQFVPKCKYYYYYYYYYY------
Consensus

Teleogryllus_occipitalis        ------------------------------------------------------------------------------------------------------------------------
Acheta_domesticus               TYKCDNYVKILFPFLSDQL*DPTTTSVT*LF*MMF*HQRFDCISQELGIHPKI*VIFLGWDSHFFEDKVIIYIIVIY*FYSRDFISKS---*GFERNLGIRLSRLYSLGGTVQSLQQVLL
Gryllus_longicercus             ------YYYYYYYYYL*LTHRPTWTTRACTLHLPLSRWQVPVTS------PR--------DQH---------LAGYDFYPPGLLALAVWKIK*****GVK----YAQGGT-----QTRN
Consensus

Teleogryllus_occipitalis        ------------------------------------------------------------------------------------------------------------------------
Acheta_domesticus               LLLLLLLLLLLLLLLLHLHDVIHTHIFF*TTACRFVCKNE*HYSLYTCADISISFLEFYIKFNFFYFSYLILYSCCIF----------------------ISIHVVLIAKEY*EPFRKE
Gryllus_longicercus             LLV---------------------------------RNRYGTTALVISAI*SAYYYYY--YYYYYYYYYCCFSCCYLGILPI*LKSDE*IHPTKLFIRTDISIHRFINCRRVVRACEKG
Consensus

Teleogryllus_occipitalis        -----------------------M-------------------------------------------------KI---------------------------------------------
Acheta_domesticus               KCHVLFA---KEI*VF*FFYFFRMNKTESEIFILST---------*RECFRKLLSTDANNIESSIR-------KI---------CYYVFRHDVIKSCNTLFFLLLINFVTTHFKNRSKR
Gryllus_longicercus             KNNVMFCMSKKDELIF----FPRMNKTESKKLIICTQIALERLFLVKCCGQTLITSRVSLARFAVKF*DIALCRIKYCNPIIMFSCSY*FRSDT----------FLNVCAIHFKNRSRR
Consensus                                              *                                                 *

Teleogryllus_occipitalis        ------------------------------------------------------------------------------------------------------------------------
Acheta_domesticus               RGVC--*LHLTKICYFIIAYISATP*PERLIFVLFCR*EALLPYRVCLECLNDSTENEMTA*RNRMYYTNIASPVLRRLG------RARVDPNKNGPVYEYALYHRRCM*SEAGA---HR
Gryllus_longicercus             RGVR*EKLRLTKICYFIIAYMSATTRVAYFGTVL*IKGDIICAWSA*MTLLEMK*LHDATGEKYSLTPTETSRRIVSRSEQEWPRLQERTQPYK--------------LSEAGALDLNT
Consensus

Teleogryllus_occipitalis        ------------------------------------------------------------------------------------------------------------------------
Acheta_domesticus               *YKF*IFSAIND------SSAIVPPHSILRDKE*RGKT----------------GAKQK-NA--*SSF*RVTSFICHYRSLIQTARYCR*K*VSLISLIYLQL*YLRTVAFDPSSCV*K
Gryllus_longicercus             V*KVFLFCS-N*RFFGYYATAFYP*RQRIA*NDGASKTVECWIVLLAYF*VSCGRSLKQHFNAYNGSMYL*FR*YVCNRSS-------CKPYRLVL-----------AVGF------RK
Consensus

Teleogryllus_occipitalis        ------------------------------------------------------------------------------------------------------------------------
Acheta_domesticus               SIINSISHENKIPLKYHGSVIVL--F*KMV*CYSI----LSLSNY***INTQNIVNSYYPITP*KIIAVPVM*FQLVRFVRREIINLRICFYFIS*P------HA-----SVAFLL----
Gryllus_longicercus             EEINSISHGNKIPLMYHCAVIVFSK*YQMLRCTLIPIPTLS-SNY*YRINTQNTANYYCPITL*KVTALPVI*VQLVSFVRPE--KKFACLFSFHFPKHTDLLHSH*YSTTAAFTI*PIR
Consensus

Teleogryllus_occipitalis        ------------------------------------------------------------------------------------------------------------------------
Acheta_domesticus               IFYNVCIHDTA*STFFLFLINVFLYISPSSFVYSMYRLHLEL--*FVL*PFSITLKRI----------CEISAICDQIEEQNVK**LLYIVN*INT*IMNKGNSDTSKYYIFYFCFCVI
Gryllus_longicercus             IFF--CSYKCVFTVLYTSPIRIFFIPLTSNIIFRFIAVFKKL*KGFVK*LHSLTRSRDRMCYTYYFTLHCELSKKFN-YEQ*NIDYSKCYFFKKIVS--L**GNSD-------------
Consensus

Teleogryllus_occipitalis        ------------------------------------------------------------------KLLSNI------------------------------------------------
Acheta_domesticus               IKMHSENDSFSFPN**LKTIEVTHSLIVLLFC*KVIGN---------FNA*LFIF**TLEHAS*LRKVISTLNLFQIQTKSESNTVSANNCSVLPSMLTNFQHFPVISADTRHGLTGRLP
Gryllus_longicercus             LKMIPS----ALSTE*LK*IEVTLGPIVLLYC*KVTGSSIMRHSLFLMNA*T*FFTPQIDFYS---KFVSNTN--KISIKY*NCKV----CAVLNKFLT-------ISSDFK--------
Consensus                                                                         *  *

Teleogryllus_occipitalis        ------------------------------------------------------------------------------------------------------------------------
Acheta_domesticus               AALQVMGHDDGTDGGKISKKWKKNLQLE*IVIILLIL*EYVAIQ*SDIS*KKVLKN*LNNKIF*RFKLTVKIVLFDH*ILLFKSIFKYIYKKFF*LKHYNFCYFLK*IRHKGS*YDFHIL
Gryllus_longicercus             ------------------------------------------CQES-CGVKRLM-----------------------------------------------------VRH----------
Consensus

Teleogryllus_occipitalis        ------------------------------------------------------------------------------------------------------------------------
Acheta_domesticus               GGGAGFVSALPLAQNFLSPTLPKISDVKNLMDKRG*WFAIFKLKLIICVPKLVNRPINLSSSL*DSTRTIKHTIIHSRRLAHLHSSLPRCRQAATRGCALQPQVKDDP----EGGEEFTL
Gryllus_longicercus             ---------F*TEDNYMRAELSKPS---------------------------NKPLEFVVRFHENNQAYDNSFQKTSSFAQ*SLLLPP-DAGATRGRALQPQVKDDPADLRRKGVRNLL
Consensus

Teleogryllus_occipitalis        ------------------------------------------------------------------------------------------------------------------------
Acheta_domesticus               ---------GAAGRARPGQGRGSPVGRRGLREA*GYPSW*RRRRRAAHSGPKGARGNNKMEGNISSNVHFSFGKQKNFQVYNINNHYRTNYRYRLLQNRKSCVIKCRFFCTY*NKFSDAV
Gryllus_longicercus             WAQRSGAGRGGAGREAAQWGGAGSVGHEAIRLGNGGGGGG-----VAEGGAKWAETTCSMG---TTKCK*TFPKMCIFTV-----------------DRKK*SFRCRI------KFSNAA
Consensus

Teleogryllus_occipitalis        ----------------------------------------------------------------I-------------------------------------------------------
Acheta_domesticus               I*SL*IDTKPNSFKQI*TFRKKRQSIGH*EKA*K*RKSF*R*NENIVTEDIEW**SVS------IKN--YK*NKC---NEMYLPGAISLYVLFIL*FQYFF--NFEICL-----------
Gryllus_longicercus             I*FL*IHTKPNSLKILQHFVRR-----GVRKG*M*RKSFSV*NEKIV*RDIETSCAI*TNIINVIDNEV*IMNECI*LDQFHSEG-------FIV--QYFFL*NFPLPLNEPLPA*LQMS
Consensus                                                                       *

Teleogryllus_occipitalis        ------------------------------------------------------------------------------------------------------------------------
Acheta_domesticus               ----CFRMF---------ACK------------------------------LASYVPLSDC-------------------MCVFKICY---------AHFCAVSFHWVVQLGQWISY*
Gryllus_longicercus             FTFAFIRLYVRV*NMLRAVCRSGGRDR*GSSAVAISYPKIAGSSPASGMVVLCSSFHPLSSSSSYQVRQRETILNNLSQENPLWVT*LPYIEDRNKEEEGRFGAVSLYCVVQPAQWVPYM
Consensus

Teleogryllus_occipitalis        ------------------------------------------------------------------------------------------------------------------------
Acheta_domesticus               QIRGQTNSSCN*VFSFIQ*KNA*PEHSRK*ADGALSAHCPGQGGRSARGGRVA------------------------------------------------------------------Q
Gryllus_longicercus             TNSW*TNSLCNFAFSFIH*K-*LTRTCRK*ASGALSAHCPARGKRSSRGGRGAGGRRYGGAFVMEIR*GLRERRLAPDVDGVKEERAAAAGAFPRGRTAFAPRRWAAAGL*GLAPCGGGG
Consensus

Teleogryllus_occipitalis        ------------------------------------------------------------------------------------------------------------------------
Acheta_domesticus               GGSGA-------------------------------------------------------------------------------------------------------------------
Gryllus_longicercus             GGGGATTMLKLFPGRLFQVCQRARRLESLSC*LFVTLPSGAGTESLLCF**TNAEGL*KIMVKKLTCLLK*LIKLYNIGYLCRTFSRIFNISSSILFFLQGNKIPKKSAIKGSNWMRK*I
Consensus

Teleogryllus_occipitalis        --------------------------------------
Acheta_domesticus               --------------------------------------
Gryllus_longicercus             FLFNYYVW**RIAFHI*RDKSLVLVL*IINILITYSFF
Consensus
```

**Tocc.13G0004800.1_exon2**

```
Teleogryllus_occipitalis        ------------------------------------------------------------------------------------------------------------------------
Acheta_domesticus               -----RLFSRRY---------NLKKIDSRN*DKSF--TFFGSVHSKS---ELKLLPCRGFLFFPPAICTEI*CVFFIIFNIVIRFTIWSHI*VR*LRENTFPVSQ*SAVRPHNY*RDV--
Gryllus_longicercus             GGKMSRQRARRTAWGCSEGEKNLRK----SGDMPFLLLFFFITYTKNISFVLEML--------IPGI--KIIASTHILLLLSIRLR-*DHI-------ETCSMSCISIFLSRNLYRSVNI
Consensus

Teleogryllus_occipitalis        ------------------------------------------------------------------------------------------------------------------------
Acheta_domesticus               ----------------IVLNDVLTSKI--*LHQPGIGYTPEDLSYFSWLGQSLL*G*SDNLYYCYLLILFSGFYLQ------------------------ELRV*AEFRHPTFPTLFTGRN
Gryllus_longicercus             IIIIIIIIIIIIIIIIIIIYDSLIDRLGPRVHAPCI---------YHYLG-----GRCL*LHQEISI*LDTTFILQGC*PWRYGK*SDDDSEG*NMPKAGLKL-AIFWSETATALLP**S
Consensus

Teleogryllus_occipitalis        ------------------------------------------------------------------------------------------------------------------------
Acheta_domesticus               SAITPASFIIIIIIIIIIIIIIIIITST*CNTHPYFL-------LNNGL*VRL*K*MTLFSLYVC*YFNFVFRILHQV*FLLFFIFDFIFVLYFYFHPRCINCQRVLRTFQKGK-MSCFV
Gryllus_longicercus             ---LPSDRLIIIIIIIIIIIIIIIVV----------FLVVI*ASYLSD--*SRMNKFIRRNSLYVPIFQSIVLLIV----------------------------EE**EPVRKEKIMSCFV
Consensus

Teleogryllus_occipitalis        ------------------------------------------------------------------------------------------------------------------------
Acheta_domesticus               CQGNIGFLIFLFF*DEQNRVRDIHTFYVERMF-------S*IAEHGRE*HREFH*KDLLLRFQT*RY-----QILQYFIFSVVN*FRN--DTF*KSIEEKGCLLIAFNENML-----FYH
Gryllus_longicercus             CQRKMN----*FFFQG*TKPSQRN**FVPKLL*RDYF**SAVDRH--**LREFH*LDLLLSFETSRYVESNIAIL**CFLVVINFVVTHF*TFARYI-----LKIALGEGVFVRKNCV*R
Consensus

Teleogryllus_occipitalis        ------------------------------------------------------------------------------------------------------------------------
Acheta_domesticus               RLHLCNPMT--RAPYFRIVL*IRGAITLSCVLGVLK*LY*KRNDCMTQ*DVLYKYSLTCTETSRKSTSRSEQEWSCL*VRTLP*K---MY-VK*SRCSSVVQILNFFSN-*RFFGYCA--
Gryllus_longicercus             KYVILSSLTCLQPPEWPILVPYCK*KATSFVLGVLK*LYLK*NDCMT----LQERNIASPLQRRLGGS*VD-----------PNKNGPVYKSALNRIS*VKQVLLI*TQFKKYFFFAAIN
Consensus

Teleogryllus_occipitalis        ------------------------------------------------------------------------------------------------------------------------
Acheta_domesticus               --TAFNP*G------------QGMTWKDGCKTEECLIVLLTCNVFHLSLSVAYSNSTLLQ-MKISIINFADIFATVVLANRSV*SVLLCLKKYN*LDLS*K*NSSQIPWFGYSFVLKNG
Gryllus_longicercus             DSSAIMPPHSILSDKE*RRMMVQARQWNAGSS---------FWRIFE-SVAVAH*NSILMHIMEVCISNFADMFATVVPANRIVWSLLLGLEK-------RK*TRSRM-----------A
Consensus

Teleogryllus_occipitalis        ------------------------------------------------------------------------------------------------------------------------
Acheta_domesticus               LMLLYPIPI*LL-IVNKYTKYSEFLLPNYTVKNYRSTRNVISVSKV---------CEAGNN*FAYLF-------------LFH--FLTSRVCSIPINILQRLHSRYRLIDFFSFPHKCV
Gryllus_longicercus             IKFLSCIIVRL*YFLNNIKCYAVL---*YLSQPYRQITNIE*IHKIQRITTAQLHCKK*LHYL*FKFSLLVL*GQKKNLRVCFHFIFLSTRICYIRTNILQRLRSRYSLYEFFSVPTNVF
Consensus

Teleogryllus_occipitalis        ------------------------------------------------------------------------------------------------------------------------
Acheta_domesticus               ---FIHLPIIIRIFYVQVTSGIIVRFI--AFFNNFKKDL*NKCNL*PDRGAEREIIITLHSELNKY----LNYE*RKLGY**ILHFLF---LFLCNNKNAF*E*FLLLSKLIT-EDNRSH
Gryllus_longicercus             LLYFIHHQFVYSLFRLR----LILYFVL*PFSKNFKRDL*NNCTP*PDLETECVILIILHYIVN*VKSLIMNNET*T-----IVNAIFLKKLFLCNKETAISK*FLQLCQLNN*NESKSH
Consensus

Teleogryllus_occipitalis        ------------------------------------------------------------------------------------------------------------------------
Acheta_domesticus               AQSDSFVILLKGNRKF*CLTLYFLMNP*TCFLTPQSNFYSKFVSNTN*IRIKYCKCK*LFSSSFNVNKFPTFSCNKCRHQAWTDRTIARGPASHGAR*WD*WRKN**KMEKEFTTRVNRY
Gryllus_longicercus             SVR*FCYIVRR*PEAL*CATLYF**T------LELDSLLLKLISTLNLCQIQ---------------------TKLASNTETARSV-----------------------QFLT-----
Consensus

Teleogryllus_occipitalis        ----------------------------------------------------------------------------------------------------------GRGGGE---------
Acheta_domesticus               HTSNFIRVCRDTVV*YFIKESIKKLAKQQN--FLAF*AHRKNCII*SLNTFI*IYF*IYLQKVFLT*TL*FLLFFEINKA*RFLI----RLSHF-----------GGGGGLRFSIAPSPE
Gryllus_longicercus             ---NF*R--------FLVISNVKSLVE*RG*WFAIFKPKIITCAQNLVN-----------RPINLLSSL-----*DSTRTIKHTIIHSRRLPHLHSSPSCCRQTRGRRGGARFS------
Consensus                                                                                                                             *  **

Teleogryllus_occipitalis        ----------------------------------------------------------------------------------------------------GRSRGRSRAGAGAGRRSRA
Acheta_domesticus               FSISNTAKDFRCQESYG*KRLMVRHF*TEANYMRTEVSKPSNKPLEFLVRFHENNQAHDNSFQKTCSFAQ*SPPLPPGGDAGVRAAASG*R*SGRG*GIYFGRS-GARQAGAGQG--QPS
Gryllus_longicercus             --------------------------------LRLKMIRP-----------------------TC-------------------------AGRG*GIYFGRS-GAGREGAGQGERQPS
Consensus                                                                                                                   *** *    *** *

Teleogryllus_occipitalis        GA-----GGRTG---EQGAGAG--------------------------------------------------------------------------------------------------
Acheta_domesticus               GAARAP*GMRLSVLVTAAAAGG----AQWAER--GAWEQQNGRKHFLKCAFFFWKTEKFSSI*Y**PLPNQLPLQITTKQEKLRN*MSILLHILKQVFRCSNLKFMNRYEA*FV*TNLNI
Gryllus_longicercus             GAARAPWGMRPSVLVTAGAGAGWPRAAQSGPRQHAAWEQQNVSKHFQKCAF---------------------LR*TGKNEVLDVGSS---------FRMQPFDFYKSIRSQIL*KYYSI
Consensus                       **      * *         *  *

Teleogryllus_occipitalis        ------------------------------------------------------------------------------------------------------------------------
Acheta_domesticus               S*EAAEYWTLRKGVKVAKIILKVKRKYCYRRH*MVIIR-----------FNKKL*IK*M-----**NVFTWGYFFVCFIHFMISVFF*L*NLPLL*NVCLQIG--LIRAFIGLYVRV*N
Gryllus_longicercus             SYEEA-----------------*EKGKCSENHSLCEMKRLFKETLKRPVPFEQIL*M*LIMKCE**MSAFNWINFILKVLLF--SIFFFKISL------CL*MSRFLRDCK*ASHLRLSD
Consensus

Teleogryllus_occipitalis        ------------------------------------------------------------------------------------------------------------------------
Acheta_domesticus               MLCALLCRVISLGCATRTMD--------ILLTNSRSNKLVV*L---------------------------GLFIYPIK-----------------------------------------
Gryllus_longicercus             YTCVFKI------C*GRFADPVAEIAKAVVP*RSRTQRSRVRVPPRAWLYCVRPFTPYHHHHHIRLGNGKPS*IIYPRKTRYGSPNFRILKIGTKKKKGGLVPCHSTVLCNPHNGYLI*Q
Consensus

Teleogryllus_occipitalis        ---------------------------------------------AGAGAGAGV-----------GAGAG----------------------------AGAG----------AGAGAGA
Acheta_domesticus               ------KCLTR--------------------TF*KVG*RSAFGPL-SGAGRAICS---------------------------------------------RRARG----------AGGEWGG
Gryllus_longicercus             IRGEPIRCVTSPFHLSIKND*PELVENRLRVPFRPTAPRGVSAPLAEGAGRAGGVMAAHLSWKSDEGCGSGDSHLMWMALKKSERLRRGRSPGAAPRLRHGAGRPRGCEALRRAGGGGGG
Consensus                                                                 ***                                                *          **   *

Teleogryllus_occipitalis        GA*---------------------------------------------------------------------------------------------------------------------
Acheta_domesticus               ------------------------------------------------------------------------------------------------------------------------
Gryllus_longicercus             GARRPCSNCSRAGSFRCVKEHGALNLSLVNYL*HYHRGQGQRVCCVFNEQMLKVYRRSWLKSLLAY*NS**NFIILVTYVVLFLVSLIFPHQFFFFYRAIKYQKNQL*KVLIGCESKYSY
Consensus

Teleogryllus_occipitalis        ------------------------------------
Acheta_domesticus               ------------------------------------
Gryllus_longicercus             LIIMCGDEGLLFIFSAIKASF*CSKL*IY**LIRFF
Consensus
```

**Tocc.14G0001810.1**

```
Teleogryllus_occipitalis      CAGTTCAGAATAA-----------------------------------------------------------------------------------------------------------
Tarbinskiellus_portentosus    TATTTCAAACGAGTCTTGAA----ACGTGTGAAGTTGA----TATTTCCCATTCAGAACATGTTCCCTTTTTCAAACTTCGATTTTTTATGTT-----------------TTTGTACTT
Gryllus_bimaculatus           TGATTCAGGAGAATTTTGAATTATATCTATAAAACCGAGGACTAACTTTCTATAGGATTAACTAGCGTTTGCTAATCTACGTTTTACTCAATTCAACGAGGACGTGACAAGTTGGTAGAA
Consensus                        ****    *

Teleogryllus_occipitalis      -------------------------------------------------------------------------------GAGAACATGGTTACA-------------TAAATGATAATTA
Tarbinskiellus_portentosus    GTAGCCAGAAAACCAAATCTCCAATATCTTCTCAAGTTCTGTTTTTTAA------TTCCTTTACATTTCAAGCCTCTTAAAAATGCTTTAACGAGCCATTCAGTGTTTCAATTTTAACCA
Gryllus_bimaculatus           AAGGCTAAGAAATACAGAAAGTGCTACCTAACTAAAACCTCTATCTTGGAAACATACATGTTGTGCATATAGTTTTTCAAGAATGCTTTAATA-------------TTAACTACAATTG
Consensus                                                                                      * **     * *                * **    **

Teleogryllus_occipitalis      -TTAT-------------AGTGGAGAGGAAGATGA--------------------------------------GTGGAAATTCTATTGAGTCGAGA---------------------
Tarbinskiellus_portentosus    TTAATCTATCGTATTACTGAAATGTTTGGAAGTTGA--CATTGTTTTAGACATCGCATGTGTGTTGCATCGCATCGGCGCTTGCTTACTTATTCAAAATGAATAATGT-----------
Gryllus_bimaculatus           TTTAT-------------GAAGGATCAGAAAATCACTCGTCGTGT-----------TTTCTGTTCTTTTCCATTATCGTAATTTTATTATTTGAAAAGAATTAAATTTTAATTTTCATT
Consensus                      * **                 *    ***  * *                                          *      ** *   *  * *

Teleogryllus_occipitalis      AATTTCTTGGCTACAGTCAAGTAGG-----------------------------------------------------------------------------------------------
Tarbinskiellus_portentosus    GGAGTTTTGAACAAGATTACCAGGACAATTA----AAGTTTCCTTCGAAAACAAATCATTACAAATCTAACACACTGAATAATTTT--------------ATTGTTACACAGTTTCAGT
Gryllus_bimaculatus           AATTTATGAATTATTATTACATGGGCAAATATTGGGAATCTGATATTTGAGTGAACATCGATGAACTGAAAAAACGAAGTAGCTCTTAAAATTGTAGCCAGATCAACCCTCACTTTCAAA
Consensus                         * *     *   * *    *

Teleogryllus_occipitalis      ------------------------TTTGGAGCTAGATGTTTGTGATGCTTTTCCAAT------------------GTATAACAAAG--------------ACAAATGAAATGAAGGAA
Tarbinskiellus_portentosus    GTA------ACAATAGAAAACAATTCAAGATTCAGAGAAATATATTAATATATTGAATATTAAATACGTAT-----ATATAAGAGGGGTGTGATTGGTTTACAGTAAACAAGTTGACAAA
Gryllus_bimaculatus           TTATAATACACAATTTACCATATTTCTAAACCGAAATAATTATCTTATCGTTTTAATTGCGAAGAAAATTTCAGTGGTGCAAAAAAG--------------TGTGATTAGAACAAGAAA
Consensus                                             *    *   * *    * *  *    *    *                    *  ** *  *                   *  *     *  **

Teleogryllus_occipitalis      GTCATCCAT-TGGTACAGTTCTTGGAATTATTTATAAA-----------TCTGTAACAAAAAATTATTAAAATATT-----------------------------ATTGCATTTG---
Tarbinskiellus_portentosus    TTCATTCATGCGTTTCAGTAATTTAAATACTTTTTTATTTTTAAACAATATATTTTAAAGCATAATTCTGCAGTACTTTTTCTGGTTTATTATAAAA-----TGTCAATCATGTTTATTT
Gryllus_bimaculatus           ATTG--CAT-TATTTAATTTCTAGAAGTTTCTTCTATATGCTGAAAGTTGTTAATATCGAAAAAAAAATTCAATATTC----GGGTTGGTTATAAAAGTTGTGGTTAGTTGTATTTACCT
Consensus                      *    ***    *  * *  *   * *   ** *               *   *      * *    *  * ** *                           *    ***

Teleogryllus_occipitalis      ----------------------------------------------------AAAGGGTTACGAGGATT------------------------------AAGAATATATGAACATACC-
Tarbinskiellus_portentosus    TTATGGAAATTAATGTGTTCACACC----------AGCAGTATGAGTTGCTCGAAGAAATATCTGGACTATTC--------------------CGAAGAGCTAGAGAACCGATATATTG
Gryllus_bimaculatus           CTTCTTAAGTGAAAATGTATTCATTTGGCTTCAGAAGTAATTTACAAAGATTAAAGACTTGCAAGGTTTACTCTCAGTGAAACACATGGCATATTGGAGAAACAATATAAAAATGTATTG
Consensus                                                                      ***   *    **  *                                  *    *   *  **

Teleogryllus_occipitalis      ---------------AGATCAAAATATTAGAACATGGTT---------------AGATAAATGTT----AATTA-----TTAAAGCGTAGGGGAAGAAGAAGAGTGA---------
Tarbinskiellus_portentosus    TAAAACGTGG---------------ACTGGAATTTGTTGGGACTATTGTGAGAAAAAATGACTATC--------------TTTGAAATACAG-------TAATAAACAATTG------
Gryllus_bimaculatus           AAAAGTATGGATACGGAGACTATAACACTGAAAAATATTTCATGTAATG---GGAAAAATAAATAGTCTGGAAATTATTTGATTTAAATCGTAGCGTGAGTTGAAAAGTAATTGAATGTA
Consensus                                               * *  **  *  *                * ** * *                  ** **    **         *  *   *

Teleogryllus_occipitalis      --------------------------------AAATTCTATTGA----------GTCGAGAAATTTCTTGGCTATTGTCCAGTAGGTTTAGAGCAGGATGATTGGAATGCATTTCCAA
Tarbinskiellus_portentosus    ---------GAAGAAAGTCACTGTC--------AGGTCTTGTTGGTC--------------------------------------------------AAATAACTGCAGTATTTATTCAA
Gryllus_bimaculatus           GAATGCAATAAAATAATTCATTTTCTCTGAATTGCATTTTATTAATTTCATCTCTGATTATAGTATTTTTTTCCAATAGTACCCTAGATTACTTGTGAAGAAATTTTTGTATTTATTTTA
Consensus                                                  *  * **                                                            * *    *   * *   *

Teleogryllus_occipitalis      ACGTTGGTGAT-----------------ATTTTGAACT------------------------TATTTATTAAA-------------------------------------------
Tarbinskiellus_portentosus    ATATTATTATAAGCATATTCTTACTGCATATTCTTAACGCATA-----------------------TTGCCAAA-CAGCTTAATCGCATATTCGTAACCAGAACAGATTCGTATTTCAA
Gryllus_bimaculatus           CTGACGATAACC-----------TCTAAATTCTAAACTAATAAGCTACTAAACAAAAAATGTGTGTCTACTAAATTGACCCTACAGCTTATCAGTAAGTAGATCATATGCGTATTTAGA
Consensus                            *                     *** * ***                           *   ***

Teleogryllus_occipitalis      -----TCTTTTCTTGA-------------------------------------------------ACTGTTGCTGCAT----TAATATCTCGCTGT----------------------
Tarbinskiellus_portentosus    GAGACTTAACTGATGA---GCAGACGTCCTGATGGCGATAGCGGAAGAGCTAAATGTGCCATCGAGGTCGCAGCAGCGTCTGTGAGGATGTGACCGCAGGAGCGGGACGCGGGAAGCGGG
Gryllus_bimaculatus           ACAACTCGTTTCTTGATAACCAAACGTTCTAAATGGATC-------------ACGAGCCATTCAGACCGCATCCGCATGTGAGGAGATCTCCCAGC----------------------
Consensus                          *    *  ***                                                *   * ** *        ** *  * *

Teleogryllus_occipitalis      ---------------------------------------------------------------------------------------------AGTGCAACA--------------
Tarbinskiellus_portentosus    GGCGGGCGGGCGGGCGGGCGGGCGGGGACAGCGGGGGCAGCGGGGGCAGCGGGGGCAGCGGGGGCAGCGGGGGCAGCGGGGGCAGCGGGGGCAGCGGGGGCAGCGGGGGCAGCGGGGGCA
Gryllus_bimaculatus           ---------------------------------------------------------------------------------------------GGAGCAGTGGGTGTGTGGGG----
Consensus                                                                                                       * ***

Teleogryllus_occipitalis      ------------------------------------------------------------------------------------------------------------------------
Tarbinskiellus_portentosus    GCGGGGGCAGCGGGGGCAGCGGGGGCAGCGGGGGTGTTCTGCACAATGGGCCGCGGCGGCGCGTAATACAGACATCGCTGGCTGCATCGCGATGGAGGCGCGTCGCTTCCTGGCCTATGC
Gryllus_bimaculatus           --GGGACGAGCGGGGGCGGGGGGAGCCCT----GCGTTCTGCA------GCCGAGACGGCGCGTAATACAGACATCGCTTGCTCCATCGCGATCAGGCCGCGTCGCGCCTATTCCTTGTC
Consensus

Teleogryllus_occipitalis      -----------------------------------------------------TTGGTTATGAACCAGGG------------------------------------------------
Tarbinskiellus_portentosus    GCCGCCCTGCGCCTCCCCGCGCCGCCC------------------AGCGCTGCCCCTCACGATCCAGAGCTCCGTAATTAAGGCGATCGGCGGCGGCGGAGGCGGAGAAGTAGGAGGAG
Gryllus_bimaculatus           CCCCCCCCGTCTCTCTCTCTCTTTCTCTCTCTCTCTCTCTCATTGTCGCTCTTCCTCACGACCCAGAGCTTCGTAATTAAGGCGATTAAGGCGATCGGA---GGAGGAGGAGGAGGAG
Consensus                                                                 * * ** **** *

Teleogryllus_occipitalis      ----------------TGCAGATGAAGTT------TT-----------------------------------------------TCTTAGACGTTTTGATTGAAA------------
Tarbinskiellus_portentosus    GAAGCGCCGGAGCGTGTCGCGGACGCGGACGCTGAGTCGAAGCCCGGCGATGCGGCTTGTCGATAGAGGCGATGCATTGCGCCAGA-TGCACGCGGCGTTGCGCGGGAAGCGCGCGATCC
Gryllus_bimaculatus           GAAGCGCCAGAGCGTGTCGCGGACGCGGCT------TCGAAGCCCGGCGATGCAGCCTGTCGATAGAGGCGATGCATTGCGCTGGGAGCCTCGCCGTGCTGCTGGAGGGAGGCGGGAGAC
Consensus                                       ** ** *  *        *                                                *  *   **   *

Teleogryllus_occipitalis      ------------------TCTTCGA-----CAGATGTGATGGGAAACTCTCACGCCGG--CCGGTTTTCGTACTTATGCTTCCAAGAGATCTGGACAACGTGTGGAAGTGGATGAAGAAT
Tarbinskiellus_portentosus    CTCCTCGAAGGAAGCCGCGCTTTAGACGCTCTCGCGTGATGGGAAATTCTCAGGTCAG-TTCCATCCACGTCCTTCTCTTCTTGGGAAAATACGACGGCGCGTGCGA------GAGAGAA
Gryllus_bimaculatus           CTTCCCTCGGAAGACCGCTTTTCGA-----CTGATGCGATGGGGAACTCTCACGACGGTTTCGCCTTTCGTTCTCCTGCTGCCATGAGATCTGGCCGACATGTGAGAGAGGACGAAGGAA
Consensus                                          **        *    * ****** ** ***** * * *   *      *** **  *  *     ** *    * *  *  ***  *      **   *

Teleogryllus_occipitalis      GTGAACACAGAGGGGGGGGGGTAAATGCGGAAAATAAAAATAGATTATACGAGATAGAATATTAGGAATGTACAGAAAATGACACAGCGAGTAT----CAATTCAAAATAGTTATTGGAA
Tarbinskiellus_portentosus    GCAAAAAGAGAGAGGTGG---------------------------AAGGAAGGTAGGAAGGTAGGAAGATAGAGAAAGT-ACGGACTAGGTATGGTGTGAGAGAAAGTAATTGGTAGGA
Gryllus_bimaculatus           GTGAACAGAGAGGGGGAG--------GCGGAAAATAAAAGTAAATTTAAGGAGGTAGAGTATCAGGAATGAAGAGAAAATGAAACACTGAGTAT----TTATTCAAAATAATAAGTAGAA
Consensus                     *  ** * **** **  *                                 ** ***      *****   * ***** * *   *    ****      *   *** ** *   * * *

Teleogryllus_occipitalis      GATGTTC-----------------------------------------------------------------------------------------------------------------
Tarbinskiellus_portentosus    TAAAAGGGTGGTGAGGAAAAAG--------------------------------------------------------------------------------------------------
Gryllus_bimaculatus           GAAGAATGTGAAGAAGAAGAAGAATGTAAAGAAGAGGAAGAAGAATATAAAGAATAGGAAGAAGAAGAACGTGAAGAAAAATGTGAAGAAGAATGTGAAGAAGAAGAGGAAGAATAGGAA
Consensus                      *

Teleogryllus_occipitalis      ------------------------------------------------------------------------------------------------------------------------
Tarbinskiellus_portentosus    ---------------------------------------------------------------------------------------------------------AGAATGGTGGT--
Gryllus_bimaculatus           TGTGAAGAAGAAGAGGAAGAAGAAGAATGTGAAGAAGAGGAAGAAGAAGAATGTGAAGAAGAGGGAGAATAAGAATGTGAAGAAGAGGAAGAATAAGAATGTGAAGAAGAATAATAATGT
Consensus

Teleogryllus_occipitalis      ------------------------------------------------------------------------------------------------------------------------
Tarbinskiellus_portentosus    ---------------------------------------------------------------------------------------------------TGAAGTAGGTTAGGTAGGT
Gryllus_bimaculatus           GAAGAAGGAGAAGAAGAAGAAGAATGTGAAGAAGAAGAATGTGAAGAAGAAGAATAATGTGAAGAAGGAGAAGAAGAAGAAGAAGAAGAAGGTGAAGAAGAAGAAGGTGAAGAAGAAGAAGAAGGT
Consensus

Teleogryllus_occipitalis      ---------------------------------------------------------------------------------------------------------------CAACGGTACA
Tarbinskiellus_portentosus    AGGTAGGTAGGTAGGTAGGTAGGTAGGTAGGTAGGTTGGTAGGTAGGTAGGTAGGTAGGTAGGTAGGTAG---------GTAGGTAGGTAGGTAGGTAGGAAGGAAGGAAGGAAGGAAGG
Gryllus_bimaculatus           GAAGAAGAAGAAGAAGAAGAAGATGAAGAAGAAGAATGTGAAGAAGAAGAAGAATGTGAAGAAGAAGAAGAATGTGAAGAAGAATGTGAAGAAGAA-GAAGAAGAAAAAGAAGA
Consensus                                                                                                                      *  * *

Teleogryllus_occipitalis      GTCTAAAGAATTGAAATCACCATAATGATACCATAAAGTCTGAAATAGAGAAAAC---------------------------ATAACGAAAATGAAAAAAAACTTGAAGT----------
Tarbinskiellus_portentosus    AAGGAAGGAAGGAAG------------------------------------------------------------------GAAGGGGTAATGAGAGGAAATATGTAAA----------
Gryllus_bimaculatus           ATGTAAAGAAGAAGAAGTTCCAGTACAAAAATGTGCGGACTGTAGTAATGAAAATTGCAAAACACCAAAAACTAAAAGAGAAAACAGAAAGATAAAAAAAAACTTAAAATTATTTTTAAA
Consensus                         ** ***                                                                                 ** * *  ***  *  *

Teleogryllus_occipitalis      -ATCTGATGTAATTATTTTTAATATAGAAGTACGAGGTGACAAGGTAAATGACTTTAATTATAGAGGCAATGGAATGTCTGTCATCTAAAGTAACATCGAATATAAATAATTGGAAGACA
Tarbinskiellus_portentosus    ----------------------------------------------------------------------------------------------------GAATATATACAATGATAAAAAA
Gryllus_bimaculatus           AATTAATTAAAATTATTTTTAATGTCAAAGAATGATGTATCCATGTAAGTACATTTAATTATAGAAGCAATGGAATATCTGTTATCTATGGTAATAGGGAACTTATACGTTTAAAAGAC-
Consensus                                                                                                                  ***  ** *   *   ** *

Teleogryllus_occipitalis      TTTCAGTAATACACAGAGATTTACTTGAAACTCCTCCCACAATTATTATTCCATTCTTTTATTAAATGTAAAATACAATCGCAGAAGACAGAGGATATGTATGTGAAAAATAAAGCAGTG
Tarbinskiellus_portentosus    TAAAGGGCAGATAAGGAAAT-------------------------------------------------GAAAATAAATGCAGAGAAGGAGGAATGAGAAAAAGAAAAGAGCAGCGTTG
Gryllus_bimaculatus           TTTCAATAACACACAGTGATTTTCTCGAAACTCCTCCCACAATTAGTATTCCATTCTTTGATTAAATGTAAGAAATAGTGACAGAGGACGGAAGAACTGGATGTGAAAAAGGAAGAAATG
Consensus                     *       * * *  *  **                                                 * * *    ****  *      *   * *   *****    **   **

Teleogryllus_occipitalis      TTGTAATAAATTTGTCTGTAAGCAAGAAGCTTCATTCGTCTCCACGAACAAGAAATTTGATAAAAATTGTGTAATTTTTACCCTCGAAA-CACGACTATATCATC-------TACATAAT
Tarbinskiellus_portentosus    TATTAATA-ATTTATCAAGGAAGCAAAAGTTTCATTTCTCGTTACGGAAAAAAGTTTGGGCTGAAATTATTTTGCTTTGATTCGAAATATCACG--TAATTGATCTTAAATTAACAGAAC
Gryllus_bimaculatus           AACGAATACATTTACCTGTAAATAAGAAGCTTCATTCATCTACACGAACAAGACATTTGATGAAAAATGTGTGTGTTTTACTAAAGAAATCACAACTACATAACCT----ATTACGTGAT
Consensus                         **** ****  *    *   * *** ******  **   *** * ** *  ** *    *** * * *   *** *      * * ***   **  * * *         **   *

Teleogryllus_occipitalis      ----------TATTCACAAC--TGTTTGCCATTTCGAGATCATTAATCTTCTAGTA-CGTAAGAGCGTAAGATGCGTACTAACGTAGAATTAAGAAAATTGACAGAATTTATTGG----
Tarbinskiellus_portentosus    ----------TTTTGGTAATTATTTTTTCCATTGCAAGATCATTAGTCTACTAATTTCCTTCCAGCAAAAGAGAC-----------AGTTCTAAAAATTGAAGACAATCATTAGGACT
Gryllus_bimaculatus           GAATTCTCGTACGTACGTAAT--GTGCTCTTATTTCAGTATCATTATTCTTATCGT---------TCGTAAGATGCATACTTACGTTAAATTACGAAAATTGAAAACATTTATTGA----
Consensus                                 *    **       *   *** *   ******* ***  *  *          *  ****  *           * **   ********    * * ***

Teleogryllus_occipitalis      ---AGCAAATAAAACAAAATGAACTTCCTTGCGCTTAAGGAGTCGAGGCGCAAACTACATCTGGCTACTGGCGAATCGGAATCCAGCCCGATGTTCCTCTCGCCGTCAGAACTGGCCCCG
Tarbinskiellus_portentosus    AATGAATAACAAGACGGCACGGATTTCGCCGTTCGTTAGG-----------ACATTACAACGGAGC-CTGGTGAATCCCGATTCTGTACGAGTTCCCTCTCGCTACCAGGGCTGGCACCG
Gryllus_bimaculatus           ---ATAAAACAAAACAAAATGAACTTCTGCACTCTTAAGGAATTGAGGTGCAAACTACATCTGGCTACCGGCGAATCGGAATCCAGGCCGGTGTTCCTCTCGCCGCCAGAACTGGCCCCG
Consensus                           ** ** **   * * * ***      * * ***           * * **** * *    * ** *****   ** * *  **   * ********   ***   ***** ***

Teleogryllus_occipitalis      AGAAGGCTAGCG-------GCCAGGCTTCGCGCCCGCGGCTTCGGCGGCGGAGCTGGCGGTGGGATCCGGTGCCCAGGCGCCGCCTC---GCCGCCTCCAGCGAGACTCCTGCCGCCGTC
Tarbinskiellus_portentosus    AGGCGGCTAGCCGGCGACCGGCAGGCGCTGCGCCTGCGGCTCCGGCAGCGGAGACGGCGGCGGGATCCGGGGCCCAGAC----------GCCGCCTCCAGCTAGGCTCCCGCTCCCT--
Gryllus_bimaculatus           AGCAGGCTAGCGGCCAAGCGCCAAGCTTTGCGCCCGCGGCTTCGGCGGCGGAGCTGGCGGTGGGATCCGGGGTCCAGGCGCCGCCGCCAAGCTGCCTCCAGCGAGGCTCCCGCCGCCG--
Consensus                     **  *******        * ** **   ***** ****** **** ******  ***** ********* * **** *           ** ********* ** **** **  **

Teleogryllus_occipitalis      GCCACCACCGCCATCGCCACCACCGCCATCGCCACCACCGCCATCGCCACCACCGCCACCGCCACCACCGCCATCGCCACCACCGCCACCGCCACAGCCACCGCCACCGCCACCGCCCCC
Tarbinskiellus_portentosus    ------------------------------------------------------------------------------------------------------------------------
Gryllus_bimaculatus           ------------------------------------------------------------------------------------------------------------------------
Consensus

Teleogryllus_occipitalis      GCCATCGCCACCACCGCCTCCGCCACCGCCATCACCGCCATCACCGCCGTCGCCACCGCCACCGCCACCACCGCCTCCGCCACCGCCATCGCCACCCGCCACCGCCACCGCTACCGCCGC
Tarbinskiellus_portentosus    ------------------------------------------------------------------------------------------------------------------------
Gryllus_bimaculatus           ----------------------------------------------------------------------CCGCCCCNNNNNNNNNNNNNNNNNNNNNNNNNNNNNNNNNNNNNNNNNNN
Consensus

Teleogryllus_occipitalis      CACCACCGCCCCCACGGCCACCACCGCCACCGCCACCGCCACCACGGCCACCACCGCCACCGCCACCGCTACCGCCACCGCCGCCACCACCGCCGCCACCGCCGCCACCACCGCCACCAC
Tarbinskiellus_portentosus    ------------------------------------------------------------------------------------------------------------------------
Gryllus_bimaculatus           NNNNNNNNNNNNNNNNNNNNNNNNNNNNNNNNNNNNNNNNNNNNNNNNNNNNNNNNNNNNNNNNNNNNNNNNNNNNNCACCACCGTCA-------------------------------------------
Consensus

Teleogryllus_occipitalis      CGCCACAGCCATCACCATCTCCGCCGCCACTACCGCCACCACCGCCACCGCCACCGCCACCGCCACCGCCGCCACCGCCGCCACCGCCGCCACCGCCGCCACCGCCGCCACCGCCGCCAC
Tarbinskiellus_portentosus    ------------------------------------------------------------------------------------------------------------------------
Gryllus_bimaculatus           ---------------------------------------------------------------------------------------------------------------------CAC
Consensus

Teleogryllus_occipitalis      CGCCGCCACCGCCGCCACCACCGCCACCGCCACCGCCACCGCCACCGCCACCGCCGCCGCCGCCGCCGCCGCCGCCGCCGCCGTCTCTGCCACAACCACAACCACAGCCACAACTACAGC
Tarbinskiellus_portentosus    --------------------------------------------CTGCTACCGC------------------------------------------------------------------
Gryllus_bimaculatus           CGCCGCCATAGCCGTCCCTGCCGTCACCGCCACCGCCACCGCCACCGCCACCGC------------------------------------------------------------------
Consensus                                                                   * ** *****

Teleogryllus_occipitalis      CACAGCCACAACAACAACCTCAACCACCGCCAACGTCACCGCCACCCCCACCGCCACAGCCACCGCCACCGCCGACGCCAACGTCACCGCCACCACCACCGTCTCCGCCTCCACCGCCAC
Tarbinskiellus_portentosus    ------------------------------------------------------------------------------------------------------------------------
Gryllus_bimaculatus           ------------------------------------------------------------------------------------------------------------------------
Consensus

Teleogryllus_occipitalis      CGCCGACGCCAACGTCACCGCCACCACCACCGTCTCCGCCTCCATCTCAACCACCTCCTCCACGTCAGCCGTCTCTGCCAACACTGCCACCGCGGCCATCGCCGCGATCGCCACCGCCGC
Tarbinskiellus_portentosus    ------------------------------------------------------------------------------------------------------------------------
Gryllus_bimaculatus           ------------------------------------------------------------------------------------------------------------------------
Consensus

Teleogryllus_occipitalis      CATCGCCACGGCTACCGCCGTCACCACCACCACCACCACCACCACCGTCGCCGCCGCGCGCCTCGCGTGTCAGGCCTAATGAATGCTAAGCACGACGCCGCCAGCCGTTCGACCTGT
Tarbinskiellus_portentosus    --------------------------------------------------CGCCGCCGCGCGCCTCGCGTGTCAGGCCTAATGAATGCTAAGCACGACGCCGCCAGCCGTTCGACCTGT
Gryllus_bimaculatus           --------------------------------------------------CACCGCCGCGCGCCTCGCGTGTCAGGCCTAATGAATGCTAAGCACGACGCCGCCAGCCGTTCGACCTGT
Consensus                                                                       * ********************************************************************

Teleogryllus_occipitalis      TGTTGTTTTCCAGTTCACTCAGCCTATAATTTTTATTGTCACGGTAAAGAAAATACCAGCAGCCGCTCCGTCTTCTTGCCCTCCGCCCCGCCTCCTCGTCCTCGTCCTCGTCCTCGCACT
Tarbinskiellus_portentosus    TGTTGTTTTCCAGTTCACTCAGCCTATAATTTTTATTGTCACGGTAAAGAAAATAACAGCAGCCGCTTCGTCTTCTTGCCCTCTGCGCAGCCTCCTCGTCCTCGT--------------
Gryllus_bimaculatus           TGTTGTTTTCCAGTTCACTCAGCCTATAATTTTTATTGTCACGGTAAAGAAAATAACAGCAGCCGCTTCGTCTTCTTGCCCTCCGCGCGGCCTCCTCGTCCTCGC------------CT
Consensus                     ******************************************************* *********** *************** ** * ***************

Teleogryllus_occipitalis      GGCTTCTCGACGCTCCTCTCTCC-CGTGCCCACCCTTCCTTCGGCCGCTCCGCGCGCCGTC----------------------CGCTGTCTCGCTTGCCCTTCTTGCAGCGTTGGATGG
Tarbinskiellus_portentosus    ---CCCCCCCCCTCCTCTCTCCGAGCCCGCTGCGTCCTGCTACCTACCCCGCGCGCTGTCCACCGCGTCGGGAAATGGCCGTCGTCTGTCTCGCTCGCTCTCCACGGAGCGTCCGGAGG
Gryllus_bimaculatus           GGCTCCTCGACGCGCCTCTCTCC-CGCGCCCTCCCTTCGCCGCGCCGCTCCGCGCGCTGTCCGCTGCGCCGA-----------GCGCTGTCTCGCTTGCTCTGCTTGTCGCGTGGGAAGG
Consensus                          * *  * * *********  *  * *  * * *      *  * ******** ***                       ********** ** ** *  *  ****  *  **

Teleogryllus_occipitalis      TCTTCTTCTCTGTCGCTCTTCCG-------------------------------------------------------------------------------------------------
Tarbinskiellus_portentosus    ATGTGTTCTCCCTCGCTCTTCCGCTTCGCCGCTCGCGCGCTGTCCGTCTGCGCGACGGGCGCCGCCTGCTGCTTGTGCGCGTCCCCTCACCTGCTTCGTCGGCCTCTTCGTGCGAGGATC
Gryllus_bimaculatus           TCGTGTTCTCTGTCGCTCTCCCGAGTC-------------------CCGCGCGGC---TACCGCTCACAGCTTGTGCACGTCCAACCGTTCG-----------ATTAGTGAGGAGATC
Consensus                        * *****  ******* ***

Teleogryllus_occipitalis      ------------------------------------------------------------------------------------------------------------------------
Tarbinskiellus_portentosus    GCTGTCGTCCTCGCCT--------CAGAGCGAAATTTTCTCCATCTAGCCGGATGAAGATGCGCTTTGCGGCTGATGTCTCTTTGAGAACGATATTGTTTTTT-------------TAAT
Gryllus_bimaculatus           GCTGACGCCTTCGCCTCTGATTCACAGGAGAAAATCTCCTCTGTTAGG--AGAAAAGCACGAGCTT---GCTTTATTAGTTGTTTAGATTGGCGCTGGTCTTCGTTCACACCCAAGATGC
Consensus

Teleogryllus_occipitalis      ------------------------------------------------------------------------------------------------------------------------
Tarbinskiellus_portentosus    GCTGGATATCACTATACTAAACATTATCACATACCCACTTTACGACAAATTCCAATA--------------------------TAAGAACAAACCATTGTTAATCAGGAAAACTTTTGT
Gryllus_bimaculatus           ACTCGATATT----TACGAATCTTTGTATTCTACGAACTTTGAGAACTATCGCCACACTCTTGGGTAGTGCTGACTGAAAAAACCGAGAAGGGACTAATGTGGATAAACAGGACGTAGGC
Consensus

Teleogryllus_occipitalis      ------------------------------------------------------------------------------------------------------------------------
Tarbinskiellus_portentosus    ATCAATTCACGAAGCTCCGATAATACTTTTCTTTAAAAAGTGTTACGAATATTTAAGAGTGGGGAAATTTTAACACAAAATTCAATTCAAAAATATAGTTTAAAATATTCCATTTTTGAA
Gryllus_bimaculatus           GTC--TTGACGGTTCT----TGATACTTTTTTTCAAAAACTTTTG-----------------------TTCACCCAACAATCCATCCACACTAGCGCCTATAAATGAGCACTT------
Consensus

Teleogryllus_occipitalis      ------------------------------------------------------------------------------------------------------------------------
Tarbinskiellus_portentosus    GATATAATAGTTTTGAATCTGGGAAATATTTTCTATTCATCGCGAATTGTAAGAAACGTTAATGGTACGACTTAAACTAAAAGTAAACATGCAATTGCACCTATCATTGCAAATAACGTC
Gryllus_bimaculatus           ----------------TCCACAAAATAGCTTCTCCCTG--------------------ATCGATATAATTGAACTCGATGCGCAACA---ATTTCAATCTGATGTTTTAA-------T
Consensus

Teleogryllus_occipitalis      ------------------------------------------------------------------------------------------------------------------------
Tarbinskiellus_portentosus    TTTAGGAAGTTTTTGTTGTAGAAACATTTTTTTACATGCCAACAGAAAATATTAATTGCGGCTTCATTCAATTAAGAACATGCTTTAAGGGTCAAGATTCAATTTGTCTGCAGAATCTTT
Gryllus_bimaculatus           TTTTAGAAGTATTTCATGTGAA-----------------------------------------------------------------------------------GTCTAGAACATTTTC
Consensus

Teleogryllus_occipitalis      ------------------------------------------------------------------------------------------------------------------------
Tarbinskiellus_portentosus    ATGACTTAAATGTAAATGACCTTCGCGAATGTAAATGATCCGTAACTATAGTATAGATACAGGATGGAGTAAAAGATGGGACAGTCGCTTACAGATACGGCCCTCAGAAATAATGGAGTT
Gryllus_bimaculatus           AAAATTTTCTTAGAATTGACT-------------------------------------------------------------------------------------------CGAGATT
Consensus

Teleogryllus_occipitalis      ------------------------------------------------------------------------------------------------------------------------
Tarbinskiellus_portentosus    TTTAAAAGACACTTAGCAGATAACGTGTTTTCATACATTATAACAAAATCTTCACTTCTCTGCTCTTGAATCACCTTTCAACCTGTGACACGATCAC----CTGATGGTAAACATTTCTG
Gryllus_bimaculatus           TTT------TCTTTAGAAAACTGTGTACTTGTAAATATGATAAT-ATATCAACATATTTCTTCTTCCTAA--------------------GACTACAGTGCTGAAGAGGAAAATTGTTT
Consensus

Teleogryllus_occipitalis      ------------------------------------------------------------------------------------------------------------------------
Tarbinskiellus_portentosus    AGATCATTCGAAAATTACCGTCACTCCTCCCTTTTCTGAAGGTGGAGAAGACGGAAACATATCTACACATAACAGAATGTCTTTATAAGCGCAAGGTTCACACCATCTTTTTGTGAGGGT
Gryllus_bimaculatus           TAGTCA------------------------TTATTTGTATGTGTGGAGGACGACAG-----------------------------------------------TTCTGACGAGCGA
Consensus

Teleogryllus_occipitalis      ------------------------------------------------------------------------------------------------------------------------
Tarbinskiellus_portentosus    AGGTCTAATTC---ACACACATTGCCCTAAAATAATGATGCATGGAATTTTTATAATTAATGACATTTACGTTCATATCATACAACTATAC---------------------------
Gryllus_bimaculatus           ACGTTTTATTCTGGAAATATTTTGTTCTGAAATA------CATCAAAGTGCTTTCACAAATTTGGTGTGCGTTCGAATTCCAATAAAATACCAAAAAATTCCAAAGCTTCATTCTTGAAA
Consensus

Teleogryllus_occipitalis      ------------------------------------------------------------------------------------------------------------------------
Tarbinskiellus_portentosus    ---ACTAGTGCTACTTTCAAGCTCTTTGTTCACTAGCCCC-------------TCTTTTAAAATCAAGTGAGATGAGAGTC-CATTAAAAAATAATCGTACAGTTGCTTCGGAGAACAA
Gryllus_bimaculatus           ATTACAAATTCAAATCTCCTACTCTTGGTTTTCTGGTAACAGGTTTACGAGAAATATTTGAAGAGTTTGTGGGATTCAAGTTGCAATACTATGATATTATATATTCGTTACAAAAAATAC
Consensus

Teleogryllus_occipitalis      ------------------------------------------------------------------------------------------------------------------------
Tarbinskiellus_portentosus    AGGAAAACTTCATCACTGAATCGC-----ATTCTACAAATCGAGCTGGAGTGAACTCGTCGAACGAATTGTAG----------------------AACGCGTGA-----CGGTCGTTTG
Gryllus_bimaculatus           AAGAAAAATGTGCAGTTGAAGCACCACTAATTTTAT---TTGAGAACAACTTTCTTTATTCATCTAATGATAGTGGTTCAATTACTATCACTTACCTACCCGTAAATGTTCCTTTATTTA
Consensus

Teleogryllus_occipitalis      ------------------------------------------------------------------------------------------------------------------------
Tarbinskiellus_portentosus    GCCTCGAAGAACGGGAGTTTGCCAAACAGCCCGAGCGCTGACCGCTGGACGCTGTCTGCAACATCGAGTGCTTGACGTGAGAAGTGGCGCCAGTCGTA-AGCTGCGATGTTCCT-TCTCC
Gryllus_bimaculatus           ACCTCAGG-------------CGAAATGCTCTGAGGGAGGTTGCAGAATGGCGGCCGGAACGTTGGAACCTGTGCCTTTATCGCGGCGTCTCCCAGACAGCCAAGCAGTTACTGACTCC
Consensus

Teleogryllus_occipitalis      ---------------------------TTTGTCTGCTCC---------------------------------------------------------------------------------
Tarbinskiellus_portentosus    GGCGGGAGTGCCCTCGCTCCGTA----ACTGACTGGTCC---CGTTAAGAGCTGTCAACTTT---ACACTCCATTATGCGTTCGTTCCATCACACACATTACAACTGCATATACTTCACA
Gryllus_bimaculatus           GGCCGTGGTAGCCTAAGCTCTAATTTCTTTATTTGTTCCTATCTTGAATAACTTTTGGTTTTTTGATAACTAGTCATAAGTTTTCTC-----------TGCGTTCTCGTATTTTTT---
Consensus                                                *   ** ***

Teleogryllus_occipitalis      ------------------------------------------------------------------------------------------------------------------------
Tarbinskiellus_portentosus    CCGTTCCGAATATTTTTCTGGCTTTCATATCGTTAATCTATCAGCAAGCGAACGTATGTGTGCTTATGACTTAAAAGCTGGCAGGCATTAACAGATTATAGACTCTGAAGCATCGCTACG
Gryllus_bimaculatus           ------------TATTTTTATTTTTTATTTTTTTTATTTTT------GTGCGCATATTAACAGTCTCAAACCAAACCCTTTCCGGCTT-------CTGTGGAACACCAATCAATGGAAGA
Consensus

Teleogryllus_occipitalis      ------------------------------------------------------------------------------------------------------------------------
Tarbinskiellus_portentosus    TTTTAGGATA---CAGACAGCAACAAATGAAAGAACT---GCGTTTCCCAAAGCAACGTCGAATAC-----------------------------GAACTTTTGCAATTACTTC-----
Gryllus_bimaculatus           TCTCAGAATAAATTAGAATATAACTGAAGGAAAAACTTACTCATTTACGAACCAATCATAAAATGTATTTATAAATATATTACAACCATTTTCAGAAAGTTTCTTTATTTATTTTCAATT
Consensus

Teleogryllus_occipitalis      -----------------------------------------------------------------CCTGTGGCGGGCAC---------------------------------------
Tarbinskiellus_portentosus    -------TCGAGTCAGGCAAAGAATAGCACATACATTATCTATGTAAGCTCGCAATGATTTGTTGTTCTTTTCGAGTTTAAGAAAATAATTAGTTTAAGAAACAAAACGTTTTCGAAAAT
Gryllus_bimaculatus           TTTGTGGTTCAGCAAGGTGATATCTATTTAGTTCATTGTTTGT-------------ACTTGCTATGCTTTCCGCGGGTACACAGATGTCTGA----------GGCATTTTTTTAAATAA
Consensus                                                                                      * *   * *   *

Teleogryllus_occipitalis      ------------------------------------------------------------------------------------------------------------------------
Tarbinskiellus_portentosus    ATAATCCAACTTTTATTACACGTTCTTTGATGTACAATTGGTTTGAATGACAAATGCATATCTCCACA--------------------GTTGTAAGA---------------------
Gryllus_bimaculatus           CTAGGTGAATATTCACTATGTTGTTTCGGTTGTACAACTGTTATGTACAGGTATTAAATAAGATTGCAACATATTTATGTGTAAGATTGGCTGTAAGAGAATGTTGGAGAAAGAGATTAG
Consensus

Teleogryllus_occipitalis      ------------------------------------------------------------------------------------------------------------------------
Tarbinskiellus_portentosus    -----------GTTCACCATCATTTT-----TACTCTATCAAGATCATGGCAAATTGTACACTAATTATTTATGACGTACATTGAAAATTTATATTTATAATTTGATTCCTGGAAACATC
Gryllus_bimaculatus           GCATGGCGAAGGTTTCAAATTATTTTATAAATATTGGACTAGAATTGAGAGGAATGCATCACTCA-GAGTTAGGAAATATTTCAAAAATAAAACT------TCTAAACGTTGGAAAAGTA
Consensus

Teleogryllus_occipitalis      ------------------------------------------------------------------------------------------------------------------------
Tarbinskiellus_portentosus    GTTGACCAT-----------AGGAATAAAGAAAACTTTG--------------GACGATTATAAAATAACCAGAC---AATTTGATTTTAATTACTTTCGAATTAGGTTTTTTTGAGGG
Gryllus_bimaculatus           ACTGATTACCTAATGAAAAAGAGAGAAAAAACAACTTTGTGTGCCAAAGTTGATGGCCTCTTCAATACTACGAGACTTTAACTTCATCTTTTTCGTTTCCCATATTTTTACTTTTCATAC
Consensus

Teleogryllus_occipitalis      -------------------------------------------------------------------GGCCACC------------------------------------------
Tarbinskiellus_portentosus    GGGTAGAAGATTT-----------CATTTTAAATA--------ATGATAACTGTTTGGAGAGATTTCTTGACTGCAATGTTTAAAAAATAAAAGTCCATTAGAATATAATTTTAGGTCTT
Gryllus_bimaculatus           TCGCAAACATTTTAAATATTTAGACATTGTAAATATAACAGTTGTAATAAATGAACAAAAA-----TTTGGCTGTCATAT----ACGACAGAGTTCTAC----AAATAACTTGGGG----
Consensus                                                                                      * *

Teleogryllus_occipitalis      -------------------
Tarbinskiellus_portentosus    TCACGTACGAAAA------
Gryllus_bimaculatus           ---GGACGAAAAAATCCC
Consensus
```

**Tocc.14G0001810.1_exon1**

```
Teleogryllus_occipitalis          ------------------------------------------------------------------------------------------------------------------------
Tarbinskiellus_portentosus        GATLTLFQTSLETCEVDISHSEH-------------------------VPFFKLRFFMFLYL*PENQ--------ISNIFSSSVF*FLYIS------SLLKML*RAIQCF---------
Gryllus_bimaculatus               -LQ*KLF*GQRNVYAVPASFTKNAKI*LDDSGEF*IISIKPRTNFL*D*LAFANLRFTQF------NEDVTSW*KRLRNTESAT*KLPLSWKHTCCAYSFSRML*Y*LQLFMKDQKITRR
Consensus

Teleogryllus_occipitalis          ------------------------------------------------------------------------------------------------------------------------
Tarbinskiellus_portentosus        ---NFNH*SIVLLKCLEVDIVLDIACVLHR--IGACLLIQNE*CGVLNKITRTIKVSFENKSLQI*HTE*FYCYTVSV*Q*KTIQDSEKYINILNIKYVYIRGV*LVYSKQVDKFIHAFQ
Gryllus_bimaculatus               VFCSFPLS*FYYLKRIKF*FSLIYELLLHGQILGI*YLSEHR*T---EKTK*LLKL*PDQPSLSNYNTQ----FTIFL-------NRNNYLIVLIAKKISVVQKSVIRTRK----LHYLI
Consensus

Teleogryllus_occipitalis          ------------------------------------------------------------------------------------------------------------------------
Tarbinskiellus_portentosus        *FKYFFIFK--------QY---ILKHNSAVLFLVYYKMSI--MFIF---MEINVFTPAV*VARRNIWTIPKS*RTDI----------------L*NVDWNLLGLL*EKMTIFEIQ**TI
Gryllus_bimaculatus               SRSFFYMLKVVNIEKKIQYSGWL*K-----LWLVVFTSS*VKMYSFGFRSNLQRLKTCKVYSQ*NTWHIGETI*KCIEKYGYGDYNTEKYFM*WEK*IVWKLFD-LNRSVS*KVIECRMQ
Consensus

Teleogryllus_occipitalis          ------------------------------------------------------------------------------------------------------------------------
Tarbinskiellus_portentosus        GRKSLS----GLVGQITAVFIQILL*AY---SYCIFLTHIAKQLNRIFVTR---------------------TDSYFKRLN**ADVLMAIAEELNVPSRSQQRL*GCDRRSGTREAGAG
Gryllus_bimaculatus               *NNSFSLNCILLISSLIIVFFSNSTLDYL*RNFCIYFTDDNL*ILN**ATKQKMCVY*IDPTAYQ*VDHMRI*NNSFLDNQTF*MD---------HEPFRPHPHV----RRS---PSGAV
Consensus

Teleogryllus_occipitalis          ------------------------------------------------------------------------------------------------------------------------
Tarbinskiellus_portentosus        GRAGGRGQRGQRGQRGQRGQRGQRGQRGQRGQRGQRGQRGQRGQRGQRGCSAQWAAAARNTDIAGCIAMEARRFLAYAPPCASPRRPALPLTIQSSVIKAIGGG----GGGEVGGGSAGA
Gryllus_bimaculatus               GVWGGRA-------------------------------------GAGGALRSAAETARNTDIACSIAIRPRRAYSLSPP-----RLSLSFSLSLSLIVALPHDPELRN*GD*GDRRR--
Consensus

Teleogryllus_occipitalis          ------------------------------------------------------------------------------------------------------------------------
Tarbinskiellus_portentosus        CRGRGR*VEARRCGLSIEAMH------CARCT-RRCAGSARSLLEGSRALDALA*WEILRSV-------PSTSFSSWENTTARAREKQ-------------------------------
Gryllus_bimaculatus               -RRRRKRQSVSRTRLRSPAMQPVDRGDALRWEPRRAAGGRRETFPRKTAF-RLMRWGTLTTVSPFVLLLP*DLADM*ERTKEVNREGEAENKSKFKEVEYQE*RENETLSIYSK**VEEE
Consensus

Teleogryllus_occipitalis          ------------------------------------------------------------------------------------------------------------------------
Tarbinskiellus_portentosus        -------------------------------------------------------KERGGRKVGR*EDRESTD*VWCER-----------K*LVG*KGGEEKENGG*SRLGR*VGR*VG
Gryllus_bimaculatus               CEEEEECKEEEEEYKE*EEEEREEKCEEECEEEEEE*ECEEEEEEEECEEEEEEECEEEGE*ECEEEEE*ECEEE**CEEGEEEECEEEECEEEE*CEEGEEEEEEGEEEEGEE------
Consensus

Teleogryllus_occipitalis          ------------------------------------------------------------------------------------------------------------------------
Tarbinskiellus_portentosus        R*VGRLVGR*VGR*VGR*VGR*VGRKEGRKEGRKEGRGNERKYVKNIYNDKK*RADKEMKINAEKEE*EKE--KSSVVLI----------------------IYQGSKSFISRYGKKFGL
Gryllus_bimaculatus               ------------------------EEEGEEEEEEEDEEEECEEEEEECEEEEEEECEEE----CEEEEEEKEECKEEEVPVQKCADCSNENCKTPKTKRENRKIKKNLKLFLKIN*NYF*C
Consensus

Teleogryllus_occipitalis          ------------------------------------------------------------------------------------------------------------------------
Tarbinskiellus_portentosus        KLFCFDSKY-------------------------HVIDLKLTELLVIIF---------------SIARSLVY*FPSSKR-DSSKN*RQSLGLMNN-----KTARISPFVR--------
Gryllus_bimaculatus               QRMMYPCKYI*L*KQWNICYLW**GTYTFKRLSITHSDFLETPPTISIPFFD*M*EIVTEDGRTGCEKGRNERIHLPVNKKLHSSTRTRH---LMKNVCVLLKKSQLHNLLRDEFSYVRN
Consensus

Teleogryllus_occipitalis          ------------------------------------------------------------------------------------------------------------------------
Tarbinskiellus_portentosus        ------------------------------------TLQR---------------SLVNPDSVRVPSRYQGWHRGG*PATGRRCACGSGSGDGGGIRGPDAASS*APA---------
Gryllus_bimaculatus               VLLFQYHYSYRS*DAYLR*ITKIENIY*IKQNKMNFCTLKELRCKLHLATGESESRPVFLSPPEL-APSRLAAKRQALRPRLRRR----SWRWDPGSRRRRQAASSEAPAAAAPXXXXXX
Consensus

Teleogryllus_occipitalis          ----------------------------------------------------------------M-------------------------------------------------------
Tarbinskiellus_portentosus        -----------------------------------------------PSATAAAARLACQA**MLSTTPPAVRPVVVFQFTQPIIFIVTVKKITAAASSSCPLRSLLVL---------
Gryllus_bimaculatus               XXXXXXXXXXXXXXXXXXXXXXXXXXXXXXXXXHHRHTAAIAVPAVTATATATATATAARLACQA**MLSTTPPAVRPVVVFQFTQPIIFIVTVKKITAAASSSCPPRGLLVLAWLLDAPLSR
Consensus                                                                                           *

Teleogryllus_occipitalis          ------------------------------------------------------------------------------------------------------------------------
Tarbinskiellus_portentosus        -VPPPSSLRARCVLLPTPRAVHRV-----GKWPSSVSLALHGASGGCVLPRSSASPLARCPSARRAPPAACARPLTCFVGLFVRGSLSSSPQSEIFSI*PDEDALCG*CLFENDIVFLML
Gryllus_bimaculatus               ALPSPRR-SARC---PLRRALSRLLCLSRGKVVFSVAL---------------------PSPARLPLTACARPTV----RLVRRSLTPSPL-----IHRRKSPLLGE---KHELALLVV
Consensus

Teleogryllus_occipitalis          ------------------------------------------------------------------------------------------------------------------------
Tarbinskiellus_portentosus        DITILNIITYPLYDKF--------------------QYKNKPLL-IRKTFVSIHEAPIILFFKK-CY-----------EYLRVGKF-------*HKIQFKNIV*NIPFLKI**F*IWEI
Gryllus_bimaculatus               *IGA-GLRSHPRCTRYLRIFVFYEL*ELSPHSWVVLTEKTEKGLMWINRT*AS*RF--LILFFKNFCSPNNPSTLAPINEHFPQNSFSLIDIIELDAQQFQS---DVLIFRSISCEV*NI
Consensus

Teleogryllus_occipitalis          ------------------------------------------------------------------------------------------------------------------------
Tarbinskiellus_portentosus        FSIHREL*ETLMVRLKLKVNMQLHLSLQITSLGSFC-CR--NIFLHANRKY*LRLHSIKNML*GSRFNLSAESL*LKCK*PSRM*MIRNYSIDTGWSKRWDSRLQIRPSEIMEFLKDT*Q
Gryllus_bimaculatus               FKIFLELTRDFFFR------------------KLCTCKYDNI---STYFFFLRLQC*RGKLF*SLFVCVEDDSSDERTFYSGNILF*NTS----------------------------
Consensus

Teleogryllus_occipitalis          ------------------------------------------------------------------------------------------------------------------------
Tarbinskiellus_portentosus        ITCFHTL*QNLHFSALESPFNL*HDHLMVNISEIIRKLPSLLPFLKVEKTETYLHITECLYKRKVHTIFL*G*V*FTHIALK**CMEFL*LMTFTFISYNYTLVLLSSSLFTSPSFKIK*
Gryllus_bimaculatus               -KCFHKF--GVRSNSNKIPKNSKASFLKITNSNLL-----LLVFW-*QVYEKYL---KSLWDSSCNTMIL------------------------YIRYK-------------------
Consensus

Teleogryllus_occipitalis          ------------------------------------------------------------------------------------------------------------------------
Tarbinskiellus_portentosus        DESPLKNNRTVASENKGKLHH*IAFYKSSWSELVERIVERVTVVWPRRTGVCQTARALTAGRCLQHRVLDVRSGASRKLRCSFSGGSALAP--*LTGPVKSCQLYTPLCVRSITH---IT
Gryllus_bimaculatus               -----KYKKNV------QLKHH*FYLRTTFFIHLMIVVQLLSLTYP------*MFLYLTSGEML*GR---LQNGGRNVGTCAFIAASPRQPSSY*LRPW*PKL*FLYLFLS*ITFGFLIT
Consensus

Teleogryllus_occipitalis          ------------------------------------------------------------------------------------------------------------------------
Tarbinskiellus_portentosus        TAYTSHRSEYFSGFHIVNLSASERMCAYDLKAGRH*QIIDSEASLRFRIQTATNERTAFPKATSNTNFCNY-FSSQAKNSTYIIYVSSQ*FVVLFEFKKIISLRNKTFSKI*SNFYYTFF
Gryllus_bimaculatus               SHKFSLRSRIFLFLFFIFF-----------------------IFVRILTVSNQ--------TLSGFCGTPINGRSQNK--LEYN*RKNLLI---YEPIIKCIYKYITTIFRKFLY-LF
Consensus

Teleogryllus_occipitalis          ------------------------------------------------------------------------------------------------------------------------
Tarbinskiellus_portentosus        DVQLV*MTNAYLHSCKSSPSFLL------------YQDHGK-----------LYTNYL*RTLKIYIYNLIPGN----------------IVDHRNKE------------NFGRL------
Gryllus_bimaculatus               SIFVVQQGDIYLVHC----LYLLCFPRVHRCLRHFFK*LGEYSLCCFGCTTVMY-RY*IRLQHIYV*DWL*ENVGERD*AWRRFQIIL*ILD*N*EECITQS*EIFQK*NF*TLEK*LIT
Consensus

Teleogryllus_occipitalis          ----------------------------------------------------------------------------------------
Tarbinskiellus_portentosus        ----*NNQTI*F*LLSN*VFLRGVEDF----------------------------ILNNDNCLERFLDCNV*KIKVH*NIILGLSRTK
Gryllus_bimaculatus               **KREKKQLCVPKLMASSI----LRDFNFIFFVSHIFTFHTRKHFKYLDIVNITVVIN-----EQKFGCHIRQSSTNN---LGGRKNP
Consensus
```

**Tocc.14G0001810.1_exon2**

```
Teleogryllus_occipitalis        -------------------------APPRRLQR---------------------------------------------------------------------------------------
Tarbinskiellus_portentosus      ----FVRERPKIIF*WTFIF*T---LQSRNLSK------------------------QLSLFKMKSSTPLKK--------------T*FESN*NQIVWLF-------YN---RPKFSL
Gryllus_bimaculatus             YNRNYFKGKE---------MCTQCQLPSRKTQRFNWMIQENFELYL*NRGLTFYRIN*RLLIYVLLNSTRT*QVGRKG*EIQKVLPN*NLYLGNIHVVHIVFQECFNINYNCL*RIRKSL
Consensus                                                *

Teleogryllus_occipitalis        ------------------------------------------------------------------------------------------------------------------------
Tarbinskiellus_portentosus      FLWSTMF--------PGIKL*I*IF--------------------------------NVRHK*LVYNLP*S**SKNDGELLQL---------W-RYAFVIQTNC---------------
Gryllus_bimaculatus             VVFSVLFHYRNFII*KELNFNFH*FMNYYYMGKYWESDI*VNIDELKKRSSS*NCSQINPHFQIIIHNLPYF*T-----EIIILSF*LRRKFQWCKKV*LEQENCII*FLEVSSIC*KLL
Consensus

Teleogryllus_occipitalis        ------------------------------------------------------------------------------------------------------------------------
Tarbinskiellus_portentosus      -TSKNV**KLDYIFENVLFLKLI----IFLNSKRTTNHCELT*IM----------------------------------------YVLFFA*LEK*LQK------------FVFDVAL
Gryllus_bimaculatus             ISKKKFNIRVGYKSCG*LYLPLLK*KCIHLASEVIYKD*RLARFTLSETHGILEKQYKNVLKSMDTETITLKNISCNGKNK*SGNYLI*IVA*VEK*LNVECNKIIHFL*IAFY*FHL*L
Consensus

Teleogryllus_occipitalis        ------------------------------------------------------------------------------------------------------------------------
Tarbinskiellus_portentosus      G-----------NAVLSFVAVCILKRSDA-----------------SESIIC*CLPAFKS*AHIRSLADRLTI*KPEKY----SERCEVYAVVMCVMERTHNGV*S*QLLTG-------
Gryllus_bimaculatus             *YFFPIVP*ITCEEIFVFILLTITSKF*TNKLLNKKCVSTKLTLQLISK*IIC----VFRTT--------RFLITKRSKWITSHSDRI-----------RM*GDLPAEQWVCGGDERGRG
Consensus

Teleogryllus_occipitalis        ----------------------------------------------------------------------DSCR---------------------------------------------
Tarbinskiellus_portentosus      -PVSYGAR------ALPPEKEHR-----------------SLRLAPLLTS-----------------STRCCRQRPAVSARAVWQTPVLRGQTTVTRSTIRSTSSLQLDL*NAIQ-**
Gryllus_bimaculatus             EPCVLQPRRRVIQTSLAPSRSGRVAPIPCPPPVSLSLSLSLSLSLFLTTQSFVIKAIKAIGGGGGGGSARACRGR-GFEAR----------RCSLSIEAMHCAGSLAVLLEGGGRPSL
Consensus                                                                                **

Teleogryllus_occipitalis        --------------------------------RRHHRHRHHRH----------------RH-------------------HRHRHHRHRHHRH----------------RHHRHRHSH
Tarbinskiellus_portentosus      SFPLFS----EATVRLFFNGLSSHLILKEGLVNKELESSTSV*LYDMNVNVINYKNS---MHHYF-------RAMCVN*TYPHKKMV*TLRL*RHSVMCRYVSVFSTFRKGRSDGNFRMI
Gryllus_bimaculatus             GRPLFD*CDGELSRRFRLSFSCCHEIWPTCERGRRK*TERGRRKIKVNLRR*SIRNEEKMKH*VFIQNNK*KKNVKKKKNVKKRKKNIKNRKKKNVKK----NVKKNVKKKRKNRNVKKK
Consensus                                                                *                            *                    *

Teleogryllus_occipitalis        RHRHRPRHRHHR-LRHRHHRHHRRRHRH--------------------------------------------------------------------------------------------
Tarbinskiellus_portentosus      SEMFTIR*SCHR-LKG-DSRAEK*RFCYNV*KHVIC-----------*VSFKNSIISEGRICKRL--------------------------------------------------------S
Gryllus_bimaculatus             RKKKNVKKRKKKNVKKRENKNVKKRKNKNVKKNNNVKKEKKKNVKKKNVKKKNNVKKEKKKKKKVKKKKVKKKKKVKKKKKKKMKKKNVKKKKKNVKKKKKNVKKNVKKKKKKKKNVKKK
Consensus                                             *

Teleogryllus_occipitalis        -------------------RHHRLRHR---------------------------------------------------------------------------------------------
Tarbinskiellus_portentosus      HLLLHPVSIL*LRIIYIREGHLHLSHKDSADKLNLDP*SMFLIE*SRN*YFLLACKKMFLQQKLPKDVICNDRCNCMF--TFSL-SRTINVSYNSR*IENISQIQNY-YIFKNGIF*TIF
Gryllus_bimaculatus             KFQYKNVRTVVMKI----AKHQKLKEKTER*KKT*N-------------YF----------*KLIK-IIFNVKE*CIHVSTFNYRSNGISVIYGNRELIRLKDFQ*HTVIFSKLLPQLVF
Consensus                                         *  *

Teleogryllus_occipitalis        -----------------------------------HRHPP--------------------------------------------------------------------------------
Tarbinskiellus_portentosus      LN*ILC*N------------------FPTLKYS*HFLKKSIIGAS*IDTKVFLINNGLFLYWNLS*SGYVI--MFSIVISSIKKTISFSKRHQPQSASSSG*MEKISL*GEDDSDPRTK
Gryllus_bimaculatus             HSLIKCKK**QRTEELDVKKEEMNEYIYL*IRSFIHLHEQDI**KMCV---FY*RNHNYITYYVMNSRTYVMCSYFSIIILIVRKMHTYVK------------LRKLKTFIE*N---KTK
Consensus                                                          *

Teleogryllus_occipitalis        --------------------PPPLPPPPPPPR-PPPPPPPPPRP----------------------------------------------------------------------------
Tarbinskiellus_portentosus      RPTKQVRGRAQAAGGARRA--DGQRASGEAEERGRTHPPDAPWRASETDDGHFPTRWTARGVGSRTQRARREEGGGTRTRRLRRGQEDEAAAVIFFTVTIKIIG*VNWKTTTGRTAGGVV
Gryllus_bimaculatus             *TSALLRN-----*GANYIWLPANRNPGRCSSR-RQNWPRAG*RPS--------AKLCARGFGGGA-------GGGIR------------------------------------------
Consensus                                                *     *    *

Teleogryllus_occipitalis        -------------------------PPPPPPLPPPPPPPPPP----PPPPPPP-------------------------------------------------------------------
Tarbinskiellus_portentosus      LSIH*A*HARRAAAAVAEGAGA*LEAASGPRIPPPSPLPEPQAQRLPVAG*PPRCQPW*REGTRTESGFTRLRCNVLTNGEIRAVLLFISPNDCLQFLELSLLLEGN**TNDLAMEKIIT
Gryllus_bimaculatus             -----------------------GPGAAAKLPPARLPPPP--------------------------------------------------------------------------------
Consensus                                                **    * *

Teleogryllus_occipitalis        ---------------------------------------------------------------------------PQPSPSPPPLPPPPPPPPPPPPPPPPPPPPPPPPPPPPPPPPP
Tarbinskiellus_portentosus      KSSVNLRSIT*YFESKQNNFSPNFFP*REMKLLLP**IINTTLLFSFSHSSFSAFIFISLSALYFLSLYIFFTYFLSLPLPSFLPSFLPSFLPTYLPTYLPTYLPTYLPTNLPTYLPTYL
Gryllus_bimaculatus             ------------------------------------------------------------------------------PXXXXXXXXXXXXXXXXXXXXXXXXXXXXXXXXXXXXXTTVT
Consensus                                                                                    *

Teleogryllus_occipitalis        PPPPPPPPPPPPPPPPPP-------------------------PP--------------------------------------------PPPSLPQPQPQPQLQPQP----------
Tarbinskiellus_portentosus      PTYLPNLLQPPFSFSSPPFYPTNYFLSHHT*SVLSLSSYLPTFLPPLSFCFSLARAVVFSQEEKDVDGTDLRISHHARASKARLPSRRDRALPAQRRVHLAQCIASIDKP----------
Gryllus_bimaculatus             PPP*PSLPSPPPPPPPPPP-------------------------PP--------------------------------------------PRASRVRPNEC*ARRRQPFDLLLFSSSL
Consensus                       *   *    **     **                         **                                                *                 *

Teleogryllus_occipitalis        -----------QQQP-------QPPPTSPPPPPP--------------------------------QPPPPPTPTSPP-------PPP-----------------------SPPP----
Tarbinskiellus_portentosus      ------HRRASTQRPRPRHAPALPPPTSPPPPPPIALITELWIVRGSAGRRGEAQGGA*ARKRRASIAMQPAMSVLRAAAAHCAEHPRCPCP-RCPRCPRCPRCPRC----PRCPRCPR
Gryllus_bimaculatus             SL*FLLSR*RK*QQPL-RLLALRAASSSSPGSST--------------------------------RLSPAPSLRRAA-------PRAVRCAERCLACSACRVGRSCSLSLSRVPRGYR
Consensus                                   * *             * *                                      *                *                             *

Teleogryllus_occipitalis        -------P------------------------------------------------------------------PPPT-------------------------PTSPP---------
Tarbinskiellus_portentosus      C------P------------------------------------------------------------------RCPRCPRCPRCPRP----------------PARPP---------
Gryllus_bimaculatus             SQLVHVQPFD**GDR*RLRL*FTGENLLC*EKSTSLLY*LFRLALVFVHTQDALDIYESLYSTNFENYRHTLG*C*LKKPRRD*CG*TGRRRLDGS*YFFSKTFVHPTIHPH*RL*MSTF
Consensus                              *                                                                    *                          *    *

Teleogryllus_occipitalis        ----PPPS----------------------------------------------------------------------------------------------------------------
Tarbinskiellus_portentosus      ----APAS-RVPLLRS------------HPHRRCCDLDGTFSSSAIAIRTSAHQLSLLKYESVLVTNMRL---------------SCLA-------------------------------
Gryllus_bimaculatus             HKIASP*SI*LNSMRNNFNLMF*FLEVFHVKSR------TFS------KFS*N*LEIFSLENCVLVNMIIYQHISSS*DYSAEEENCFSHYLYVWRTTVLTSERFILEIFCSEIHQSAFT
Consensus                            * *

Teleogryllus_occipitalis        -------PPPSQPPP----------------------------------------------------------PRQPSLPT----------LPPR------PSPR-------------SPP
Tarbinskiellus_portentosus      --ICVK-------------NMQ*EYAYN----------NI----*INTAVI-------------------------------------------*PTR--------------------------
Gryllus_bimaculatus             NLVCVRIPIKYQKIPKLHS*KLQIQISYSWFSGNRFTRNI*RVCGIQVAIL*YYIFVTKNTRKMCS*STTNFI*EQLSLFI***WFNYYHLPTRKCSFI*PQAKCSEGGCRMAAGTLEPV
Consensus                                                                                                                                 * *

Teleogryllus_occipitalis        PPSPRLPPS---------------------------------------------------------PPPPPPPPPPSPPRAS-------------------------------------------
Tarbinskiellus_portentosus      PDSDFLP--------------------------------IVYYCISKIVIFS-------------HNSPNKFQSTFYNISVL*LFGIVQIFLRATHTAGVNTLISIKINMIDIL**TRK
Gryllus_bimaculatus             PLSRRLPDSQAVTDSGRGSLSSNFFICSYLE*LLVF**LVISFLCV--LVFFYFYFLFFLFLCAY*QSQTKPFPASVEHQSMEDLRIN*NITEGKTYSF-TNQS*NVFINILQPFSES--
Consensus                       * *  **                                                                                     *

Teleogryllus_occipitalis        ----------------------------------------------------------------------------------------------------------------RVR------
Tarbinskiellus_portentosus      STAELCFKIYCLKIKKYLNY*NA*MNLSTCLL*TNHTPLIYTYLIFNILIYFSES*IVFYCYTETV*Q*NYSVC*I-CNDLFSKETL--IVLVILFKTPHYSF*ISKQAPMRCNTHAMSK
Gryllus_bimaculatus             ------FFIY-FQFLWFSKVISI*FIVCTCYAFRGYTDV*GIFLNN*VNIHYVVSVVQLLC-TGIK*DCNIFMCKIGCKRMLEKEIRHGEGFKLFYK---YWTRIERNASLRVRKYFKNK
Consensus                                                                                                                                   *

Teleogryllus_occipitalis        --------------------------------------------------------PNE--------C*------------------------------------------------
Tarbinskiellus_portentosus      TMSTSKHFSNTID*WLKLKH*MAR*SIFKRLEM*RN*KTELEKILEIWFSGYKYKNIKNRSLKKGT--CS-EW--------------------------EISTSHVSRLV*NNVNVA---
Gryllus_bimaculatus             TS-------------------------KR---------------------WKSN*LPNEKERKNNFVCQS*WPLQYYETLTSSFSFPIFLLFILANILNI*TL*I*QL**MNKNLAVIY
Consensus                                                                              *         *

Teleogryllus_occipitalis        -------------
Tarbinskiellus_portentosus      -------------
Gryllus_bimaculatus             DRVLQIIWGDEKI
Consensus
```

**Tocc.367G0000010.1**

```
Teleogryllus_occipitalis      ATATTTTGAAATTAAAATATTTTGGGTGCGATATTTTAACATCGACCCCCATAGGTTGTGCTCTCTTCGCCATGTCAGAAATCGTCTTTCTGACTGGAATATTAAATAGTCAGTGATACG
Acheta_domesticus             ATATTATGAAAATAAAATATTTTCAGTTAAGGATTTTAATATCGACCCTCATAGGTCG--------TCGCCACGTTAGGTGTCGTCTCT----CTGGAATGT----TAGTCAATGATAGG
Consensus                     ***** ***** ***********  **     ******* ******** ******* *        ****** ** **   ****** *    ******* *    ****** ***** *

Teleogryllus_occipitalis      AGGCGATCTCAGAGTTGAAACAGGTCTTAATGTCTTCTCGGGAAGGAGAAAGAAT-----AAATATAACAAGAAACGAGAAATAAACCAGGAGTC-AGAAAATACGTCTGTTAAAGCCTC
Acheta_domesticus             AGGCGGTCTCTGAGTTGAAAGAGACCTTAATATGTCCTCTGGAAGGAGGAAAAGTACAAGAAATATAAAAGGAATTGAGAAGCAAAGGAGGAGTCAAAAAAATACGTCTGCTAAAGCC-C
Consensus                     ***** **** ********* **  ****** * * *** ******** ** * *     ******** * ***  *****  ***  ******* * ************ ******* *

Teleogryllus_occipitalis      CAACAGTGTTATATTCCTTTTCTTGTGTCGTCGGGCGTTACTTTCAAGTATCCTTGACTCAGACAAATGCGTCTTTGATCTTTTTCGAGTCGTTTTCCTTTCCTCAATTTTTTTCTCTCA
Acheta_domesticus             CAACACTGTTATATCTTTCCT----TGTCGTCGGGCGTTGGTTTCAAGTATCCTTGACTCTGACAAATGCGTCTTTGATCTTCTTCGAATCGTTTTCCTTTCTTCAAATTTTCCCCCTCA
Consensus                     ***** ********   *  *    **************  ****************** ******************** ***** ************ **** ****  * ****

Teleogryllus_occipitalis      ACTTTCGCTCTTTCTTAATTTTCTTCTCGACGTCAGGCTCTCCGATGTATCAACATCTGAAAATGAAACCCTTTATCTGCGCTGTATATCTAATTTATCGCTCTTCGTCGGCCCTGTCAC
Acheta_domesticus             ACTTTCGCTCTTTCTTAATTTTCTTCTCGACTTCAGGCTCTCCAATGCATCAACATCTGAAAATGAAACCCTTTATCTGCGTTGTATATCTAATTTATCGCTCTTCGTCAGCCCCGTCAC
Consensus                     ***************************** *********** *** ********************************* ************************** **** *****

Teleogryllus_occipitalis      CTCTCACGCGCACT-CCCCGGGCAGCACCGCCTTCACAGCGGCCGCCGCCGCCACCGCCACCGCCACCGCCACCGCCACCGCCGCCGCCACCCCCGCCGCCACCCCCGCCACCGCCGCCA
Acheta_domesticus             CTCTCACGTGCACTCCCCCGGACAGCGCCGCTTTCACAGCGGCCGCCGCCGCCCCCG---------------------------------------------------------------
Consensus                     ******** ***** ****** **** **** ******************** ***

Teleogryllus_occipitalis      CCGCCGCCACCGCCGCCACCCCCGCCGCCACCGCCGCCACCGCCGCCACCGCCGCCACCGCCACCGCCACCGCCGCCACCGCCACCGCCACCGCCACCGCCACCGCCAATT
Acheta_domesticus             ---------------------------------------CCGCCGGCGCCGCTGCCACCGCCGCCACCGCCACTGCCGTCGCCGCCGCCTCCCGCGACACCGACACCGCCGCCG------
Consensus                                                            ****** * **** ******************** ***  ******* ** **  ** ***** ******* ***

Teleogryllus_occipitalis      CCACCGCCACCGCTACCGCCACCGCCAATTCCGCCGCCACCACCGCTACCGCCACCACGCCGCCCGCCGCCACCCGCCACCGCCACCGCCACCGCCACCACCACCACCGCCACCGCCGCC
Acheta_domesticus             ------------------------------------------------------------------------------------------------------------------------
Consensus

Teleogryllus_occipitalis      ACCGCCACCGCCGCCACCGTCACCGCCGCAACCGTCACCACCGCCATCGCCGTCTCCACCGCCACCGCCACCGCCACCGCCACCGCCACCGCCACCGCCACCGCCACCGCCACCGCCACC
Acheta_domesticus             ------------------------------------------------------------------------------------------------------------------------
Consensus

Teleogryllus_occipitalis      GCCACCGCCACCGCCTCTACACCCGCCACCGCCACCACCACAGCCACCGCCGCCACCGTCACCACCGCCTCCACCACCCCACCGCCATCGCCATCGCCATCGCCACCACCACCACCGCCA
Acheta_domesticus             ------------------------------------------------------------------------------------------------------------------------
Consensus

Teleogryllus_occipitalis      CCACCACCACCACCACCACCACCACCACCACCACCACCACCACCACCACCACCACCACCACCACCGCCACCACCACCACCACCACCACCGTCACCACCGACACCACCACCACCACCACCGTCA
Acheta_domesticus             ------------------------------------------------------------------------------------------------------------------------
Consensus

Teleogryllus_occipitalis      CCACCGCCACCGTCACCGCCGCAACCGTCACCACCGCCACCGCCACCGCCACTACCCCCGCCACCGCCACCACCACCGCCGCCATCACCACCGCCACCGCCACCACCGCCGCCGCCGCCG
Acheta_domesticus             ------------------------------------------------------------------------------------------------------------------------
Consensus

Teleogryllus_occipitalis      CCACCGCCACCACCACCGCCACCACCACCACCGCCGCCATCACCACCGCCACCGCCACCACCACCGCCACCACCGCCGCCACCGACACCACCGCCATCACCACCCCCACCGCCAACGCCA
Acheta_domesticus             ------------------------------------------------------------------------------------------------------------------------
Consensus

Teleogryllus_occipitalis      TCGCCATCACCACCACCGCCACCACCACCACCACCACCATCACCACCACCACCGTCACCACCGCCACCGTCACCACCGCCACCGTCACCACCGCCACCGTCACCGCCGCAACCGTCACCA
Acheta_domesticus             ------------------------------------------------------------------------------------------------------------------------
Consensus

Teleogryllus_occipitalis      CCGCCACCGCCACCGCCGCCGCCACCGCCACCGCCACCGCCACTACCCCCGCCCCCGCCACCACCACCGCCGCCATCACCACCGCCACCGCCACCGCCACCACCGCCGCCGCCGCCGCCG
Acheta_domesticus             ------------------------------------------------------------------------------------------------------------------------
Consensus

Teleogryllus_occipitalis      CCGCCGCCGCCGCCGCCGCCGCCGCCGCCACCGCCACCGCCACCACCACCACCGCCGCCATCACCACCGCCACCGCCACCGCCACCACCGCCGCCACCGCCACCGCCACCGCCACTA
Acheta_domesticus             ------------------------------------------------------------------CCACCGTCGCCACCGCCGCCACCGCCGCCGCCACCGCCACCGCCGCCGCCACCA
Consensus                                                                                       ****** * ** ** ******** ******************** ******* *

Teleogryllus_occipitalis      CCCCCGCCACCGCCGCCATCACCACCGCCACCGCCACCGCCACCGCCACCGCCACCGCCACCACCGCCACCACCGCCGCCGCCGCCGCCGCCGCCGCCGCCGCCGCCATCAGCA
Acheta_domesticus             CCGCCGCCACCGCCGCCGCCACCGCCGCCACCGCCGCCGCCACCGCCGCCACCGCCGCCACCGCCGCCACCGCCGCCACCGCCGCCACCGCCGCCACCGCCGCCACCGCCGCCGTCGGCG
Consensus                     ** **************  **** *********** *********** ** ** *********** ******** *********** ******** ******** ******** ** **

Teleogryllus_occipitalis      CCGACGAGTCGTTCCTTCCCCTCCAAGGGTAAAAGGATTCGTCAATTATGCAAATGTTTTAAGTGAAAACCAGAGCGCGCTGTTAGTGGCCGGGCGCCATGCAAATCGGAGCTGCCGCCC
Acheta_domesticus             CCGGCGAGTCGTTCCTTCCCCTCCAAGGGTAAAAGGATTCGTCAATTATGCAAATGTTTTAAGTGAAAACCAGAGCGCGCTGTTAGTGGCCAGGCGCCATGCAAATCGGAGCCGCCGCCC
Consensus                     *** ***************************************************************************** ******************* *******

Teleogryllus_occipitalis      CCGTCGTCCCCCACGCTCCCGAAGCATACAACGCCTACGACATCCCCTCTCCCTCCACATCCTCCCGCAGGCTGAGGTGGCTCAGTGCAGCAACTGATTTCACTGCAGTGCCTCTGACCT
Acheta_domesticus             TC-TCGTCTCCCACGCTCCCGACGCCTACAACGCCTACGACATCCCCGCTTCCTGCACATCCTCCCCCCGGCTGGGGTGGCTCAGAGCATCAACTGATATCACTGCAGTGC--CTGTCCT
Consensus                      * ***** ************* ** ********************* ** *** *********** * ***** ********** *** ******** ************  *** ***

Teleogryllus_occipitalis      AGAGGTCCTTGGGATCGAACCT---TGAAAATTCAAGCGACAGGCTGCTCTGCTTGGTAGTTTTCAGTGGTTTCTCTGATCAATACAAAAAGAGTACCTCCAGAATGCTGGAGGAGTTCC
Acheta_domesticus             GGAGGTCCTAGGGTTCGGGTCTTGATGAGAATTCAAACGCCTGGCTGCTCAAATTGGTGGTTTTCCTTGGTTT-TCTGATCAATACCGGAAGAATACCACCAGAATTCTGGAGCAGGACC
Consensus                      ******** *** ***   **   *** ******* ** * ********   ***** ******   ****** ************   **** **** ******* ****** **  **

Teleogryllus_occipitalis      CTATCAAAG---------------------------------------------------------------------------------------------------------------
Acheta_domesticus             CCATCAAAGAGAACATAATCCATGCGTGATCTTTCCTTCTTCATCCATATACCTCCTCTGCCATACATAAGTATAAGTTATACTTTCATGCACTATATGCCCACACGTCGGGAGAGCAGT
Consensus                     * *******

Teleogryllus_occipitalis      ------------------------------------------------------------------------------------------------------------------------
Acheta_domesticus             CGAGGGCTTAGAACTTGTCTCCGCTGAGGCGGTCCTGGCCTACTGGGGGTGATACCTGGAGGAGAATCCCCGGAAACCTATGTGTATGTCGGTGCCGGATCTCCCCATCCAGCTGGAGAA
Consensus

Teleogryllus_occipitalis      ------------------------------------------------------------------------------------------------------------------------
Acheta_domesticus             GAACACTGGCATCTCCCGTGATAAAAAGGAAAGTAGCTGCGTATTAGAAGAATCACTTCACTTACTAATGAGGACTTGACATGCTCAAAAACGAAATAATGAATCTCCATCCTCCACCCC
Consensus

Teleogryllus_occipitalis      ------------------------------------------------------------------------------------------------------------------------
Acheta_domesticus             GACTCTGCTTCCACCTGTCTGACGTTTCCGTTCTCCGTCACCGCCACAACACTATCTCAAGCTTGTCCTCCTCCCTCGCTGACCCTGCGACCACGTCCCCATTGCCTTCCACCTTTGCTC
Consensus

Teleogryllus_occipitalis      ------------------------------------------------------------------------------------------------------------------------
Acheta_domesticus             CCATTCGACCTCACACTTGTGCTTGCCCTTCGCCGCCATCTCACCCGTCTTCGCCCCGTGCTCACCCCATACTCACATCAACCTCTCCCGACATCGCTCCGTCATCGCCCCATCCTCGTC
Consensus

Teleogryllus_occipitalis      ------------------------------------------------------------------------------------------------------------------------
Acheta_domesticus             CCGCCCTCGCTCCTATCTCAGGCCCGTCCTCACTCCTCTTCGTACGGTTGTCATTACATGGAATTACTTCGAATTGCATACAGTAATAAAACTAATCTTAGCACAACAGTGTTTTCTAAT
Consensus

Teleogryllus_occipitalis      ------------------------------------------------------------------------------------------------------------------------
Acheta_domesticus             ATGAAACAAAAAATTGAAACACTTGACCAAAAAAATGTAATATACCAAATACCATGTAATGACAGTGAATCTACCAATAGGCCAAACTTCACAAATGCTAAAGAAAAGGATTTCACAACA
Consensus

Teleogryllus_occipitalis      ------------------------------------------------------------------------------------------------------------------------
Acheta_domesticus             CAAAAATGACATTAAAAACTTTATTTCTGTGCACTAGCCAAACACTCTCCAGACCATATACACACTTTCGACTGTGAACACACAAAAATCCTAGCCAAAGAAAAGGAACTCAAAAAAGAT
Consensus

Teleogryllus_occipitalis      ------------------------------------------------------------------------------------------------------------------------
Acheta_domesticus             TAACTAAAGAGATGTTGATCATAAATAAAGACAAAAACAAAGCCTGCTATCACAAAACTAATATAGATCAAGTCGGTGCCATCTATAATGGTATTAGCATGACTAGACAATAGAATAGGA
Consensus

Teleogryllus_occipitalis      ------------------------------------------------------------------------------------------------------------------------
Acheta_domesticus             AAAAAGCAACTCAATATGTAATTAGTAGAAATAATTACATATAATGTAATTCTTTATATATCTAAAAATAATGTATATAATAAAAAATTAATATAAAAAAGGAAATATATCAAAAATATA
Consensus

Teleogryllus_occipitalis      ------------------------------------------------------------------------------------------------------------------------
Acheta_domesticus             ATATATGTAATAATAAATAAATTAATAAAAAGAATGGGCAATATATAATAATATGGCCATGTAAATAATAAGTTATAATATTTATGTGTTAAAGATTTTCAACAAGAAAAGTATGTGTTA
Consensus

Teleogryllus_occipitalis      ------------------------------------------------------------------------------------------------------------------------
Acheta_domesticus             TGTTGGTAGGACTAGCCACAAGAGATGTGATCTGTGTAGGAATGGTGGTCGAGATGAGGTGCAGTAAAGGAGGAGAGAAGGGAAATTAAAGATGATACATGATCATTTCGTTAAACAGGA
Consensus

Teleogryllus_occipitalis      ------------------------------------------------------------------------------------------------------------------------
Acheta_domesticus             TTAATACTAAGAATTATACAATAGACTTGTACATTAGATATTTTAGATTAACCTGCCGGCGGGCGCGCGGCAGCACCCCGTGCTGCCTTCGTGTTTTTATTGTTCTCAAAGGGAGTTCGG
Consensus

Teleogryllus_occipitalis      -----------------------------------------------------
Acheta_domesticus             CTGTTTGACATGTCAGGCCGCTTCAGTATTCTCACTTGGCAGCATTCTAAACG
Consensus
```

**Tocc.367G0000010.1_exon1**

```
Teleogryllus_occipitalis    ------------------------------------------------------------------------------------------------------------------------
Acheta_domesticus           NQN*EYLLKLQRKII*LSLIPQYFFNYSSHLHSCTHILSINISYLILISTFFIIINHPLAHVTGLETRRCLYSDTIFNTNCIAKSNHPLCGGNSRSRESSWL*LIVLLGFNASVES*QTG
Consensus

Teleogryllus_occipitalis    ------------------------------------------------------------------------------------------------------------------------
Acheta_domesticus           RKPRQFLQVLYPILSSVTHLHM*YMNTHVRKTNWSCELVPVEAEPRWWEIPG*EHPLGDV*ISAKYHCPAEEELLCLPRYVMFT*LVS*RARRHEIISDTISRILAR*ITEKNINKRAK*
Consensus

Teleogryllus_occipitalis    ------------------------------------------------------------------------------------------------------------------------
Acheta_domesticus           HTQSFLPLPRSVLSPYVMVKSILQLIFNKTVYCNVQYYTMNTSVAVHSISCHSLLEL*SRAQQVPWKLKKTSLYCYMT*SFTLTSHDIHVYK**HLTHILTYLFYHTWQETSQKIRDCHG
Consensus

Teleogryllus_occipitalis    ------------------------------------------------------------------------------------------------------------------------
Acheta_domesticus           GRRYANPPQDDMYVGARPPHPV*EAHRHCTG*NIHSTNGSSEQQLSCPETTTINQFRCVSFSSCNV*RYNLAVETELCTFSAKSDCSAVFITLPLKPL*PRWNITWHFASFSLVPVDVA*
Consensus

Teleogryllus_occipitalis    ------------------------------------------------------------------------------------------------------------------------
Acheta_domesticus           SQTLLHTVAIIRYLAASLKKTRNTATYSIVKTGLEGQSYKNRTIE**TSH*H*AAVM*LSMIMKY**VMQWKERKTTQT**RPP*YYENKIFSVKDFNIDPHRSSPR*VSSLWNVSQ**E
Consensus

Teleogryllus_occipitalis    ------------------------------------------------------------------------------------------------------------------------
Acheta_domesticus           AVSELKETLICPLEGGKVQEI*KELRSKGGVKKIRLLKPQHCYIFPCRRALVSSILDSDKCVFDLLRIVFLSSNFPPQLSLFLNFLLDFRLSNASTSENETLYLRCISNLSLFVSPVTSH
Consensus

Teleogryllus_occipitalis    ------------------------------------------------------------------------------------------------------------------------
Acheta_domesticus           VHSPGQRRFHSGRRRPRRRRRCHRRHRHCRRRRLPRHRHRRRHRRHRRHRRRHRHRRRHHRRHRRRHRRHRRRHRRHRRHRRHRRHRRHRRHRRRRRRRVVPSPPRVKGFVNYANVL
Consensus

Teleogryllus_occipitalis    ------------------------------------------------------------------------------------------------------------------------
Acheta_domesticus           SENQSALLVARRHANRSRRPLVSHAPDAYNAYDIPASCTSSPRLGWLRASTDITAVPVLEVLGFGS**EFKRLAAQIGGFPWFSDQYRKNTTRILEQDPIKENIIHA*SFLLHPYTSSAI
Consensus

Teleogryllus_occipitalis    ------------------------------------------------------------------------------------------------------------------------
Acheta_domesticus           HKYKLYFHALYAHTSGEQSRA*NLSPLRRSWPTGGDTWRRIPGNLCVCRCRISPSSWRRTLASPVIKRKVAAY*KNHFTY**GLDMLKNEIMNLHPPPRLCFHLSDVSVLRHRHNTISSL
Consensus

Teleogryllus_occipitalis    ------------------------------------------------------------------------------------------------------------------------
Acheta_domesticus           SSSLADPATTSPLPSTFAPIRPHTCACPSPPSHPSSPRAHPILTSTSPDIAPSSPHPRPALAPISGPSSLLFVRLSLHGITSNCIQ**N*S*HNSVF*YETKN*NT*PKKCNIPNTM**Q
Consensus

Teleogryllus_occipitalis    ------------------------------------------------------------------------------------------------------------------------
Acheta_domesticus           *IYQ*AKLHKC*RKGFHNTKMTLKTLFLCTSQTLSRPYTHFRL*THKNPSQRKGTQKRLTKEMLIINKDKNKACYHKTNIDQVGAIYNGISMTRQ*NRKKATQYVISRNNYI*CNSLYI*
Consensus

Teleogryllus_occipitalis    -------------------------------------------------------------------------------------------M----------------------------
Acheta_domesticus           K*CI**KINIKKEIYQKYNICNNK*INKKNGQYIIIWPCK**VIIFMC*RFSTRKVCVMLVGLATRDVICVGMVVEMRCSKGGEKGN*R*YMIISLNRINTKNYTIDLYIRYFRLTCRRA
Consensus                                                                                               *

Teleogryllus_occipitalis    ------------------------------------
Acheta_domesticus           RGSTPCCLRVFIVLKGSSAL*HVRPLQYSHLAAF*T
Consensus
```

**Tocc.367G0000010.1_exon2**

```
Teleogryllus_occipitalis    -----------------------------------------------------------------------------------------------------------------------P
Acheta_domesticus           PKLRISFKTSEKDNLTIINPSIFF*LFFTSSFLYPHT*YKYLILNINQHVLYHNKPSPSSCYRVGDPTLPV*RYNFQH*LYS*VKSSTVWGEF*KSGKQLALIDSSPWF*CFSRIITNGP
Consensus                                                                                                                                  *

Teleogryllus_occipitalis    TAATVTTAS---------------------------------------------------------------------------------------------------------------
Acheta_domesticus           QTAPIPSSSLPYPLLCYPFTHVVHEHTCQENKLELRACPS*GGTSLVGDTWIGTPSR*RVDQRQISLSSGGRTPLPPAIRYVYMTSELTRPET*NNQ*HNQSNSS*VNNREEH*QTS*IA
Consensus                     *     *

Teleogryllus_occipitalis    ------------------------------------------------------------------------------------------------------------------------
Acheta_domesticus           HAIFPSTSSLCTFSLCYG*INTSANFQ*NSLL*RSILHYEHICSSTFHQLPLSFGTIVTSPTGTLETEKNIVILLHDVIIYTDVSRHSCLQIITPYTYTYILILSHMAGDQSEN*RLSRR
Consensus

Teleogryllus_occipitalis    ------------------------------------------------------------------------------------------------------------------------
Acheta_domesticus           QEIRKSSTRRHVCWCPTSPSSVGSTPPLHGIKHSQH*WL**AAVVVPGNNNNQSIQMRVIFILQRLAI*SGC*NRAMYF*R*I*L*CGFYNFAPQAPMTTMEYHMAFRVFLARAGRCRMI
Consensus

Teleogryllus_occipitalis    ---------------------------------------------------------------------------------TT-------------------------------------
Acheta_domesticus           ANTASYGRDHTISSGKSEKDAKYRDIFHREDGP*GAKLQKSHYRIVNIPLTLSCSDVIINDNEVLIGDAMEGKENDSNLIKTTLIL*K*NIFS*GF*YRPS*VVATLGVVSLEC*SMIGG
Consensus                                                                                     **

Teleogryllus_occipitalis    ------------------------------------------------------------------------------------------------------------------------
Acheta_domesticus           GL*VERDLNMSSGRRKSTRNIKGIEKQRRSQKNTSAKAPTLLYLSLSSGVGFKYP*L*QMRL*SSSNRFPFFKFSPSTFALS*FSSRLQALQCINI*K*NPLSALYI*FIALRQPRHLSR
Consensus

Teleogryllus_occipitalis    -------------PPPSPSPSPPPPPPPPPPPPPPPPPPPPPPPPPPPPPPPPPPPPPPPPSPPTPPPPPPSPPPPSPPQPSPPPPPPPLPPPPPPPPPPSPPPPPPPPPPPPPPPPPPPPPPPPP
Acheta_domesticus           ALPRTAPLSQRPPPPPPPAPLPPPPPPLPSPPPPATPTPPPPPSPPPPPPPPPPPPP---P-----------------------------------PPPPPPPPPPPPPPPPPPPPPPPP
Consensus                                *** * * ****** * ****  * ***** ************    *                                  ************************

Teleogryllus_occipitalis    SPPPPPPPPPPPPPPPTPPPSPPPPP------------TPSPSPPPPPPPPPPPSPPPPSPPPPS---------------PPP-------------PSPP--------------------
Acheta_domesticus           PPPPPPPPPPPPPPPPSAPASRSFPSKGKRIRQLCKCFK*KPERAVSGQAPCKSEPPPSRLPRSRRLQRLRHPRFLHILPPAGVAQSIN*YHCSACPGGPRVRVLMRIQTPGCSNWWFSLV
Consensus                    **************  * *   *                *        *  * ****  * *               **               *  *

Teleogryllus_occipitalis    ----PPSPPQPSPPPPPPPPPPP-------------------------------------------------------PPPPLPPPPPPPP-----------------------------
Acheta_domesticus           F*SIPEEYHQNSGAGPHQREHNPCVIFPSSSIYLLCHT*V*VILSCTICPHVGRAVEGLELVSAEAVLAYWG*YLEENPRKPMCMSVPDLPIQLEKNTGISRDKKESSCVLEESLHLLMR
Consensus                       *    * *   *      *                                                       *  *     *  *

Teleogryllus_occipitalis    --------PPSPPPPPPPPPPPPPPPPPPPPPP--------PPPPPPPPP--------------PPPPPSPPPPPPPP-------------PPPPPPPPPLPPPPPP-----------
Acheta_domesticus           T*HAQKRNNESPSSTPTLLPPV*RFRSPSPPQHYLKLVLLPR*PCDHVPIAFHLCSHSTSHLCLPFAAISPVFAPCSPHTHINLSRHRSVIAPSSSRPRSYLRPVLTPLRTVVITWNYFE
Consensus                             **   *   **      * **         *  *    *               *    **   *  *              *    *   * *   *

Teleogryllus_occipitalis    ------------------------------------------------------------------------------SPPPPPPPPPPPPPPPPPPPPPPPPPPPPPPPPPPSAPTSRSFPS
Acheta_domesticus           LHTVIKLILAQQCFLI*NKKLKHLTKKM*YTKYHVMTVNLPIGQTSQMLKKRISQHKNDIKNFISVH*PNTLQTIYTLS------------------------------TVNTQKS*PK
Consensus                                                                                                 *                                  *  * *

Teleogryllus_occipitalis    K--GKRI---------RQ------------------------------------------------------------------------------------------------------
Acheta_domesticus           KRNSKKIN*RDVDHK*RQKQSLLSQN*YRSSRCHL*WY*HD*TIE*EKSNSICN**K*LHIM*FFIYLKIMYIIKN*YKKGNISKI*YM***IN**KEWAIYNNMAM*IISYNIYVLKIF
Consensus                   *   * *         **

Teleogryllus_occipitalis    -----------------LCK---------------------------------------------------------CFK*----------------------
Acheta_domesticus           NKKSMCYVGRTSHKRCDLCRNGGRDEVQ*RRREGKLKMIHDHFVKQD*Y*ELYNRLVH*IF*INLPAGARQHPVLPSCFYCSQREFGSLTCQAASVFSLGSILN
Consensus                                    **                                                           **
```

### Supplemental Fig. S6

**Supplemental Fig. S6. AlphaFold3-predicted structures of *de novo* gene candidate proteins in *Te. occipitalis*.** The predicted protein products from all 41 loci (gene IDs shown) were modelled with AlphaFold 3 (default settings) to obtain putative tertiary structures. For each locus, the predicted structure is shown as a ribbon representation colored by model confidence, alongside the corresponding predicted aligned error (PAE) heatmap summarizing positional uncertainty between residue pairs.

### Tocc.1G0004690.1

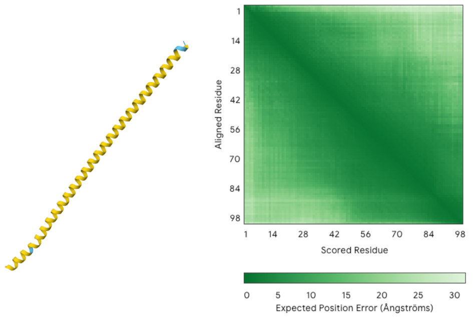


### Tocc.1G0007340.1

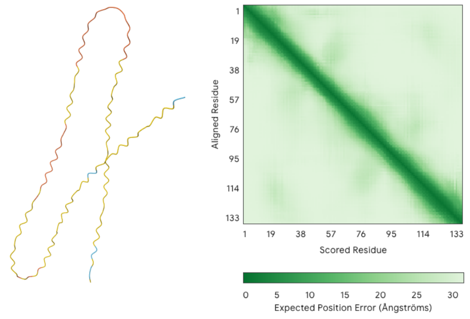


### Tocc.1G0017610.1

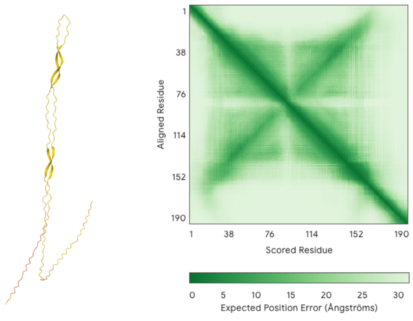


### Tocc.1G0019510.1

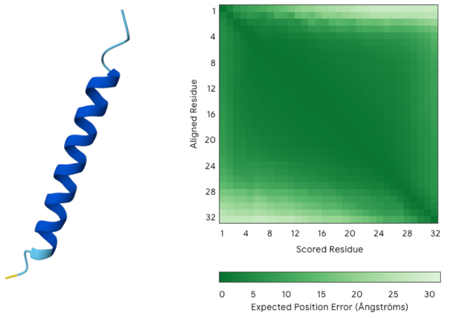


### Tocc.1G0021480.1

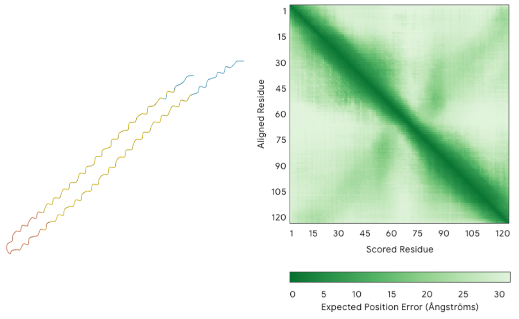


### Tocc.2G0002070.1

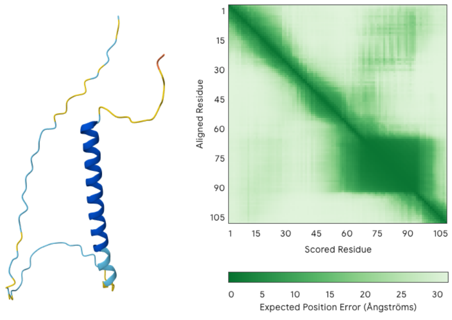


### Tocc.2G0004080.1

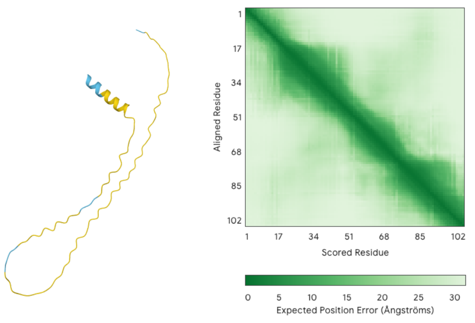


### Tocc.2G0008730.1

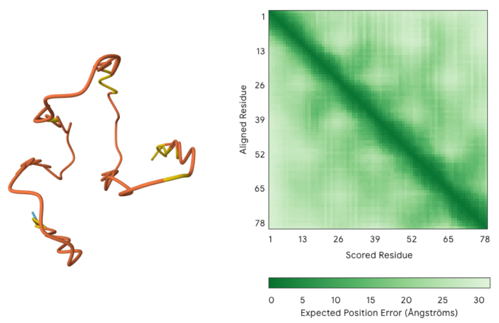


### Tocc.2G0013240.1

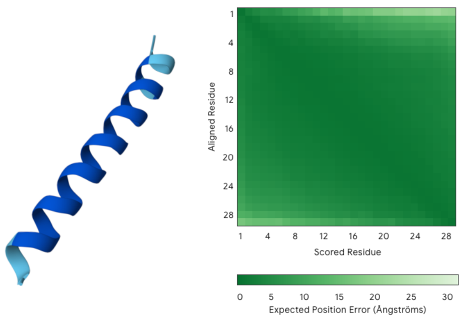


### Tocc.2G0020270.1

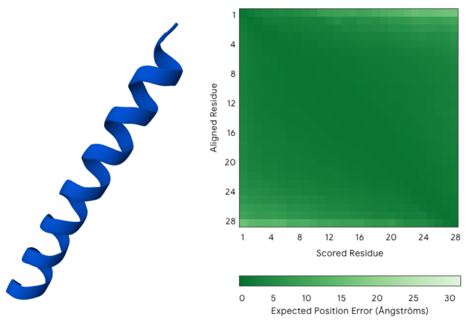


### Tocc.2G0021750.1

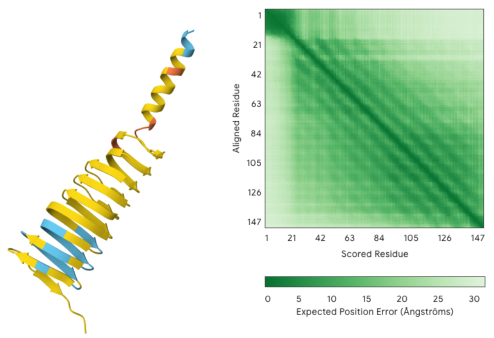


### Tocc.2G0023520.1

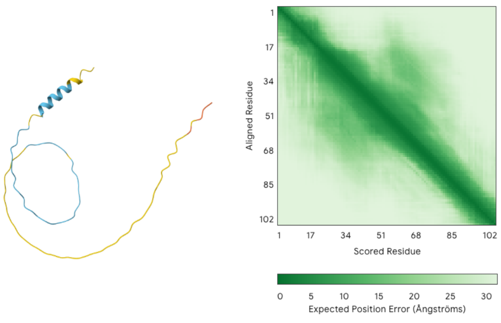


### Tocc.3G0002060.1

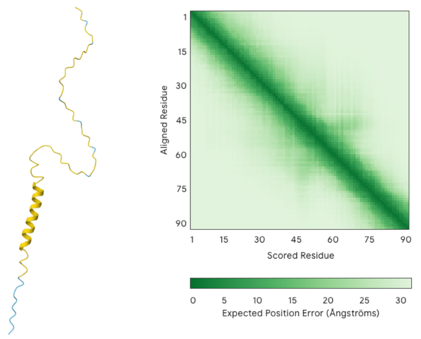


### Tocc.3G0015530.1

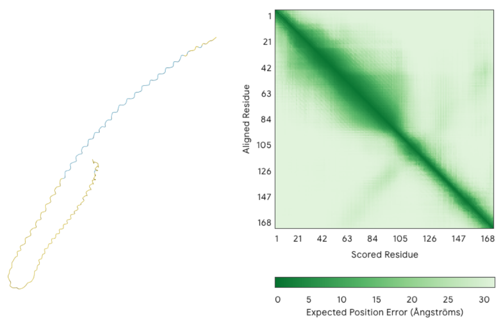


### Tocc.4G0008930.1

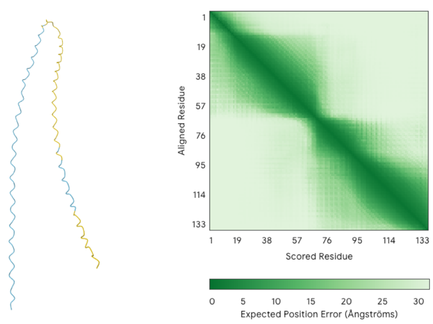

### Tocc.4G0010540.1

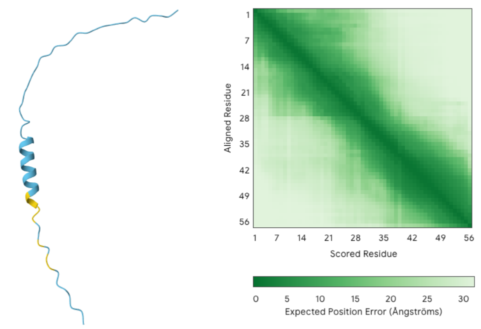


### Tocc.4G0011610.1

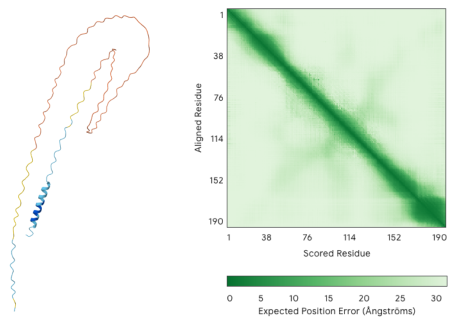


### Tocc.4G0012180.1

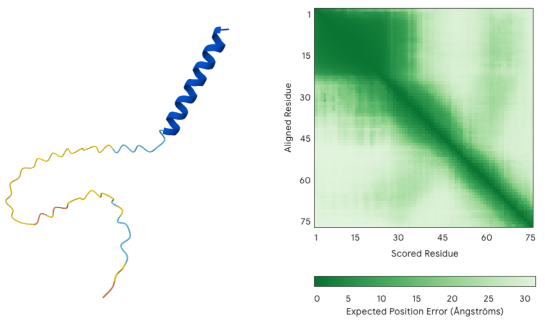


### Tocc.4G0012190.1

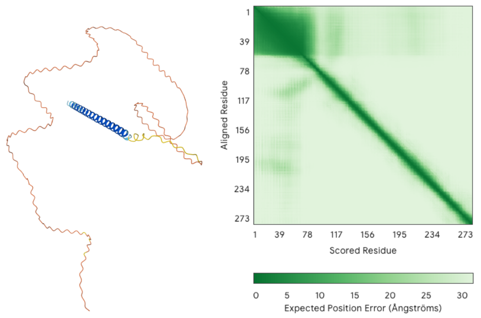


### Tocc.5G0008020.1

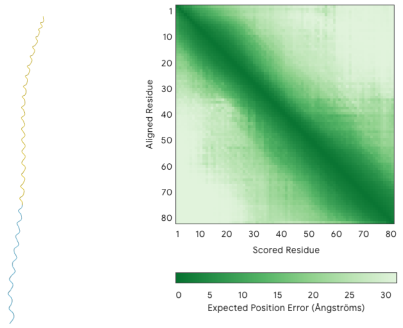


### Tocc.5G0008120.1

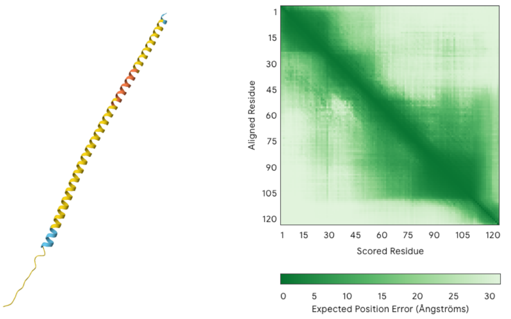


### Tocc.5G0008290.1

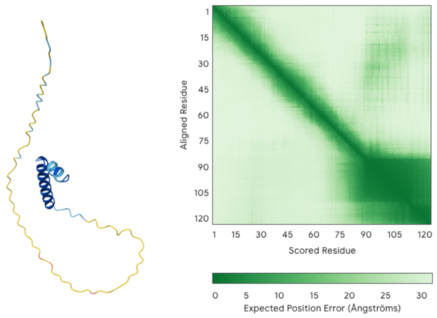


### Tocc.6G0002000.1

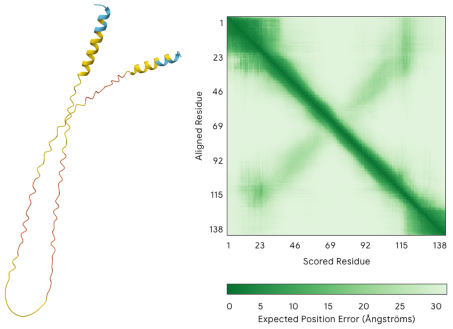


### Tocc.6G0004470.1

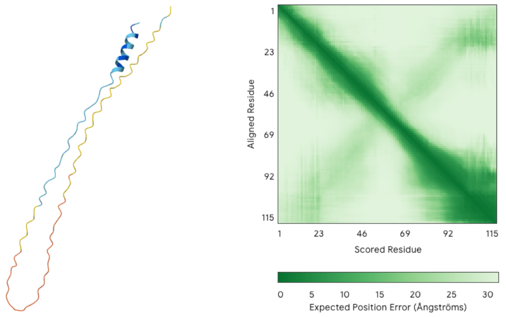


### Tocc.6G0004530.1

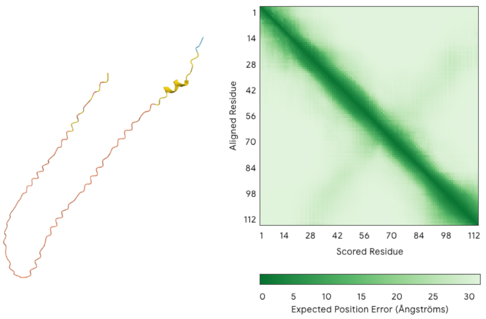


### Tocc.6G0011210.1

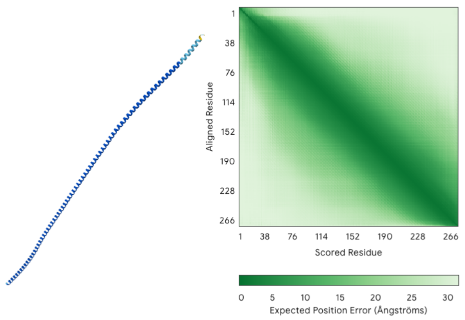


### Tocc.6G0014840.1

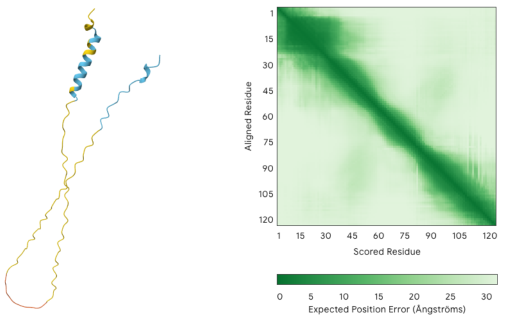


### Tocc.7G0009600.1

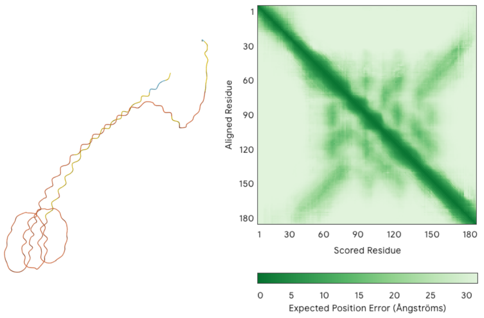


### Tocc.7G0010080.1

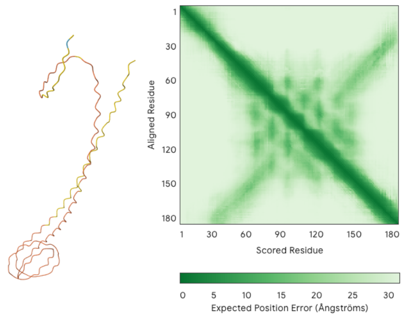


### Tocc.8G0000430.1

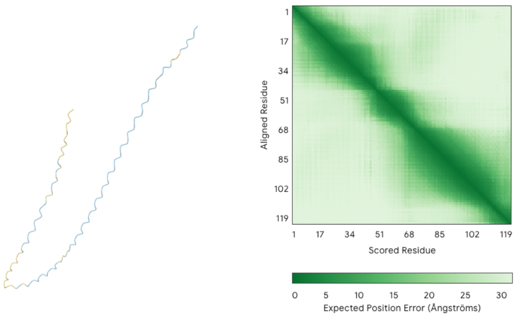

### Tocc.8G0002370.1

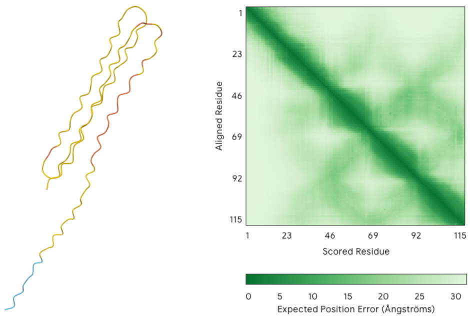


### Tocc.8G0009300.1

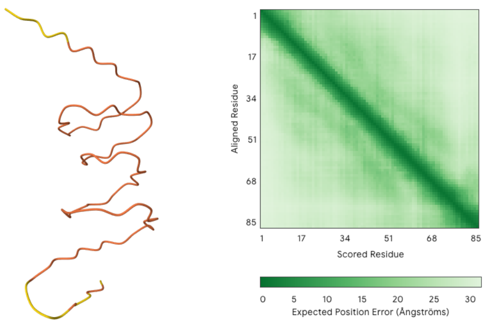


### Tocc.9G0000100.1

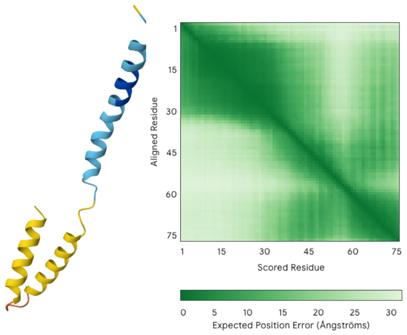


### Tocc.9G0009060.1

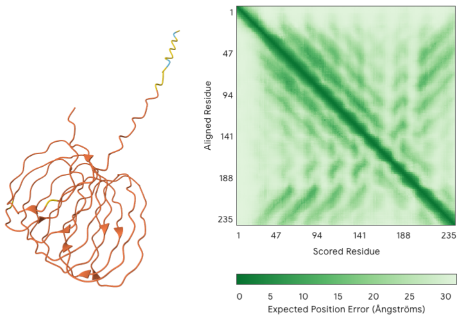


### Tocc.10G0002420.1

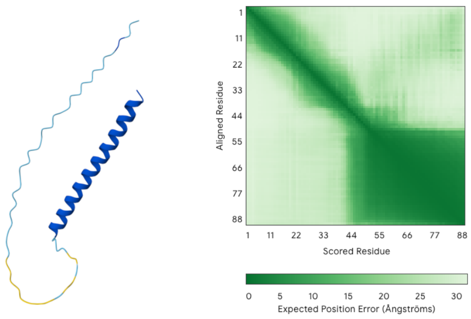


### Tocc.11G0001970.1

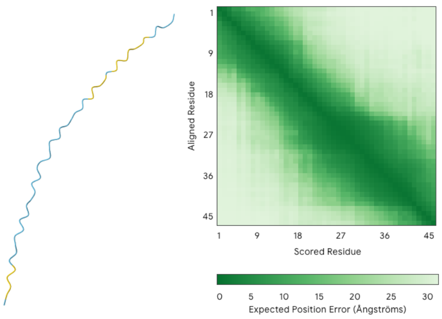


### Tocc.11G0007880.1

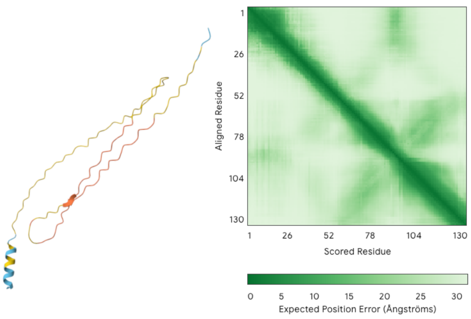


### Tocc.12G0010430.1

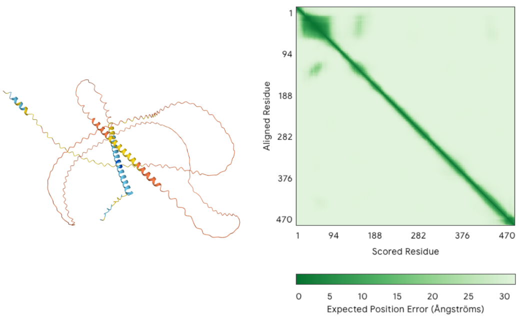


### Tocc.13G0004800.1

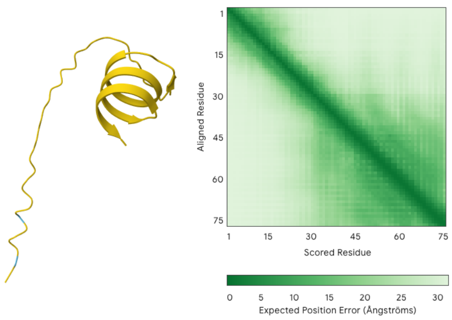


### Tocc.14G0001810.1

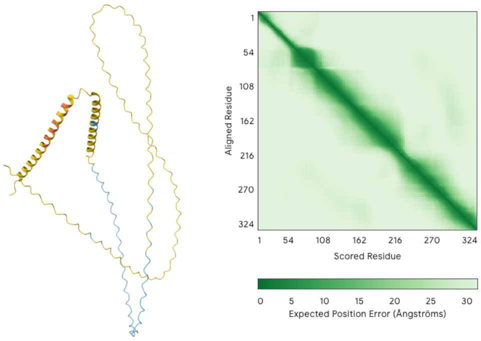


### Tocc.367G0000010.1

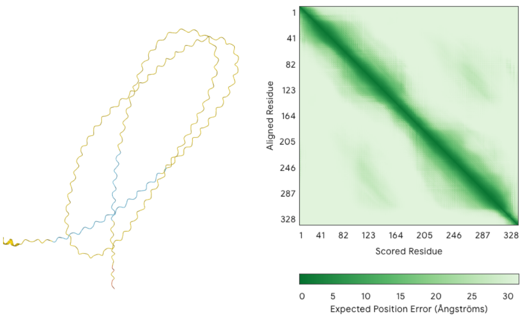

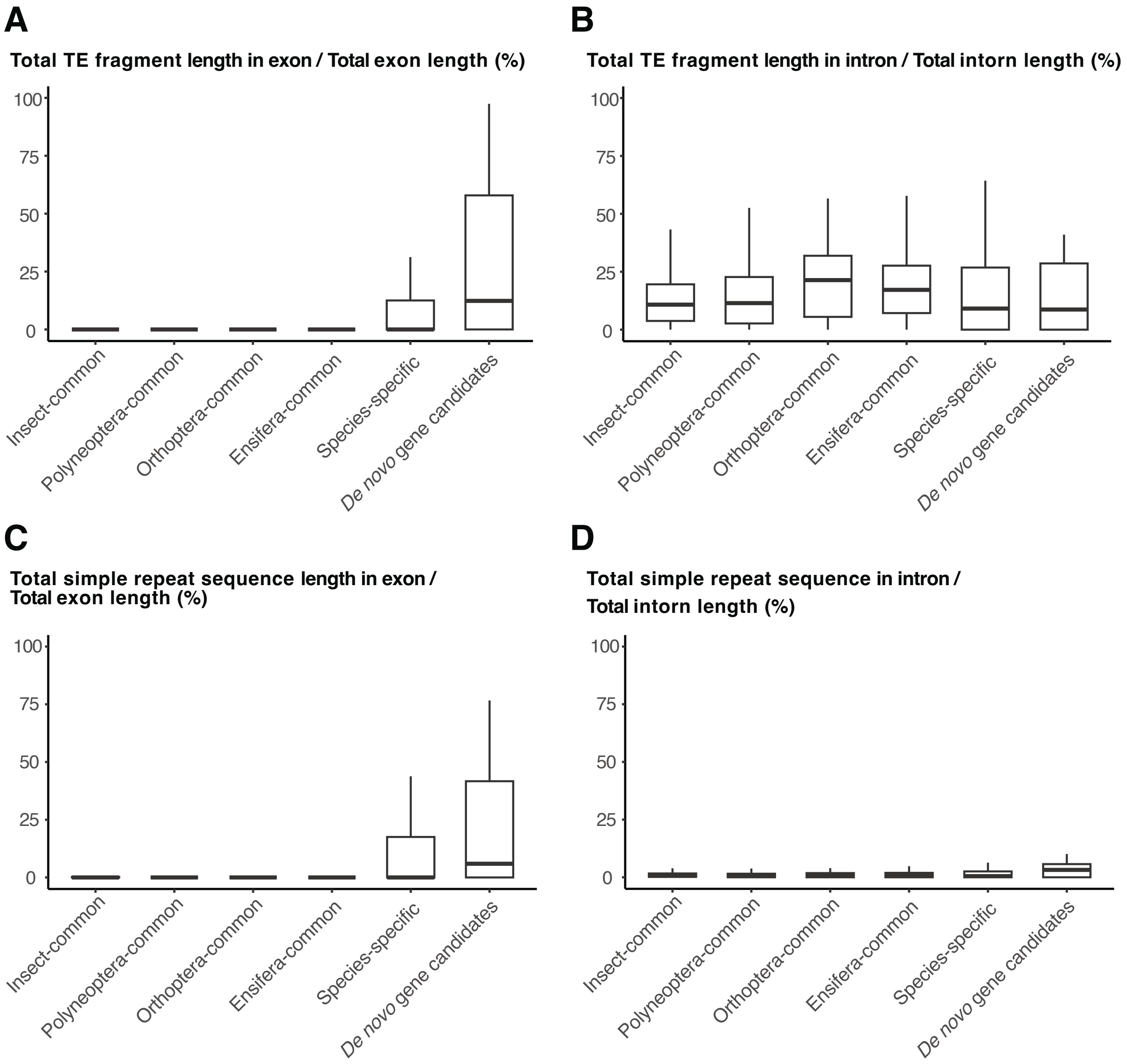


**Supplemental Fig. S7. Enrichment of transposable element fragments and simple repeats in exons and introns across gene age classes.** The proportion of repeat-derived sequence within gene features is shown for gene groups stratified by conservation category (Insect-common, Polyneoptera-common, Orthoptera-common, Ensifera-common, Species-specific) and for *de novo* gene candidates. (**A**) Total TE fragment length in exons divided by total exon length (%). (**B**) Total TE fragment length in introns divided by total intron length (%). (**C**) Total simple repeat sequence length in exons divided by total exon length (%). (**D**) Total simple repeat sequence length in introns divided by total intron length (%).

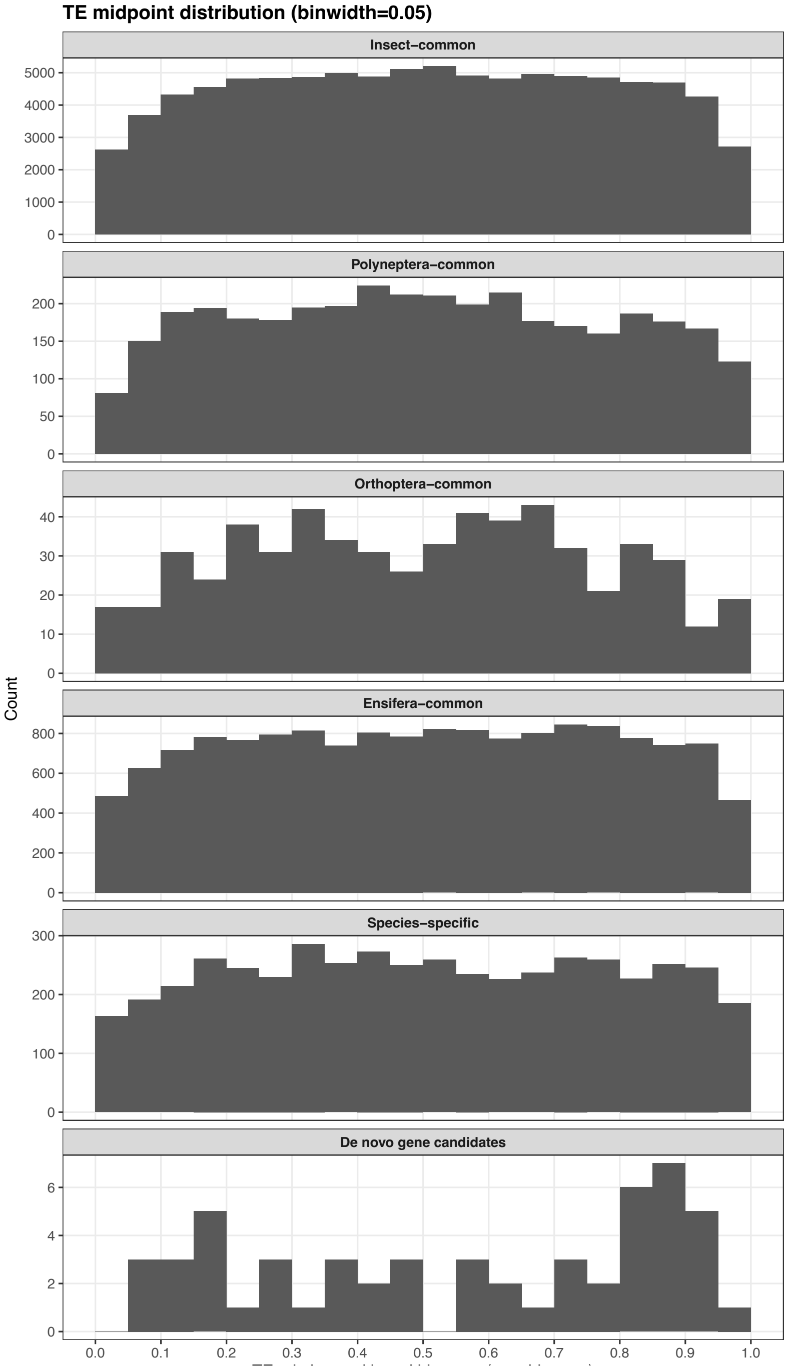


**Supplemental Fig. S8. Distribution of TE midpoints along genes across conservation categories.** Histograms show the distribution of TE fragment midpoints along genes for each conservation category (Insect-common, Polyneoptera-common, Orthoptera-common, Ensifera-common, Species-specific) and for *de novo* gene candidates. The TE midpoint position was normalized to gene length (te_mid_norm), where 0 and 1 correspond to the 5′ and 3′ ends of the gene, respectively. Counts represent the number of TE midpoints in each bin (binwidth = 0.05).

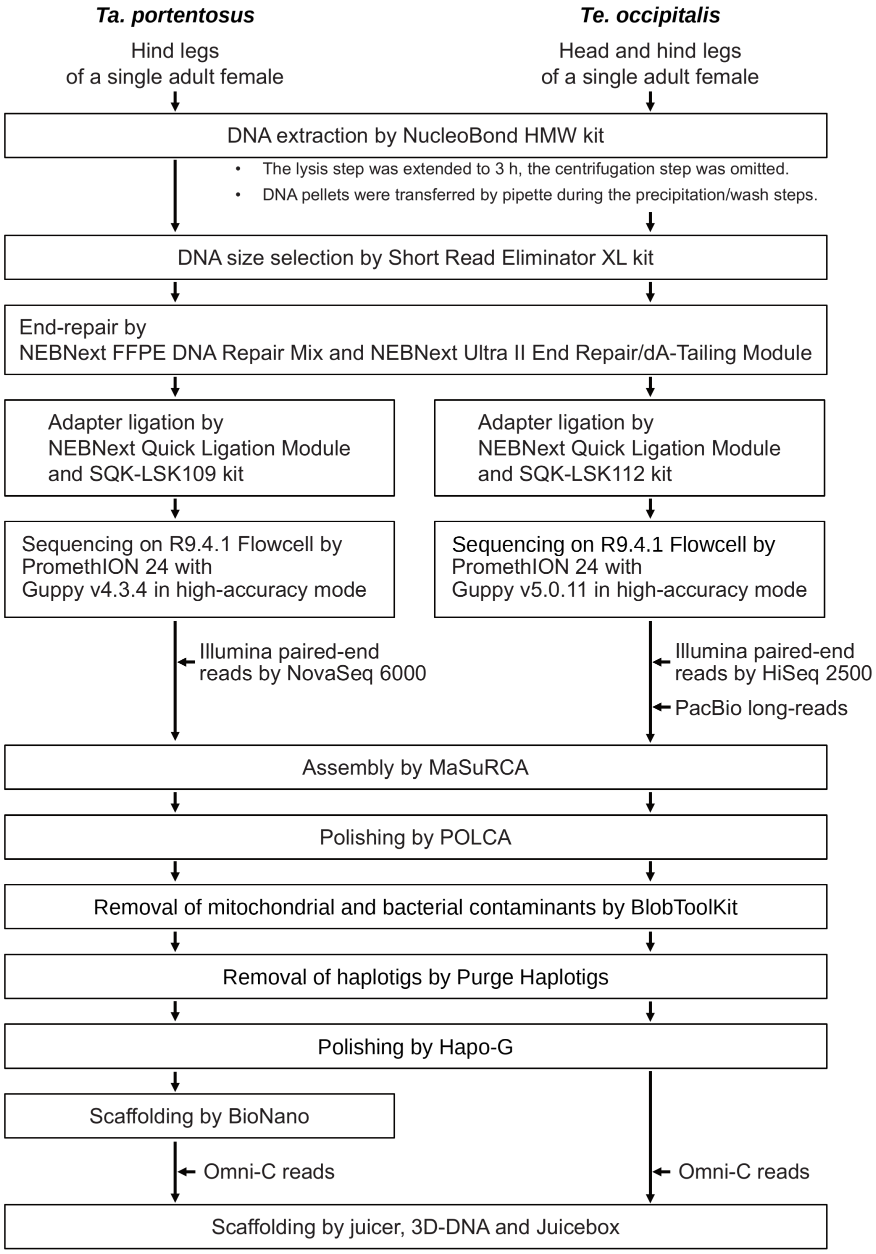


**Supplemental Fig. S9. Genome sequencing and assembly workflow for *Ta. portentosus* and *Te. occipitalis*.** Schematic overview of the pipeline used for high-molecular-weight (HMW) DNA preparation, long-/short-read sequencing, and genome assembly. HMW DNA was extracted (NucleoBond HMW kit) with extended lysis and modified handling steps, followed by size selection (Short Read Eliminator XL). Assemblies were generated with MaSuRCA and polished with POLCA and Hapo-G. Scaffolding was performed using BioNano maps and Omni-C–based scaffolding with Juicer, 3D-DNA, and Juicebox. Redundant haplotigs were removed with Purge Haplotigs, and mitochondrial/bacterial contaminants were removed using BlobToolKit. The workflow also summarizes library-preparation steps for nanopore sequencing (end-repair and adapter ligation) and sequencing configurations (PromethION; Guppy high-accuracy mode), as well as input tissues and datasets (Illumina HiSeq 2500/NovaSeq 6000 paired-end reads, PacBio long reads, and Omni-C reads) for *Ta. portentosus* and *Te. occipitalis*.

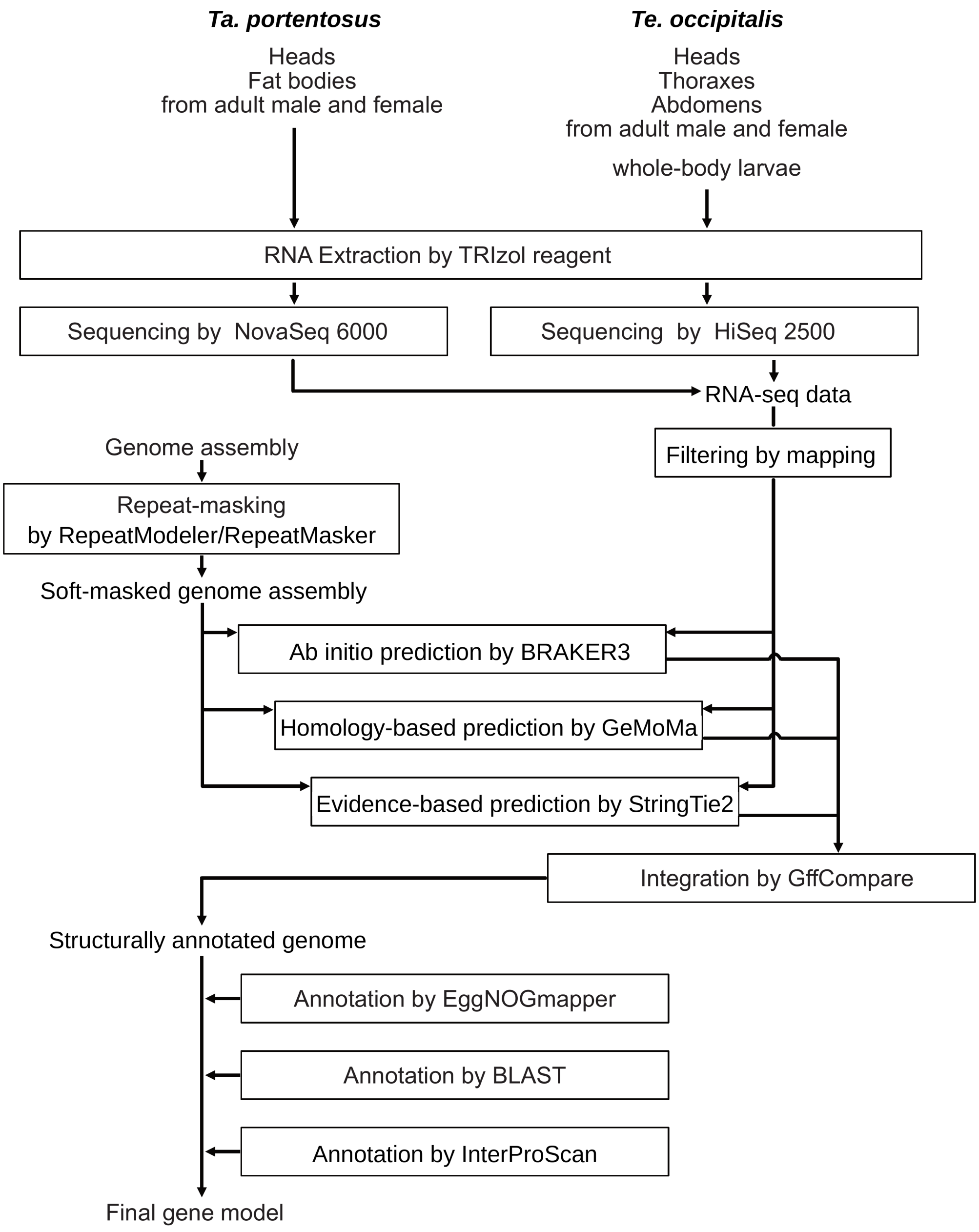


**Supplemental Fig. S10. Transcriptome-based genome annotation workflow for *Ta. portentosus* and *Te. occipitalis*.** Workflow for generating and integrating RNA-seq evidence to produce final gene models. RNA was extracted using TRIzol from multiple tissues and stages (adult male/female heads, thoraxes, and abdomens, and whole-body larvae for *Te. occipitalis*; adult heads and fat bodies for *Ta. portentosus*), followed by sequencing (NovaSeq 6000 and HiSeq 2500) and read filtering by mapping. Structural annotation integrated *ab initio* prediction (BRAKER3), evidence-based transcript prediction (StringTie2), and homology-based prediction (GeMoMa), with gene models merged using GffCompare. Repeat masking was performed with RepeatModeler/RepeatMasker prior to prediction, yielding a soft-masked genome assembly and final gene models. Functional annotation was assigned using eggNOG-mapper, BLAST, and InterProScan.

### Supplemental Table legends

Supplemental Tables S1–S12 are provided as separate Excel files.

**Supplemental Table S1. Statistics for the DNA-seq data for genome *de novo* assembly of two crickets.** Raw yield (Gbp) and read-length metrics for Illumina, ONT, PacBio, and Hi-C datasets used in genome assembly of *Ta. portentosus* and *Te. occipitalis*. Illumina/Hi-C read length refers to paired-end reads (150 bp). N50 is defined as the read length at which 50% of sequenced bases are in reads of that length or longer.

**Supplemental Table S2. Comparison of the quality of initial assemblies among *de novo* assemblers.** Summary statistics of draft assemblies generated with multiple long-read *de novo* assemblers for *Ta. portentosus* and *Te. occipitalis*. For each assembler (version indicated), we report the number of contigs, contig N50 (Mbp), total assembled contig length (Gbp), and longest contig length (Mbp). Assembly completeness was assessed with BUSCO and is shown as the percentage of Complete single-copy, Complete duplicated, Fragmented, and Missing BUSCO genes. Higher contig N50 and BUSCO Complete single-copy values indicate improved contiguity and completeness, respectively.

**Supplemental Table S3. Statistics for the genome assemblies in each step.** Assembly metrics for *Ta. portentosus* and *Te. occipitalis* at successive steps: initial assembly (MaSuRCA), polishing (POLCA, HapoG), decontamination (BLAST-based), heterozygosity/haplotig removal (Purge Haplotigs), and scaffolding (BioNano and Hi-C via juicer/3D-DNA/Juicebox). Hyphens indicate steps not performed.

**Supplemental Table S4. Sources of genomic data of insect species used in this study.** List of insect taxa included in comparative analyses, with their higher-level taxonomy and the corresponding GenBank/RefSeq assembly accession IDs. “This study” indicates genome assemblies generated as part of the present work (*Tarbinskiellus portentosus* and *Teleogryllus occipitalis*). For publicly available genomes, accession IDs are provided as listed in NCBI.

**Supplemental Table S5. Evolutionary divergence categories and tissue-specific transcript expression (TPM) of *Te. occipitalis*.** List of *Te. occipitalis* transcript IDs, their assigned evolutionary divergence categories, and transcript expression levels (TPM) across eight tissues in males and females. Categories (Category_1–Category_9) represent discrete divergence points defined based on the deepest phylogenetic level at which an orthologous/syntenic region was detected (see Methods for category definitions and criteria). Expression values are reported as transcripts per million (TPM) for brain, foregut, flight muscle, gonad, hindgut, midgut, Malpighian tubule, and femur in both sexes.

**Supplemental Table S6. BLASTN similarity of *Te. occipitalis de novo* gene candidates to intergenic regions of other crickets.** For each *Te. occipitalis de novo* gene candidate (rows), BLASTN searches were performed against intergenic regions of each comparative cricket genome. For each target species, columns labeled “CDS_best_bitscore” and “CDS_best_pident” report the bitscore and percent identity of the best BLASTN hit obtained when using the candidate’s CDS sequence as the query. Columns labeled “gene_best_bitscore” and “gene_best_pident” report the best-hit statistics when using the candidate’s gene sequence as the query. Blank cells indicate no hit passing the reporting threshold defined in Methods.

**Supplemental Table S7. Cross-species summary of TE-segment homology within coding regions and associated duplicated gene clusters.** For each TE-derived segment identified in *Te. occipitalis* (TE segment ID; including segment index and genomic span), the table summarizes (i) the genomic context of the segment within the source locus (exon, intron, or both), (ii) TE classes showing sequence homology to the segment, and (iii) homologous gene/transcript matches in other cricket species. For each matched gene/transcript, we report CDS and protein lengths, the coordinates of the TE-homologous region within the CDS (bp) and protein (aa), and the fraction (%) of the CDS/protein covered by TE-segment–homologous sequence. “Copy number” indicates the number of paralogous copies detected for the matched gene within the target species under the study’s clustering criterion, and “Duplicated gene cluster ID” denotes the corresponding duplicate gene group identifier.

**Supplemental Table S8. Summary of duplicated gene clusters by species: counts with/without TE-segment overlap.** For each duplicated gene cluster (DGG) in each species, the table reports the total number of duplicated genes assigned to the cluster and the number of genes that do or do not overlap TE-derived segments. "Overlapping TE segments" indicates genes whose annotated gene body intersects at least one TE-segment identified in this study. "Without TE segments" indicates duplicated genes in the same cluster with no overlap with any TE segment. Counts satisfy: (overlapping TE segments) + (without TE segments) = (total duplicated genes).

**Supplemental Table S9. Evolutionary conservation of repetitive sequence elements identified in *de novo* genes.** Pairwise summary matrix describing the conservation status of repetitive-sequence segments detected within *Te. occipitalis de novo* gene candidates when compared across insect genomes. For each focal species (rows) and target species (column groups), counts of repetitive segments are classified into four categories—Conserved, Neutral, Diverged, and Non-shared—based on sequence similarity of the homologous regions identified in the target species. "Non-shared" indicates that no homologous region meeting the reporting threshold was detected in the target genome. Category definitions and similarity thresholds used to assign segments to the four classes are provided in Methods.

**Supplemental Table S10. Proportion of "overlap-rich" orthogroups in each expected gene age.** For each expected gene-age class, we report the proportion of orthogroups classified as "overlap-rich" with respect to simple repeats or transposable elements (TEs) within exons or introns. Percentages are calculated relative to the total number of orthogroups in the corresponding gene-age class, and the number of overlap-rich orthogroups is shown in parentheses. Definitions of "overlap-rich" and repeat-overlap criteria (simple repeats and TE fragments; exon vs intron) are described in Methods.

**Supplemental Table S11. Conservation of early- (N0) and late- (PostN0) inserted repetitive elements in Category 9 gene families across insects.** Counts of orthologous genes in Category 9 gene families (i.e., gene families maintained since the insect last common ancestor) classified by inferred insertion timing and conservation status of repetitive elements at orthologous loci. "Early (N0)" indicates insertions inferred to have occurred in the insect last common ancestral lineage; "Late (PostN0)" indicates insertions inferred to have occurred after divergence among insect taxa. For each species, genes are categorized as conserved (repeat present at the orthologous locus), neutral or diverged (repeat detected but sequence similarity indicates neutral-like or diverged states), non_shared (repeat present but not shared at the orthologous locus across taxa), no_alignment (orthologous locus not alignable), or insufficient_background (insufficient comparative context to classify). TOTAL is the sum across categories for each timing class.

**Supplemental Table S12. Neutrally conserved TE fragments acquired by genes after the insect radiation.** List of *Te. occipitalis* genes containing TE-derived fragments inferred to represent post-radiation acquisitions that show neutral-level conservation across species comparisons. For each gene, the genomic coordinates of the TE fragment are reported, along with its relative position within the gene, calculated as the normalized midpoint of the TE fragment along the gene body (0 = 5′ end; 1 = 3′ end; 0.5 = gene center).
